# Spatial transcriptomics analysis identifies a unique tumor-promoting function of the meningeal stroma in melanoma leptomeningeal disease

**DOI:** 10.1101/2023.12.18.572266

**Authors:** Hasan Alhaddad, Oscar E. Ospina, Mariam Lotfy Khaled, Yuan Ren, Peter Forsyth, Yolanda Pina, Robert Macaulay, Vincent Law, Kenneth Y. Tsai, W Douglas Cress, Brooke Fridley, Inna Smalley

## Abstract

Leptomeningeal disease (LMD) remains a rapidly lethal complication for late-stage melanoma patients. The inaccessible nature of the disease site and lack of understanding of the biology of this unique metastatic site are major barriers to developing efficacious therapies for patients with melanoma LMD. Here, we characterize the tumor microenvironment of the leptomeningeal tissues and patient-matched extra-cranial metastatic sites using spatial transcriptomic analyses with *in vitro* and *in vivo* validation. We show the spatial landscape of melanoma LMD to be characterized by a lack of immune infiltration and instead exhibit a higher level of stromal involvement. We show that the tumor-stroma interactions at the leptomeninges activate pathways implicated in tumor-promoting signaling, mediated through upregulation of SERPINA3 at the tumor-stroma interface. Our functional experiments establish that the meningeal stroma is required for melanoma cells to survive in the CSF environment and that these interactions lead to a lack of MAPK inhibitor sensitivity in the tumor. We show that knocking down SERPINA3 or inhibiting the downstream IGR1R/PI3K/AKT axis results in re-sensitization of the tumor to MAPK-targeting therapy and tumor cell death in the leptomeningeal environment. Our data provides a spatial atlas of melanoma LMD, identifies the tumor-promoting role of meningeal stroma, and demonstrates a mechanism for overcoming microenvironment-mediated drug resistance unique to this metastatic site.

## INTRODUCTION

Leptomeningeal metastasis is a devastating terminal complication occurring in 5-8% of melanoma patients and remains a major clinical challenge to the treatment of late-stage melanoma patients^1–3^. Even with aggressive treatment, the overall survival for melanoma patients with leptomeningeal disease (LMD) has not altered in several decades and is typically measured in weeks to a few months^1,3^. Despite significant advancements in the treatment of metastatic melanoma using MAPK-targeting therapies and checkpoint inhibitors at extra-cranial sites or even within the brain parenchyma, the majority of tumors at the leptomeninges do not respond to therapy^3,4^. Due to the diffuse nature of the disease, inaccessible site of the tumor, rapid tumor progression, and technological limitations, little is known about the biology of melanoma LMD. We have previously demonstrated that cerebrospinal fluid (CSF) from patients with LMD can modulate BRAF inhibitor responses and may contribute to drug resistance^5^. Single-cell RNAseq characterization of the CSF microenvironment has shown an immune-suppressed cellular landscape in LMD compared to other sites of disease^6^. Therefore, the leptomeninges likely serve as a “sanctuary” site for melanoma cells treated with targeted kinase inhibitors and checkpoint inhibitor therapies. However, these studies only show us a glimpse into the fluid microenvironment of LMD and do not provide a characterization of the leptomeningeal tissues. No previous studies have examined the spatial microenvironment of LMD, and none have compared melanoma tumors adherent to the leptomeninges to extra-cranial sites.

In melanoma LMD, the tumor cells seed to the CSF space and the membrane coverings of the brain and spinal cord termed the leptomeninges^1,2^. The leptomeninges consist of the pia mater, the arachnoid mater, and the CSF space. The pia mater covers the brain’s surface and consists of ICAM1- and SLC38A2-expressing fibroblast-like meningeal cells held together by tight junctions on top of a basement membrane^7,8^. Like melanocytes, these specialized meningeal cells arise from mesenchymal tissues in the neural tube^8^. Many studies have examined the roles of the meningeal cells in neural development and inflammation, but their role in cancer has never been explored^9–12^. The CSF environment is metabolically harsh compared to other sites of disease. Healthy CSF generally lacks micronutrients or proteins^13^, leading us to question how the rapidly growing tumor cells could survive in such an environment. We wondered if the specialized fibroblast-like cells forming the meninges could play a pro-tumorigenic role similar to cancer-associated fibroblasts (CAF) in the LMD microenvironment. In this manuscript, we perform spatial transcriptomic profiling comparing leptomeningeal tissues to patient-matched extra-cranial melanoma metastases. We show that CAF-like activation of the LMD stroma promotes the survival of the tumor cells in the CSF environment and renders them insensitive to MAPK-targeting therapies. This data identifies a mechanism for overcoming the stroma-induced MAPK inhibitor resistance.

## RESULTS

### The spatial landscape of melanoma LMD

To define the cellular landscape of melanoma LMD, we performed an analysis of spatial transcriptomic data from twelve tissue specimens from five patients, including patient-matched tissues from leptomeningeal metastases (5 tissues) and extra-cranial sites (4 tissues) from two melanoma patients with leptomeningeal involvement (**Figure 1A, Supplemental Table 1).** Gene expression analysis immediately highlighted striking differences in the ecosystem of the tumor at the leptomeninges and those at extra-cranial sites from the same patients, including the gene expression profiles of the tumor cells themselves (**Supplemental Figure 1A, B**). Cell type deconvolution analysis identified unique tissue niches at different disease sites, which correlated well with H&E staining of the same tissues (as determined by a qualitative assessment from two independent, experienced pathologists, **Figure 1B, Supplemental Figure 2**). We noted striking differences in stromal and immune infiltration between the leptomeninges and extra-cranial sites (**Figure 1C**). Most extra-cranial samples (3/4) demonstrated some level of immune infiltration, while the majority (4/5) of LMD samples lacked measurable immune infiltration. Meanwhile, tumors at the leptomeninges demonstrate a more profound stromal involvement instead (**Figure 1C, D**).

**Figure 1.**
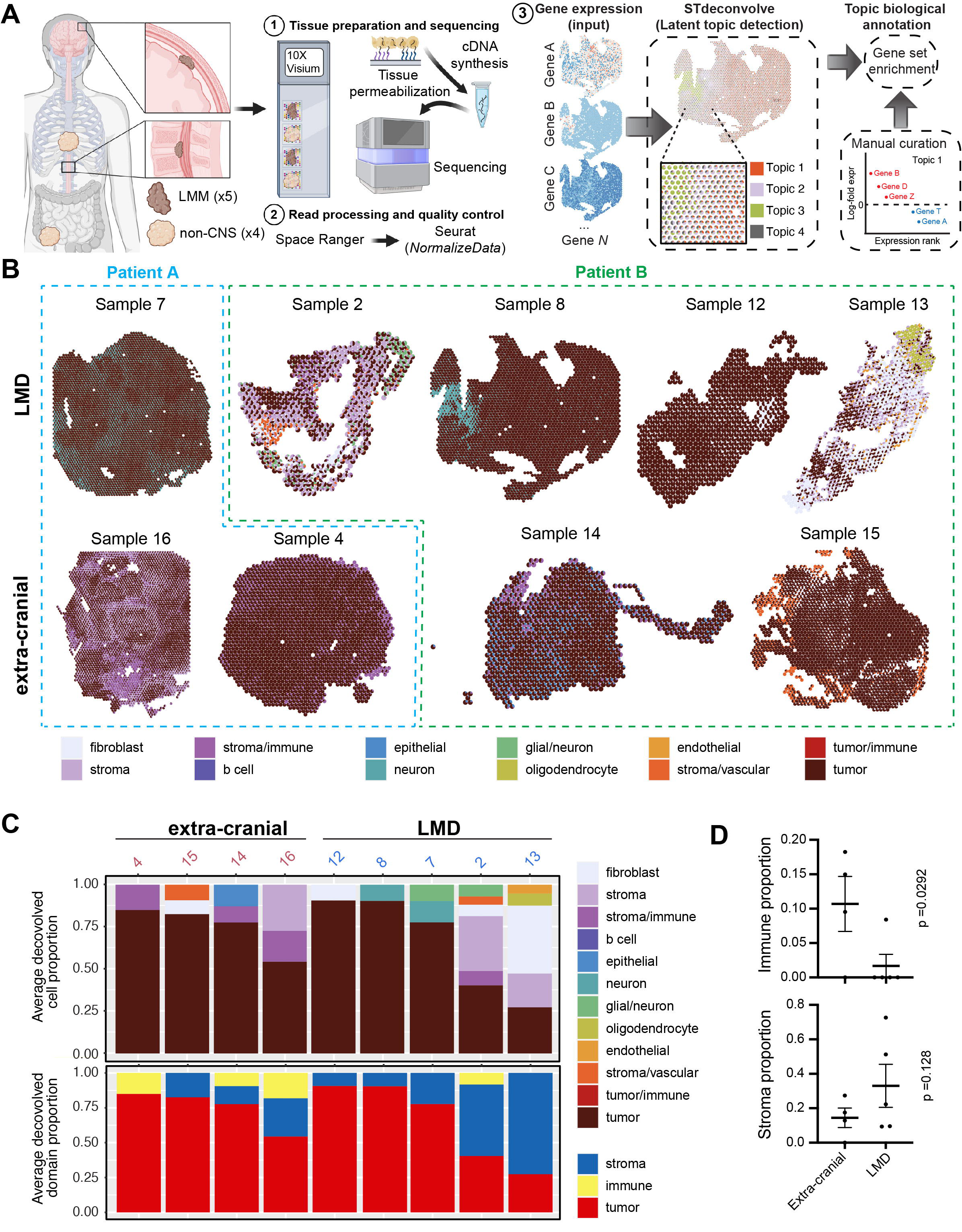
Spatial atlas of melanoma LMD compared to other sites of disease. **A.** Schematic of the spatial transcriptomic analysis workflow, including tissue selection, preparation, sequencing, data quality control, and cell type deconvolution analysis. Created with BioRender.com **B.** Tissue maps showing spatial cell type deconvolution for each spot. The cell type color key is the same for panels B and C. **C.** Bar graphs showing the average deconvolved cell proportion for the major cell types identified in each sample, organized by tissue of origin. **D.** Scatter plots showing the mean proportion of spots predominantly comprised of immune and stromal cells in extra-cranial versus leptomeningeal metastasis samples.

### Spatially driven signaling in melanoma LMD

To identify potential spatially driven signaling in melanoma LMD, we examined KEGG pathway enrichment that exhibited significant spatial distribution patterns. We defined spatial patterns as areas of high expression with respect to the entire tissue (i.e., expression hotspots, see Methods) (**Figure 2A**). The Spearman coefficients, p-values, and spatial pathway maps for all analyzed pathways in all samples are included in **Supplemental Table 2** and **Supplemental Figure 3**. To better visualize regions of the tumor, stroma, and immune infiltration on the spatial maps for consecutive visualizations, we condensed the cell subtype categories into the “tumor,” “stroma,” and “immune” categories (**Supplemental Figures 4,5**). Overall, we identified differences in the spatial expression of several pathways in LMD and extra-cranial metastases (**Figure 2B, Supplemental Figure 3, Supplemental Table 2**). Notably, we have found spatial patterns of signaling previously associated with LMD biology, such as complement and coagulation and TGFβ signaling (**Figure 2B**)^5^. Importantly, we have noted upregulation of several pathways critical for melanoma biology and MAPK inhibitor resistance in areas of tumor-stroma interface, including ECM receptor interaction, insulin signaling, MAPK signaling, phosphatidylinositol signaling, and neurotrophin signaling (**Figure 2B**)^5,14–20^.

**Figure 2.**
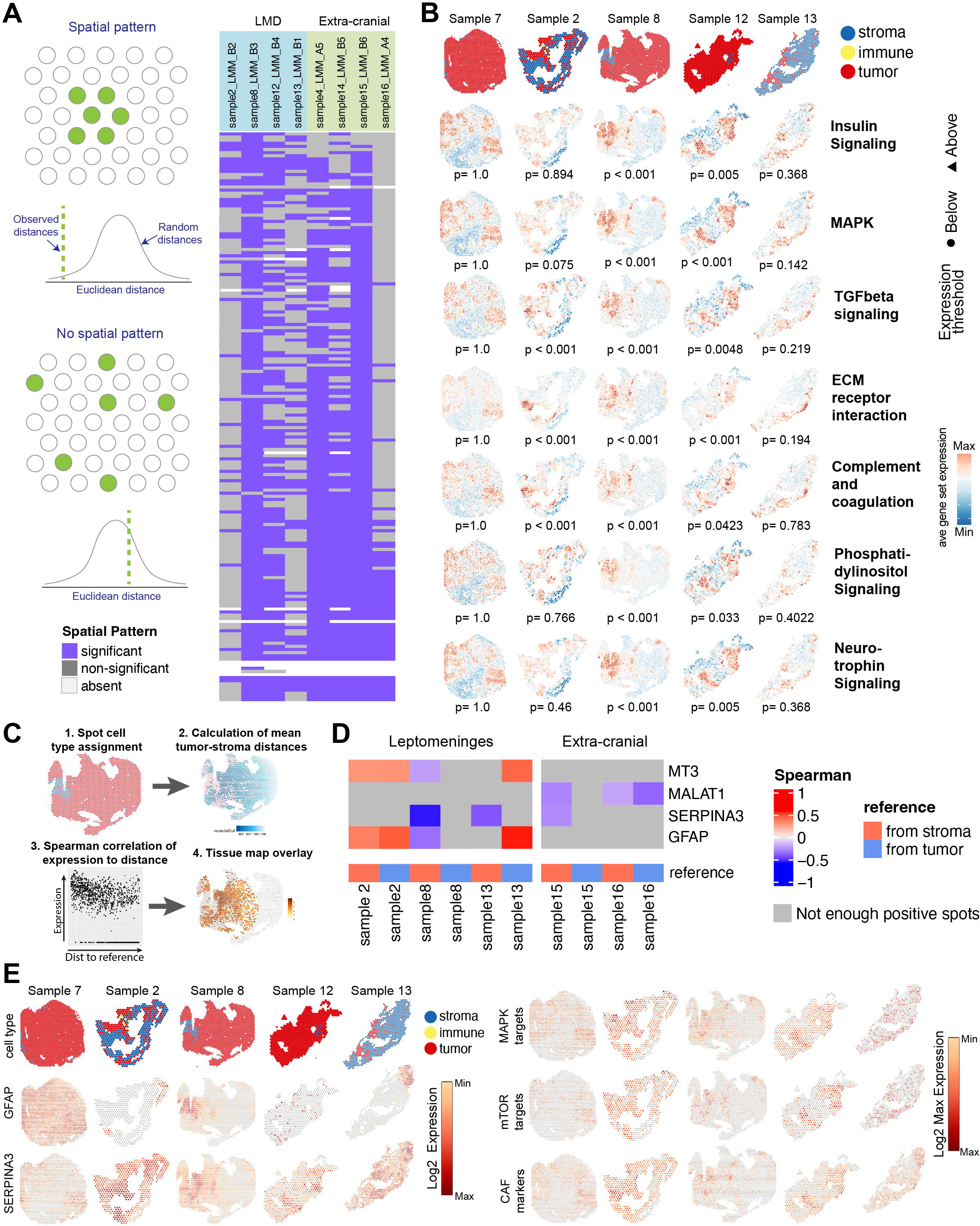
Spatially driven signaling in melanoma LMD. **A.** Schematic showing the basic principles behind the algorithm used to identify KEGG pathway enrichment that have significant spatial patterns (left) and a heatmap highlighting the quantity and similarities/differences of KEGG pathways showing significant spatial pattern within each tissue sample (right). **B.** Spatial tissue maps showing the position of the stroma, immune, and tumor spots along with tissue maps visualizing the average gene set expression for major pathways important in melanoma biology and drug resistance for each LMD sample. **C.** A schematic showing the workflow for determining which genes show a correlation between the level of expression and the distance between tumor and stroma spots. **D.** A heatmap of the genes showing a significant correlation between expression and the distance between tumor and stroma spots across multiple LMD samples. **E.** Spatial tissue maps showing the position of the stroma, immune, and tumor spots along with tissue maps visualizing the log 2 gene expression for GFAP and SERPINA3, and the log2 max expression for sets of genes that are MAPK targets, mTOR targets, and CAF markers (Supplemental Table 4).

### Tumor-stroma interactions in melanoma LMD

The greater extent of tumor-stroma interfaces and the apparent upregulation of several processes integral to melanoma growth and drug resistance in the leptomeningeal tissues prompted us to examine the spatially distinct functional interactions among the melanoma tumor cells and the leptomeningeal stroma. Instead of relying on a subjective determination of a tumor/stroma interface, we developed a novel analysis algorithm that leveraged the spatial distances between tumor and stroma spots and their correlation to the expression of individual genes (**Figure 2C**). Genes demonstrating a spatial correlation for all samples are shown in **Supplemental Table 3 and 4**. The genes with a significant correlation between the level of expression and the spatial proximity between tumor and stromal spots that are enriched in leptomeningeal tissues (patterns observed in multiple samples) include MT3, SERPINA3, and glial fibrillary acidic protein (GFAP) (**Figure 2D**). GFAP upregulation is a known marker of glial scarring related to several neurodegenerative conditions^21,22^. MT3 is well implicated in several neurodegenerative processes, including cerebral ischemia, Parkinson’s, and Alzheimer’s diseases^23^. These findings are consistent with the clinical experience that most patients with leptomeningeal disease develop debilitating multi-focal neurological symptoms, including cranial neuropathies, headaches, ataxia, limb weakness, numbness, pain, or paralysis^2,24–26^. Ischemic infarction and other disruptions caused by increased intracranial pressure due to the impedance of CSF flow may contribute to some of these symptoms. However, the spatial expression gradient we observed in GFAP and MT3 expression, with upregulation at the tumor-stroma interface in leptomeningeal metastases, suggests that neural damage occurs directly at the tumor/stroma interface (**Figure 2E**).

The expression of SERPINA3 correlated with intervals of distance between the tumor and the stromal spots in the leptomeningeal samples. Overall, LMD tissues had a higher proportion of non-zero SERPINA3 expressing spots per sample, although this difference was not statistically significant (**Supplemental Figure 6A**). Three LMD tissues had a sufficient number of stromal cells to examine the changes in the tumor SERPINA3 expression at increasing distance intervals from the stroma, and all three tissues showed a trend of expression decreasing at intervals further away from the stroma (**Supplemental Figure 6B**). Meanwhile, the only extra-cranial metastasis sample with sufficient stromal spots for analysis did not show this trend (**Supplemental Figure 6B**). Re-analysis of our previously published mass-spectrometry-based proteomic profiling of CSF from patients with melanoma LMD shows SERPINA3 to be enriched in patients with LMD compared to no LMD controls (1.21 Log2 ratio, p < 0.001)^5^. SERPINA3 regulates the PI3K and MAPK pathways by activating the IGF1R/integrinα5β1 signaling^27^ (**Supplemental Figure 7A**). Knowing that melanoma tumors at the leptomeninges rarely respond to MAPK inhibitor therapy and grow rapidly, we examined the spatial expression of MAPK targets and targets of mTOR in the leptomeningeal tissues (**Supplemental Table 5**). Not surprisingly, we see a strong upregulation of MAPK and mTOR target expression in the tumor regions at the leptomeninges closer to the tumor-stroma interface (**Figure 2E**). We noted that the expression of SERPINA3 and IGF1R/integrinα5β1 co-localized at areas of greater stromal interface (**Supplemental Figure 7B**), highlighting the potential interaction between the stroma and tumor cells via the SERPINA3 and IGF1R/Integrinα5β1 axis. Together, this data suggests a pro-tumorigenic interaction between the tumor and stroma at the leptomeninges.

The tumor cells at the leptomeninges are typically found floating in the CSF fluid or attached to the pia mater tissues on the surface of the brain^2^. Since the pia mater is comprised predominantly of specialized meningeal fibroblast-like cells^2^, we set to investigate if these unique cells could become cancer-associated fibroblasts. Our data demonstrates an enrichment of CAF marker expression (**Supplemental Table 5**) at the tumor-stroma interface (**Figure 2E**).

### Meningeal stroma supports melanoma growth and drug resistance

The majority of melanoma LMD patients do not respond to MAPK-targeting therapy, and we recently showed that CSF from these patients could modulate BRAF inhibitor responses and contribute to drug resistance^3,5^ To test whether the meningeal stroma could support melanoma survival in the context of MAPK inhibition, we treated GFP-tagged human melanoma cells with vemurafenib (BRAFi) in monoculture or co-culture with human primary meningeal cells. We noted a significant increase in human melanoma cell survival with human meningeal cell co-cultures (**Figure 3A**). The same effect was observed in murine melanoma cells with murine primary meningeal cell cultures (**Supplemental Figure 8A**). Similarly, the culture of melanoma cells with conditioned media from melanoma-meningeal co-cultures also demonstrated a protective effect over a range of vemurafenib concentrations in an MTT assay (**Figure 3B**). Direct co-culture with meningeal cells and stimulation with conditioned media showed similar levels of protection from MAPK inhibition (**Supplemental Figure 8B**). While these experiments were carried out in the context of established normal melanoma cell culture conditions, the leptomeningeal environment is unique because the cells are surrounded by CSF^3^. Utilizing physiological CSF formulated to mimic the composition of healthy human CSF, we have found that not only do meningeal cells promote MAPK drug resistance, but they are also required for melanoma cell survival in CSF. Microscopy images of GFP-positive melanoma cells and MTT assays following treatment in the context of CSF conditioned by melanoma/meningeal cells co-culture (C-CSF) both show profound survival-promoting effects and lack of BRAF inhibitor sensitivity with conditioned CSF (**Figure 3C-D**). We confirmed these findings with an additional melanoma cell line and the BRAF + MEK inhibitor combination using dabrafenib and trametinib (**Supplemental Figure 8C**). Cell cycle analysis of melanoma cells grown in monoculture and co-culture shows no significant changes in BRAF inhibitor-mediated G1 arrest in the context of either CSF or standard media, suggesting the effects are independent of the cell cycle (**Figure 3E, Supplemental Figure 8D**). Western blot analysis validated the results observed in the spatial gene expression data, which indicated that co-localized melanoma and primary meningeal cell regions upregulate SERPINA3 when stimulated with co-culture-conditioned CSF (**Figure 3F, G**). Consistent with an increase in MAPK and PI3K-related signaling observed at the tumor-meningeal stroma interface, we also observed the conditioned CSF to promote recovery of pERK and amplify pAKT following BRAF inhibitor treatment (**Figure 3F**). In meningeal cells, exposure to CSF conditioned by melanoma/meningeal cell co-cultures also promoted cyclin D1, fibronectin, TGFβ1, and TGFβR expression, suggesting that exposure to tumor cells in the CSF space activates pro-tumorigenic, CAF-associated programs in meningeal stroma (**Figure 3G**). We observed similar signaling effects under standard complete media with serum. The basal levels of pERK and pAKT in melanoma cells were much lower when cells were cultured in CSF compared to standard complete medium, and therefore, the meningeal cell-mediated signaling effects are much more pronounced in the CSF co-cultures than under normal media conditions (**Supplemental Figure 9**).

**Figure 3.**
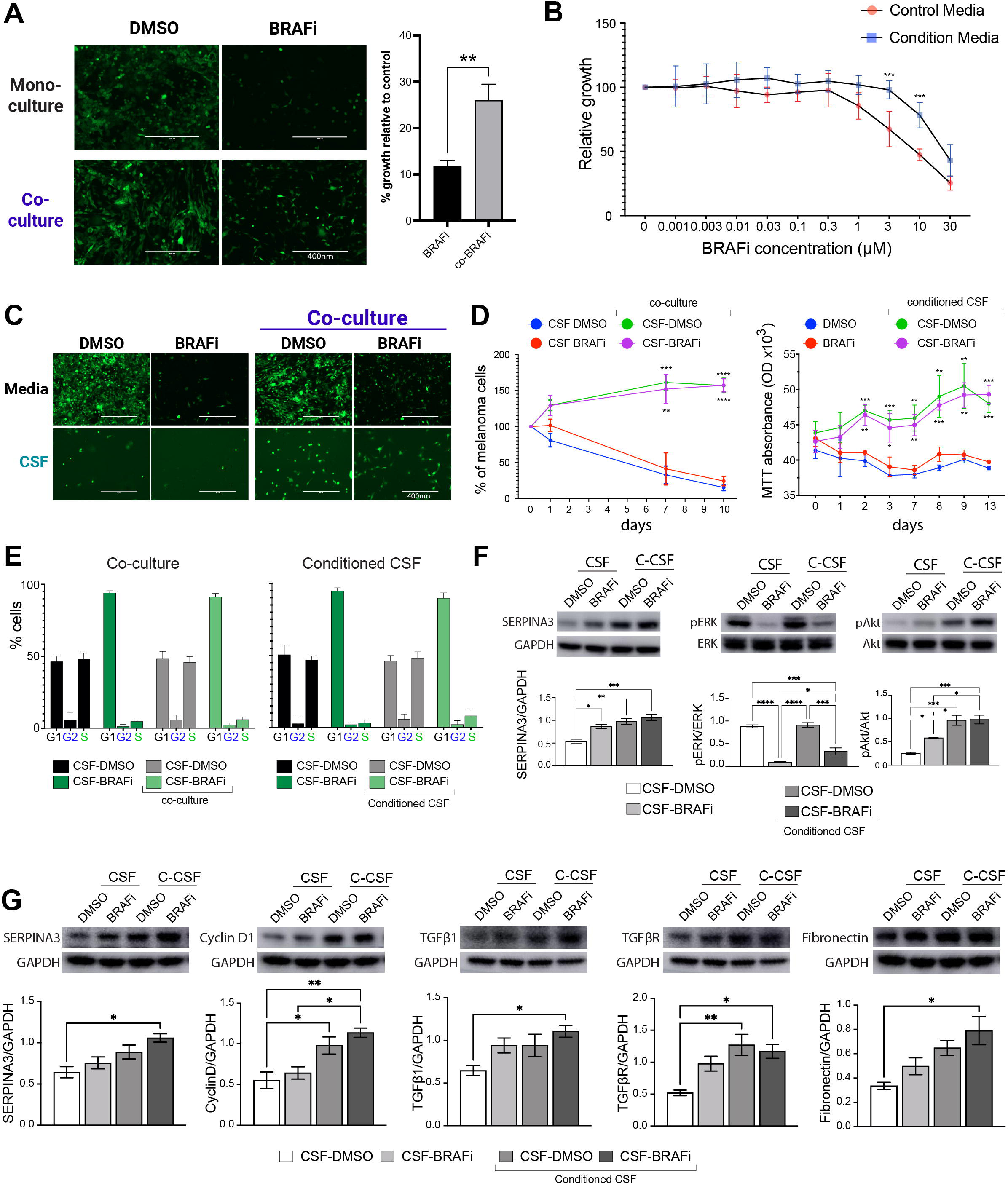
Meningeal cells are required for melanoma survival in the CSF environment. **A.** Representative microscopy images showing GFP-tagged WM164 melanoma cells treated with 3μM vemurafenib (BRAFi) or DMSO control in monoculture versus co-culture with primary meningeal cells (left). Bar graphs show the quantification of GFP+ cells across ten biological replicates relative to the respective vehicle-treated controls (right). **B.** MTT assay showing the relative growth of WM164 melanoma cells treated with increasing doses of vemurafenib (BRAFi) in control media versus media conditioned by the co-culture of melanoma with primary meningeal cells. **C.** Representative microscopy images showing GFP-tagged WM164 melanoma cells treated with 3μM vemurafenib (BRAFi) or DMSO control in monoculture versus co-culture with primary meningeal cells in the context of CSF. **D.** Quantification of the GFP+ WM164 melanoma cells treated with 3μM vemurafenib (BRAFi) or DMSO control in monoculture versus co-culture with primary meningeal cells for ten days in CSF (left). WM164 melanoma cells treated with 3μM vemurafenib (BRAFi) or DMSO control in with regular CSF vs CSF conditioned by co-culture of melanoma with primary meningeal cells over 13 days (right). Significance was calculated between the co-culture or conditioned media arm and the respective monoculture control arm **E.** Flow cytometry assessment of cell cycle using PI staining in WM164 melanoma cells treated with 3μM vemurafenib or DMSO in the context of direct co-culture with primary meningeal cells or with CSF conditioned by co-culture of melanoma with primary meningeal cells. **F.** Western blot analysis of WM164 melanoma cells treated with 3μM vemurafenib or DMSO control in fresh CSF vs conditioned CSF showing abundance of SERPINA3, pERK(Thr202/Tyr204), ERK, pAKT (Ser473) and AKT. **G.** Western blot analysis of primary meningeal cells stimulated with WM164-conditioned CSF showing the abundance of SERPINA3, cyclin D1, TGFβ1, TGFβR, and fibronectin. All statistical significance was assessed using Student’s t-test *p < 0.05, **p < 0.01, ***p < 0.001, ****p < 0.0001.

### Meningeal cells promote tumor growth and MAPKi resistance *in vivo*

We have previously shown that primary human dermal fibroblasts promote melanoma BRAFi resistance^19^. To compare how well meningeal cells promote tumor growth and drug resistance compared to dermal fibroblasts, we injected human melanoma cells into flanks of animals, either by themselves or in combination with primary dermal fibroblasts or primary meningeal cells in a 1:1 ratio (**Figure 4A**). Twenty-four hours following implantation, we placed all animals on either control or drug-formulated chow containing the dabrafenib and trametinib combination (MAPKi). As expected, co-injection of melanoma with dermal fibroblasts promoted rapid tumor growth and diminished sensitivity to MAPKi (**Figure 4B**). Surprisingly, the tumor growth of melanoma cells co-injected with meningeal cells was even faster than those injected with dermal fibroblasts. These tumors were also even more resistant to MAPKi than those co-injected with dermal fibroblasts. Pairwise analyses of covariance among linear fit models of each experimental cohort show that the speed of growth (slope) was indeed significantly different among tumor cells injected by themselves, those co-injected with dermal fibroblasts, and those co-injected with meningeal cells (**Figures 4C-D**). At day 21 post-injection, the animals with melanoma cells injected on their own had not started to form measurable tumors yet, while the animals harboring melanoma cells co-injected with meningeal cells were at or approaching the endpoint (**Figure 4E**).

**Figure 4.**
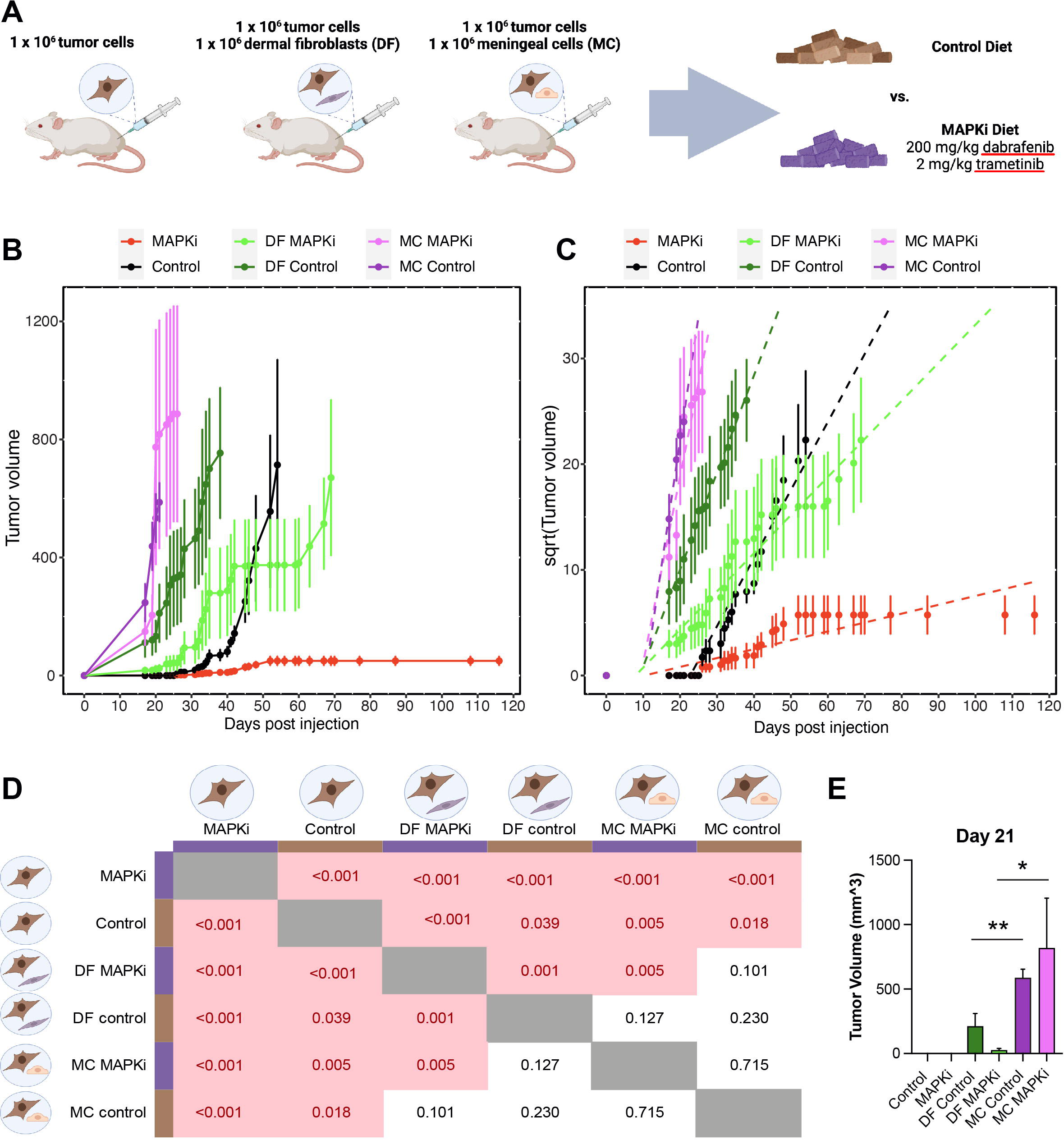
Meningeal cells promote growth and resistance to MAPKi *in vivo.* **A.** A schematic outlining the experimental cohorts of the animal experiment. Mice were injected subcutaneously with 1 million WM164 cells alone, 1 million WM164 cells with 1 million primary meningeal cells (MC), or 1 million WM164 cells with 1 million dermal fibroblast (DF) cells. Mice were fed a control rodent diet (control) or a rodent diet formulated with 200 mg/kg dabrafenib and 2 mg/kg trametinib (MAPKi). Created with BioRender.com **B.** Line graph showing the average growth of tumors in each experimental cohort over time, with error bars depicting standard error. Tumor size was fixed following the endpoint of animals until the last animal in the cohort reached the endpoint. **C.** Square root transformation and slope visualization for each experimental cohort in panel B. **D.** *p*-values for each two-way comparison of slope among experimental cohorts (ANCOVA). Icons created with BioRender.com **E.** Bar graph visualizing the average tumor volume in each experimental cohort on day 21. Data represent mean ± standard error. Statistical significance was assessed using Student’s t-test *p < 0.05, **p < 0.01.

### Inhibition of tumor-meningeal cell crosstalk abrogates tumor survival and drug resistance

Since we observed amplification of AKT phosphorylation during MAPK inhibitor treatment in the context of conditioned CSF, and it has been shown that SERPINA3 promotes AKT signaling via activation of IGF1R^27^, we tested to see if inhibition of IGF1R would re-sensitize cells to MAPK inhibitor therapy. We noted that the meningeal cultures protected melanoma tumor cells similarly in the context of either single-agent BRAF inhibitor or dual BRAF/MEK inhibitor therapy (**Figures 3A-D, Supplemental Figure 8C, Figures 5A-B**, and **Figures 5D-E**). Inhibition of IGF1R using linsitinib successfully blocked the survival-promoting effects of meningeal cell co-culture in CSF (**Figure 5A, B**). The MAPK inhibitor resistance effects were also blocked under standard complete medium conditions (**Supplemental Figure 10**). We then tested whether SERPINA3 is required for the survival-promoting effects of meningeal cell co-culture using siRNA-mediated knockdown of SERPINA3 (**Figure 5C**). Knockdown of SERPINA3 significantly reduced melanoma cell growth and MAPK inhibitor resistance in CSF (**Figure 5D-E**). Once again, the MAPK inhibitor resistance effects were blocked during siRNA-mediated knockdown under normal complete medium conditions (**Supplemental Figure 11A-B**).

**Figure 5.**
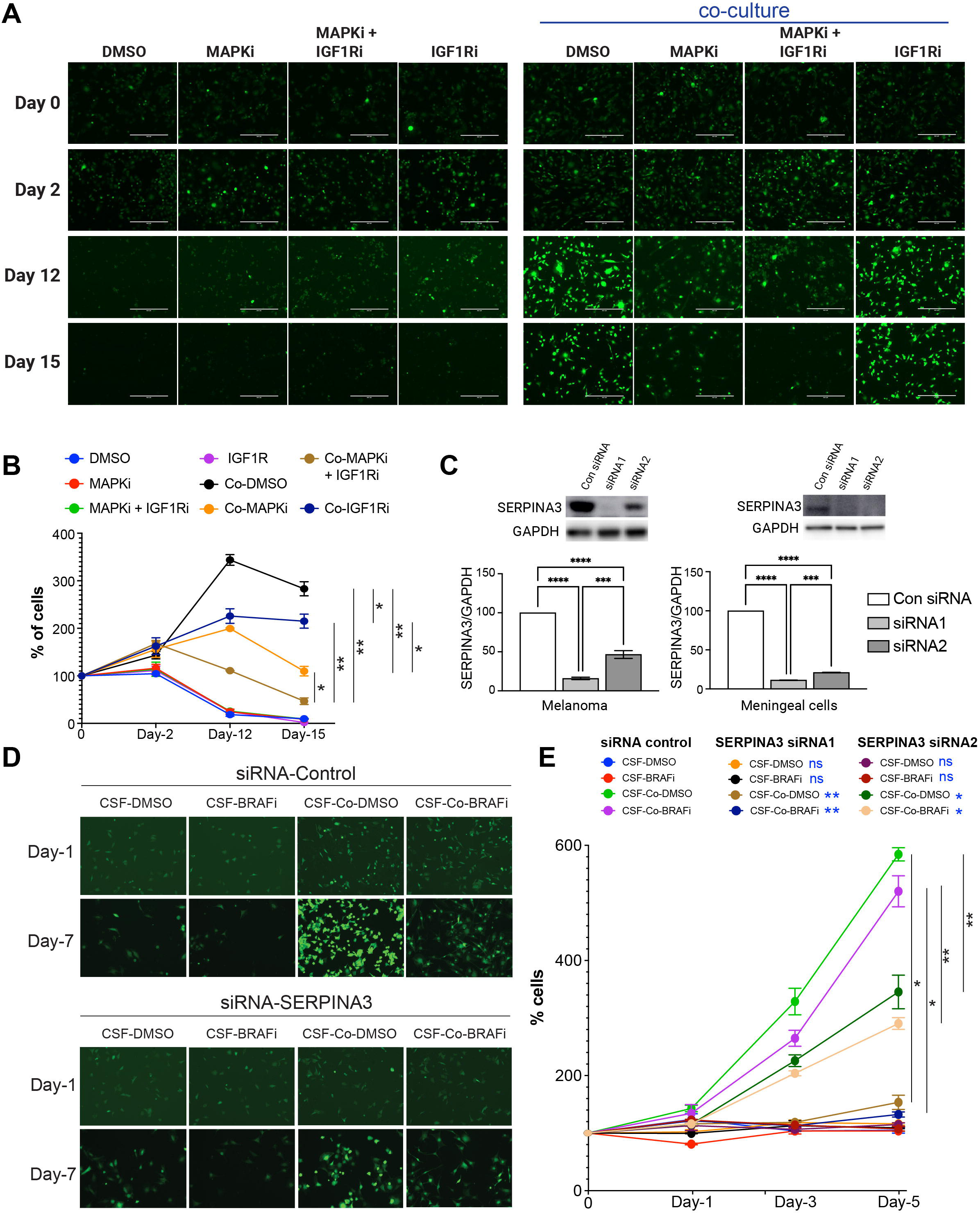
Abrogating SERPINA3/IGF1R signaling sensitized melanoma to MAPKi in the CSF environment. **A.** Representative microscopy images showing GFP-tagged WM164 melanoma cells treated with DMSO control, 100nM dabrafenib/10nM trametinib (MAPKi), 1μM linsitinib (IGF1Ri), or the triple combination in monoculture versus co-culture with primary meningeal cells in the context of CSF. **B.** Quantification of the GFP+ WM164 melanoma cells treated with 3 DMSO control, 100nM dabrafenib/10nM trametinib (MAPKi), 1μM linsitinib (IGF1Ri), or the triple combination in monoculture versus co-culture with primary meningeal cells for 15 days in CSF. **C.** Western blots showing knockdown of SERPINA3 with two individual SERPINA3 siRNAs (siRNA1 and siRNA2) in the WM164 melanoma cells and the HMC meningeal cells. Scrambled sequence siRNA was used as control (Con siRNA) **D.** Representative microscopy images showing GFP-tagged WM164 melanoma cells treated with 3μM vemurafenib (BRAFi) or DMSO (Control) in monoculture versus co-culture (Co) with primary meningeal cells in the context of CSF following knockdown of SERPINA3 (siRNA 1 and siRNA2) or Control siRNA (Con siRNA) in both cell types. **E.** Quantification of GFP-tagged WM164 cells from panel D. Statistical significance was assessed using Student’s t-tests (panels B, C and E). Significance is denoted as *p < 0.05, **p < 0.01, ***p < 0.001, ****p < 0.0001, ns = not significant

## DISCUSSION

Although first described a century and a half ago^28^, little progress has been made in our understanding of the biology, aggressive nature, and vulnerabilities of leptomeningeal disease^1^. Most recently, technological advancements such as scRNAseq have finally allowed us to glimpse the biology of this difficult-to-sample site^6,29,30^. Although these methods uncovered valuable information, the tumor microenvironment of the cells suspended in CSF may not reflect the tumor microenvironment of the melanoma tumors adherent to the leptomeningeal tissues and the potential interactions occurring among those elements of the tumor microenvironment. It is also becoming increasingly clear that the spatial context of the tumor microenvironment is critical in understanding the functional heterogeneity of the tumor and the tumor’s interactions within the LMD ecosystem. We had previously observed that the CSF of patients with melanoma LMD harbored much fewer immune infiltrates, but the analysis of the CSF component did not provide any insights into the stromal compartment of the tumor microenvironment. Here, we utilized spatial transcriptomics technologies and applied novel analyses^31^ to show that the melanoma tumors at the leptomeninges demonstrate a greater interaction with the stroma than those at extra-cranial sites of disease. We confirm that gene expression involved in pathways that are integral for melanoma growth and drug resistance, including MAPK and phosphatidylinositol signaling^20,32–36^, and processes that we have previously identified in CSF fluid to be important for melanoma LMD, such as TGFβ signaling and complement and coagulation^5,37^, are not distributed randomly throughout the tissue but are rather correlated to the proximity of the tumor to the stroma. Instead of relying on a more arbitrary determination of the tumor/stroma interface as has been done in other spatial transcriptomics studies^38–41^, we have developed a novel algorithm for calculating the correlation of gene expression to the distance between spots containing tumor and stromal cells, allowing the data to ascertain if areas of high expression occur at the interface is between the tumor and stromal cells.

The initial impression of the leptomeningeal environment is that it should be inhospitable to rapidly growing tumor cells. Typically, CSF contains few nutrients or proteins, suggesting few growth factors and metabolites to sustain tumor survival and growth. However, the rapid disease progression of melanoma LMD in most patients suggests otherwise. The stromal component of the leptomeningeal tissues is very unique, comprising of vasculature and specialized fibroblast-like meningeal cells^8^. Our data shows that these meningeal cells turn on pro-tumorigenic signaling when exposed to melanoma cells in the CSF environment and support tumor survival and drug resistance in this otherwise hostile environment. Many previous studies have demonstrated pro-tumorigenic roles of cancer-associated fibroblasts in melanoma tumors, generally utilizing dermal fibroblast cultures^19,42–44^. In the central nervous system microenvironment, astrocytes have been implicated in aiding melanoma cell metastasis and survival^45–47^. We now demonstrate that the meningeal stroma is required for melanoma cells to survive in the CSF environment through upregulation of SERPINA3 expression and subsequent activation of the PI3K/AKT and MAPK signaling. The role of SERPINA3 in regulating the PI3K/AKT and ERK signaling was initially identified in adipogenesis, where SERPINA3 was shown to modulate IGF1R and Integrin signaling by inhibiting serine proteases^27^. Although a great number of intrinsic and acquired MAPK inhibitor resistance mechanisms have now been identified, many of them converge on either the reactivation of ERK signaling or the activation of compensatory pathways through PI3K/AKT^32^. Here, we show that meningeal cell-induced SERPINA3 expression at the tumor/stroma interface promotes the amplification of PI3K/AKT signaling and the reactivation of ERK in the tumor cells, thereby overcoming BRAF/MEK inhibition.

Most patients with melanoma LMD do not respond to MAPK-targeting therapies such as the BRAF/MEK inhibitor combination^1^. Our data identifies an LMD-specific vulnerability in melanoma cells, which can be exploited therapeutically to improve sensitivity to MAPK-targeting therapies. Some of the major difficulties faced by patients with melanoma LMD are the severe neurological symptoms that negatively impact their quality of life, including pain, nausea, diplopia, weakness, and paralysis^1,2^. These symptoms have primarily been attributed to the development of hydrocephalus and increased intracranial pressure due to the tumor obstructing normal CSF flow^48,49^. However, our spatial analysis of patient specimens shows increased MT3 expression and expression of the reactive gliosis marker GFAP at the tumor/stroma interface. Previously, GFAP and MT3 have been linked to trauma, ischemia, and neurodegenerative diseases, including encephalomyelitis, multiple sclerosis, Parkinson’s disease, Alexander disease, Alzheimer’s disease, and Amyotrophic lateral sclerosis^21,23,50–52^. Our data provides evidence supporting a direct tumor-mediated reactive gliosis and possible neuronal damage occurring at the cellular level. This data is in line with the observations that interventions relieving hydrocephalus, such as ventroperitoneal or lumbar shunting, can result in a reduction of some neurological symptoms like headache and nausea but may have no effect on other symptoms like cognitive dysfunction or gait dysfunction^48^.

One limitation of our work is that we initially profiled relatively few tissues from few patients, and it is not entirely clear if all melanoma LMD tumors will exhibit upregulation of SERPINA3. However, the rarity of this patient population (5-7% of metastatic melanoma patients), the sizable gaps in our understanding of melanoma biology at the leptomeninges, and the preclinical *in vitro* and *in vivo* validation of our findings highlight the unparallel value of this work. Furthermore, several small molecule and antibody-based inhibitors of IGF1R and the PI3K/AKT/mTOR pathway are currently being tested in the clinic^53–56^, highlighting the potential translatability of combining these drugs with BRAF/MEK inhibitor therapy for the treatment of melanoma LMD.

## METHODS

### Patient specimens and Spatial transcriptomics

This study was conducted in accordance with recognized ethical guidelines (e.g., Declaration of Helsinki, CIOMS, Belmont Report, U.S. Common Rule). Twelve human melanoma specimens from five patients were procured under protocols approved by the Institutional Review Board (MCC 21044 and MCC 20779). Frozen tissue specimens were processed according to the Visium Spatial Gene Expression User Guide using reagents from the Visium Spatial Gene Expression Kit (10X Genomics, Pleasanton, CA).

### Spatial transcriptomics analysis

Sequence .fastq files were generated using the default settings with *spaceranger mkfastq* from base call intensities outputs from the two Illumina NextSeq 2000 runs. Then, the reads were passed to *spaceranger count* along with H&E .tif images from each sample. We used the automatic spots to images in *spaceranger count*. In cases where the tissues featured holes or tears and noise background counts were retrieved from those areas without tissue, we used 10X Loupe v4 to align the spots to the tissue images manually and passed the resulting .json file to *spaceranger count* via the *--loupe-alignment* option. The resulting sparse read count matrices and associated spot coordinates were used in downstream analyses. All spatial RNAseq datasets will be uploaded to GEO and made publicly available upon publication. All code utilized for analysis is uploaded to GitHub and will be made publicly available upon publication.

### Cell typing and spatial deconvolution

To infer the cell type composition within each spot, we used Latent Dirichlet Allocation (LDA) as implemented in the R package STdeconvolve ^57^. The algorithm detects latent topics in the gene expression data. The latent topics represent transcriptional profiles of cell types potentially present in the spots. To infer the number of topics (*K*) present in each ST sample, we ran the algorithm from *K*=3 to *K*=15. Identification of the *K* value with the best fit to the data was done by examining the LDA model’s perplexity and the number of topics with low representation in the spots. We then selected the *K* value corresponding to the model with the lowest perplexity and the lowest number of low-abundance topics (**Supplemental Figure 12**). To assign a biological identity to the topics, we applied gene set enrichment analysis (GSEA) on each topic gene expression (*annotateCellTypesGSEA* function). The cell type gene markers identifying cell types were extracted from the ENCODE and BluePrint databases ^58,59^ using the R package celldex ^60^. Manual curation of topic biological identities was done based on examination of the genes for each cell type compared to the average expression across all cell types. Genes with log-fold > 1 for a given cell type were selected as markers. We manually annotated cell types by examining the log-fold gene expression of each topic of the top 15 genes with the highest log-fold change and the bottom 15 genes with the lowest log-fold change in each topic, compared to the average expression of those same genes across all other topics. The changes made using the manual curation procedure are presented in the supplemental materials (**Supplemental Table 6**).

### Spatial pathway enrichment analysis

We aimed to identify gene sets that showed evidence of spatial expression “hotspots.” We modified the *STenrich* function from the R package spatialGE^61,62^ to test for hotspots in the average expression of KEGG pathway genes. The method is a permutation-based adaptation of a previously published method ^63^. The algorithm identifies spots with average log-normalized gene set expression above the average plus one standard deviation across the entire tissue. Next, Euclidean distances among the high-expression spots are calculated, an equal number of spots is randomly chosen from the entire tissue, and Euclidean distances among the randomly selected spots are calculated. The random selection procedure was conducted 1000 times. Finally, the sum of distances among the high expression spots was compared against the 1000 sums of distances from the randomly selected spots (null distribution). If the sum of distances among high expression spots was higher than the null distribution, it indicated that the spots expressing the pathway are closer to each other than expected by chance (i.e., spots are aggregated in a hotspot).

### Gene expression gradients at tissue niche interfaces

The spatial patterns of gene expression at the interface of two niches are often studied by performing differential gene expression analysis on clusters that lie between the niches. Here, we opted for an alternative approach to study gradients of gene expression, hence overcoming the necessity of defining an interface cluster. In this approach, we first collapsed the biological identities of each spot to the most abundant cell type (topic/cell type proportion ≥ 50%). Then, further collapsing was done by classifying spots within tumor, immune, and stromal compartments, with the latter defined as any spot not classified as a tumor or immune. Next, we calculated the Euclidean distances of each spot to a reference niche (e.g., stroma compartment). For each spot, the average distance to the reference niche was calculated. Spearman’s correlations are calculated between the average distance to the reference and log-transformed normalized gene expression. The p-values associated with the Spearman coefficients were used to ascertain whether a given gene exhibited an expression gradient.

### Cell culture

The identities of all cell lines were confirmed through short tandem repeat validation analysis, and all cell cultures were routinely tested for mycoplasma contamination. Melanoma cell lines were maintained in 5% FBS/RPMI-1640, and primary meningeal cells were maintained in a Meningeal Cell Medium (Sciencell, catalog No.#1401). Assays in this manuscript were performed in either standard medium or cerebrospinal fluid (CSF) obtained from Ecocyte bioscience (LRE-S-LSG-1000-1) supplemented with 1% BSA.

### In vitro cell growth/viability analysis

Human melanoma (WM164) and mouse melanoma (SM1) cell lines were plated on cell culture plates as a monolayer or in co-culture with human meningeal cells (HMC) or primary mice meningeal cells (MMC). After 24 hours, all plates were treated with either BRAF inhibitor (vemurafenib 3 μM monotherapy or trametinib 10nM plus dabrafenib 100nM combination), BRAF inhibitor plus insulin growth factor 1 inhibitor (linsitinib 1μM) or vehicle (DMSO). The number of the GFP-tagged cells was quantified using ImageJ software.

### Conditioned medium and CSF

Conditioned media was generated by adding 10 ml of meningeal medium or CSF to a co-culture of WM164 plus HMC or SM1 plus primary mice meningeal cells in 10-cm culture plates. After 24 hours of incubation, the medium/CSF was removed and centrifuged at 25000 rcf for 5 minutes at room temperature to remove any cell debris. The medium was filtered through a 0.22-micron filter before mixing with fresh meningeal medium or fresh CSF at a ratio of 1:1.

### MTT assays

MTT assays were carried out as described previously^64^. Briefly, one thousand WM164 cells were cultured in each well of a 96-well plate in either fresh or conditioned CSF. The next day, the cells were exposed to different concentrations of vemurafenib (0 to 30 mM). The plate was incubated at 37°C and 5% CO^2^ for three days. On day 4, 20uL of MTT reagent was added to each well and incubated for 3 hours, then CSF was removed, and MTT crystals were solubilized in 50μL of DMSO. Absorbance was measured at 490 nm.

### Cell cycle assays

Cell cycle analysis was carried out as described previously^64^. Cells were plated in a 10-cm culture plate and allowed to grow to 70% confluency. Cells were harvested and washed with 2 ml of phosphate-buffered saline (PBS). The cell pellet was re-suspended with 2 mL ice-cold 70% ethanol and fixed overnight. The next day, the cells were rehydrated with 20 mL PBS for 30 minutes at room temperature and then centrifuged and washed with PBS. Cells were stained with propidium iodide. Data was acquired using the FACSCanto (BD) instrument and analyzed using ModFit software.

### Western blots

Western blot analysis was carried out as described previously^20^. Protein extracts were immunoblotted using antibodies against SERPINA3 (1:1000, Genetex), phospho-ERK (Thr202/Tyr204, 1:1000, Cell Signaling Technology), ERK(1:1000, Cell Signaling Technology), phospho-Akt (1:1000, Cell Signaling Technology), Akt (Ser473, 1:1000, Cell Signaling Technology), CyclinD1 (1:1000, Cell Signaling Technology), TGF β1(1:1000, Cell Signaling Technology), Fibronectin (1:1000, BD) and GAPDH (1:5000, Sigma Aldrich). Digitalized blot images were developed using Amersham Imager 600 (Amersham Bioscience). Quantification of the protein expression was performed using ImageJ software.

### siRNA knockdowns

Electroporation-assisted siRNA knockdown of SERPINA3 (or scrambled control) was carried out using the Neon transfection system 100 uL kit following the manufacturer’s protocol (Invitrogen). Electroporation was performed at a pulse voltage of 1200. Cells were then seeded in 10 cm cell culture plates containing pre-warmed medium. SERPINA3 knockdown was confirmed by Western Blot.

### Animals

All mouse work was conducted in accordance with recognized ethical guidelines and IACUC approval. SCID female mice were used to evaluate the effect of HMC on the development of melanoma tumors. Mice were injected subcutaneously with 1 million WM164 cells alone, 1 million WM164 cells with 1 million primary meningeal cells (MC), or 1 million WM164 cells with 1 million dermal fibroblast (DF) cells. Once the tumors were palpable (on day 17), animals were started on either a control rodent diet or a rodent diet formulated with 200 mg/kg dabrafenib and 2 mg/kg trametinib (MAPKi) as previously described (Research Diets)^65^.

### Statistical analysis

Western blot data was analyzed by GraphPad Prism using ordinary one-way ANOVA followed by Tukey’s multiple comparison test. Cell counting was analyzed using an unpaired parametric student’s T-test. P-values were considered significant if less than 0.05. Significance is annotated on figures as follows: *p < 0.05, **p < 0.01, ***p < 0.001, ****p < 0.0001.

## ACKNOWLEDGEMENTS

We thank the patients and their families for their valuable and generous contributions to this study. This work was generously supported by the Tara Miller Melanoma Foundation Young Investigator Award from the Melanoma Research Alliance (932727 to IS) and a T32 Training Grant (T32CA23339 to BF and WDC). The work in the Smalley lab is also supported by a Research Scholar Grant from the American Cancer Society (RSG-23-1040487-01-MM), an R00 from the National Institutes of Health (R00 CA226679), and an R21 from the National Institutes of Health (R21 CA274060). This work has been supported in part by the Molecular Genomics, Flow Cytometry, Microscopy, and Bioinformatics and Biostatistics Shared Resources at the H. Lee Moffitt Cancer Center & Research Institute, an NCI designated Comprehensive Cancer Center (P30-CA076292).

## REFERENCES CITED

1 Glitza, I. C. et al. Leptomeningeal disease in melanoma patients: An update to treatment, challenges, and future directions. Pigment Cell Melanoma Res (2020). 10.1111/pcmr.12861

2 Khaled, M. L., Tarhini, A. A., Forsyth, P. A., Smalley, I. & Piña, Y. Leptomeningeal Disease (LMD) in Patients with Melanoma Metastases. Cancers (Basel) 15 (2023). 10.3390/cancers15061884

3 Smalley, K. S., Fedorenko, I. V., Kenchappa, R. S., Sahebjam, S. & Forsyth, P. A. Managing leptomeningeal melanoma metastases in the era of immune and targeted therapy. Int J Cancer 139, 1195–1201 (2016). 10.1002/ijc.30147

4 Ferguson, S. D. et al. Predictors of survival in metastatic melanoma patients with leptomeningeal disease (LMD). J Neurooncol 142, 499–509 (2019). 10.1007/s11060-019-03121-2

5 Smalley, I. et al. Proteomic Analysis of CSF from Patients with Leptomeningeal Melanoma Metastases Identifies Signatures Associated with Disease Progression and Therapeutic Resistance. Clin Cancer Res 26, 2163–2175 (2020). 10.1158/1078-0432.Ccr-19-2840

6 Smalley, I. et al. Single-Cell Characterization of the Immune Microenvironment of Melanoma Brain and Leptomeningeal Metastases. Clin Cancer Res 27, 4109–4125 (2021). 10.1158/1078-0432.Ccr-21-1694

7 Remsik, J. et al. Characterization, isolation, and in vitro culture of leptomeningeal fibroblasts. J Neuroimmunol 361, 577727 (2021). 10.1016/j.jneuroim.2021.577727

8 Adeeb, N. et al. The pia mater: a comprehensive review of literature. Child’s Nervous System 29, 1803–1810 (2013). 10.1007/s00381-013-2044-5

9 Remsik, J. et al. Characterization, isolation, and in vitro culture of leptomeningeal fibroblasts. Journal of Neuroimmunology 361, 577727 (2021). 10.1016/j.jneuroim.2021.577727

10 Choe, Y. & Pleasure, S. J. Meningeal Bmps regulate cortical layer formation. Brain plasticity 4, 169–183 (2018).

11 Pinho-Ribeiro, F. A. et al. Bacteria hijack a meningeal neuroimmune axis to facilitate brain invasion. Nature 615, 472–481 (2023).

12 Rua, R. & McGavern, D. B. Advances in Meningeal Immunity. Trends in Molecular Medicine 24, 542–559 (2018). 10.1016/j.molmed.2018.04.003

13. Fishman, R. (Philadelphia press, 1992).

14 Chi, M., Ye, Y., Zhang, X. D. & Chen, J. Insulin induces drug resistance in melanoma through activation of the PI3K/Akt pathway. Drug Des Devel Ther 8, 255–262 (2014). 10.2147/dddt.S53568

15 Das, I. et al. Inhibiting insulin and mTOR signaling by afatinib and crizotinib combination fosters broad cytotoxic effects in cutaneous malignant melanoma. Cell Death & Disease 11, 882 (2020). 10.1038/s41419-020-03097-2

16 Pimiento, J. M. et al. Melanoma genotypes and phenotypes get personal. Laboratory Investigation 93, 858–867 (2013). 10.1038/labinvest.2013.84

17 Smalley, I. & Smalley, K. ERK inhibition: A new front in the war against MAPK pathway-driven cancers? Cancer Discov. 2018; 8: 140–142. doi: 10.1158/2159-8290.(CD-17-1355.[Abstract][CrossRef][Google Scholar]).

18 Truzzi, F. et al. Neurotrophins and their receptors stimulate melanoma cell proliferation and migration. J Invest Dermatol 128, 2031–2040 (2008). 10.1038/jid.2008.21

19 Fedorenko, I. V., Wargo, J. A., Flaherty, K. T., Messina, J. L. & Smalley, K. S. BRAF Inhibition Generates a Host-Tumor Niche that Mediates Therapeutic Escape. J Invest Dermatol 135, 3115–3124 (2015). 10.1038/jid.2015.329

20 Fedorenko, I. V. et al. Fibronectin induction abrogates the BRAF inhibitor response of BRAF V600E/PTEN-null melanoma cells. Oncogene 35, 1225–1235 (2016). 10.1038/onc.2015.188

21 Middeldorp, J. & Hol, E. M. GFAP in health and disease. Progress in Neurobiology 93, 421–443 (2011). 10.1016/j.pneurobio.2011.01.005

22 Smith, M. E. & Eng, L. F. Glial fibrillary acidic protein in chronic relapsing experimental allergic encephalomyelitis in SJL/J mice. J Neurosci Res 18, 203–208 (1987). 10.1002/jnr.490180129

23 Juárez-Rebollar, D., Rios, C., Nava-Ruíz, C. & Méndez-Armenta, M. Metallothionein in Brain Disorders. Oxid Med Cell Longev 2017, 5828056 (2017). 10.1155/2017/5828056

24 Taylor, J. W. et al. Primary leptomeningeal lymphoma: International Primary CNS Lymphoma Collaborative Group report. Neurology 81, 1690–1696 (2013). 10.1212/01.wnl.0000435302.02895.f3

25 Nolan, C. P. & Abrey, L. E. Leptomeningeal metastases from leukemias and lymphomas. Cancer Treat Res 125, 53–69 (2005).

26 Leal, T., Chang, J. E., Mehta, M. & Robins, H. I. Leptomeningeal Metastasis: Challenges in Diagnosis and Treatment. Curr Cancer Ther Rev 7, 319–327 (2011). 10.2174/157339411797642597

27 Choi, Y. et al. Serpina3c Regulates Adipogenesis by Modulating Insulin Growth Factor 1 and Integrin Signaling. iScience 23, 100961 (2020). 10.1016/j.isci.2020.100961

28 Eberth, C. Zur entwickelung des epithelioms (cholesteatoms) der pia und der lunge. Archiv für pathologische Anatomie und Physiologie und für klinische Medicin 49, 51–63 (1869).

29 Prakadan, S. M. et al. Genomic and transcriptomic correlates of immunotherapy response within the tumor microenvironment of leptomeningeal metastases. Nat Commun 12, 5955 (2021). 10.1038/s41467-021-25860-5

30 Chi, Y. et al. Cancer cells deploy lipocalin-2 to collect limiting iron in leptomeningeal metastasis. Science 369, 276–282 (2020). 10.1126/science.aaz2193

31 Ospina, O. E. et al. spatialGE: quantification and visualization of the tumor microenvironment heterogeneity using spatial transcriptomics. Bioinformatics 38, 2645–2647 (2022). 10.1093/bioinformatics/btac145

32 Fedorenko, I. V., Paraiso, K. H. & Smalley, K. S. Acquired and intrinsic BRAF inhibitor resistance in BRAF V600E mutant melanoma. Biochem Pharmacol 82, 201–209 (2011). 10.1016/j.bcp.2011.05.015

33 Wu, H., Goel, V. & Haluska, F. G. PTEN signaling pathways in melanoma. Oncogene 22, 3113–3122 (2003). 10.1038/sj.onc.1206451

34 Cohen, C. et al. Mitogen-actived protein kinase activation is an early event in melanoma progression. Clin Cancer Res 8, 3728–3733 (2002).

35 Inamdar, G. S., Madhunapantula, S. V. & Robertson, G. P. Targeting the MAPK pathway in melanoma: why some approaches succeed and other fail. Biochem Pharmacol 80, 624–637 (2010). 10.1016/j.bcp.2010.04.029

36 Davies, M. A. The role of the PI3K-AKT pathway in melanoma. Cancer J 18, 142–147 (2012). 10.1097/PPO.0b013e31824d448c

37 Boire, A. et al. Complement Component 3 Adapts the Cerebrospinal Fluid for Leptomeningeal Metastasis. Cell 168, 1101–1113 e1113 (2017). 10.1016/j.cell.2017.02.025

38 Hunter, M. V., Moncada, R., Weiss, J. M., Yanai, I. & White, R. M. Spatially resolved transcriptomics reveals the architecture of the tumor-microenvironment interface. Nature Communications 12, 6278 (2021). 10.1038/s41467-021-26614-z

39 Peng, Z., Ye, M., Ding, H., Feng, Z. & Hu, K. Spatial transcriptomics atlas reveals the crosstalk between cancer-associated fibroblasts and tumor microenvironment components in colorectal cancer. Journal of Translational Medicine 20, 302 (2022). 10.1186/s12967-022-03510-8

40 Moncada, R. et al. Integrating microarray-based spatial transcriptomics and single-cell RNA-seq reveals tissue architecture in pancreatic ductal adenocarcinomas. Nat Biotechnol 38, 333–342 (2020). 10.1038/s41587-019-0392-8

41 Andersson, A. et al. Spatial deconvolution of HER2-positive breast cancer delineates tumor-associated cell type interactions. Nature Communications 12, 6012 (2021). 10.1038/s41467-021-26271-2

42 Balsamo, M. et al. Melanoma-associated fibroblasts modulate NK cell phenotype and antitumor cytotoxicity. Proc Natl Acad Sci U S A 106, 20847–20852 (2009). 10.1073/pnas.0906481106

43 Li, G. et al. Function and regulation of melanoma-stromal fibroblast interactions: when seeds meet soil. Oncogene 22, 3162–3171 (2003). 10.1038/sj.onc.1206455

44 Straussman, R. et al. Tumour micro-environment elicits innate resistance to RAF inhibitors through HGF secretion. Nature 487, 500–504 (2012). 10.1038/nature11183

45 Marchetti, D., Li, J. & Shen, R. Astrocytes contribute to the brain-metastatic specificity of melanoma cells by producing heparanase. Cancer Res 60, 4767–4770 (2000).

46 Klein, A. et al. Astrocytes facilitate melanoma brain metastasis via secretion of IL-23. J Pathol 236, 116–127 (2015). 10.1002/path.4509

47 Niessner, H. et al. Targeting hyperactivation of the AKT survival pathway to overcome therapy resistance of melanoma brain metastases. Cancer medicine 2, 76–85 (2013). 10.1002/cam4.50

48 Lamba, N., Fick, T., Nandoe Tewarie, R. & Broekman, M. L. Management of hydrocephalus in patients with leptomeningeal metastases: an ethical approach to decision-making. Journal of Neuro-Oncology 140, 5–13 (2018). 10.1007/s11060-018-2949-7

49 Kim, H. S. et al. Clinical outcome of cerebrospinal fluid shunts in patients with leptomeningeal carcinomatosis. World J Surg Oncol 17, 59 (2019). 10.1186/s12957-019-1595-7

50 Chatterjee, P. et al. Plasma glial fibrillary acidic protein is elevated in cognitively normal older adults at risk of Alzheimer’s disease. Translational Psychiatry 11, 27 (2021). 10.1038/s41398-020-01137-1

51 Messing, A. et al. Fatal encephalopathy with astrocyte inclusions in GFAP transgenic mice. Am J Pathol 152, 391–398 (1998).

52 Damier, P., Hirsch, E. C., Zhang, P., Agid, Y. & Javoy-Agid, F. Glutathione peroxidase, glial cells and Parkinson’s disease. Neuroscience 52, 1–6 (1993). 10.1016/0306-4522(93)90175-f

53 Werner, H., Sarfstein, R. & Bruchim, I. Investigational IGF1R inhibitors in early stage clinical trials for cancer therapy. Expert Opin Investig Drugs 28, 1101–1112 (2019). 10.1080/13543784.2019.1694660

54 Peng, Y., Wang, Y., Zhou, C., Mei, W. & Zeng, C. PI3K/Akt/mTOR Pathway and Its Role in Cancer Therapeutics: Are We Making Headway? Front Oncol 12, 819128 (2022). 10.3389/fonc.2022.819128

55 Yang, J. et al. Targeting PI3K in cancer: mechanisms and advances in clinical trials. Molecular Cancer 18, 26 (2019). 10.1186/s12943-019-0954-x

56 Dienstmann, R., Rodon, J., Serra, V. & Tabernero, J. Picking the Point of Inhibition: A Comparative Review of PI3K/AKT/mTOR Pathway Inhibitors. Molecular Cancer Therapeutics 13, 1021–1031 (2014). 10.1158/1535-7163.Mct-13-0639

57 Miller, B. F., Huang, F., Atta, L., Sahoo, A. & Fan, J. Reference-free cell type deconvolution of multi-cellular pixel-resolution spatially resolved transcriptomics data. Nat Commun 13, 2339 (2022). 10.1038/s41467-022-30033-z

58 Consortium, E. P. An integrated encyclopedia of DNA elements in the human genome. Nature 489, 57–74 (2012). 10.1038/nature11247

59 Martens, J. H. & Stunnenberg, H. G. BLUEPRINT: mapping human blood cell epigenomes. Haematologica 98, 1487–1489 (2013). 10.3324/haematol.2013.094243

60 Aran, D. et al. Reference-based analysis of lung single-cell sequencing reveals a transitional profibrotic macrophage. Nat Immunol 20, 163–172 (2019). 10.1038/s41590-018-0276-y

61 spatialGE: An R package for visualization and analysis of spatially-resolved gene expression v. 1.0 (GitHub, 2023).

62 Ospina, O. E. et al. spatialGE: Quantification and visualization of the tumor microenvironment heterogeneity using spatial transcriptomics. Bioinformatics 38, 2645– 2647 (2022). 10.1093/bioinformatics/btac145

63 Hunter, M. V., Moncada, R., Weiss, J. M., Yanai, I. & White, R. M. Spatially resolved transcriptomics reveals the architecture of the tumor-microenvironment interface. Nat Commun 12, 6278 (2021). 10.1038/s41467-021-26614-z

64 Fedorenko, I. V., Fang, B., Koomen, J. M., Gibney, G. T. & Smalley, K. S. Amuvatinib has cytotoxic effects against NRAS-mutant melanoma but not BRAF-mutant melanoma. Melanoma Res 24, 448–453 (2014). 10.1097/cmr.0000000000000103

65. Phadke, M., et al. Targeted therapy given after anti-PD-1 leads to prolonged responses in mouse melanoma models through sustained antitumor immunity. Cancer Immunology Research In Press (2021).

