## Supplemental Figures 1-2 and 4-12 for "Spatial transcriptomics analysis identifies a unique tumor-promoting function of the meningeal stroma in melanoma leptomeningeal disease"

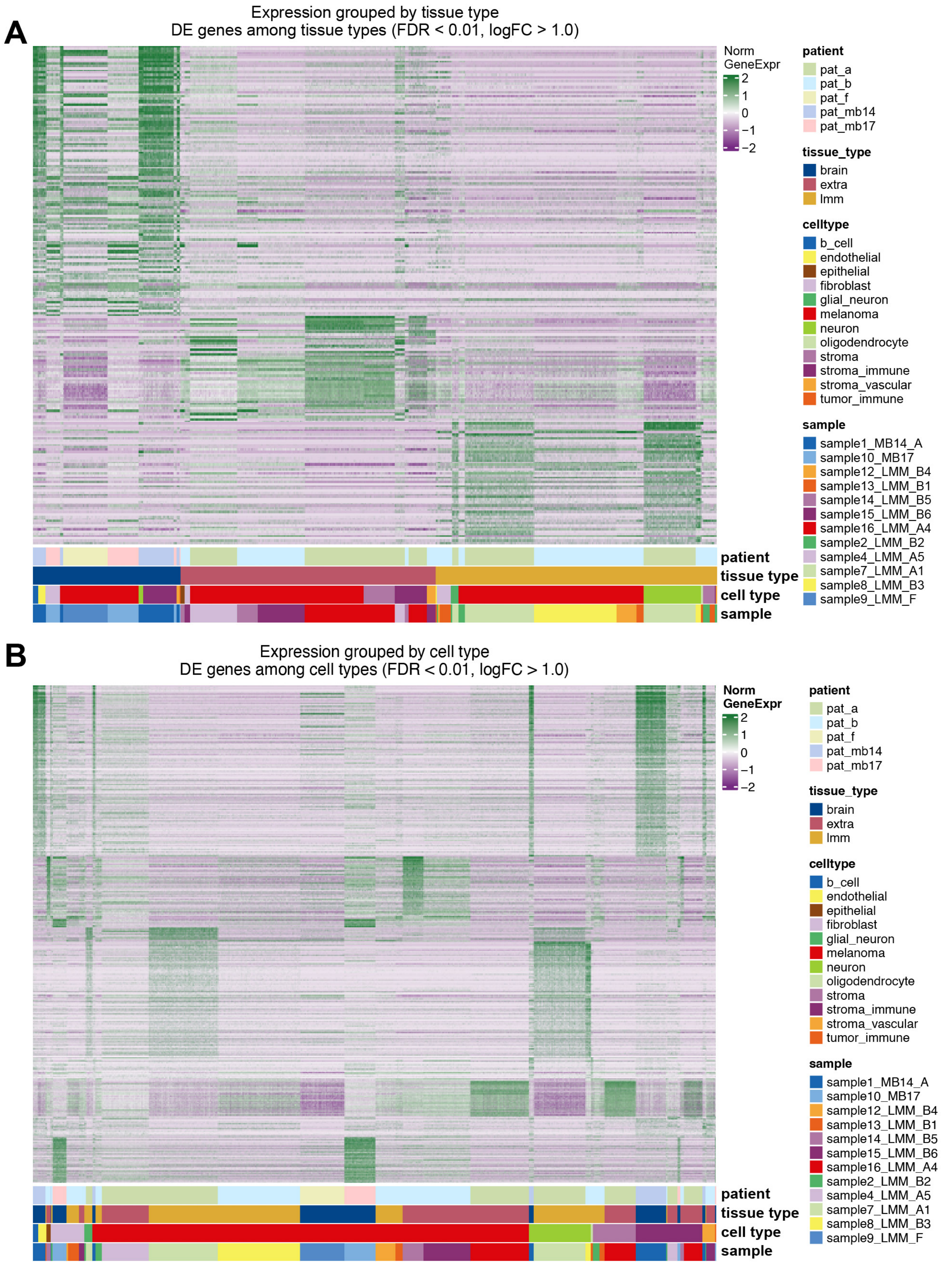


**Supplemental Figure 1. A.** Heatmap of the differentially expressed genes in spots grouped by tissue type **B.** Heatmap of the differentially expressed genes in spots grouped by cell type type.

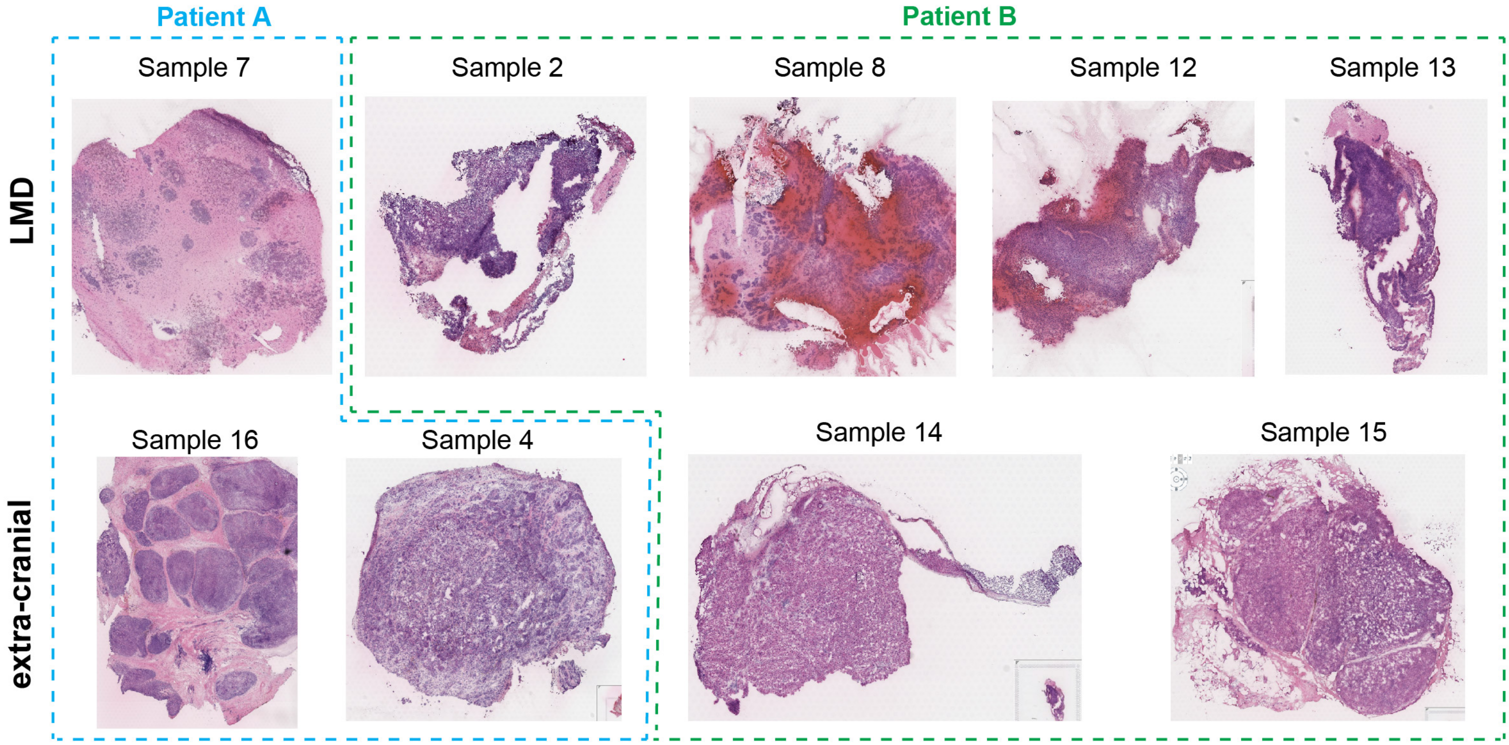


**Supplemental Figure 2.** H & E images for the melanoma leptomeningeal metastasis samples and patient-matched extra-cranial metastasis samples matching the cell type deconvolution in Figure 1B.

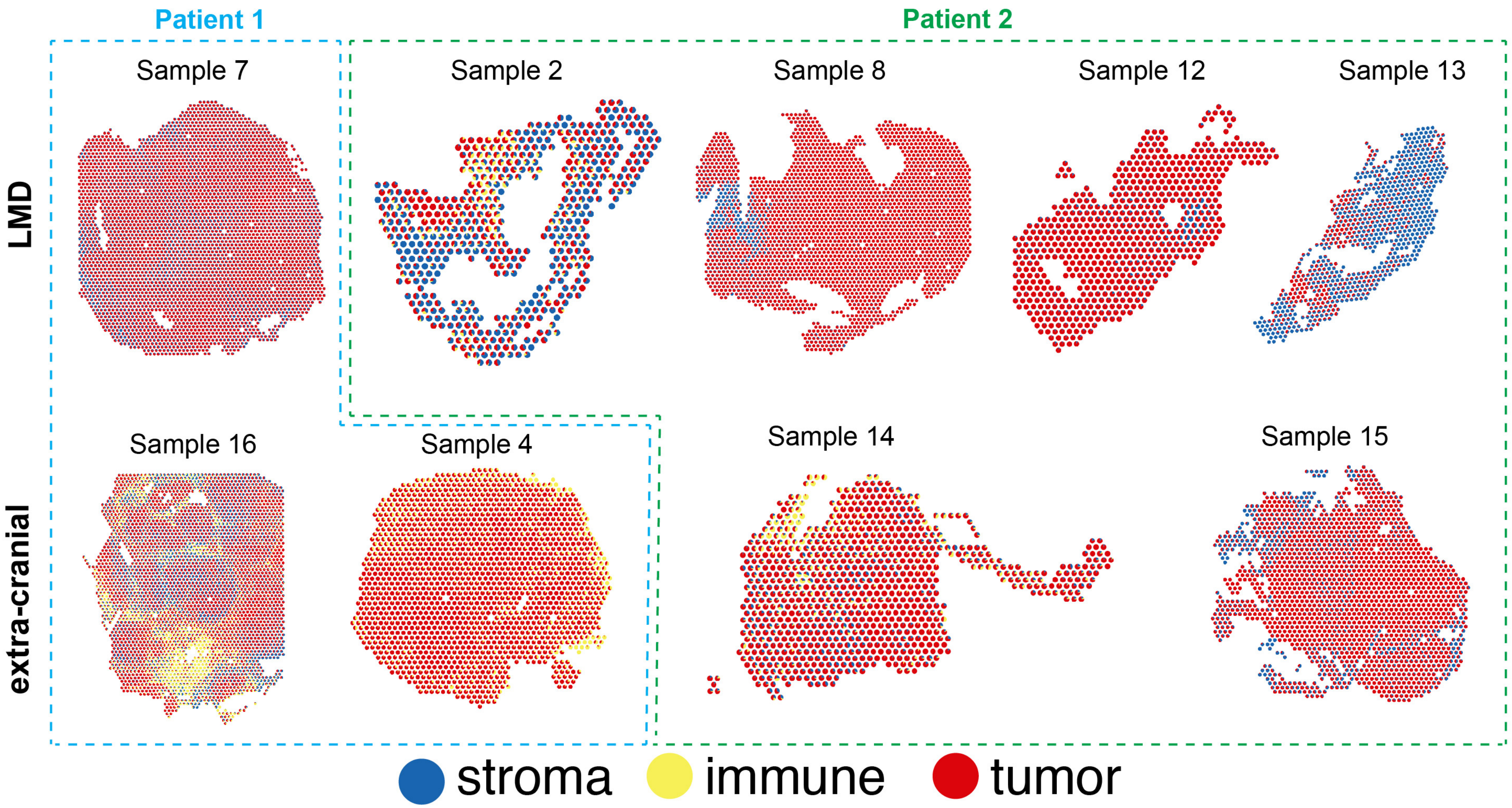


**Supplemental Figure 4.** Spatial tissue maps showing the deconvolution of the stroma, immune, and tumor cell types within each spot. To better visualize regions of the tumor, stroma and immune infiltration on the spatial maps for consecutive visualizations, the cell sub-type categories were condensed into the "tumor", "stroma" and "immune" categories. Each spot was divided into piecharts for the proportion of the stroma, immune or tumor cells present.

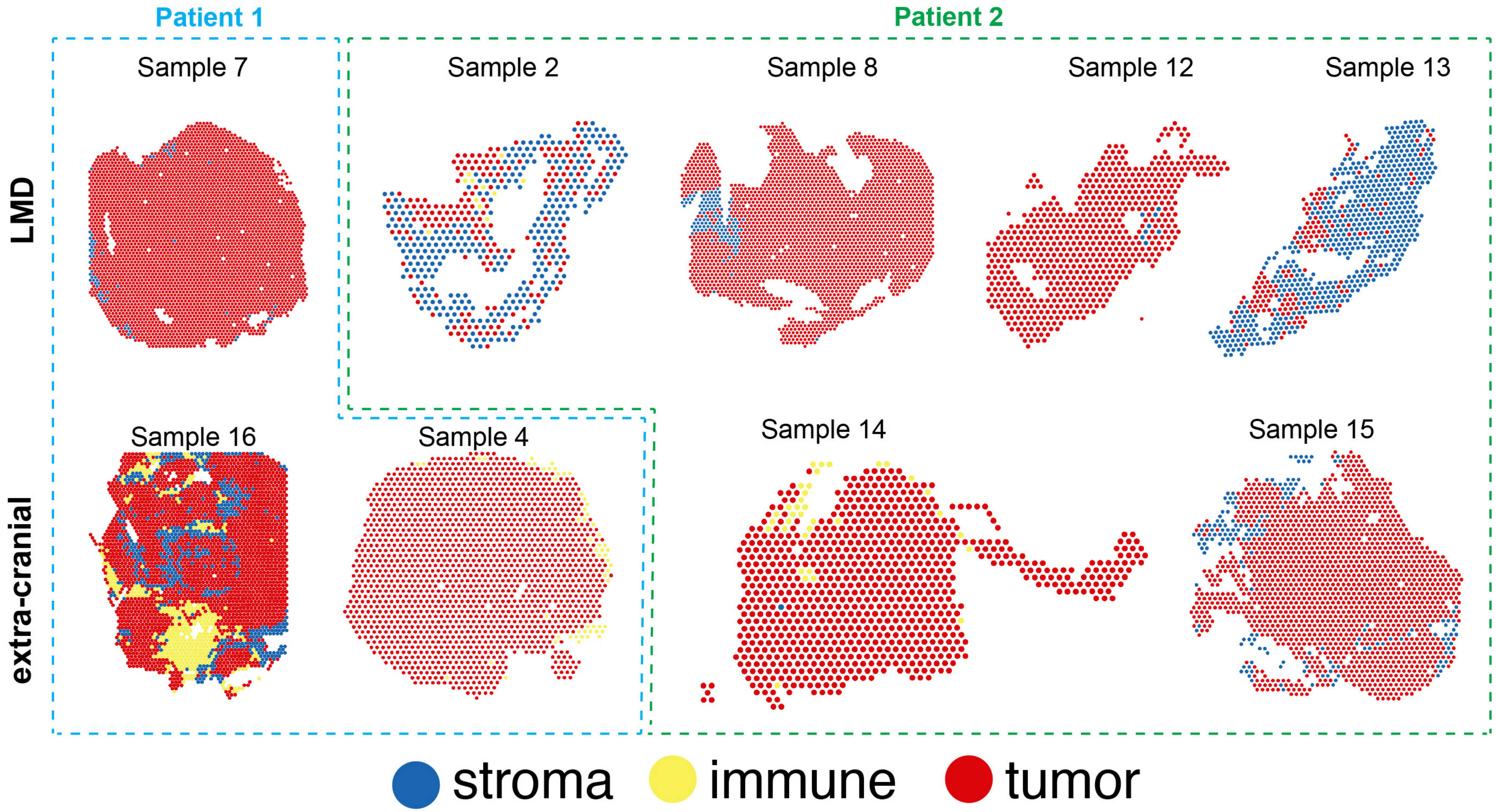


**Supplemental Figure 5.** Spatial tissue maps showing the position of the stroma, immune, and tumor spots. To better visualize regions of the tumor, stroma and immune infiltration on the spatial maps for consecutive visualizations, the cell subtype categories were condensed into the “tumor”, “stroma” and “immune” categories. Each spot was assigned the stroma, immune or tumor category based on the predominant cell type present.

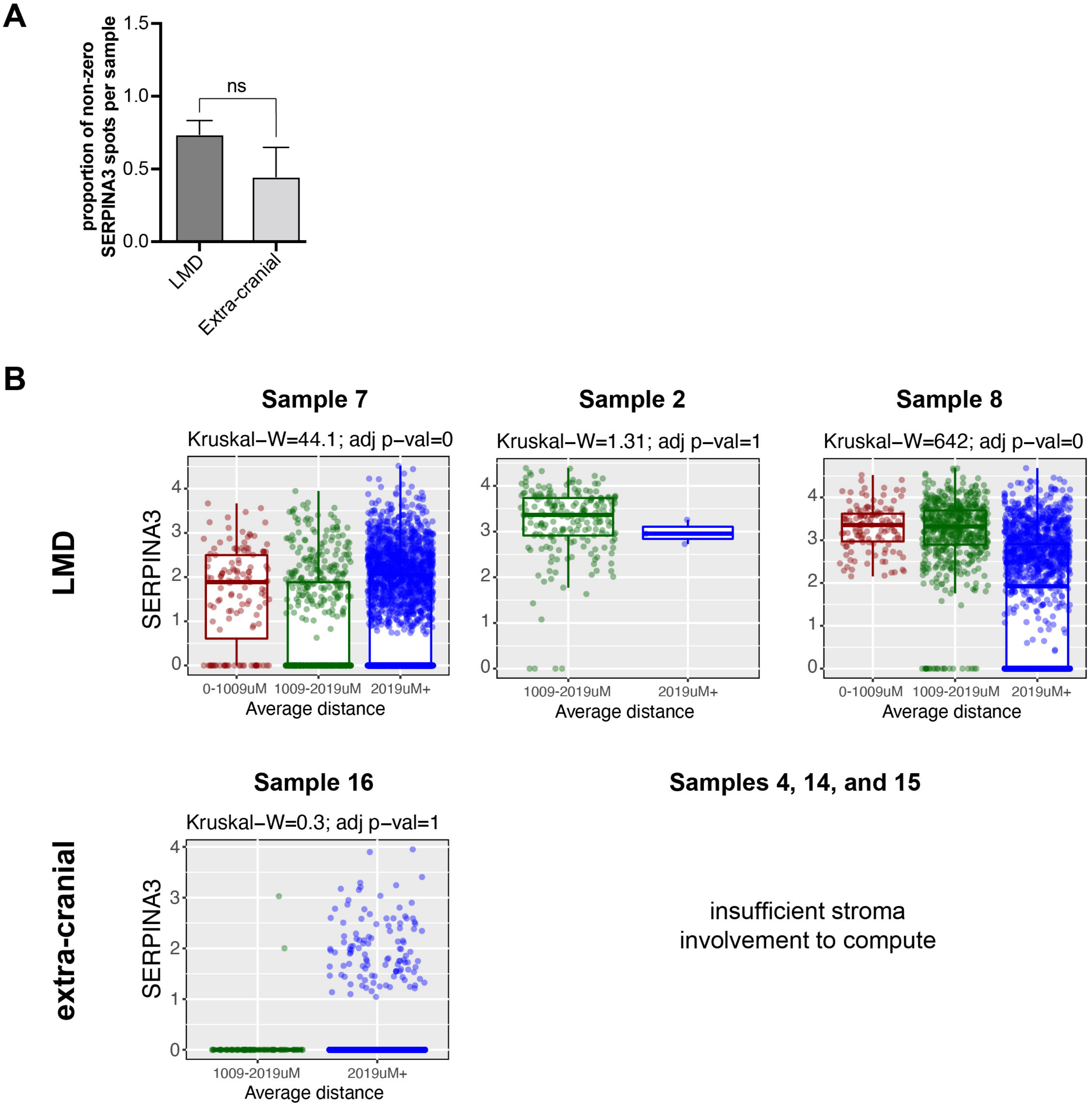


**Supplemental Figure 6. A.** Bar graph showing the proportion of non-zero expression of SERPINA3 spots per sample. **B.** Boxplots showing the normalized expression of SERPINA3 gene in each tumor spot based on the average distance between the tumor spot and stroma spots. No enough stromal involvement was found in extra-cranial tissue from Samples 4, 14 and 15 and therefore it was not possible to calculate expression of SERPINA3 relative to distance of tumor to stroma.

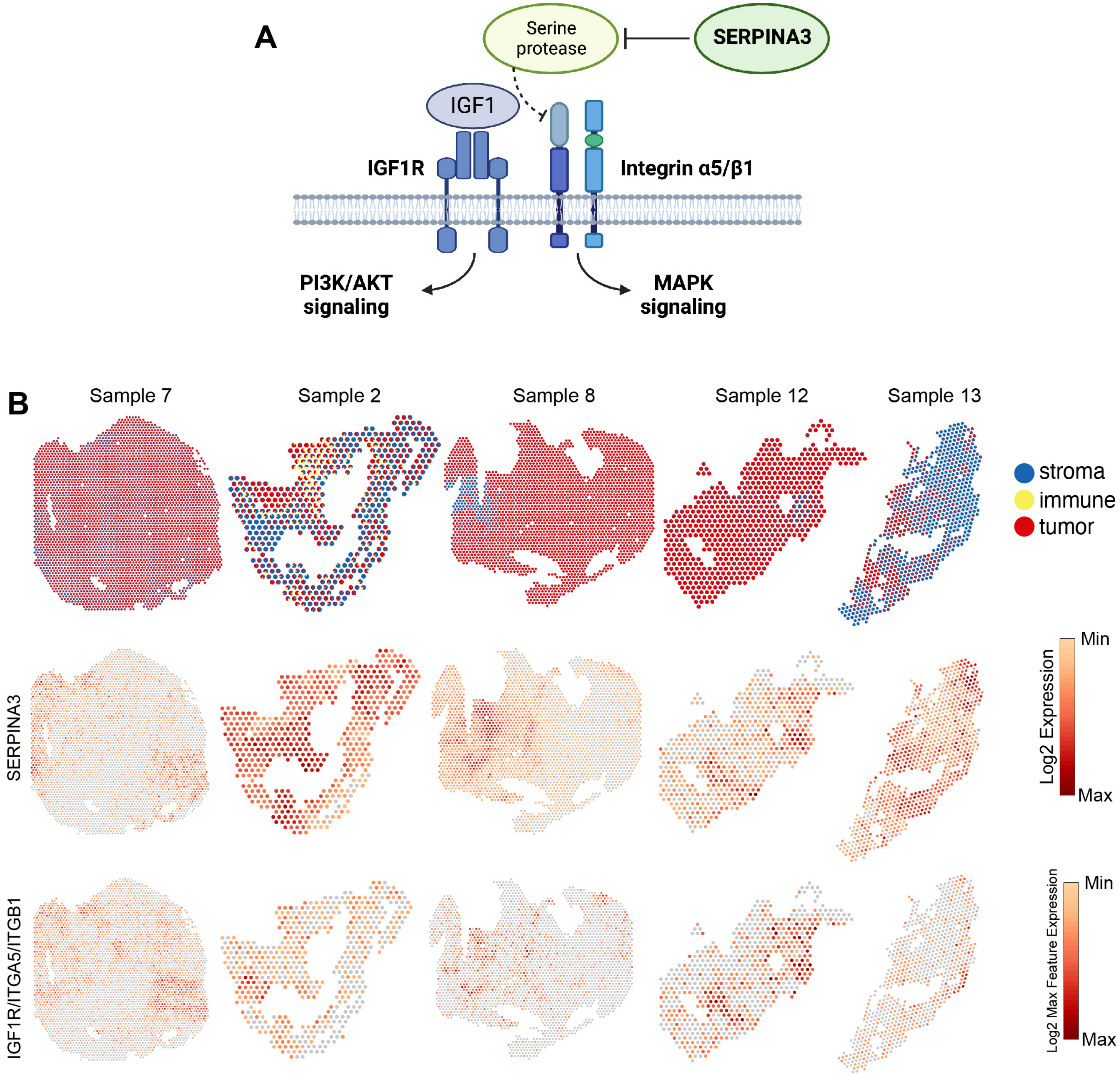


**Supplemental Figure 7. A.** SERPINA3 was previously shown to regulate PI3K and MAPK pathways via activation of the IGF1R/integrinα5β1 signaling. B. Spatial tissue maps showing the position of the stroma, immune, and tumor spots along with tissue maps visualizing the log 2 gene expression for SERPINA3, and the log2 max expression for the IGF1R/ITGA5/ITGB1sets of genes.

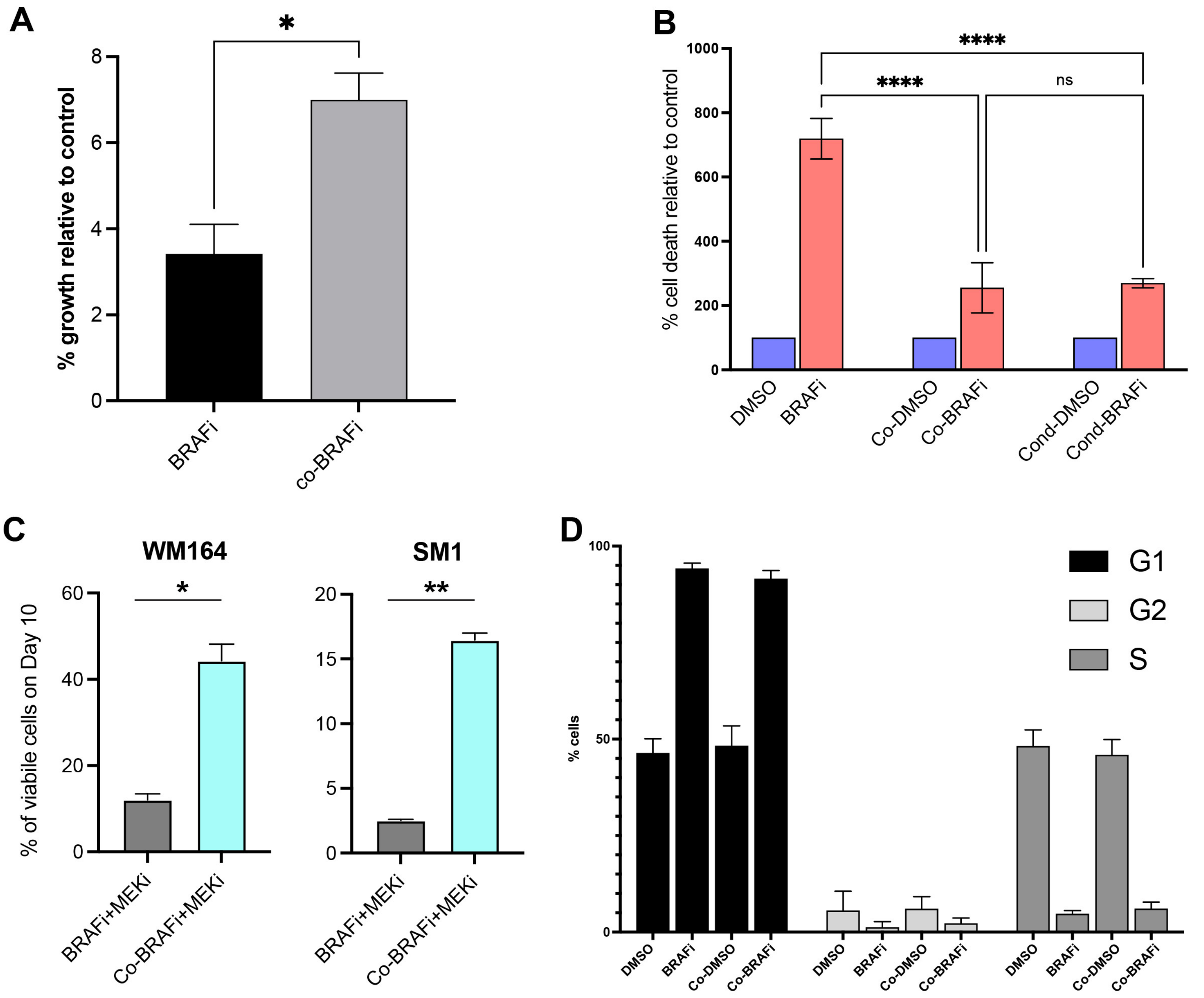


**Supplemental Figure 8. A.** Quantification of % growth following 3μM vemurafenib treatment (BRAFi) in SM1 mono-culture and co-culture with primary murine meningeal cells, relative to day 1 control. **B.** Quantification of viable cells following 3μM vemurafenib treatment (BRAFi) in WM164 mono-culture, co-culture with meningeal cells (Co) and with conditioned media from meningeal cells (Cond). **C.** Bar graphs showing the % of viabile cells left on Day 10 relative to Day 0 for SM1 and WM164 cell lines treated with 100nM dabrafenib + 10nM trametinib. **D.** Flow cytometry assessment of cell cycle using PI staining in WM164 melanoma cells treated with 3μM vemurafenib (BRAFi) or DMSO in the context of monoculture or direct co-culture with primary meningeal cells (Co) under normal media conditions.

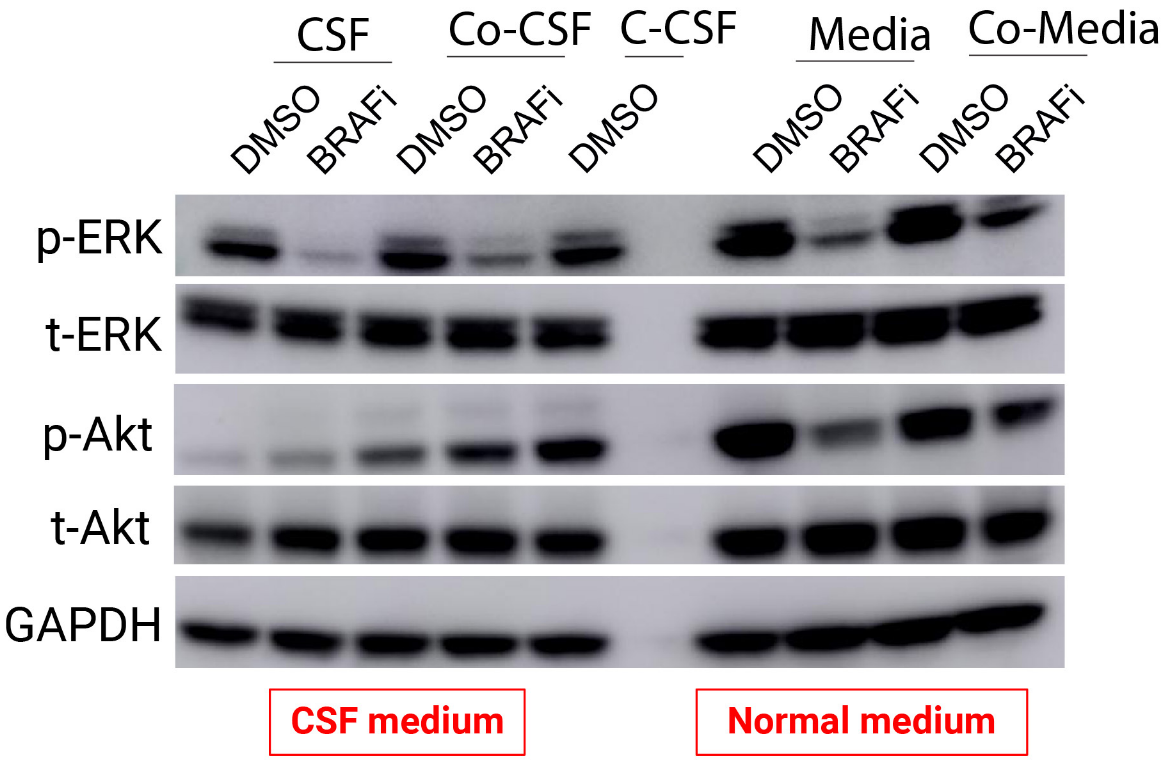


**Supplemental Figure 9.** Western blot analysis of WM164 melanoma cells treated with 3μM vemurafenib or DMSO control in regular or conditioned media or artificial CSF showing abundance of pERK(Thr202/Tyr204), ERK, pAKT (Ser473) and AKT. Co-CSF is conditioned CSF from co-cultures of melanoma and primary meningeal cells, whereas C-CSF is CSF conditioned by primary meningeal cells only.Co-Media is conditioned media from co-cultures of melanoma and primary meningeal cells.

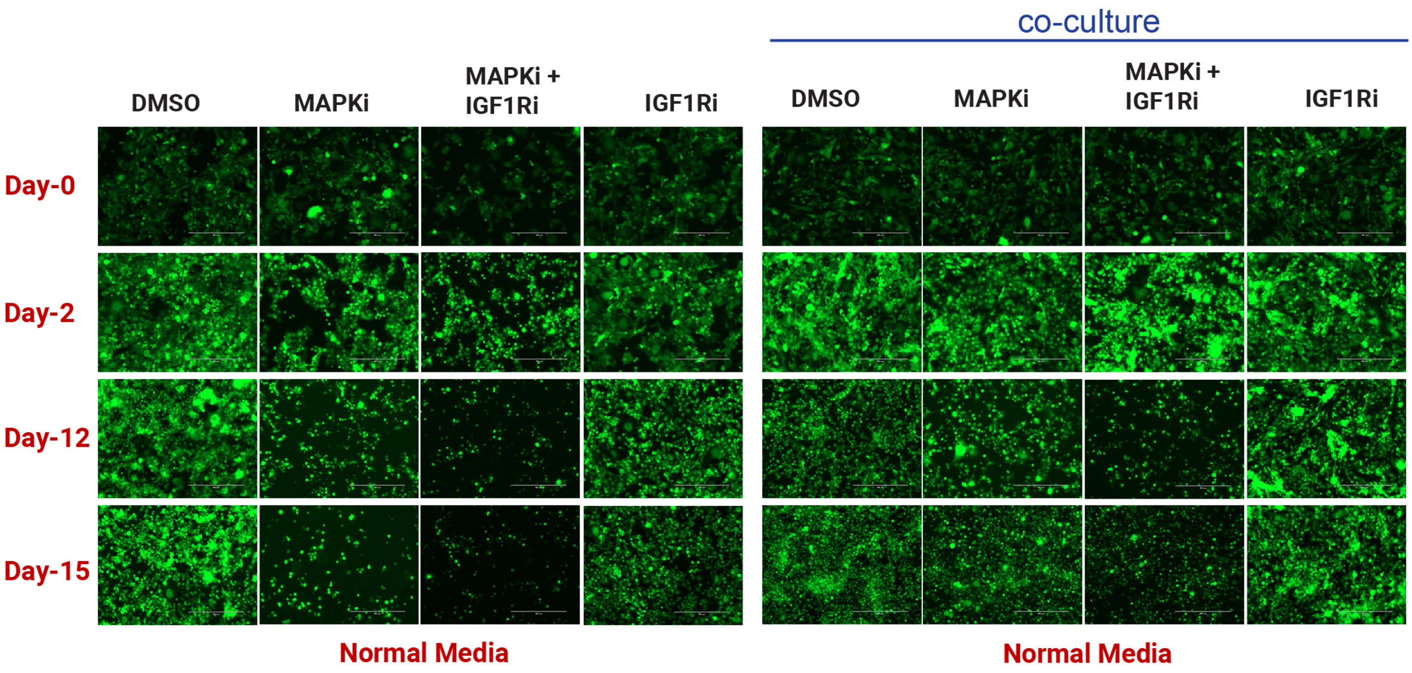


**Supplemental Figure 10.** Representative microscopy images showing GFP-tagged WM164 melanoma cells treated with DMSO control, 100nM dabrafenib/10nM trametinib, 1μM linsitinib, or the triple combination in monoculture versus co-culture with primary meningeal cells in the context of normal media conditions.

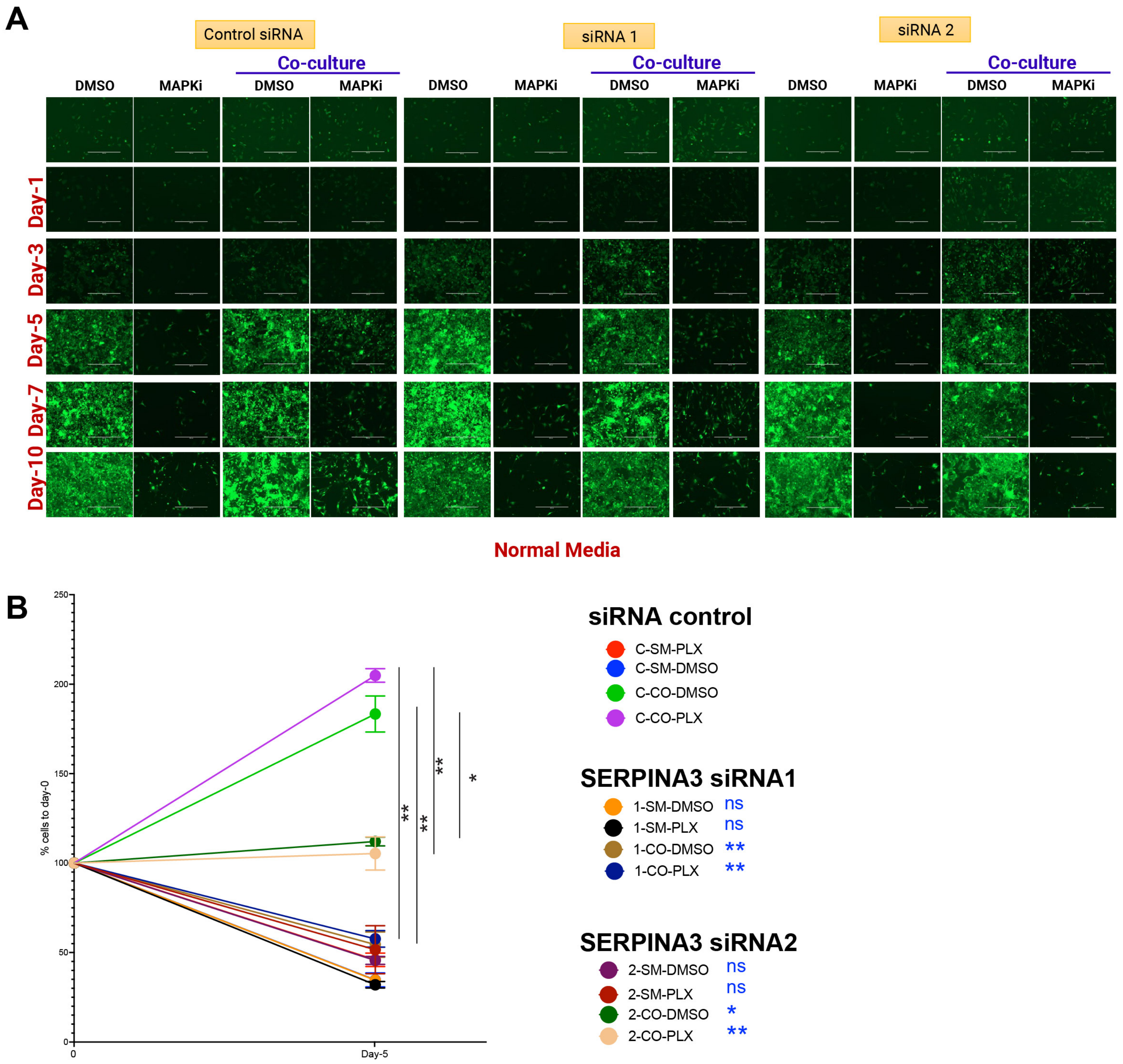


**Supplemental Figure 11. A.** Representative microscopy images showing GFP-tagged WM164 melanoma cells treated with 3μM vemurafenib or DMSO control in monoculture versus co-culture with primary meningeal cells in the context of normal media conditions following knockdown of SERPINA3 or Control siRNA (siRNAc) in both cell types.

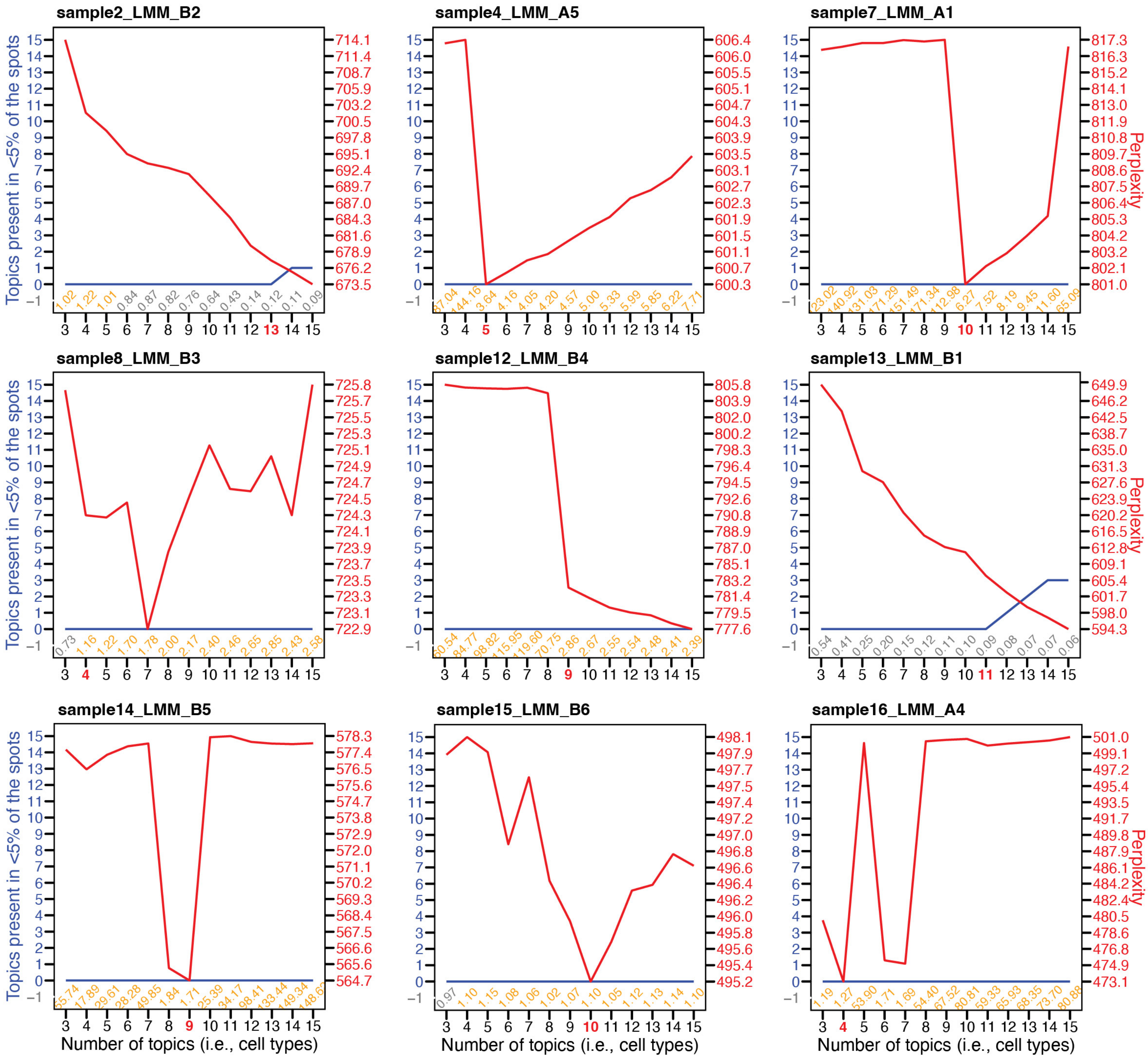


**Supplemental Figure 12.** The perplexity (red line) for a series of LDA models fit to each Visium sample assuming a number of topics ranging from 3 to 15. Models with low perplexity are preferred. To assist our selection of the most likely number of topics, the number of topics scarcely repesented (<5% of spots; blue line) in the model was also examined. A compromise between low perplexity and low number of rare topics was selecte for each sample. The numbers below the curves represent the models' alpha values. Values closer to lower than 1 (gray) are preferred when possible, resulting in each spot having more than one topic. The selected model is shown with a bold red number.
