## Supplemental Figure 3 for "Spatial transcriptomics analysis identifies a unique tumor-promoting function of the meningeal stroma in melanoma leptomeningeal disease"

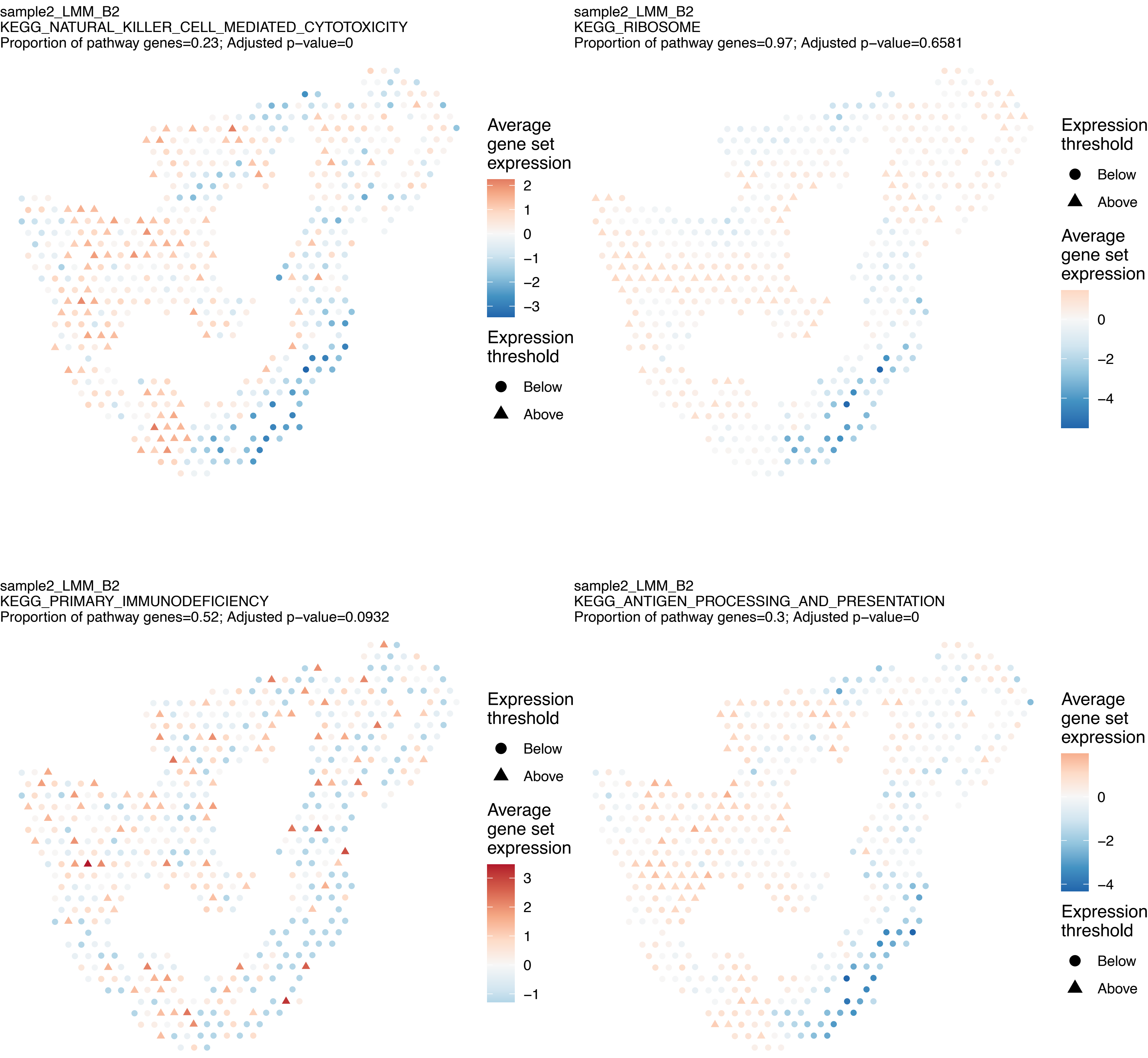
sample2_LMM_B2
KEGG_NATURAL_KILLER_CELL_MEDIATED_CYTOTOXICITY
Proportion of pathway genes=0.23; Adjusted p−value=0
Average gene set expression
2
1
0
−1
−2
−3
Expression threshold
Below
Above
sample2_LMM_B2
KEGG_RIBOSOME
Proportion of pathway genes=0.97; Adjusted p−value=0.6581
Expression threshold
Below
Above
Average gene set expression
0
−2
−4
sample2_LMM_B2
KEGG_PRIMARY_IMMUNODEFICIENCY
Proportion of pathway genes=0.52; Adjusted p−value=0.0932
Expression threshold
Below
Above
Average gene set expression
3
2
1
0
−1
sample2_LMM_B2
KEGG_ANTIGEN_PROCESSING_AND_PRESENTATION
Proportion of pathway genes=0.3; Adjusted p−value=0
Average gene set expression
0
−2
−4
Expression threshold
Below
Above

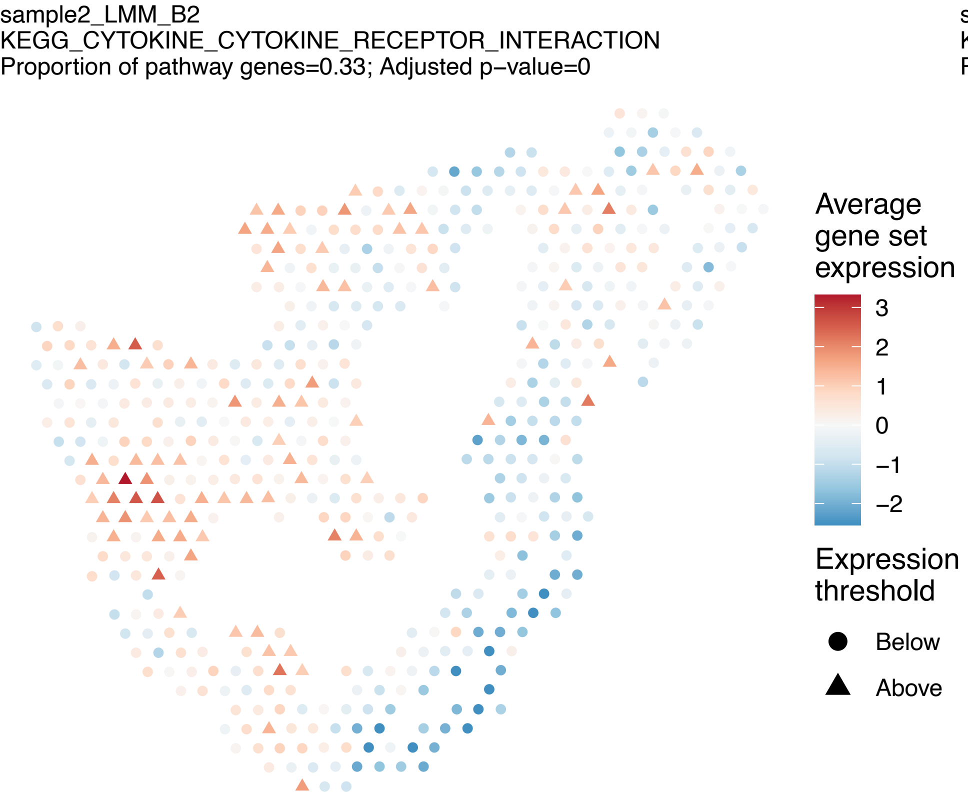
sample2_LMM_B2
KEGG_CYTOKINE_CYTOKINE_RECEPTOR_INTERACTION
Proportion of pathway genes=0.33; Adjusted p−value=0
Average gene set expression
3
2
1
0
−1
−2
Expression threshold
Below
Above

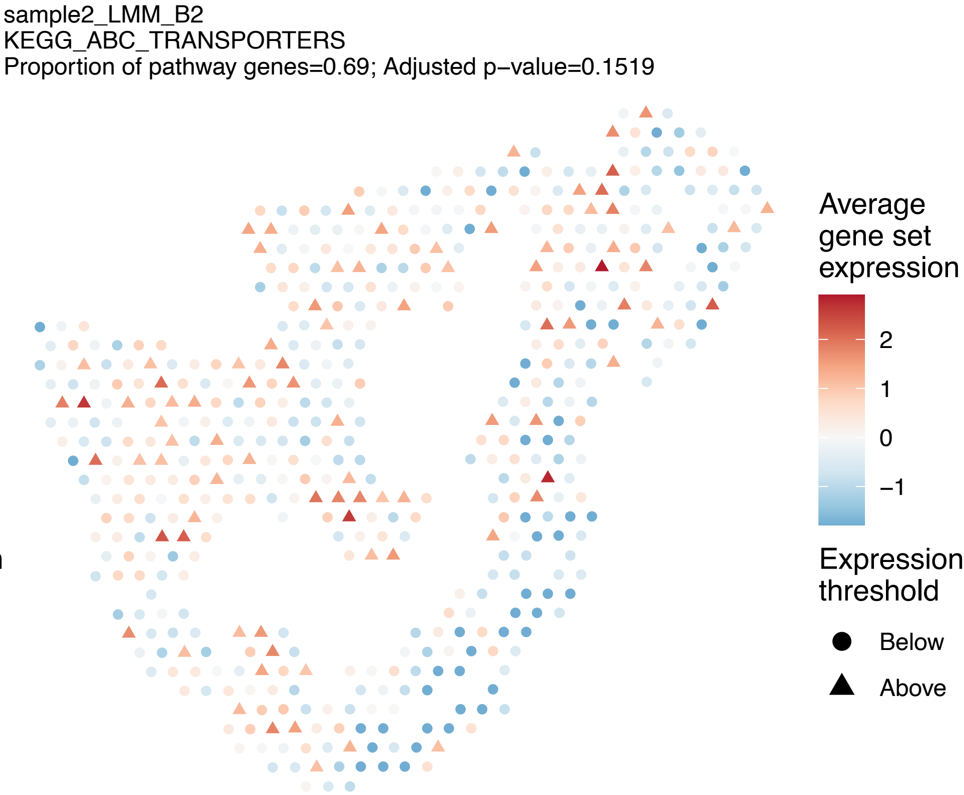
sample2_LMM_B2
KEGG_ABC_TRANSPORTERS
Proportion of pathway genes=0.69; Adjusted p−value=0.1519
Average gene set expression
2
1
0
−1
Expression threshold
Below
Above

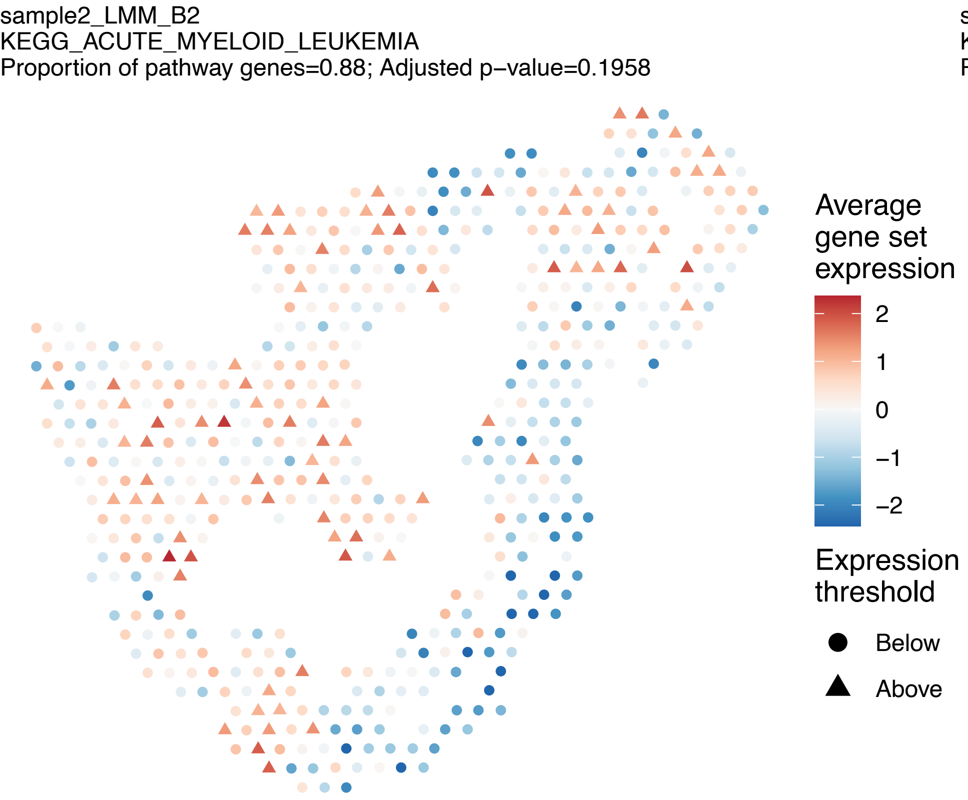
sample2_LMM_B2
KEGG_ACUTE_MYELOID_LEUKEMIA
Proportion of pathway genes=0.88; Adjusted p−value=0.1958
Average gene set expression
2
1
0
−1
−2
Expression threshold
Below
Above

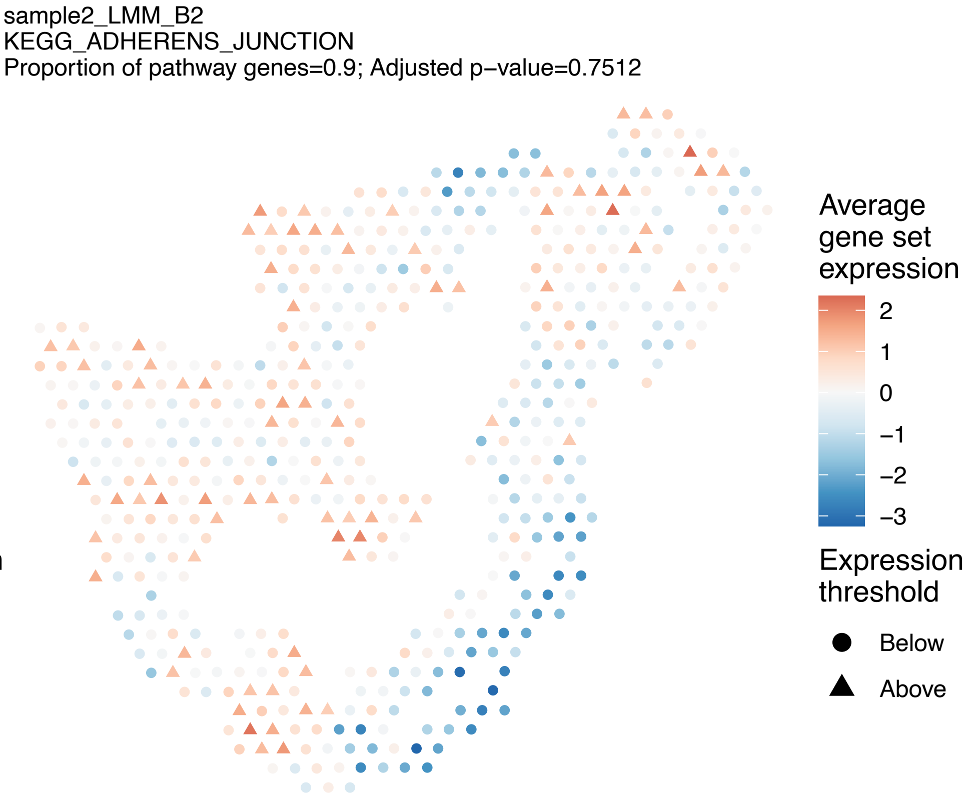
sample2_LMM_B2
KEGG_ADHERENS_JUNCTION
Proportion of pathway genes=0.9; Adjusted p−value=0.7512
Average gene set expression
2
1
0
−1
−2
−3
Expression threshold
Below
Above

sample2_LMM_B2
KEGG_ADIPOCYTOKINE_SIGNALING_PATHWAY
Proportion of pathway genes=0.69; Adjusted p−value=0.1516

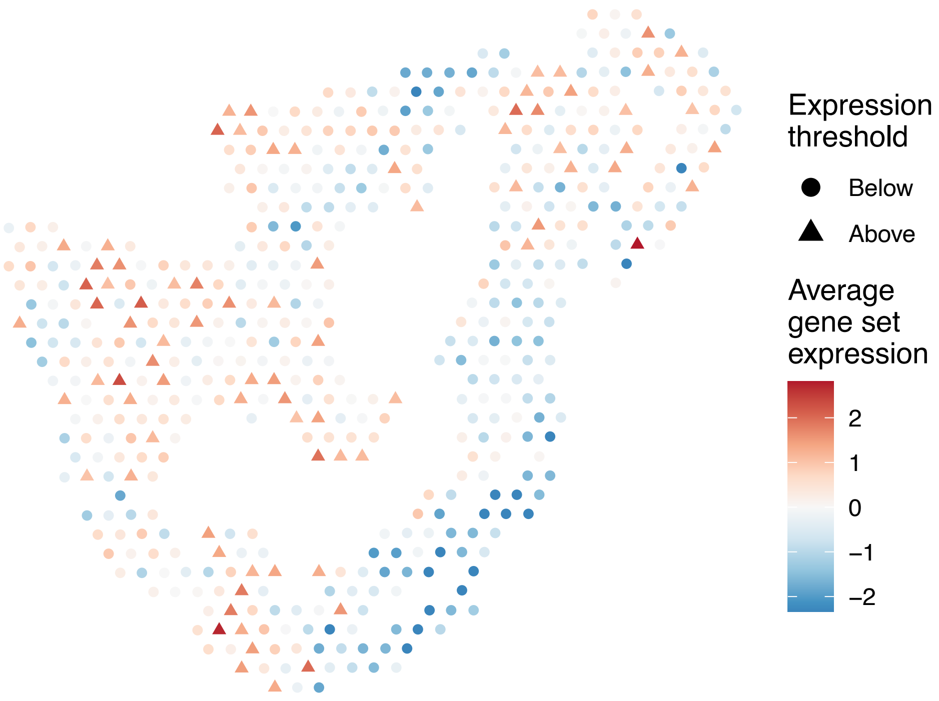


sample2_LMM_B2
KEGG_ALANINE_ASPARTATE_AND_GLUTAMATE_METABOLISM
Proportion of pathway genes=0.64; Adjusted p−value=0.7757

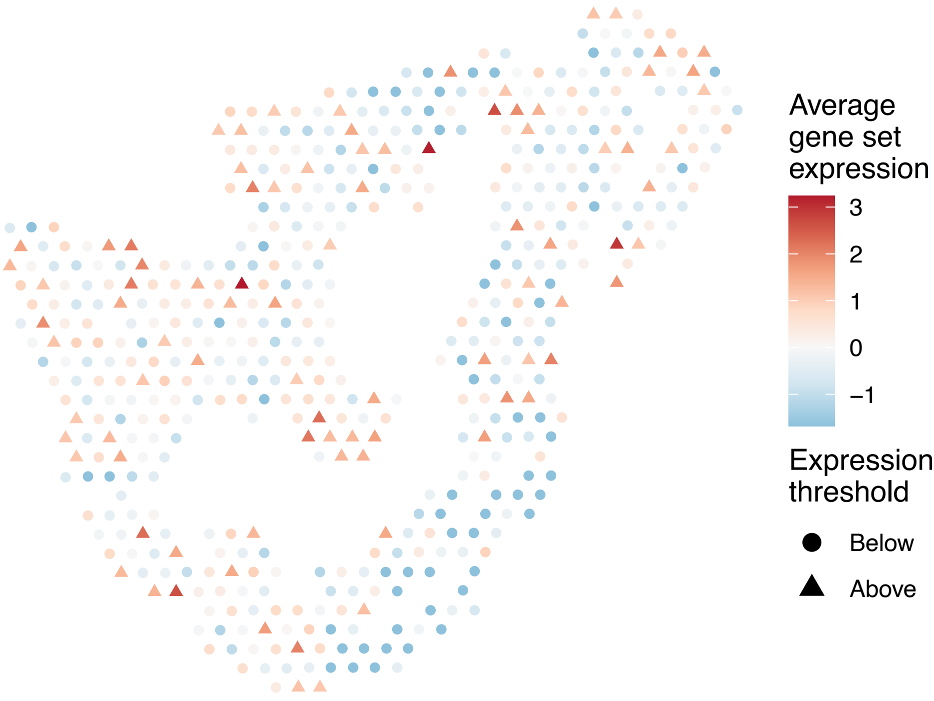


sample2_LMM_B2
KEGG_ALDOSTERONE_REGULATED_SODIUM_REABSORPTION
Proportion of pathway genes=0.5; Adjusted p−value=0.2669

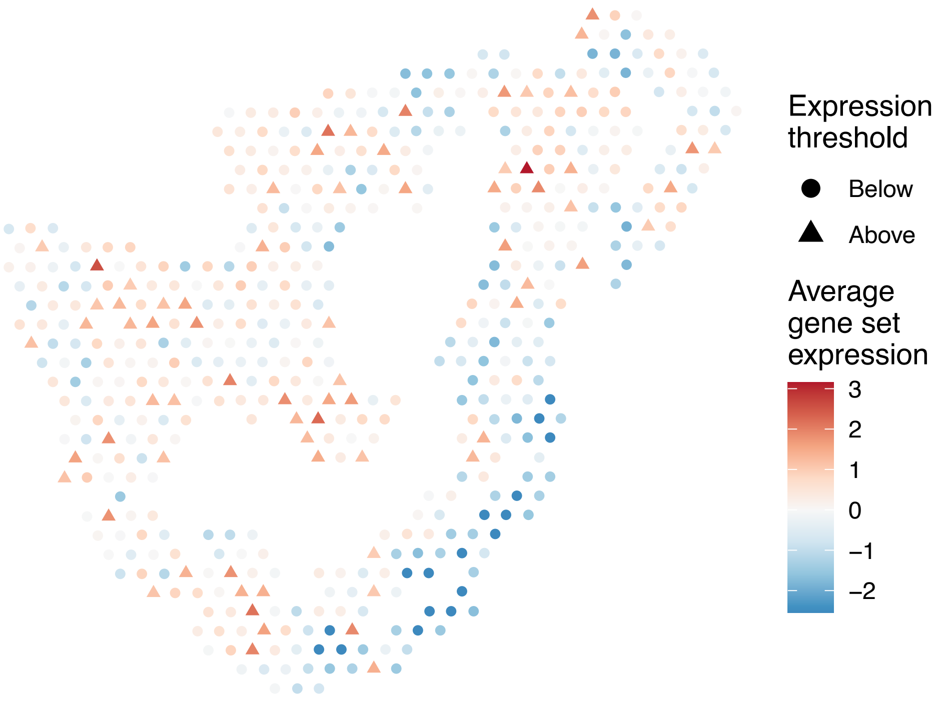


sample2_LMM_B2
KEGG_ALLOGRAFT_REJECTION
Proportion of pathway genes=0.62; Adjusted p−value=0

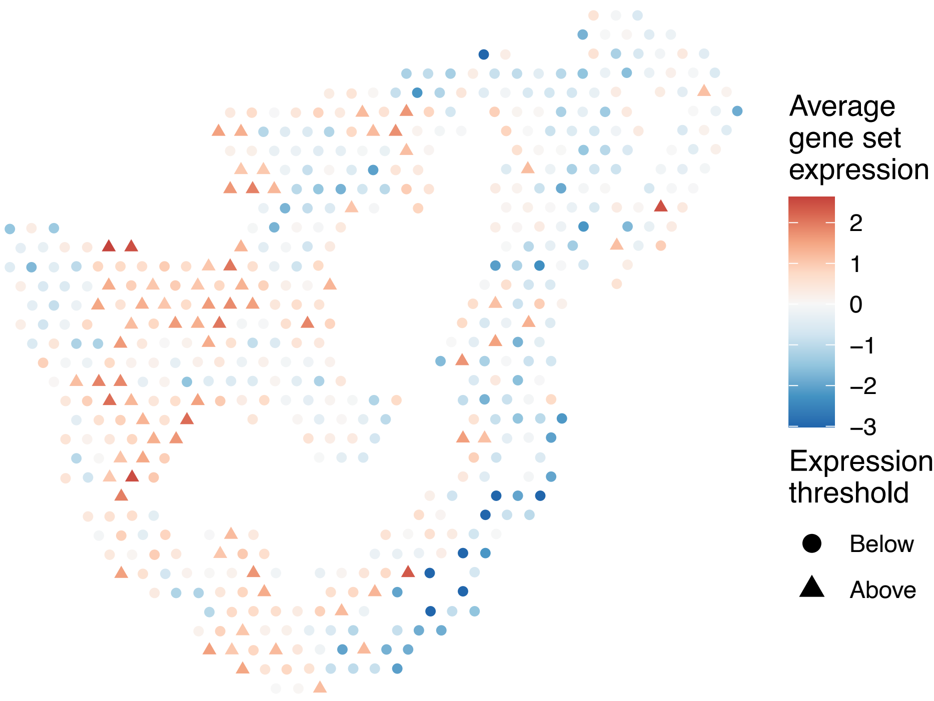

sample2_LMM_B2
KEGG_ALZHEIMERS_DISEASE
Proportion of pathway genes=0.82; Adjusted p−value=0

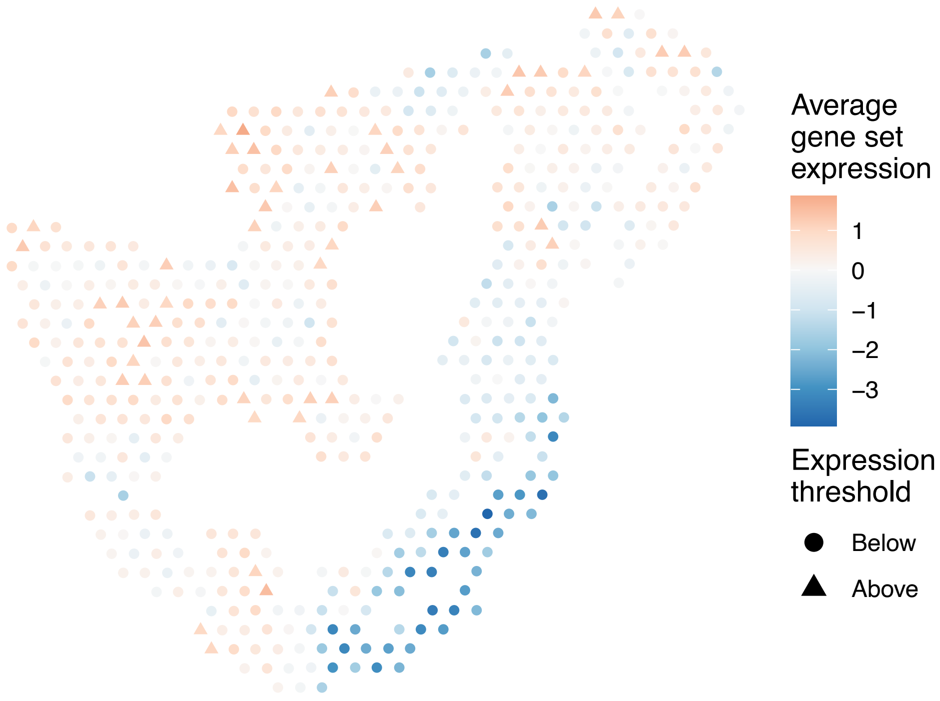


sample2_LMM_B2
KEGG_AMINO_SUGAR_AND_NUCLEOTIDE_SUGAR_METABOLISM
Proportion of pathway genes=0.85; Adjusted p−value=0.3292

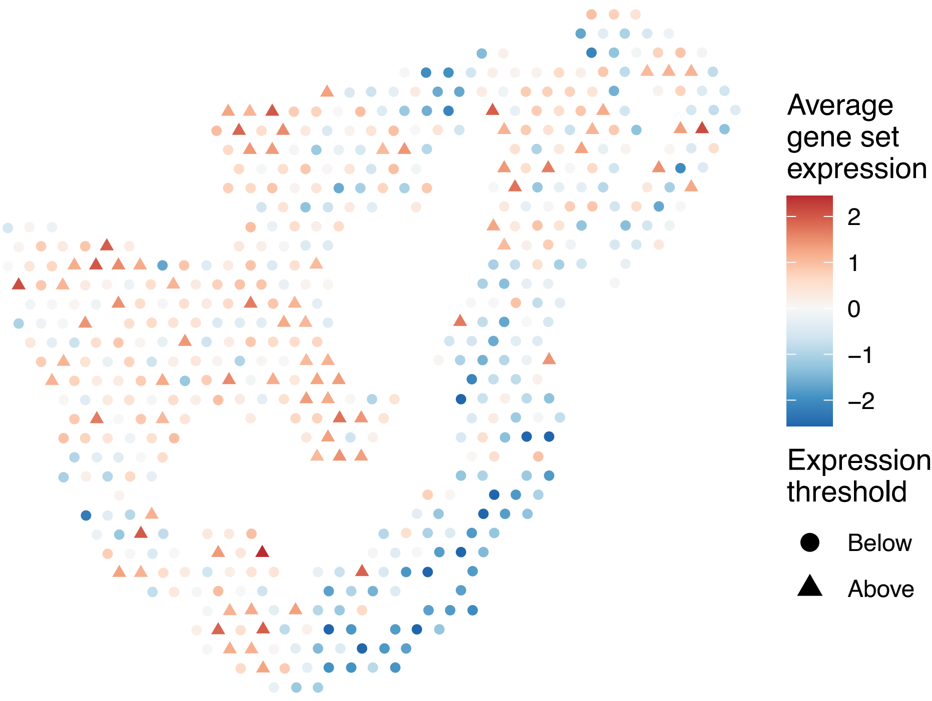


sample2_LMM_B2
KEGG_AMINOACYL_TRNA_BIOSYNTHESIS
Proportion of pathway genes=0.51; Adjusted p−value=0.1151

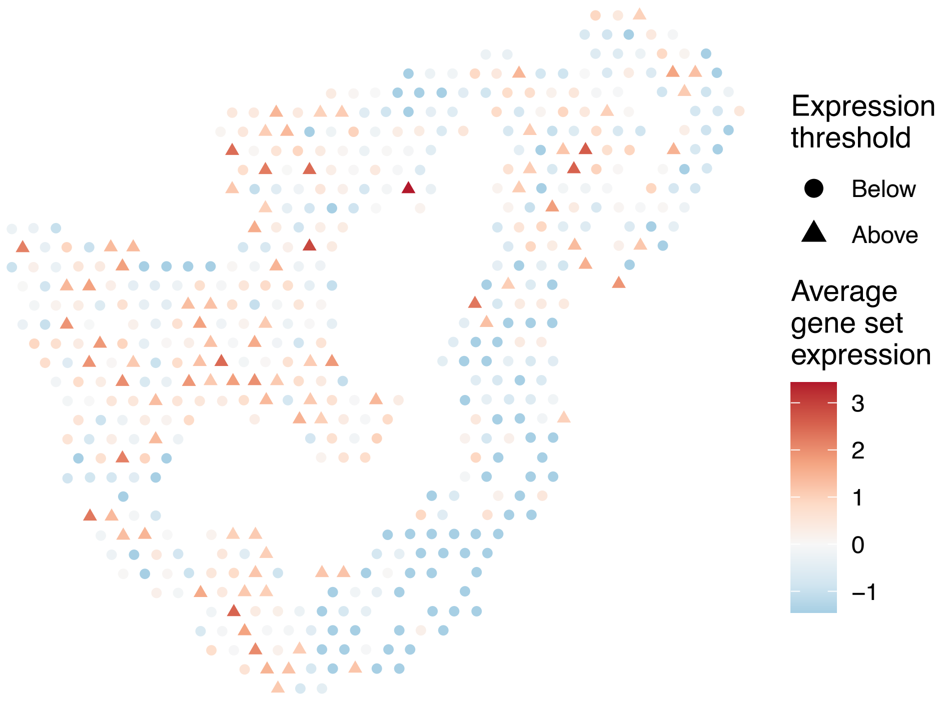


sample2_LMM_B2
KEGG_AMYOTROPHIC_LATERAL_SCLEROSIS_ALS
Proportion of pathway genes=0.6; Adjusted p−value=0.4558

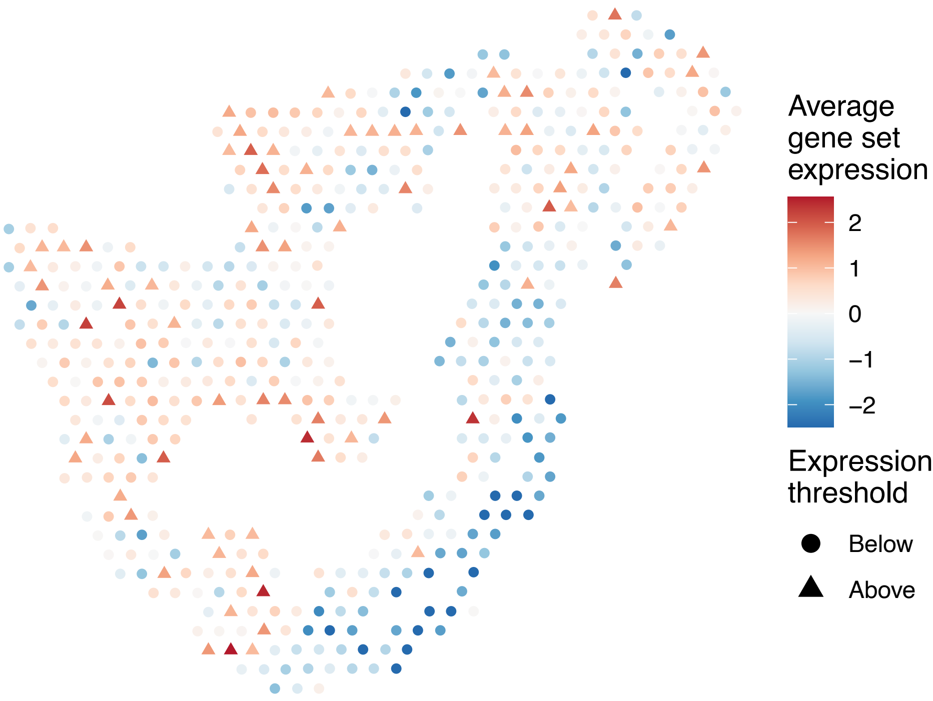

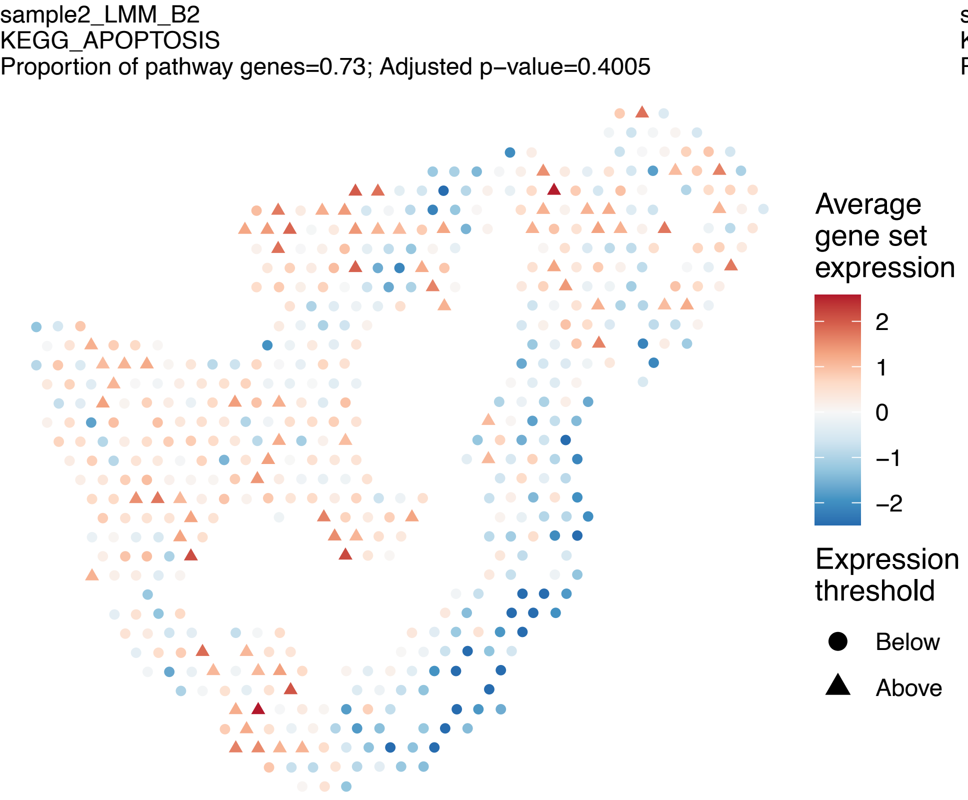
sample2_LMM_B2
KEGG_APOPTOSIS
Proportion of pathway genes=0.73; Adjusted p−value=0.4005
Average gene set expression
2
1
0
−1
−2
Expression threshold
Below
Above

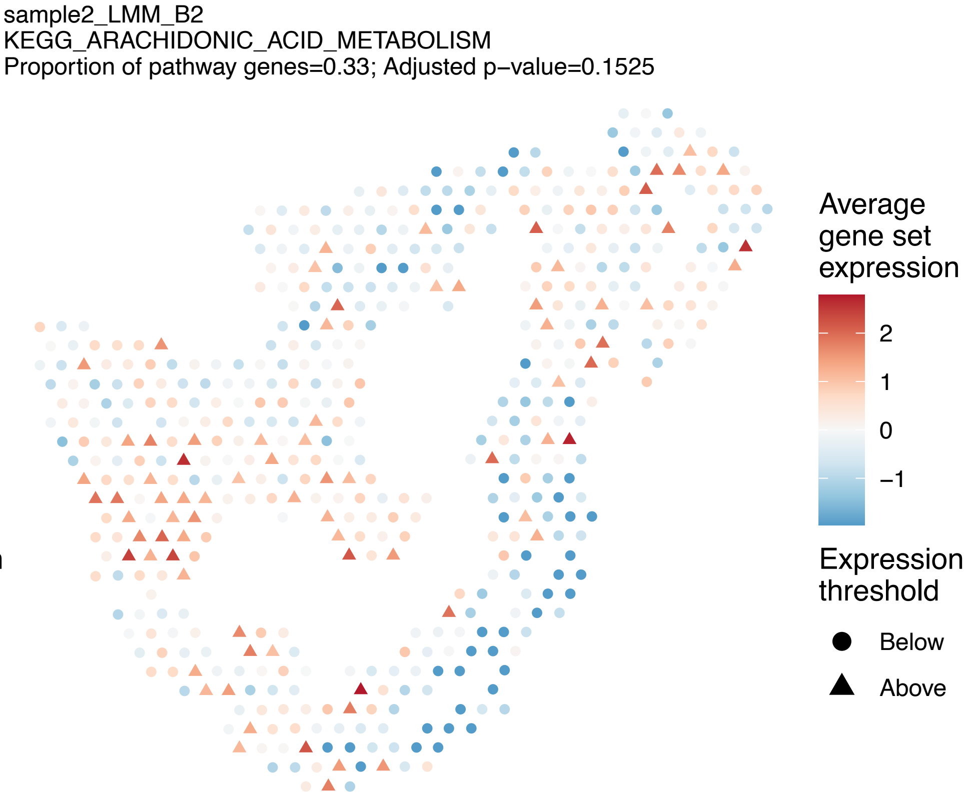
sample2_LMM_B2
KEGG_ARACHIDONIC_ACID_METABOLISM
Proportion of pathway genes=0.33; Adjusted p−value=0.1525
Average gene set expression
2
1
0
−1
Expression threshold
Below
Above

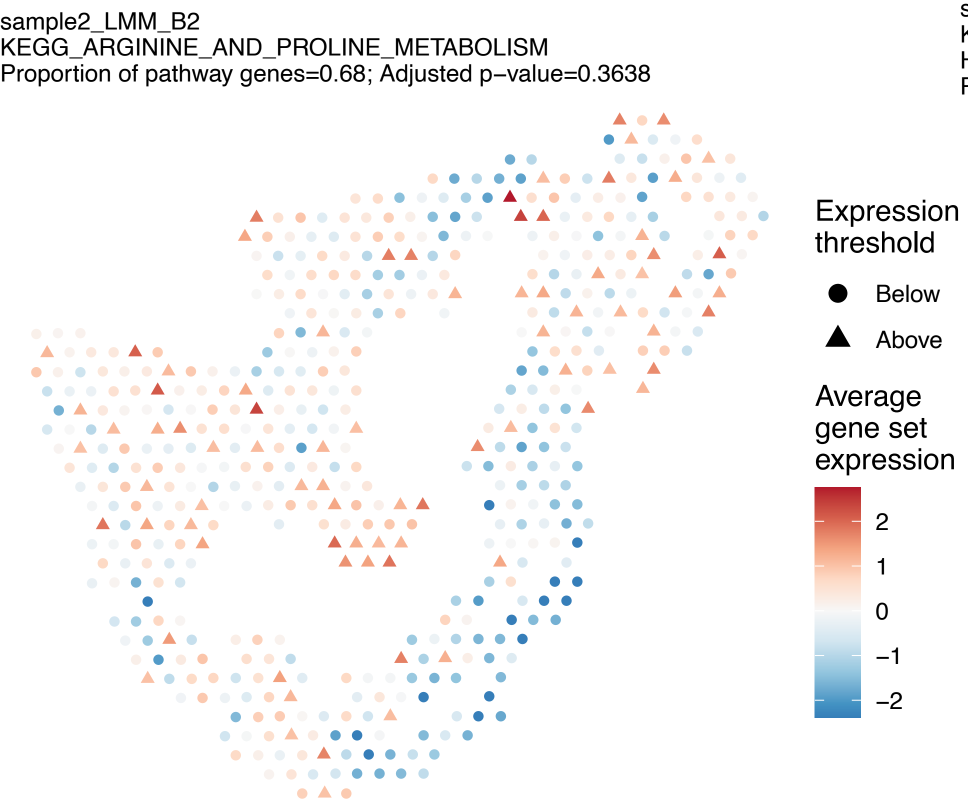
sample2_LMM_B2
KEGG_ARGININE_AND_PROLINE_METABOLISM
Proportion of pathway genes=0.68; Adjusted p−value=0.3638
Expression threshold
Below
Above
Average gene set expression
2
1
0
−1
−2

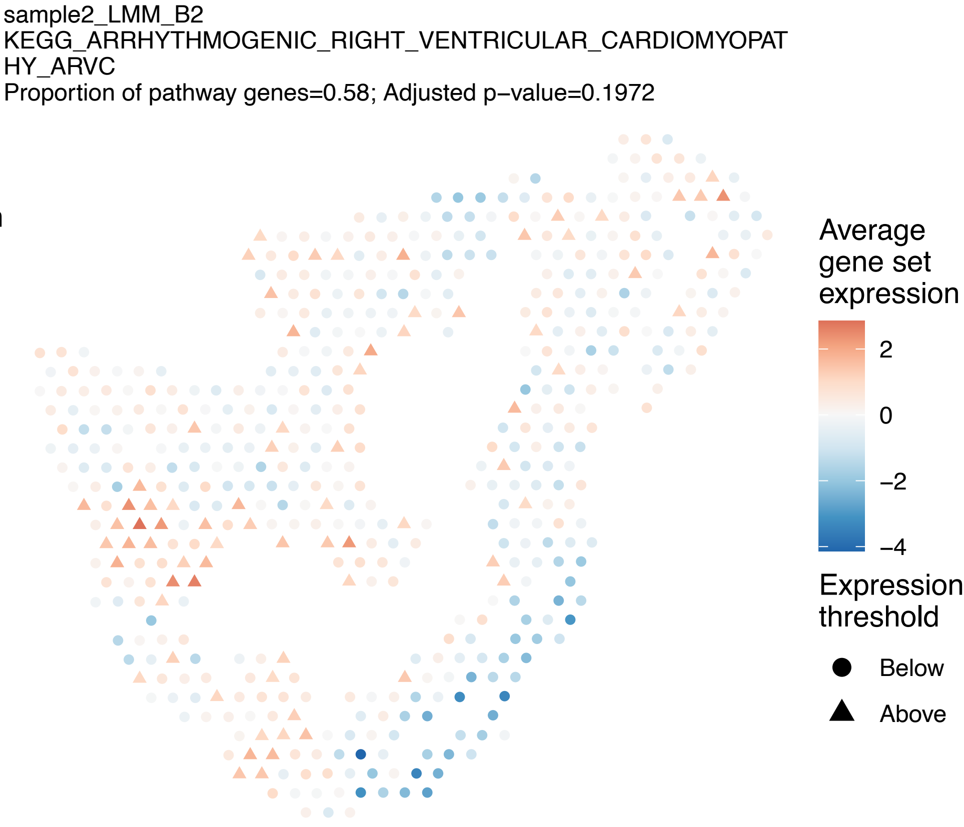
sample2_LMM_B2
KEGG_ARRHYTHMOGENIC_RIGHT_VENTRICULAR_CARDIOMYOPAT
HY_ARVC
Proportion of pathway genes=0.58; Adjusted p−value=0.1972
Average gene set expression
2
0
−2
−4
Expression threshold
Below
Above

sample2_LMM_B2
KEGG_ASTHMA
Proportion of pathway genes=0.56; Adjusted p−value=0.115

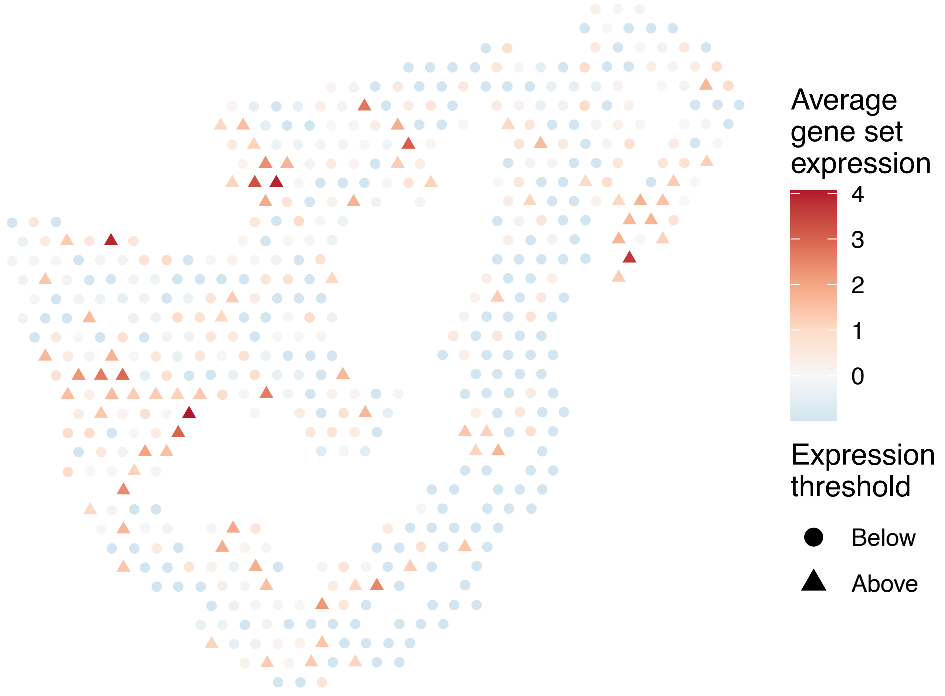


sample2_LMM_B2
KEGG_AUTOIMMUNE_THYROID_DISEASE
Proportion of pathway genes=0.59; Adjusted p−value=0

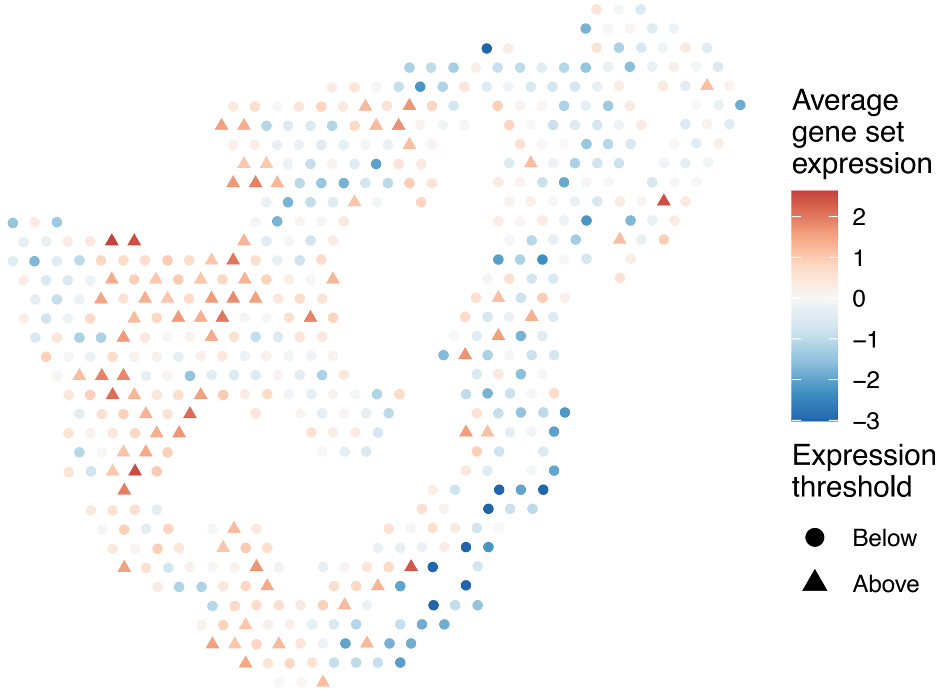


sample2_LMM_B2
KEGG_AXON_GUIDANCE
Proportion of pathway genes=0.7; Adjusted p−value=0.0514

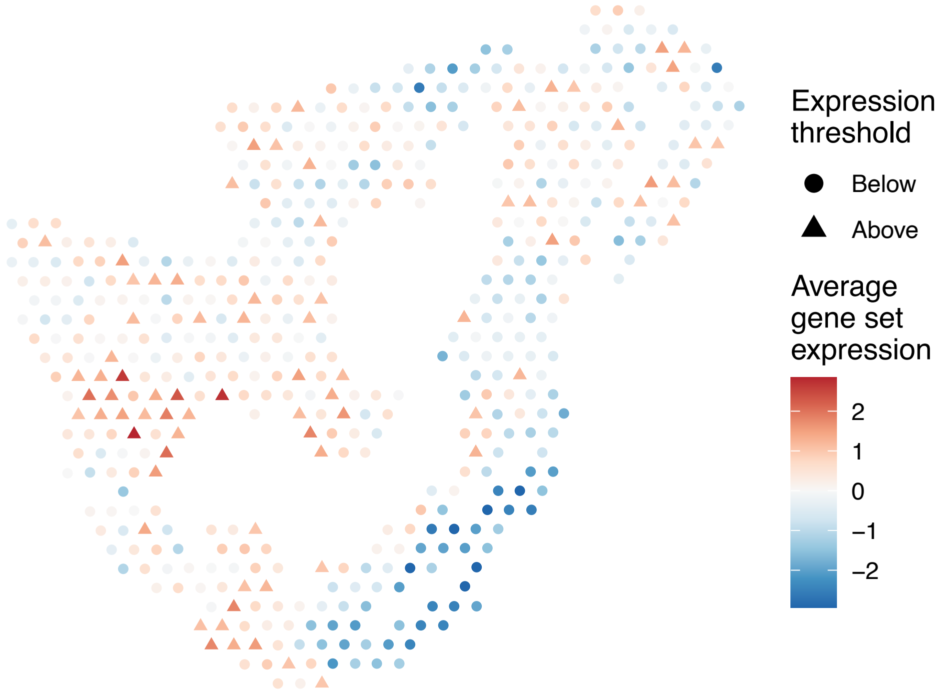


sample2_LMM_B2
KEGG_B_CELL_RECEPTOR_SIGNALING_PATHWAY
Proportion of pathway genes=0.81; Adjusted p−value=0.4377

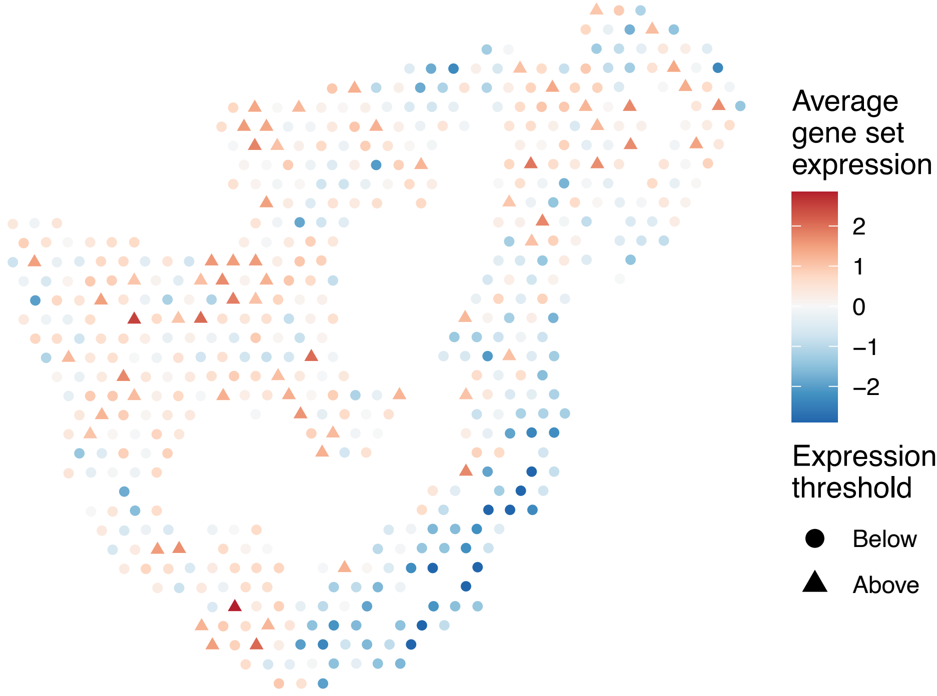

sample2_LMM_B2
KEGG_BASAL_CELL_CARCINOMA
Proportion of pathway genes=0.47; Adjusted p−value=0.3177

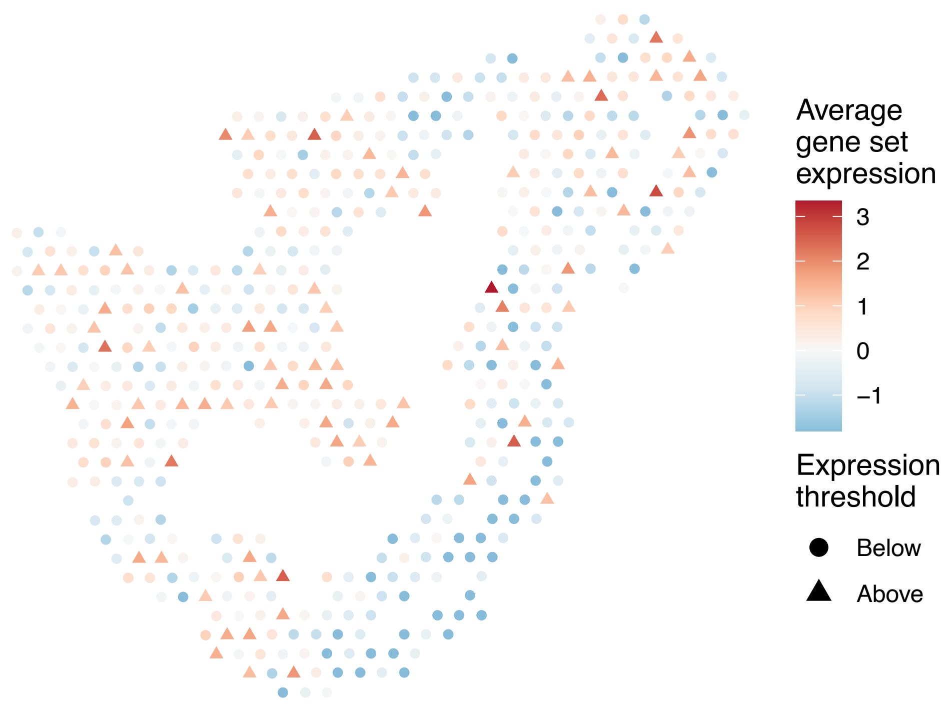


sample2_LMM_B2
KEGG_BASAL_TRANSCRIPTION_FACTORS
Proportion of pathway genes=0.87; Adjusted p−value=0.0696

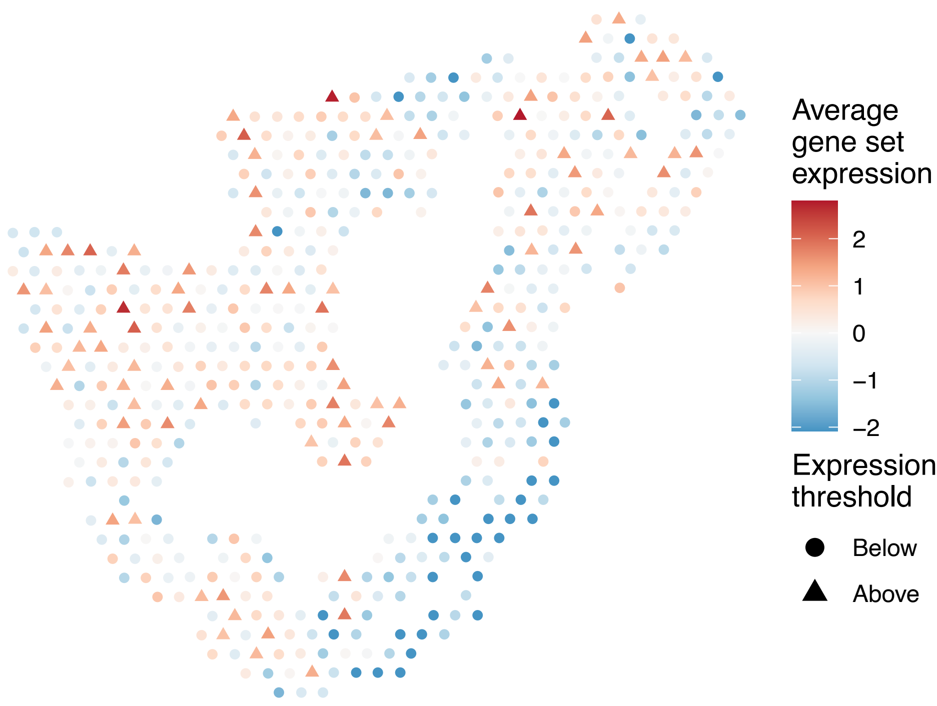


sample2_LMM_B2
KEGG_BASE_EXCISION_REPAIR
Proportion of pathway genes=0.94; Adjusted p−value=0.5638

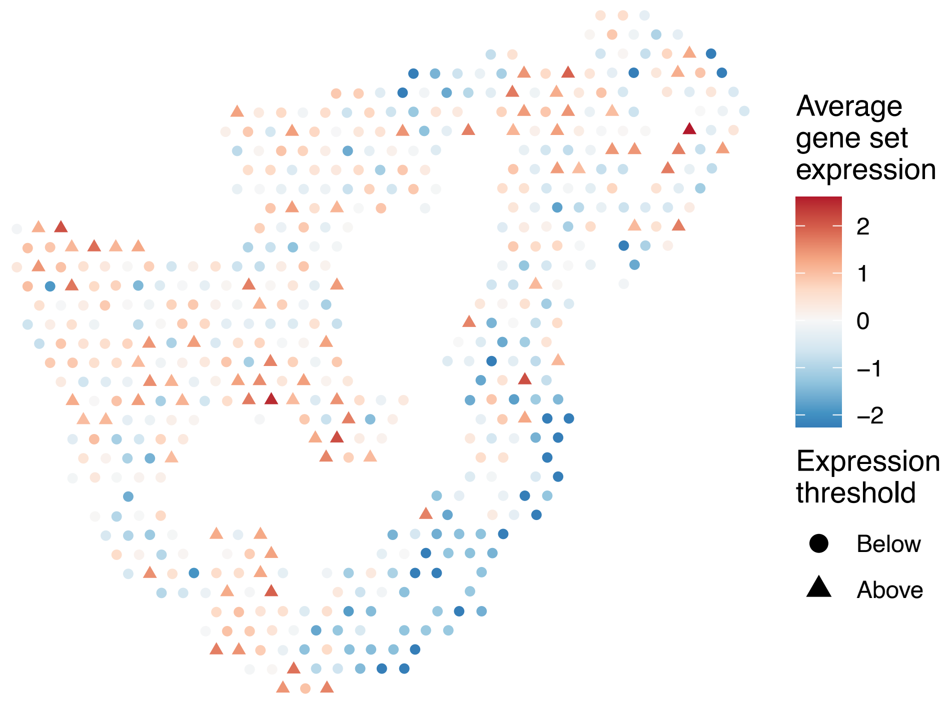


sample2_LMM_B2
KEGG_BETA_ALANINE_METABOLISM
Proportion of pathway genes=0.64; Adjusted p−value=0.2278

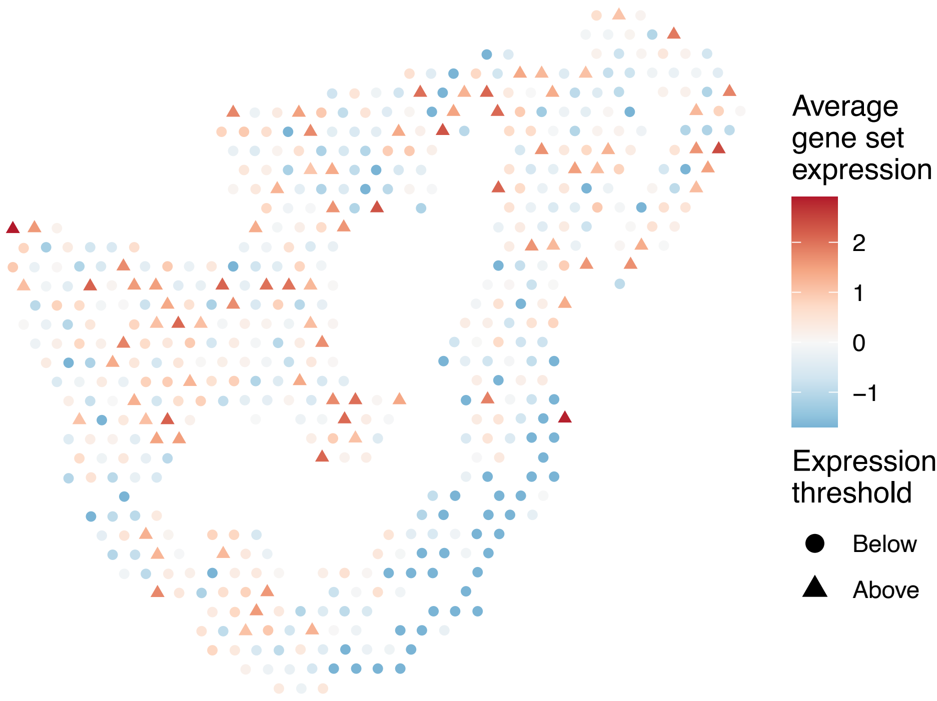

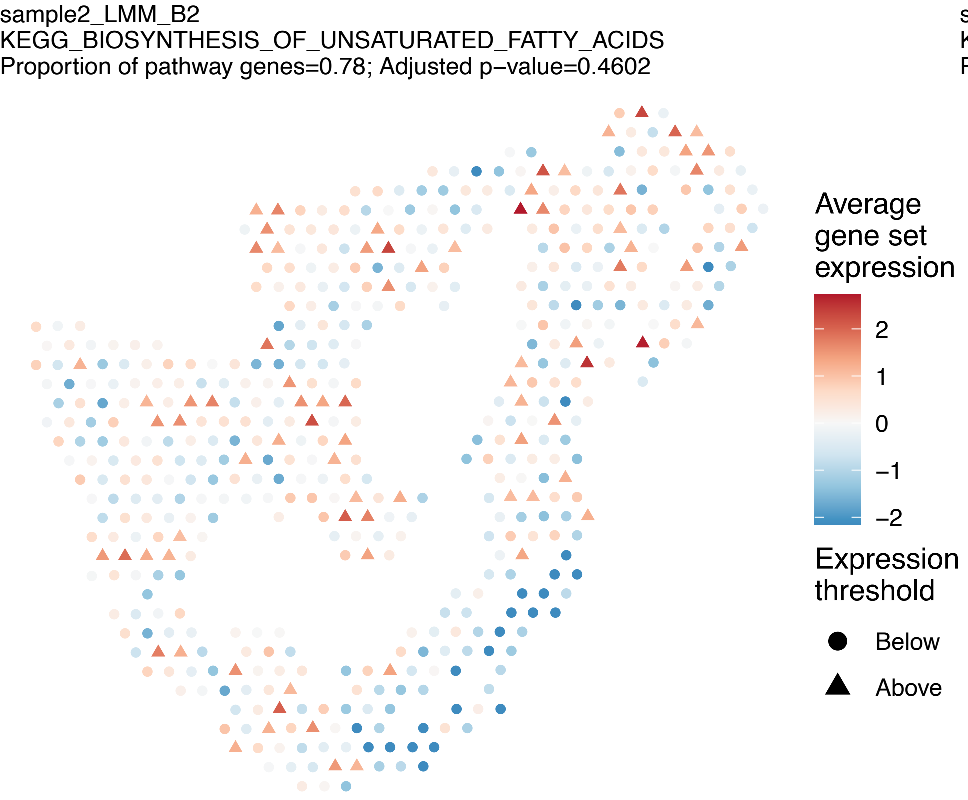
sample2_LMM_B2
KEGG_BIOSYNTHESIS_OF_UNSATURATED_FATTY_ACIDS
Proportion of pathway genes=0.78; Adjusted p−value=0.4602
Average gene set expression
2
1
0
−1
−2
Expression threshold
Below
Above

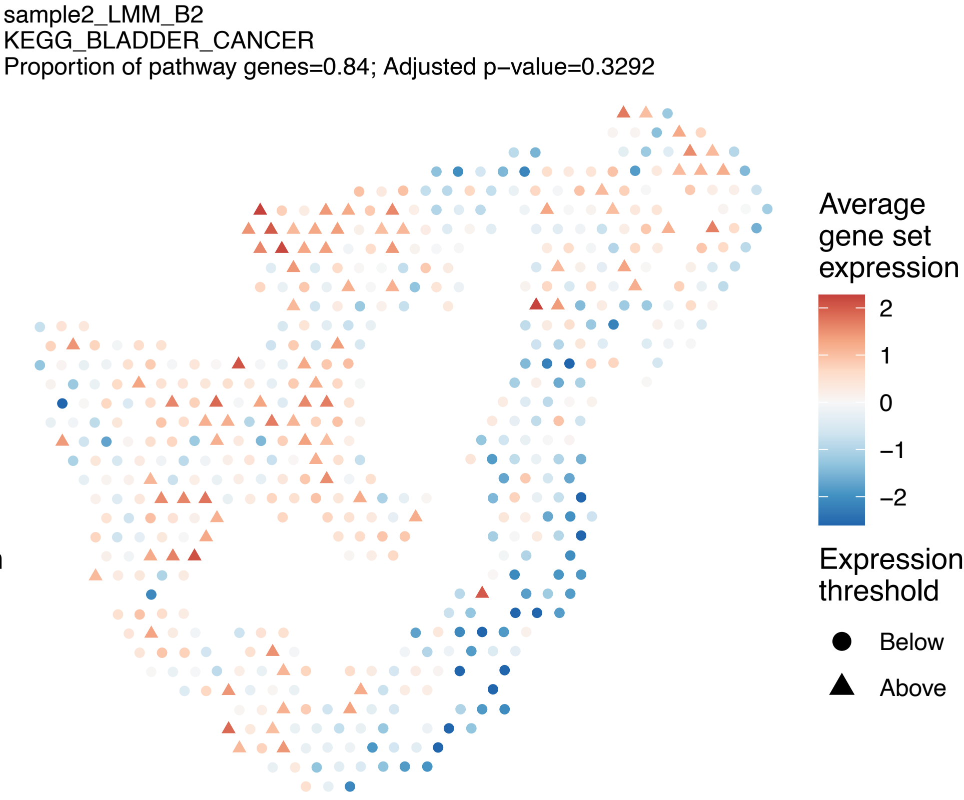
sample2_LMM_B2
KEGG_BLADDER_CANCER
Proportion of pathway genes=0.84; Adjusted p−value=0.3292
Average gene set expression
2
1
0
−1
−2
Expression threshold
Below
Above

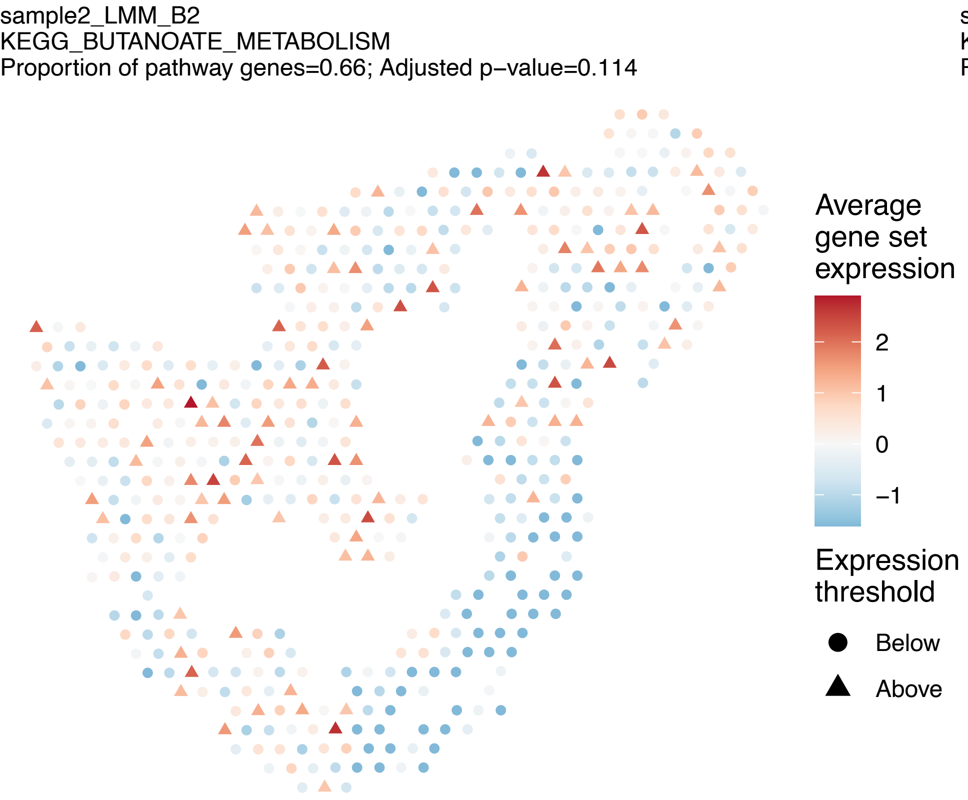
sample2_LMM_B2
KEGG_BUTANOATE_METABOLISM
Proportion of pathway genes=0.66; Adjusted p−value=0.114
Average gene set expression
2
1
0
−1
Expression threshold
Below
Above

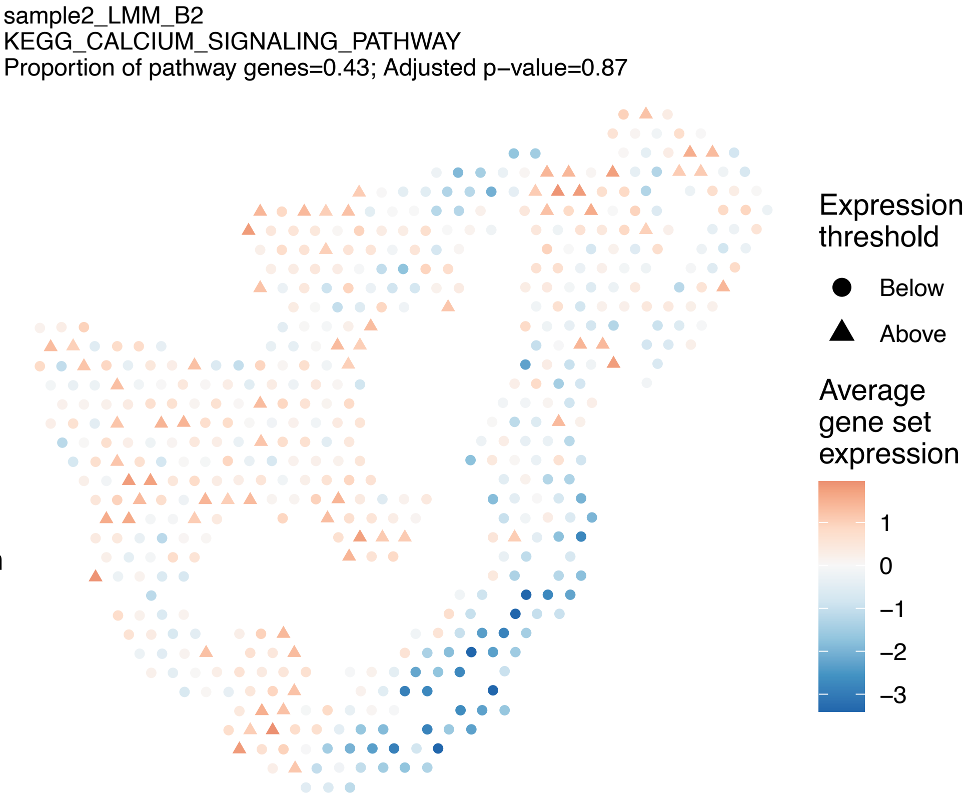
sample2_LMM_B2
KEGG_CALCIUM_SIGNALING_PATHWAY
Proportion of pathway genes=0.43; Adjusted p−value=0.87
Expression threshold
Below
Above
Average gene set expression
1
0
−1
−2
−3

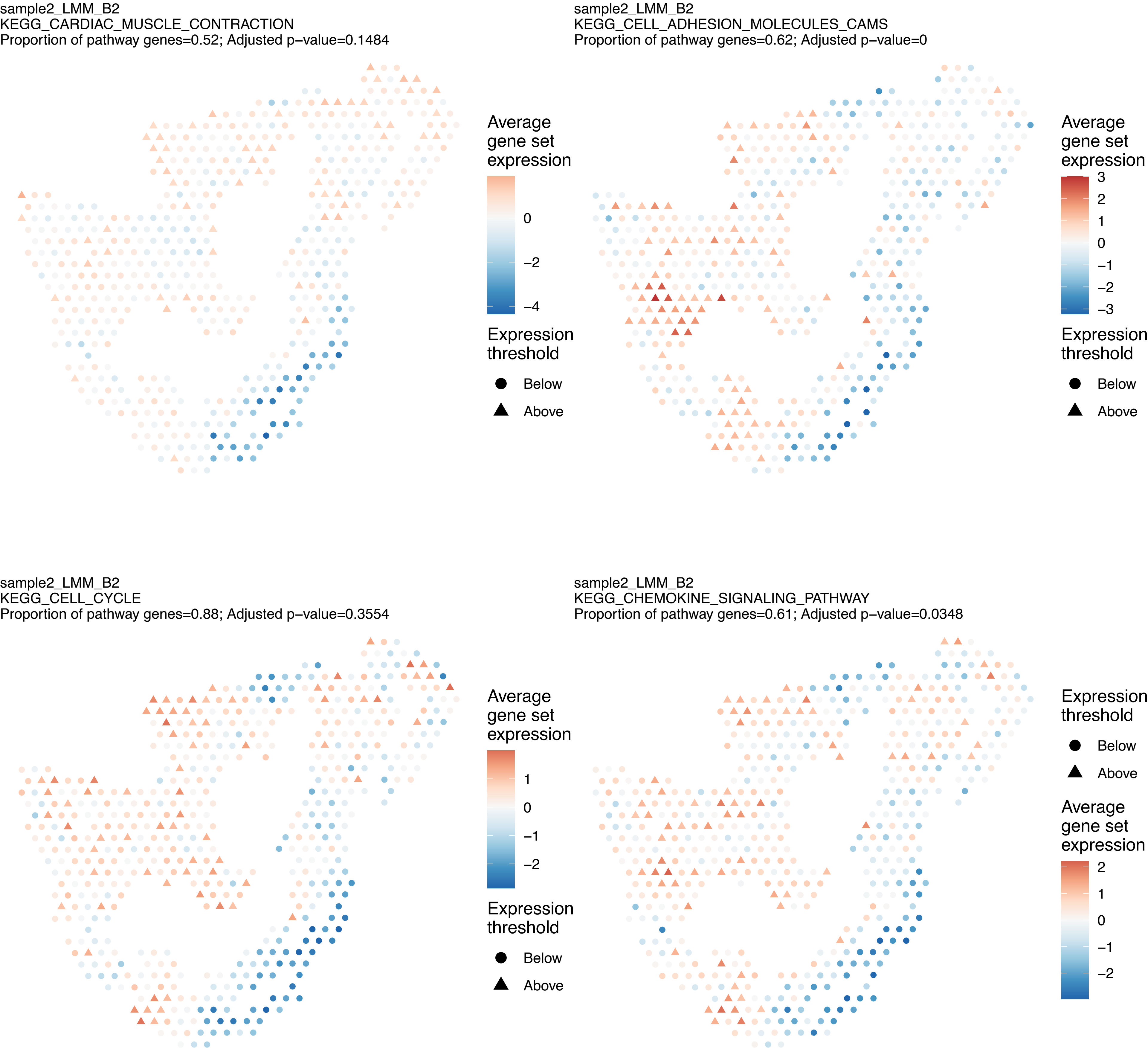
sample2_LMM_B2
KEGG_CARDIAC_MUSCLE_CONTRACTION
Proportion of pathway genes=0.52; Adjusted p−value=0.1484
Average gene set expression
0
−2
−4
Expression threshold
Below
Above
sample2_LMM_B2
KEGG_CELL_ADHESION_MOLECULES_CAMS
Proportion of pathway genes=0.62; Adjusted p−value=0
Average gene set expression
3
2
1
0
−1
−2
−3
Expression threshold
Below
Above
sample2_LMM_B2
KEGG_CELL_CYCLE
Proportion of pathway genes=0.88; Adjusted p−value=0.3554
Average gene set expression
1
0
−1
−2
Expression threshold
Below
Above
sample2_LMM_B2
KEGG_CHEMOKINE_SIGNALING_PATHWAY
Proportion of pathway genes=0.61; Adjusted p−value=0.0348
Expression threshold
Below
Above
Average gene set expression
2
1
0
−1
−2

sample2_LMM_B2
KEGG_CHRONIC_MYELOID_LEUKEMIA
Proportion of pathway genes=0.91; Adjusted p–value=0.2532

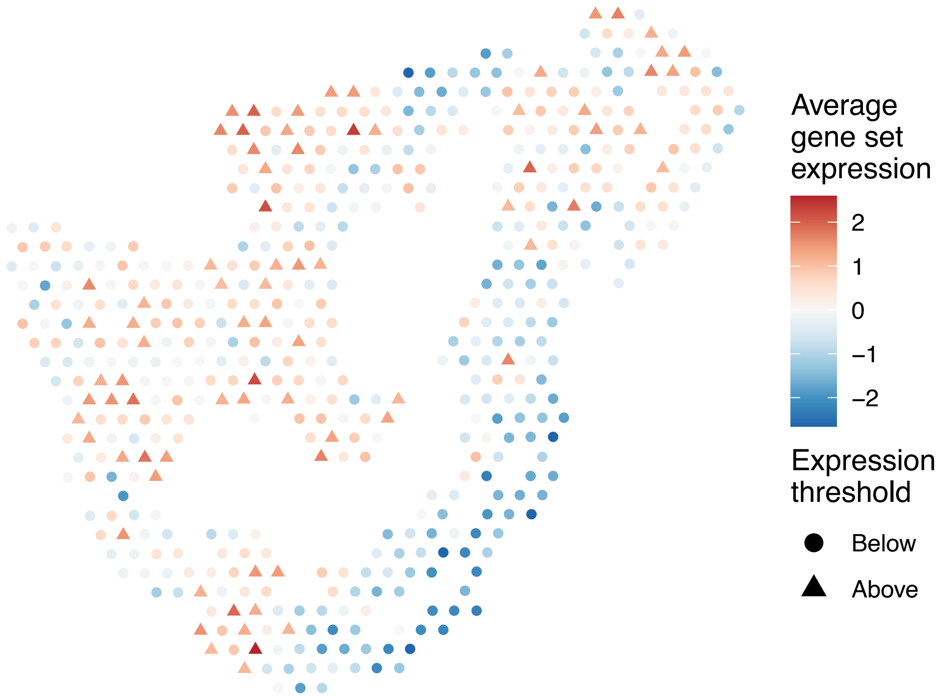


sample2_LMM_B2
KEGG_CIRCADIAN_RHYTHM_MAMMAL
Proportion of pathway genes=0.85; Adjusted p–value=0.986

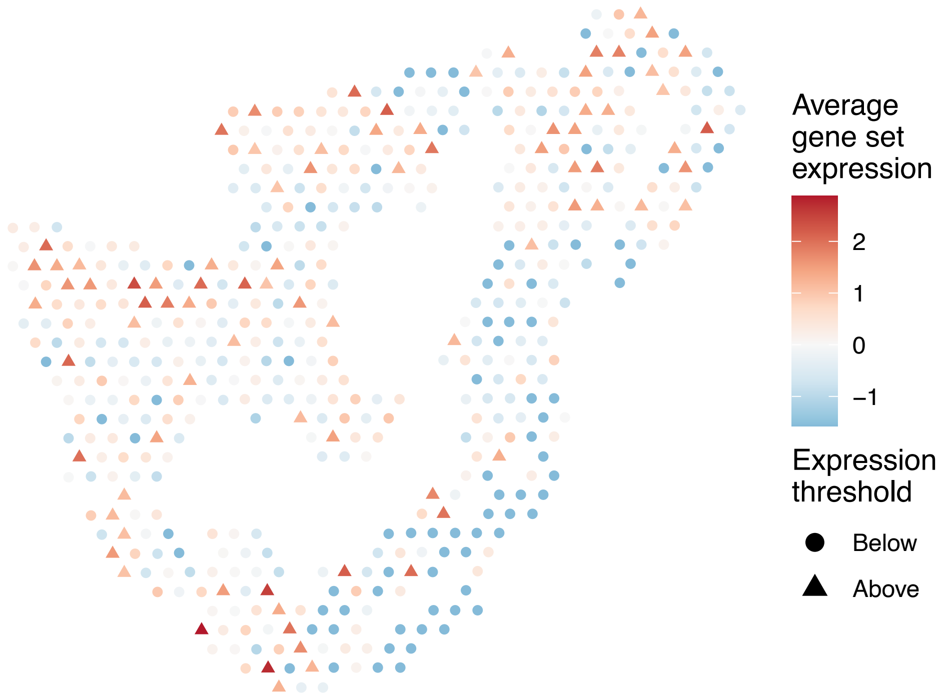


sample2_LMM_B2
KEGG_CITRATE_CYCLE_TCA_CYCLE
Proportion of pathway genes=0.88; Adjusted p–value=0.0504

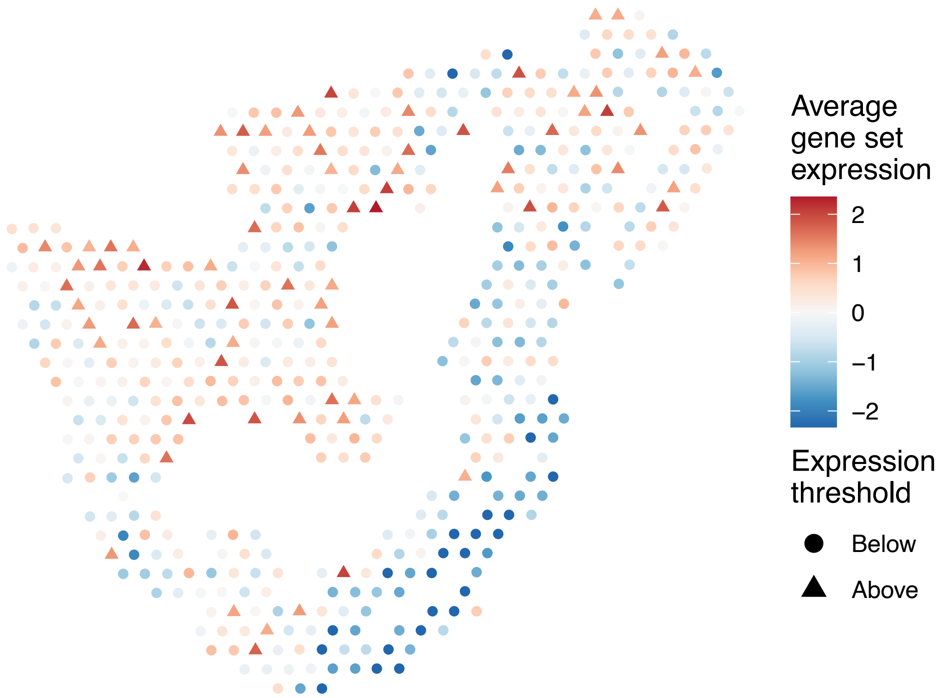


sample2_LMM_B2
KEGG_COLORECTAL_CANCER
Proportion of pathway genes=0.87; Adjusted p–value=0.1824

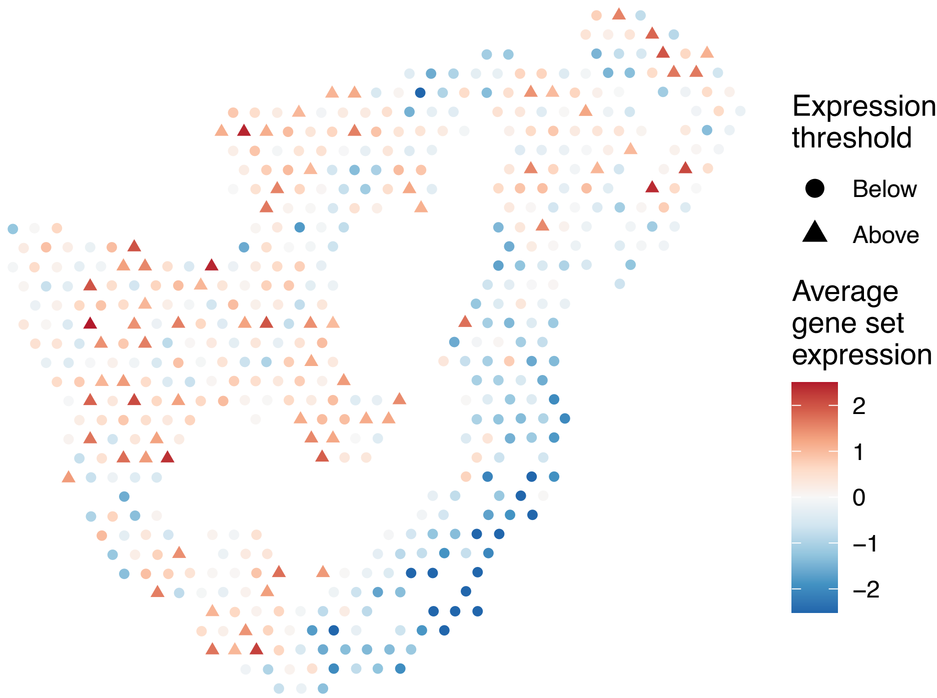

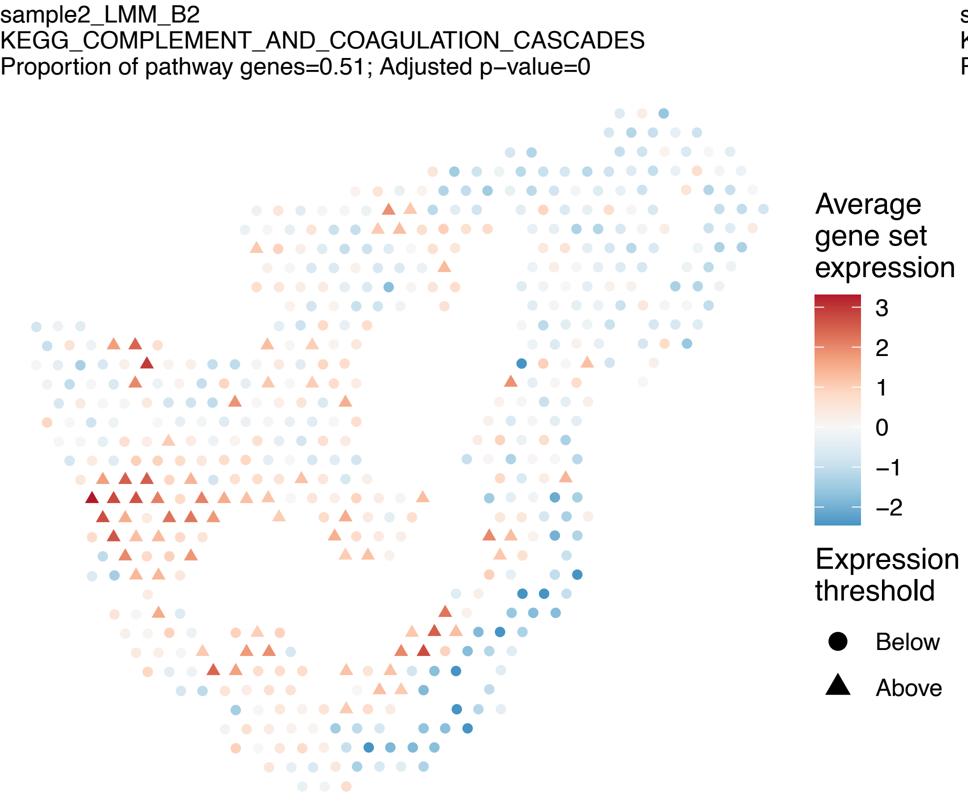
sample2_LMM_B2
KEGG_COMPLEMENT_AND_COAGULATION_CASCADES
Proportion of pathway genes=0.51; Adjusted p−value=0
Average gene set expression
3
2
1
0
−1
−2
Expression threshold
Below
Above

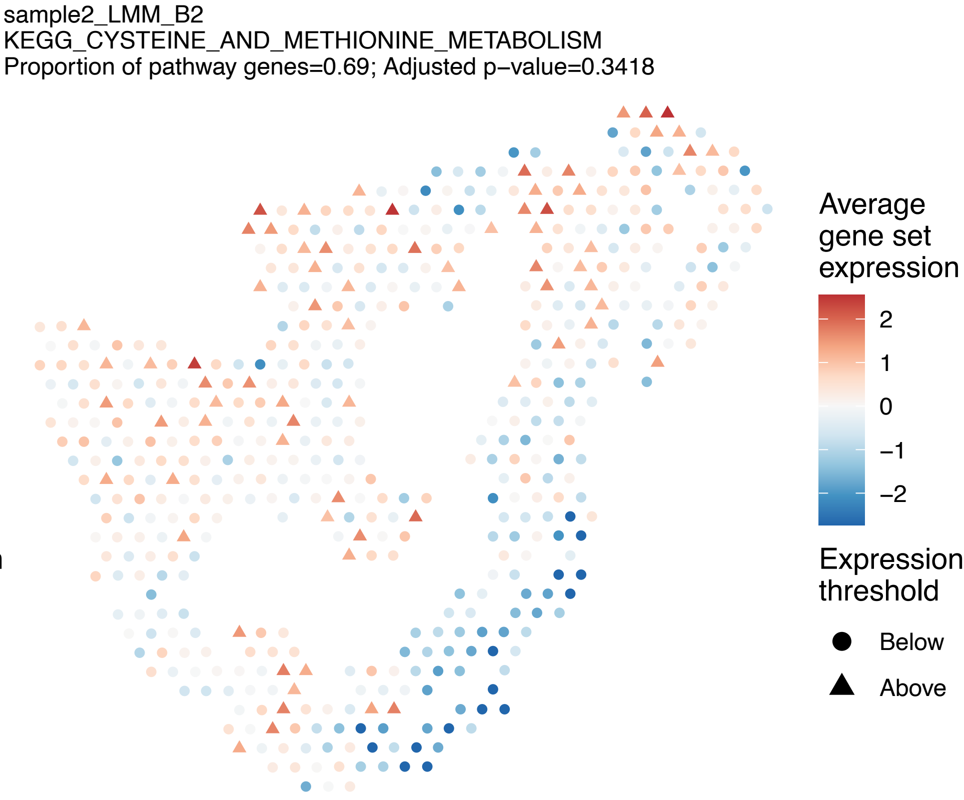
sample2_LMM_B2
KEGG_CYSTEINE_AND_METHIONINE_METABOLISM
Proportion of pathway genes=0.69; Adjusted p−value=0.3418
Average gene set expression
2
1
0
−1
−2
Expression threshold
Below
Above

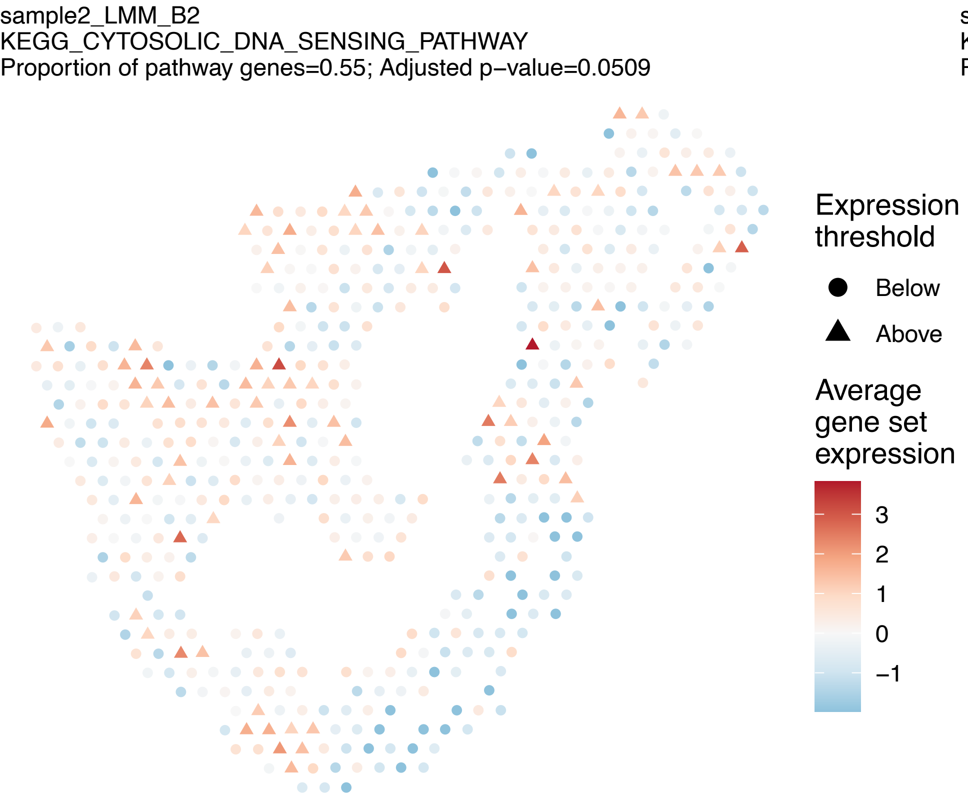
sample2_LMM_B2
KEGG_CYTOSOLIC_DNA_SENSING_PATHWAY
Proportion of pathway genes=0.55; Adjusted p−value=0.0509
Expression threshold
Below
Above
Average gene set expression
3
2
1
0
−1

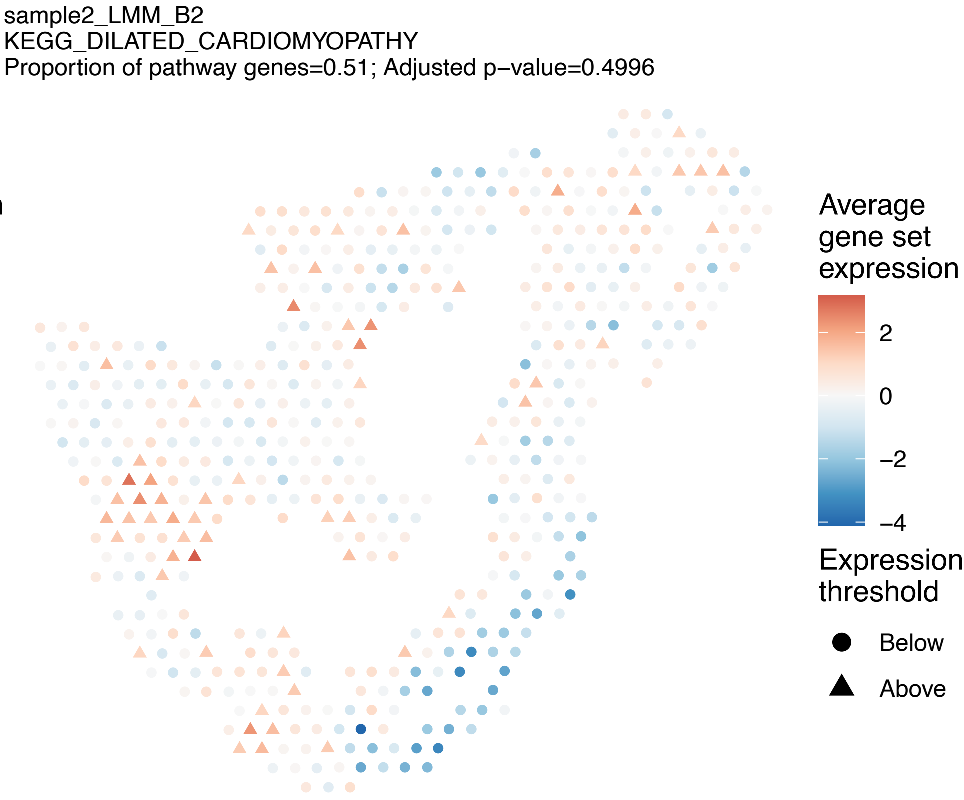
sample2_LMM_B2
KEGG_DILATED_CARDIOMYOPATHY
Proportion of pathway genes=0.51; Adjusted p−value=0.4996
Average gene set expression
2
0
−2
−4
Expression threshold
Below
Above

sample2_LMM_B2
KEGG_DNA_REPLICATION
Proportion of pathway genes=0.95; Adjusted p−value=0.0666

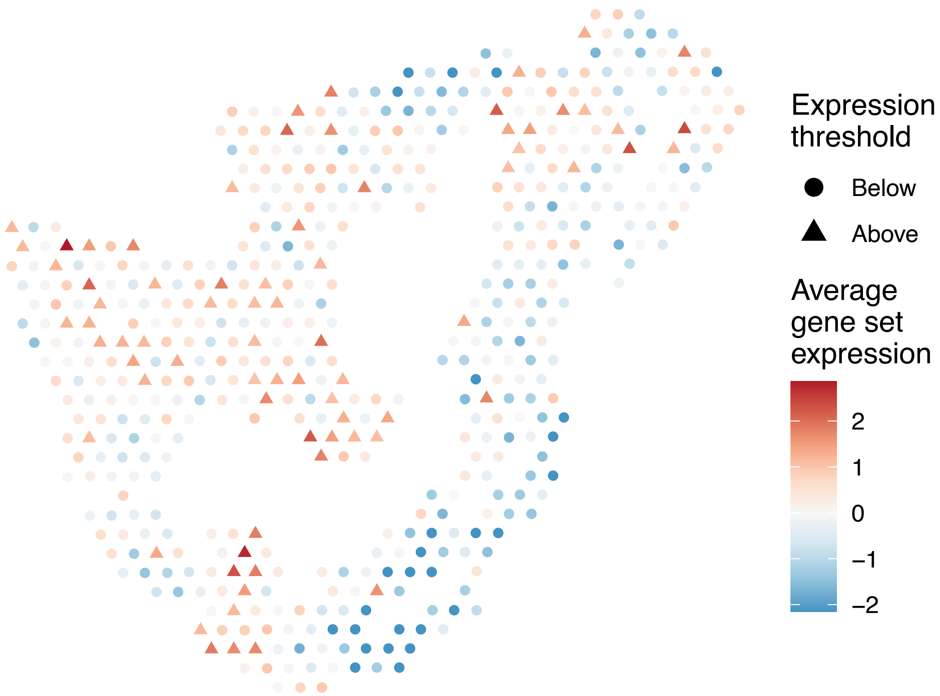


sample2_LMM_B2
KEGG_DORSO_VENTRAL_AXIS_FORMATION
Proportion of pathway genes=0.78; Adjusted p−value=0.0509

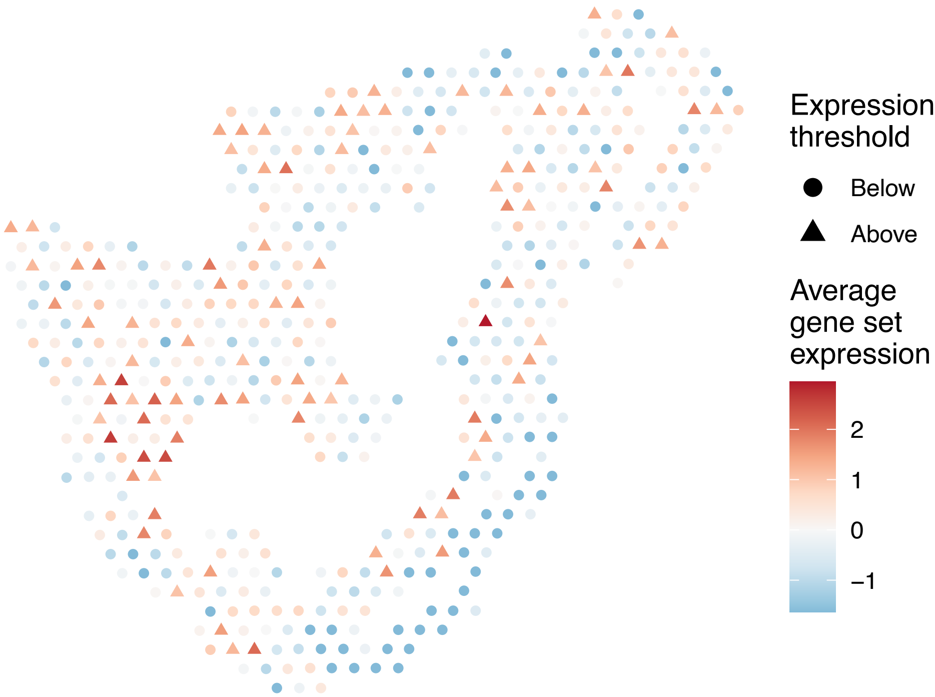


sample2_LMM_B2
KEGG_DRUG_METABOLISM_CYTOCHROME_P450
Proportion of pathway genes=0.28; Adjusted p−value=0.5365

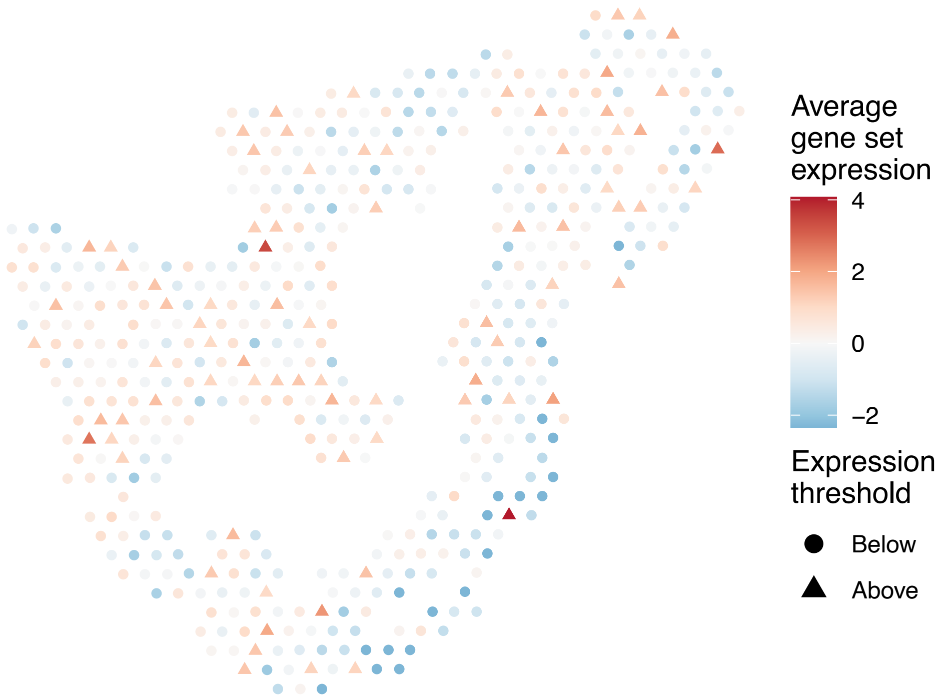


sample2_LMM_B2
KEGG_DRUG_METABOLISM_OTHER_ENZYMES
Proportion of pathway genes=0.3; Adjusted p−value=0.1151

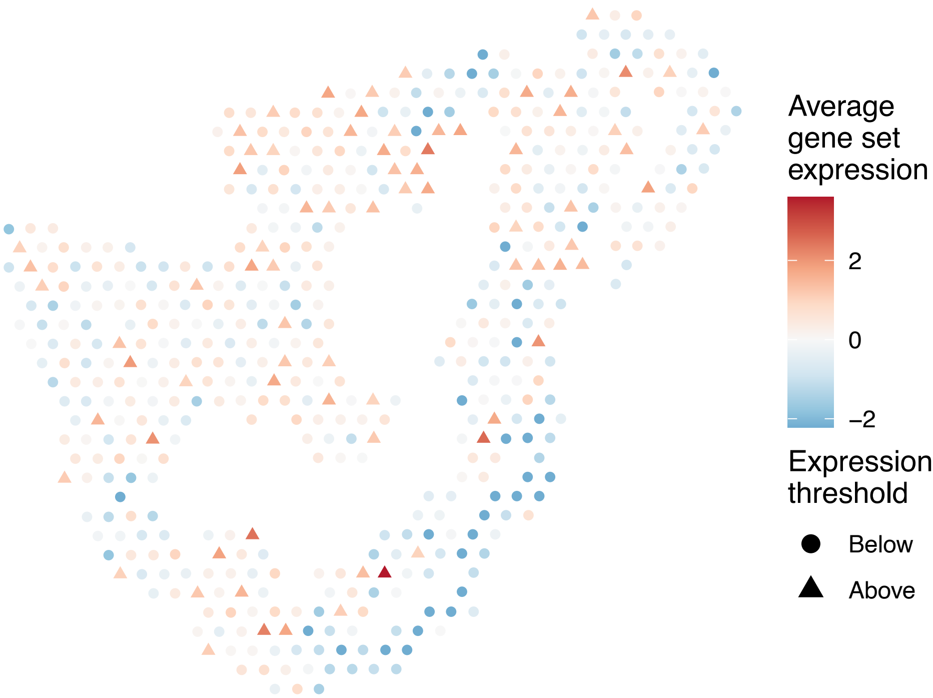

sample2_LMM_B2
KEGG_ECM_RECEPTOR_INTERACTION
Proportion of pathway genes=0.68; Adjusted p−value=0

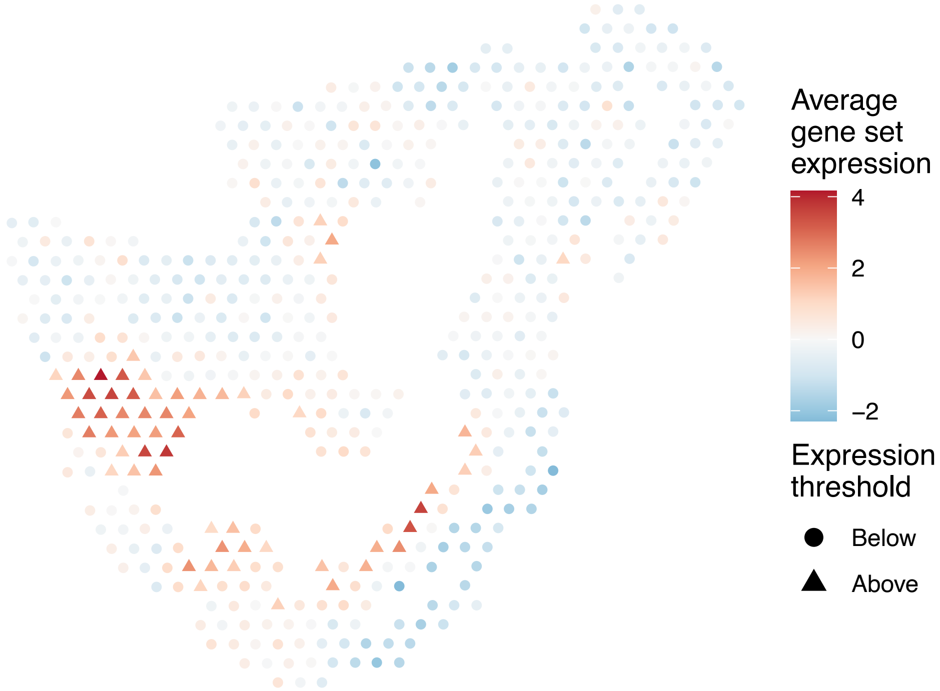


sample2_LMM_B2
KEGG_ENDOCYTOSIS
Proportion of pathway genes=0.82; Adjusted p−value=0

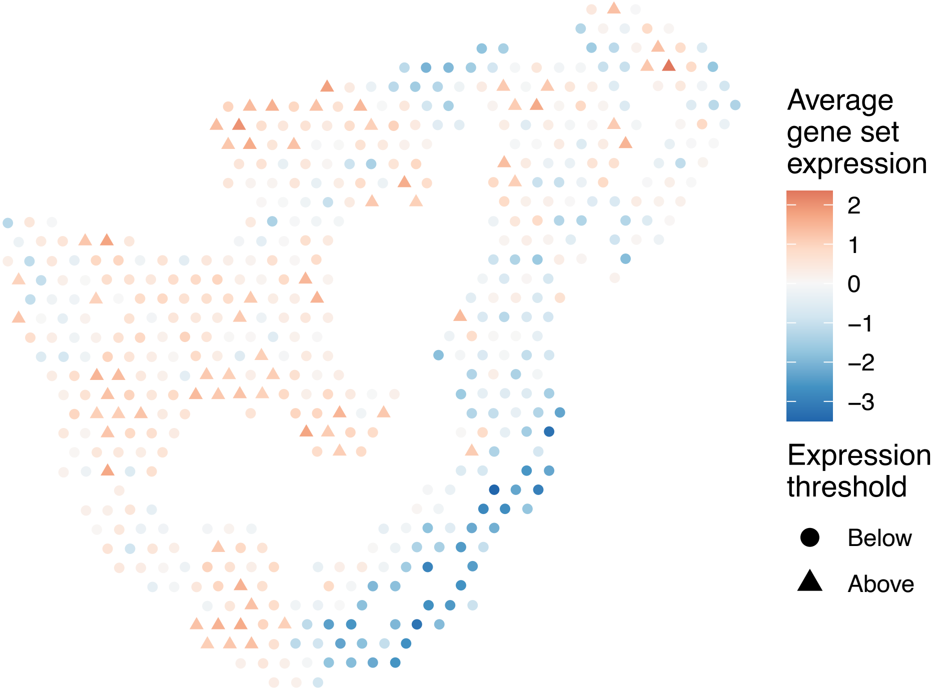


sample2_LMM_B2
KEGG_ENDOMETRIAL_CANCER
Proportion of pathway genes=0.85; Adjusted p−value=0.1824

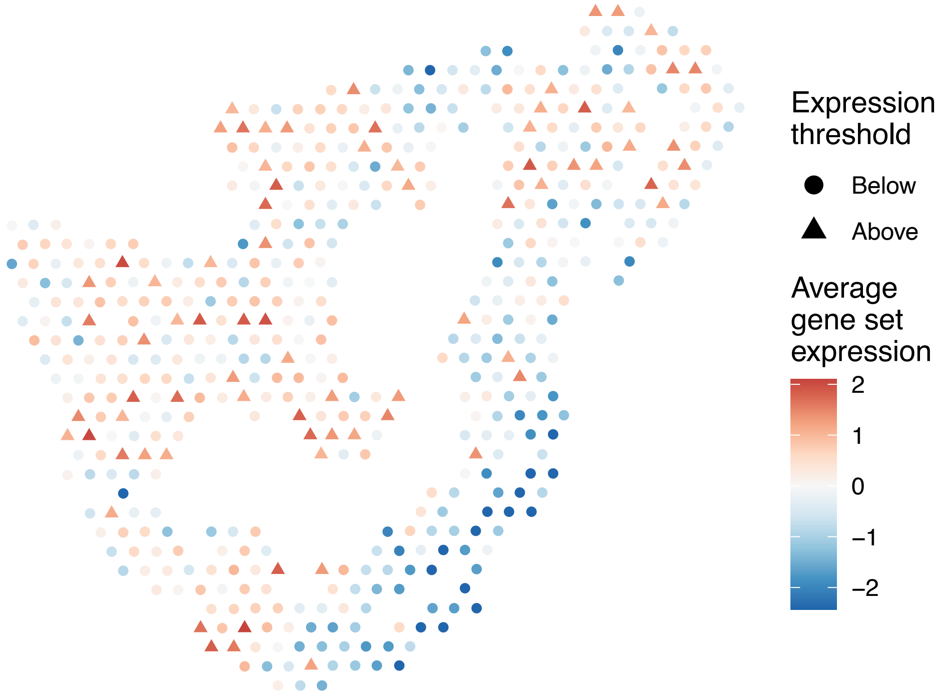


sample2_LMM_B2
KEGG_EPITHELIAL_CELL_SIGNALING_IN_HELICOBACTER_PYL
ORI_INFECTION
Proportion of pathway genes=0.76; Adjusted p−value=0.1552

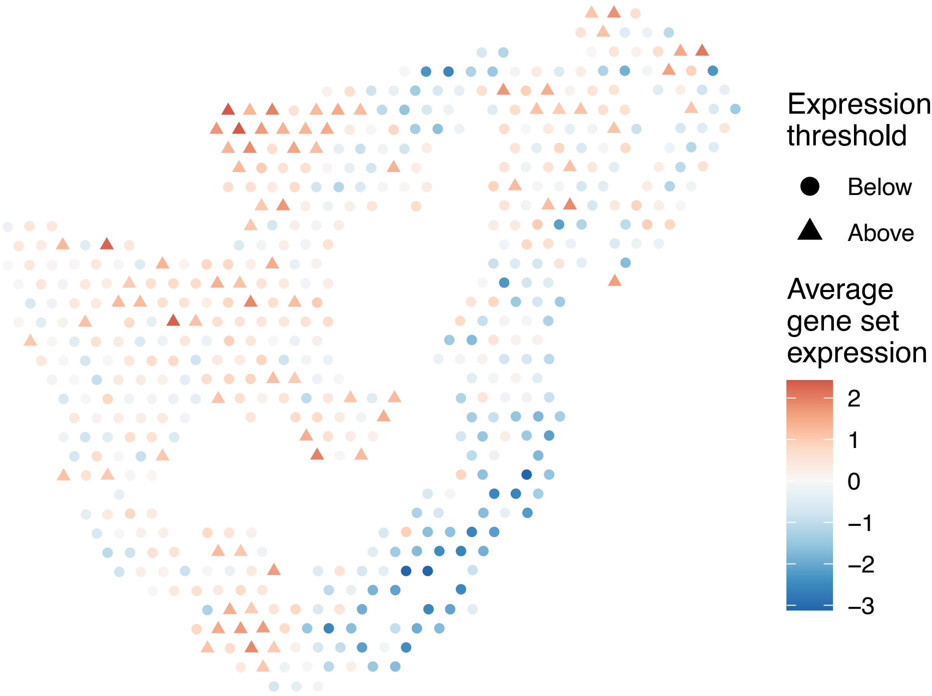

sample2_LMM_B2
KEGG_ERBB_SIGNALING_PATHWAY
Proportion of pathway genes=0.76; Adjusted p−value=0.3554

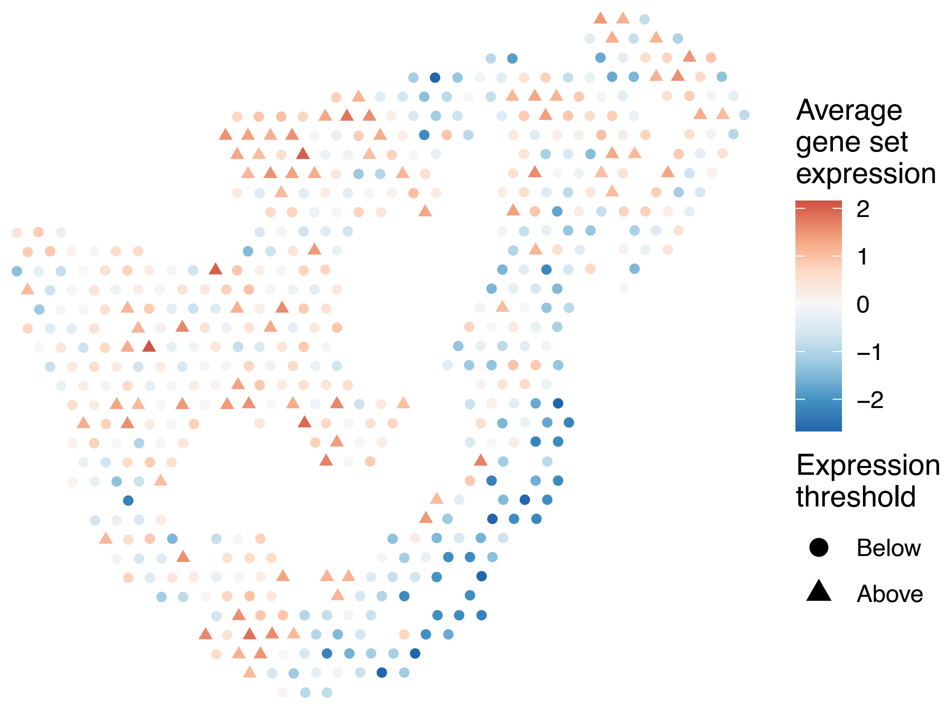


sample2_LMM_B2
KEGG_ETHER_LIPID_METABOLISM
Proportion of pathway genes=0.54; Adjusted p−value=0.0514

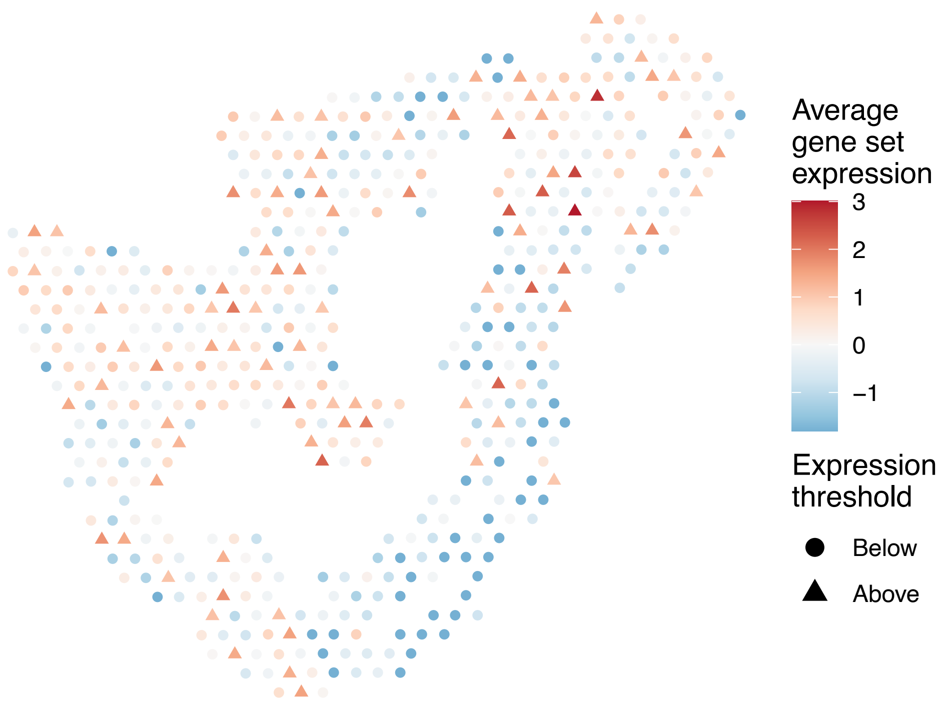


sample2_LMM_B2
KEGG_FATTY_ACID_METABOLISM
Proportion of pathway genes=0.79; Adjusted p−value=0.7004

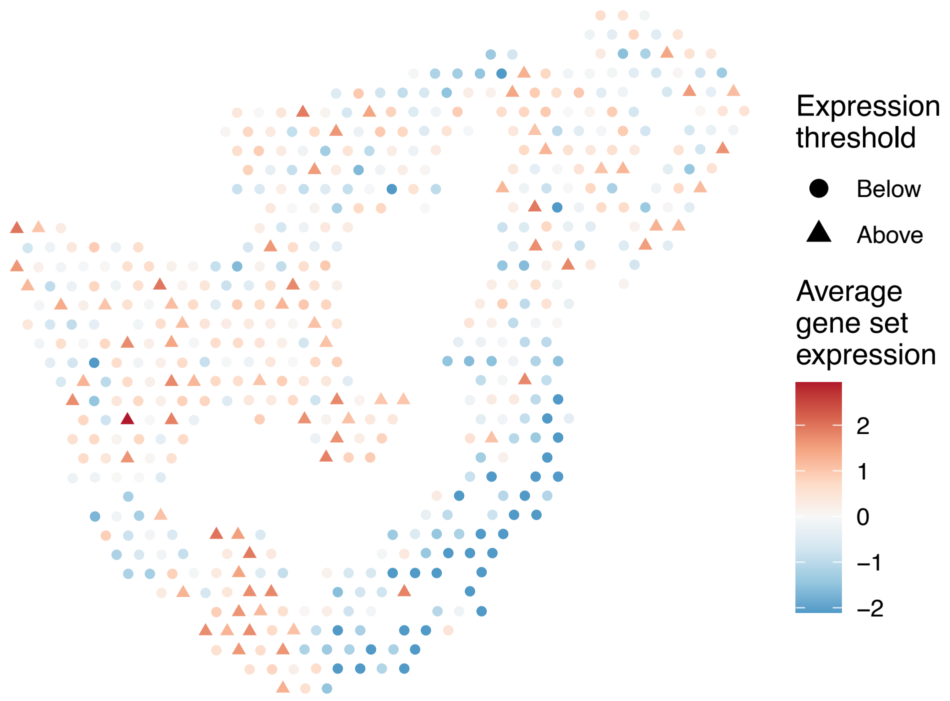


sample2_LMM_B2
KEGG_FC_EPSILON_RI_SIGNALING_PATHWAY
Proportion of pathway genes=0.6; Adjusted p−value=0.4757

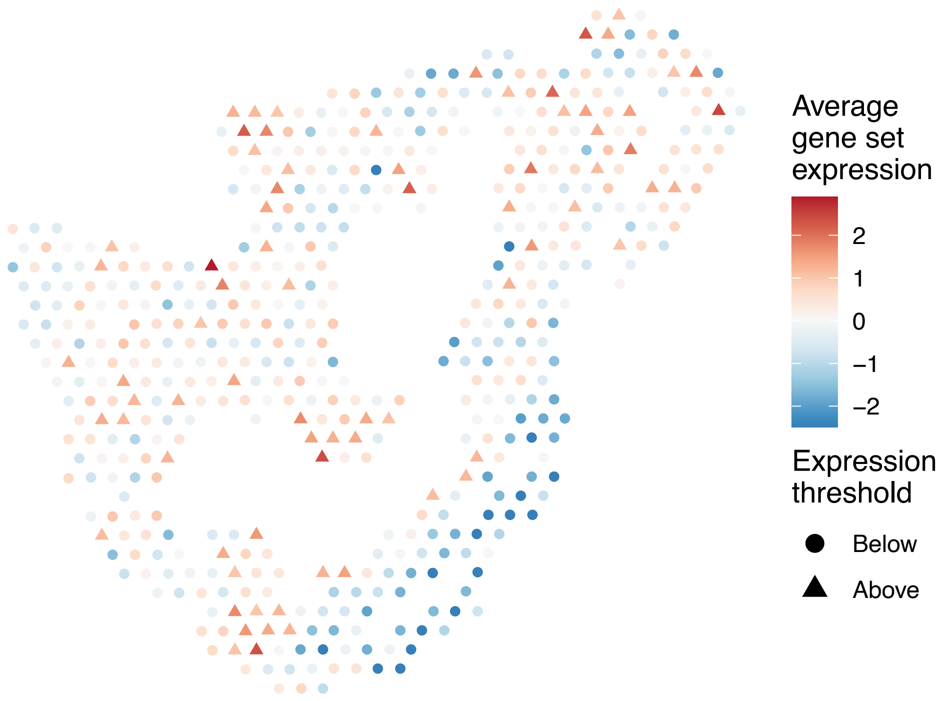

sample2_LMM_B2
KEGG_FC_GAMMA_R_MEDIATED_PHAGOCYTOSIS
Proportion of pathway genes=0.83; Adjusted p–value=0.443

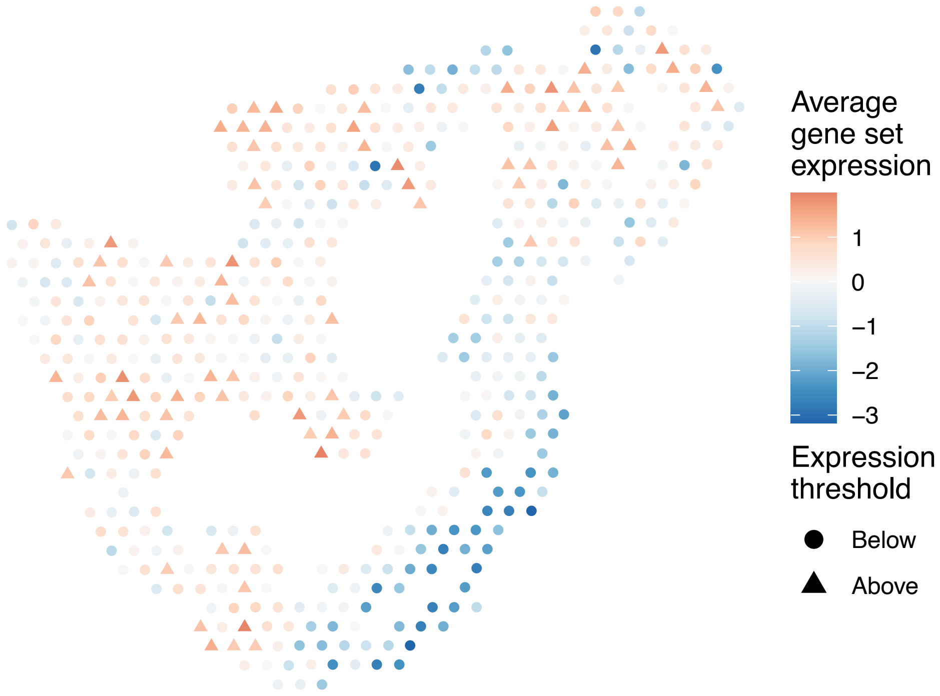


sample2_LMM_B2
KEGG_FOCAL_ADHESION
Proportion of pathway genes=0.81; Adjusted p–value=0

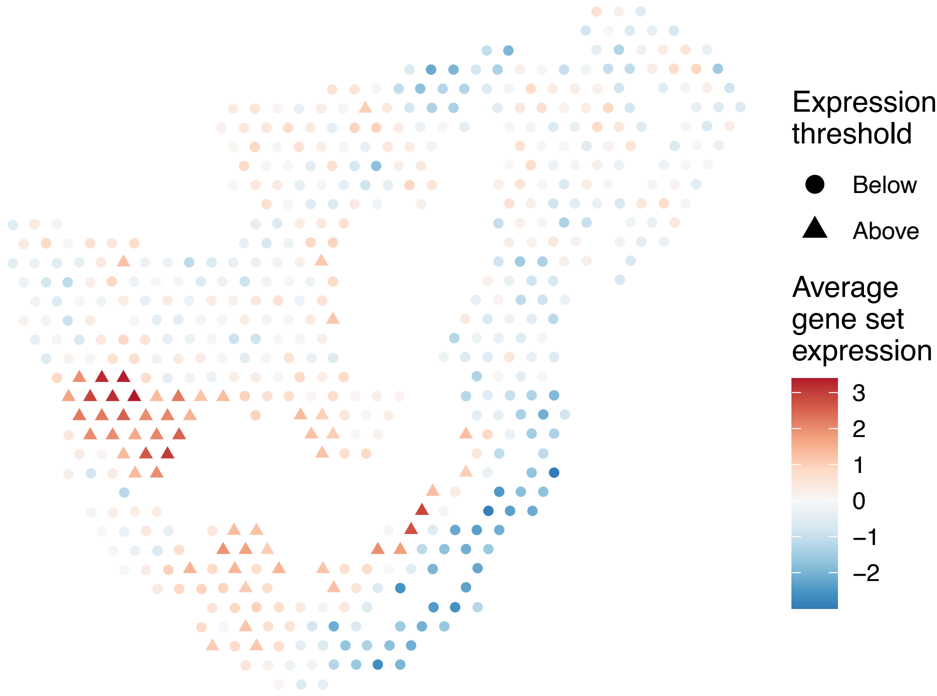


sample2_LMM_B2
KEGG_FRUCTOSE_AND_MANNOSE_METABOLISM
Proportion of pathway genes=0.71; Adjusted p–value=0.5544

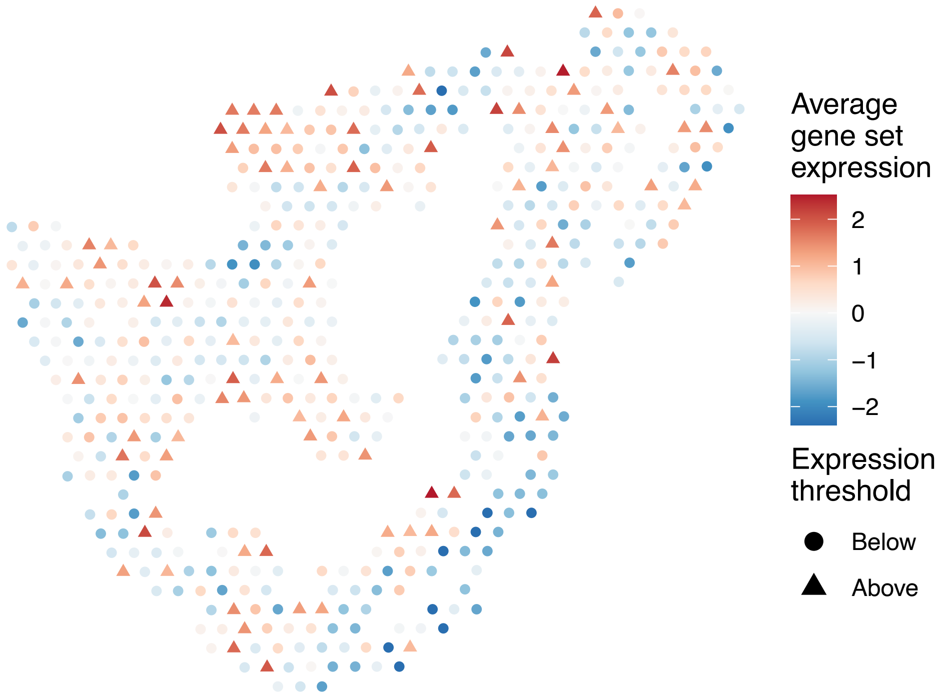


sample2_LMM_B2
KEGG_GALACTOSE_METABOLISM
Proportion of pathway genes=0.66; Adjusted p–value=0.0225

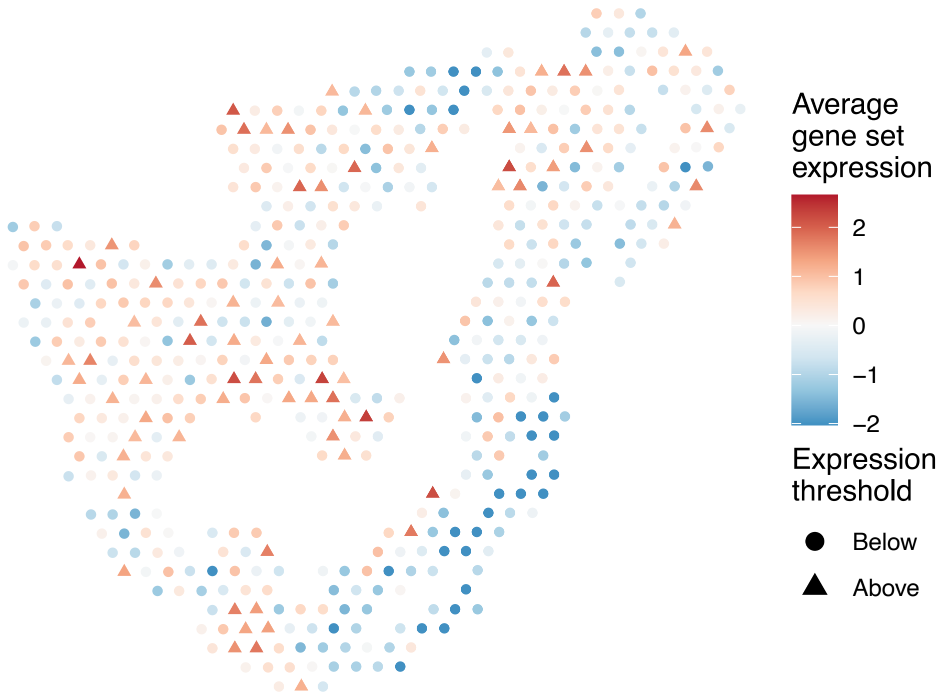

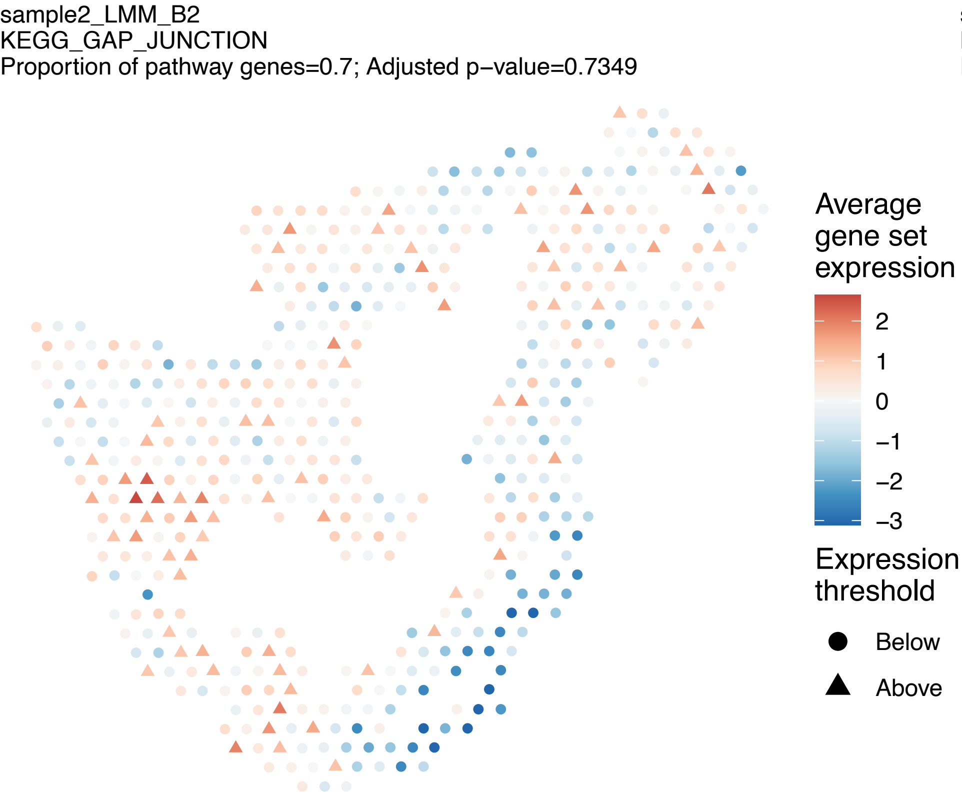
sample2_LMM_B2
KEGG_GAP_JUNCTION
Proportion of pathway genes=0.7; Adjusted p−value=0.7349
Average gene set expression
2
1
0
−1
−2
−3
Expression threshold
Below
Above

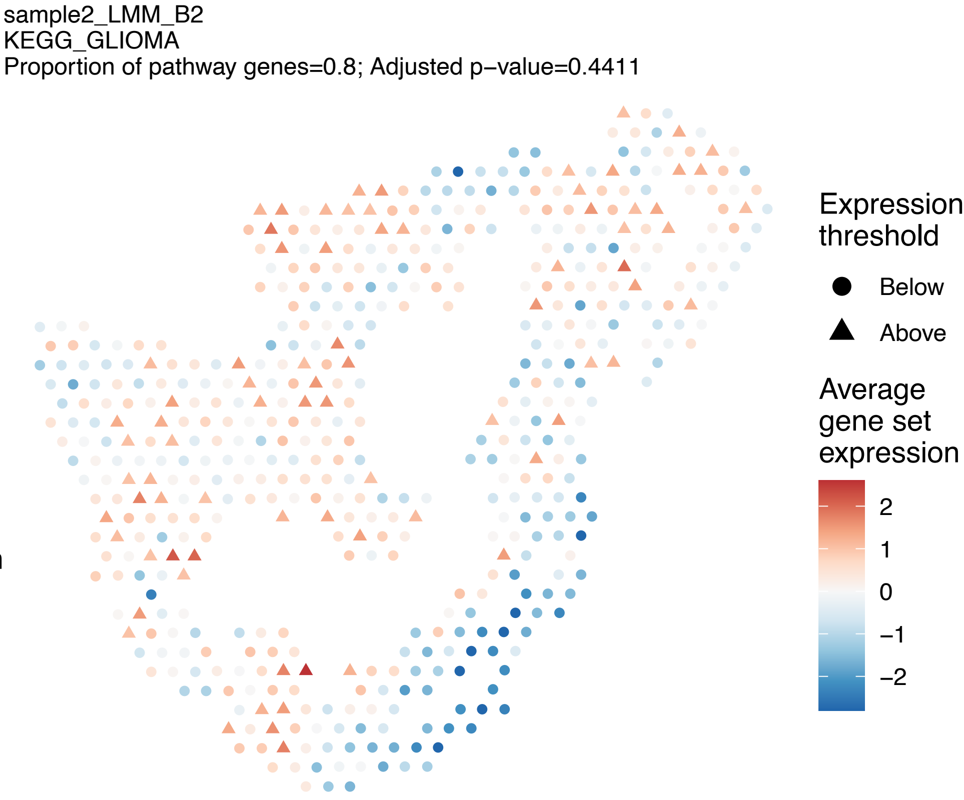
sample2_LMM_B2
KEGG_GLIOMA
Proportion of pathway genes=0.8; Adjusted p−value=0.4411
Expression threshold
Below
Above
Average gene set expression
2
1
0
−1
−2

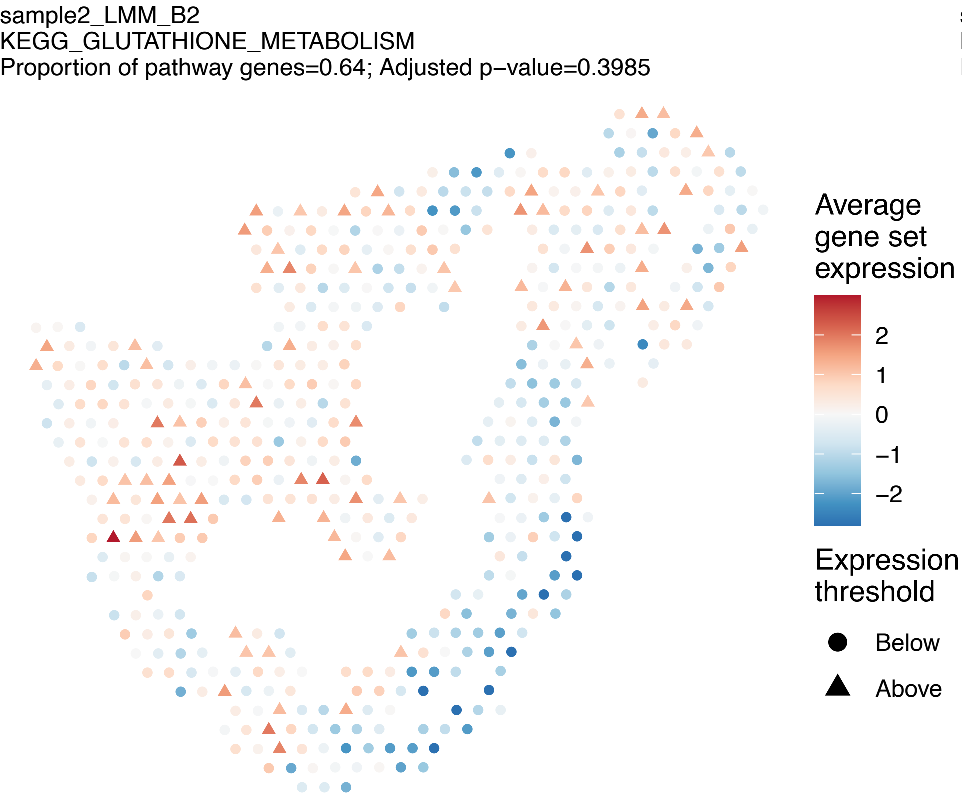
sample2_LMM_B2
KEGG_GLUTATHIONE_METABOLISM
Proportion of pathway genes=0.64; Adjusted p−value=0.3985
Average gene set expression
2
1
0
−1
−2
Expression threshold
Below
Above

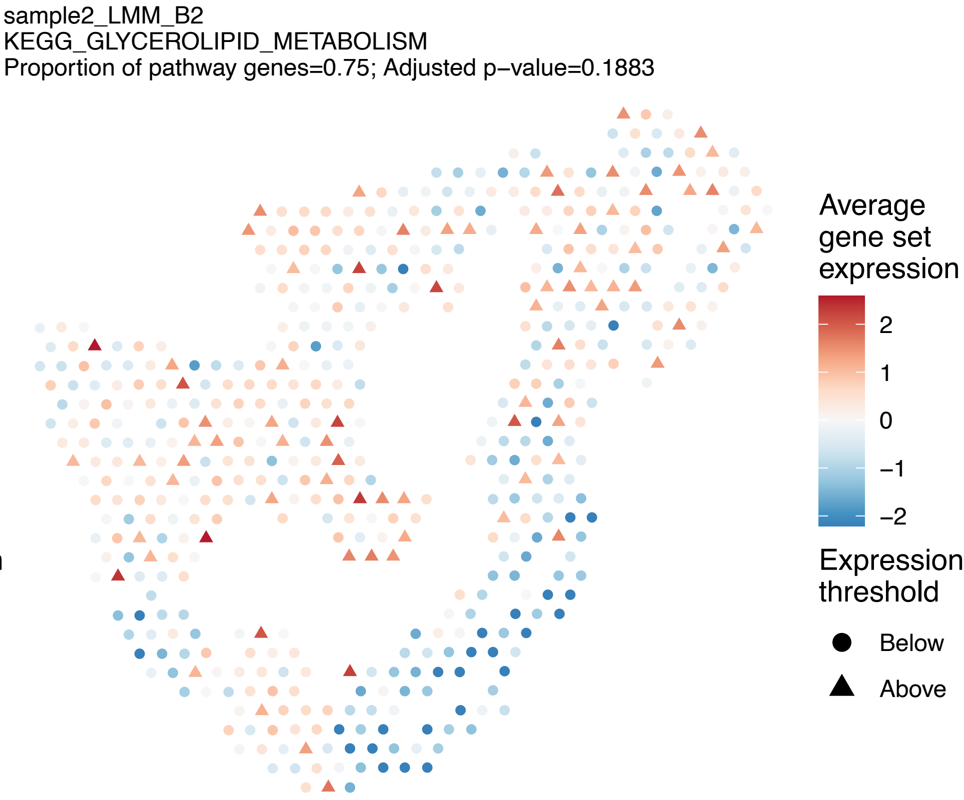
sample2_LMM_B2
KEGG_GLYCEROLIPID_METABOLISM
Proportion of pathway genes=0.75; Adjusted p−value=0.1883
Average gene set expression
2
1
0
−1
−2
Expression threshold
Below
Above

sample2_LMM_B2
KEGG_GLYCEROPHOSPHOLIPID_METABOLISM
Proportion of pathway genes=0.74; Adjusted p−value=0.3371

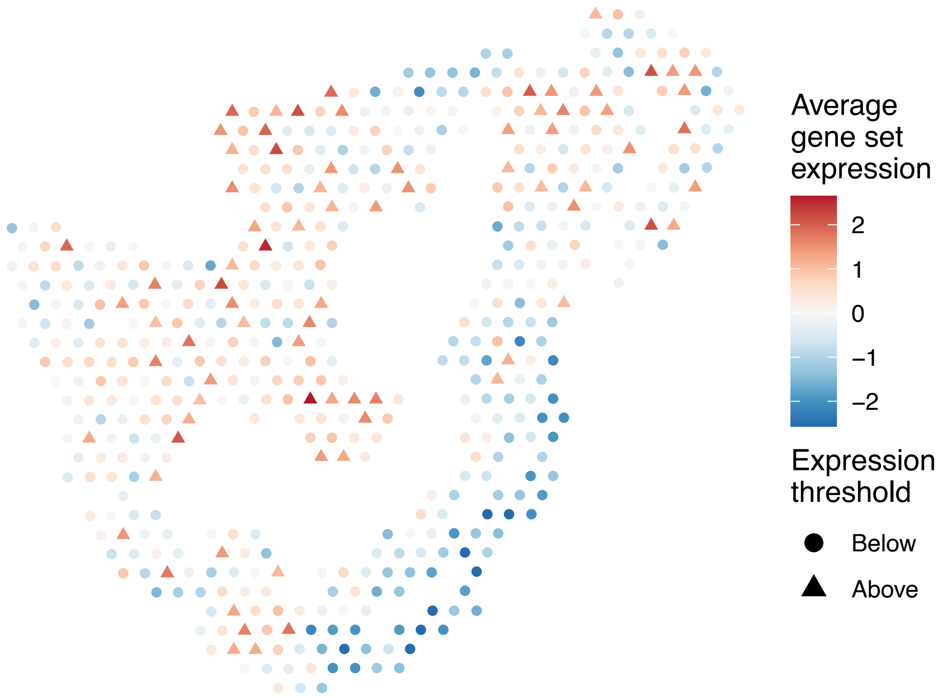


sample2_LMM_B2
KEGG_GLYCINE_SERINE_AND_THREONINE_METABOLISM
Proportion of pathway genes=0.53; Adjusted p−value=0.174

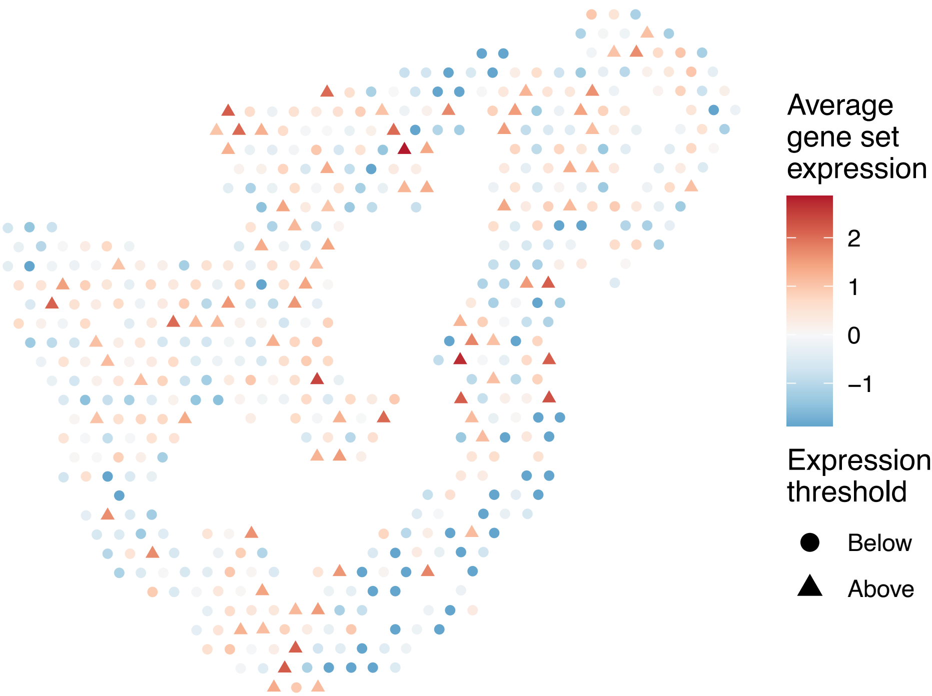


sample2_LMM_B2
KEGG_GLYCOLYSIS_GLUCONEOGENESIS
Proportion of pathway genes=0.63; Adjusted p−value=0.986

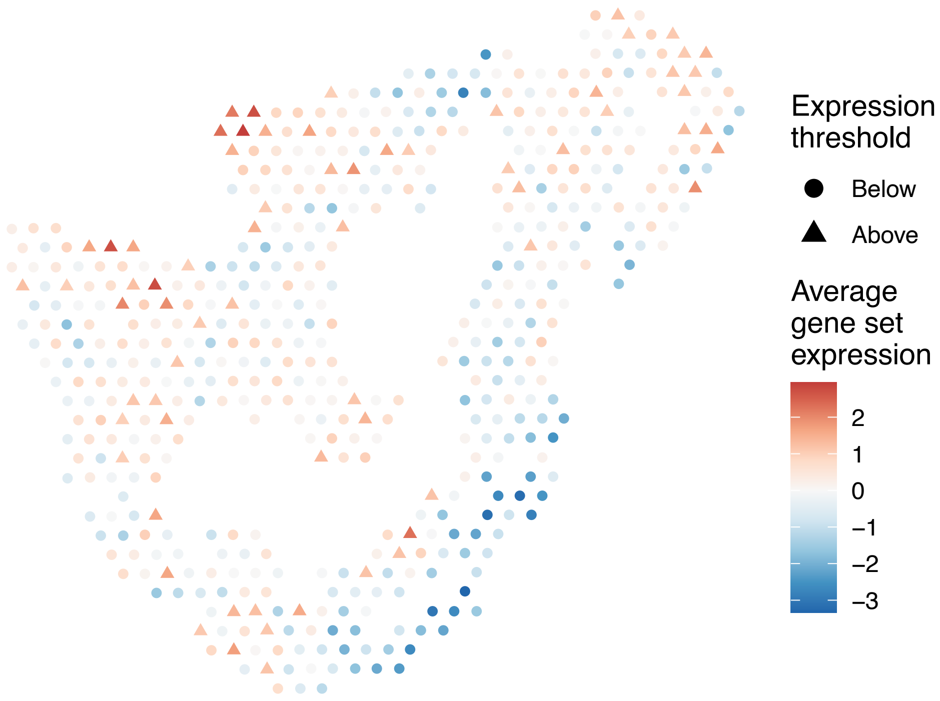


sample2_LMM_B2
KEGG_GLYCOSAMINOGLYCAN_BIOSYNTHESIS_CHONDROITIN_SU
LFATE
Proportion of pathway genes=0.83; Adjusted p−value=0.0946

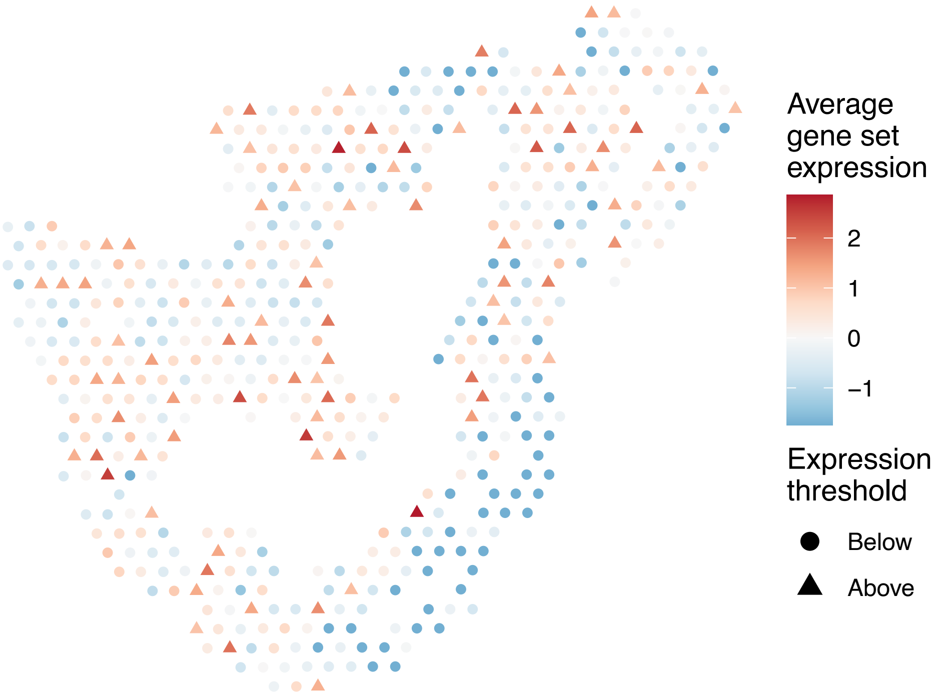

sample2_LMM_B2
KEGG_GLYCOSAMINOGLYCAN_BIOSYNTHESIS_HEPARAN_SULFAT
E
Proportion of pathway genes=0.63; Adjusted p–value=0.1169

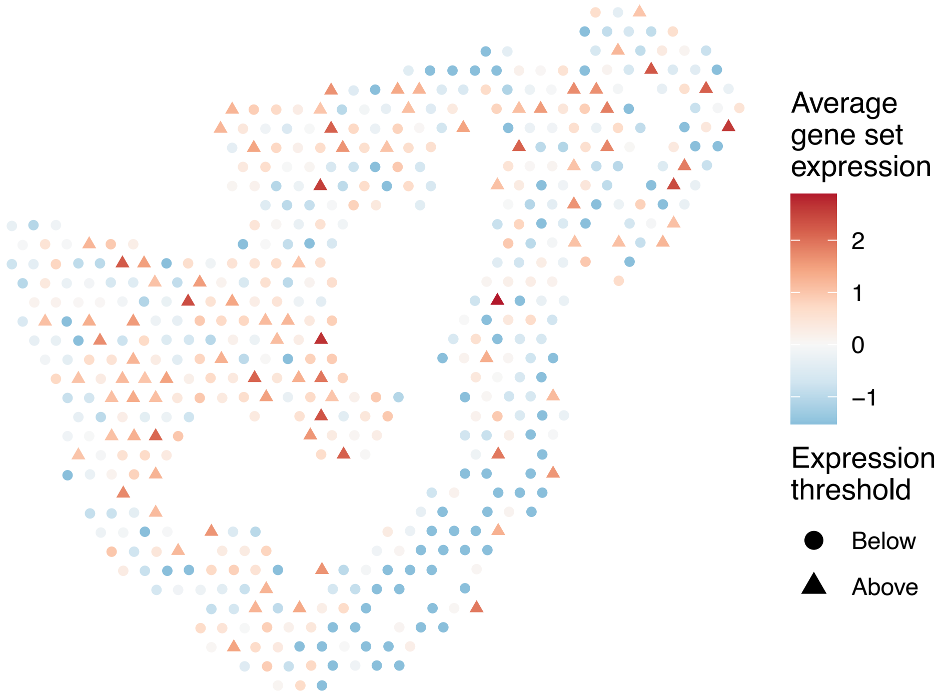


sample2_LMM_B2
KEGG_GLYCOSAMINOGLYCAN_BIOSYNTHESIS_KERATAN_SULFAT
E
Proportion of pathway genes=0.93; Adjusted p–value=0

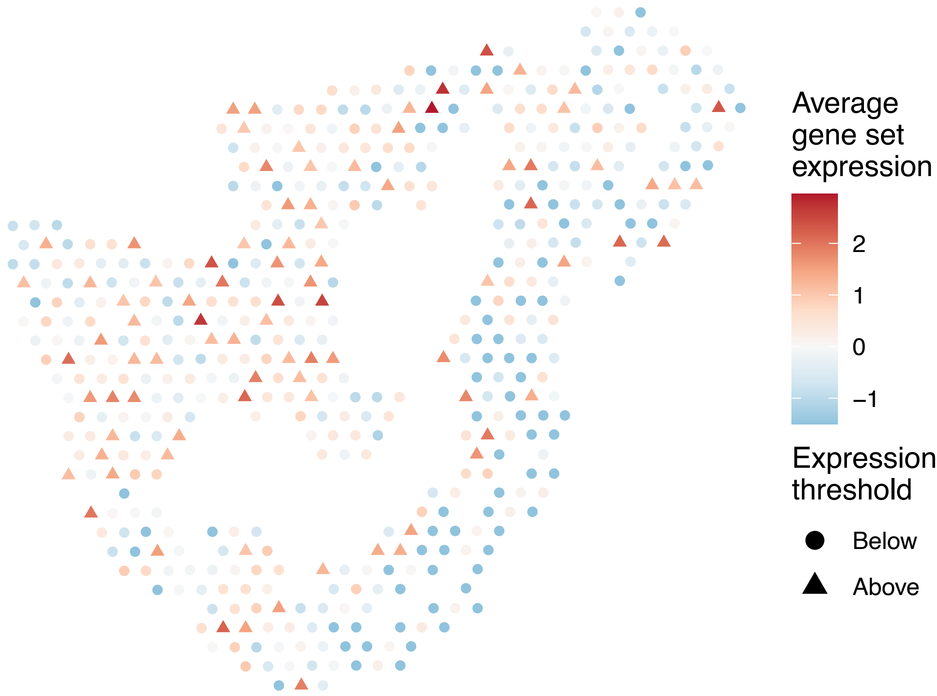


sample2_LMM_B2
KEGG_GLYCOSAMINOGLYCAN_DEGRADATION
Proportion of pathway genes=0.67; Adjusted p–value=0.0696

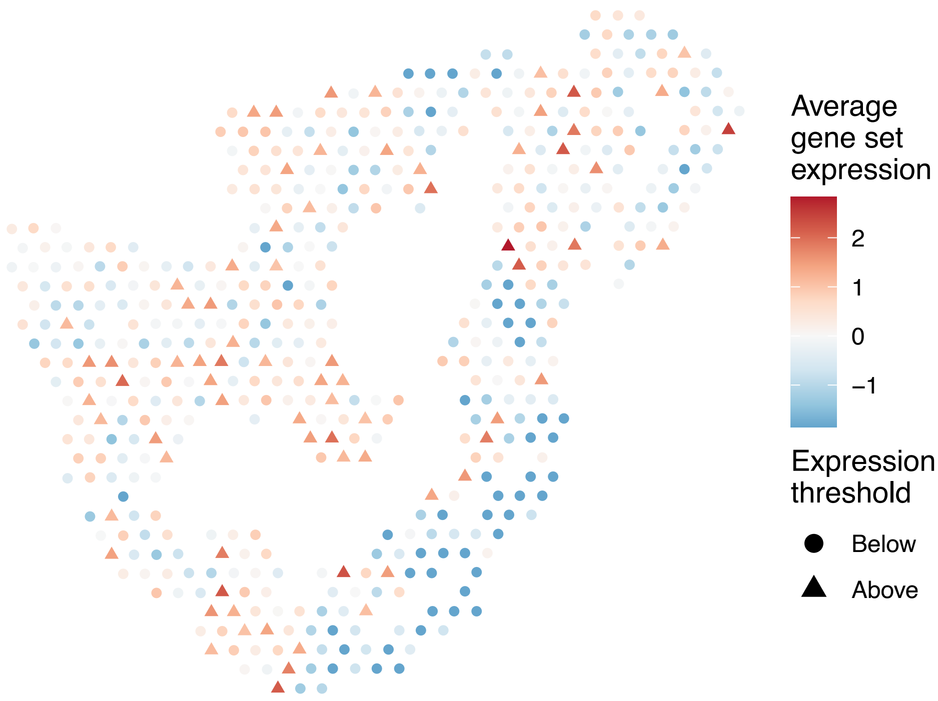


sample2_LMM_B2
KEGG_GLYCOSPHINGOLIPID_BIOSYNTHESIS_GANGLIO_SERIES
Proportion of pathway genes=0.79; Adjusted p–value=0.0514

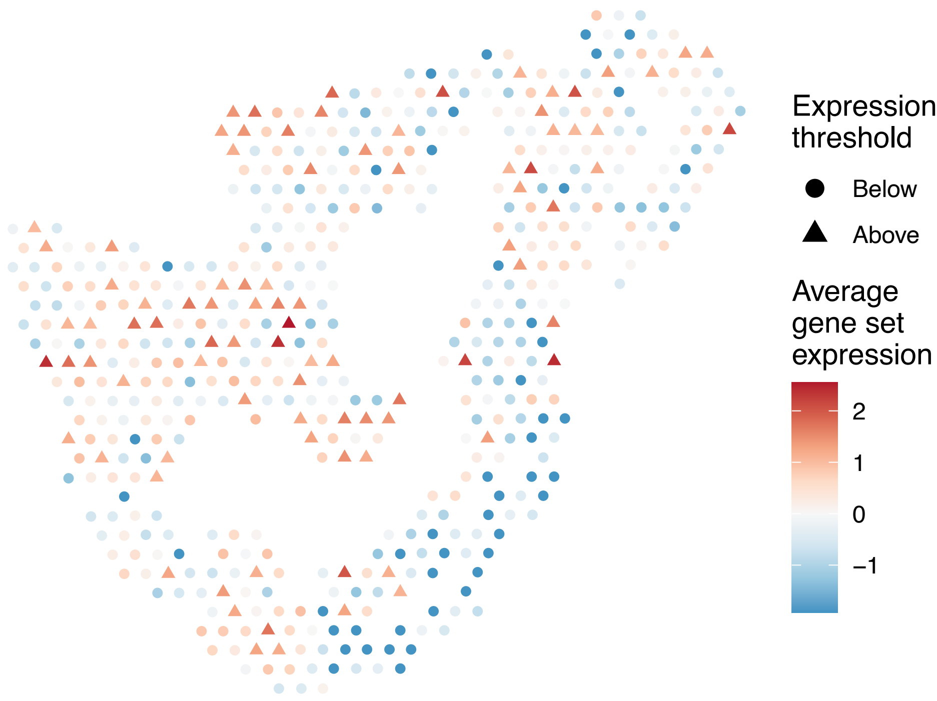

sample2_LMM_B2
KEGG_GLYCOSPHINGOLIPID_BIOSYNTHESIS_GLOBO_SERIES
Proportion of pathway genes=0.57; Adjusted p−value=NA

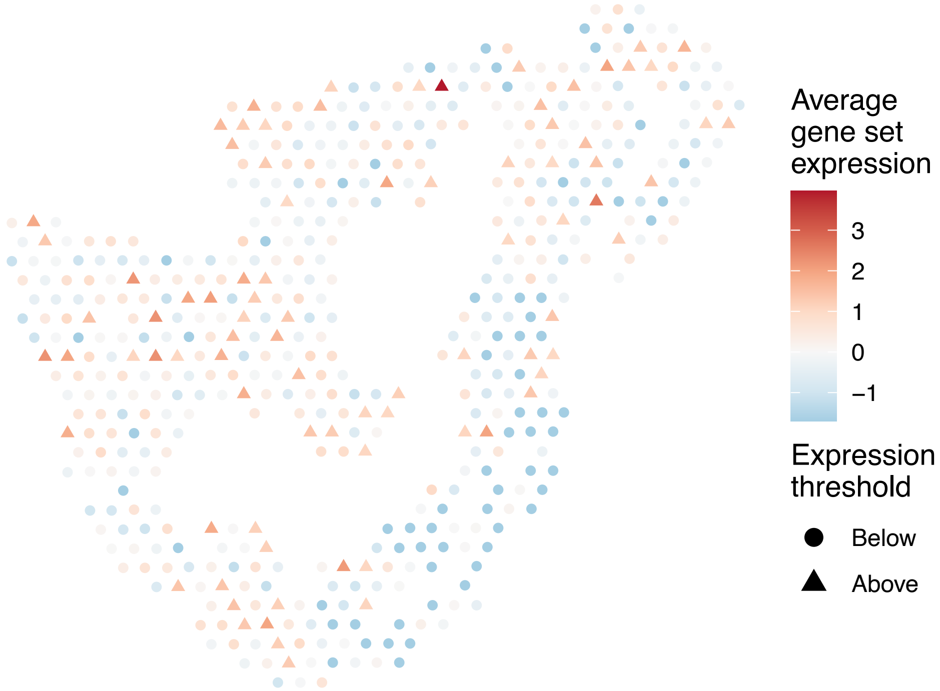


sample2_LMM_B2
KEGG_GLYCOSPHINGOLIPID_BIOSYNTHESIS_LACTO_AND_NEOL
ACTO_SERIES
Proportion of pathway genes=0.45; Adjusted p−value=0.1169

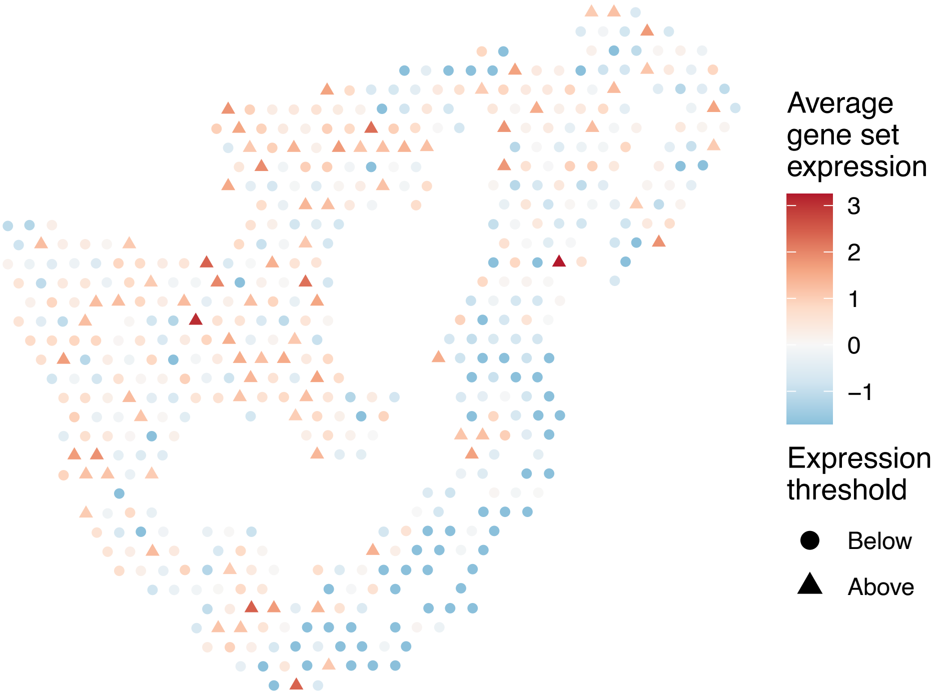


sample2_LMM_B2
KEGG_GLYCOSYLPHOSPHATIDYLINOSITOL_GPI_ANCHOR_BIOSY
NTHESIS
Proportion of pathway genes=0.93; Adjusted p−value=0.59

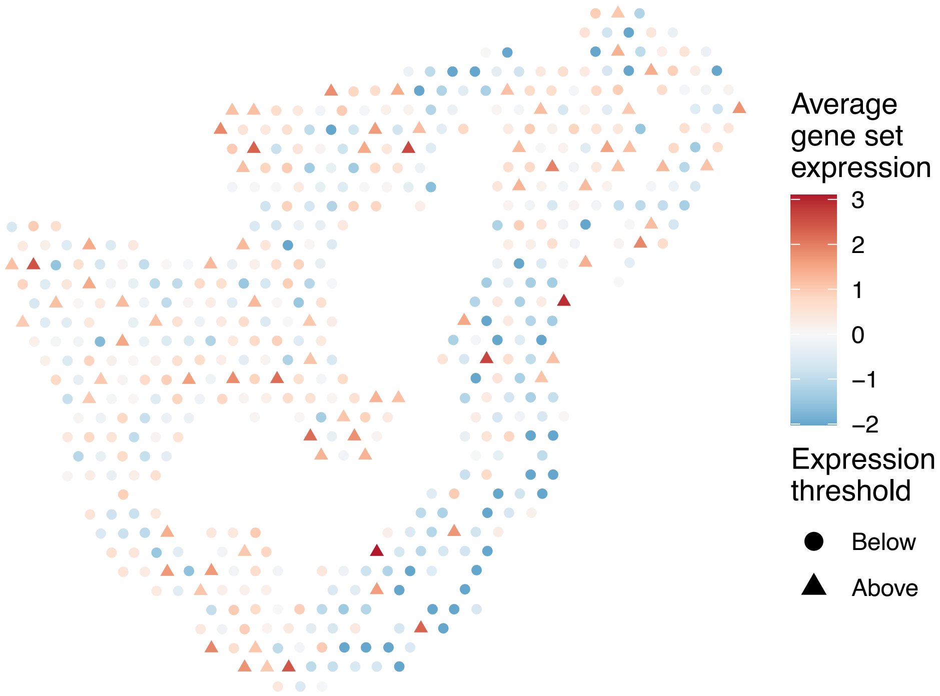


sample2_LMM_B2
KEGG_GLYOXYLATE_AND_DICARBOXYLATE_METABOLISM
Proportion of pathway genes=0.88; Adjusted p−value=0

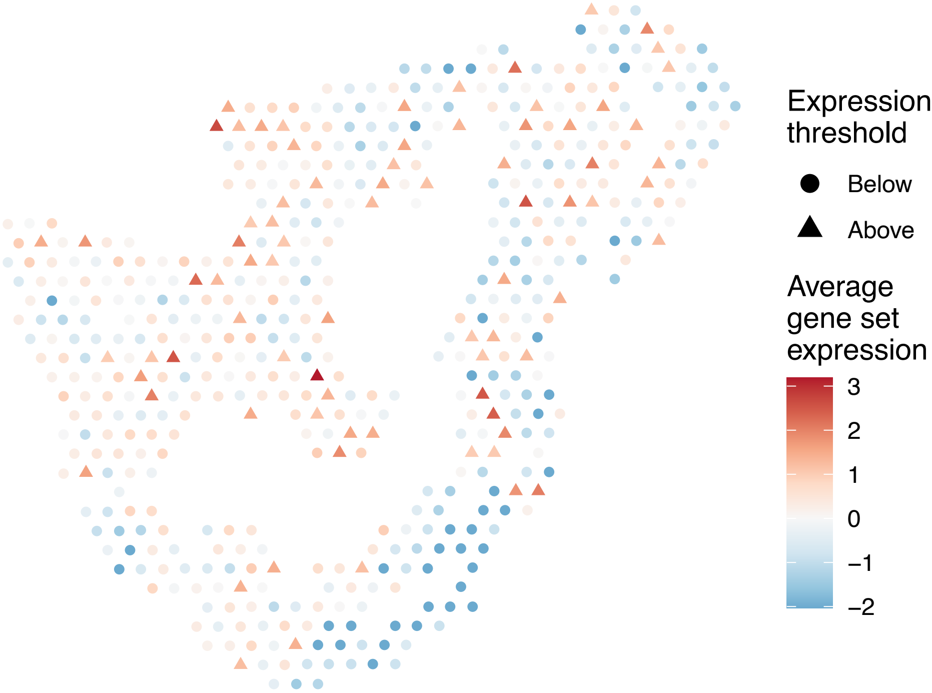

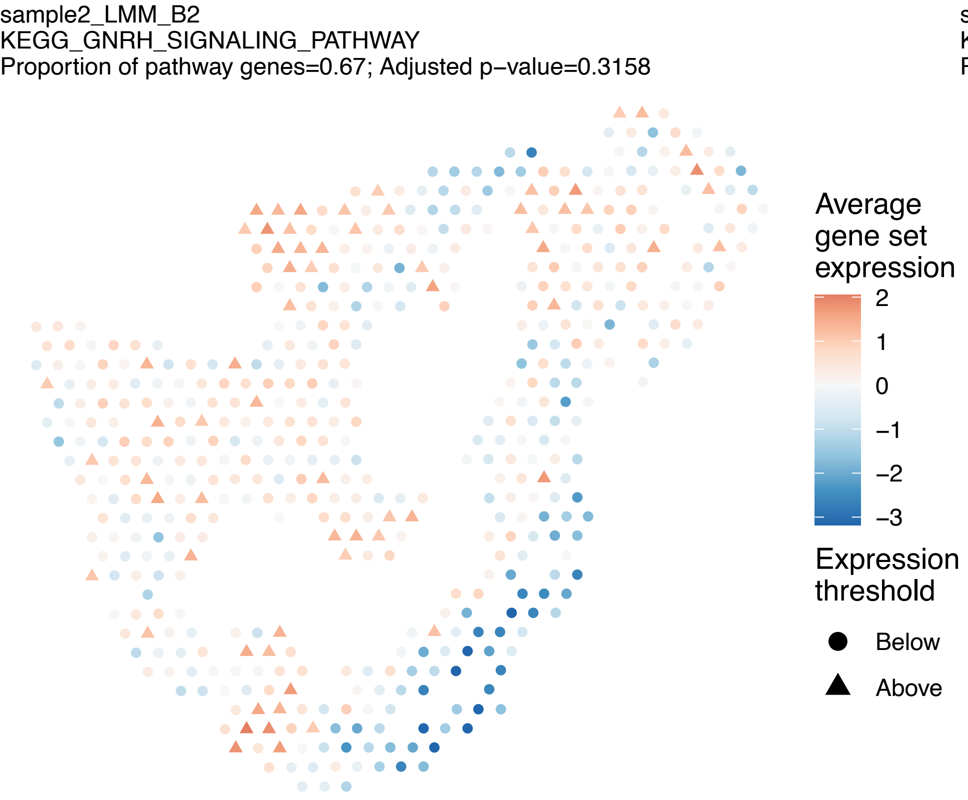
sample2_LMM_B2
KEGG_GNRH_SIGNALING_PATHWAY
Proportion of pathway genes=0.67; Adjusted p−value=0.3158
Average gene set expression
2
1
0
−1
−2
−3
Expression threshold
Below
Above

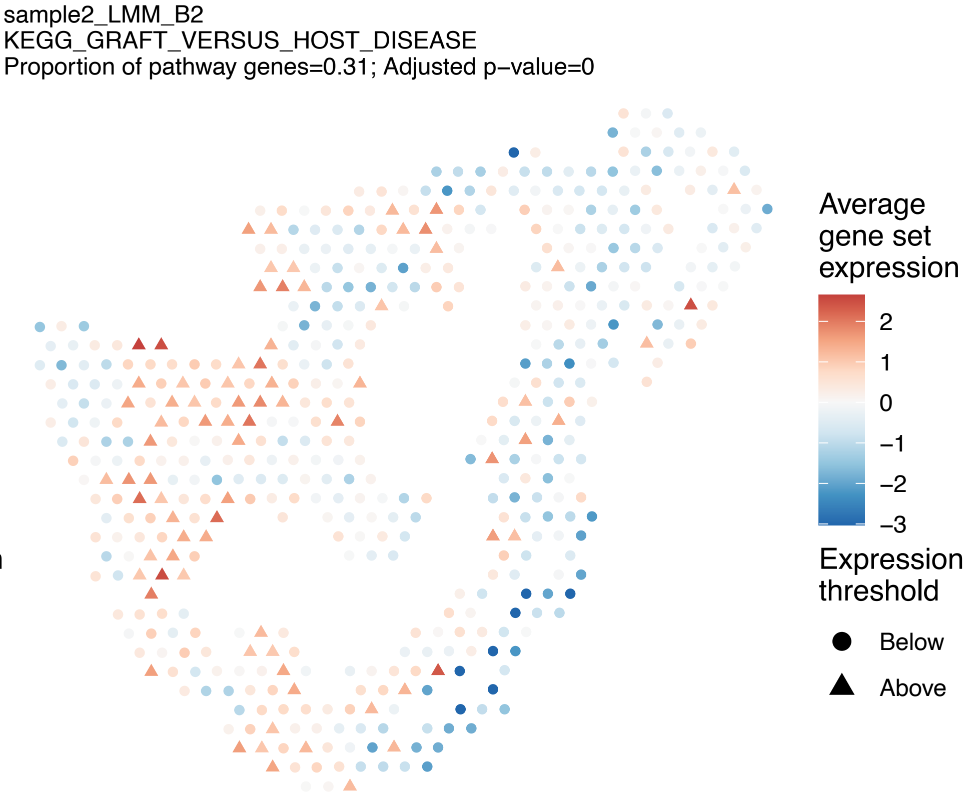
sample2_LMM_B2
KEGG_GRAFT_VERSUS_HOST_DISEASE
Proportion of pathway genes=0.31; Adjusted p−value=0
Average gene set expression
2
1
0
−1
−2
−3
Expression threshold
Below
Above

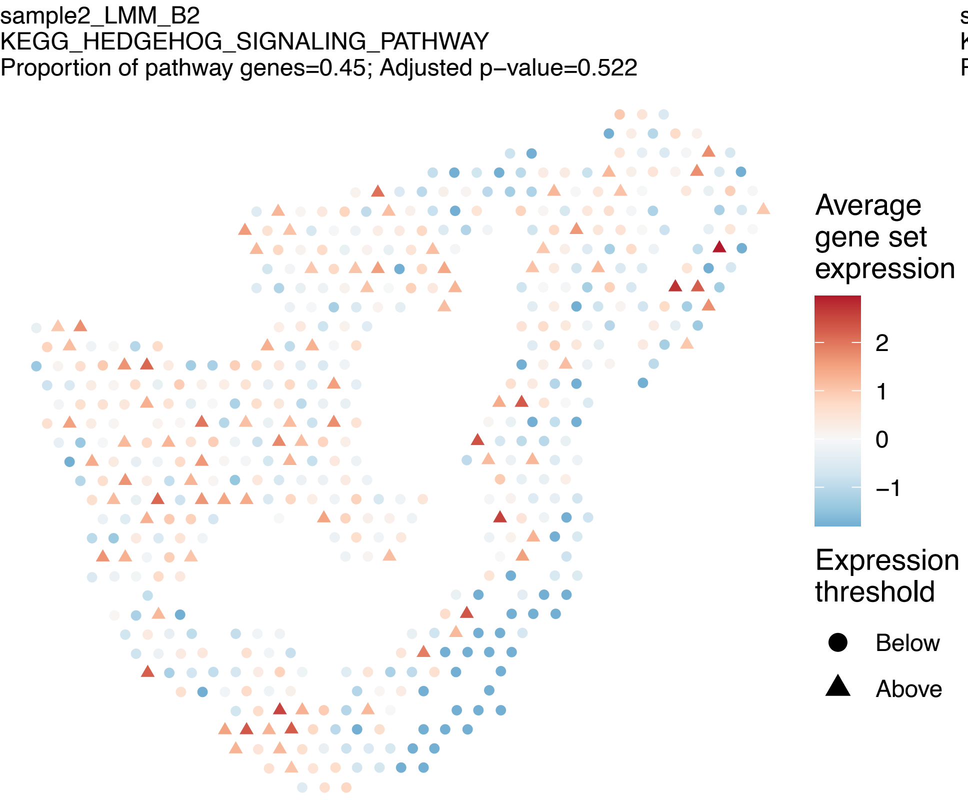
sample2_LMM_B2
KEGG_HEDGEHOG_SIGNALING_PATHWAY
Proportion of pathway genes=0.45; Adjusted p−value=0.522
Average gene set expression
2
1
0
−1
Expression threshold
Below
Above

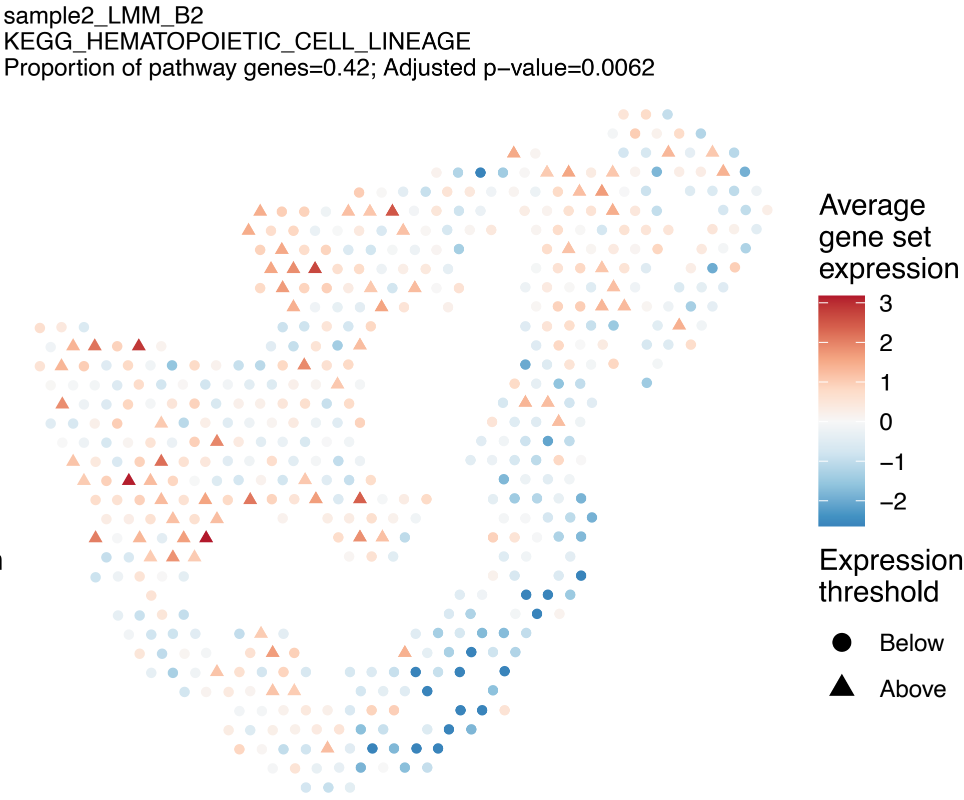
sample2_LMM_B2
KEGG_HEMATOPOIETIC_CELL_LINEAGE
Proportion of pathway genes=0.42; Adjusted p−value=0.0062
Average gene set expression
3
2
1
0
−1
−2
Expression threshold
Below
Above

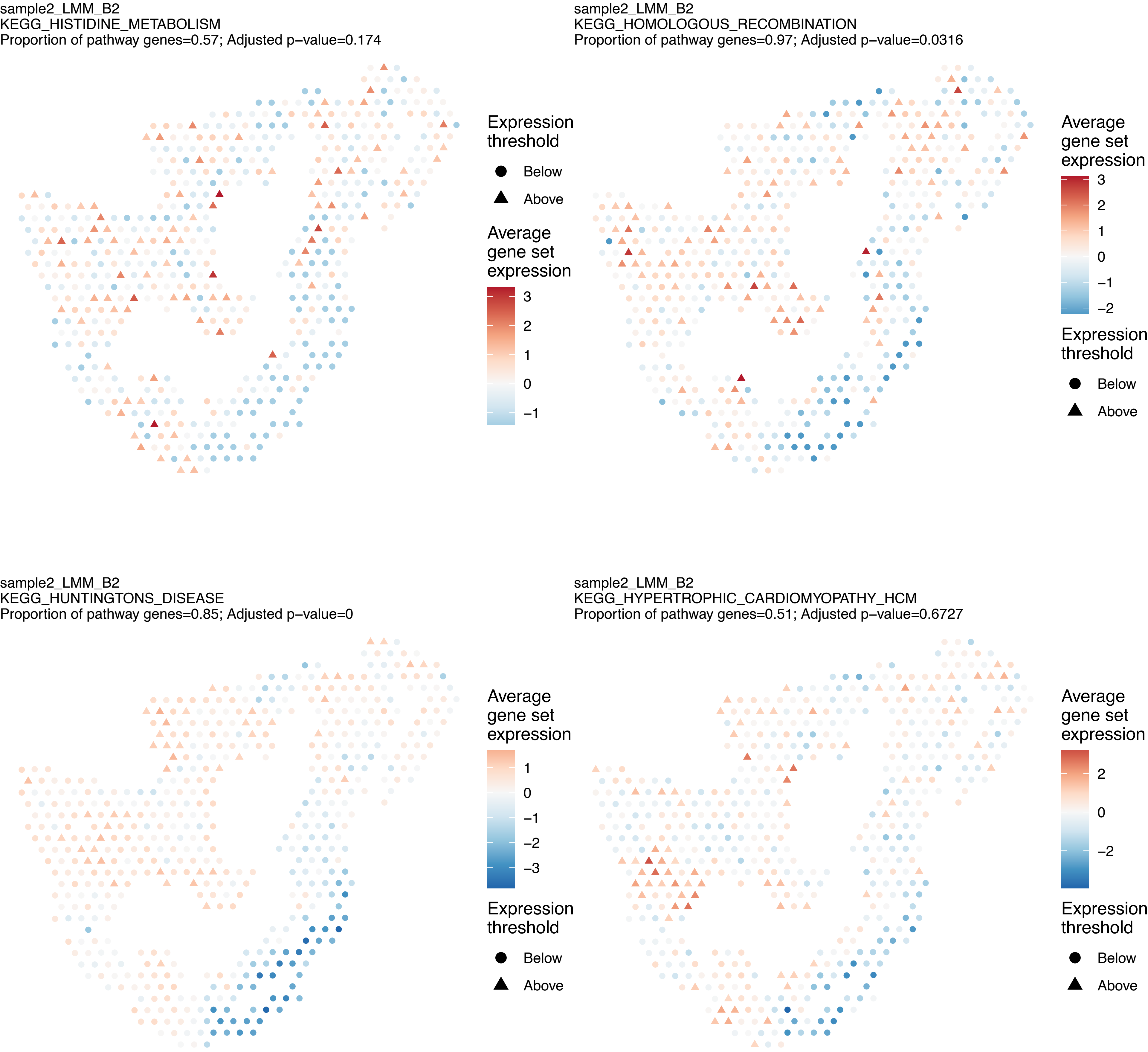
sample2_LMM_B2
KEGG_HISTIDINE_METABOLISM
Proportion of pathway genes=0.57; Adjusted p−value=0.174
Expression threshold
Below
Above
Average gene set expression
3
2
1
0
−1
sample2_LMM_B2
KEGG_HOMOLOGOUS_RECOMBINATION
Proportion of pathway genes=0.97; Adjusted p−value=0.0316
Average gene set expression
3
2
1
0
−1
−2
Expression threshold
Below
Above
sample2_LMM_B2
KEGG_HUNTINGTONS_DISEASE
Proportion of pathway genes=0.85; Adjusted p−value=0
Average gene set expression
1
0
−1
−2
−3
Expression threshold
Below
Above
sample2_LMM_B2
KEGG_HYPERTROPHIC_CARDIOMYOPATHY_HCM
Proportion of pathway genes=0.51; Adjusted p−value=0.6727
Average gene set expression
2
0
−2
Expression threshold
Below
Above

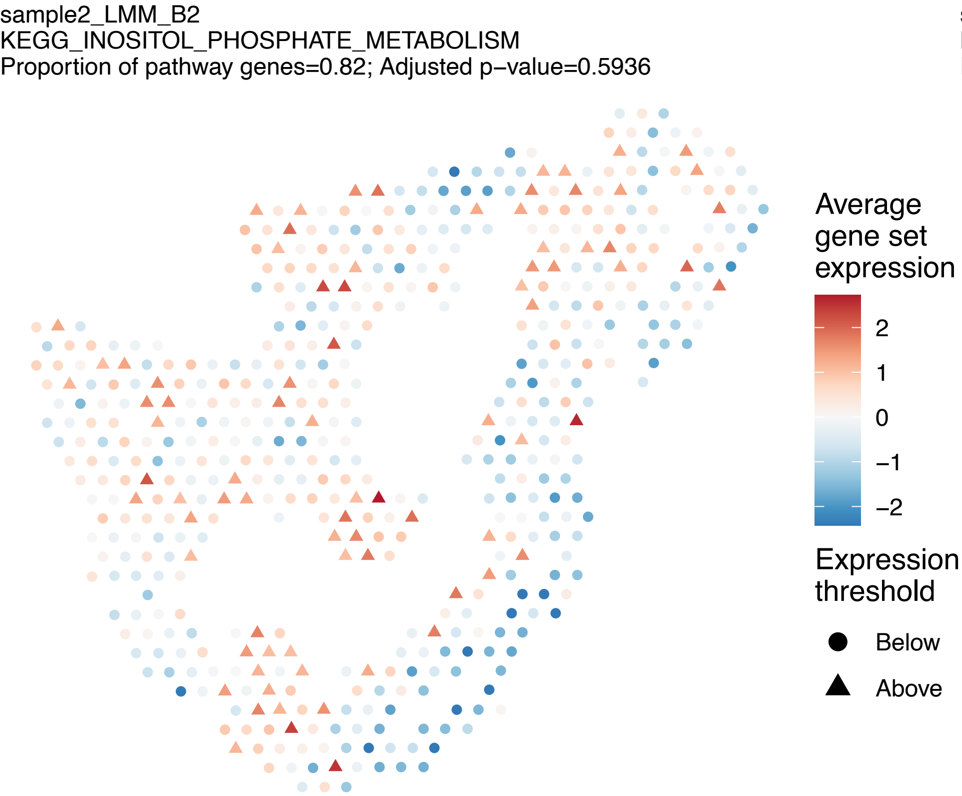
sample2_LMM_B2
KEGG_INOSITOL_PHOSPHATE_METABOLISM
Proportion of pathway genes=0.82; Adjusted p−value=0.5936
Average gene set expression
2
1
0
−1
−2
Expression threshold
Below
Above

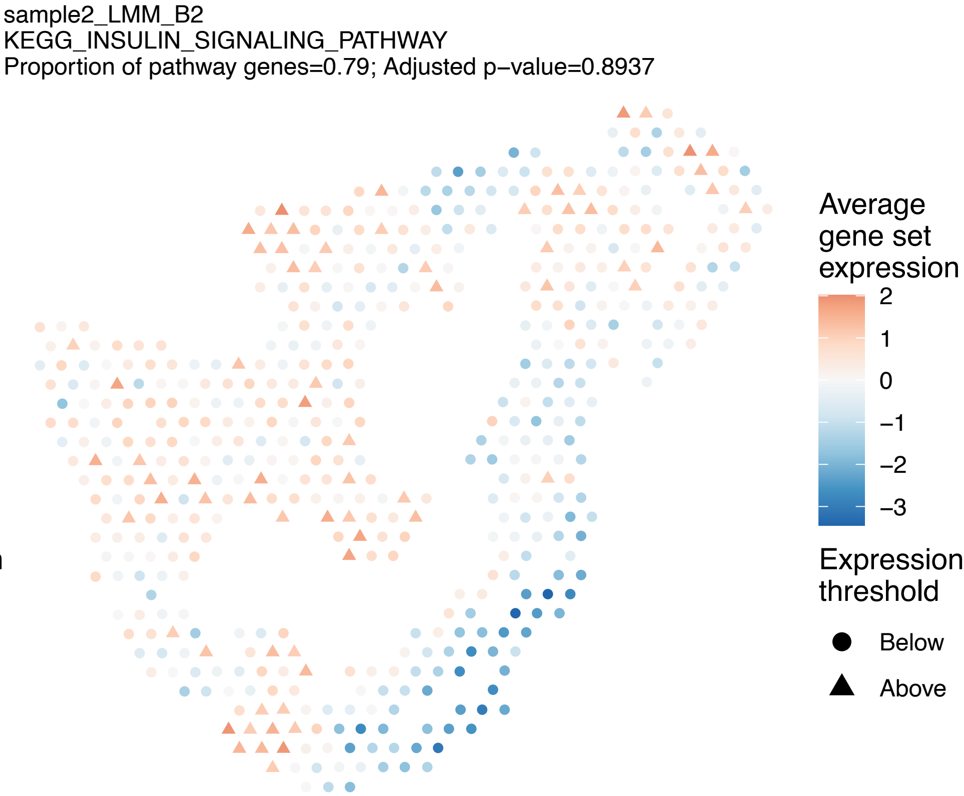
sample2_LMM_B2
KEGG_INSULIN_SIGNALING_PATHWAY
Proportion of pathway genes=0.79; Adjusted p−value=0.8937
Average gene set expression
2
1
0
−1
−2
−3
Expression threshold
Below
Above

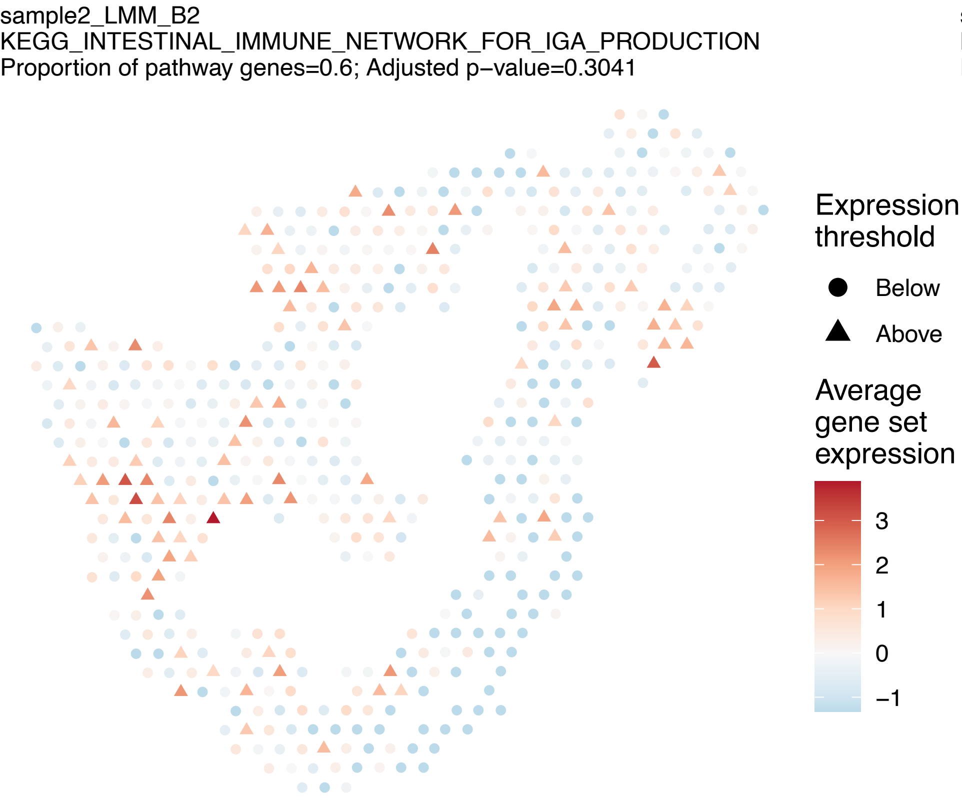
sample2_LMM_B2
KEGG_INTESTINAL_IMMUNE_NETWORK_FOR_IGA_PRODUCTION
Proportion of pathway genes=0.6; Adjusted p−value=0.3041
Expression threshold
Below
Above
Average gene set expression
3
2
1
0
−1

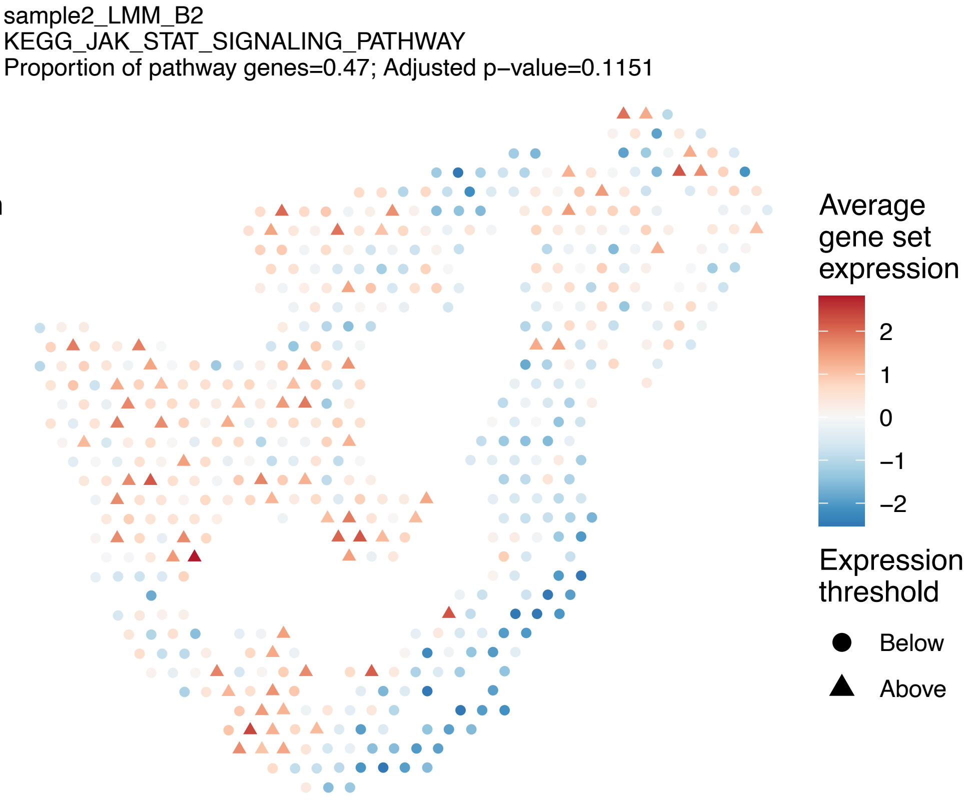
sample2_LMM_B2
KEGG_JAK_STAT_SIGNALING_PATHWAY
Proportion of pathway genes=0.47; Adjusted p−value=0.1151
Average gene set expression
2
1
0
−1
−2
Expression threshold
Below
Above

sample2_LMM_B2
KEGG_LEISHMANIA_INFECTION
Proportion of pathway genes=0.65; Adjusted p−value=0.2397

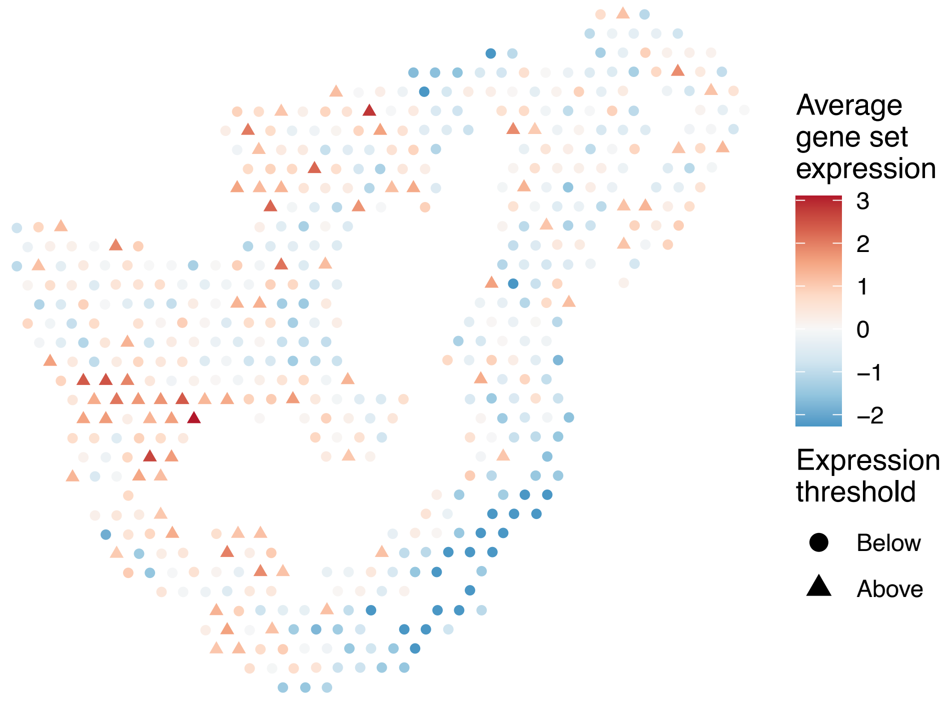


sample2_LMM_B2
KEGG_LEUKOCYTE_TRANSENDOTHELIAL_MIGRATION
Proportion of pathway genes=0.65; Adjusted p−value=0

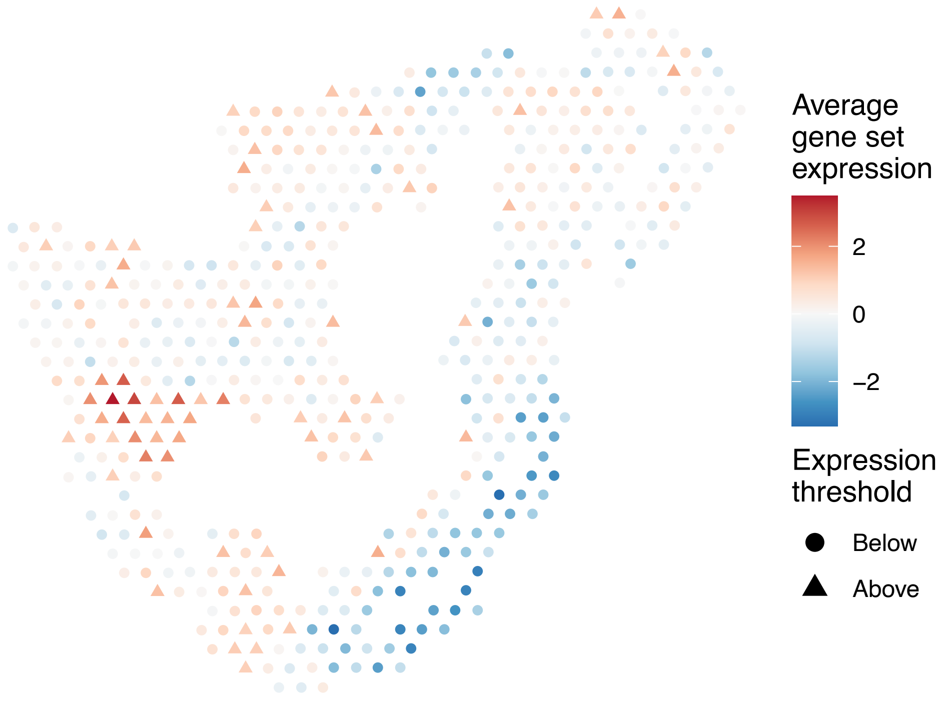


sample2_LMM_B2
KEGG_LIMONENE_AND_PINENE_DEGRADATION
Proportion of pathway genes=0.9; Adjusted p−value=NA

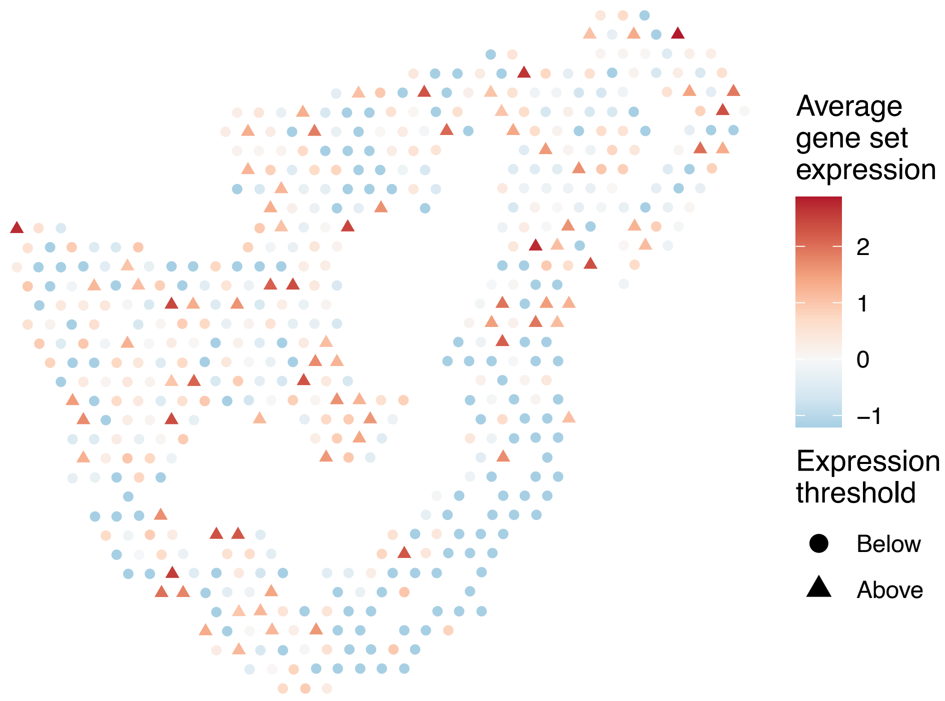


sample2_LMM_B2
KEGG_LONG_TERM_DEPRESSION
Proportion of pathway genes=0.55; Adjusted p−value=0.1864

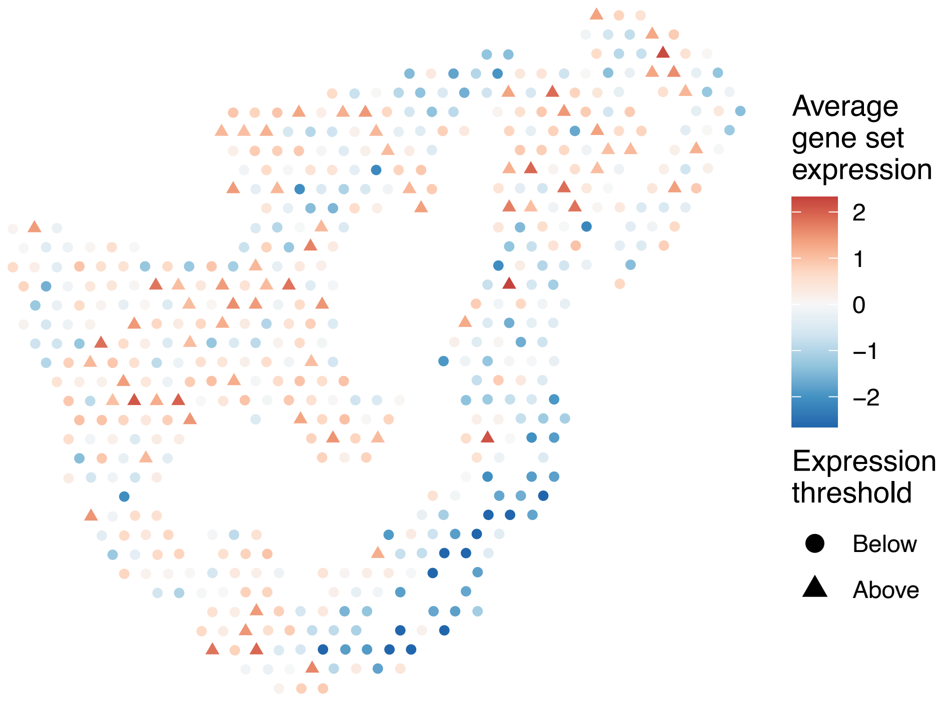

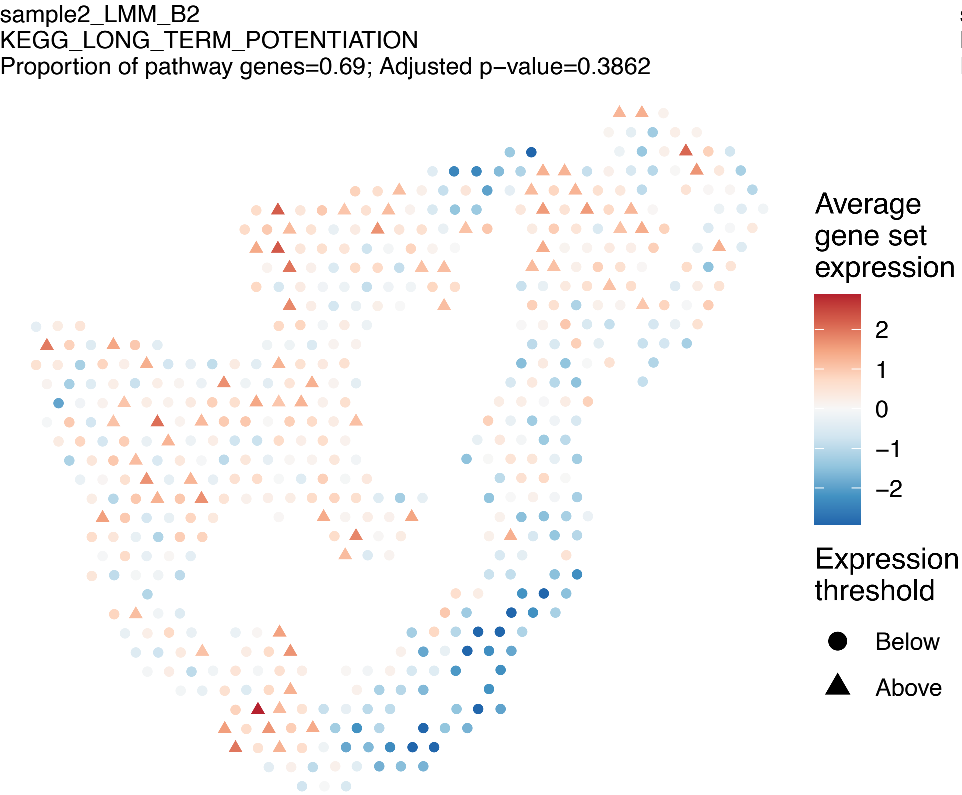
sample2_LMM_B2
KEGG_LONG_TERM_POTENTIATION
Proportion of pathway genes=0.69; Adjusted p−value=0.3862
Average gene set expression
2
1
0
−1
−2
Expression threshold
Below
Above

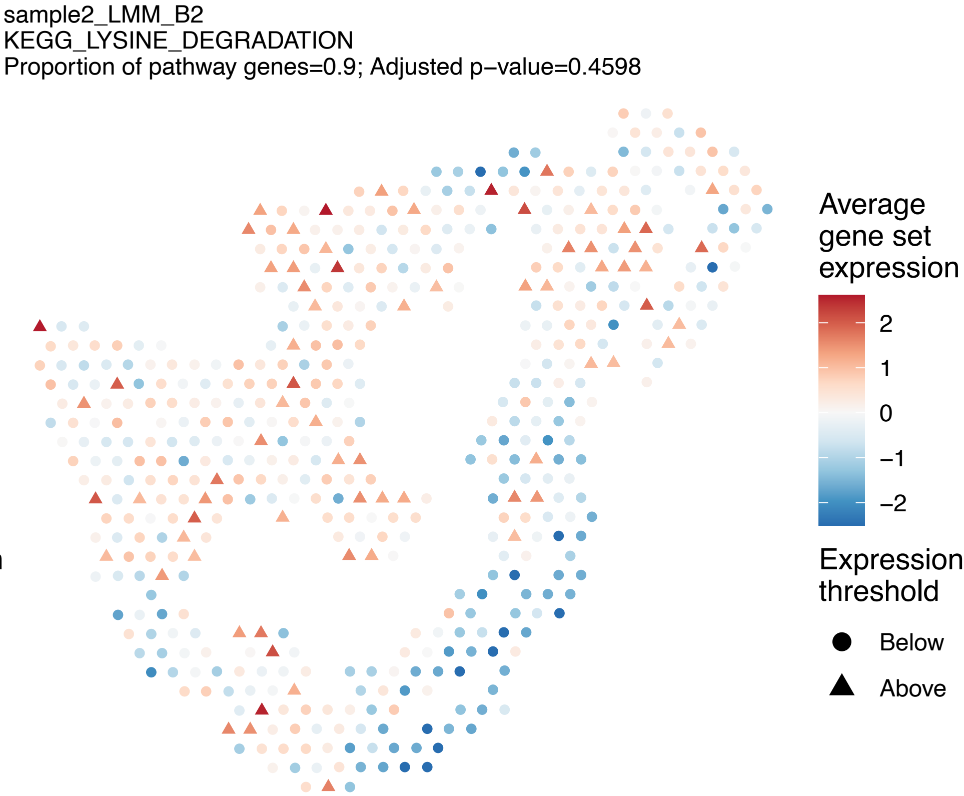
sample2_LMM_B2
KEGG_LYSINE_DEGRADATION
Proportion of pathway genes=0.9; Adjusted p−value=0.4598
Average gene set expression
2
1
0
−1
−2
Expression threshold
Below
Above

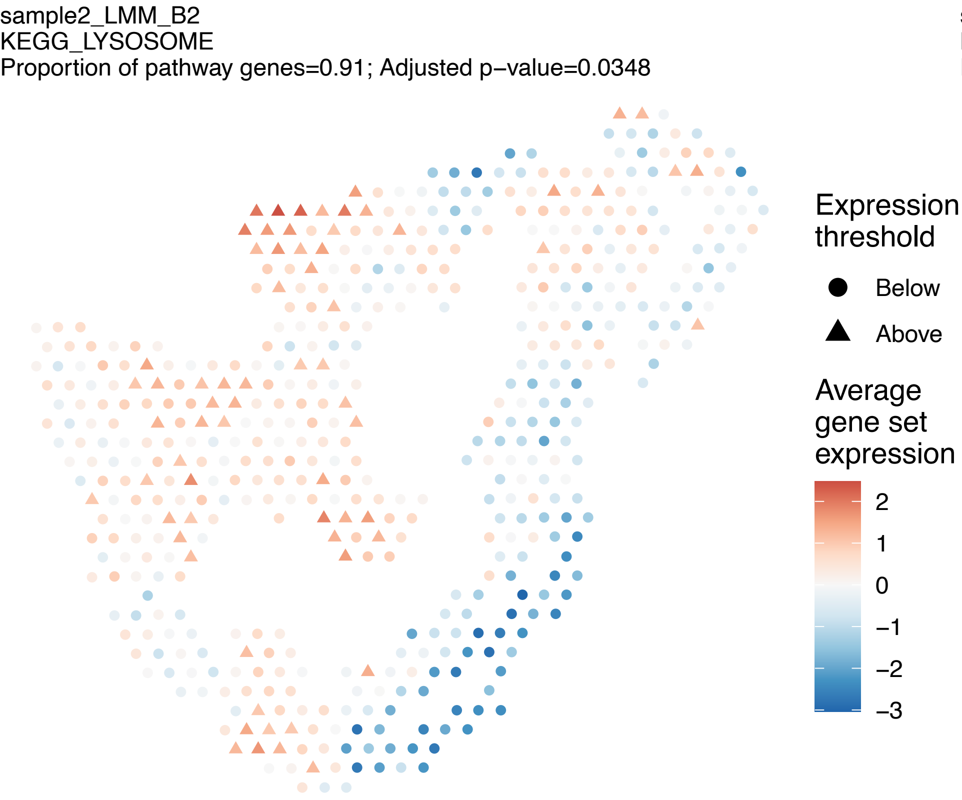
sample2_LMM_B2
KEGG_LYSOSOME
Proportion of pathway genes=0.91; Adjusted p−value=0.0348
Expression threshold
Below
Above
Average gene set expression
2
1
0
−1
−2
−3

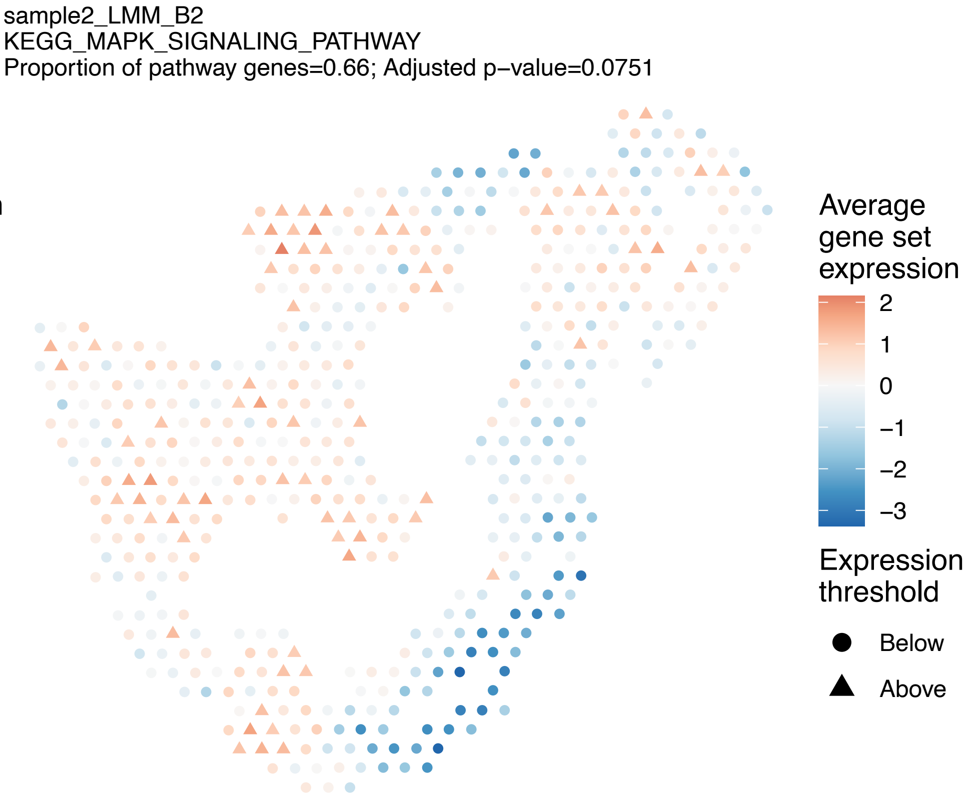
sample2_LMM_B2
KEGG_MAPK_SIGNALING_PATHWAY
Proportion of pathway genes=0.66; Adjusted p−value=0.0751
Average gene set expression
2
1
0
−1
−2
−3
Expression threshold
Below
Above

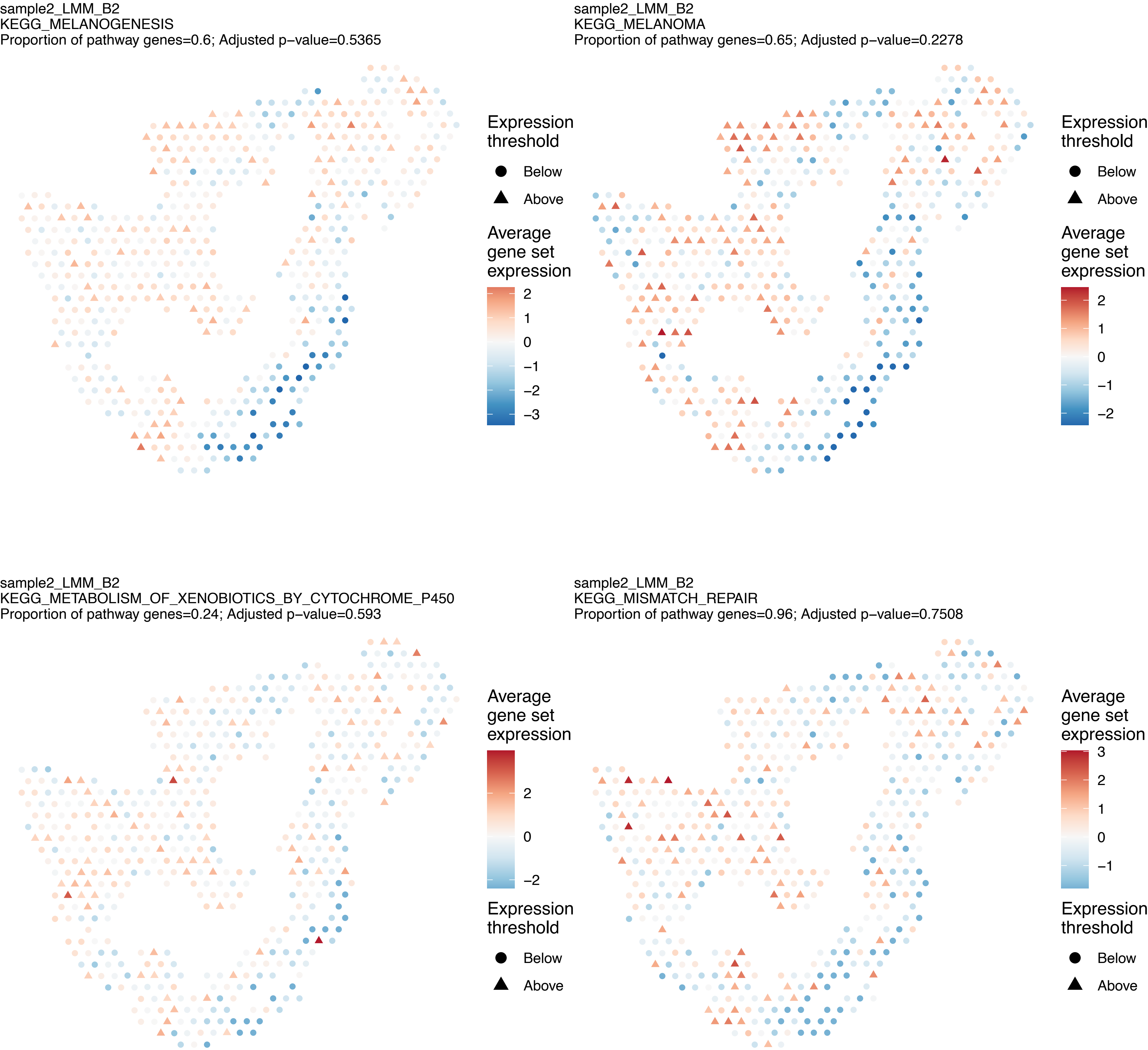

sample2_LMM_B2
KEGG_MELANOGENESIS
Proportion of pathway genes=0.6; Adjusted p−value=0.5365
Expression threshold
Below
Above
Average gene set expression
2
1
0
−1
−2
−3
sample2_LMM_B2
KEGG_MELANOMA
Proportion of pathway genes=0.65; Adjusted p−value=0.2278
Expression threshold
Below
Above
Average gene set expression
2
1
0
−1
−2
sample2_LMM_B2
KEGG_METABOLISM_OF_XENOBIOTICS_BY_CYTOCHROME_P450
Proportion of pathway genes=0.24; Adjusted p−value=0.593
Average gene set expression
2
0
−2
Expression threshold
Below
Above
sample2_LMM_B2
KEGG_MISMATCH_REPAIR
Proportion of pathway genes=0.96; Adjusted p−value=0.7508
Average gene set expression
3
2
1
0
−1
Expression threshold
Below
Above

sample2_LMM_B2
KEGG_MTOR_SIGNALING_PATHWAY
Proportion of pathway genes=0.83; Adjusted p−value=0.302

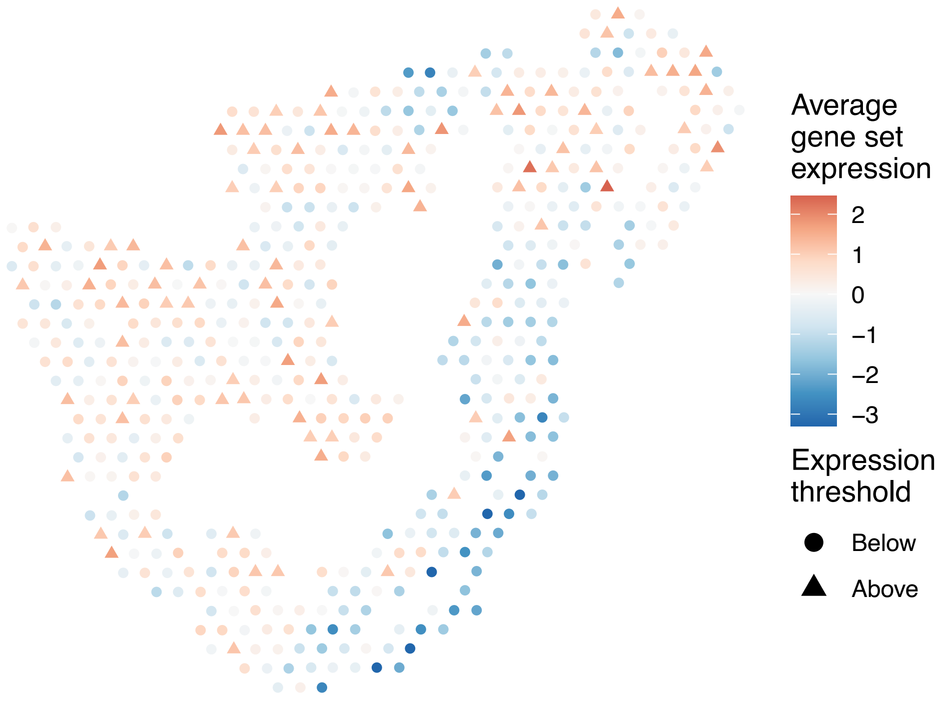


sample2_LMM_B2
KEGG_N_GLYCAN_BIOSYNTHESIS
Proportion of pathway genes=0.91; Adjusted p−value=0

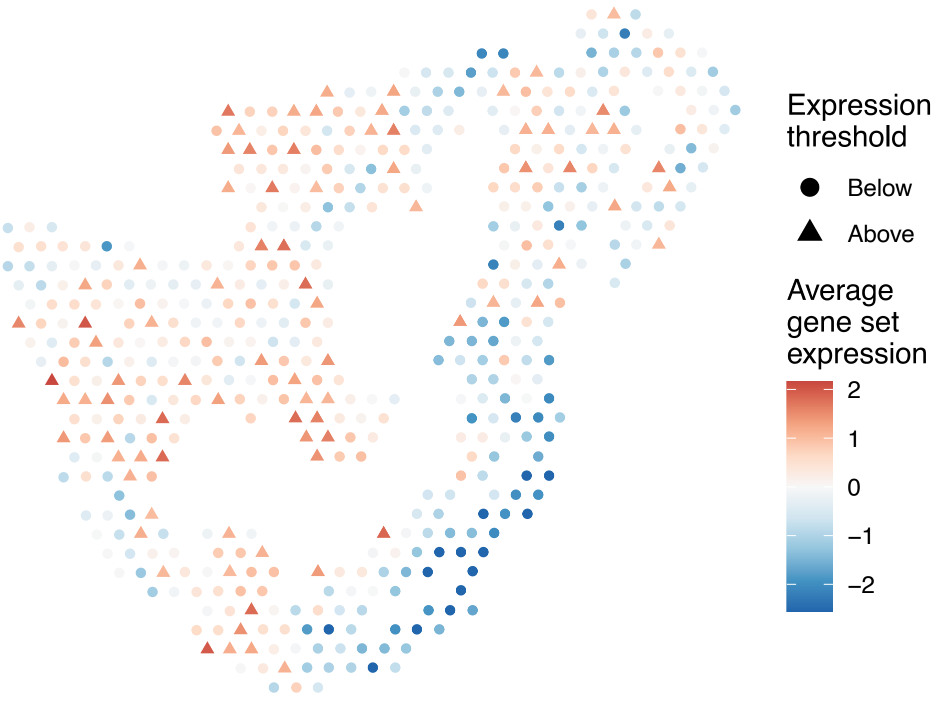


sample2_LMM_B2
KEGG_NEUROACTIVE_LIGAND_RECEPTOR_INTERACTION
Proportion of pathway genes=0.19; Adjusted p−value=0.1972

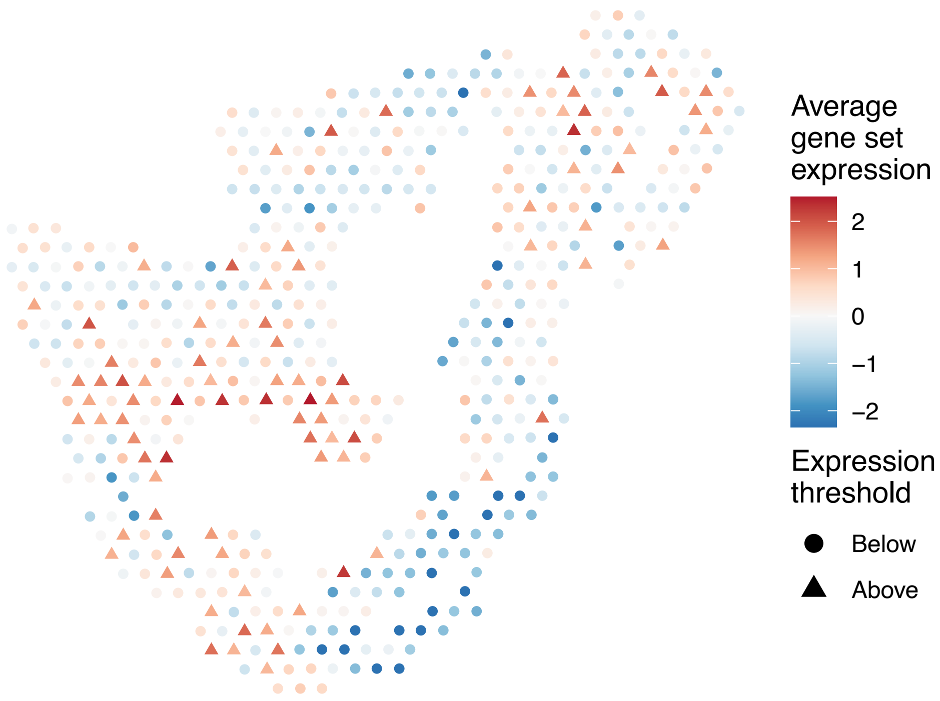


sample2_LMM_B2
KEGG_NEUROTROPHIN_SIGNALING_PATHWAY
Proportion of pathway genes=0.83; Adjusted p−value=0.4598

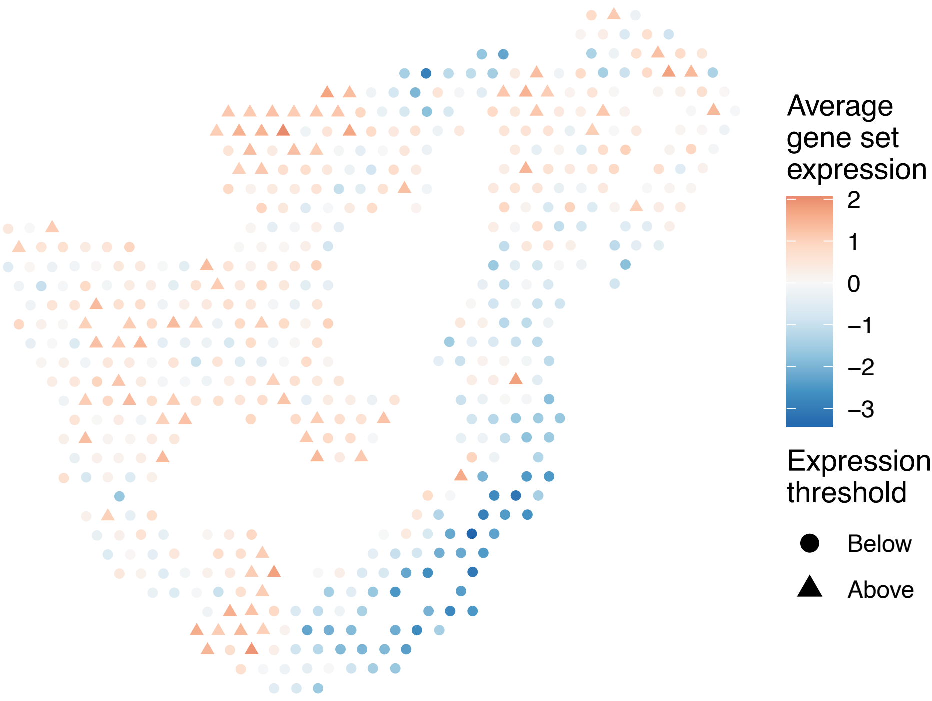

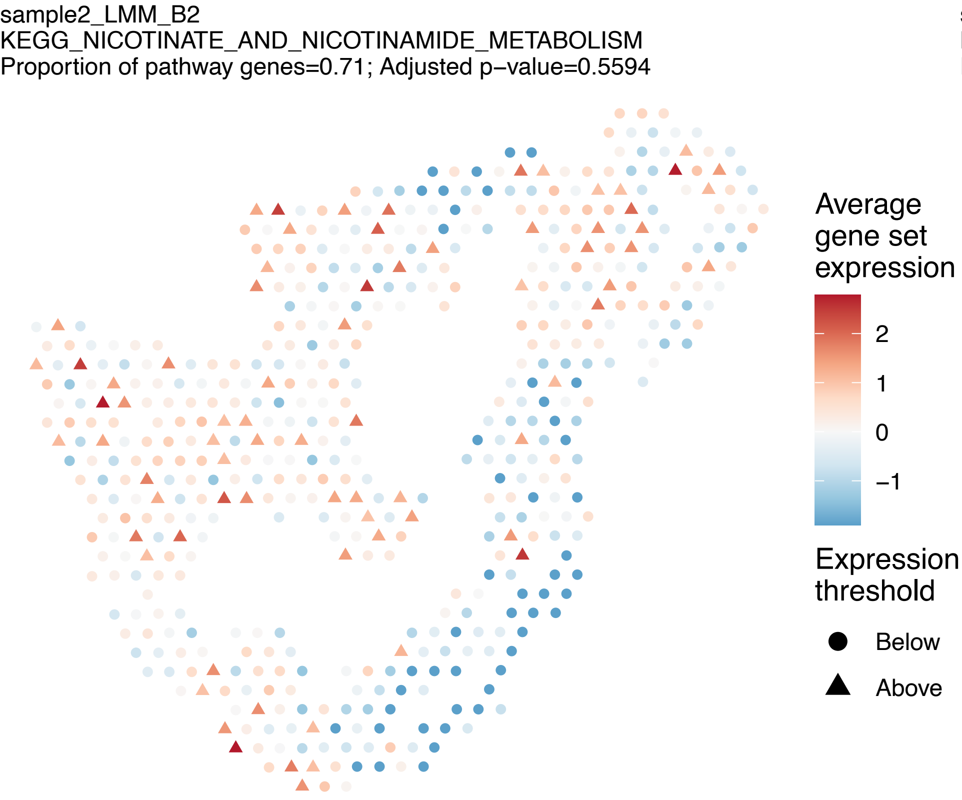
sample2_LMM_B2
KEGG_NICOTINATE_AND_NICOTINAMIDE_METABOLISM
Proportion of pathway genes=0.71; Adjusted p−value=0.5594
Average gene set expression
2
1
0
−1
Expression threshold
Below
Above

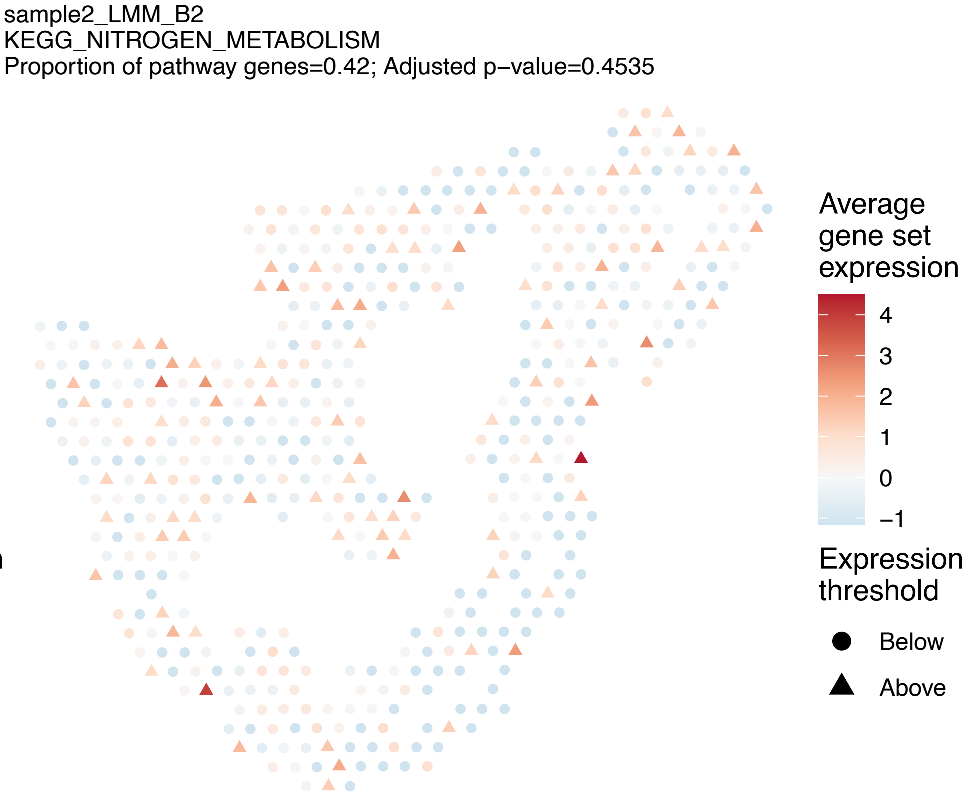
sample2_LMM_B2
KEGG_NITROGEN_METABOLISM
Proportion of pathway genes=0.42; Adjusted p−value=0.4535
Average gene set expression
4
3
2
1
0
−1
Expression threshold
Below
Above

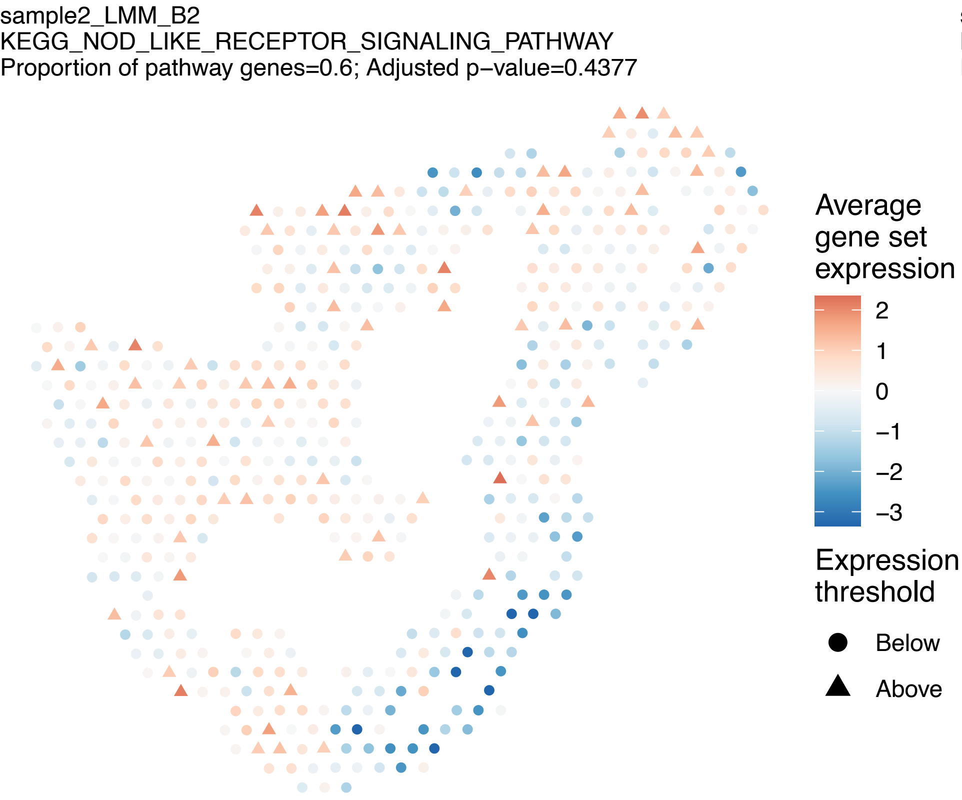
sample2_LMM_B2
KEGG_NOD_LIKE_RECEPTOR_SIGNALING_PATHWAY
Proportion of pathway genes=0.6; Adjusted p−value=0.4377
Average gene set expression
2
1
0
−1
−2
−3
Expression threshold
Below
Above

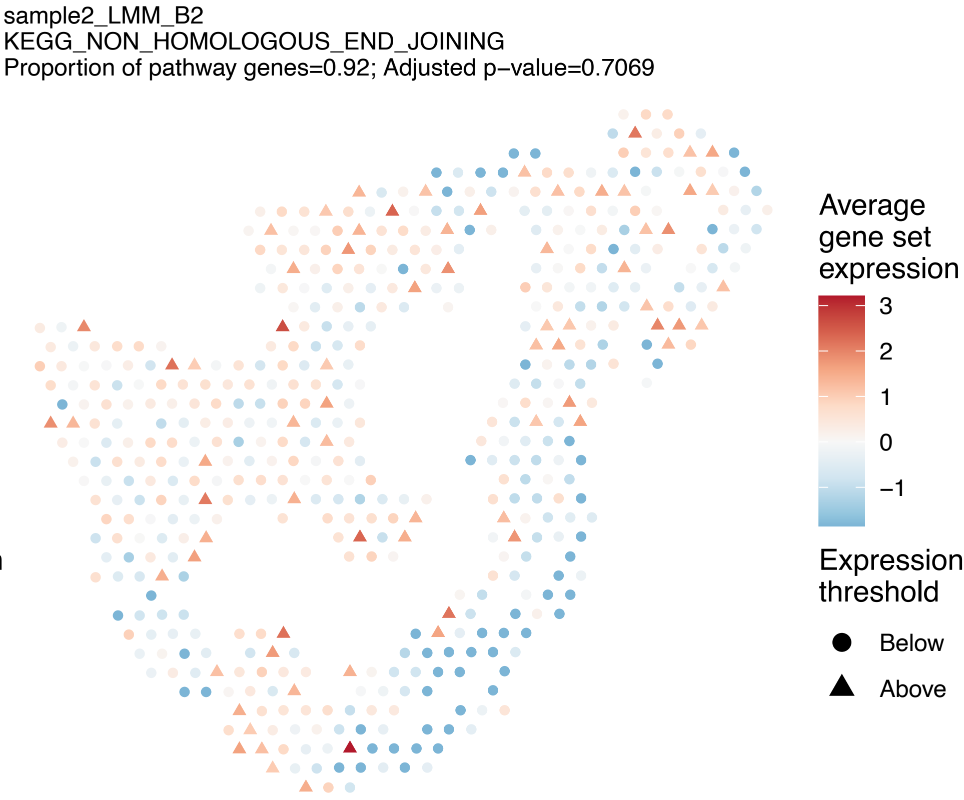
sample2_LMM_B2
KEGG_NON_HOMOLOGOUS_END_JOINING
Proportion of pathway genes=0.92; Adjusted p−value=0.7069
Average gene set expression
3
2
1
0
−1
Expression threshold
Below
Above

sample2_LMM_B2
KEGG_NON_SMALL_CELL_LUNG_CANCER
Proportion of pathway genes=0.87; Adjusted p−value=0

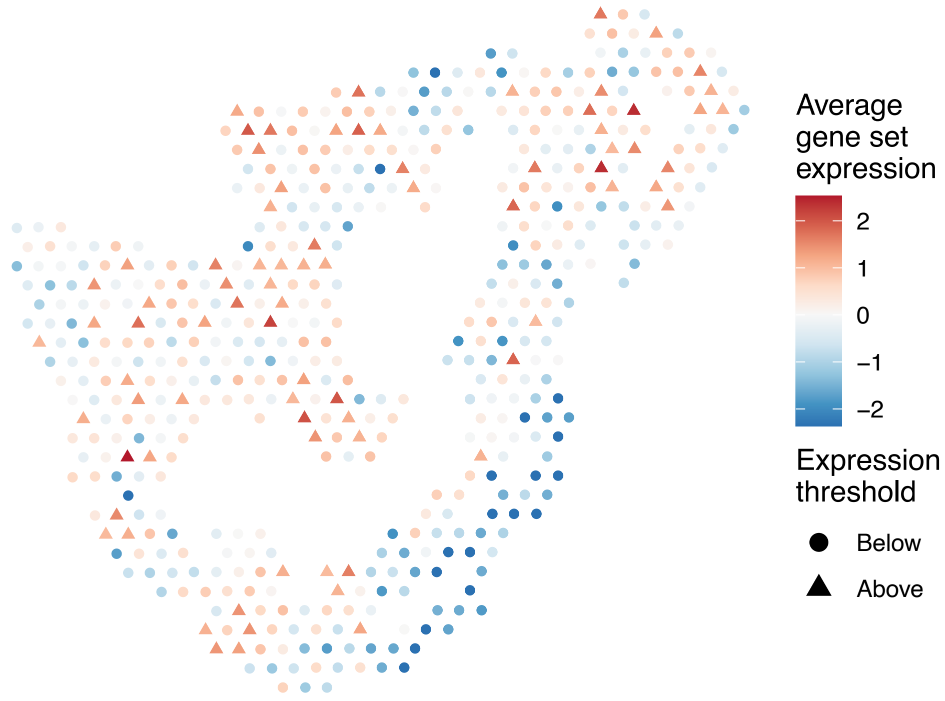


sample2_LMM_B2
KEGG_NOTCH_SIGNALING_PATHWAY
Proportion of pathway genes=0.84; Adjusted p−value=0.0755

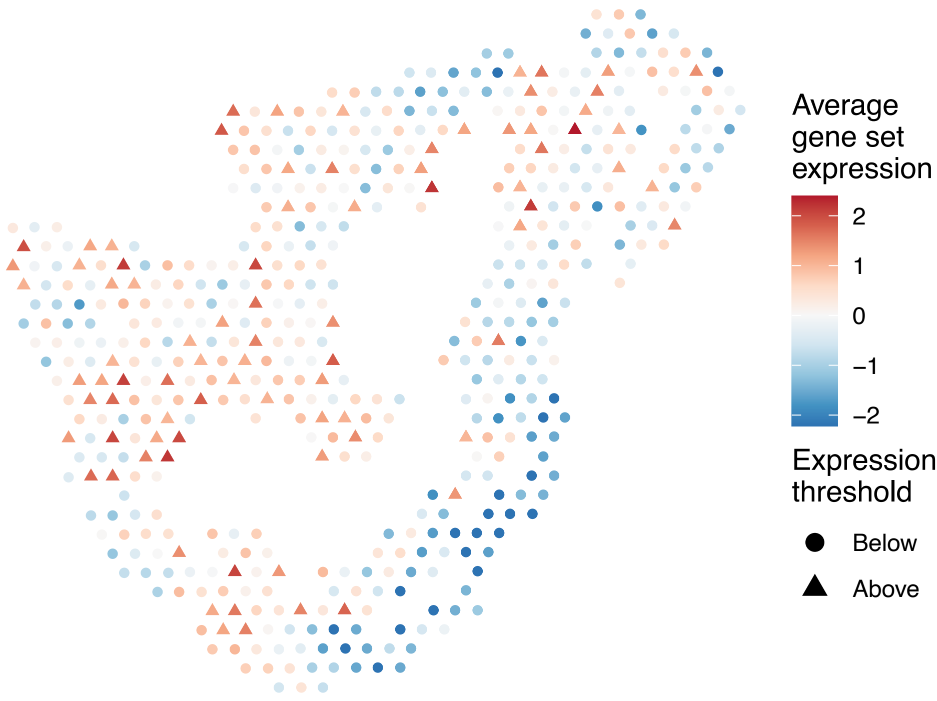


sample2_LMM_B2
KEGG_NUCLEOTIDE_EXCISION_REPAIR
Proportion of pathway genes=0.98; Adjusted p−value=0.012

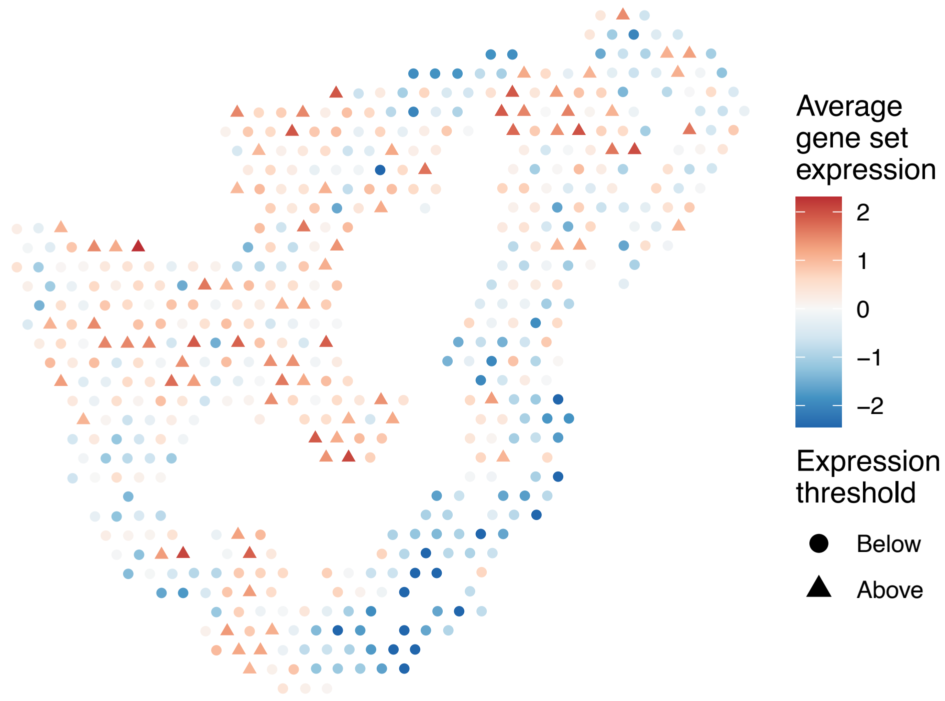


sample2_LMM_B2
KEGG_O_GLYCAN_BIOSYNTHESIS
Proportion of pathway genes=0.48; Adjusted p−value=0.1958

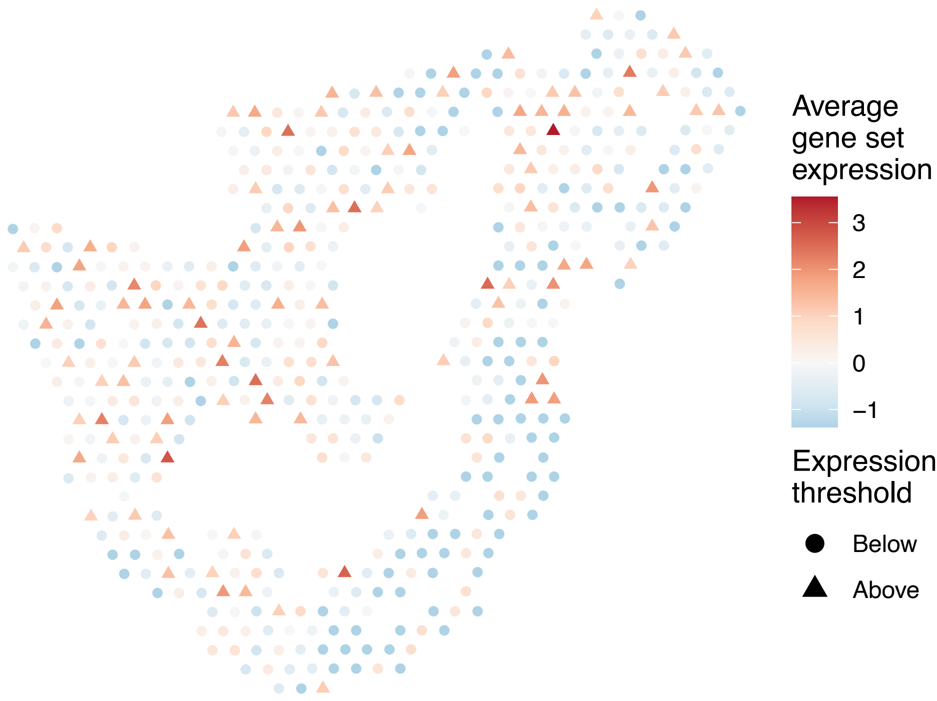

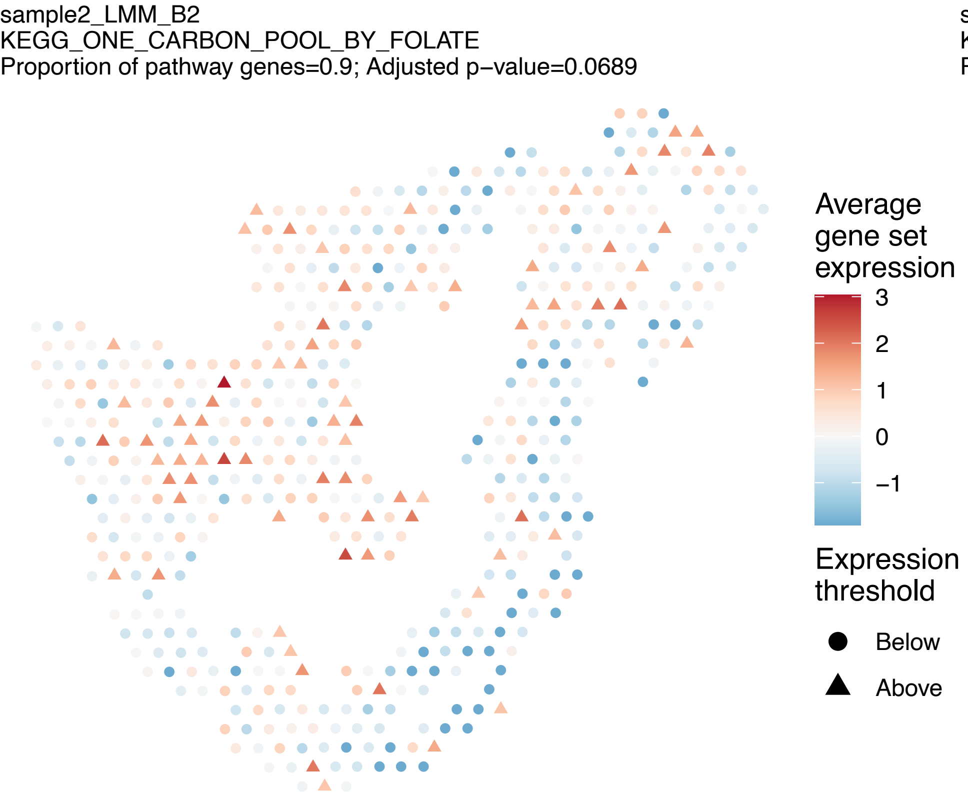
sample2_LMM_B2
KEGG_ONE_CARBON_POOL_BY_FOLATE
Proportion of pathway genes=0.9; Adjusted p−value=0.0689
Average gene set expression
3
2
1
0
−1
Expression threshold
Below
Above

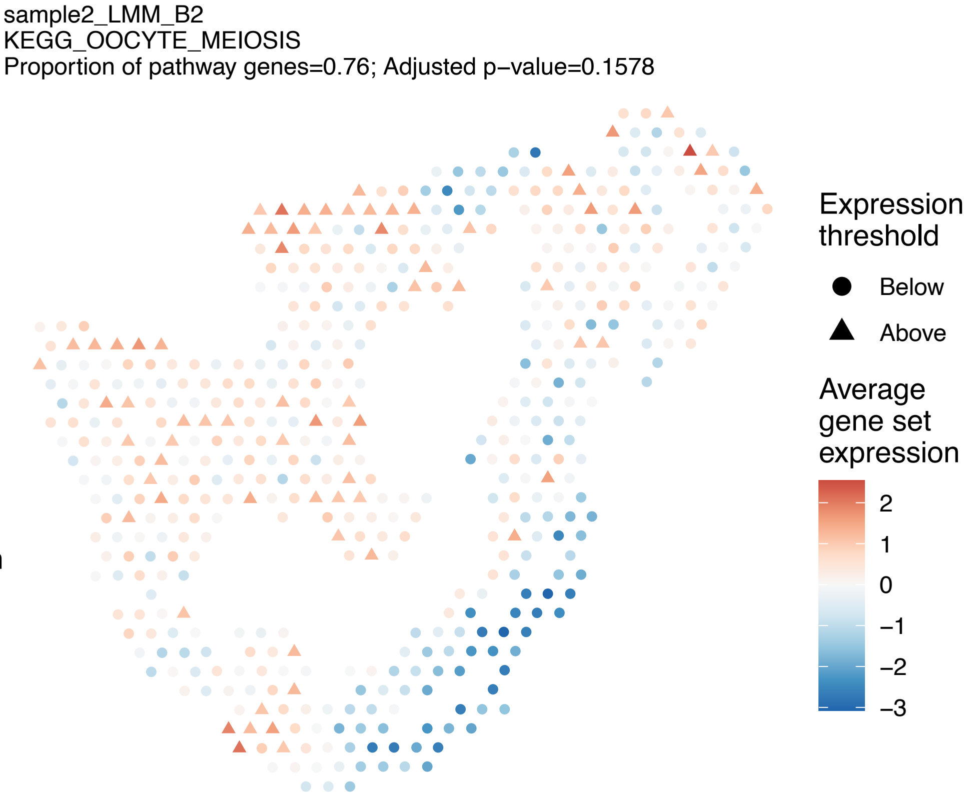
sample2_LMM_B2
KEGG_OOCYTE_MEIOSIS
Proportion of pathway genes=0.76; Adjusted p−value=0.1578
Expression threshold
Below
Above
Average gene set expression
2
1
0
−1
−2
−3

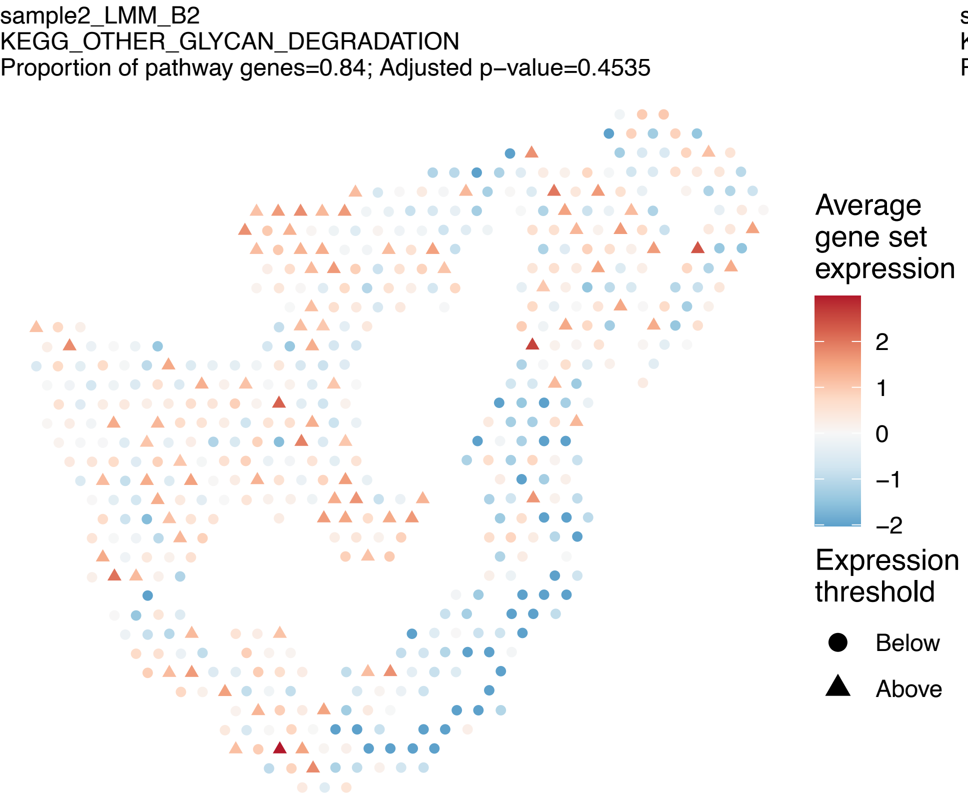
sample2_LMM_B2
KEGG_OTHER_GLYCAN_DEGRADATION
Proportion of pathway genes=0.84; Adjusted p−value=0.4535
Average gene set expression
2
1
0
−1
−2
Expression threshold
Below
Above

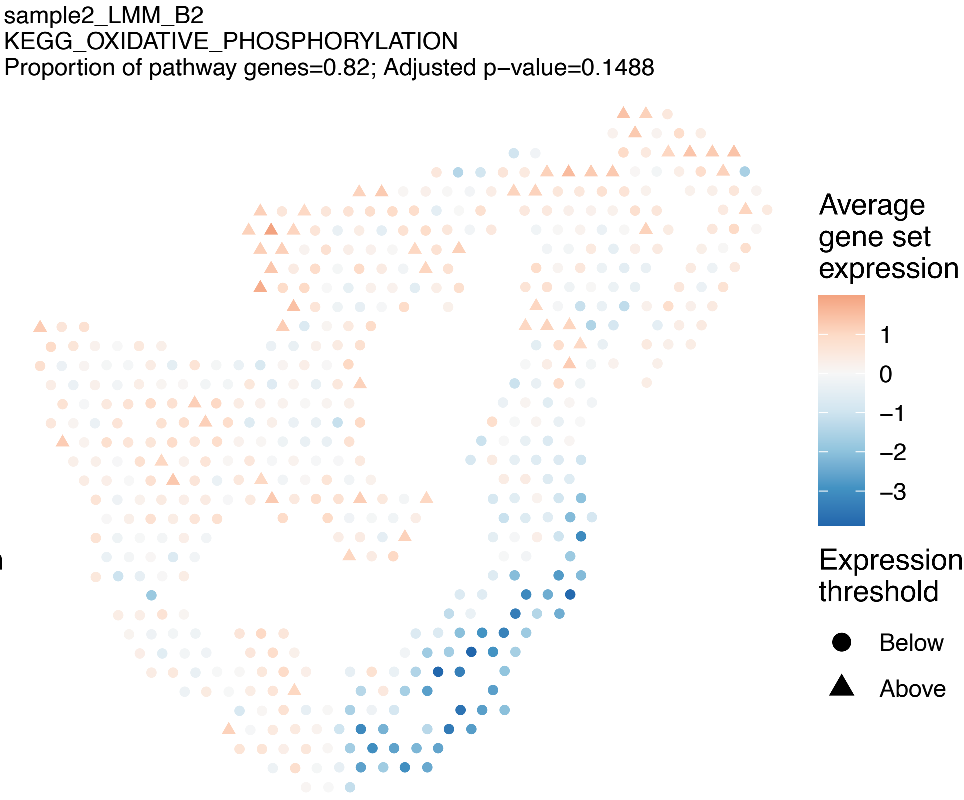
sample2_LMM_B2
KEGG_OXIDATIVE_PHOSPHORYLATION
Proportion of pathway genes=0.82; Adjusted p−value=0.1488
Average gene set expression
1
0
−1
−2
−3
Expression threshold
Below
Above

sample2_LMM_B2
KEGG_P53_SIGNALING_PATHWAY
Proportion of pathway genes=0.86; Adjusted p−value=0.0643

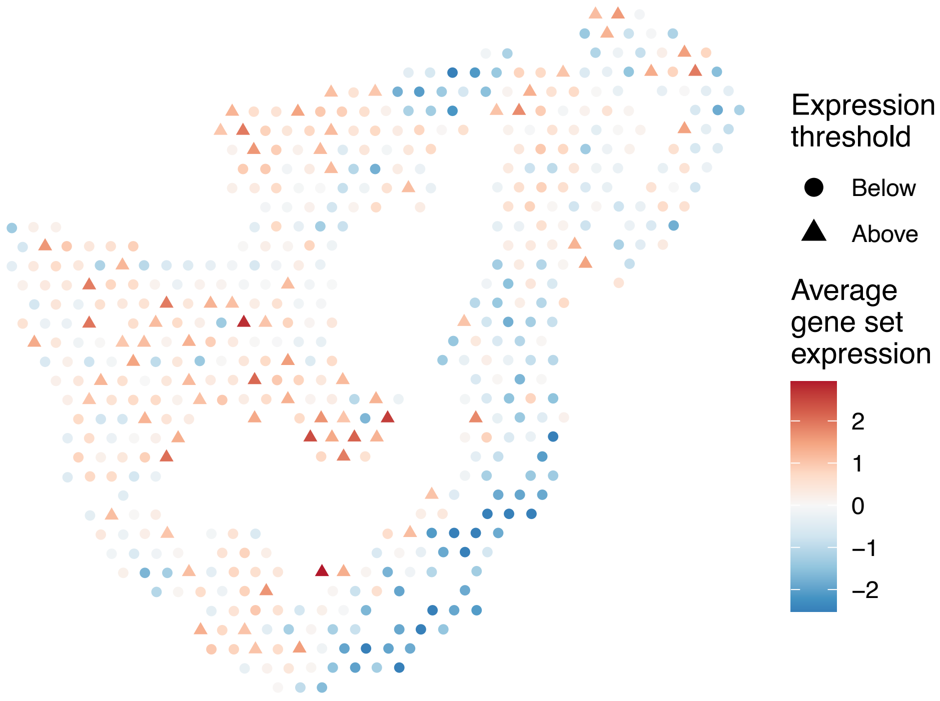


sample2_LMM_B2
KEGG_PANCREATIC_CANCER
Proportion of pathway genes=0.86; Adjusted p−value=0.3638

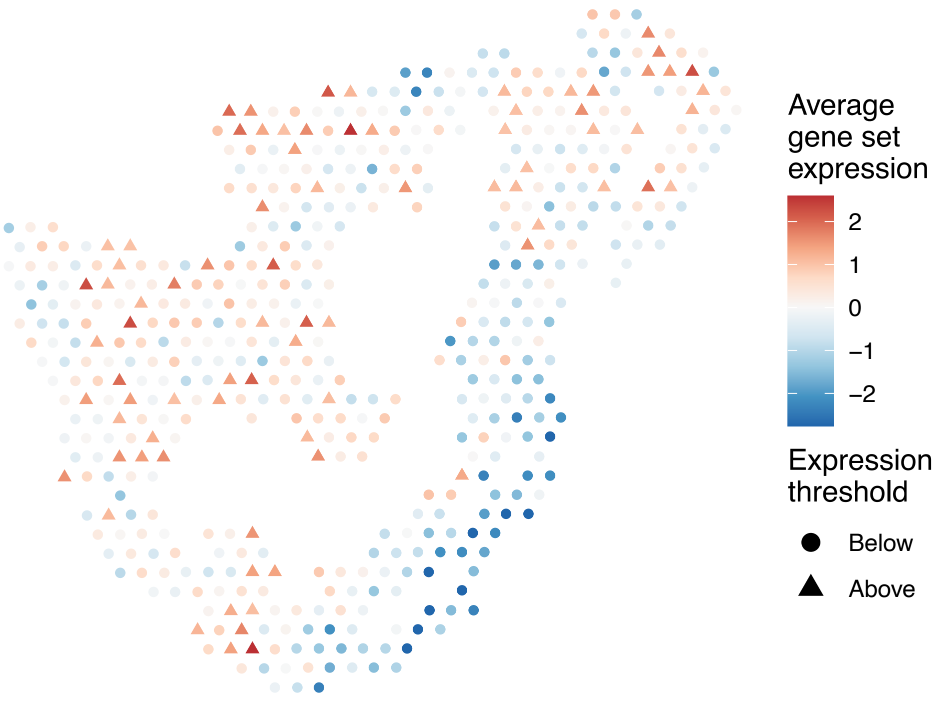


sample2_LMM_B2
KEGG_PANTOTHENATE_AND_COA_BIOSYNTHESIS
Proportion of pathway genes=0.59; Adjusted p−value=0.4558

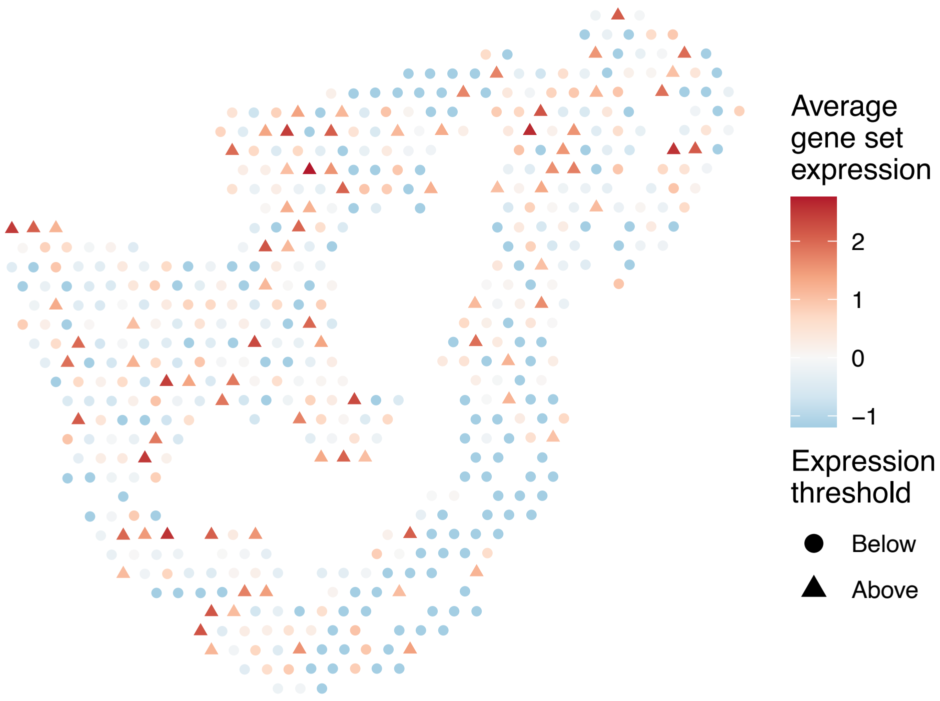


sample2_LMM_B2
KEGG_PARKINSONS_DISEASE
Proportion of pathway genes=0.85; Adjusted p−value=0.1648

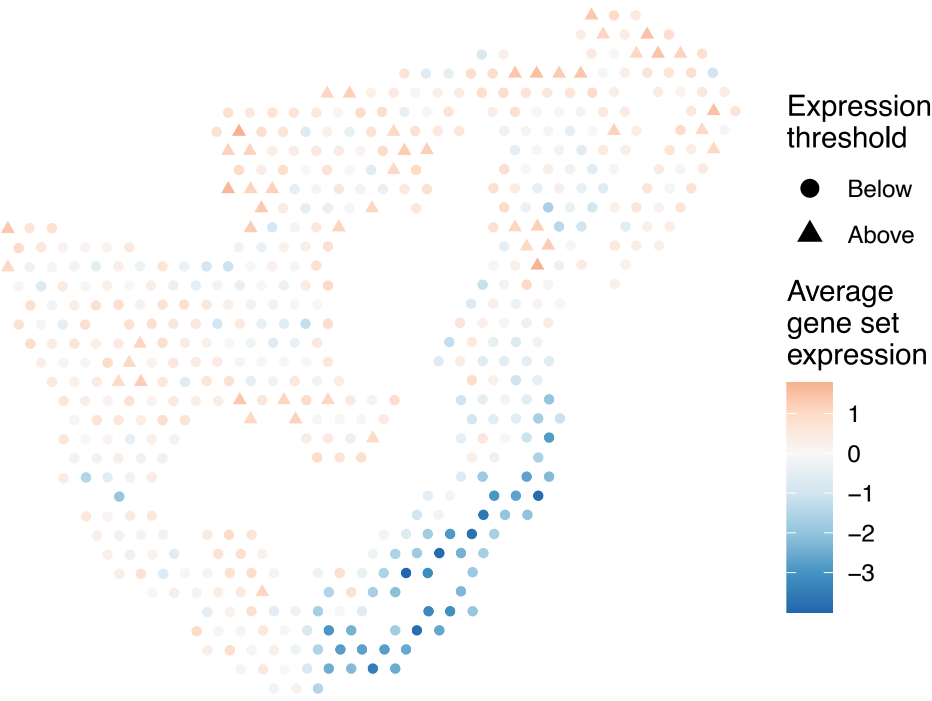

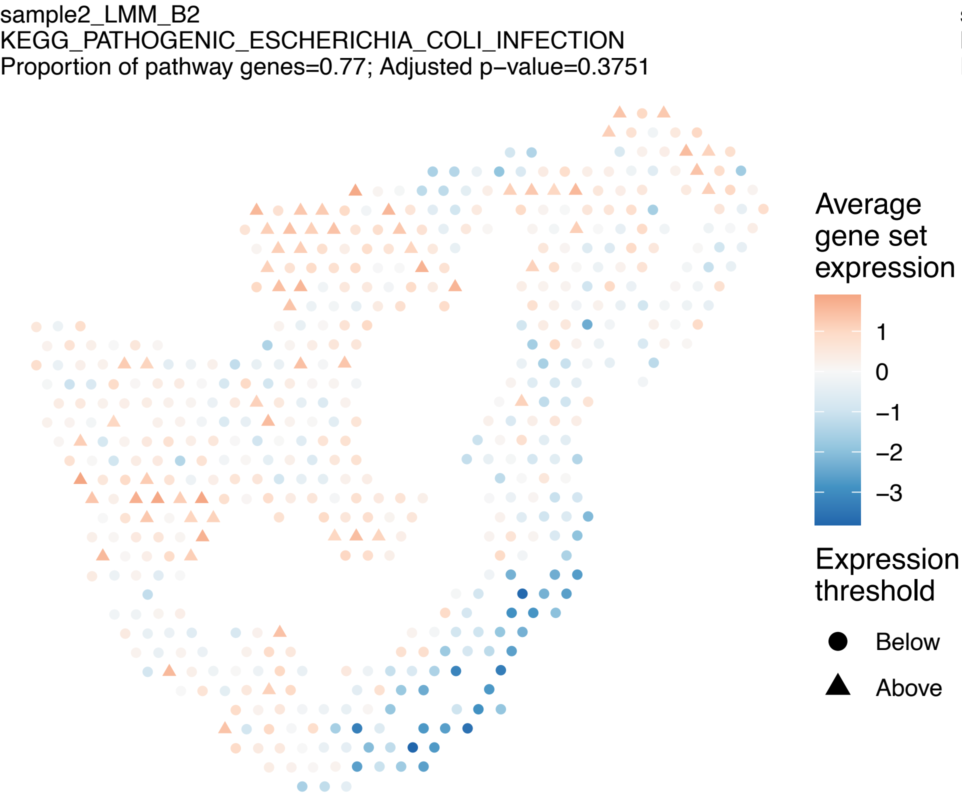
sample2_LMM_B2
KEGG_PATHOGENIC_ESCHERICHIA_COLI_INFECTION
Proportion of pathway genes=0.77; Adjusted p−value=0.3751
Average gene set expression
1
0
−1
−2
−3
Expression threshold
Below
Above

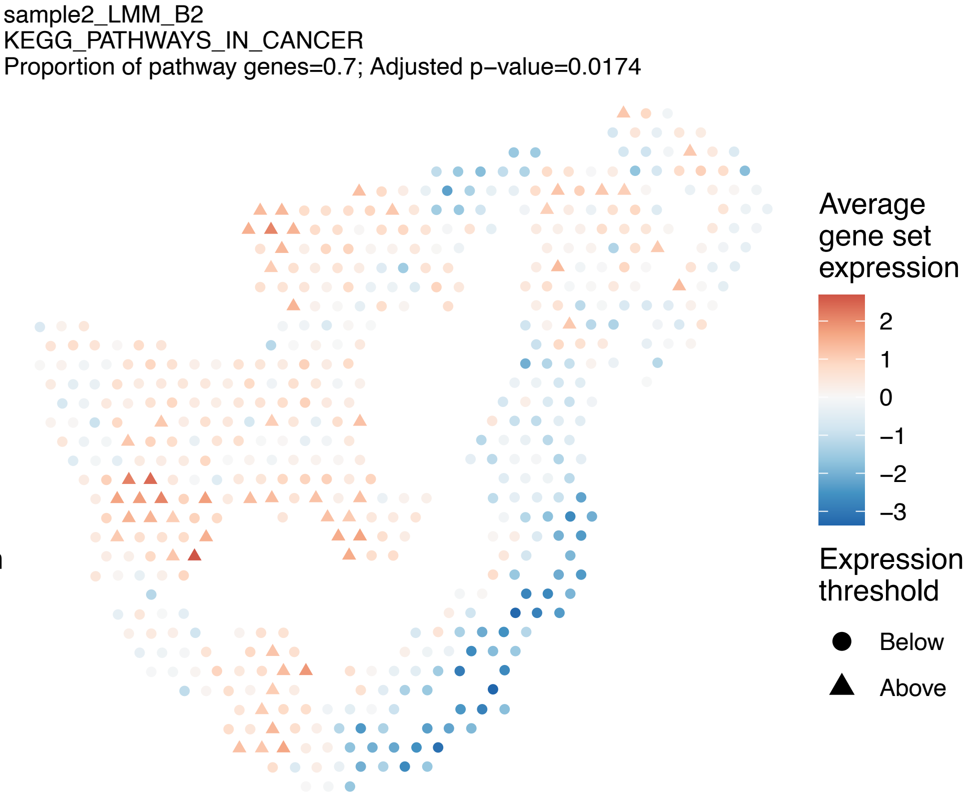
sample2_LMM_B2
KEGG_PATHWAYS_IN_CANCER
Proportion of pathway genes=0.7; Adjusted p−value=0.0174
Average gene set expression
2
1
0
−1
−2
−3
Expression threshold
Below
Above

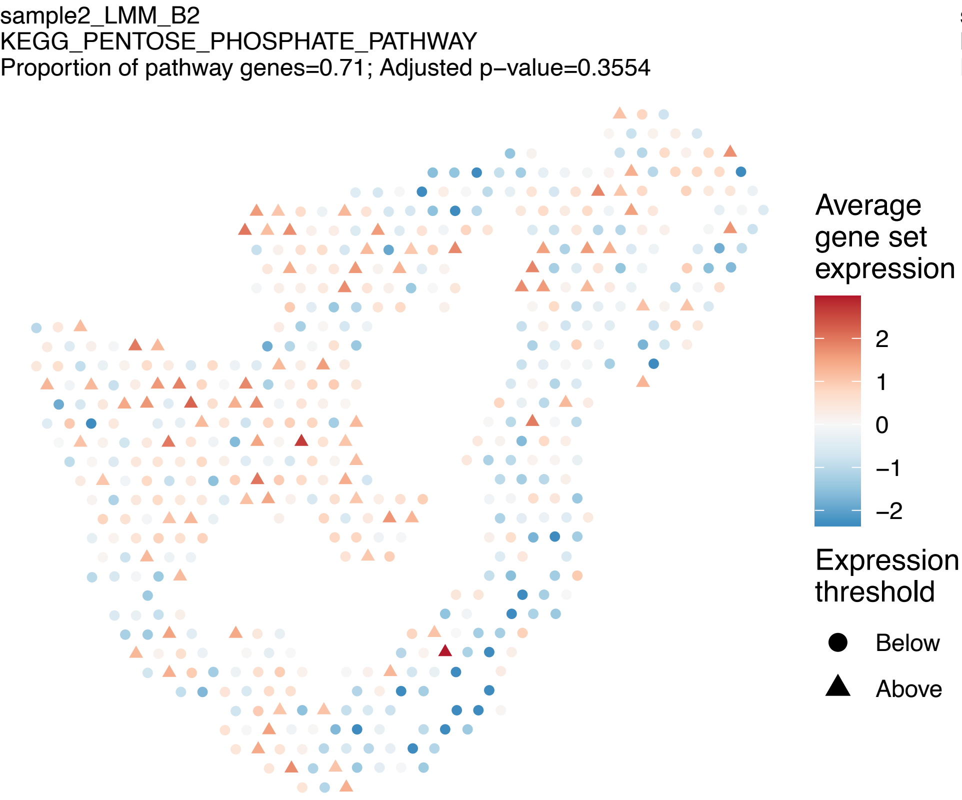
sample2_LMM_B2
KEGG_PENTOSE_PHOSPHATE_PATHWAY
Proportion of pathway genes=0.71; Adjusted p−value=0.3554
Average gene set expression
2
1
0
−1
−2
Expression threshold
Below
Above

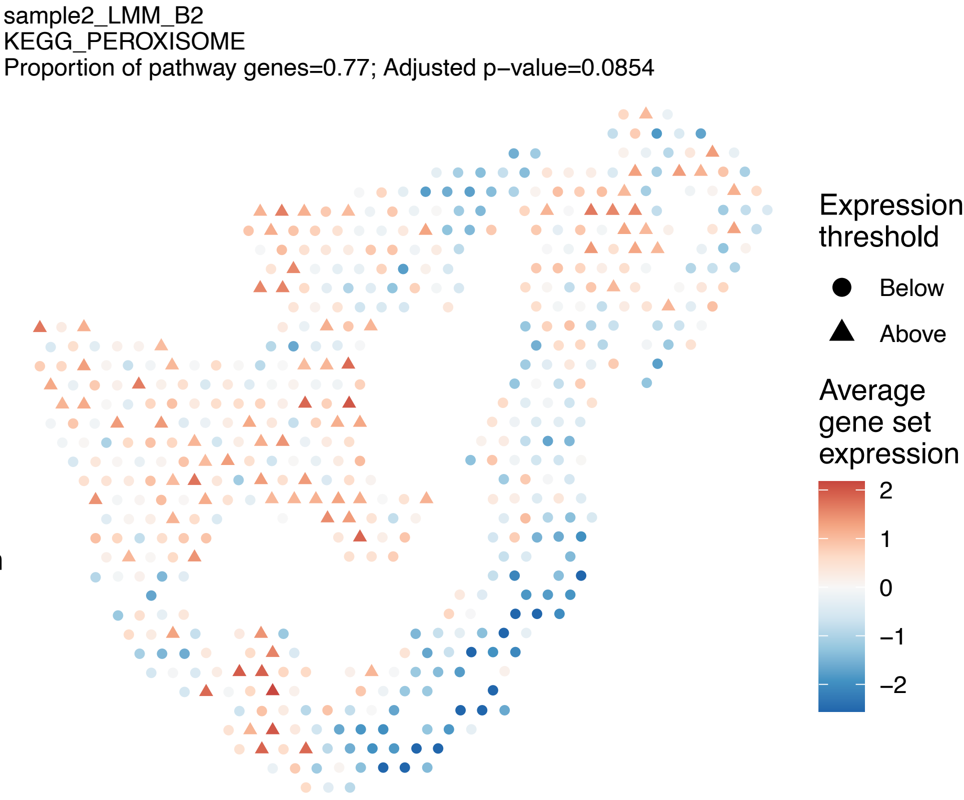
sample2_LMM_B2
KEGG_PEROXISOME
Proportion of pathway genes=0.77; Adjusted p−value=0.0854
Expression threshold
Below
Above
Average gene set expression
2
1
0
−1
−2

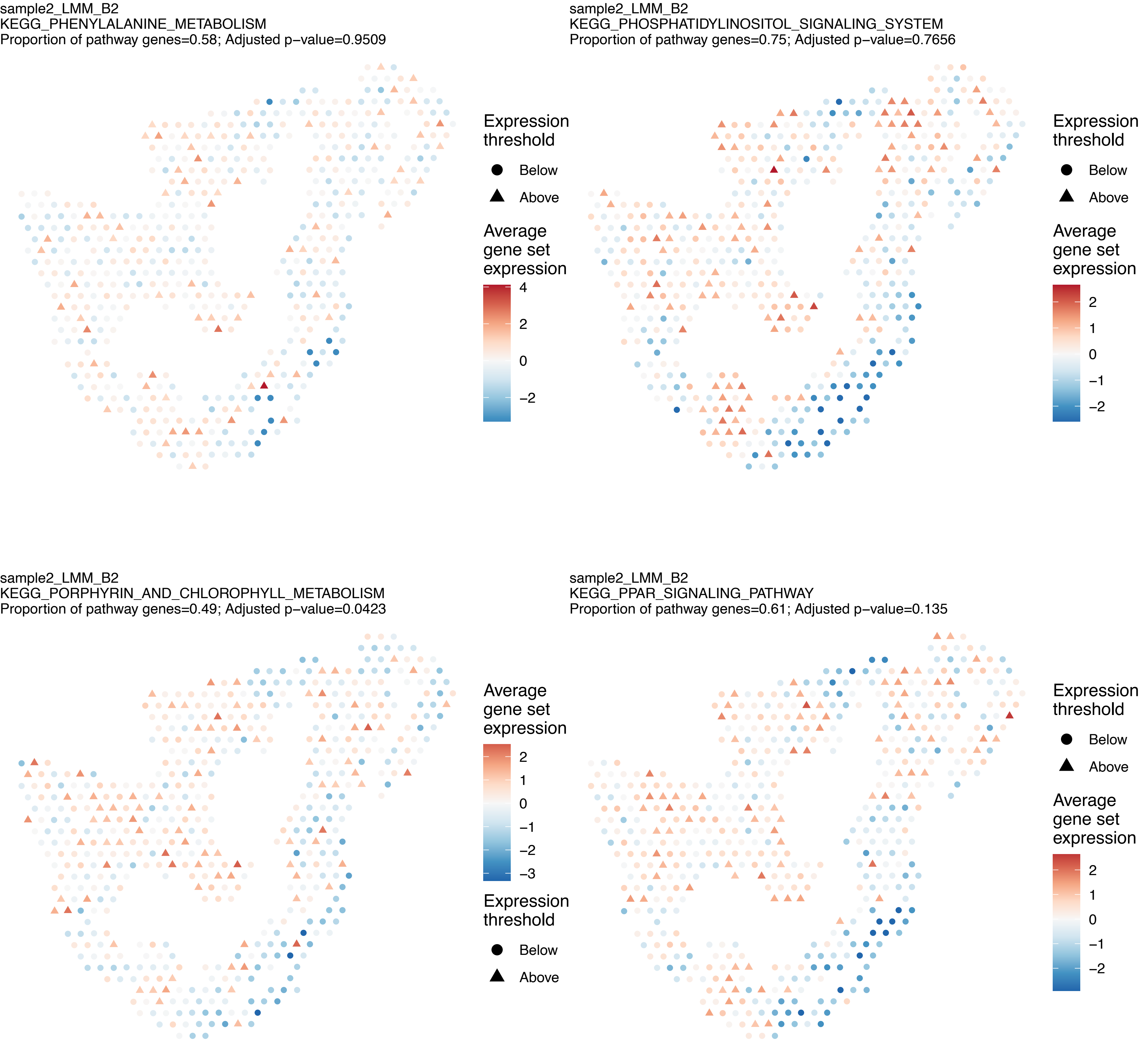
sample2_LMM_B2
KEGG_PHENYLALANINE_METABOLISM
Proportion of pathway genes=0.58; Adjusted p−value=0.9509
Expression threshold
Below
Above
Average gene set expression
4
2
0
−2
sample2_LMM_B2
KEGG_PHOSPHATIDYLINOSITOL_SIGNALING_SYSTEM
Proportion of pathway genes=0.75; Adjusted p−value=0.7656
Expression threshold
Below
Above
Average gene set expression
2
1
0
−1
−2
sample2_LMM_B2
KEGG_PORPHYRIN_AND_CHLOROPHYLL_METABOLISM
Proportion of pathway genes=0.49; Adjusted p−value=0.0423
Average gene set expression
2
1
0
−1
−2
−3
Expression threshold
Below
Above
sample2_LMM_B2
KEGG_PPAR_SIGNALING_PATHWAY
Proportion of pathway genes=0.61; Adjusted p−value=0.135
Expression threshold
Below
Above
Average gene set expression
2
1
0
−1
−2

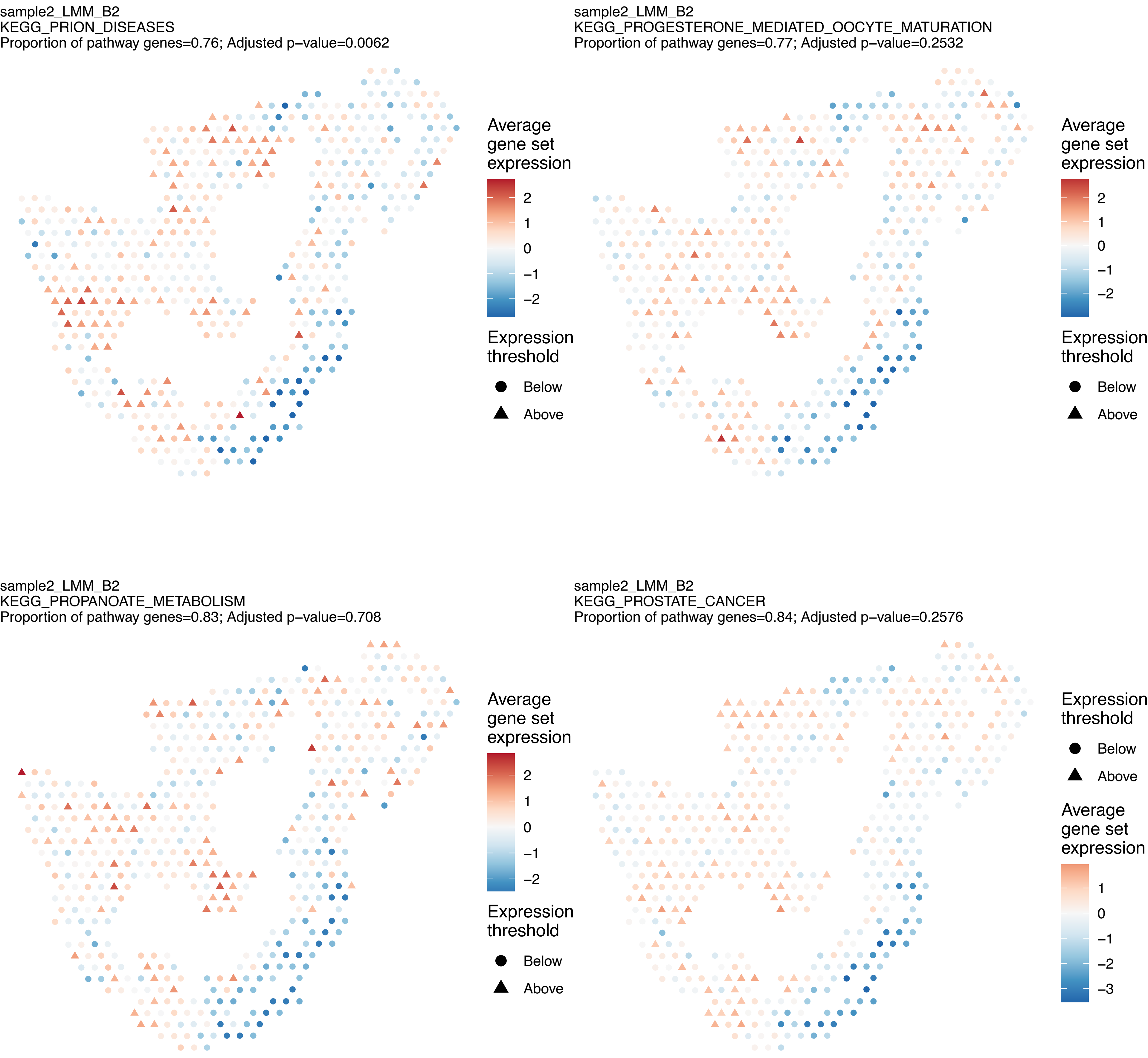
sample2_LMM_B2
KEGG_PRION_DISEASES
Proportion of pathway genes=0.76; Adjusted p−value=0.0062
Average gene set expression
2
1
0
−1
−2
Expression threshold
Below
Above
sample2_LMM_B2
KEGG_PROGESTERONE_MEDIATED_OOCYTE_MATURATION
Proportion of pathway genes=0.77; Adjusted p−value=0.2532
Average gene set expression
2
1
0
−1
−2
Expression threshold
Below
Above
sample2_LMM_B2
KEGG_PROPANOATE_METABOLISM
Proportion of pathway genes=0.83; Adjusted p−value=0.708
Average gene set expression
2
1
0
−1
−2
Expression threshold
Below
Above
sample2_LMM_B2
KEGG_PROSTATE_CANCER
Proportion of pathway genes=0.84; Adjusted p−value=0.2576
Expression threshold
Below
Above
Average gene set expression
1
0
−1
−2
−3

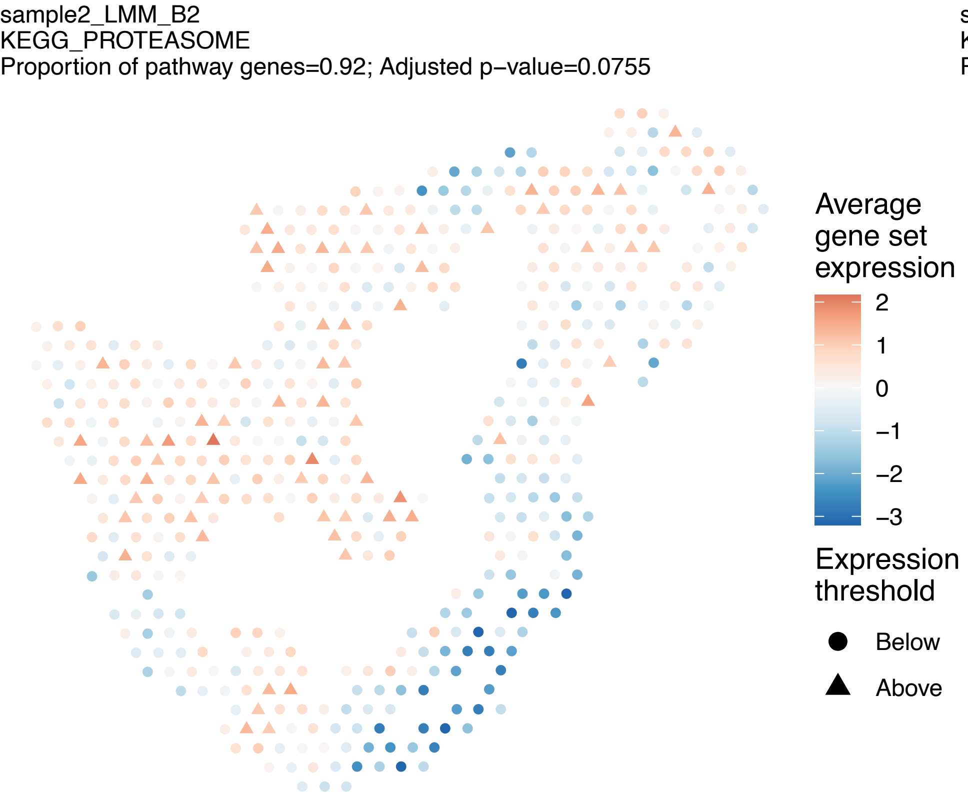
sample2_LMM_B2
KEGG_PROTEASOME
Proportion of pathway genes=0.92; Adjusted p−value=0.0755
Average gene set expression
2
1
0
−1
−2
−3
Expression threshold
Below
Above

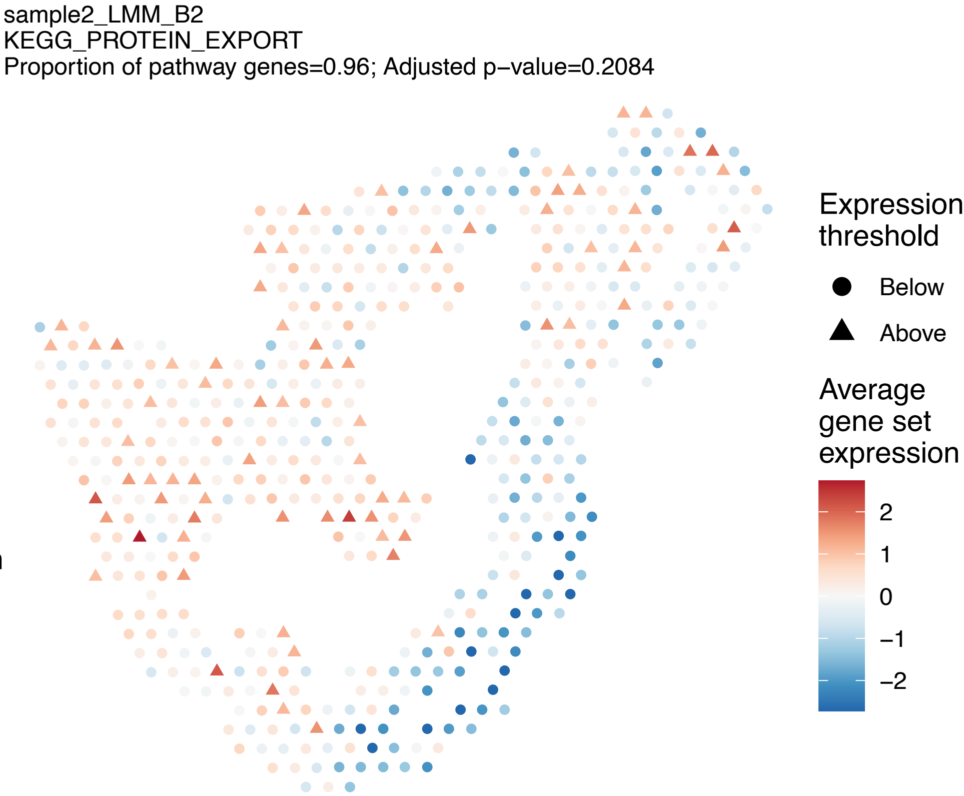
sample2_LMM_B2
KEGG_PROTEIN_EXPORT
Proportion of pathway genes=0.96; Adjusted p−value=0.2084
Expression threshold
Below
Above
Average gene set expression
2
1
0
−1
−2

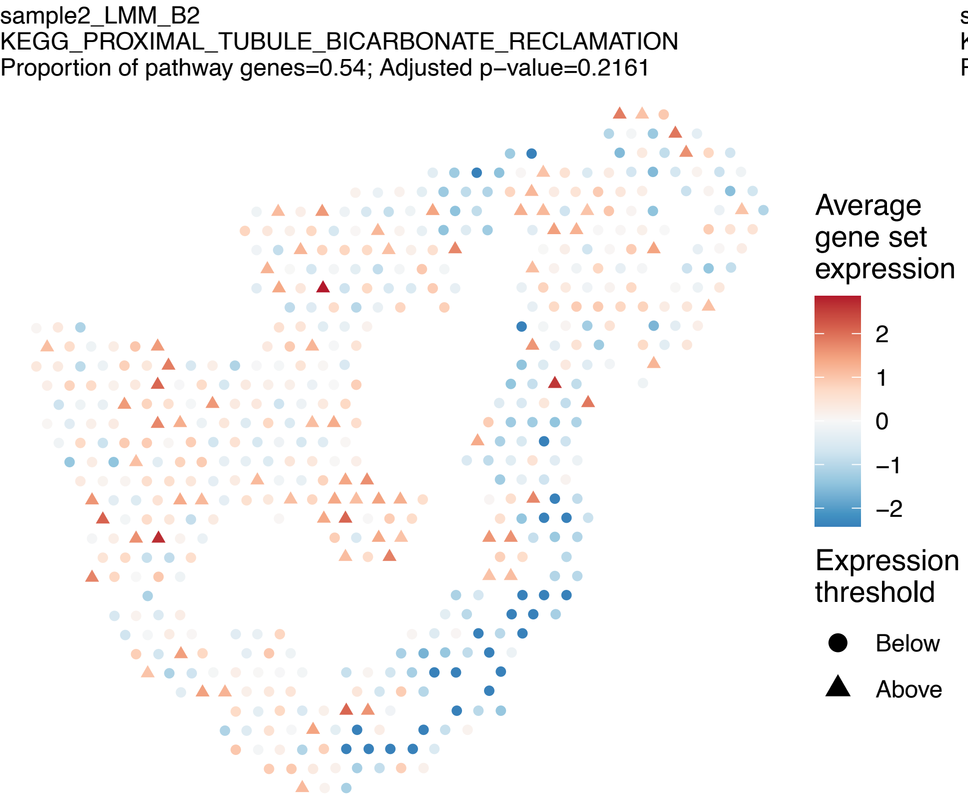
sample2_LMM_B2
KEGG_PROXIMAL_TUBULE_BICARBONATE_RECLAMATION
Proportion of pathway genes=0.54; Adjusted p−value=0.2161
Average gene set expression
2
1
0
−1
−2
Expression threshold
Below
Above

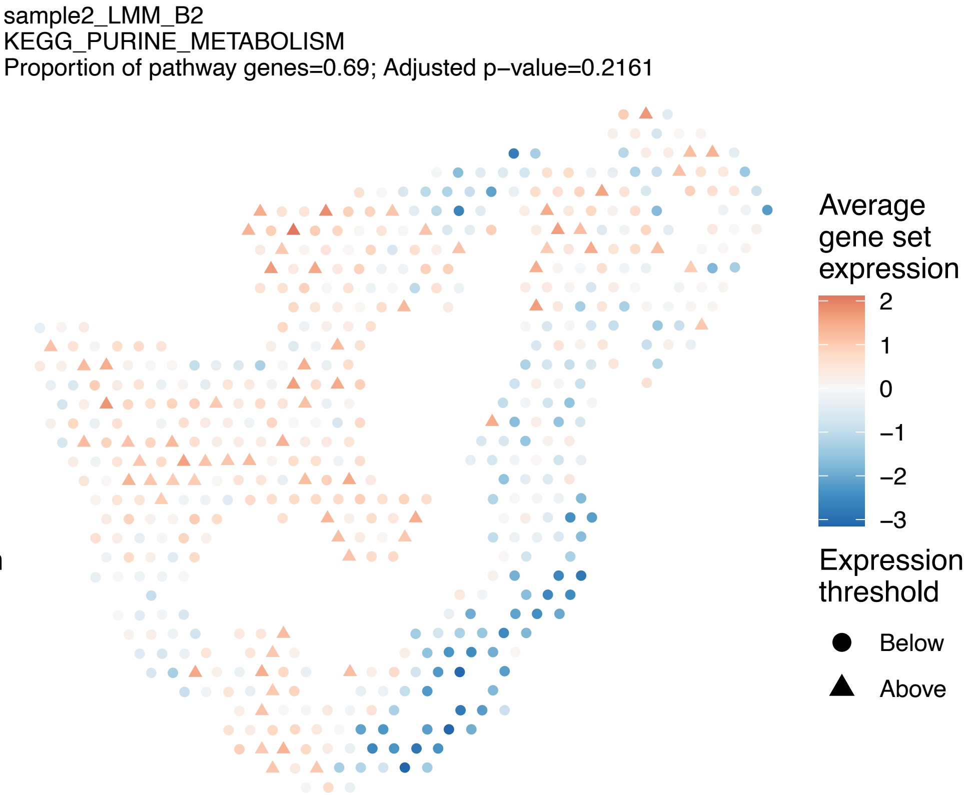
sample2_LMM_B2
KEGG_PURINE_METABOLISM
Proportion of pathway genes=0.69; Adjusted p−value=0.2161
Average gene set expression
2
1
0
−1
−2
−3
Expression threshold
Below
Above

sample2_LMM_B2
KEGG_PYRIMIDINE_METABOLISM
Proportion of pathway genes=0.79; Adjusted p−value=0

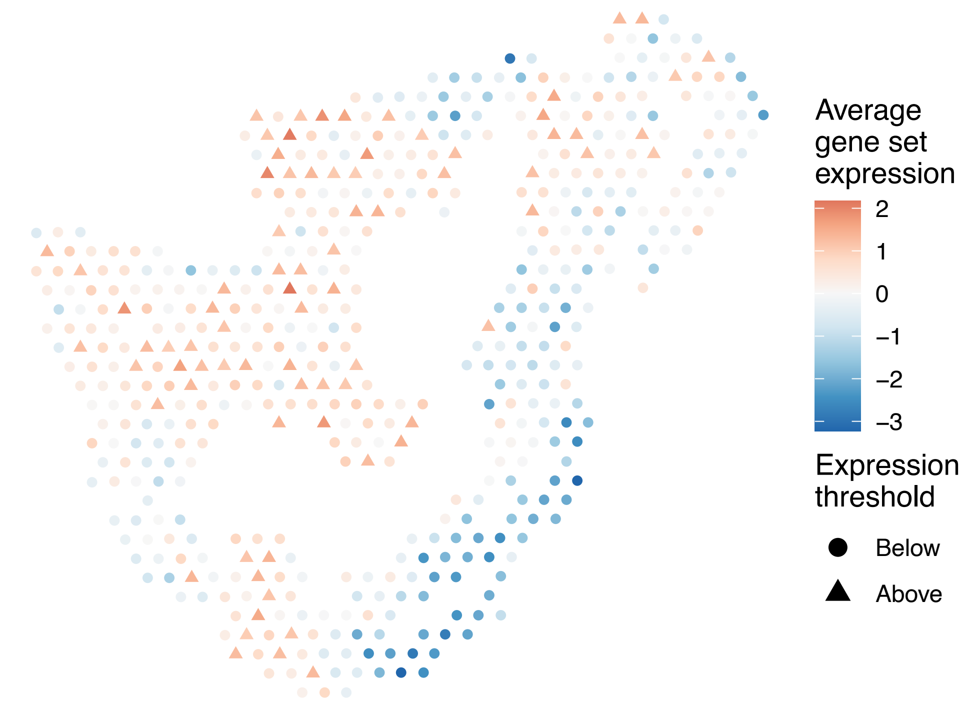


sample2_LMM_B2
KEGG_PYRUVATE_METABOLISM
Proportion of pathway genes=0.77; Adjusted p−value=0.5988

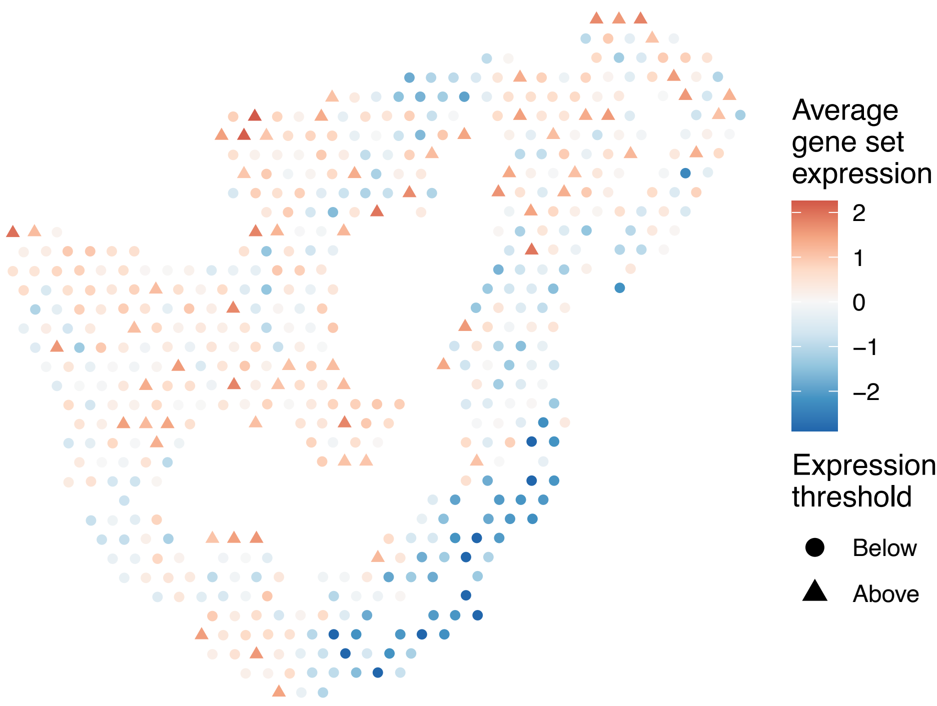


sample2_LMM_B2
KEGG_REGULATION_OF_ACTIN_CYTOSKELETON
Proportion of pathway genes=0.72; Adjusted p−value=0.2095

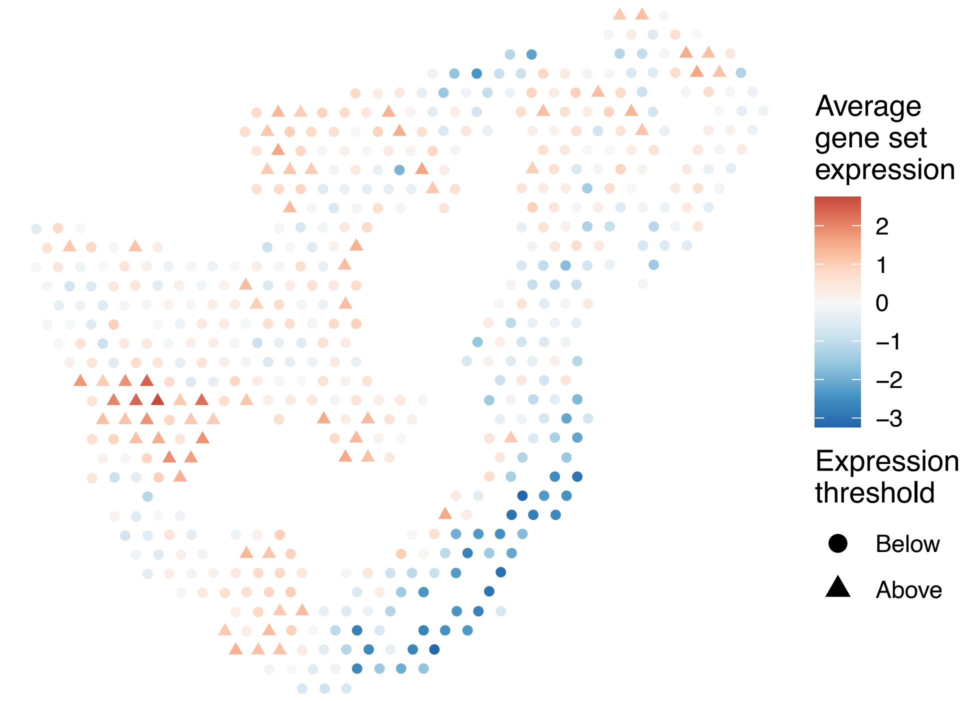


sample2_LMM_B2
KEGG_REGULATION_OF_AUTOPHAGY
Proportion of pathway genes=0.51; Adjusted p−value=0.1922

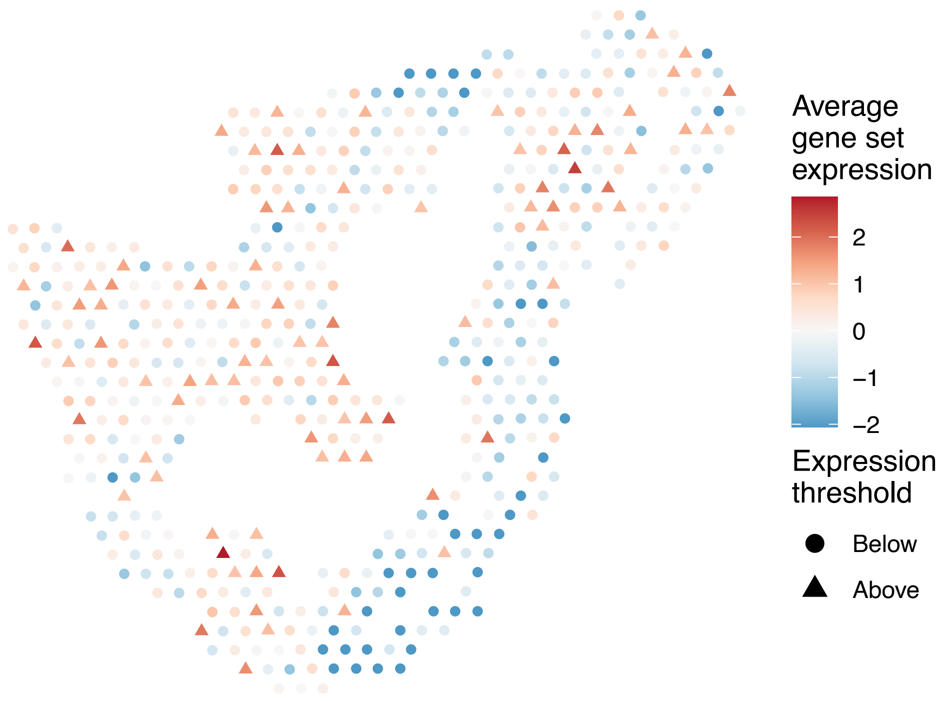

sample2_LMM_B2
KEGG_RENAL_CELL_CARCINOMA
Proportion of pathway genes=0.85; Adjusted p−value=0.2478

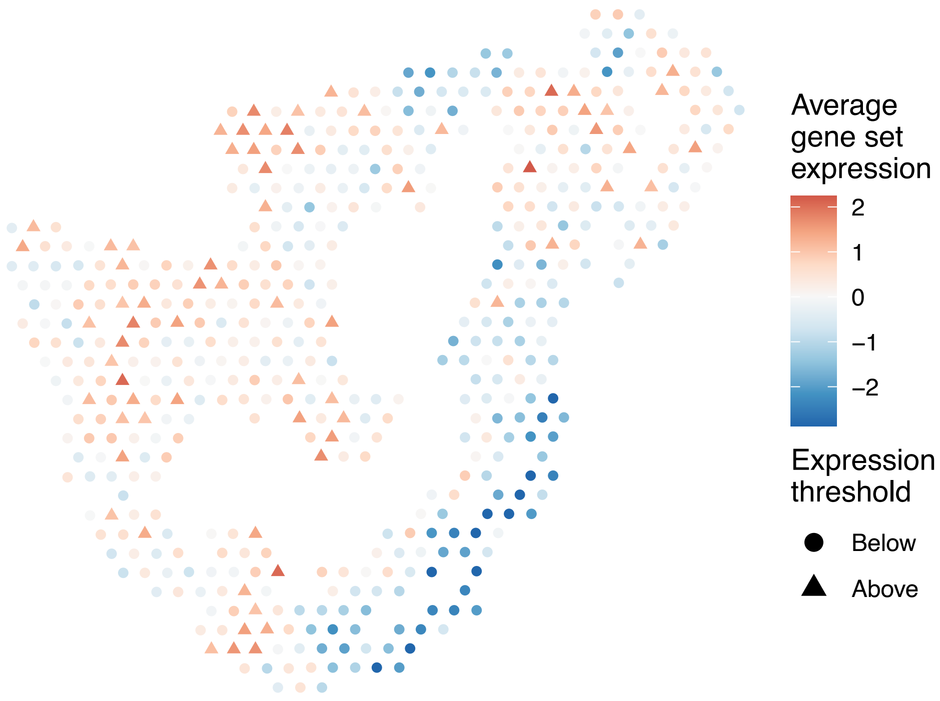


sample2_LMM_B2
KEGG_RETINOL_METABOLISM
Proportion of pathway genes=0.27; Adjusted p−value=0.0509

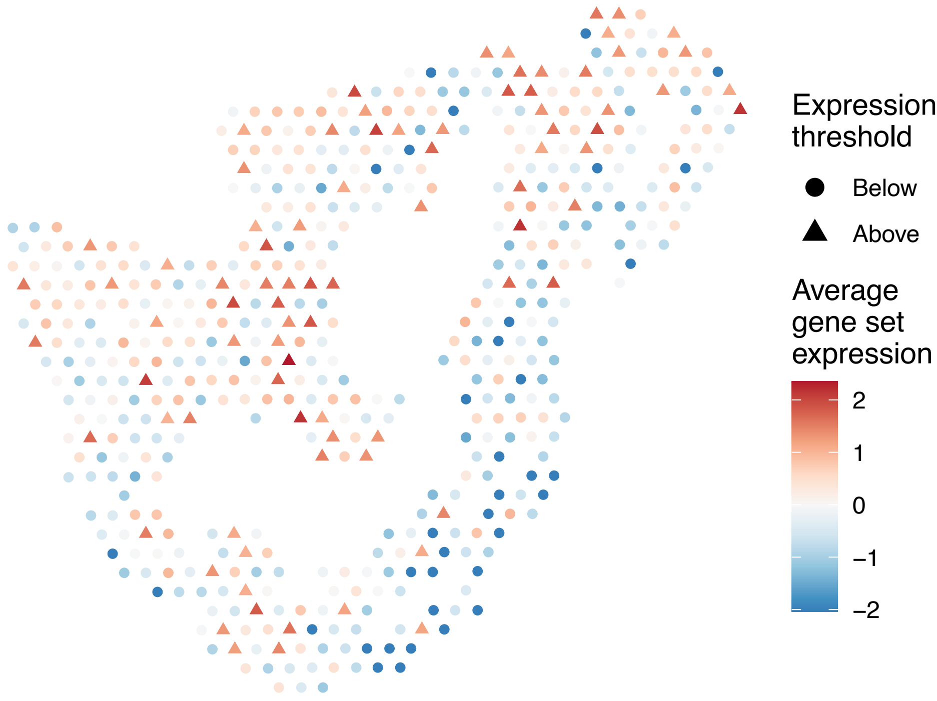


sample2_LMM_B2
KEGG_RIBOFLAVIN_METABOLISM
Proportion of pathway genes=0.75; Adjusted p−value=0.0423

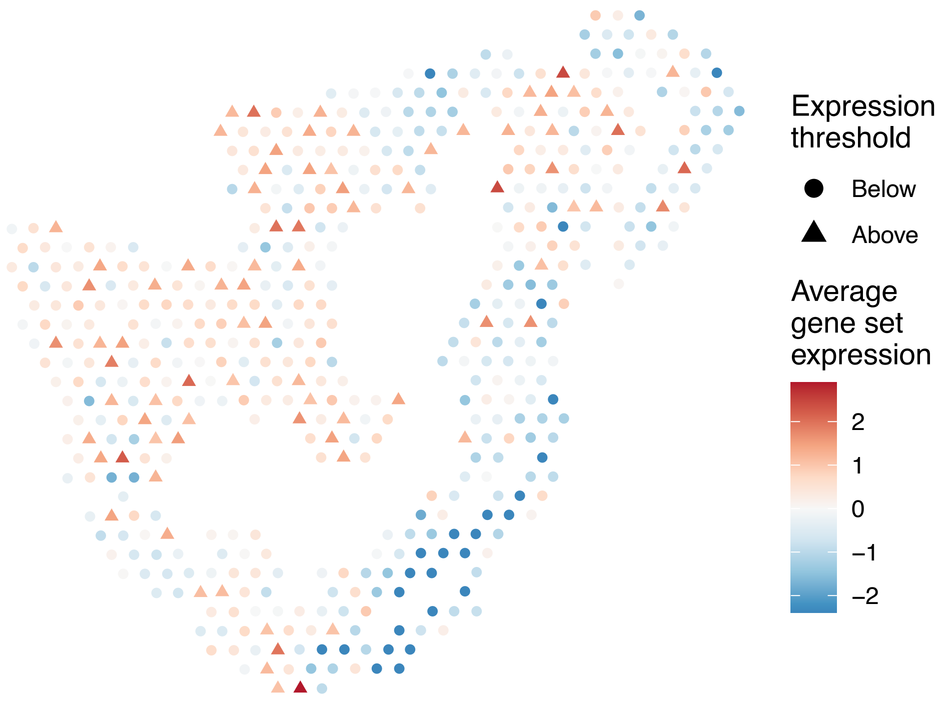


sample2_LMM_B2
KEGG_RIG_I_LIKE_RECEPTOR_SIGNALING_PATHWAY
Proportion of pathway genes=0.58; Adjusted p−value=0.0643

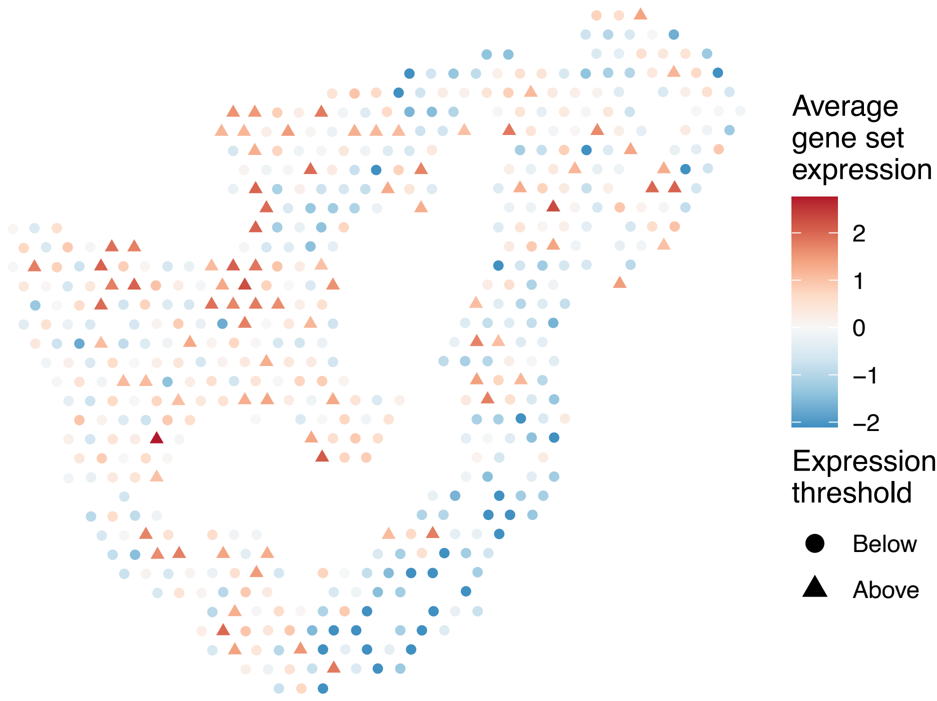

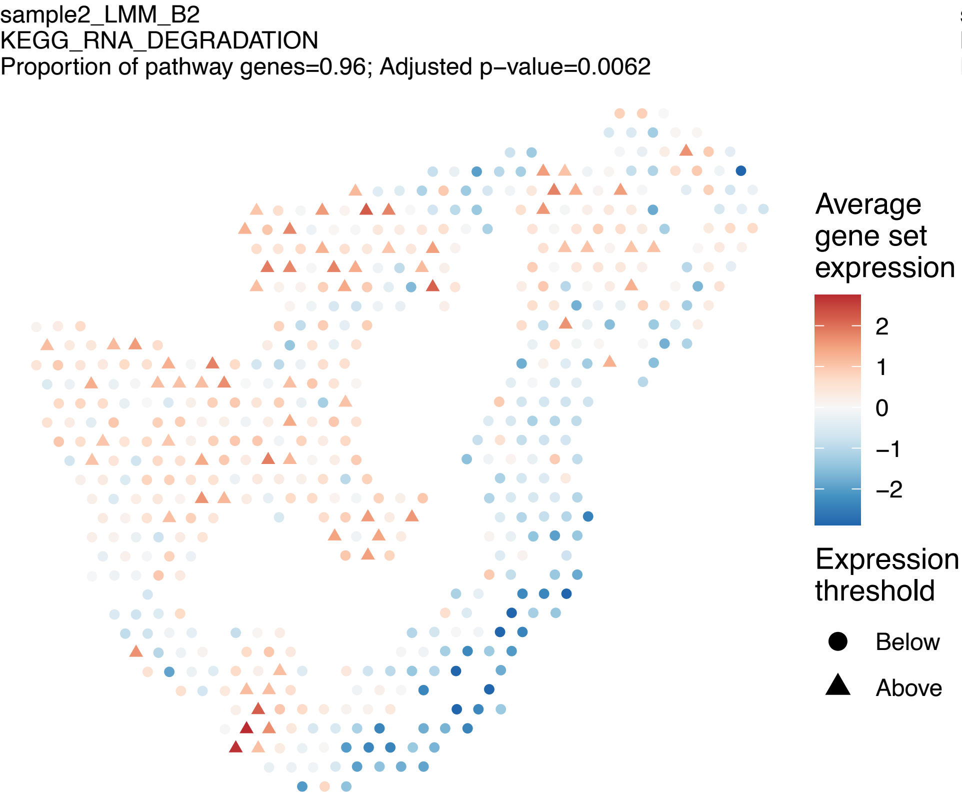
sample2_LMM_B2
KEGG_RNA_DEGRADATION
Proportion of pathway genes=0.96; Adjusted p−value=0.0062
Average gene set expression
2
1
0
−1
−2
Expression threshold
Below
Above

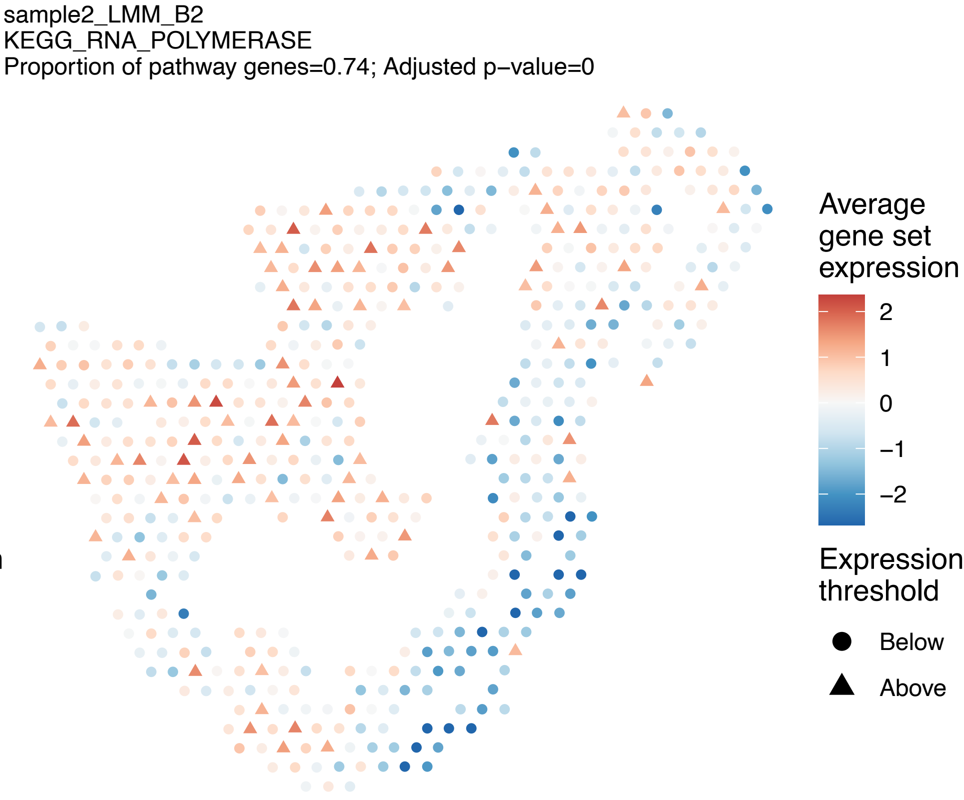
sample2_LMM_B2
KEGG_RNA_POLYMERASE
Proportion of pathway genes=0.74; Adjusted p−value=0
Average gene set expression
2
1
0
−1
−2
Expression threshold
Below
Above

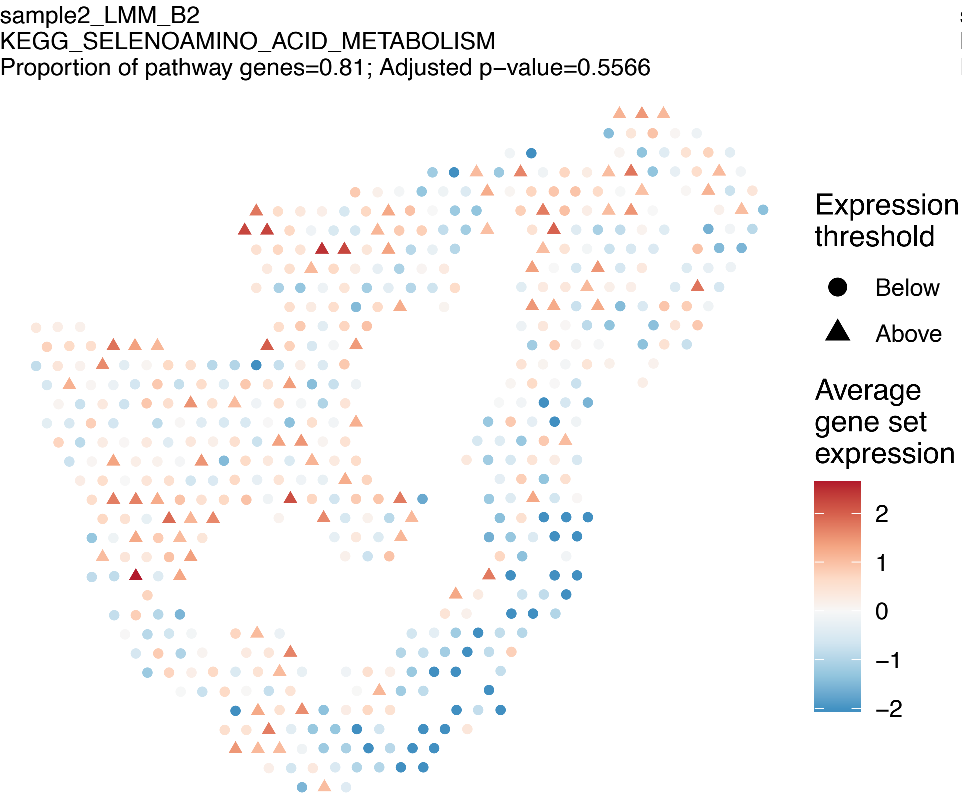
sample2_LMM_B2
KEGG_SELENOAMINO_ACID_METABOLISM
Proportion of pathway genes=0.81; Adjusted p−value=0.5566
Expression threshold
Below
Above
Average gene set expression
2
1
0
−1
−2

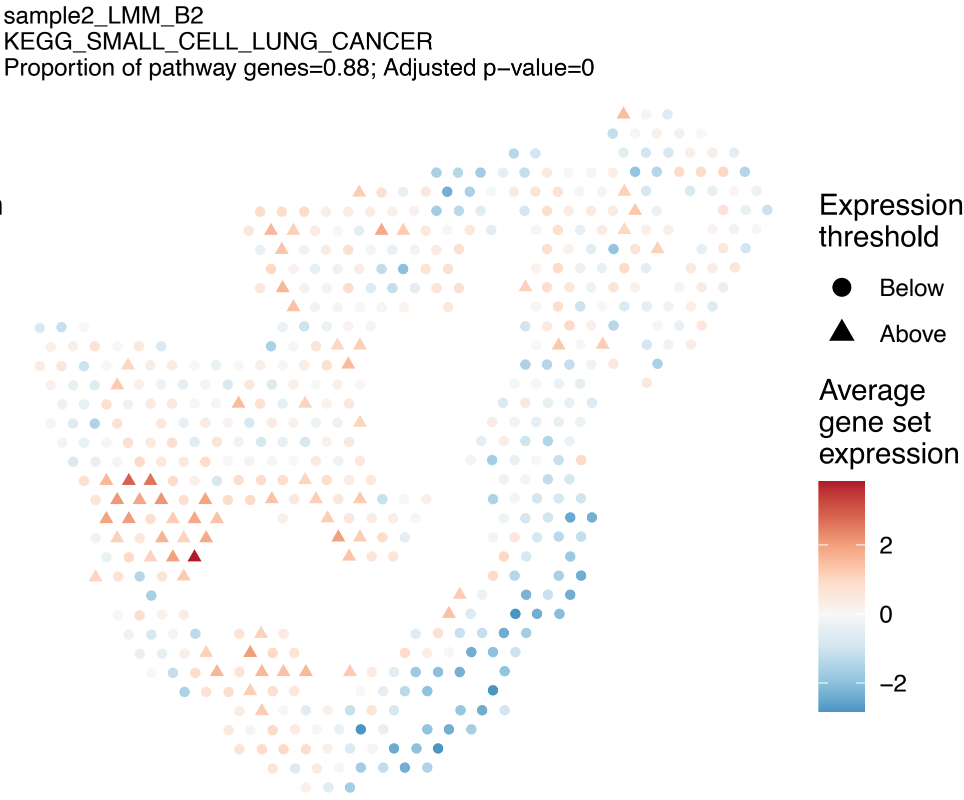
sample2_LMM_B2
KEGG_SMALL_CELL_LUNG_CANCER
Proportion of pathway genes=0.88; Adjusted p−value=0
Expression threshold
Below
Above
Average gene set expression
2
0
−2

sample2_LMM_B2
KEGG_SNARE_INTERACTIONS_IN_VESICULAR_TRANSPORT
Proportion of pathway genes=0.87; Adjusted p−value=0.1824

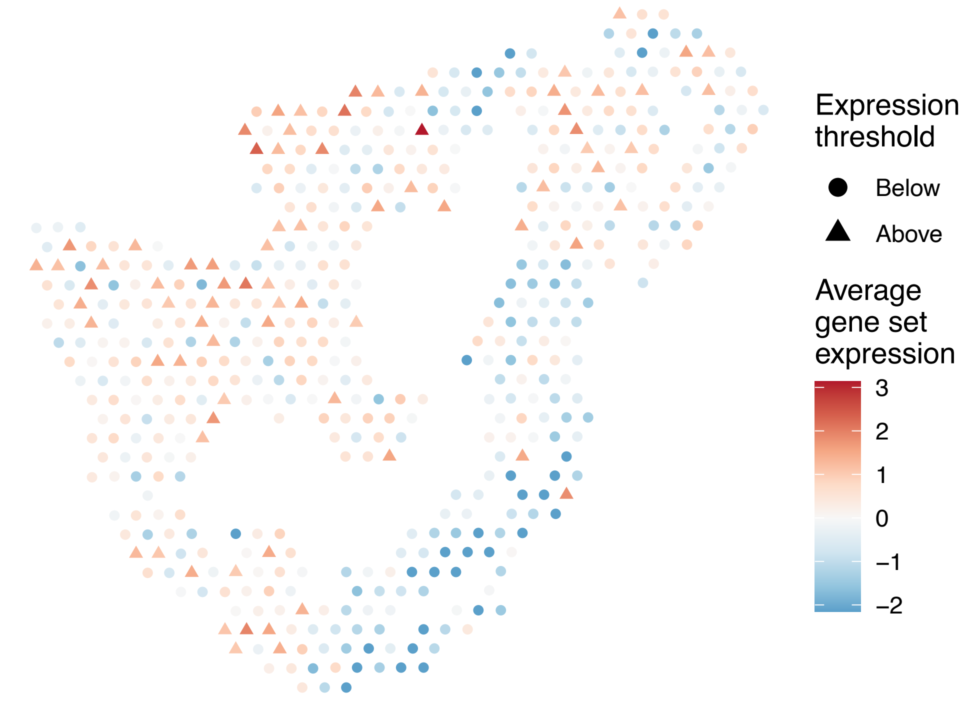


sample2_LMM_B2
KEGG_SPHINGOLIPID_METABOLISM
Proportion of pathway genes=0.74; Adjusted p−value=0.5281

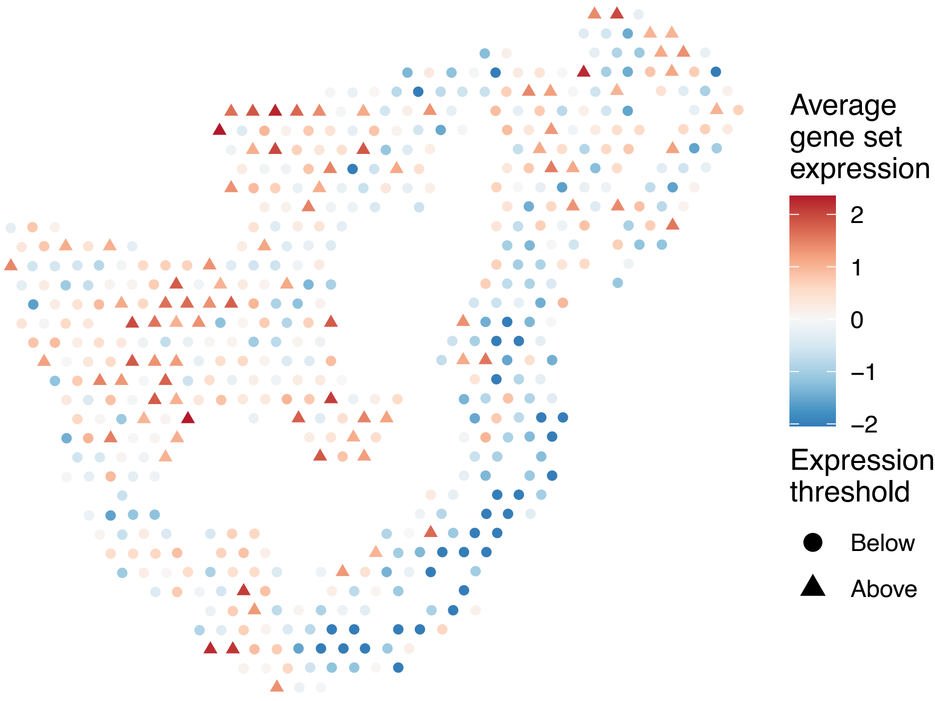


sample2_LMM_B2
KEGG_SPLICEOSOME
Proportion of pathway genes=0.95; Adjusted p−value=0.0806

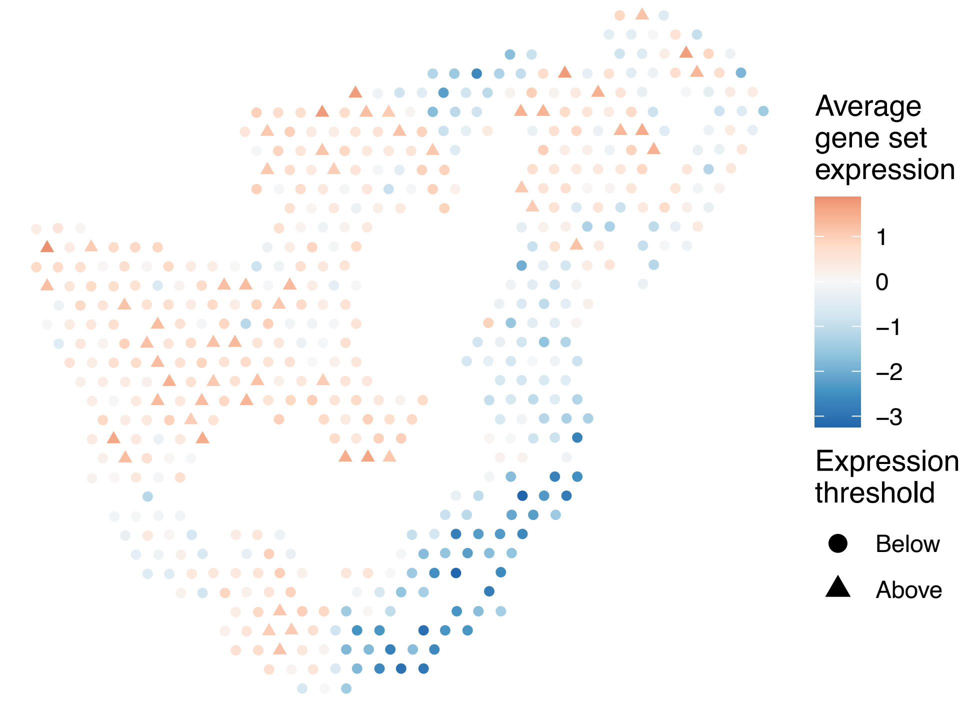


sample2_LMM_B2
KEGG_STARCH_AND_SUCROSE_METABOLISM
Proportion of pathway genes=0.34; Adjusted p−value=0.1717

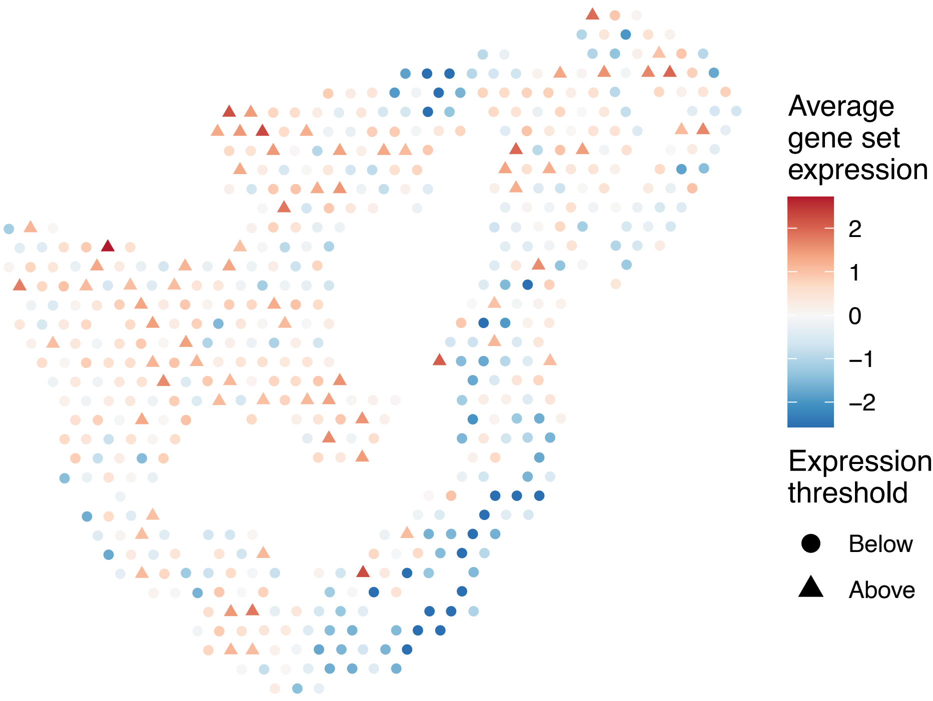

sample2_LMM_B2
KEGG_STEROID_BIOSYNTHESIS
Proportion of pathway genes=0.79; Adjusted p−value=0.9813

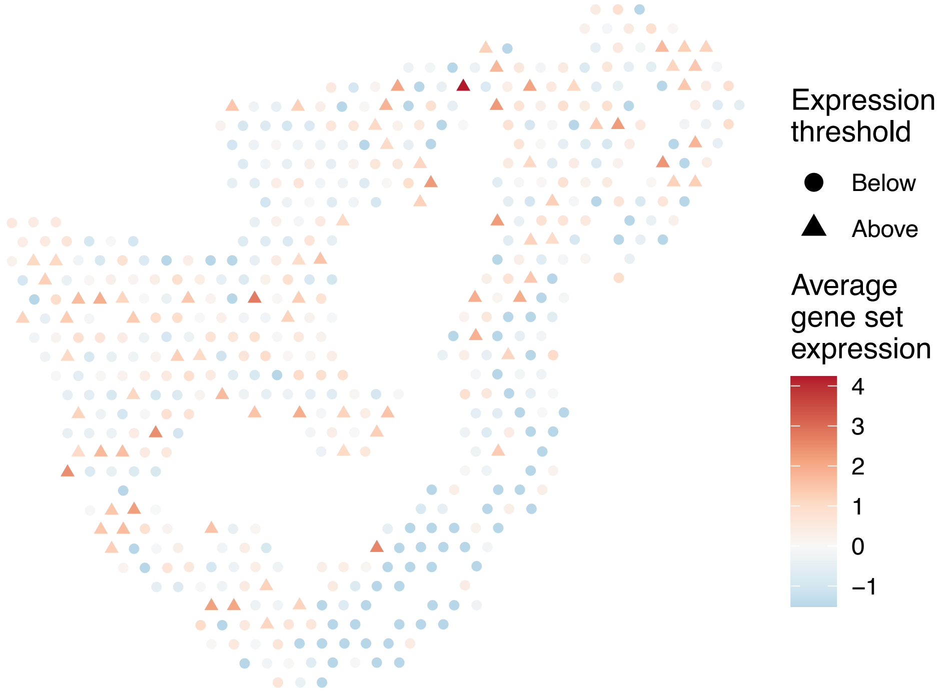


sample2_LMM_B2
KEGG_STEROID_HORMONE_BIOSYNTHESIS
Proportion of pathway genes=0.22; Adjusted p−value=0.2095

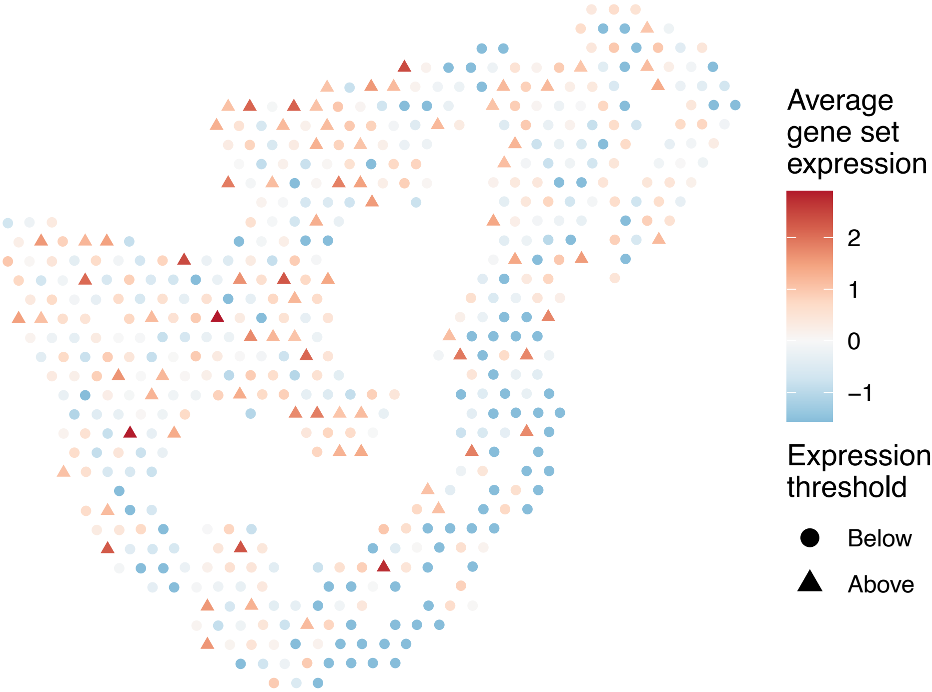


sample2_LMM_B2
KEGG_SULFUR_METABOLISM
Proportion of pathway genes=0.54; Adjusted p−value=NA

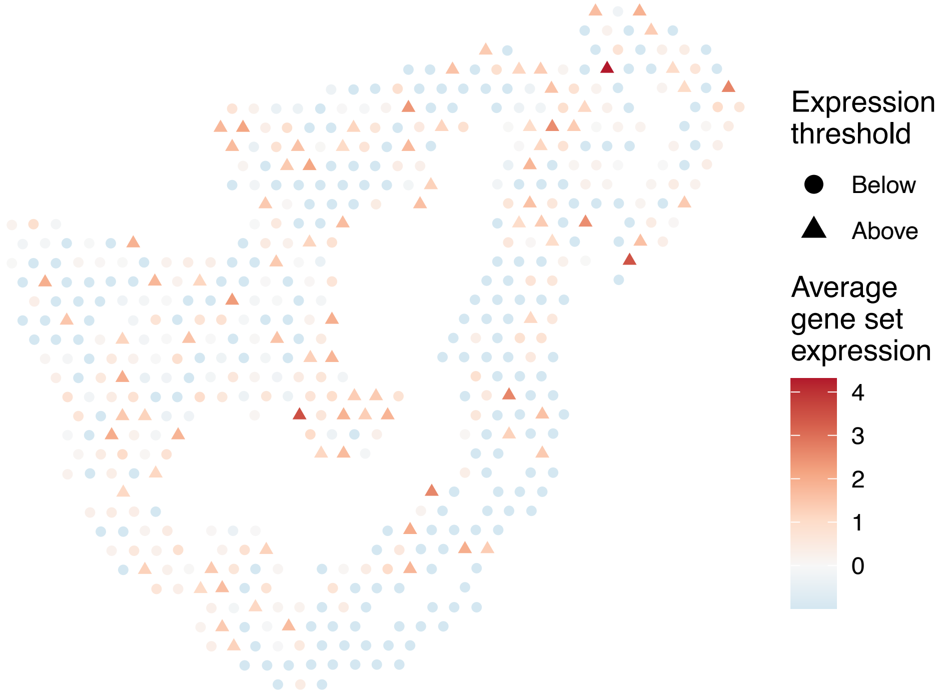


sample2_LMM_B2
KEGG_SYSTEMIC_LUPUS_ERYTHEMATOSUS
Proportion of pathway genes=0.34; Adjusted p−value=0.1522

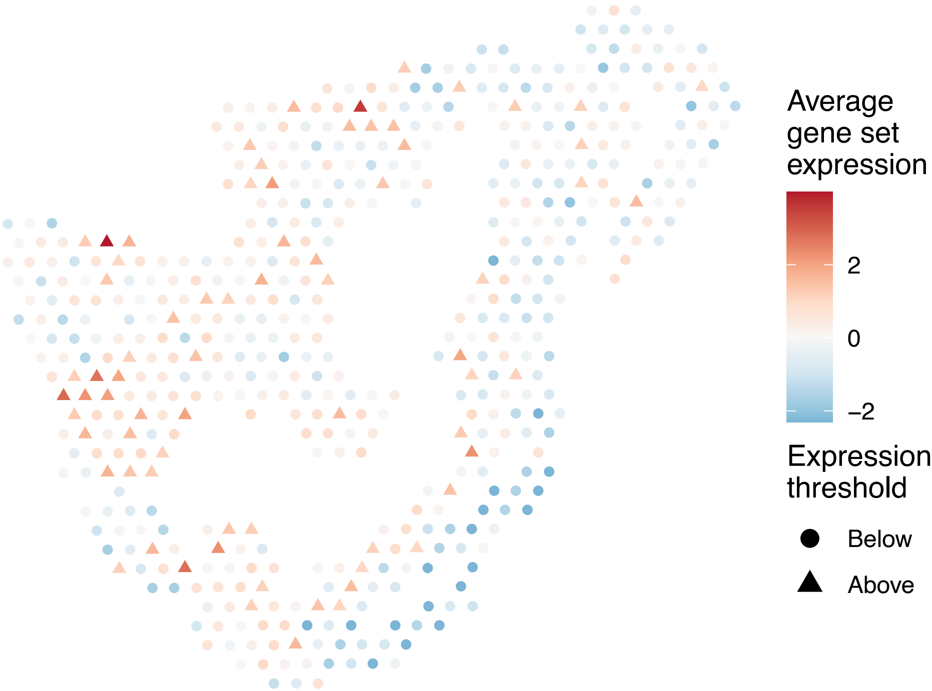

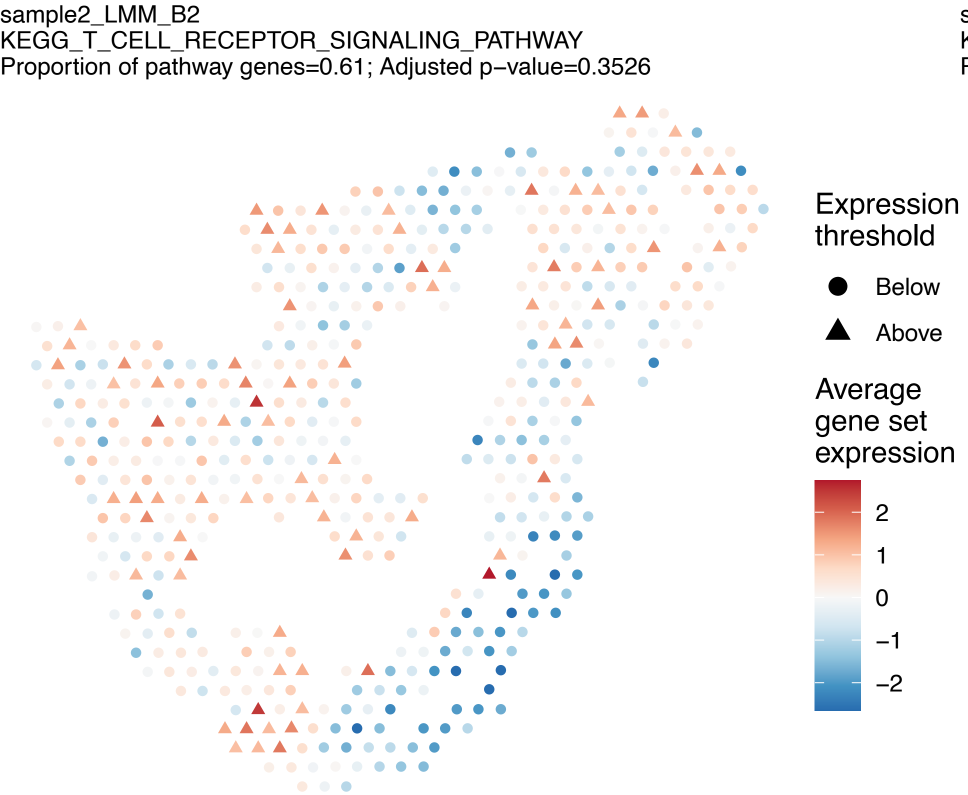
sample2_LMM_B2
KEGG_T_CELL_RECEPTOR_SIGNALING_PATHWAY
Proportion of pathway genes=0.61; Adjusted p−value=0.3526
Expression threshold
Below
Above
Average gene set expression
2
1
0
−1
−2

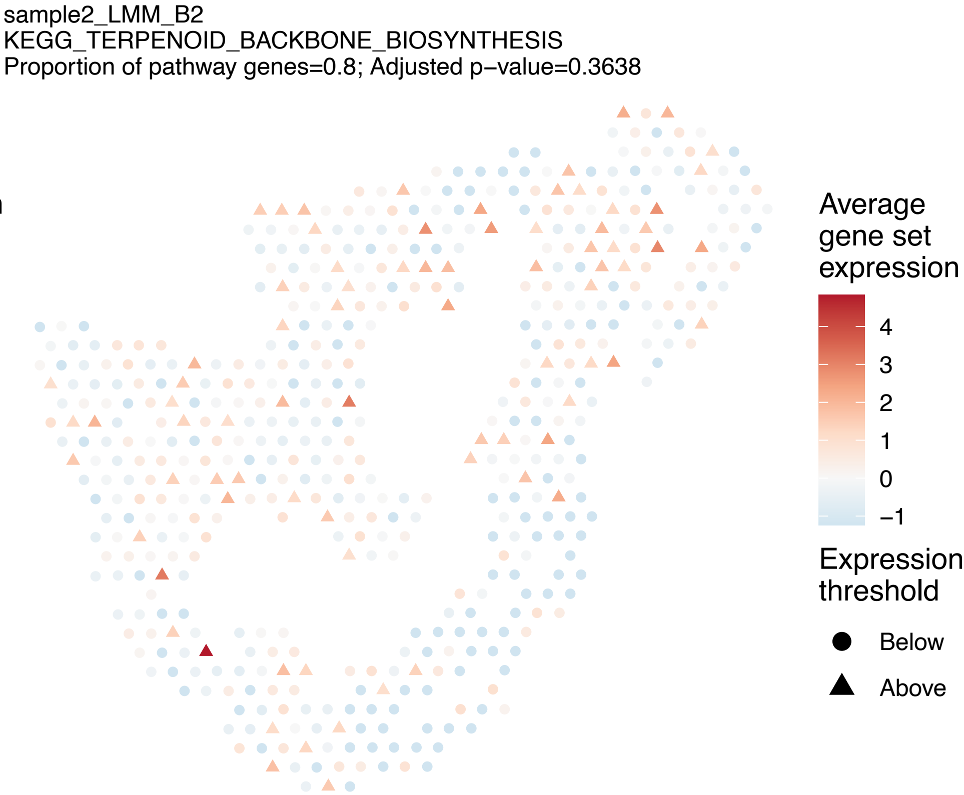
sample2_LMM_B2
KEGG_TERPENOID_BACKBONE_BIOSYNTHESIS
Proportion of pathway genes=0.8; Adjusted p−value=0.3638
Average gene set expression
4
3
2
1
0
−1
Expression threshold
Below
Above

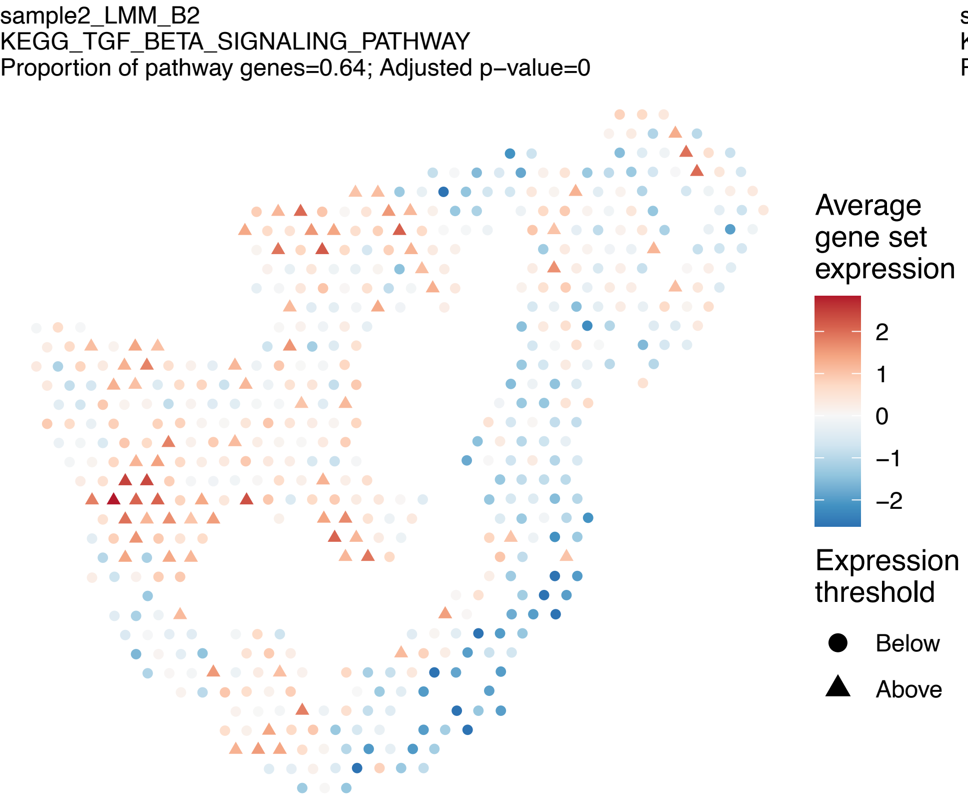
sample2_LMM_B2
KEGG_TGF_BETA_SIGNALING_PATHWAY
Proportion of pathway genes=0.64; Adjusted p−value=0
Average gene set expression
2
1
0
−1
−2
Expression threshold
Below
Above

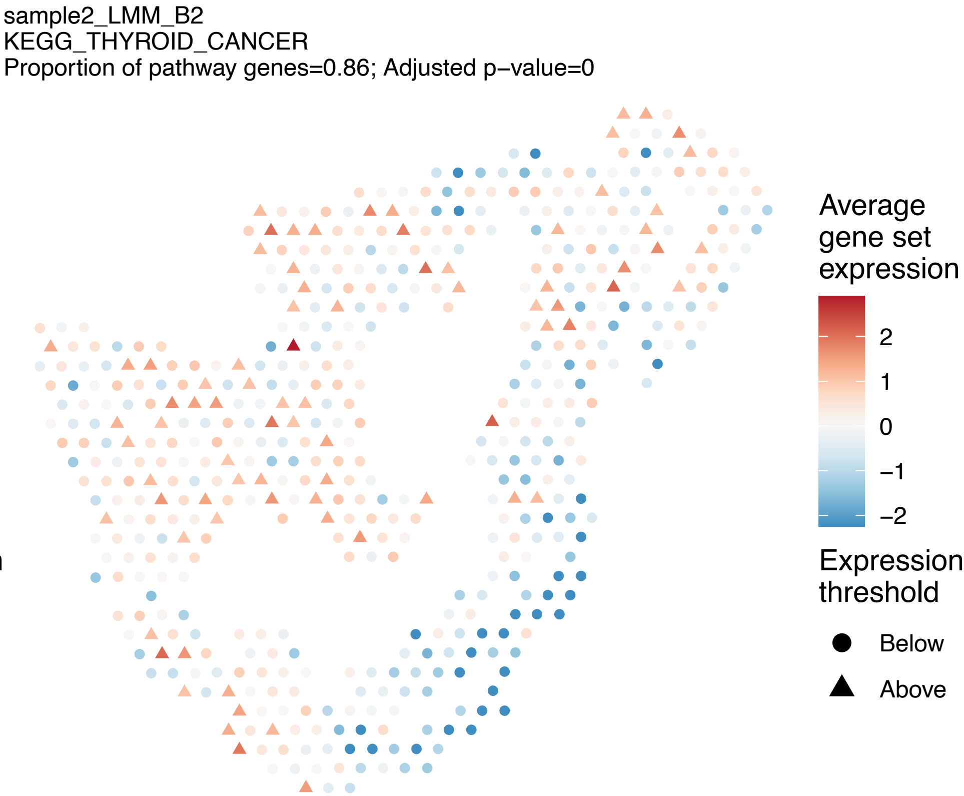
sample2_LMM_B2
KEGG_THYROID_CANCER
Proportion of pathway genes=0.86; Adjusted p−value=0
Average gene set expression
2
1
0
−1
−2
Expression threshold
Below
Above

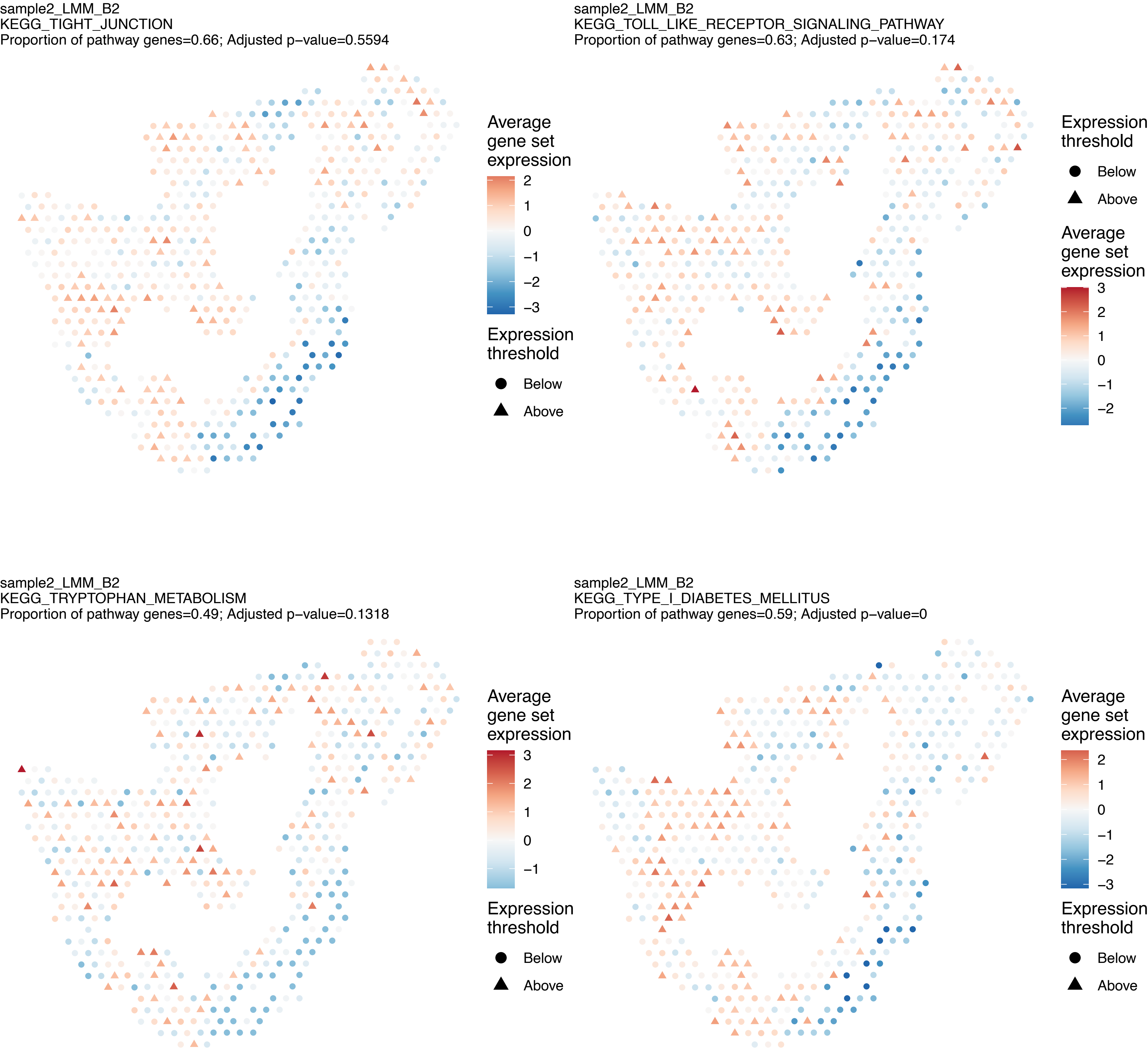
sample2_LMM_B2
KEGG_TIGHT_JUNCTION
Proportion of pathway genes=0.66; Adjusted p−value=0.5594
Average gene set expression
2
1
0
−1
−2
−3
Expression threshold
Below
Above
sample2_LMM_B2
KEGG_TOLL_LIKE_RECEPTOR_SIGNALING_PATHWAY
Proportion of pathway genes=0.63; Adjusted p−value=0.174
Expression threshold
Below
Above
Average gene set expression
3
2
1
0
−1
−2
sample2_LMM_B2
KEGG_TRYPTOPHAN_METABOLISM
Proportion of pathway genes=0.49; Adjusted p−value=0.1318
Average gene set expression
3
2
1
0
−1
Expression threshold
Below
Above
sample2_LMM_B2
KEGG_TYPE_I_DIABETES_MELLITUS
Proportion of pathway genes=0.59; Adjusted p−value=0
Average gene set expression
2
1
0
−1
−2
−3
Expression threshold
Below
Above

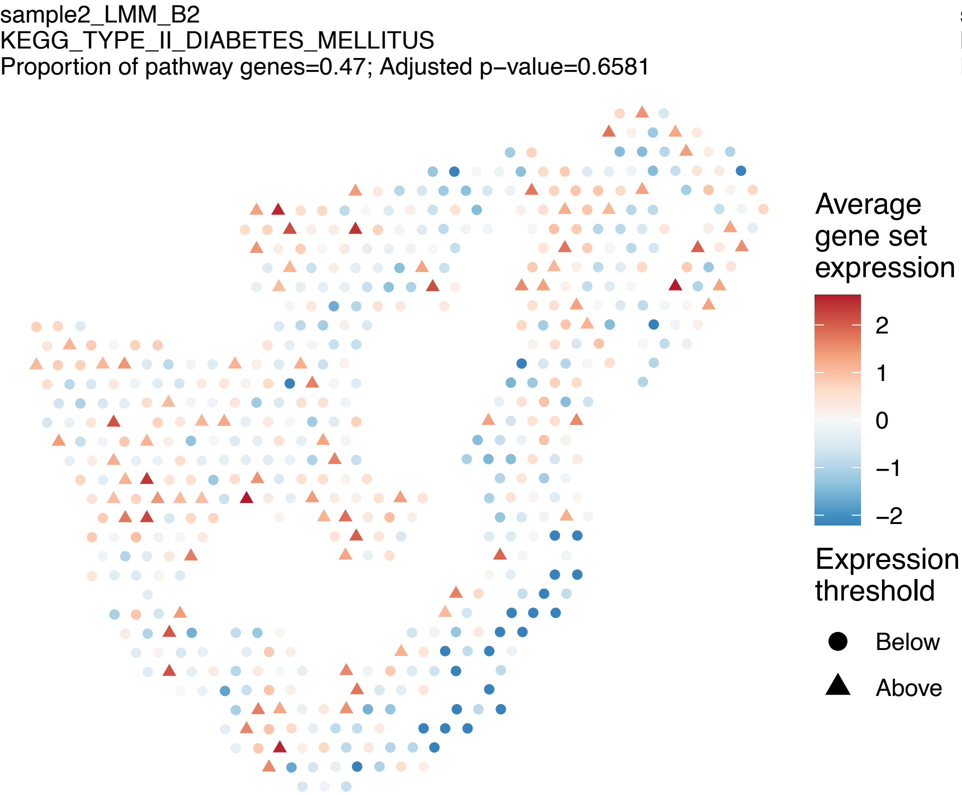
sample2_LMM_B2
KEGG_TYPE_II_DIABETES_MELLITUS
Proportion of pathway genes=0.47; Adjusted p−value=0.6581
Average gene set expression
2
1
0
−1
−2
Expression threshold
Below
Above

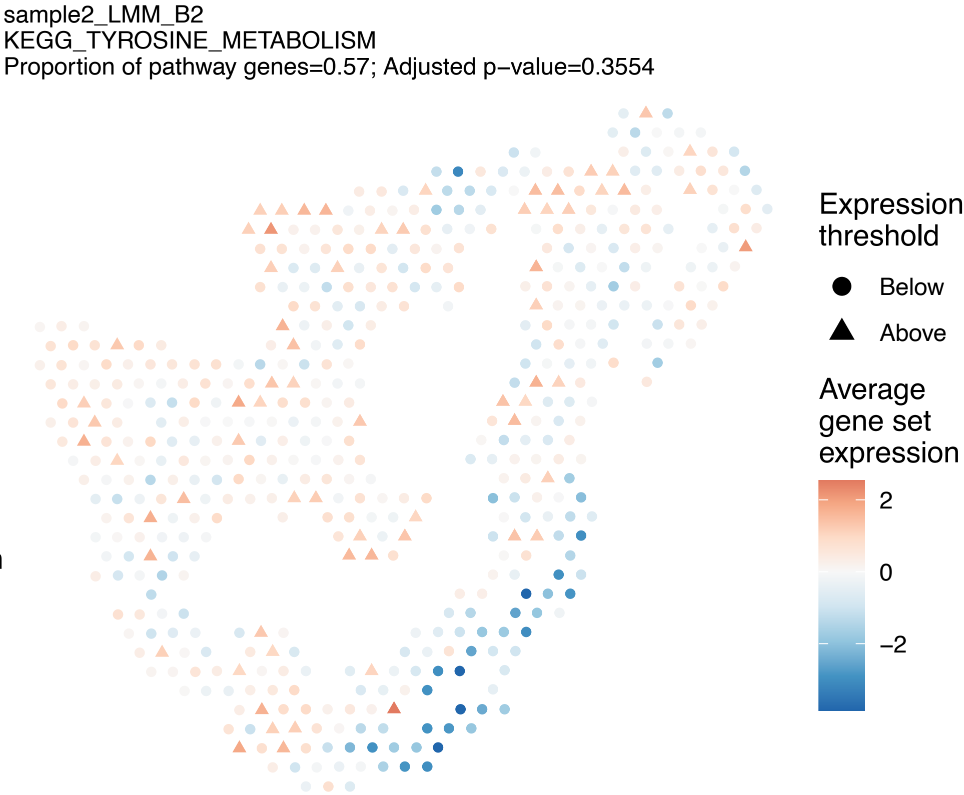
sample2_LMM_B2
KEGG_TYROSINE_METABOLISM
Proportion of pathway genes=0.57; Adjusted p−value=0.3554
Expression threshold
Below
Above
Average gene set expression
2
0
−2

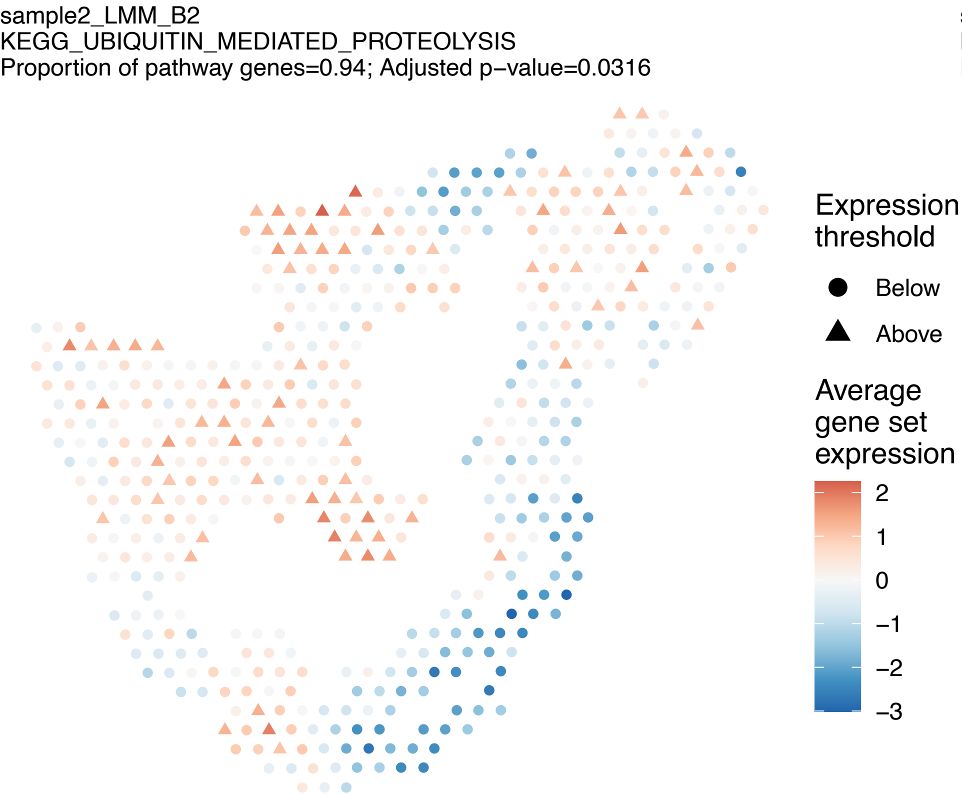
sample2_LMM_B2
KEGG_UBIQUITIN_MEDIATED_PROTEOLYSIS
Proportion of pathway genes=0.94; Adjusted p−value=0.0316
Expression threshold
Below
Above
Average gene set expression
2
1
0
−1
−2
−3

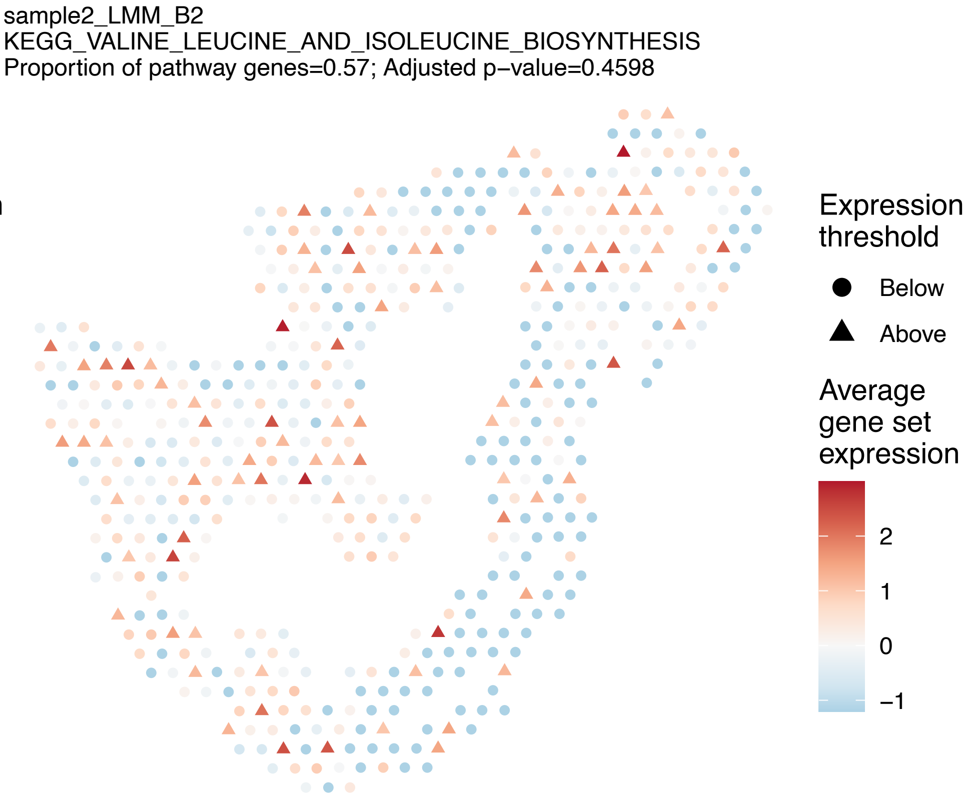
sample2_LMM_B2
KEGG_VALINE_LEUCINE_AND_ISOLEUCINE_BIOSYNTHESIS
Proportion of pathway genes=0.57; Adjusted p−value=0.4598
Expression threshold
Below
Above
Average gene set expression
2
1
0
−1

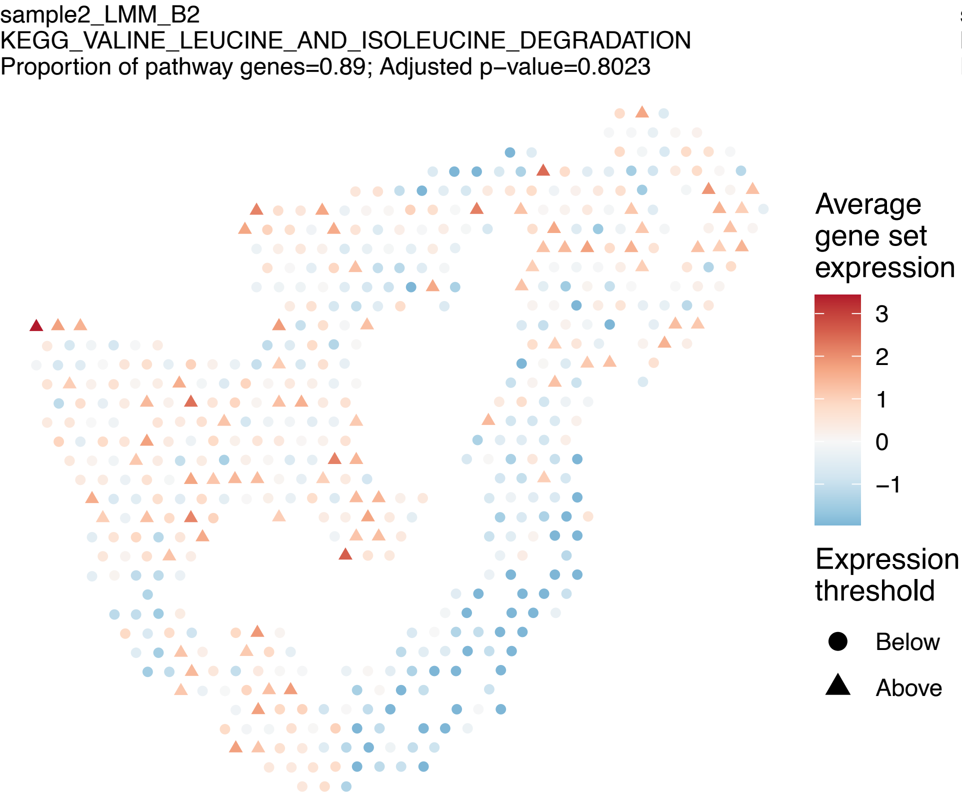
sample2_LMM_B2
KEGG_VALINE_LEUCINE_AND_ISOLEUCINE_DEGRADATION
Proportion of pathway genes=0.89; Adjusted p−value=0.8023
Average gene set expression
3
2
1
0
−1
Expression threshold
Below
Above

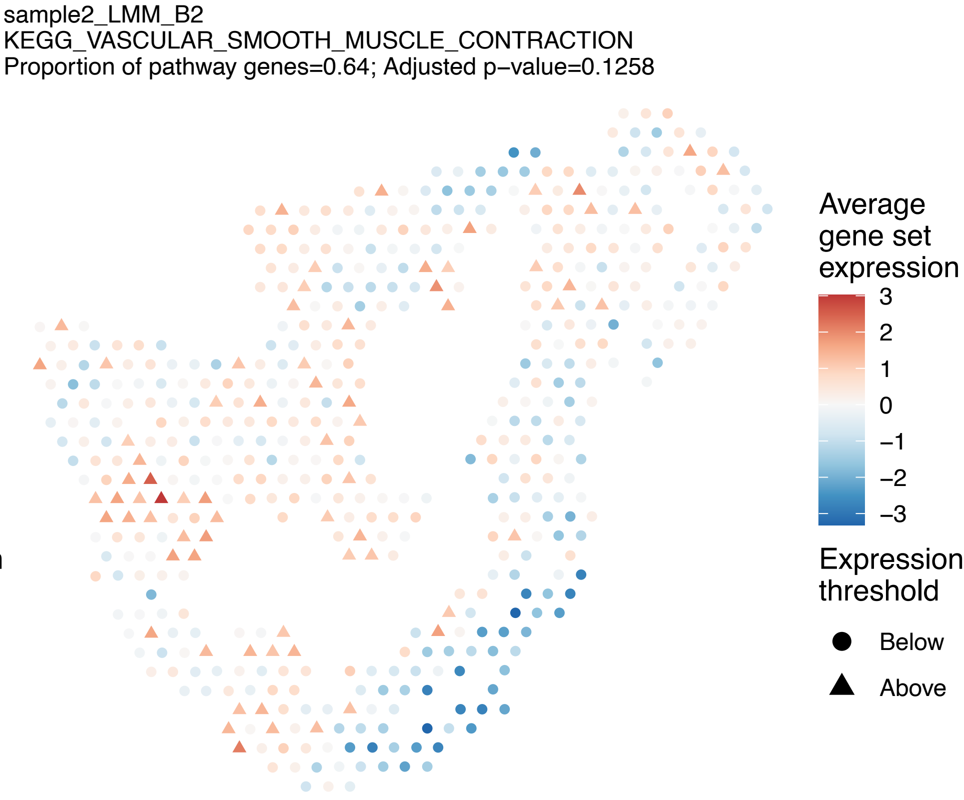
sample2_LMM_B2
KEGG_VASCULAR_SMOOTH_MUSCLE_CONTRACTION
Proportion of pathway genes=0.64; Adjusted p−value=0.1258
Average gene set expression
3
2
1
0
−1
−2
−3
Expression threshold
Below
Above

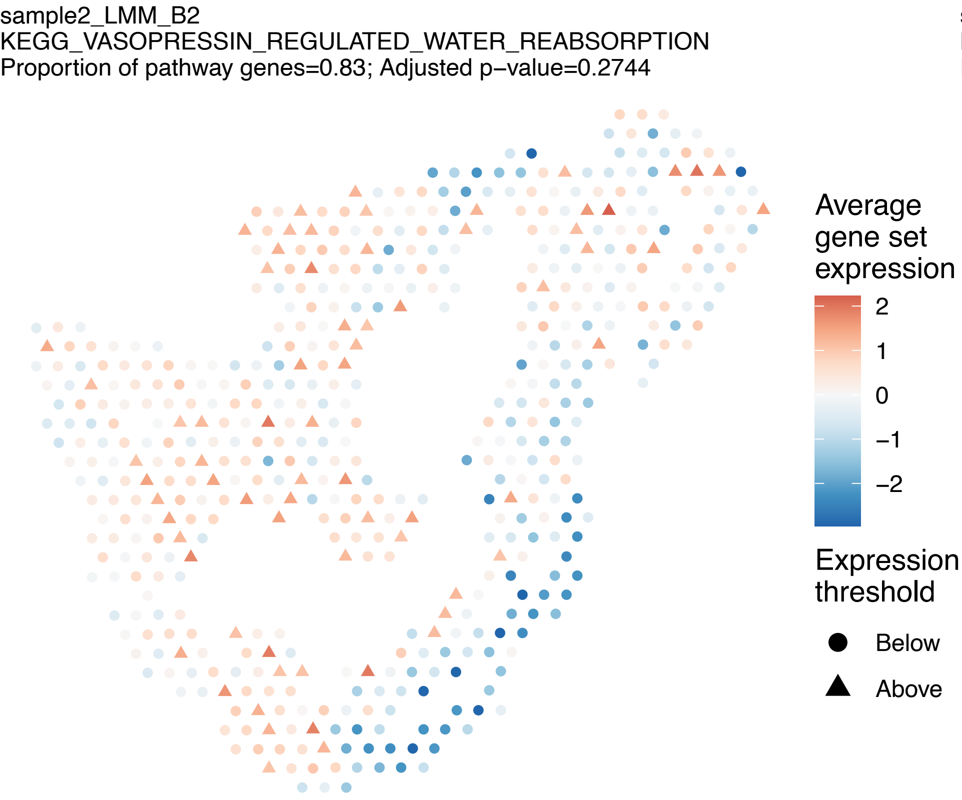
sample2_LMM_B2
KEGG_VASOPRESSIN_REGULATED_WATER_REABSORPTION
Proportion of pathway genes=0.83; Adjusted p−value=0.2744
Average gene set expression
2
1
0
−1
−2
Expression threshold
Below
Above

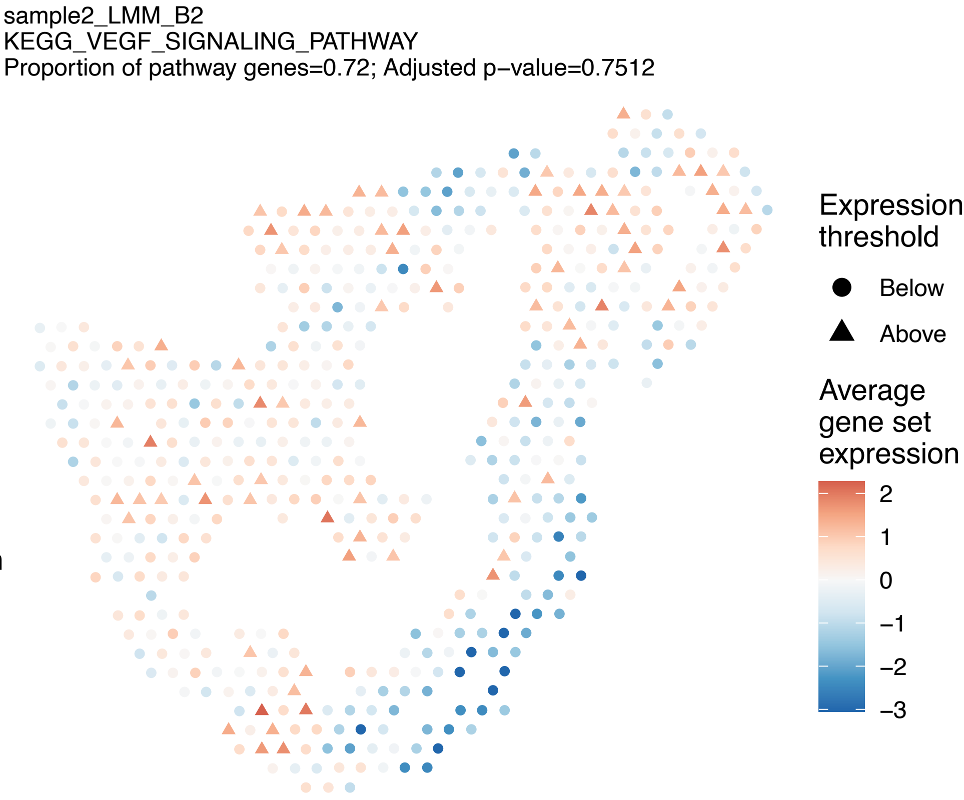
sample2_LMM_B2
KEGG_VEGF_SIGNALING_PATHWAY
Proportion of pathway genes=0.72; Adjusted p−value=0.7512
Expression threshold
Below
Above
Average gene set expression
2
1
0
−1
−2
−3

sample2_LMM_B2
KEGG_VIBRIO_CHOLERAE_INFECTION
Proportion of pathway genes=0.67; Adjusted p−value=0.302

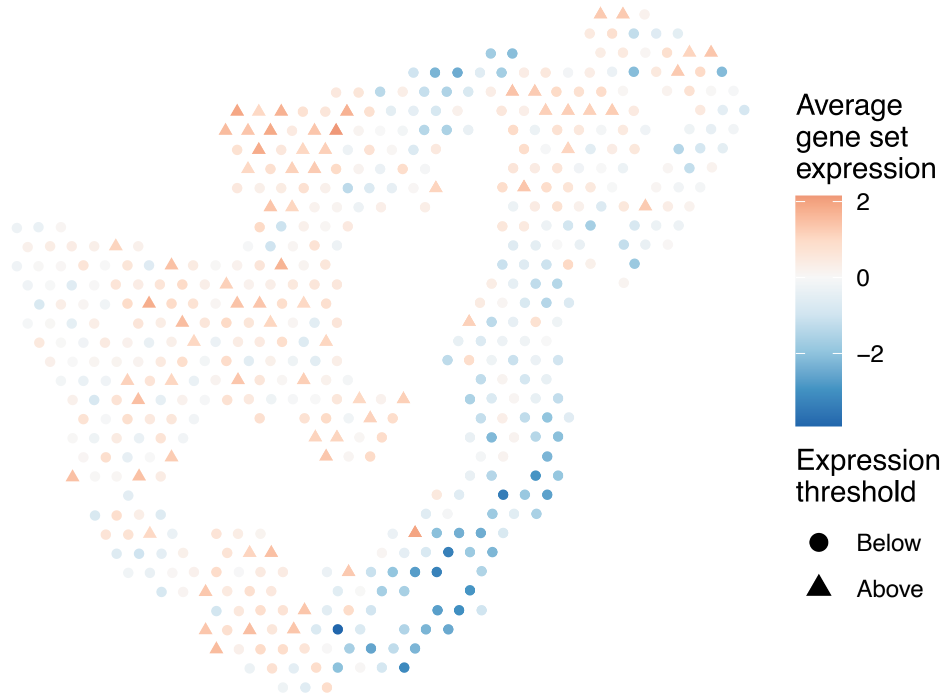


sample2_LMM_B2
KEGG_VIRAL_MYOCARDITIS
Proportion of pathway genes=0.7; Adjusted p−value=0

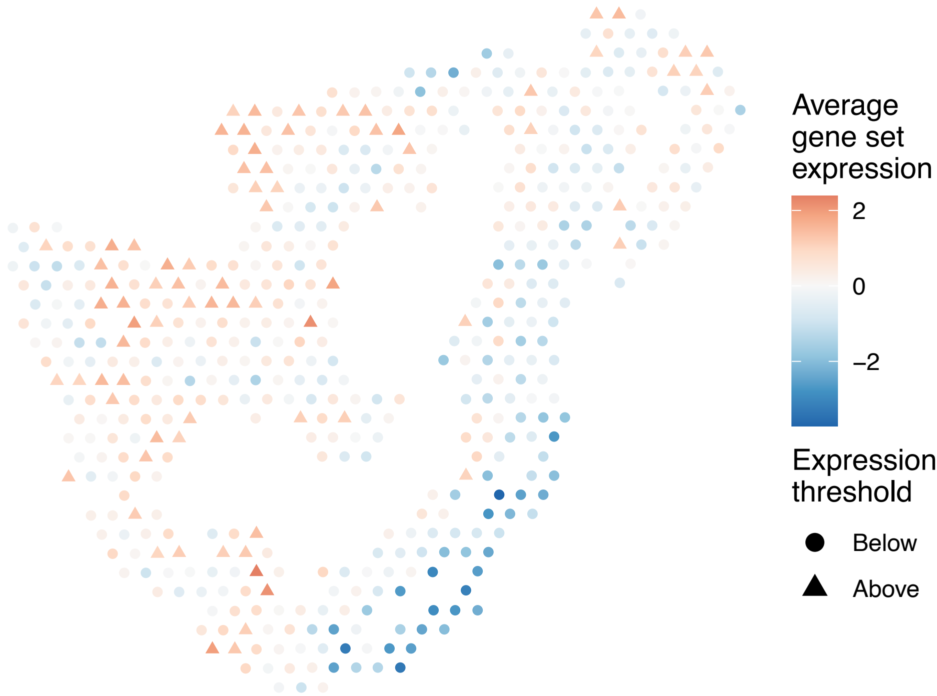


sample2_LMM_B2
KEGG_WNT_SIGNALING_PATHWAY
Proportion of pathway genes=0.71; Adjusted p−value=0.4005

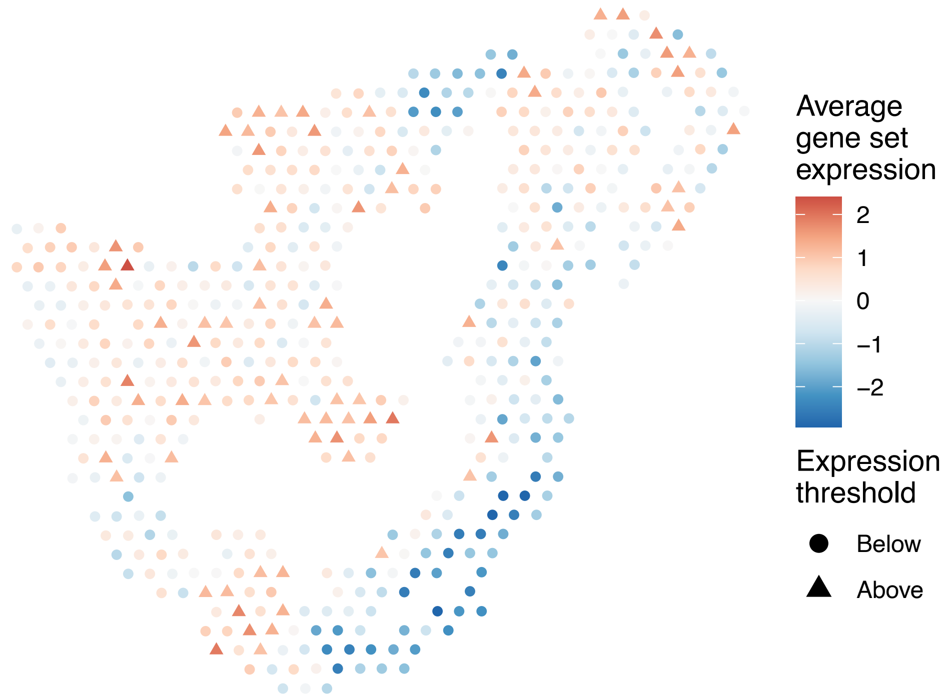


sample2_LMM_B2
KEGG_PENTOSE_AND_GLUCURONATE_INTERCONVERSIONS
Proportion of pathway genes=0.29; Adjusted p−value=NA

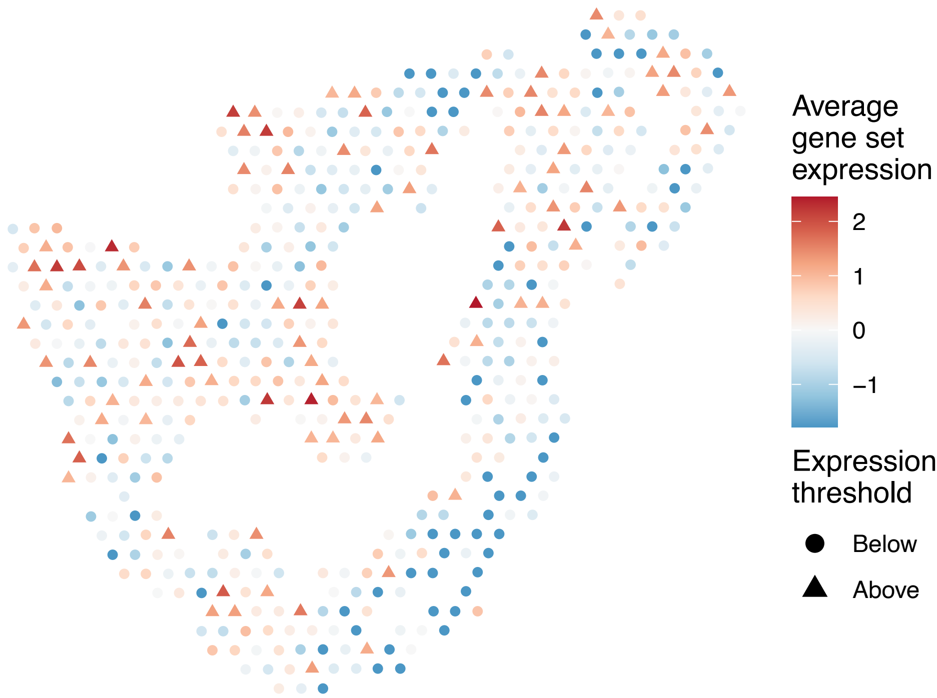

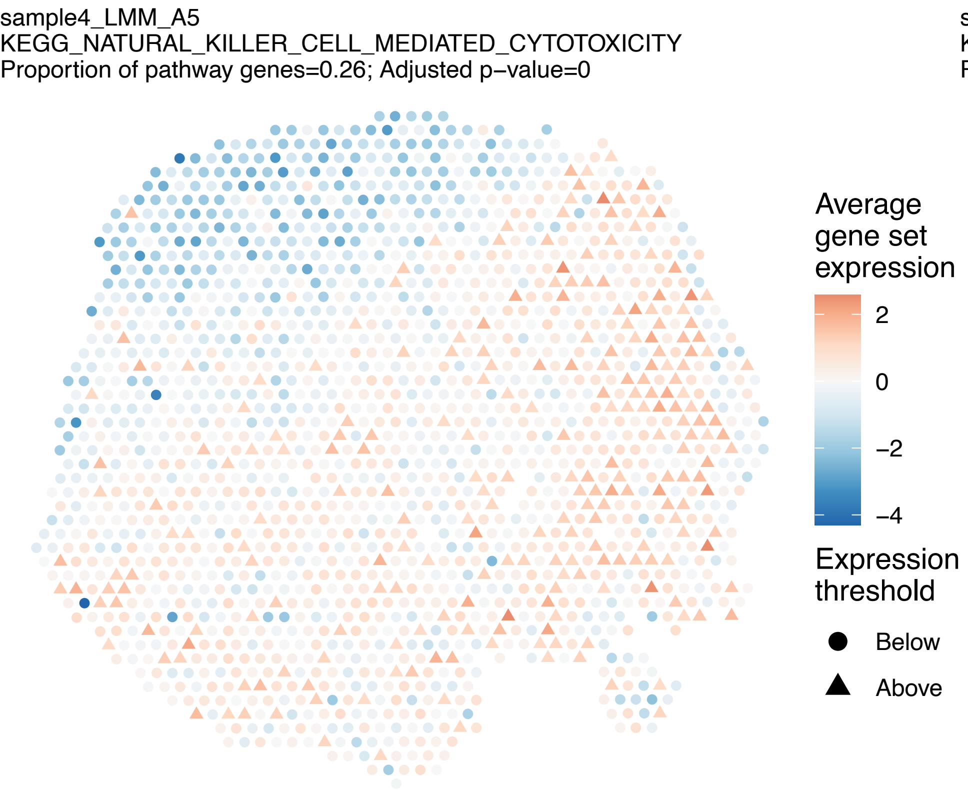
sample4_LMM_A5
KEGG_NATURAL_KILLER_CELL_MEDIATED_CYTOTOXICITY
Proportion of pathway genes=0.26; Adjusted p−value=0
Average gene set expression
2
0
−2
−4
Expression threshold
Below
Above

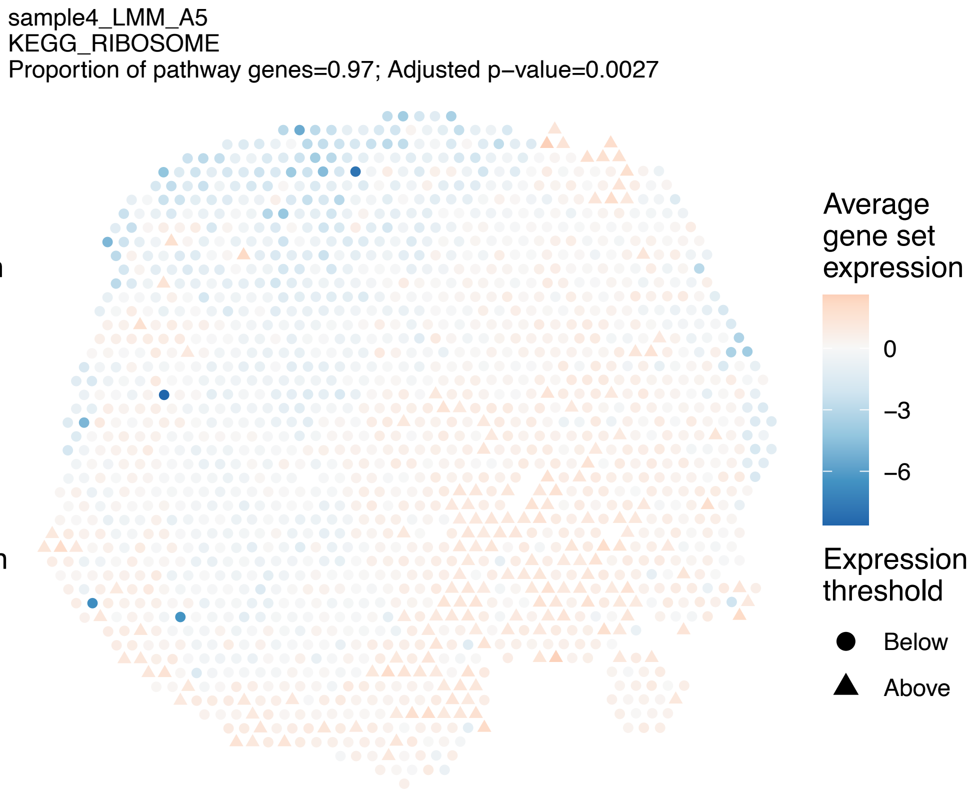
sample4_LMM_A5
KEGG_RIBOSOME
Proportion of pathway genes=0.97; Adjusted p−value=0.0027
Average gene set expression
0
−3
−6
Expression threshold
Below
Above

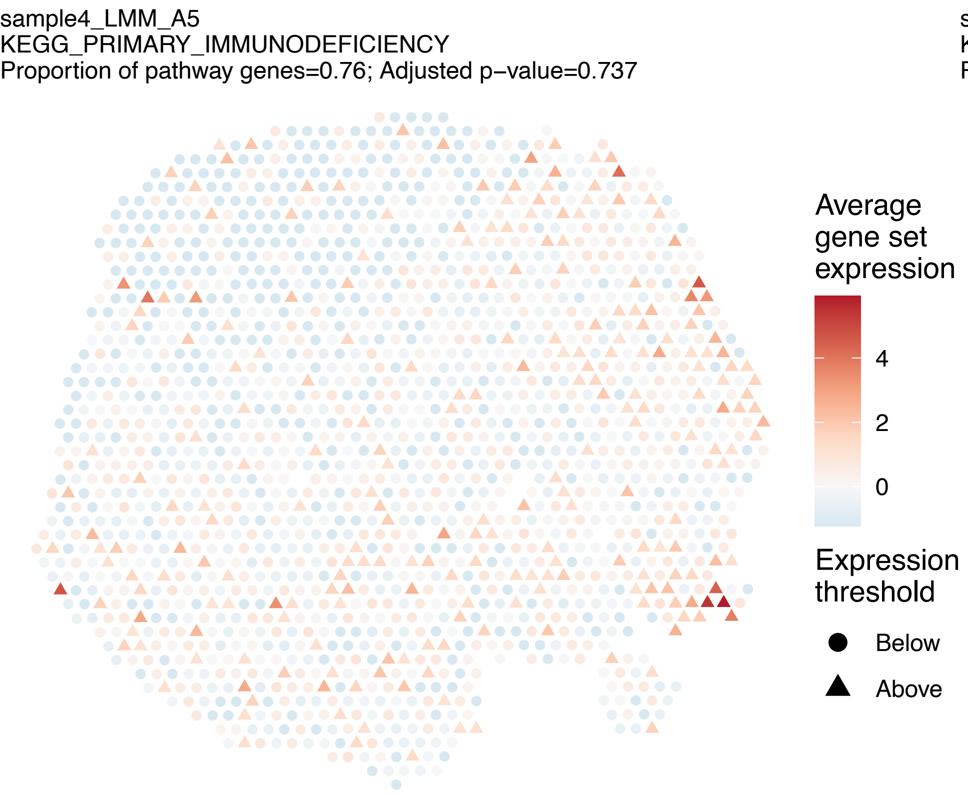
sample4_LMM_A5
KEGG_PRIMARY_IMMUNODEFICIENCY
Proportion of pathway genes=0.76; Adjusted p−value=0.737
Average gene set expression
4
2
0
Expression threshold
Below
Above

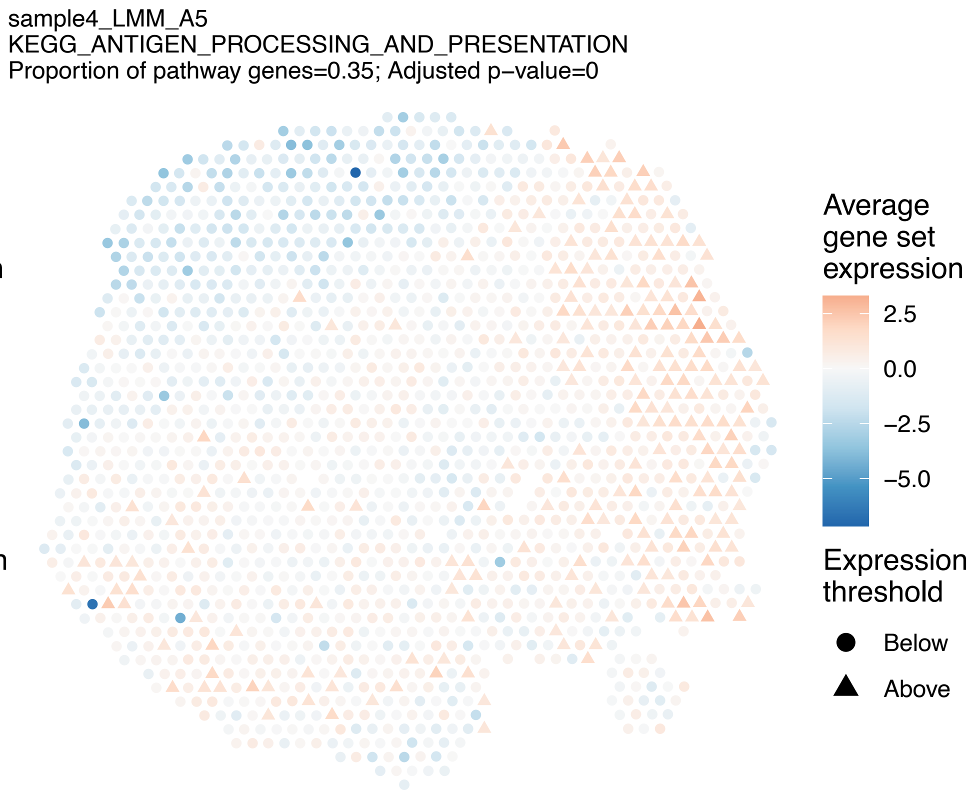
sample4_LMM_A5
KEGG_ANTIGEN_PROCESSING_AND_PRESENTATION
Proportion of pathway genes=0.35; Adjusted p−value=0
Average gene set expression
2.5
0.0
−2.5
−5.0
Expression threshold
Below
Above

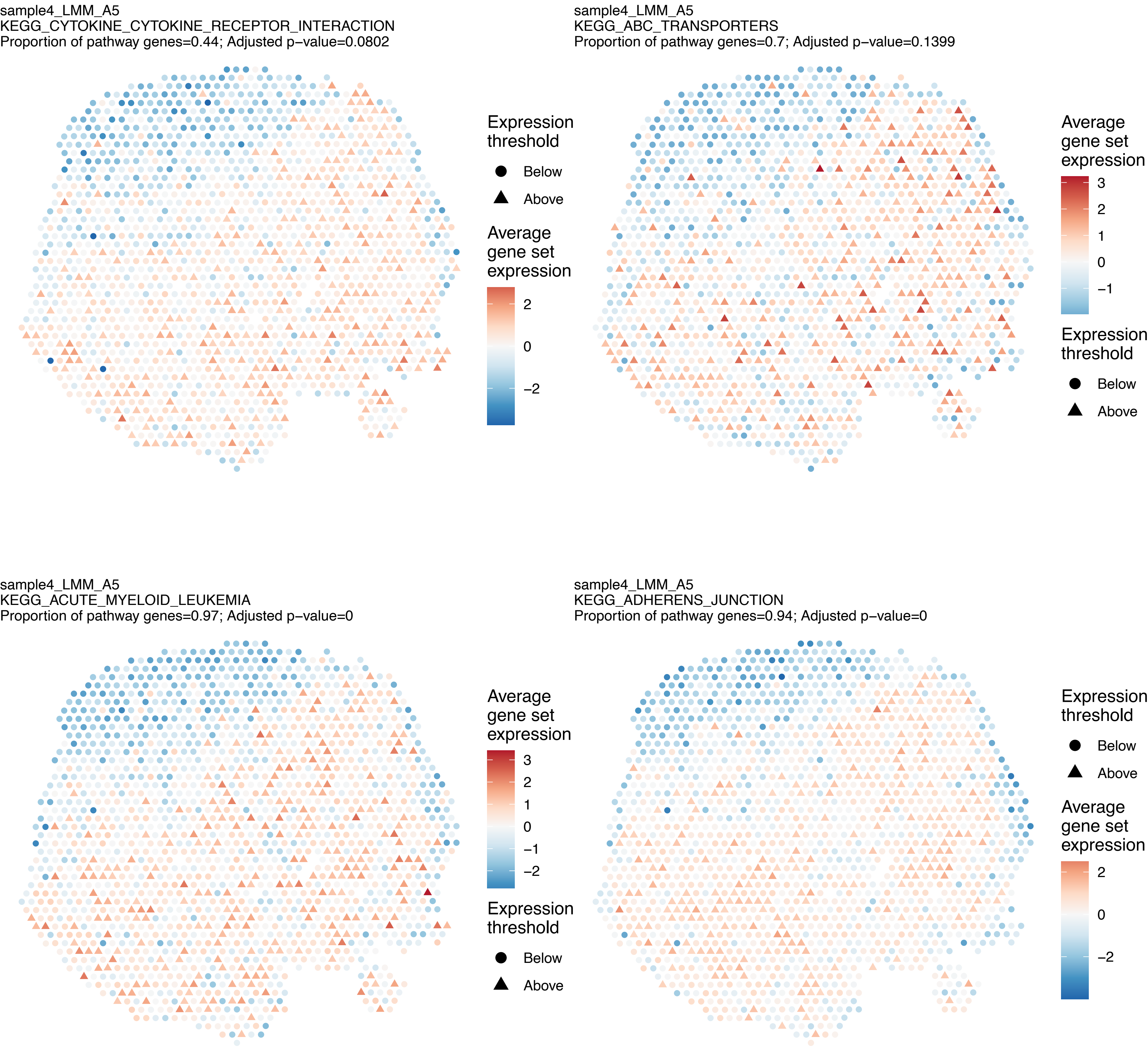
sample4_LMM_A5
KEGG_CYTOKINE_CYTOKINE_RECEPTOR_INTERACTION
Proportion of pathway genes=0.44; Adjusted p−value=0.0802
Expression threshold
Below
Above
Average gene set expression
2
0
−2
sample4_LMM_A5
KEGG_ABC_TRANSPORTERS
Proportion of pathway genes=0.7; Adjusted p−value=0.1399
Average gene set expression
3
2
1
0
−1
Expression threshold
Below
Above
sample4_LMM_A5
KEGG_ACUTE_MYELOID_LEUKEMIA
Proportion of pathway genes=0.97; Adjusted p−value=0
Average gene set expression
3
2
1
0
−1
−2
Expression threshold
Below
Above
sample4_LMM_A5
KEGG_ADHERENS_JUNCTION
Proportion of pathway genes=0.94; Adjusted p−value=0
Expression threshold
Below
Above
Average gene set expression
2
0
−2

sample4_LMM_A5
KEGG_ADIPOCYTOKINE_SIGNALING_PATHWAY
Proportion of pathway genes=0.75; Adjusted p−value=0.0075

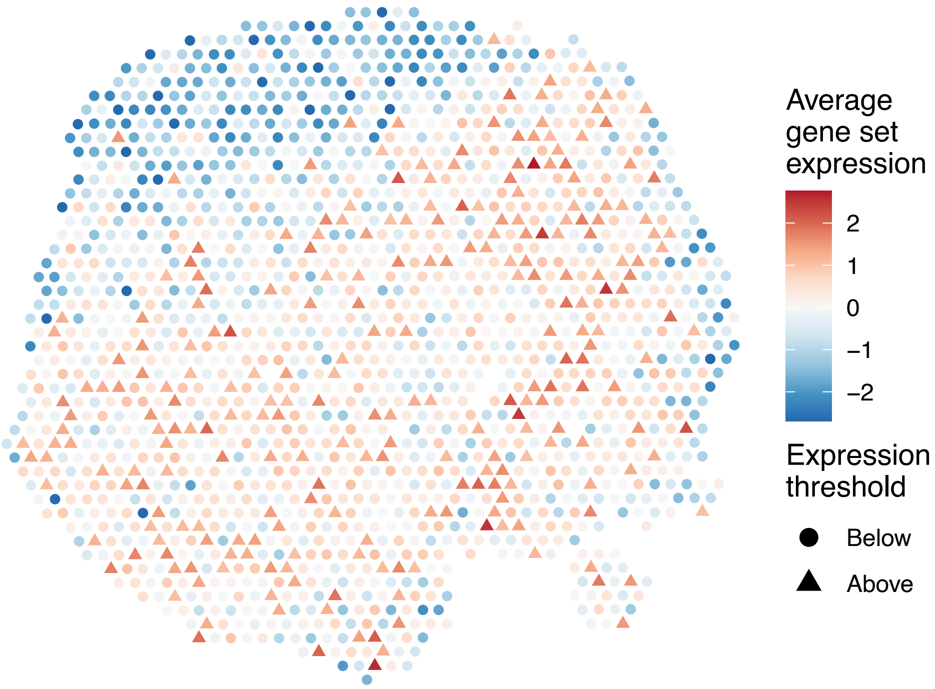


sample4_LMM_A5
KEGG_ALANINE_ASPARTATE_AND_GLUTAMATE_METABOLISM
Proportion of pathway genes=0.7; Adjusted p−value=0

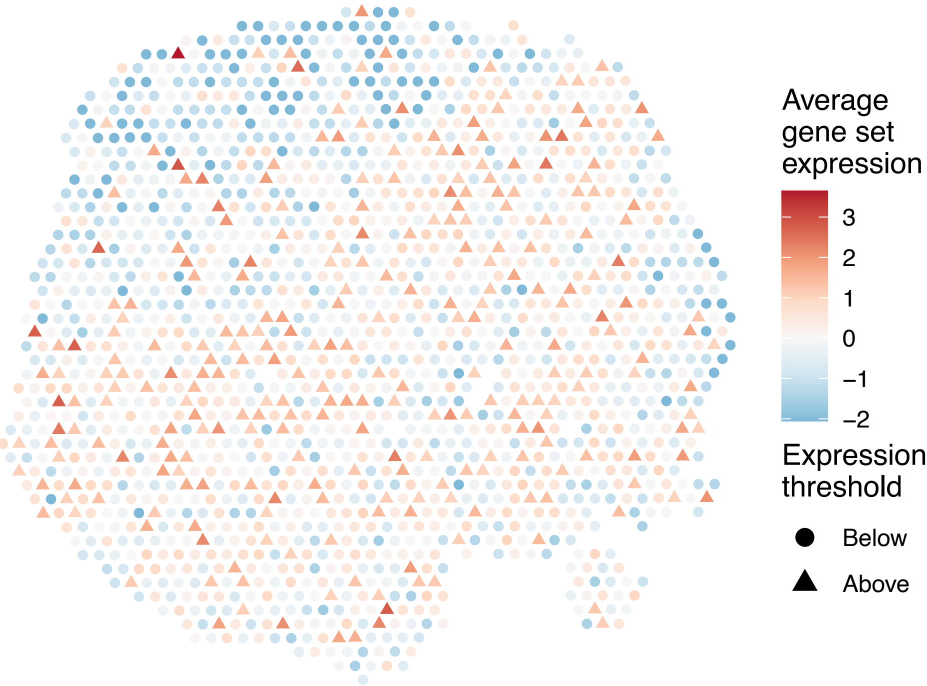


sample4_LMM_A5
KEGG_ALDOSTERONE_REGULATED_SODIUM_REABSORPTION
Proportion of pathway genes=0.67; Adjusted p−value=0

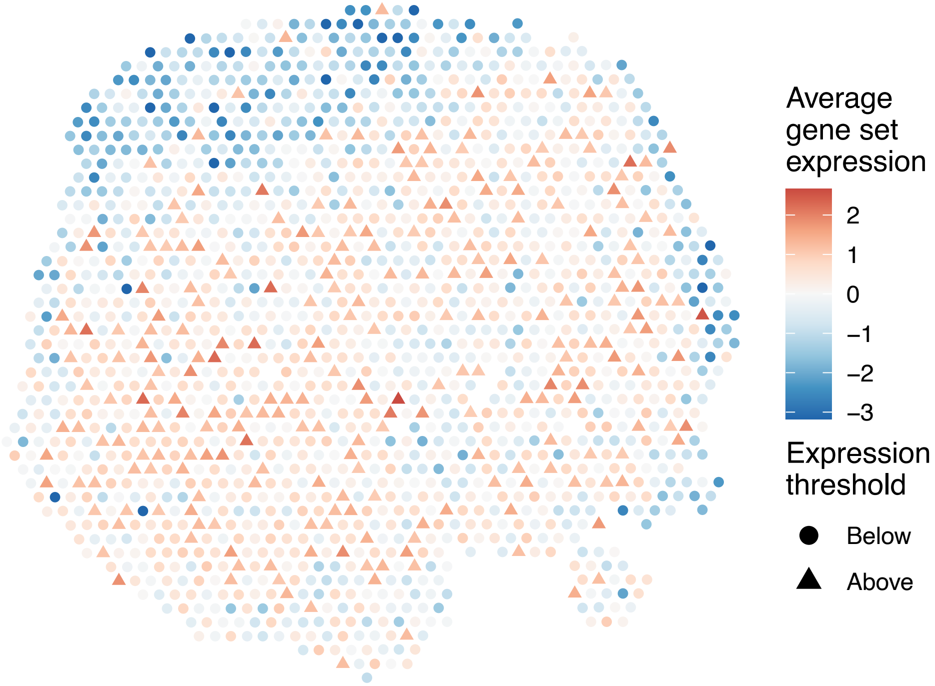


sample4_LMM_A5
KEGG_ALLOGRAFT_REJECTION
Proportion of pathway genes=0.76; Adjusted p−value=0

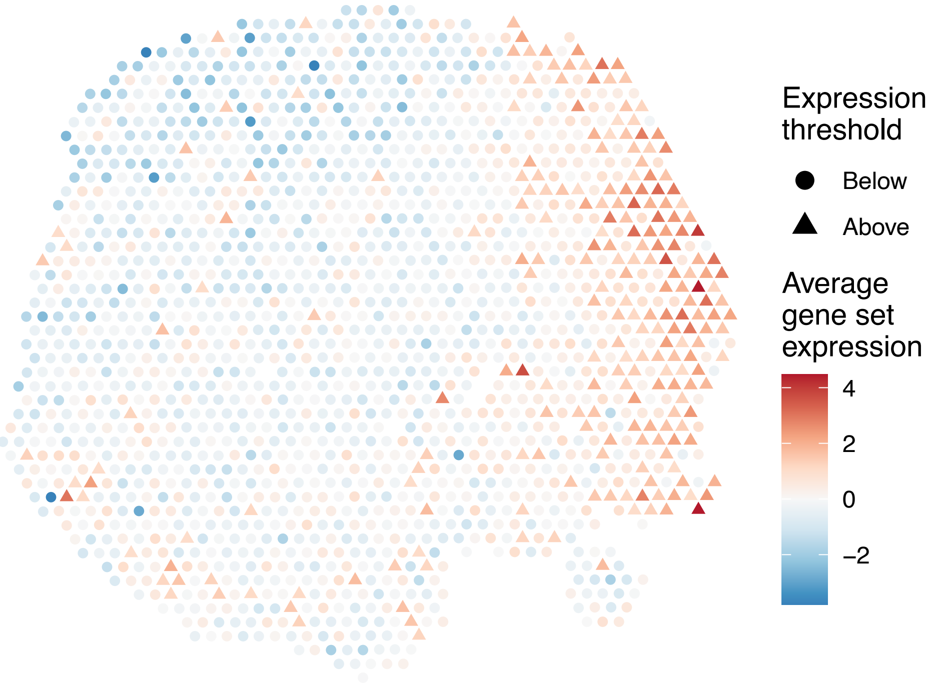

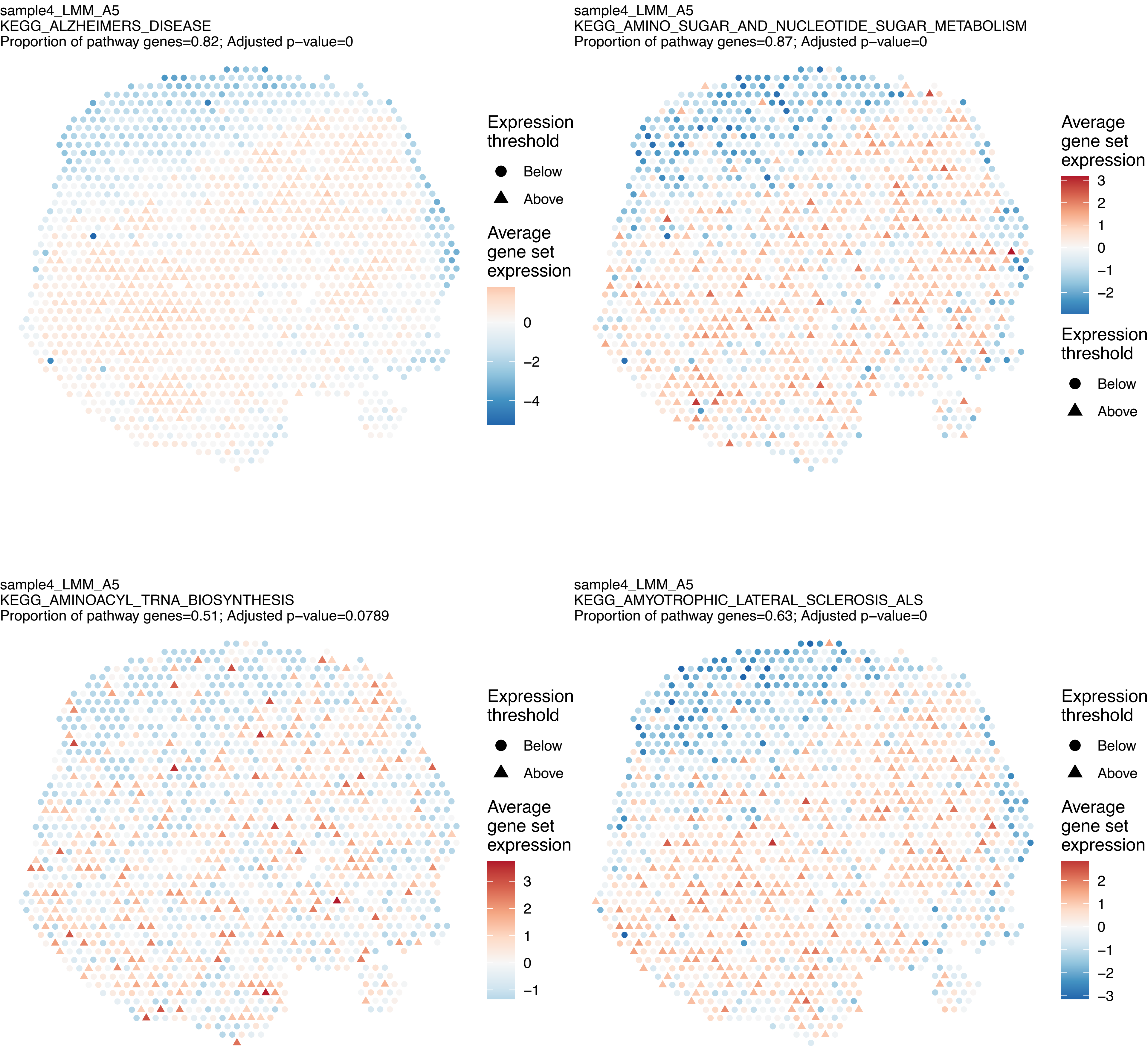
sample4_LMM_A5
KEGG_ALZHEIMERS_DISEASE
Proportion of pathway genes=0.82; Adjusted p−value=0
Expression threshold
Below
Above
Average gene set expression
0
−2
−4
sample4_LMM_A5
KEGG_AMINO_SUGAR_AND_NUCLEOTIDE_SUGAR_METABOLISM
Proportion of pathway genes=0.87; Adjusted p−value=0
Average gene set expression
3
2
1
0
−1
−2
Expression threshold
Below
Above
sample4_LMM_A5
KEGG_AMINOACYL_TRNA_BIOSYNTHESIS
Proportion of pathway genes=0.51; Adjusted p−value=0.0789
Expression threshold
Below
Above
Average gene set expression
3
2
1
0
−1
sample4_LMM_A5
KEGG_AMYOTROPHIC_LATERAL_SCLEROSIS_ALS
Proportion of pathway genes=0.63; Adjusted p−value=0
Expression threshold
Below
Above
Average gene set expression
2
1
0
−1
−2
−3

sample4_LMM_A5
KEGG_APOPTOSIS
Proportion of pathway genes=0.81; Adjusted p−value=0

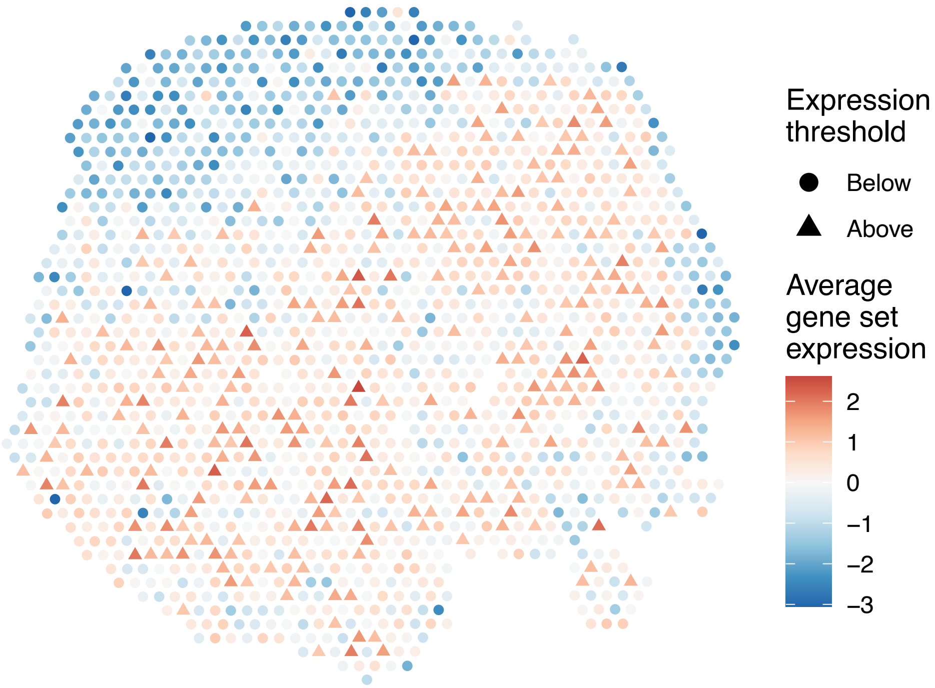


sample4_LMM_A5
KEGG_ARACHIDONIC_ACID_METABOLISM
Proportion of pathway genes=0.42; Adjusted p−value=0.0795

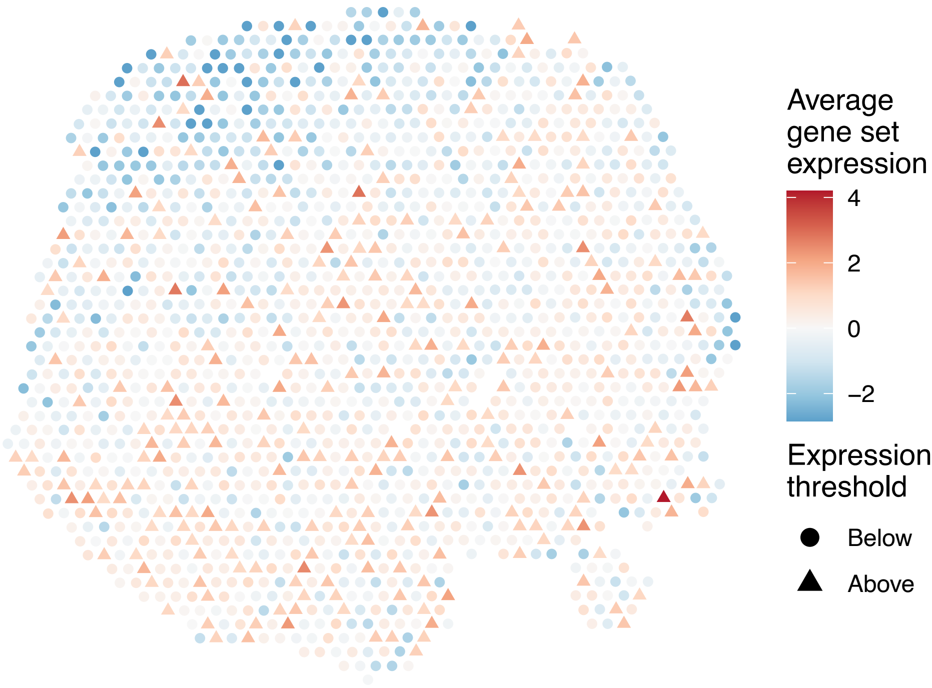


sample4_LMM_A5
KEGG_ARGININE_AND_PROLINE_METABOLISM
Proportion of pathway genes=0.73; Adjusted p−value=0

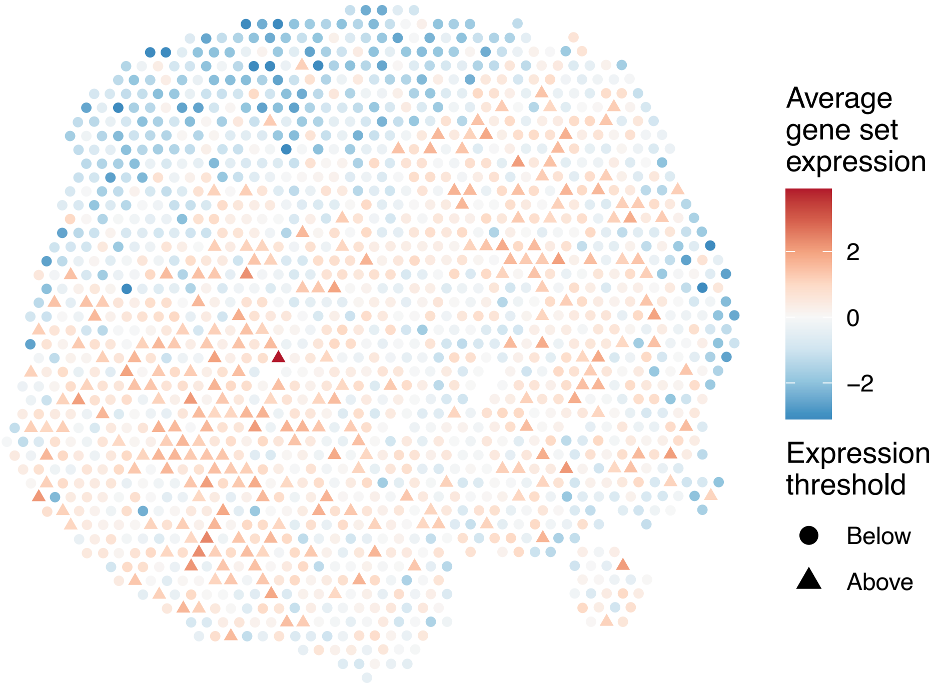


sample4_LMM_A5
KEGG_ARRHYTHMOGENIC_RIGHT_VENTRICULAR_CARDIOMYOPAT
HY_ARVC
Proportion of pathway genes=0.73; Adjusted p−value=0

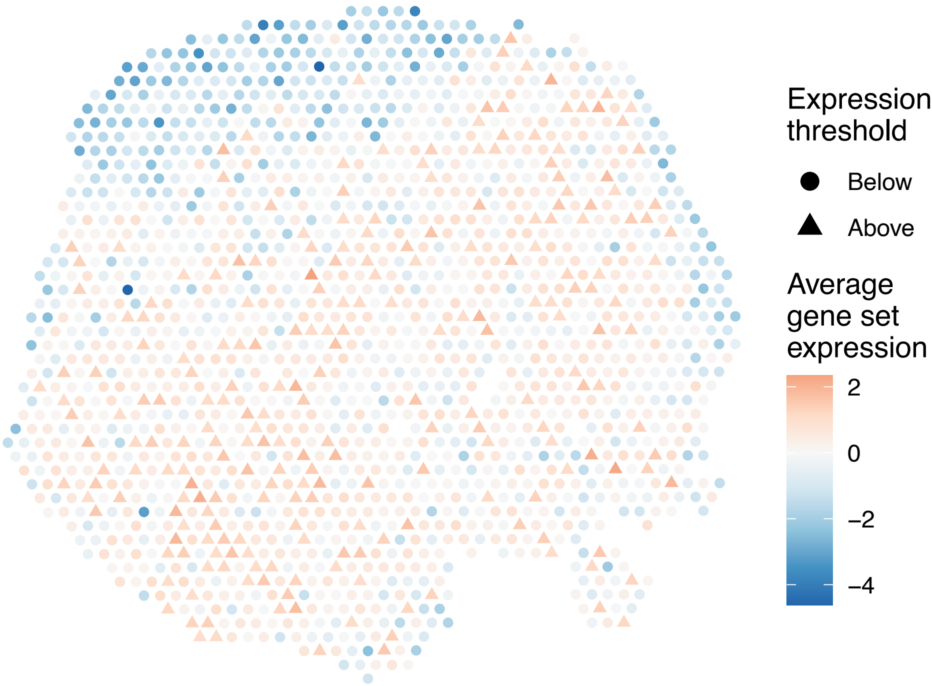

sample4_LMM_A5
KEGG_ASTHMA
Proportion of pathway genes=0.72; Adjusted p−value=1

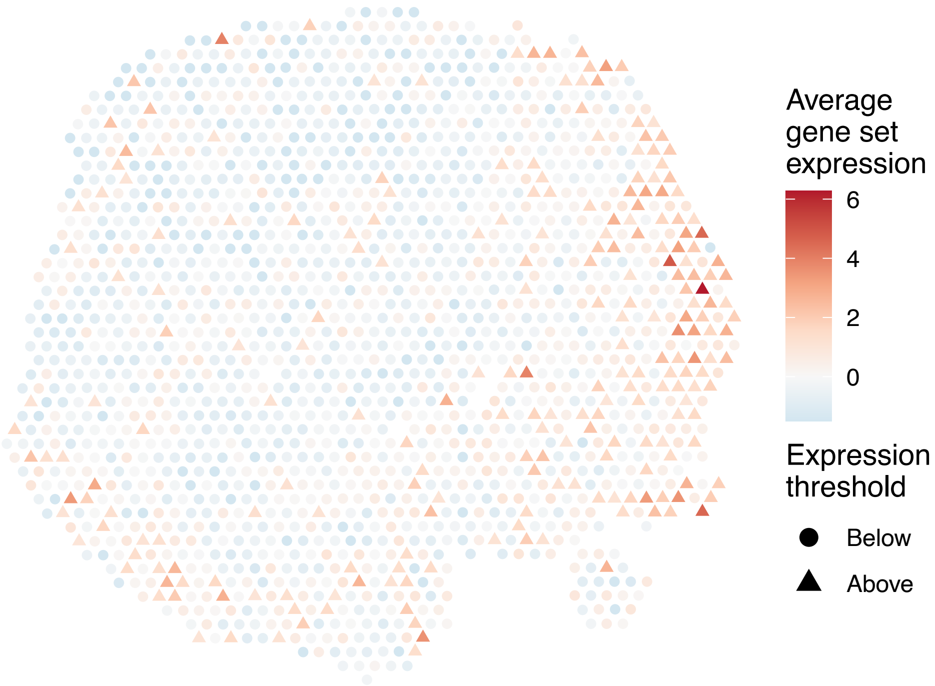


sample4_LMM_A5
KEGG_AUTOIMMUNE_THYROID_DISEASE
Proportion of pathway genes=0.72; Adjusted p−value=0

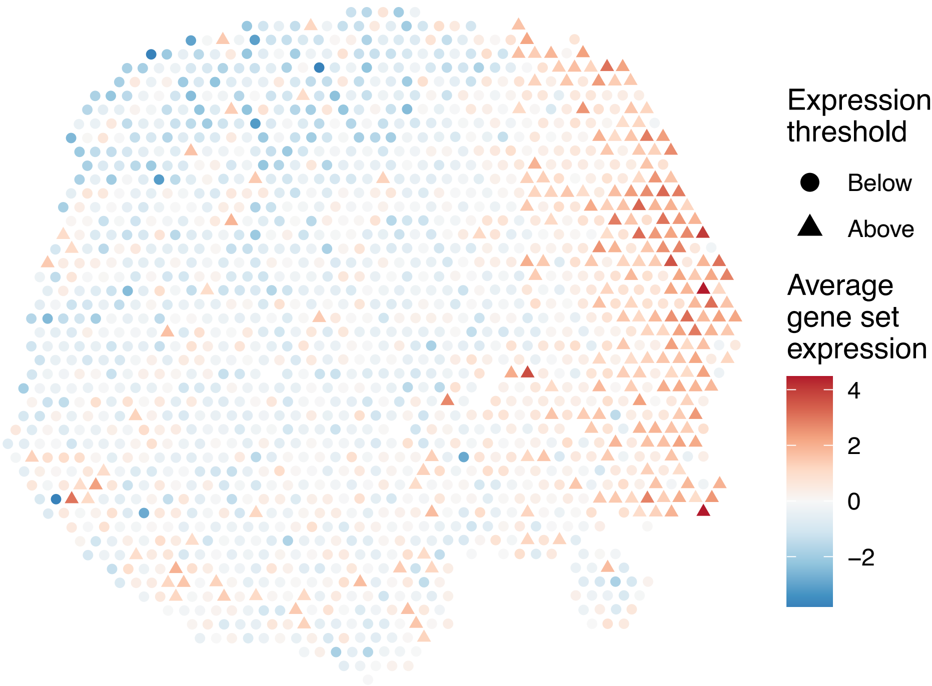


sample4_LMM_A5
KEGG_AXON_GUIDANCE
Proportion of pathway genes=0.82; Adjusted p−value=0

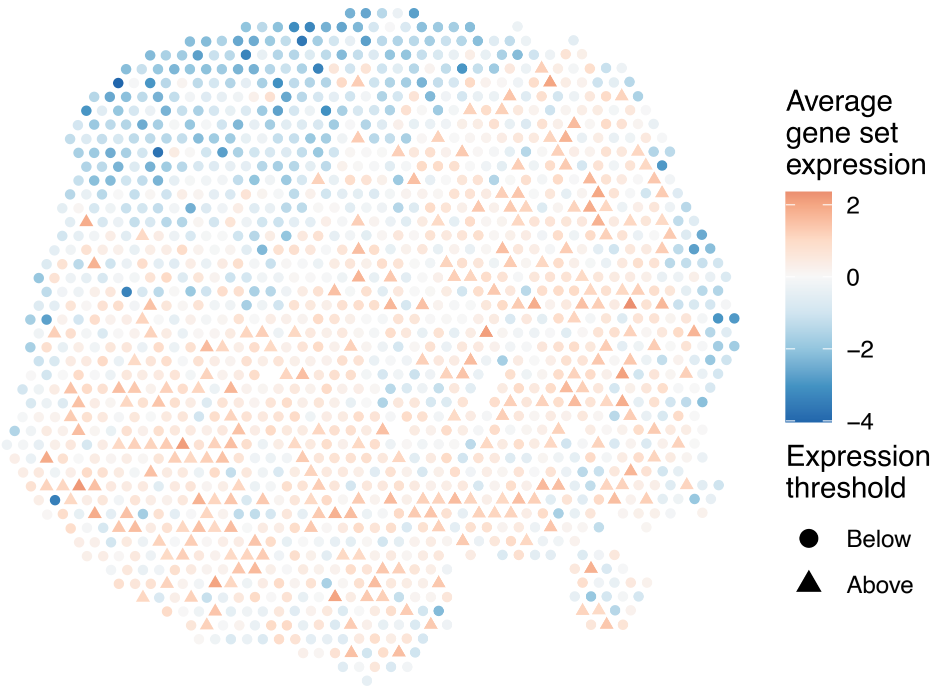


sample4_LMM_A5
KEGG_B_CELL_RECEPTOR_SIGNALING_PATHWAY
Proportion of pathway genes=0.9; Adjusted p−value=0

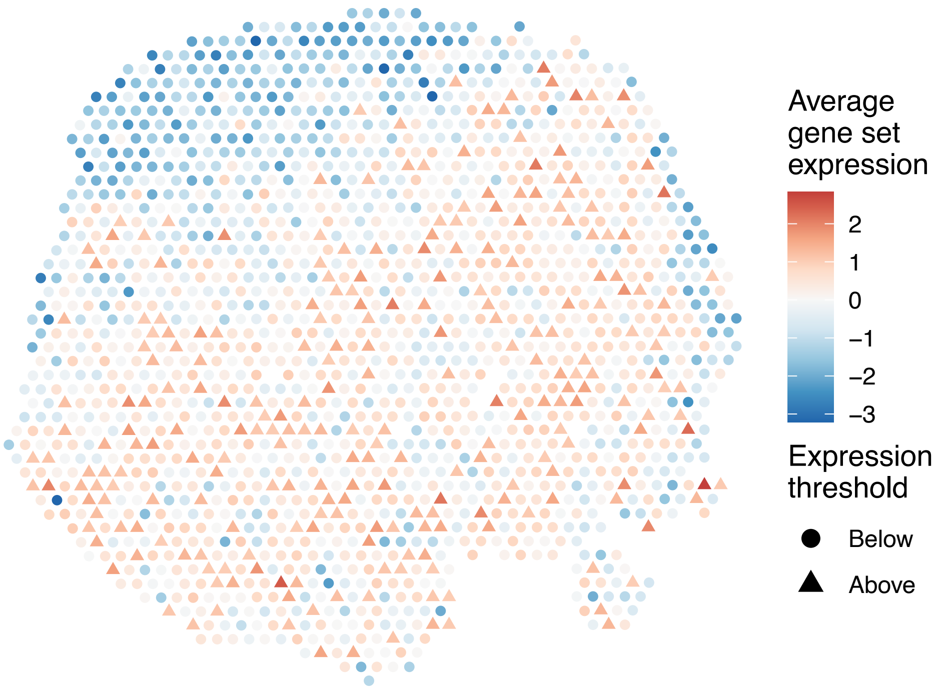

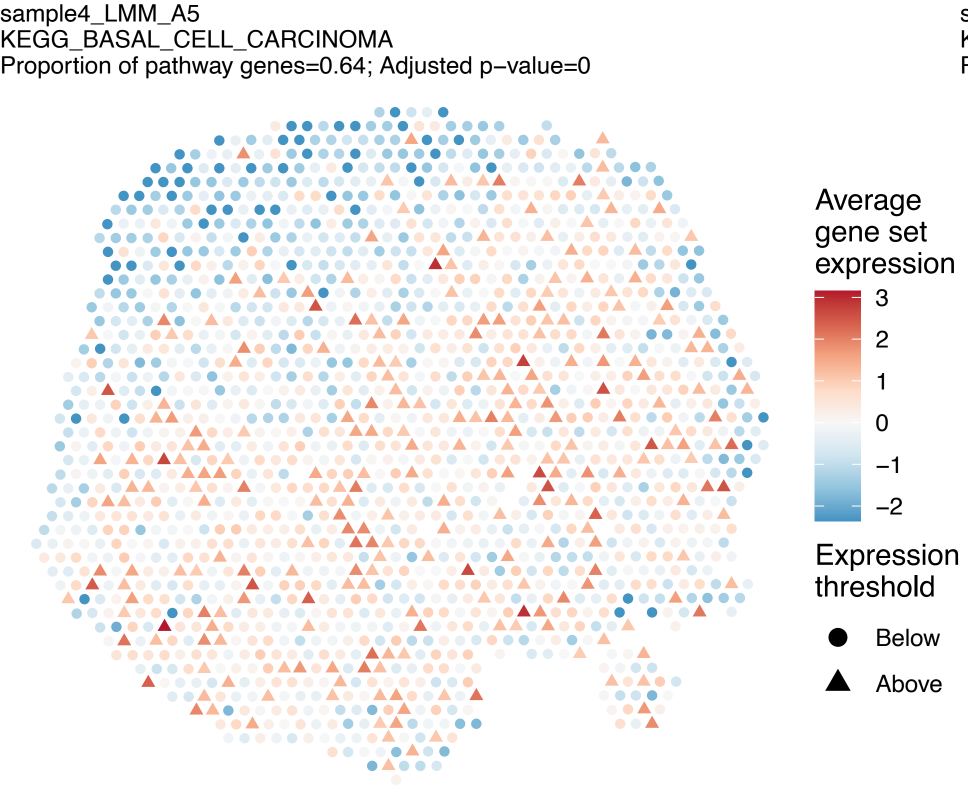
sample4_LMM_A5
KEGG_BASAL_CELL_CARCINOMA
Proportion of pathway genes=0.64; Adjusted p−value=0
Average gene set expression
3
2
1
0
−1
−2
Expression threshold
Below
Above

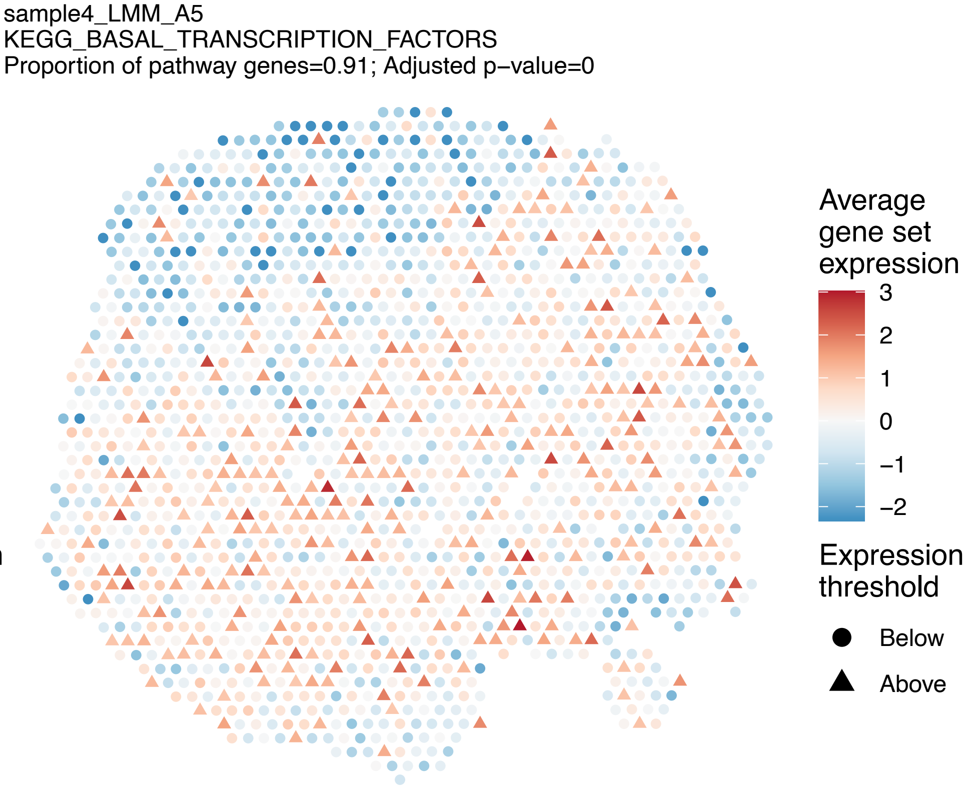
sample4_LMM_A5
KEGG_BASAL_TRANSCRIPTION_FACTORS
Proportion of pathway genes=0.91; Adjusted p−value=0
Average gene set expression
3
2
1
0
−1
−2
Expression threshold
Below
Above

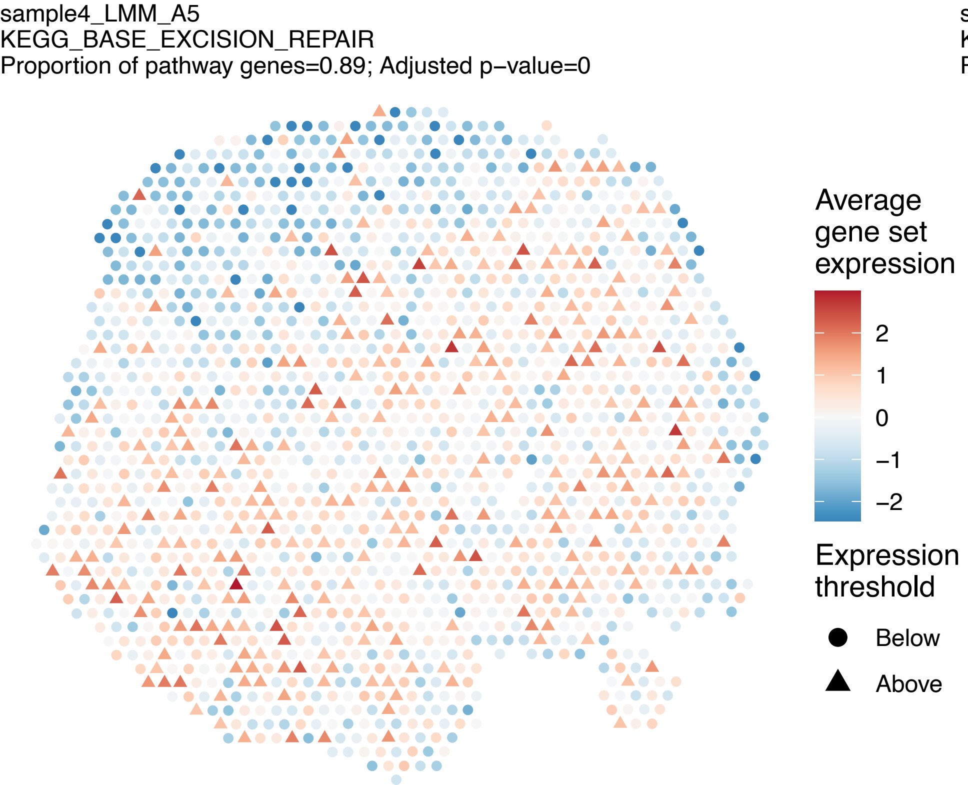
sample4_LMM_A5
KEGG_BASE_EXCISION_REPAIR
Proportion of pathway genes=0.89; Adjusted p−value=0
Average gene set expression
2
1
0
−1
−2
Expression threshold
Below
Above

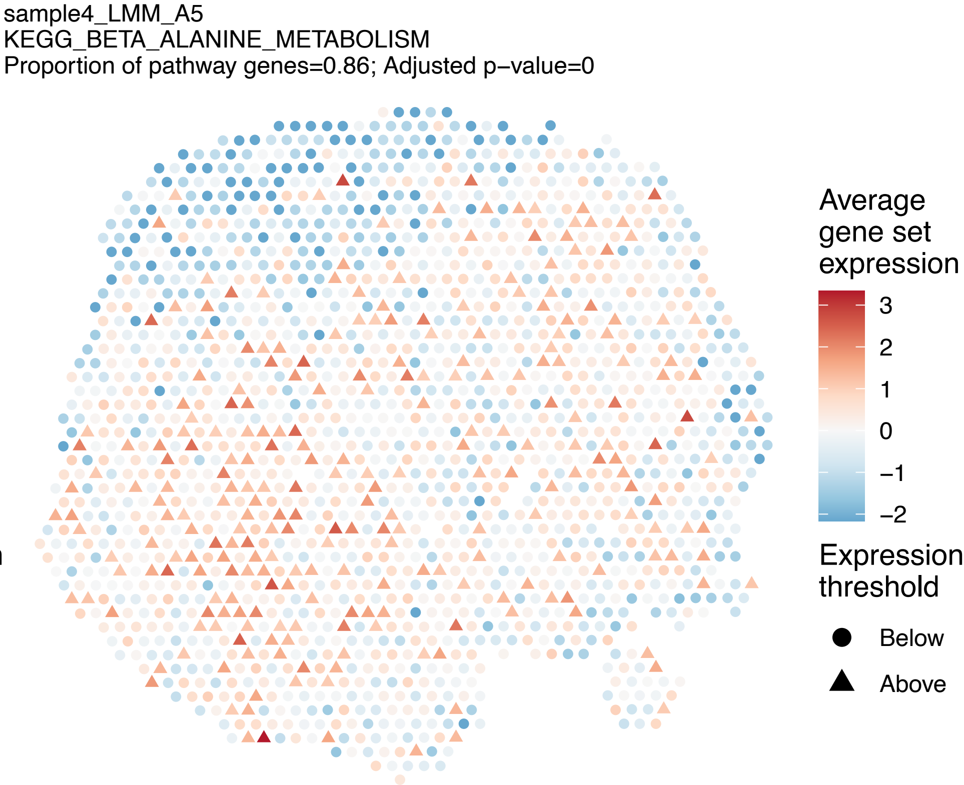
sample4_LMM_A5
KEGG_BETA_ALANINE_METABOLISM
Proportion of pathway genes=0.86; Adjusted p−value=0
Average gene set expression
3
2
1
0
−1
−2
Expression threshold
Below
Above

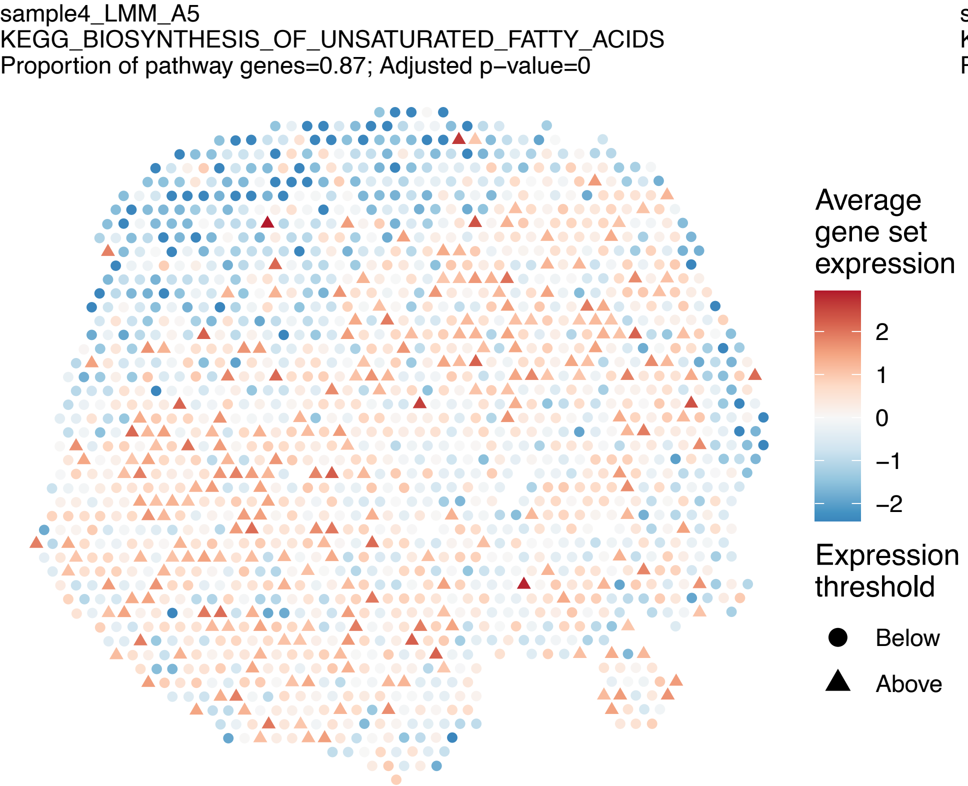
sample4_LMM_A5
KEGG_BIOSYNTHESIS_OF_UNSATURATED_FATTY_ACIDS
Proportion of pathway genes=0.87; Adjusted p−value=0
Average gene set expression
2
1
0
−1
−2
Expression threshold
Below
Above

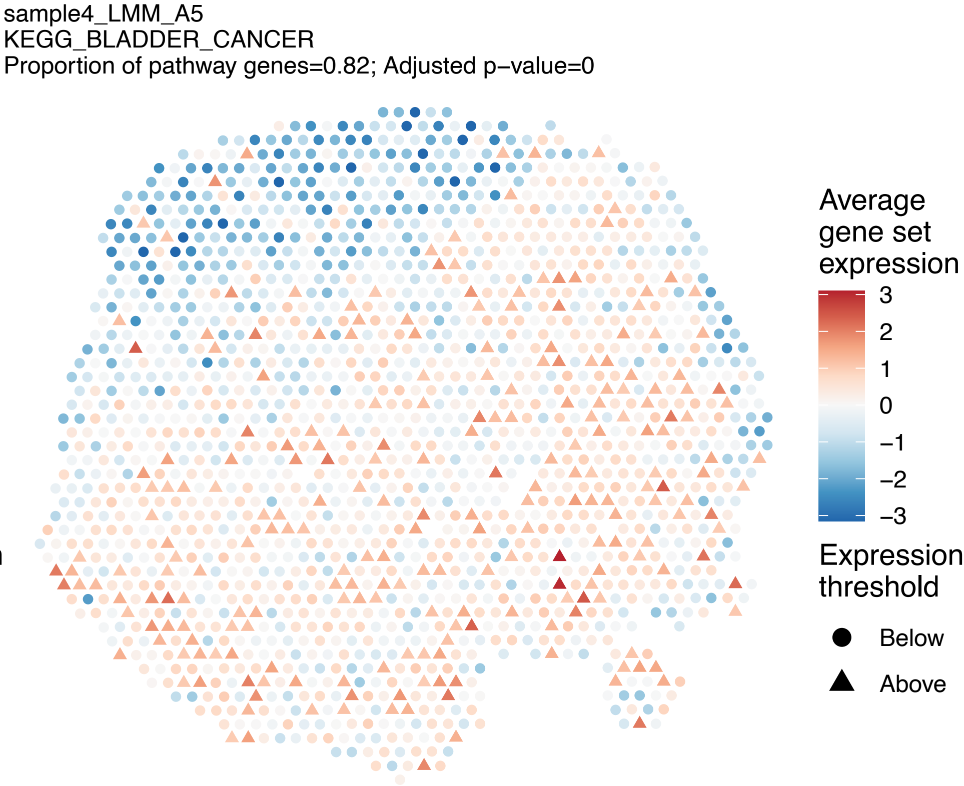
sample4_LMM_A5
KEGG_BLADDER_CANCER
Proportion of pathway genes=0.82; Adjusted p−value=0
Average gene set expression
3
2
1
0
−1
−2
−3
Expression threshold
Below
Above

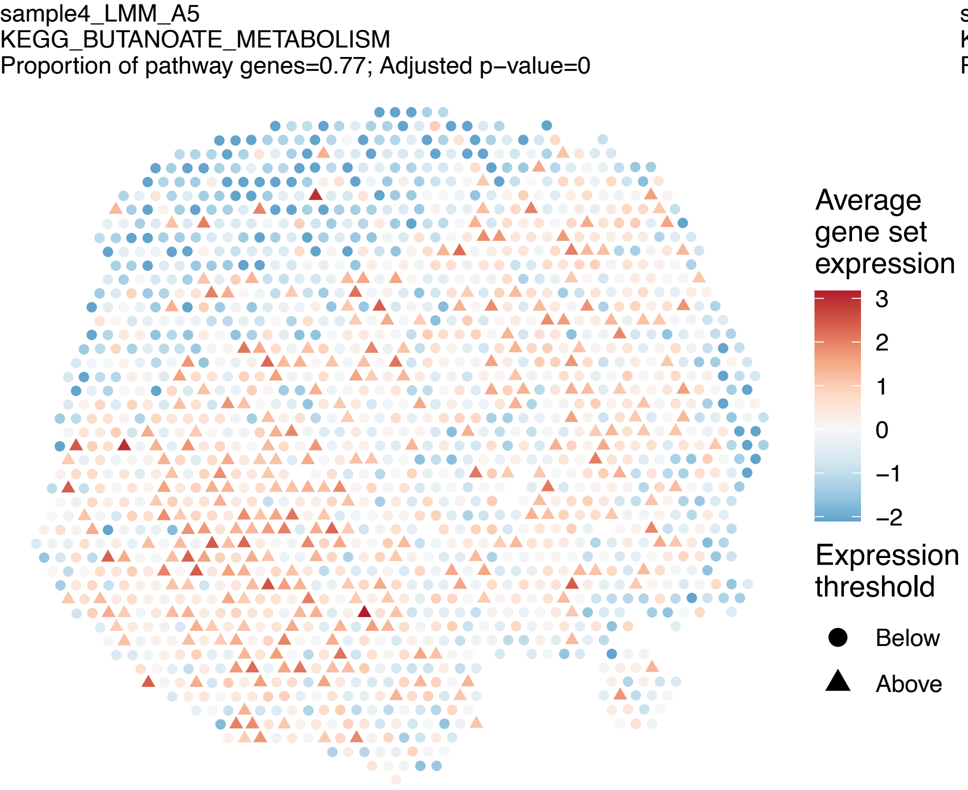
sample4_LMM_A5
KEGG_BUTANOATE_METABOLISM
Proportion of pathway genes=0.77; Adjusted p−value=0
Average gene set expression
3
2
1
0
−1
−2
Expression threshold
Below
Above

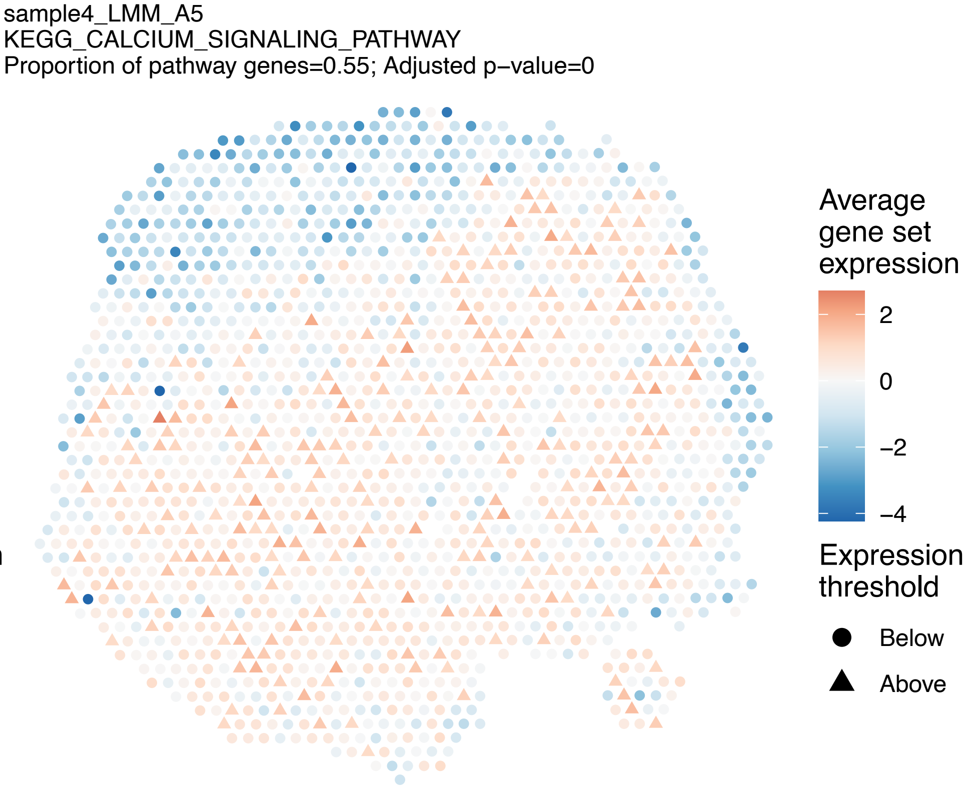
sample4_LMM_A5
KEGG_CALCIUM_SIGNALING_PATHWAY
Proportion of pathway genes=0.55; Adjusted p−value=0
Average gene set expression
2
0
−2
−4
Expression threshold
Below
Above

sample4_LMM_A5
KEGG_CARDIAC_MUSCLE_CONTRACTION
Proportion of pathway genes=0.63; Adjusted p−value=0

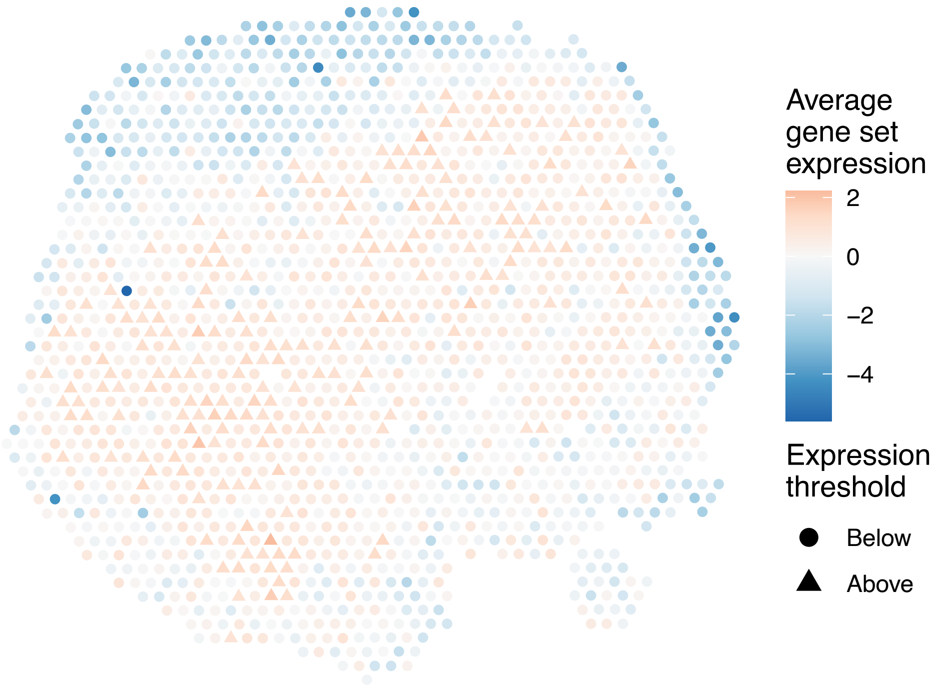


sample4_LMM_A5
KEGG_CELL_ADHESION_MOLECULES_CAMS
Proportion of pathway genes=0.78; Adjusted p−value=0

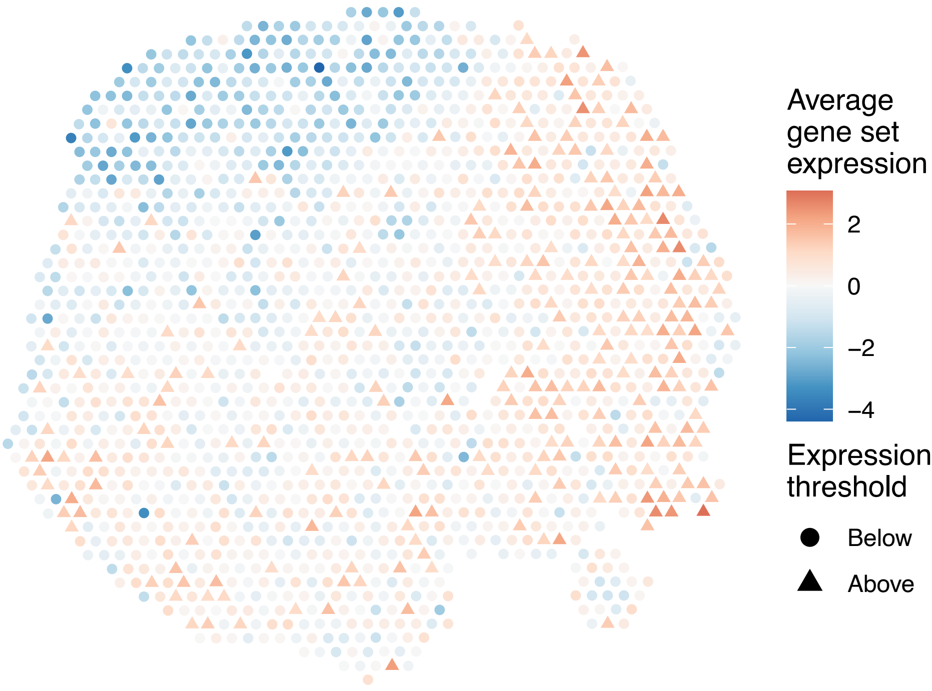


sample4_LMM_A5
KEGG_CELL_CYCLE
Proportion of pathway genes=0.78; Adjusted p−value=0

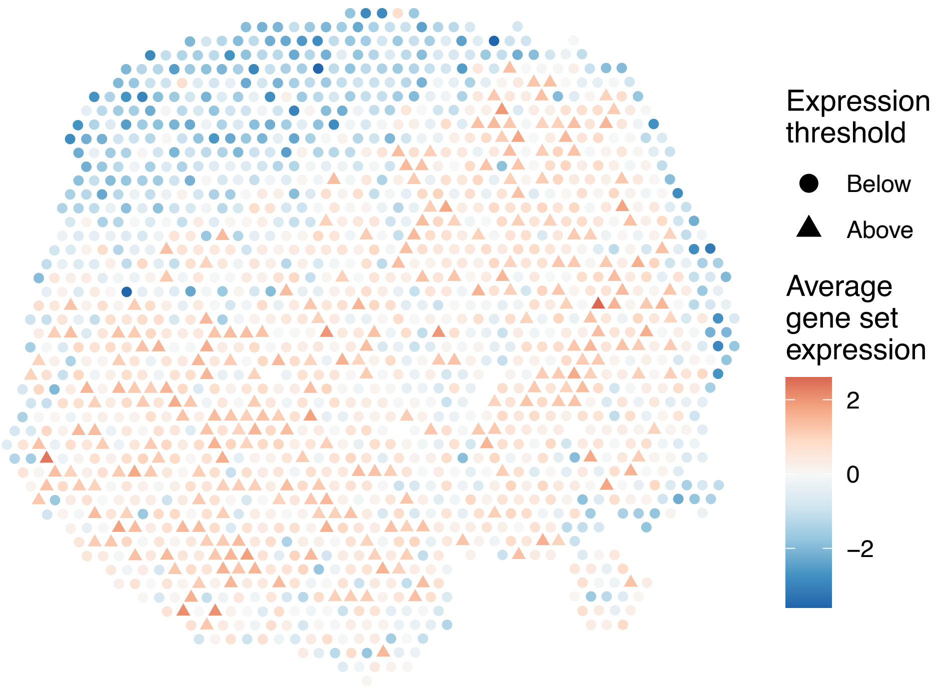


sample4_LMM_A5
KEGG_CHEMOKINE_SIGNALING_PATHWAY
Proportion of pathway genes=0.69; Adjusted p−value=0.0027

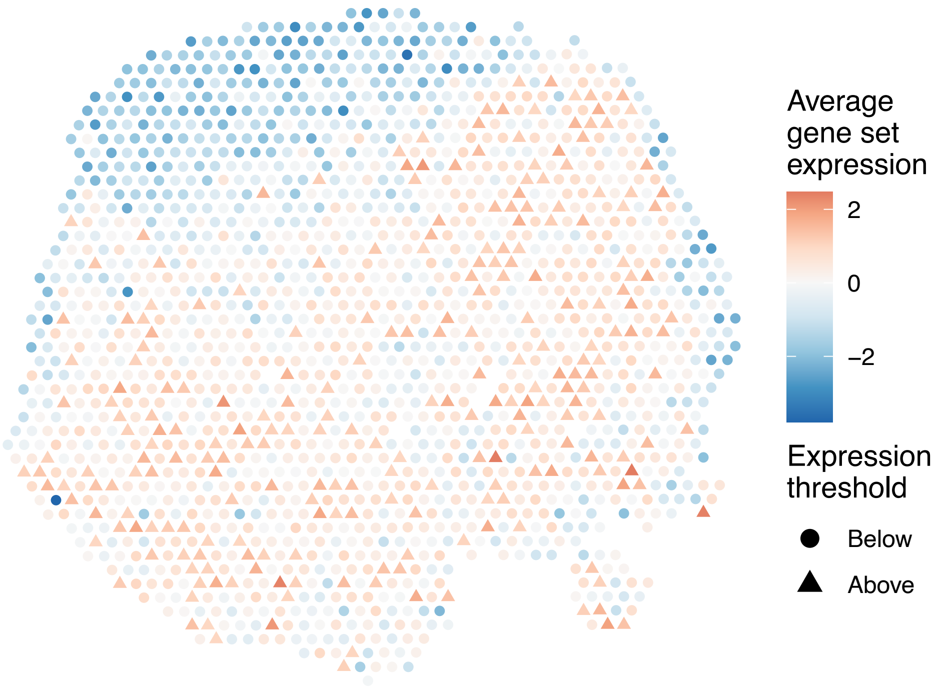

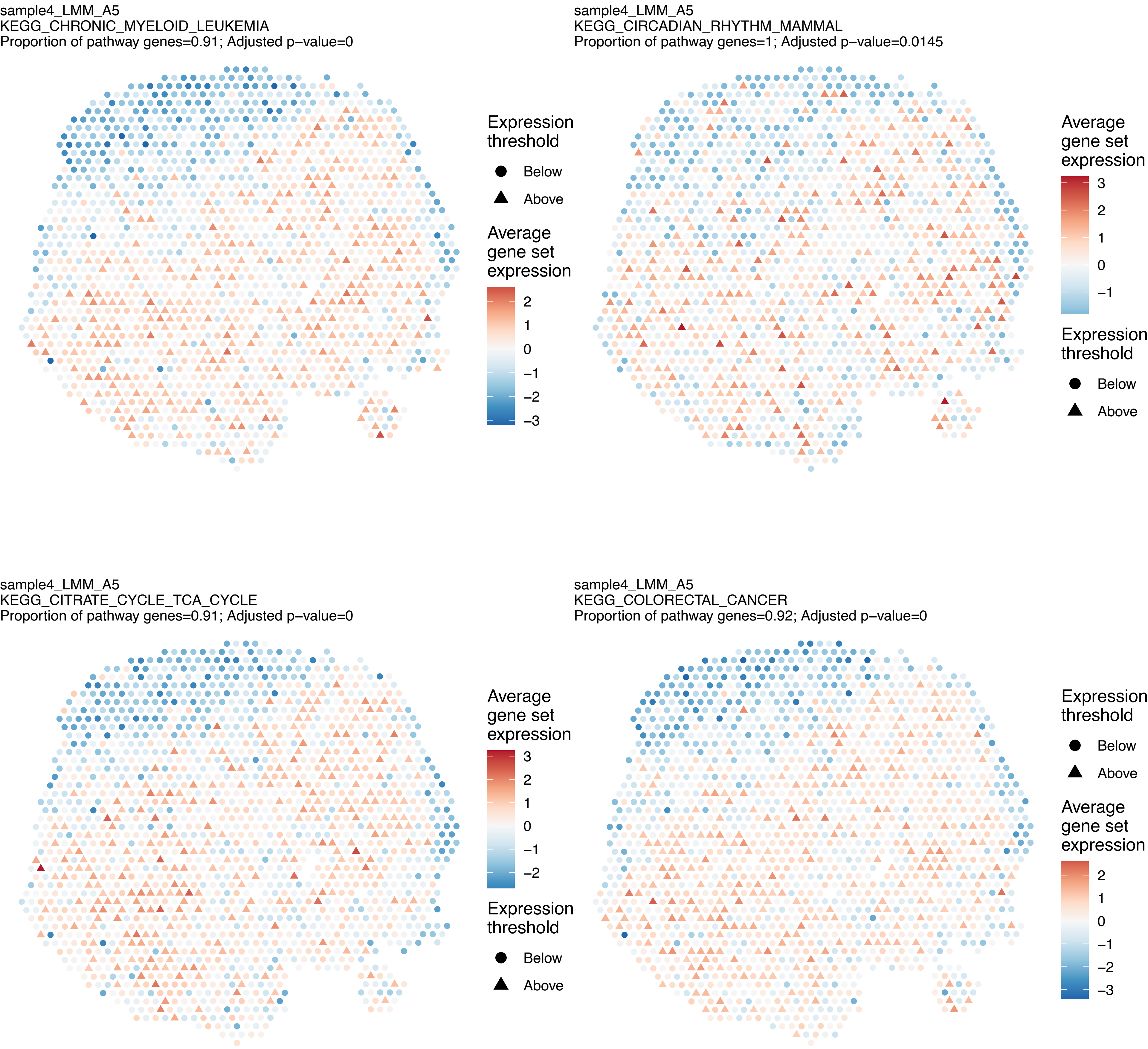

sample4_LMM_A5
KEGG_CHRONIC_MYELOID_LEUKEMIA
Proportion of pathway genes=0.91; Adjusted p−value=0
Expression threshold
Below
Above
Average gene set expression
2
1
0
−1
−2
−3
sample4_LMM_A5
KEGG_CIRCADIAN_RHYTHM_MAMMAL
Proportion of pathway genes=1; Adjusted p−value=0.0145
Average gene set expression
3
2
1
0
−1
Expression threshold
Below
Above
sample4_LMM_A5
KEGG_CITRATE_CYCLE_TCA_CYCLE
Proportion of pathway genes=0.91; Adjusted p−value=0
Average gene set expression
3
2
1
0
−1
−2
Expression threshold
Below
Above
sample4_LMM_A5
KEGG_COLORECTAL_CANCER
Proportion of pathway genes=0.92; Adjusted p−value=0
Expression threshold
Below
Above
Average gene set expression
2
1
0
−1
−2
−3

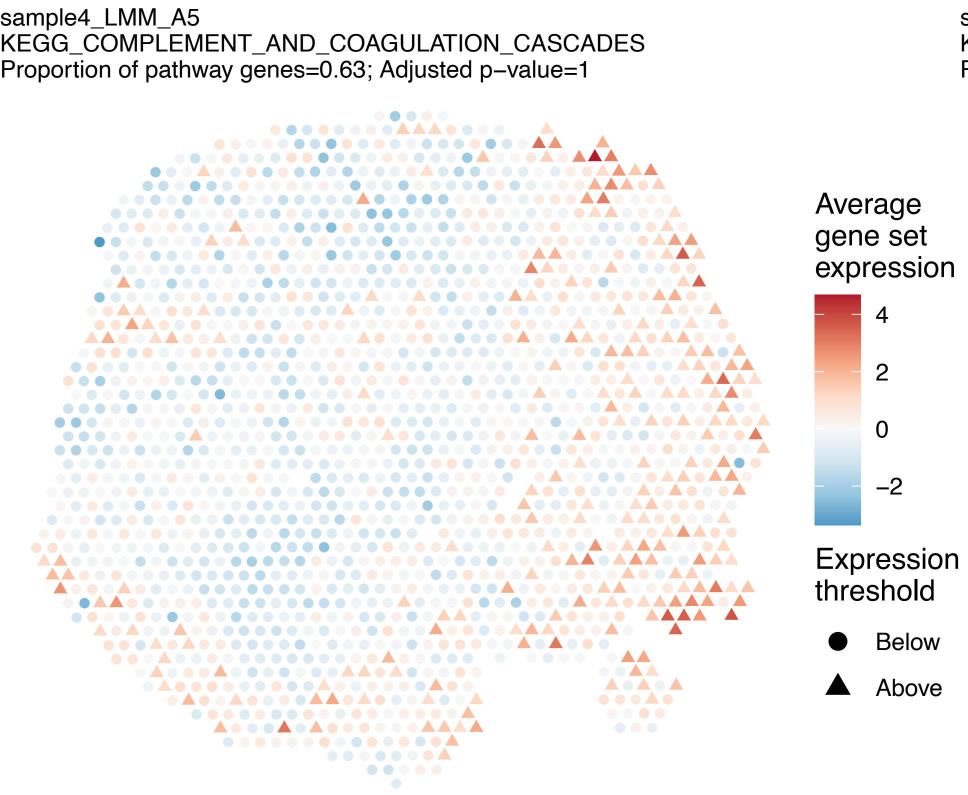

sample4_LMM_A5
KEGG_COMPLEMENT_AND_COAGULATION_CASCADES
Proportion of pathway genes=0.63; Adjusted p−value=1
Average gene set expression
4
2
0
−2
Expression threshold
Below
Above


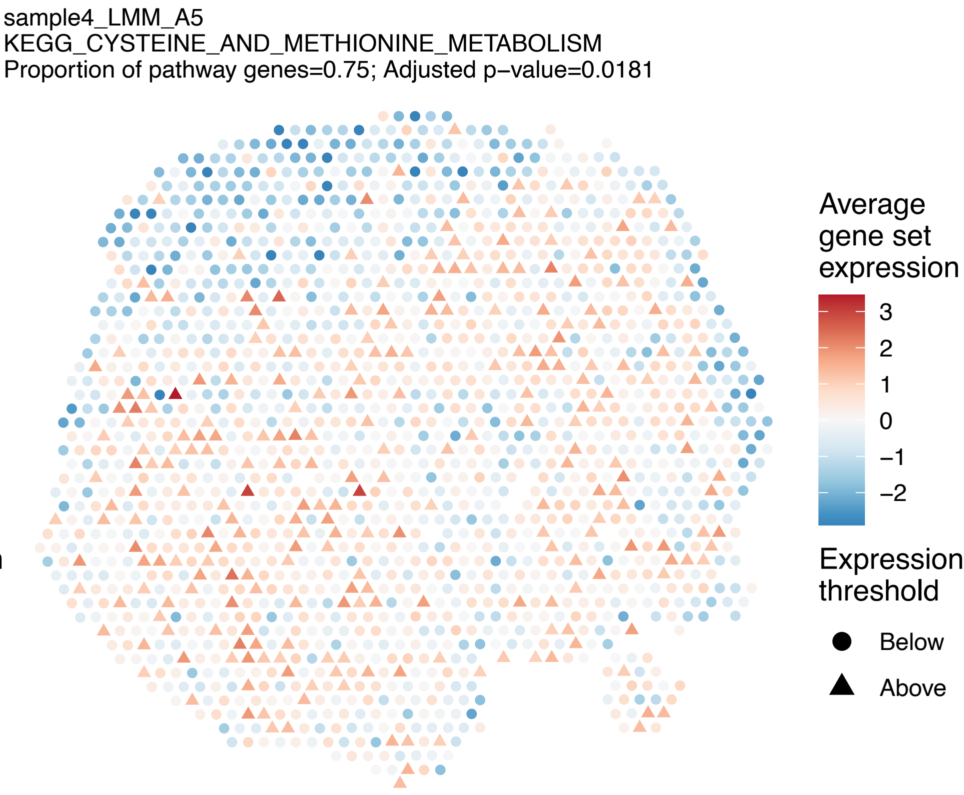

sample4_LMM_A5
KEGG_CYSTEINE_AND_METHIONINE_METABOLISM
Proportion of pathway genes=0.75; Adjusted p−value=0.0181
Average gene set expression
3
2
1
0
−1
−2
Expression threshold
Below
Above


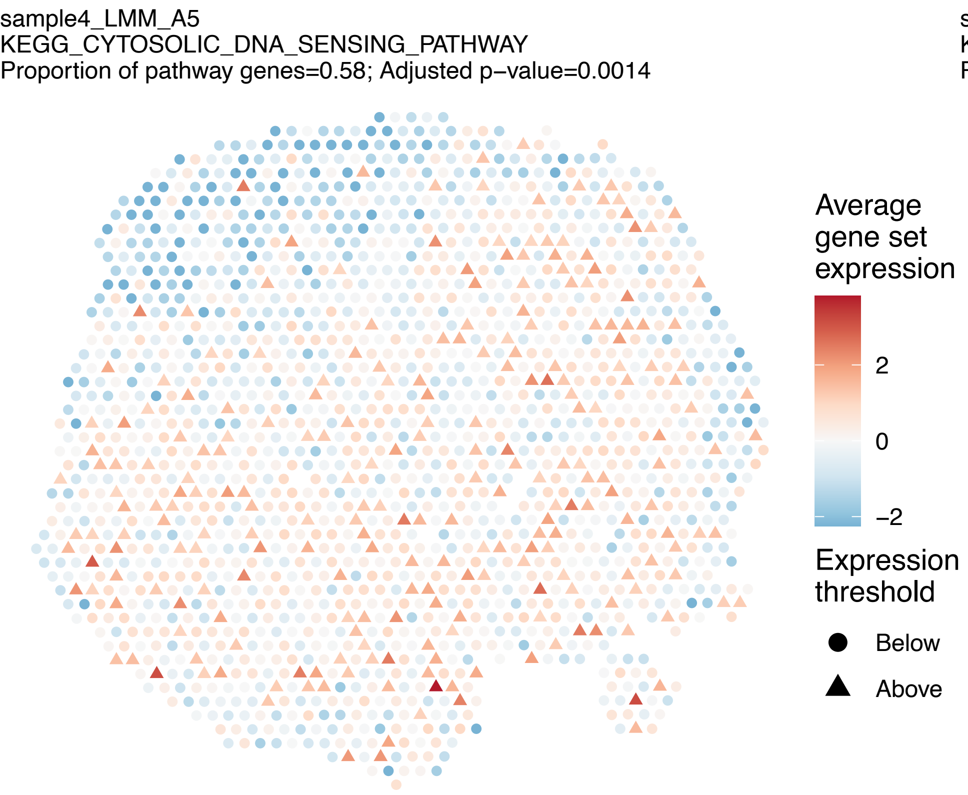

sample4_LMM_A5
KEGG_CYTOSOLIC_DNA_SENSING_PATHWAY
Proportion of pathway genes=0.58; Adjusted p−value=0.0014
Average gene set expression
2
0
−2
Expression threshold
Below
Above


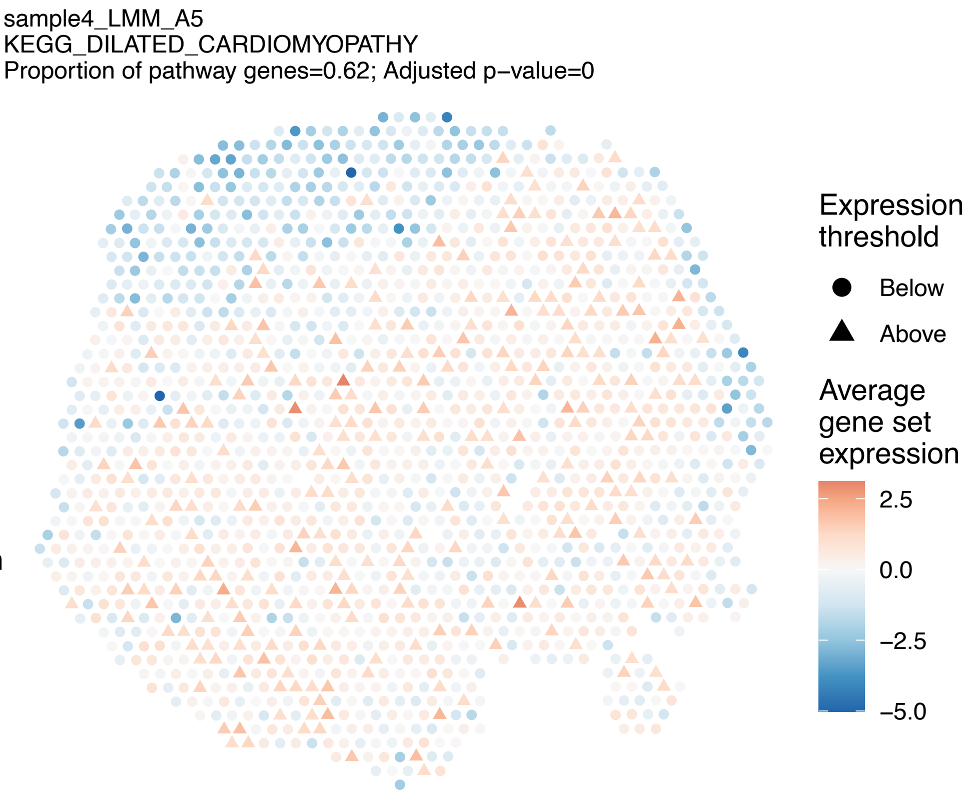

sample4_LMM_A5
KEGG_DILATED_CARDIOMYOPATHY
Proportion of pathway genes=0.62; Adjusted p−value=0
Expression threshold
Below
Above
Average gene set expression
2.5
0.0
−2.5
−5.0

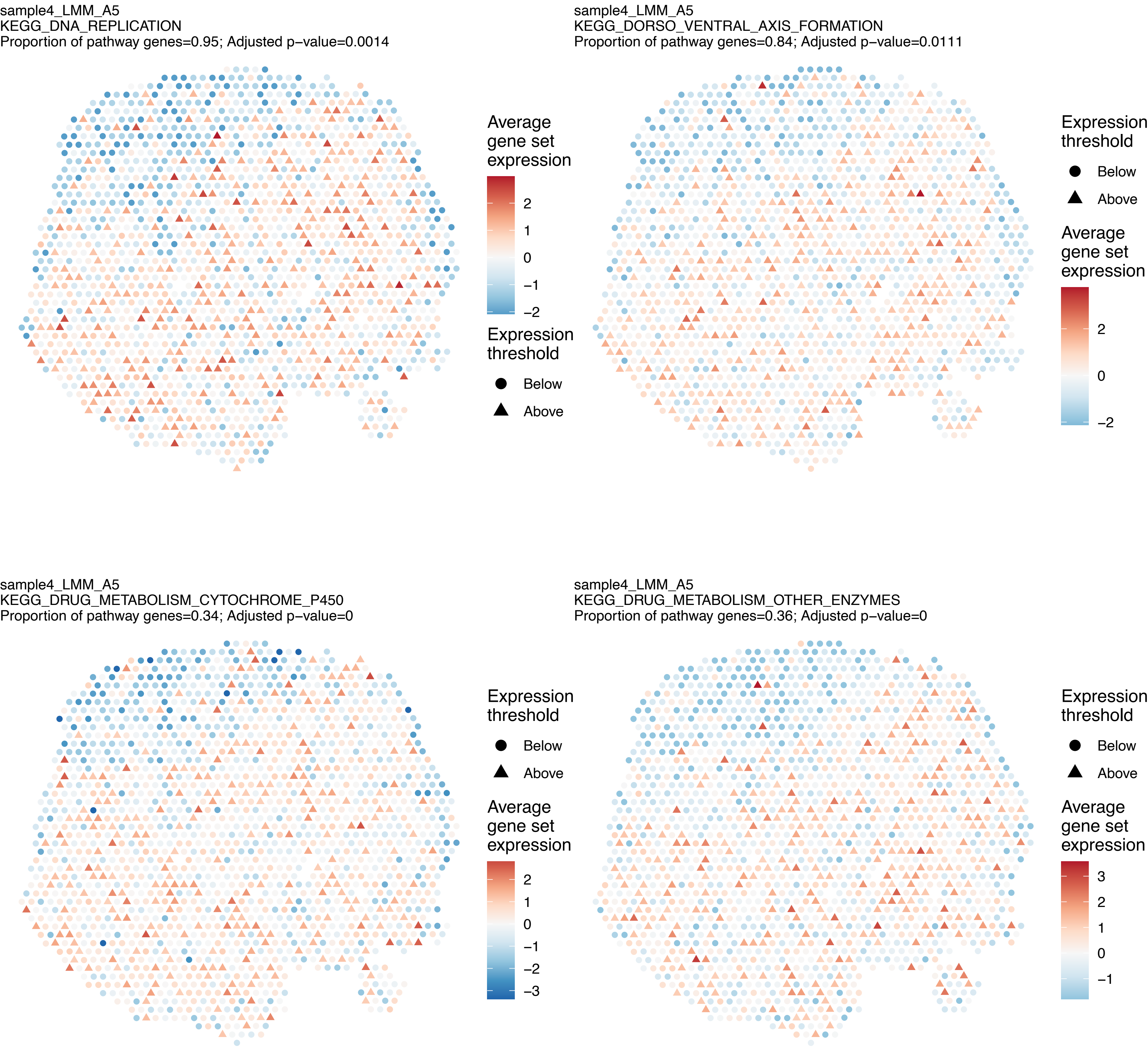

sample4_LMM_A5
KEGG_DNA_REPLICATION
Proportion of pathway genes=0.95; Adjusted p−value=0.0014
Average gene set expression
2
1
0
−1
−2
Expression threshold
Below
Above
sample4_LMM_A5
KEGG_DORSO_VENTRAL_AXIS_FORMATION
Proportion of pathway genes=0.84; Adjusted p−value=0.0111
Expression threshold
Below
Above
Average gene set expression
2
0
−2
sample4_LMM_A5
KEGG_DRUG_METABOLISM_CYTOCHROME_P450
Proportion of pathway genes=0.34; Adjusted p−value=0
Expression threshold
Below
Above
Average gene set expression
2
1
0
−1
−2
−3
sample4_LMM_A5
KEGG_DRUG_METABOLISM_OTHER_ENZYMES
Proportion of pathway genes=0.36; Adjusted p−value=0
Expression threshold
Below
Above
Average gene set expression
3
2
1
0
−1

sample4_LMM_A5
KEGG_ECM_RECEPTOR_INTERACTION
Proportion of pathway genes=0.72; Adjusted p−value=0.9653

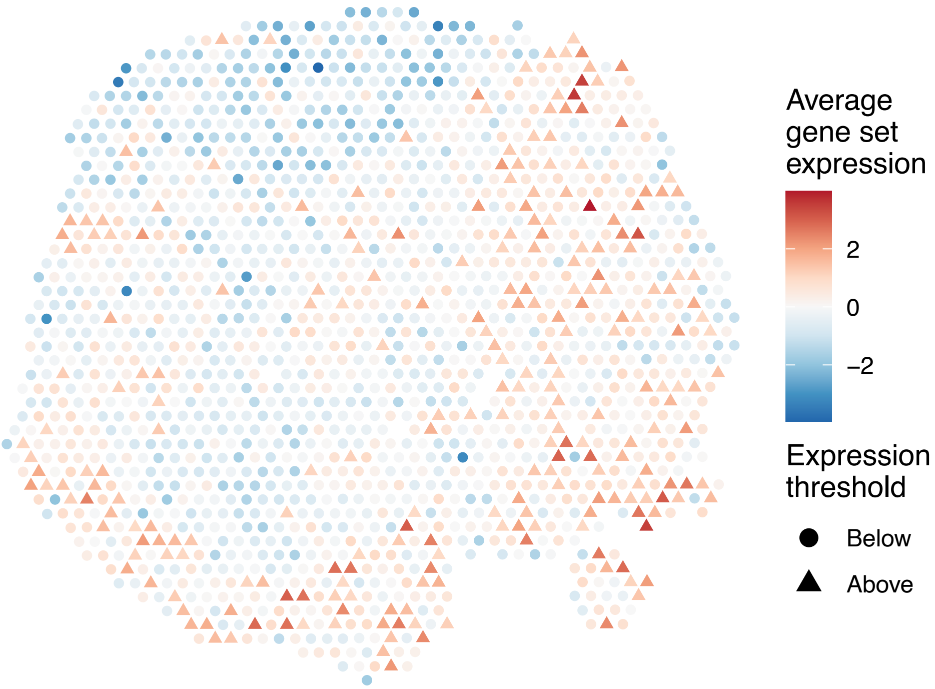


sample4_LMM_A5
KEGG_ENDOCYTOSIS
Proportion of pathway genes=0.87; Adjusted p−value=0

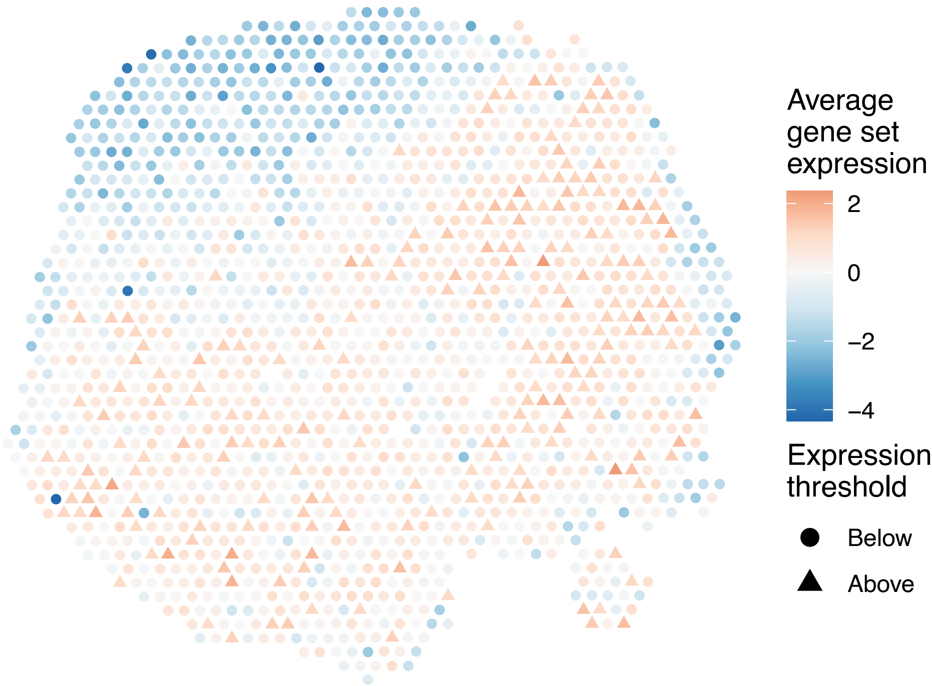


sample4_LMM_A5
KEGG_ENDOMETRIAL_CANCER
Proportion of pathway genes=0.93; Adjusted p−value=0

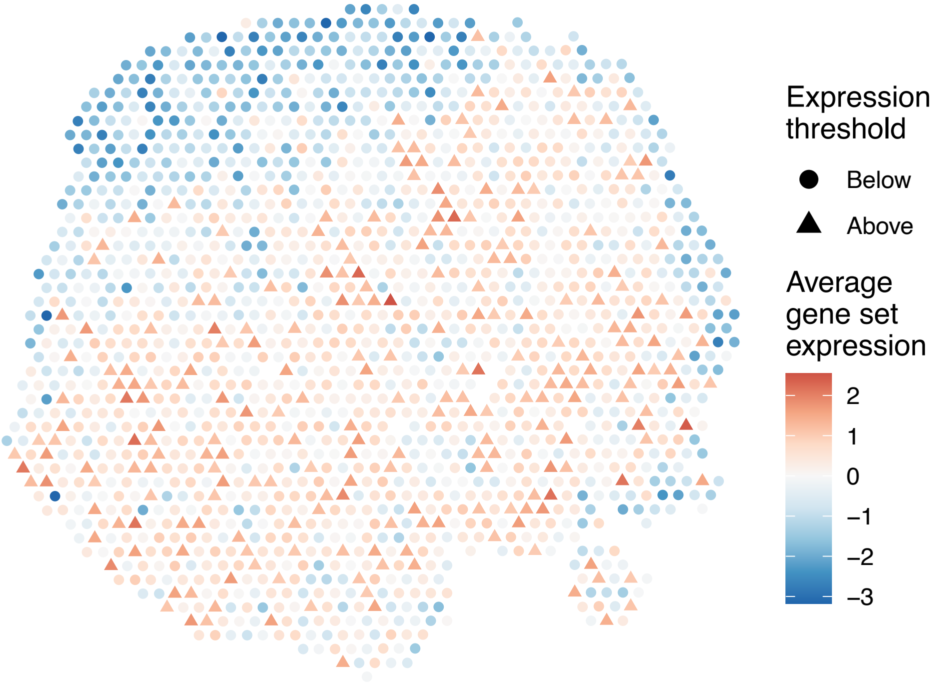


sample4_LMM_A5
KEGG_EPITHELIAL_CELL_SIGNALING_IN_HELICOBACTER_PYLORI_INFECTION
Proportion of pathway genes=0.77; Adjusted p−value=0

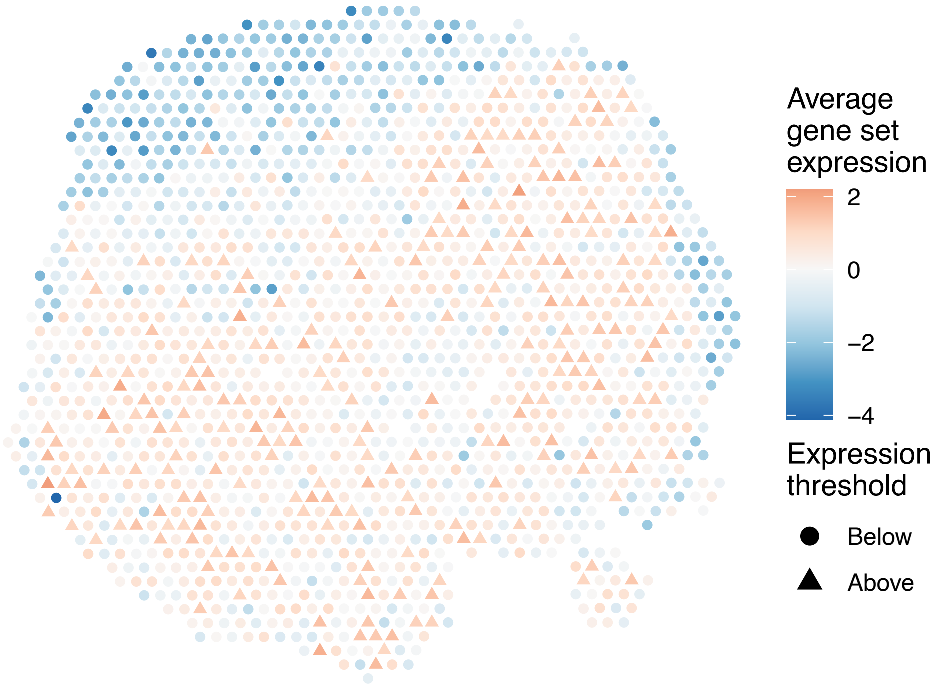

sample4_LMM_A5
KEGG_ERBB_SIGNALING_PATHWAY
Proportion of pathway genes=0.84; Adjusted p−value=0

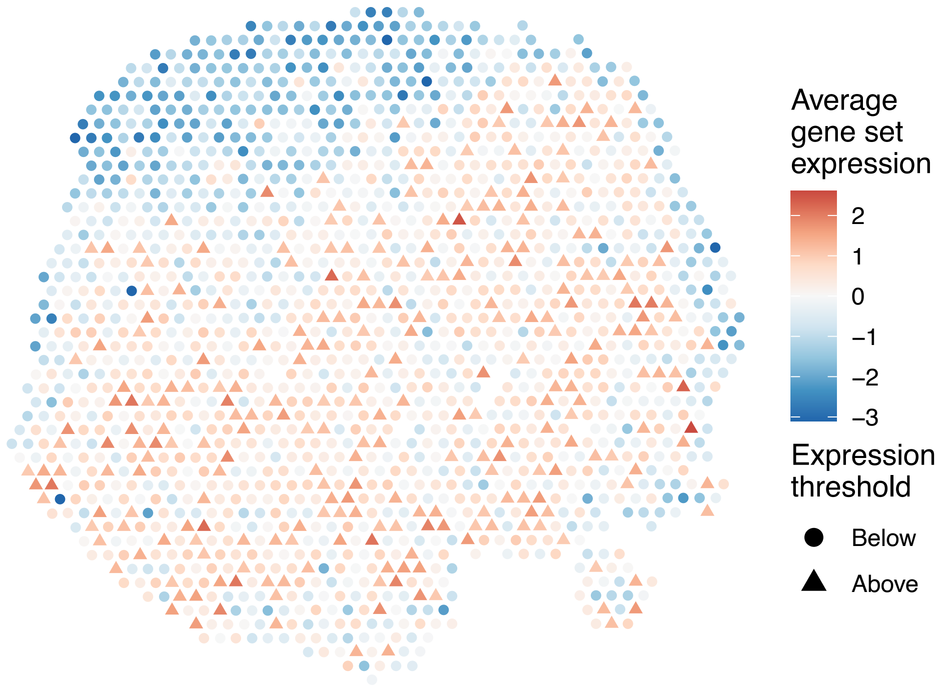


sample4_LMM_A5
KEGG_ETHER_LIPID_METABOLISM
Proportion of pathway genes=0.6; Adjusted p−value=0.2559

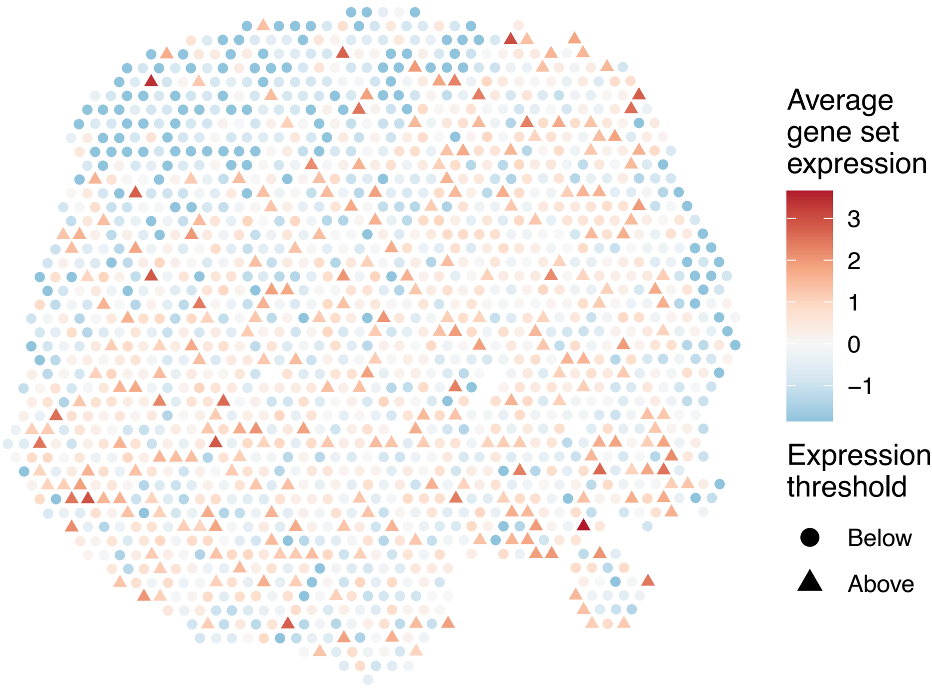


sample4_LMM_A5
KEGG_FATTY_ACID_METABOLISM
Proportion of pathway genes=0.81; Adjusted p−value=0

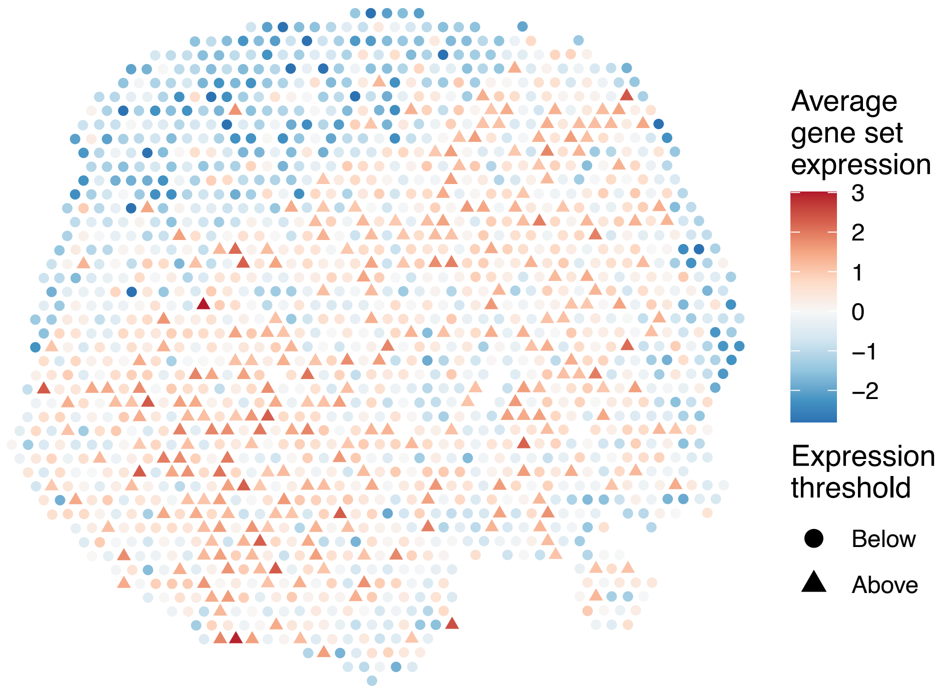


sample4_LMM_A5
KEGG_FC_EPSILON_RI_SIGNALING_PATHWAY
Proportion of pathway genes=0.68; Adjusted p−value=0.0052

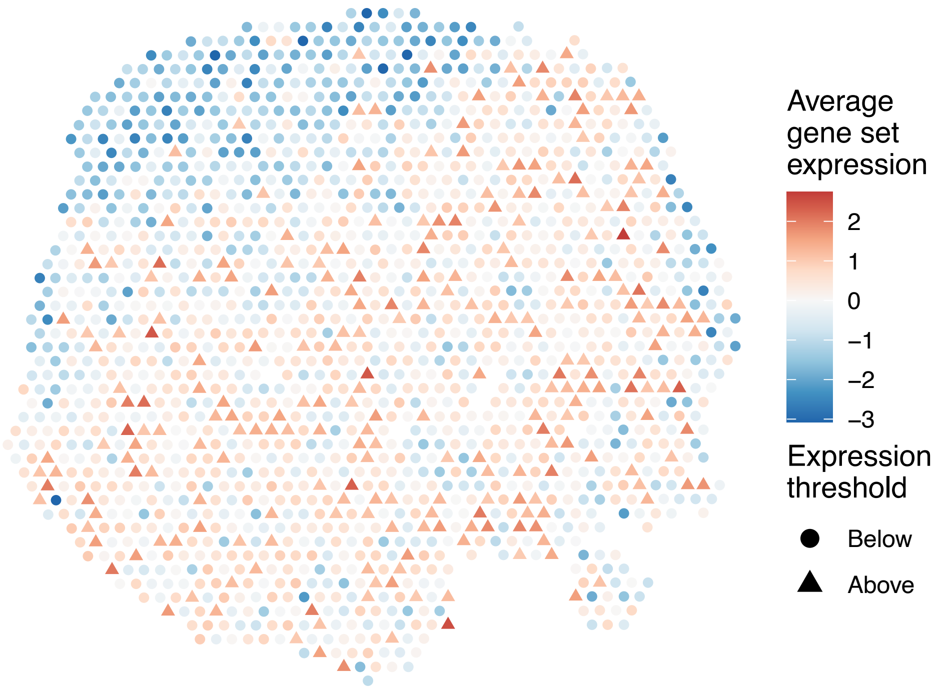

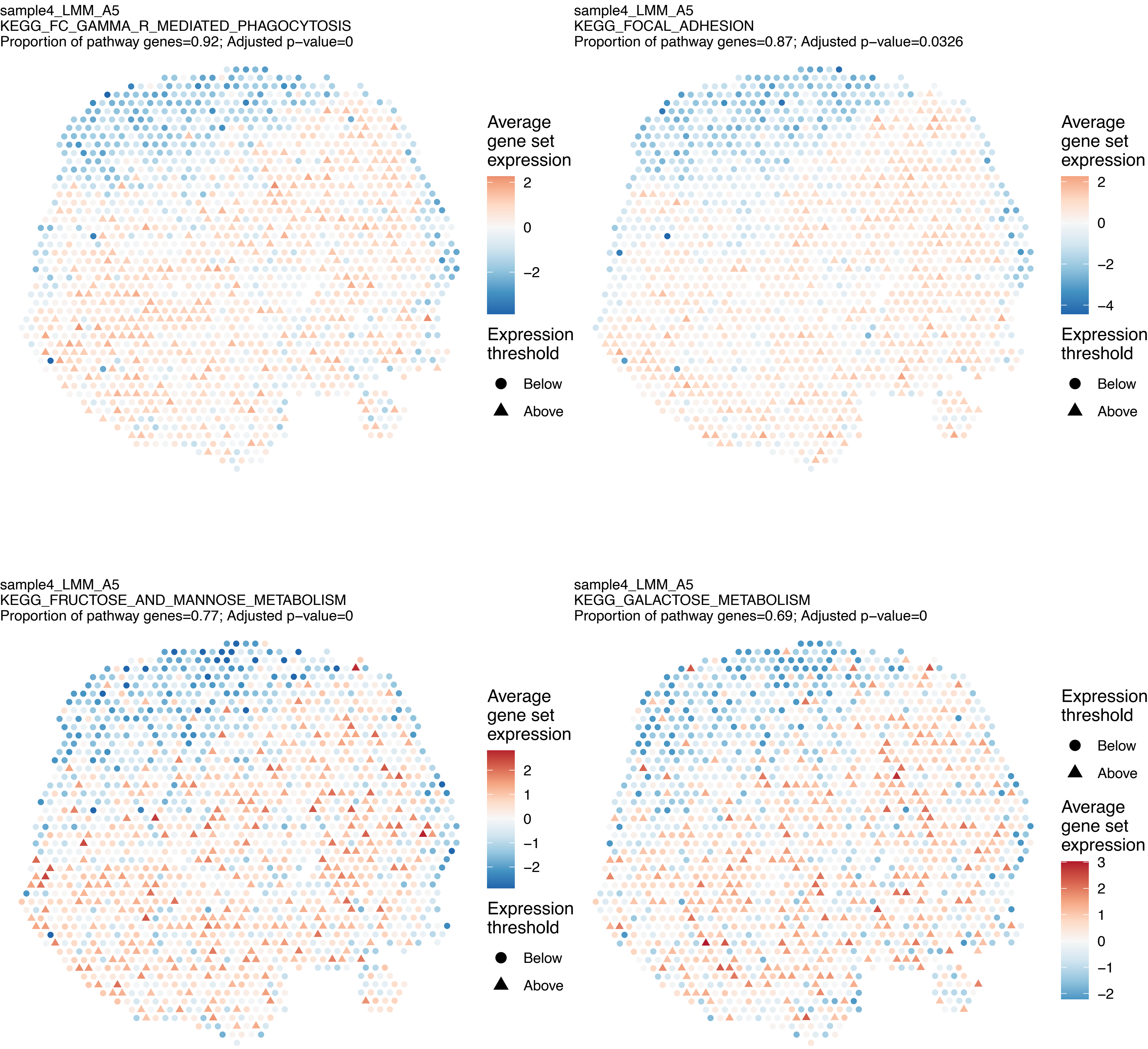
sample4_LMM_A5
KEGG_FC_GAMMA_R_MEDIATED_PHAGOCYTOSIS
Proportion of pathway genes=0.92; Adjusted p−value=0
Average gene set expression
2
0
−2
Expression threshold
Below
Above
sample4_LMM_A5
KEGG_FOCAL_ADHESION
Proportion of pathway genes=0.87; Adjusted p−value=0.0326
Average gene set expression
2
0
−2
−4
Expression threshold
Below
Above
sample4_LMM_A5
KEGG_FRUCTOSE_AND_MANNOSE_METABOLISM
Proportion of pathway genes=0.77; Adjusted p−value=0
Average gene set expression
2
1
0
−1
−2
Expression threshold
Below
Above
sample4_LMM_A5
KEGG_GALACTOSE_METABOLISM
Proportion of pathway genes=0.69; Adjusted p−value=0
Expression threshold
Below
Above
Average gene set expression
3
2
1
0
−1
−2

sample4_LMM_A5
KEGG_GAP_JUNCTION
Proportion of pathway genes=0.74; Adjusted p−value=0

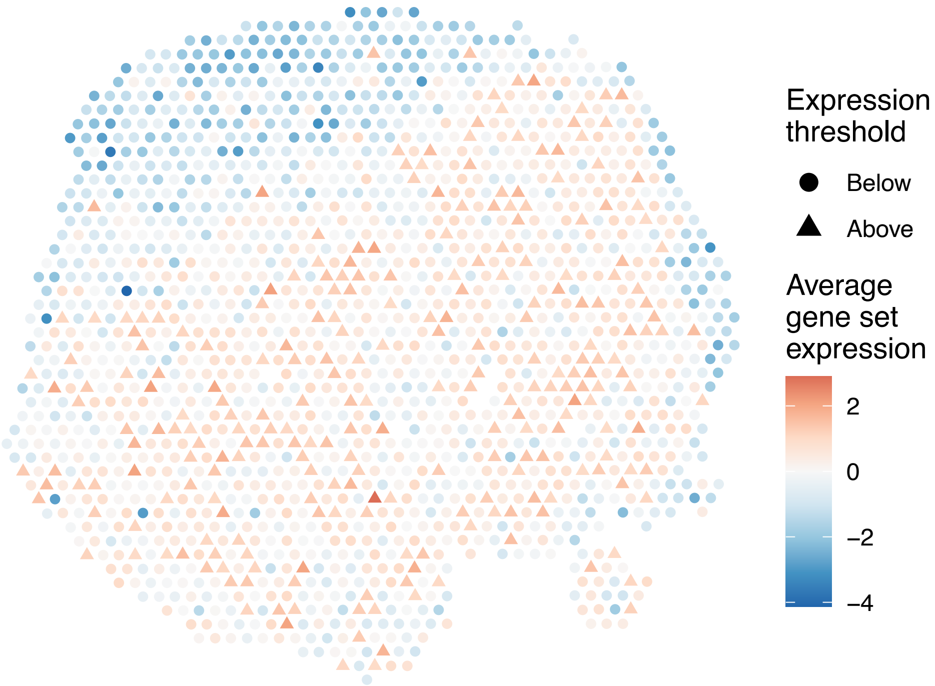


sample4_LMM_A5
KEGG_GLIOMA
Proportion of pathway genes=0.86; Adjusted p−value=0

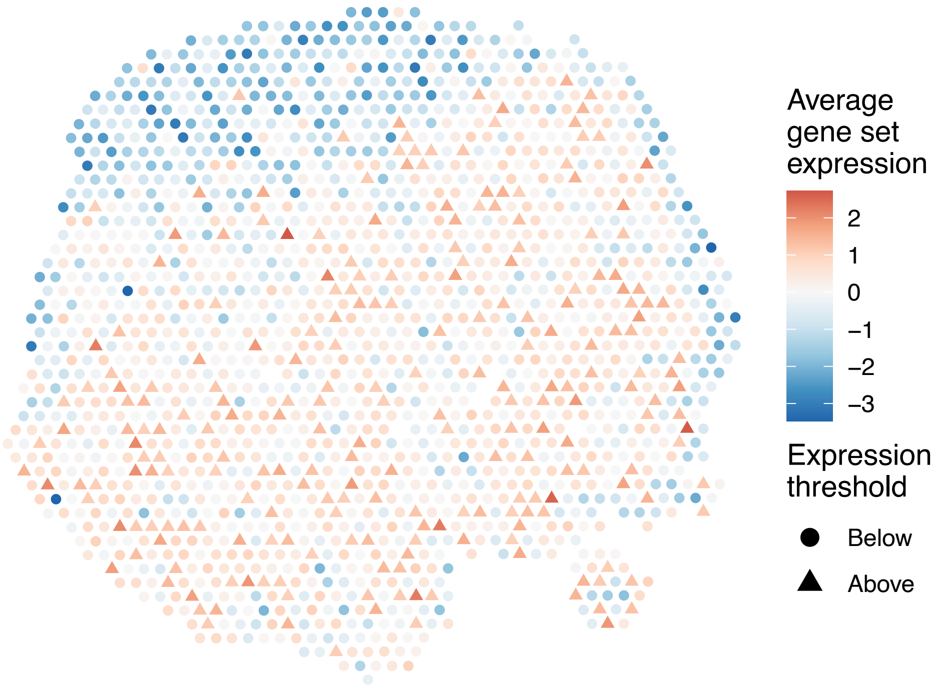


sample4_LMM_A5
KEGG_GLUTATHIONE_METABOLISM
Proportion of pathway genes=0.64; Adjusted p−value=0

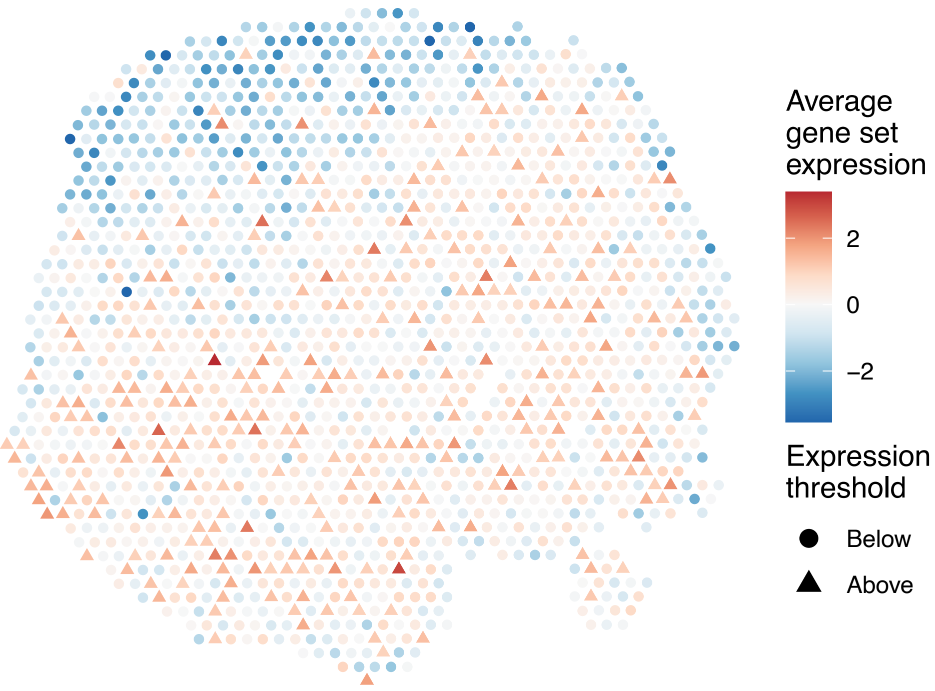


sample4_LMM_A5
KEGG_GLYCEROLIPID_METABOLISM
Proportion of pathway genes=0.76; Adjusted p−value=0

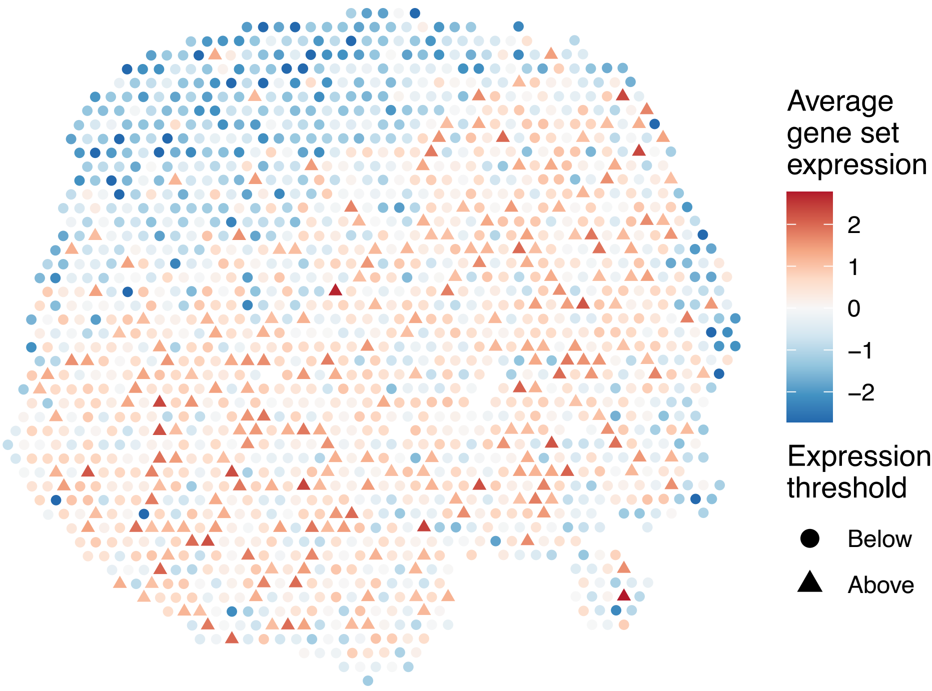

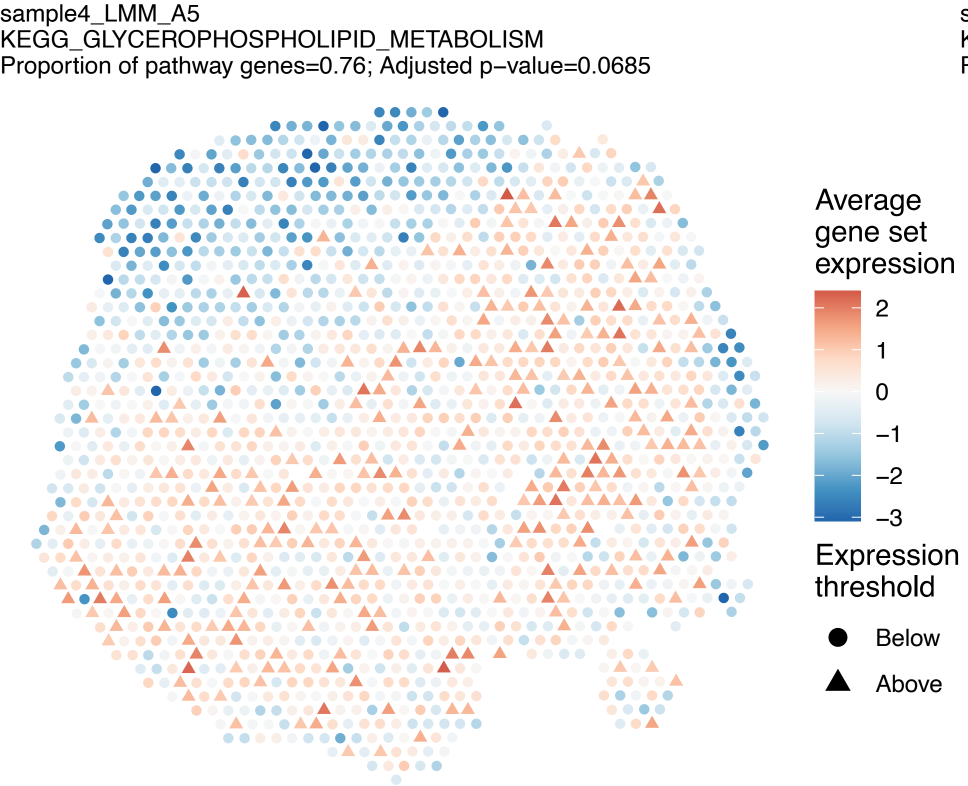
sample4_LMM_A5
KEGG_GLYCEROPHOSPHOLIPID_METABOLISM
Proportion of pathway genes=0.76; Adjusted p−value=0.0685
Average gene set expression
2
1
0
−1
−2
−3
Expression threshold
Below
Above

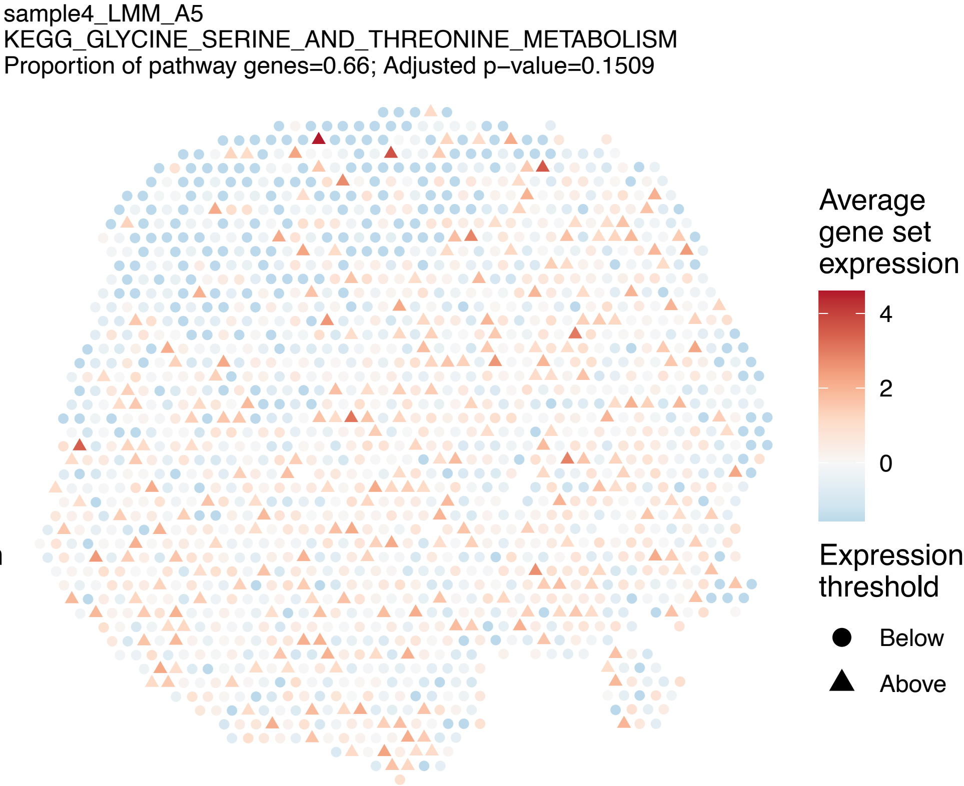
sample4_LMM_A5
KEGG_GLYCINE_SERINE_AND_THREONINE_METABOLISM
Proportion of pathway genes=0.66; Adjusted p−value=0.1509
Average gene set expression
4
2
0
Expression threshold
Below
Above

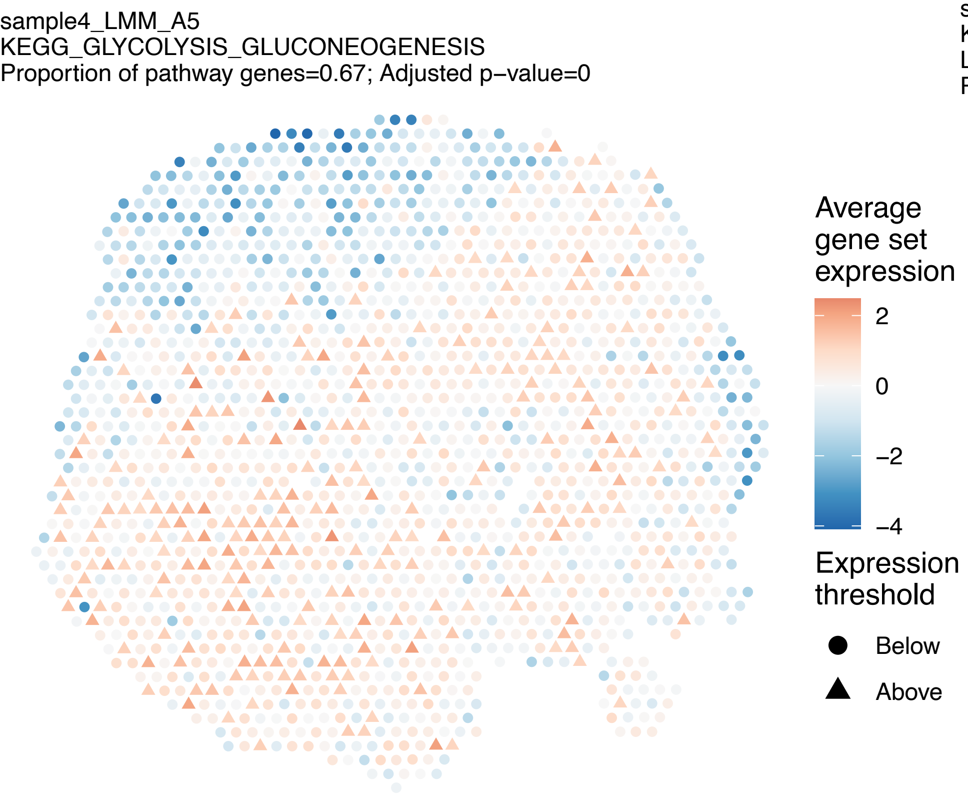
sample4_LMM_A5
KEGG_GLYCOLYSIS_GLUCONEOGENESIS
Proportion of pathway genes=0.67; Adjusted p−value=0
Average gene set expression
2
0
−2
−4
Expression threshold
Below
Above

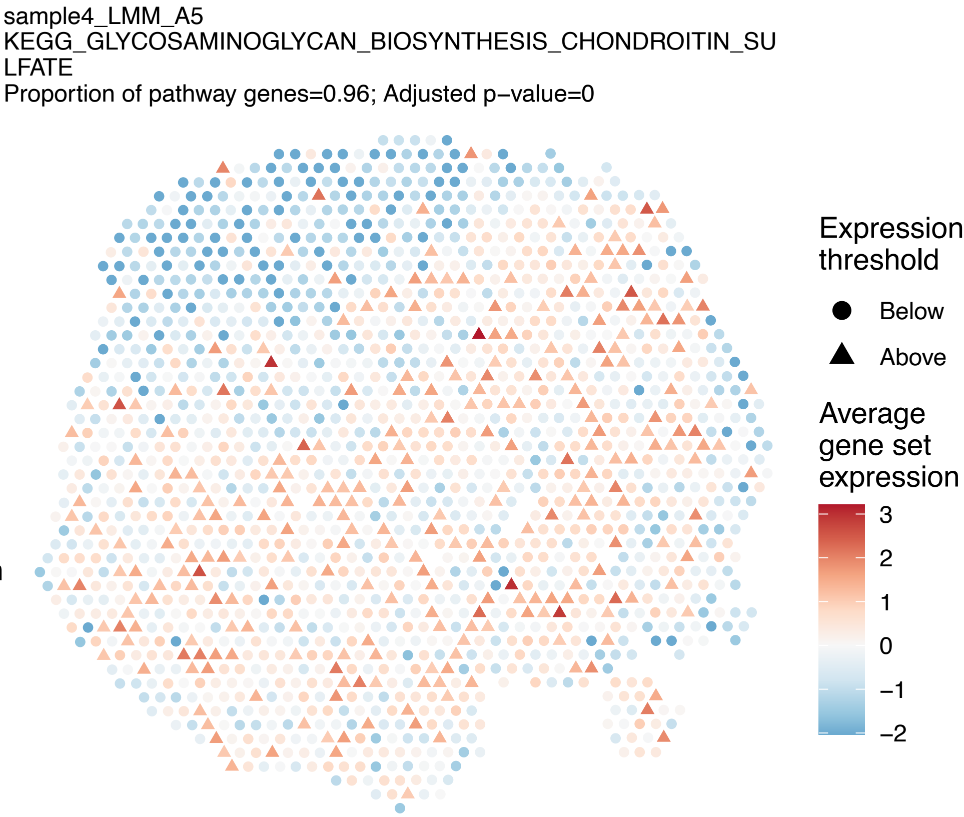
sample4_LMM_A5
KEGG_GLYCOSAMINOGLYCAN_BIOSYNTHESIS_CHONDROITIN_SU
LFATE
Proportion of pathway genes=0.96; Adjusted p−value=0
Expression threshold
Below
Above
Average gene set expression
3
2
1
0
−1
−2

sample4_LMM_A5
KEGG_GLYCOSAMINOGLYCAN_BIOSYNTHESIS_HEPARAN_SULFAT
E
Proportion of pathway genes=0.7; Adjusted p−value=0

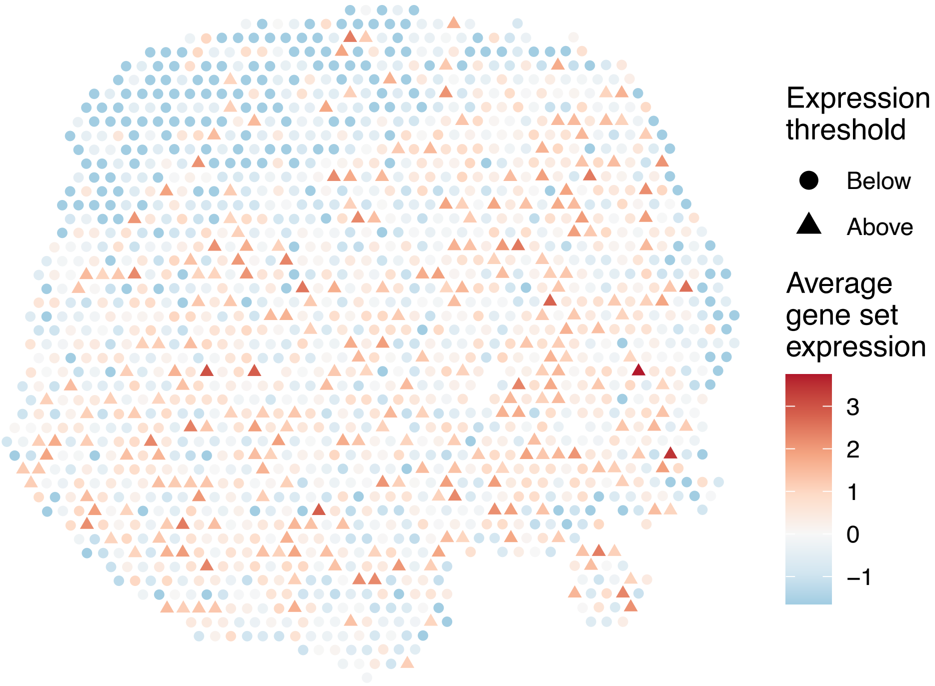


sample4_LMM_A5
KEGG_GLYCOSAMINOGLYCAN_BIOSYNTHESIS_KERATAN_SULFAT
E
Proportion of pathway genes=0.93; Adjusted p−value=0.0261

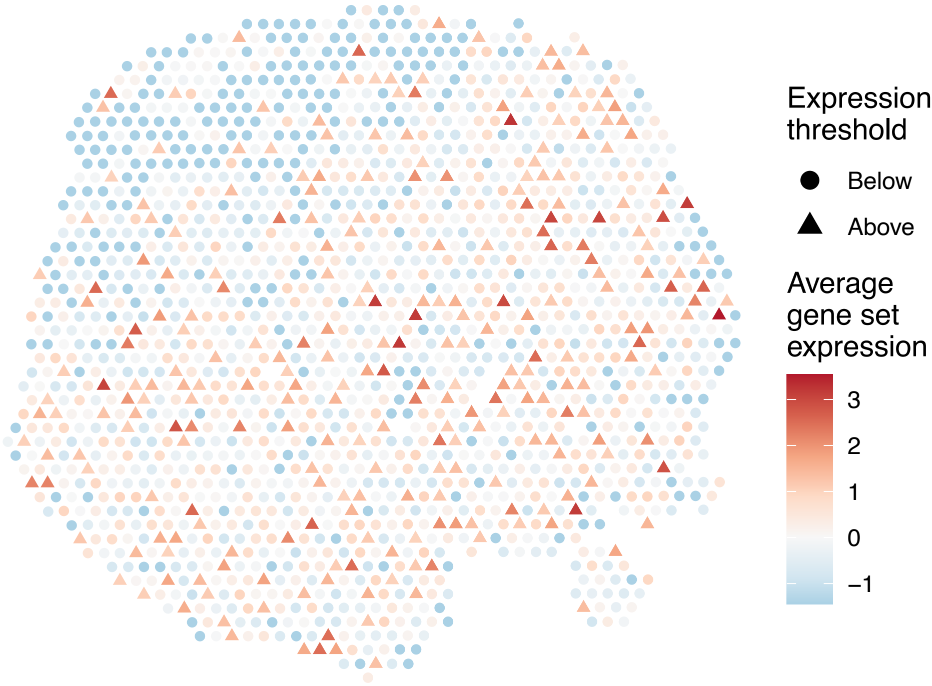


sample4_LMM_A5
KEGG_GLYCOSAMINOGLYCAN_DEGRADATION
Proportion of pathway genes=0.71; Adjusted p−value=0.0052

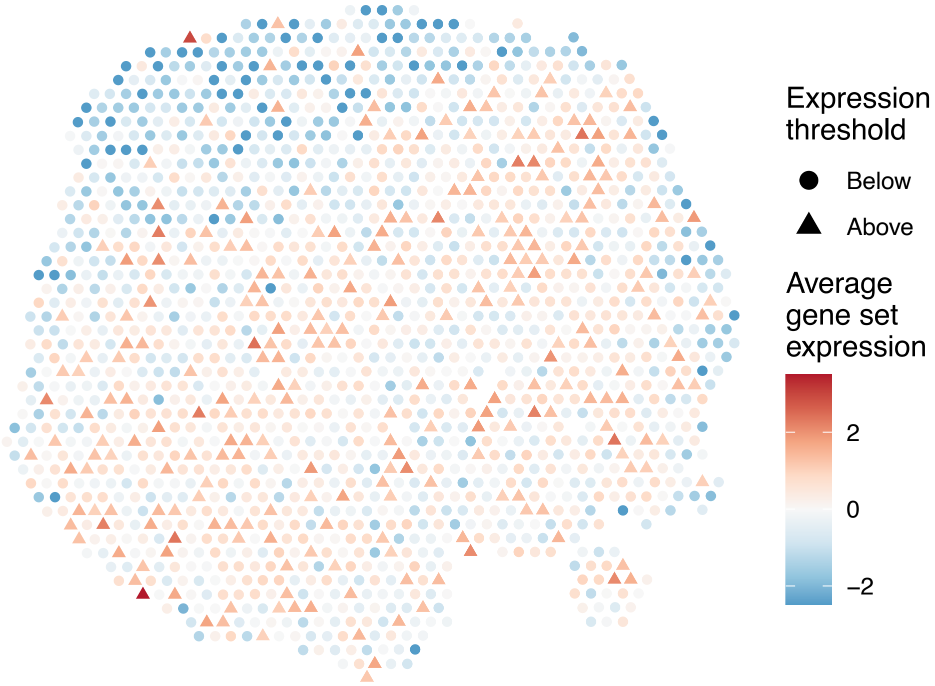


sample4_LMM_A5
KEGG_GLYCOSPHINGOLIPID_BIOSYNTHESIS_GANGLIO_SERIES
Proportion of pathway genes=0.84; Adjusted p−value=0.0951

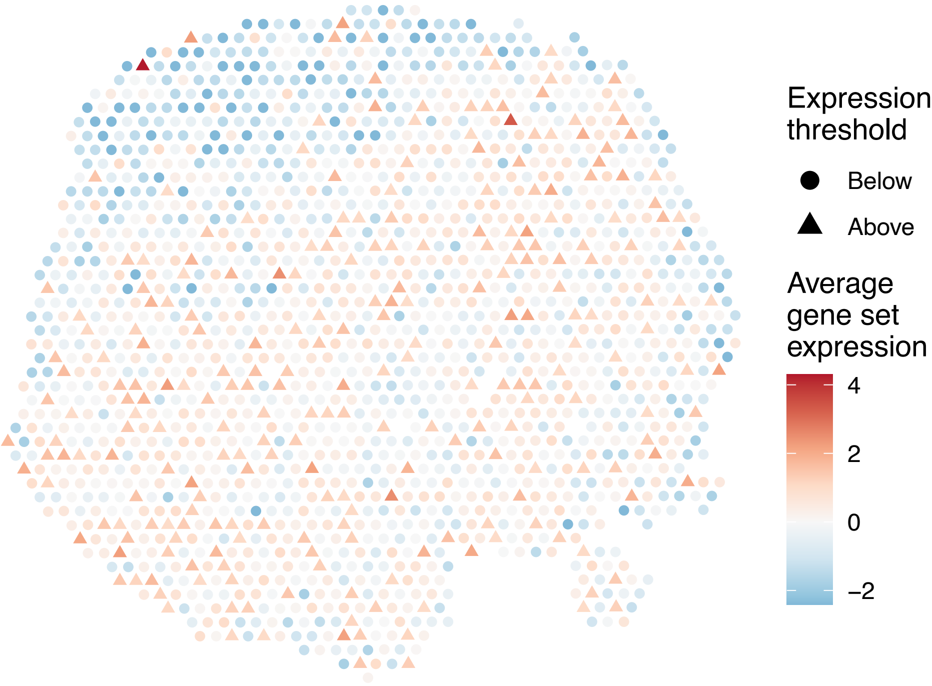

sample4_LMM_A5
KEGG_GLYCOSPHINGOLIPID_BIOSYNTHESIS_GLOBO_SERIES
Proportion of pathway genes=0.71; Adjusted p−value=0.0052

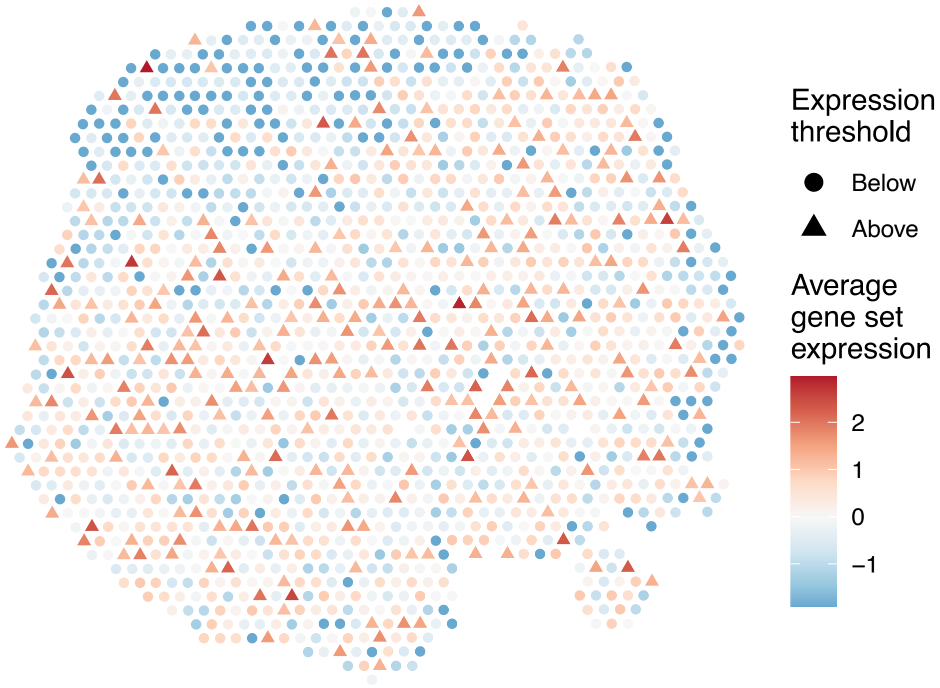


sample4_LMM_A5
KEGG_GLYCOSPHINGOLIPID_BIOSYNTHESIS_LACTO_AND_NEOL
ACTO_SERIES
Proportion of pathway genes=0.52; Adjusted p−value=0.0075

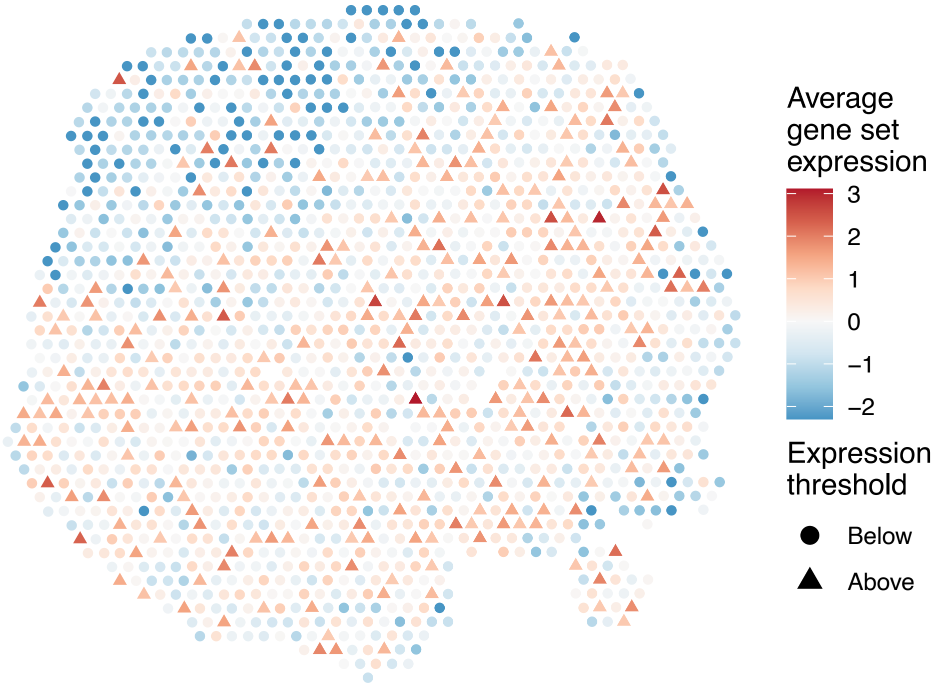


sample4_LMM_A5
KEGG_GLYCOSYLPHOSPHATIDYLINOSITOL_GPI_ANCHOR_BIOSY
NTHESIS
Proportion of pathway genes=0.96; Adjusted p−value=0.0014

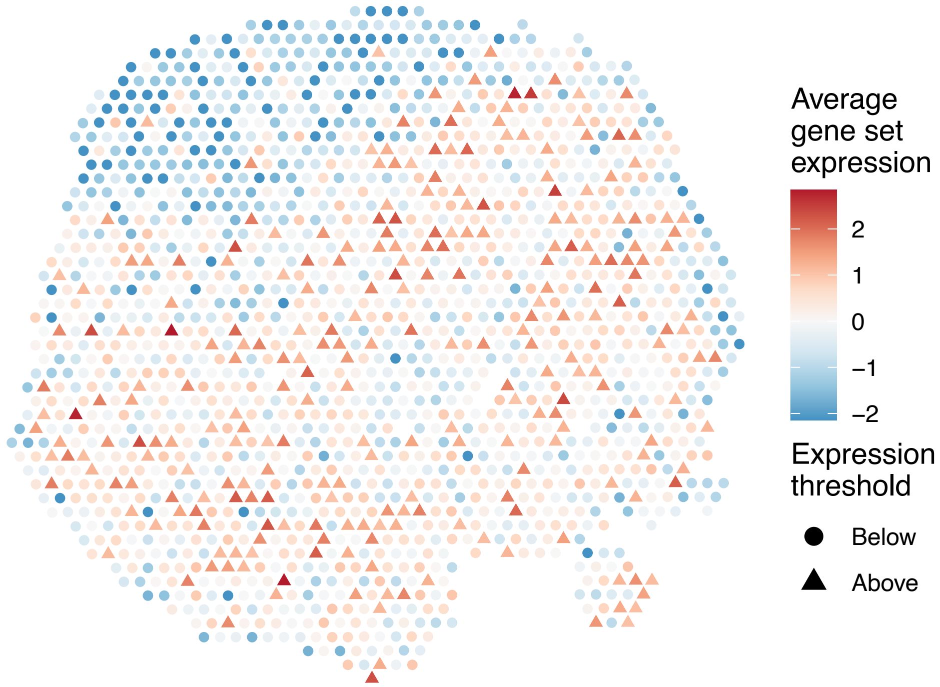


sample4_LMM_A5
KEGG_GLYOXYLATE_AND_DICARBOXYLATE_METABOLISM
Proportion of pathway genes=0.88; Adjusted p−value=0

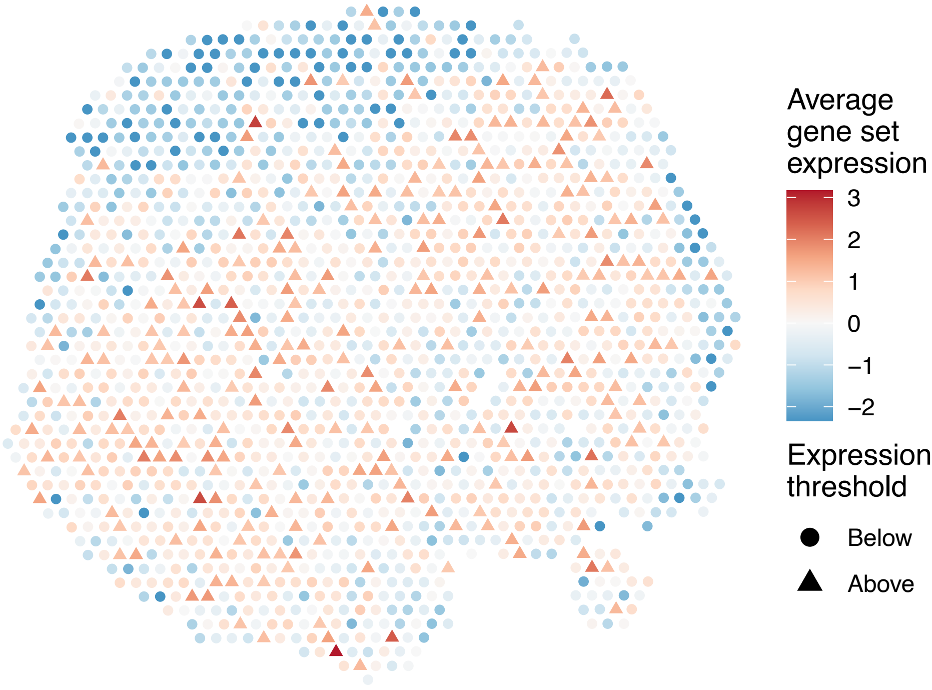

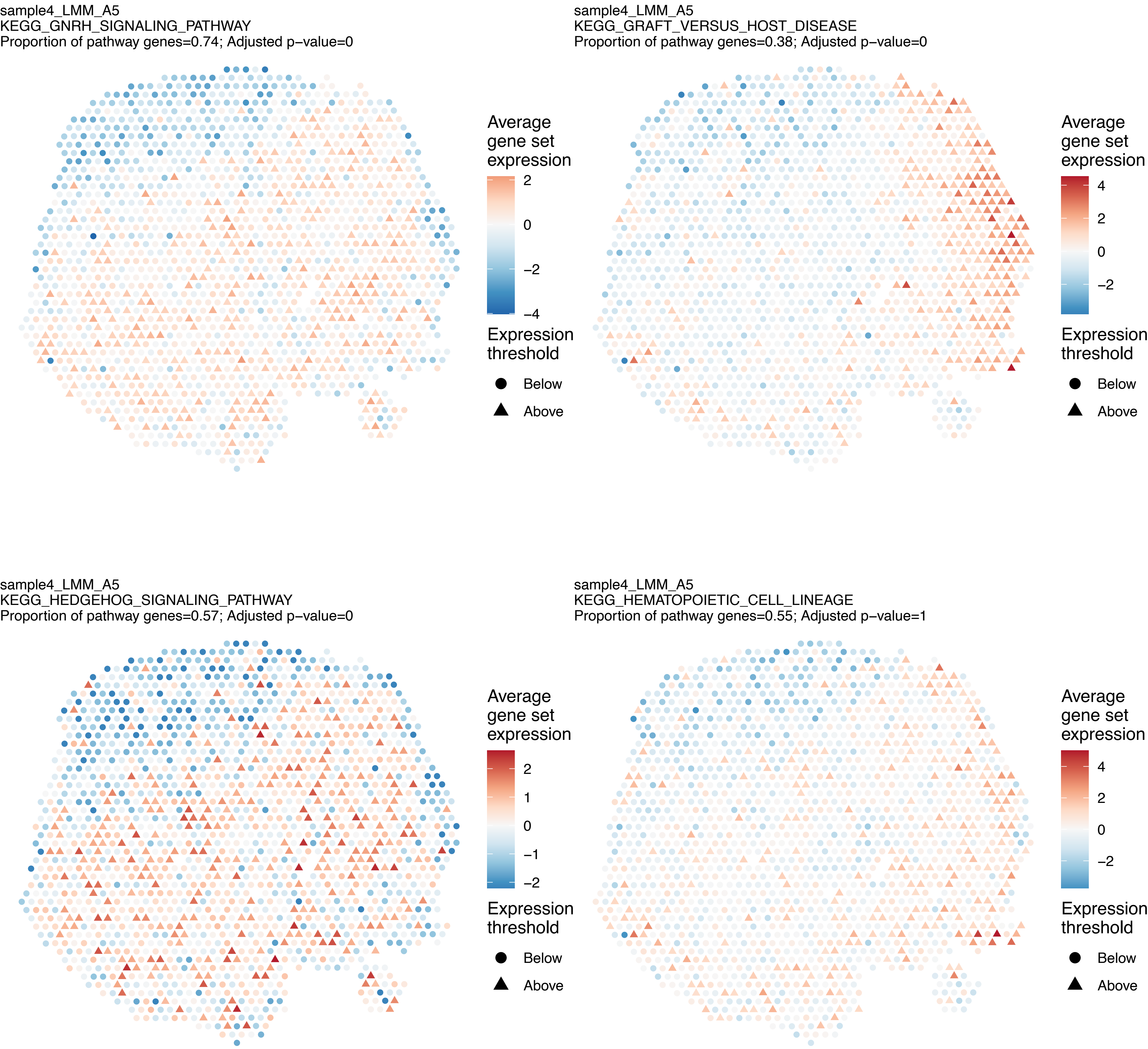
sample4_LMM_A5
KEGG_GNRH_SIGNALING_PATHWAY
Proportion of pathway genes=0.74; Adjusted p−value=0
Average gene set expression
2
0
−2
−4
Expression threshold
Below
Above
sample4_LMM_A5
KEGG_GRAFT_VERSUS_HOST_DISEASE
Proportion of pathway genes=0.38; Adjusted p−value=0
Average gene set expression
4
2
0
−2
Expression threshold
Below
Above
sample4_LMM_A5
KEGG_HEDGEHOG_SIGNALING_PATHWAY
Proportion of pathway genes=0.57; Adjusted p−value=0
Average gene set expression
2
1
0
−1
−2
Expression threshold
Below
Above
sample4_LMM_A5
KEGG_HEMATOPOIETIC_CELL_LINEAGE
Proportion of pathway genes=0.55; Adjusted p−value=1
Average gene set expression
4
2
0
−2
Expression threshold
Below
Above

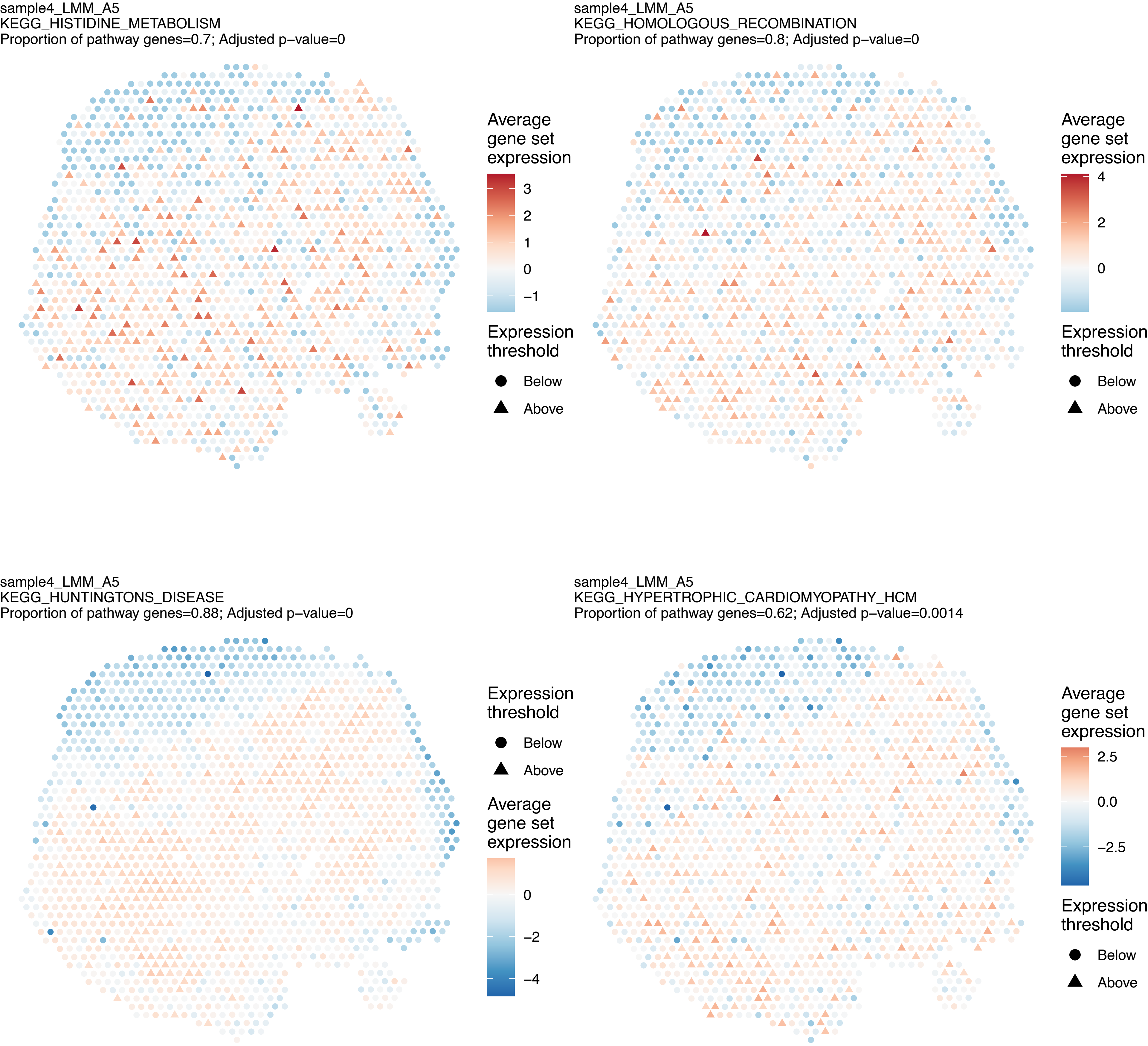
sample4_LMM_A5
KEGG_HISTIDINE_METABOLISM
Proportion of pathway genes=0.7; Adjusted p−value=0
Average gene set expression
3
2
1
0
−1
Expression threshold
Below
Above
sample4_LMM_A5
KEGG_HOMOLOGOUS_RECOMBINATION
Proportion of pathway genes=0.8; Adjusted p−value=0
Average gene set expression
4
2
0
Expression threshold
Below
Above
sample4_LMM_A5
KEGG_HUNTINGTONS_DISEASE
Proportion of pathway genes=0.88; Adjusted p−value=0
Expression threshold
Below
Above
Average gene set expression
0
−2
−4
sample4_LMM_A5
KEGG_HYPERTROPHIC_CARDIOMYOPATHY_HCM
Proportion of pathway genes=0.62; Adjusted p−value=0.0014
Average gene set expression
2.5
0.0
−2.5
Expression threshold
Below
Above

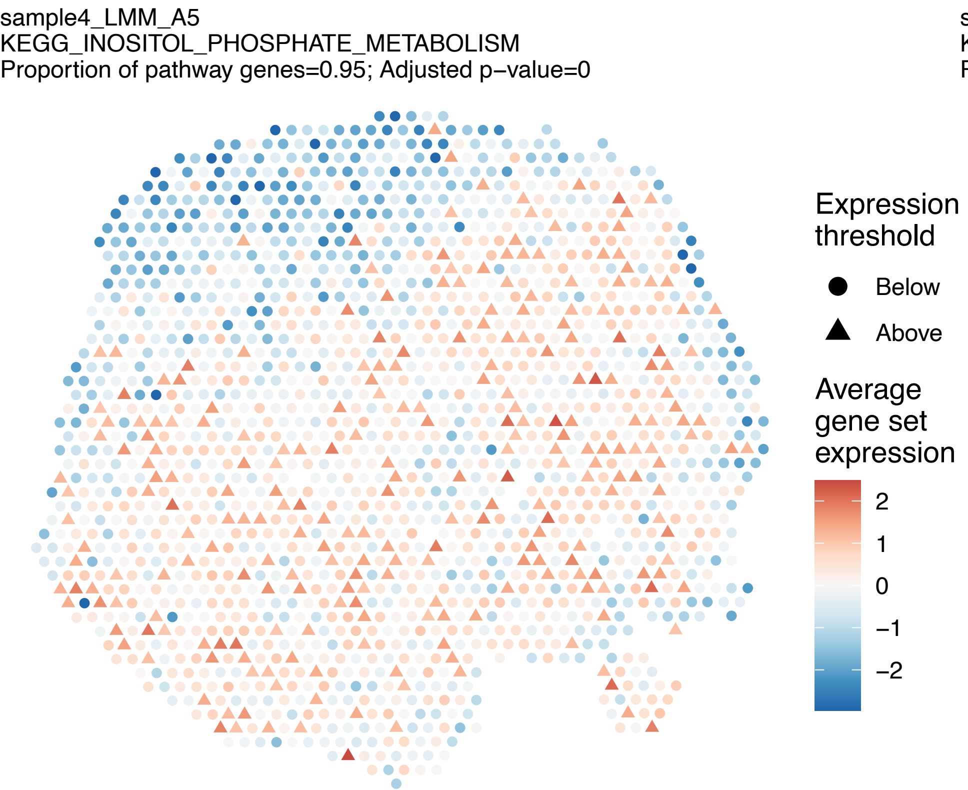
sample4_LMM_A5
KEGG_INOSITOL_PHOSPHATE_METABOLISM
Proportion of pathway genes=0.95; Adjusted p−value=0
Expression threshold
Below
Above
Average gene set expression
2
1
0
−1
−2

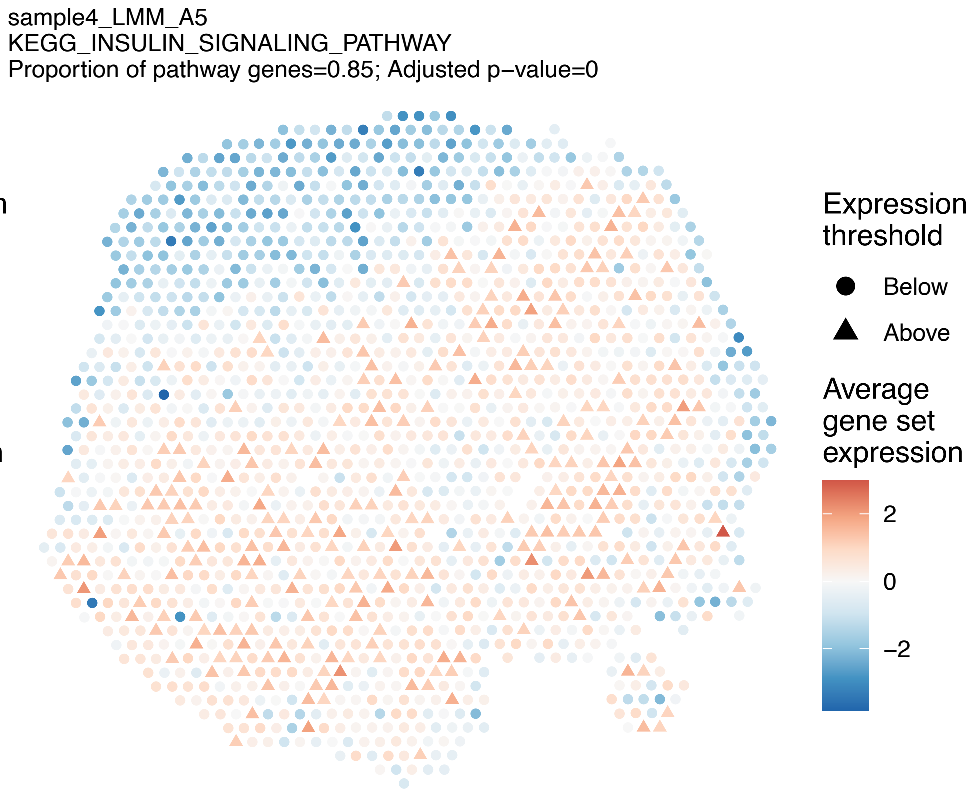
sample4_LMM_A5
KEGG_INSULIN_SIGNALING_PATHWAY
Proportion of pathway genes=0.85; Adjusted p−value=0
Expression threshold
Below
Above
Average gene set expression
2
0
−2

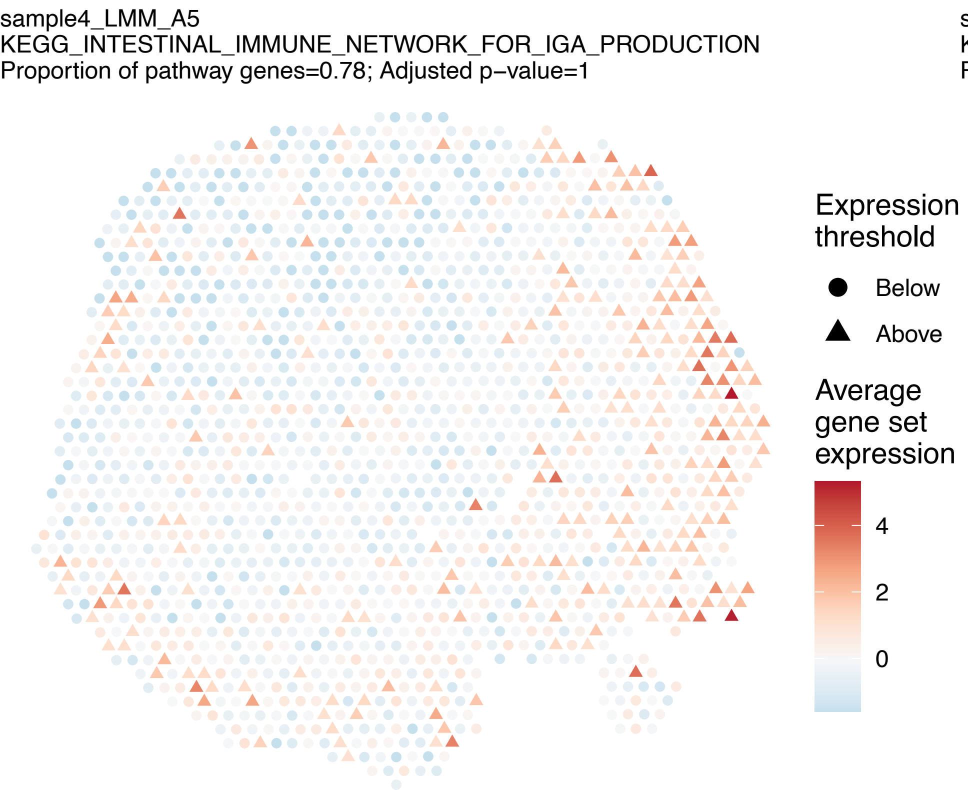
sample4_LMM_A5
KEGG_INTESTINAL_IMMUNE_NETWORK_FOR_IGA_PRODUCTION
Proportion of pathway genes=0.78; Adjusted p−value=1
Expression threshold
Below
Above
Average gene set expression
4
2
0

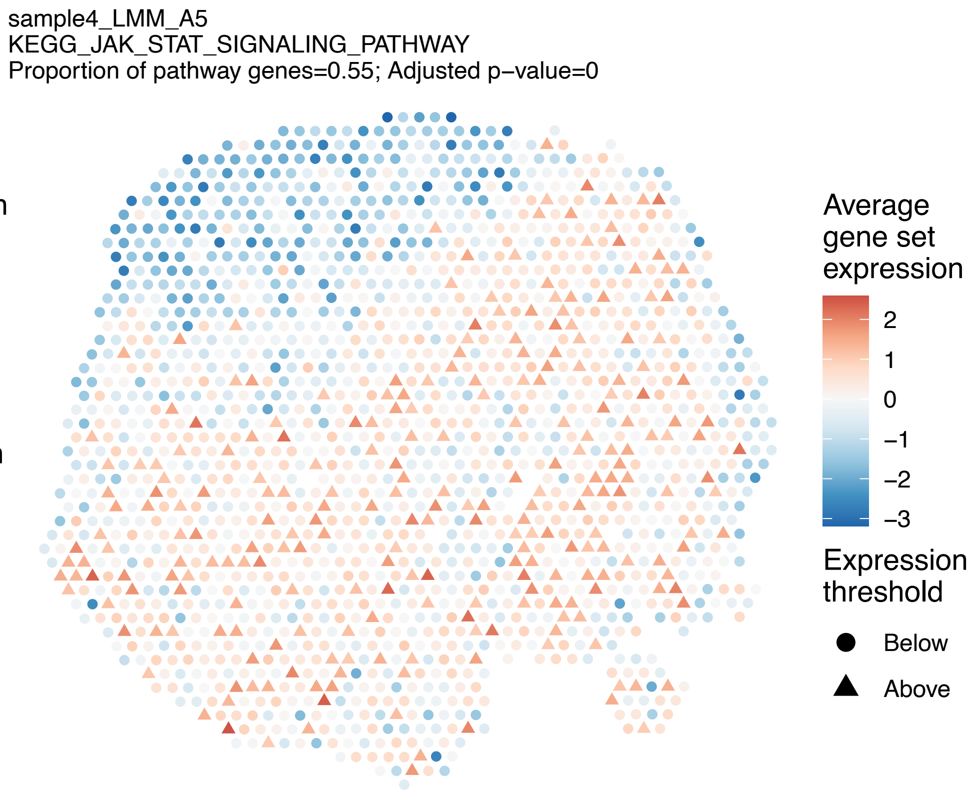
sample4_LMM_A5
KEGG_JAK_STAT_SIGNALING_PATHWAY
Proportion of pathway genes=0.55; Adjusted p−value=0
Average gene set expression
2
1
0
−1
−2
−3
Expression threshold
Below
Above

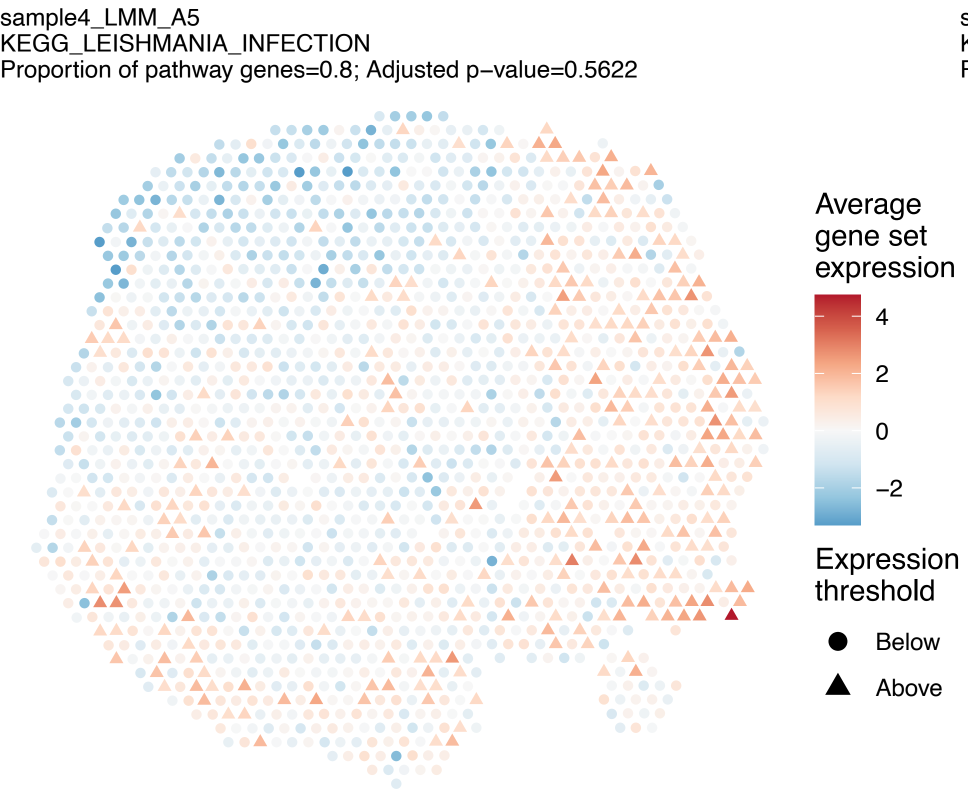
sample4_LMM_A5
KEGG_LEISHMANIA_INFECTION
Proportion of pathway genes=0.8; Adjusted p−value=0.5622
Average gene set expression
4
2
0
−2
Expression threshold
Below
Above

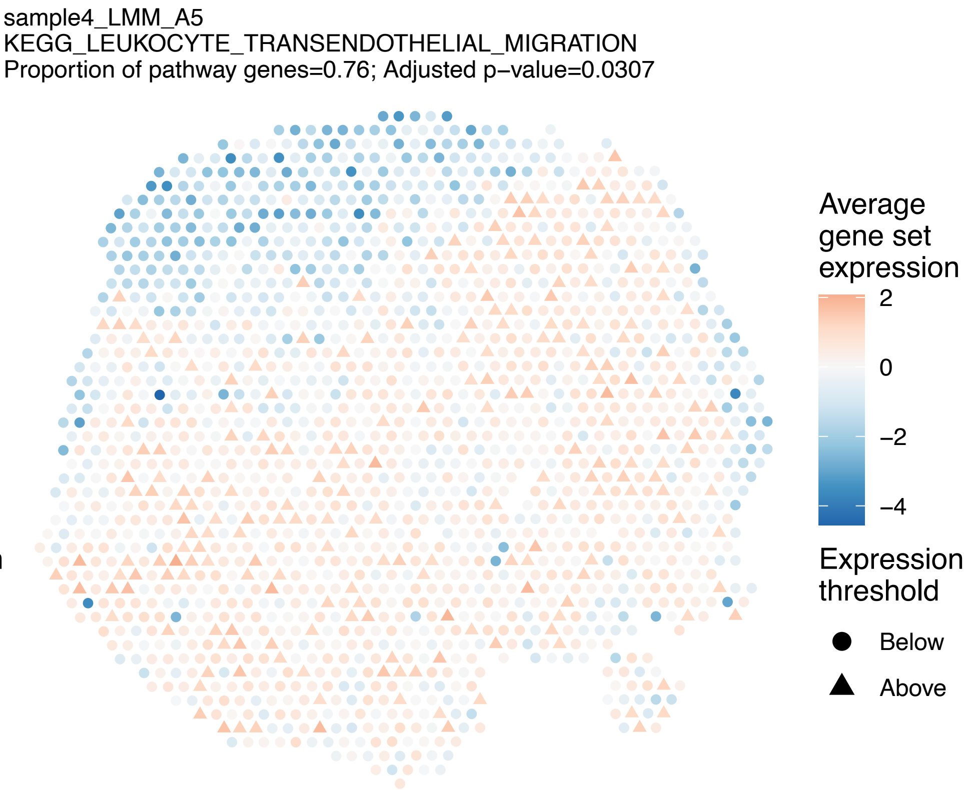
sample4_LMM_A5
KEGG_LEUKOCYTE_TRANSENDOTHELIAL_MIGRATION
Proportion of pathway genes=0.76; Adjusted p−value=0.0307
Average gene set expression
2
0
−2
−4
Expression threshold
Below
Above

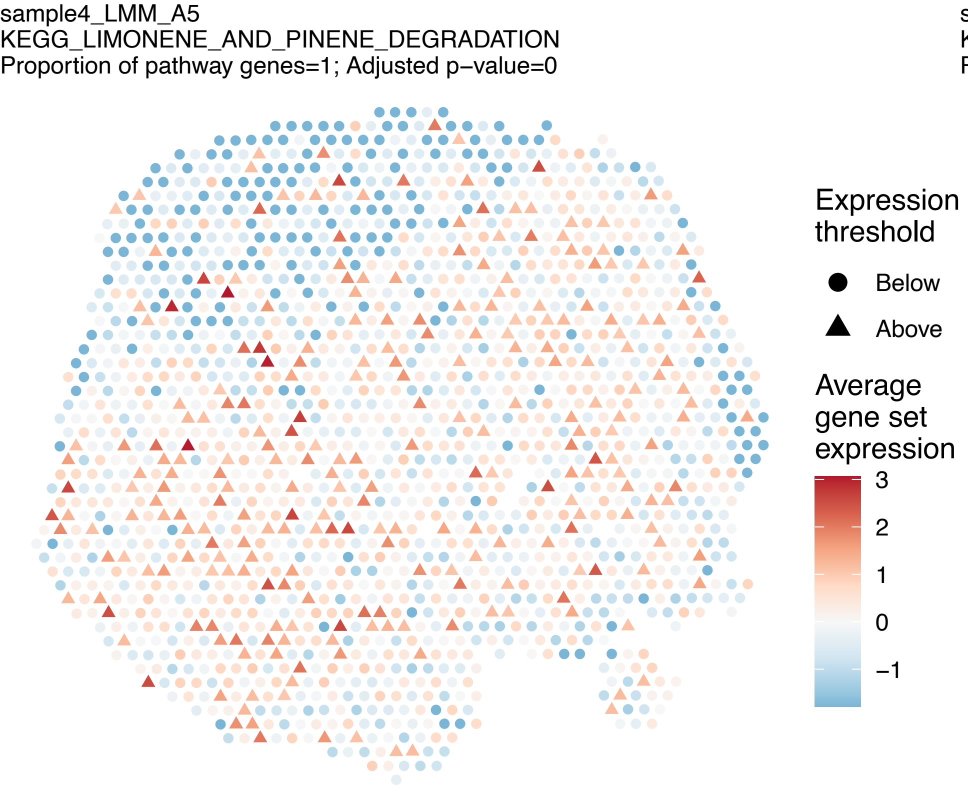
sample4_LMM_A5
KEGG_LIMONENE_AND_PINENE_DEGRADATION
Proportion of pathway genes=1; Adjusted p−value=0
Expression threshold
Below
Above
Average gene set expression
3
2
1
0
−1

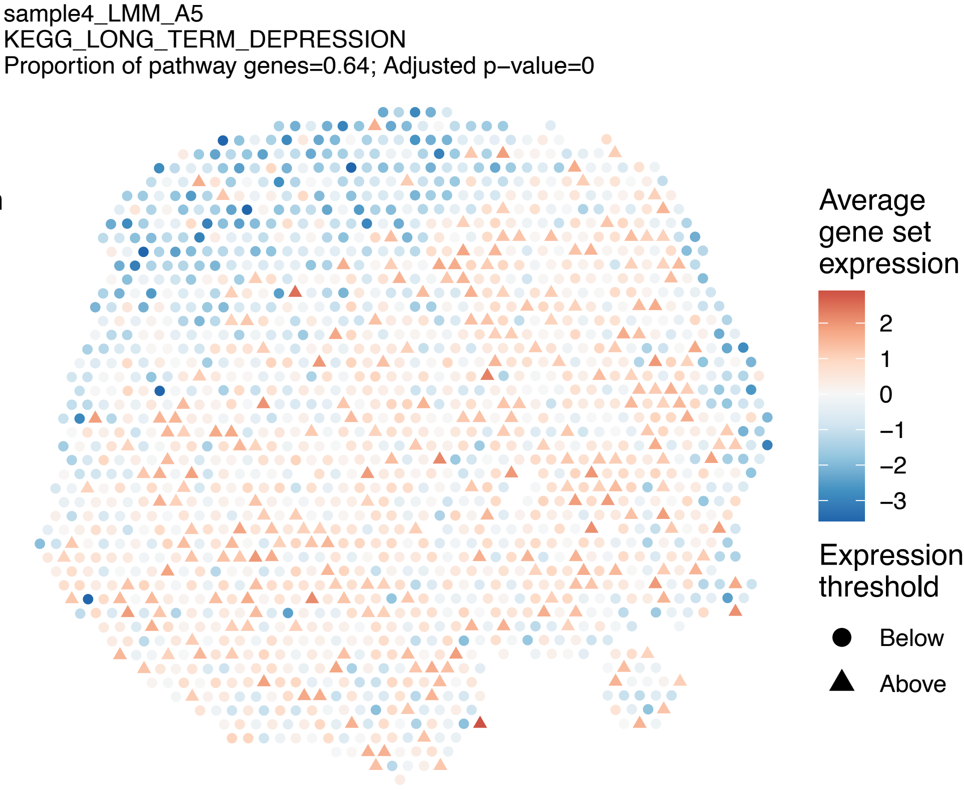
sample4_LMM_A5
KEGG_LONG_TERM_DEPRESSION
Proportion of pathway genes=0.64; Adjusted p−value=0
Average gene set expression
2
1
0
−1
−2
−3
Expression threshold
Below
Above

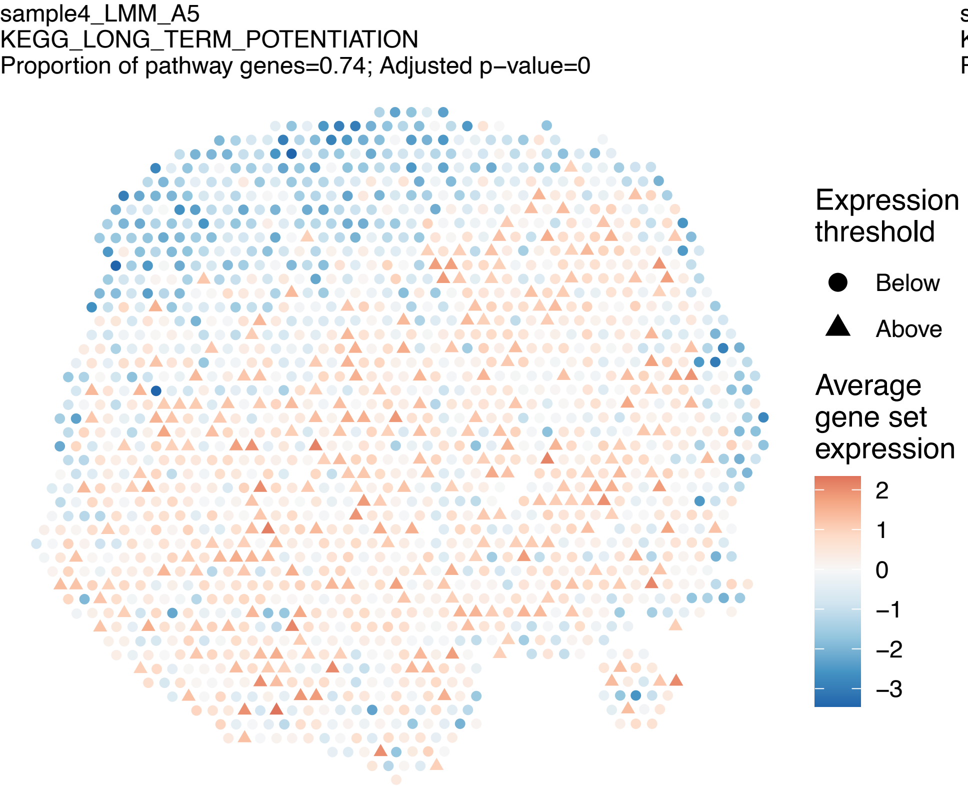
sample4_LMM_A5
KEGG_LONG_TERM_POTENTIATION
Proportion of pathway genes=0.74; Adjusted p−value=0
Expression threshold
Below
Above
Average gene set expression
2
1
0
−1
−2
−3

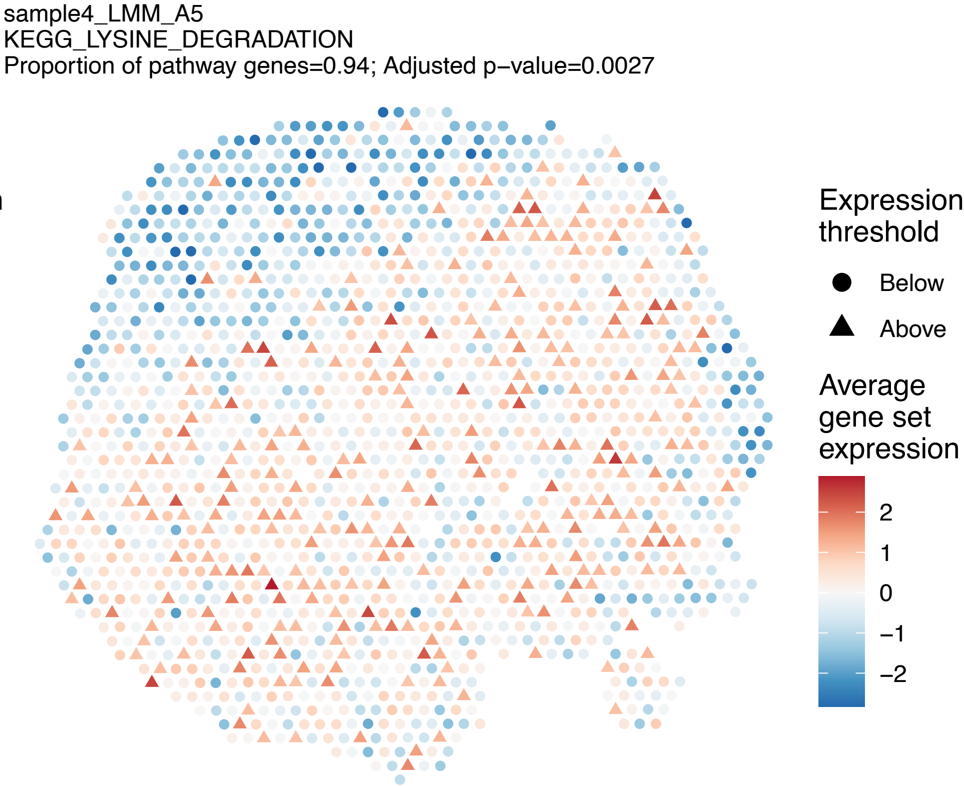
sample4_LMM_A5
KEGG_LYSINE_DEGRADATION
Proportion of pathway genes=0.94; Adjusted p−value=0.0027
Expression threshold
Below
Above
Average gene set expression
2
1
0
−1
−2

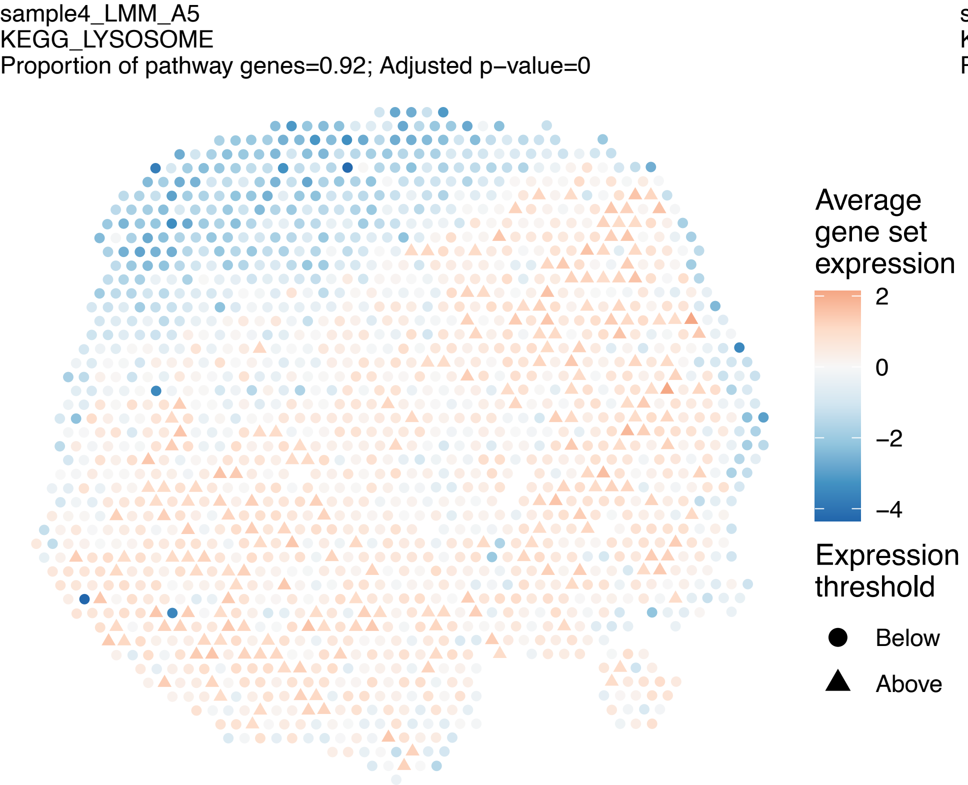
sample4_LMM_A5
KEGG_LYSOSOME
Proportion of pathway genes=0.92; Adjusted p−value=0
Average gene set expression
2
0
−2
−4
Expression threshold
Below
Above

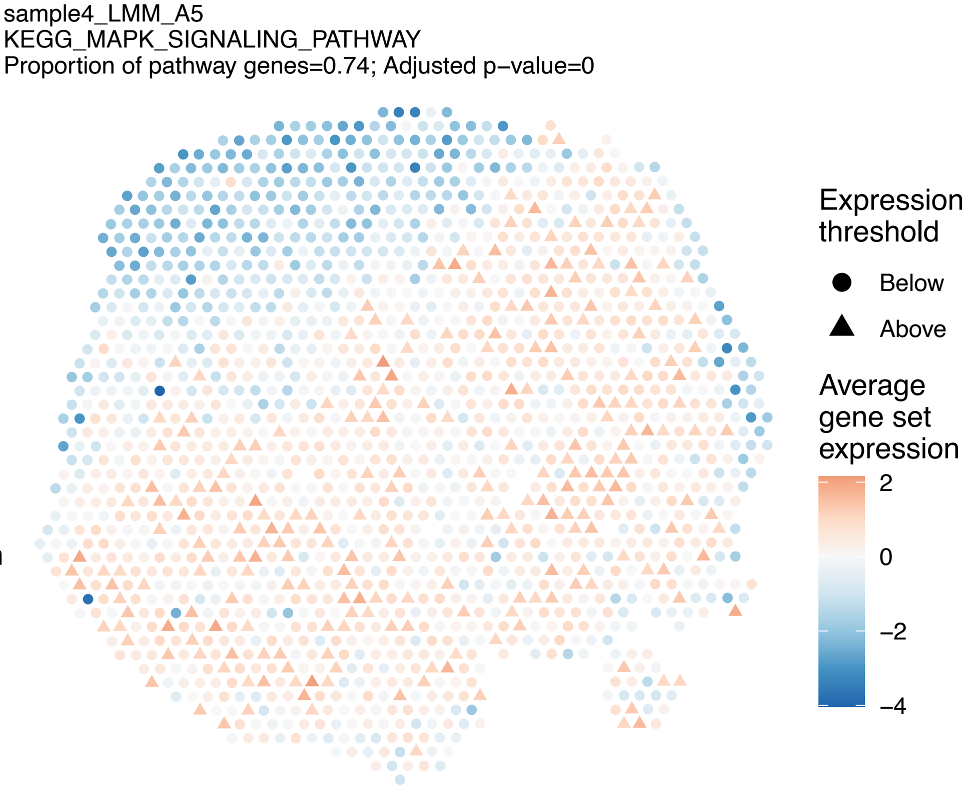
sample4_LMM_A5
KEGG_MAPK_SIGNALING_PATHWAY
Proportion of pathway genes=0.74; Adjusted p−value=0
Expression threshold
Below
Above
Average gene set expression
2
0
−2
−4

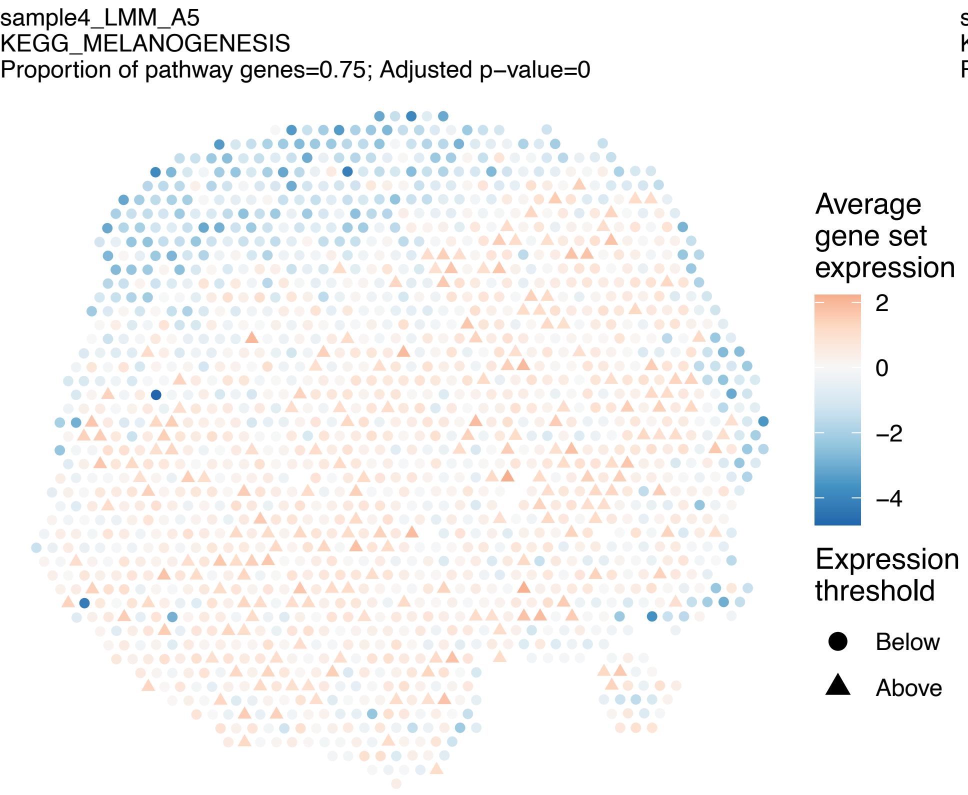
sample4_LMM_A5
KEGG_MELANOGENESIS
Proportion of pathway genes=0.75; Adjusted p−value=0
Average gene set expression
2
0
−2
−4
Expression threshold
Below
Above

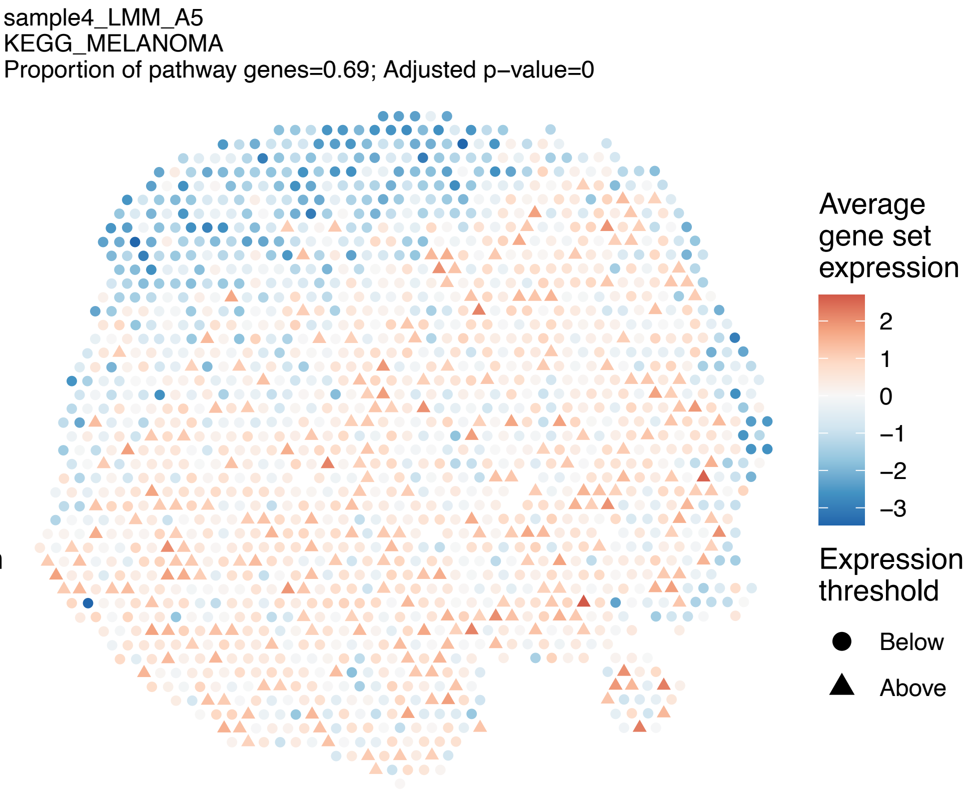
sample4_LMM_A5
KEGG_MELANOMA
Proportion of pathway genes=0.69; Adjusted p−value=0
Average gene set expression
2
1
0
−1
−2
−3
Expression threshold
Below
Above

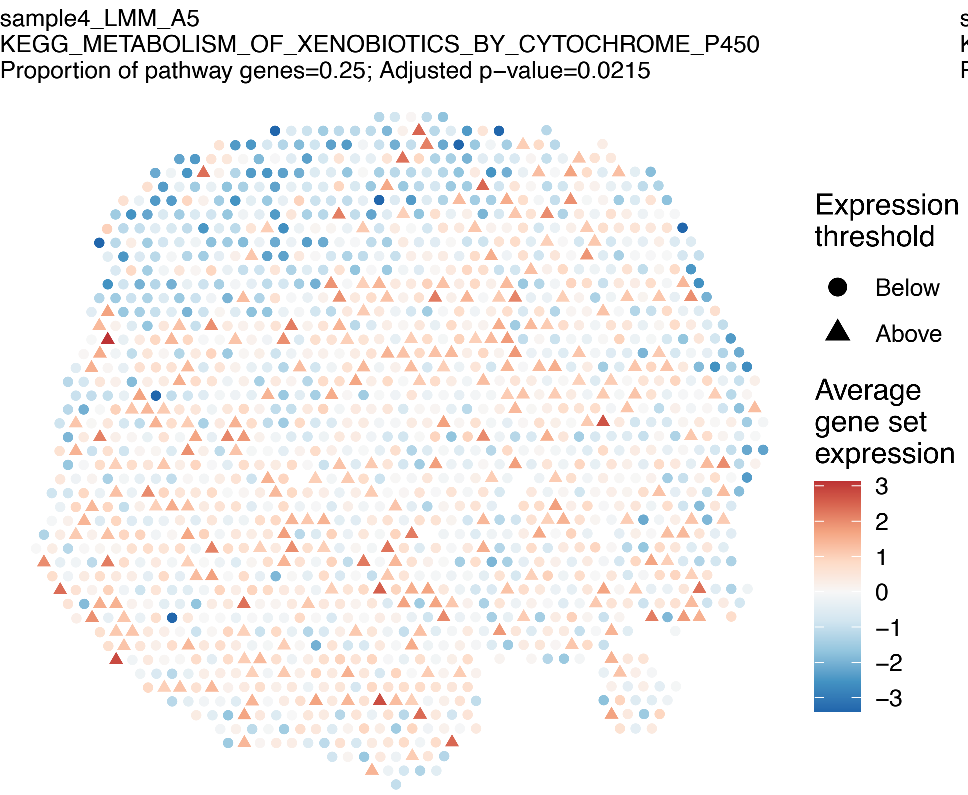
sample4_LMM_A5
KEGG_METABOLISM_OF_XENOBIOTICS_BY_CYTOCHROME_P450
Proportion of pathway genes=0.25; Adjusted p−value=0.0215
Expression threshold
Below
Above
Average gene set expression
3
2
1
0
−1
−2
−3

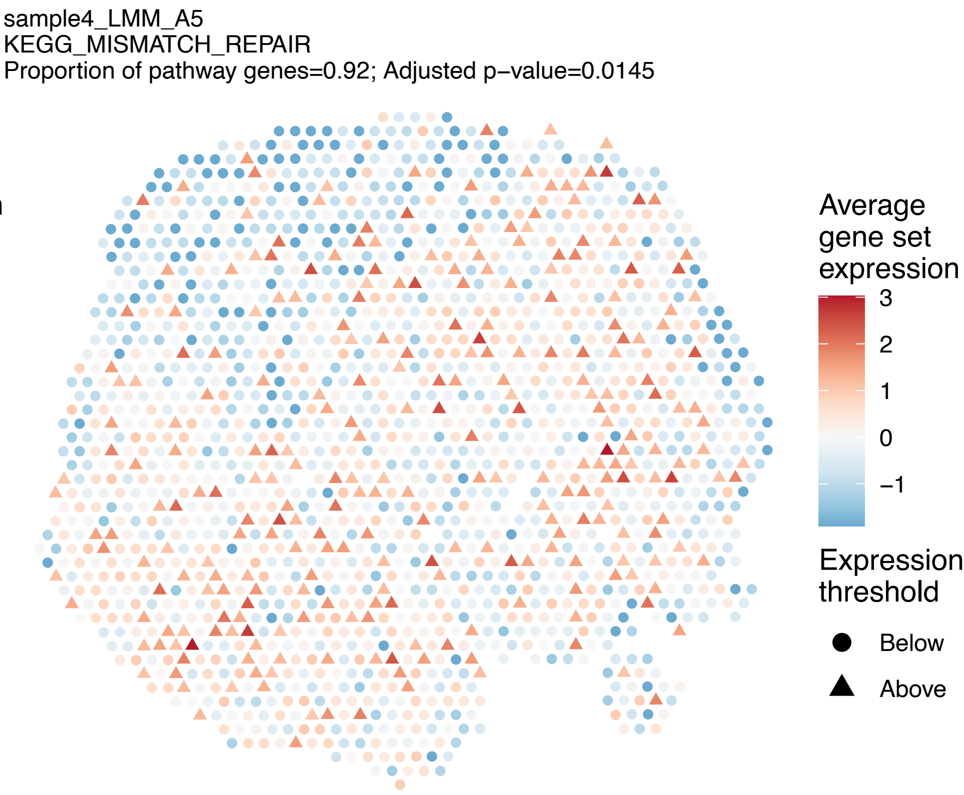
sample4_LMM_A5
KEGG_MISMATCH_REPAIR
Proportion of pathway genes=0.92; Adjusted p−value=0.0145
Average gene set expression
3
2
1
0
−1
Expression threshold
Below
Above

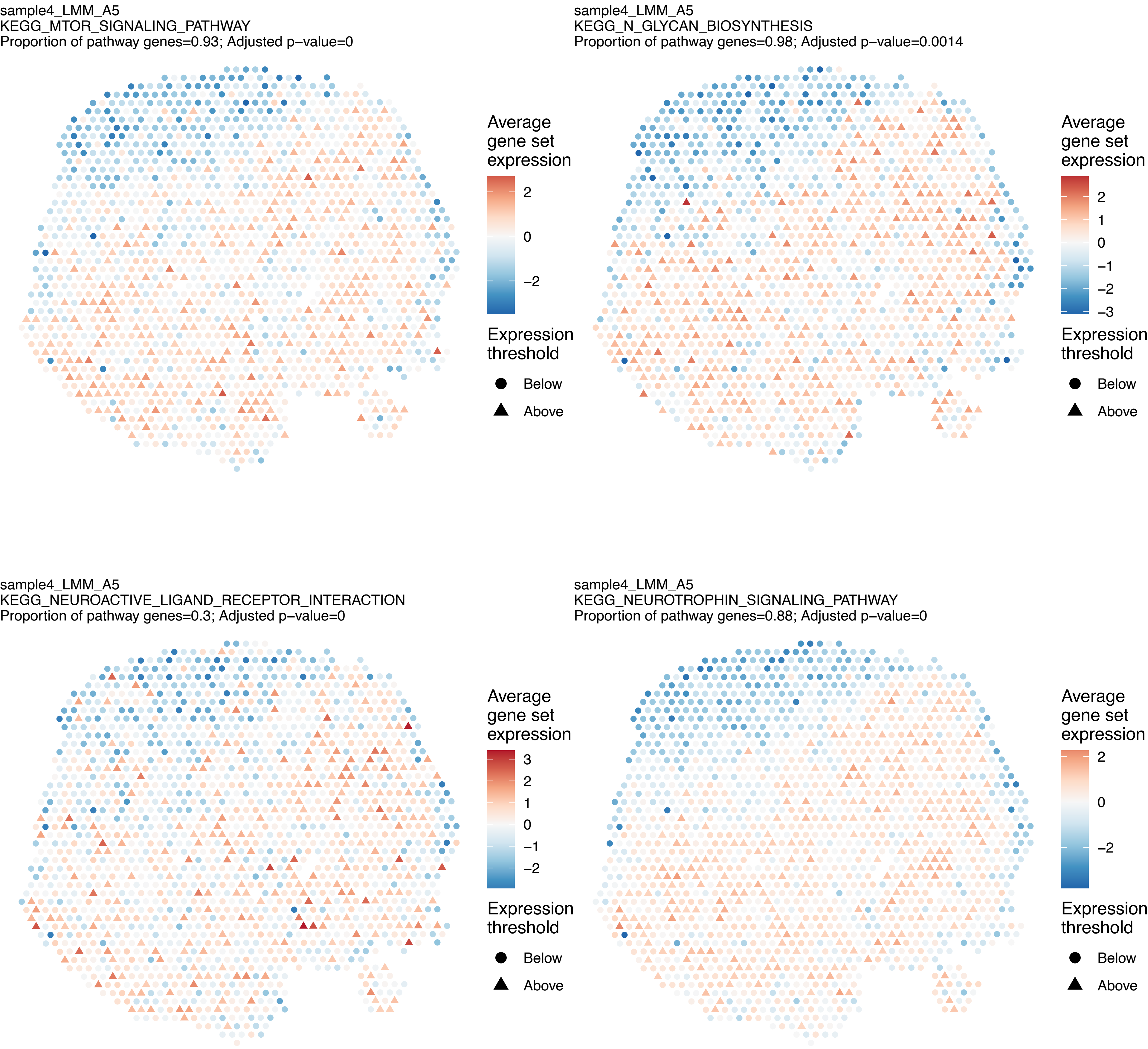

sample4_LMM_A5
KEGG_MTOR_SIGNALING_PATHWAY
Proportion of pathway genes=0.93; Adjusted p−value=0
Average gene set expression
2
0
−2
Expression threshold
Below
Above
sample4_LMM_A5
KEGG_N_GLYCAN_BIOSYNTHESIS
Proportion of pathway genes=0.98; Adjusted p−value=0.0014
Average gene set expression
2
1
0
−1
−2
−3
Expression threshold
Below
Above
sample4_LMM_A5
KEGG_NEUROACTIVE_LIGAND_RECEPTOR_INTERACTION
Proportion of pathway genes=0.3; Adjusted p−value=0
Average gene set expression
3
2
1
0
−1
−2
Expression threshold
Below
Above
sample4_LMM_A5
KEGG_NEUROTROPHIN_SIGNALING_PATHWAY
Proportion of pathway genes=0.88; Adjusted p−value=0
Average gene set expression
2
0
−2
Expression threshold
Below
Above

sample4_LMM_A5
KEGG_NICOTINATE_AND_NICOTINAMIDE_METABOLISM
Proportion of pathway genes=0.83; Adjusted p−value=0.3446

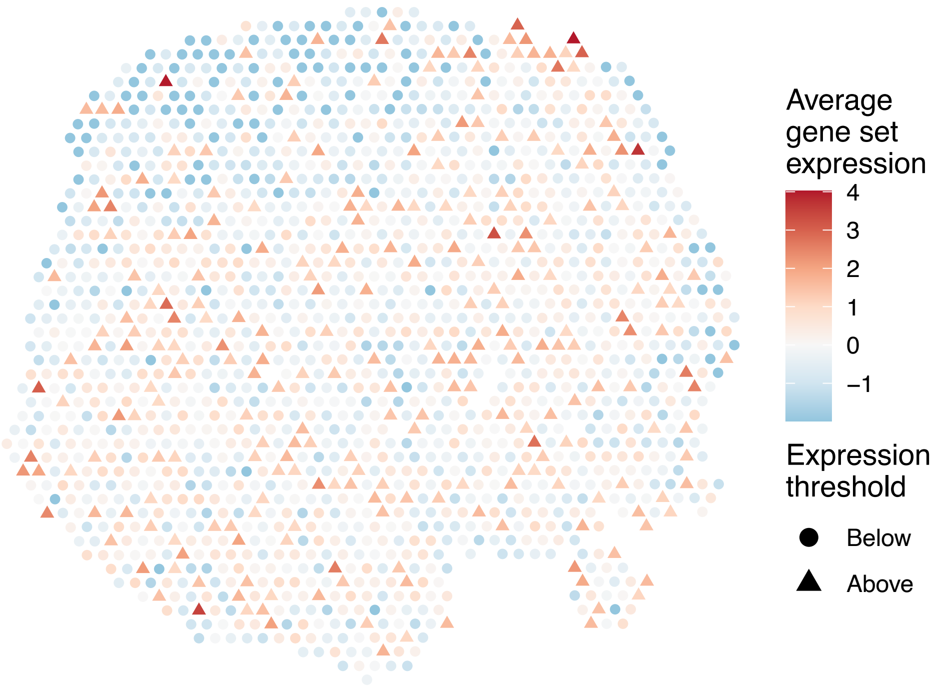


sample4_LMM_A5
KEGG_NITROGEN_METABOLISM
Proportion of pathway genes=0.54; Adjusted p−value=0.0075

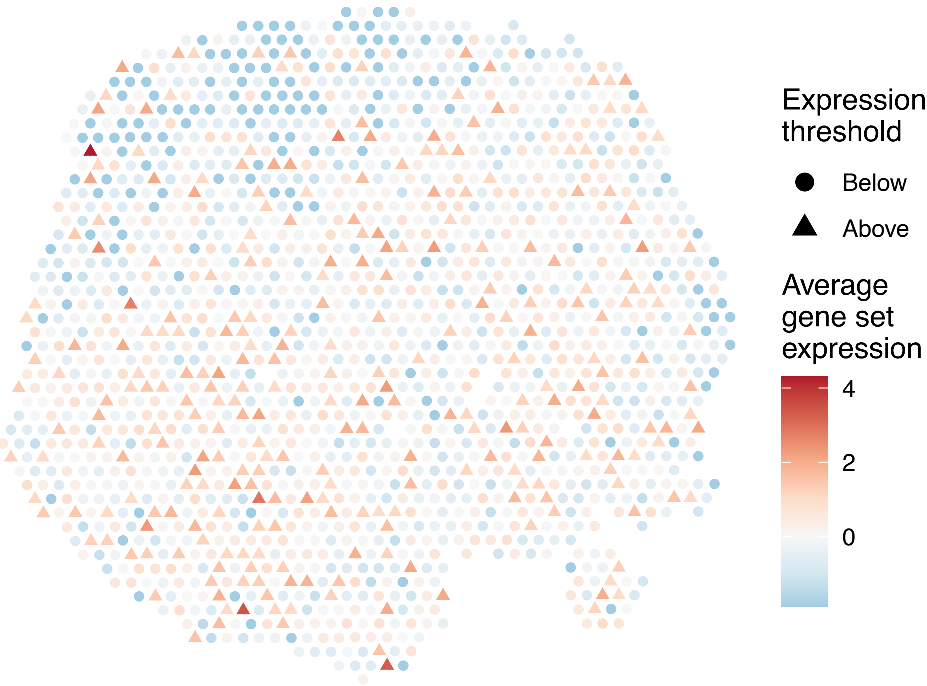


sample4_LMM_A5
KEGG_NOD_LIKE_RECEPTOR_SIGNALING_PATHWAY
Proportion of pathway genes=0.68; Adjusted p−value=0

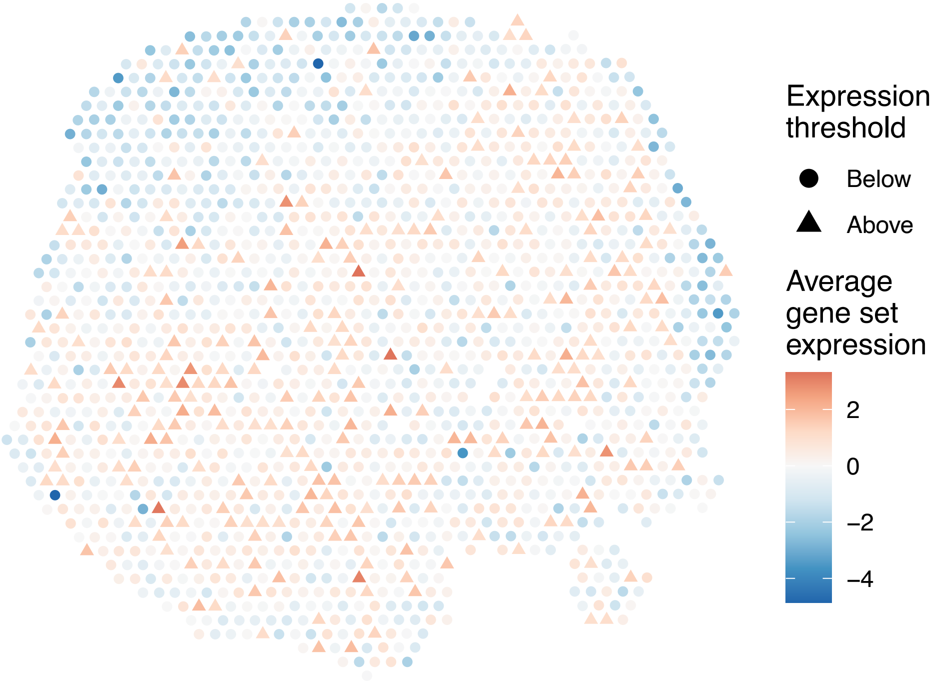


sample4_LMM_A5
KEGG_NON_HOMOLOGOUS_END_JOINING
Proportion of pathway genes=0.92; Adjusted p−value=0.0655

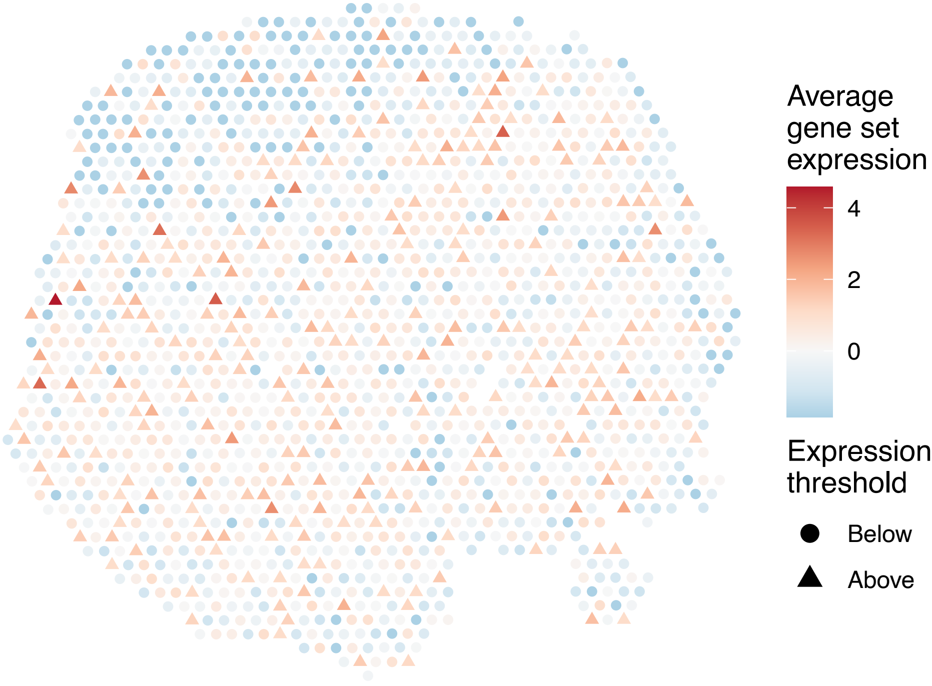

sample4_LMM_A5
KEGG_NON_SMALL_CELL_LUNG_CANCER
Proportion of pathway genes=0.92; Adjusted p−value=0

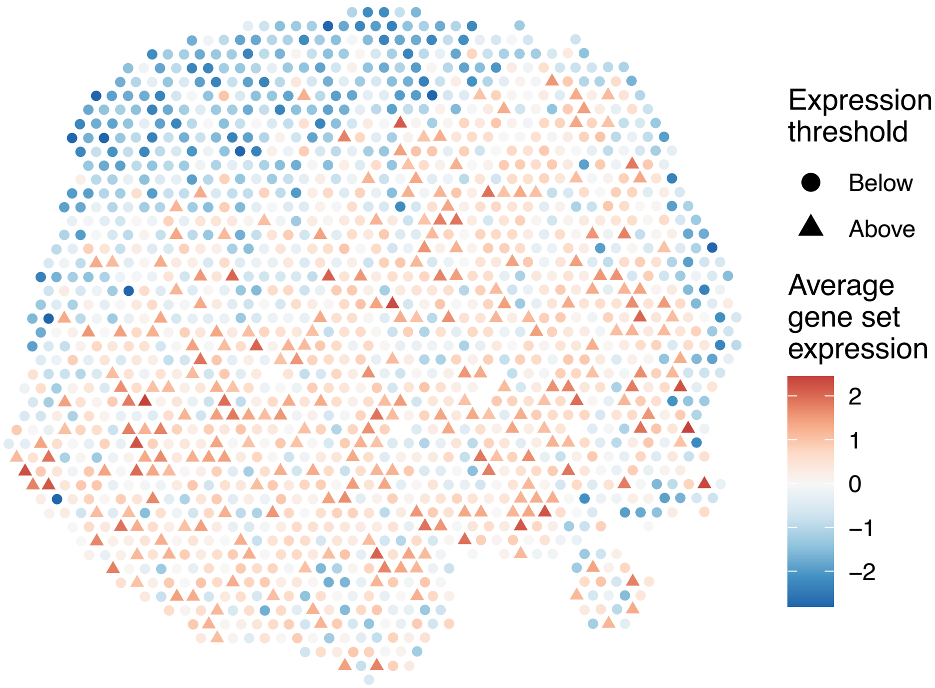


sample4_LMM_A5
KEGG_NOTCH_SIGNALING_PATHWAY
Proportion of pathway genes=0.93; Adjusted p−value=0

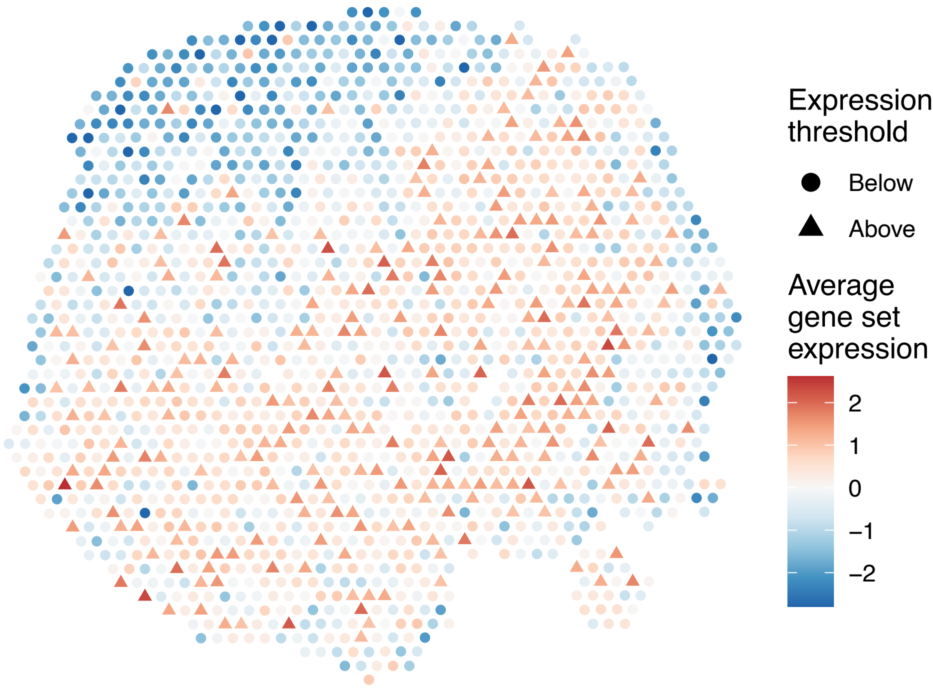


sample4_LMM_A5
KEGG_NUCLEOTIDE_EXCISION_REPAIR
Proportion of pathway genes=0.96; Adjusted p−value=0

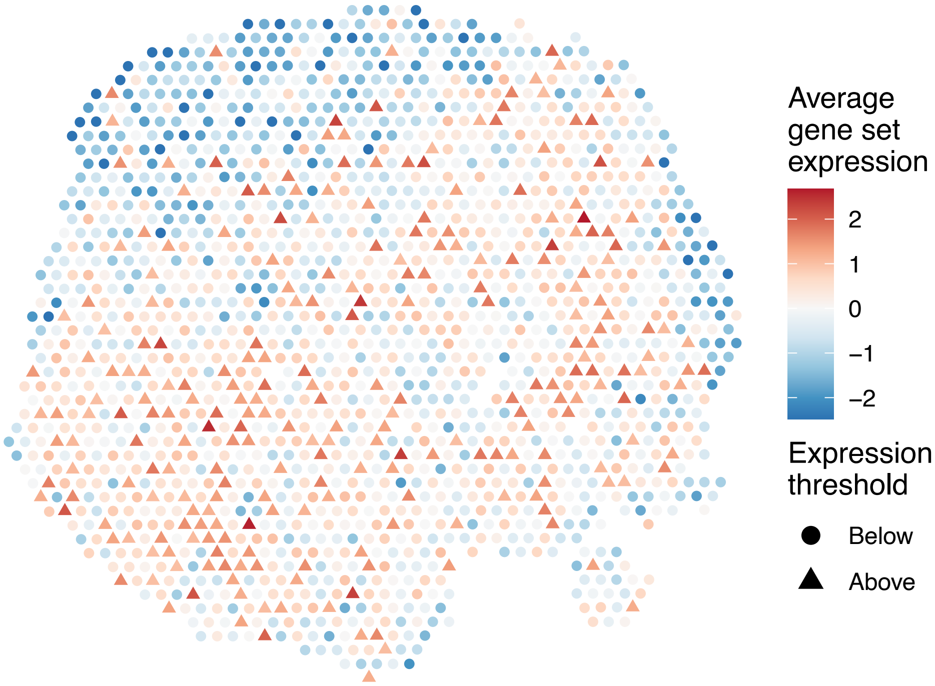


sample4_LMM_A5
KEGG_O_GLYCAN_BIOSYNTHESIS
Proportion of pathway genes=0.67; Adjusted p−value=0

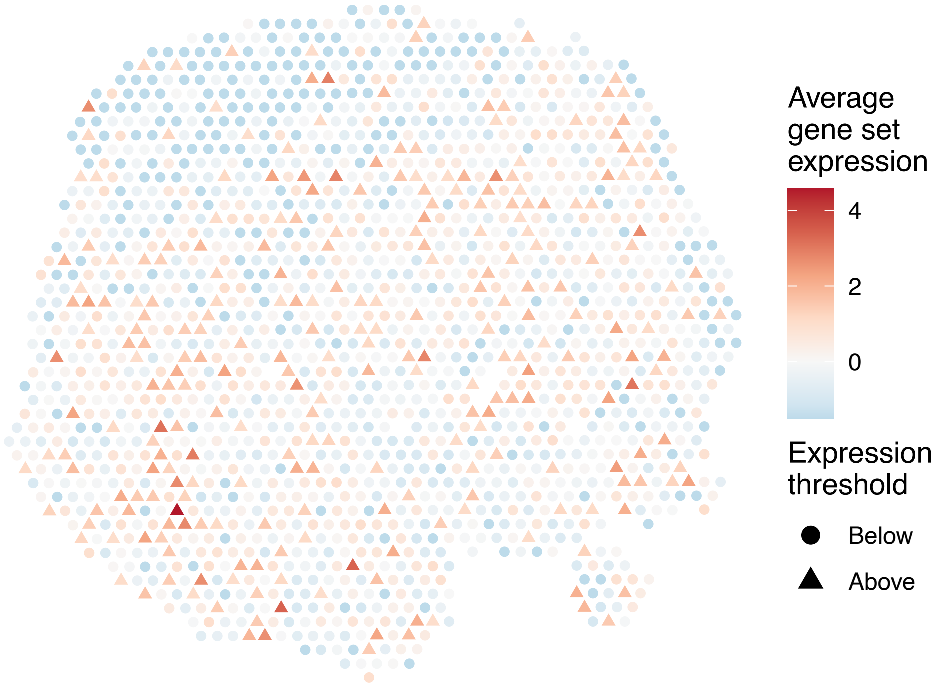

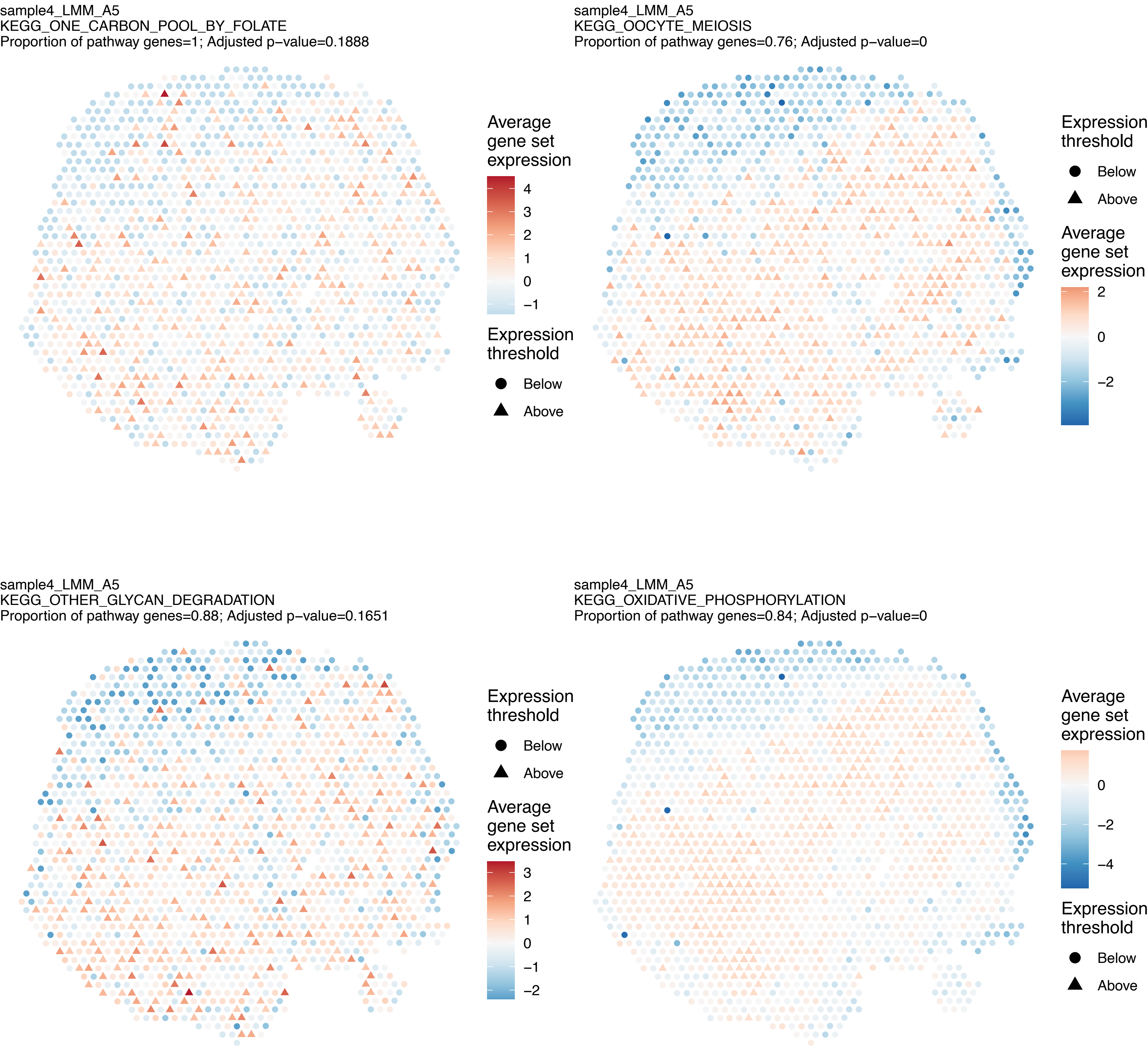

sample4_LMM_A5
KEGG_ONE_CARBON_POOL_BY_FOLATE
Proportion of pathway genes=1; Adjusted p−value=0.1888
Average gene set expression
4
3
2
1
0
−1
Expression threshold
Below
Above
sample4_LMM_A5
KEGG_OOCYTE_MEIOSIS
Proportion of pathway genes=0.76; Adjusted p−value=0
Expression threshold
Below
Above
Average gene set expression
2
0
−2
sample4_LMM_A5
KEGG_OTHER_GLYCAN_DEGRADATION
Proportion of pathway genes=0.88; Adjusted p−value=0.1651
Expression threshold
Below
Above
Average gene set expression
3
2
1
0
−1
−2
sample4_LMM_A5
KEGG_OXIDATIVE_PHOSPHORYLATION
Proportion of pathway genes=0.84; Adjusted p−value=0
Average gene set expression
0
−2
−4
Expression threshold
Below
Above

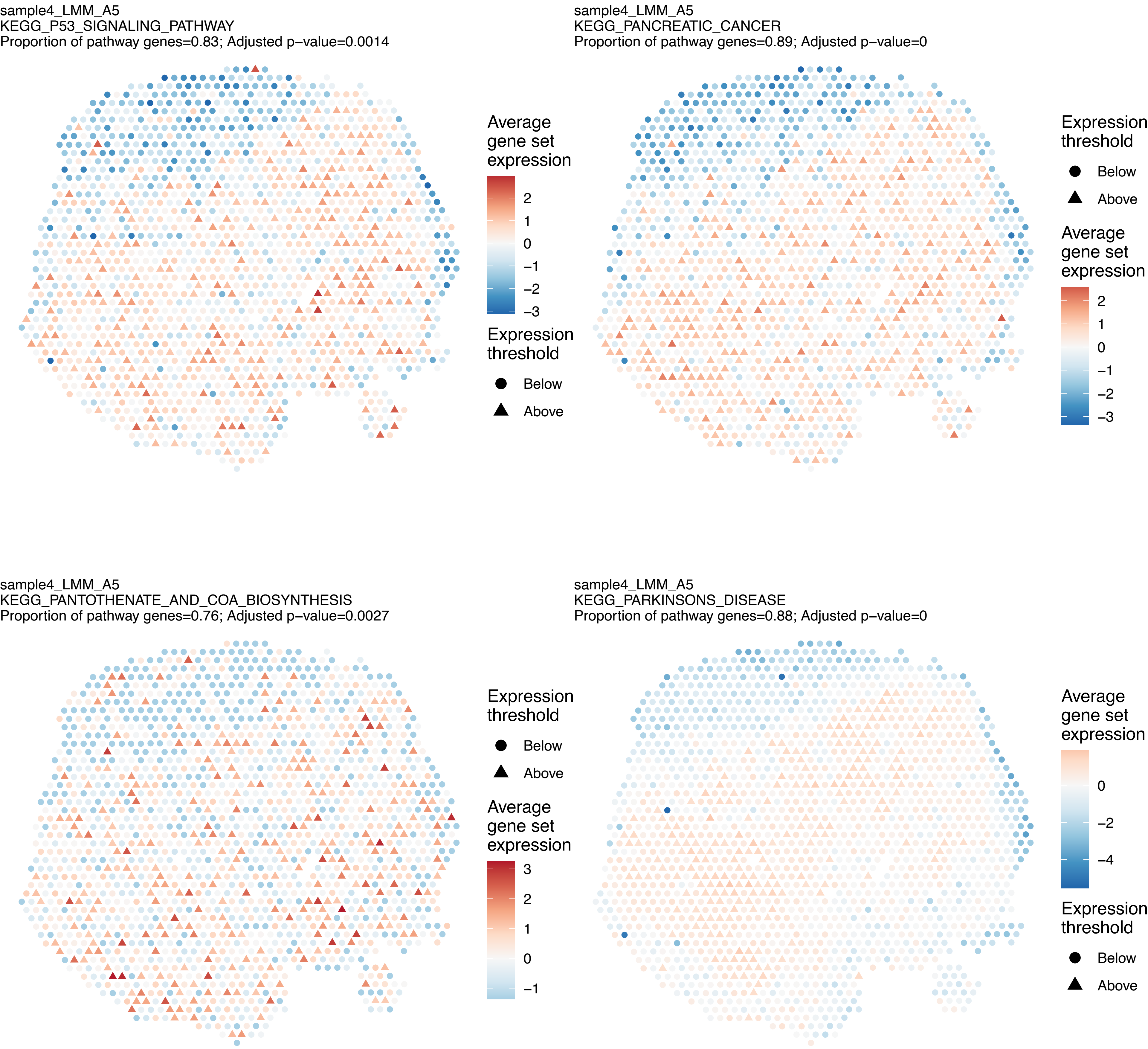
sample4_LMM_A5
KEGG_P53_SIGNALING_PATHWAY
Proportion of pathway genes=0.83; Adjusted p−value=0.0014
Average gene set expression
2
1
0
−1
−2
−3
Expression threshold
Below
Above
sample4_LMM_A5
KEGG_PANCREATIC_CANCER
Proportion of pathway genes=0.89; Adjusted p−value=0
Expression threshold
Below
Above
Average gene set expression
2
1
0
−1
−2
−3
sample4_LMM_A5
KEGG_PANTOTHENATE_AND_COA_BIOSYNTHESIS
Proportion of pathway genes=0.76; Adjusted p−value=0.0027
Expression threshold
Below
Above
Average gene set expression
3
2
1
0
−1
sample4_LMM_A5
KEGG_PARKINSONS_DISEASE
Proportion of pathway genes=0.88; Adjusted p−value=0
Average gene set expression
0
−2
−4
Expression threshold
Below
Above

sample4_LMM_A5
KEGG_PATHOGENIC_ESCHERICHIA_COLI_INFECTION
Proportion of pathway genes=0.86; Adjusted p−value=0

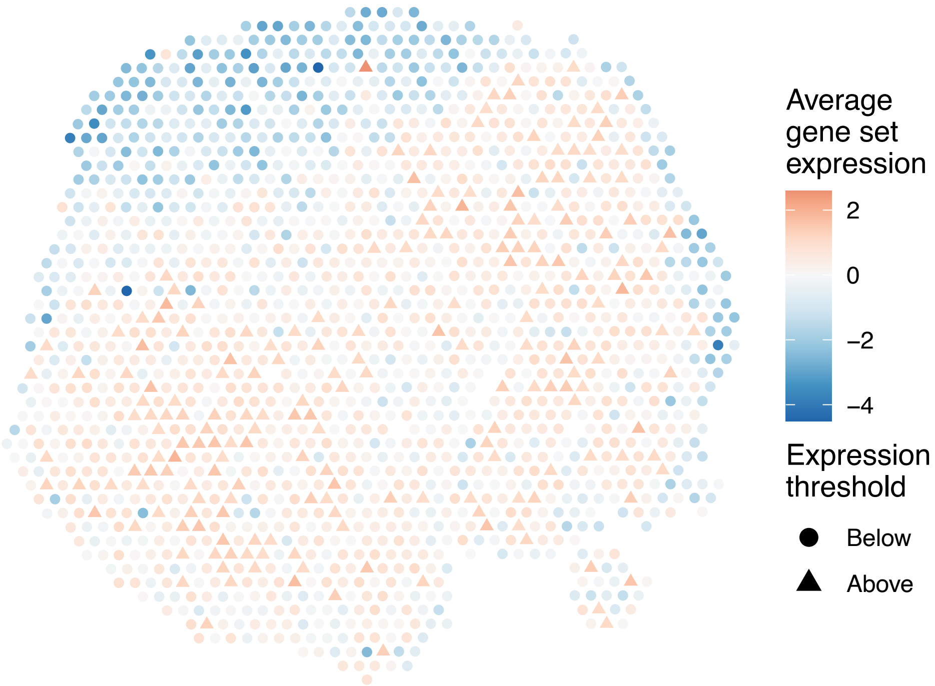


sample4_LMM_A5
KEGG_PATHWAYS_IN_CANCER
Proportion of pathway genes=0.77; Adjusted p−value=0

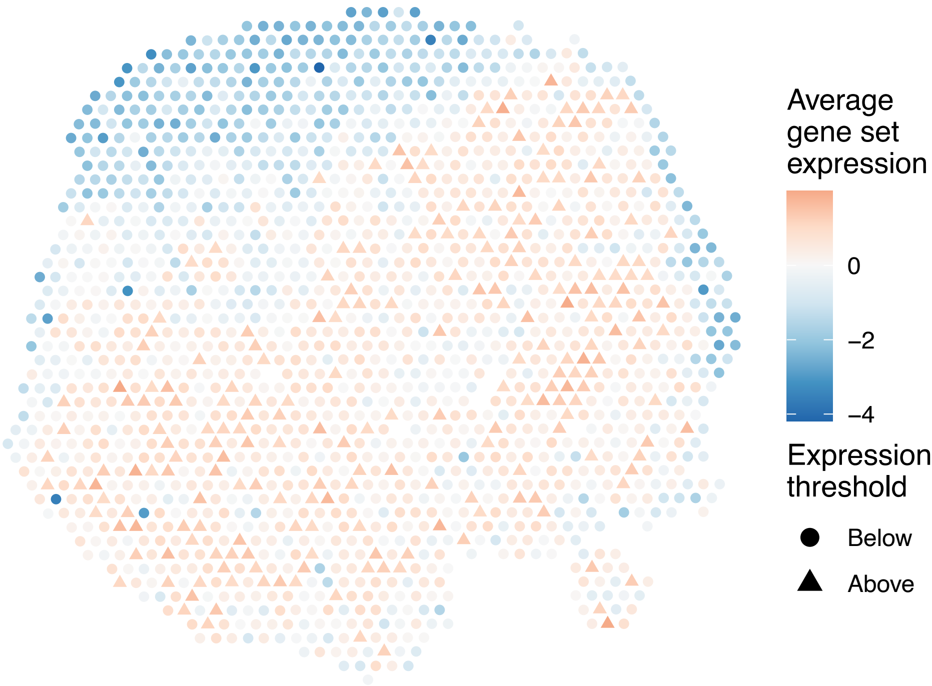


sample4_LMM_A5
KEGG_PENTOSE_PHOSPHATE_PATHWAY
Proportion of pathway genes=0.79; Adjusted p−value=0

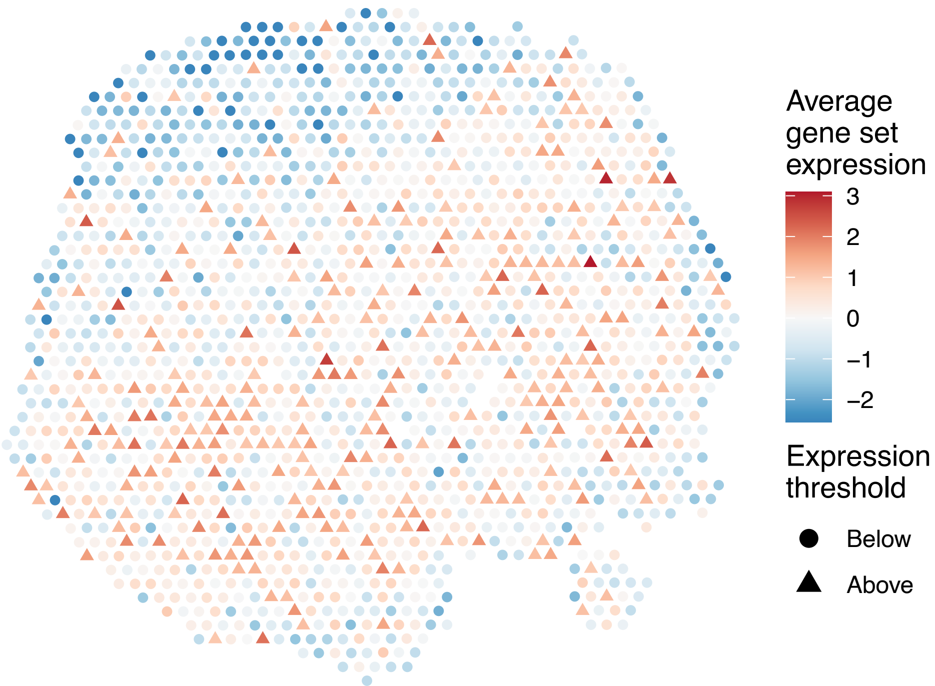


sample4_LMM_A5
KEGG_PEROXISOME
Proportion of pathway genes=0.85; Adjusted p−value=0

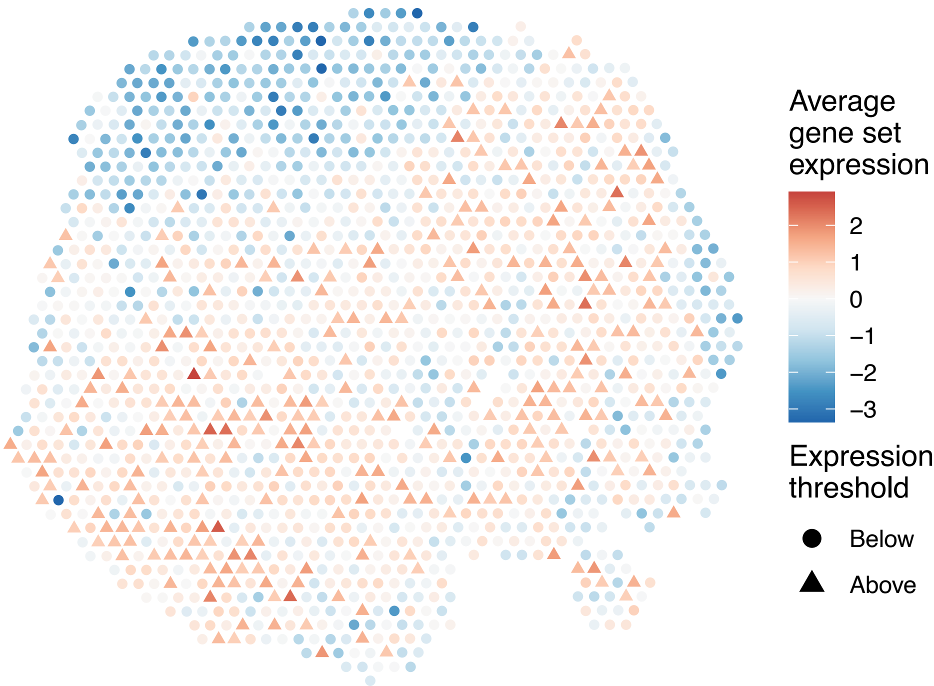

sample4_LMM_A5
KEGG_PHENYLALANINE_METABOLISM
Proportion of pathway genes=0.68; Adjusted p−value=0.2639

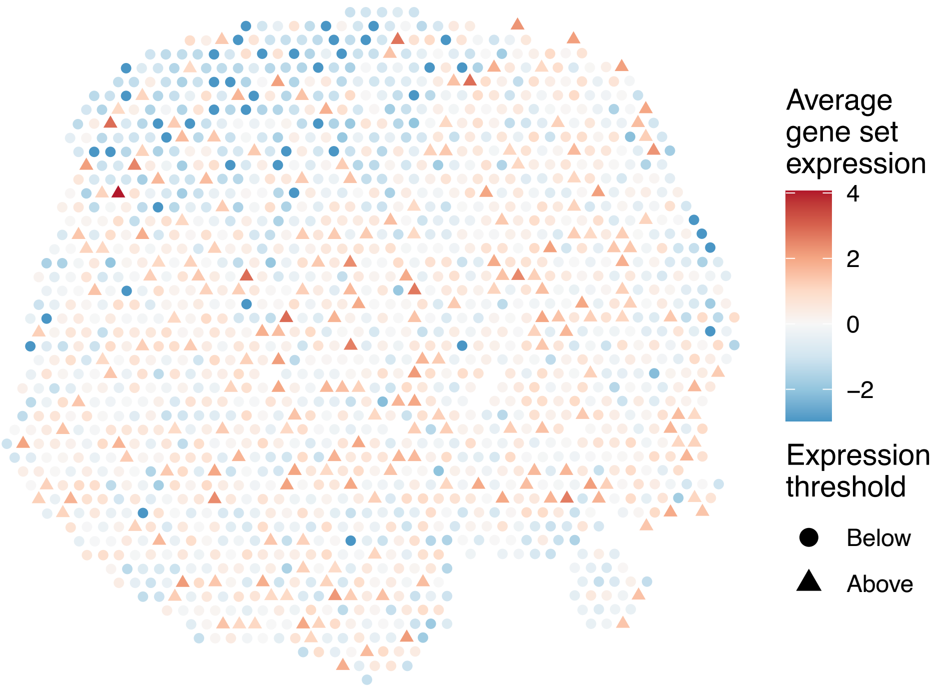


sample4_LMM_A5
KEGG_PHOSPHATIDYLINOSITOL_SIGNALING_SYSTEM
Proportion of pathway genes=0.89; Adjusted p−value=0

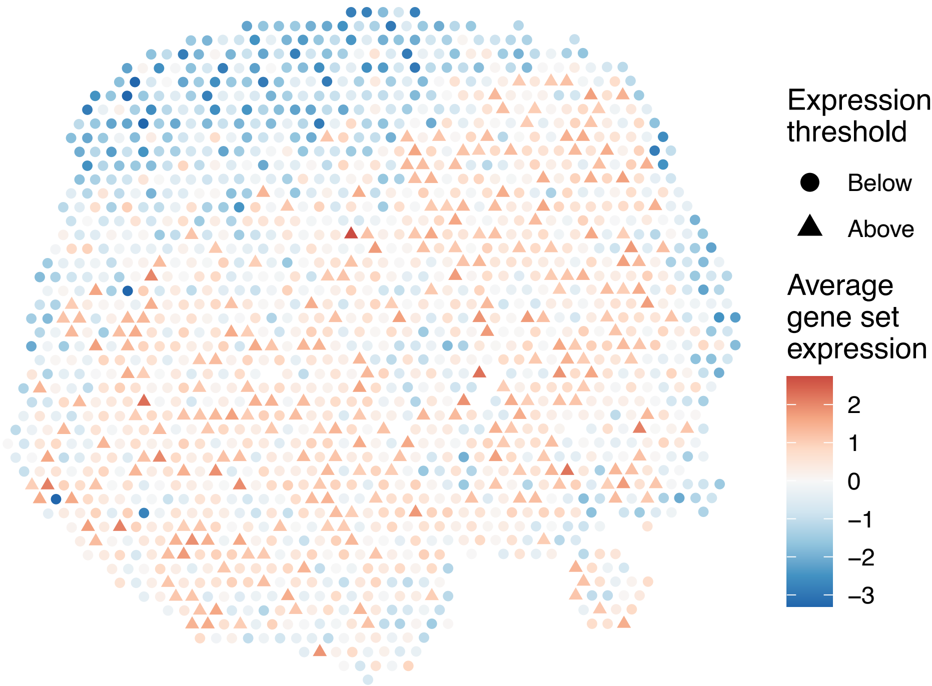


sample4_LMM_A5
KEGG_PORPHYRIN_AND_CHLOROPHYLL_METABOLISM
Proportion of pathway genes=0.44; Adjusted p−value=0

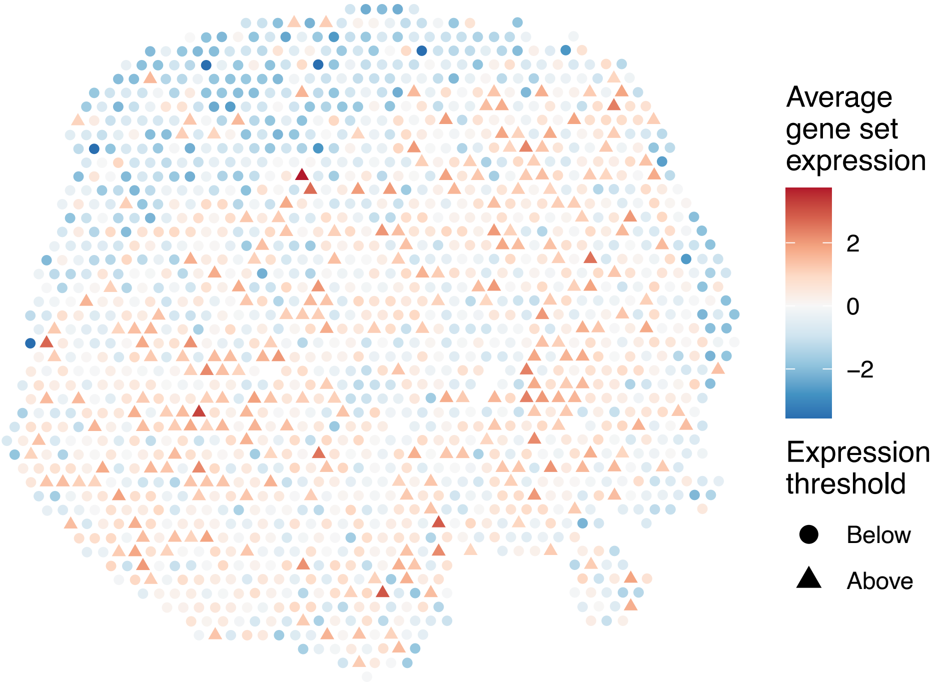


sample4_LMM_A5
KEGG_PPAR_SIGNALING_PATHWAY
Proportion of pathway genes=0.67; Adjusted p−value=0

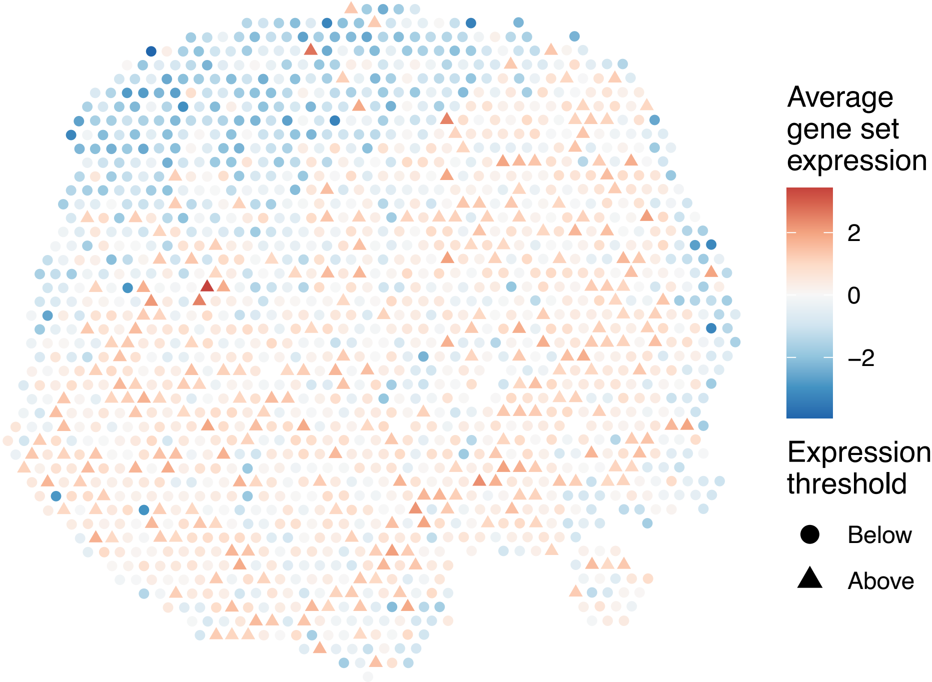

sample4_LMM_A5
KEGG_PRION_DISEASES
Proportion of pathway genes=0.83; Adjusted p−value=0.0692

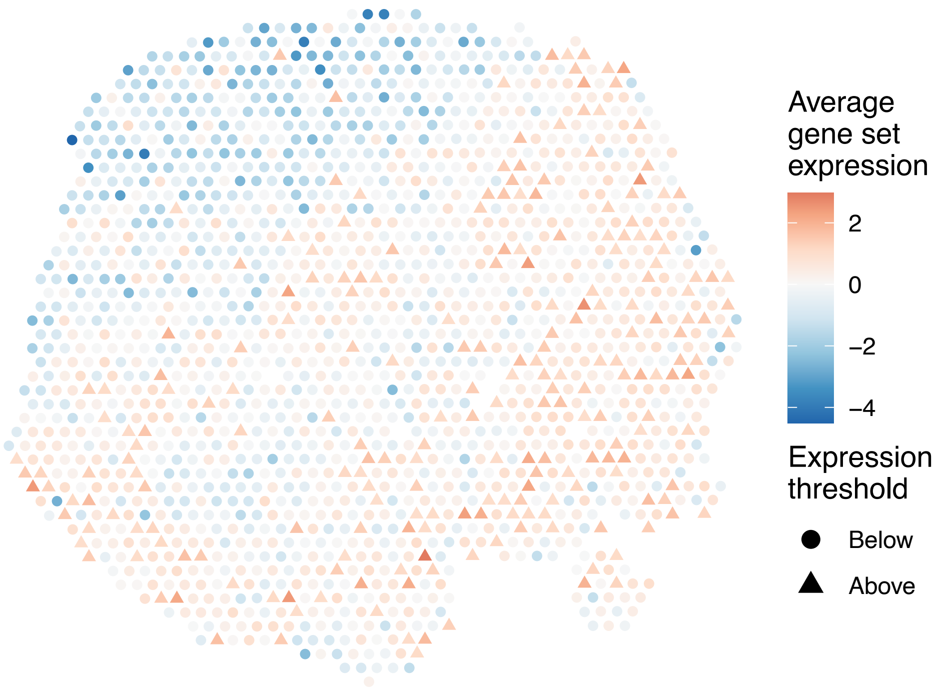


sample4_LMM_A5
KEGG_PROGESTERONE_MEDIATED_OOCYTE_MATURATION
Proportion of pathway genes=0.81; Adjusted p−value=0

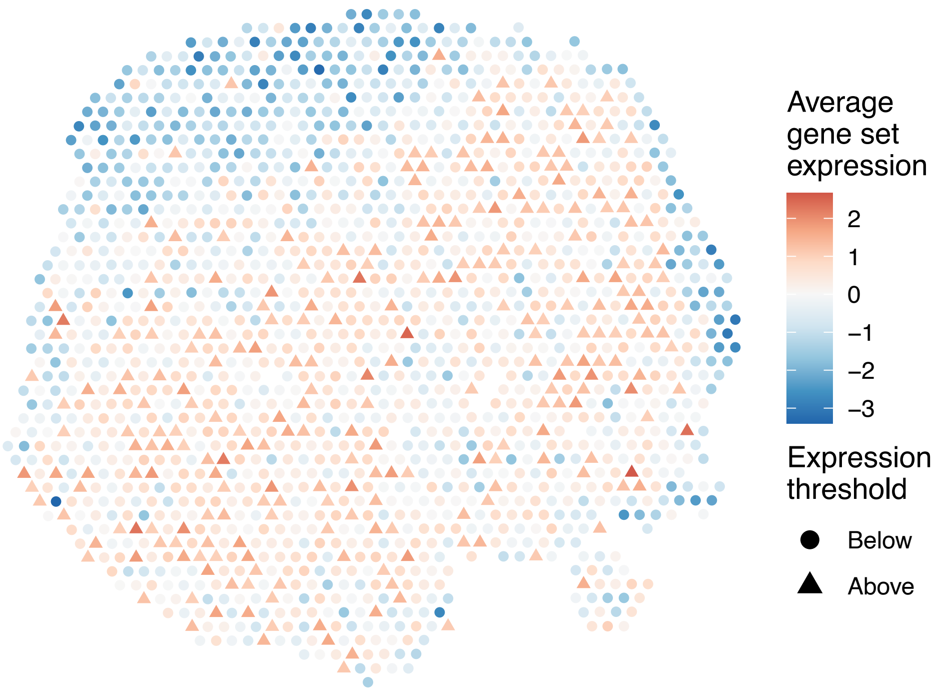


sample4_LMM_A5
KEGG_PROPANOATE_METABOLISM
Proportion of pathway genes=0.89; Adjusted p−value=0

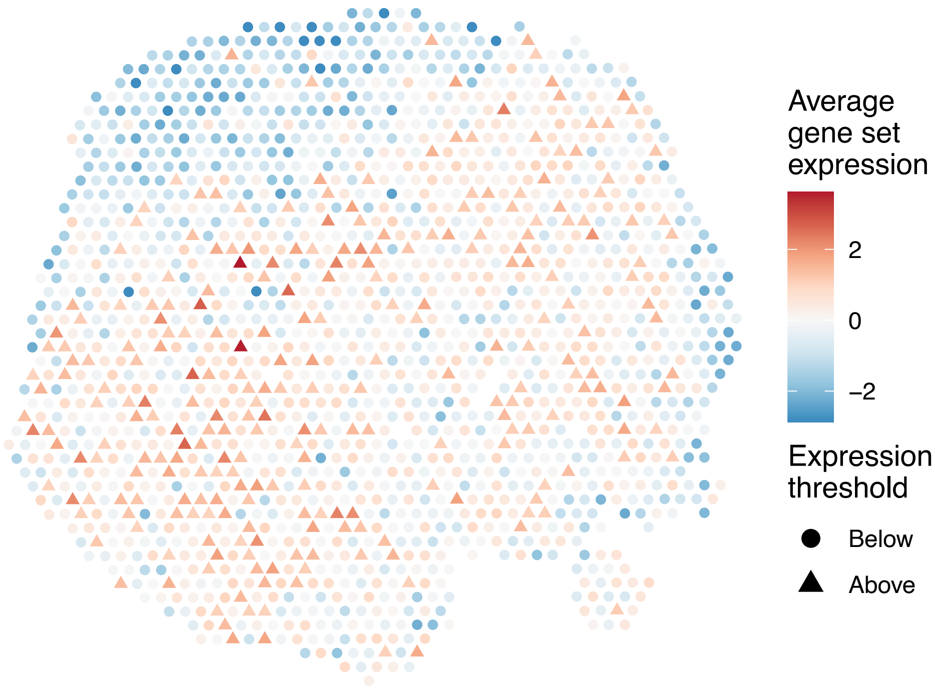


sample4_LMM_A5
KEGG_PROSTATE_CANCER
Proportion of pathway genes=0.88; Adjusted p−value=0

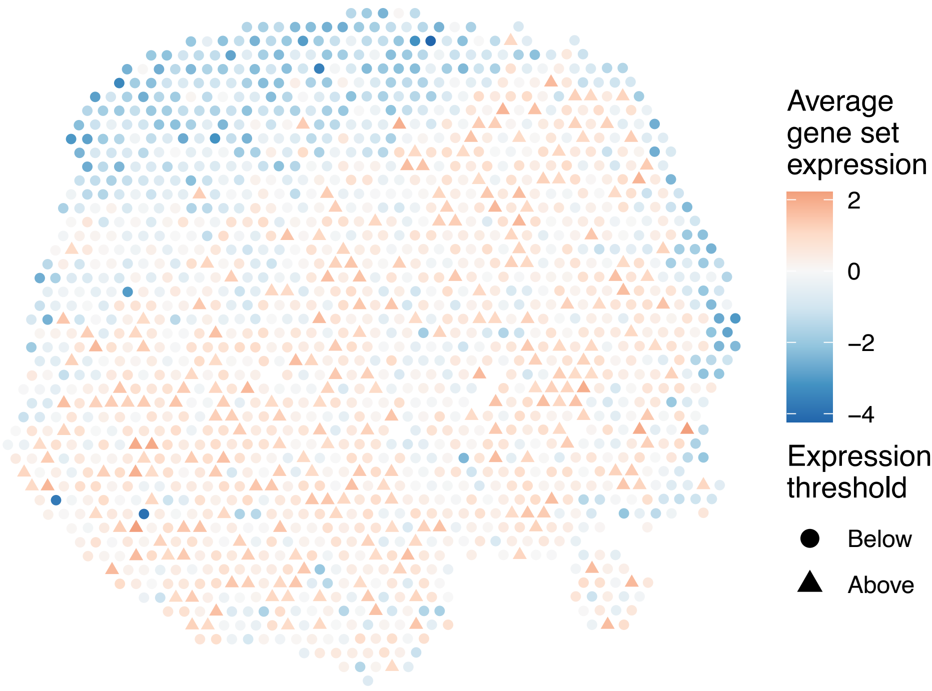

sample4_LMM_A5
KEGG_PROTEASOME
Proportion of pathway genes=0.92; Adjusted p−value=0.0052

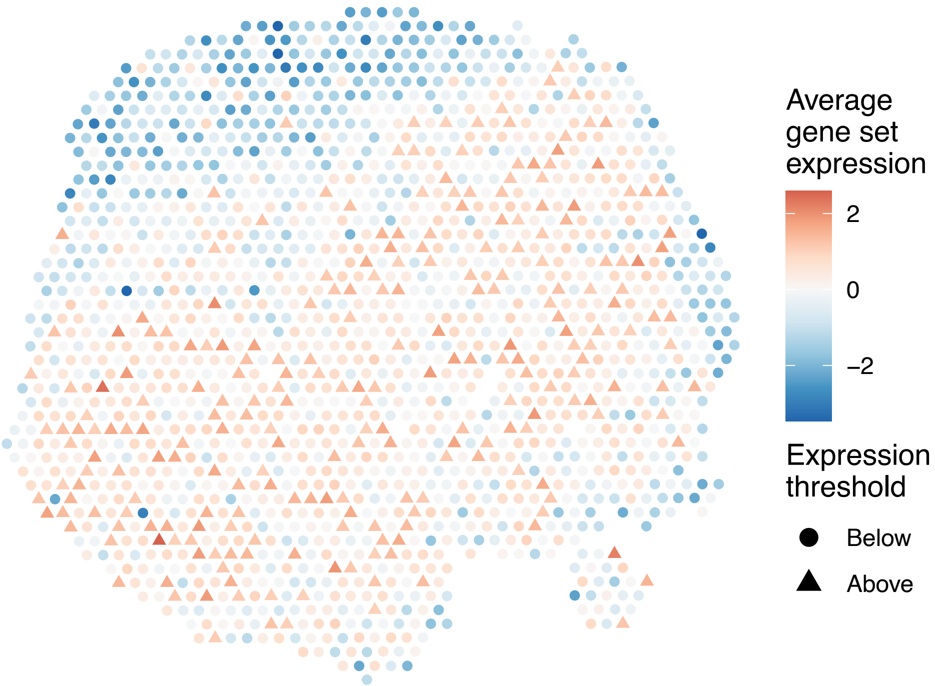


sample4_LMM_A5
KEGG_PROTEIN_EXPORT
Proportion of pathway genes=0.96; Adjusted p−value=0

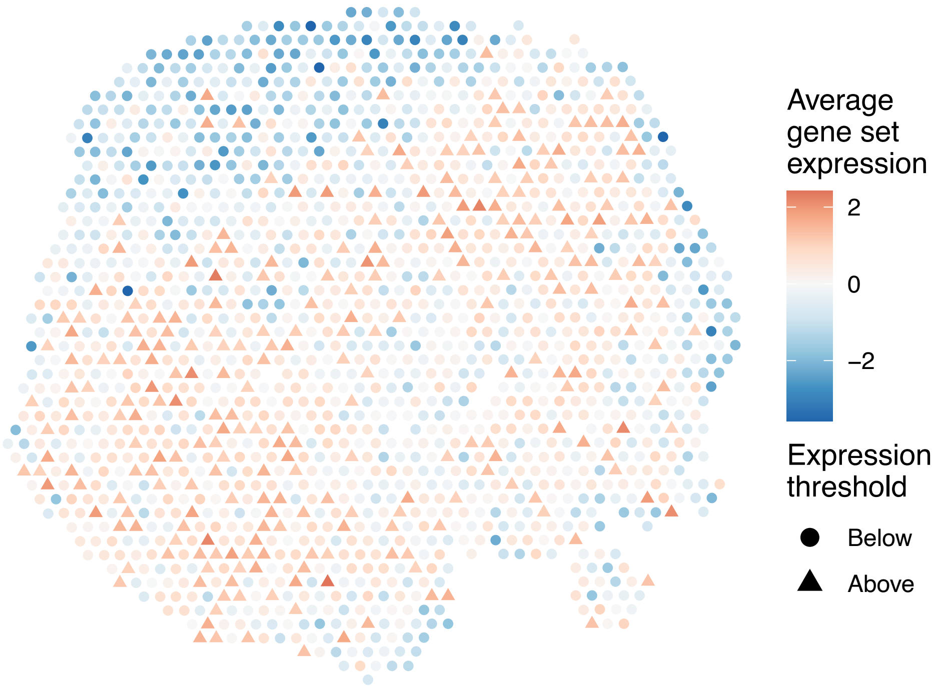


sample4_LMM_A5
KEGG_PROXIMAL_TUBULE_BICARBONATE_RECLAMATION
Proportion of pathway genes=0.65; Adjusted p−value=0

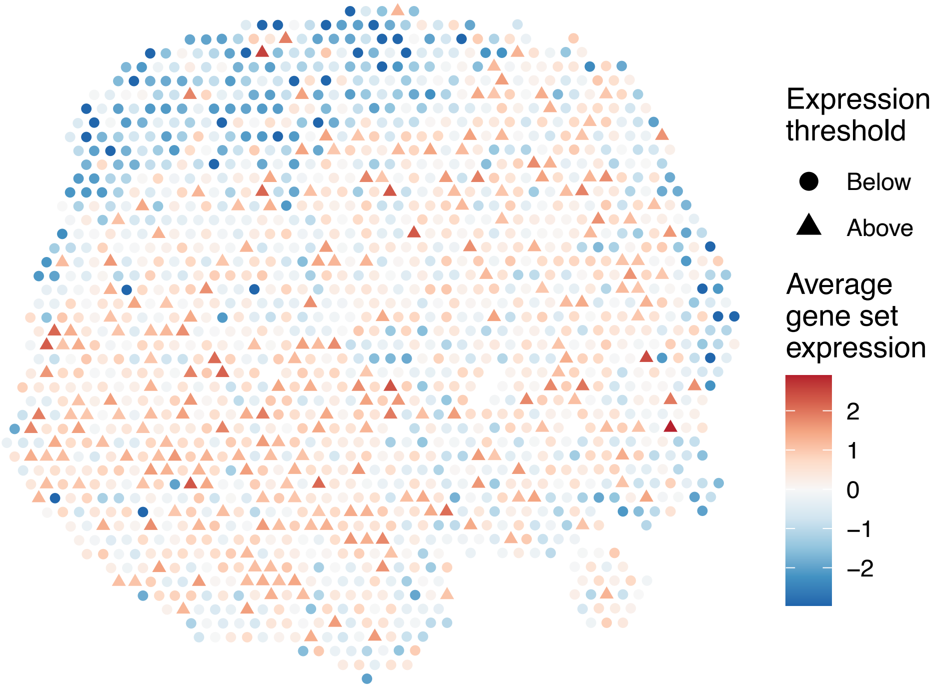


sample4_LMM_A5
KEGG_PURINE_METABOLISM
Proportion of pathway genes=0.75; Adjusted p−value=0

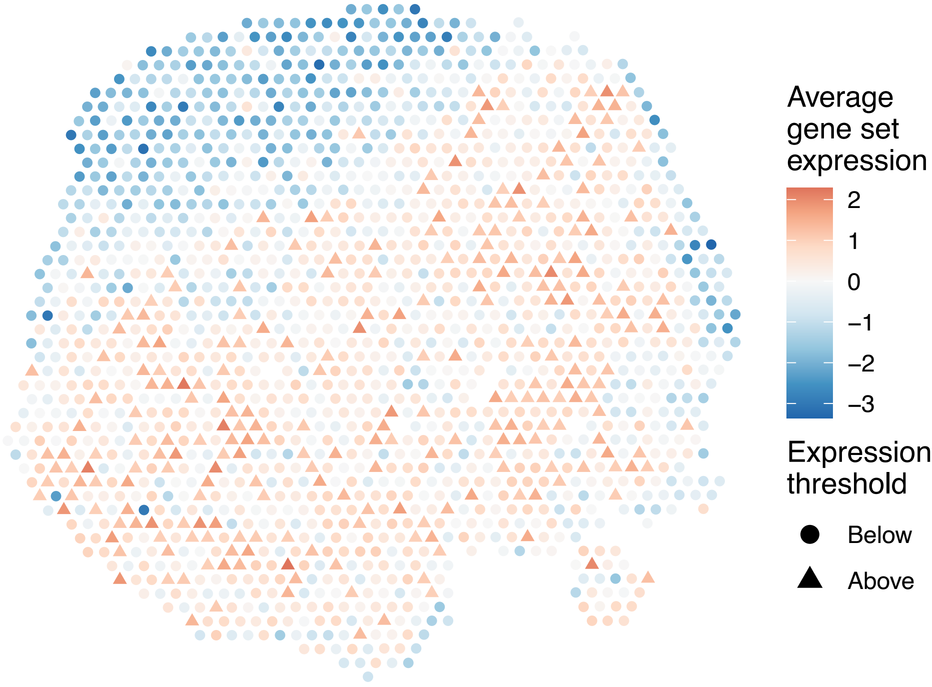

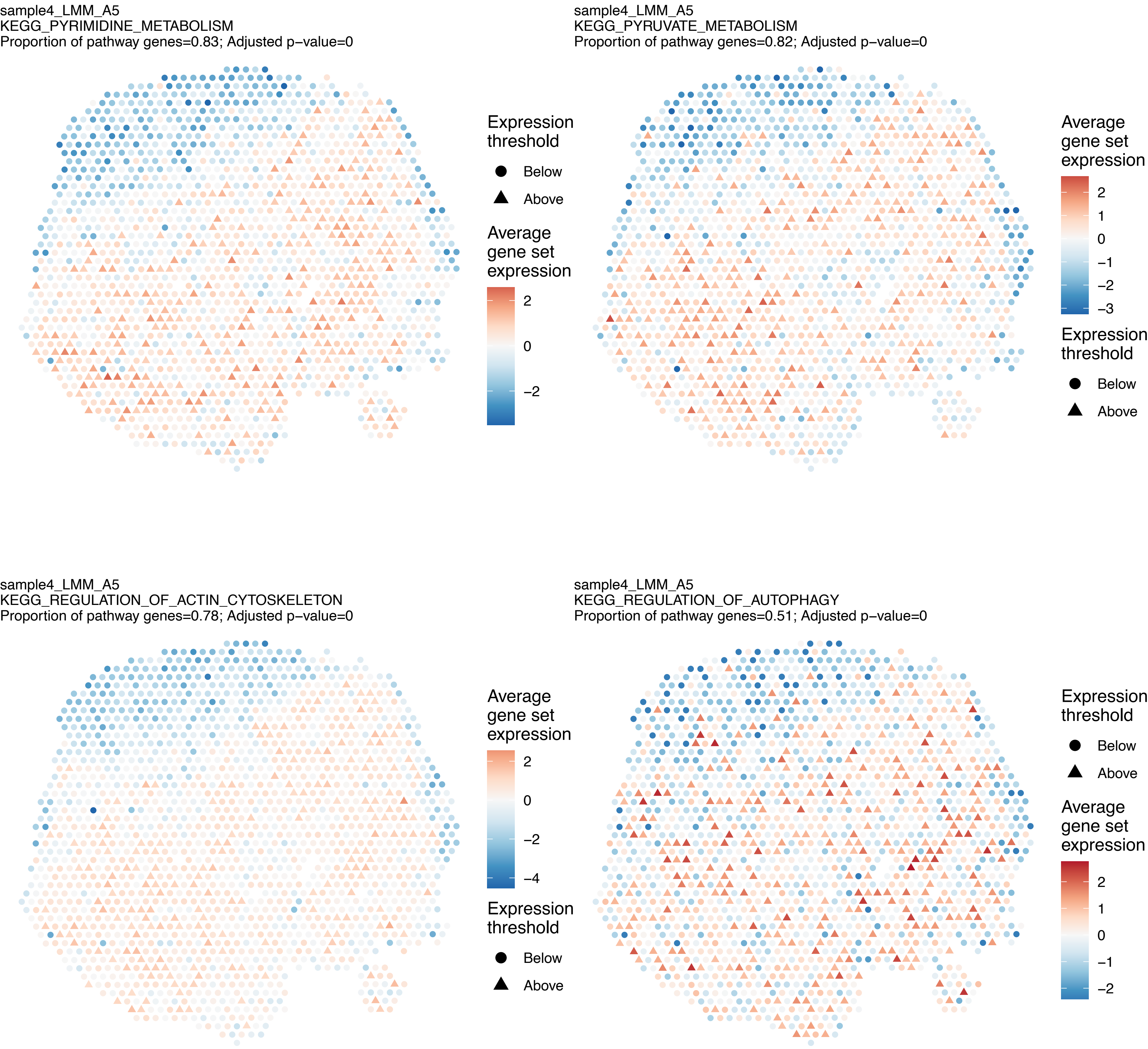

sample4_LMM_A5
KEGG_PYRIMIDINE_METABOLISM
Proportion of pathway genes=0.83; Adjusted p−value=0
Expression threshold
Below
Above
Average gene set expression
2
0
−2
sample4_LMM_A5
KEGG_PYRUVATE_METABOLISM
Proportion of pathway genes=0.82; Adjusted p−value=0
Average gene set expression
2
1
0
−1
−2
−3
Expression threshold
Below
Above
sample4_LMM_A5
KEGG_REGULATION_OF_ACTIN_CYTOSKELETON
Proportion of pathway genes=0.78; Adjusted p−value=0
Average gene set expression
2
0
−2
−4
Expression threshold
Below
Above
sample4_LMM_A5
KEGG_REGULATION_OF_AUTOPHAGY
Proportion of pathway genes=0.51; Adjusted p−value=0
Expression threshold
Below
Above
Average gene set expression
2
1
0
−1
−2

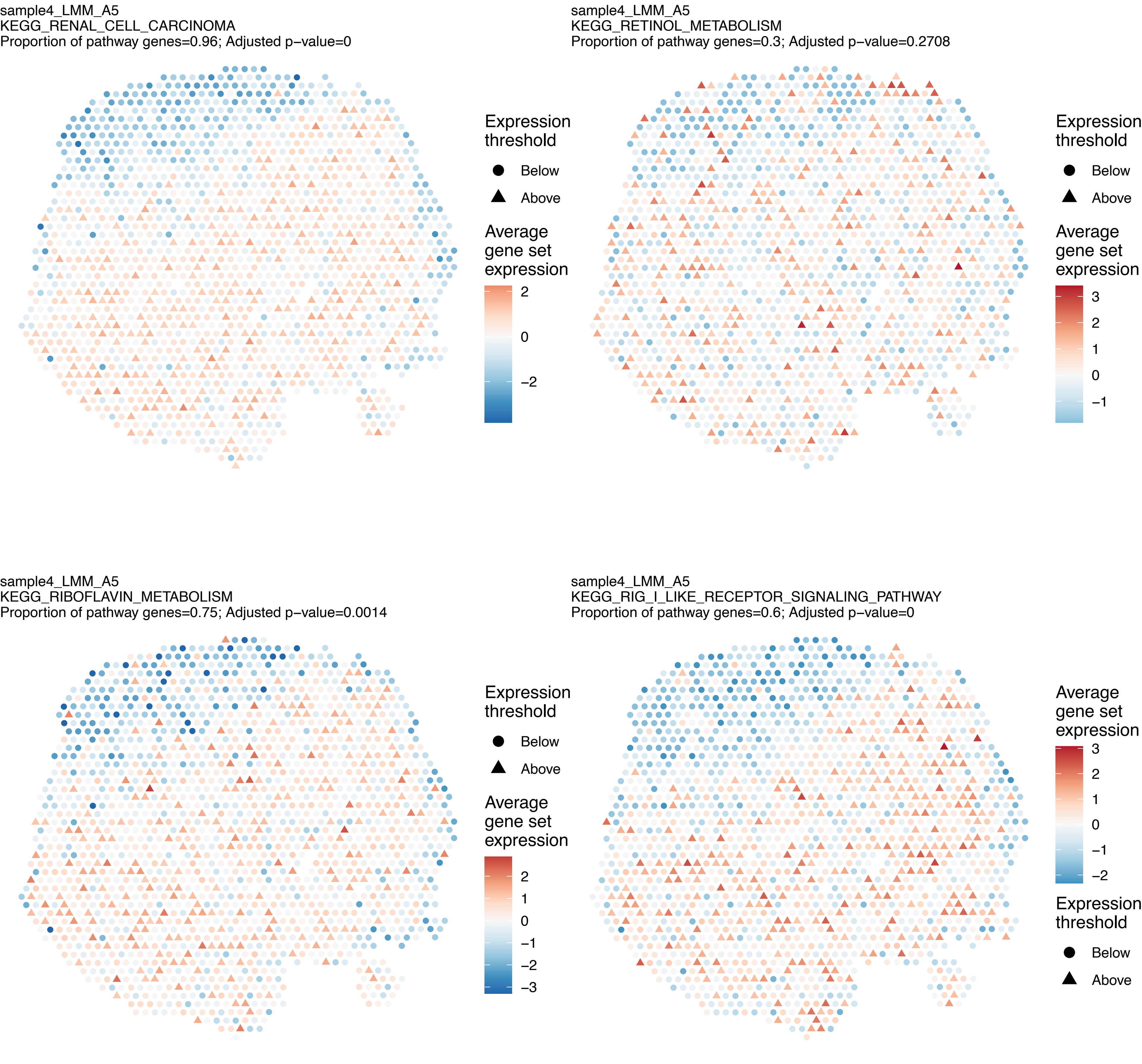
sample4_LMM_A5
KEGG_RENAL_CELL_CARCINOMA
Proportion of pathway genes=0.96; Adjusted p−value=0
Expression threshold
Below
Above
Average gene set expression
2
0
−2
sample4_LMM_A5
KEGG_RETINOL_METABOLISM
Proportion of pathway genes=0.3; Adjusted p−value=0.2708
Expression threshold
Below
Above
Average gene set expression
3
2
1
0
−1
sample4_LMM_A5
KEGG_RIBOFLAVIN_METABOLISM
Proportion of pathway genes=0.75; Adjusted p−value=0.0014
Expression threshold
Below
Above
Average gene set expression
2
1
0
−1
−2
−3
sample4_LMM_A5
KEGG_RIG_I_LIKE_RECEPTOR_SIGNALING_PATHWAY
Proportion of pathway genes=0.6; Adjusted p−value=0
Average gene set expression
3
2
1
0
−1
−2
Expression threshold
Below
Above

sample4_LMM_A5
KEGG_RNA_DEGRADATION
Proportion of pathway genes=0.96; Adjusted p−value=0

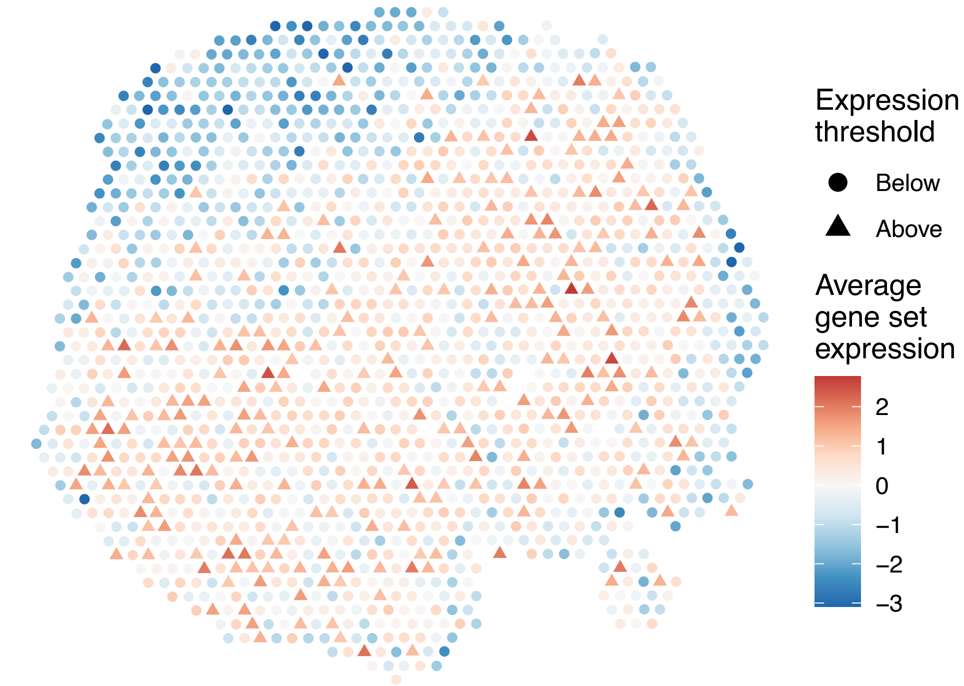


sample4_LMM_A5
KEGG_RNA_POLYMERASE
Proportion of pathway genes=0.77; Adjusted p−value=0

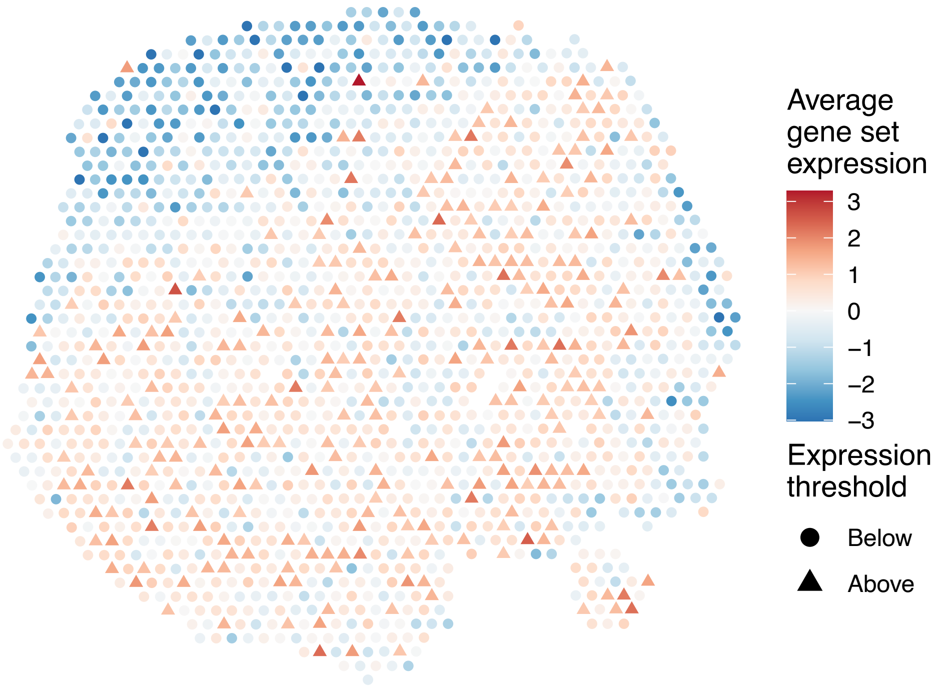


sample4_LMM_A5
KEGG_SELENOAMINO_ACID_METABOLISM
Proportion of pathway genes=0.81; Adjusted p−value=0.0014

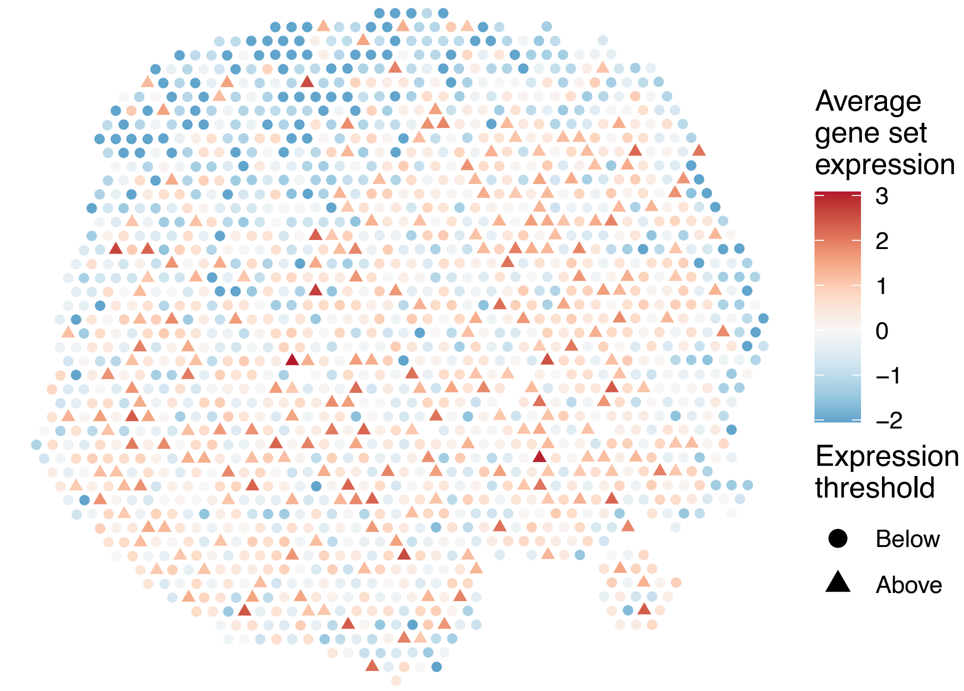


sample4_LMM_A5
KEGG_SMALL_CELL_LUNG_CANCER
Proportion of pathway genes=0.88; Adjusted p−value=0

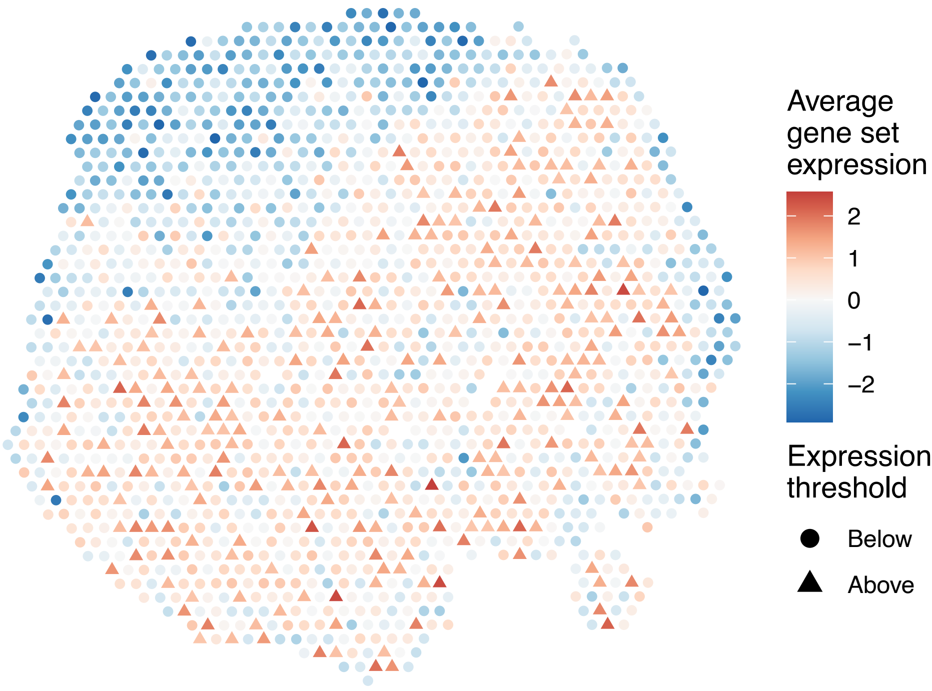

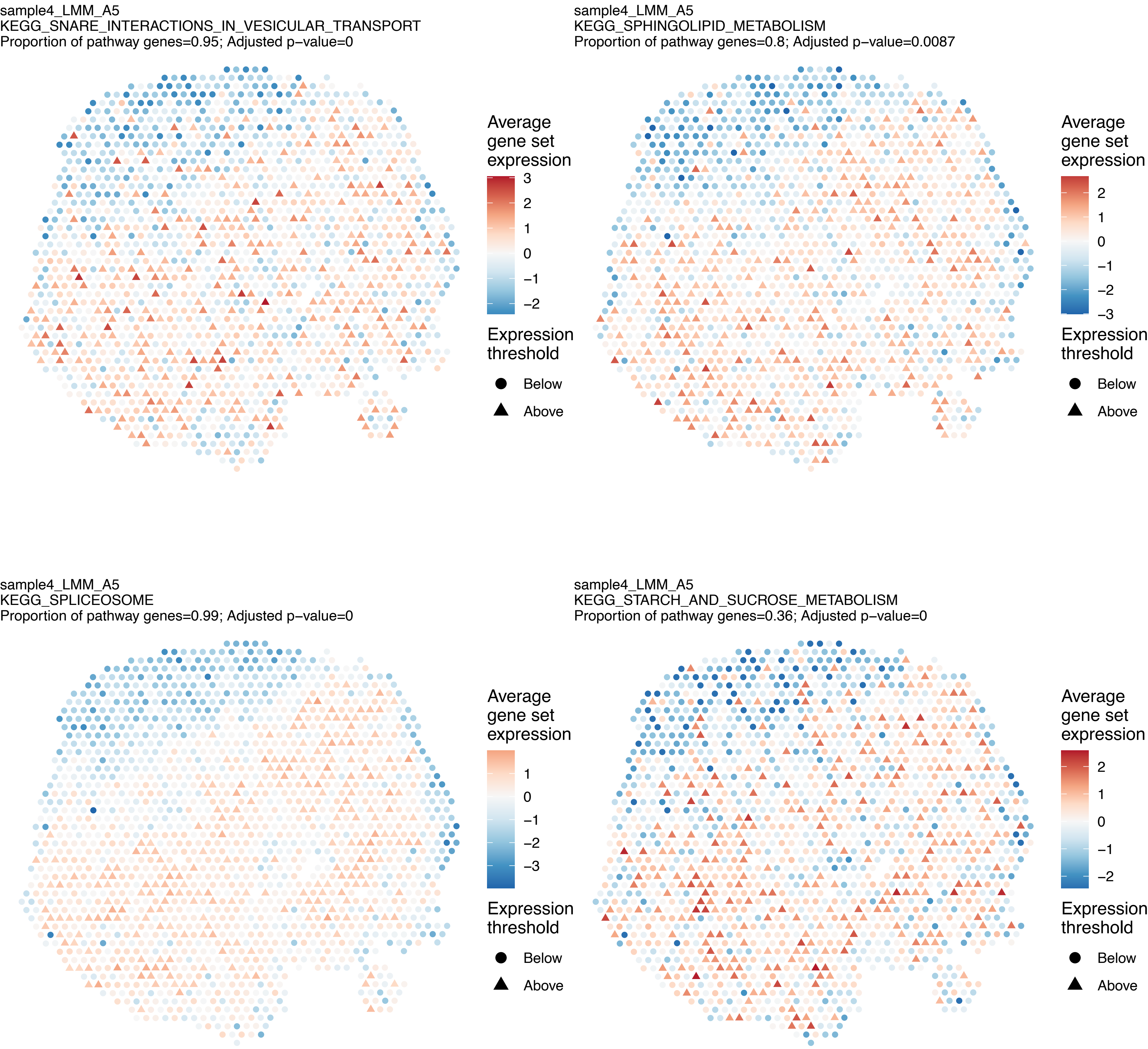
sample4_LMM_A5
KEGG_SNARE_INTERACTIONS_IN_VESICULAR_TRANSPORT
Proportion of pathway genes=0.95; Adjusted p−value=0
Average gene set expression
3
2
1
0
−1
−2
Expression threshold
Below
Above
sample4_LMM_A5
KEGG_SPHINGOLIPID_METABOLISM
Proportion of pathway genes=0.8; Adjusted p−value=0.0087
Average gene set expression
2
1
0
−1
−2
−3
Expression threshold
Below
Above
sample4_LMM_A5
KEGG_SPLICEOSOME
Proportion of pathway genes=0.99; Adjusted p−value=0
Average gene set expression
1
0
−1
−2
−3
Expression threshold
Below
Above
sample4_LMM_A5
KEGG_STARCH_AND_SUCROSE_METABOLISM
Proportion of pathway genes=0.36; Adjusted p−value=0
Average gene set expression
2
1
0
−1
−2
Expression threshold
Below
Above

sample4_LMM_A5
KEGG_STEROID_BIOSYNTHESIS
Proportion of pathway genes=0.89; Adjusted p−value=0.0027

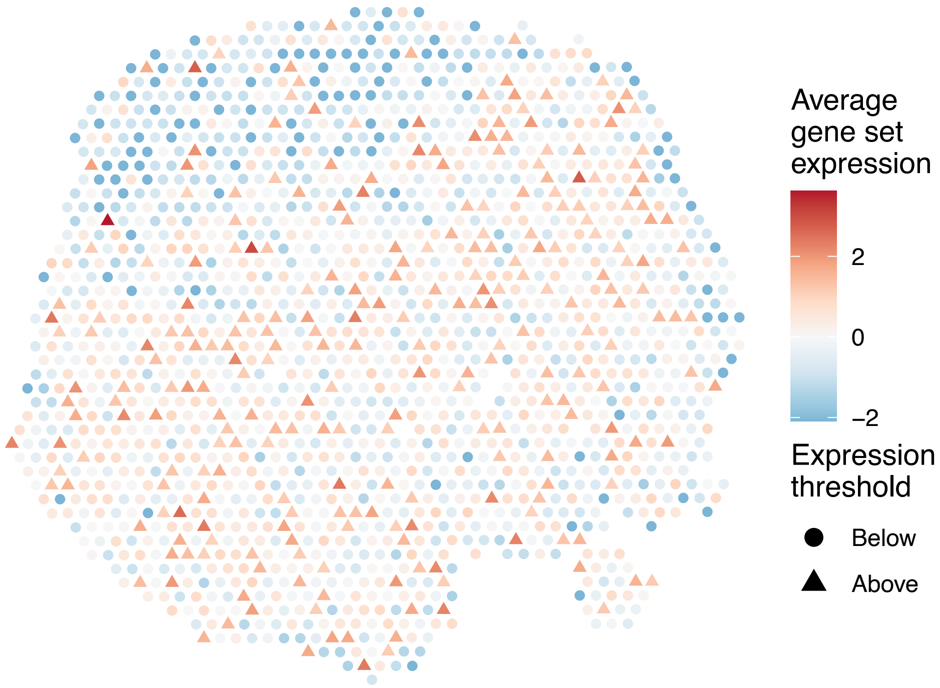


sample4_LMM_A5
KEGG_STEROID_HORMONE_BIOSYNTHESIS
Proportion of pathway genes=0.25; Adjusted p−value=0.1297

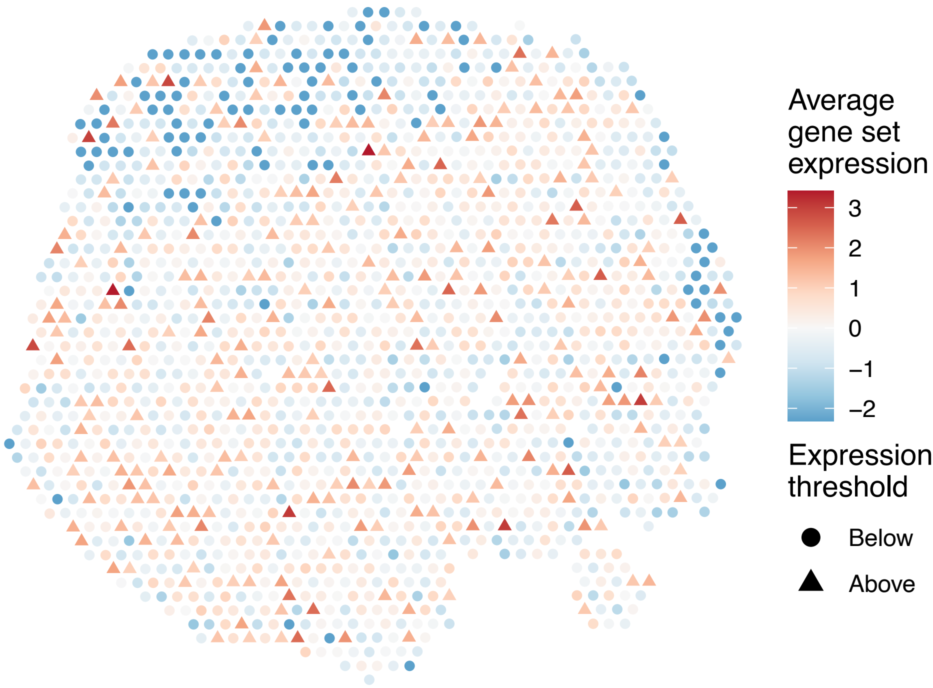


sample4_LMM_A5
KEGG_SULFUR_METABOLISM
Proportion of pathway genes=0.85; Adjusted p−value=0.0087

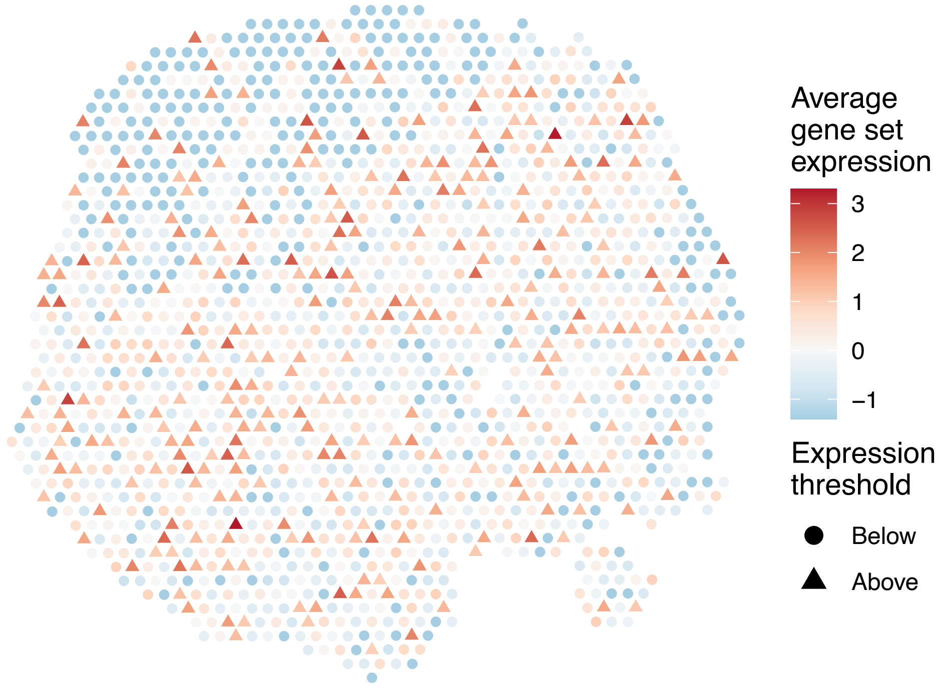


sample4_LMM_A5
KEGG_SYSTEMIC_LUPUS_ERYTHEMATOSUS
Proportion of pathway genes=0.42; Adjusted p−value=0.9873

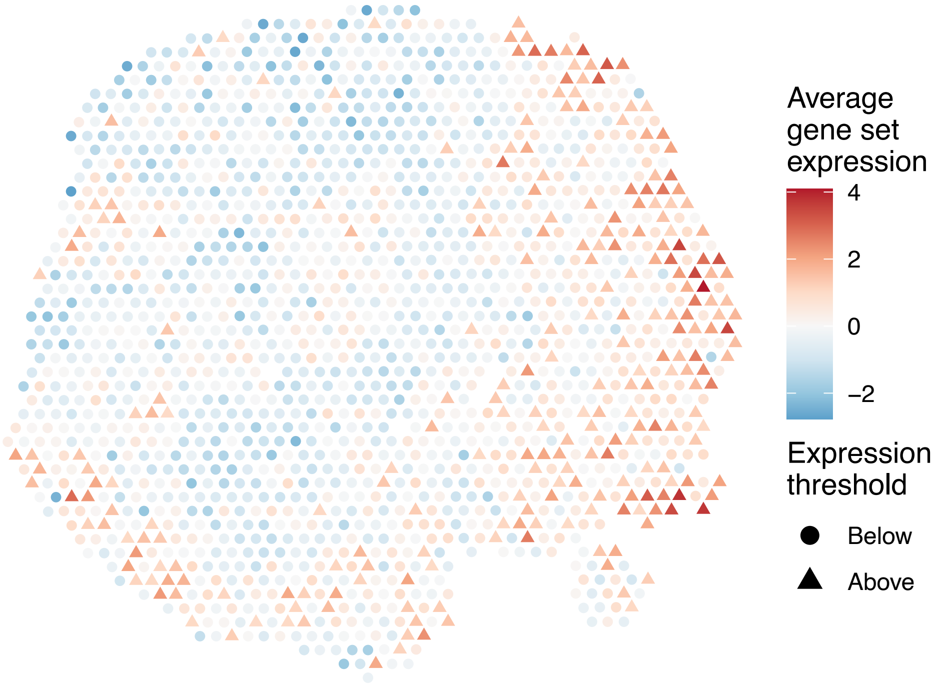

sample4_LMM_A5
KEGG_T_CELL_RECEPTOR_SIGNALING_PATHWAY
Proportion of pathway genes=0.76; Adjusted p−value=0

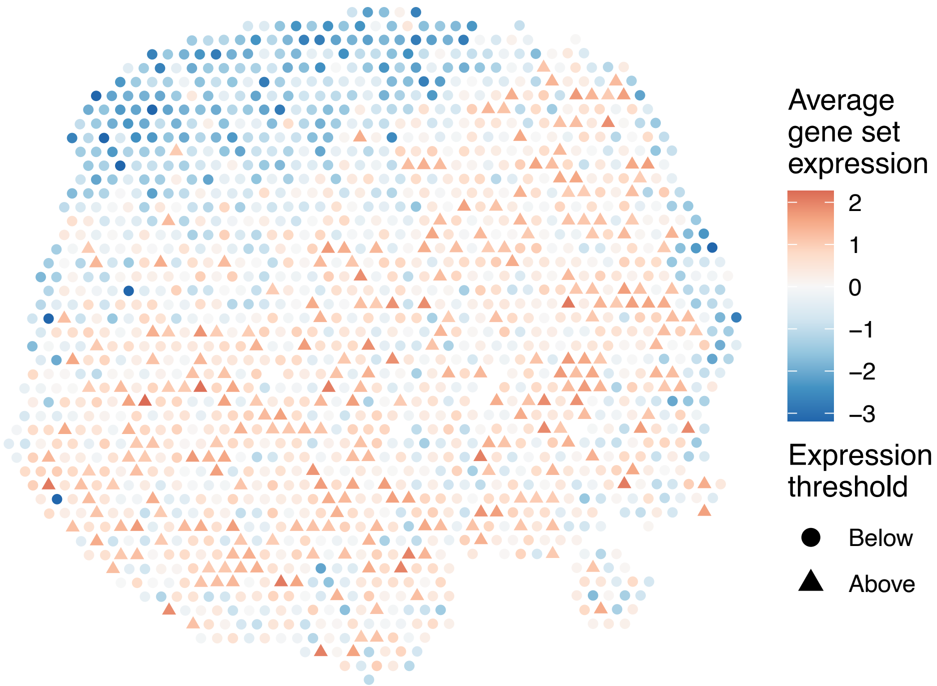


sample4_LMM_A5
KEGG_TERPENOID_BACKBONE_BIOSYNTHESIS
Proportion of pathway genes=0.87; Adjusted p−value=0

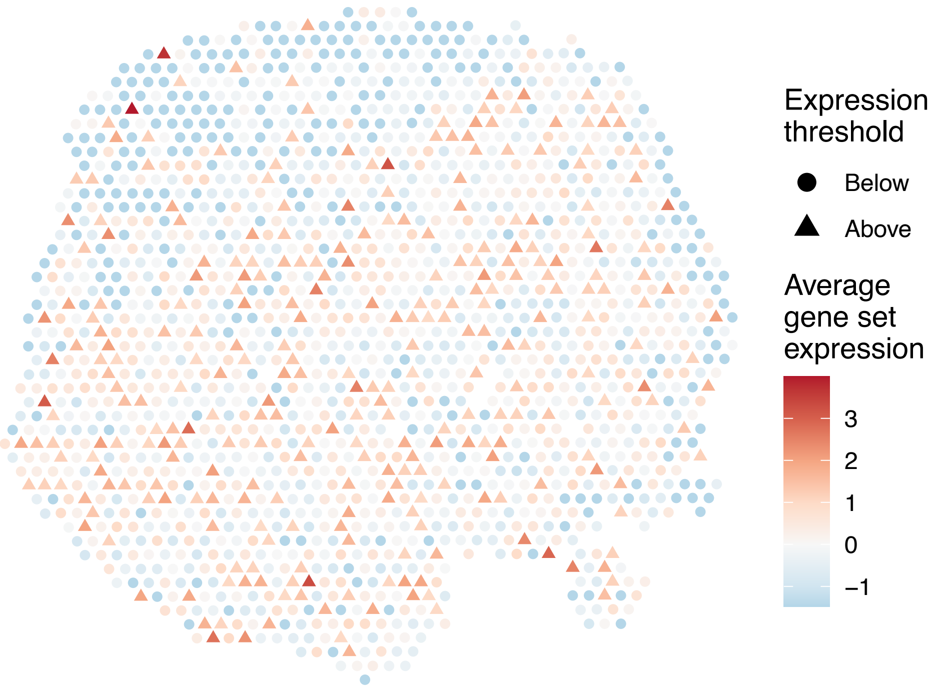


sample4_LMM_A5
KEGG_TGF_BETA_SIGNALING_PATHWAY
Proportion of pathway genes=0.73; Adjusted p−value=0

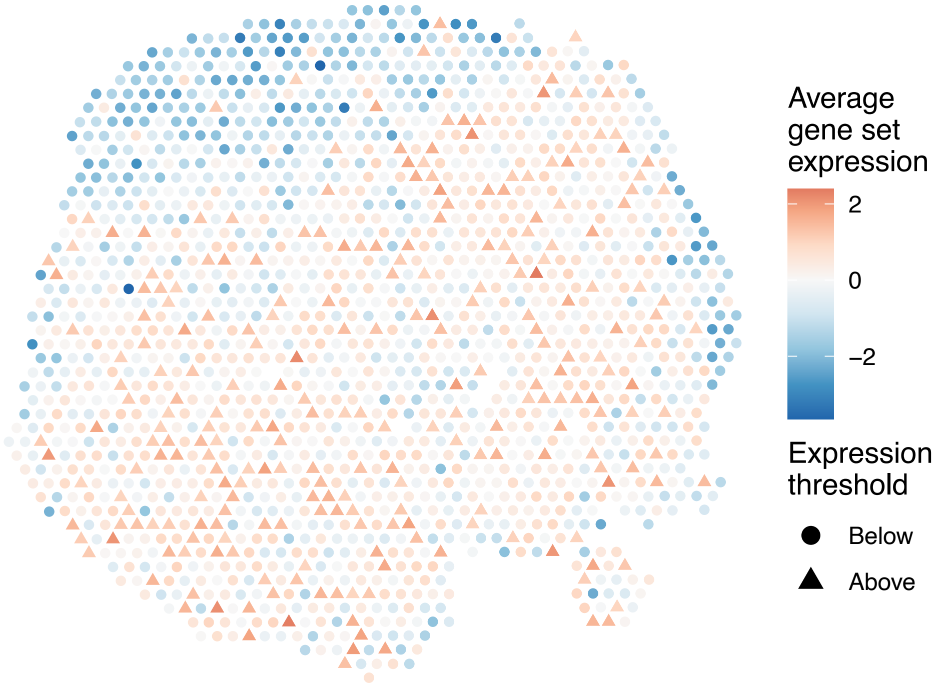


sample4_LMM_A5
KEGG_THYROID_CANCER
Proportion of pathway genes=0.91; Adjusted p−value=0.0347

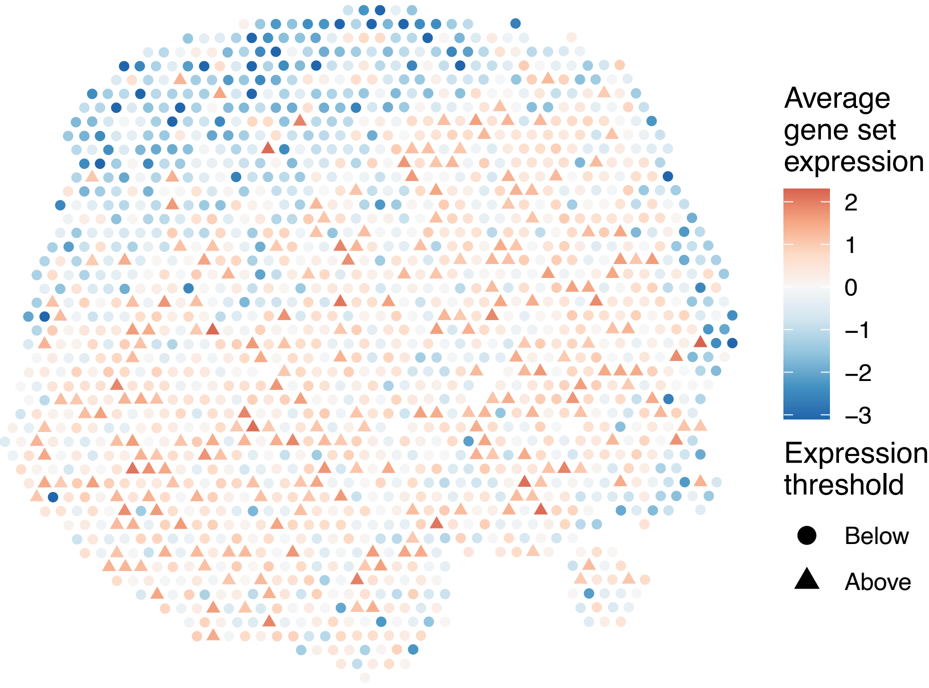

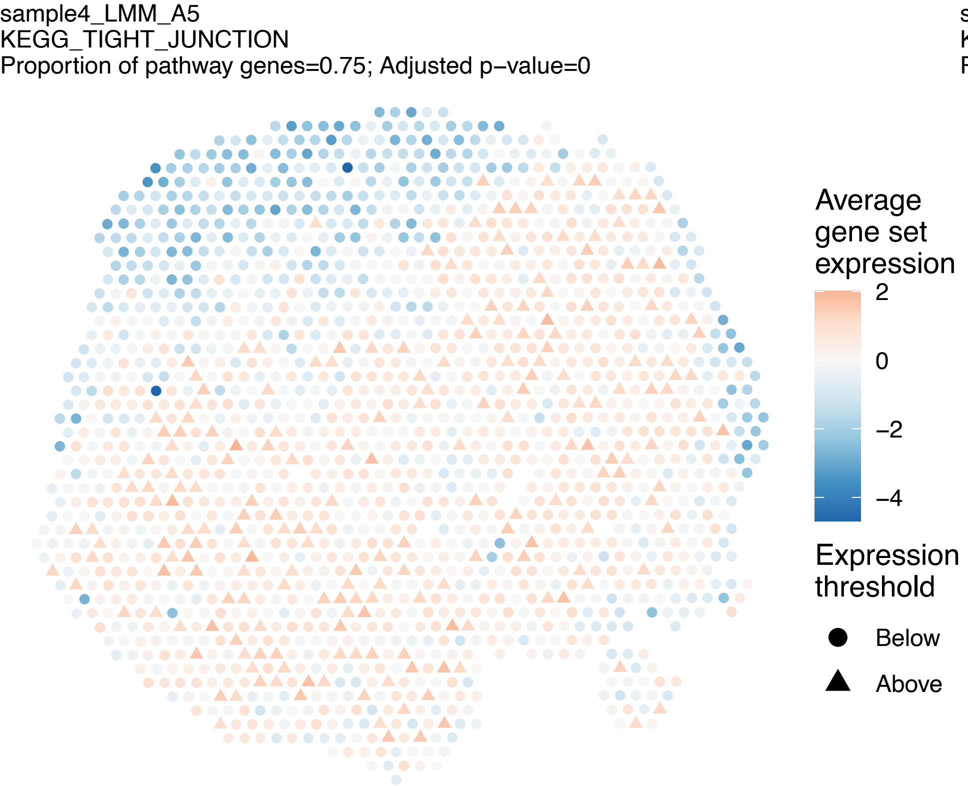
sample4_LMM_A5
KEGG_TIGHT_JUNCTION
Proportion of pathway genes=0.75; Adjusted p−value=0
Average gene set expression
2
0
−2
−4
Expression threshold
Below
Above

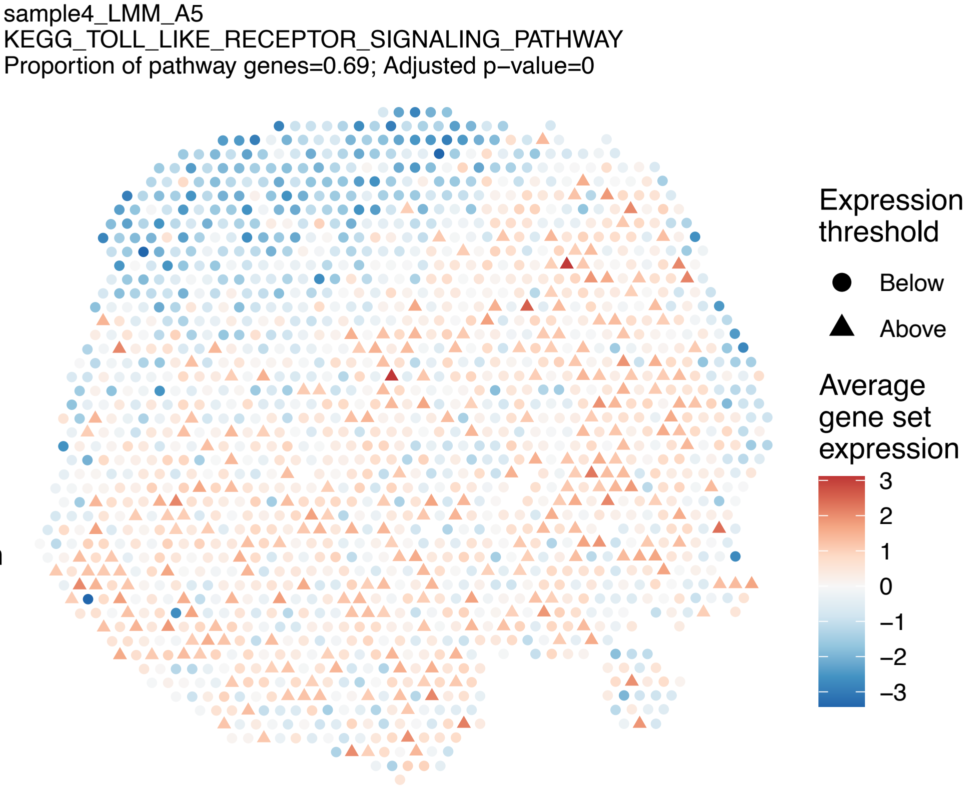
sample4_LMM_A5
KEGG_TOLL_LIKE_RECEPTOR_SIGNALING_PATHWAY
Proportion of pathway genes=0.69; Adjusted p−value=0
Expression threshold
Below
Above
Average gene set expression
3
2
1
0
−1
−2
−3

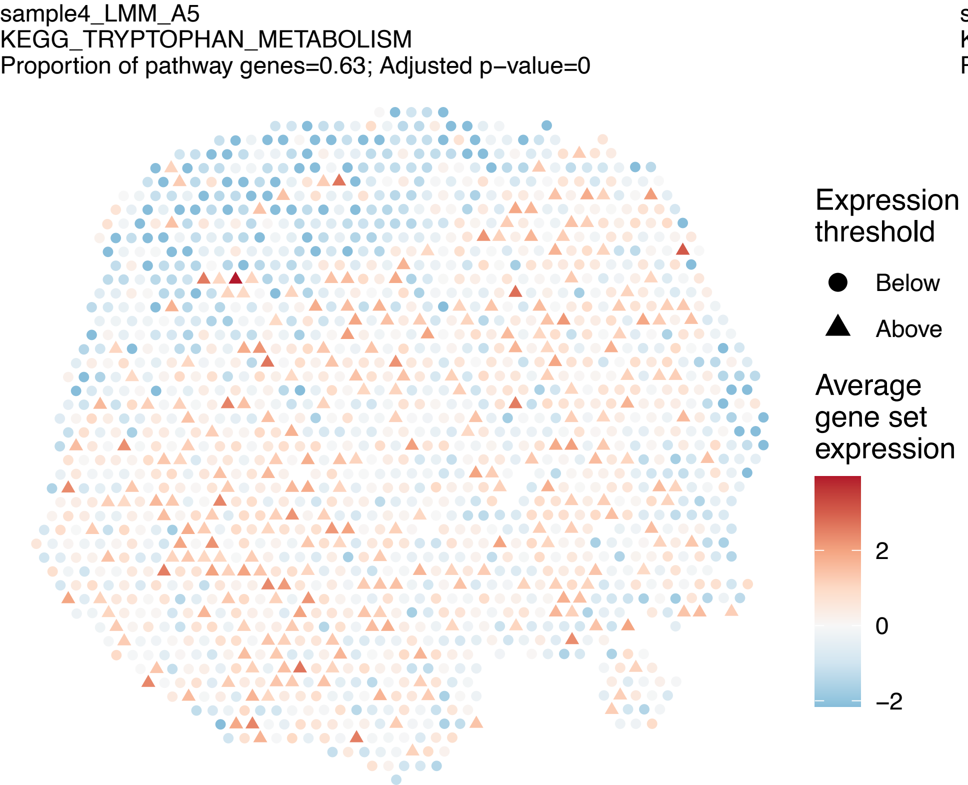
sample4_LMM_A5
KEGG_TRYPTOPHAN_METABOLISM
Proportion of pathway genes=0.63; Adjusted p−value=0
Expression threshold
Below
Above
Average gene set expression
2
0
−2

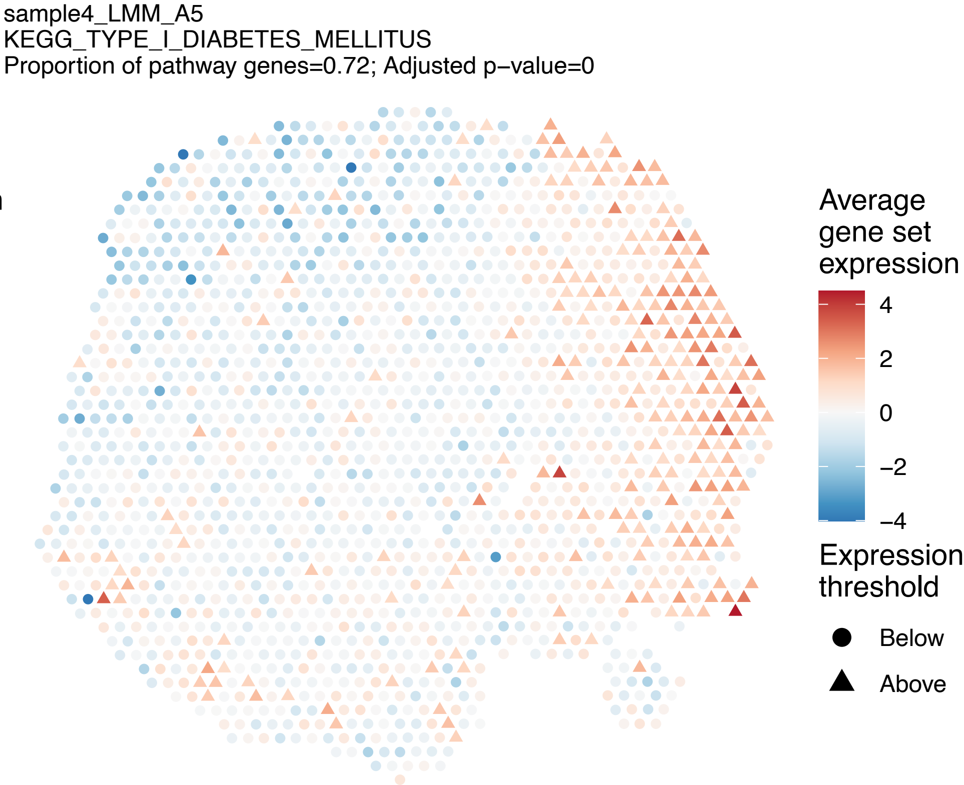
sample4_LMM_A5
KEGG_TYPE_I_DIABETES_MELLITUS
Proportion of pathway genes=0.72; Adjusted p−value=0
Average gene set expression
4
2
0
−2
−4
Expression threshold
Below
Above

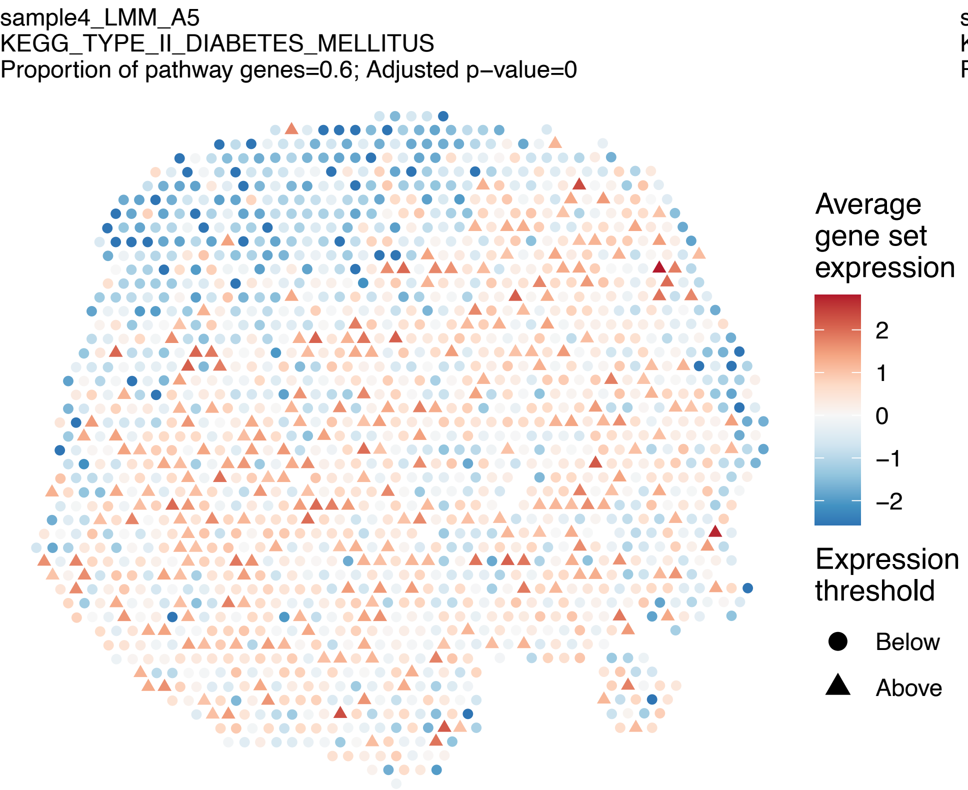
sample4_LMM_A5
KEGG_TYPE_II_DIABETES_MELLITUS
Proportion of pathway genes=0.6; Adjusted p−value=0
Average gene set expression
2
1
0
−1
−2
Expression threshold
Below
Above

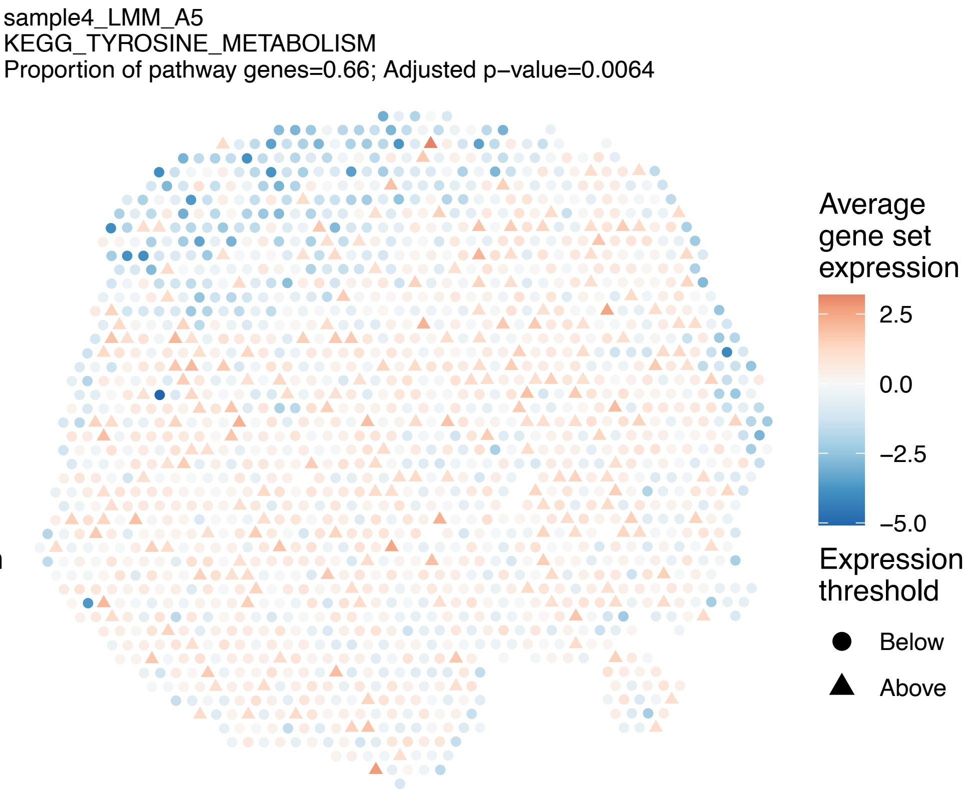
sample4_LMM_A5
KEGG_TYROSINE_METABOLISM
Proportion of pathway genes=0.66; Adjusted p−value=0.0064
Average gene set expression
2.5
0.0
−2.5
−5.0
Expression threshold
Below
Above

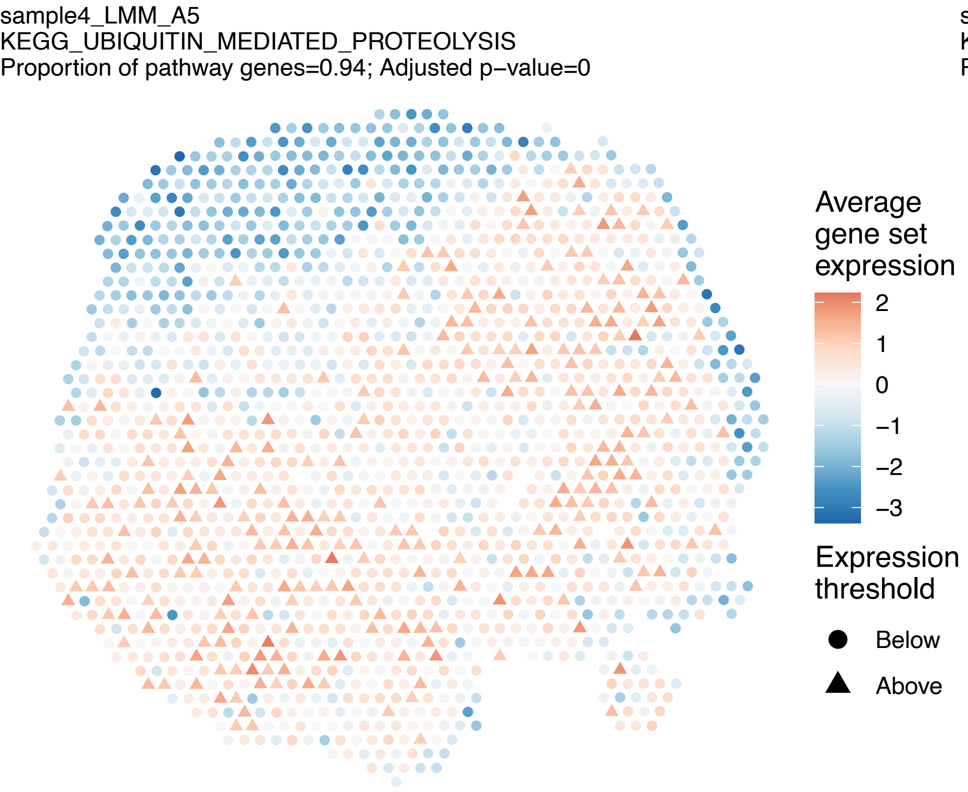
sample4_LMM_A5
KEGG_UBIQUITIN_MEDIATED_PROTEOLYSIS
Proportion of pathway genes=0.94; Adjusted p−value=0
Average gene set expression
2
1
0
−1
−2
−3
Expression threshold
Below
Above

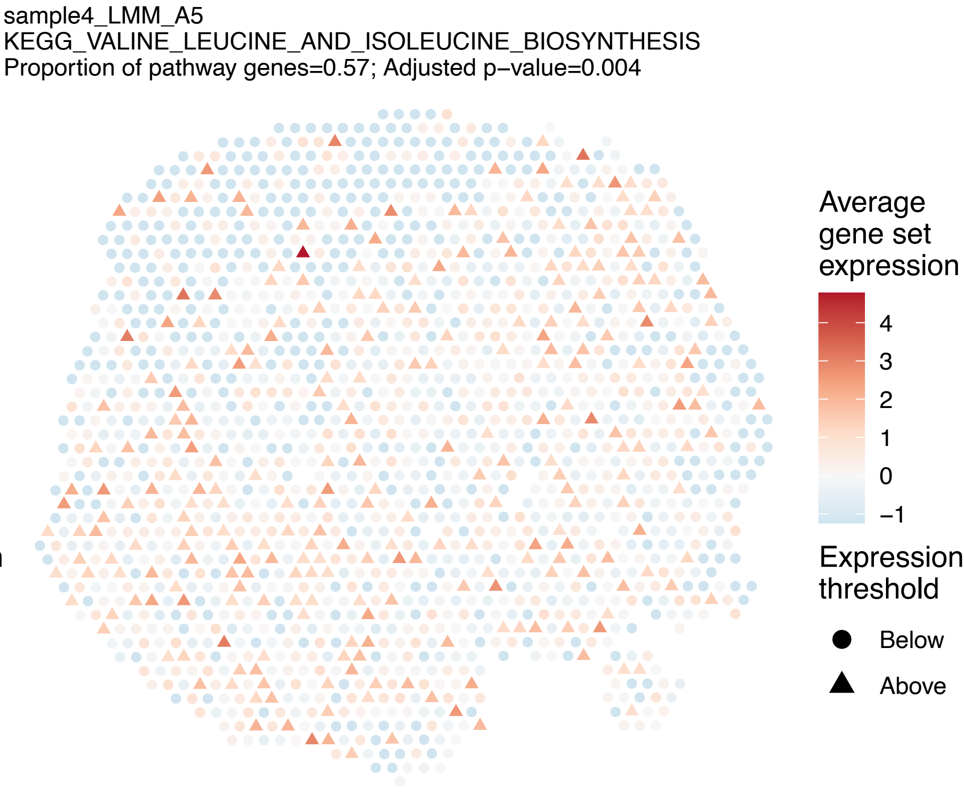
sample4_LMM_A5
KEGG_VALINE_LEUCINE_AND_ISOLEUCINE_BIOSYNTHESIS
Proportion of pathway genes=0.57; Adjusted p−value=0.004
Average gene set expression
4
3
2
1
0
−1
Expression threshold
Below
Above

sample4_LMM_A5
KEGG_VALINE_LEUCINE_AND_ISOLEUCINE_DEGRADATION
Proportion of pathway genes=0.98; Adjusted p−value=0

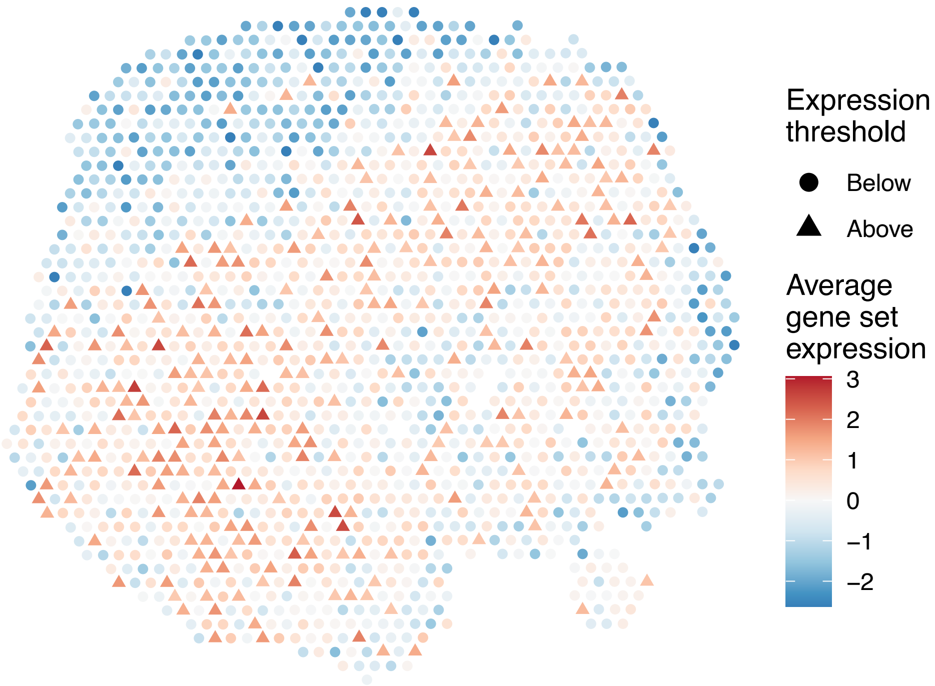


sample4_LMM_A5
KEGG_VASCULAR_SMOOTH_MUSCLE_CONTRACTION
Proportion of pathway genes=0.66; Adjusted p−value=0

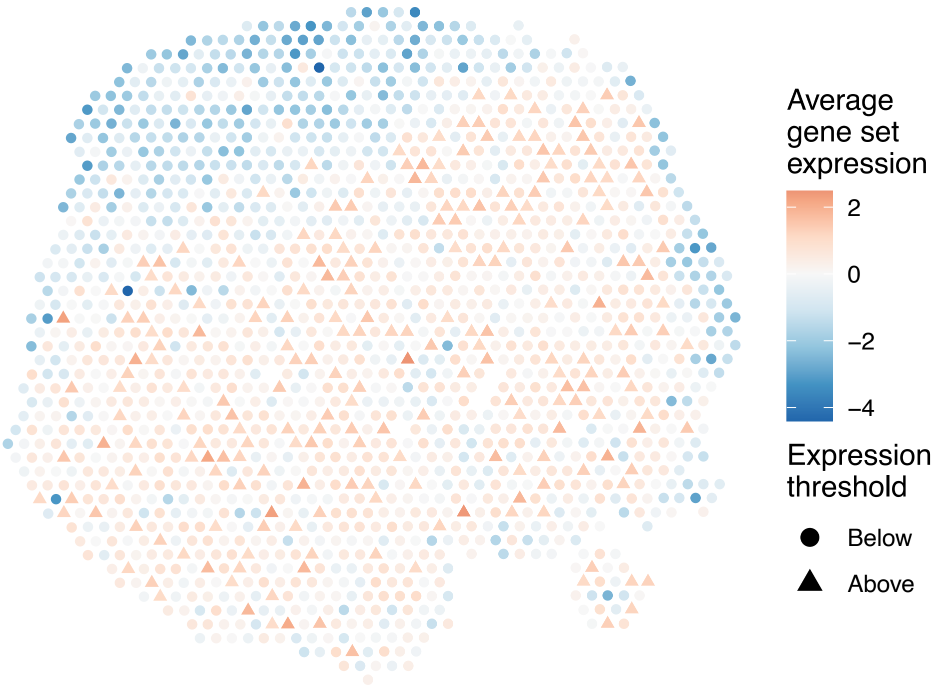


sample4_LMM_A5
KEGG_VASOPRESSIN_REGULATED_WATER_REABSORPTION
Proportion of pathway genes=0.89; Adjusted p−value=0

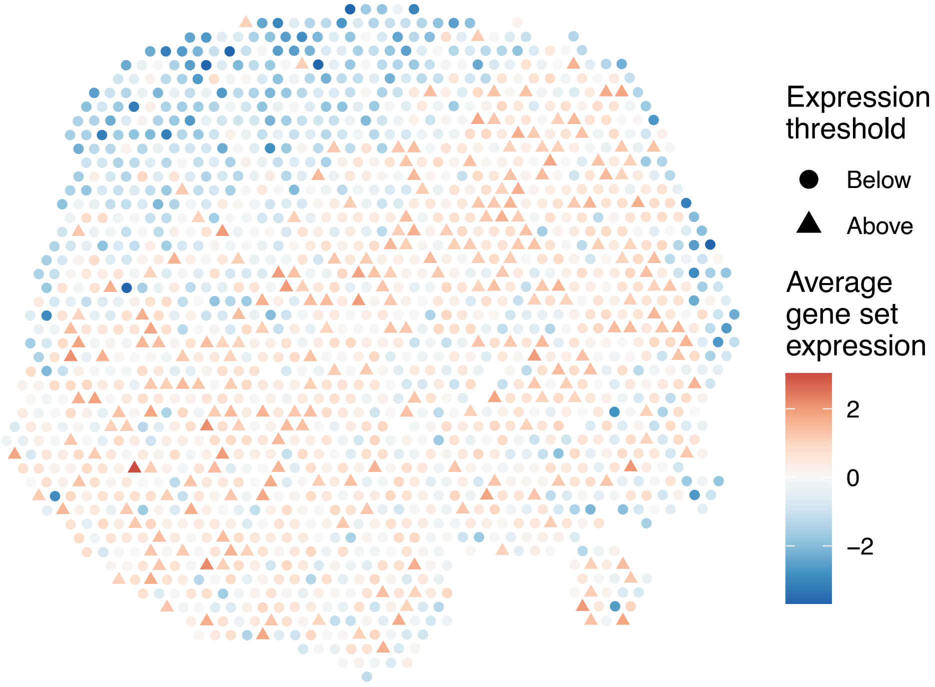


sample4_LMM_A5
KEGG_VEGF_SIGNALING_PATHWAY
Proportion of pathway genes=0.79; Adjusted p−value=0

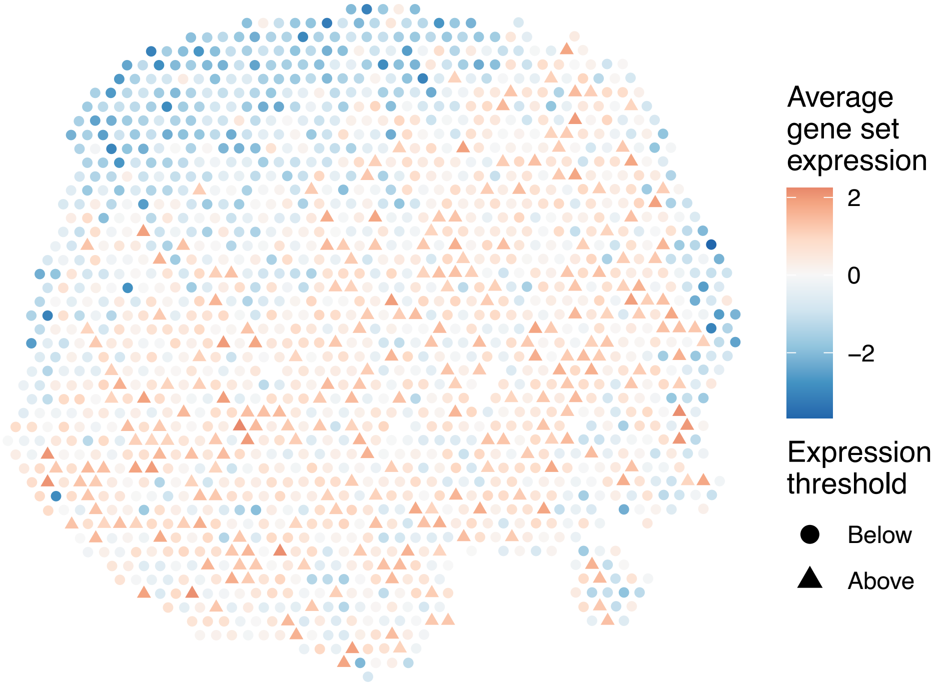

sample4_LMM_A5
KEGG_VIBRIO_CHOLERAE_INFECTION
Proportion of pathway genes=0.73; Adjusted p−value=0

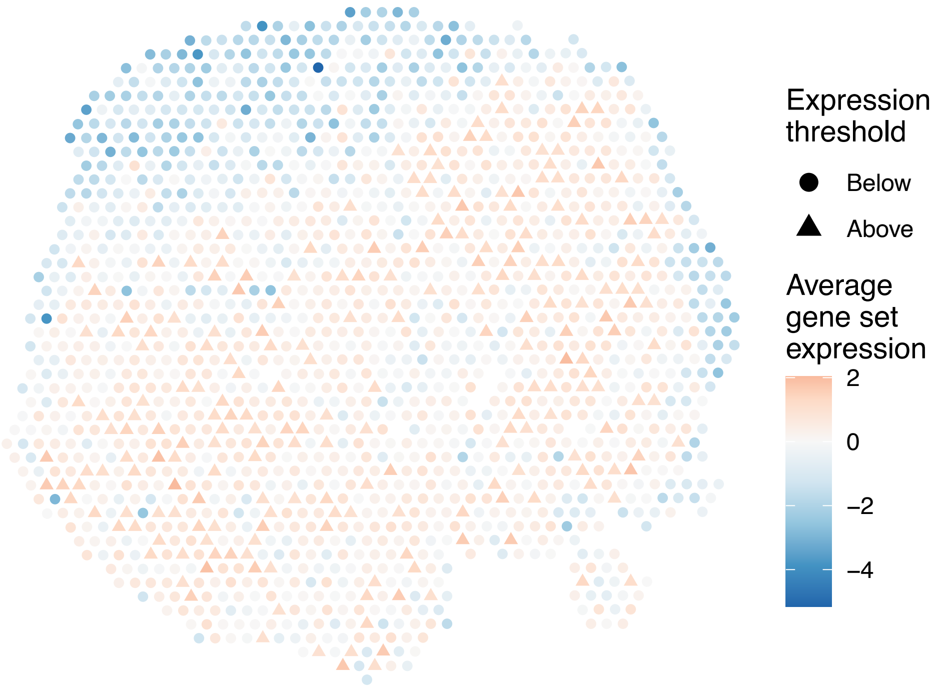


sample4_LMM_A5
KEGG_VIRAL_MYOCARDITIS
Proportion of pathway genes=0.81; Adjusted p−value=0

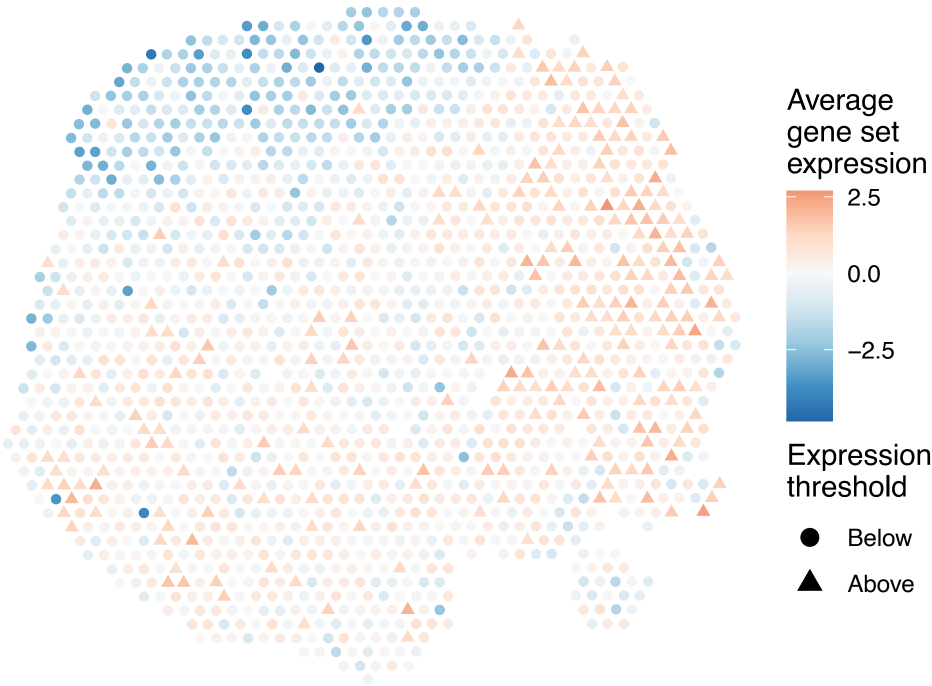


sample4_LMM_A5
KEGG_WNT_SIGNALING_PATHWAY
Proportion of pathway genes=0.8; Adjusted p−value=0

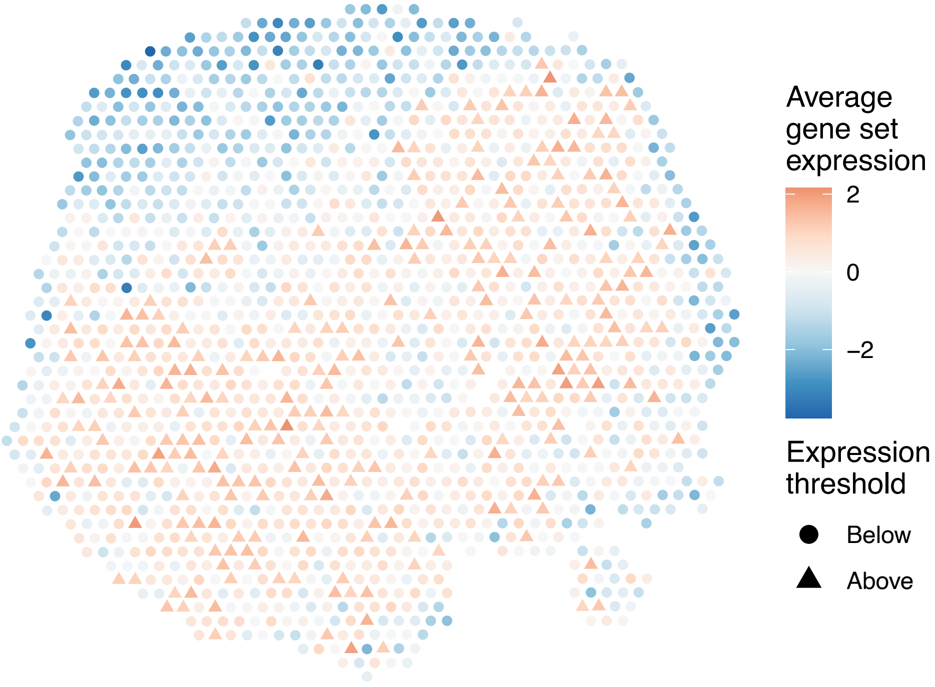


sample4_LMM_A5
KEGG_PENTOSE_AND_GLUCURONATE_INTERCONVERSIONS
Proportion of pathway genes=0.29; Adjusted p−value=NA

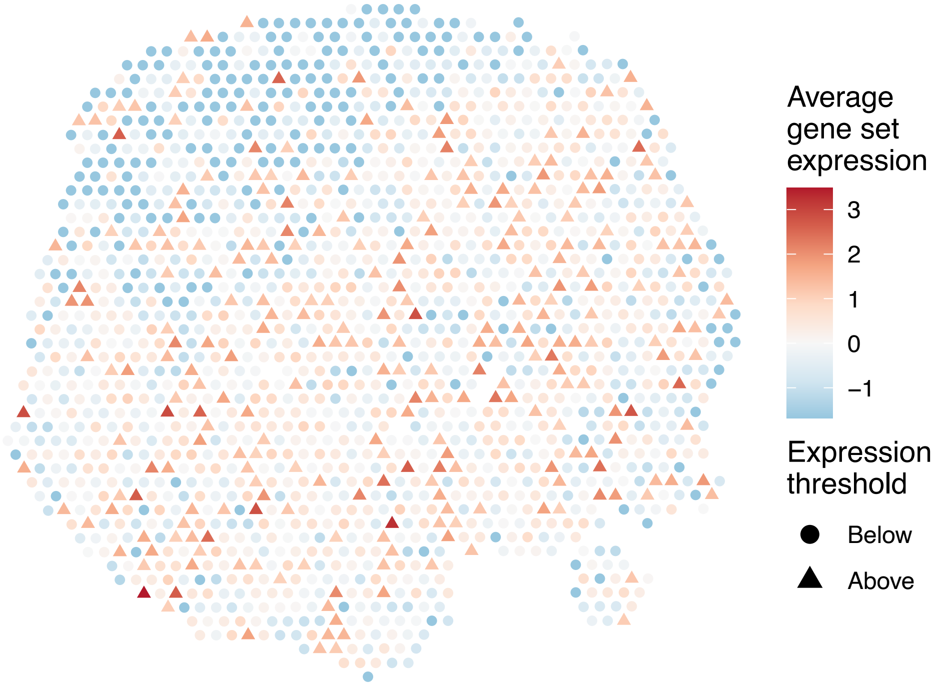

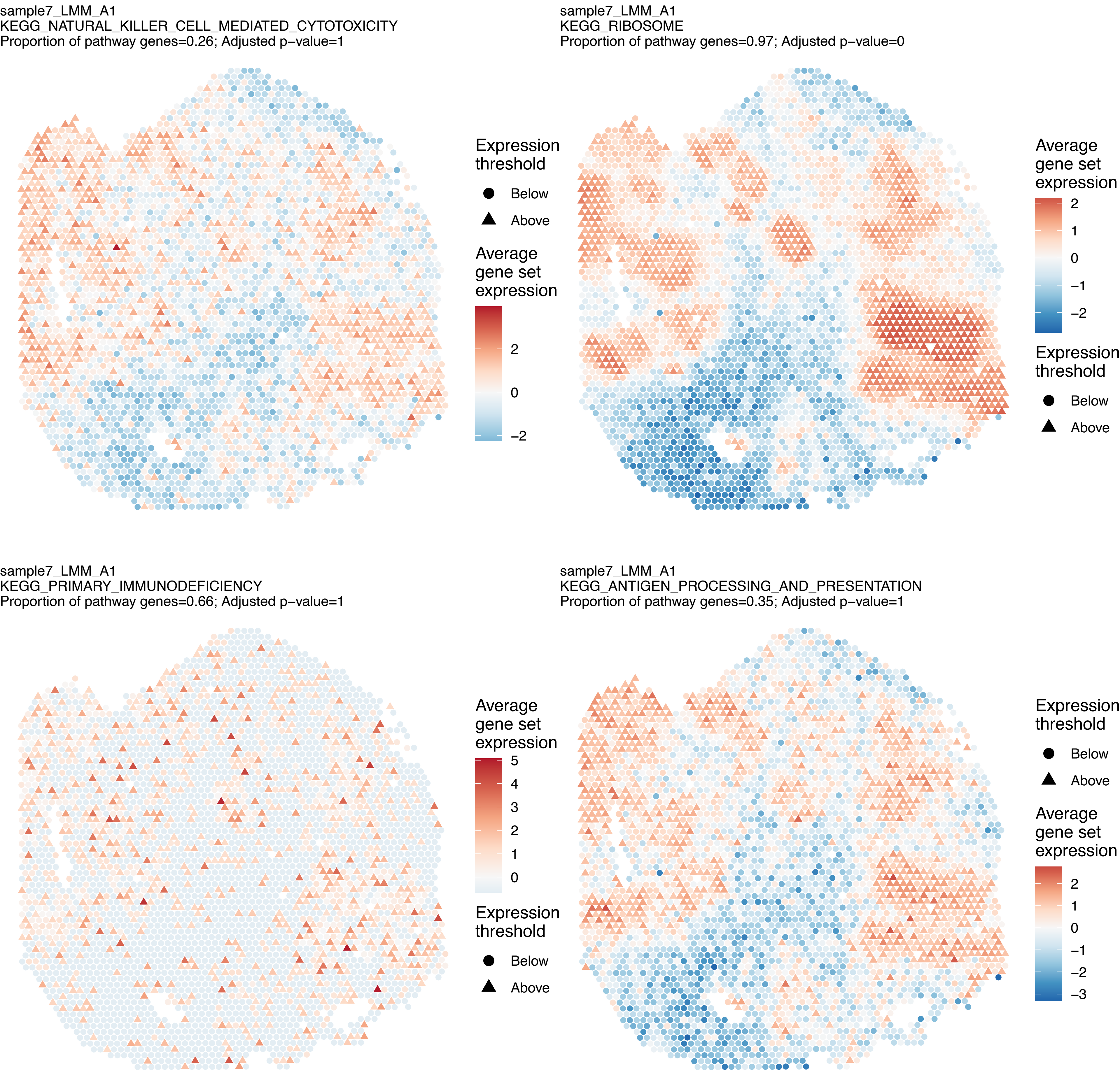
sample7_LMM_A1
KEGG_NATURAL_KILLER_CELL_MEDIATED_CYTOTOXICITY
Proportion of pathway genes=0.26; Adjusted p–value=1
Expression threshold
Below
Above
Average gene set expression
2
0
–2
sample7_LMM_A1
KEGG_RIBOSOME
Proportion of pathway genes=0.97; Adjusted p–value=0
Average gene set expression
2
1
0
–1
–2
Expression threshold
Below
Above
sample7_LMM_A1
KEGG_PRIMARY_IMMUNODEFICIENCY
Proportion of pathway genes=0.66; Adjusted p–value=1
Average gene set expression
5
4
3
2
1
0
Expression threshold
Below
Above
sample7_LMM_A1
KEGG_ANTIGEN_PROCESSING_AND_PRESENTATION
Proportion of pathway genes=0.35; Adjusted p–value=1
Expression threshold
Below
Above
Average gene set expression
2
1
0
–1
–2
–3

sample7_LMM_A1
KEGG_CYTOKINE_CYTOKINE_RECEPTOR_INTERACTION
Proportion of pathway genes=0.37; Adjusted p−value=1

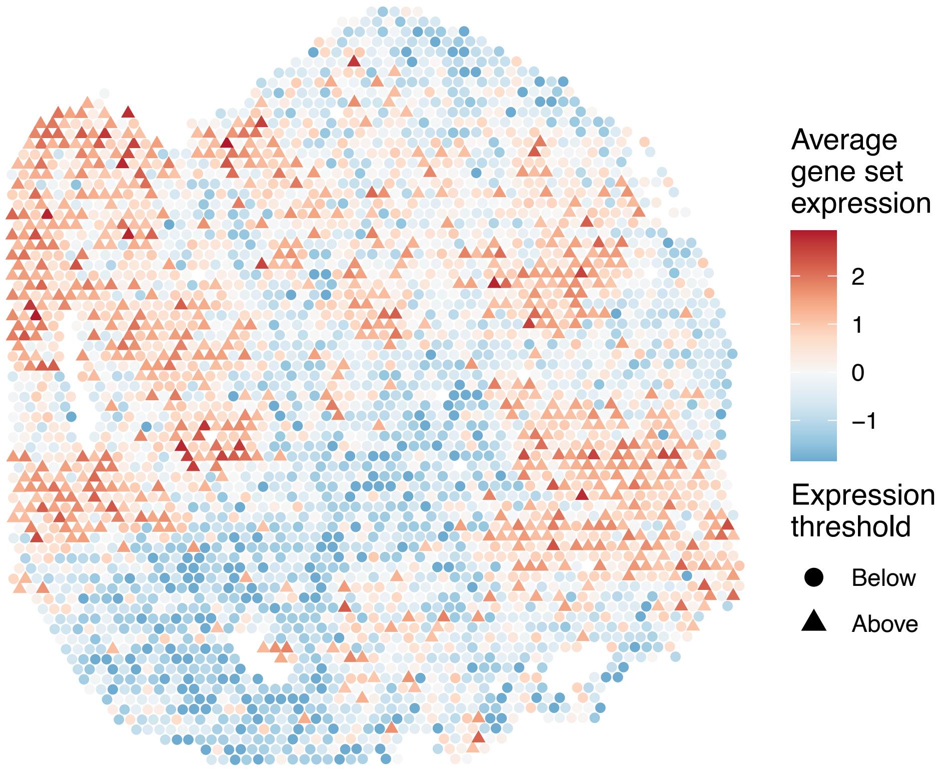


sample7_LMM_A1
KEGG_ABC_TRANSPORTERS
Proportion of pathway genes=0.82; Adjusted p−value=1

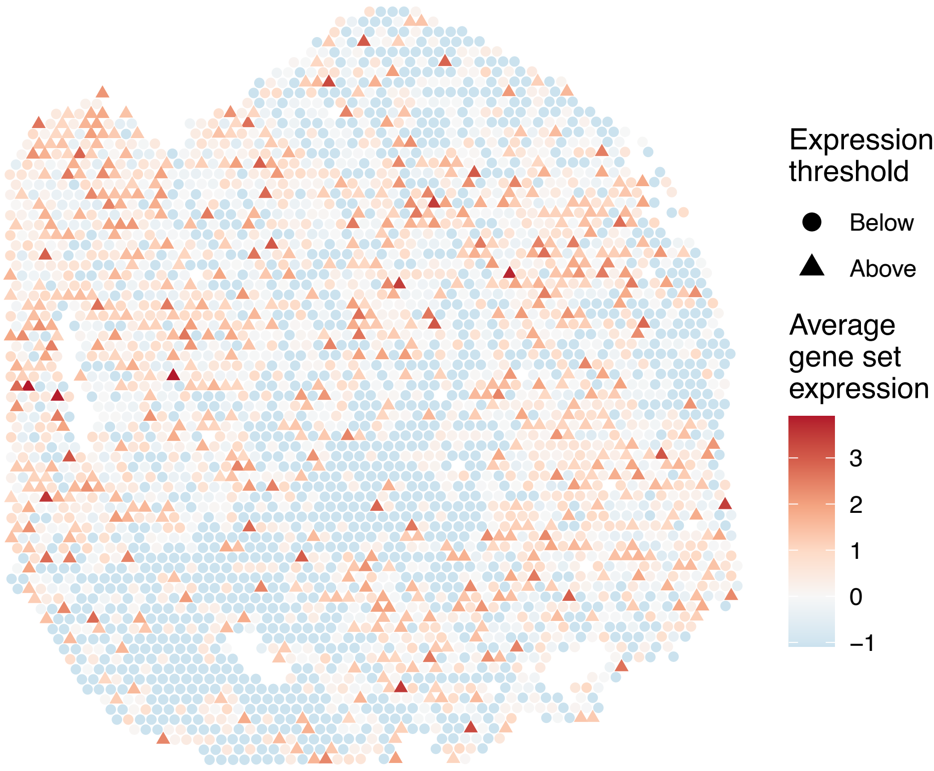


sample7_LMM_A1
KEGG_ACUTE_MYELOID_LEUKEMIA
Proportion of pathway genes=0.97; Adjusted p−value=1

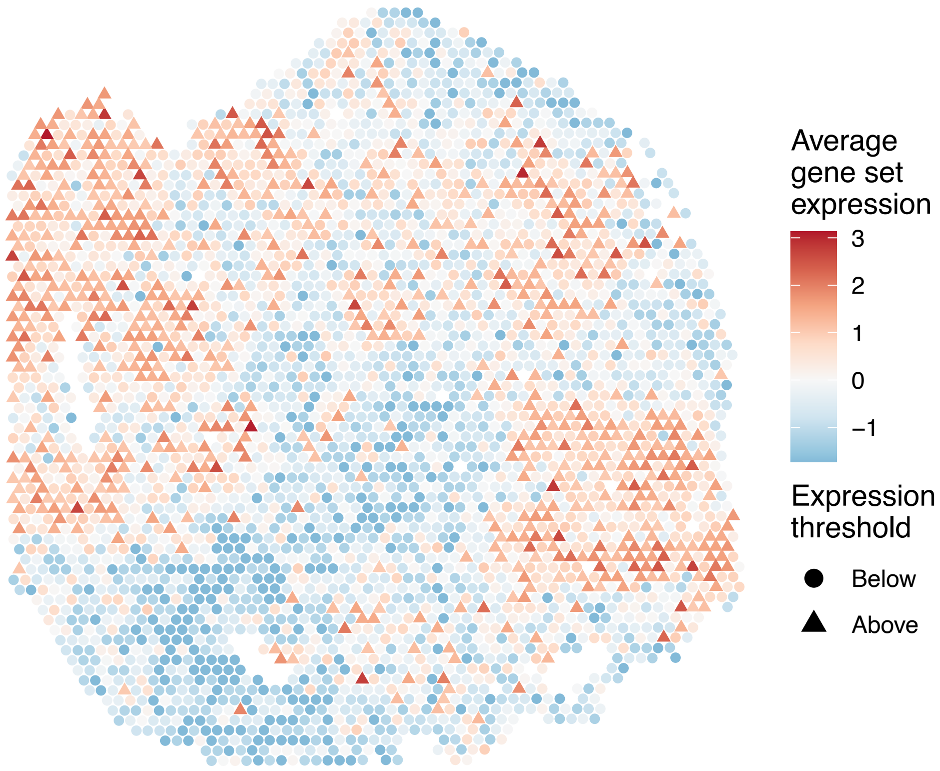


sample7_LMM_A1
KEGG_ADHERENS_JUNCTION
Proportion of pathway genes=0.95; Adjusted p−value=1

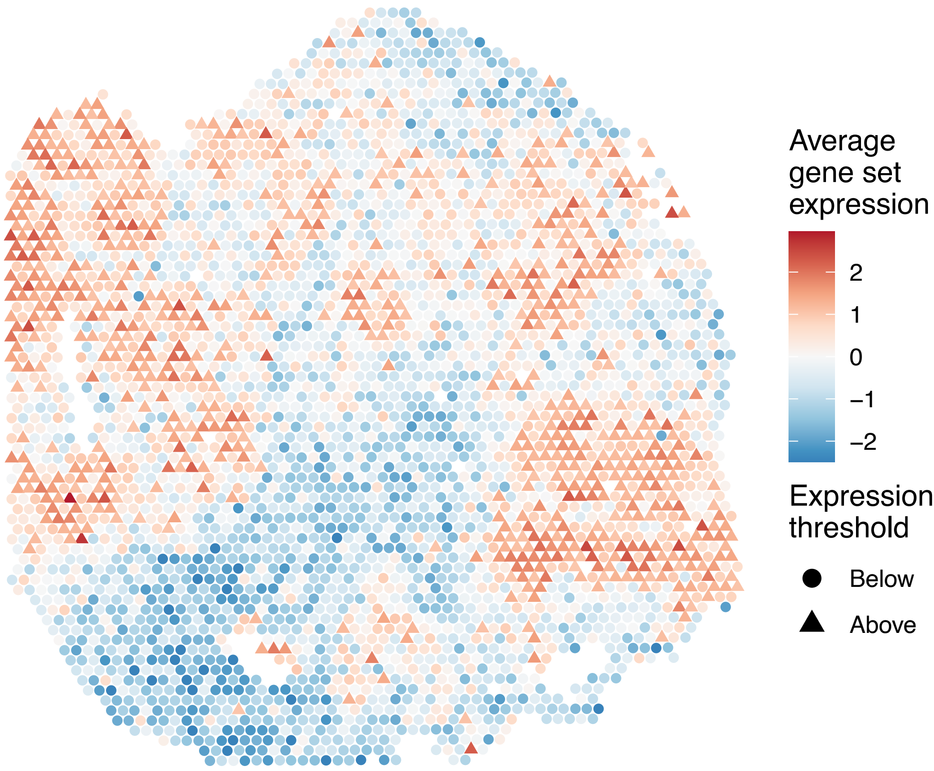

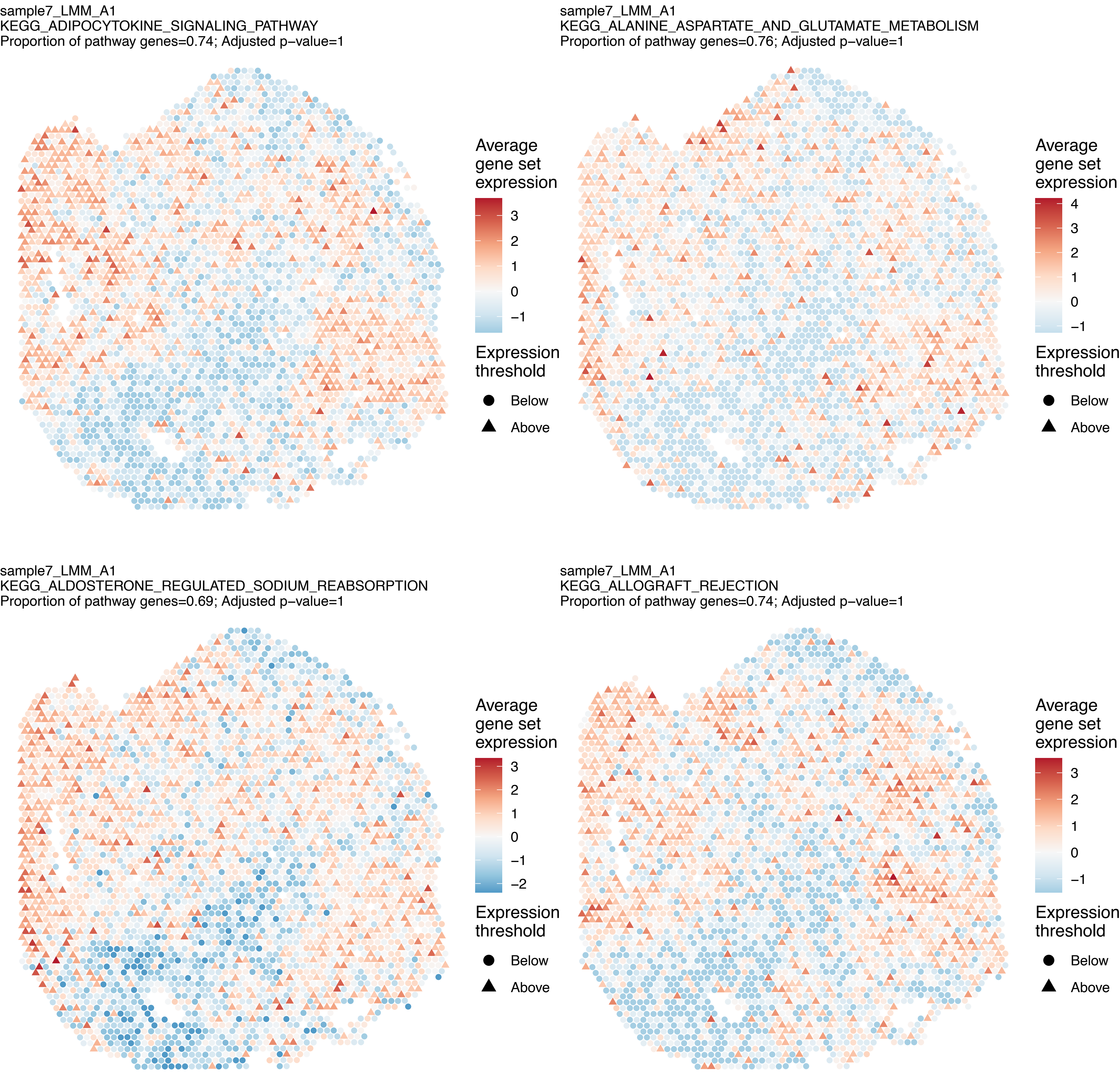
sample7_LMM_A1
KEGG_ADIPOCYTOKINE_SIGNALING_PATHWAY
Proportion of pathway genes=0.74; Adjusted p−value=1
Average gene set expression
3
2
1
0
−1
Expression threshold
Below
Above
sample7_LMM_A1
KEGG_ALANINE_ASPARTATE_AND_GLUTAMATE_METABOLISM
Proportion of pathway genes=0.76; Adjusted p−value=1
Average gene set expression
4
3
2
1
0
−1
Expression threshold
Below
Above
sample7_LMM_A1
KEGG_ALDOSTERONE_REGULATED_SODIUM_REABSORPTION
Proportion of pathway genes=0.69; Adjusted p−value=1
Average gene set expression
3
2
1
0
−1
−2
Expression threshold
Below
Above
sample7_LMM_A1
KEGG_ALLOGRAFT_REJECTION
Proportion of pathway genes=0.74; Adjusted p−value=1
Average gene set expression
3
2
1
0
−1
Expression threshold
Below
Above

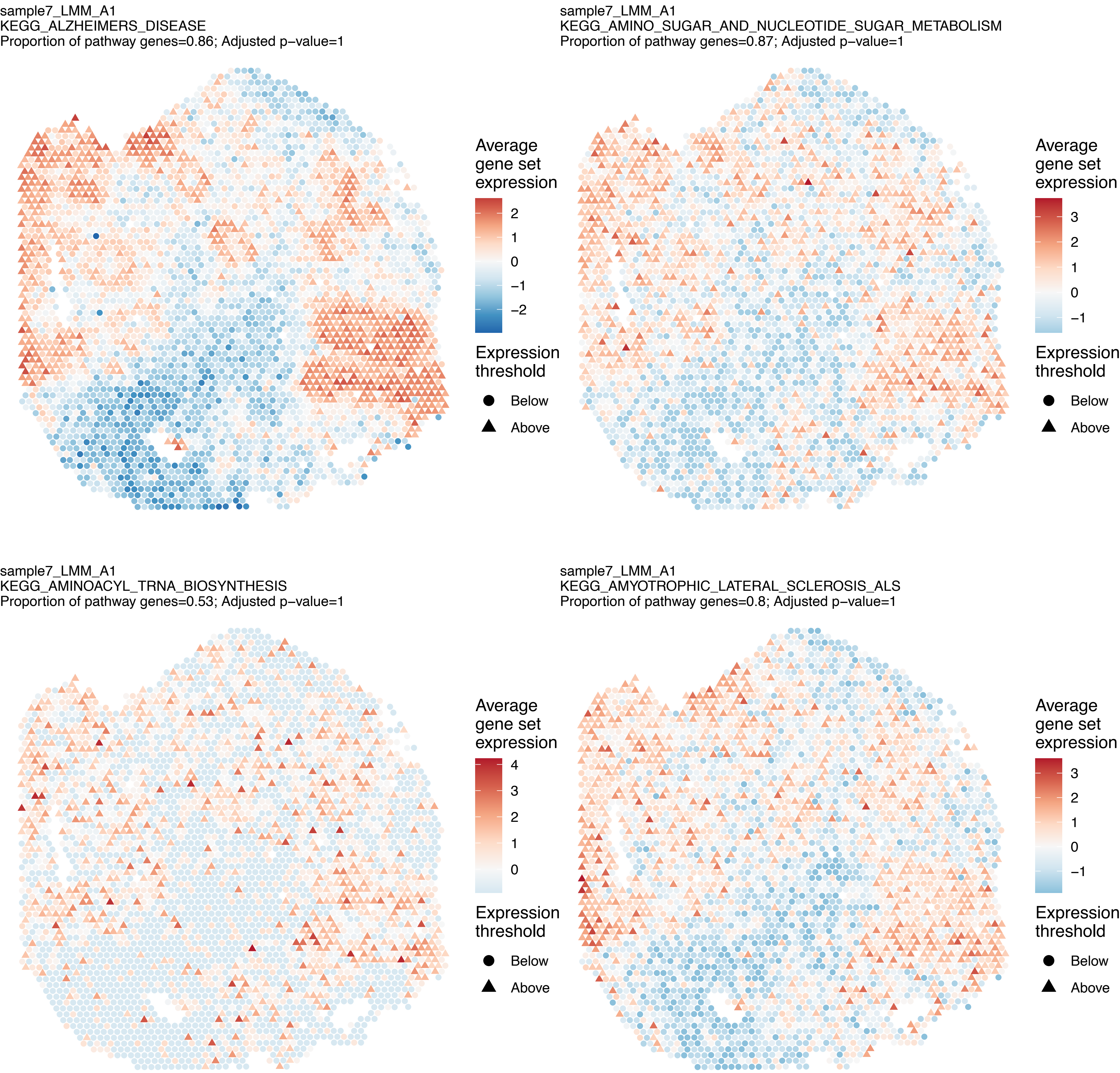
sample7_LMM_A1
KEGG_ALZHEIMERS_DISEASE
Proportion of pathway genes=0.86; Adjusted p−value=1
Average gene set expression
2
1
0
−1
−2
Expression threshold
Below
Above
sample7_LMM_A1
KEGG_AMINO_SUGAR_AND_NUCLEOTIDE_SUGAR_METABOLISM
Proportion of pathway genes=0.87; Adjusted p−value=1
Average gene set expression
3
2
1
0
−1
Expression threshold
Below
Above
sample7_LMM_A1
KEGG_AMINOACYL_TRNA_BIOSYNTHESIS
Proportion of pathway genes=0.53; Adjusted p−value=1
Average gene set expression
4
3
2
1
0
Expression threshold
Below
Above
sample7_LMM_A1
KEGG_AMYOTROPHIC_LATERAL_SCLEROSIS_ALS
Proportion of pathway genes=0.8; Adjusted p−value=1
Average gene set expression
3
2
1
0
−1
Expression threshold
Below
Above

sample7_LMM_A1
KEGG_APOPTOSIS
Proportion of pathway genes=0.8; Adjusted p–value=1

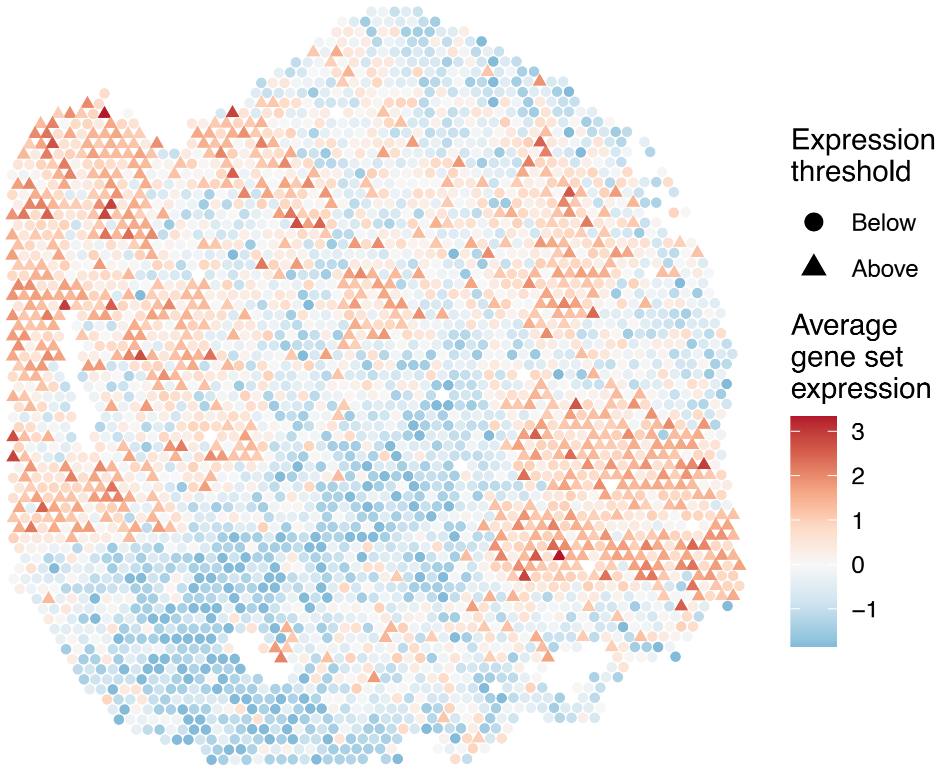


sample7_LMM_A1
KEGG_ARACHIDONIC_ACID_METABOLISM
Proportion of pathway genes=0.39; Adjusted p–value=1

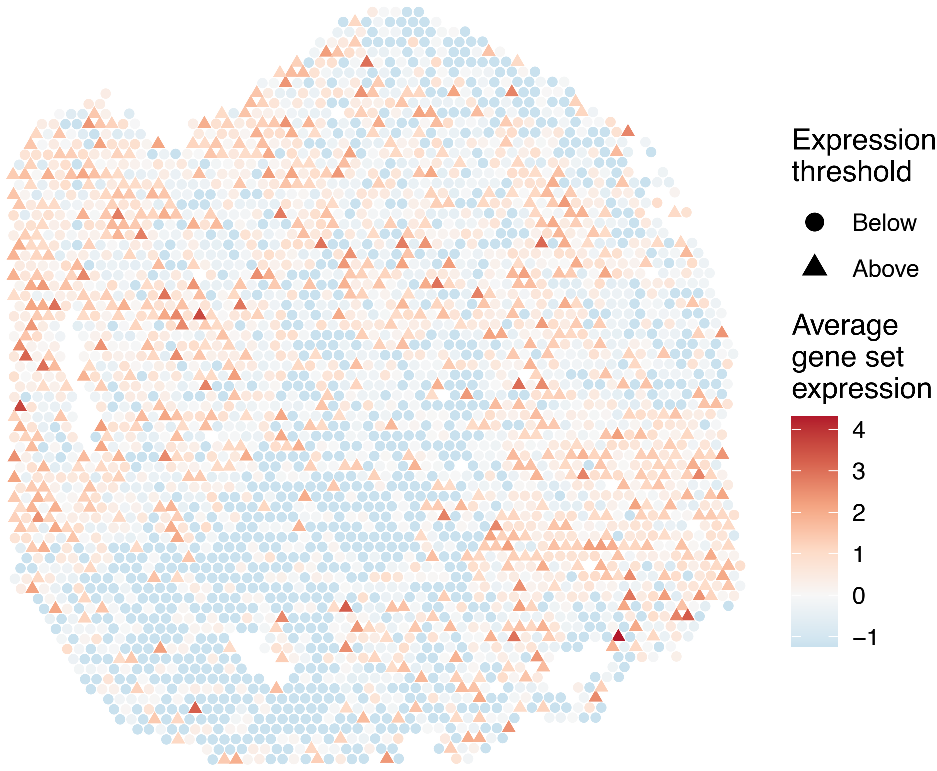


sample7_LMM_A1
KEGG_ARGININE_AND_PROLINE_METABOLISM
Proportion of pathway genes=0.8; Adjusted p–value=1

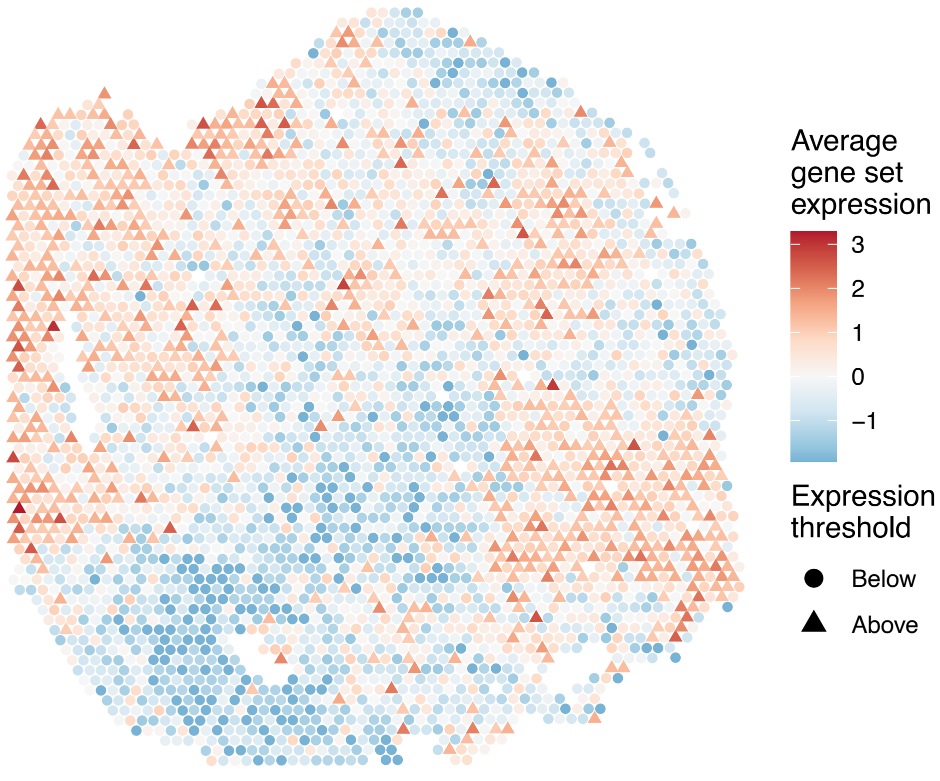


sample7_LMM_A1
KEGG_ARRHYTHMOGENIC_RIGHT_VENTRICULAR_CARDIOMYOPAT
HY_ARVC
Proportion of pathway genes=0.83; Adjusted p–value=1

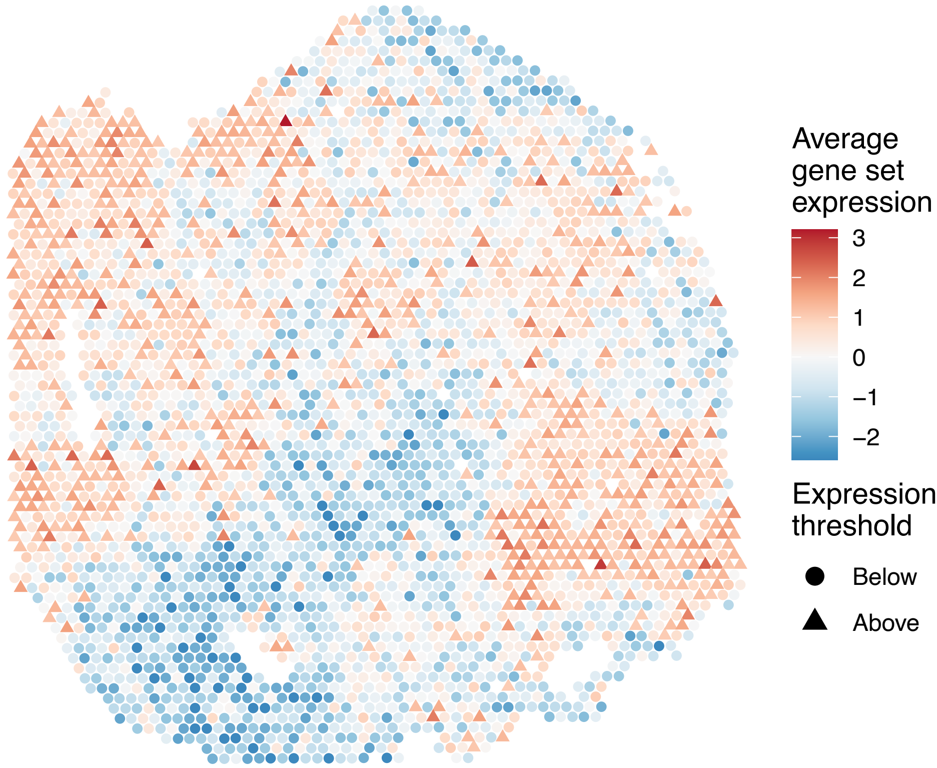

sample7_LMM_A1
KEGG_ASTHMA
Proportion of pathway genes=0.72; Adjusted p−value=1

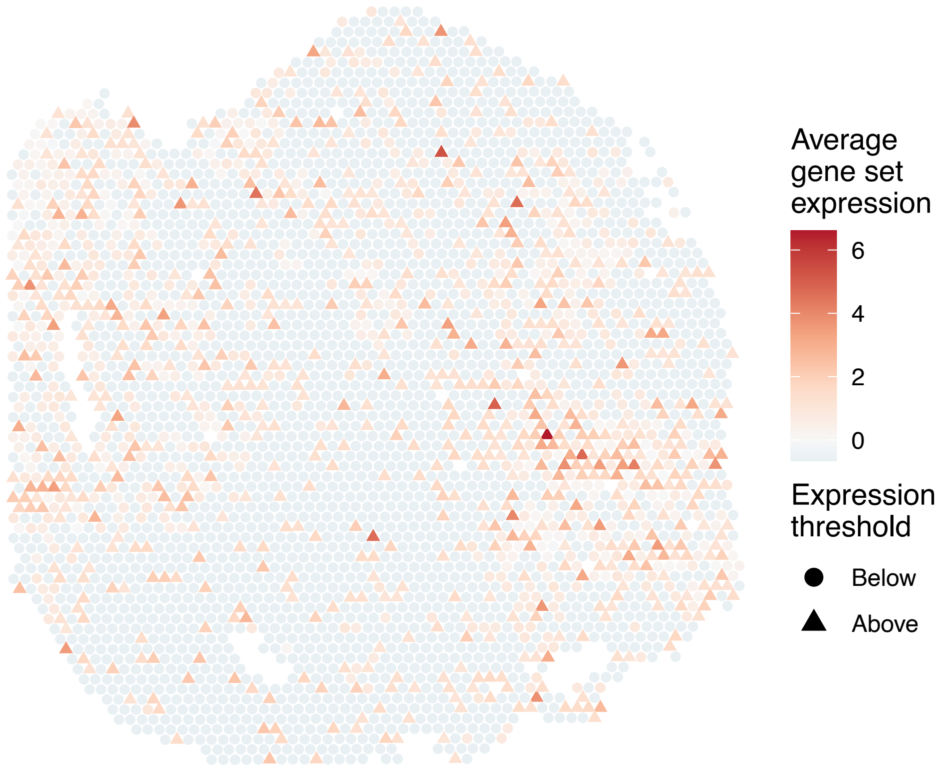


sample7_LMM_A1
KEGG_AUTOIMMUNE_THYROID_DISEASE
Proportion of pathway genes=0.7; Adjusted p−value=1

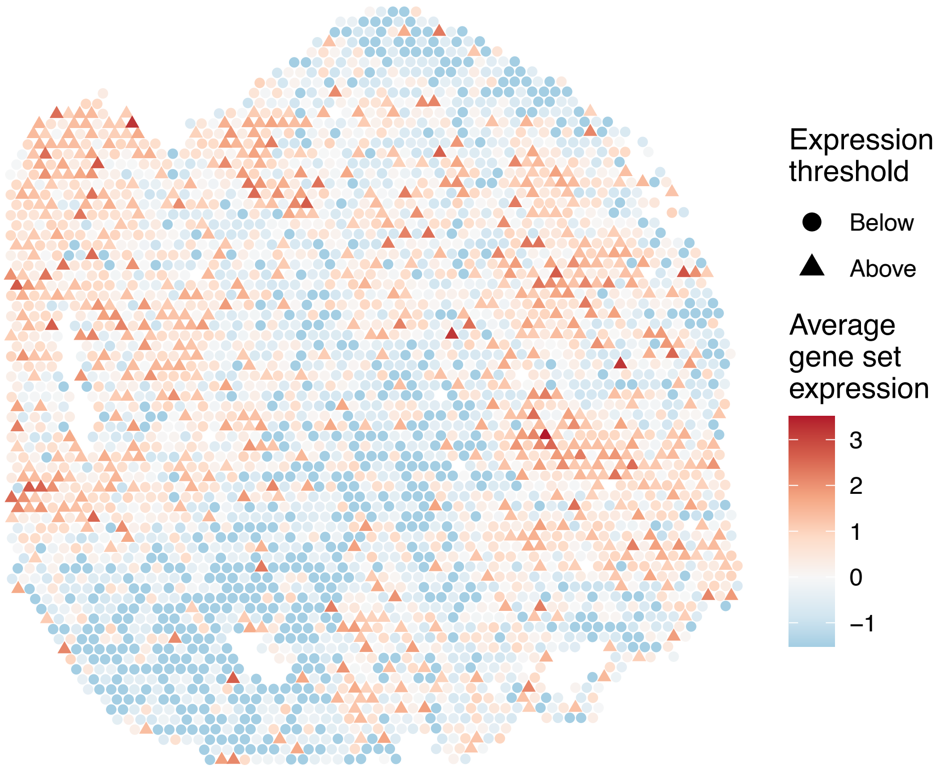


sample7_LMM_A1
KEGG_AXON_GUIDANCE
Proportion of pathway genes=0.9; Adjusted p−value=1

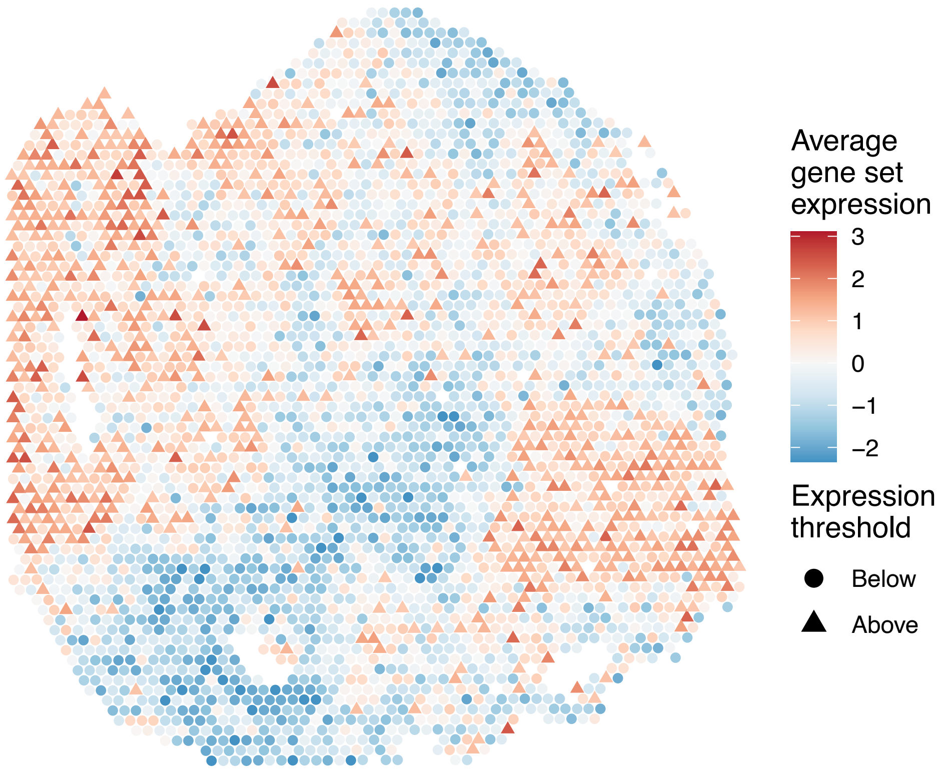


sample7_LMM_A1
KEGG_B_CELL_RECEPTOR_SIGNALING_PATHWAY
Proportion of pathway genes=0.86; Adjusted p−value=1

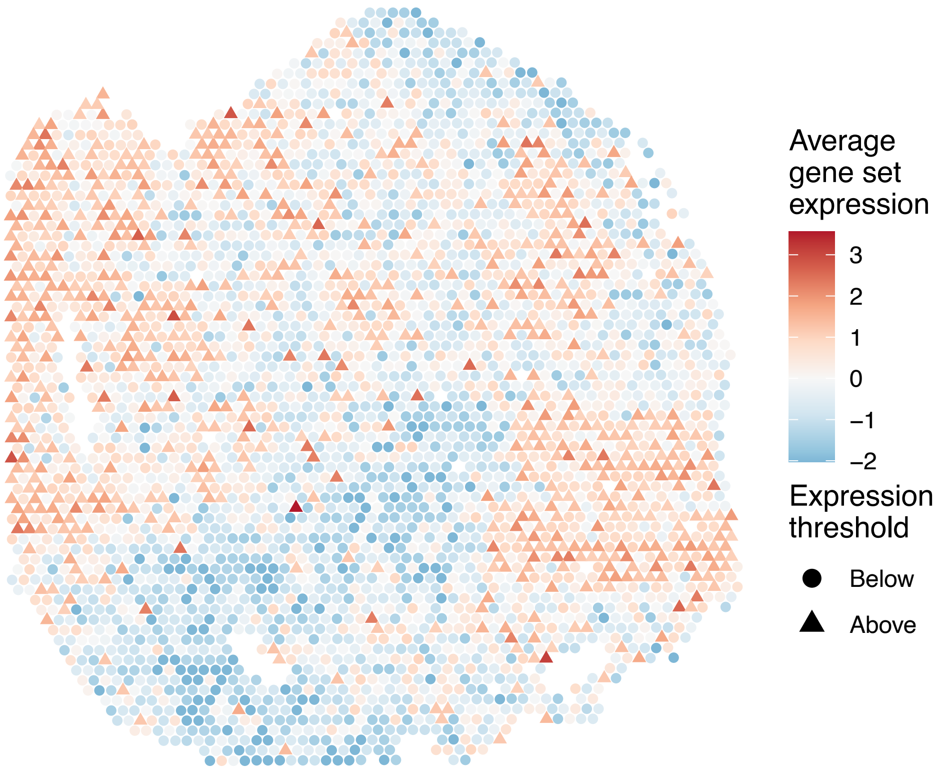

sample7_LMM_A1
KEGG_BASAL_CELL_CARCINOMA
Proportion of pathway genes=0.76; Adjusted p–value=1

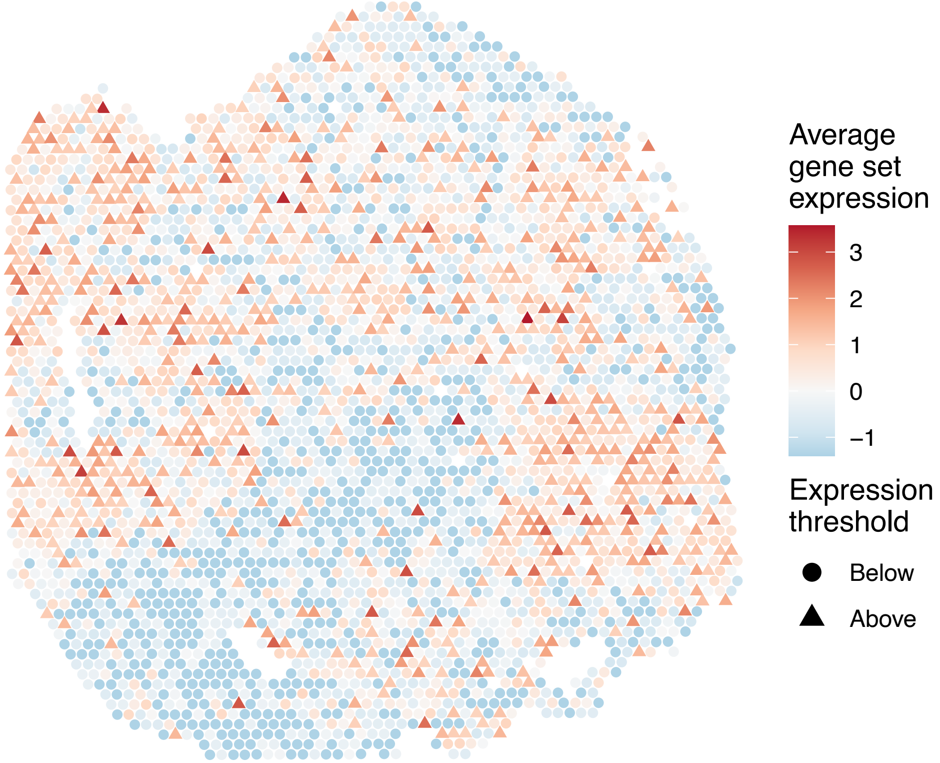


sample7_LMM_A1
KEGG_BASAL_TRANSCRIPTION_FACTORS
Proportion of pathway genes=0.91; Adjusted p–value=1

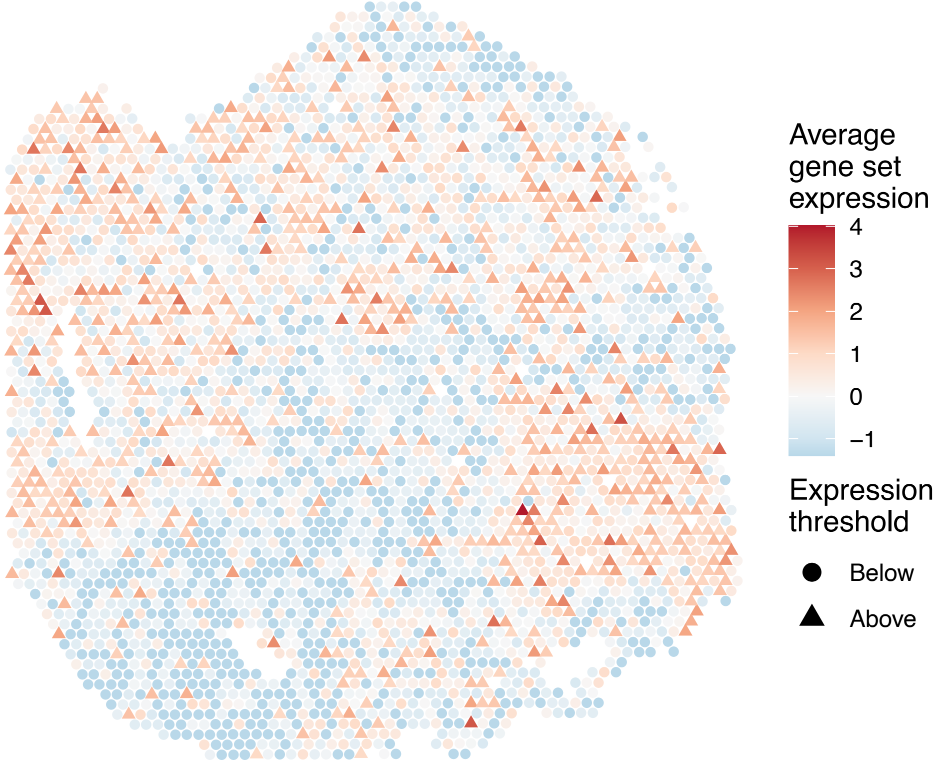


sample7_LMM_A1
KEGG_BASE_EXCISION_REPAIR
Proportion of pathway genes=0.94; Adjusted p–value=1

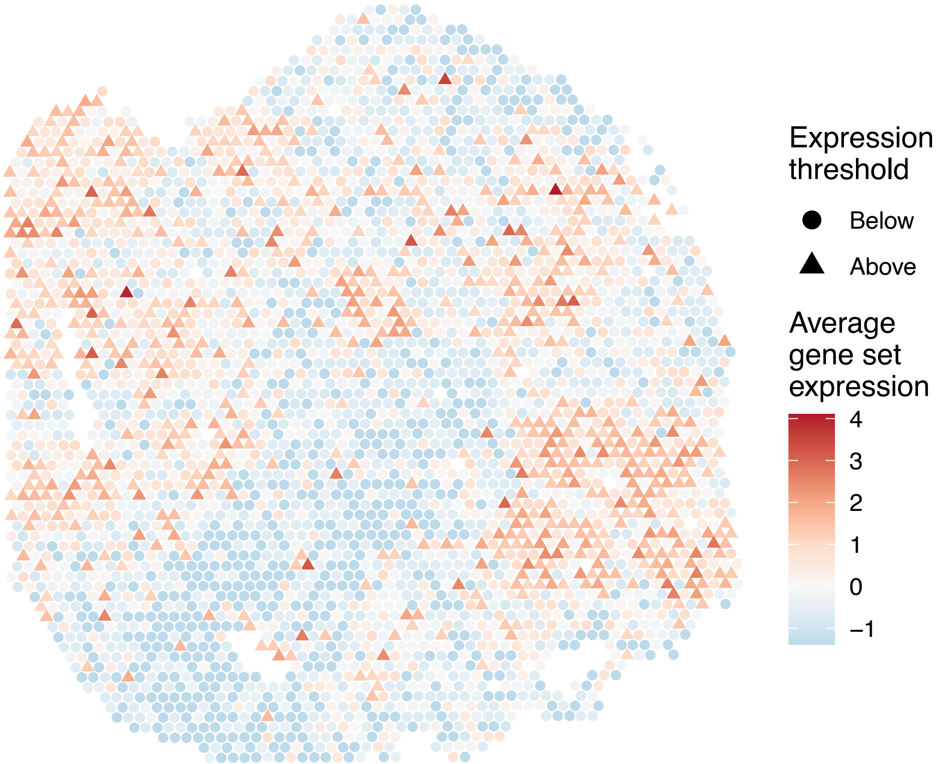


sample7_LMM_A1
KEGG_BETA_ALANINE_METABOLISM
Proportion of pathway genes=0.91; Adjusted p–value=1

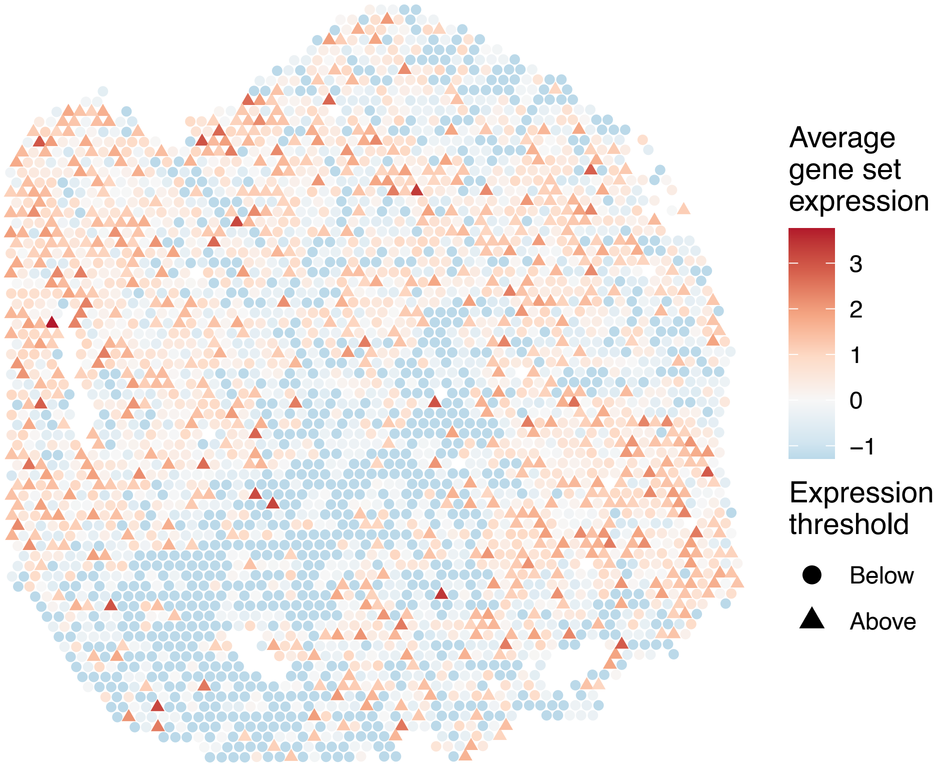

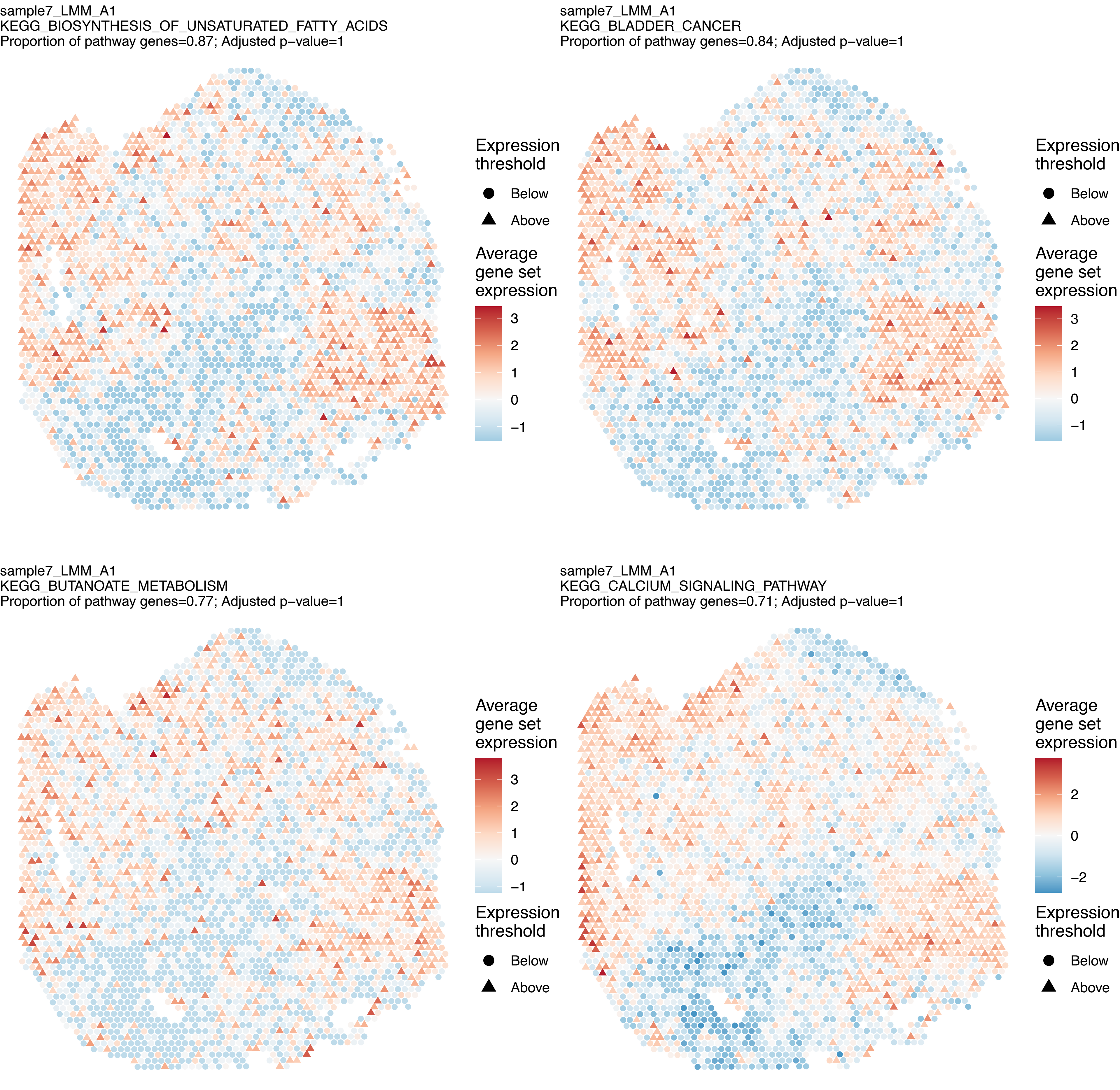

sample7_LMM_A1
KEGG_BIOSYNTHESIS_OF_UNSATURATED_FATTY_ACIDS
Proportion of pathway genes=0.87; Adjusted p–value=1
Expression threshold
Below
Above
Average gene set expression
3
2
1
0
−1
sample7_LMM_A1
KEGG_BLADDER_CANCER
Proportion of pathway genes=0.84; Adjusted p–value=1
Expression threshold
Below
Above
Average gene set expression
3
2
1
0
−1
sample7_LMM_A1
KEGG_BUTANOATE_METABOLISM
Proportion of pathway genes=0.77; Adjusted p–value=1
Average gene set expression
3
2
1
0
−1
Expression threshold
Below
Above
sample7_LMM_A1
KEGG_CALCIUM_SIGNALING_PATHWAY
Proportion of pathway genes=0.71; Adjusted p–value=1
Average gene set expression
2
0
−2
Expression threshold
Below
Above

sample7_LMM_A1
KEGG_CARDIAC_MUSCLE_CONTRACTION
Proportion of pathway genes=0.72; Adjusted p–value=1

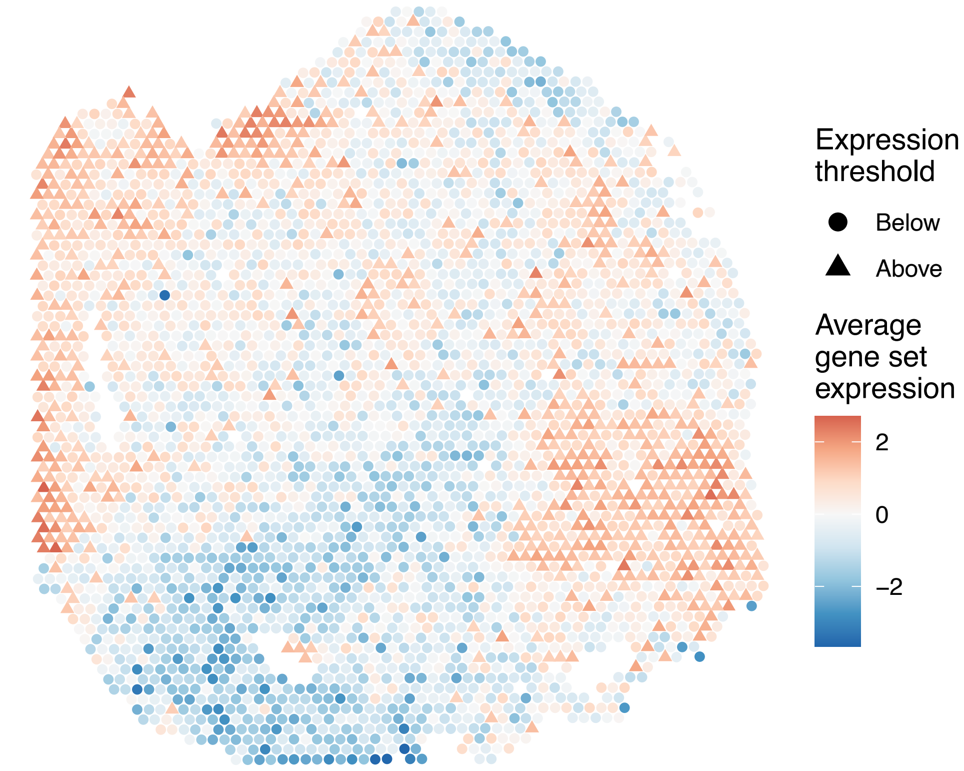


sample7_LMM_A1
KEGG_CELL_ADHESION_MOLECULES_CAMS
Proportion of pathway genes=0.8; Adjusted p–value=1

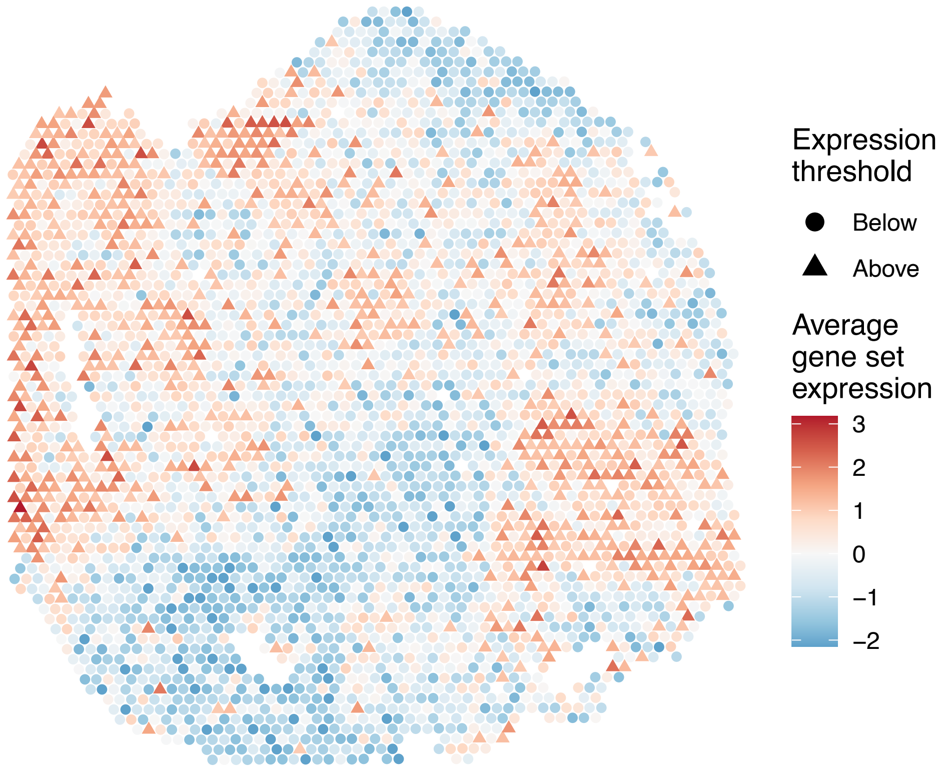


sample7_LMM_A1
KEGG_CELL_CYCLE
Proportion of pathway genes=0.94; Adjusted p–value=1

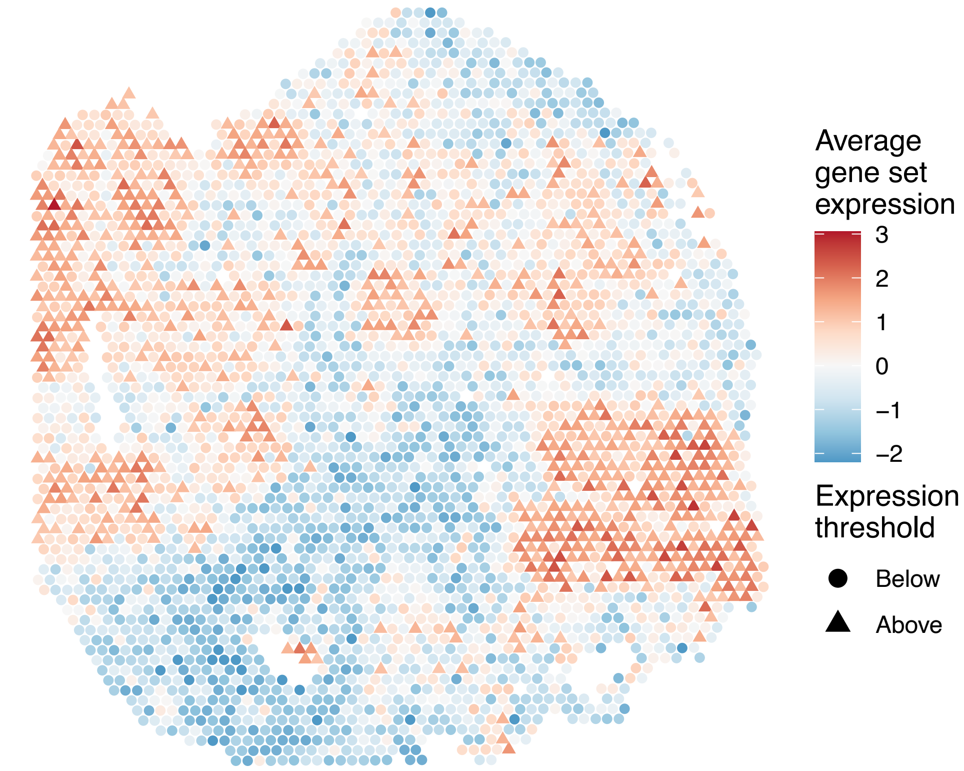


sample7_LMM_A1
KEGG_CHEMOKINE_SIGNALING_PATHWAY
Proportion of pathway genes=0.65; Adjusted p–value=1

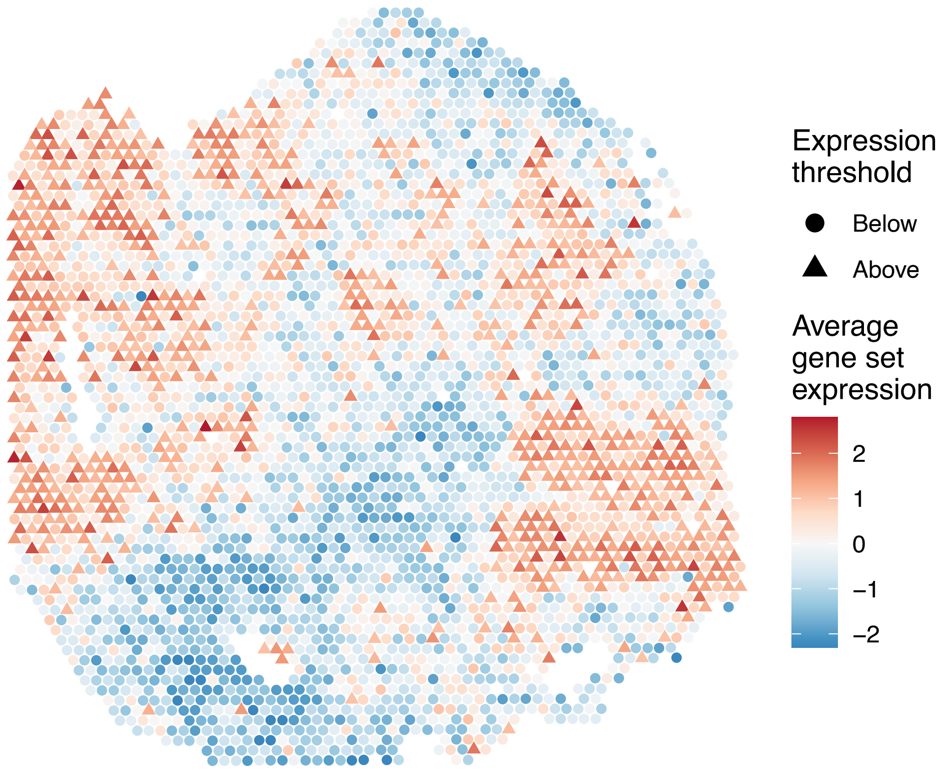

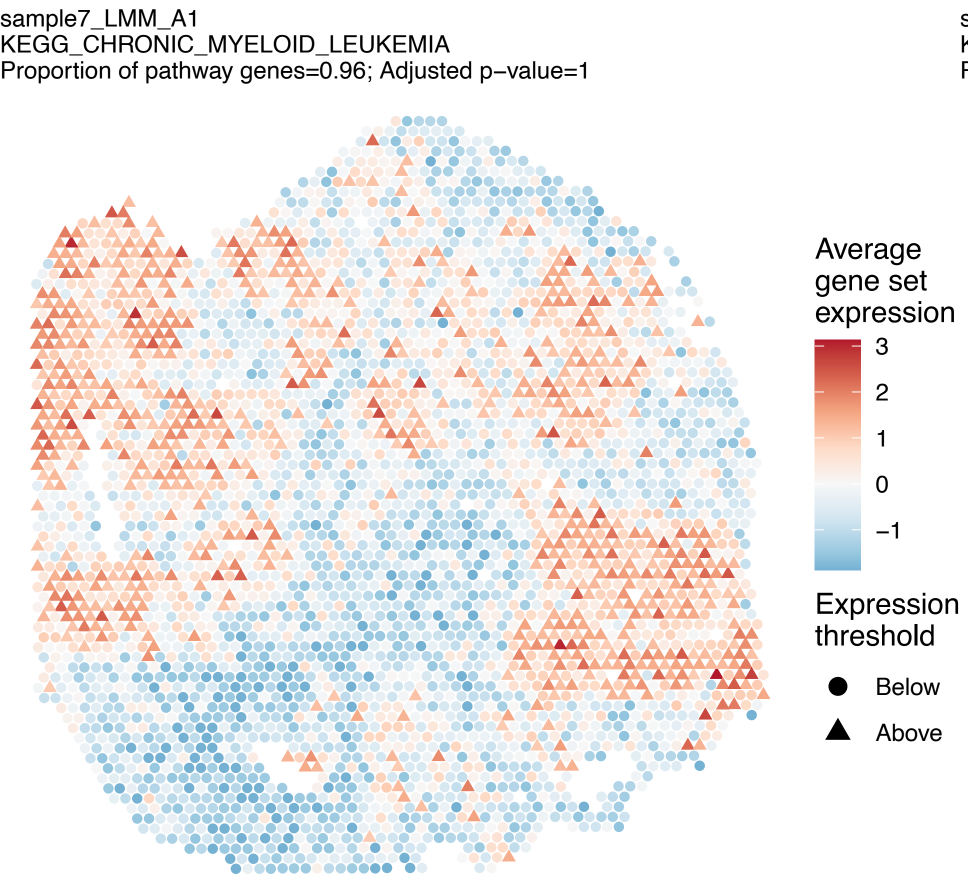
sample7_LMM_A1
KEGG_CHRONIC_MYELOID_LEUKEMIA
Proportion of pathway genes=0.96; Adjusted p−value=1
Average gene set expression
3
2
1
0
−1
Expression threshold
Below
Above

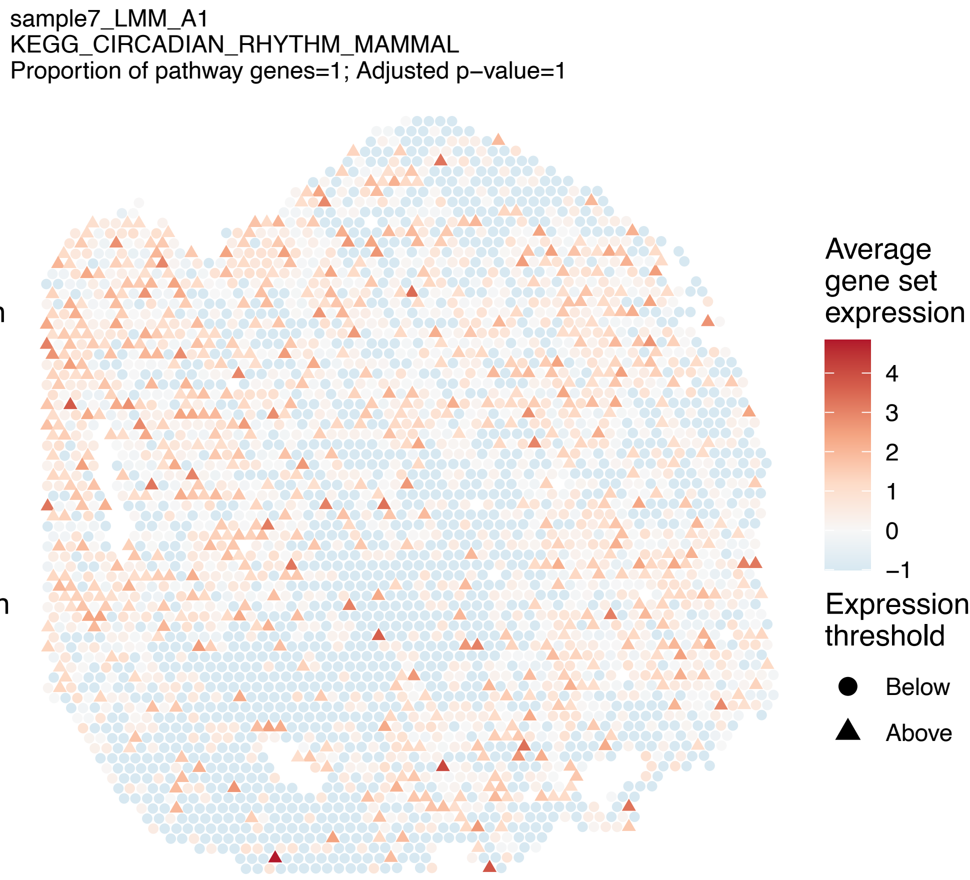
sample7_LMM_A1
KEGG_CIRCADIAN_RHYTHM_MAMMAL
Proportion of pathway genes=1; Adjusted p−value=1
Average gene set expression
4
3
2
1
0
−1
Expression threshold
Below
Above

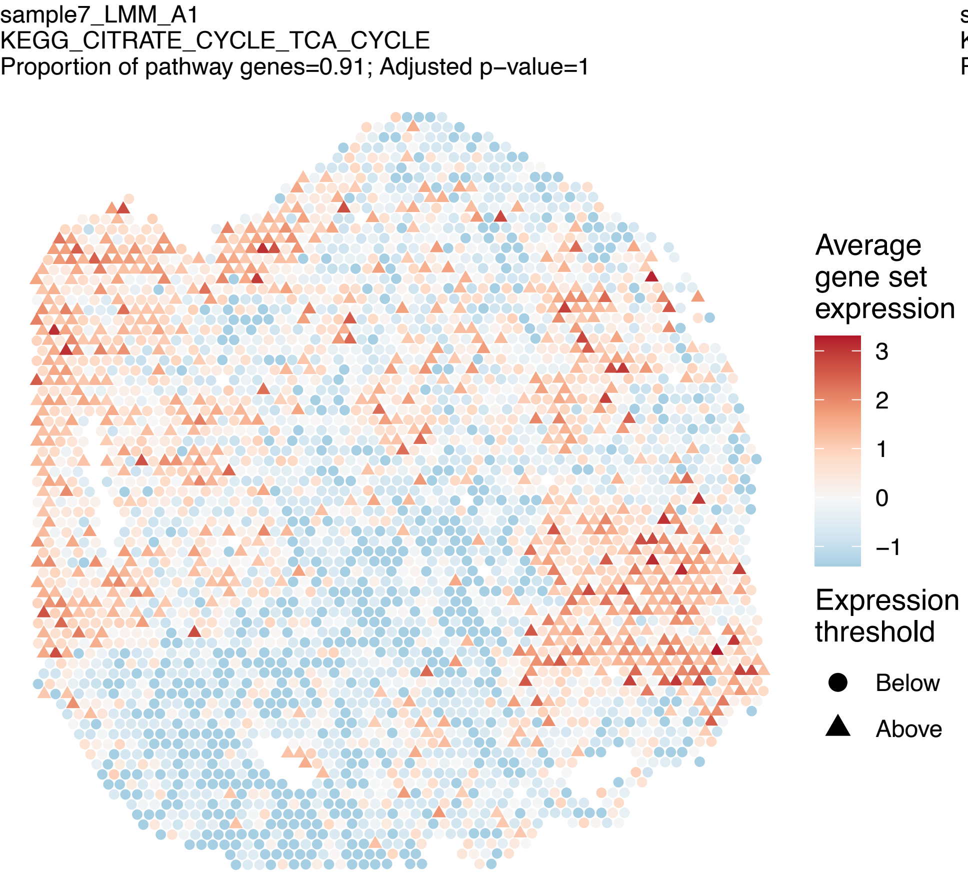
sample7_LMM_A1
KEGG_CITRATE_CYCLE_TCA_CYCLE
Proportion of pathway genes=0.91; Adjusted p−value=1
Average gene set expression
3
2
1
0
−1
Expression threshold
Below
Above

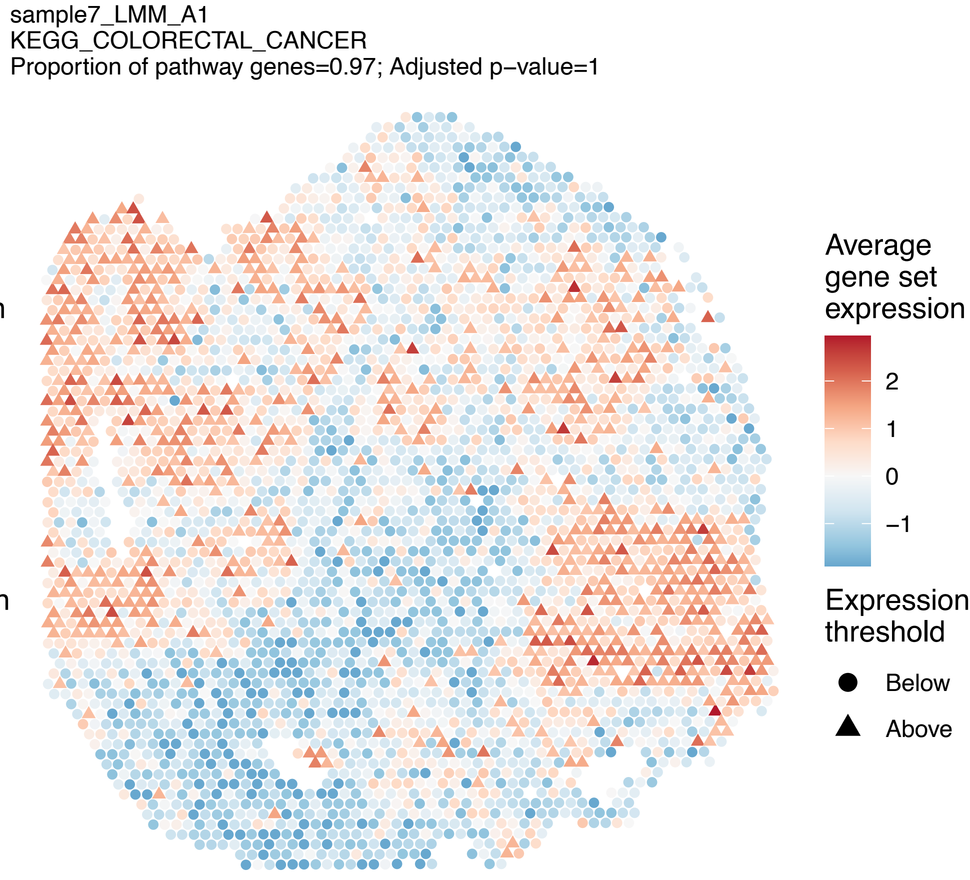
sample7_LMM_A1
KEGG_COLORECTAL_CANCER
Proportion of pathway genes=0.97; Adjusted p−value=1
Average gene set expression
2
1
0
−1
Expression threshold
Below
Above

sample7_LMM_A1
KEGG_COMPLEMENT_AND_COAGULATION_CASCADES
Proportion of pathway genes=0.59; Adjusted p−value=1

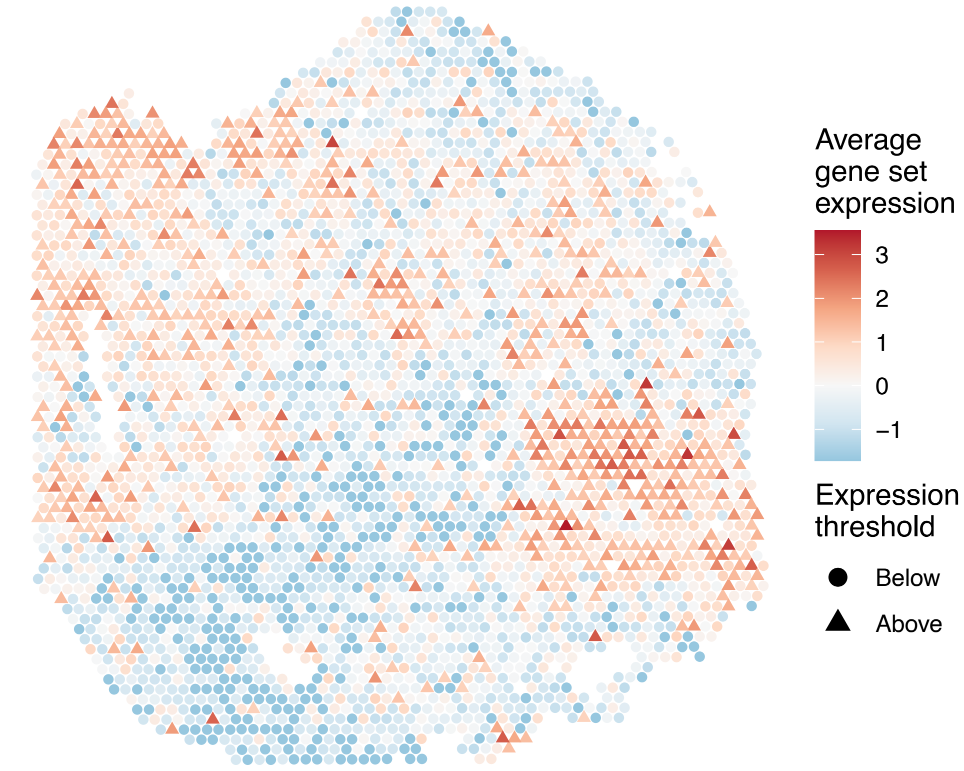


sample7_LMM_A1
KEGG_CYSTEINE_AND_METHIONINE_METABOLISM
Proportion of pathway genes=0.75; Adjusted p−value=1

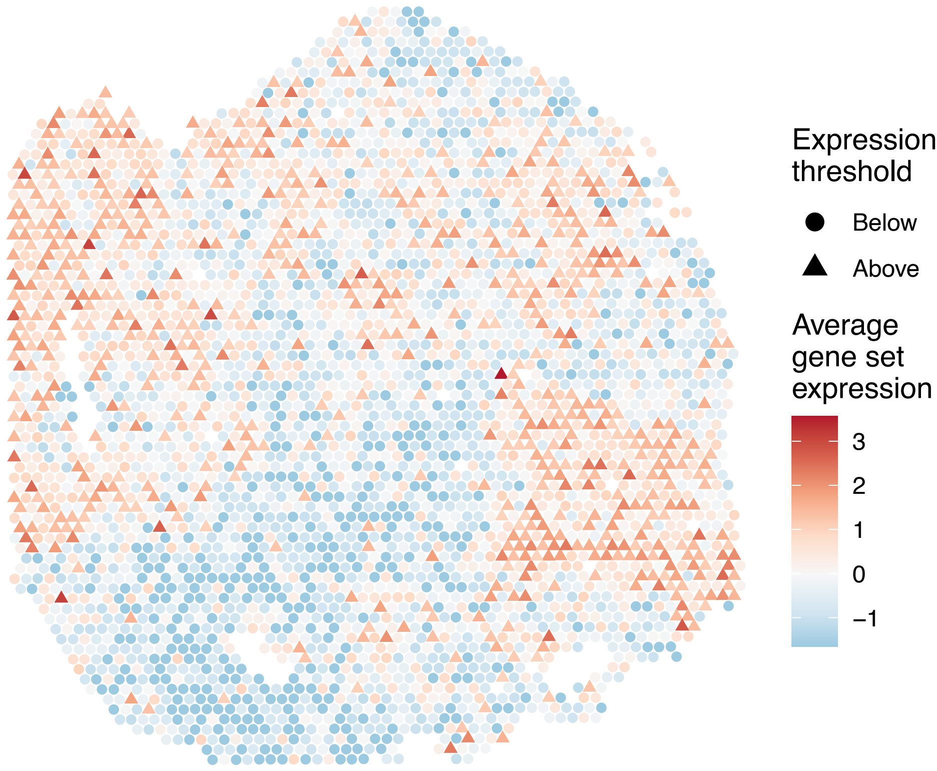


sample7_LMM_A1
KEGG_CYTOSOLIC_DNA_SENSING_PATHWAY
Proportion of pathway genes=0.49; Adjusted p−value=1

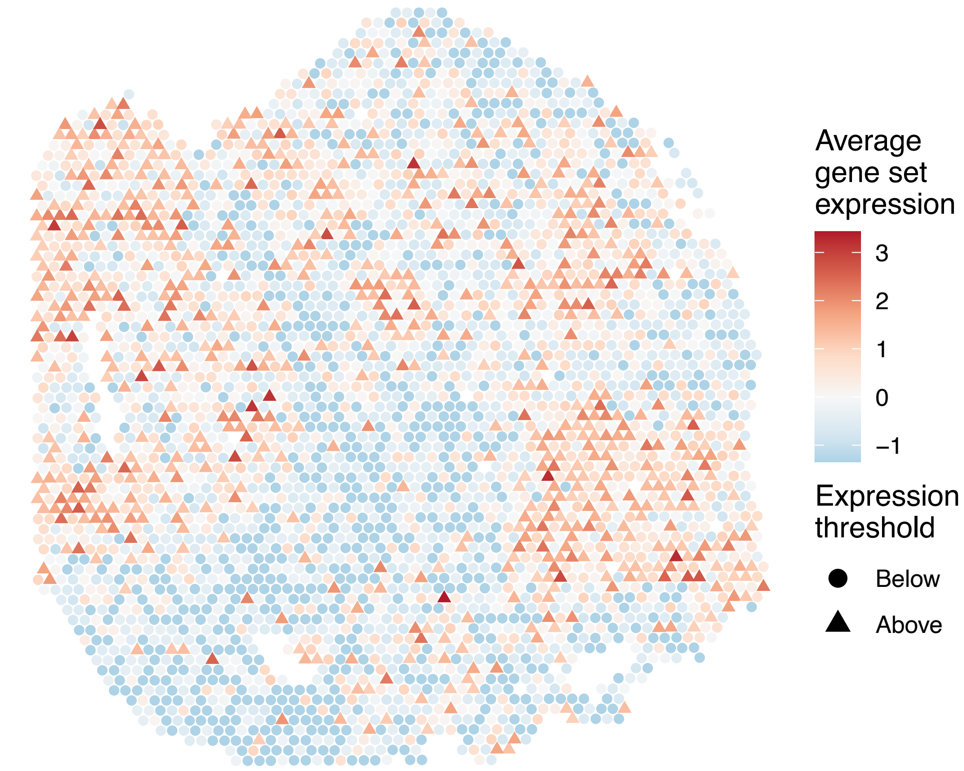


sample7_LMM_A1
KEGG_DILATED_CARDIOMYOPATHY
Proportion of pathway genes=0.73; Adjusted p−value=1

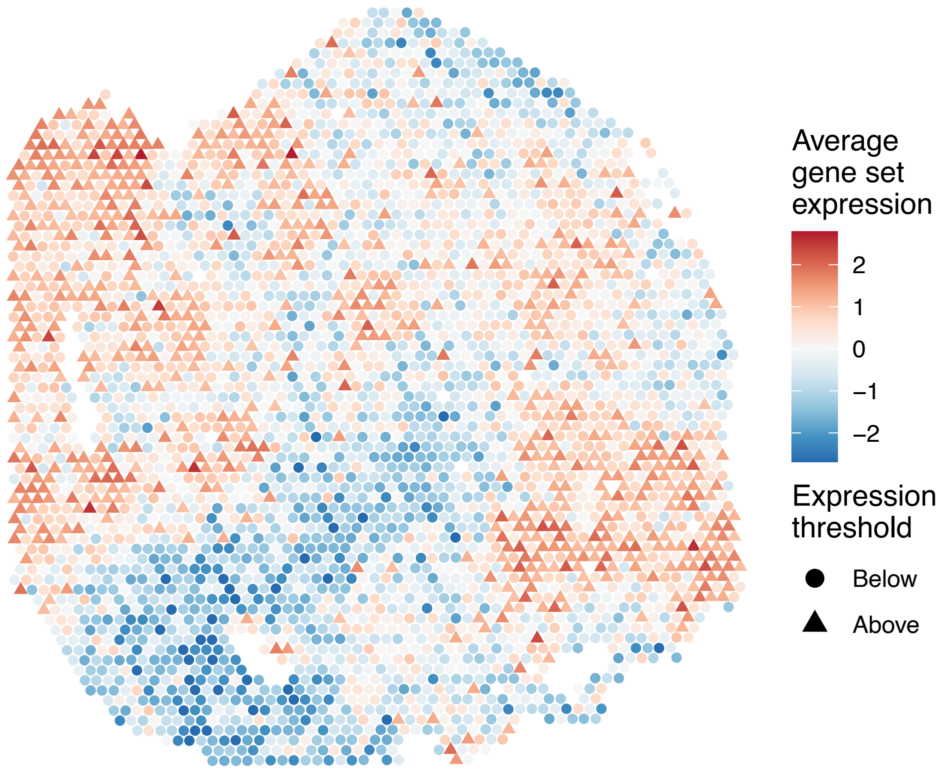

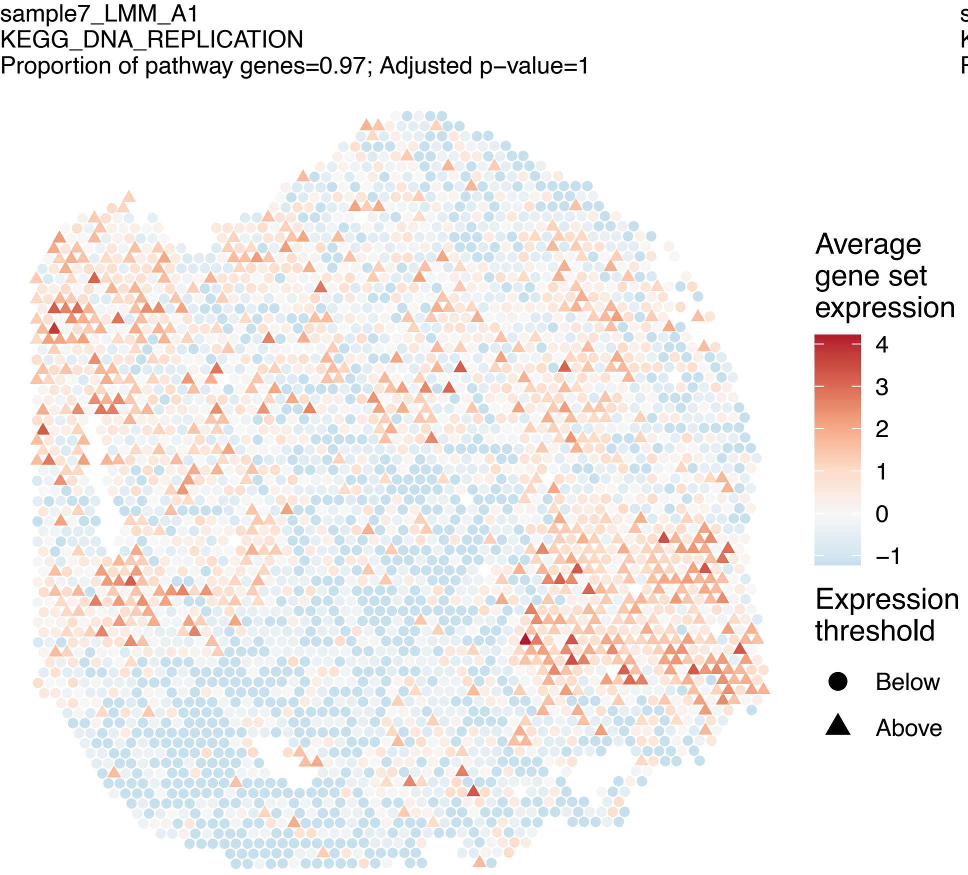
sample7_LMM_A1
KEGG_DNA_REPLICATION
Proportion of pathway genes=0.97; Adjusted p–value=1
Average gene set expression
4
3
2
1
0
−1
Expression threshold
Below
Above

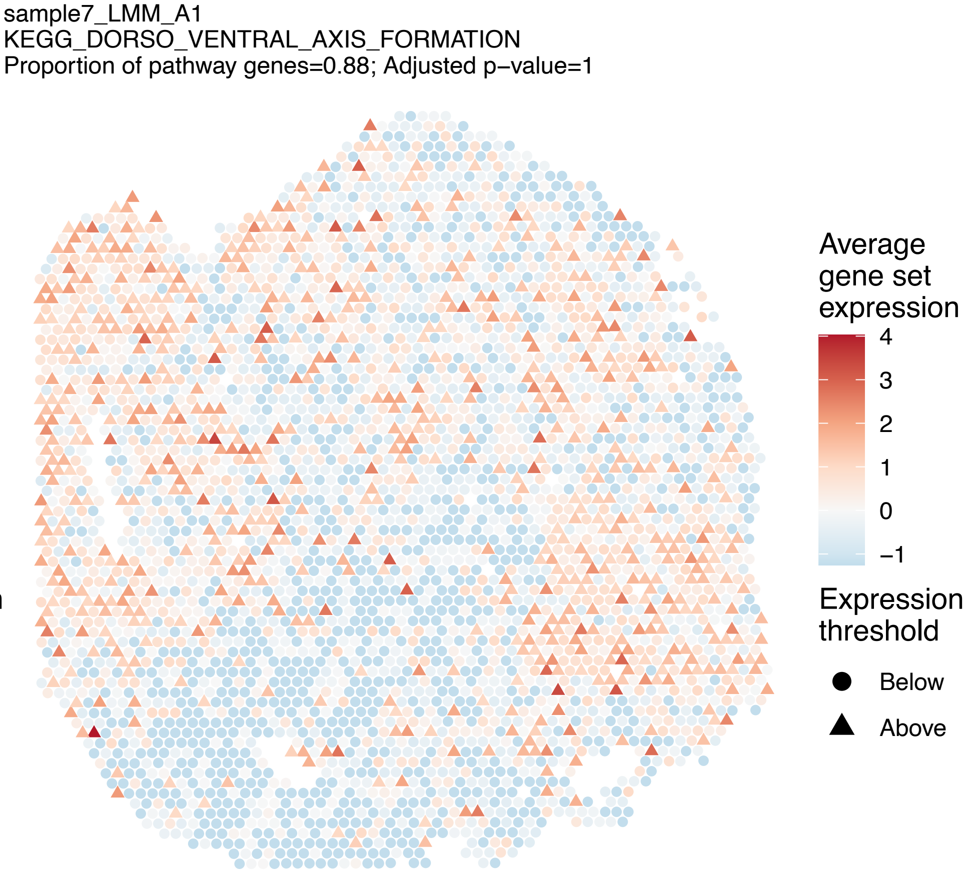
sample7_LMM_A1
KEGG_DORSO_VENTRAL_AXIS_FORMATION
Proportion of pathway genes=0.88; Adjusted p–value=1
Average gene set expression
4
3
2
1
0
−1
Expression threshold
Below
Above

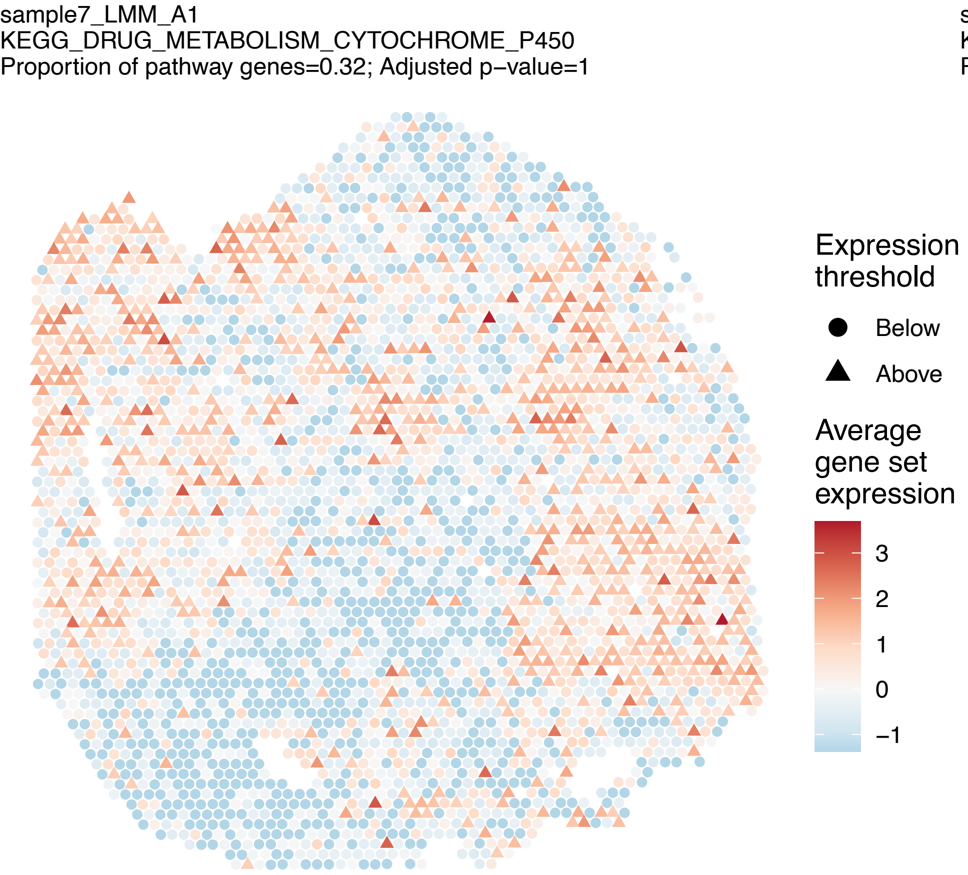
sample7_LMM_A1
KEGG_DRUG_METABOLISM_CYTOCHROME_P450
Proportion of pathway genes=0.32; Adjusted p–value=1
Expression threshold
Below
Above
Average gene set expression
3
2
1
0
−1

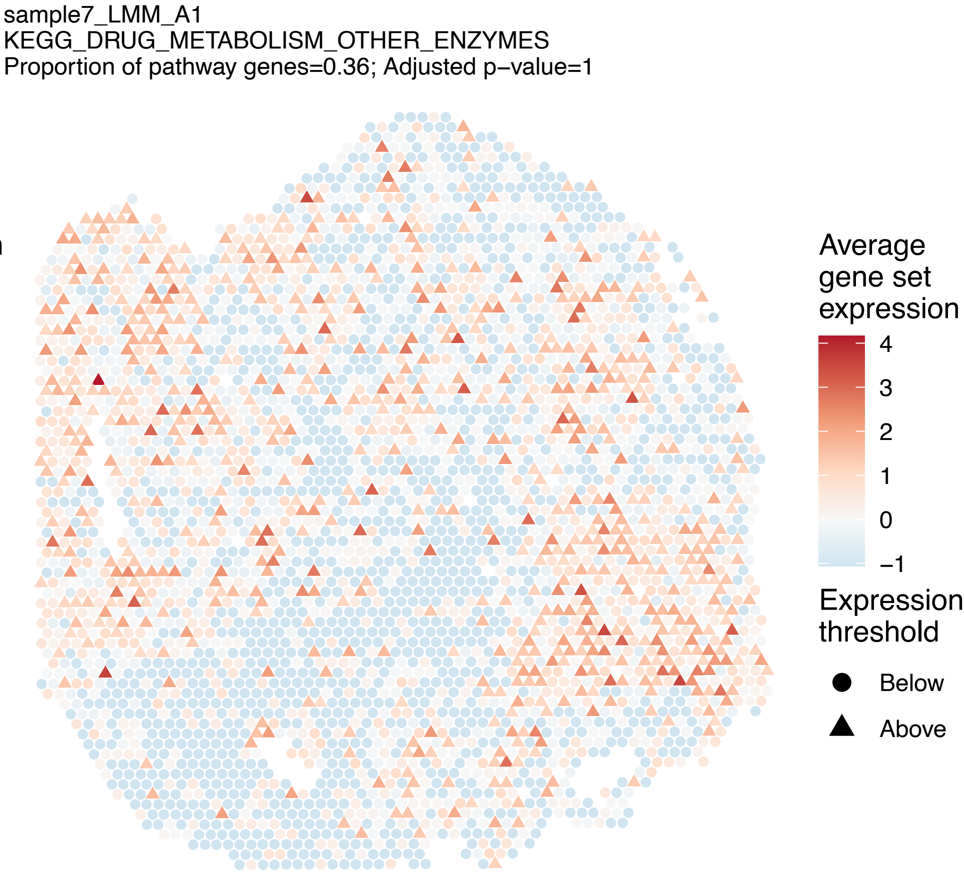
sample7_LMM_A1
KEGG_DRUG_METABOLISM_OTHER_ENZYMES
Proportion of pathway genes=0.36; Adjusted p–value=1
Average gene set expression
4
3
2
1
0
−1
Expression threshold
Below
Above

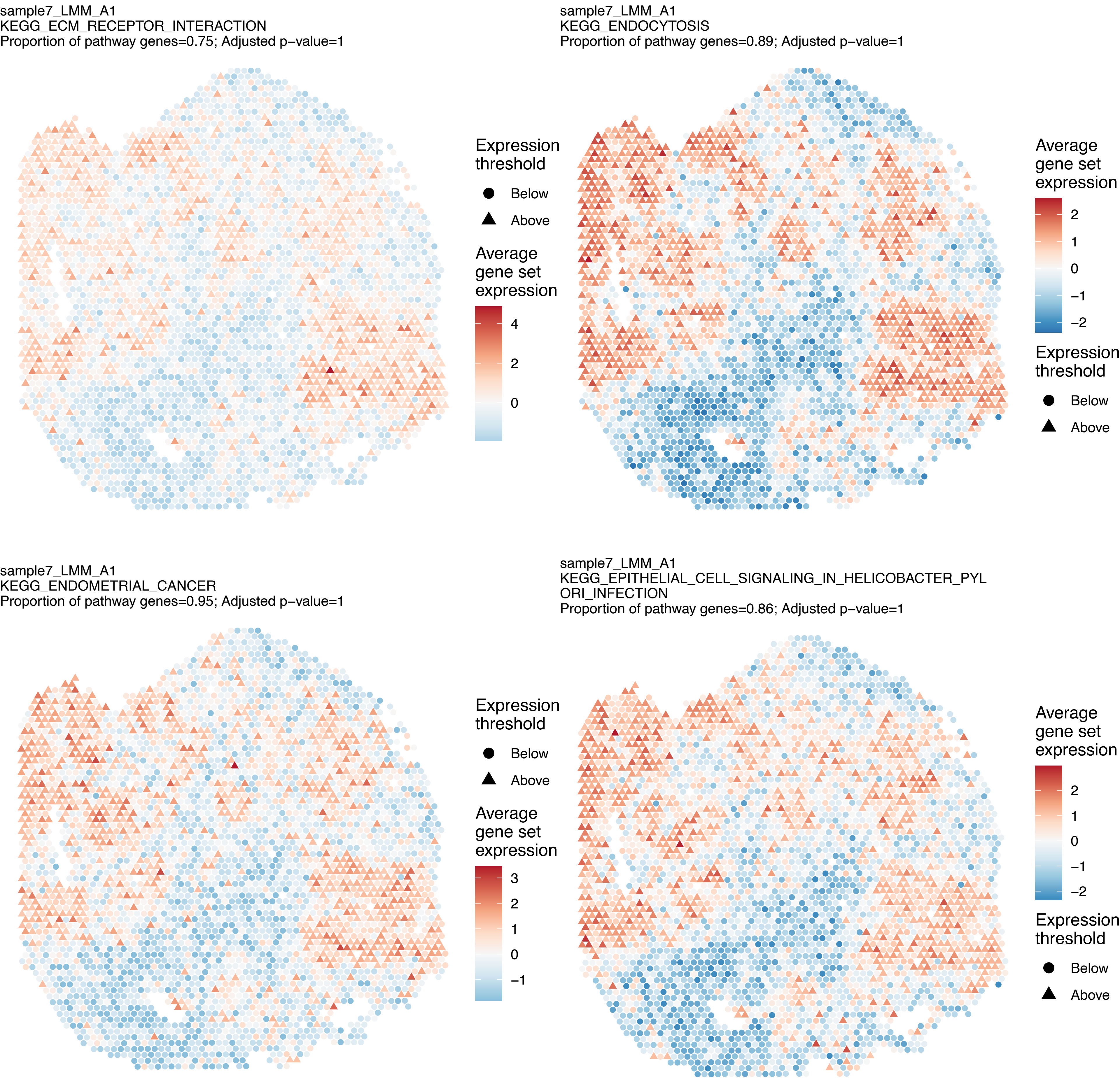

sample7_LMM_A1
KEGG_ECM_RECEPTOR_INTERACTION
Proportion of pathway genes=0.75; Adjusted p–value=1
Expression threshold
Below
Above
Average gene set expression
4
2
0
sample7_LMM_A1
KEGG_ENDOCYTOSIS
Proportion of pathway genes=0.89; Adjusted p–value=1
Average gene set expression
2
1
0
−1
−2
Expression threshold
Below
Above
sample7_LMM_A1
KEGG_ENDOMETRIAL_CANCER
Proportion of pathway genes=0.95; Adjusted p–value=1
Expression threshold
Below
Above
Average gene set expression
3
2
1
0
−1
sample7_LMM_A1
KEGG_EPITHELIAL_CELL_SIGNALING_IN_HELICOBACTER_PYL
ORI_INFECTION
Proportion of pathway genes=0.86; Adjusted p–value=1
Average gene set expression
2
1
0
−1
−2
Expression threshold
Below
Above

sample7_LMM_A1
KEGG_ERBB_SIGNALING_PATHWAY
Proportion of pathway genes=0.92; Adjusted p−value=1

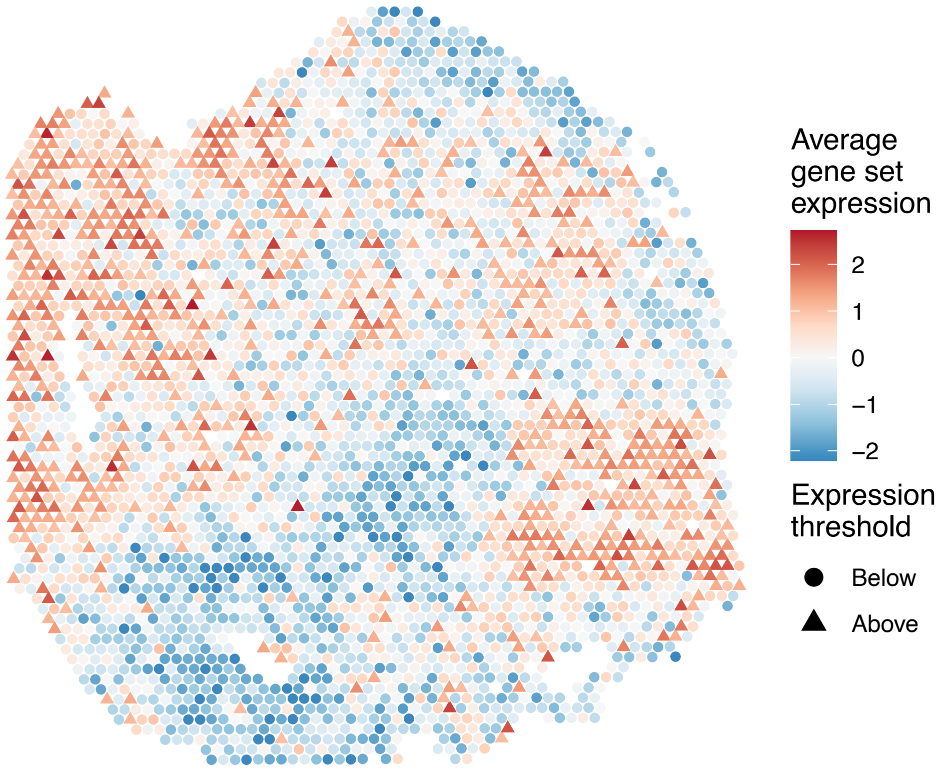


sample7_LMM_A1
KEGG_ETHER_LIPID_METABOLISM
Proportion of pathway genes=0.63; Adjusted p−value=1

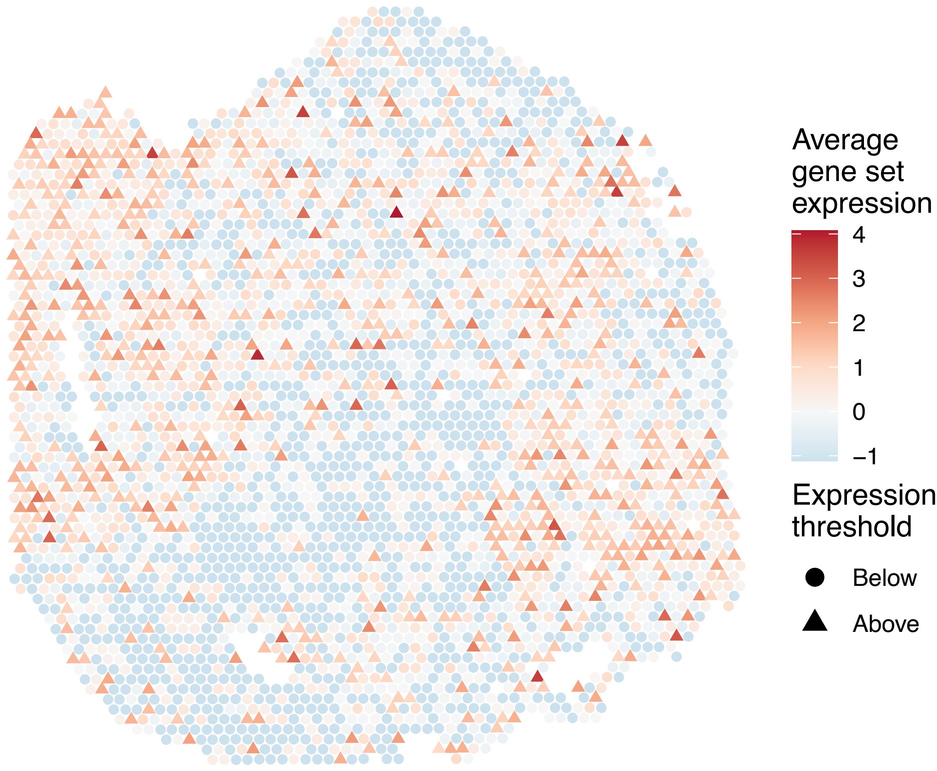


sample7_LMM_A1
KEGG_FATTY_ACID_METABOLISM
Proportion of pathway genes=0.79; Adjusted p−value=1

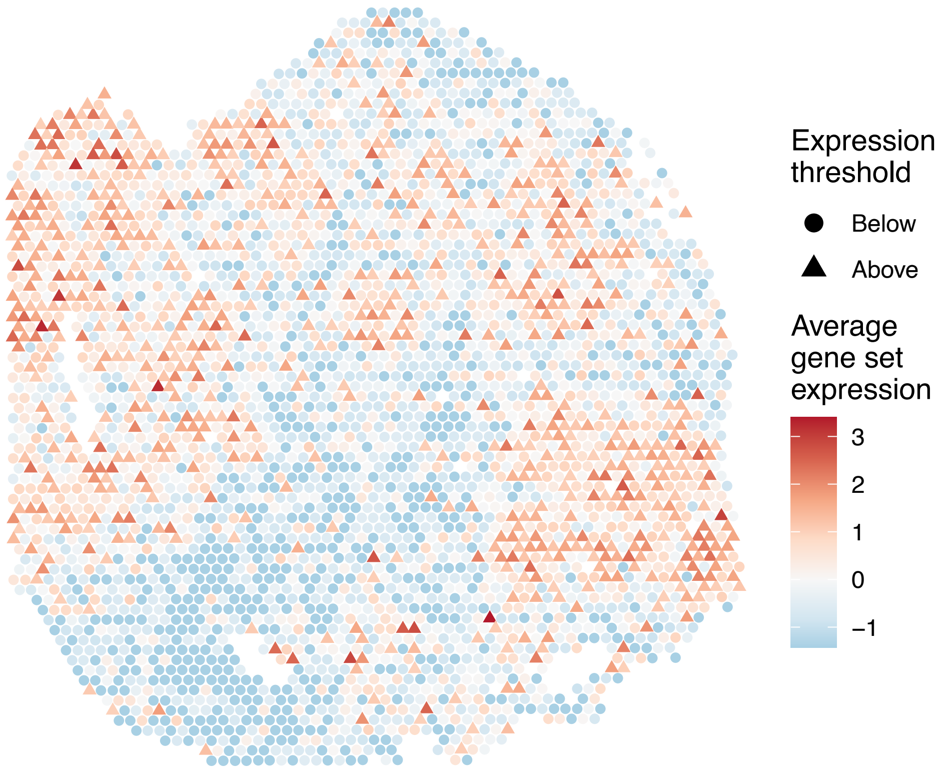


sample7_LMM_A1
KEGG_FC_EPSILON_RI_SIGNALING_PATHWAY
Proportion of pathway genes=0.68; Adjusted p−value=1

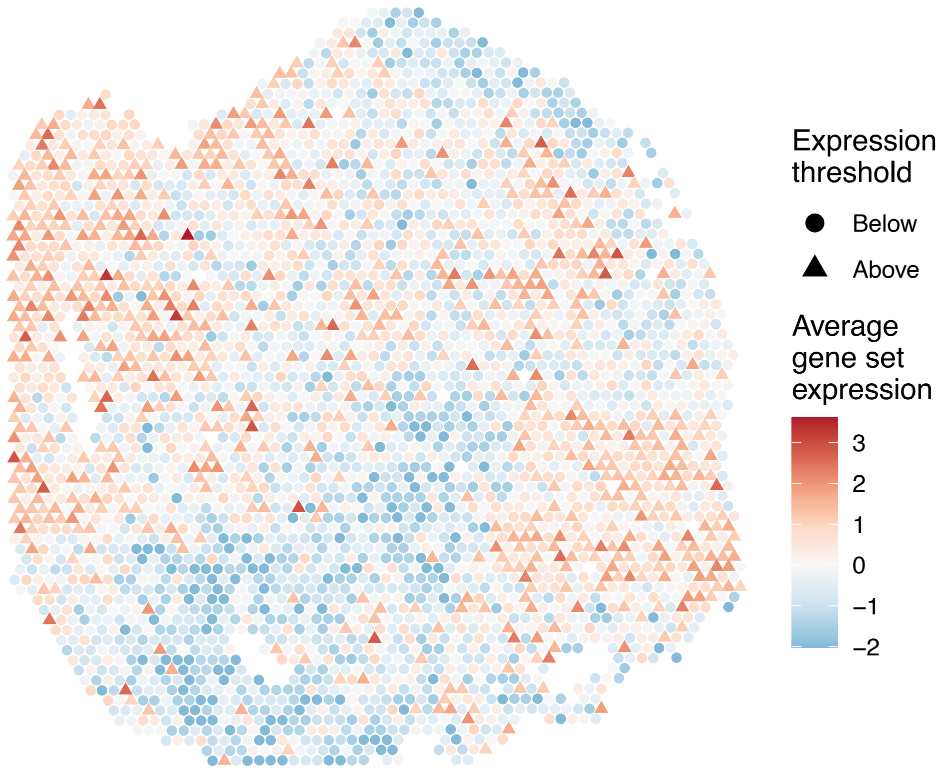

sample7_LMM_A1
KEGG_FC_GAMMA_R_MEDIATED_PHAGOCYTOSIS
Proportion of pathway genes=0.95; Adjusted p−value=1

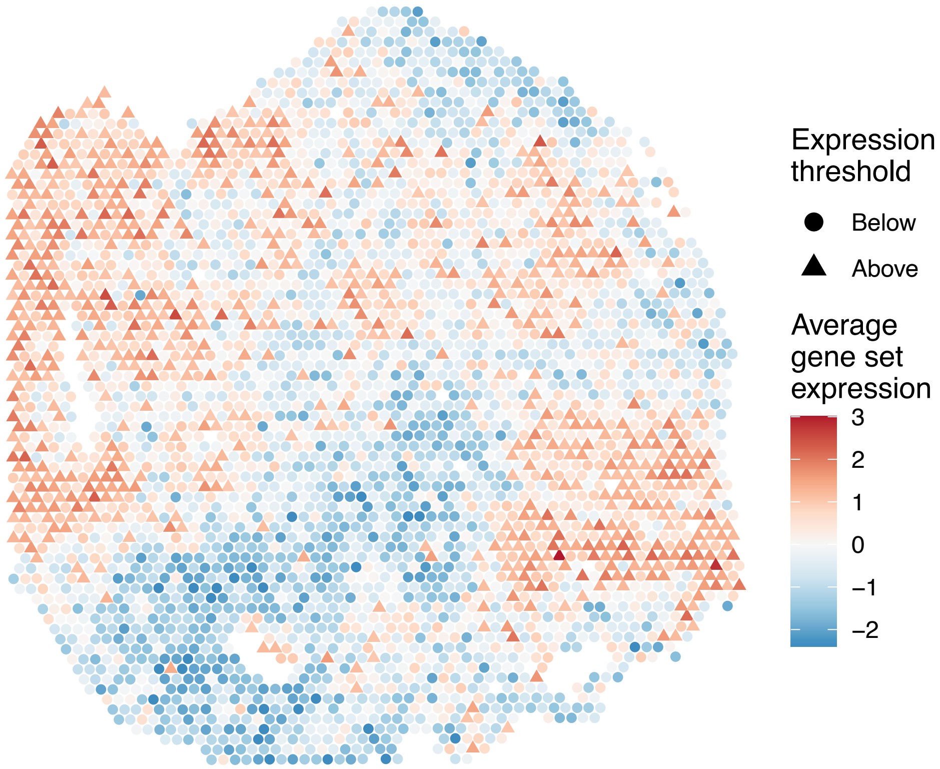


sample7_LMM_A1
KEGG_FOCAL_ADHESION
Proportion of pathway genes=0.9; Adjusted p−value=1

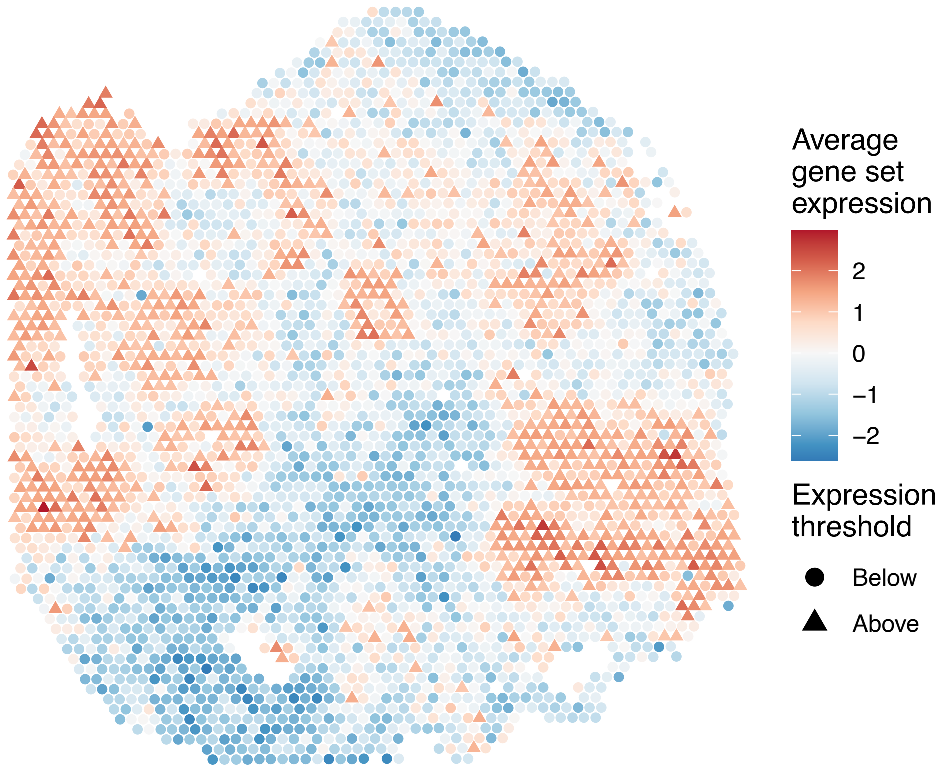


sample7_LMM_A1
KEGG_FRUCTOSE_AND_MANNOSE_METABOLISM
Proportion of pathway genes=0.74; Adjusted p−value=1

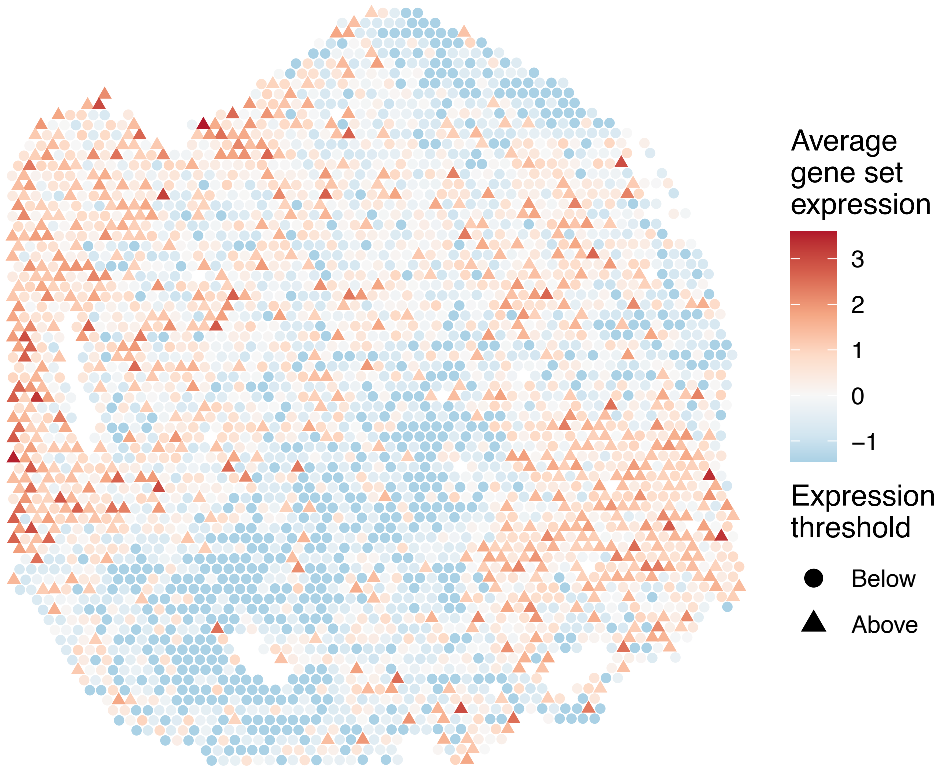


sample7_LMM_A1
KEGG_GALACTOSE_METABOLISM
Proportion of pathway genes=0.69; Adjusted p−value=1

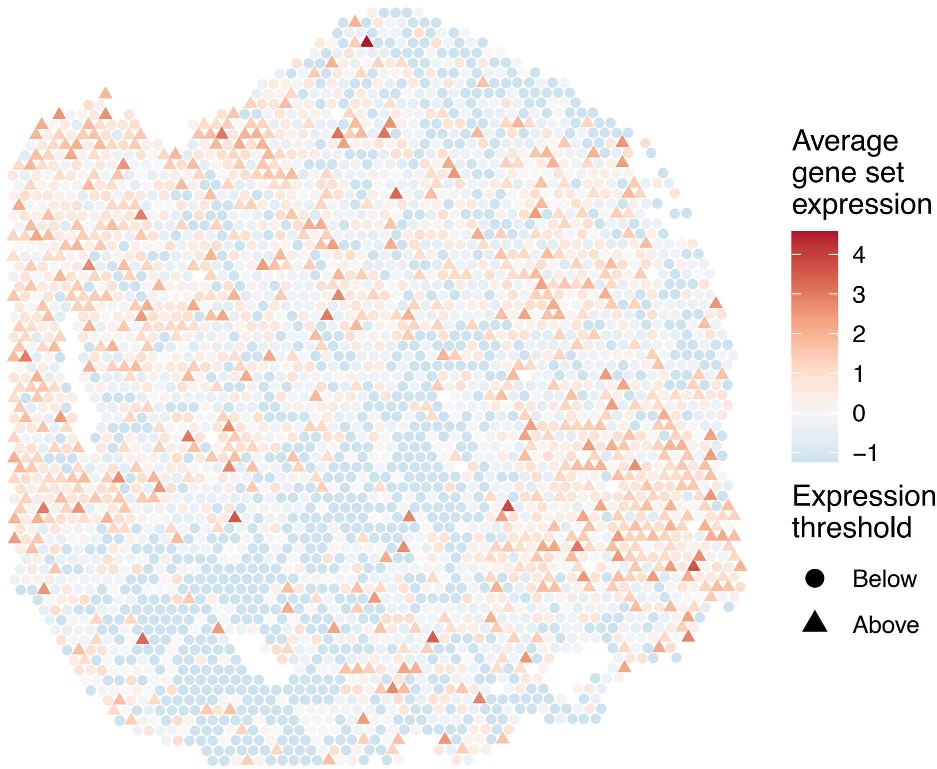

sample7_LMM_A1
KEGG_GAP_JUNCTION
Proportion of pathway genes=0.87; Adjusted p−value=1

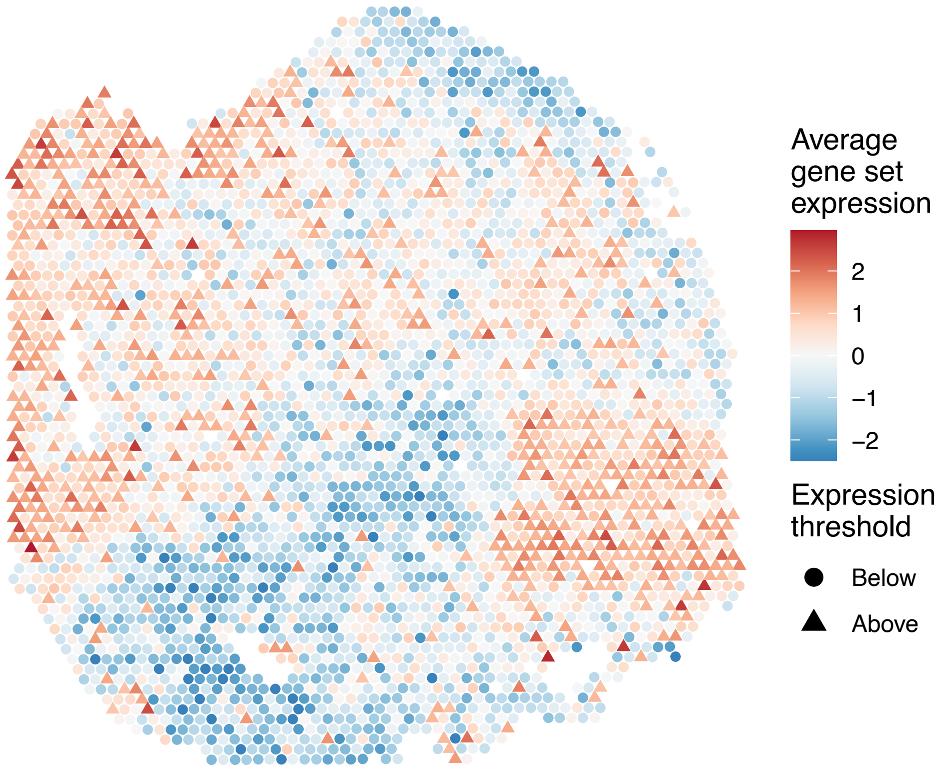


sample7_LMM_A1
KEGG_GLIOMA
Proportion of pathway genes=0.91; Adjusted p−value=1

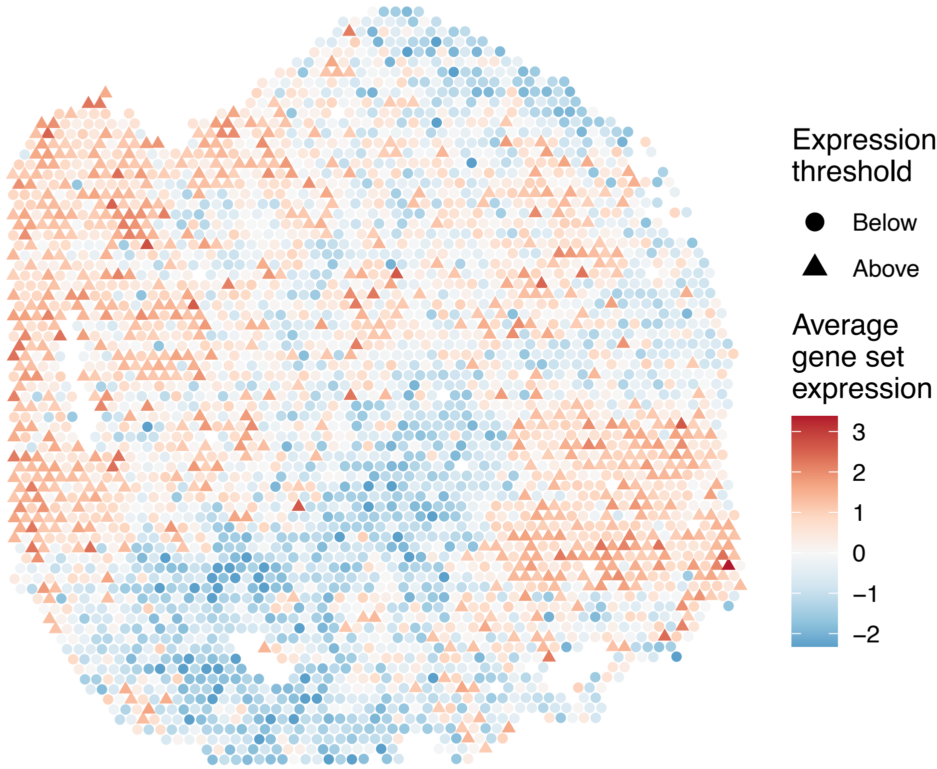


sample7_LMM_A1
KEGG_GLUTATHIONE_METABOLISM
Proportion of pathway genes=0.68; Adjusted p−value=1

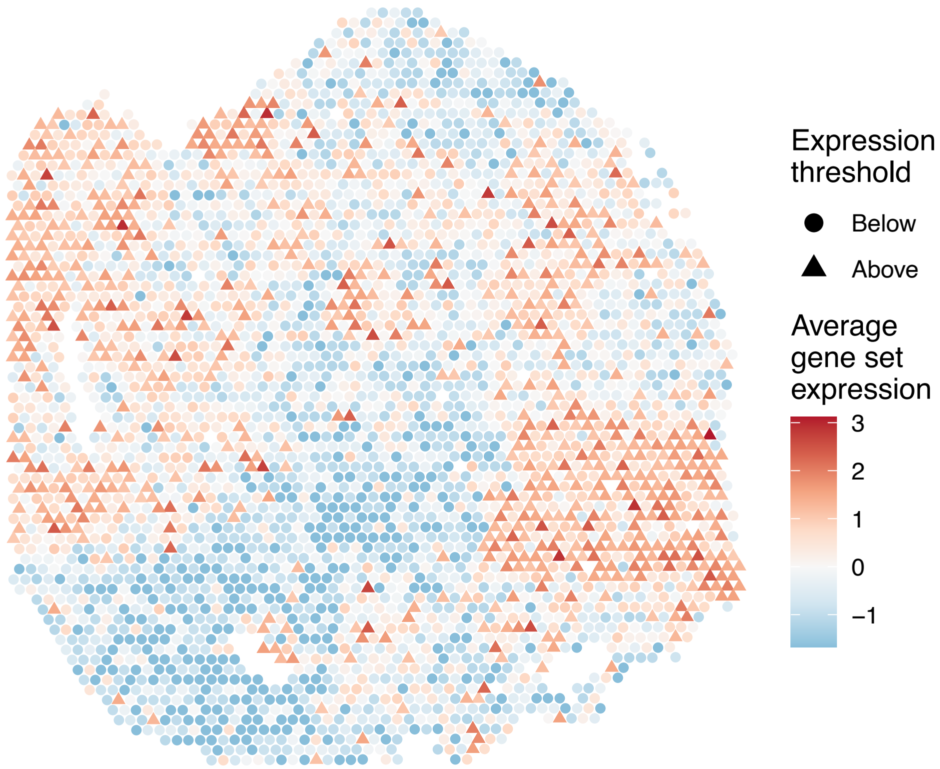


sample7_LMM_A1
KEGG_GLYCEROLIPID_METABOLISM
Proportion of pathway genes=0.83; Adjusted p−value=1

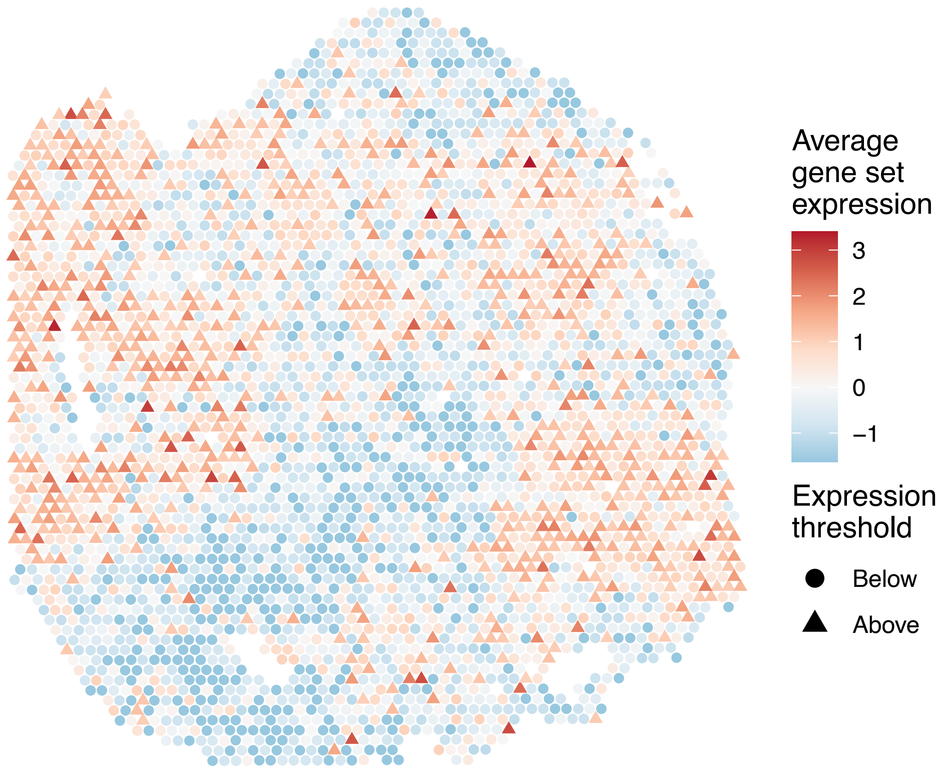

sample7_LMM_A1
KEGG_GLYCEROPHOSPHOLIPID_METABOLISM
Proportion of pathway genes=0.82; Adjusted p−value=1

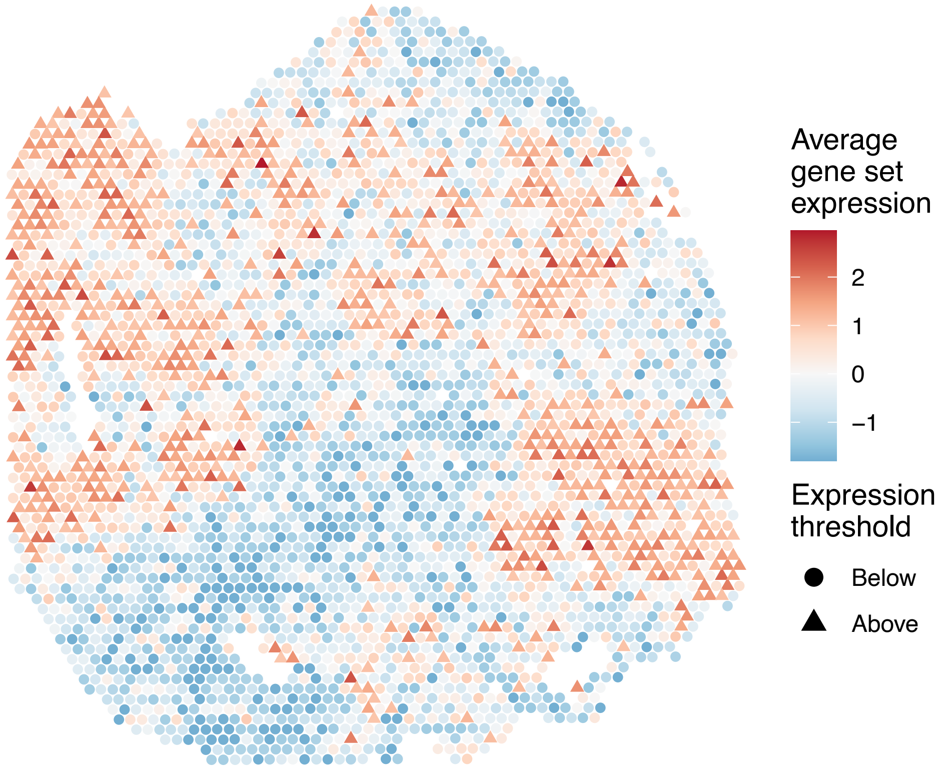


sample7_LMM_A1
KEGG_GLYCINE_SERINE_AND_THREONINE_METABOLISM
Proportion of pathway genes=0.72; Adjusted p−value=1

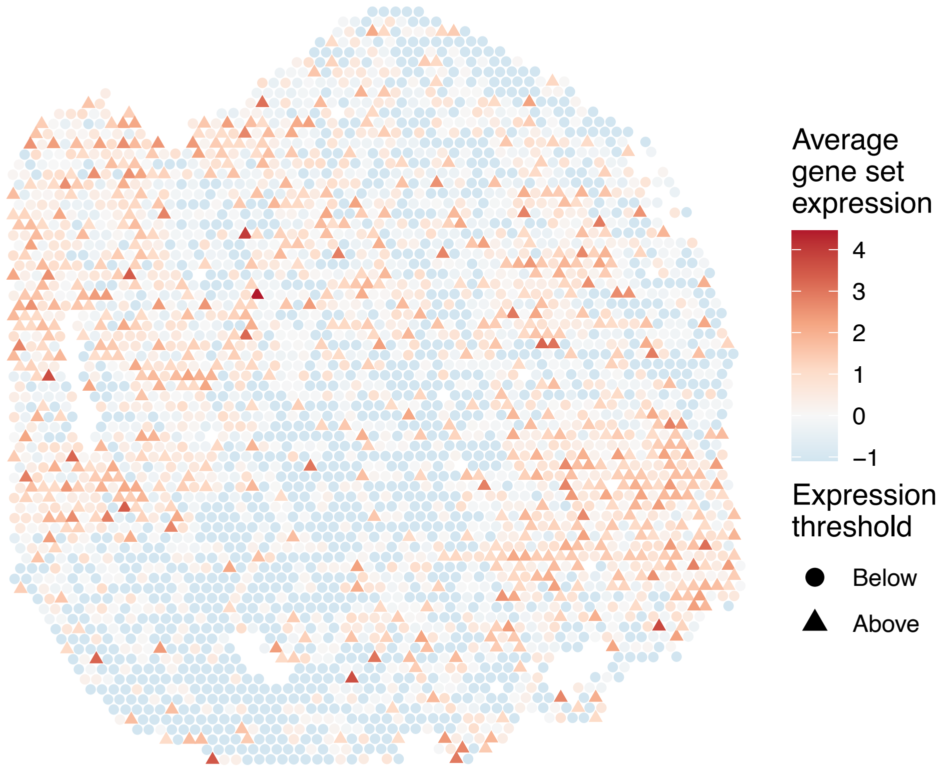


sample7_LMM_A1
KEGG_GLYCOLYSIS_GLUCONEOGENESIS
Proportion of pathway genes=0.69; Adjusted p−value=1

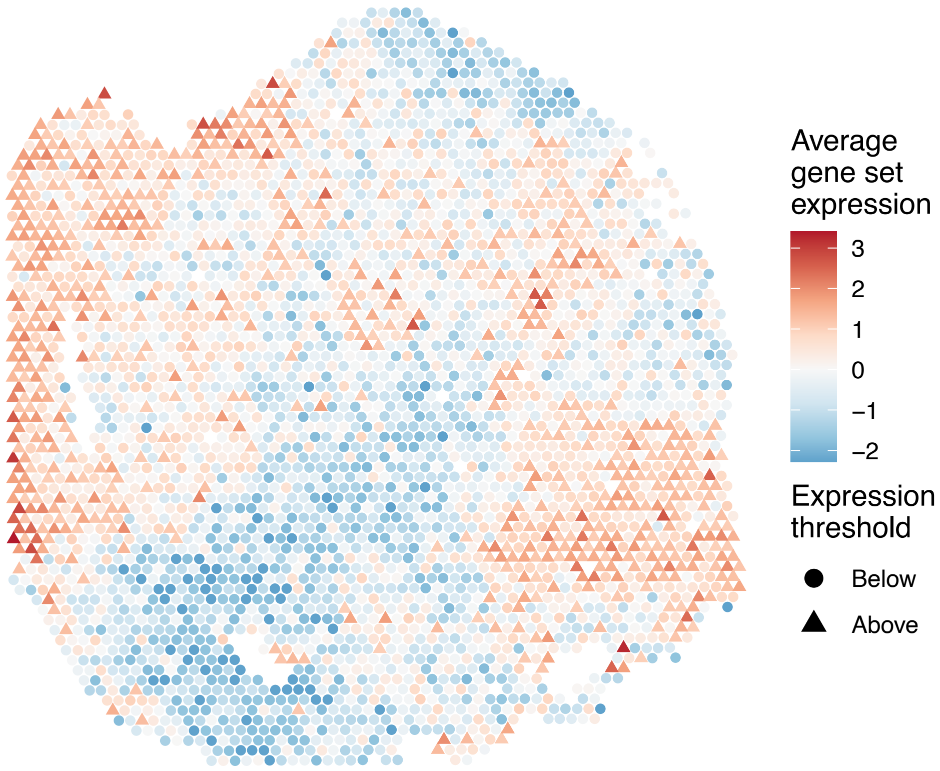


sample7_LMM_A1
KEGG_GLYCOSAMINOGLYCAN_BIOSYNTHESIS_CHONDROITIN_SU
LFATE
Proportion of pathway genes=0.96; Adjusted p−value=1

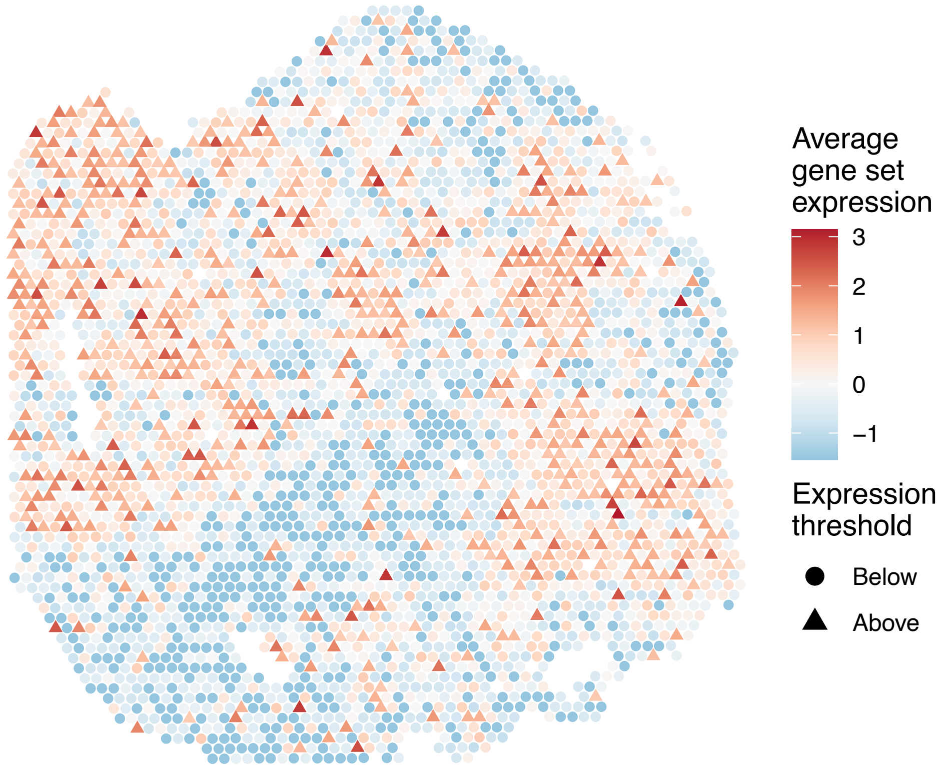

sample7_LMM_A1
KEGG_GLYCOSAMINOGLYCAN_BIOSYNTHESIS_HEPARAN_SULFAT
E
Proportion of pathway genes=0.89; Adjusted p−value=1

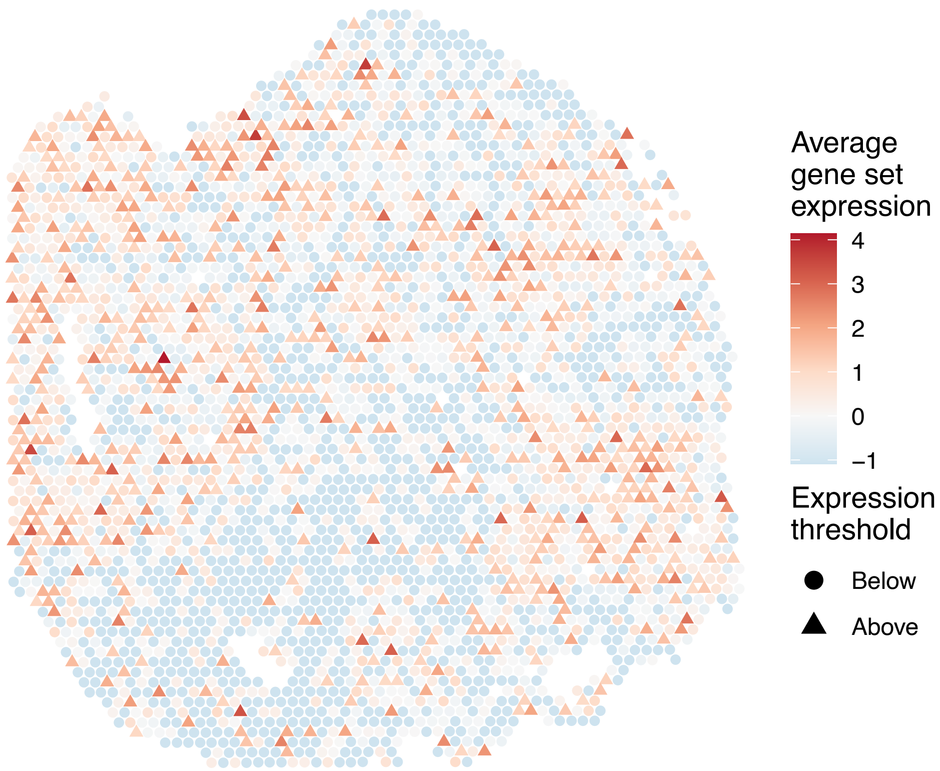


sample7_LMM_A1
KEGG_GLYCOSAMINOGLYCAN_BIOSYNTHESIS_KERATAN_SULFAT
E
Proportion of pathway genes=0.87; Adjusted p−value=1

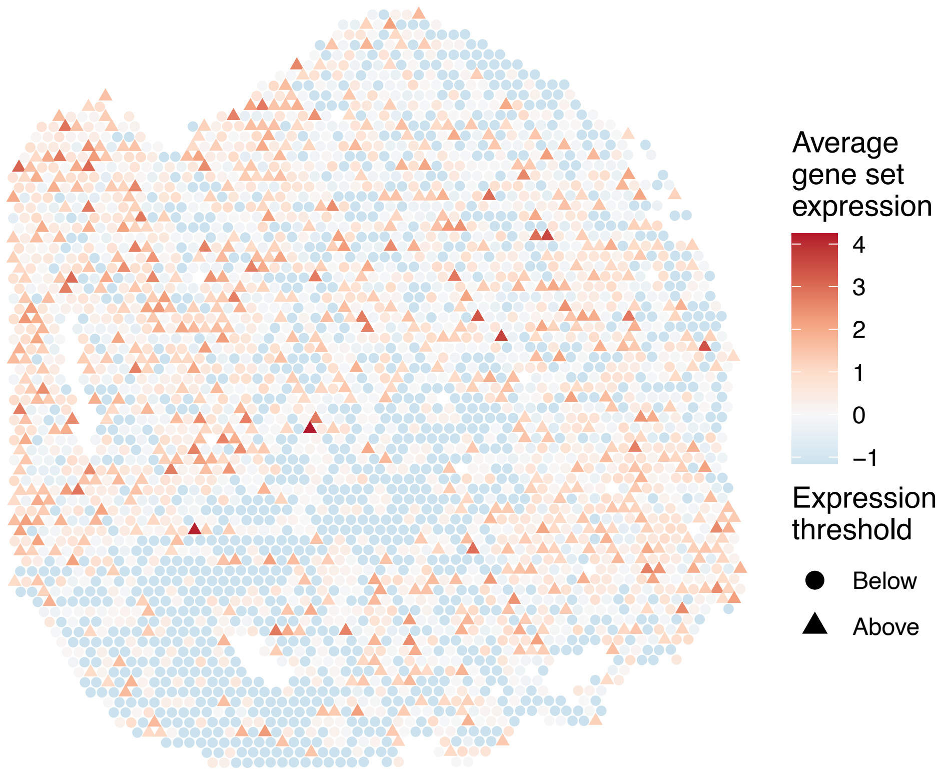


sample7_LMM_A1
KEGG_GLYCOSAMINOGLYCAN_DEGRADATION
Proportion of pathway genes=0.81; Adjusted p−value=1

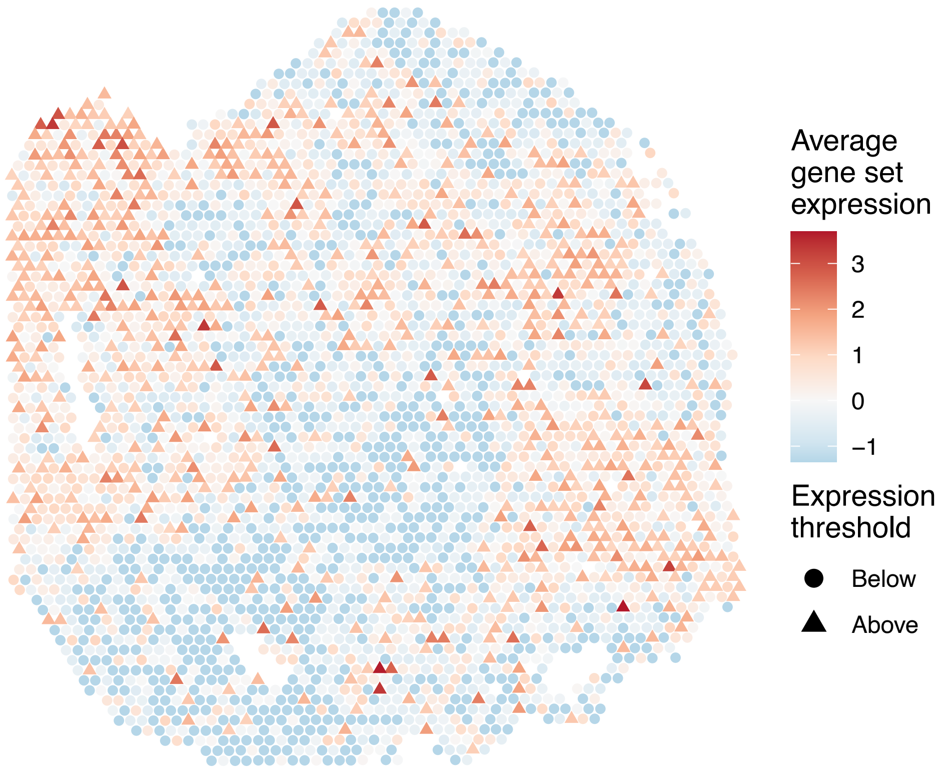


sample7_LMM_A1
KEGG_GLYCOSPHINGOLIPID_BIOSYNTHESIS_GANGLIO_SERIES
Proportion of pathway genes=1; Adjusted p−value=1

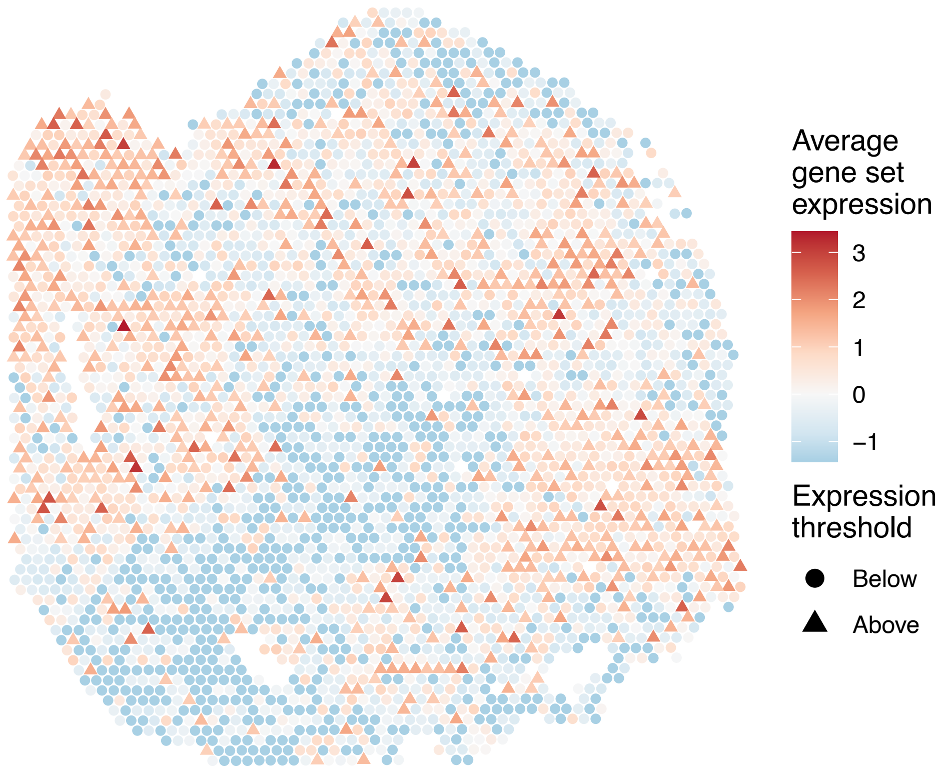

sample7_LMM_A1
KEGG_GLYCOSPHINGOLIPID_BIOSYNTHESIS_GLOBO_SERIES
Proportion of pathway genes=0.86; Adjusted p−value=1

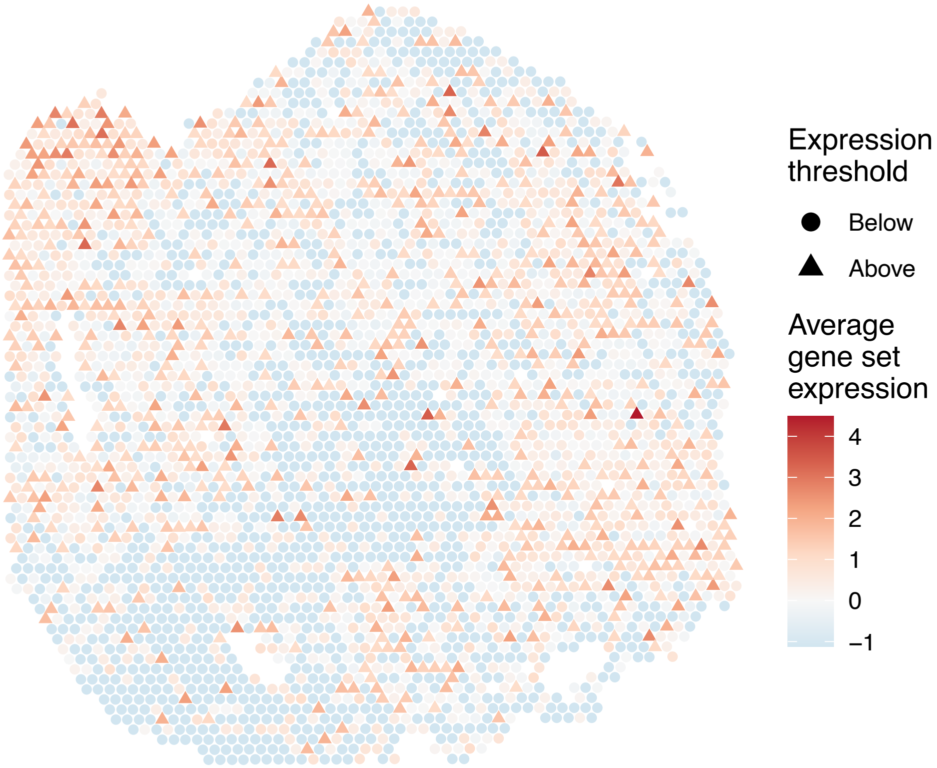


sample7_LMM_A1
KEGG_GLYCOSPHINGOLIPID_BIOSYNTHESIS_LACTO_AND_NEOLACTO_SERIES
Proportion of pathway genes=0.69; Adjusted p−value=1

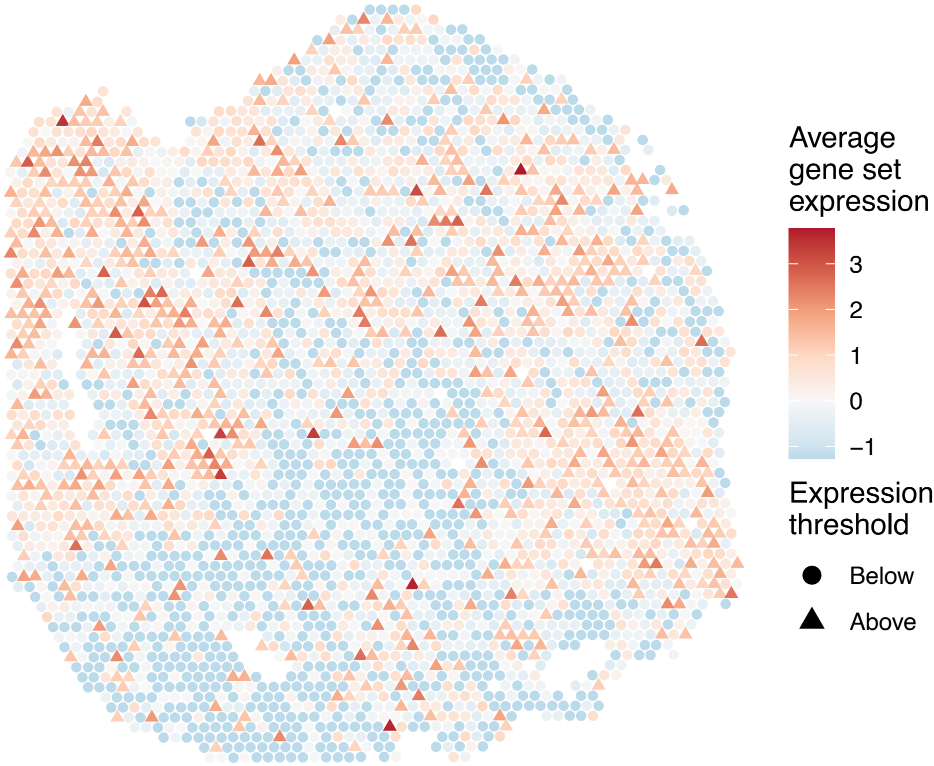


sample7_LMM_A1
KEGG_GLYCOSYLPHOSPHATIDYLINOSITOL_GPI_ANCHOR_BIOSYNTHESIS
Proportion of pathway genes=0.96; Adjusted p−value=1

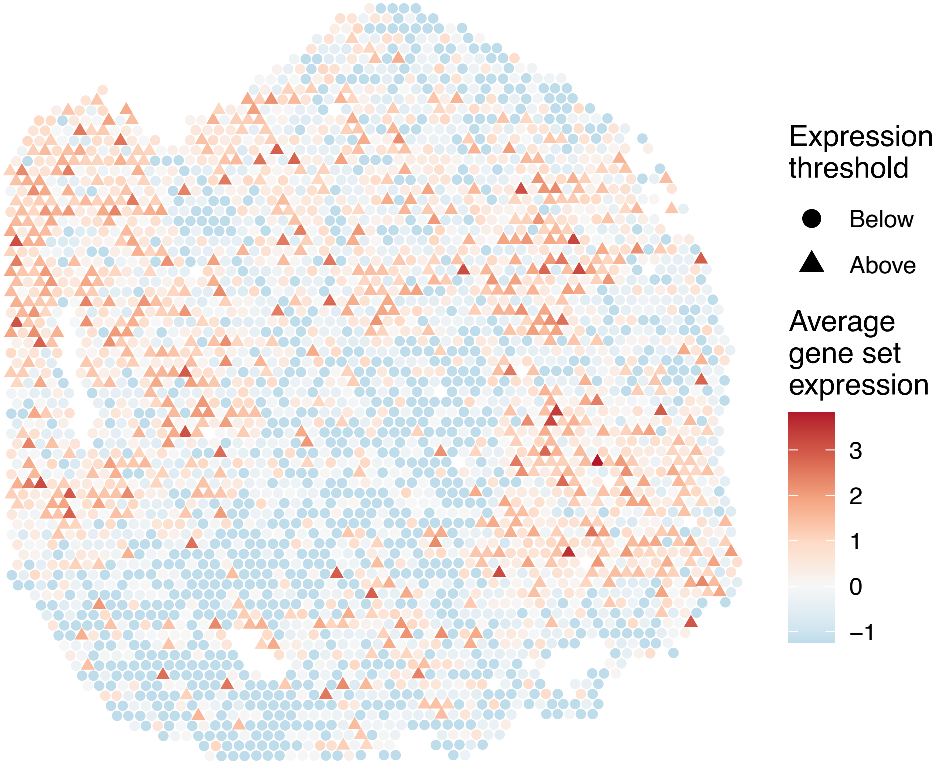


sample7_LMM_A1
KEGG_GLYOXYLATE_AND_DICARBOXYLATE_METABOLISM
Proportion of pathway genes=0.88; Adjusted p−value=1

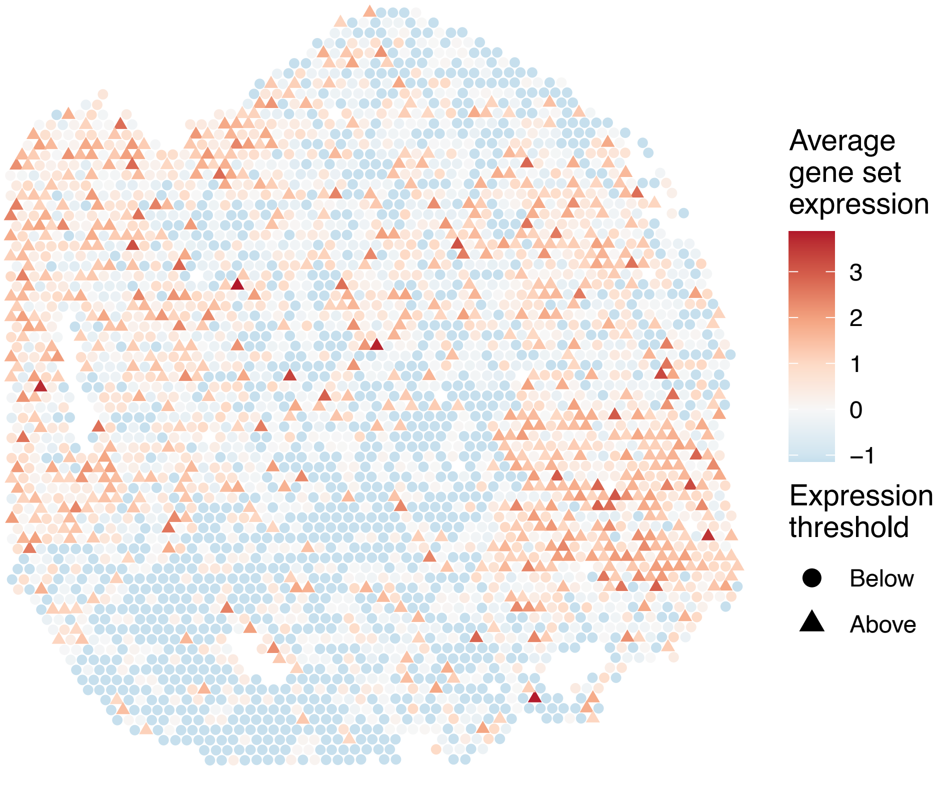

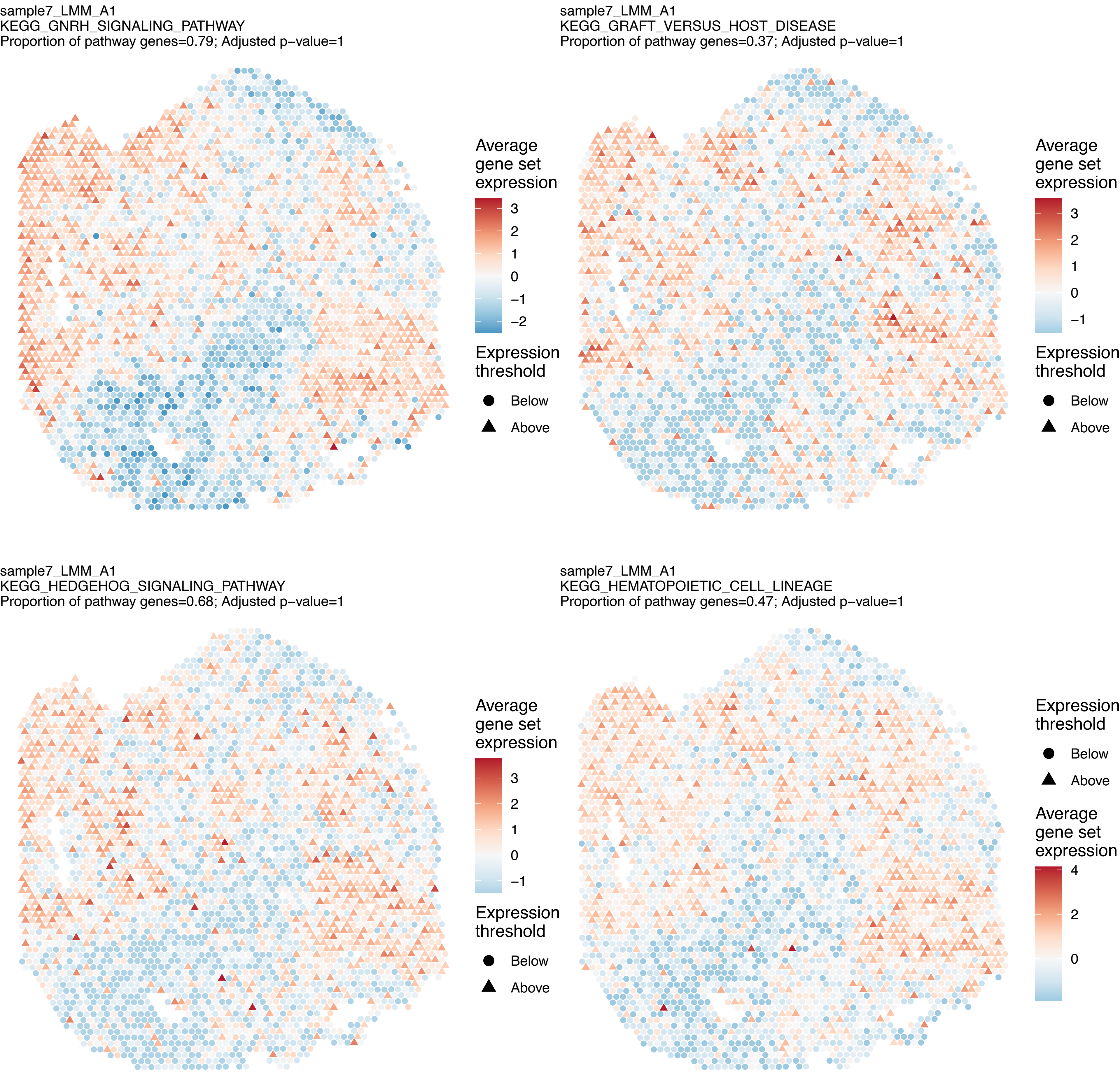
sample7_LMM_A1
KEGG_GNRH_SIGNALING_PATHWAY
Proportion of pathway genes=0.79; Adjusted p−value=1
Average gene set expression
3
2
1
0
−1
−2
Expression threshold
Below
Above
sample7_LMM_A1
KEGG_GRAFT_VERSUS_HOST_DISEASE
Proportion of pathway genes=0.37; Adjusted p−value=1
Average gene set expression
3
2
1
0
−1
Expression threshold
Below
Above
sample7_LMM_A1
KEGG_HEDGEHOG_SIGNALING_PATHWAY
Proportion of pathway genes=0.68; Adjusted p−value=1
Average gene set expression
3
2
1
0
−1
Expression threshold
Below
Above
sample7_LMM_A1
KEGG_HEMATOPOIETIC_CELL_LINEAGE
Proportion of pathway genes=0.47; Adjusted p−value=1
Expression threshold
Below
Above
Average gene set expression
4
2
0

sample7_LMM_A1
KEGG_HISTIDINE_METABOLISM
Proportion of pathway genes=0.73; Adjusted p−value=1

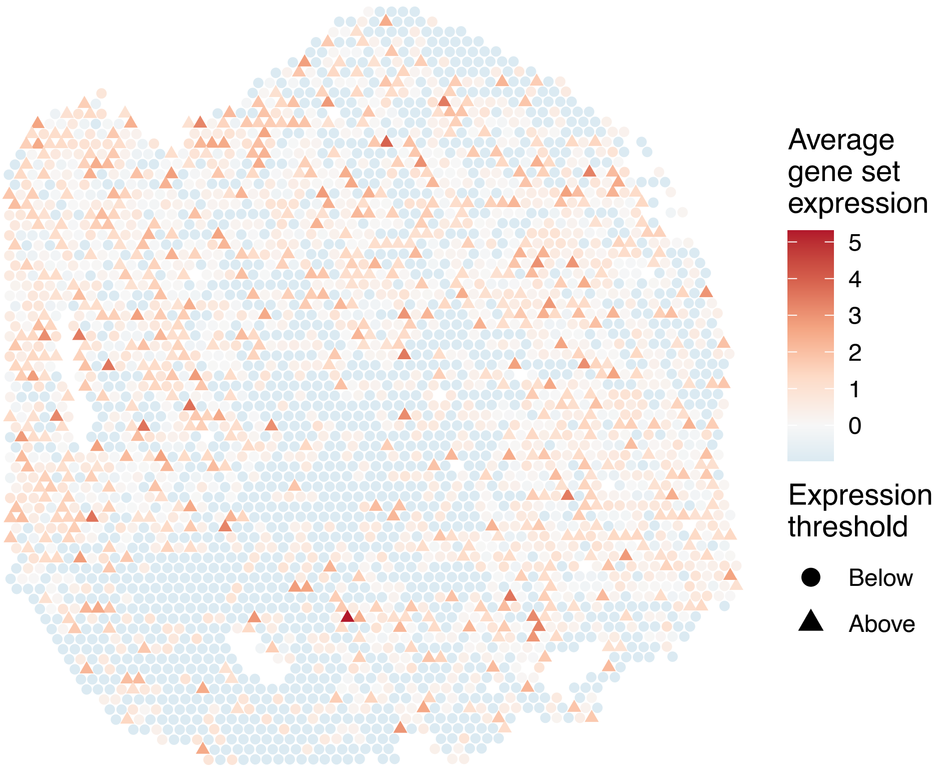


sample7_LMM_A1
KEGG_HOMOLOGOUS_RECOMBINATION
Proportion of pathway genes=0.97; Adjusted p−value=1

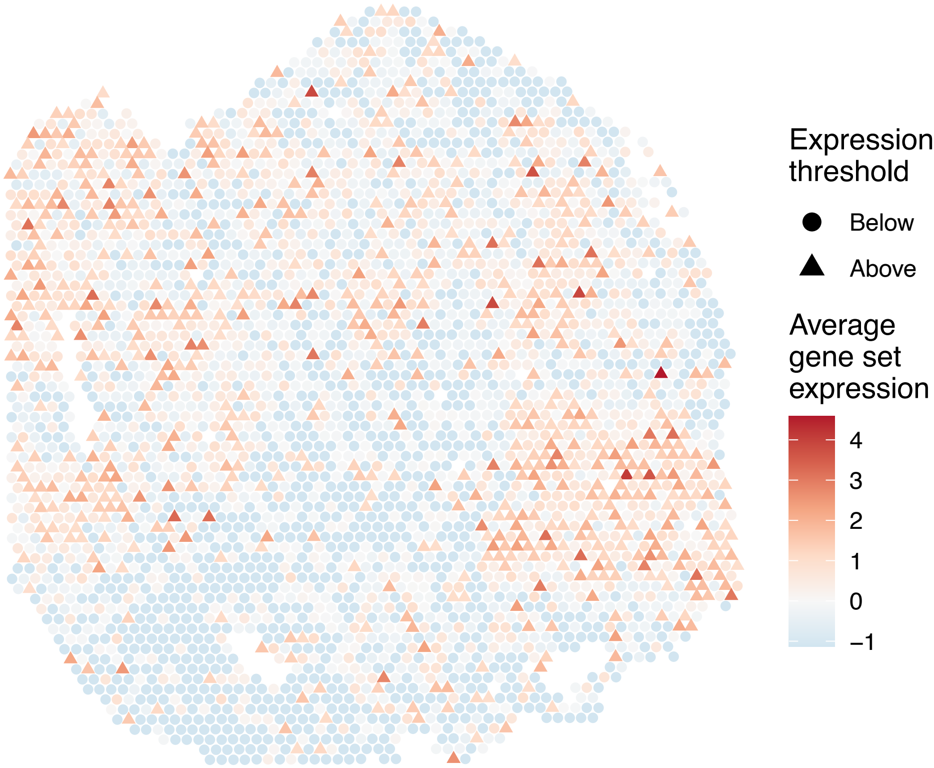


sample7_LMM_A1
KEGG_HUNTINGTONS_DISEASE
Proportion of pathway genes=0.89; Adjusted p−value=1

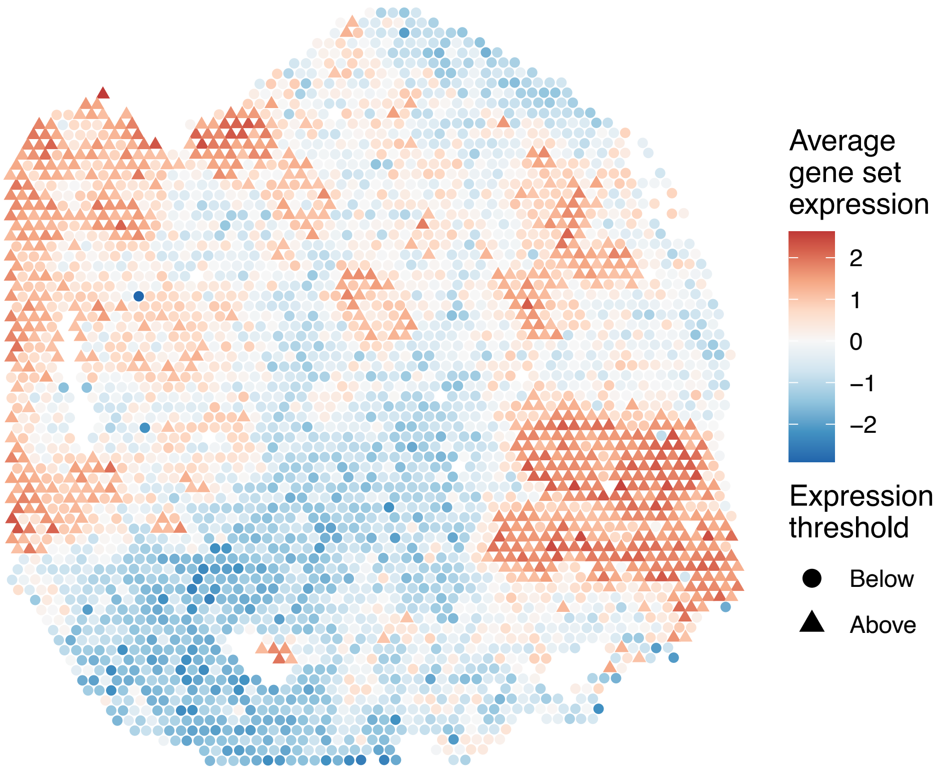


sample7_LMM_A1
KEGG_HYPERTROPHIC_CARDIOMYOPATHY_HCM
Proportion of pathway genes=0.71; Adjusted p−value=1

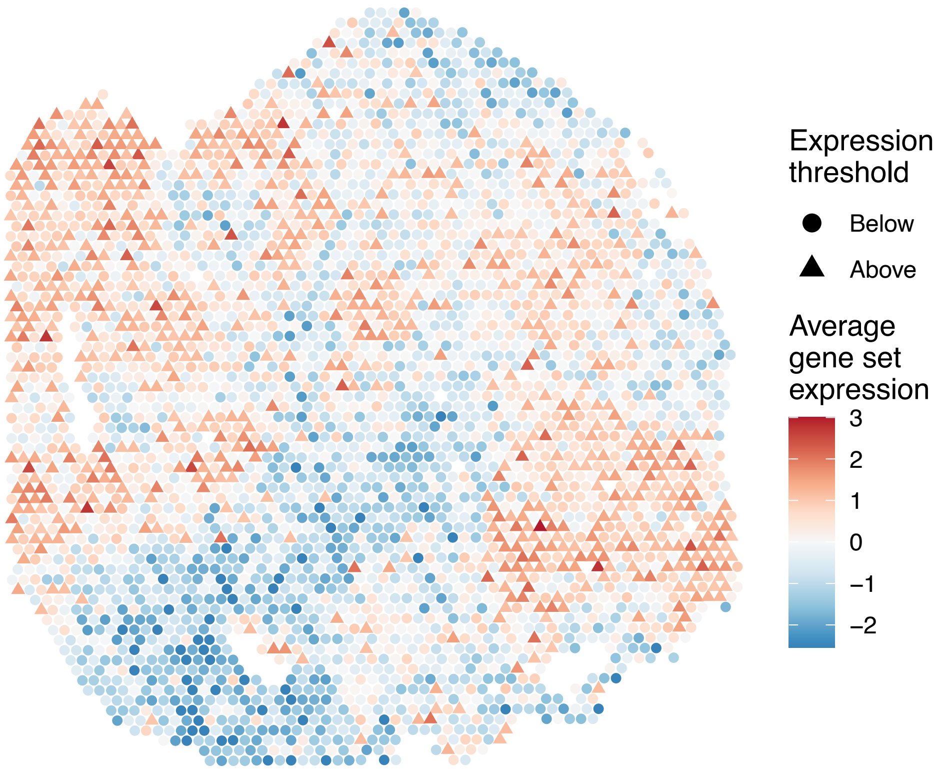

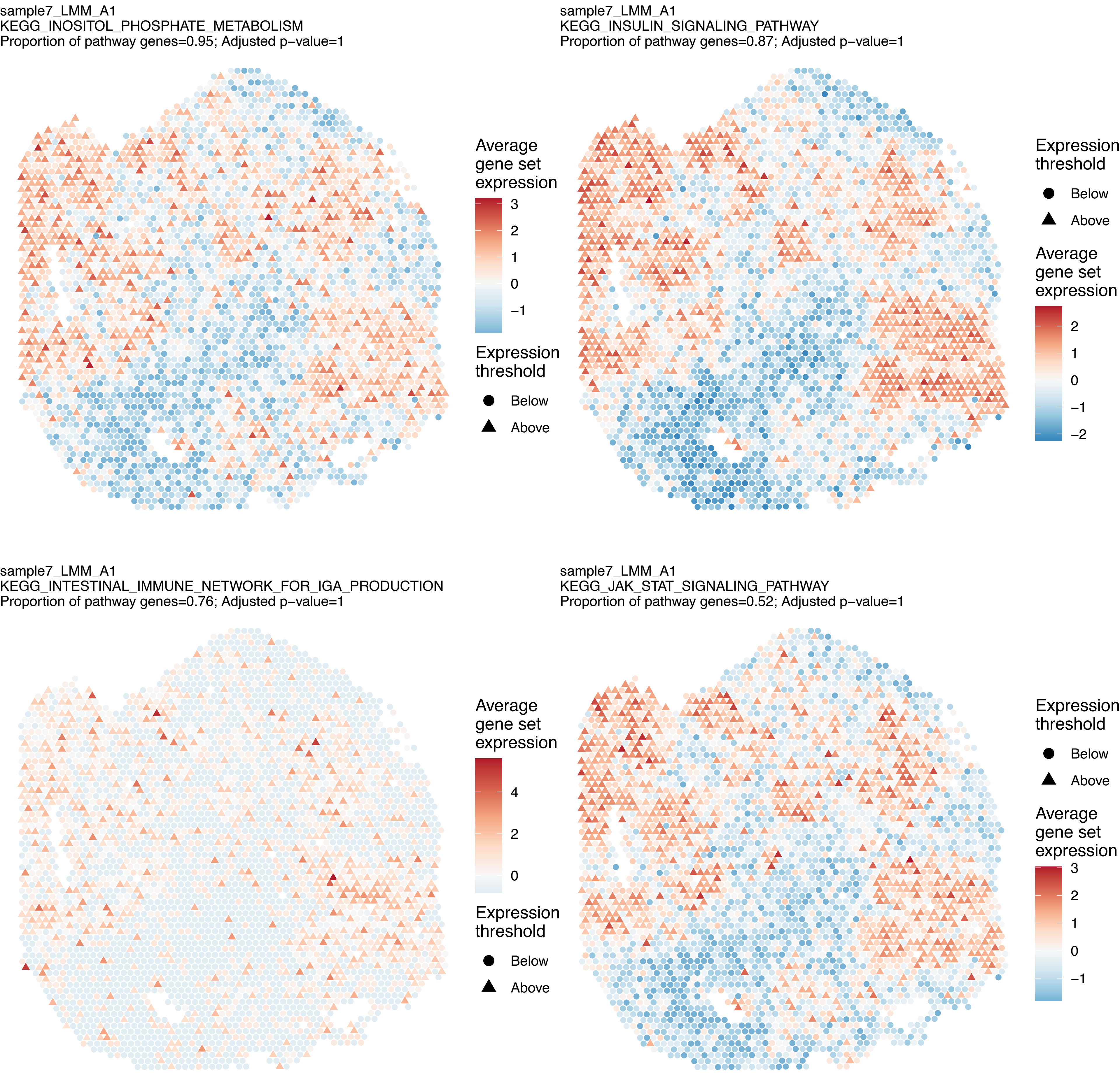
sample7_LMM_A1
KEGG_INOSITOL_PHOSPHATE_METABOLISM
Proportion of pathway genes=0.95; Adjusted p−value=1
Average gene set expression
3
2
1
0
−1
Expression threshold
Below
Above
sample7_LMM_A1
KEGG_INSULIN_SIGNALING_PATHWAY
Proportion of pathway genes=0.87; Adjusted p−value=1
Expression threshold
Below
Above
Average gene set expression
2
1
0
−1
−2
sample7_LMM_A1
KEGG_INTESTINAL_IMMUNE_NETWORK_FOR_IGA_PRODUCTION
Proportion of pathway genes=0.76; Adjusted p−value=1
Average gene set expression
4
2
0
Expression threshold
Below
Above
sample7_LMM_A1
KEGG_JAK_STAT_SIGNALING_PATHWAY
Proportion of pathway genes=0.52; Adjusted p−value=1
Expression threshold
Below
Above
Average gene set expression
3
2
1
0
−1

sample7_LMM_A1
KEGG_LEISHMANIA_INFECTION
Proportion of pathway genes=0.79; Adjusted p–value=1

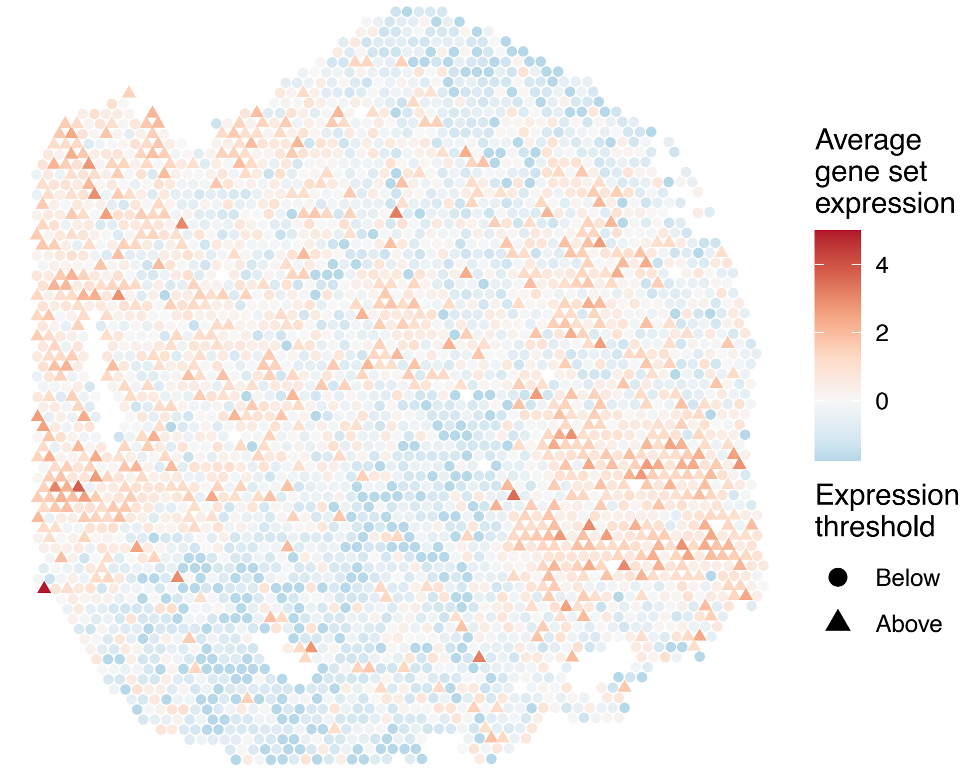


sample7_LMM_A1
KEGG_LEUKOCYTE_TRANSENDOTHELIAL_MIGRATION
Proportion of pathway genes=0.78; Adjusted p–value=1

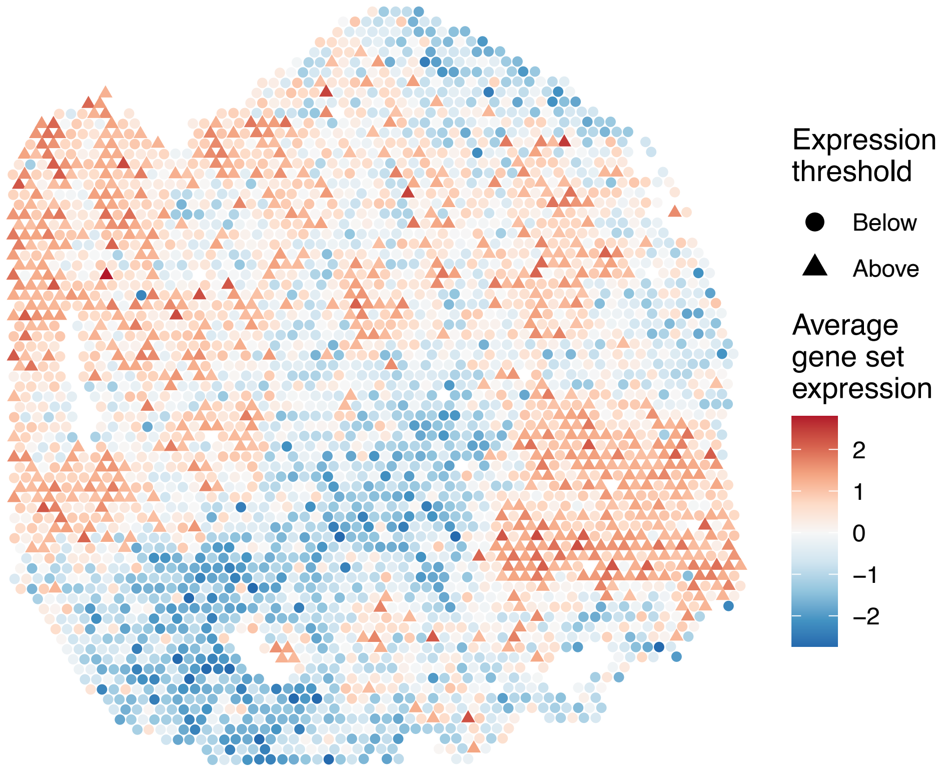


sample7_LMM_A1
KEGG_LIMONENE_AND_PINENE_DEGRADATION
Proportion of pathway genes=1; Adjusted p–value=1

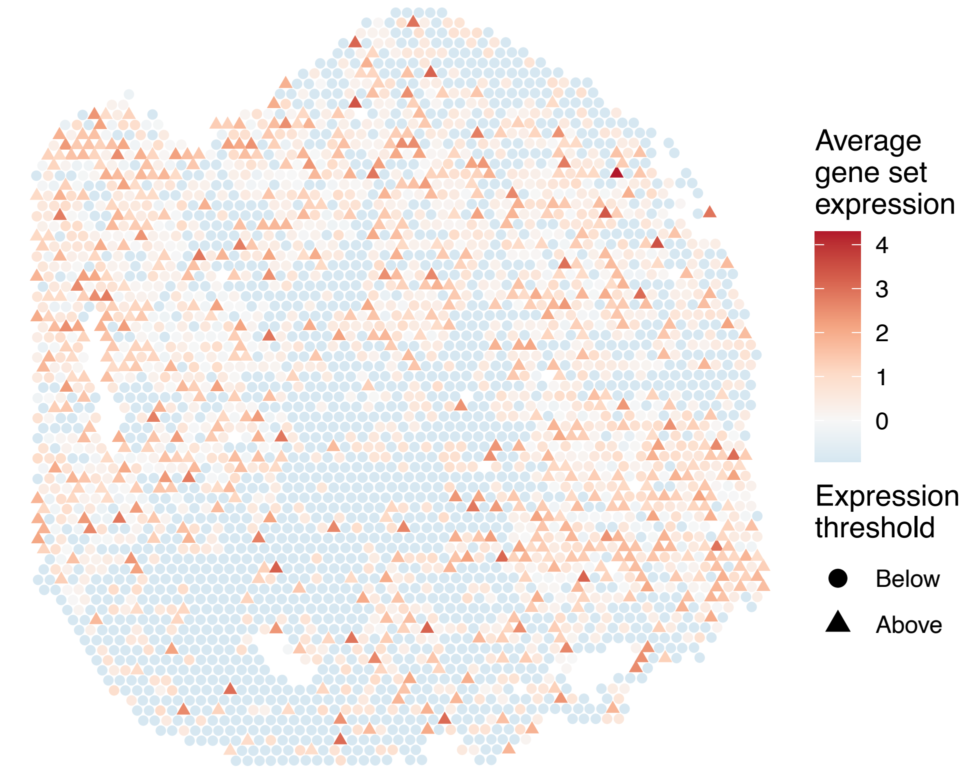


sample7_LMM_A1
KEGG_LONG_TERM_DEPRESSION
Proportion of pathway genes=0.81; Adjusted p–value=1

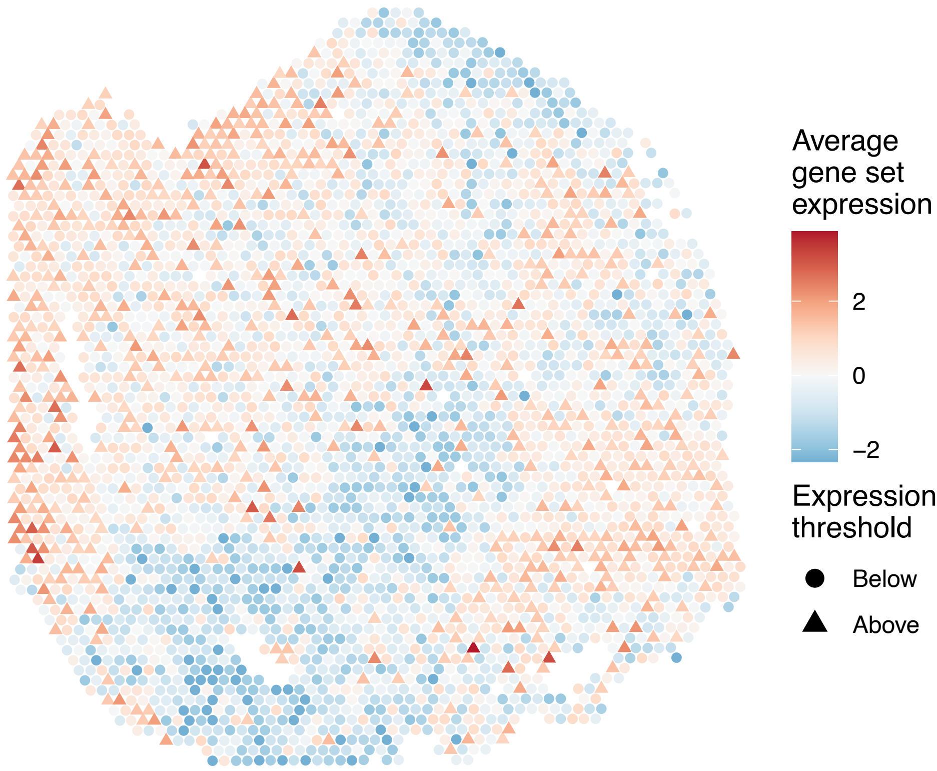

sample7_LMM_A1
KEGG_LONG_TERM_POTENTIATION
Proportion of pathway genes=0.92; Adjusted p−value=1

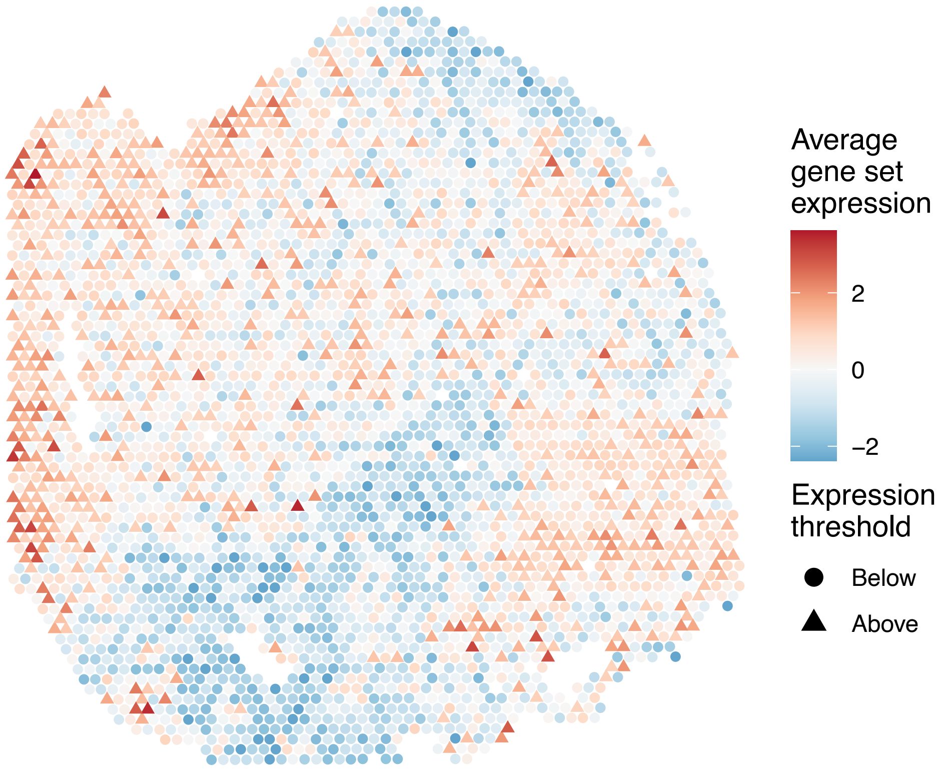


sample7_LMM_A1
KEGG_LYSINE_DEGRADATION
Proportion of pathway genes=0.98; Adjusted p−value=1

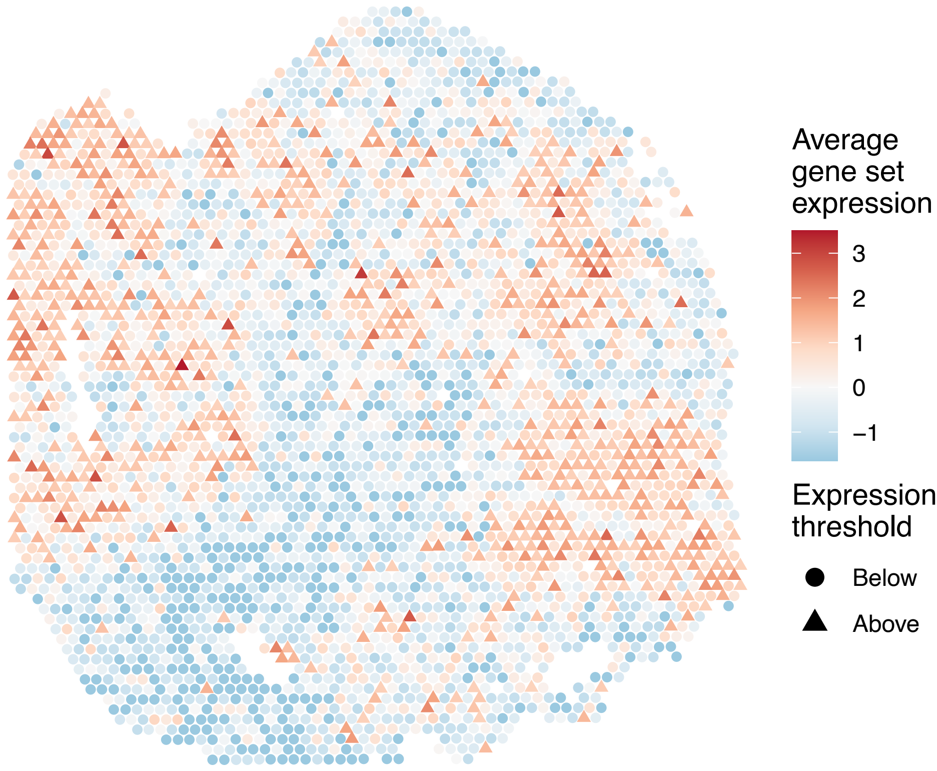


sample7_LMM_A1
KEGG_LYSOSOME
Proportion of pathway genes=0.94; Adjusted p−value=1

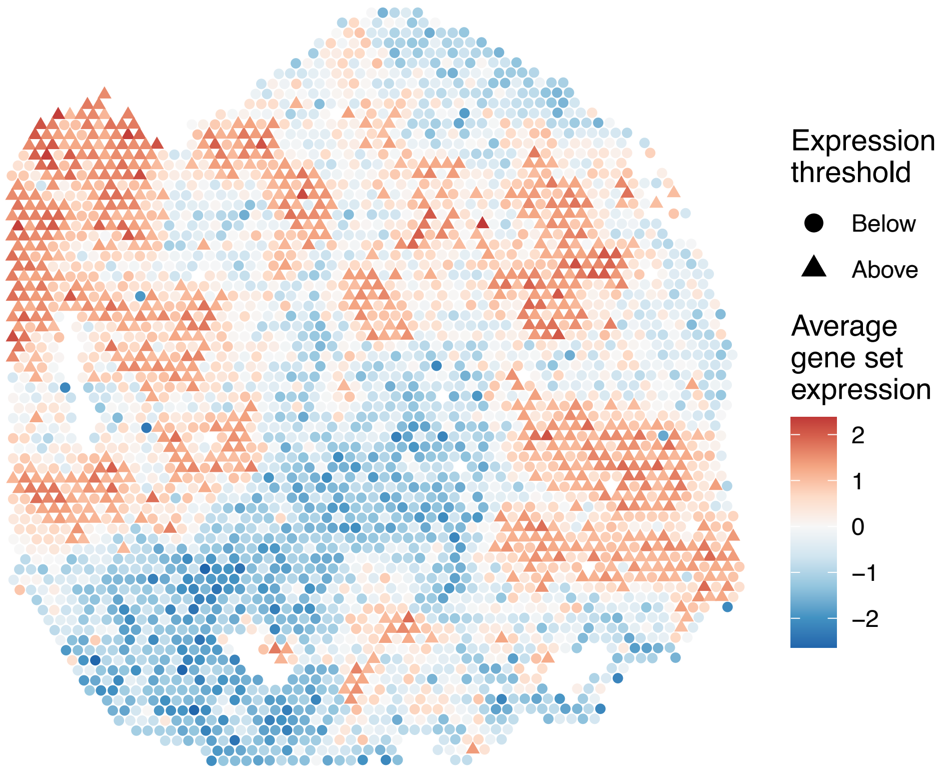


sample7_LMM_A1
KEGG_MAPK_SIGNALING_PATHWAY
Proportion of pathway genes=0.82; Adjusted p−value=1

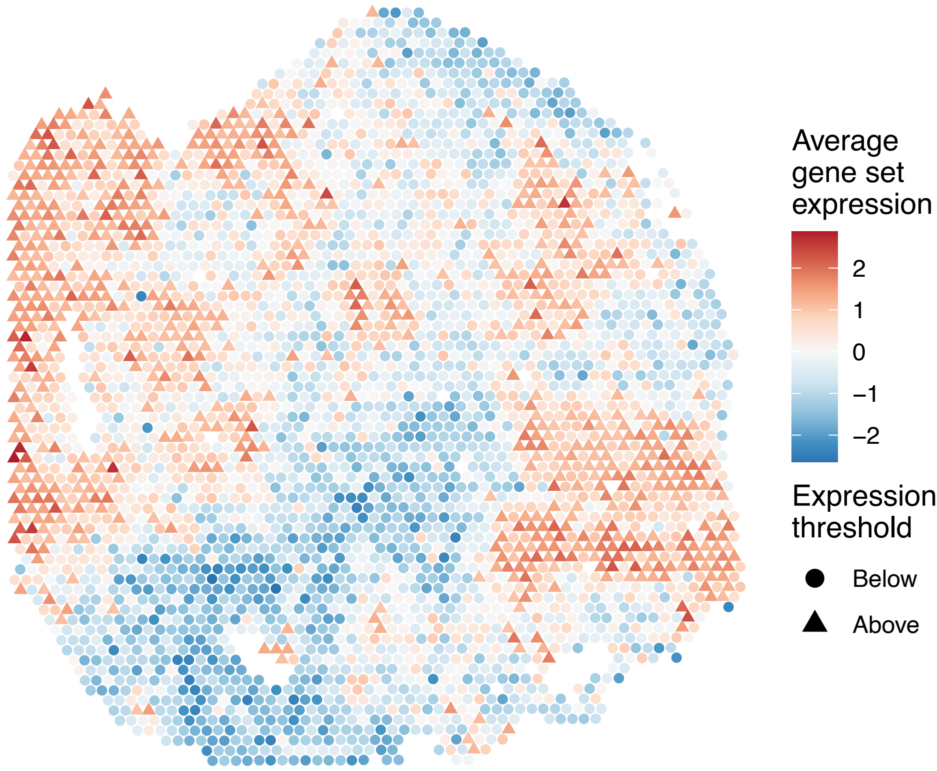

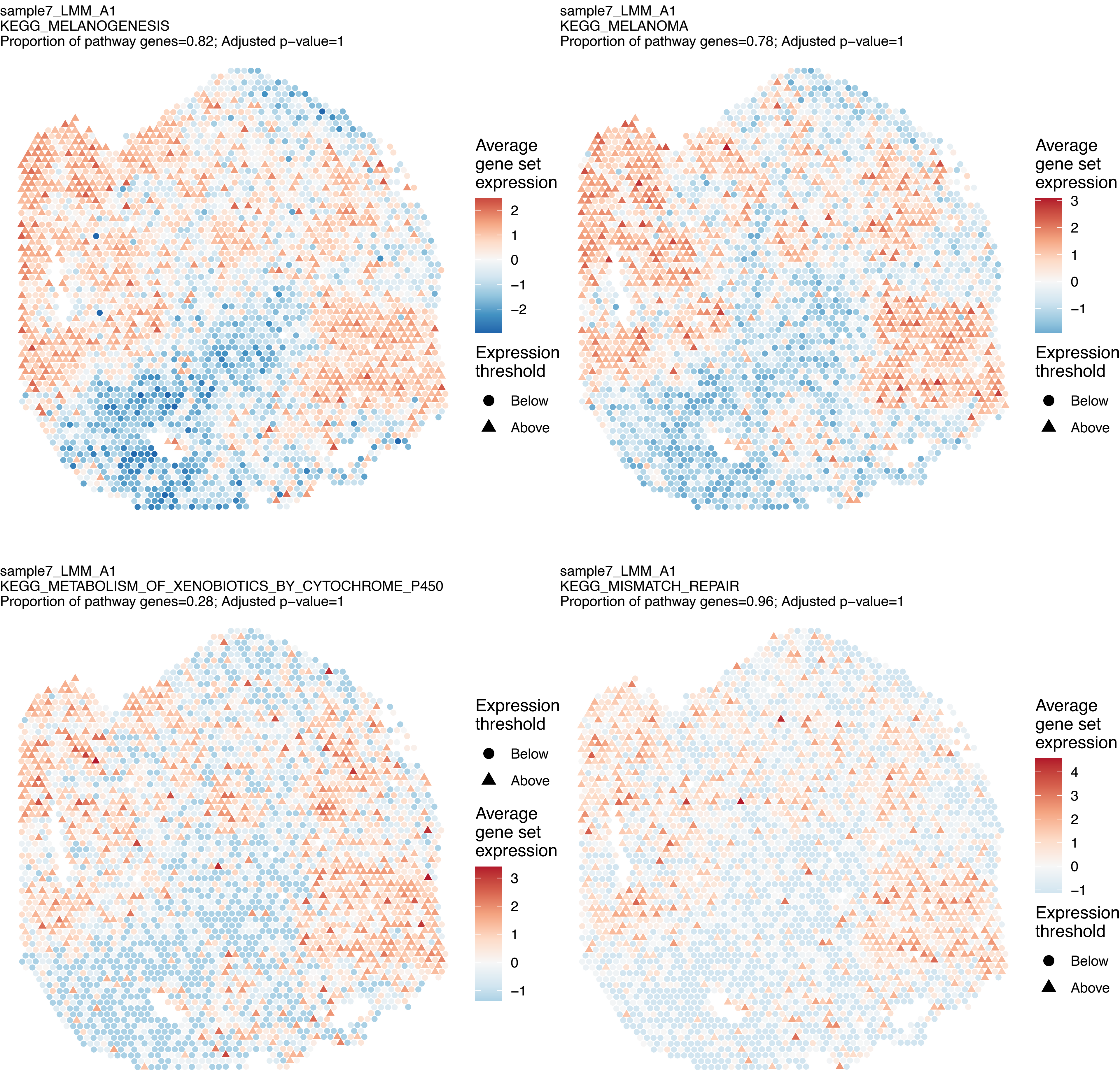
sample7_LMM_A1
KEGG_MELANOGENESIS
Proportion of pathway genes=0.82; Adjusted p−value=1
Average gene set expression
2
1
0
−1
−2
Expression threshold
Below
Above
sample7_LMM_A1
KEGG_MELANOMA
Proportion of pathway genes=0.78; Adjusted p−value=1
Average gene set expression
3
2
1
0
−1
Expression threshold
Below
Above
sample7_LMM_A1
KEGG_METABOLISM_OF_XENOBIOTICS_BY_CYTOCHROME_P450
Proportion of pathway genes=0.28; Adjusted p−value=1
Expression threshold
Below
Above
Average gene set expression
3
2
1
0
−1
sample7_LMM_A1
KEGG_MISMATCH_REPAIR
Proportion of pathway genes=0.96; Adjusted p−value=1
Average gene set expression
4
3
2
1
0
−1
Expression threshold
Below
Above

sample7_LMM_A1
KEGG_MTOR_SIGNALING_PATHWAY
Proportion of pathway genes=0.94; Adjusted p−value=1

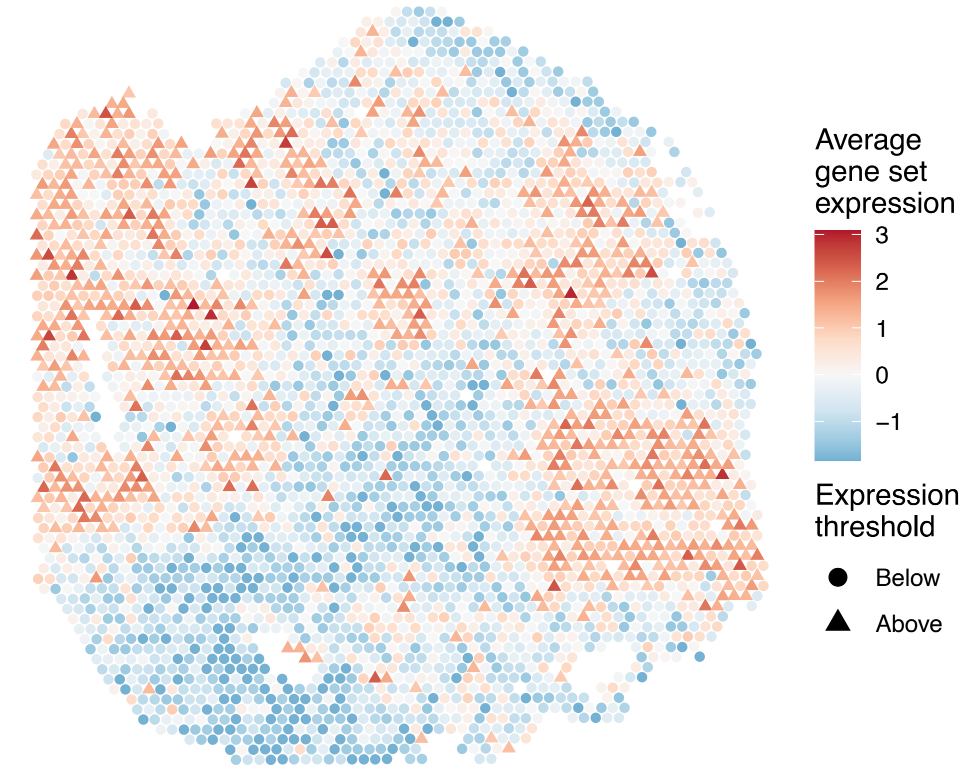


sample7_LMM_A1
KEGG_N_GLYCAN_BIOSYNTHESIS
Proportion of pathway genes=1; Adjusted p−value=1

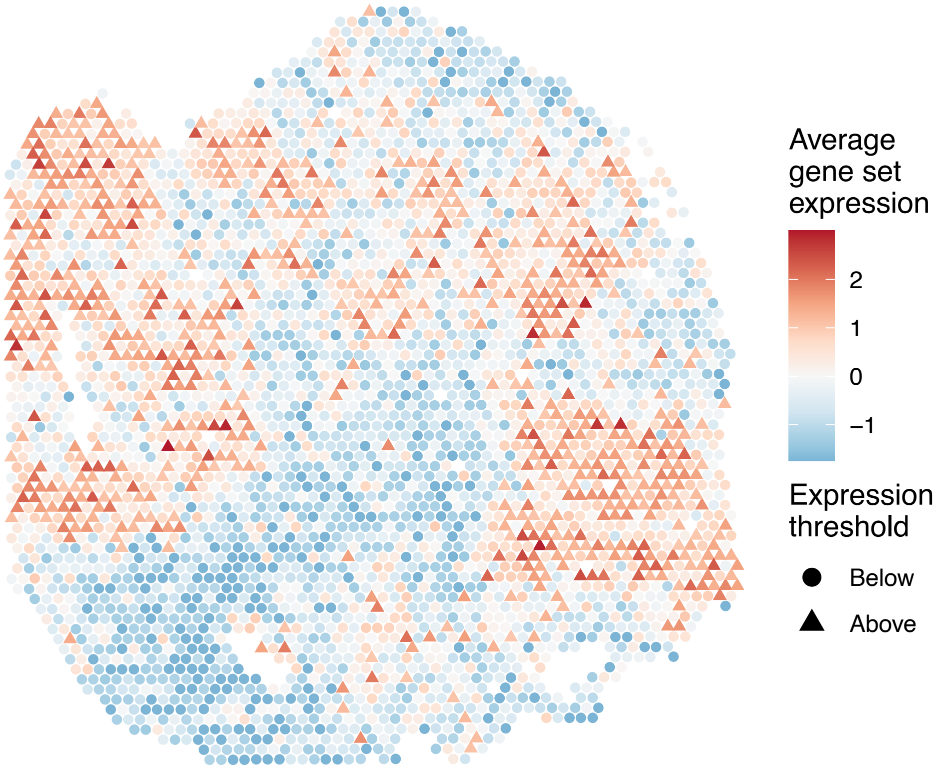


sample7_LMM_A1
KEGG_NEUROACTIVE_LIGAND_RECEPTOR_INTERACTION
Proportion of pathway genes=0.49; Adjusted p−value=1

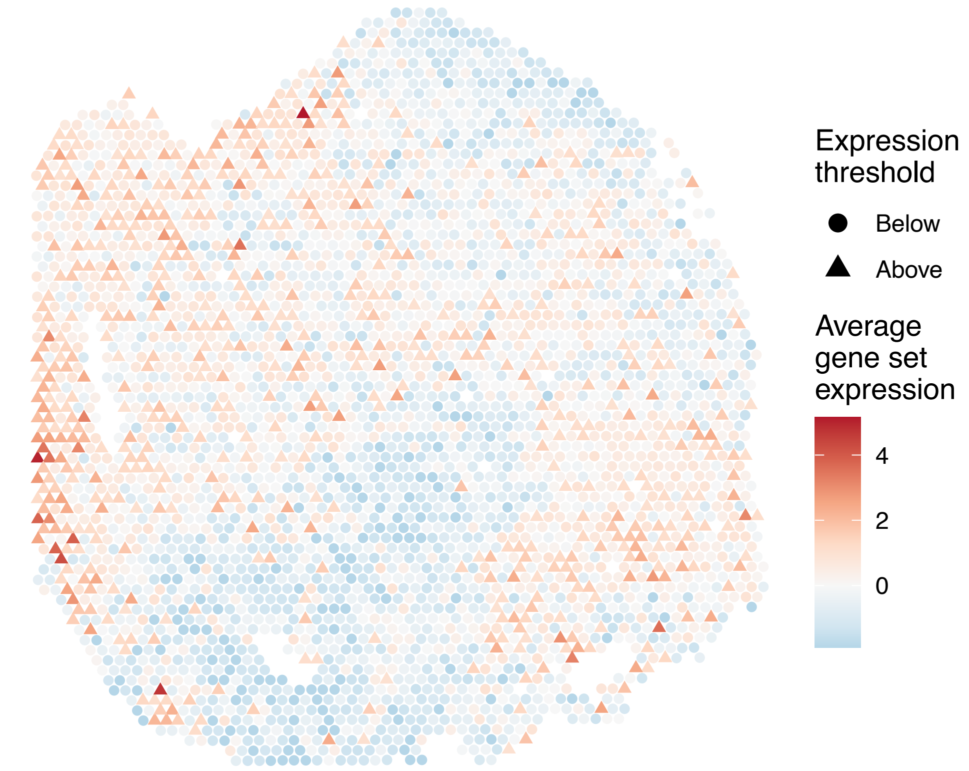


sample7_LMM_A1
KEGG_NEUROTROPHIN_SIGNALING_PATHWAY
Proportion of pathway genes=0.92; Adjusted p−value=1

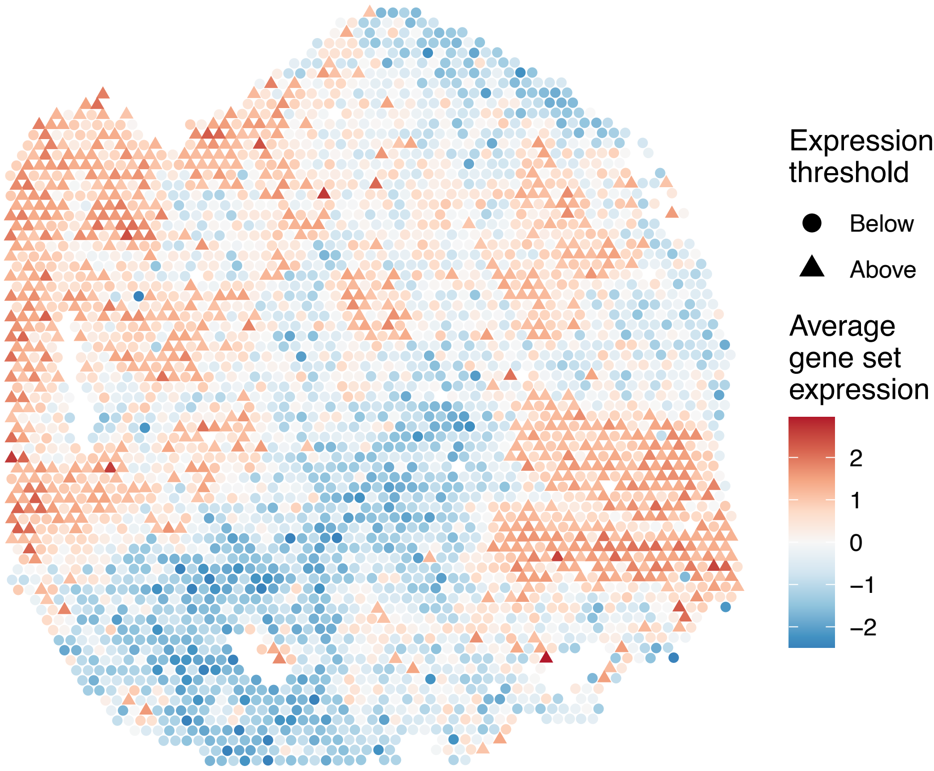

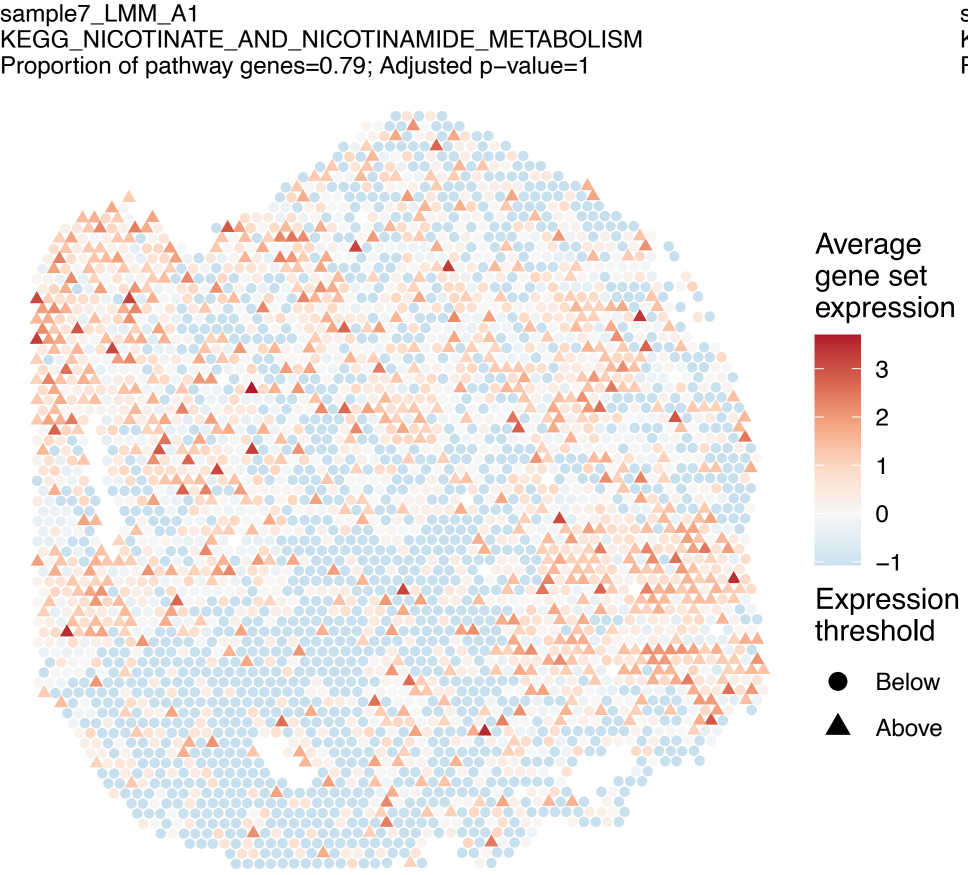
sample7_LMM_A1
KEGG_NICOTINATE_AND_NICOTINAMIDE_METABOLISM
Proportion of pathway genes=0.79; Adjusted p−value=1
Average gene set expression
3
2
1
0
−1
Expression threshold
Below
Above

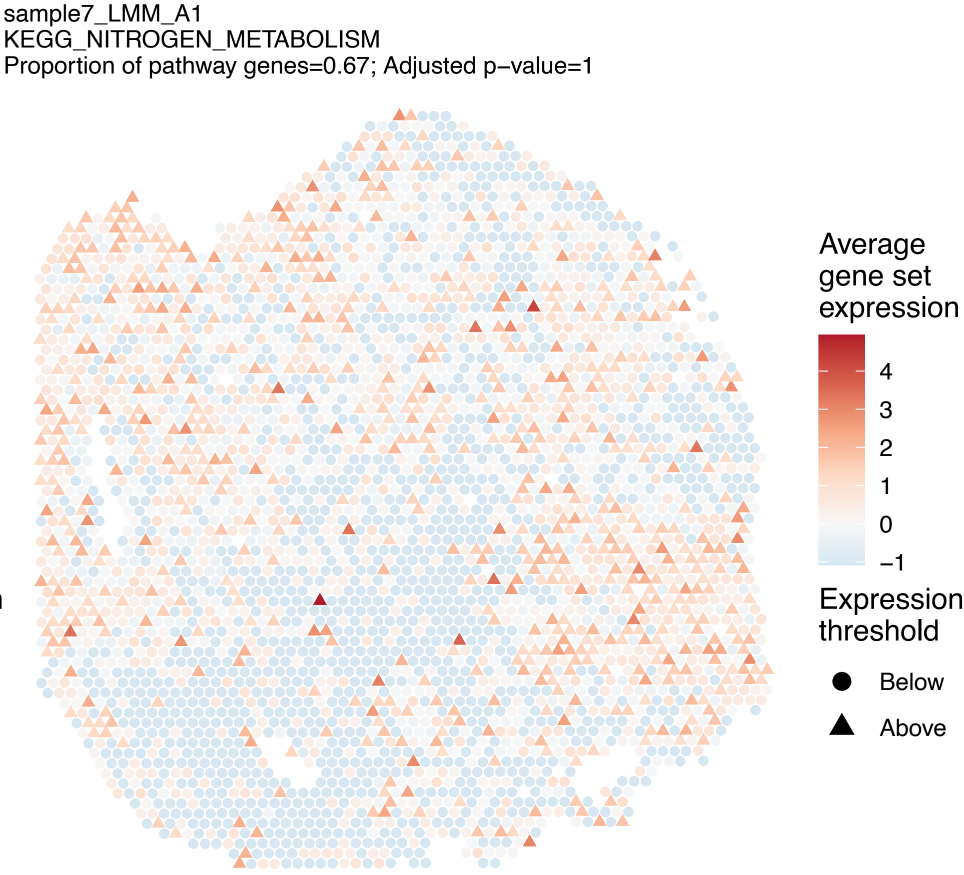
sample7_LMM_A1
KEGG_NITROGEN_METABOLISM
Proportion of pathway genes=0.67; Adjusted p−value=1
Average gene set expression
4
3
2
1
0
−1
Expression threshold
Below
Above

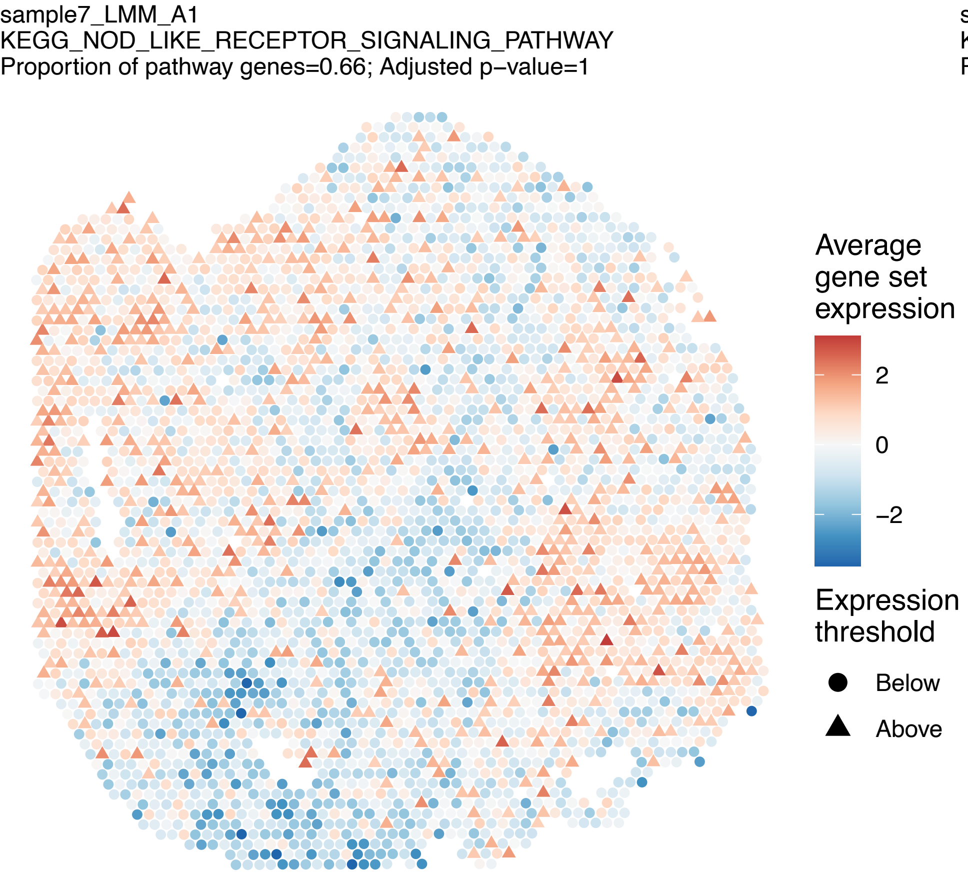
sample7_LMM_A1
KEGG_NOD_LIKE_RECEPTOR_SIGNALING_PATHWAY
Proportion of pathway genes=0.66; Adjusted p−value=1
Average gene set expression
2
0
−2
Expression threshold
Below
Above

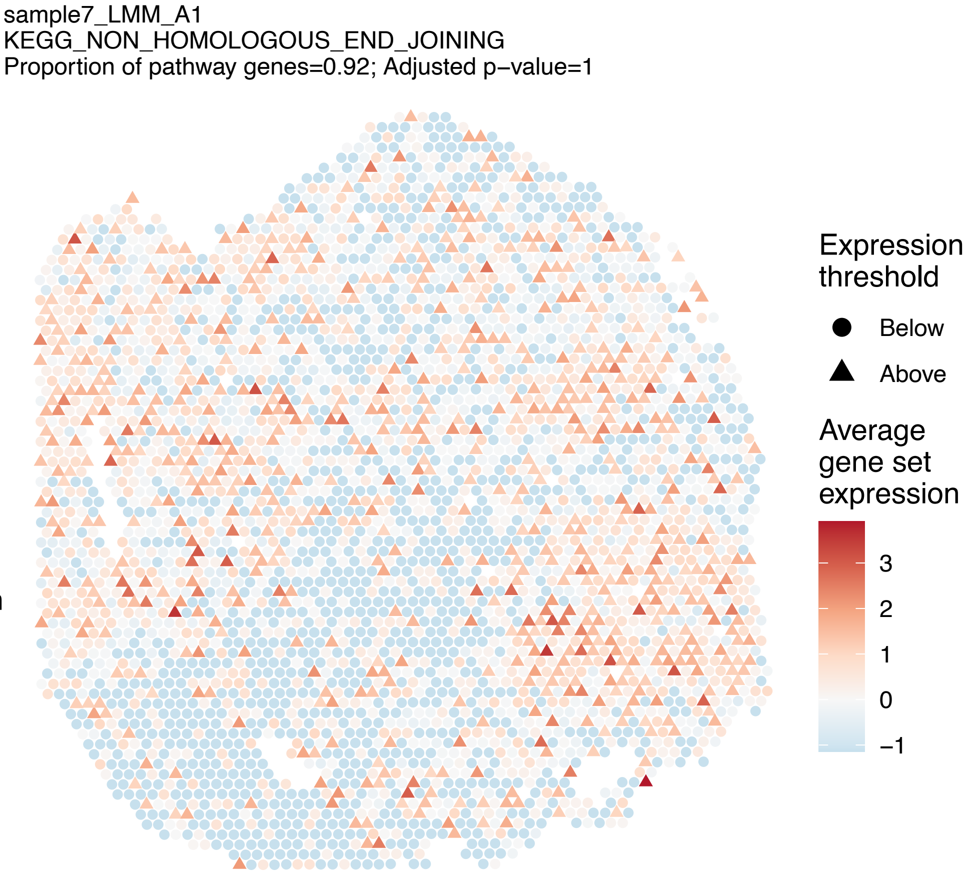
sample7_LMM_A1
KEGG_NON_HOMOLOGOUS_END_JOINING
Proportion of pathway genes=0.92; Adjusted p−value=1
Expression threshold
Below
Above
Average gene set expression
3
2
1
0
−1

sample7_LMM_A1
KEGG_NON_SMALL_CELL_LUNG_CANCER
Proportion of pathway genes=0.94; Adjusted p−value=1

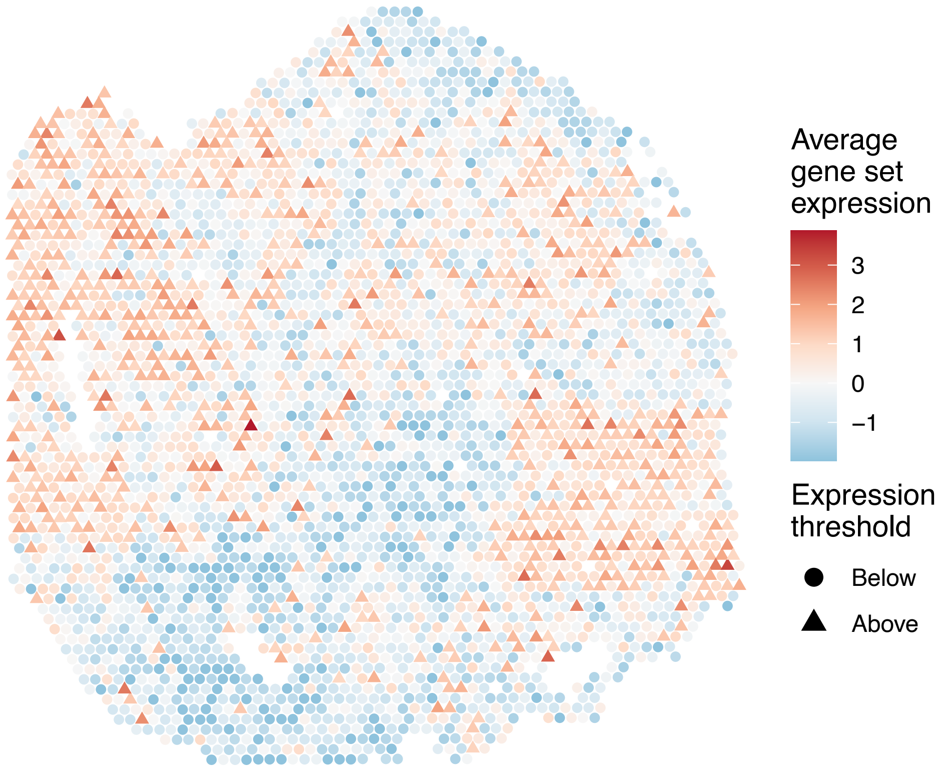


sample7_LMM_A1
KEGG_NOTCH_SIGNALING_PATHWAY
Proportion of pathway genes=0.91; Adjusted p−value=1

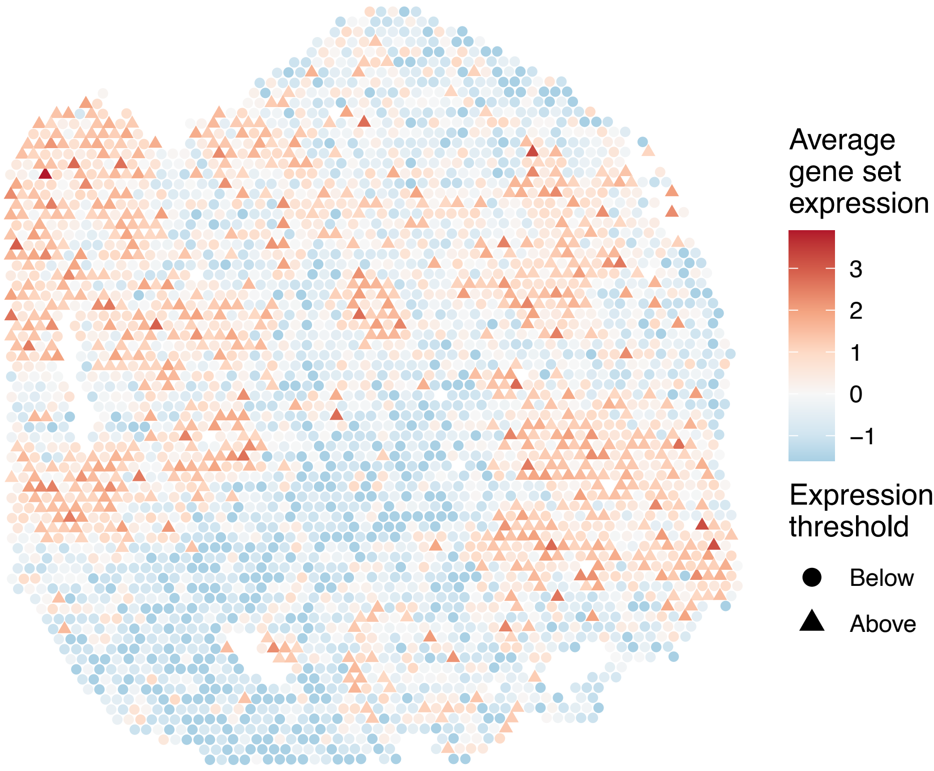


sample7_LMM_A1
KEGG_NUCLEOTIDE_EXCISION_REPAIR
Proportion of pathway genes=0.98; Adjusted p−value=1

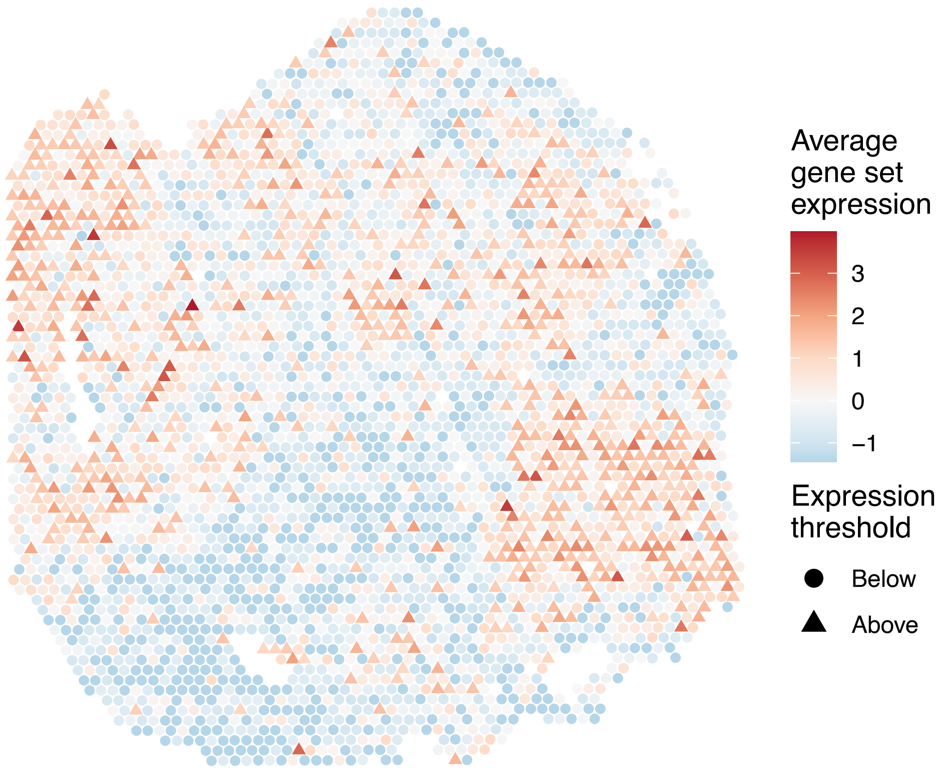


sample7_LMM_A1
KEGG_O_GLYCAN_BIOSYNTHESIS
Proportion of pathway genes=0.82; Adjusted p−value=1

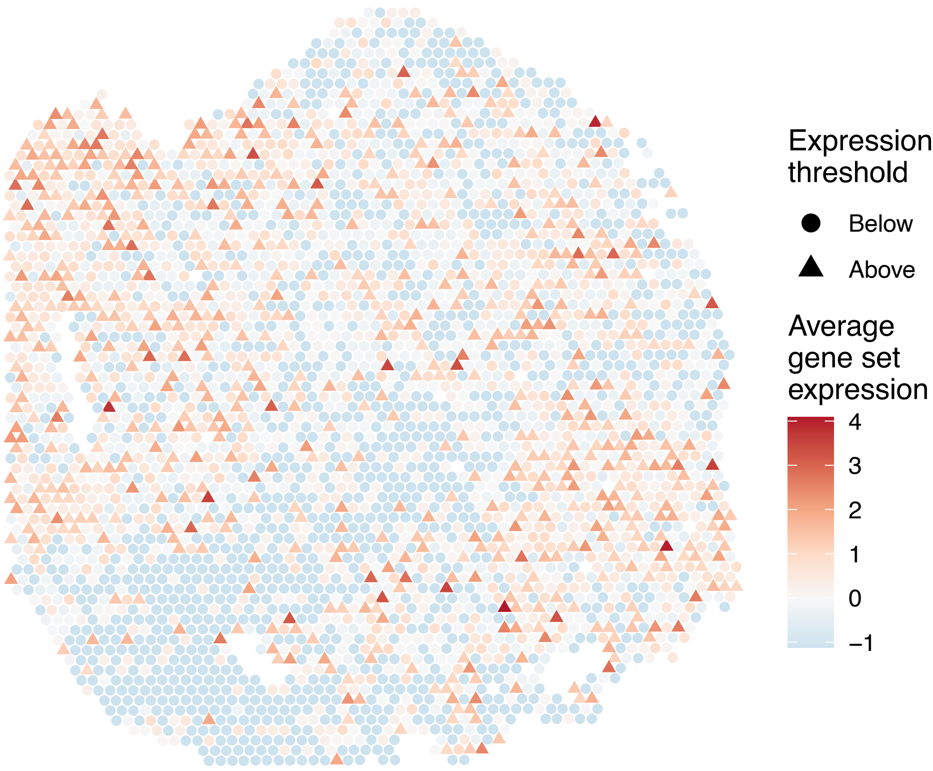

sample7_LMM_A1
KEGG_ONE_CARBON_POOL_BY_FOLATE
Proportion of pathway genes=1; Adjusted p−value=1

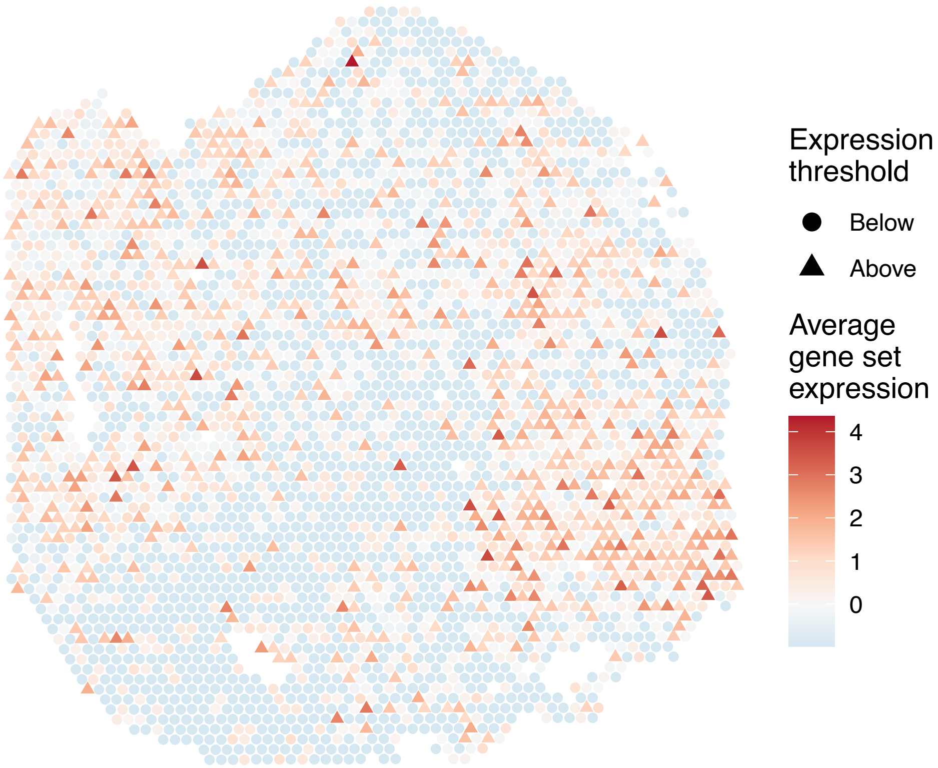


sample7_LMM_A1
KEGG_OOCYTE_MEIOSIS
Proportion of pathway genes=0.88; Adjusted p−value=1

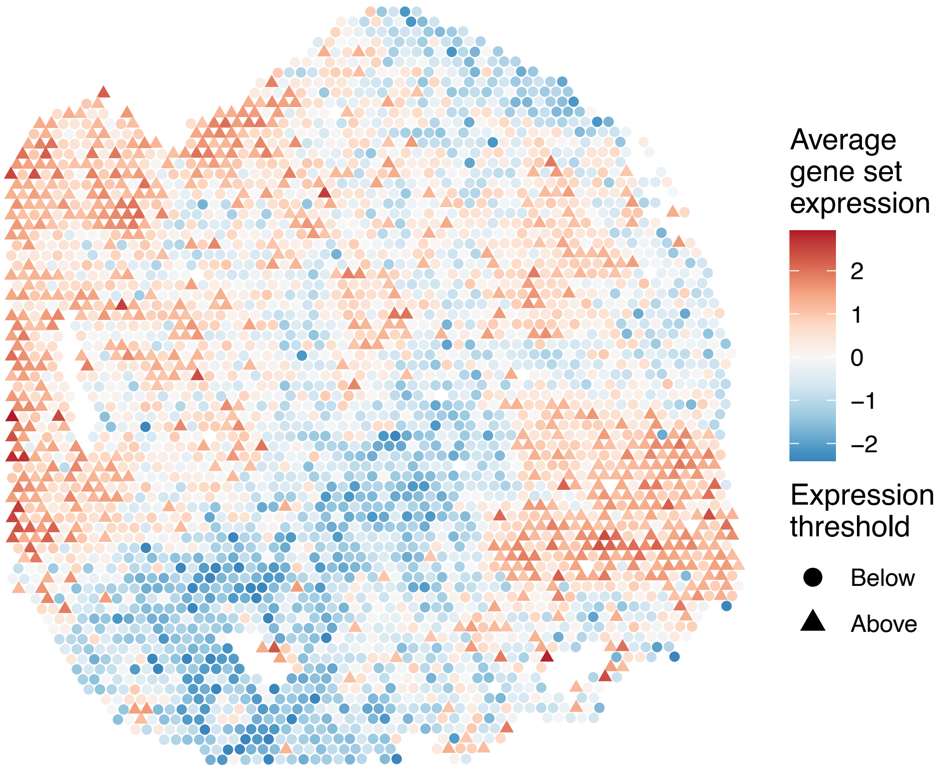


sample7_LMM_A1
KEGG_OTHER_GLYCAN_DEGRADATION
Proportion of pathway genes=0.96; Adjusted p−value=1

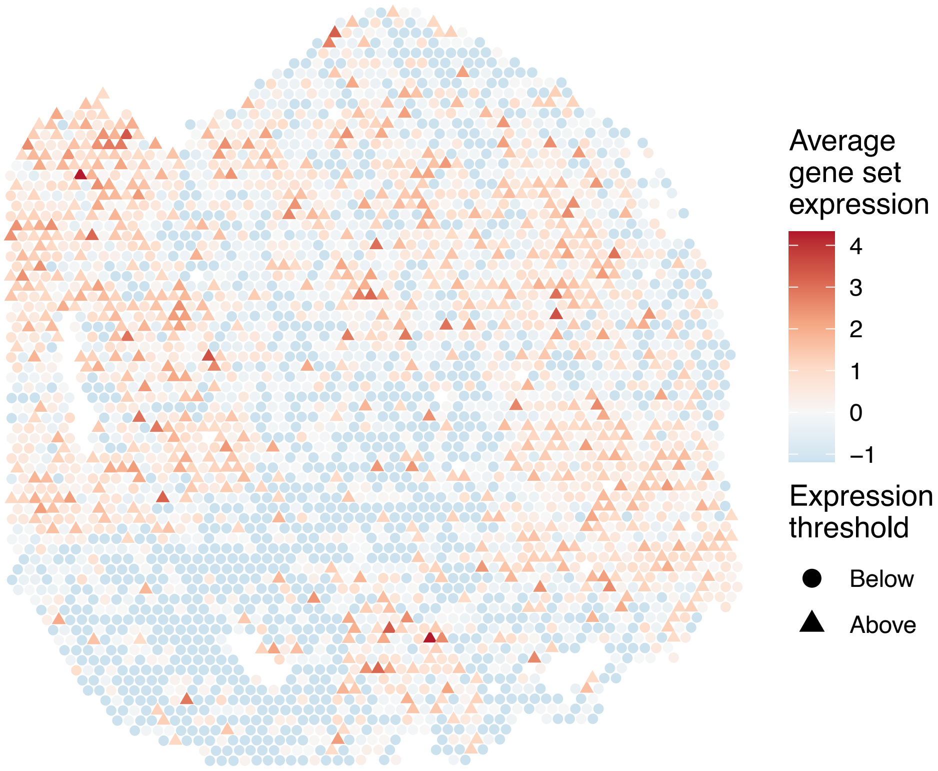


sample7_LMM_A1
KEGG_OXIDATIVE_PHOSPHORYLATION
Proportion of pathway genes=0.88; Adjusted p−value=1

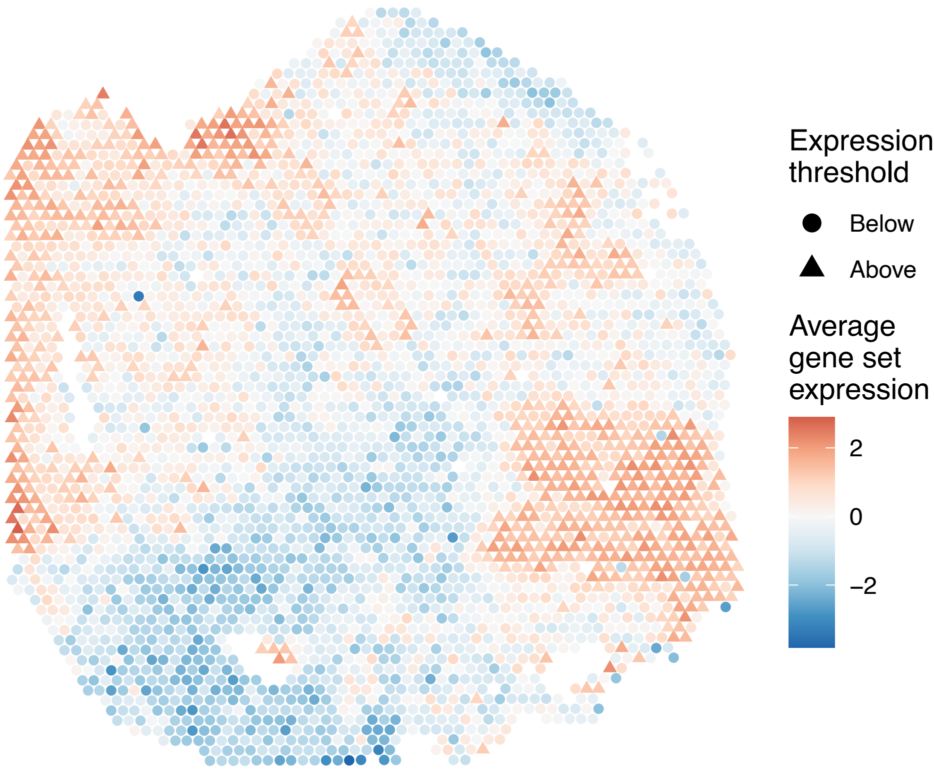

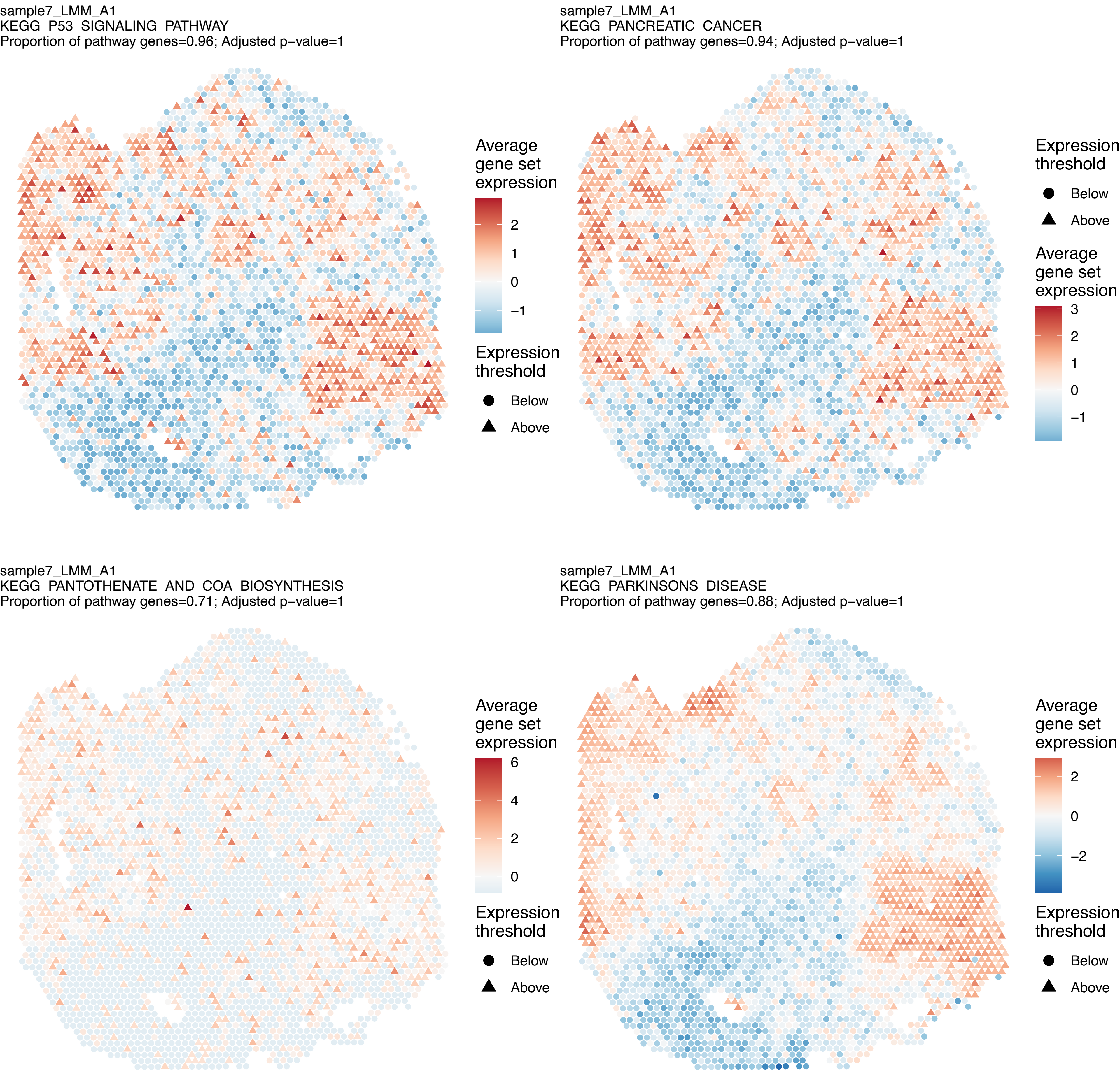
sample7_LMM_A1
KEGG_P53_SIGNALING_PATHWAY
Proportion of pathway genes=0.96; Adjusted p−value=1
Average gene set expression
2
1
0
−1
Expression threshold
Below
Above
sample7_LMM_A1
KEGG_PANCREATIC_CANCER
Proportion of pathway genes=0.94; Adjusted p−value=1
Expression threshold
Below
Above
Average gene set expression
3
2
1
0
−1
sample7_LMM_A1
KEGG_PANTOTHENATE_AND_COA_BIOSYNTHESIS
Proportion of pathway genes=0.71; Adjusted p−value=1
Average gene set expression
6
4
2
0
Expression threshold
Below
Above
sample7_LMM_A1
KEGG_PARKINSONS_DISEASE
Proportion of pathway genes=0.88; Adjusted p−value=1
Average gene set expression
2
0
−2
Expression threshold
Below
Above

sample7_LMM_A1
KEGG_PATHOGENIC_ESCHERICHIA_COLI_INFECTION
Proportion of pathway genes=0.83; Adjusted p−value=1

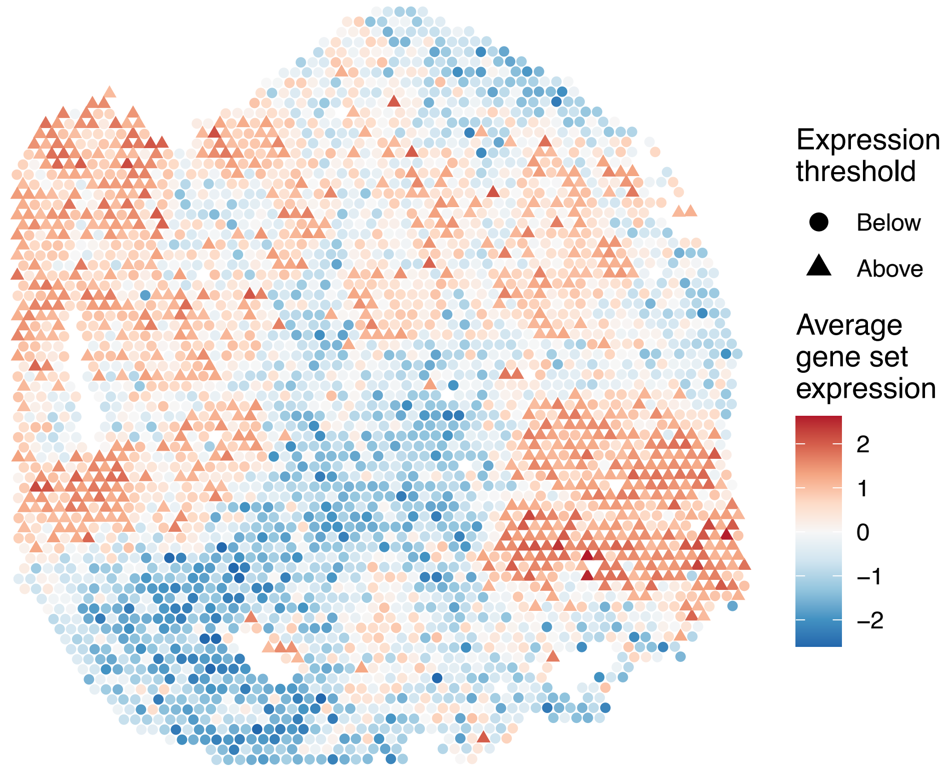


sample7_LMM_A1
KEGG_PATHWAYS_IN_CANCER
Proportion of pathway genes=0.84; Adjusted p−value=1

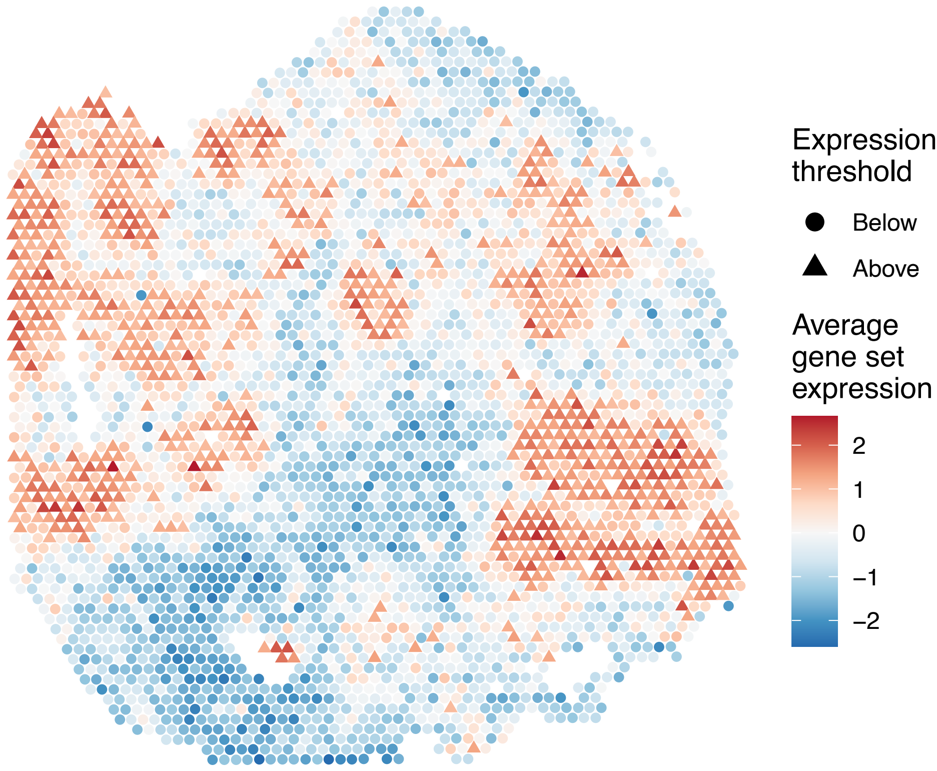


sample7_LMM_A1
KEGG_PENTOSE_PHOSPHATE_PATHWAY
Proportion of pathway genes=0.75; Adjusted p−value=1

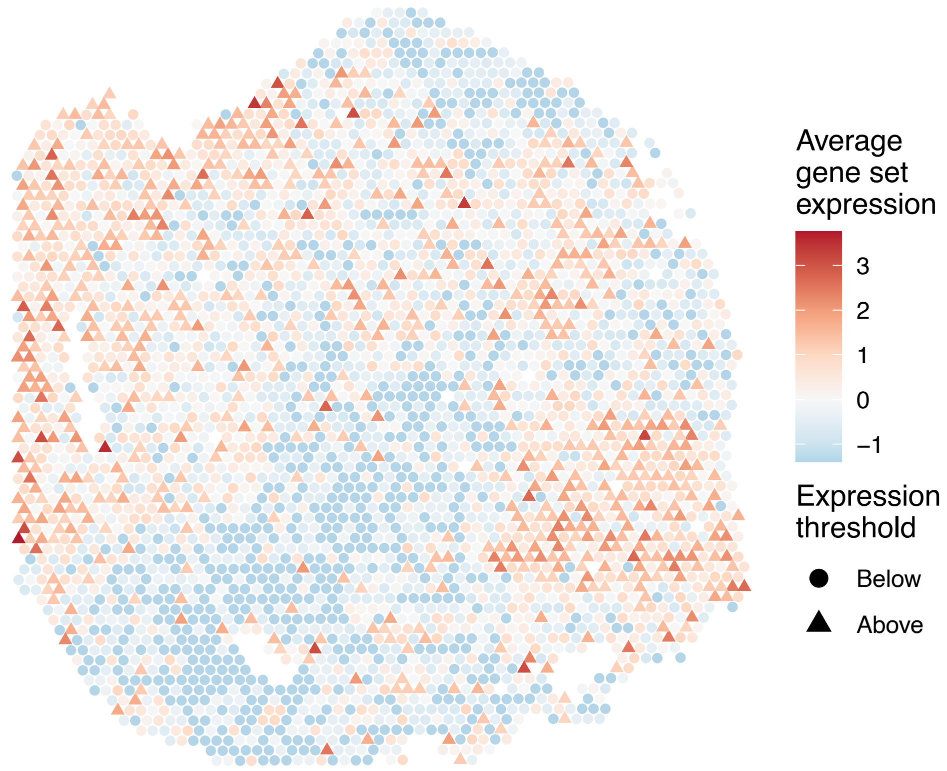


sample7_LMM_A1
KEGG_PEROXISOME
Proportion of pathway genes=0.85; Adjusted p−value=1

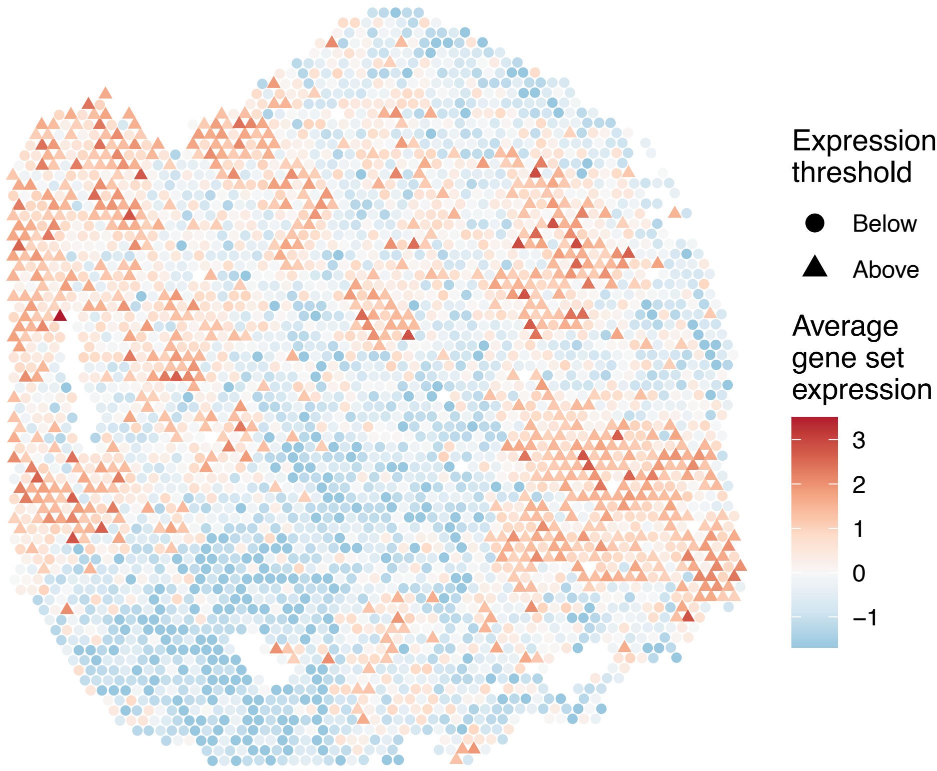

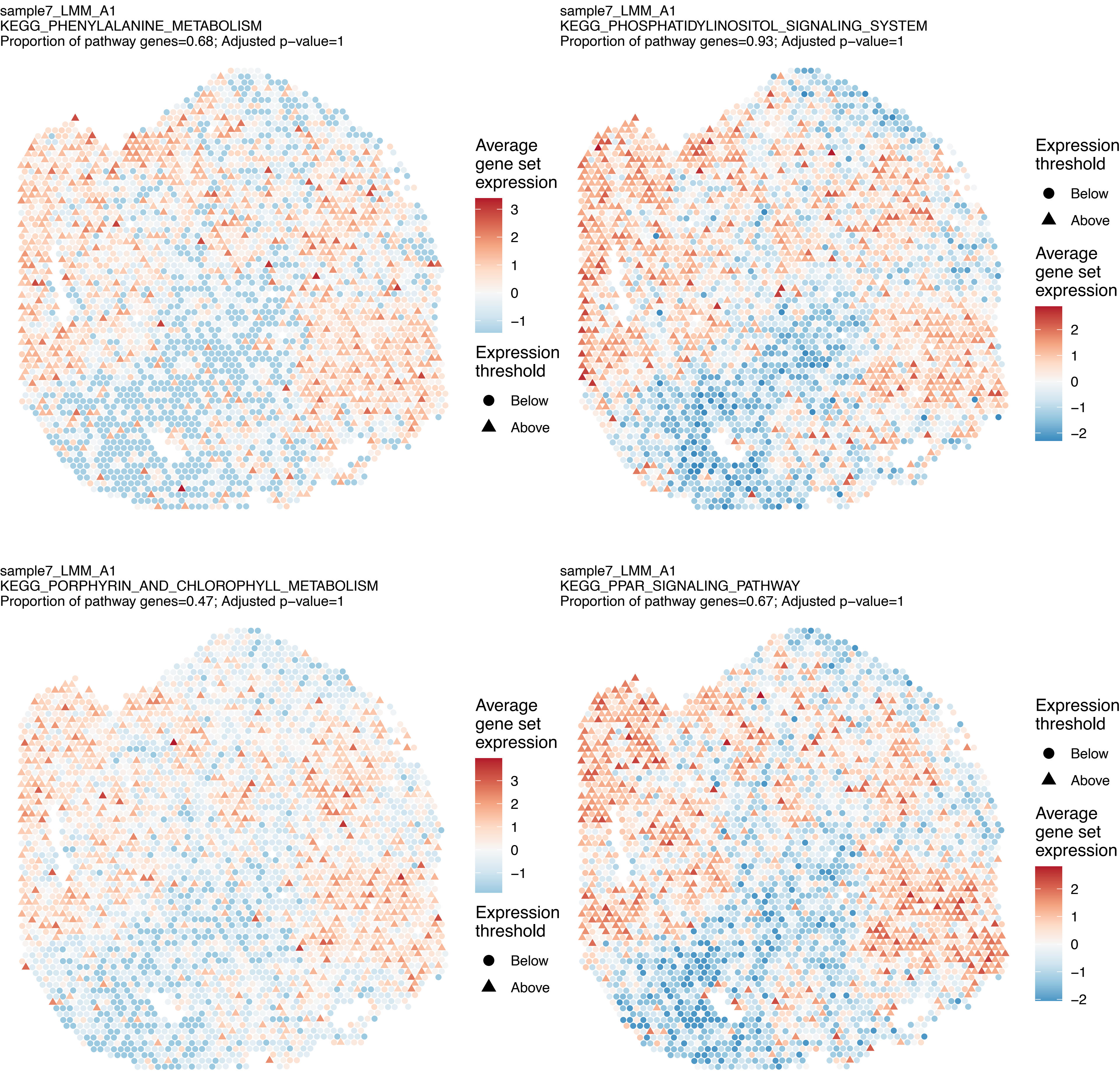

sample7_LMM_A1
KEGG_PHENYLALANINE_METABOLISM
Proportion of pathway genes=0.68; Adjusted p−value=1
Average gene set expression
3
2
1
0
−1
Expression threshold
Below
Above
sample7_LMM_A1
KEGG_PHOSPHATIDYLINOSITOL_SIGNALING_SYSTEM
Proportion of pathway genes=0.93; Adjusted p−value=1
Expression threshold
Below
Above
Average gene set expression
2
1
0
−1
−2
sample7_LMM_A1
KEGG_PORPHYRIN_AND_CHLOROPHYLL_METABOLISM
Proportion of pathway genes=0.47; Adjusted p−value=1
Average gene set expression
3
2
1
0
−1
Expression threshold
Below
Above
sample7_LMM_A1
KEGG_PPAR_SIGNALING_PATHWAY
Proportion of pathway genes=0.67; Adjusted p−value=1
Expression threshold
Below
Above
Average gene set expression
2
1
0
−1
−2

sample7_LMM_A1
KEGG_PRION_DISEASES
Proportion of pathway genes=0.76; Adjusted p–value=1

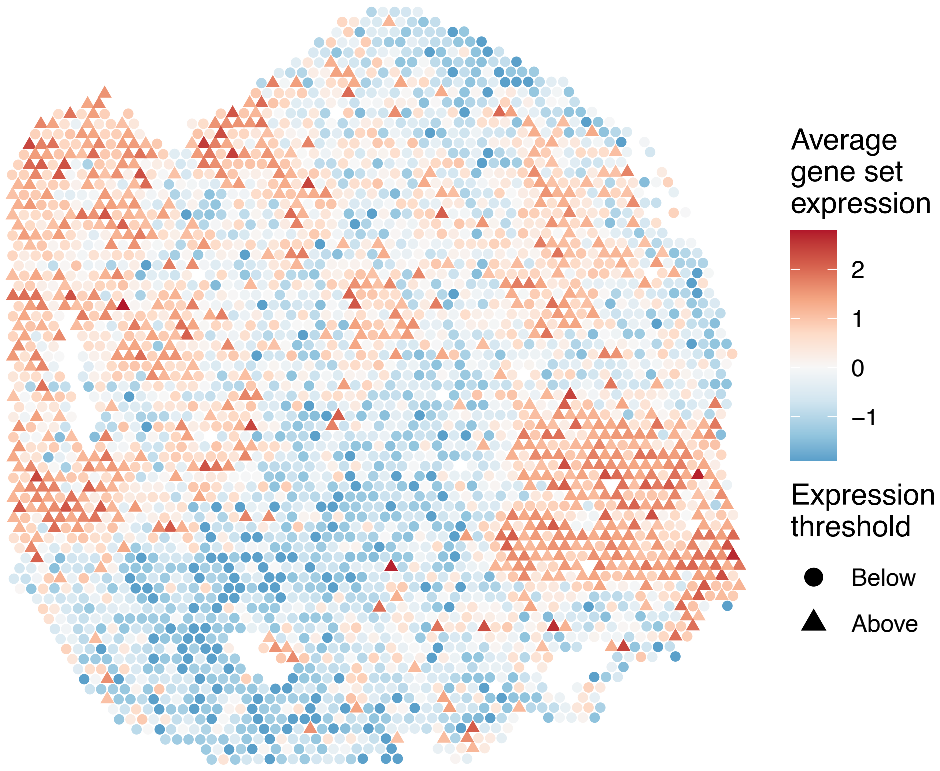


sample7_LMM_A1
KEGG_PROGESTERONE_MEDIATED_OOCYTE_MATURATION
Proportion of pathway genes=0.91; Adjusted p–value=1

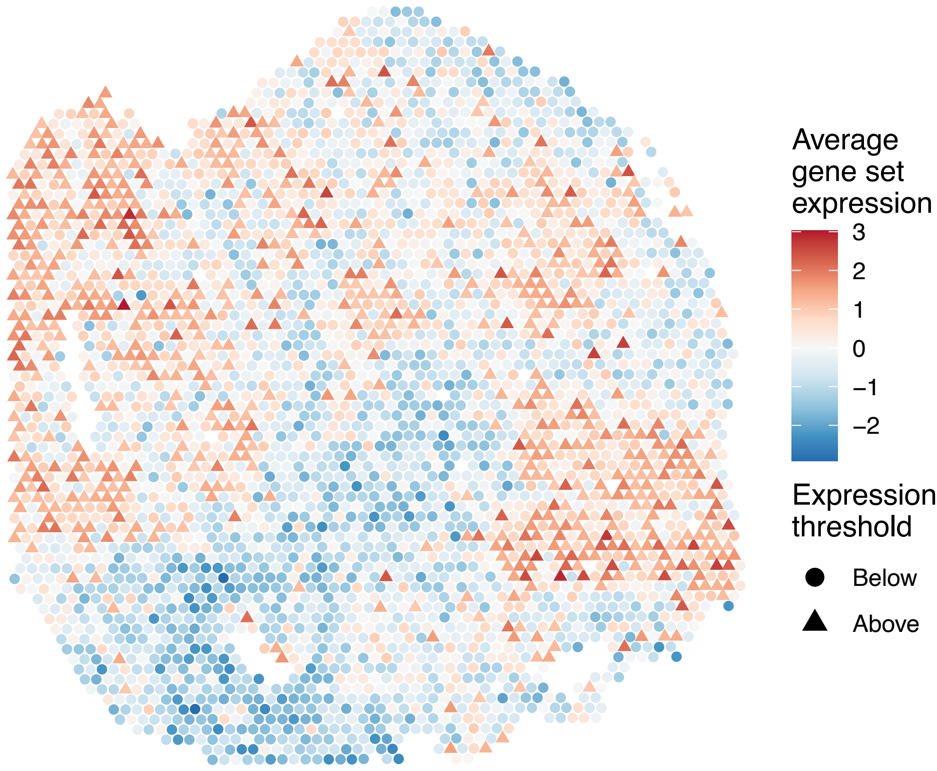


sample7_LMM_A1
KEGG_PROPANOATE_METABOLISM
Proportion of pathway genes=0.89; Adjusted p–value=1

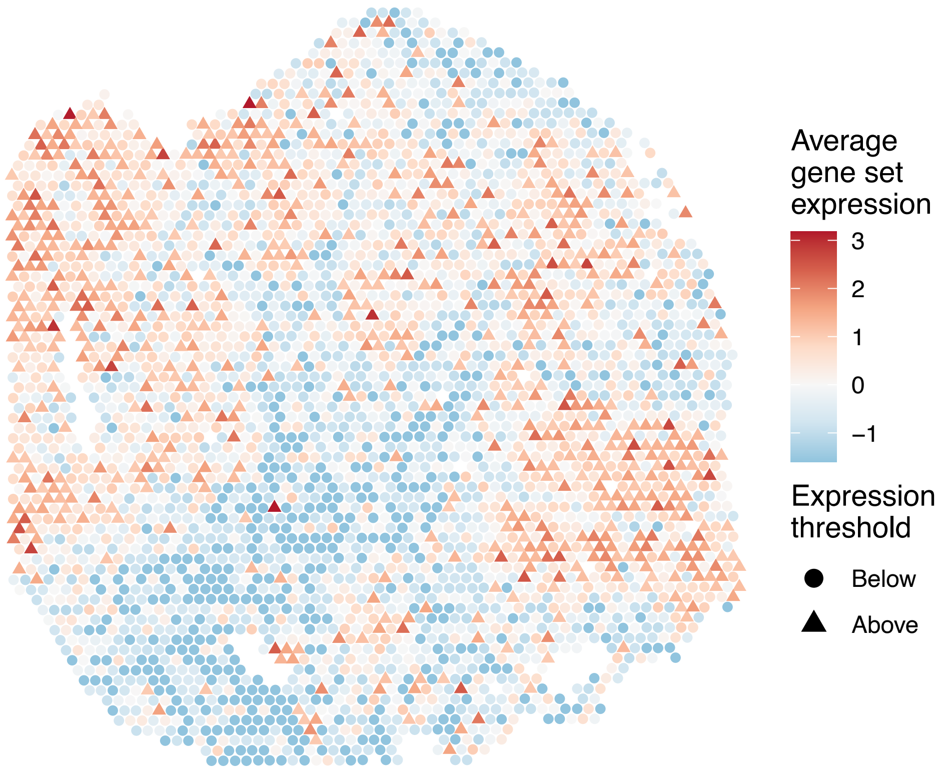


sample7_LMM_A1
KEGG_PROSTATE_CANCER
Proportion of pathway genes=0.9; Adjusted p–value=1

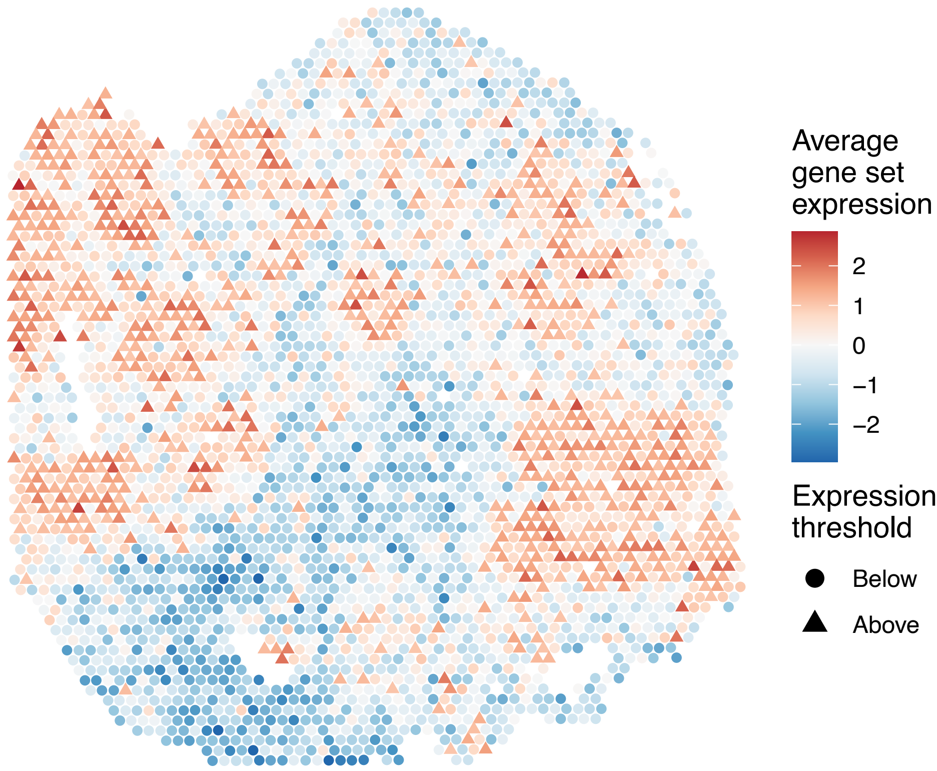

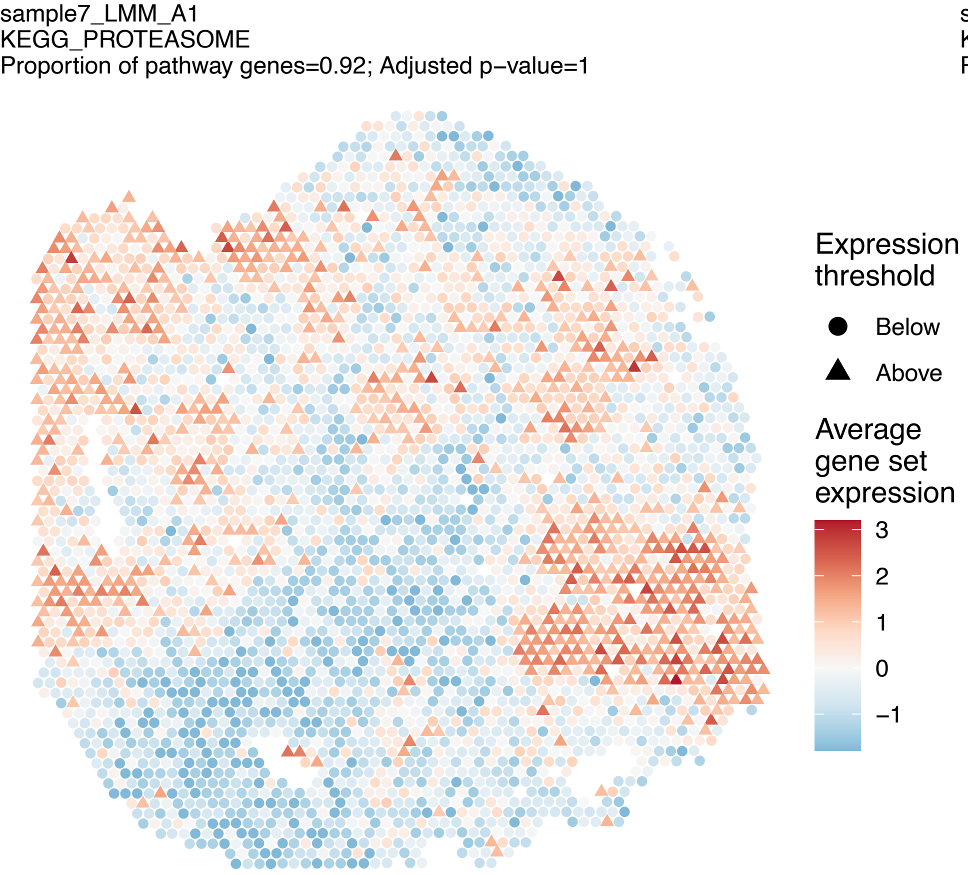
sample7_LMM_A1
KEGG_PROTEASOME
Proportion of pathway genes=0.92; Adjusted p–value=1
Expression threshold
Below
Above
Average gene set expression
3
2
1
0
–1

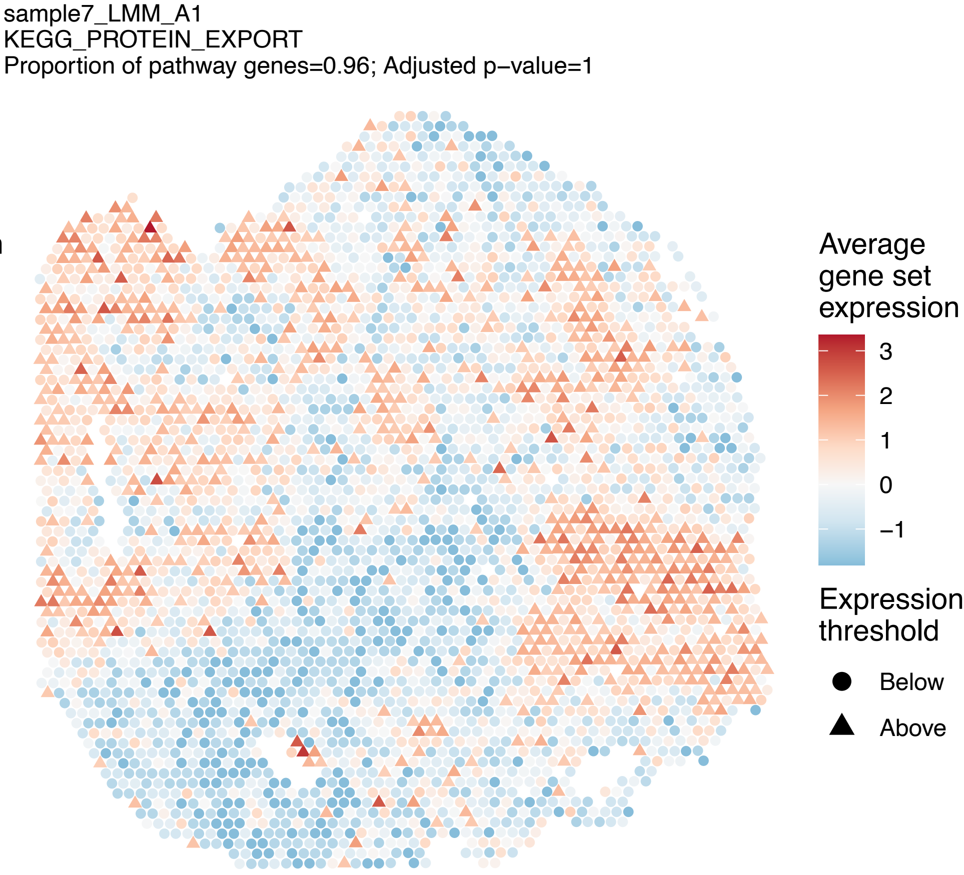
sample7_LMM_A1
KEGG_PROTEIN_EXPORT
Proportion of pathway genes=0.96; Adjusted p–value=1
Average gene set expression
3
2
1
0
–1
Expression threshold
Below
Above

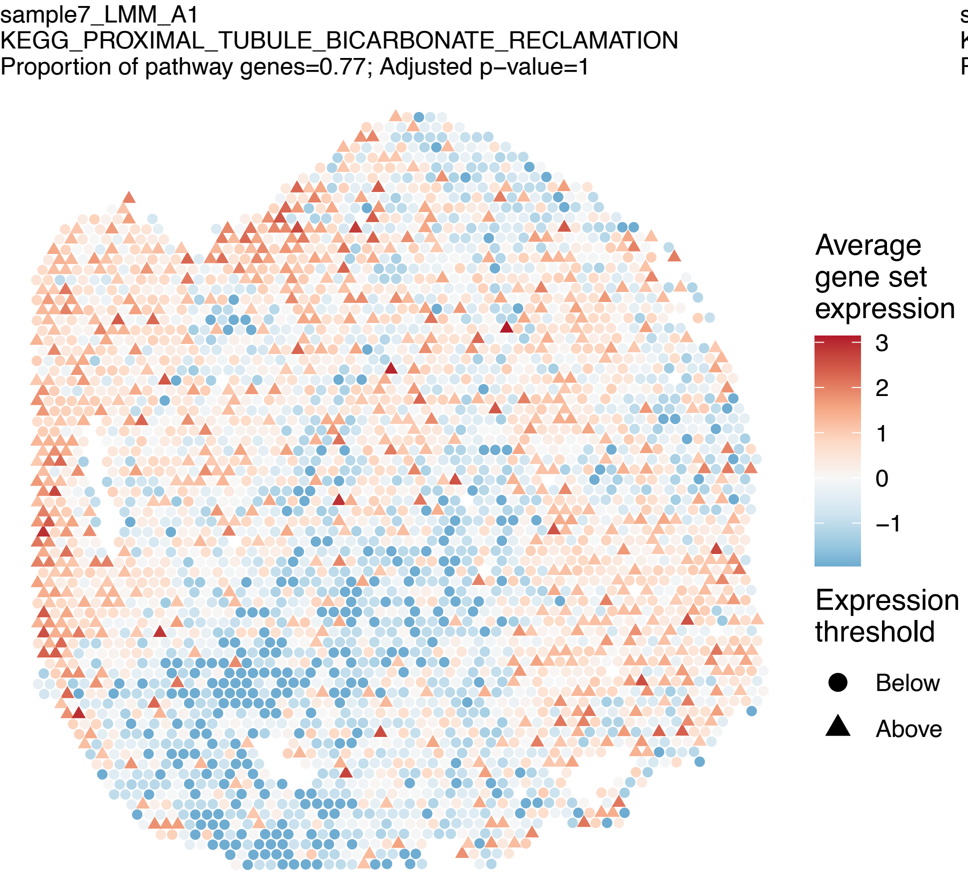
sample7_LMM_A1
KEGG_PROXIMAL_TUBULE_BICARBONATE_RECLAMATION
Proportion of pathway genes=0.77; Adjusted p–value=1
Average gene set expression
3
2
1
0
–1
Expression threshold
Below
Above

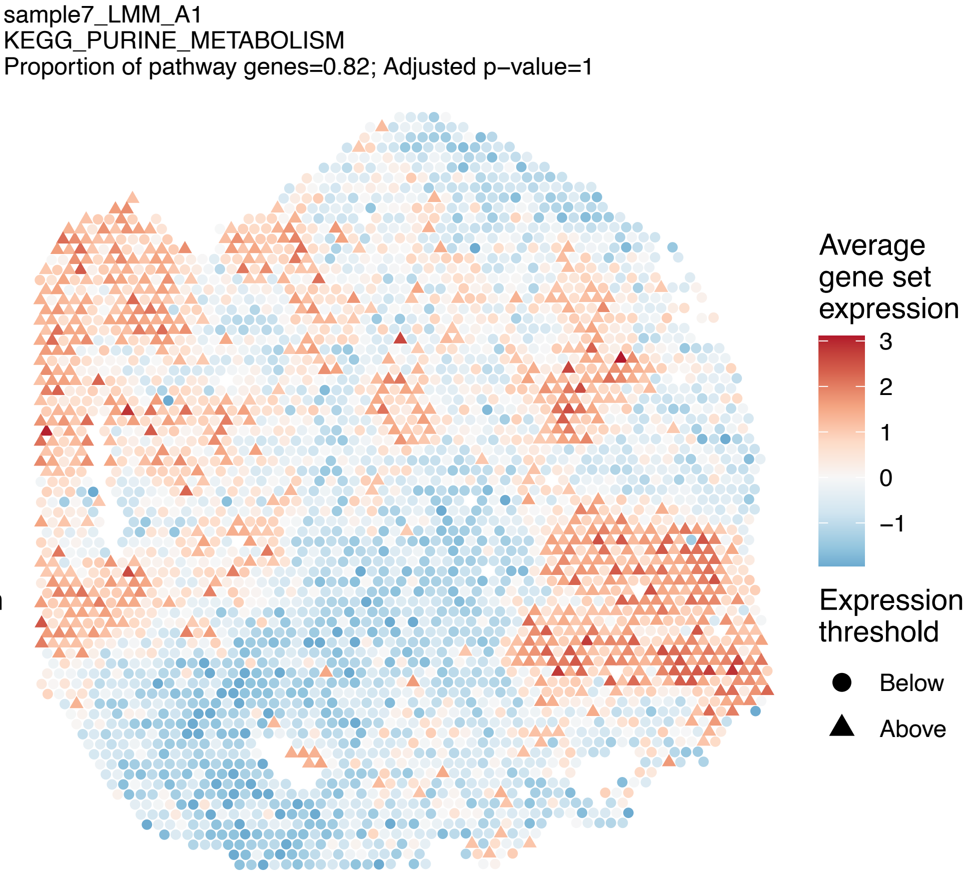
sample7_LMM_A1
KEGG_PURINE_METABOLISM
Proportion of pathway genes=0.82; Adjusted p–value=1
Average gene set expression
3
2
1
0
–1
Expression threshold
Below
Above

sample7_LMM_A1
KEGG_PYRIMIDINE_METABOLISM
Proportion of pathway genes=0.83; Adjusted p−value=1

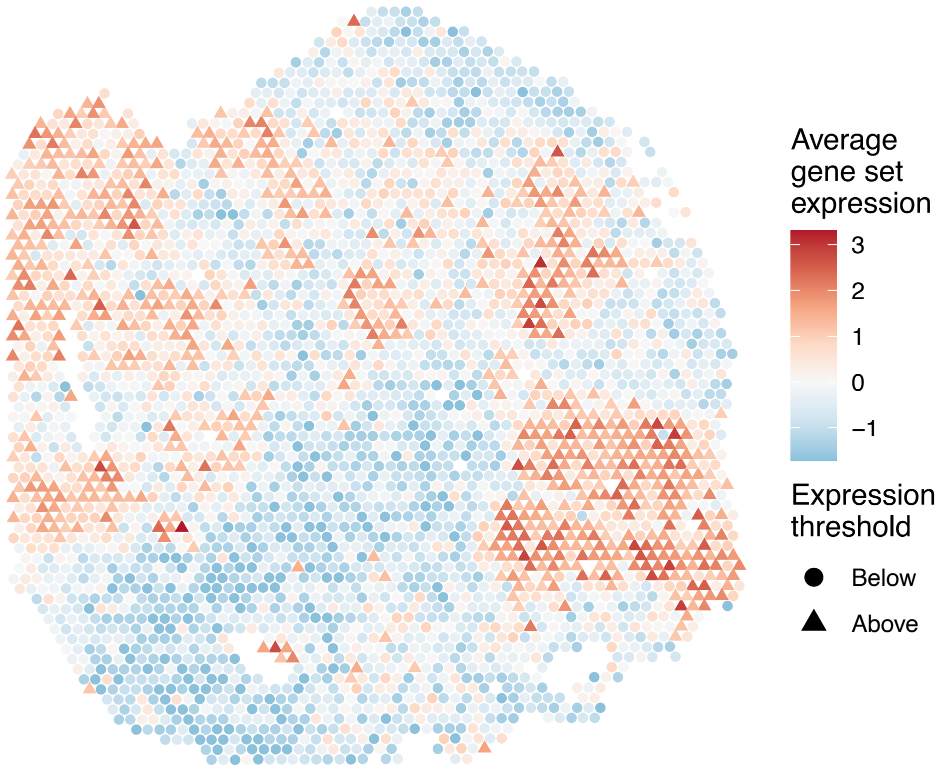


sample7_LMM_A1
KEGG_PYRUVATE_METABOLISM
Proportion of pathway genes=0.86; Adjusted p−value=1

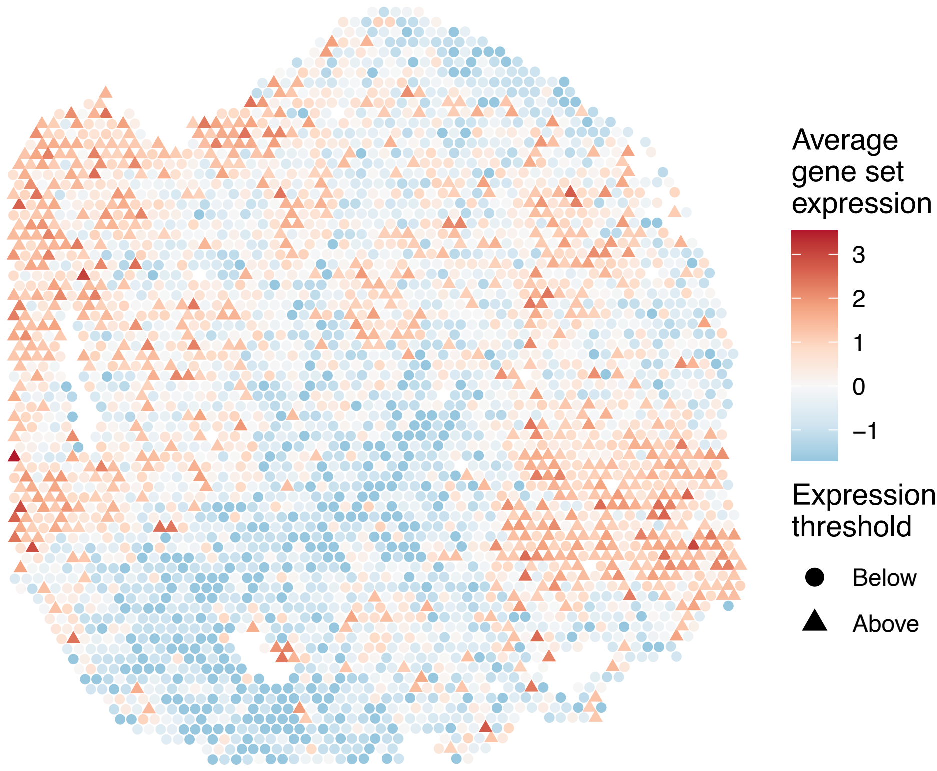


sample7_LMM_A1
KEGG_REGULATION_OF_ACTIN_CYTOSKELETON
Proportion of pathway genes=0.84; Adjusted p−value=1

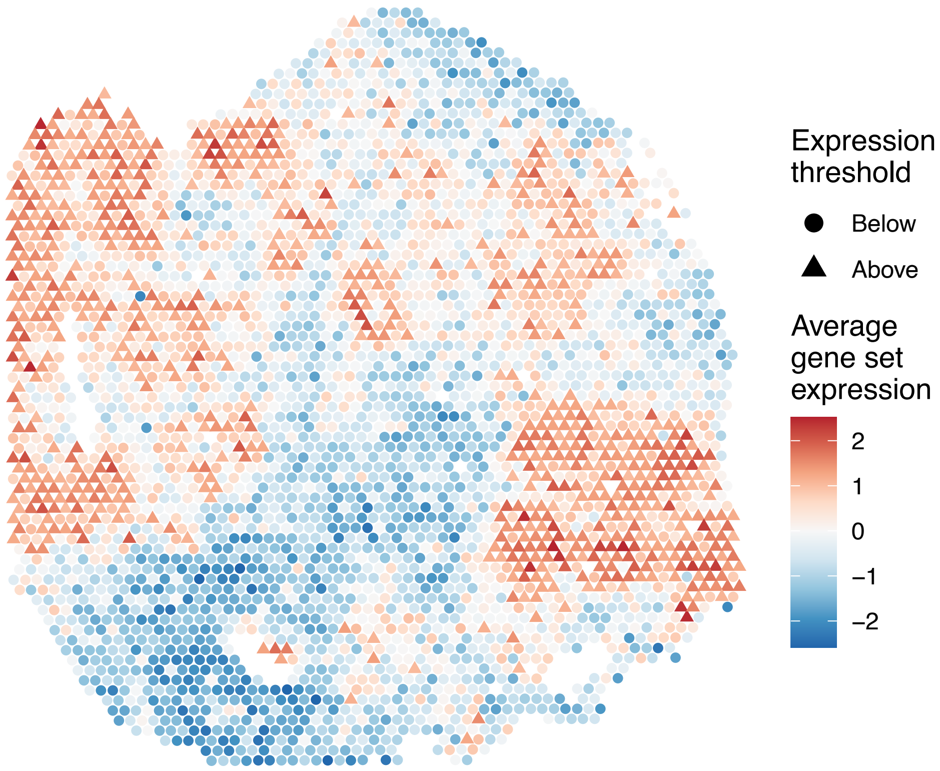


sample7_LMM_A1
KEGG_REGULATION_OF_AUTOPHAGY
Proportion of pathway genes=0.54; Adjusted p−value=1

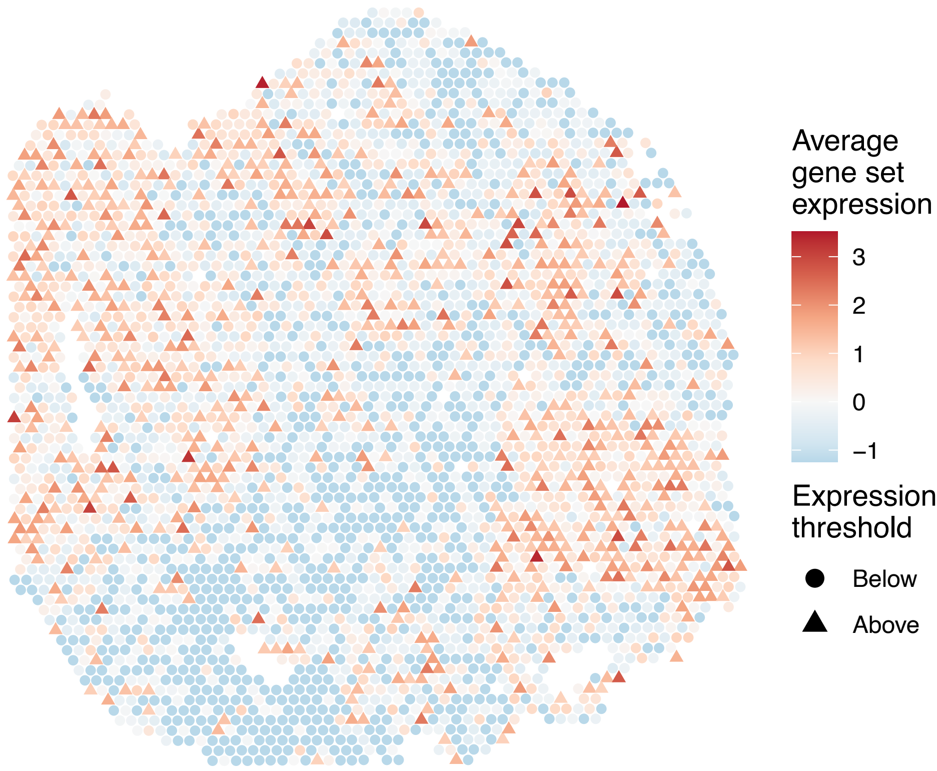

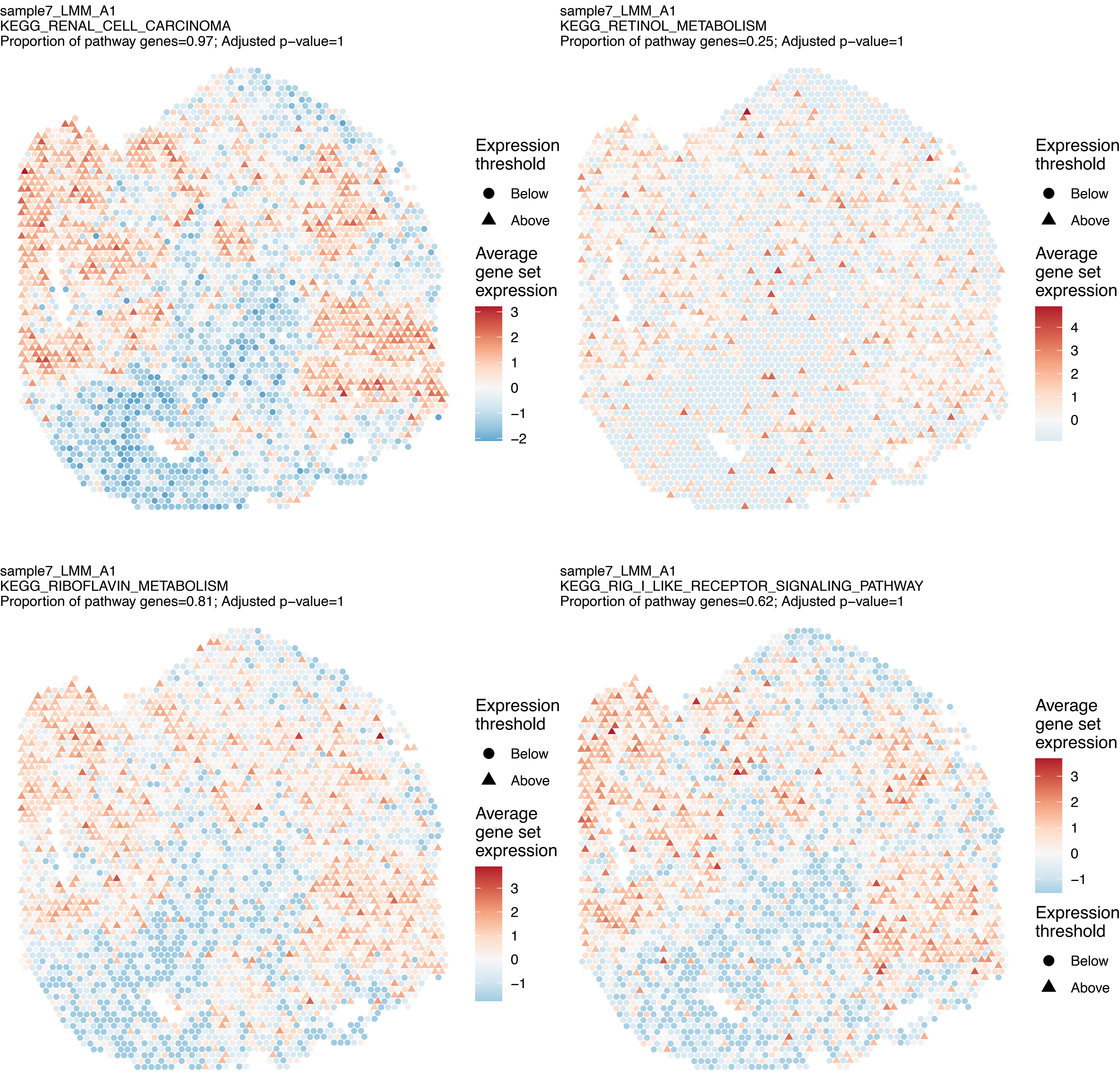
sample7_LMM_A1
KEGG_RENAL_CELL_CARCINOMA
Proportion of pathway genes=0.97; Adjusted p–value=1
Expression threshold
Below
Above
Average gene set expression
3
2
1
0
−1
−2
sample7_LMM_A1
KEGG_RETINOL_METABOLISM
Proportion of pathway genes=0.25; Adjusted p–value=1
Expression threshold
Below
Above
Average gene set expression
4
3
2
1
0
sample7_LMM_A1
KEGG_RIBOFLAVIN_METABOLISM
Proportion of pathway genes=0.81; Adjusted p–value=1
Expression threshold
Below
Above
Average gene set expression
3
2
1
0
−1
sample7_LMM_A1
KEGG_RIG_I_LIKE_RECEPTOR_SIGNALING_PATHWAY
Proportion of pathway genes=0.62; Adjusted p–value=1
Average gene set expression
3
2
1
0
−1
Expression threshold
Below
Above

sample7_LMM_A1
KEGG_RNA_DEGRADATION
Proportion of pathway genes=0.96; Adjusted p−value=1

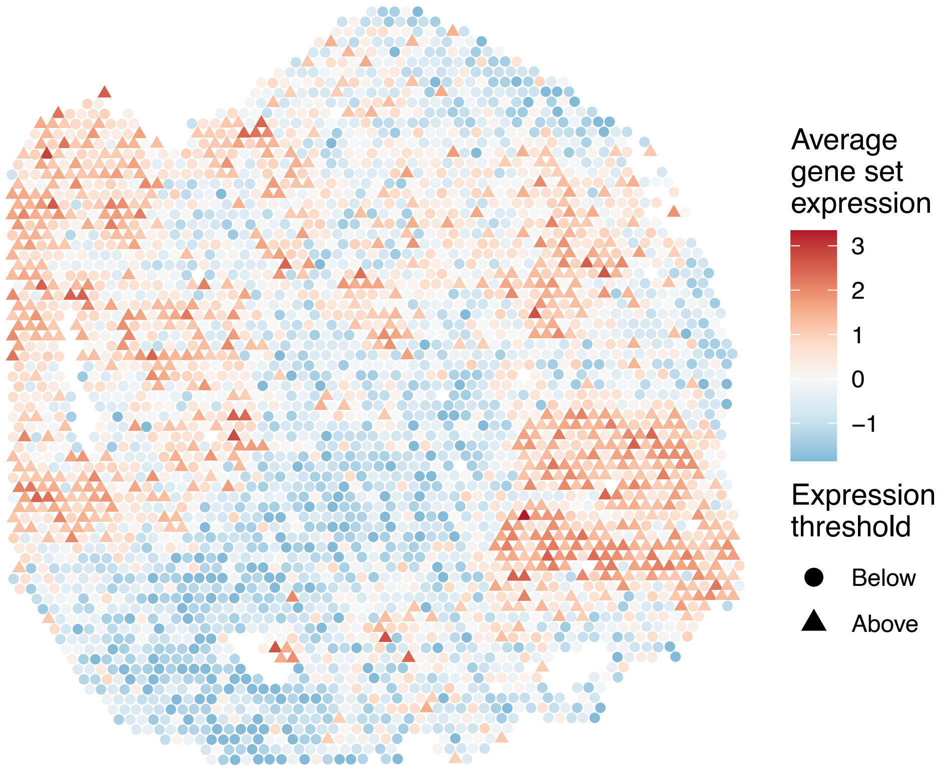


sample7_LMM_A1
KEGG_RNA_POLYMERASE
Proportion of pathway genes=0.74; Adjusted p−value=1

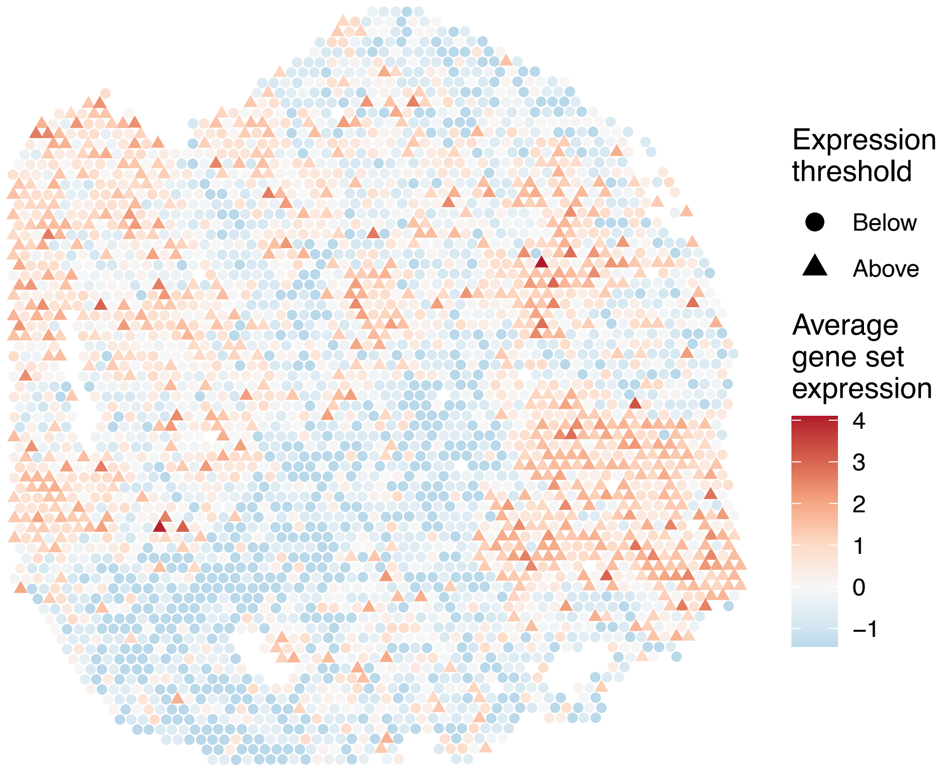


sample7_LMM_A1
KEGG_SELENOAMINO_ACID_METABOLISM
Proportion of pathway genes=0.85; Adjusted p−value=1

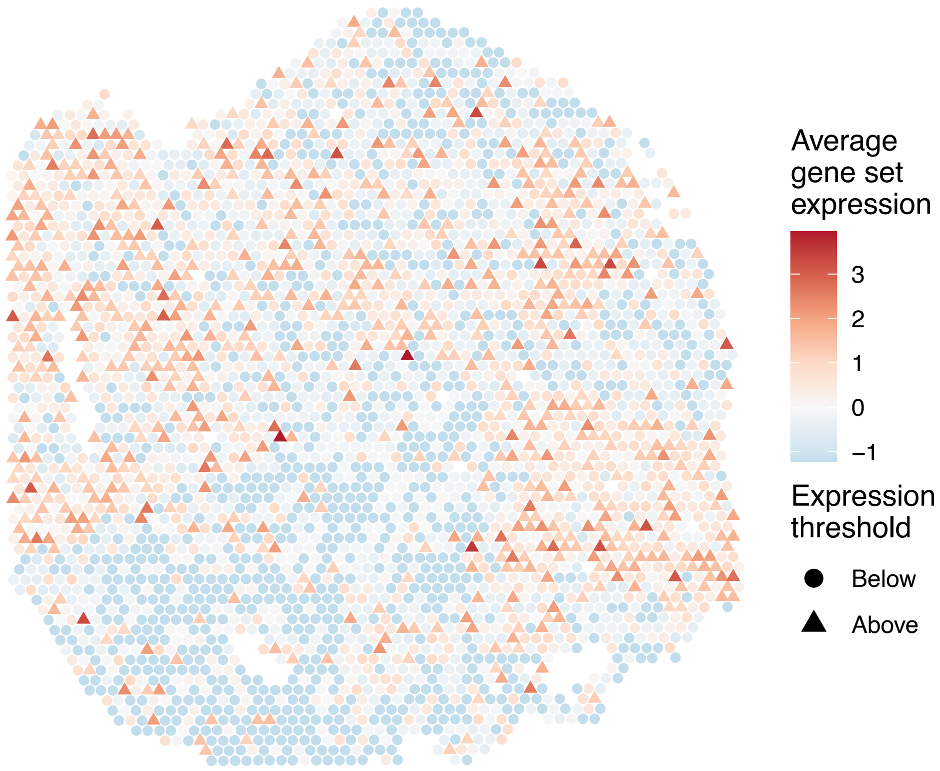


sample7_LMM_A1
KEGG_SMALL_CELL_LUNG_CANCER
Proportion of pathway genes=0.9; Adjusted p−value=1

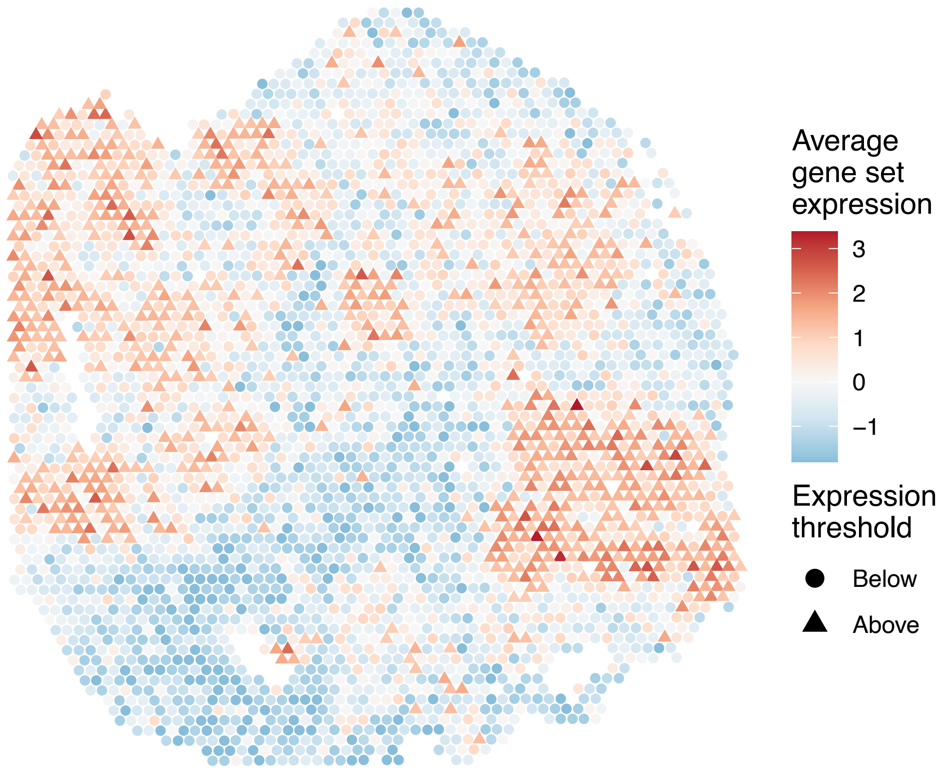

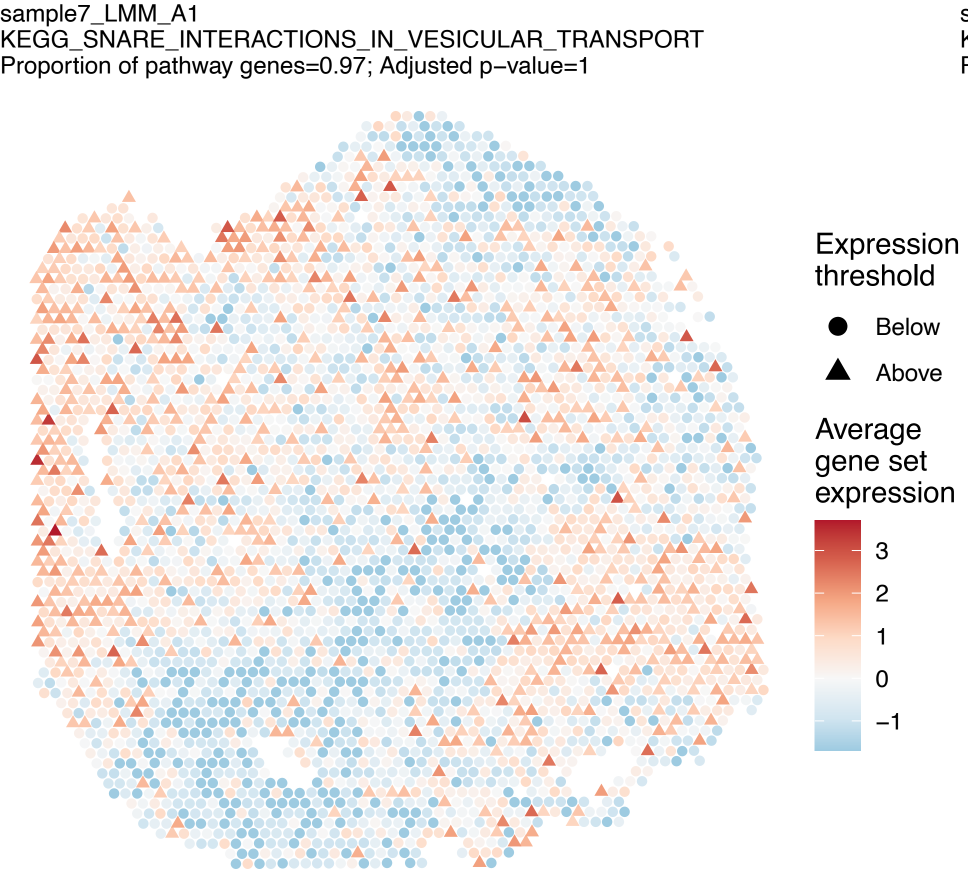
sample7_LMM_A1
KEGG_SNARE_INTERACTIONS_IN_VESICULAR_TRANSPORT
Proportion of pathway genes=0.97; Adjusted p–value=1
Expression threshold
Below
Above
Average gene set expression
3
2
1
0
–1

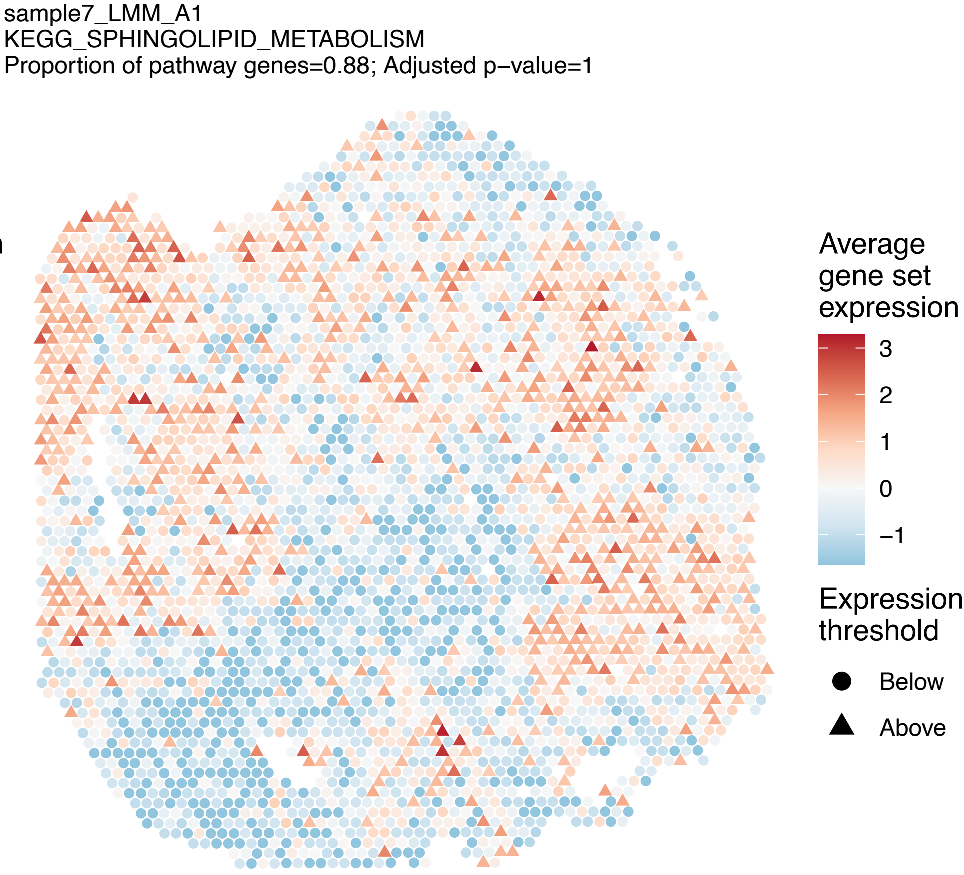
sample7_LMM_A1
KEGG_SPHINGOLIPID_METABOLISM
Proportion of pathway genes=0.88; Adjusted p–value=1
Average gene set expression
3
2
1
0
–1
Expression threshold
Below
Above

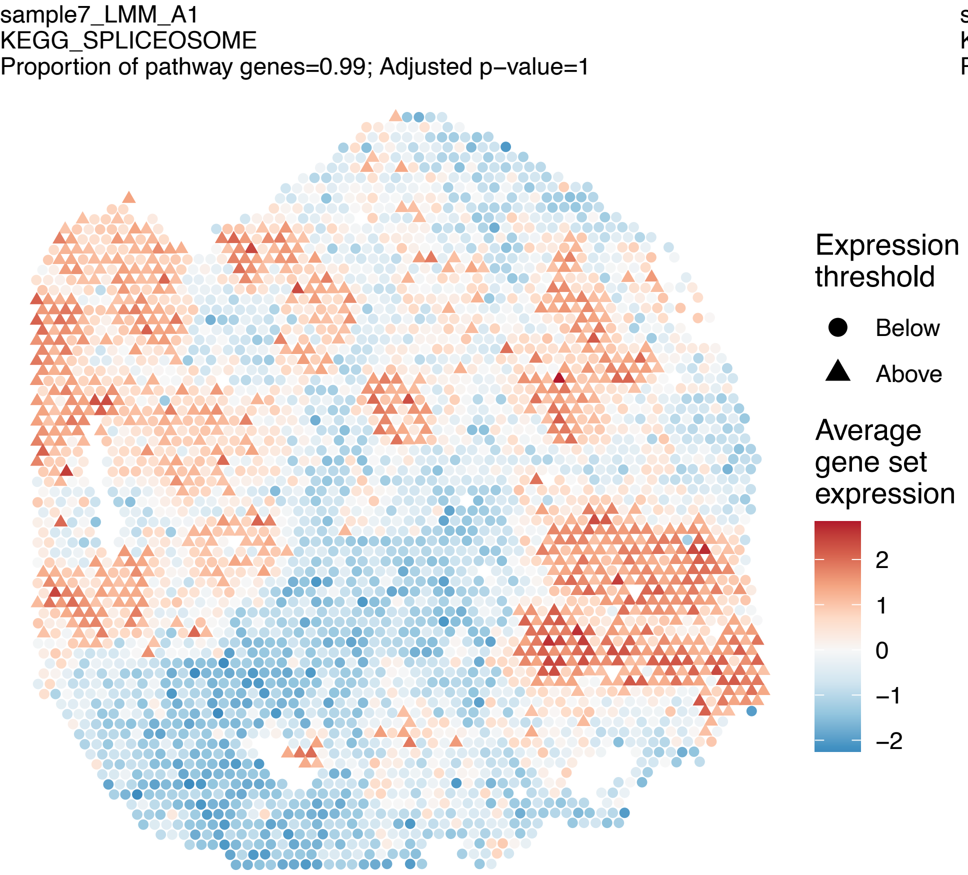
sample7_LMM_A1
KEGG_SPLICEOSOME
Proportion of pathway genes=0.99; Adjusted p–value=1
Expression threshold
Below
Above
Average gene set expression
2
1
0
–1
–2

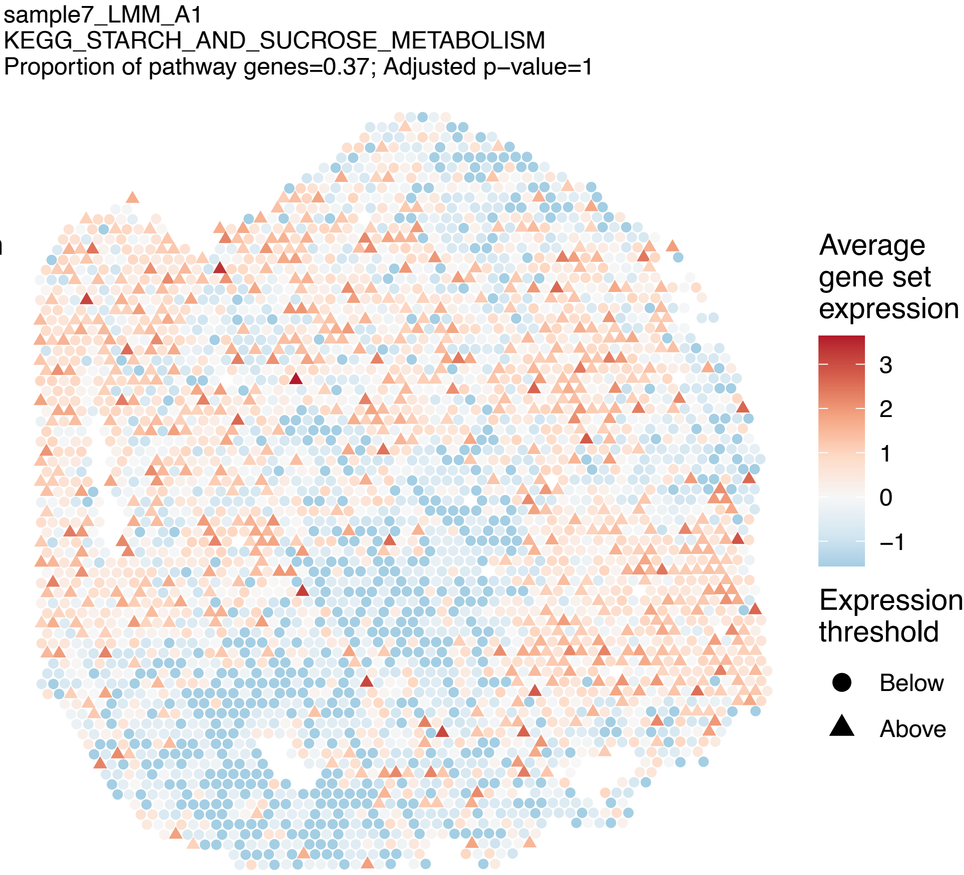
sample7_LMM_A1
KEGG_STARCH_AND_SUCROSE_METABOLISM
Proportion of pathway genes=0.37; Adjusted p–value=1
Average gene set expression
3
2
1
0
–1
Expression threshold
Below
Above

sample7_LMM_A1
KEGG_STEROID_BIOSYNTHESIS
Proportion of pathway genes=0.89; Adjusted p−value=1

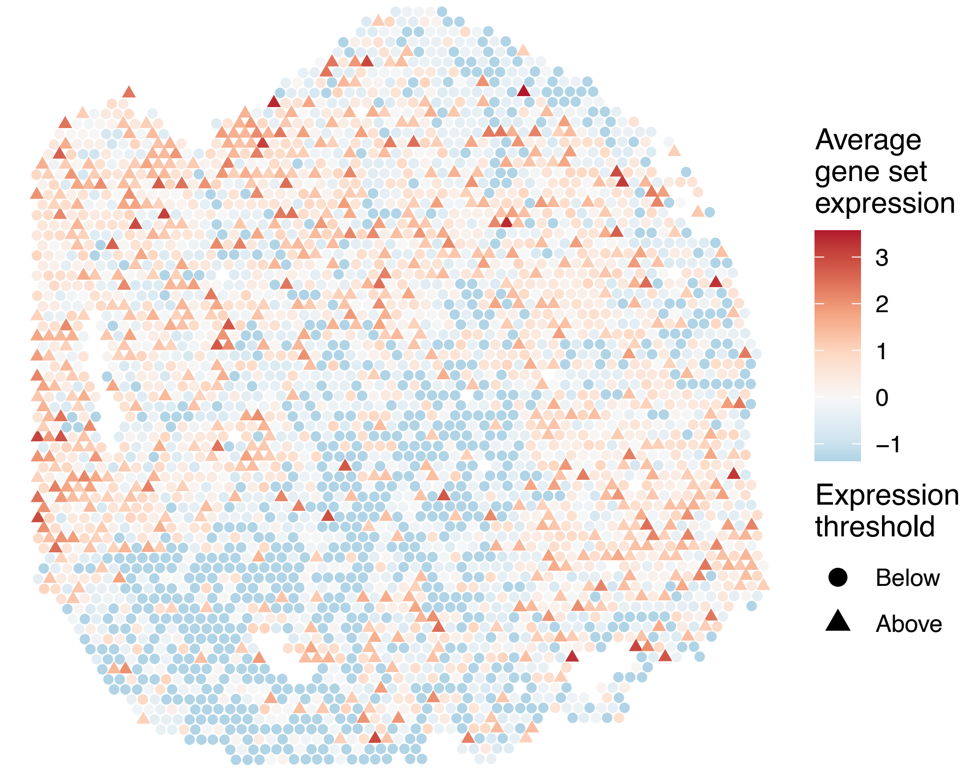


sample7_LMM_A1
KEGG_STEROID_HORMONE_BIOSYNTHESIS
Proportion of pathway genes=0.35; Adjusted p−value=1

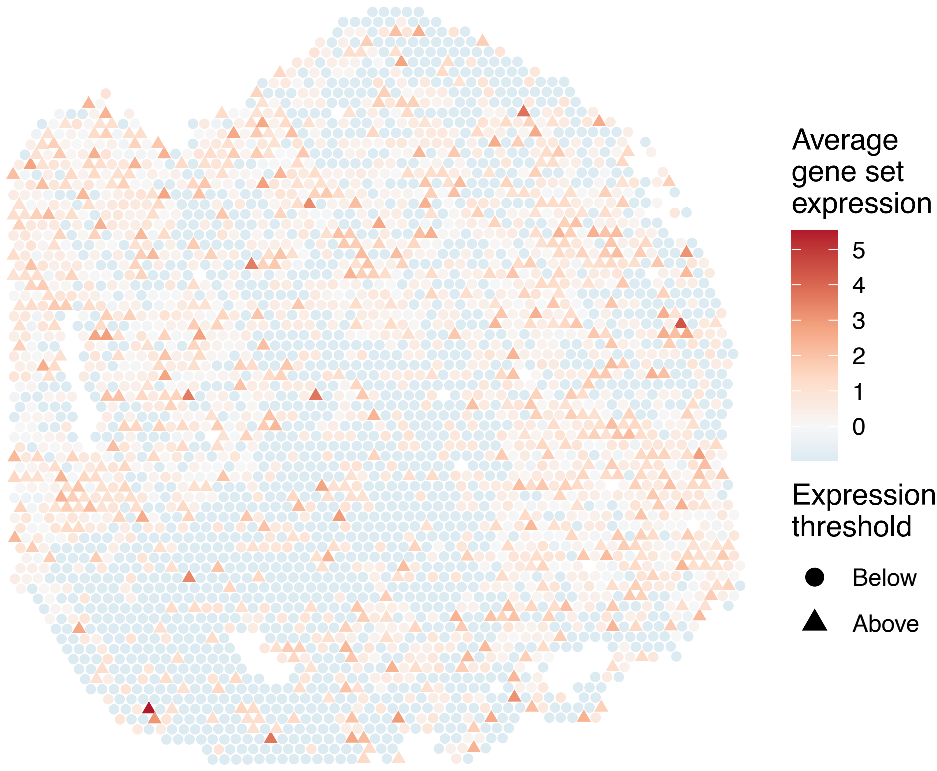


sample7_LMM_A1
KEGG_SULFUR_METABOLISM
Proportion of pathway genes=0.77; Adjusted p−value=1

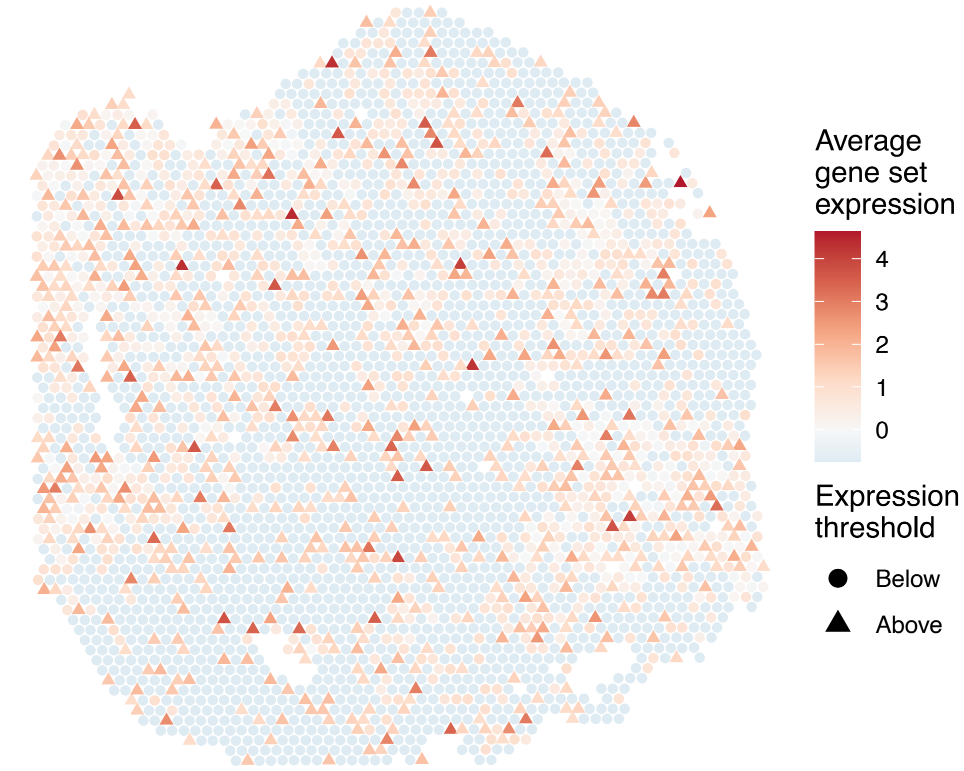


sample7_LMM_A1
KEGG_SYSTEMIC_LUPUS_ERYTHEMATOSUS
Proportion of pathway genes=0.43; Adjusted p−value=1

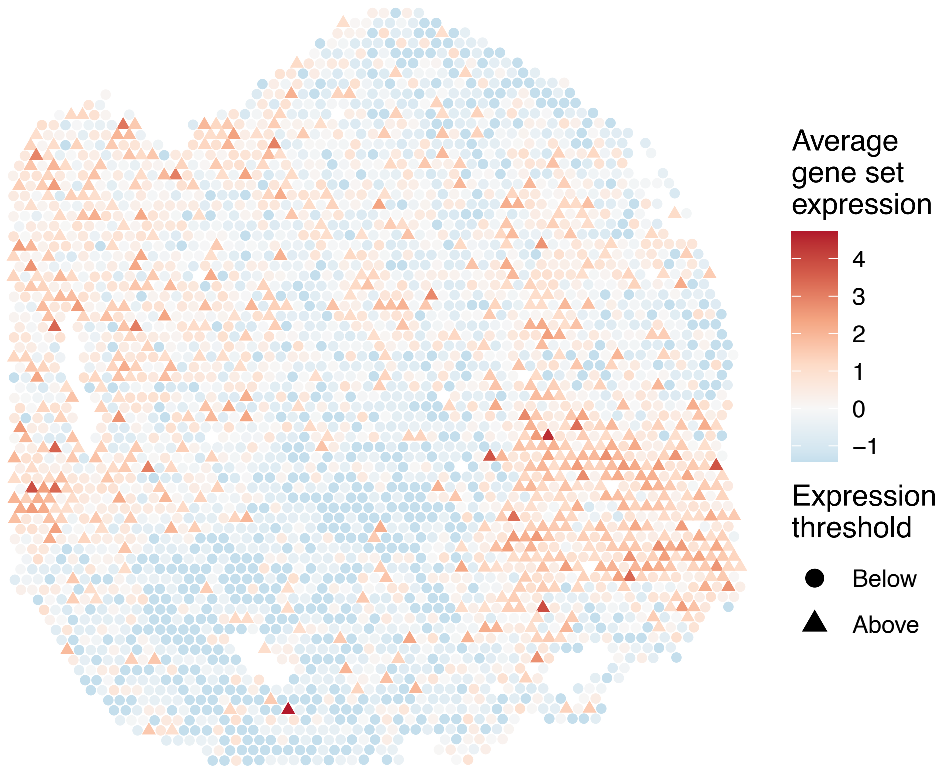

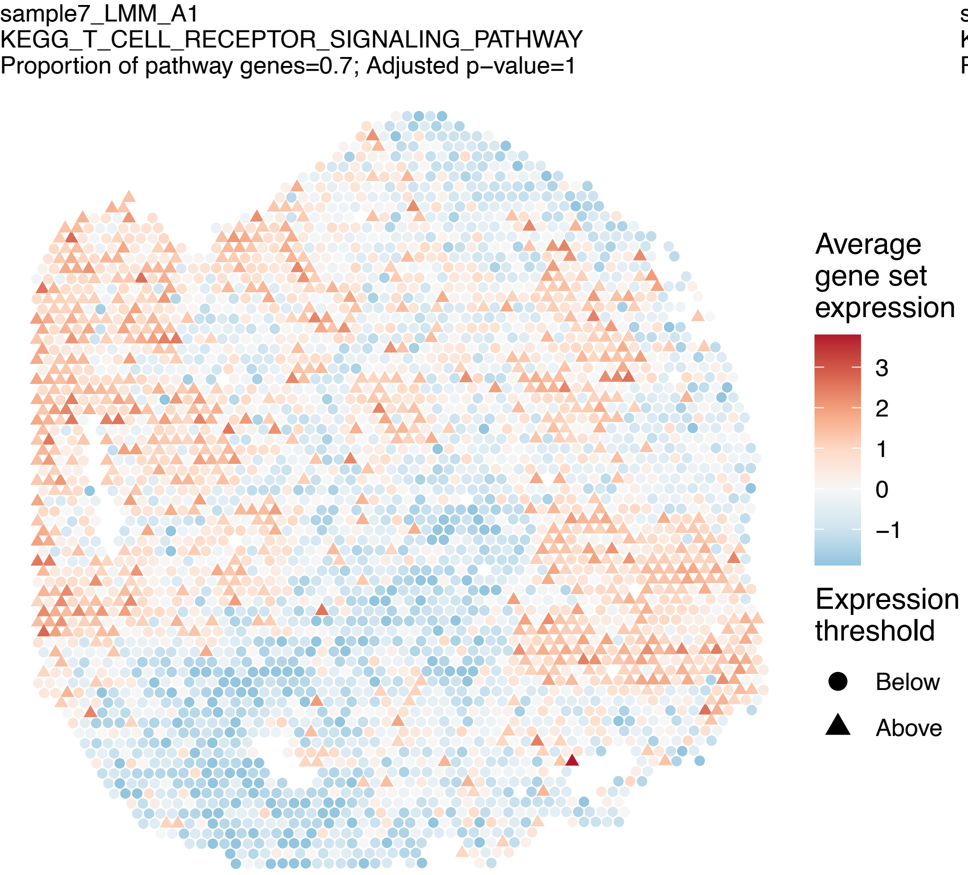
sample7_LMM_A1
KEGG_T_CELL_RECEPTOR_SIGNALING_PATHWAY
Proportion of pathway genes=0.7; Adjusted p−value=1
Average gene set expression
3
2
1
0
−1
Expression threshold
Below
Above

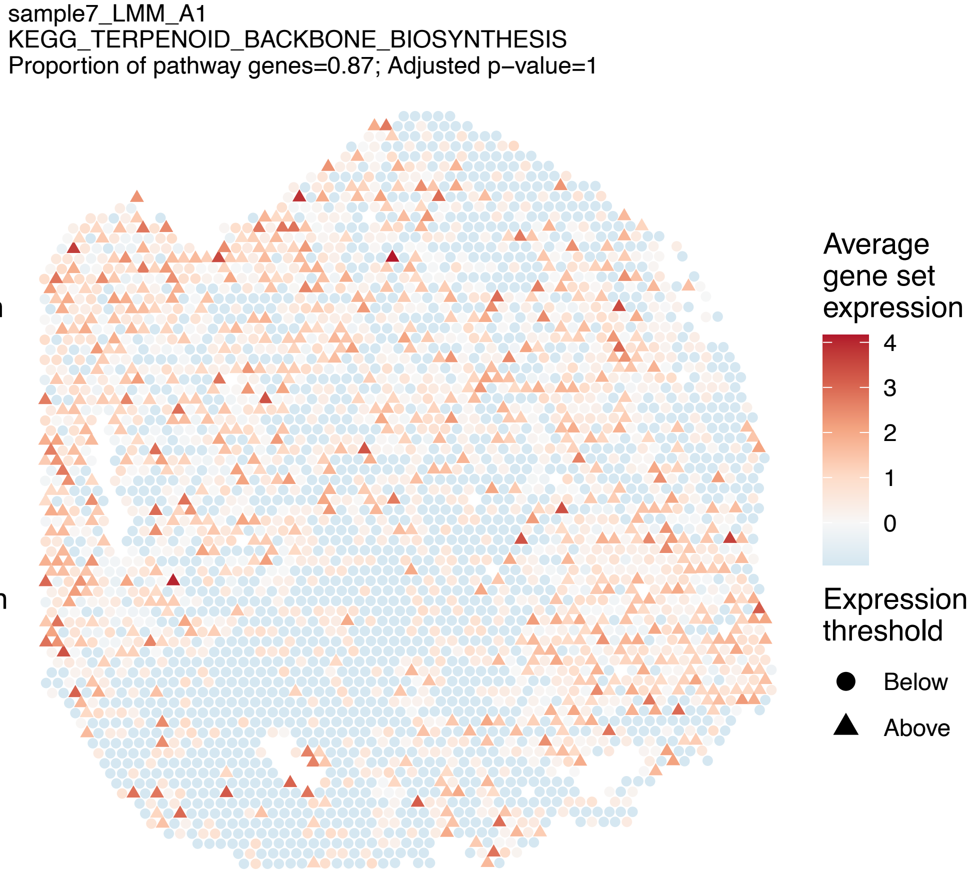
sample7_LMM_A1
KEGG_TERPENOID_BACKBONE_BIOSYNTHESIS
Proportion of pathway genes=0.87; Adjusted p−value=1
Average gene set expression
4
3
2
1
0
Expression threshold
Below
Above

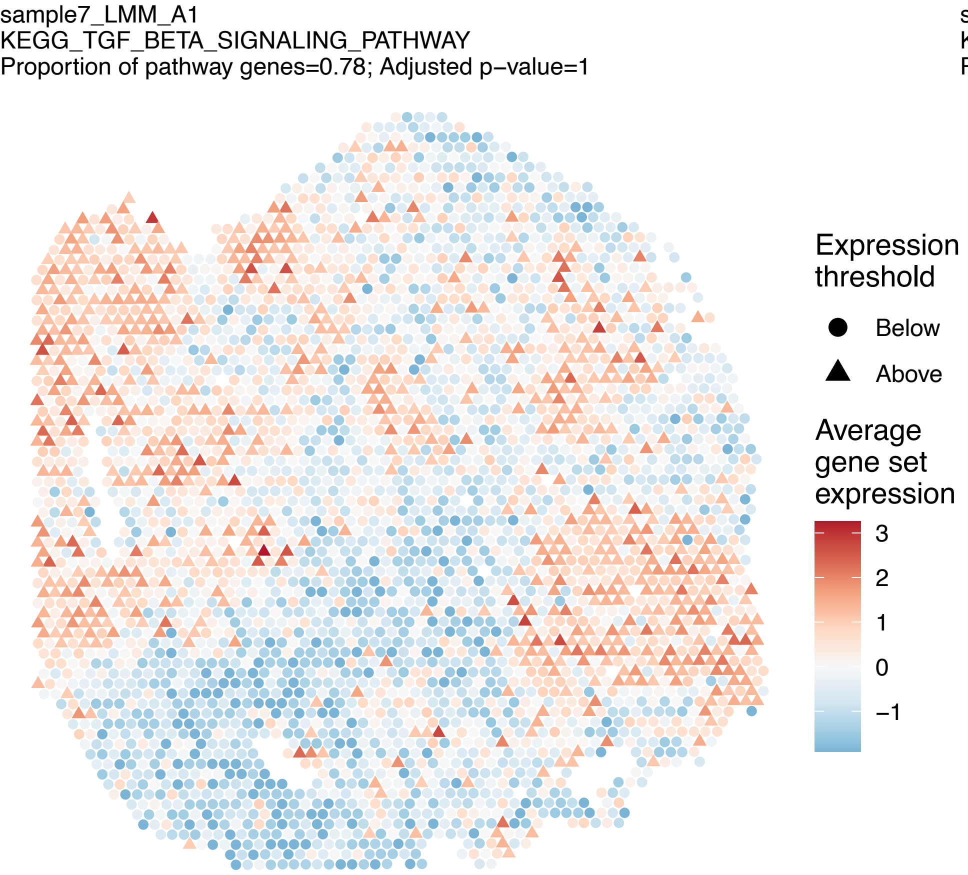
sample7_LMM_A1
KEGG_TGF_BETA_SIGNALING_PATHWAY
Proportion of pathway genes=0.78; Adjusted p−value=1
Expression threshold
Below
Above
Average gene set expression
3
2
1
0
−1

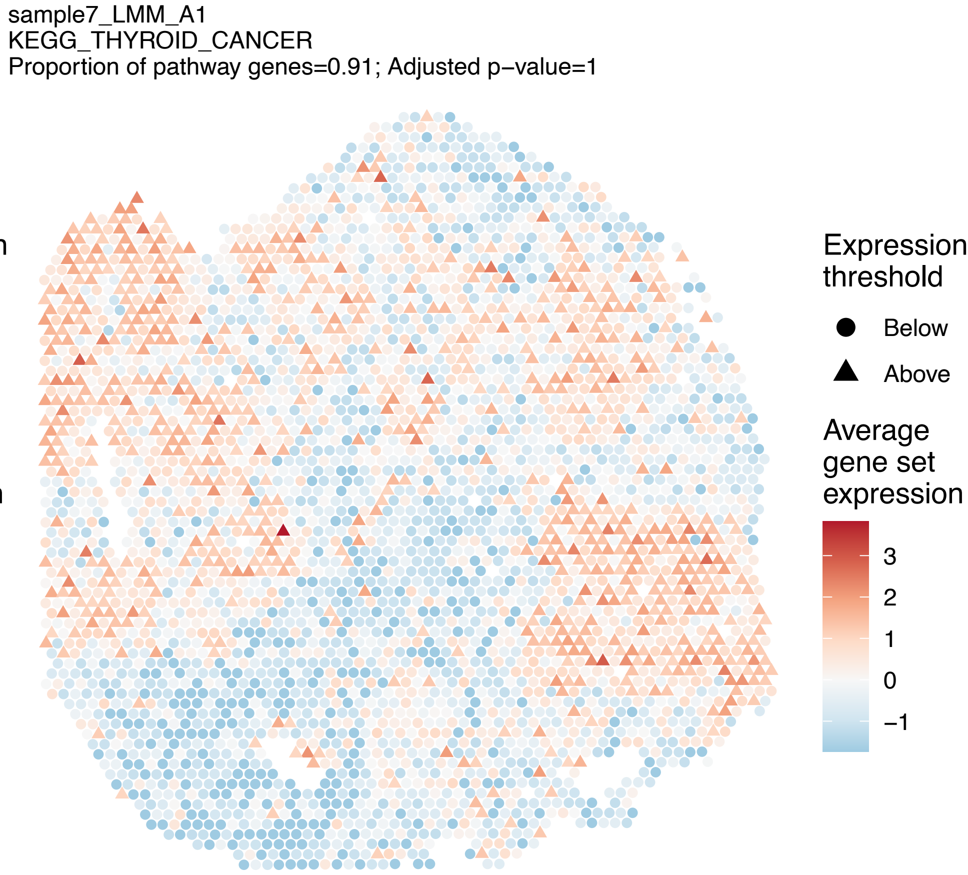
sample7_LMM_A1
KEGG_THYROID_CANCER
Proportion of pathway genes=0.91; Adjusted p−value=1
Expression threshold
Below
Above
Average gene set expression
3
2
1
0
−1

sample7_LMM_A1
KEGG_TIGHT_JUNCTION
Proportion of pathway genes=0.78; Adjusted p−value=1

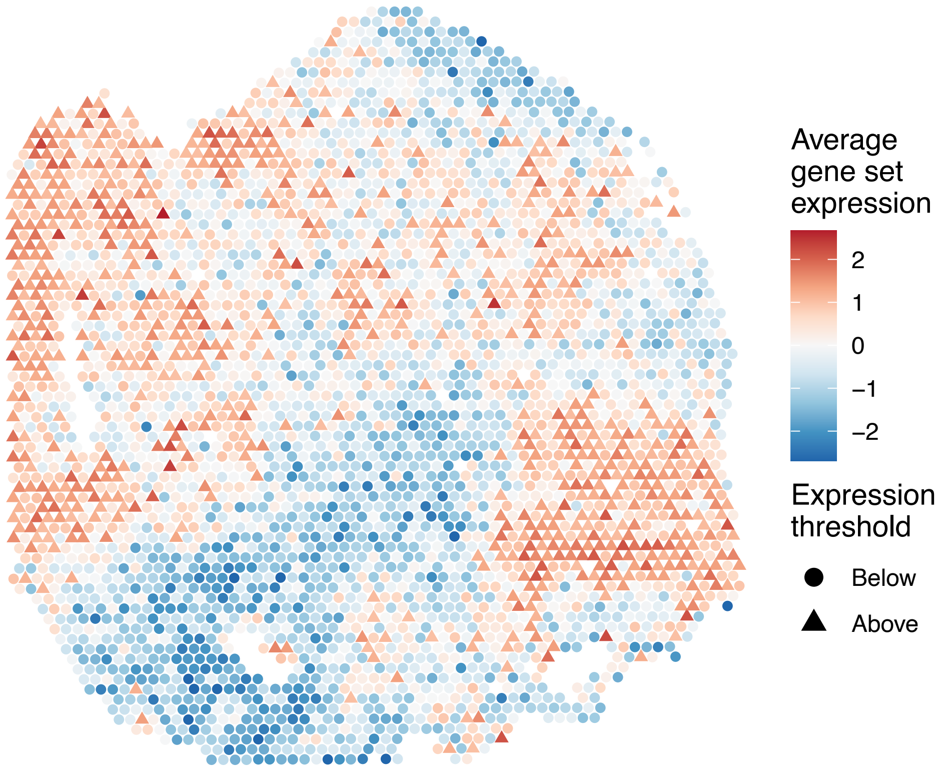


sample7_LMM_A1
KEGG_TOLL_LIKE_RECEPTOR_SIGNALING_PATHWAY
Proportion of pathway genes=0.62; Adjusted p−value=1

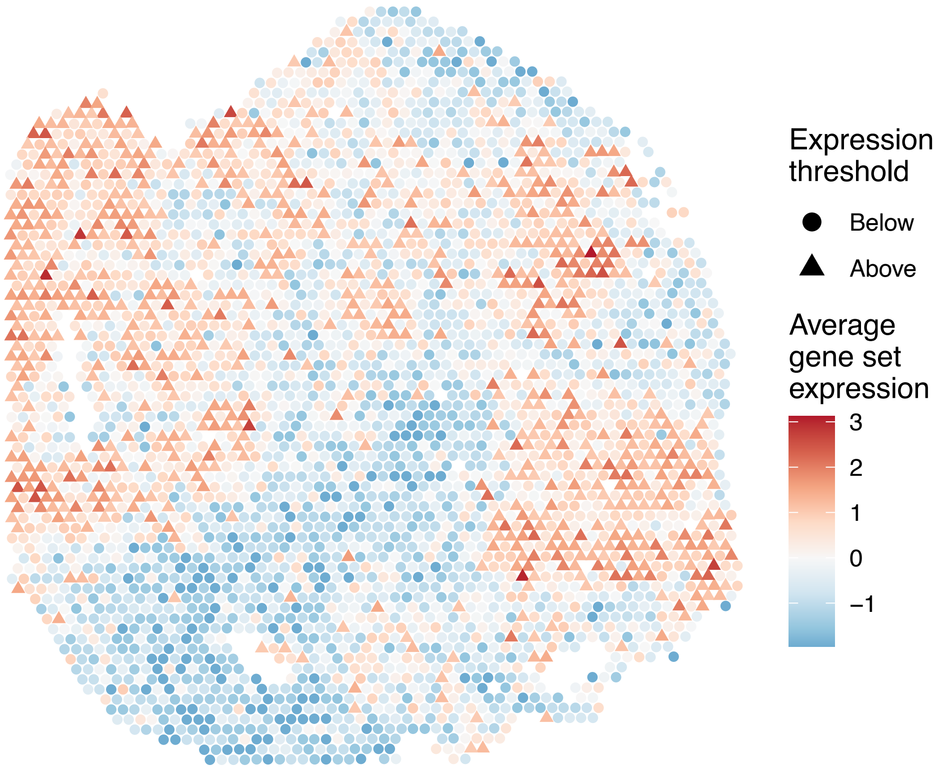


sample7_LMM_A1
KEGG_TRYPTOPHAN_METABOLISM
Proportion of pathway genes=0.51; Adjusted p−value=1

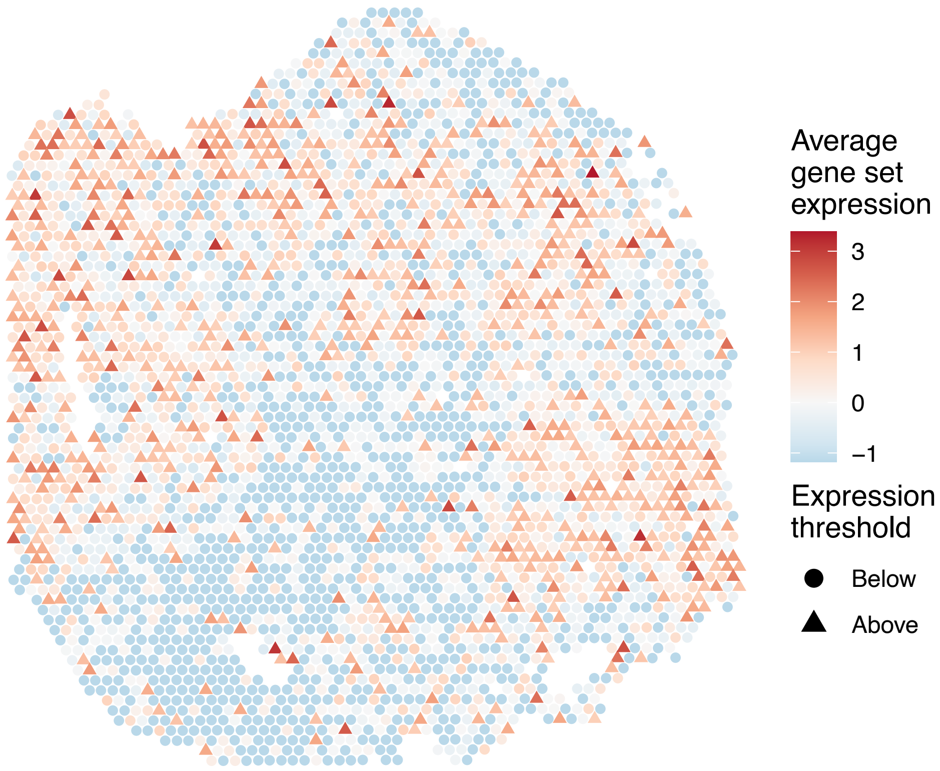


sample7_LMM_A1
KEGG_TYPE_I_DIABETES_MELLITUS
Proportion of pathway genes=0.72; Adjusted p−value=1

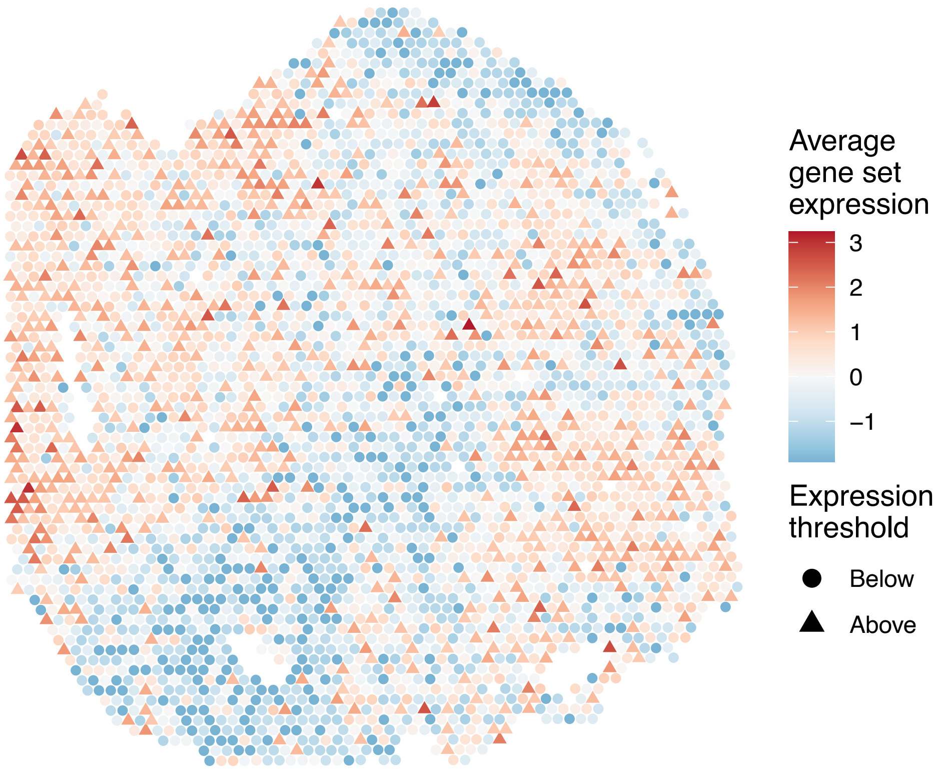

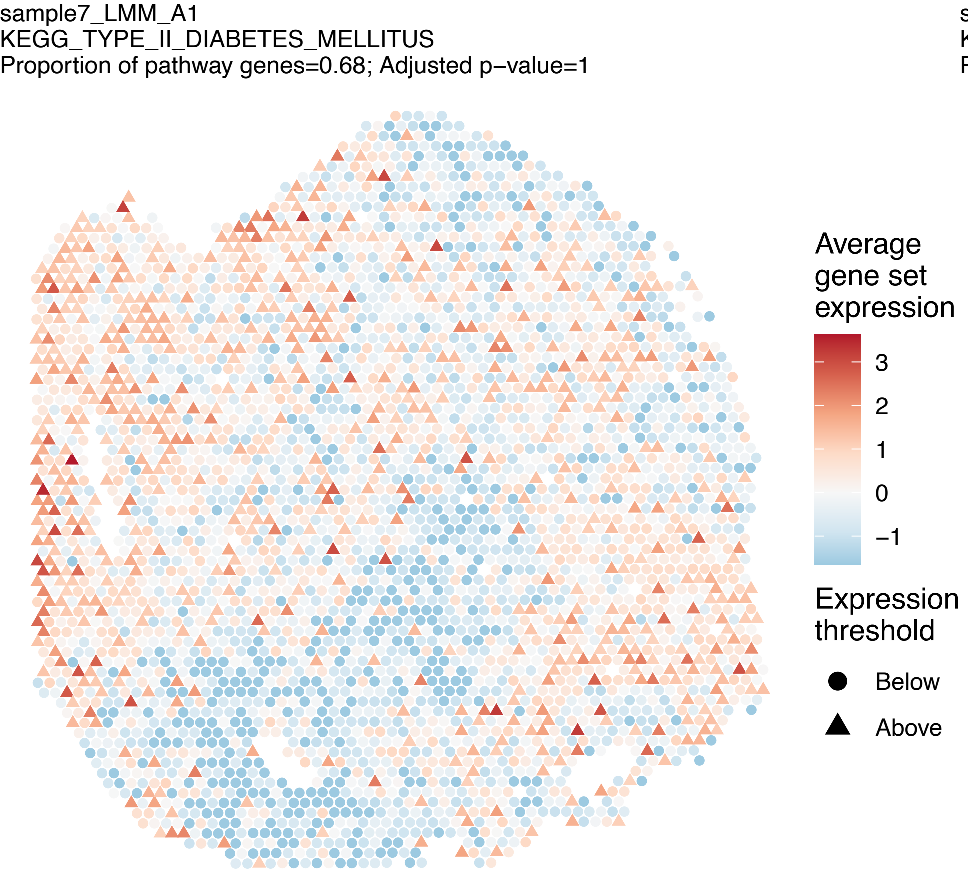
sample7_LMM_A1
KEGG_TYPE_II_DIABETES_MELLITUS
Proportion of pathway genes=0.68; Adjusted p−value=1
Average gene set expression
3
2
1
0
−1
Expression threshold
Below
Above

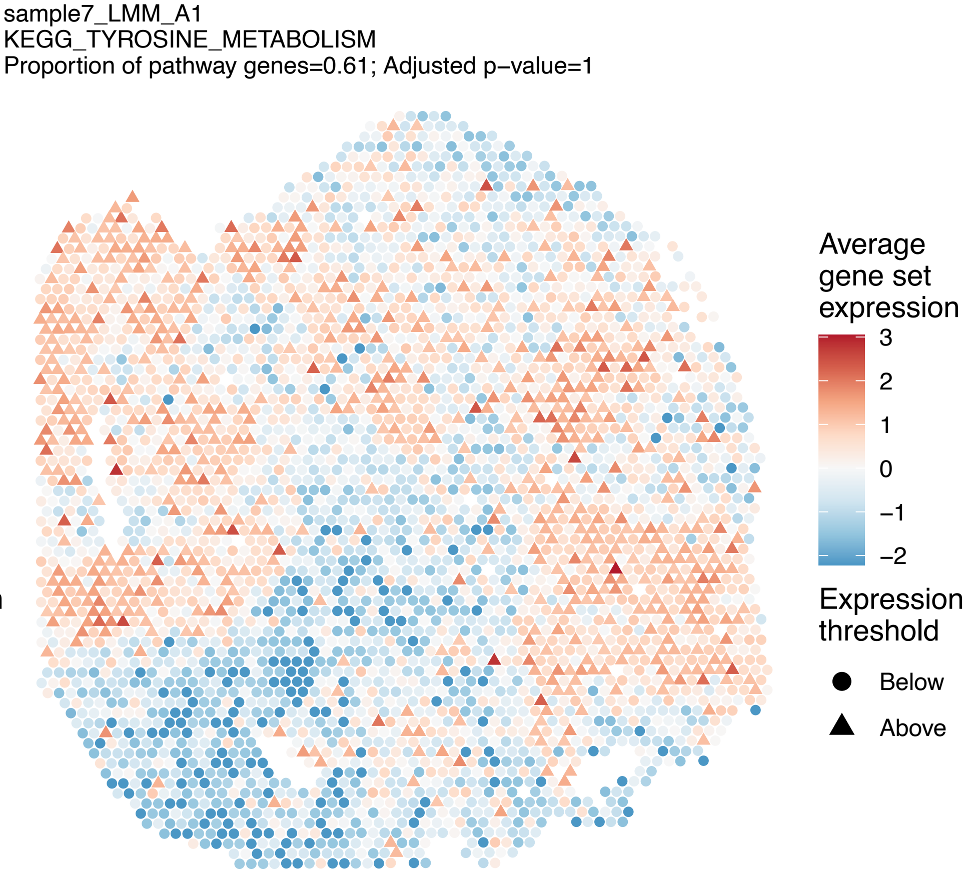
sample7_LMM_A1
KEGG_TYROSINE_METABOLISM
Proportion of pathway genes=0.61; Adjusted p−value=1
Average gene set expression
3
2
1
0
−1
−2
Expression threshold
Below
Above

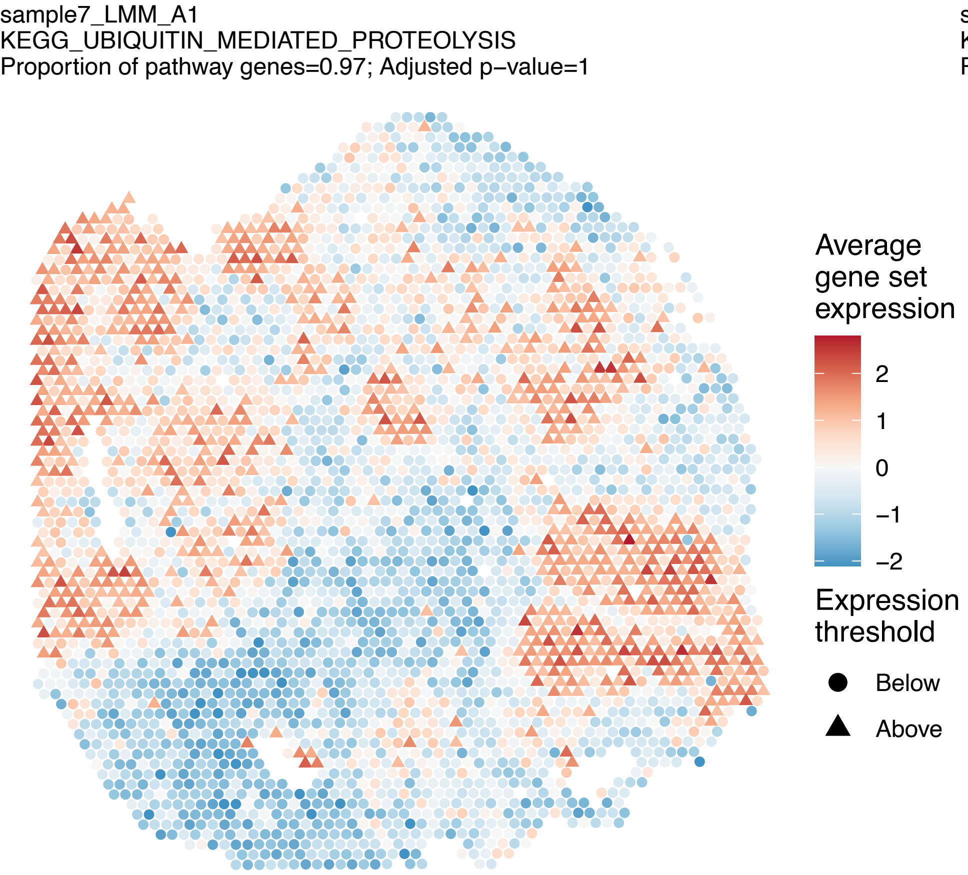
sample7_LMM_A1
KEGG_UBIQUITIN_MEDIATED_PROTEOLYSIS
Proportion of pathway genes=0.97; Adjusted p−value=1
Average gene set expression
2
1
0
−1
−2
Expression threshold
Below
Above

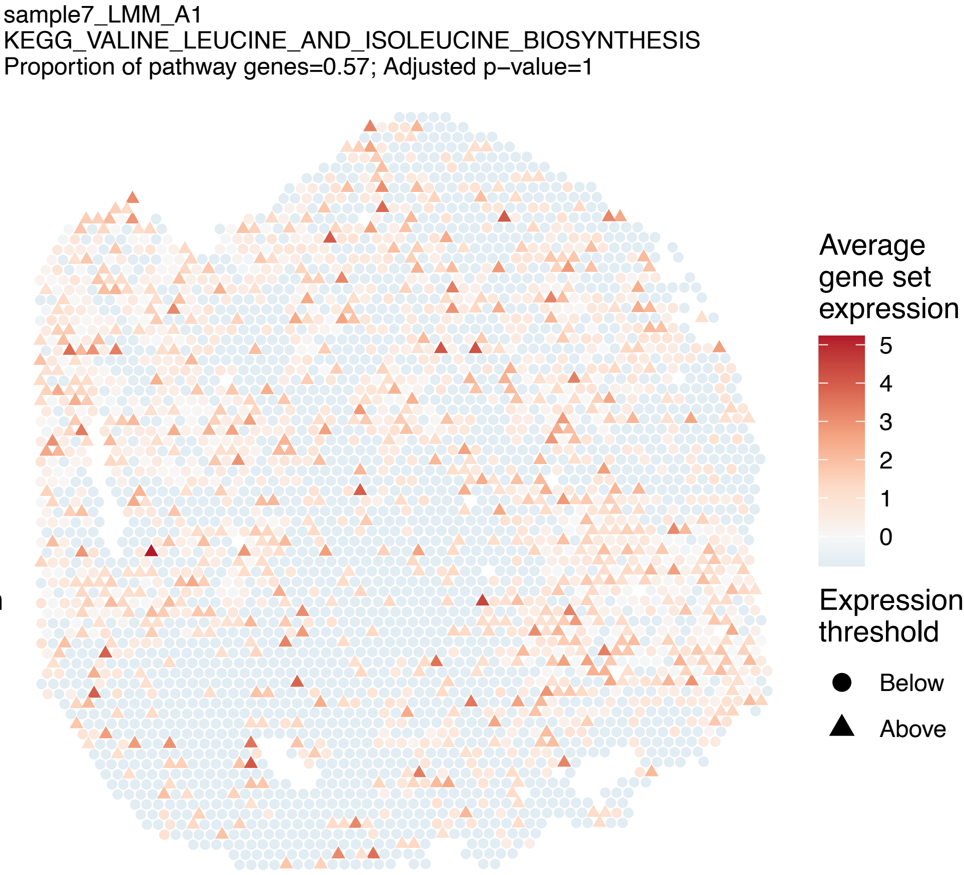
sample7_LMM_A1
KEGG_VALINE_LEUCINE_AND_ISOLEUCINE_BIOSYNTHESIS
Proportion of pathway genes=0.57; Adjusted p−value=1
Average gene set expression
5
4
3
2
1
0
Expression threshold
Below
Above

sample7_LMM_A1
KEGG_VALINE_LEUCINE_AND_ISOLEUCINE_DEGRADATION
Proportion of pathway genes=0.93; Adjusted p−value=1

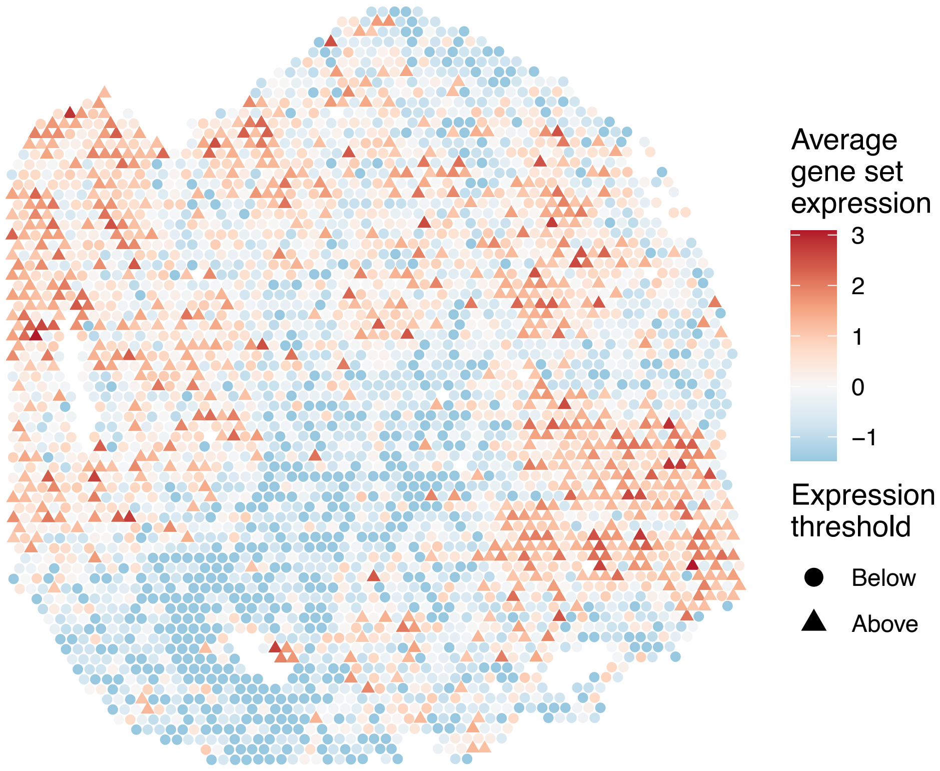


sample7_LMM_A1
KEGG_VASCULAR_SMOOTH_MUSCLE_CONTRACTION
Proportion of pathway genes=0.72; Adjusted p−value=1

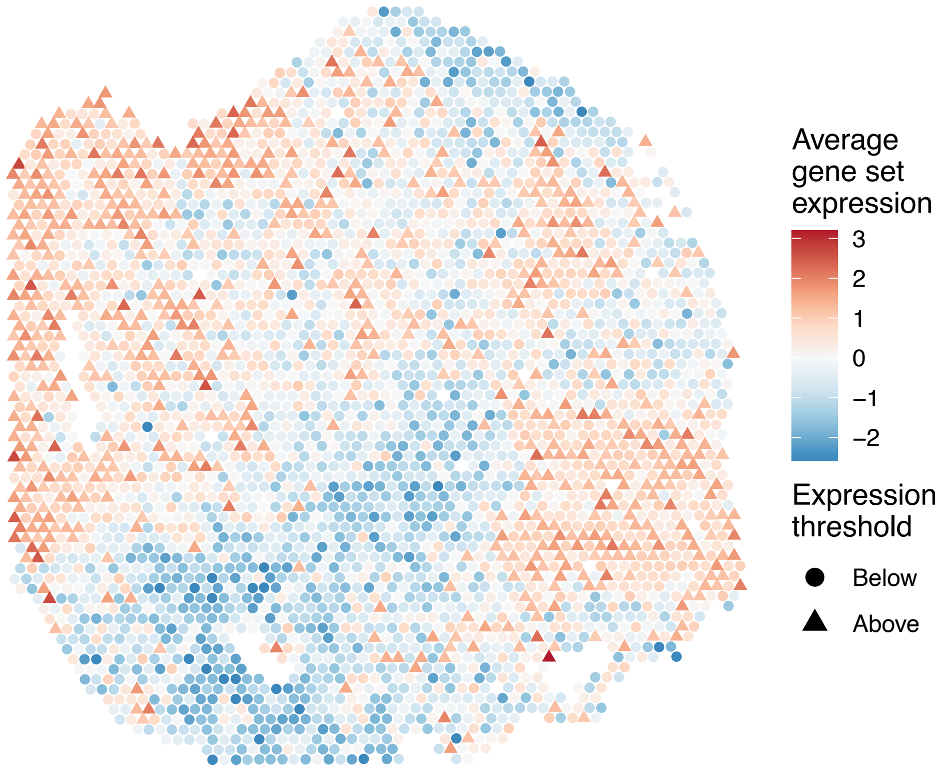


sample7_LMM_A1
KEGG_VASOPRESSIN_REGULATED_WATER_REABSORPTION
Proportion of pathway genes=0.87; Adjusted p−value=1

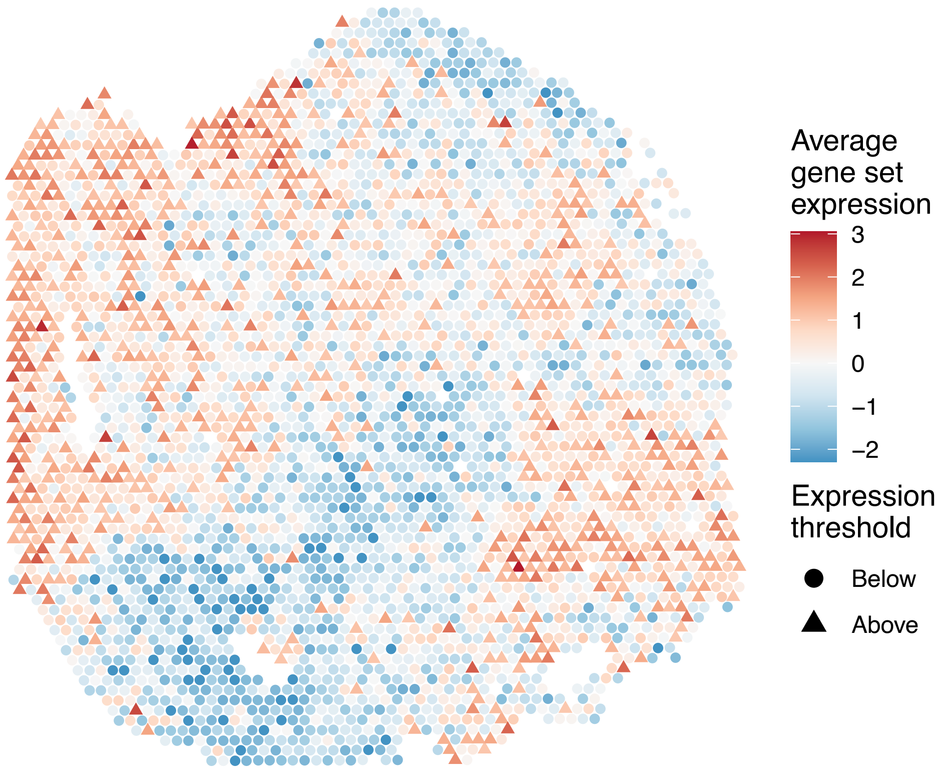


sample7_LMM_A1
KEGG_VEGF_SIGNALING_PATHWAY
Proportion of pathway genes=0.79; Adjusted p−value=1

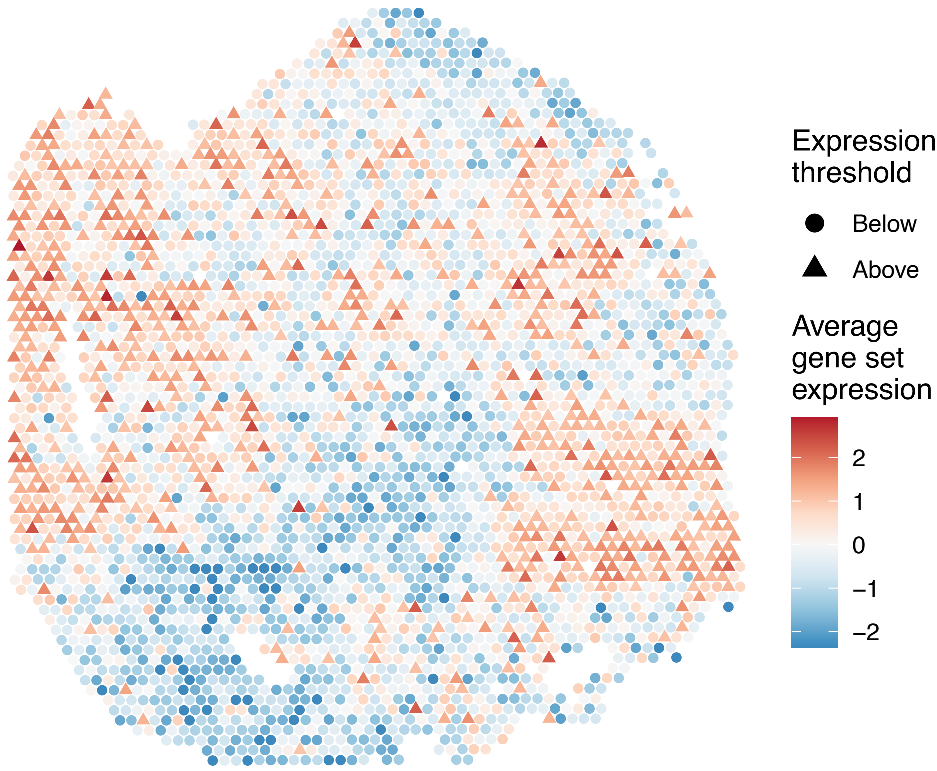

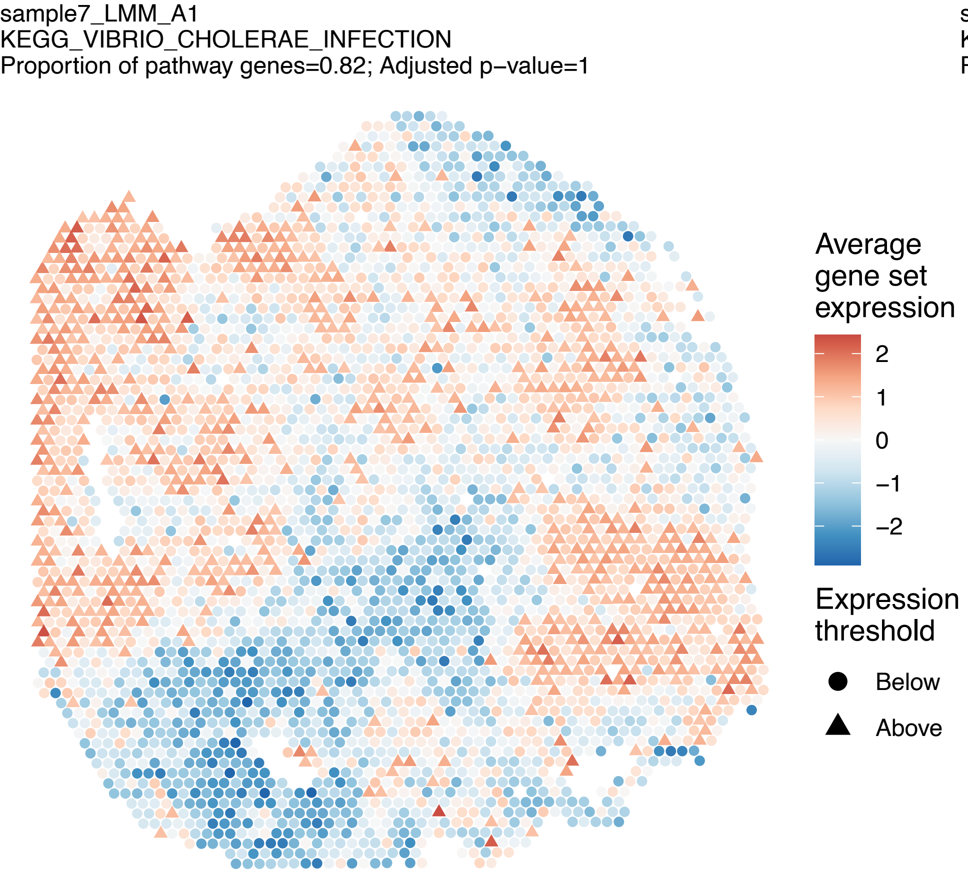
sample7_LMM_A1
KEGG_VIBRIO_CHOLERAE_INFECTION
Proportion of pathway genes=0.82; Adjusted p−value=1
Average gene set expression
2
1
0
−1
−2
Expression threshold
Below
Above

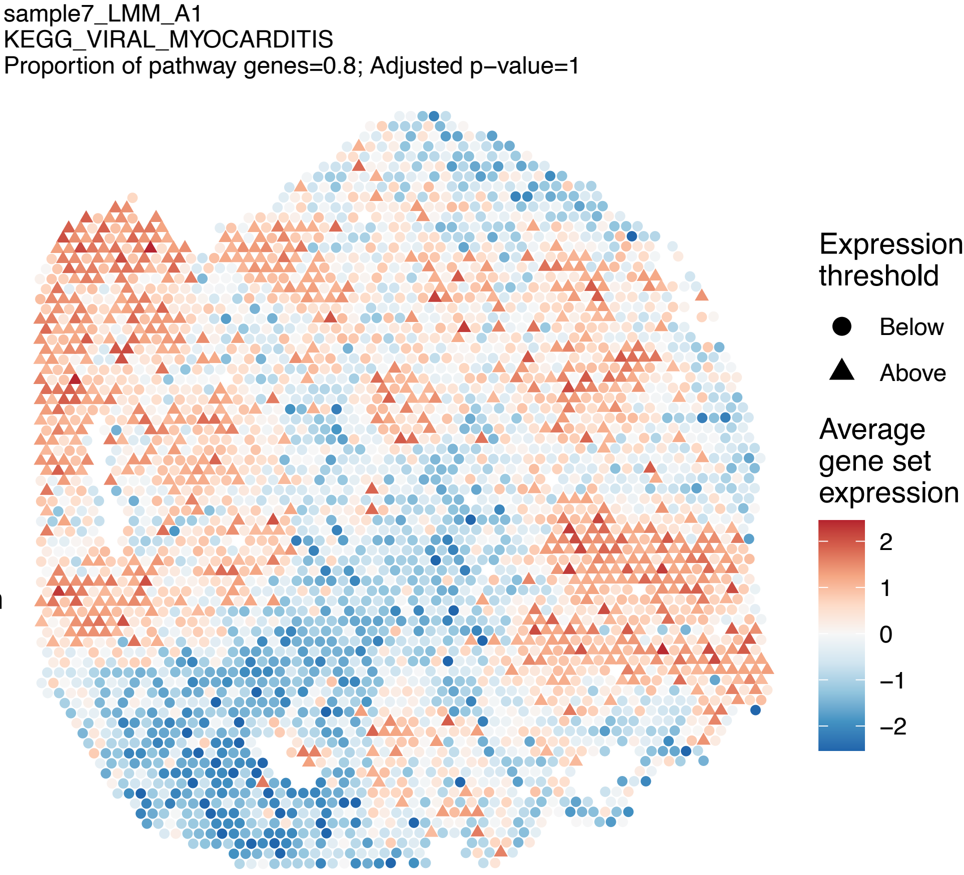
sample7_LMM_A1
KEGG_VIRAL_MYOCARDITIS
Proportion of pathway genes=0.8; Adjusted p−value=1
Expression threshold
Below
Above
Average gene set expression
2
1
0
−1
−2

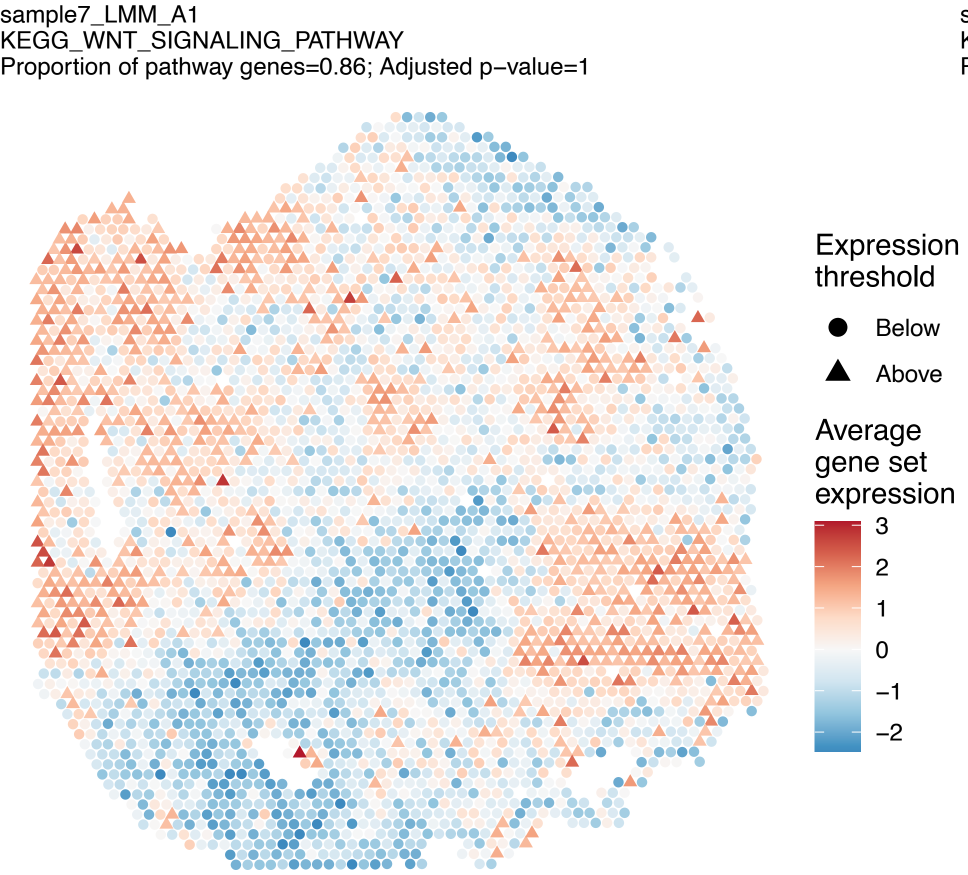
sample7_LMM_A1
KEGG_WNT_SIGNALING_PATHWAY
Proportion of pathway genes=0.86; Adjusted p−value=1
Expression threshold
Below
Above
Average gene set expression
3
2
1
0
−1
−2

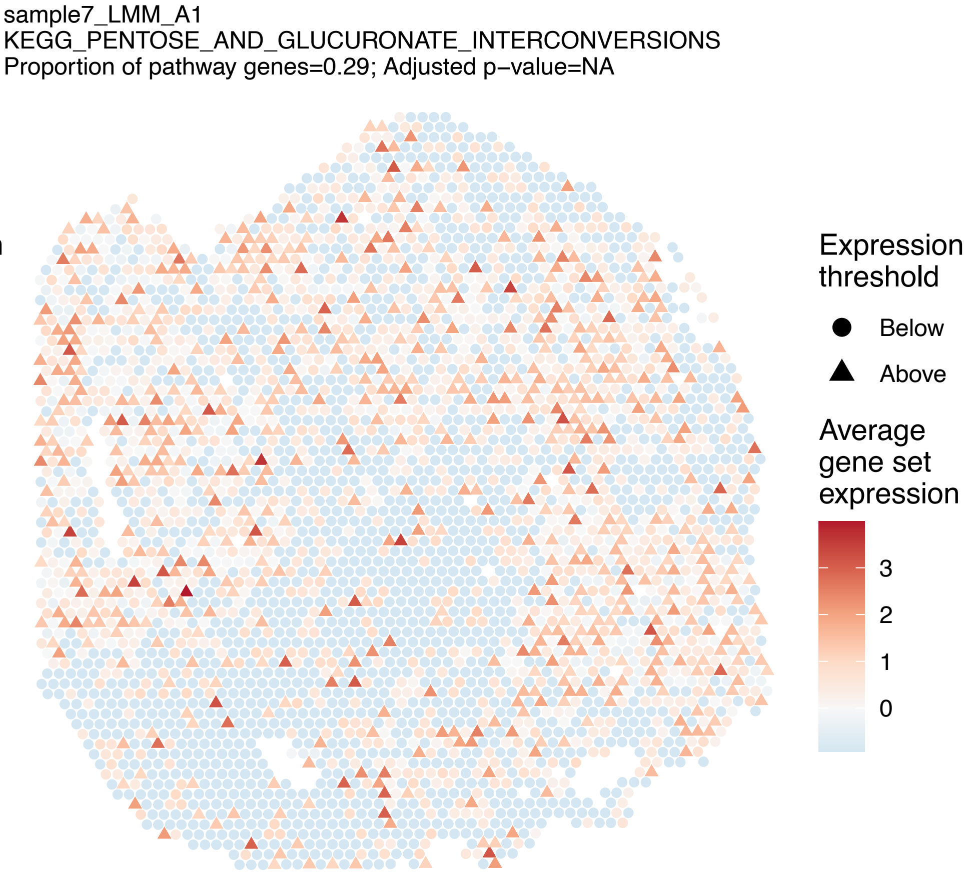
sample7_LMM_A1
KEGG_PENTOSE_AND_GLUCURONATE_INTERCONVERSIONS
Proportion of pathway genes=0.29; Adjusted p−value=NA
Expression threshold
Below
Above
Average gene set expression
3
2
1
0

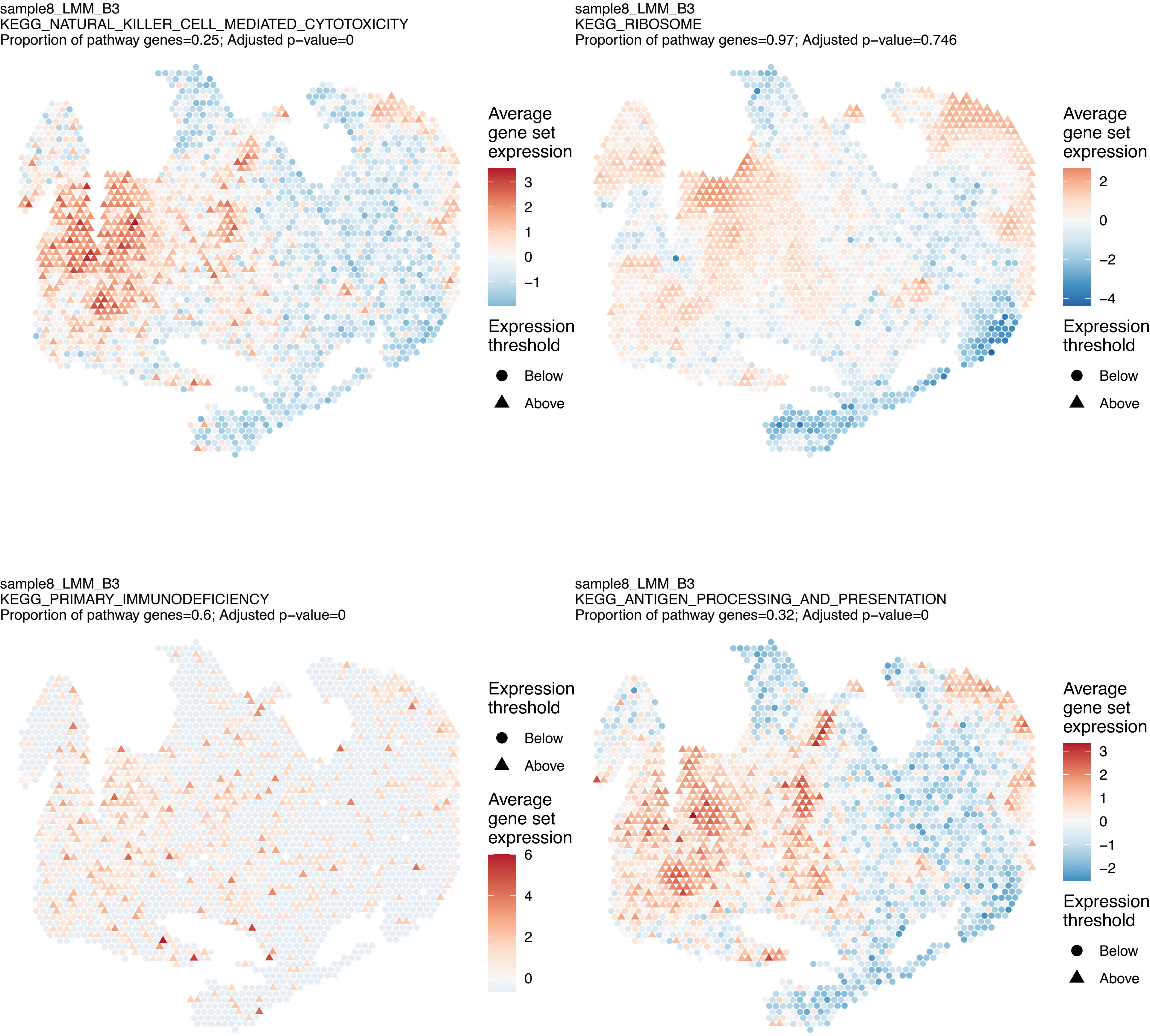
sample8_LMM_B3
KEGG_NATURAL_KILLER_CELL_MEDIATED_CYTOTOXICITY
Proportion of pathway genes=0.25; Adjusted p−value=0
Average gene set expression
3
2
1
0
−1
Expression threshold
Below
Above
sample8_LMM_B3
KEGG_RIBOSOME
Proportion of pathway genes=0.97; Adjusted p−value=0.746
Average gene set expression
2
0
−2
−4
Expression threshold
Below
Above
sample8_LMM_B3
KEGG_PRIMARY_IMMUNODEFICIENCY
Proportion of pathway genes=0.6; Adjusted p−value=0
Expression threshold
Below
Above
Average gene set expression
6
4
2
0
sample8_LMM_B3
KEGG_ANTIGEN_PROCESSING_AND_PRESENTATION
Proportion of pathway genes=0.32; Adjusted p−value=0
Average gene set expression
3
2
1
0
−1
−2
Expression threshold
Below
Above

sample8_LMM_B3
KEGG_CYTOKINE_CYTOKINE_RECEPTOR_INTERACTION
Proportion of pathway genes=0.32; Adjusted p−value=0

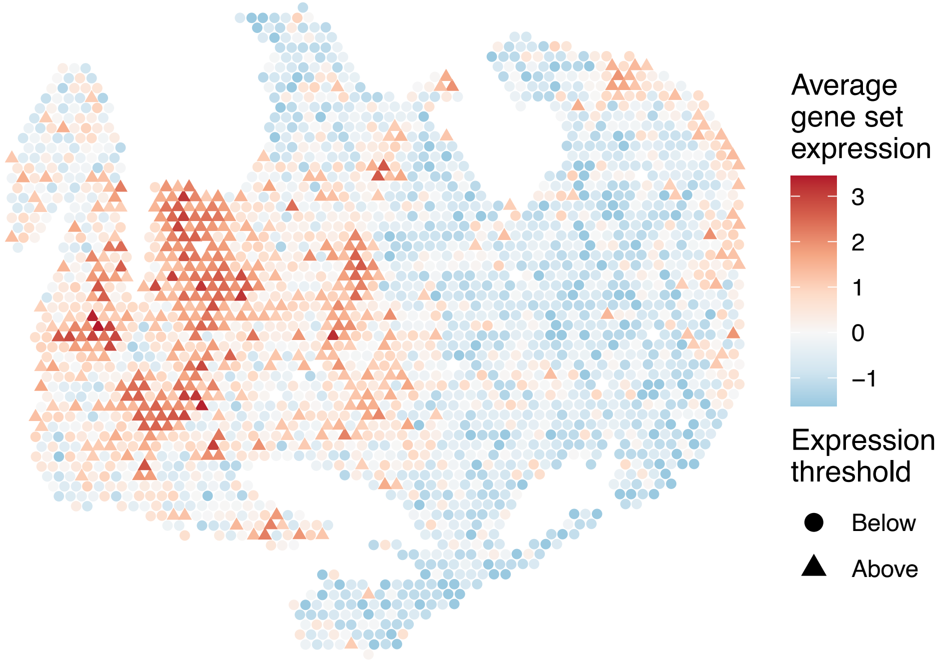


sample8_LMM_B3
KEGG_ABC_TRANSPORTERS
Proportion of pathway genes=0.7; Adjusted p−value=0

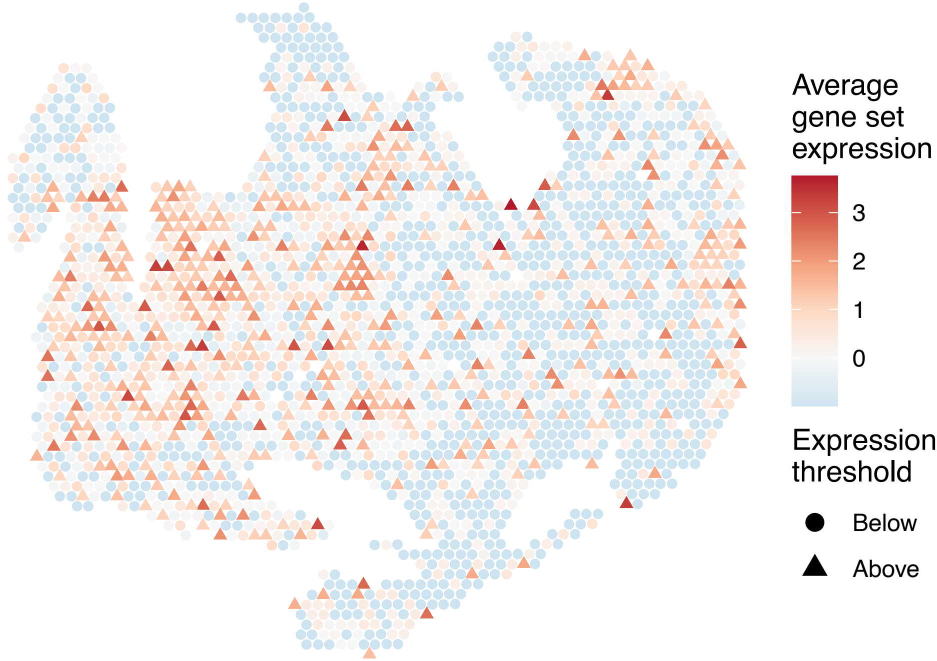


sample8_LMM_B3
KEGG_ACUTE_MYELOID_LEUKEMIA
Proportion of pathway genes=0.93; Adjusted p−value=0

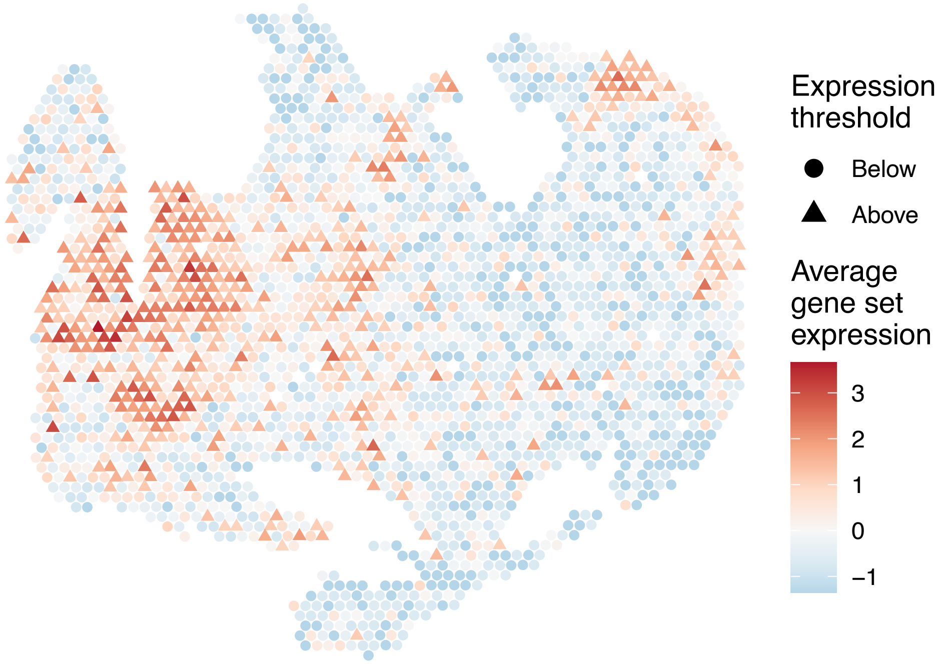


sample8_LMM_B3
KEGG_ADHERENS_JUNCTION
Proportion of pathway genes=0.94; Adjusted p−value=0

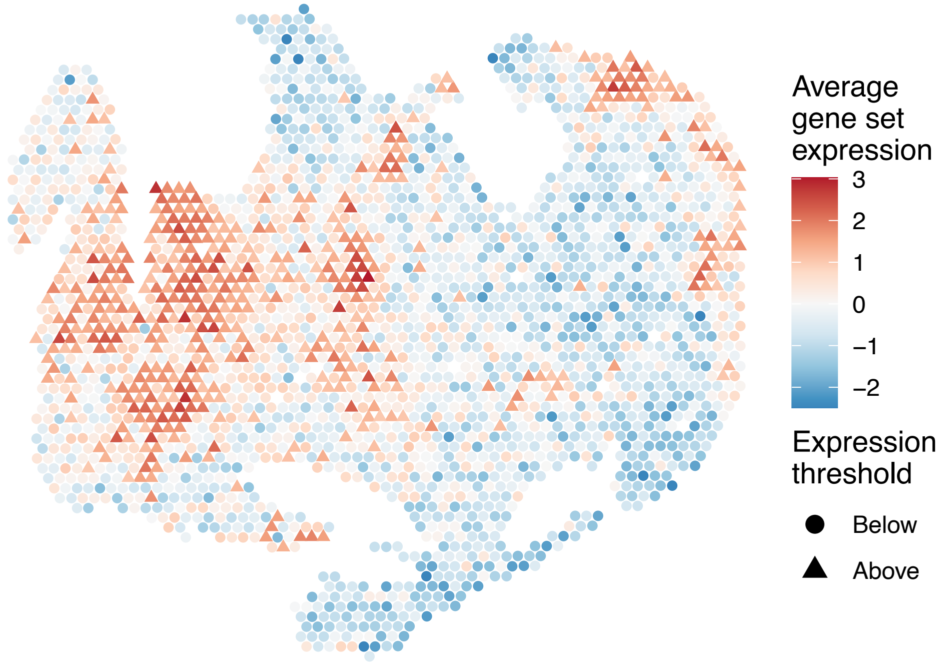

sample8_LMM_B3
KEGG_ADIPOCYTOKINE_SIGNALING_PATHWAY
Proportion of pathway genes=0.74; Adjusted p−value=0

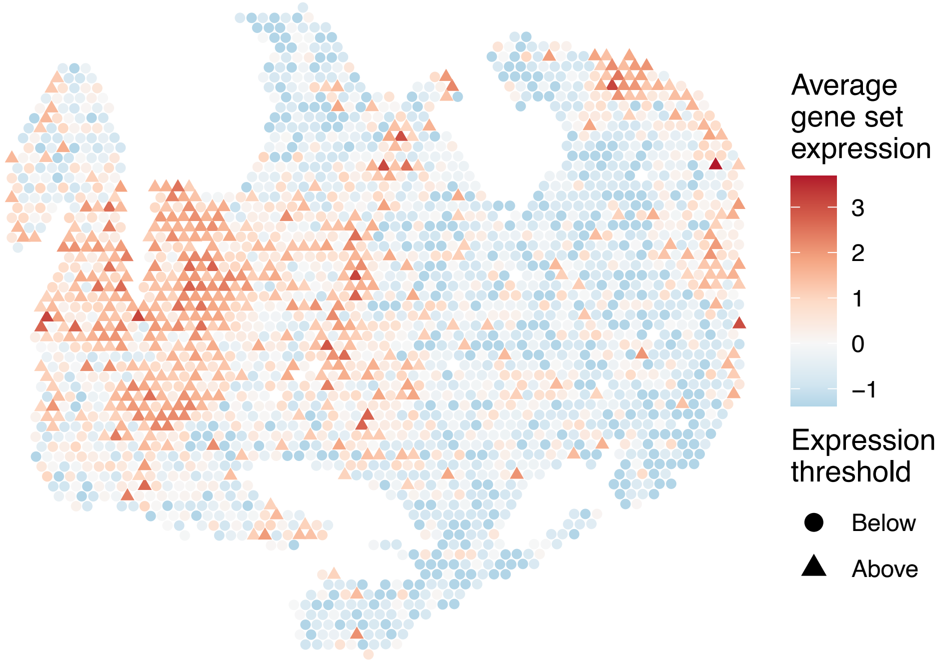


sample8_LMM_B3
KEGG_ALANINE_ASPARTATE_AND_GLUTAMATE_METABOLISM
Proportion of pathway genes=0.82; Adjusted p−value=0.0102

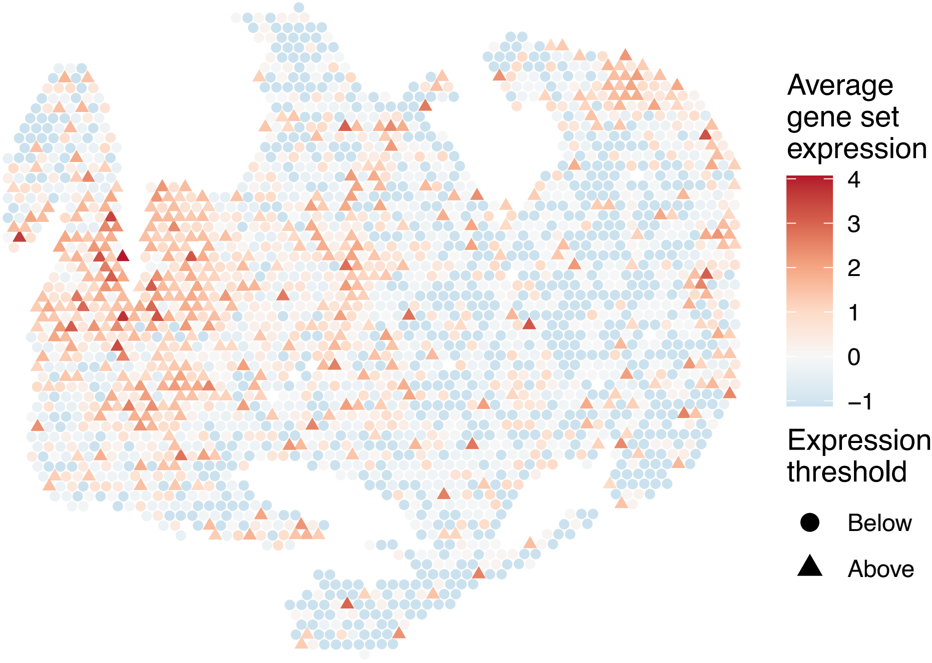


sample8_LMM_B3
KEGG_ALDOSTERONE_REGULATED_SODIUM_REABSORPTION
Proportion of pathway genes=0.64; Adjusted p−value=0

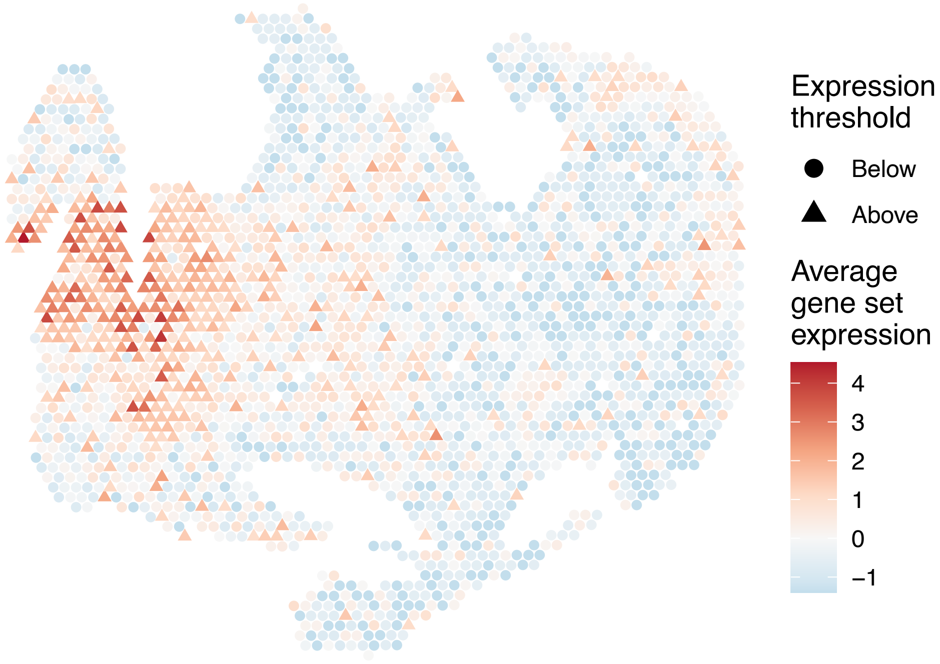


sample8_LMM_B3
KEGG_ALLOGRAFT_REJECTION
Proportion of pathway genes=0.62; Adjusted p−value=0

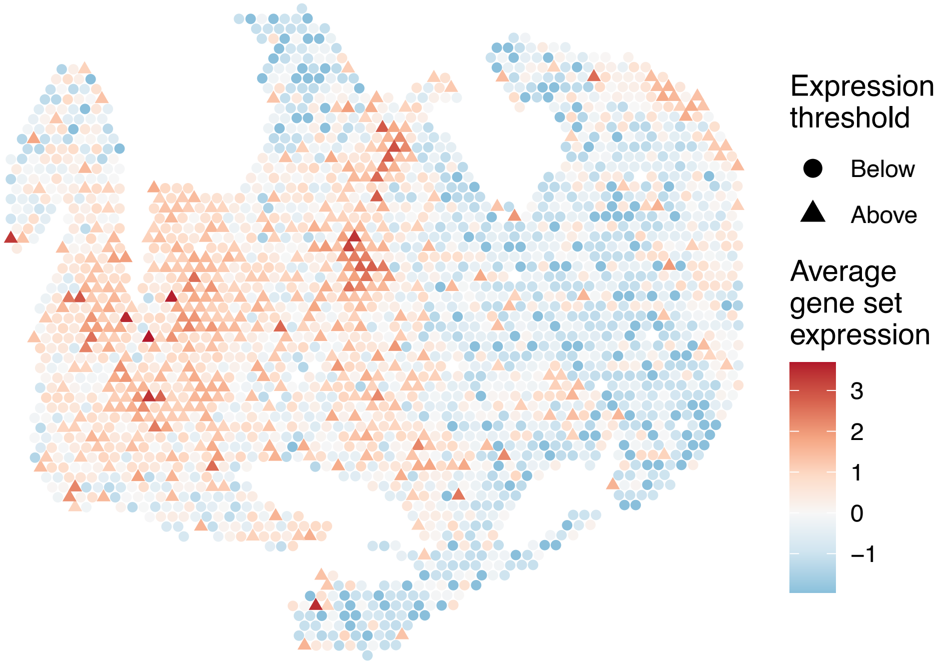

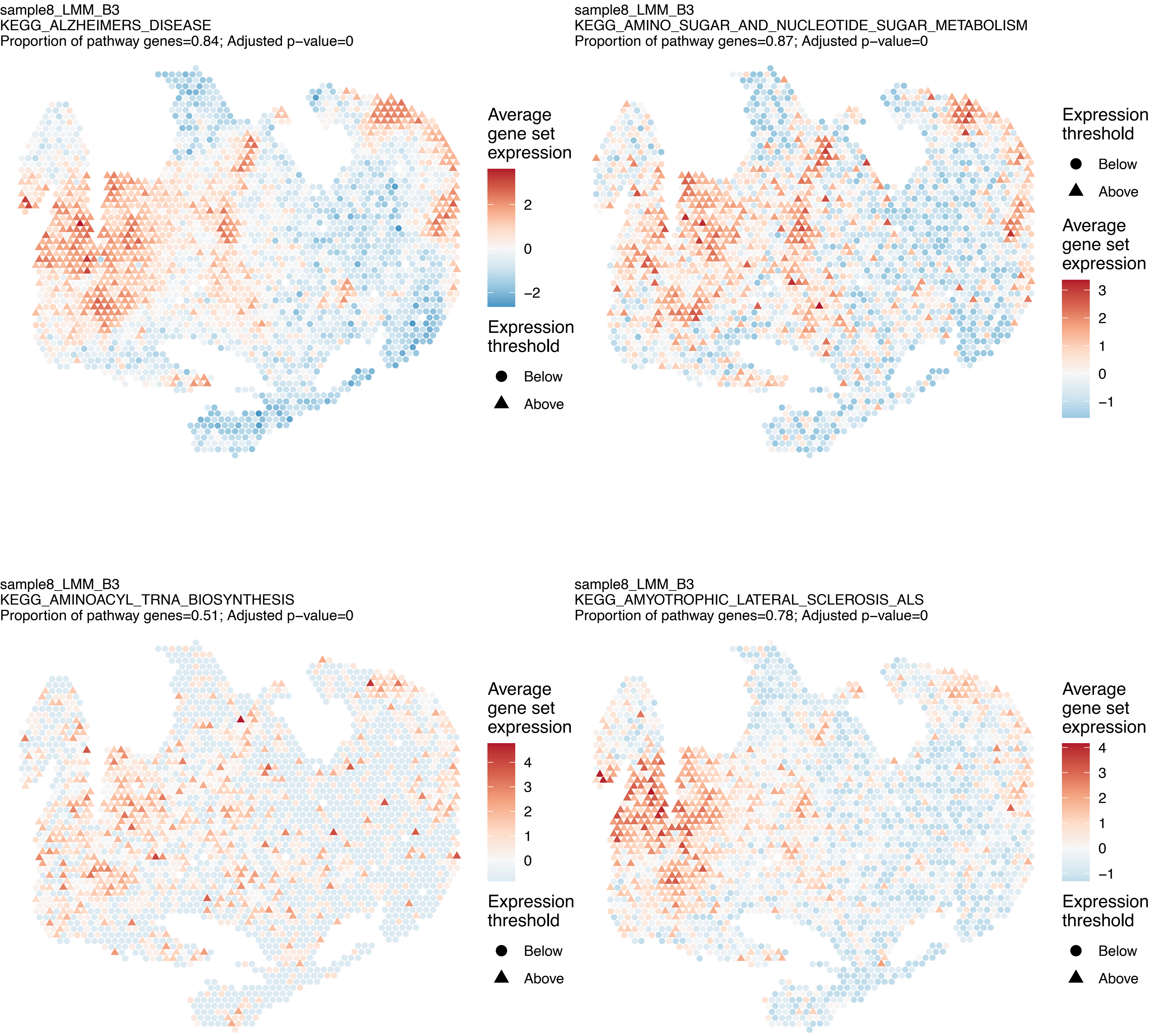
sample8_LMM_B3
KEGG_ALZHEIMERS_DISEASE
Proportion of pathway genes=0.84; Adjusted p−value=0
Average gene set expression
2
0
−2
Expression threshold
Below
Above
sample8_LMM_B3
KEGG_AMINO_SUGAR_AND_NUCLEOTIDE_SUGAR_METABOLISM
Proportion of pathway genes=0.87; Adjusted p−value=0
Expression threshold
Below
Above
Average gene set expression
3
2
1
0
−1
sample8_LMM_B3
KEGG_AMINOACYL_TRNA_BIOSYNTHESIS
Proportion of pathway genes=0.51; Adjusted p−value=0
Average gene set expression
4
3
2
1
0
Expression threshold
Below
Above
sample8_LMM_B3
KEGG_AMYOTROPHIC_LATERAL_SCLEROSIS_ALS
Proportion of pathway genes=0.78; Adjusted p−value=0
Average gene set expression
4
3
2
1
0
−1
Expression threshold
Below
Above

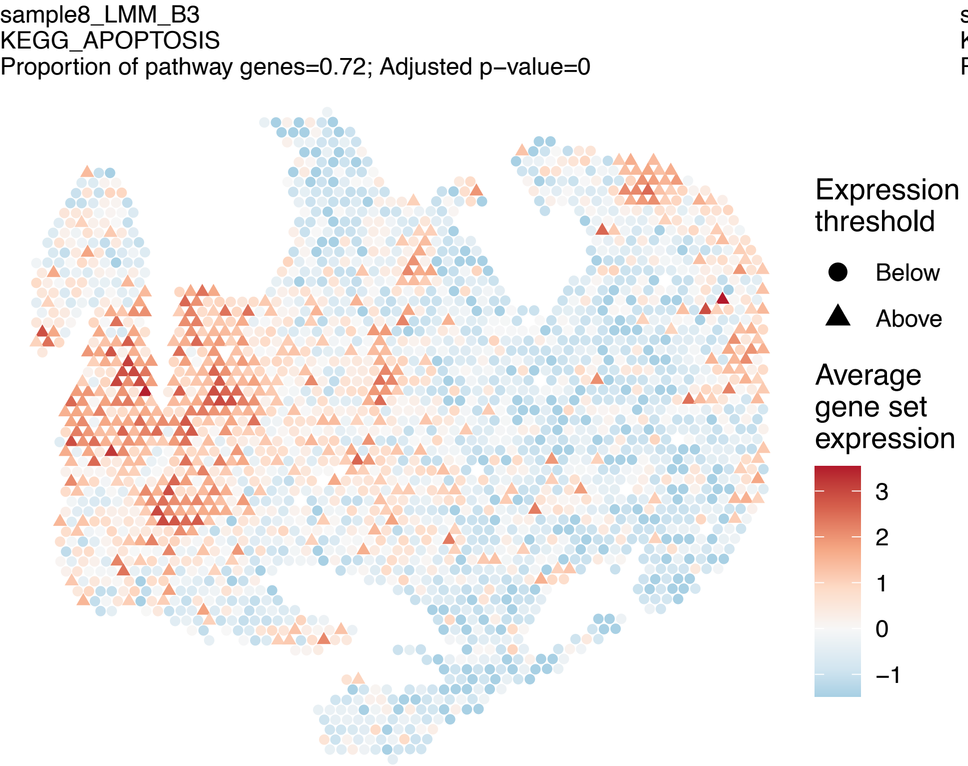

sample8_LMM_B3
KEGG_APOPTOSIS
Proportion of pathway genes=0.72; Adjusted p–value=0
Expression threshold
Below
Above
Average gene set expression
3
2
1
0
−1


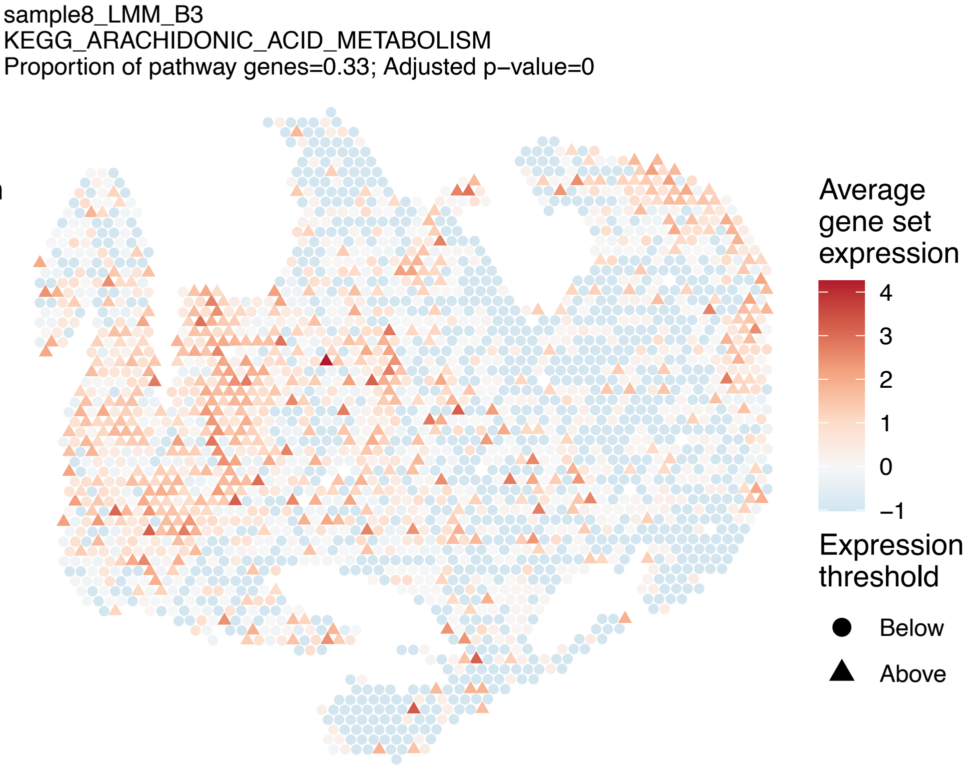

sample8_LMM_B3
KEGG_ARACHIDONIC_ACID_METABOLISM
Proportion of pathway genes=0.33; Adjusted p–value=0
Average gene set expression
4
3
2
1
0
−1
Expression threshold
Below
Above


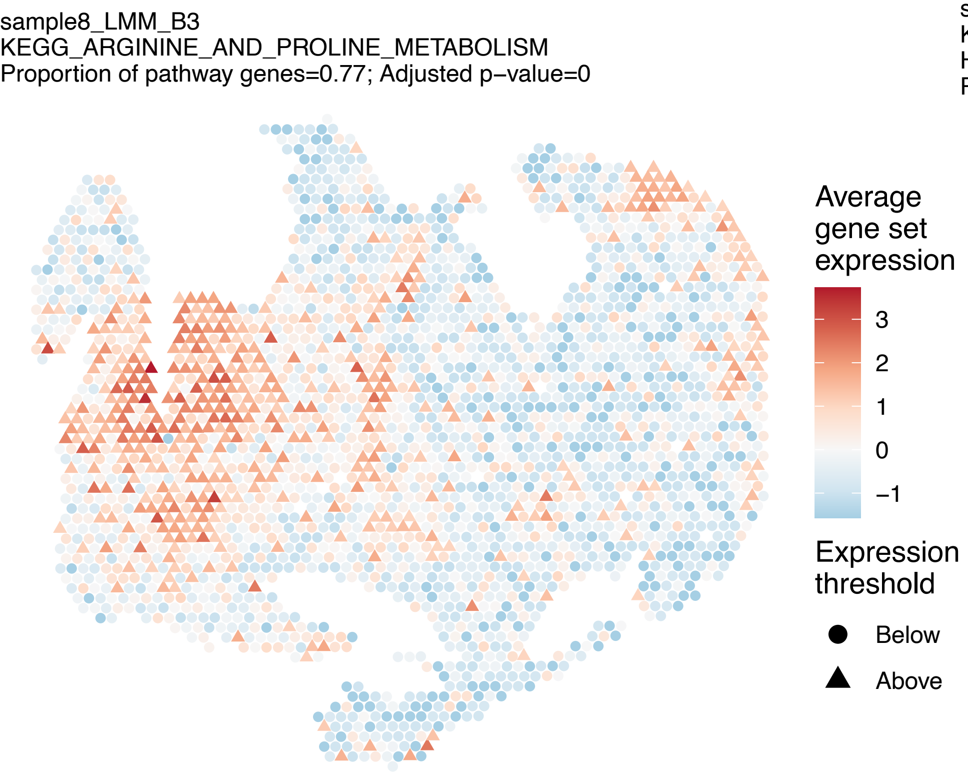

sample8_LMM_B3
KEGG_ARGININE_AND_PROLINE_METABOLISM
Proportion of pathway genes=0.77; Adjusted p–value=0
Average gene set expression
3
2
1
0
−1
Expression threshold
Below
Above


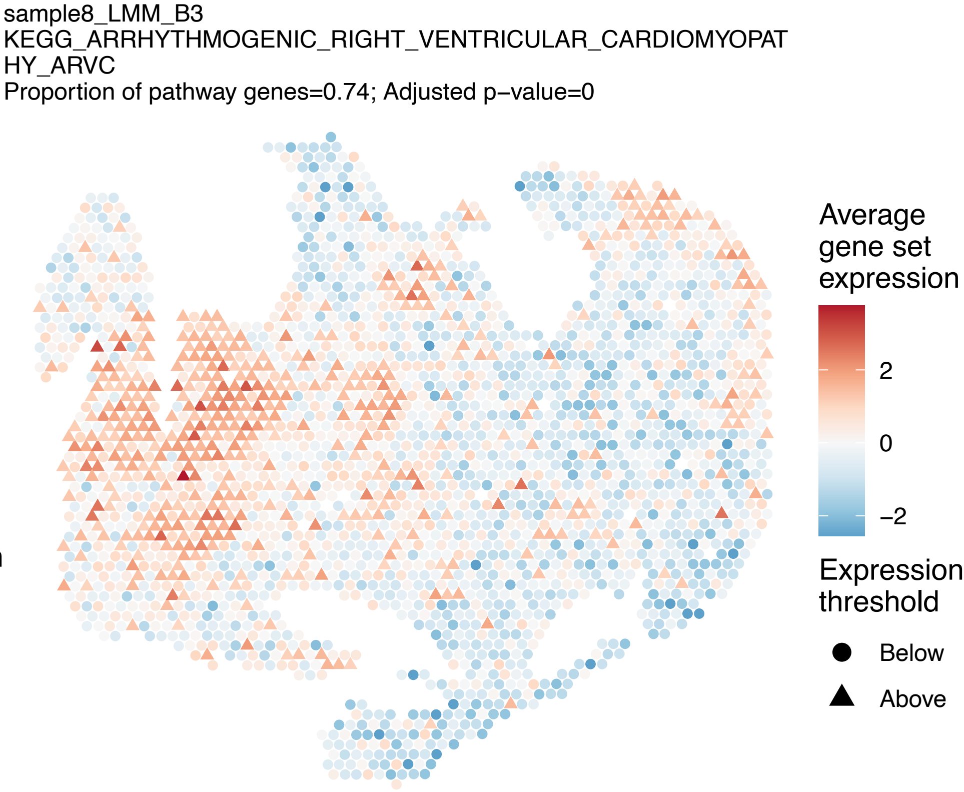

sample8_LMM_B3
KEGG_ARRHYTHMOGENIC_RIGHT_VENTRICULAR_CARDIOMYOPAT
HY_ARVC
Proportion of pathway genes=0.74; Adjusted p–value=0
Average gene set expression
2
0
−2
Expression threshold
Below
Above

sample8_LMM_B3
KEGG_ASTHMA
Proportion of pathway genes=0.56; Adjusted p−value=0

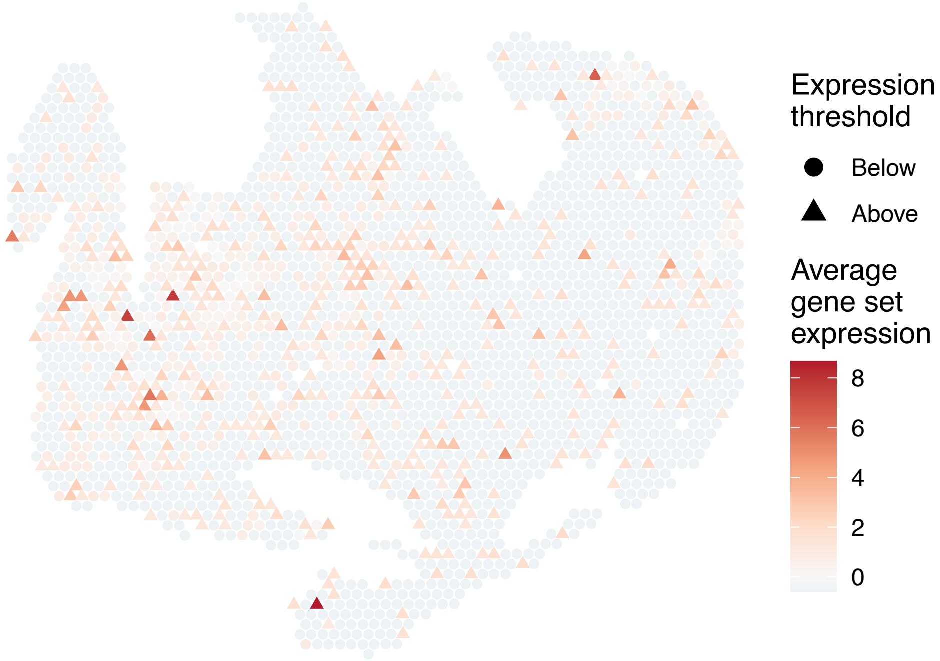


sample8_LMM_B3
KEGG_AUTOIMMUNE_THYROID_DISEASE
Proportion of pathway genes=0.59; Adjusted p−value=0

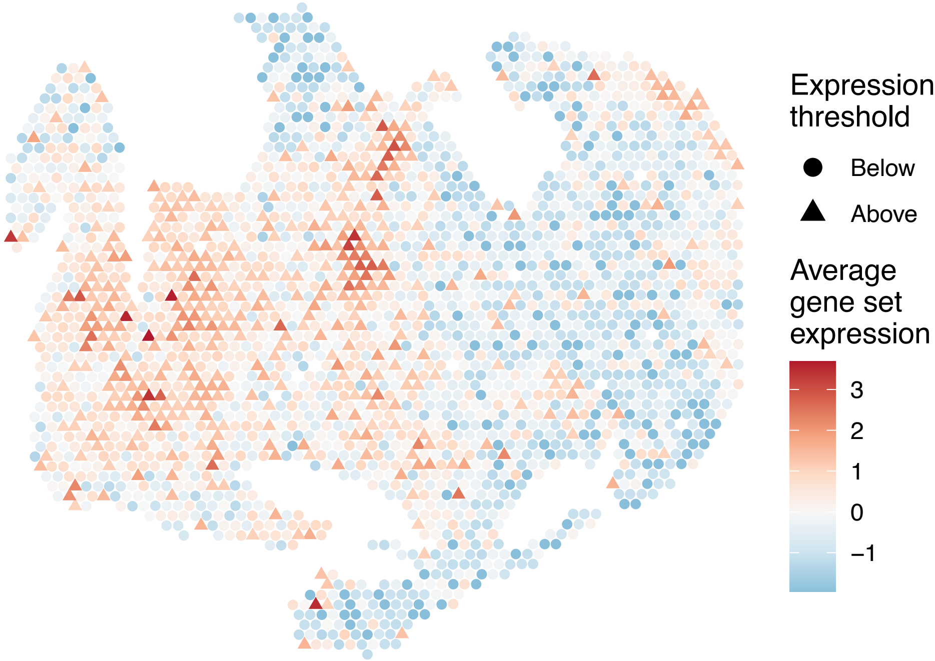


sample8_LMM_B3
KEGG_AXON_GUIDANCE
Proportion of pathway genes=0.88; Adjusted p−value=0

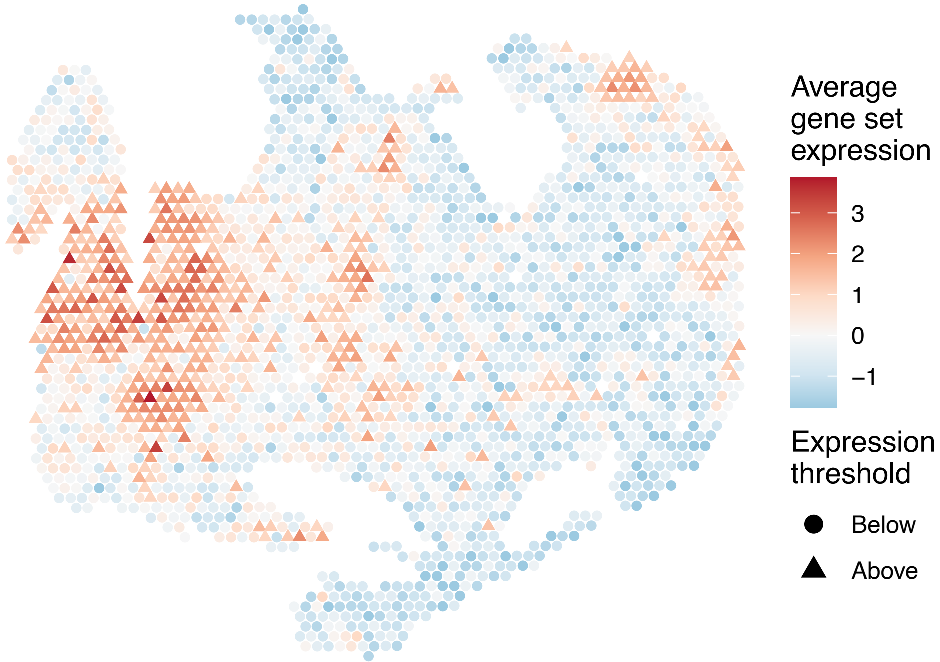


sample8_LMM_B3
KEGG_B_CELL_RECEPTOR_SIGNALING_PATHWAY
Proportion of pathway genes=0.84; Adjusted p−value=0

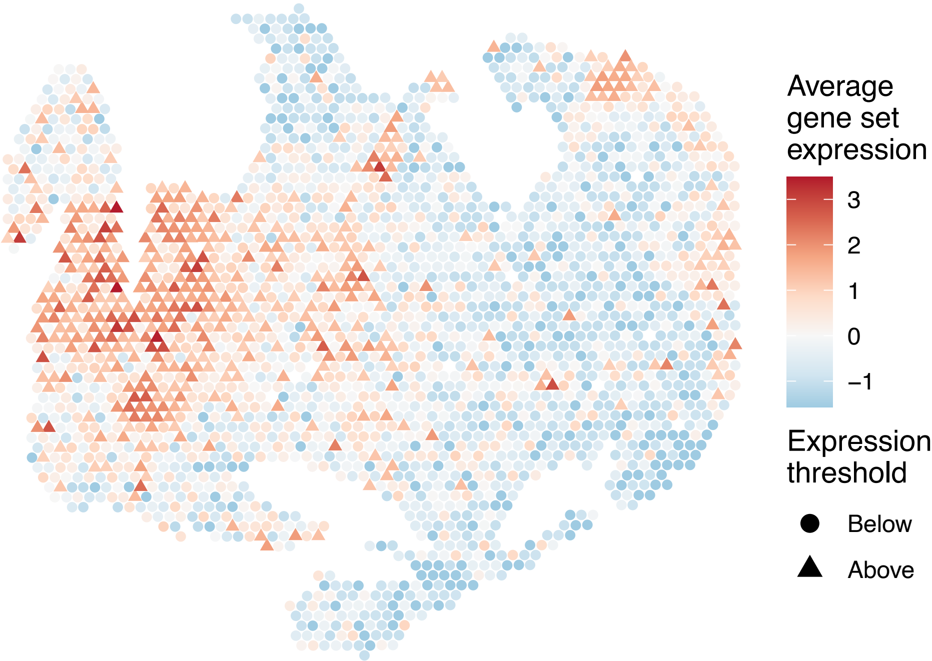

sample8_LMM_B3
KEGG_BASAL_CELL_CARCINOMA
Proportion of pathway genes=0.64; Adjusted p−value=0

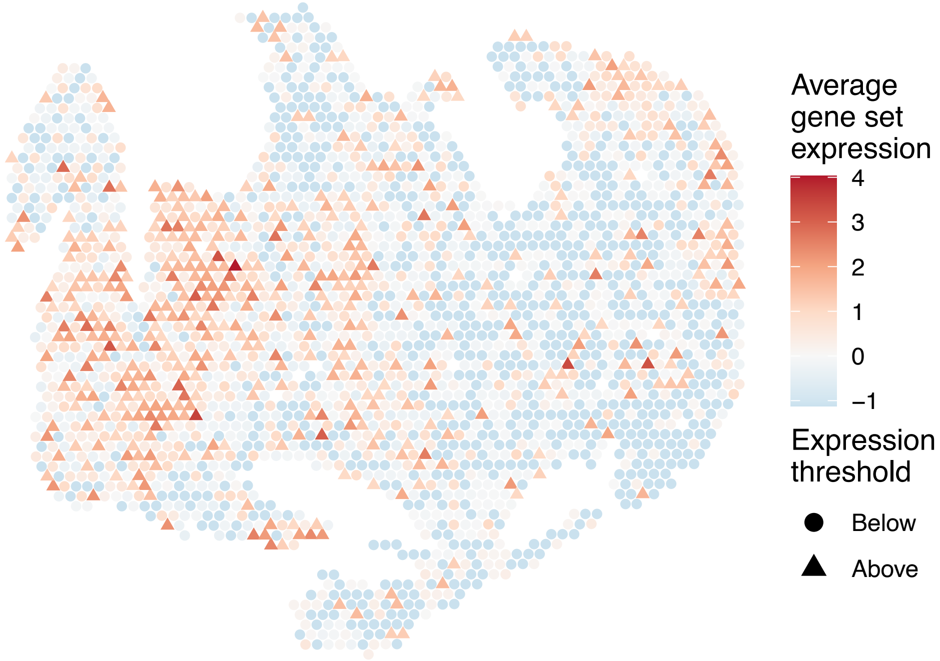


sample8_LMM_B3
KEGG_BASAL_TRANSCRIPTION_FACTORS
Proportion of pathway genes=0.91; Adjusted p−value=0

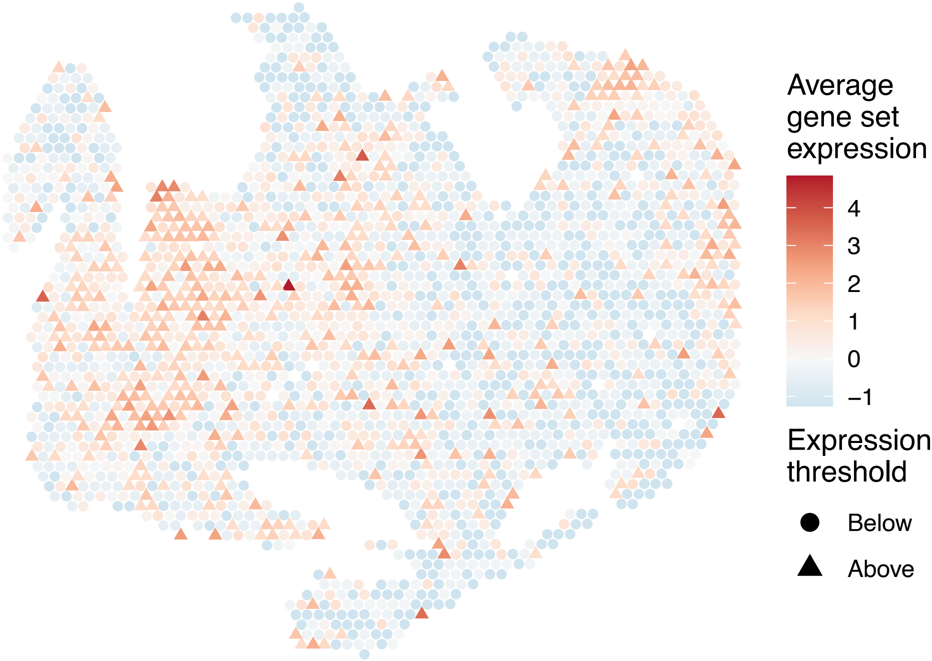


sample8_LMM_B3
KEGG_BASE_EXCISION_REPAIR
Proportion of pathway genes=0.94; Adjusted p−value=0

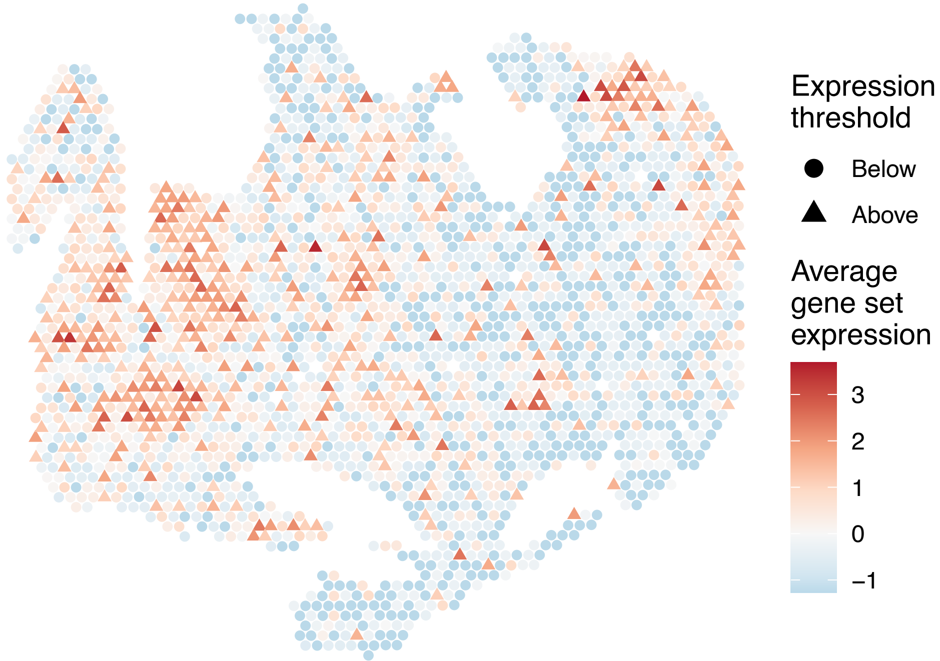


sample8_LMM_B3
KEGG_BETA_ALANINE_METABOLISM
Proportion of pathway genes=0.77; Adjusted p−value=0.6685

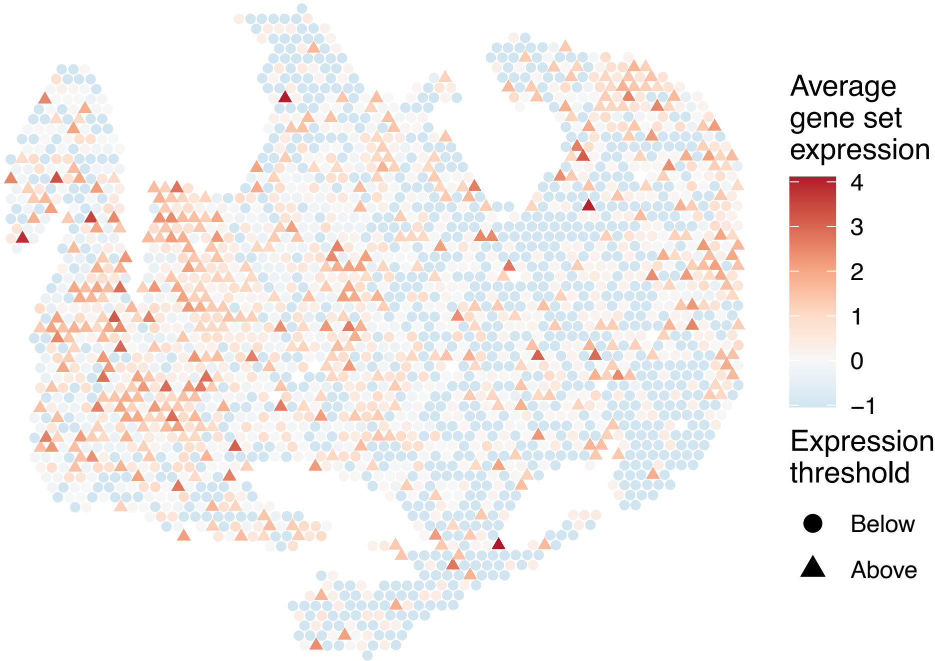

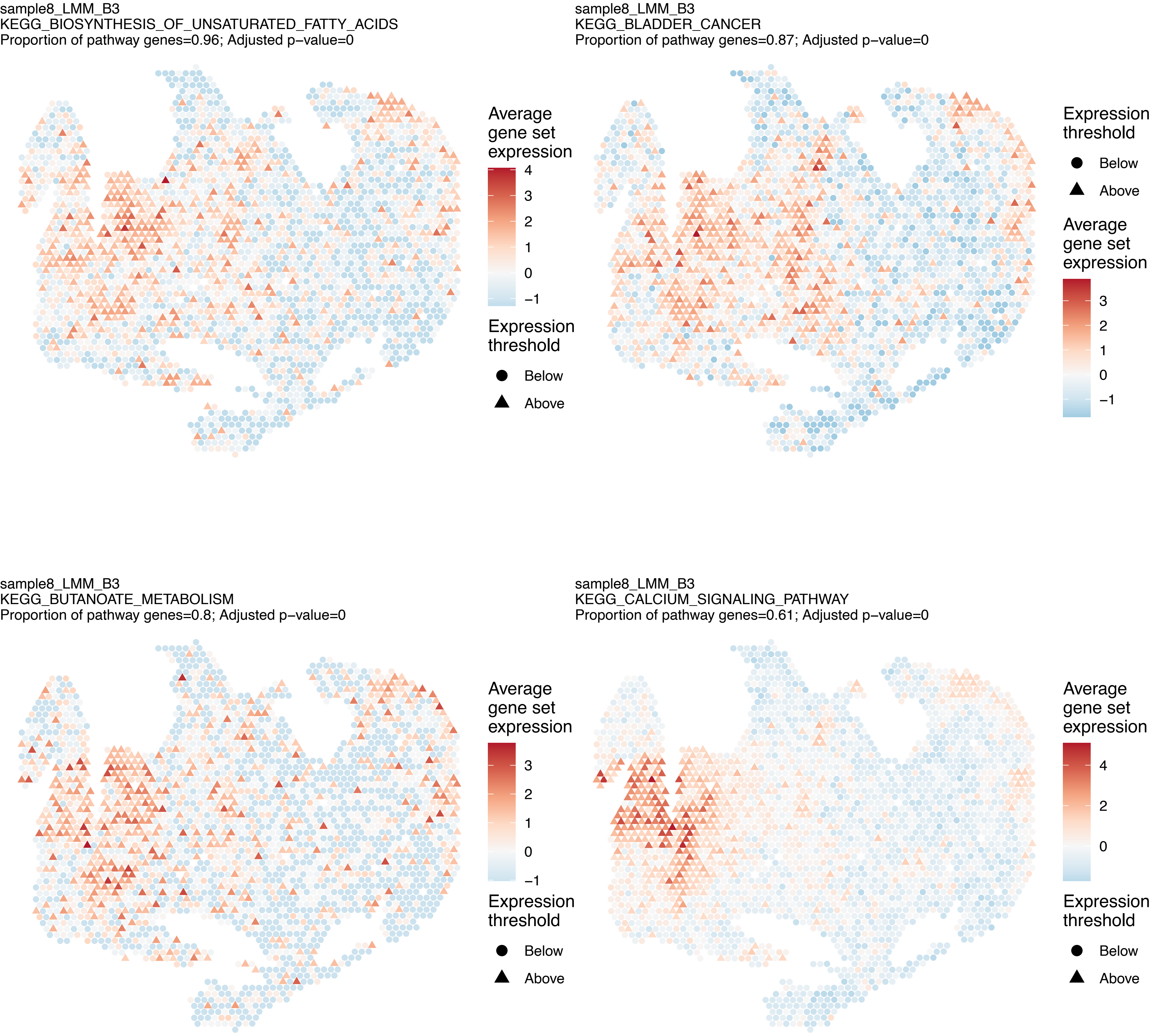
sample8_LMM_B3
KEGG_BIOSYNTHESIS_OF_UNSATURATED_FATTY_ACIDS
Proportion of pathway genes=0.96; Adjusted p−value=0
Average gene set expression
4
3
2
1
0
−1
Expression threshold
Below
Above
sample8_LMM_B3
KEGG_BLADDER_CANCER
Proportion of pathway genes=0.87; Adjusted p−value=0
Expression threshold
Below
Above
Average gene set expression
3
2
1
0
−1
sample8_LMM_B3
KEGG_BUTANOATE_METABOLISM
Proportion of pathway genes=0.8; Adjusted p−value=0
Average gene set expression
3
2
1
0
−1
Expression threshold
Below
Above
sample8_LMM_B3
KEGG_CALCIUM_SIGNALING_PATHWAY
Proportion of pathway genes=0.61; Adjusted p−value=0
Average gene set expression
4
2
0
Expression threshold
Below
Above

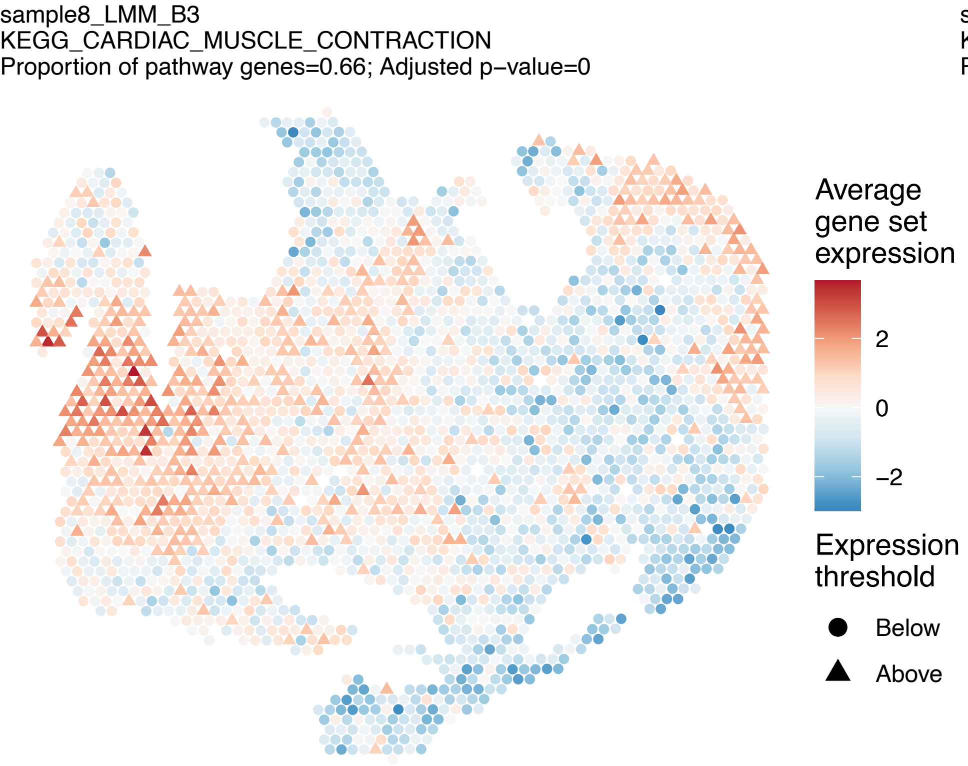
sample8_LMM_B3
KEGG_CARDIAC_MUSCLE_CONTRACTION
Proportion of pathway genes=0.66; Adjusted p−value=0
Average gene set expression
2
0
−2
Expression threshold
Below
Above

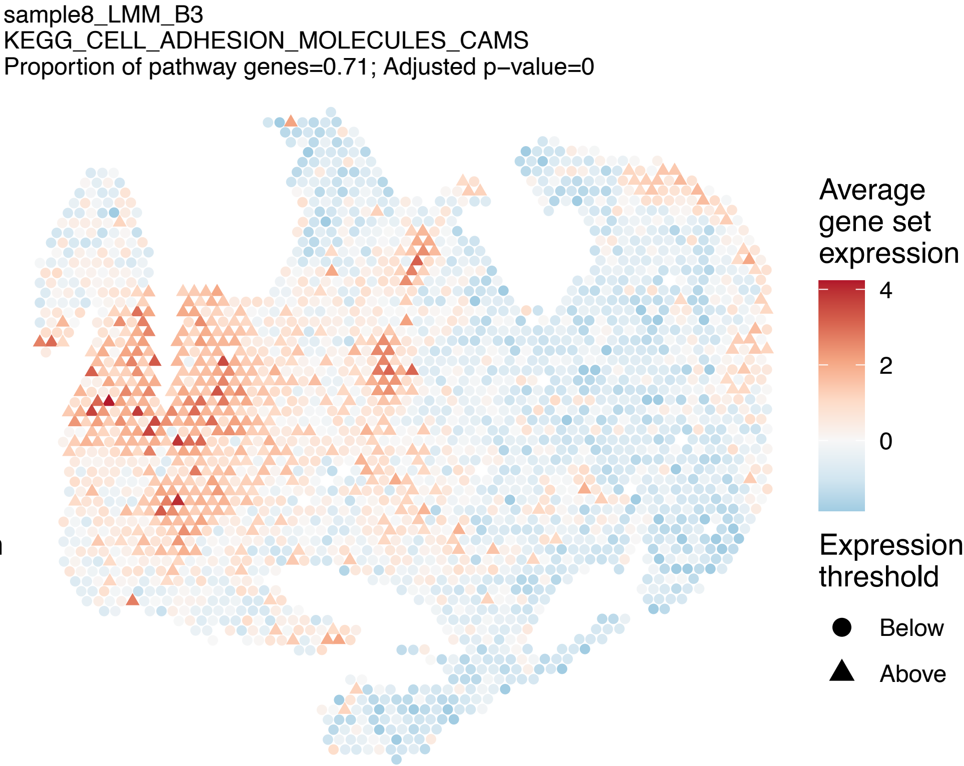
sample8_LMM_B3
KEGG_CELL_ADHESION_MOLECULES_CAMS
Proportion of pathway genes=0.71; Adjusted p−value=0
Average gene set expression
4
2
0
Expression threshold
Below
Above

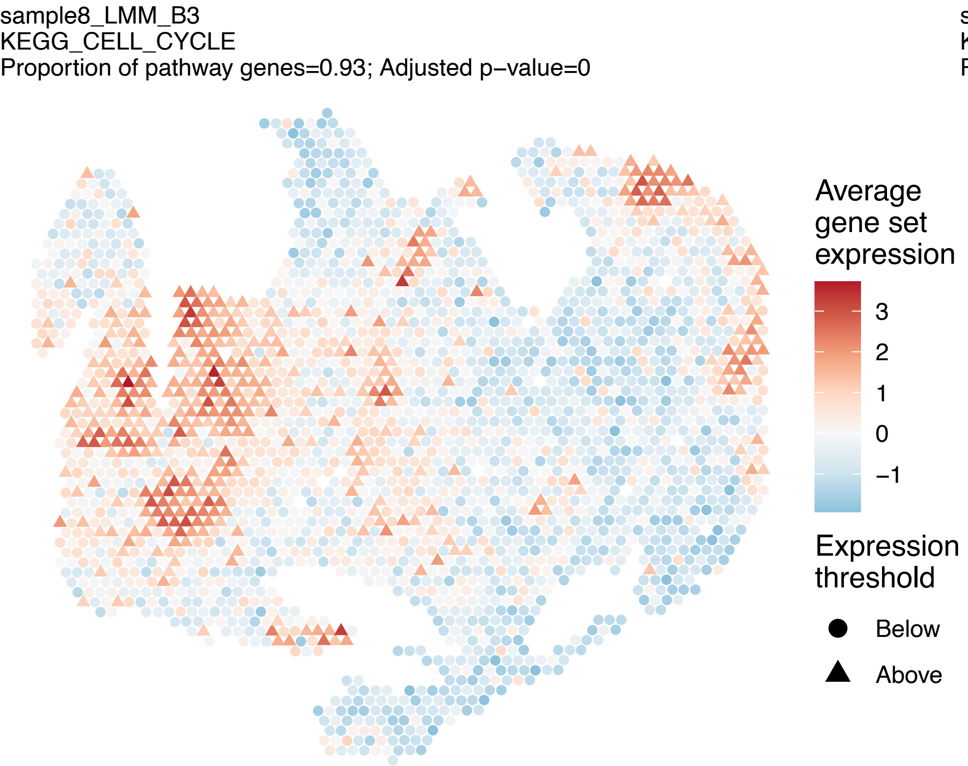
sample8_LMM_B3
KEGG_CELL_CYCLE
Proportion of pathway genes=0.93; Adjusted p−value=0
Average gene set expression
3
2
1
0
−1
Expression threshold
Below
Above

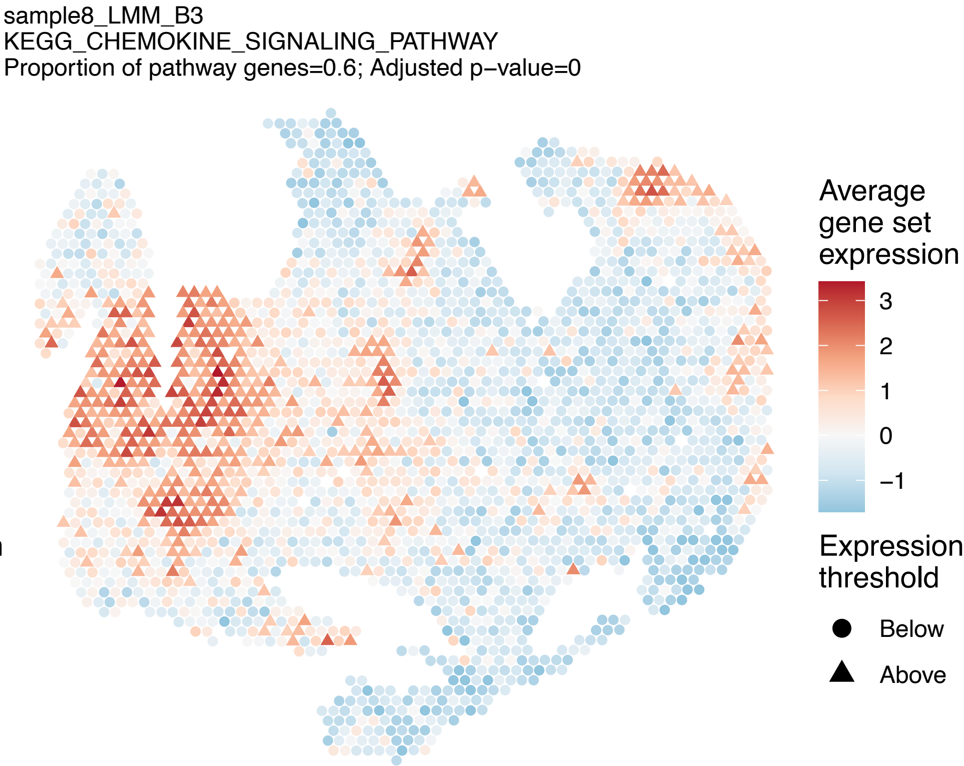
sample8_LMM_B3
KEGG_CHEMOKINE_SIGNALING_PATHWAY
Proportion of pathway genes=0.6; Adjusted p−value=0
Average gene set expression
3
2
1
0
−1
Expression threshold
Below
Above

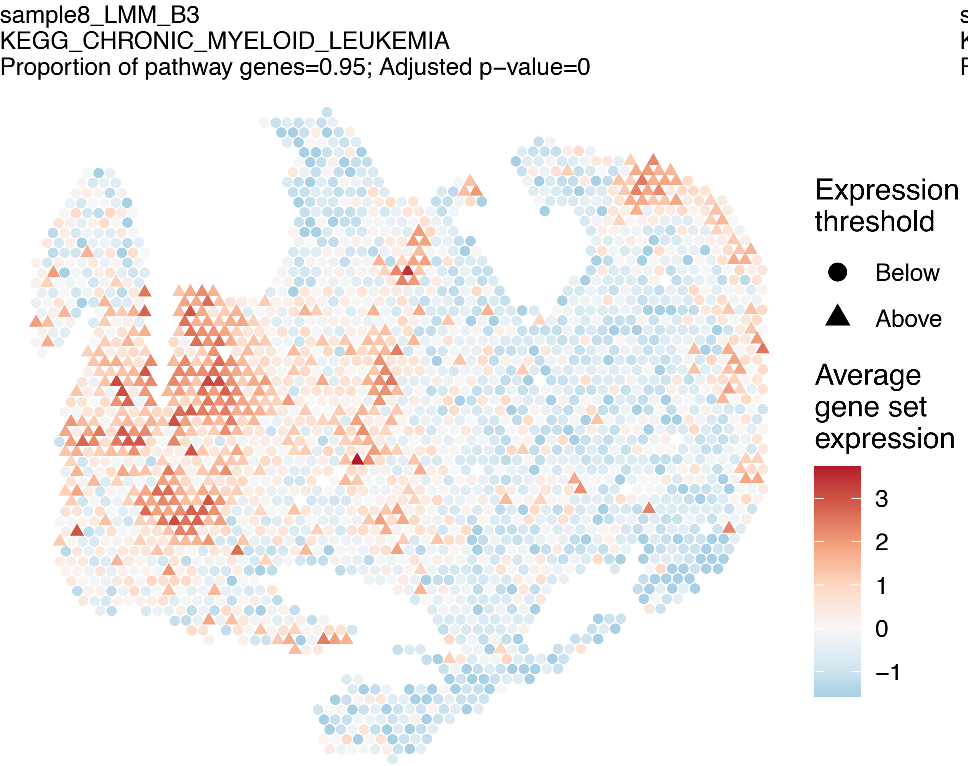
sample8_LMM_B3
KEGG_CHRONIC_MYELOID_LEUKEMIA
Proportion of pathway genes=0.95; Adjusted p−value=0
Expression threshold
Below
Above
Average gene set expression
3
2
1
0
−1

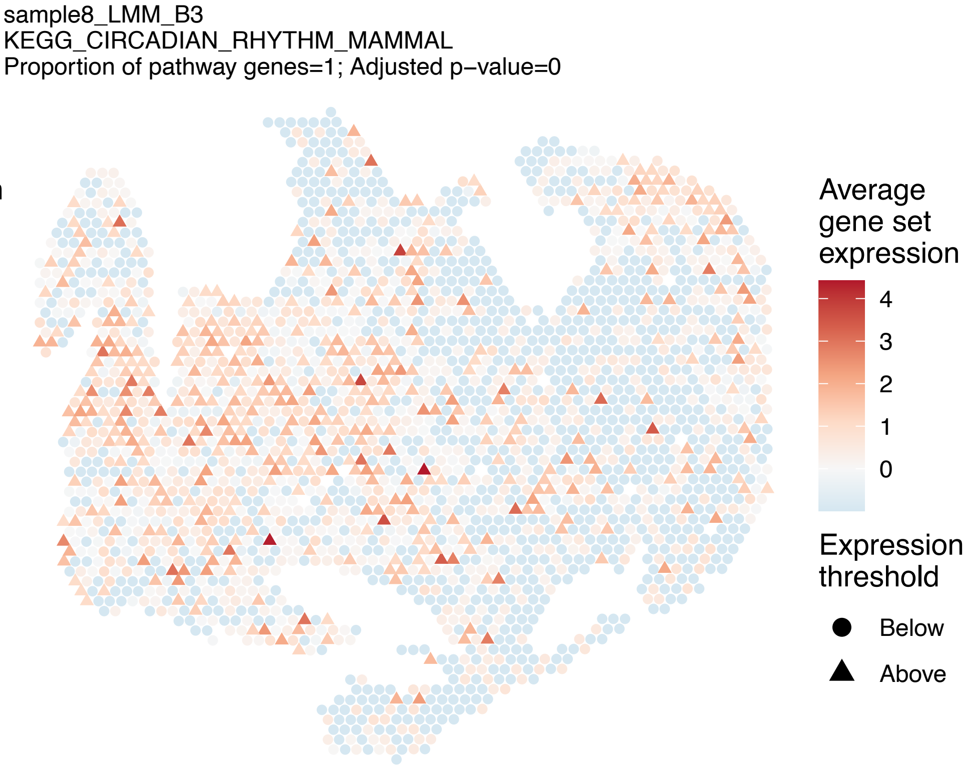
sample8_LMM_B3
KEGG_CIRCADIAN_RHYTHM_MAMMAL
Proportion of pathway genes=1; Adjusted p−value=0
Average gene set expression
4
3
2
1
0
Expression threshold
Below
Above

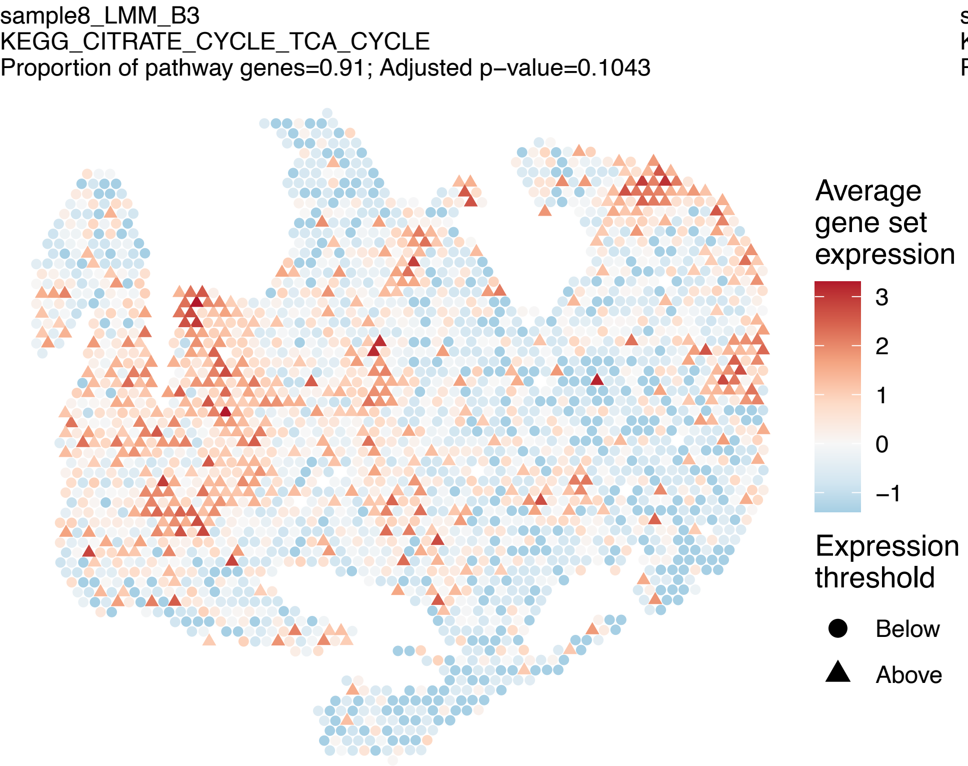
sample8_LMM_B3
KEGG_CITRATE_CYCLE_TCA_CYCLE
Proportion of pathway genes=0.91; Adjusted p−value=0.1043
Average gene set expression
3
2
1
0
−1
Expression threshold
Below
Above

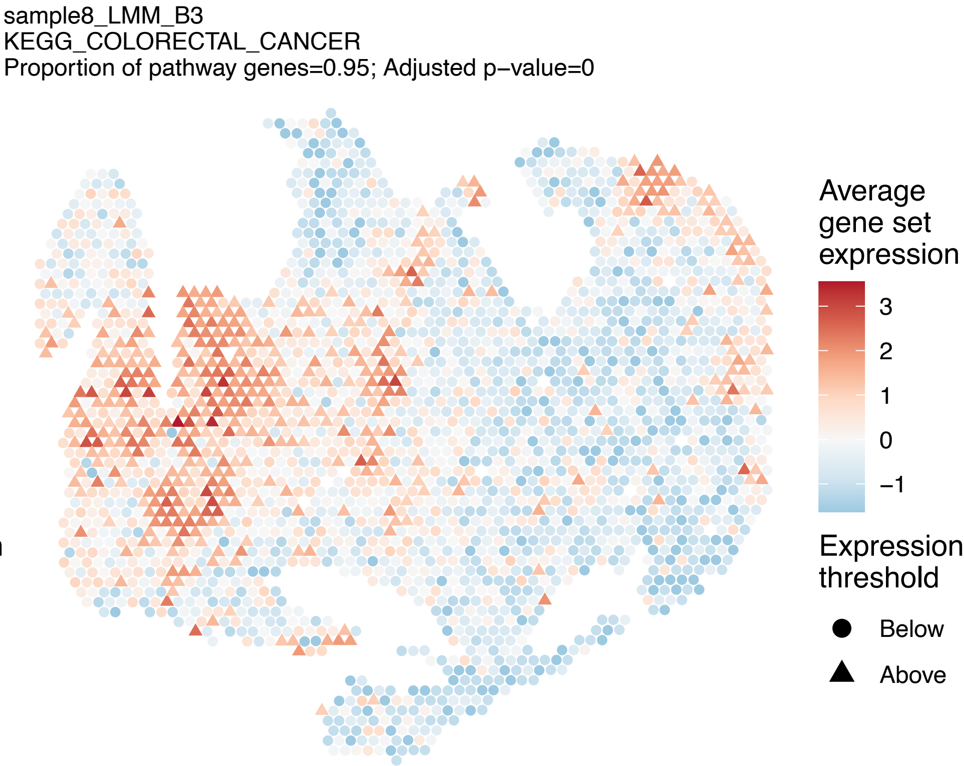
sample8_LMM_B3
KEGG_COLORECTAL_CANCER
Proportion of pathway genes=0.95; Adjusted p−value=0
Average gene set expression
3
2
1
0
−1
Expression threshold
Below
Above

sample8_LMM_B3
KEGG_COMPLEMENT_AND_COAGULATION_CASCADES
Proportion of pathway genes=0.54; Adjusted p−value=0

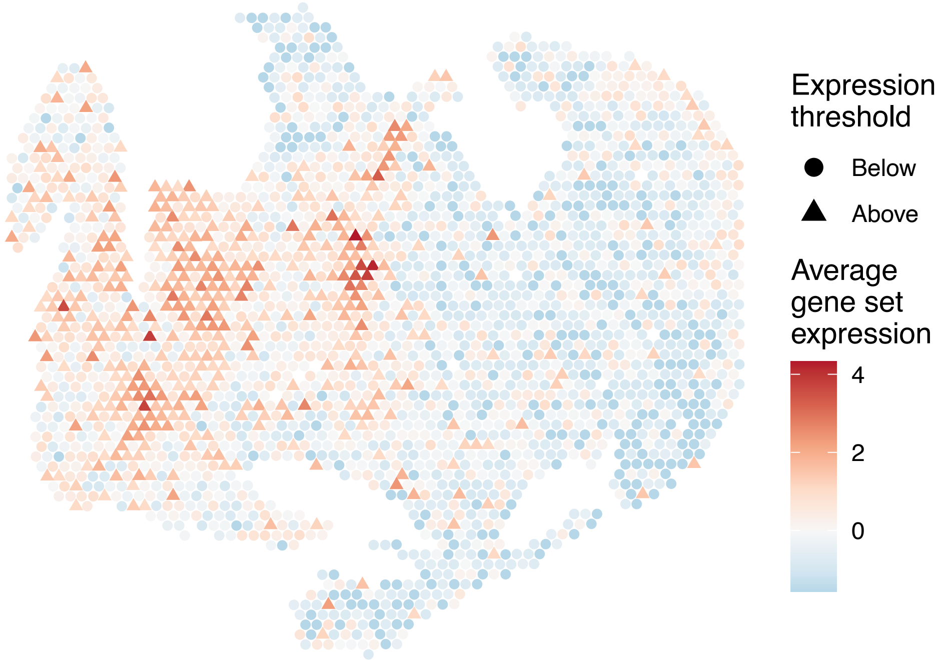


sample8_LMM_B3
KEGG_CYSTEINE_AND_METHIONINE_METABOLISM
Proportion of pathway genes=0.81; Adjusted p−value=0.0012

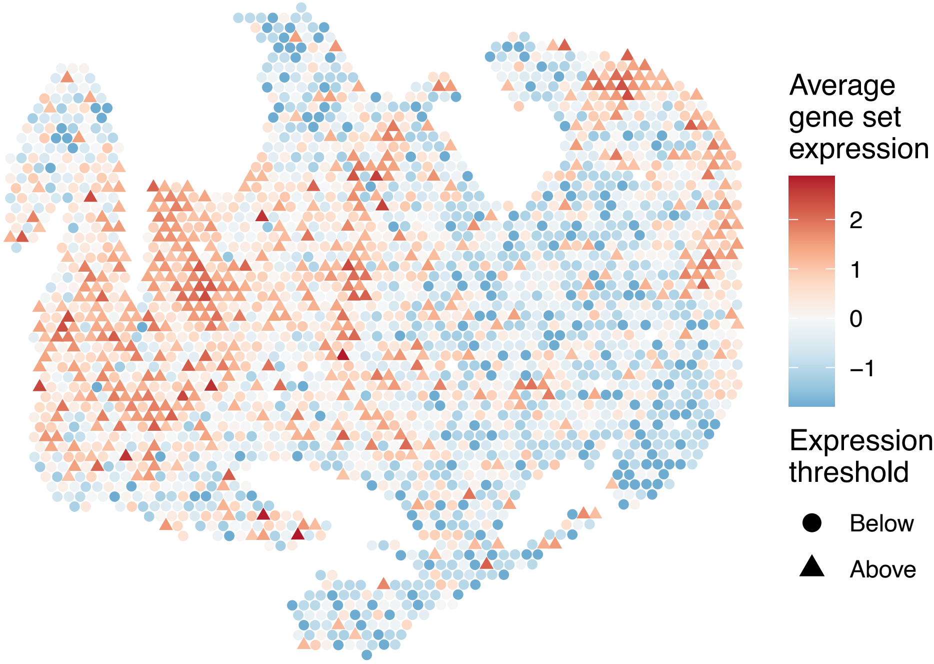


sample8_LMM_B3
KEGG_CYTOSOLIC_DNA_SENSING_PATHWAY
Proportion of pathway genes=0.52; Adjusted p−value=0

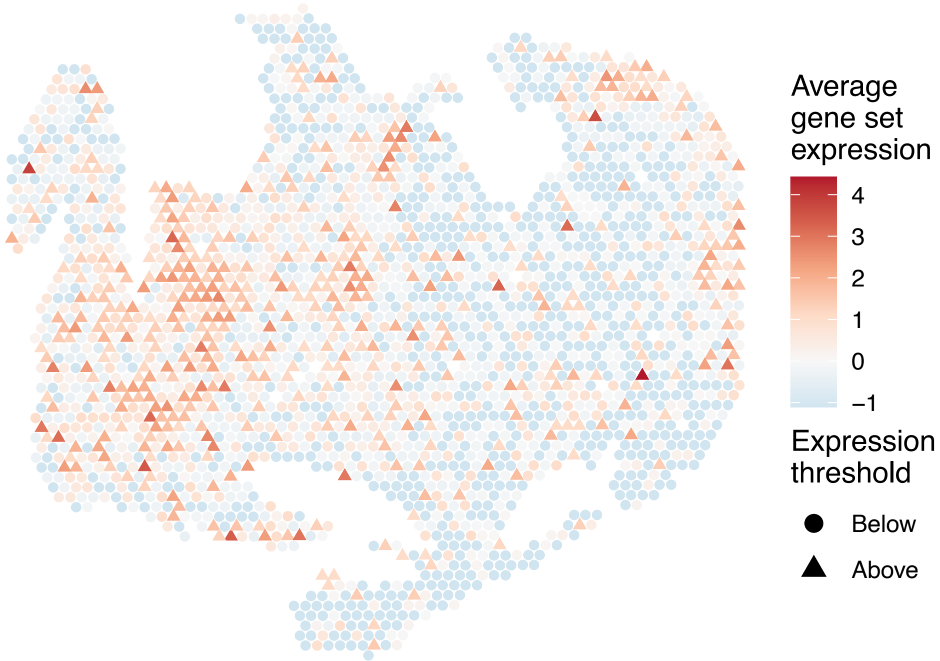


sample8_LMM_B3
KEGG_DILATED_CARDIOMYOPATHY
Proportion of pathway genes=0.62; Adjusted p−value=0

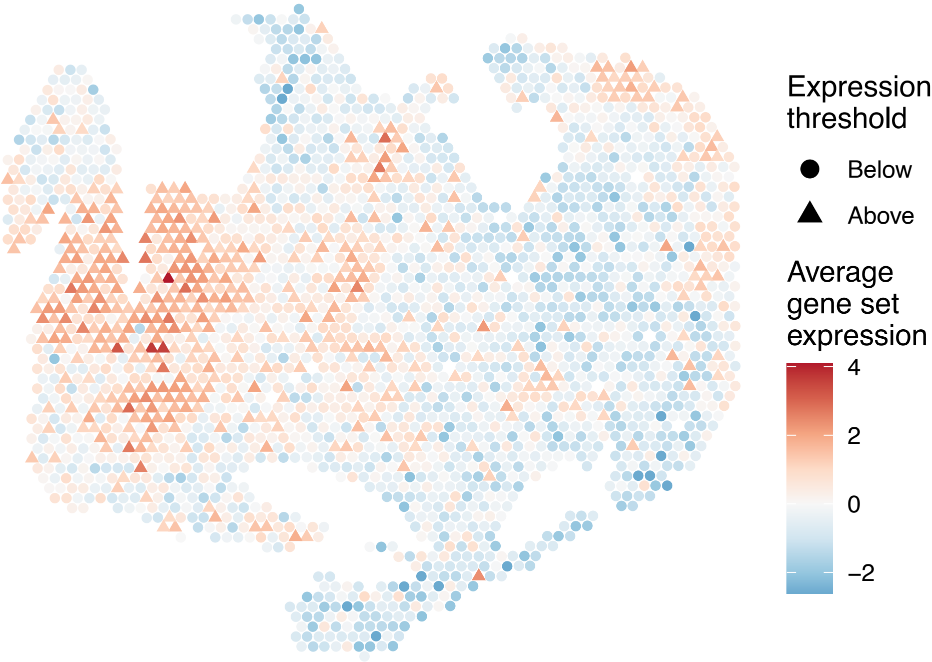

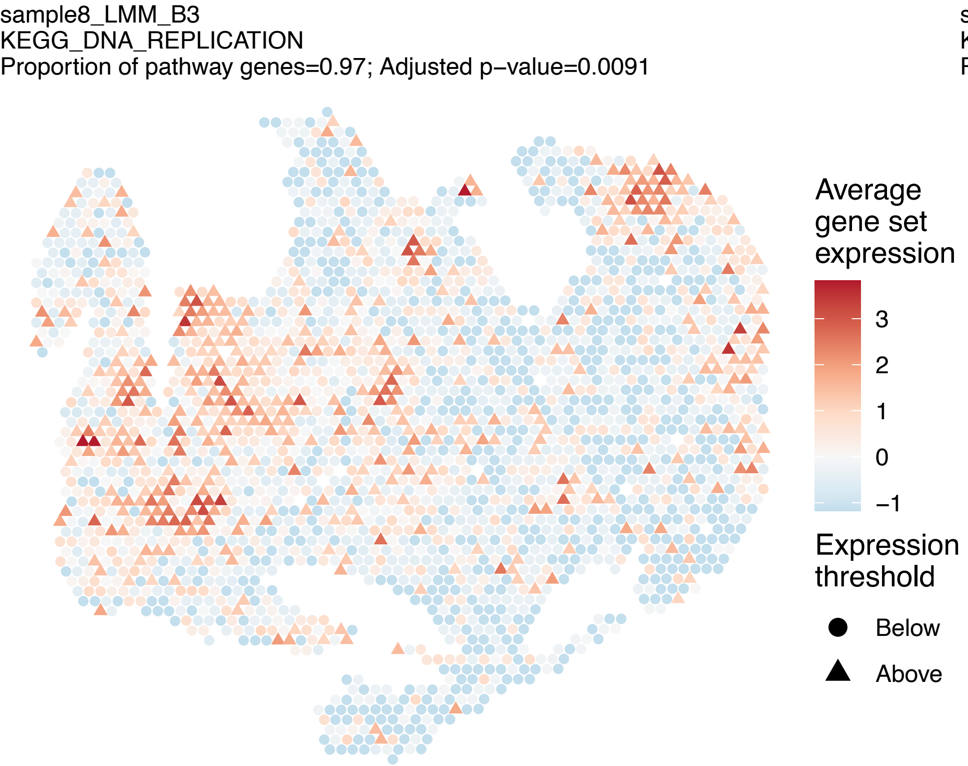
sample8_LMM_B3
KEGG_DNA_REPLICATION
Proportion of pathway genes=0.97; Adjusted p−value=0.0091
Average gene set expression
3
2
1
0
−1
Expression threshold
Below
Above

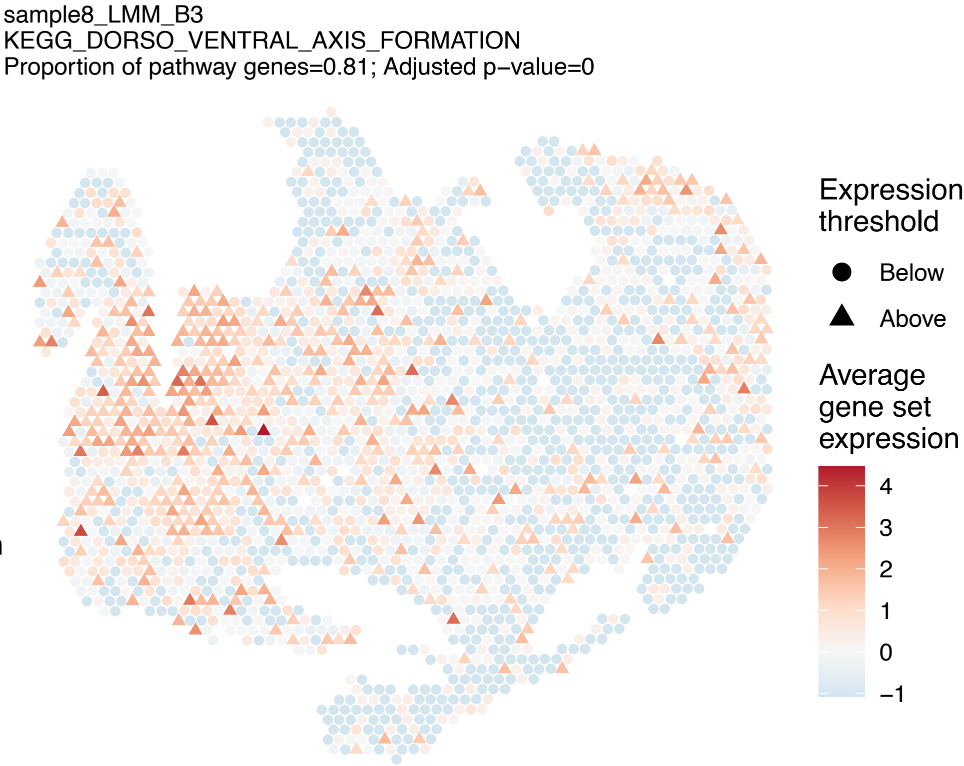
sample8_LMM_B3
KEGG_DORSO_VENTRAL_AXIS_FORMATION
Proportion of pathway genes=0.81; Adjusted p−value=0
Expression threshold
Below
Above
Average gene set expression
4
3
2
1
0
−1

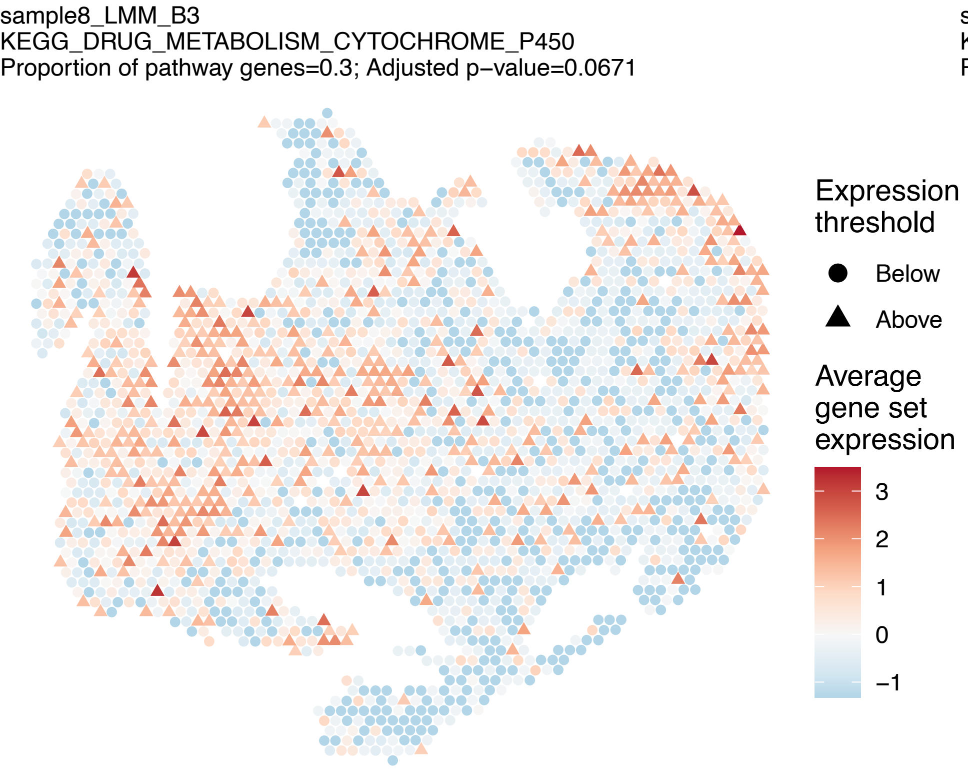
sample8_LMM_B3
KEGG_DRUG_METABOLISM_CYTOCHROME_P450
Proportion of pathway genes=0.3; Adjusted p−value=0.0671
Expression threshold
Below
Above
Average gene set expression
3
2
1
0
−1

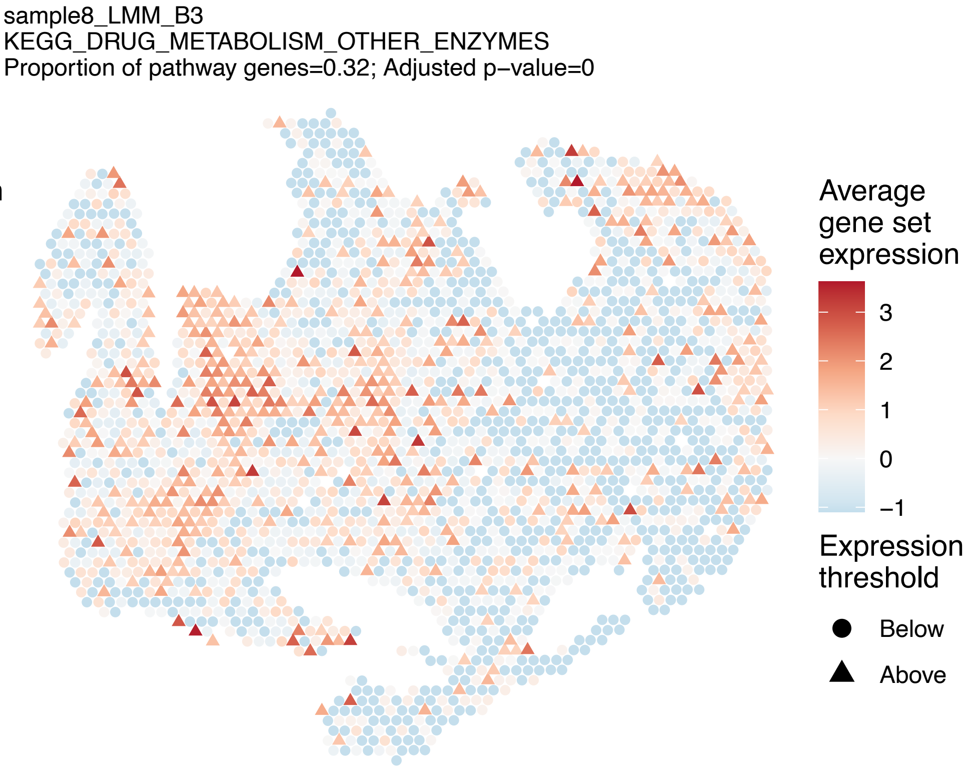
sample8_LMM_B3
KEGG_DRUG_METABOLISM_OTHER_ENZYMES
Proportion of pathway genes=0.32; Adjusted p−value=0
Average gene set expression
3
2
1
0
−1
Expression threshold
Below
Above

sample8_LMM_B3
KEGG_ECM_RECEPTOR_INTERACTION
Proportion of pathway genes=0.71; Adjusted p−value=0

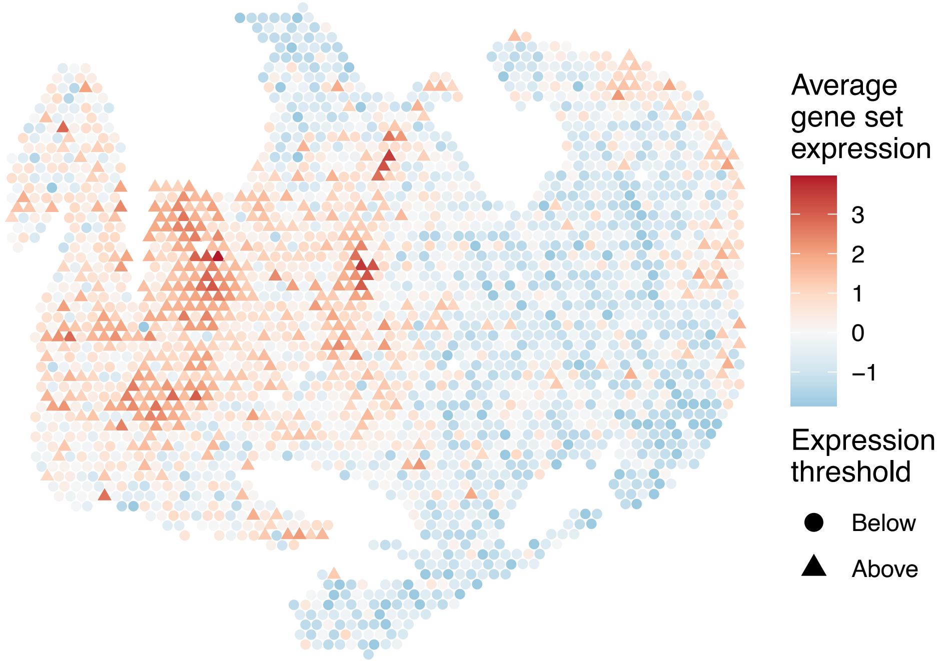


sample8_LMM_B3
KEGG_ENDOCYTOSIS
Proportion of pathway genes=0.89; Adjusted p−value=0

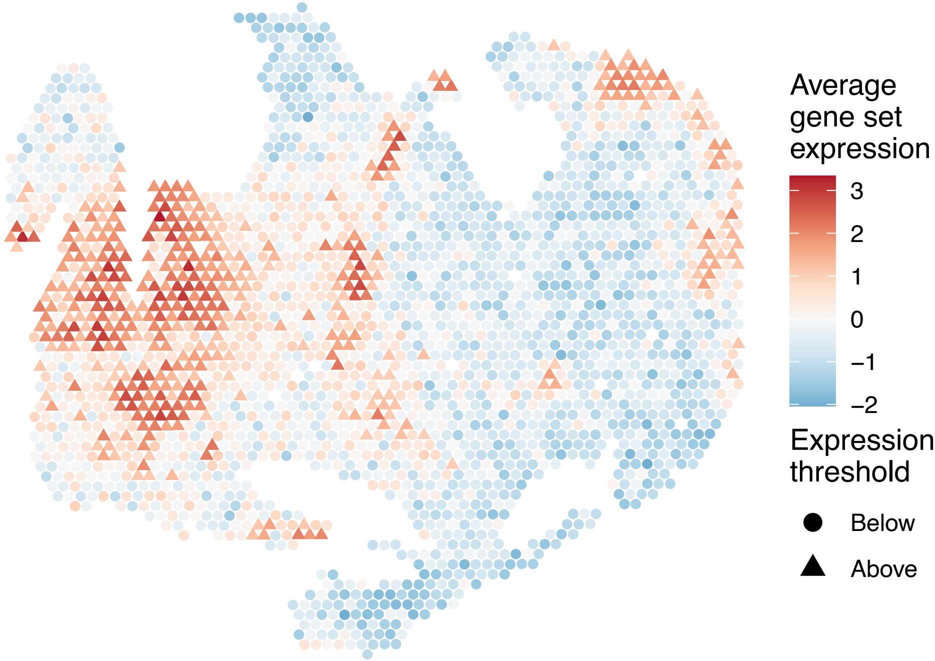


sample8_LMM_B3
KEGG_ENDOMETRIAL_CANCER
Proportion of pathway genes=0.91; Adjusted p−value=0

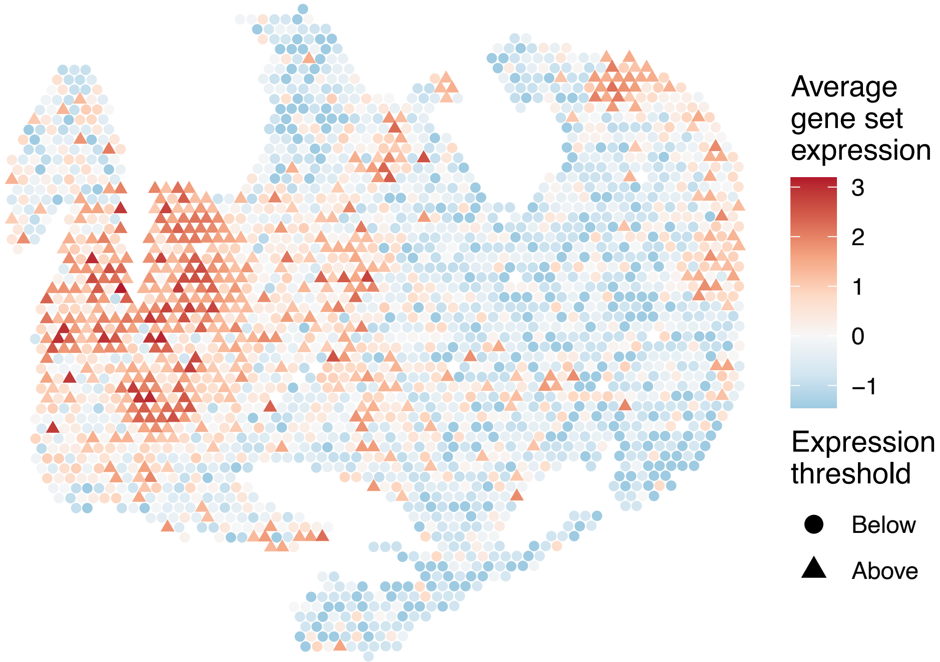


sample8_LMM_B3
KEGG_EPITHELIAL_CELL_SIGNALING_IN_HELICOBACTER_PYL
ORI_INFECTION
Proportion of pathway genes=0.89; Adjusted p−value=0

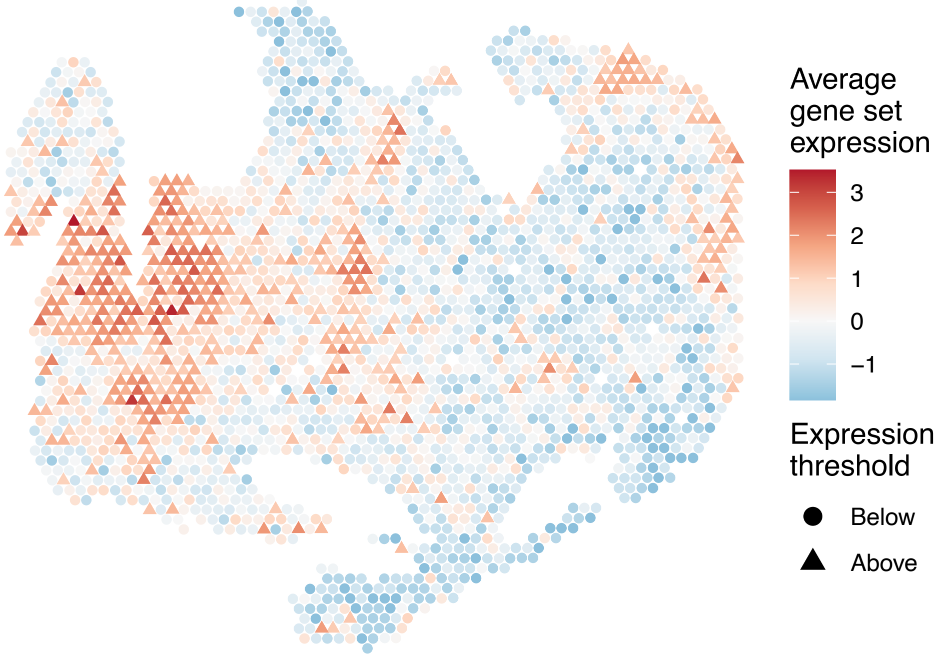

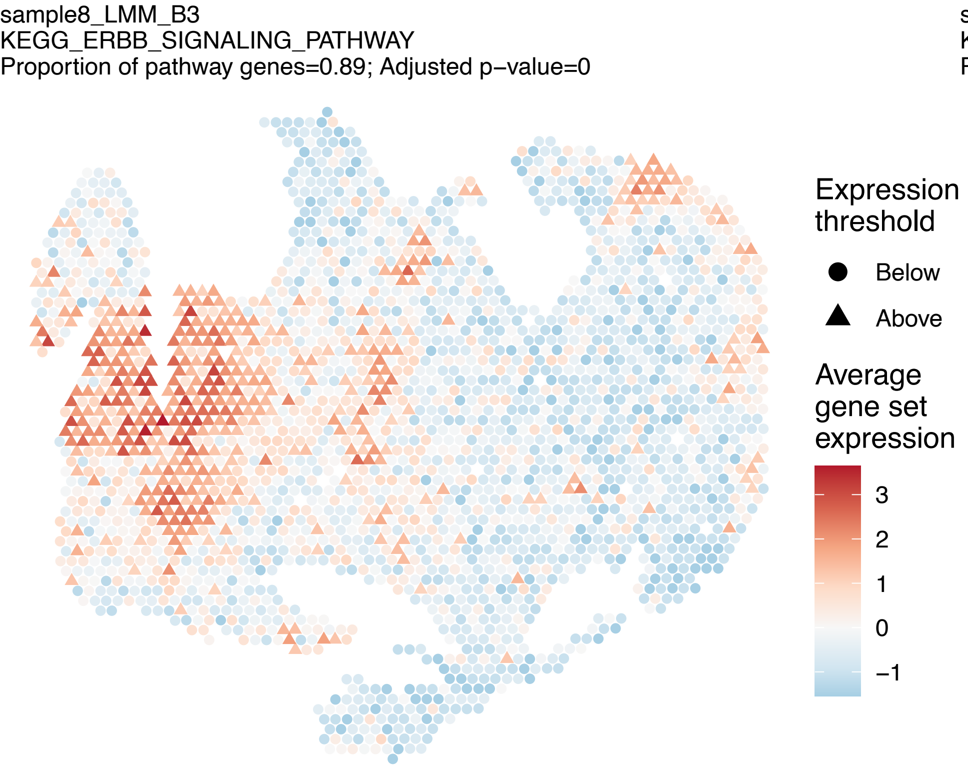
sample8_LMM_B3
KEGG_ERBB_SIGNALING_PATHWAY
Proportion of pathway genes=0.89; Adjusted p−value=0
Expression threshold
Below
Above
Average gene set expression
3
2
1
0
−1

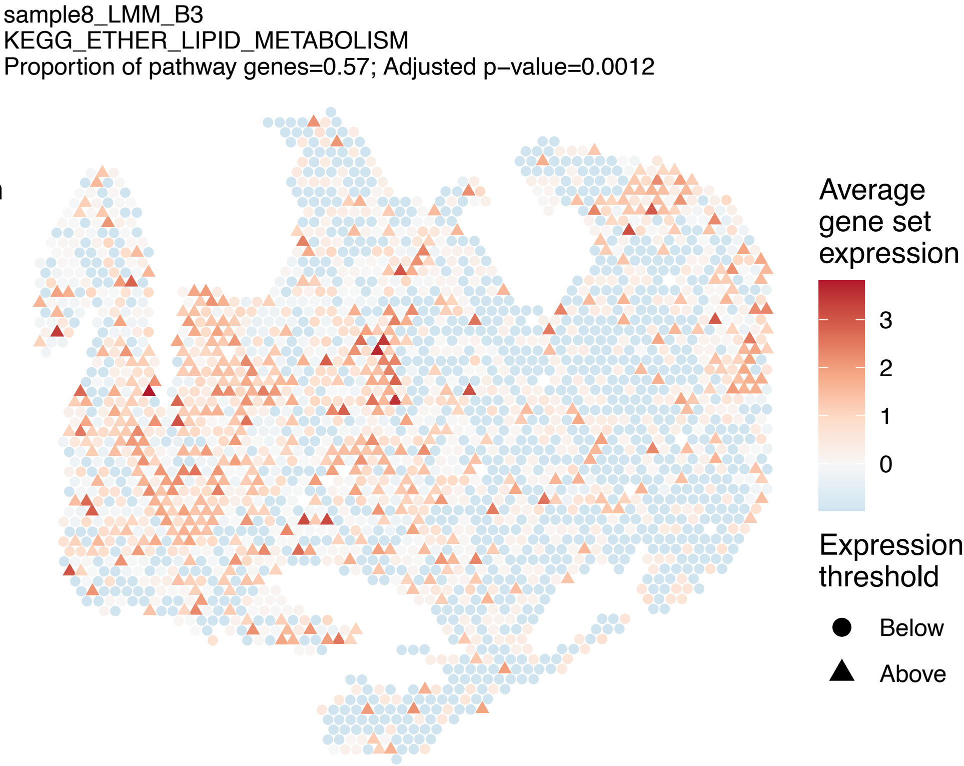
sample8_LMM_B3
KEGG_ETHER_LIPID_METABOLISM
Proportion of pathway genes=0.57; Adjusted p−value=0.0012
Average gene set expression
3
2
1
0
Expression threshold
Below
Above

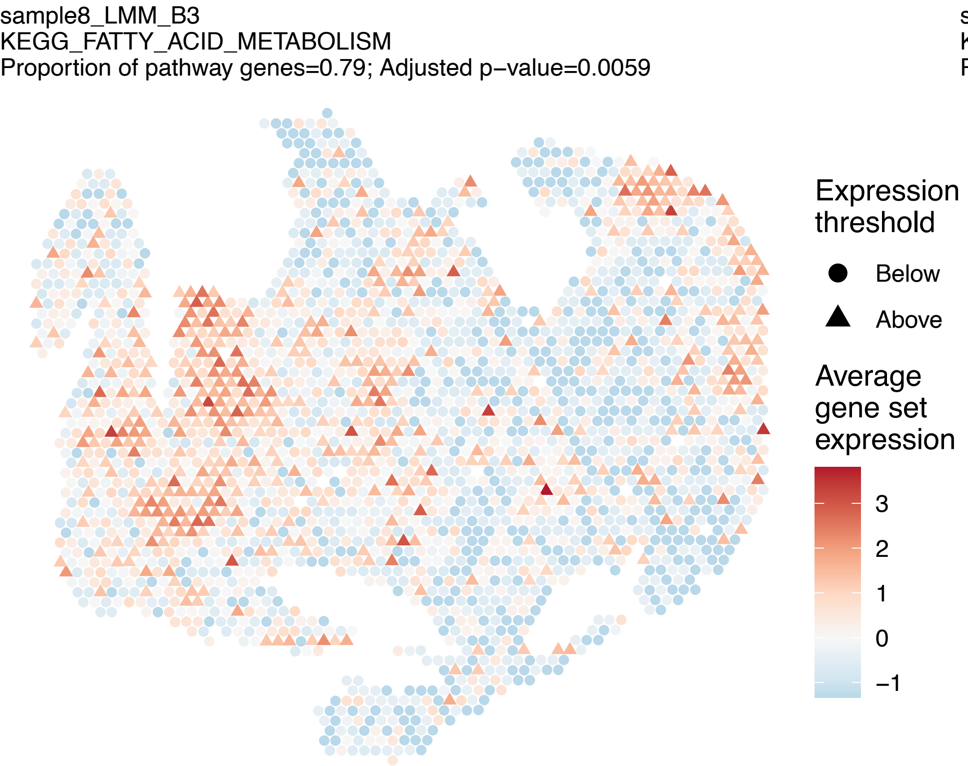
sample8_LMM_B3
KEGG_FATTY_ACID_METABOLISM
Proportion of pathway genes=0.79; Adjusted p−value=0.0059
Expression threshold
Below
Above
Average gene set expression
3
2
1
0
−1

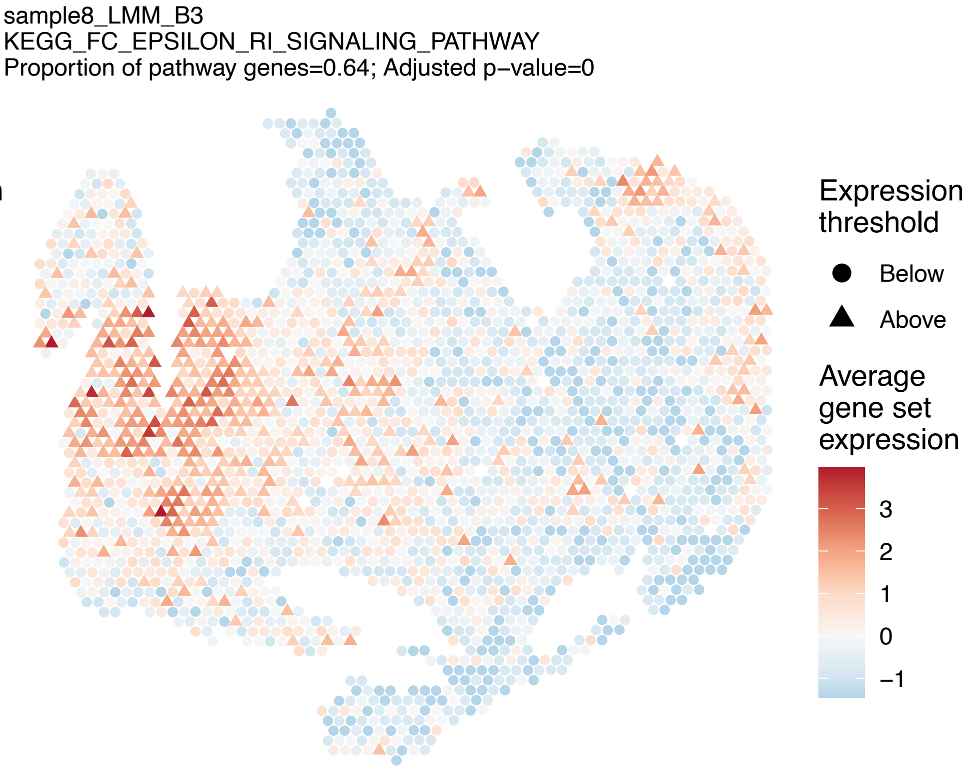
sample8_LMM_B3
KEGG_FC_EPSILON_RI_SIGNALING_PATHWAY
Proportion of pathway genes=0.64; Adjusted p−value=0
Expression threshold
Below
Above
Average gene set expression
3
2
1
0
−1

sample8_LMM_B3
KEGG_FC_GAMMA_R_MEDIATED_PHAGOCYTOSIS
Proportion of pathway genes=0.9; Adjusted p−value=0

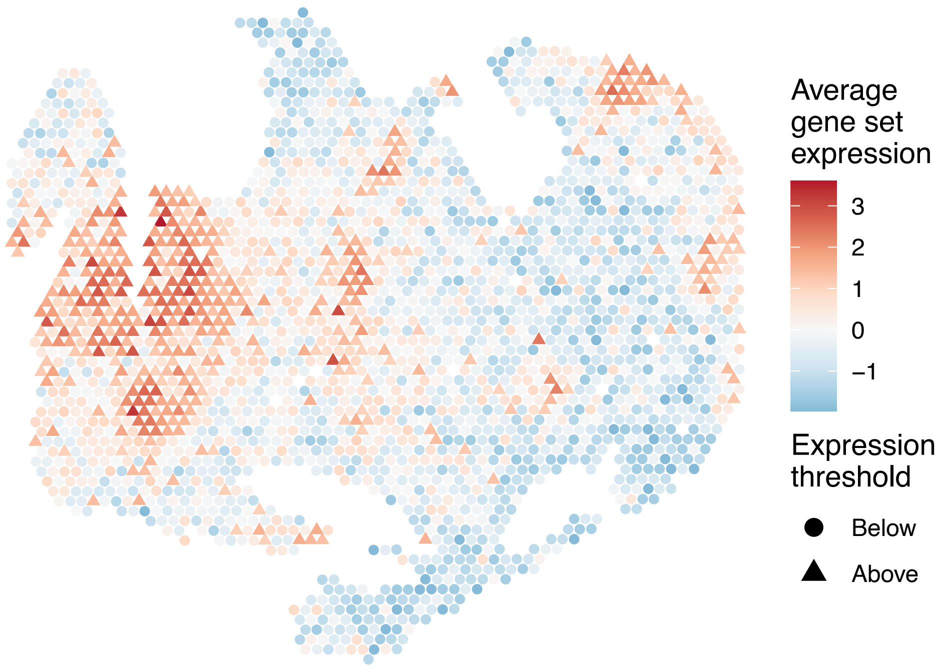


sample8_LMM_B3
KEGG_FOCAL_ADHESION
Proportion of pathway genes=0.85; Adjusted p−value=0

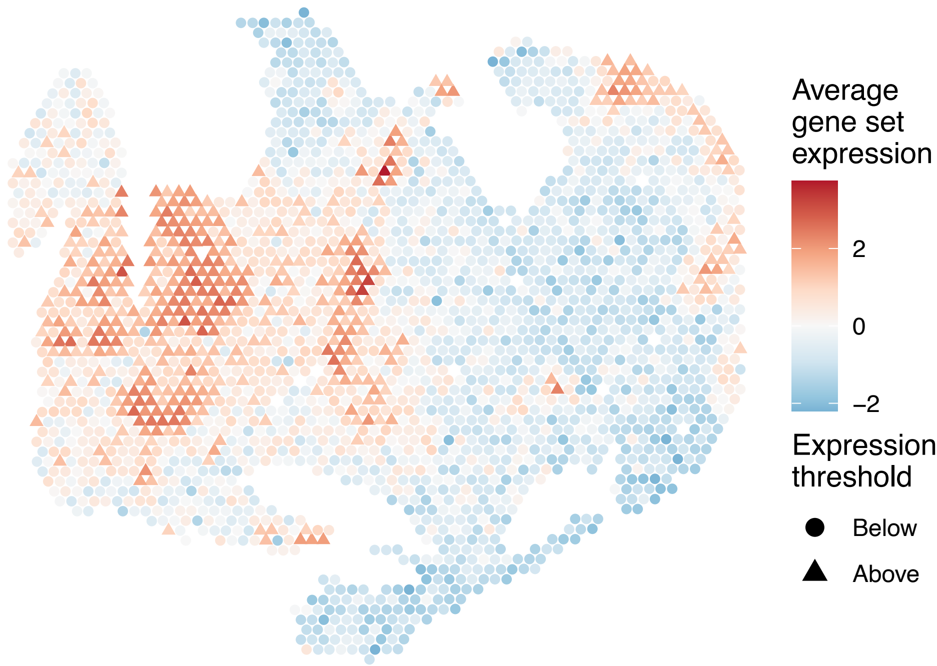


sample8_LMM_B3
KEGG_FRUCTOSE_AND_MANNOSE_METABOLISM
Proportion of pathway genes=0.8; Adjusted p−value=0

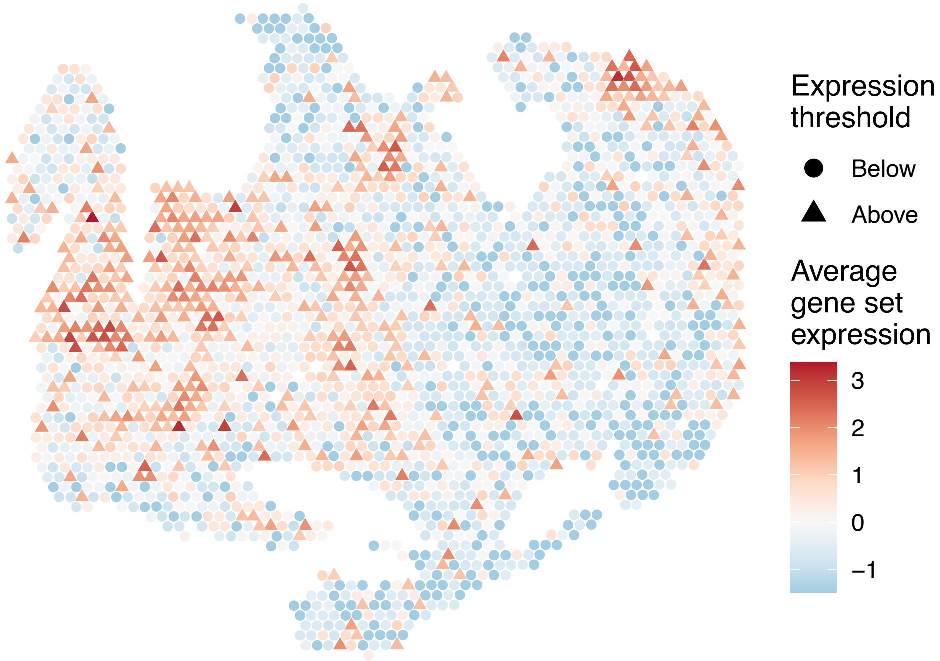


sample8_LMM_B3
KEGG_GALACTOSE_METABOLISM
Proportion of pathway genes=0.66; Adjusted p−value=0

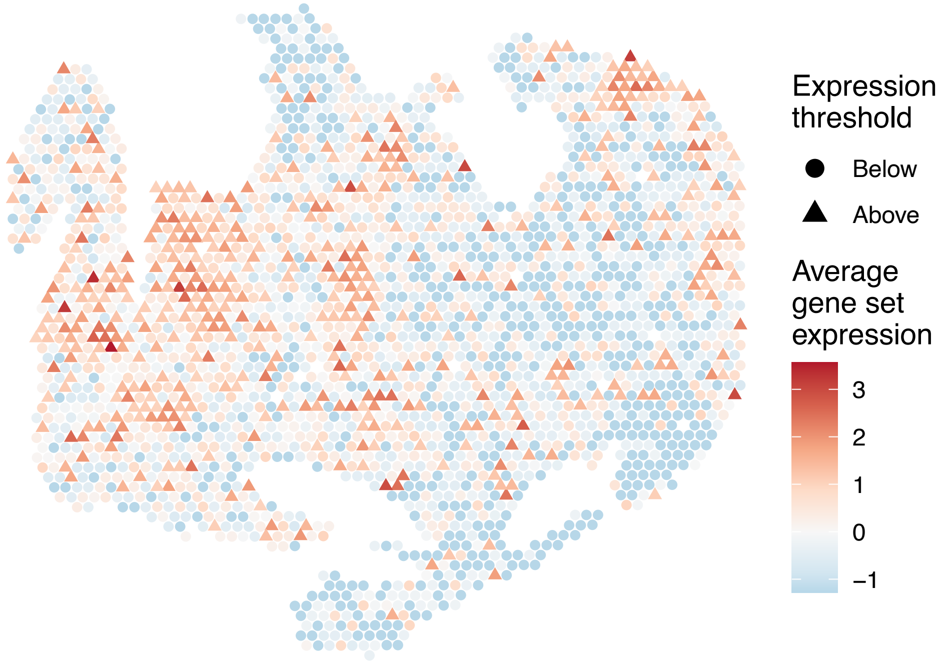

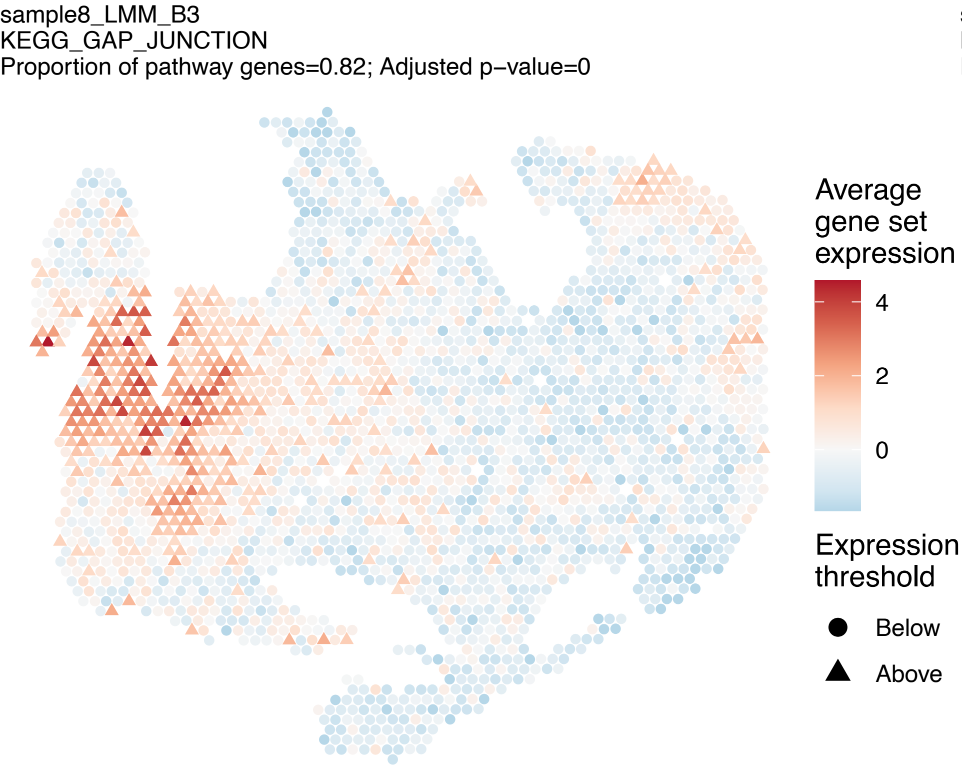

sample8_LMM_B3
KEGG_GAP_JUNCTION
Proportion of pathway genes=0.82; Adjusted p−value=0
Average gene set expression
4
2
0
Expression threshold
Below
Above


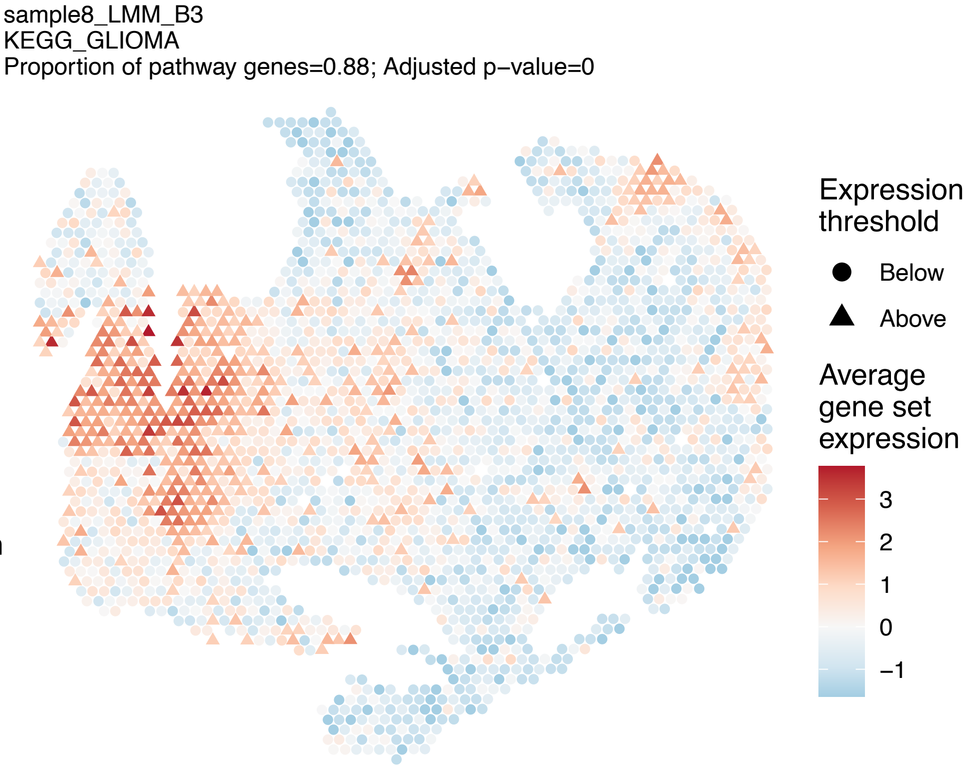

sample8_LMM_B3
KEGG_GLIOMA
Proportion of pathway genes=0.88; Adjusted p−value=0
Expression threshold
Below
Above
Average gene set expression
3
2
1
0
−1


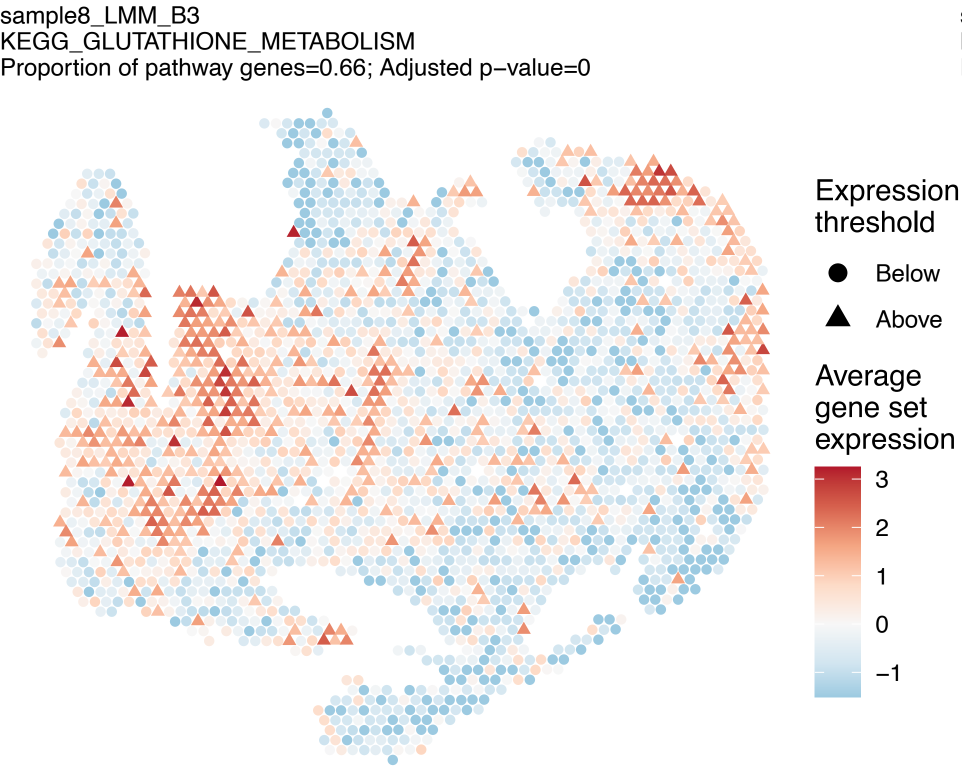

sample8_LMM_B3
KEGG_GLUTATHIONE_METABOLISM
Proportion of pathway genes=0.66; Adjusted p−value=0
Expression threshold
Below
Above
Average gene set expression
3
2
1
0
−1


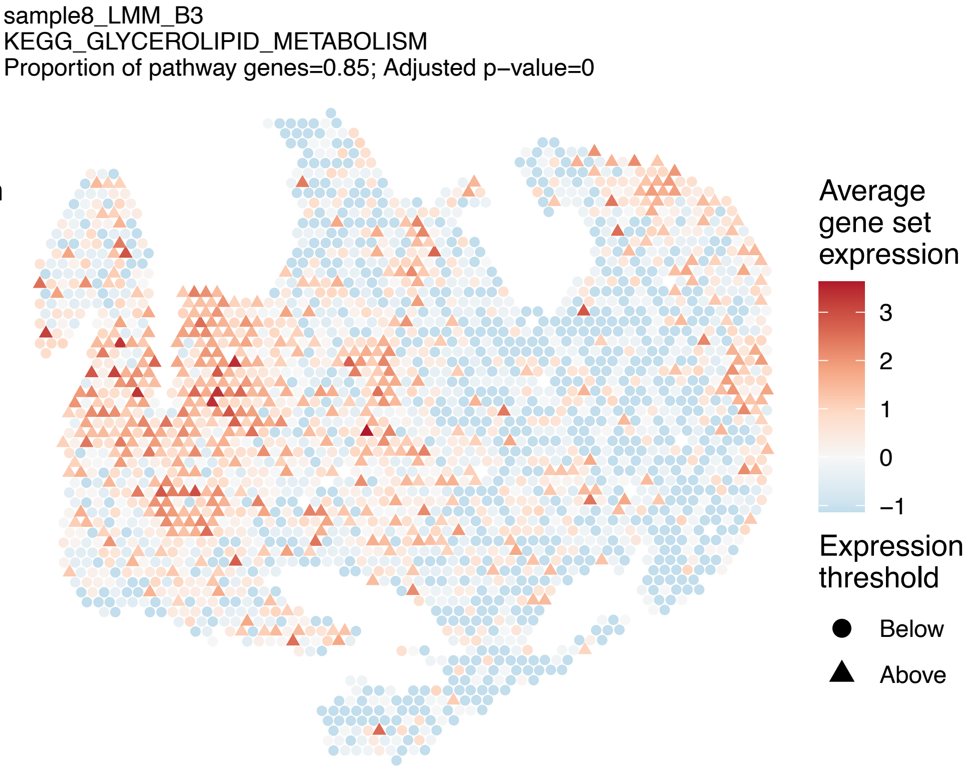

sample8_LMM_B3
KEGG_GLYCEROLIPID_METABOLISM
Proportion of pathway genes=0.85; Adjusted p−value=0
Average gene set expression
3
2
1
0
−1
Expression threshold
Below
Above

sample8_LMM_B3
KEGG_GLYCEROPHOSPHOLIPID_METABOLISM
Proportion of pathway genes=0.8; Adjusted p−value=0

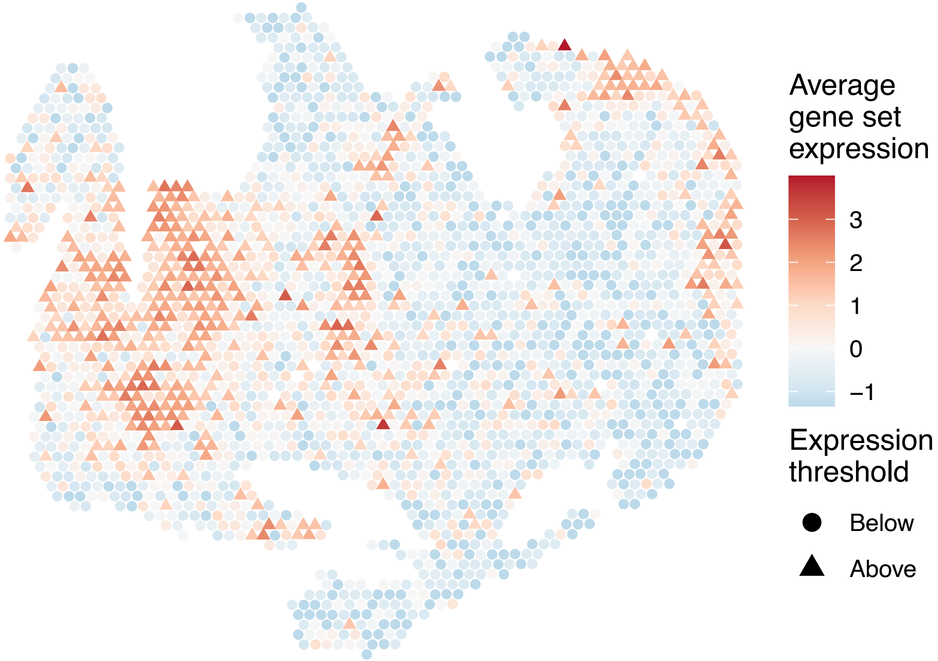


sample8_LMM_B3
KEGG_GLYCINE_SERINE_AND_THREONINE_METABOLISM
Proportion of pathway genes=0.62; Adjusted p−value=0.0012

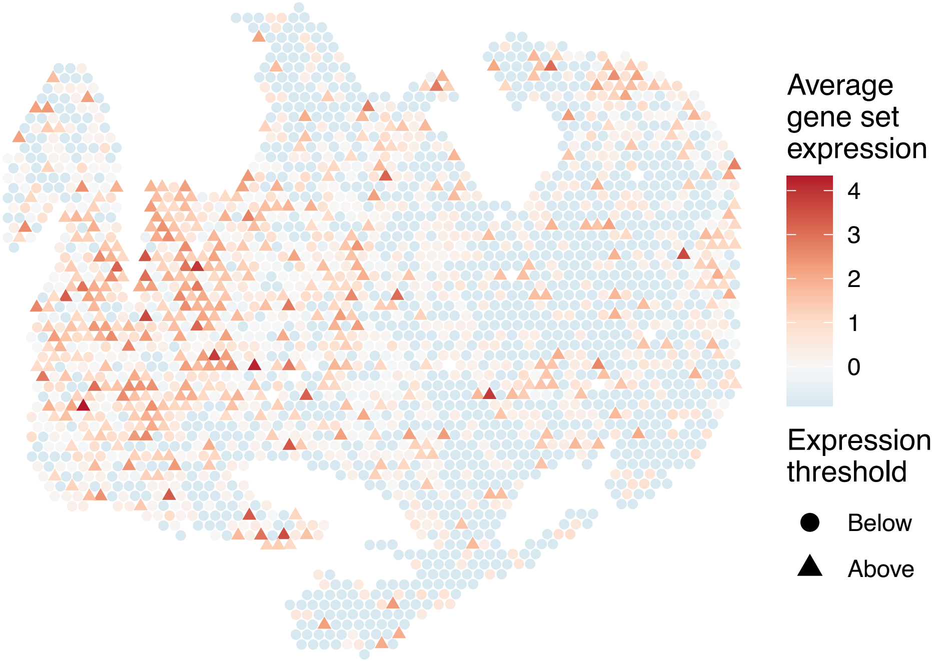


sample8_LMM_B3
KEGG_GLYCOLYSIS_GLUCONEOGENESIS
Proportion of pathway genes=0.61; Adjusted p−value=0

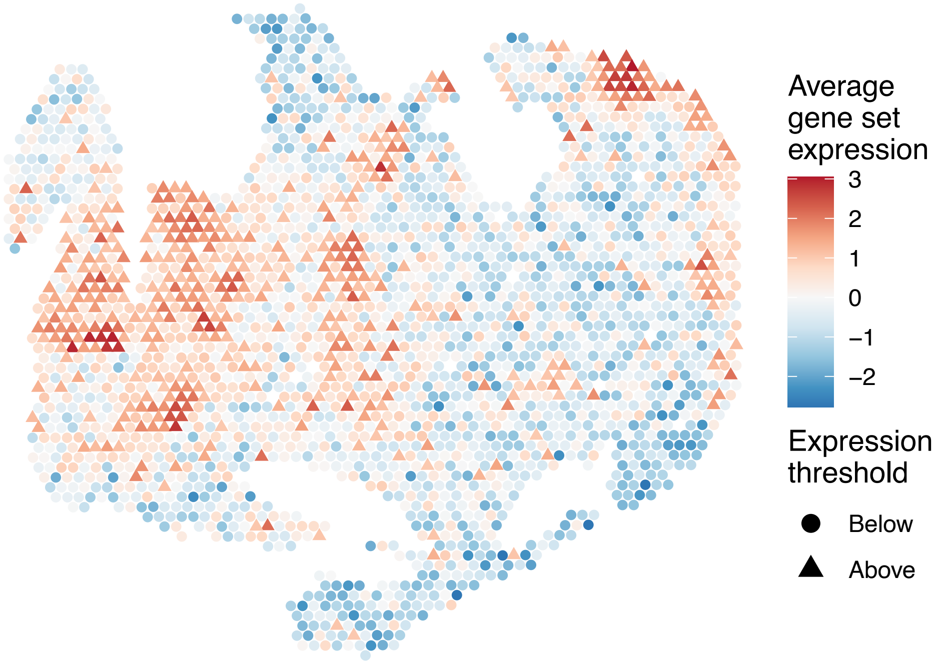


sample8_LMM_B3
KEGG_GLYCOSAMINOGLYCAN_BIOSYNTHESIS_CHONDROITIN_SU
LFATE
Proportion of pathway genes=0.91; Adjusted p−value=0

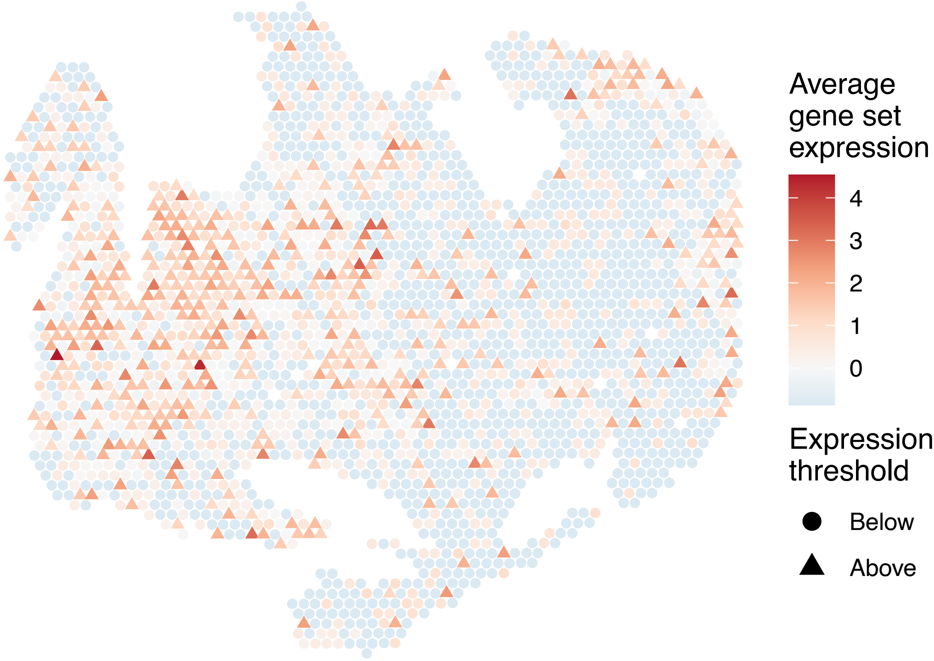

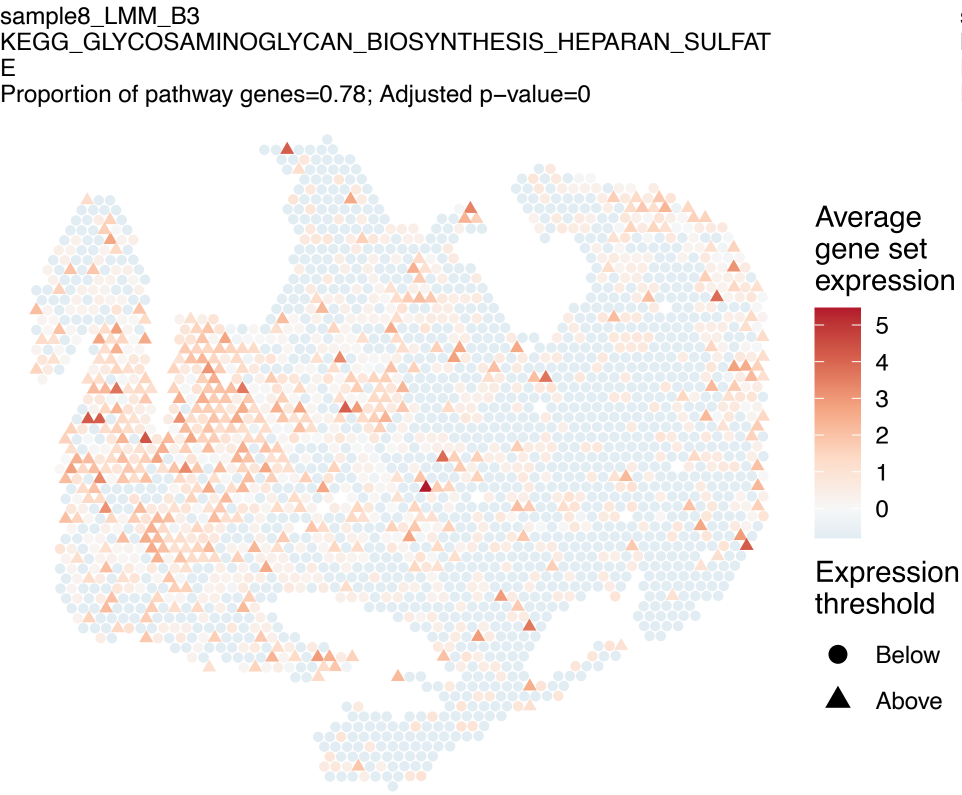
sample8_LMM_B3
KEGG_GLYCOSAMINOGLYCAN_BIOSYNTHESIS_HEPARAN_SULFAT
E
Proportion of pathway genes=0.78; Adjusted p−value=0
Average gene set expression
5
4
3
2
1
0
Expression threshold
Below
Above

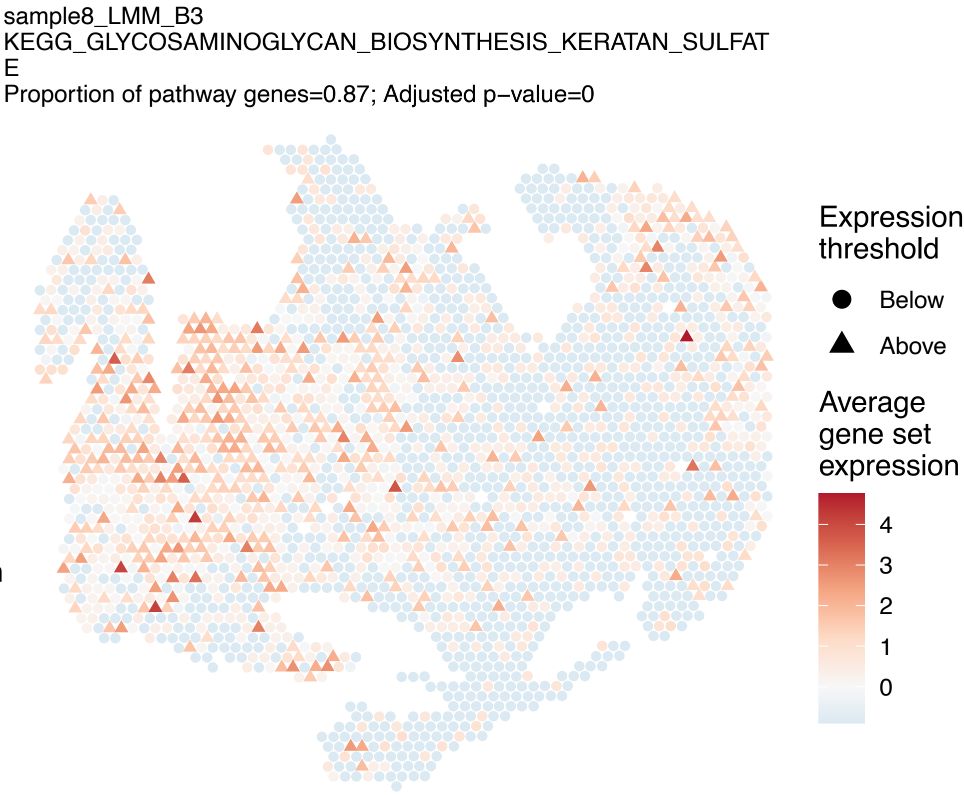
sample8_LMM_B3
KEGG_GLYCOSAMINOGLYCAN_BIOSYNTHESIS_KERATAN_SULFAT
E
Proportion of pathway genes=0.87; Adjusted p−value=0
Expression threshold
Below
Above
Average gene set expression
4
3
2
1
0

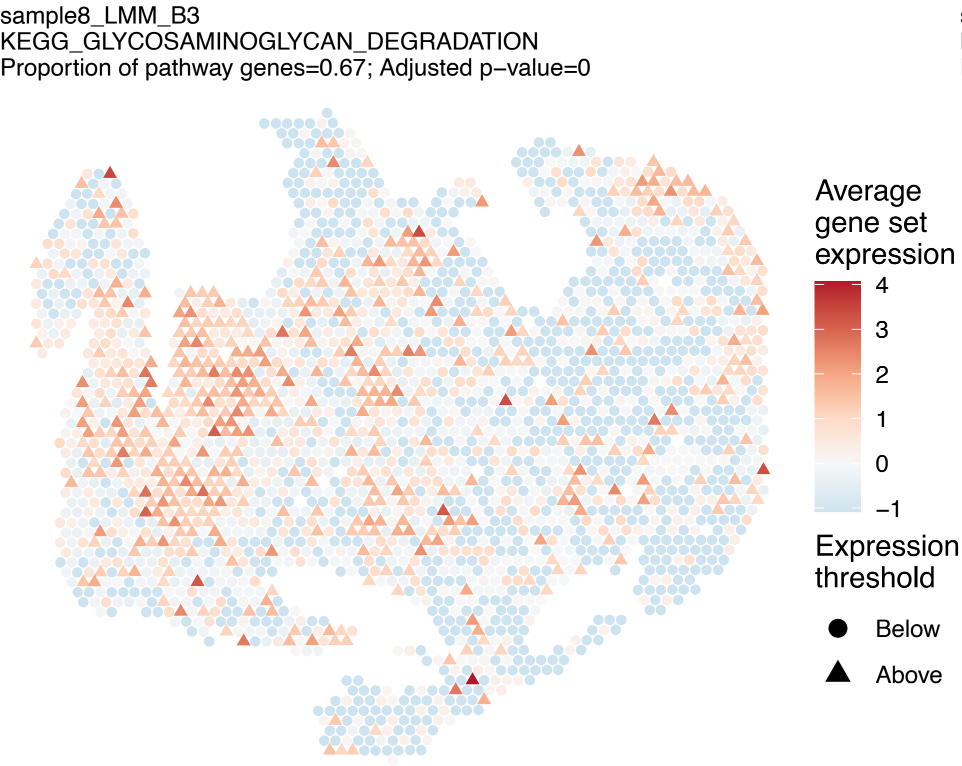
sample8_LMM_B3
KEGG_GLYCOSAMINOGLYCAN_DEGRADATION
Proportion of pathway genes=0.67; Adjusted p−value=0
Average gene set expression
4
3
2
1
0
−1
Expression threshold
Below
Above

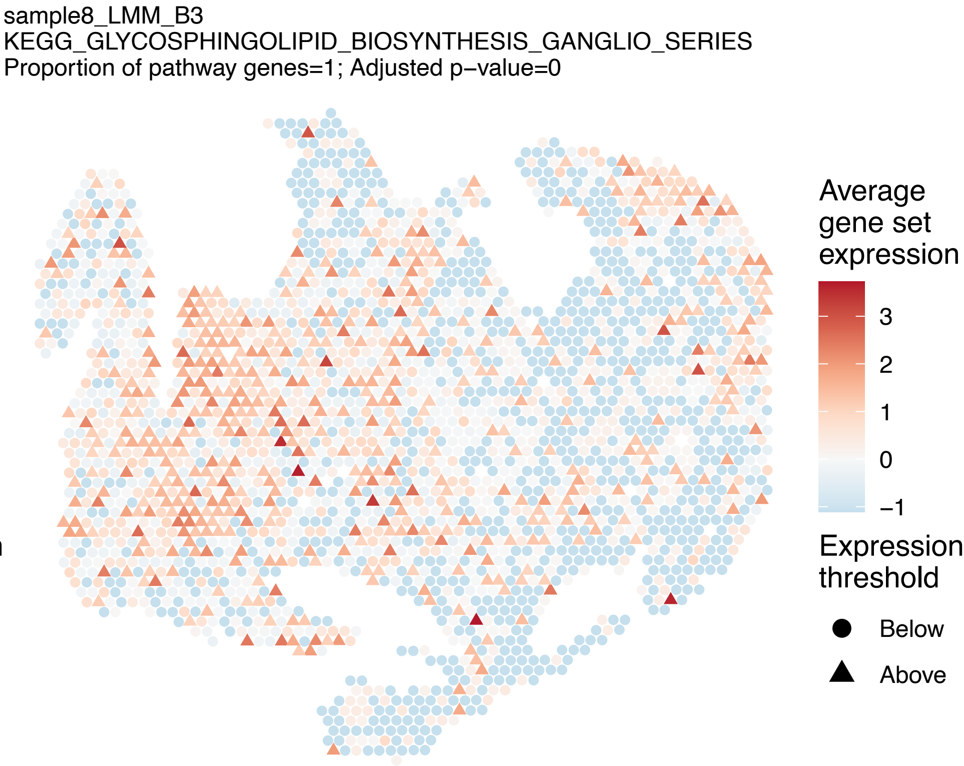
sample8_LMM_B3
KEGG_GLYCOSPHINGOLIPID_BIOSYNTHESIS_GANGLIO_SERIES
Proportion of pathway genes=1; Adjusted p−value=0
Average gene set expression
3
2
1
0
−1
Expression threshold
Below
Above

sample8_LMM_B3
KEGG_GLYCOSPHINGOLIPID_BIOSYNTHESIS_GLOBO_SERIES
Proportion of pathway genes=0.79; Adjusted p–value=0

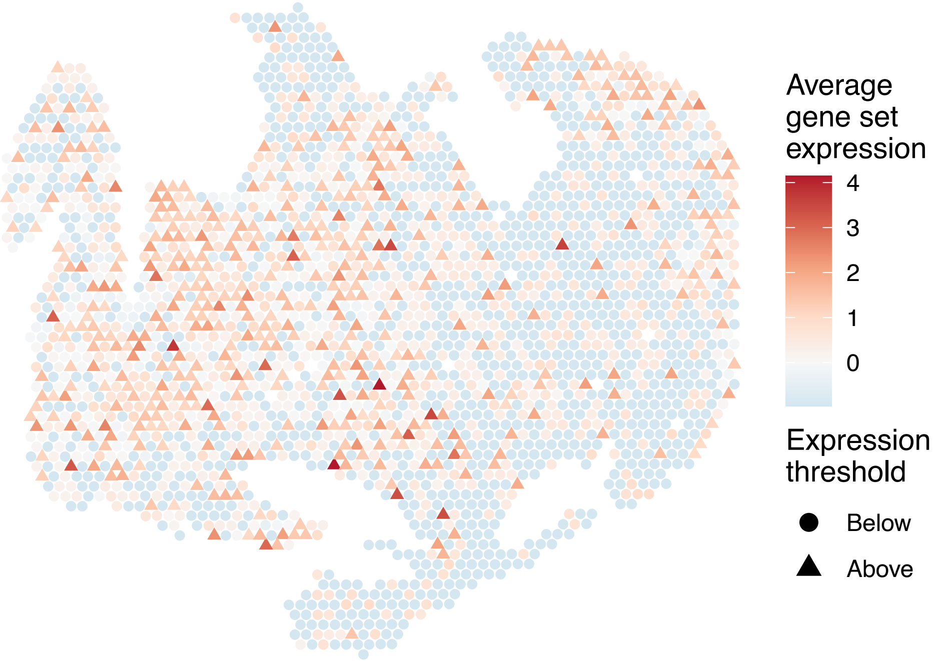


sample8_LMM_B3
KEGG_GLYCOSPHINGOLIPID_BIOSYNTHESIS_LACTO_AND_NEOL
ACTO_SERIES
Proportion of pathway genes=0.55; Adjusted p–value=0.007

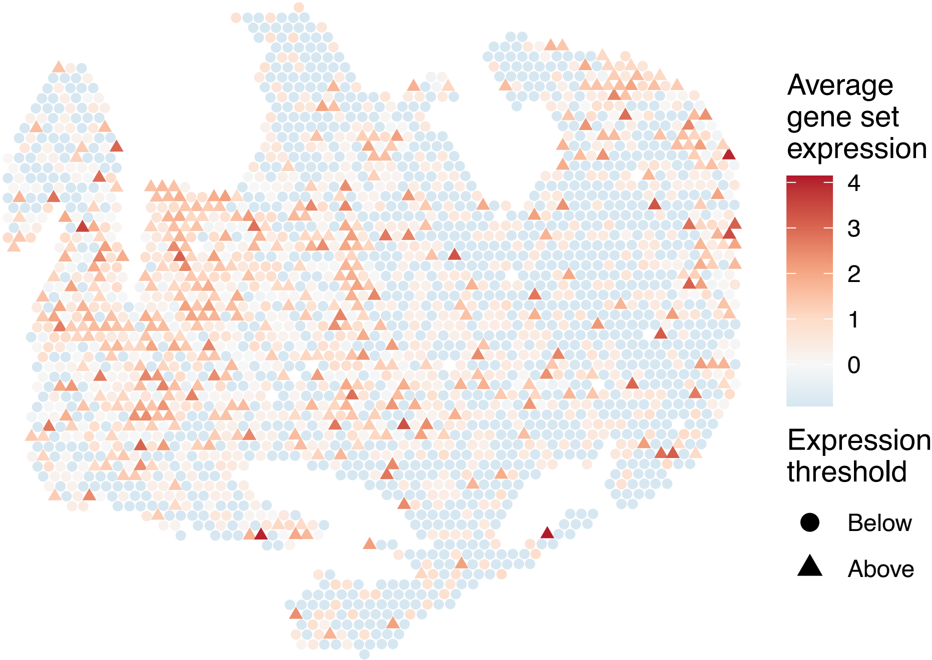


sample8_LMM_B3
KEGG_GLYCOSYLPHOSPHATIDYLINOSITOL_GPI_ANCHOR_BIOSY
NTHESIS
Proportion of pathway genes=0.96; Adjusted p–value=0

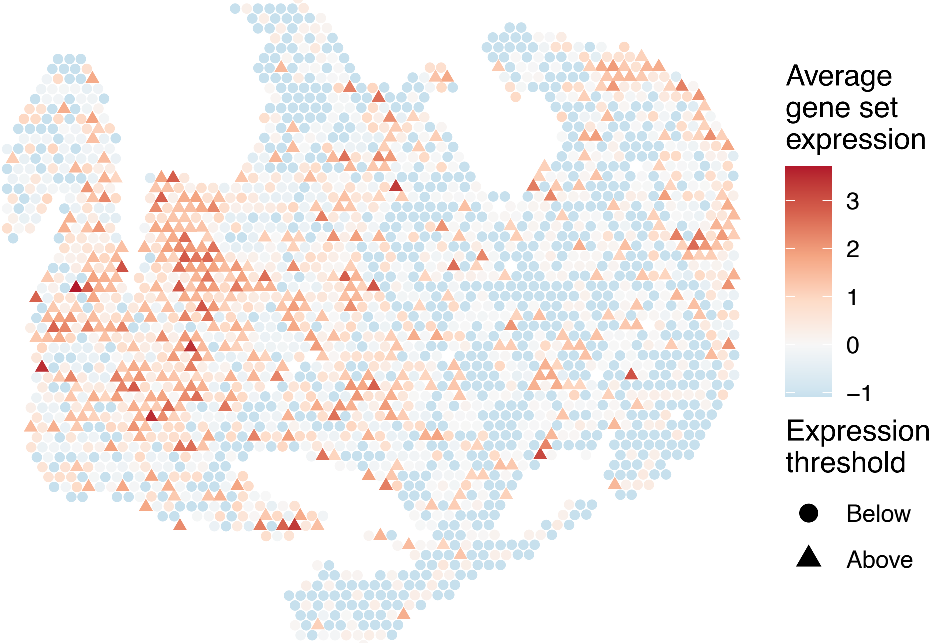


sample8_LMM_B3
KEGG_GLYOXYLATE_AND_DICARBOXYLATE_METABOLISM
Proportion of pathway genes=0.88; Adjusted p–value=0.1811

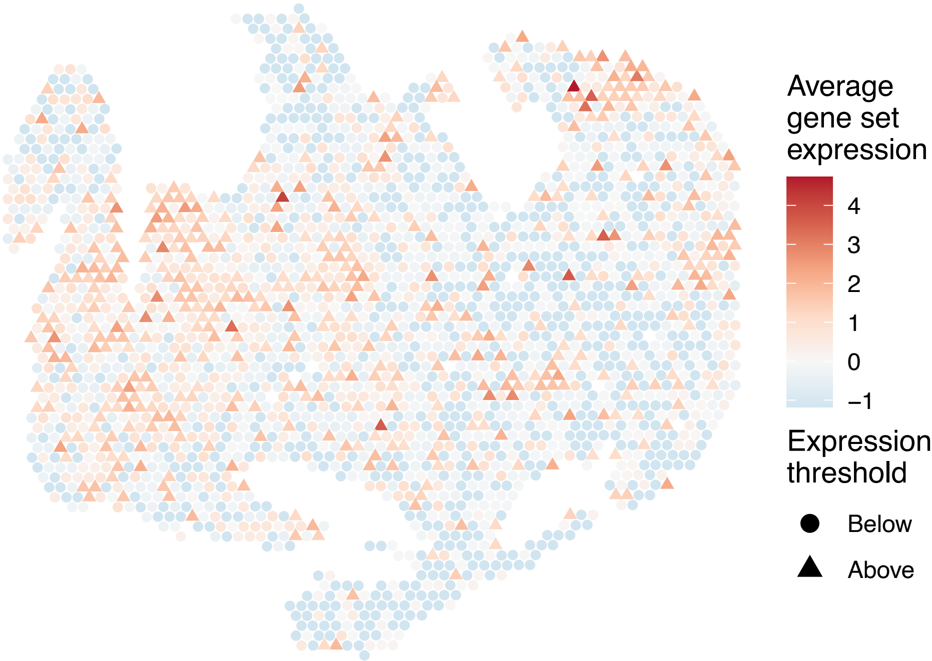

sample8_LMM_B3
KEGG_GNRH_SIGNALING_PATHWAY
Proportion of pathway genes=0.74; Adjusted p–value=0

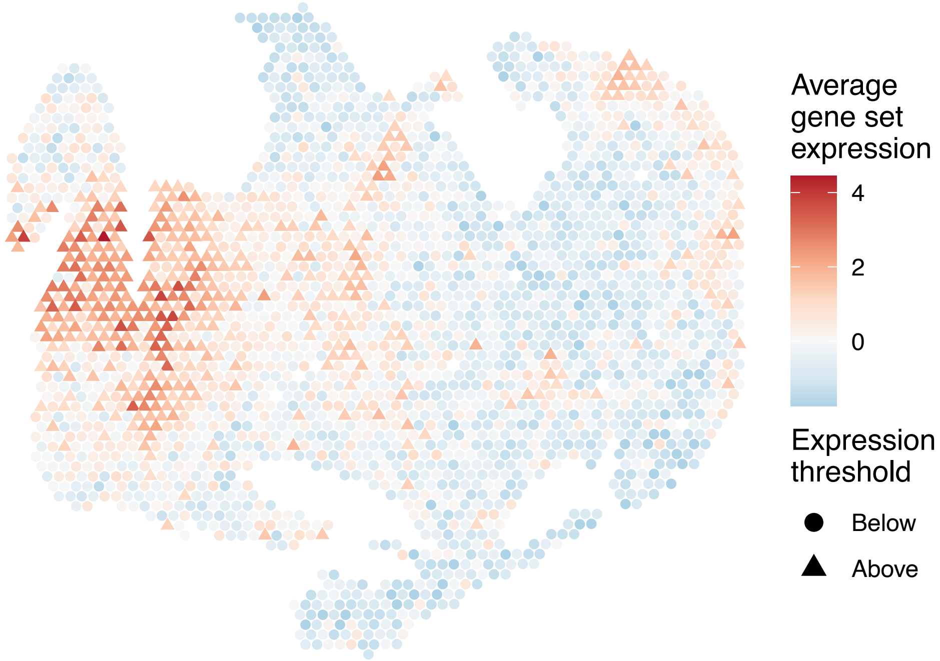


sample8_LMM_B3
KEGG_GRAFT_VERSUS_HOST_DISEASE
Proportion of pathway genes=0.31; Adjusted p–value=0

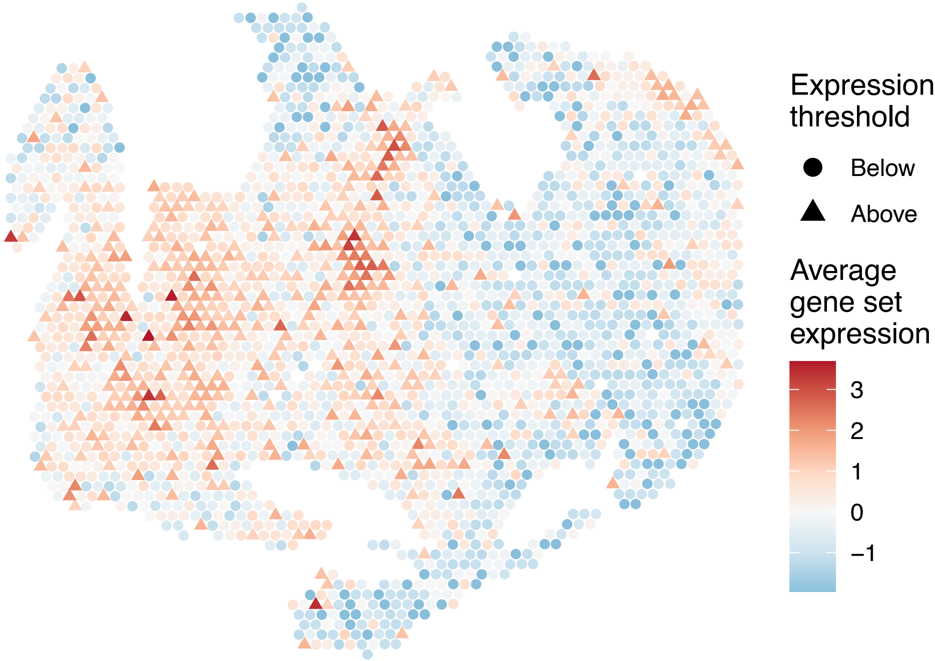


sample8_LMM_B3
KEGG_HEDGEHOG_SIGNALING_PATHWAY
Proportion of pathway genes=0.58; Adjusted p–value=0

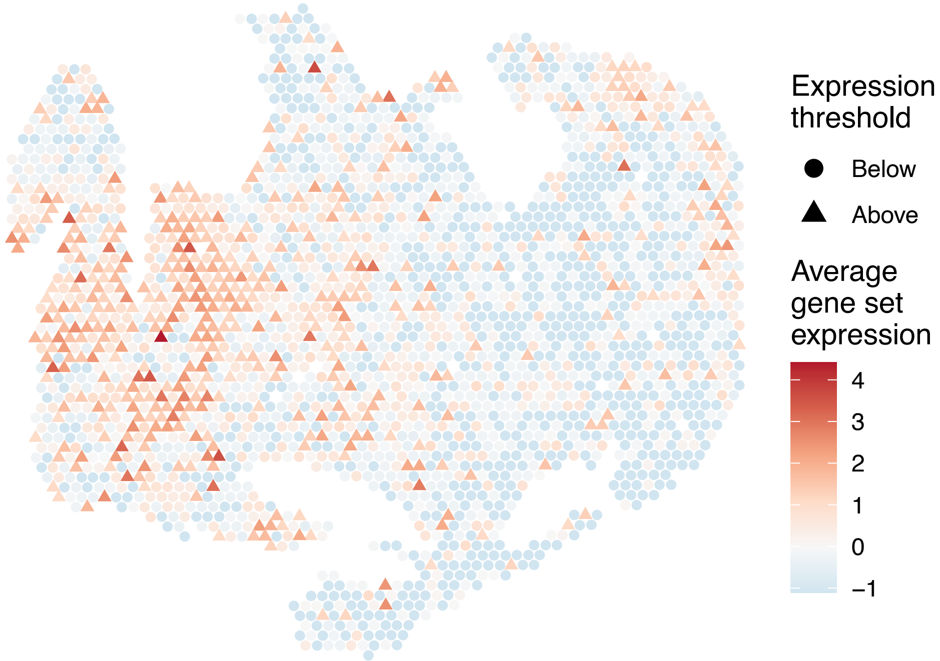


sample8_LMM_B3
KEGG_HEMATOPOIETIC_CELL_LINEAGE
Proportion of pathway genes=0.45; Adjusted p–value=0

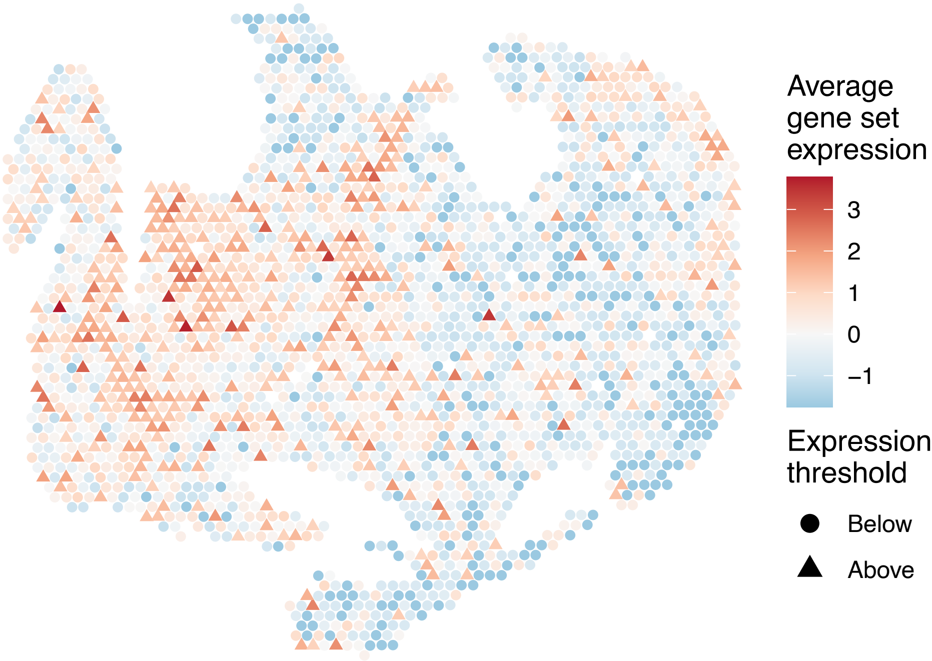

sample8_LMM_B3
KEGG_HISTIDINE_METABOLISM
Proportion of pathway genes=0.57; Adjusted p−value=0

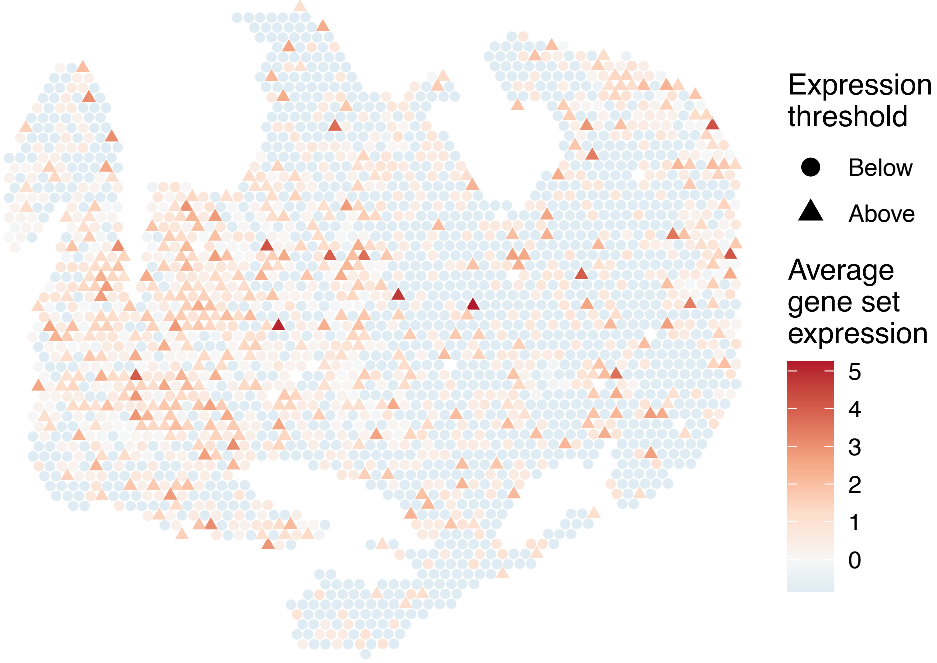


sample8_LMM_B3
KEGG_HOMOLOGOUS_RECOMBINATION
Proportion of pathway genes=0.97; Adjusted p−value=0.0262

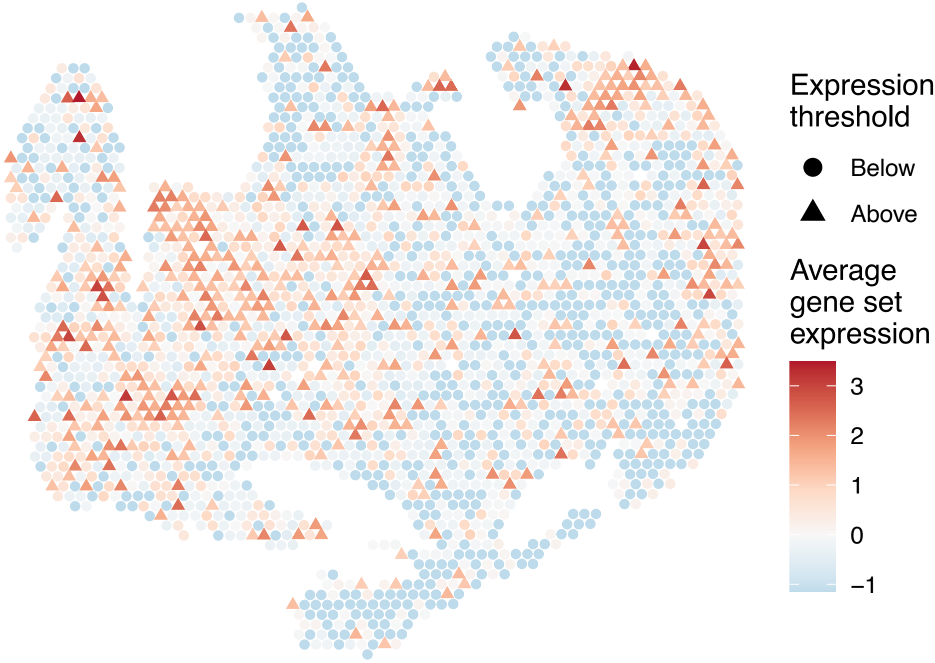


sample8_LMM_B3
KEGG_HUNTINGTONS_DISEASE
Proportion of pathway genes=0.89; Adjusted p−value=0

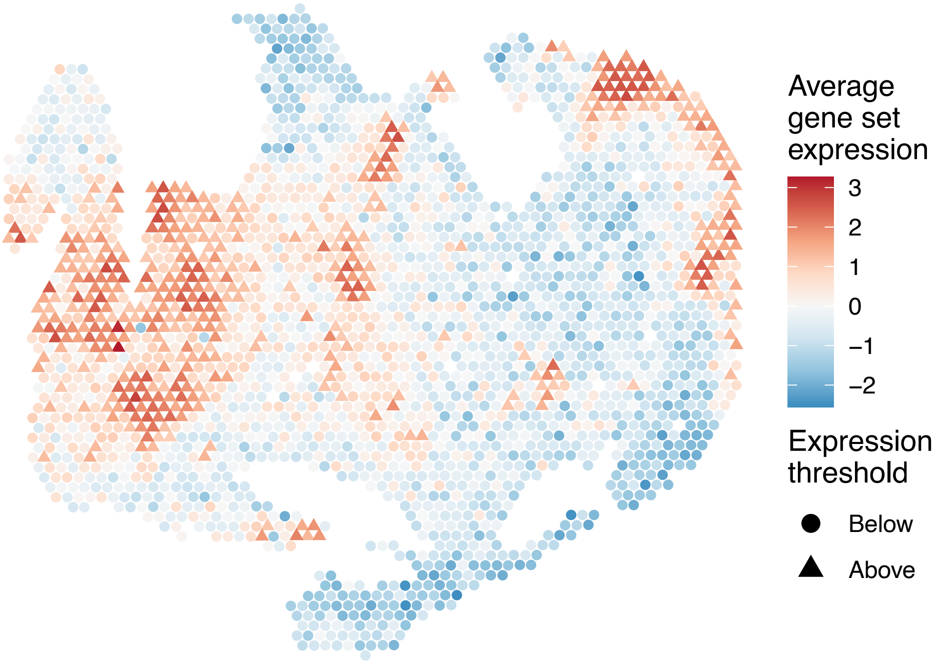


sample8_LMM_B3
KEGG_HYPERTROPHIC_CARDIOMYOPATHY_HCM
Proportion of pathway genes=0.61; Adjusted p−value=0

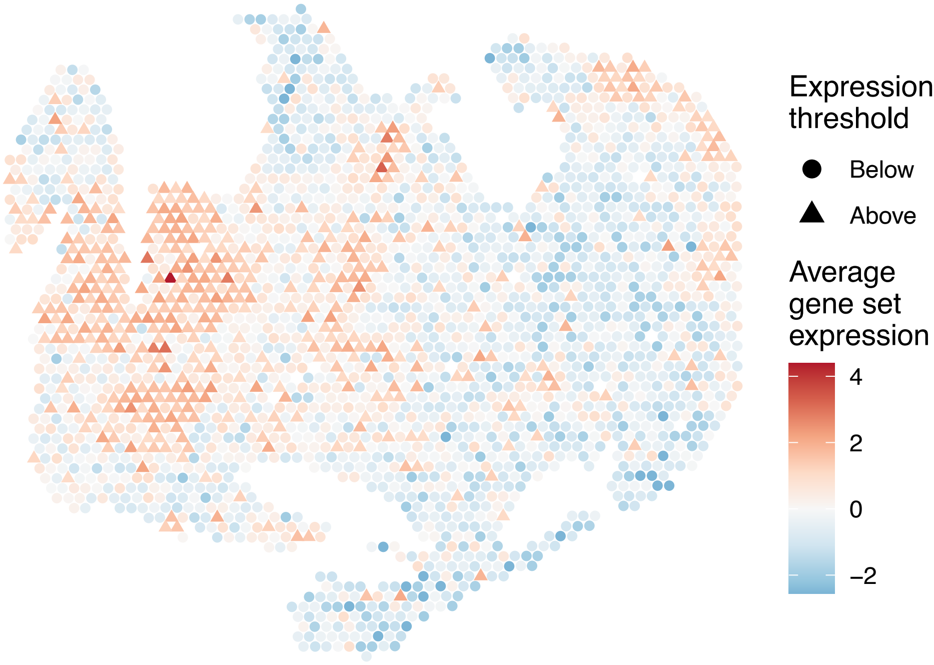

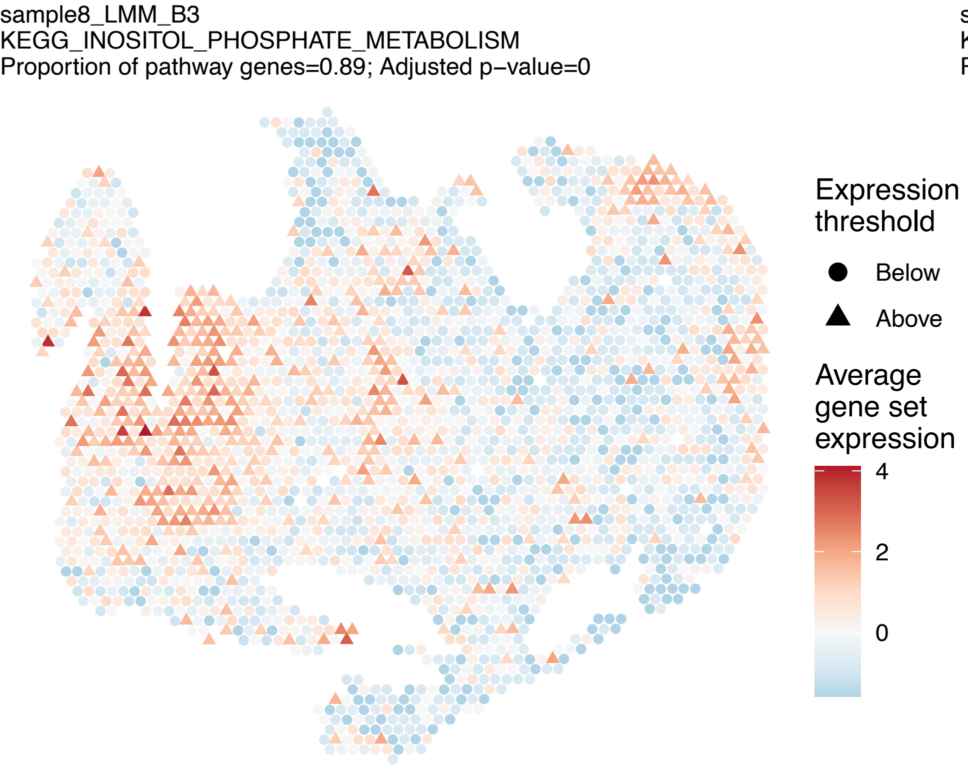
sample8_LMM_B3
KEGG_INOSITOL_PHOSPHATE_METABOLISM
Proportion of pathway genes=0.89; Adjusted p−value=0
Expression threshold
Below
Above
Average gene set expression
4
2
0

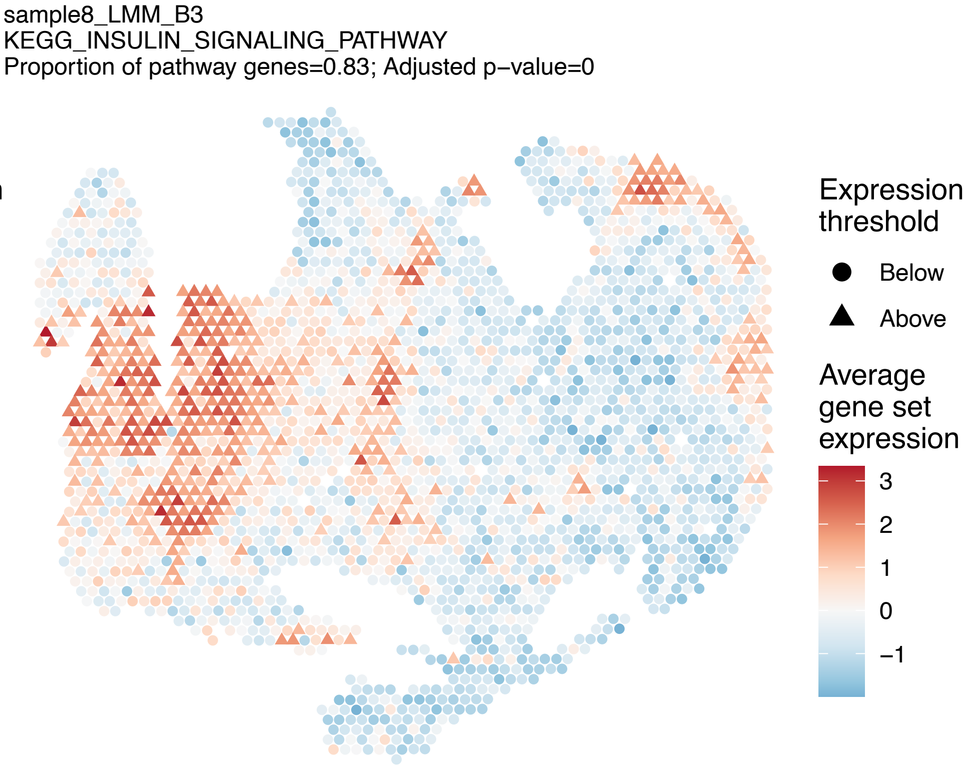
sample8_LMM_B3
KEGG_INSULIN_SIGNALING_PATHWAY
Proportion of pathway genes=0.83; Adjusted p−value=0
Expression threshold
Below
Above
Average gene set expression
3
2
1
0
−1

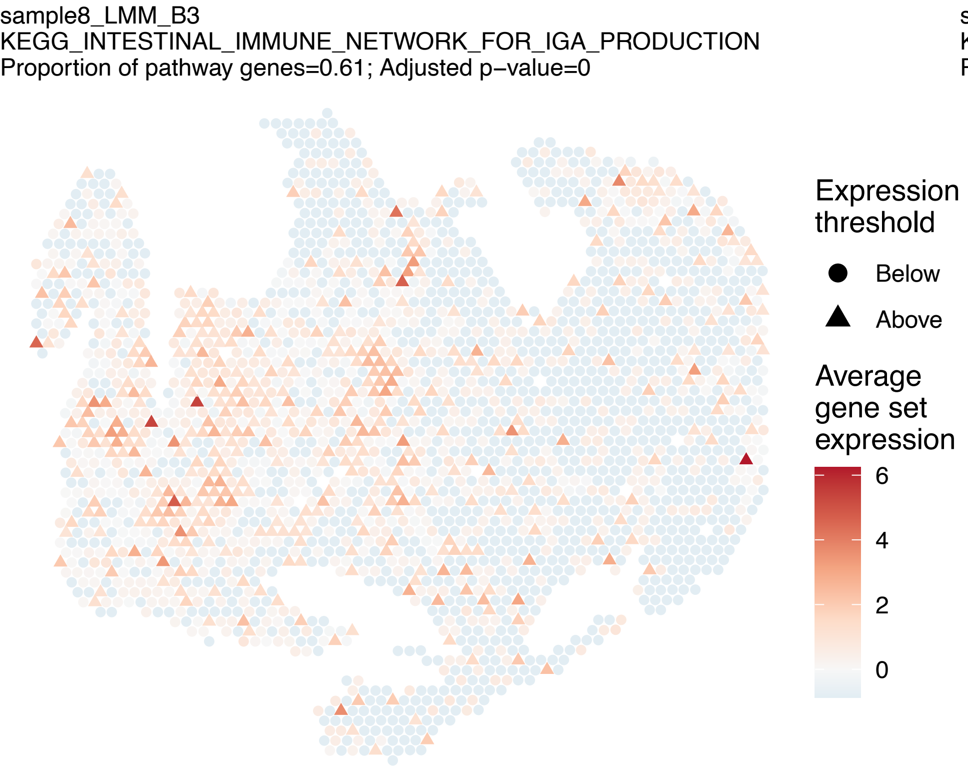
sample8_LMM_B3
KEGG_INTESTINAL_IMMUNE_NETWORK_FOR_IGA_PRODUCTION
Proportion of pathway genes=0.61; Adjusted p−value=0
Expression threshold
Below
Above
Average gene set expression
6
4
2
0

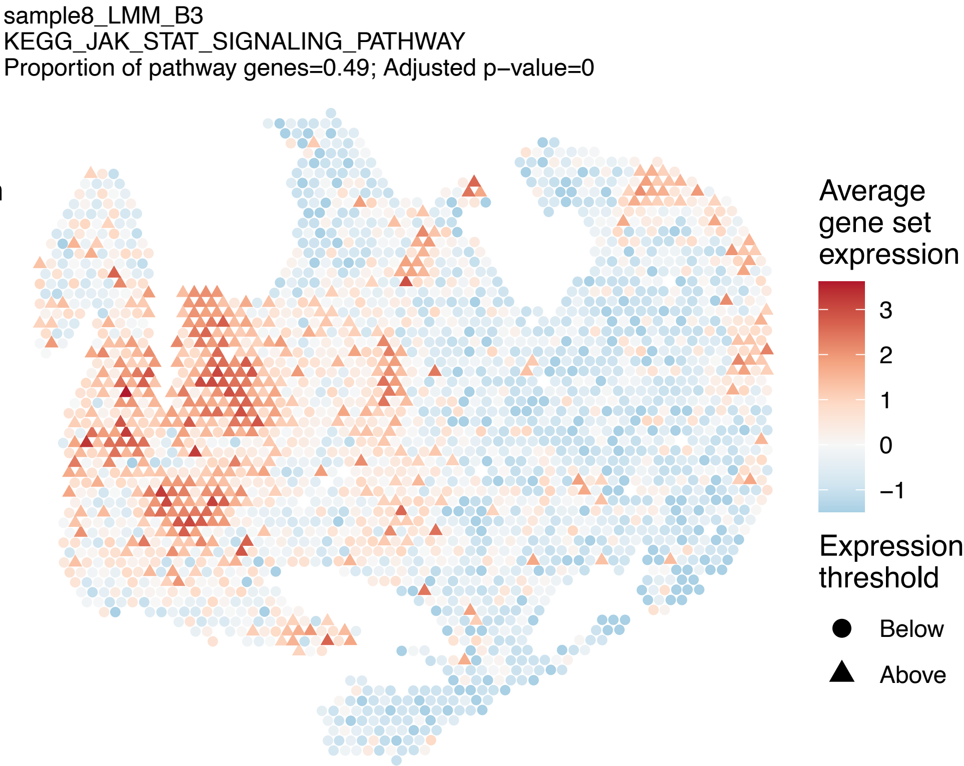
sample8_LMM_B3
KEGG_JAK_STAT_SIGNALING_PATHWAY
Proportion of pathway genes=0.49; Adjusted p−value=0
Average gene set expression
3
2
1
0
−1
Expression threshold
Below
Above

sample8_LMM_B3
KEGG_LEISHMANIA_INFECTION
Proportion of pathway genes=0.66; Adjusted p−value=0

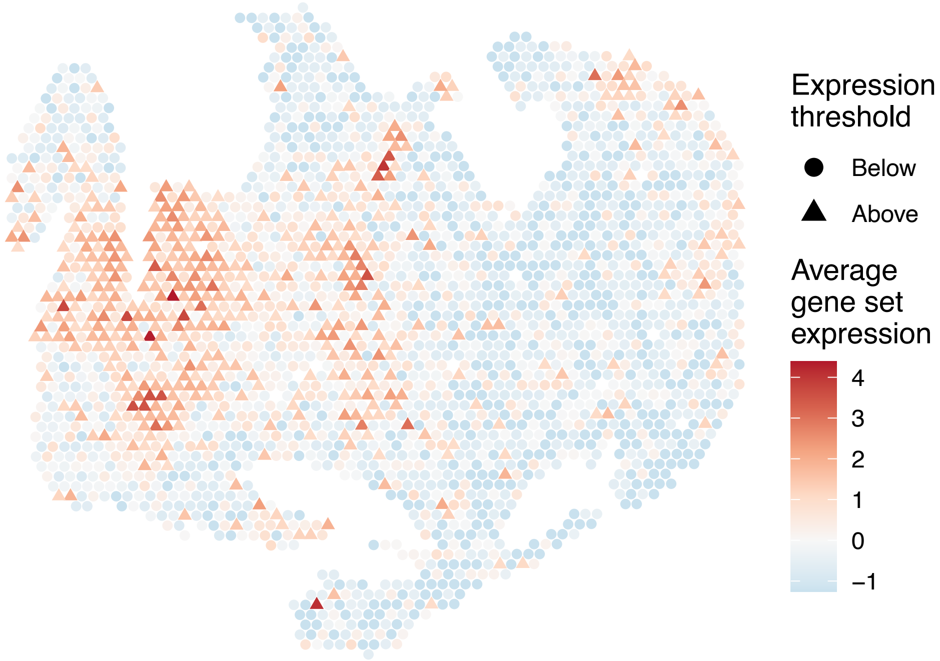


sample8_LMM_B3
KEGG_LEUKOCYTE_TRANSENDOTHELIAL_MIGRATION
Proportion of pathway genes=0.7; Adjusted p−value=0

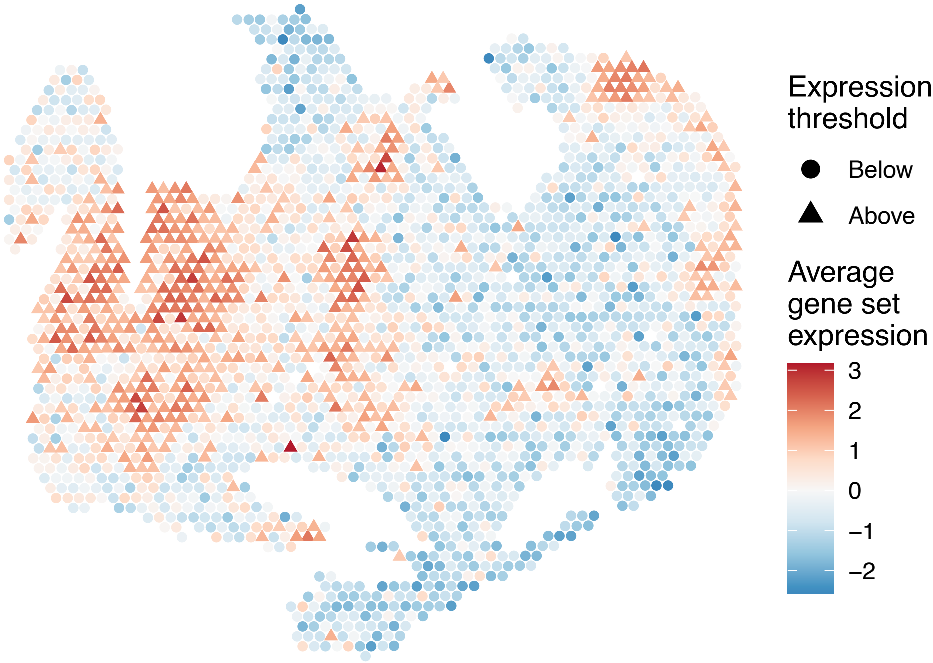


sample8_LMM_B3
KEGG_LIMONENE_AND_PINENE_DEGRADATION
Proportion of pathway genes=1; Adjusted p−value=0.7371

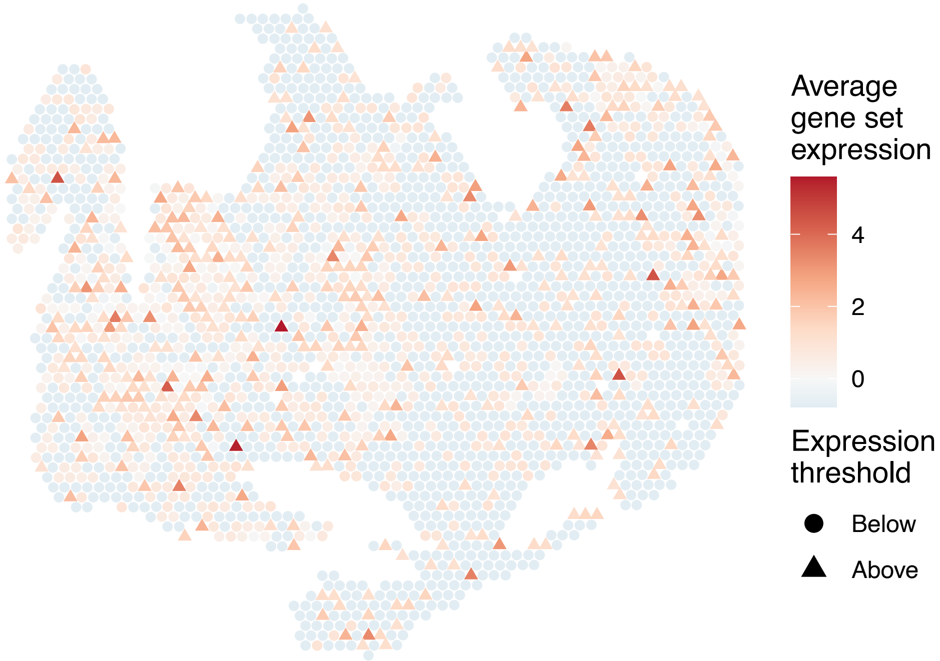


sample8_LMM_B3
KEGG_LONG_TERM_DEPRESSION
Proportion of pathway genes=0.69; Adjusted p−value=0

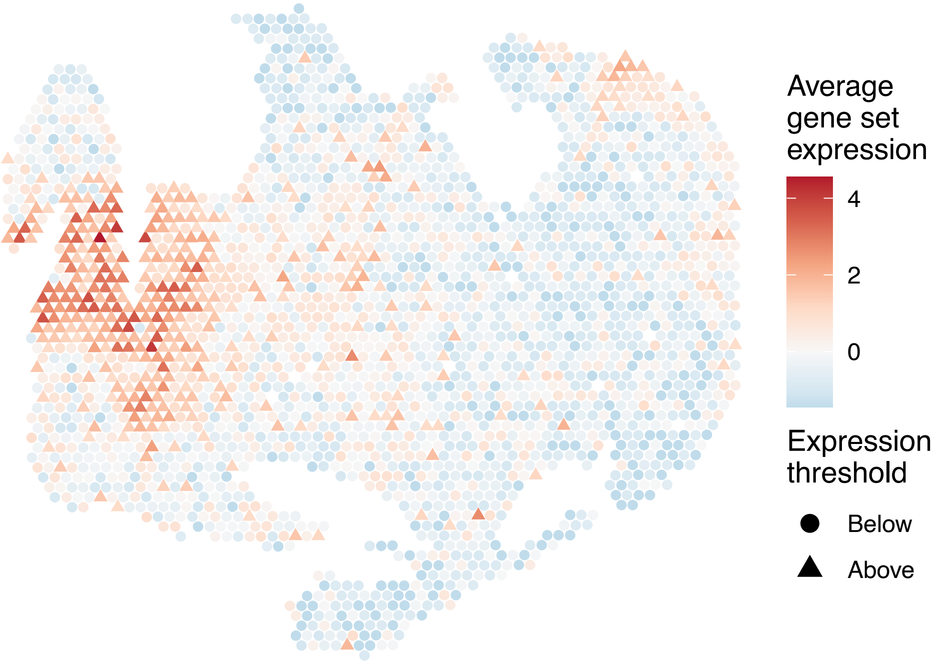

sample8_LMM_B3
KEGG_LONG_TERM_POTENTIATION
Proportion of pathway genes=0.89; Adjusted p−value=0

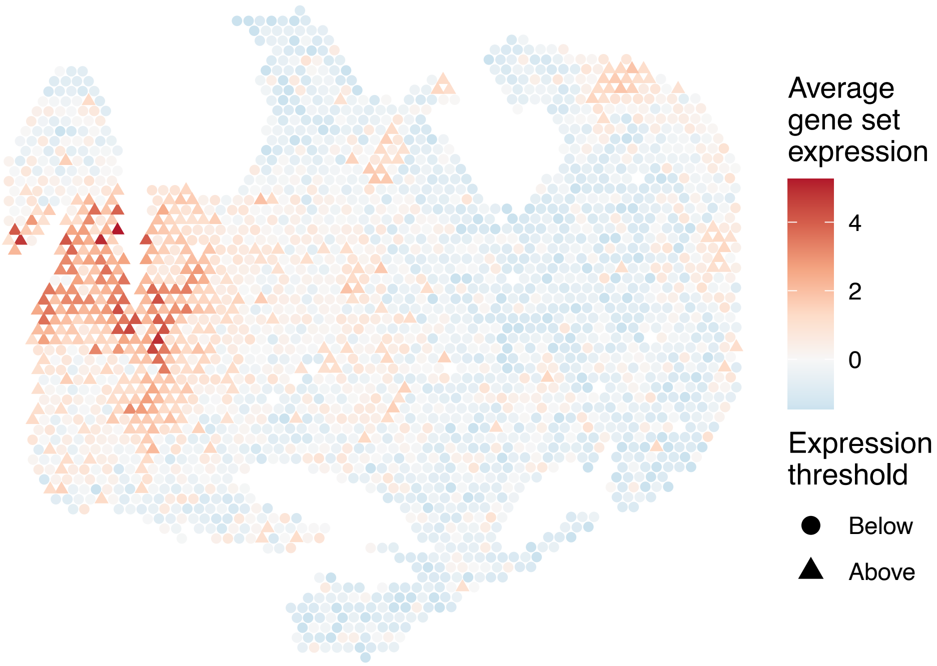


sample8_LMM_B3
KEGG_LYSINE_DEGRADATION
Proportion of pathway genes=0.96; Adjusted p−value=0

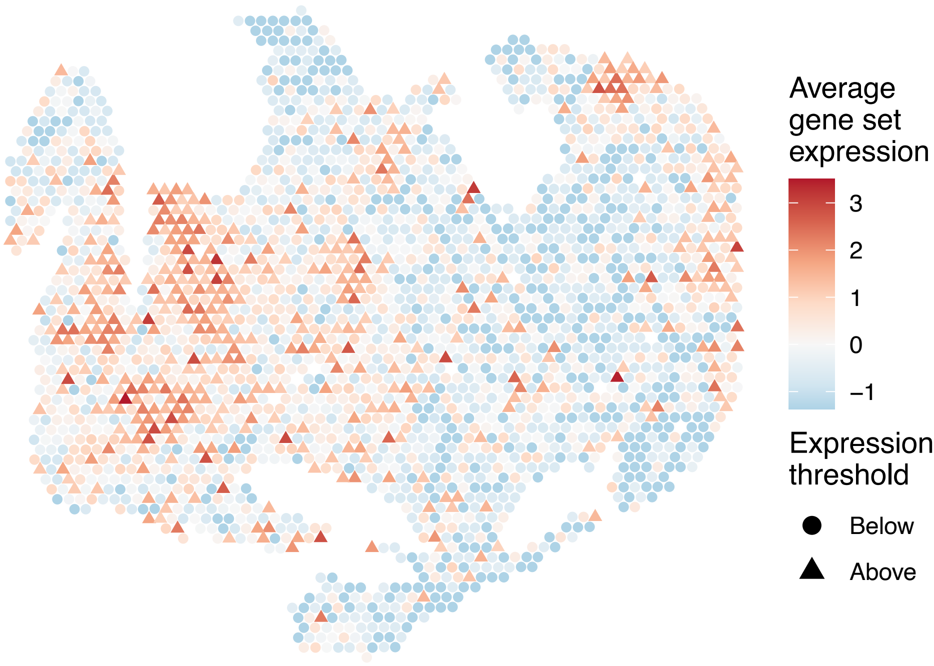


sample8_LMM_B3
KEGG_LYSOSOME
Proportion of pathway genes=0.93; Adjusted p−value=0

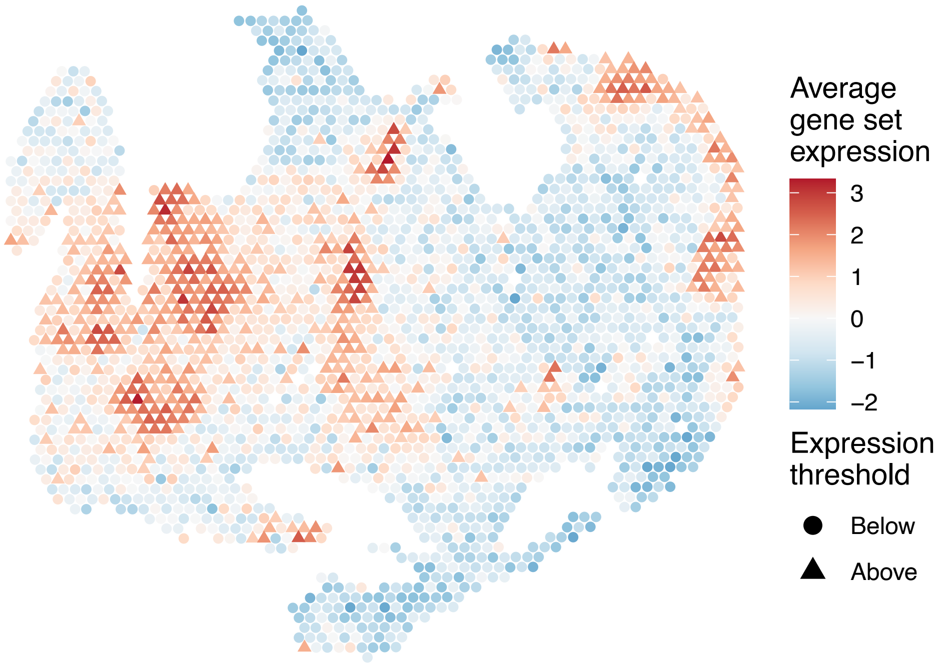


sample8_LMM_B3
KEGG_MAPK_SIGNALING_PATHWAY
Proportion of pathway genes=0.79; Adjusted p−value=0

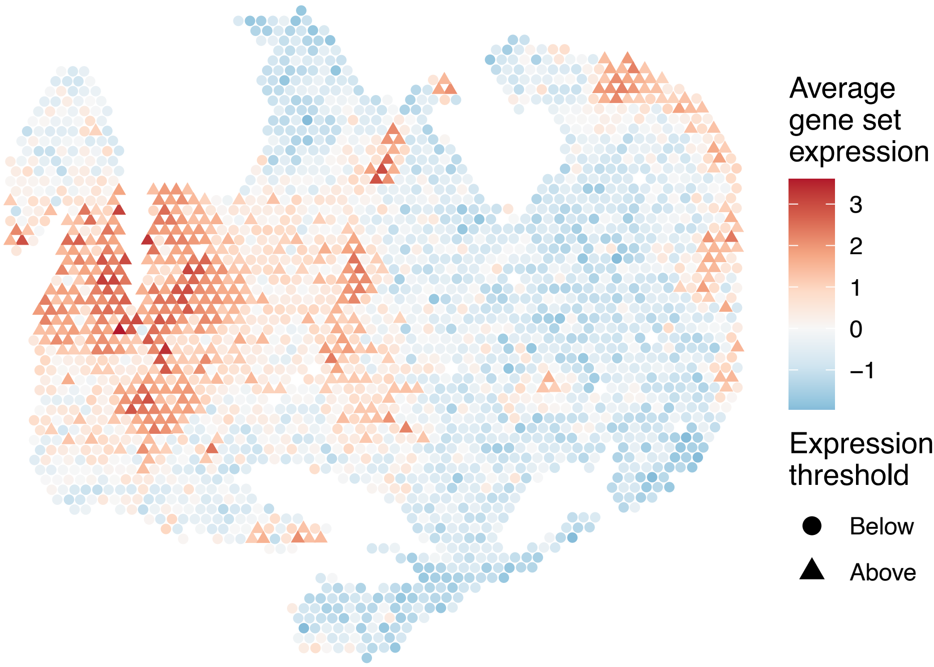

sample8_LMM_B3
KEGG_MELANOGENESIS
Proportion of pathway genes=0.75; Adjusted p–value=0

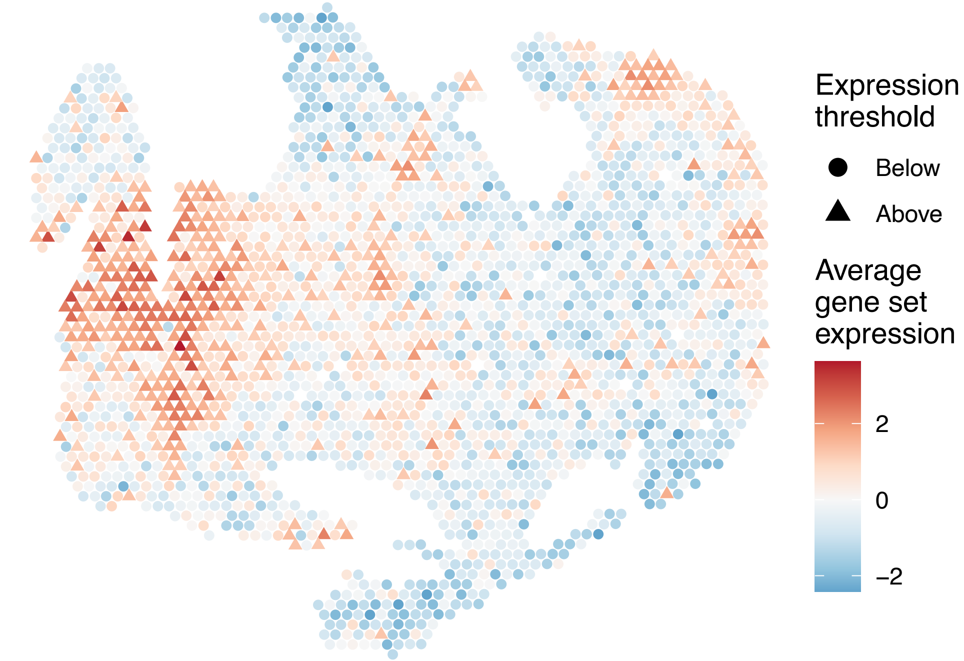


sample8_LMM_B3
KEGG_MELANOMA
Proportion of pathway genes=0.73; Adjusted p–value=0

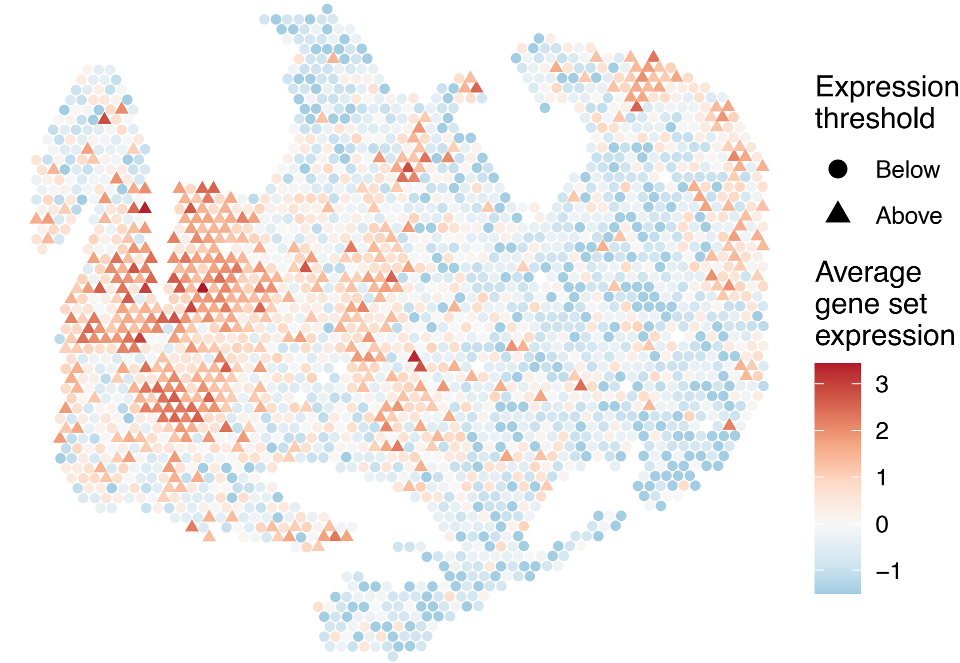


sample8_LMM_B3
KEGG_METABOLISM_OF_XENOBIOTICS_BY_CYTOCHROME_P450
Proportion of pathway genes=0.24; Adjusted p–value=0.0177

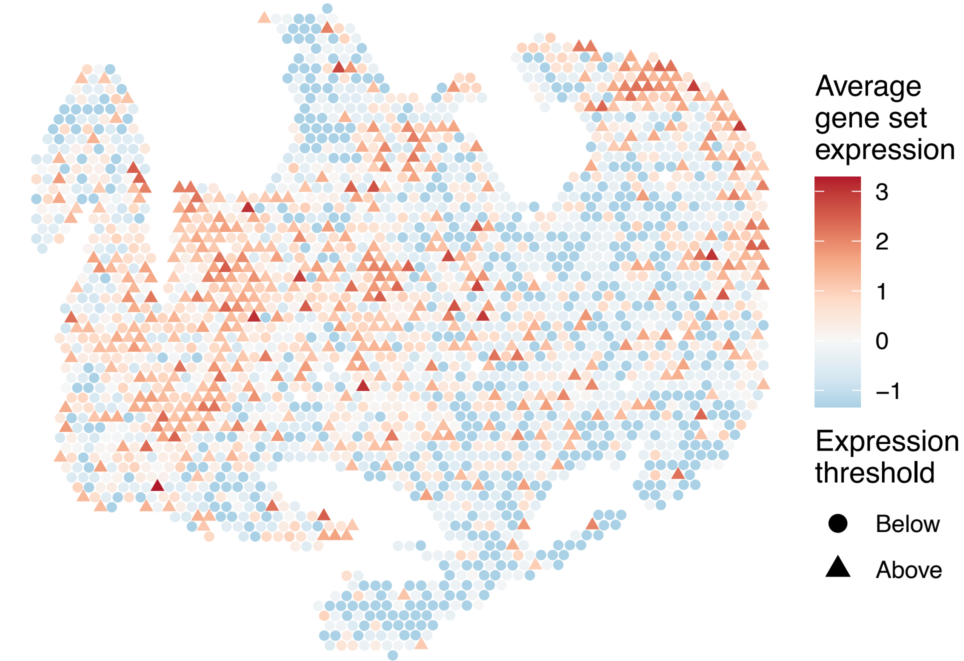


sample8_LMM_B3
KEGG_MISMATCH_REPAIR
Proportion of pathway genes=0.96; Adjusted p–value=0.0633

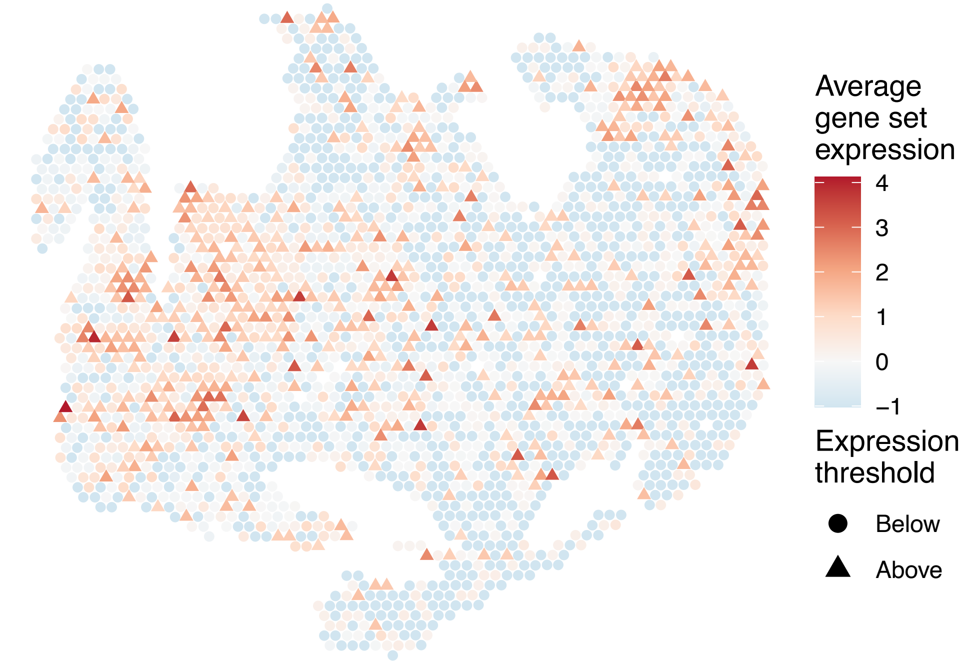

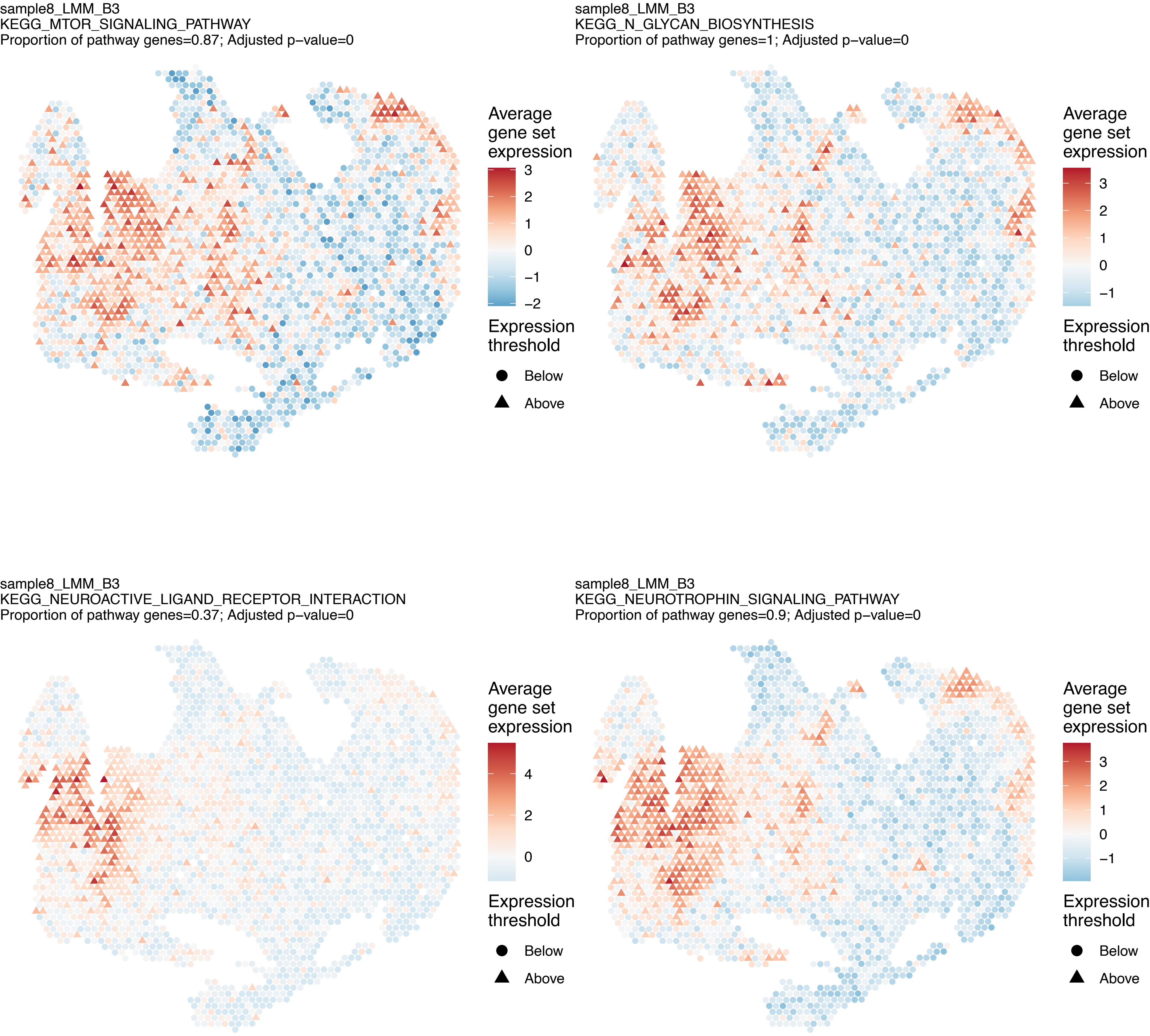

sample8_LMM_B3
KEGG_MTOR_SIGNALING_PATHWAY
Proportion of pathway genes=0.87; Adjusted p−value=0
Average gene set expression
3
2
1
0
−1
−2
Expression threshold
Below
Above
sample8_LMM_B3
KEGG_N_GLYCAN_BIOSYNTHESIS
Proportion of pathway genes=1; Adjusted p−value=0
Average gene set expression
3
2
1
0
−1
Expression threshold
Below
Above
sample8_LMM_B3
KEGG_NEUROACTIVE_LIGAND_RECEPTOR_INTERACTION
Proportion of pathway genes=0.37; Adjusted p−value=0
Average gene set expression
4
2
0
Expression threshold
Below
Above
sample8_LMM_B3
KEGG_NEUROTROPHIN_SIGNALING_PATHWAY
Proportion of pathway genes=0.9; Adjusted p−value=0
Average gene set expression
3
2
1
0
−1
Expression threshold
Below
Above

sample8_LMM_B3
KEGG_NICOTINATE_AND_NICOTINAMIDE_METABOLISM
Proportion of pathway genes=0.71; Adjusted p−value=0

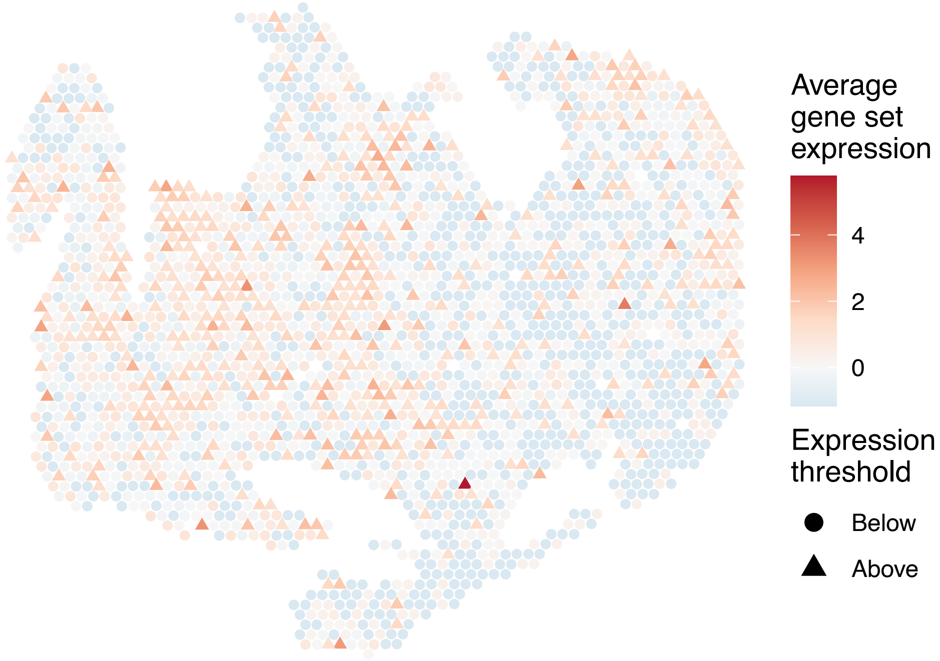


sample8_LMM_B3
KEGG_NITROGEN_METABOLISM
Proportion of pathway genes=0.58; Adjusted p−value=0.1636

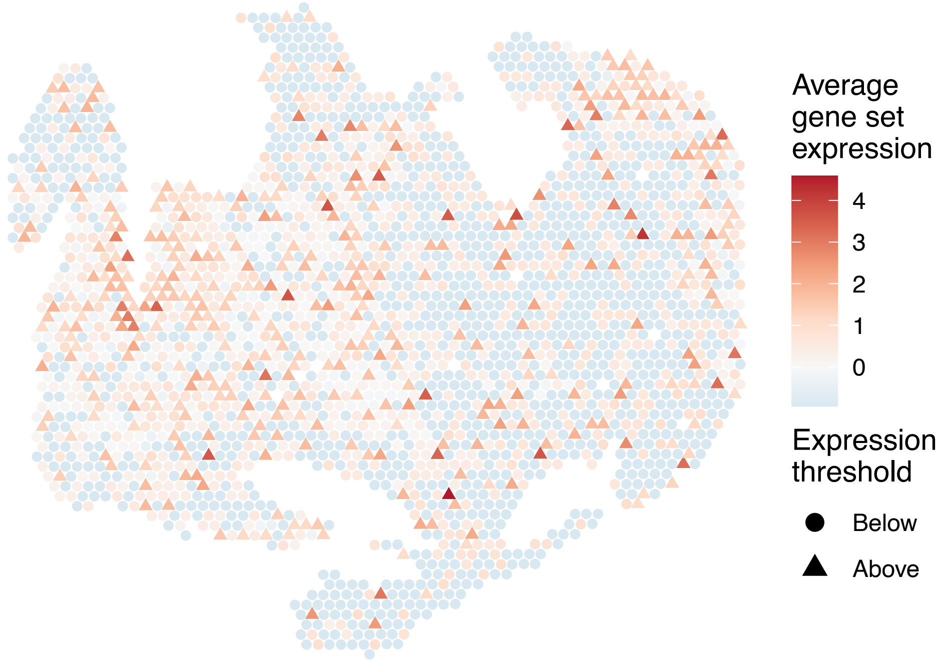


sample8_LMM_B3
KEGG_NOD_LIKE_RECEPTOR_SIGNALING_PATHWAY
Proportion of pathway genes=0.66; Adjusted p−value=0

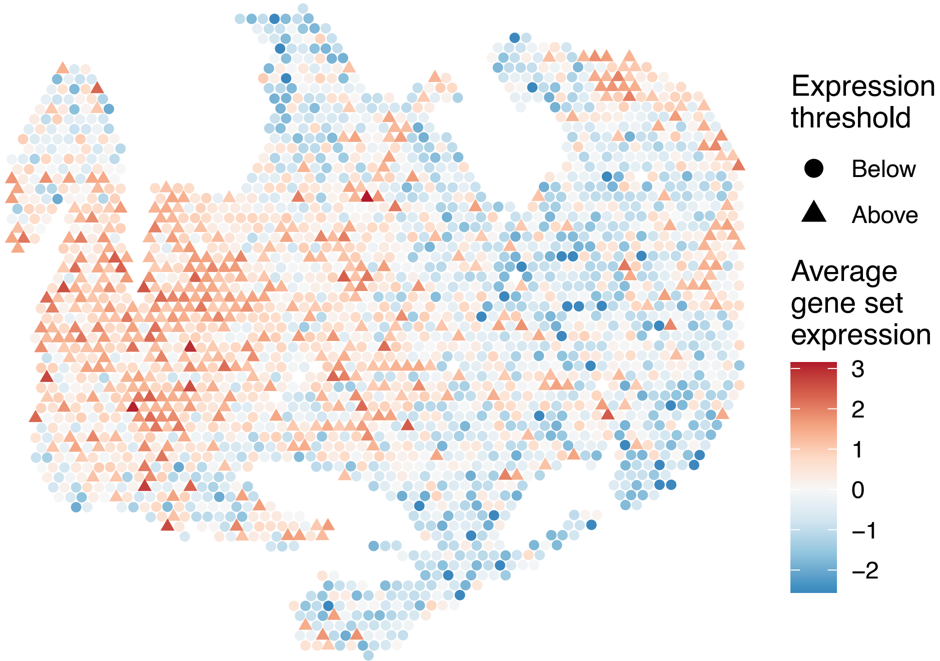


sample8_LMM_B3
KEGG_NON_HOMOLOGOUS_END_JOINING
Proportion of pathway genes=0.92; Adjusted p−value=0

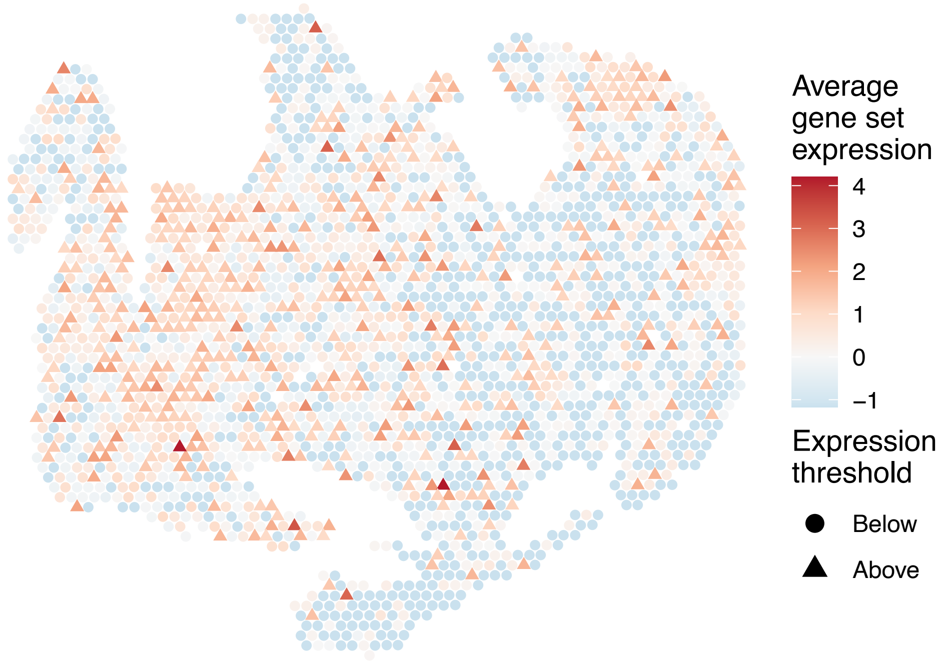

sample8_LMM_B3
KEGG_NON_SMALL_CELL_LUNG_CANCER
Proportion of pathway genes=0.94; Adjusted p−value=0

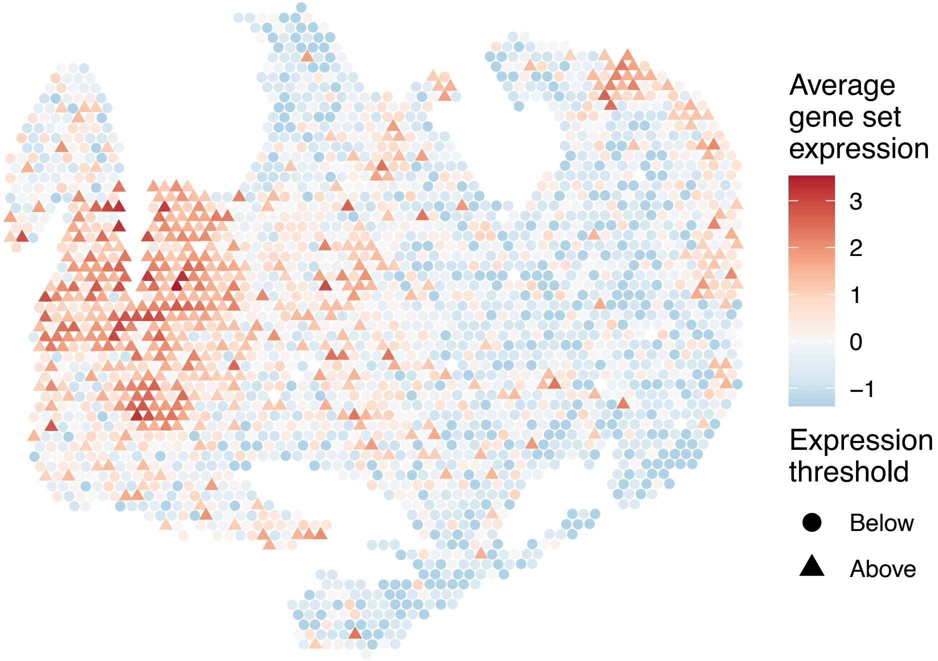


sample8_LMM_B3
KEGG_NOTCH_SIGNALING_PATHWAY
Proportion of pathway genes=0.88; Adjusted p−value=0

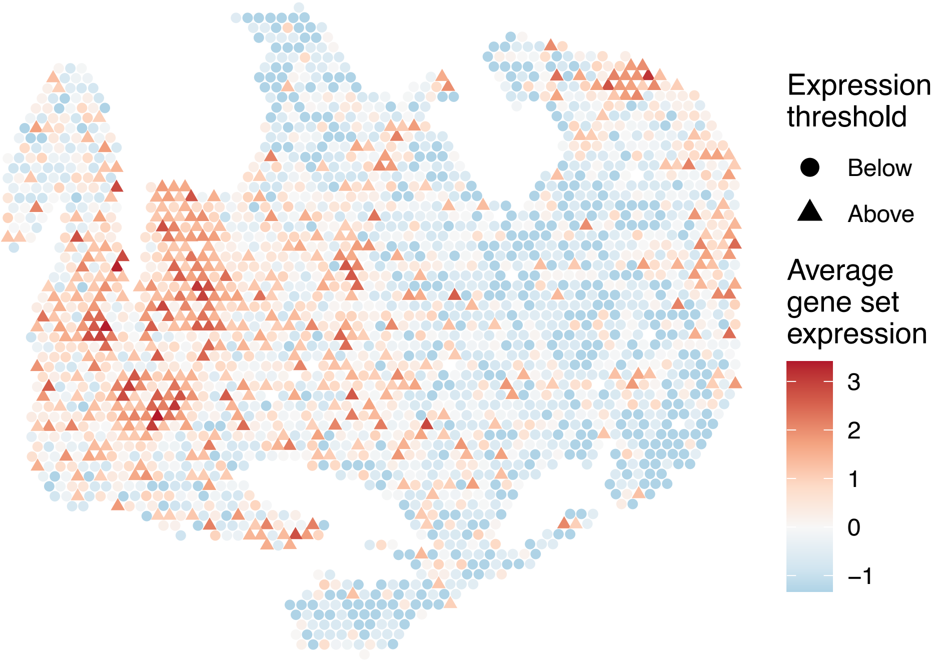


sample8_LMM_B3
KEGG_NUCLEOTIDE_EXCISION_REPAIR
Proportion of pathway genes=0.98; Adjusted p−value=0

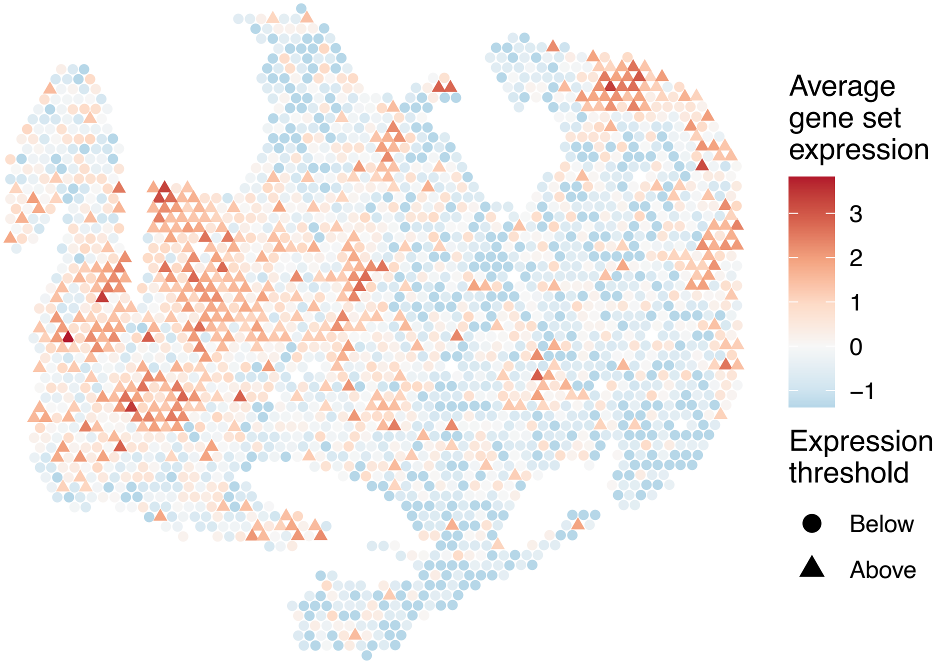


sample8_LMM_B3
KEGG_O_GLYCAN_BIOSYNTHESIS
Proportion of pathway genes=0.73; Adjusted p−value=0

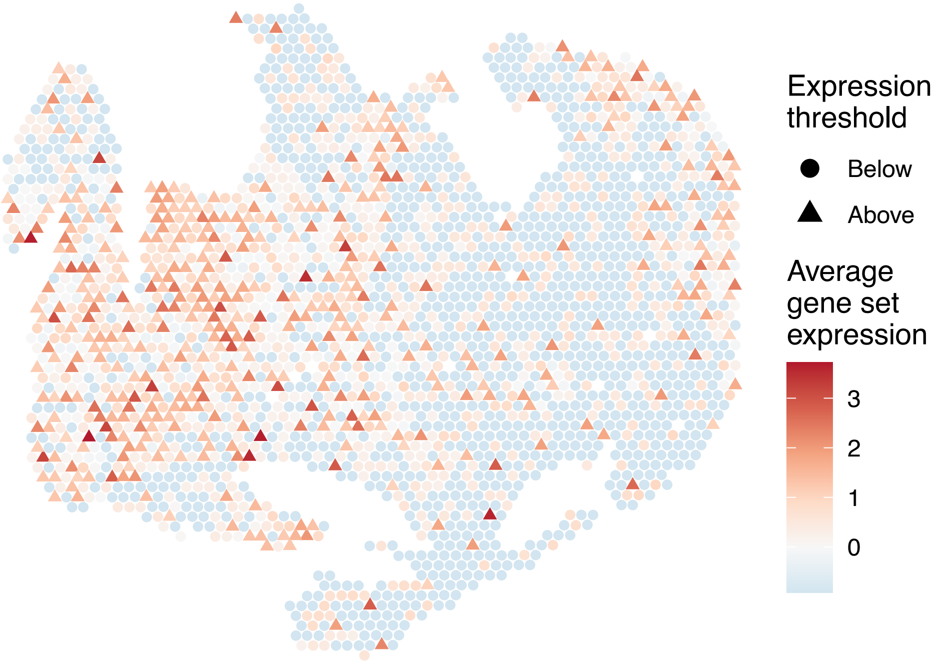

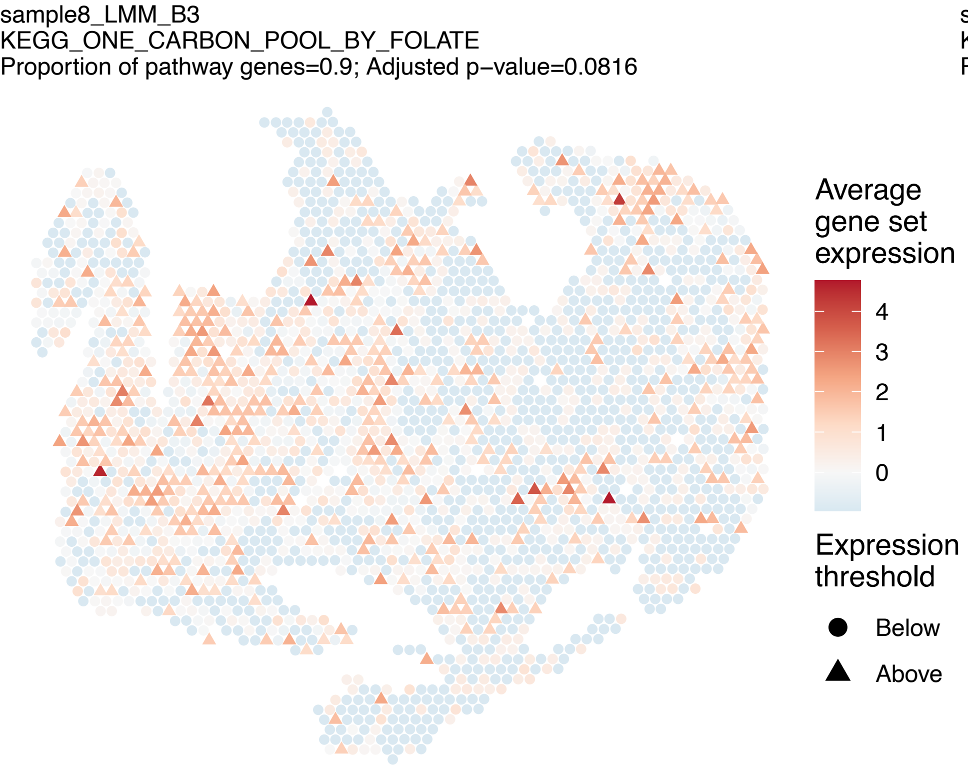
sample8_LMM_B3
KEGG_ONE_CARBON_POOL_BY_FOLATE
Proportion of pathway genes=0.9; Adjusted p−value=0.0816
Average gene set expression
4
3
2
1
0
Expression threshold
Below
Above

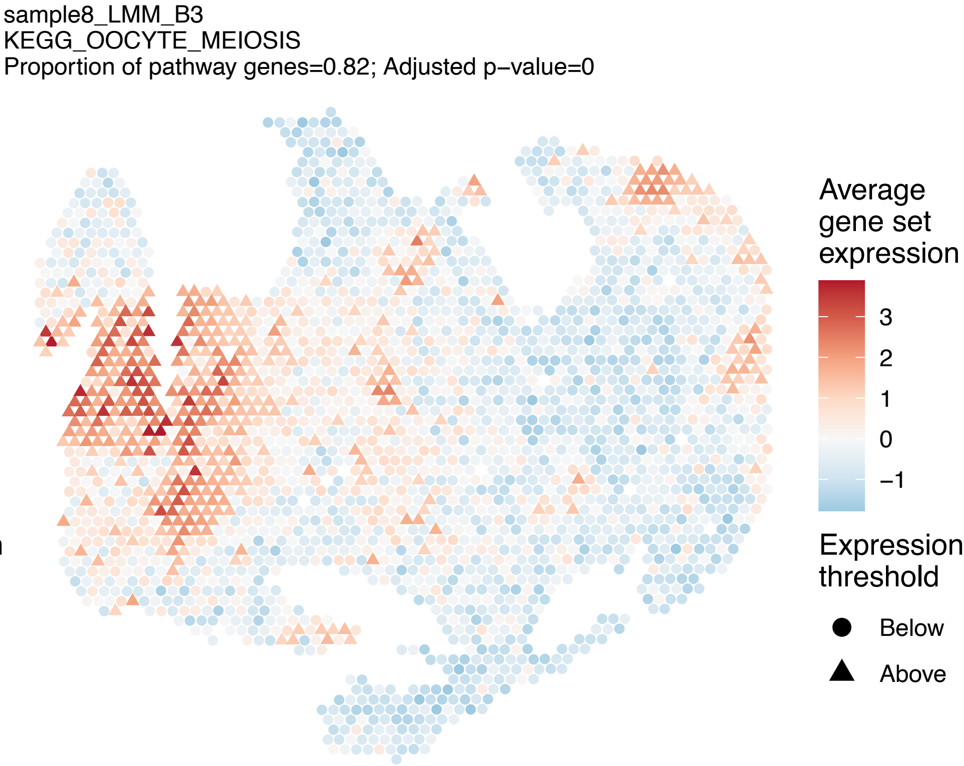
sample8_LMM_B3
KEGG_OOCYTE_MEIOSIS
Proportion of pathway genes=0.82; Adjusted p−value=0
Average gene set expression
3
2
1
0
−1
Expression threshold
Below
Above

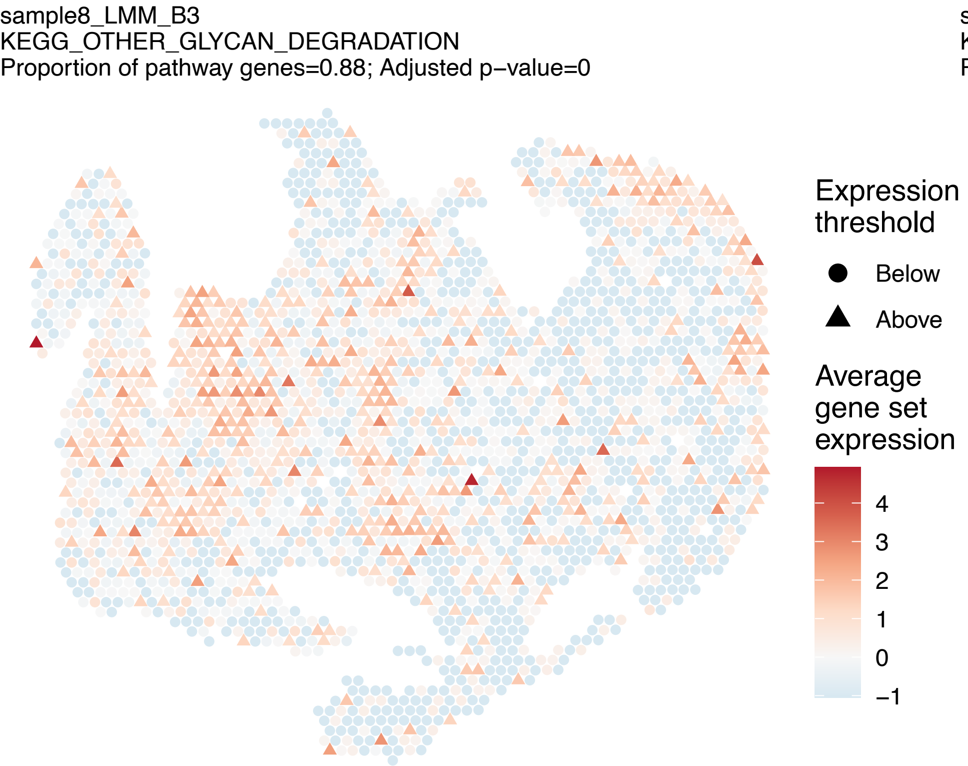
sample8_LMM_B3
KEGG_OTHER_GLYCAN_DEGRADATION
Proportion of pathway genes=0.88; Adjusted p−value=0
Expression threshold
Below
Above
Average gene set expression
4
3
2
1
0
−1

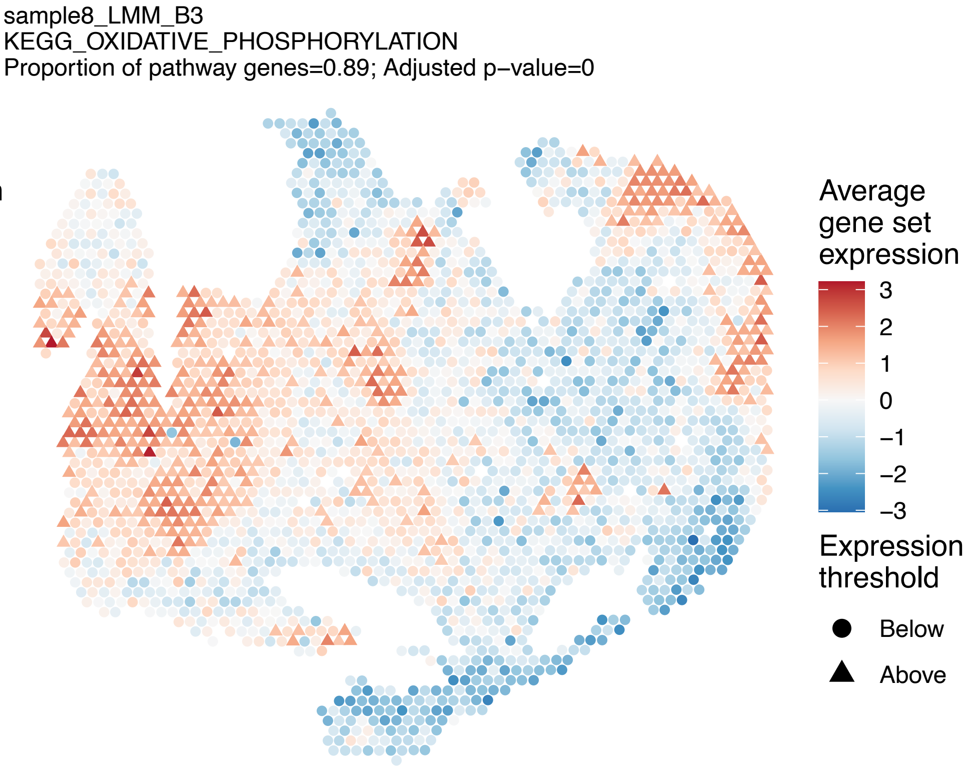
sample8_LMM_B3
KEGG_OXIDATIVE_PHOSPHORYLATION
Proportion of pathway genes=0.89; Adjusted p−value=0
Average gene set expression
3
2
1
0
−1
−2
−3
Expression threshold
Below
Above

sample8_LMM_B3
KEGG_P53_SIGNALING_PATHWAY
Proportion of pathway genes=0.89; Adjusted p–value=0

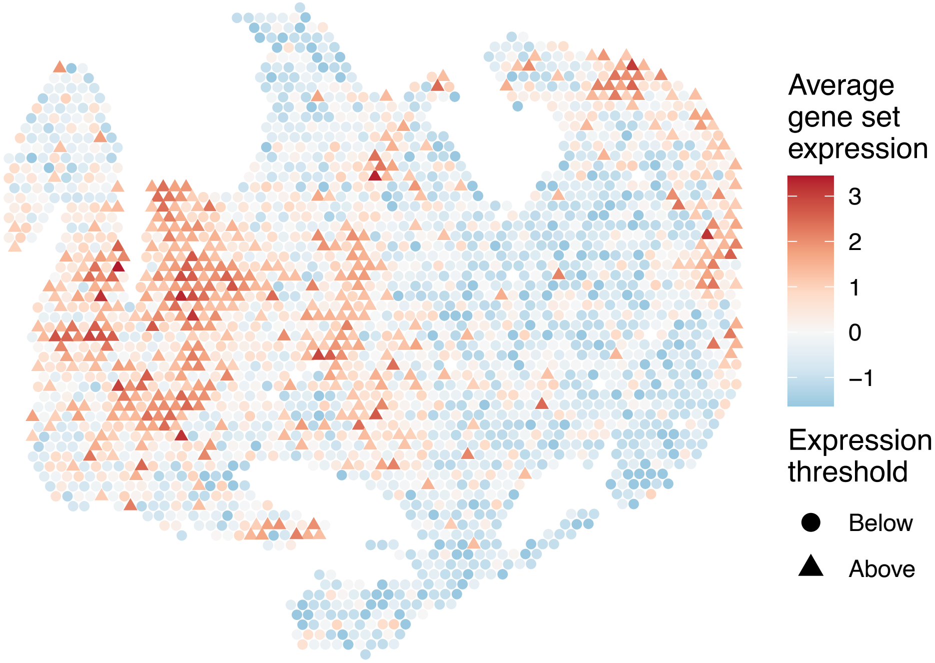


sample8_LMM_B3
KEGG_PANCREATIC_CANCER
Proportion of pathway genes=0.92; Adjusted p–value=0

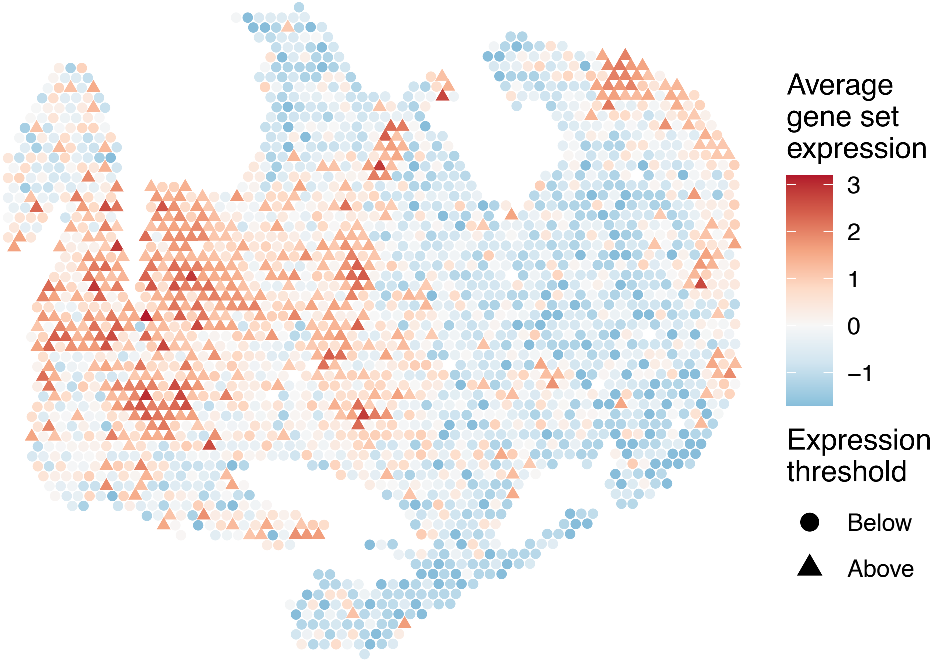


sample8_LMM_B3
KEGG_PANTOTHENATE_AND_COA_BIOSYNTHESIS
Proportion of pathway genes=0.71; Adjusted p–value=0.1532

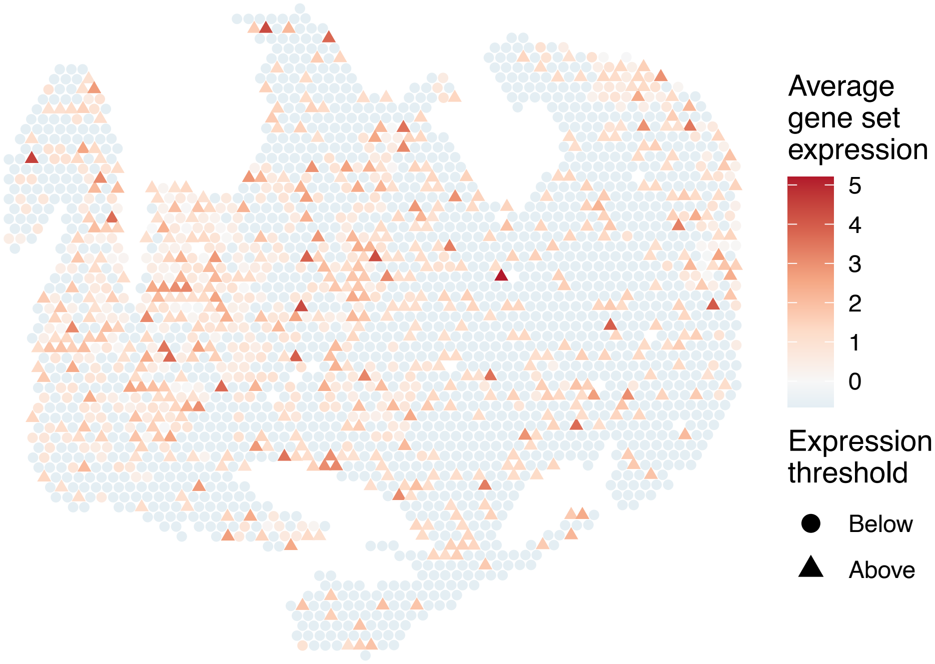


sample8_LMM_B3
KEGG_PARKINSONS_DISEASE
Proportion of pathway genes=0.89; Adjusted p–value=0

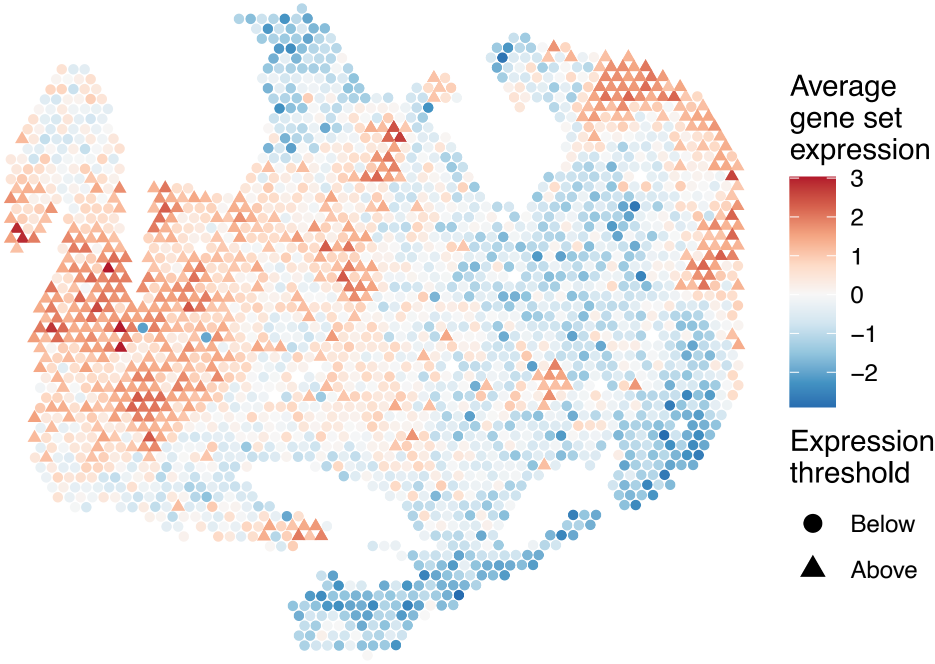

sample8_LMM_B3
KEGG_PATHOGENIC_ESCHERICHIA_COLI_INFECTION
Proportion of pathway genes=0.8; Adjusted p−value=0

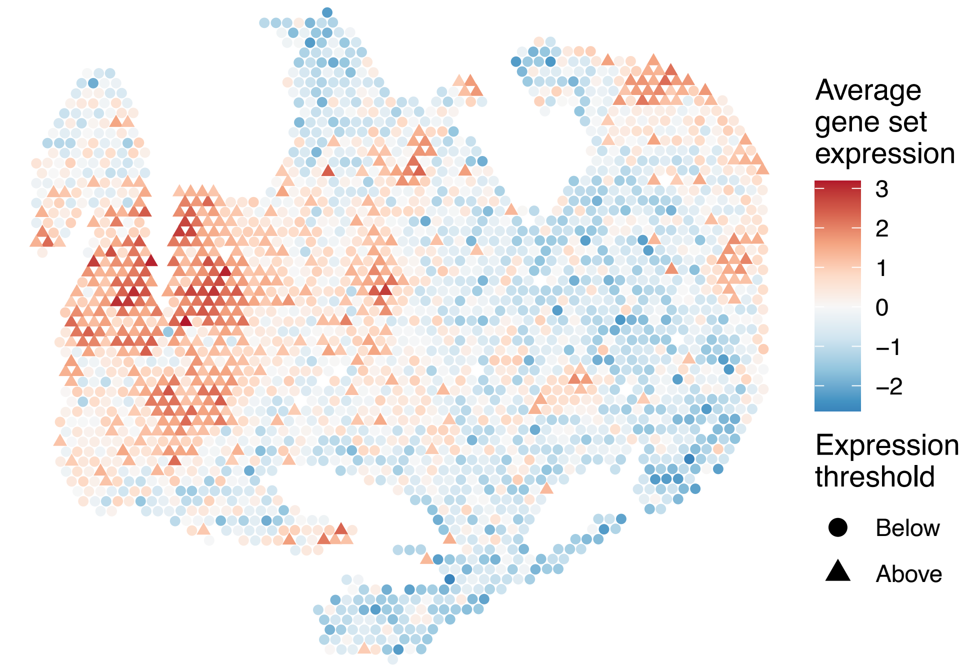


sample8_LMM_B3
KEGG_PATHWAYS_IN_CANCER
Proportion of pathway genes=0.8; Adjusted p−value=0

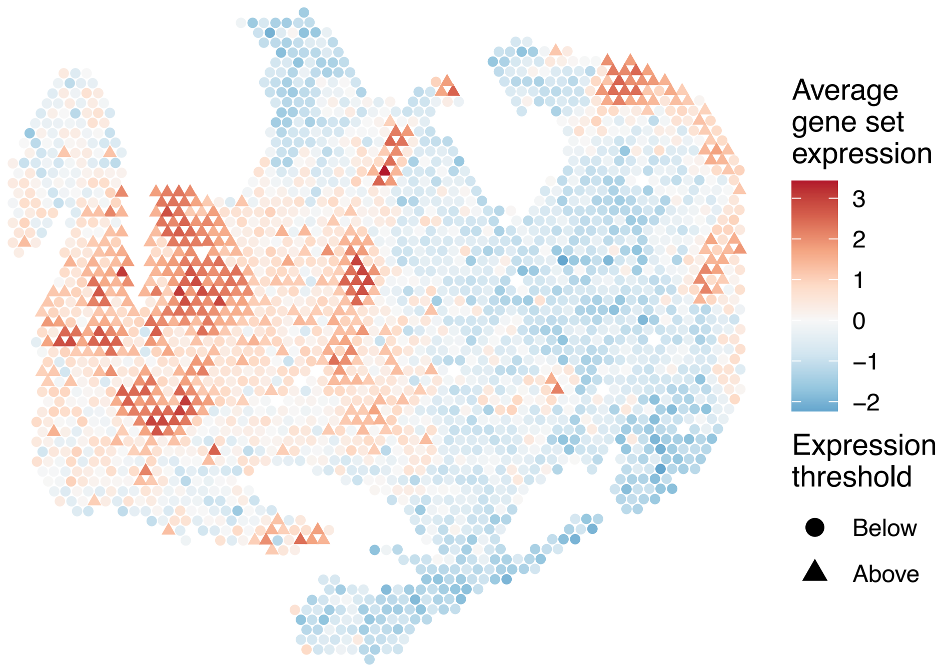


sample8_LMM_B3
KEGG_PENTOSE_PHOSPHATE_PATHWAY
Proportion of pathway genes=0.75; Adjusted p−value=0.0012

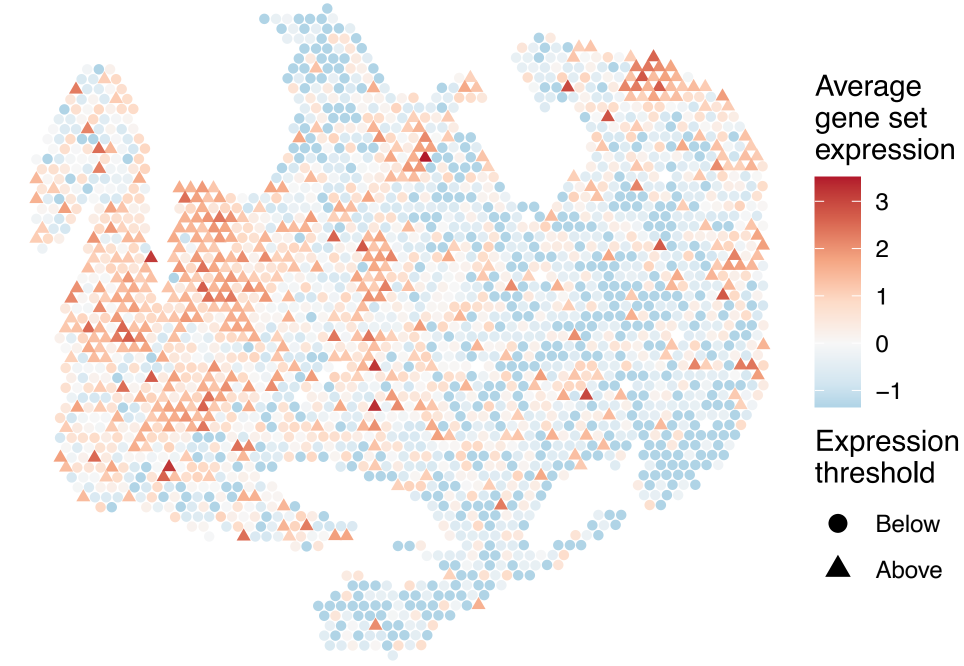


sample8_LMM_B3
KEGG_PEROXISOME
Proportion of pathway genes=0.88; Adjusted p−value=0

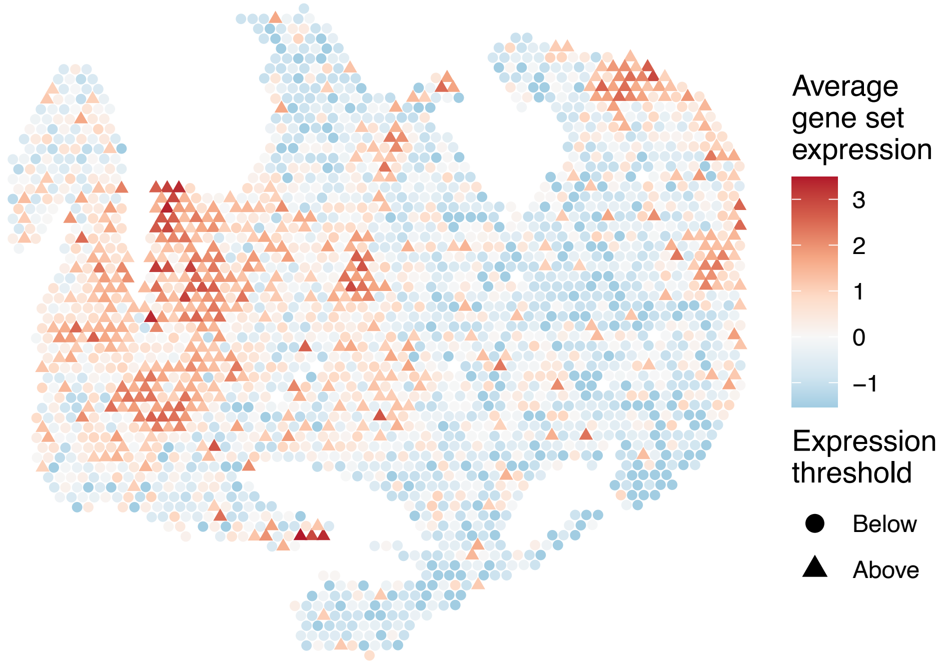

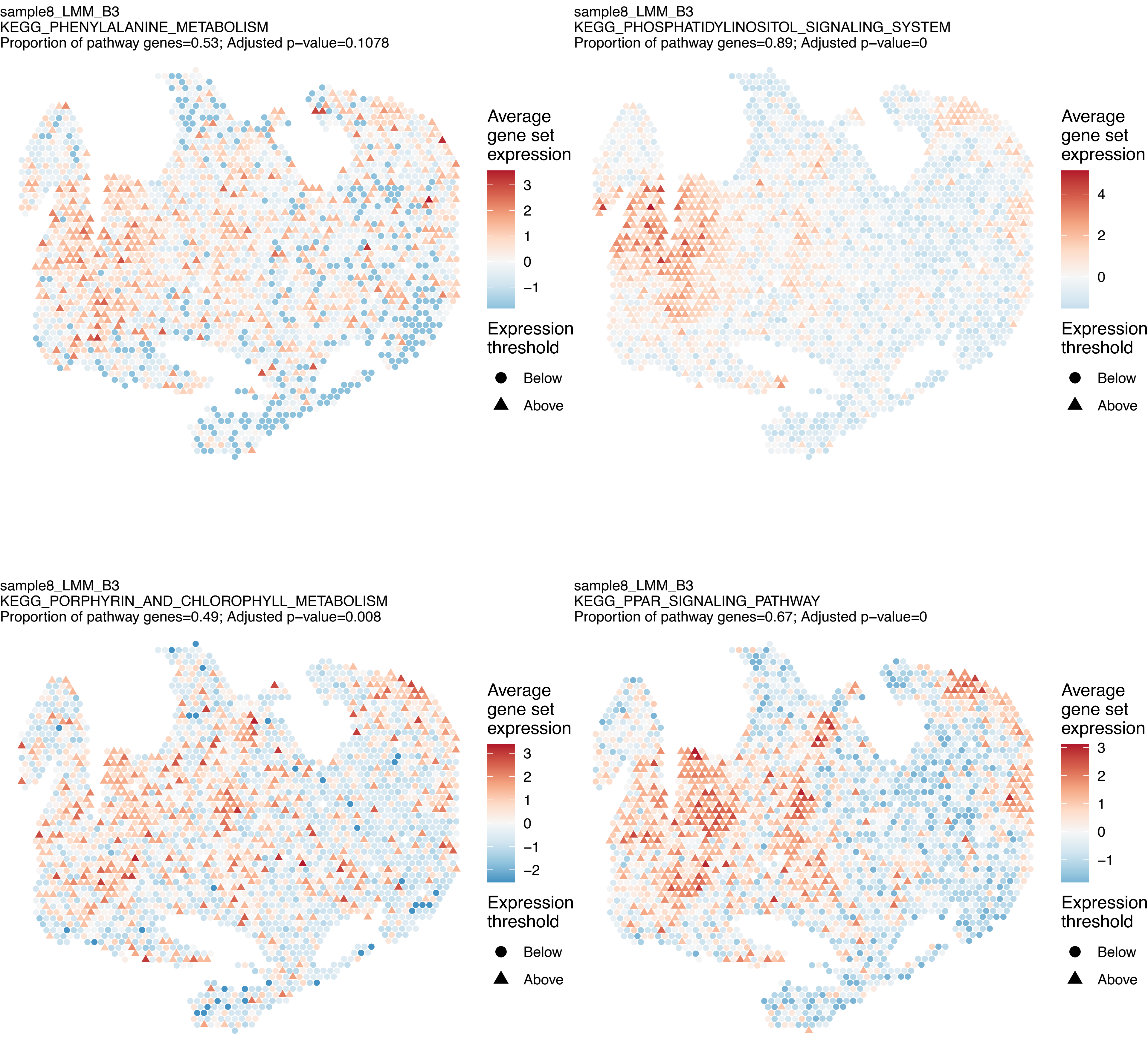
sample8_LMM_B3
KEGG_PHENYLALANINE_METABOLISM
Proportion of pathway genes=0.53; Adjusted p−value=0.1078
Average gene set expression
3
2
1
0
−1
Expression threshold
Below
Above
sample8_LMM_B3
KEGG_PHOSPHATIDYLINOSITOL_SIGNALING_SYSTEM
Proportion of pathway genes=0.89; Adjusted p−value=0
Average gene set expression
4
2
0
Expression threshold
Below
Above
sample8_LMM_B3
KEGG_PORPHYRIN_AND_CHLOROPHYLL_METABOLISM
Proportion of pathway genes=0.49; Adjusted p−value=0.008
Average gene set expression
3
2
1
0
−1
−2
Expression threshold
Below
Above
sample8_LMM_B3
KEGG_PPAR_SIGNALING_PATHWAY
Proportion of pathway genes=0.67; Adjusted p−value=0
Average gene set expression
3
2
1
0
−1
Expression threshold
Below
Above

sample8_LMM_B3
KEGG_PRION_DISEASES
Proportion of pathway genes=0.79; Adjusted p−value=0

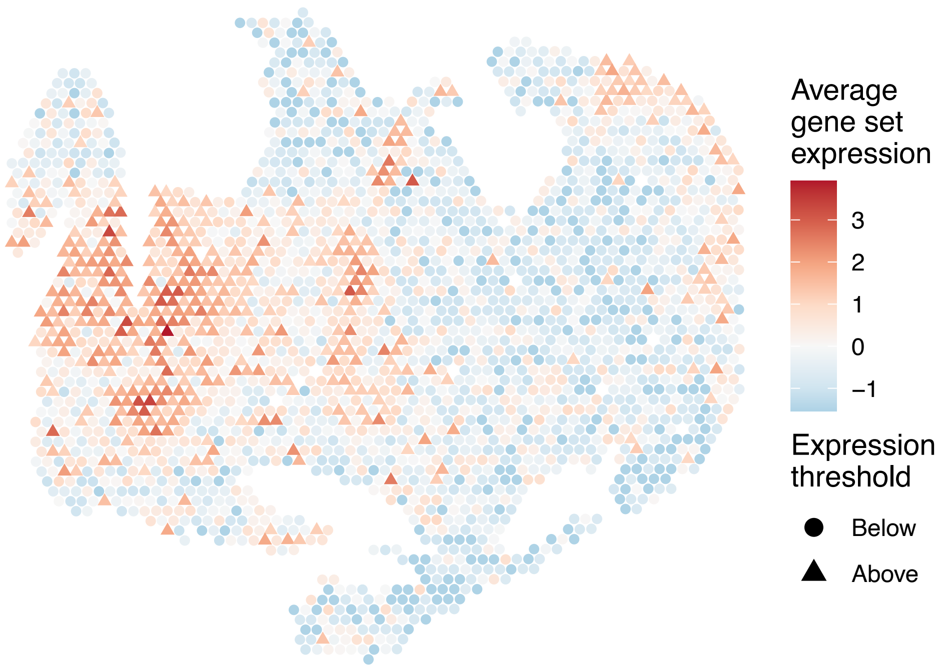


sample8_LMM_B3
KEGG_PROGESTERONE_MEDIATED_OOCYTE_MATURATION
Proportion of pathway genes=0.82; Adjusted p−value=0

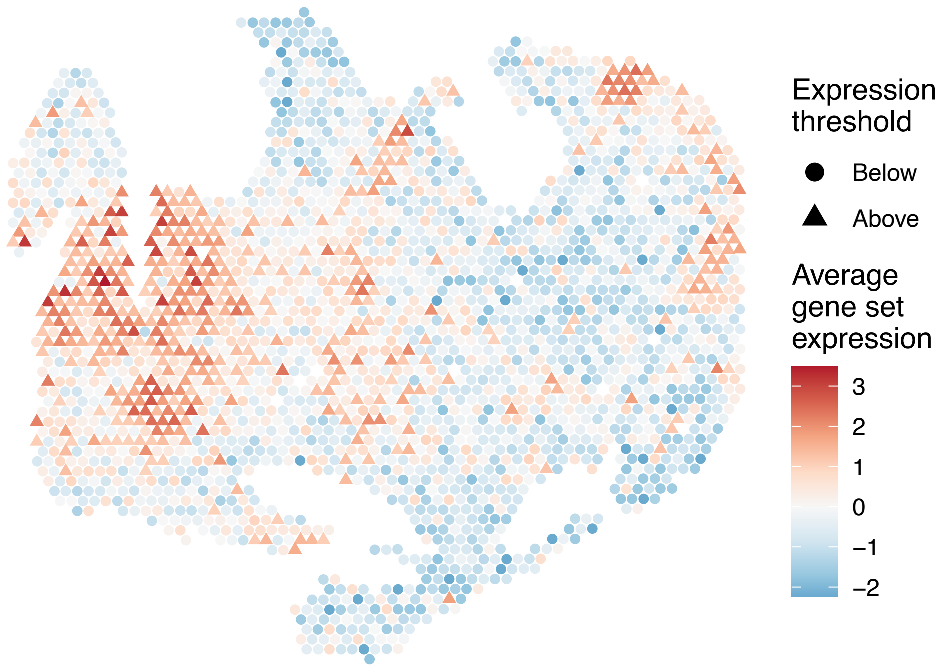


sample8_LMM_B3
KEGG_PROPANOATE_METABOLISM
Proportion of pathway genes=0.86; Adjusted p−value=0.0158

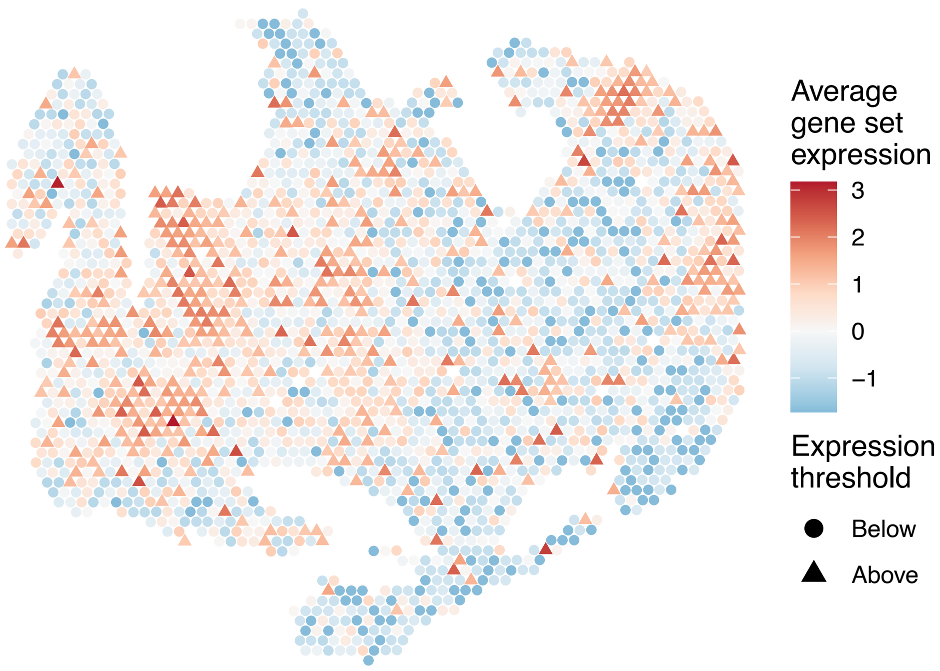


sample8_LMM_B3
KEGG_PROSTATE_CANCER
Proportion of pathway genes=0.86; Adjusted p−value=0

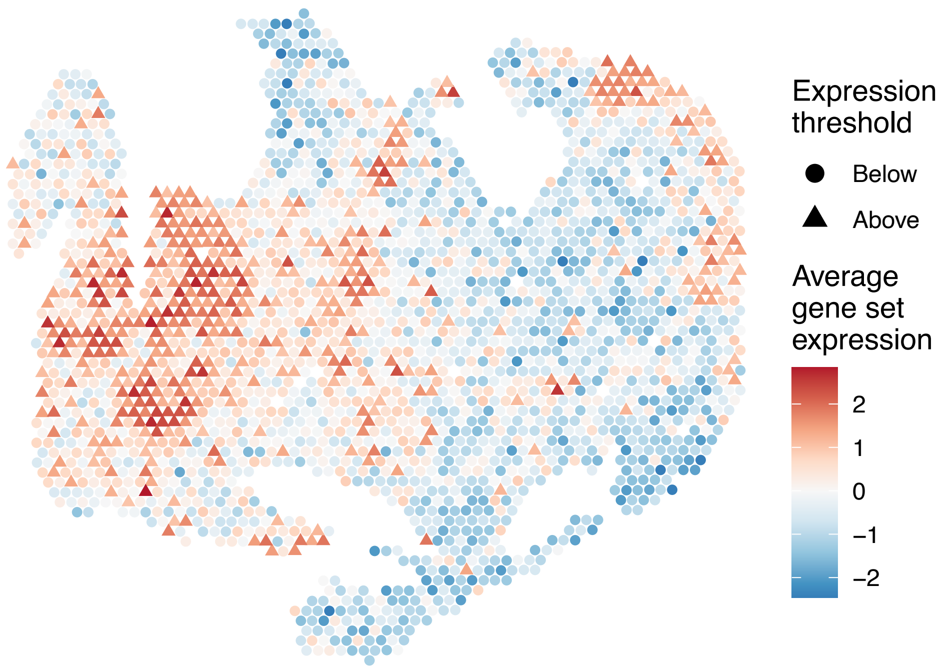

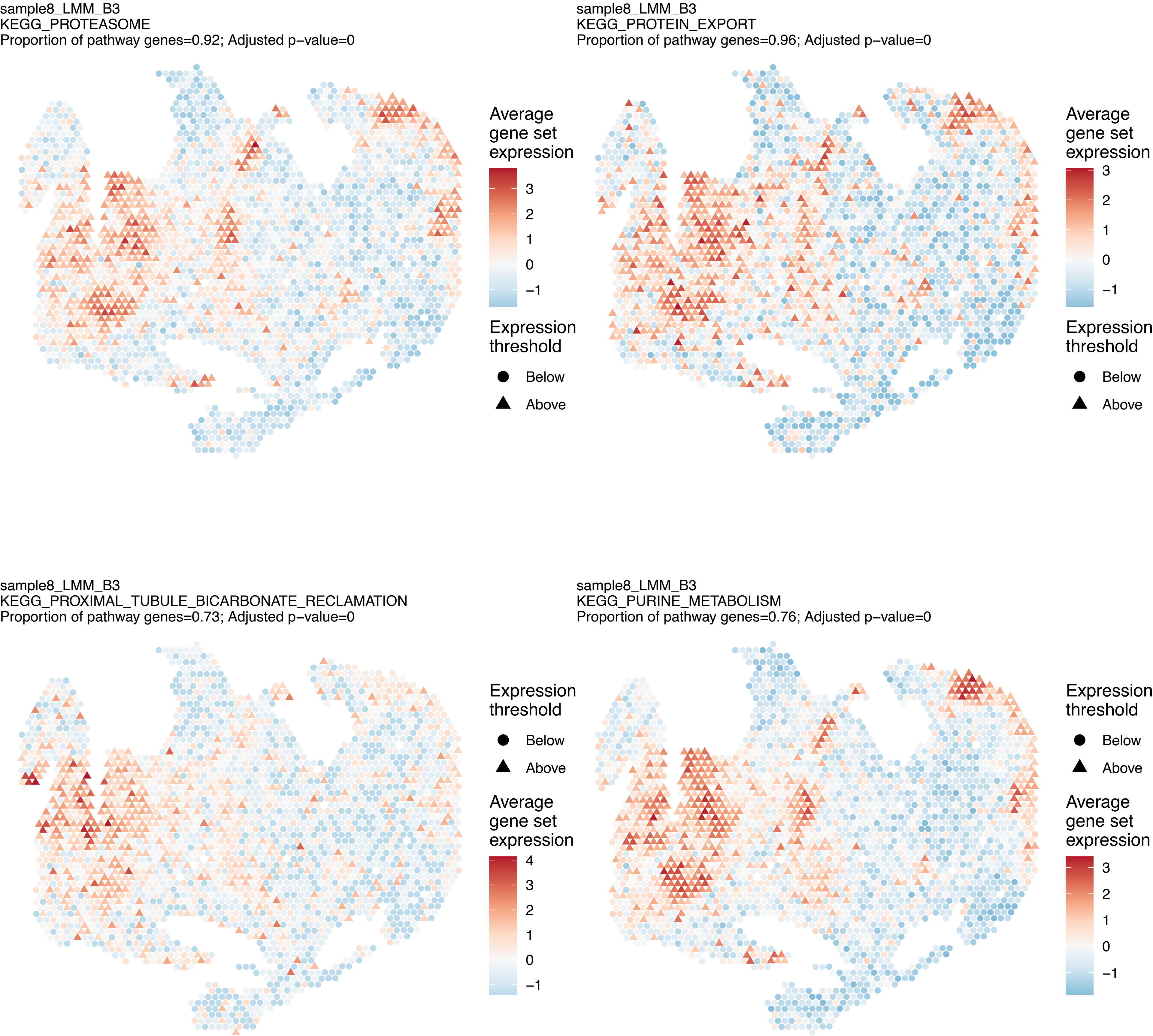
sample8_LMM_B3
KEGG_PROTEASOME
Proportion of pathway genes=0.92; Adjusted p−value=0
Average gene set expression
3
2
1
0
−1
Expression threshold
Below
Above
sample8_LMM_B3
KEGG_PROTEIN_EXPORT
Proportion of pathway genes=0.96; Adjusted p−value=0
Average gene set expression
3
2
1
0
−1
Expression threshold
Below
Above
sample8_LMM_B3
KEGG_PROXIMAL_TUBULE_BICARBONATE_RECLAMATION
Proportion of pathway genes=0.73; Adjusted p−value=0
Expression threshold
Below
Above
Average gene set expression
4
3
2
1
0
−1
sample8_LMM_B3
KEGG_PURINE_METABOLISM
Proportion of pathway genes=0.76; Adjusted p−value=0
Expression threshold
Below
Above
Average gene set expression
3
2
1
0
−1

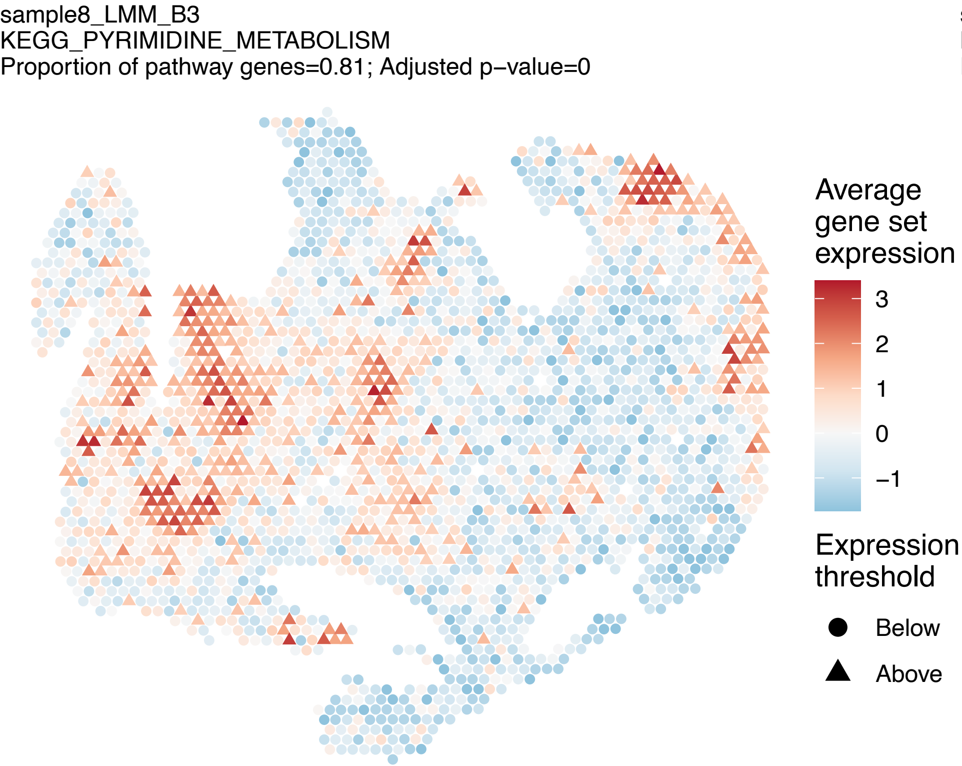
sample8_LMM_B3
KEGG_PYRIMIDINE_METABOLISM
Proportion of pathway genes=0.81; Adjusted p−value=0
Average gene set expression
3
2
1
0
−1
Expression threshold
Below
Above

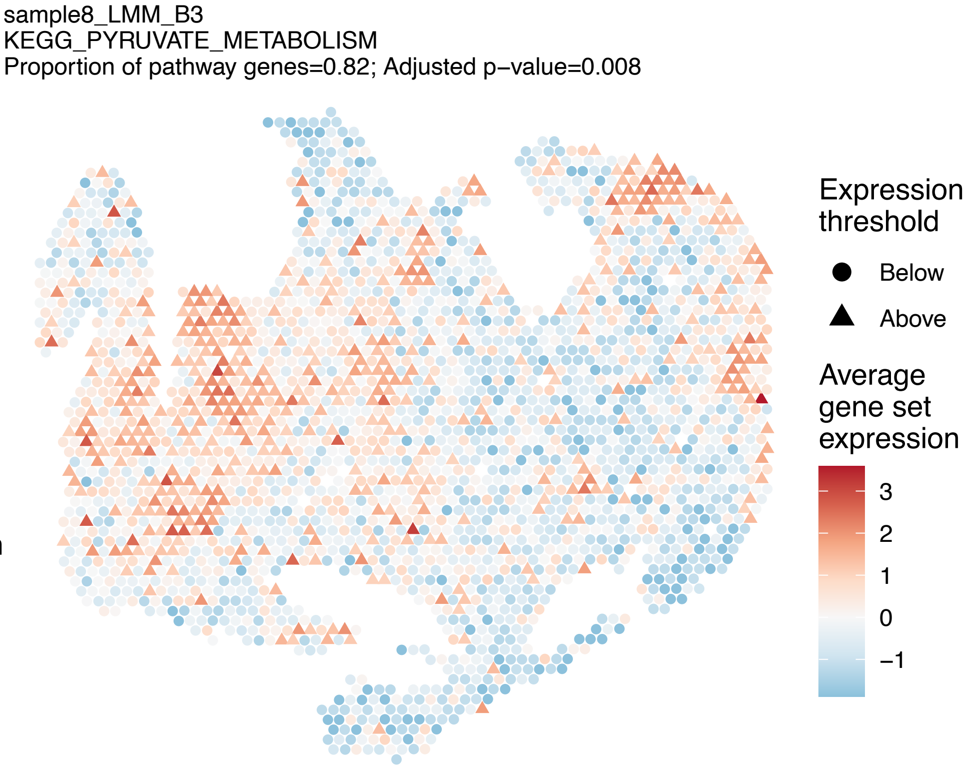
sample8_LMM_B3
KEGG_PYRUVATE_METABOLISM
Proportion of pathway genes=0.82; Adjusted p−value=0.008
Expression threshold
Below
Above
Average gene set expression
3
2
1
0
−1

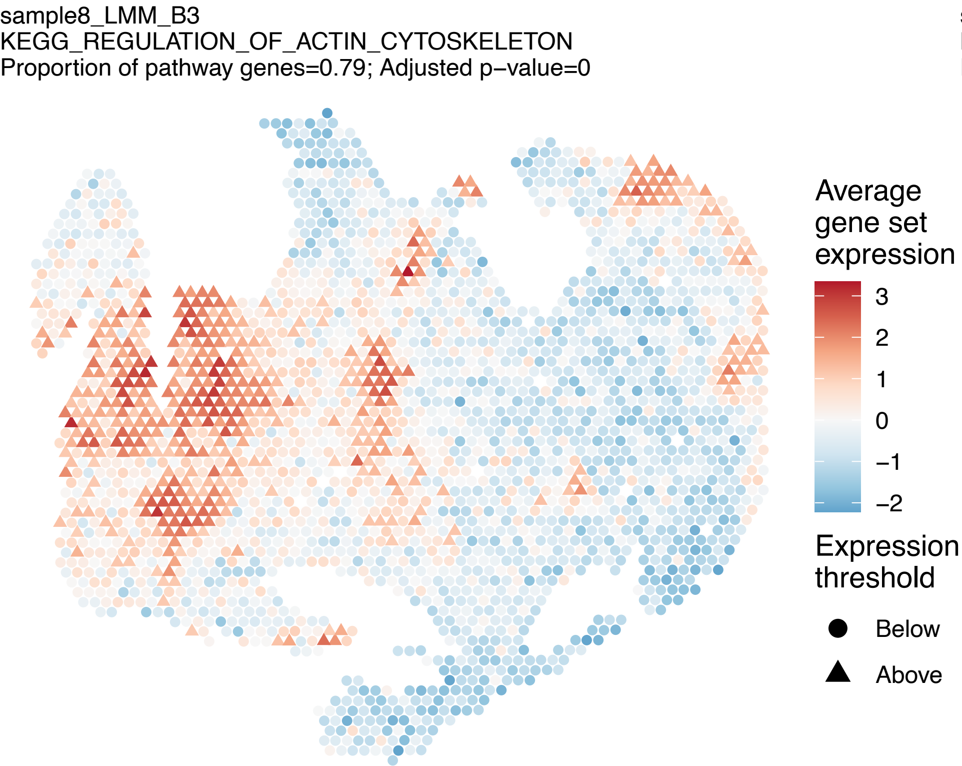
sample8_LMM_B3
KEGG_REGULATION_OF_ACTIN_CYTOSKELETON
Proportion of pathway genes=0.79; Adjusted p−value=0
Average gene set expression
3
2
1
0
−1
−2
Expression threshold
Below
Above

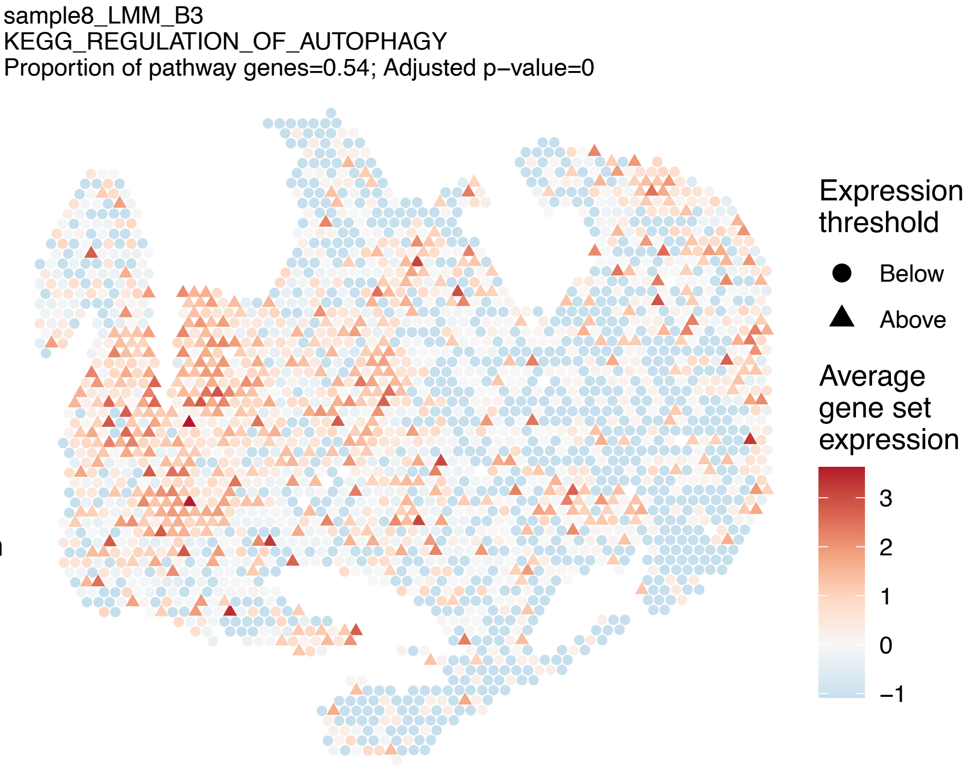
sample8_LMM_B3
KEGG_REGULATION_OF_AUTOPHAGY
Proportion of pathway genes=0.54; Adjusted p−value=0
Expression threshold
Below
Above
Average gene set expression
3
2
1
0
−1

sample8_LMM_B3
KEGG_RENAL_CELL_CARCINOMA
Proportion of pathway genes=0.94; Adjusted p−value=0

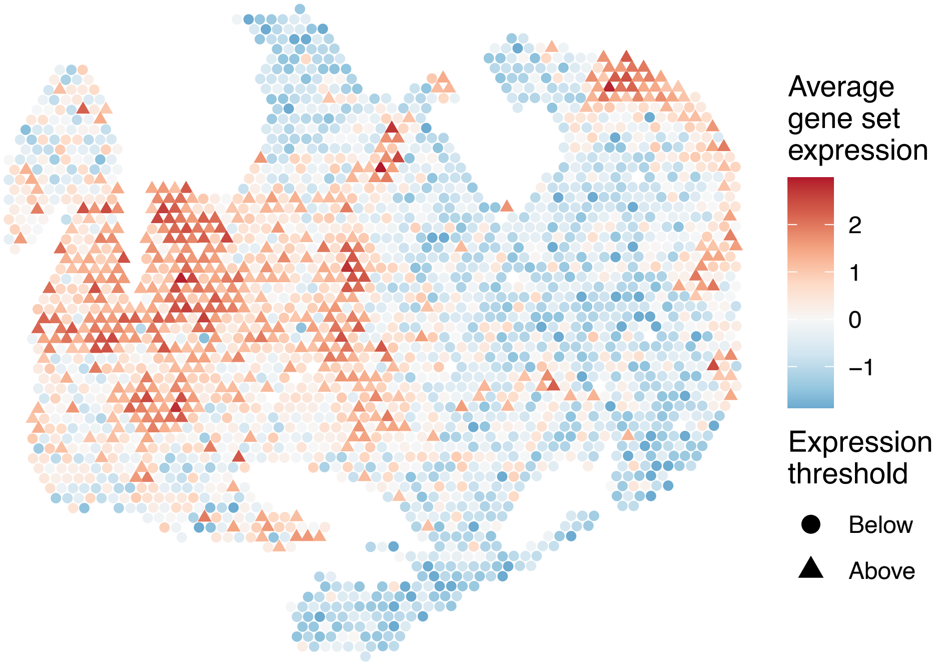


sample8_LMM_B3
KEGG_RETINOL_METABOLISM
Proportion of pathway genes=0.28; Adjusted p−value=0

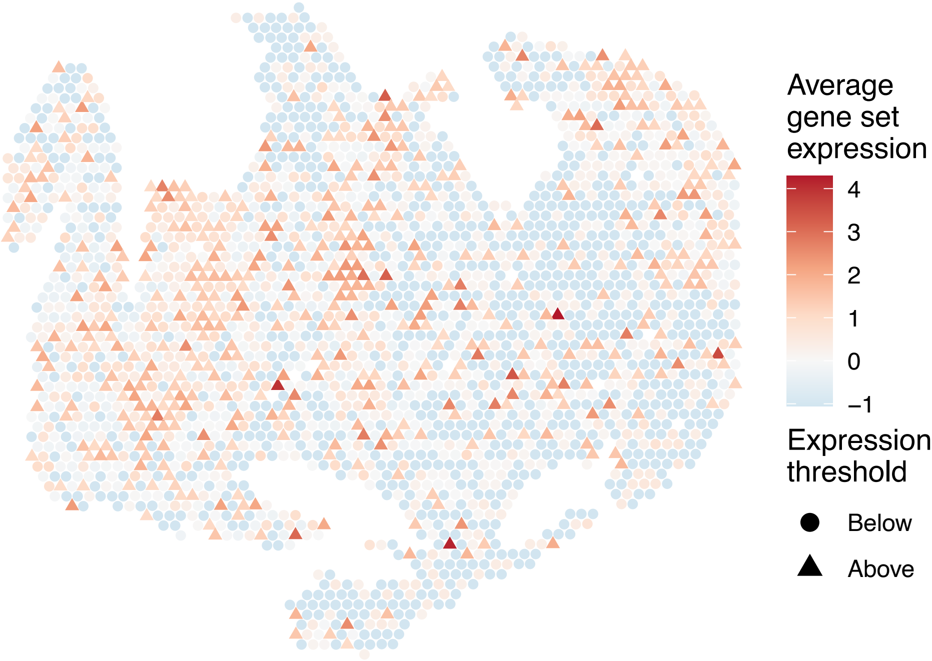


sample8_LMM_B3
KEGG_RIBOFLAVIN_METABOLISM
Proportion of pathway genes=0.81; Adjusted p−value=0.0432

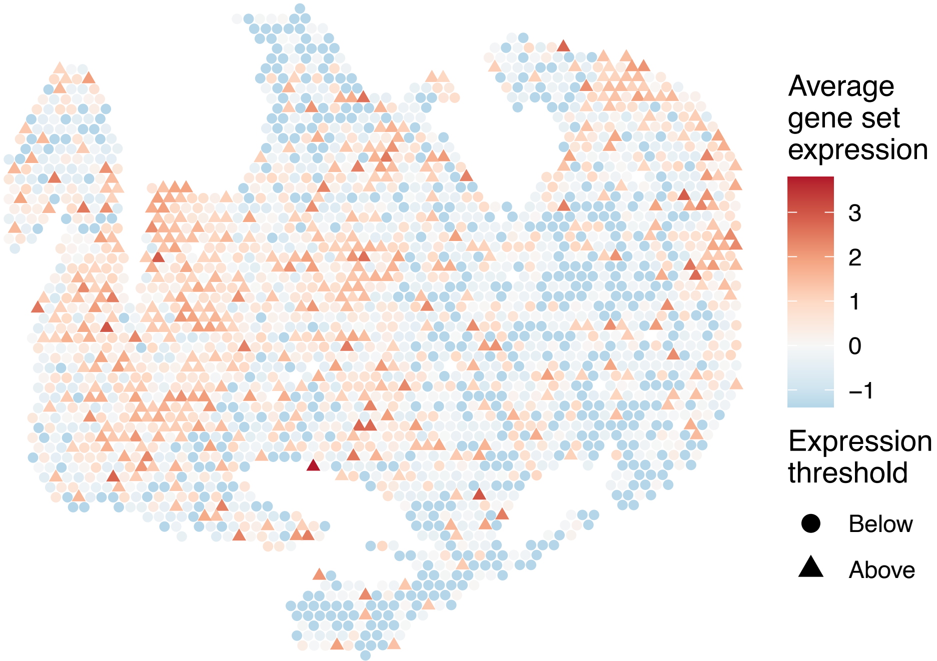


sample8_LMM_B3
KEGG_RIG_I_LIKE_RECEPTOR_SIGNALING_PATHWAY
Proportion of pathway genes=0.62; Adjusted p−value=0

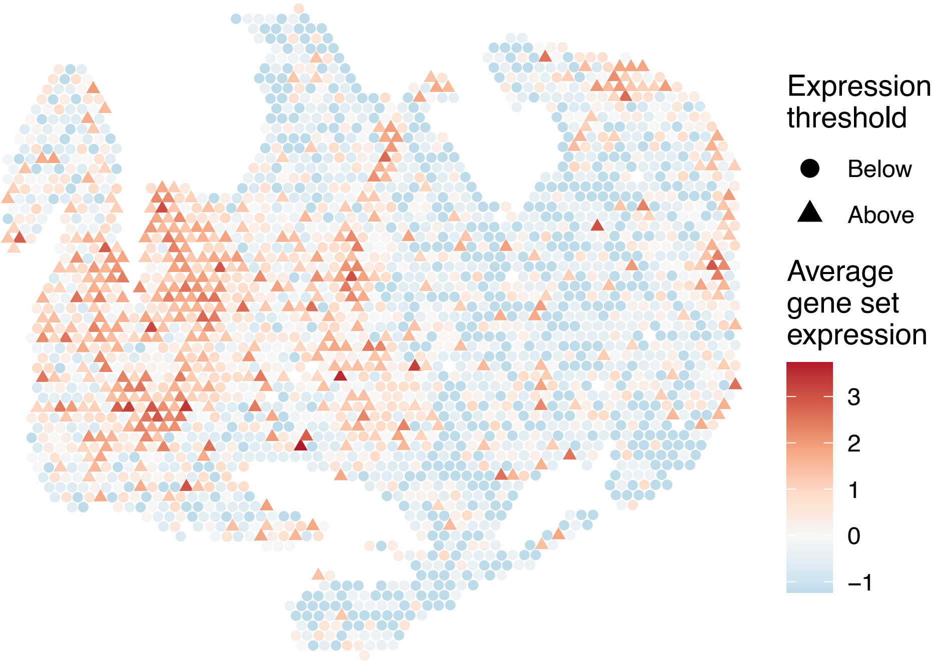

sample8_LMM_B3
KEGG_RNA_DEGRADATION
Proportion of pathway genes=0.96; Adjusted p−value=0

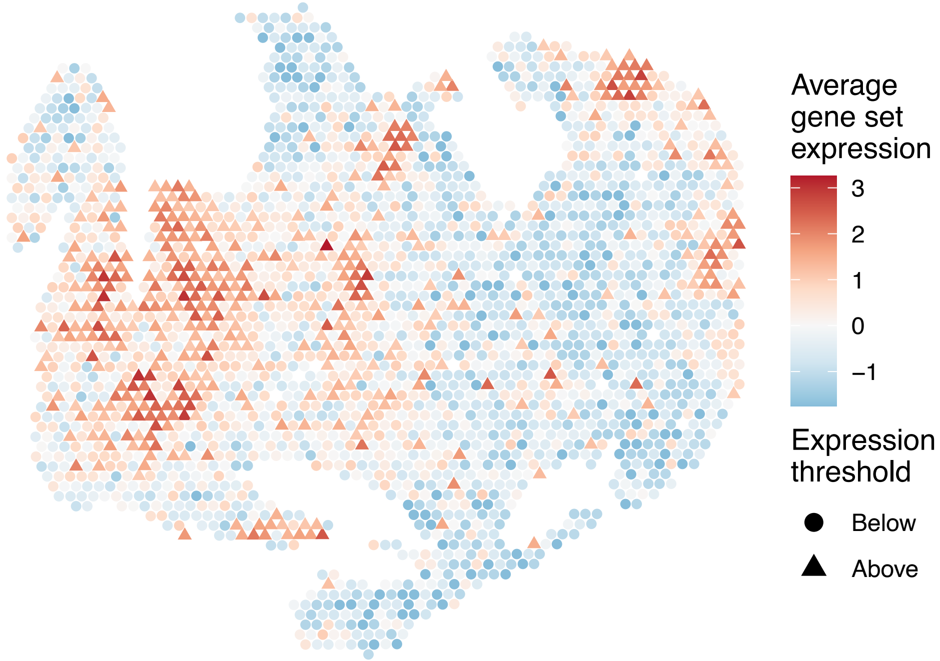


sample8_LMM_B3
KEGG_RNA_POLYMERASE
Proportion of pathway genes=0.74; Adjusted p−value=0.0177

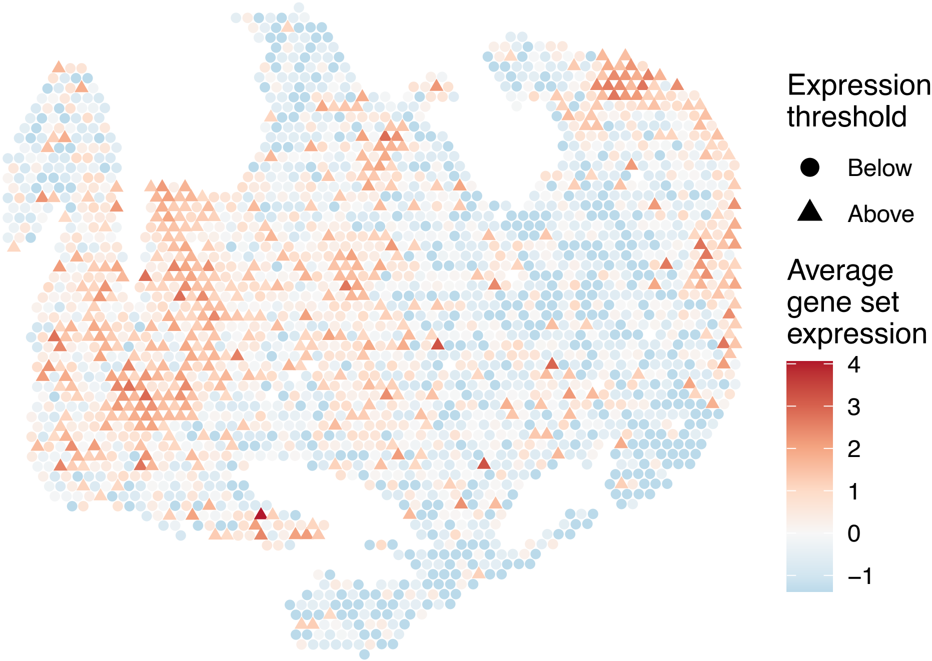


sample8_LMM_B3
KEGG_SELENOAMINO_ACID_METABOLISM
Proportion of pathway genes=0.85; Adjusted p−value=0.0012

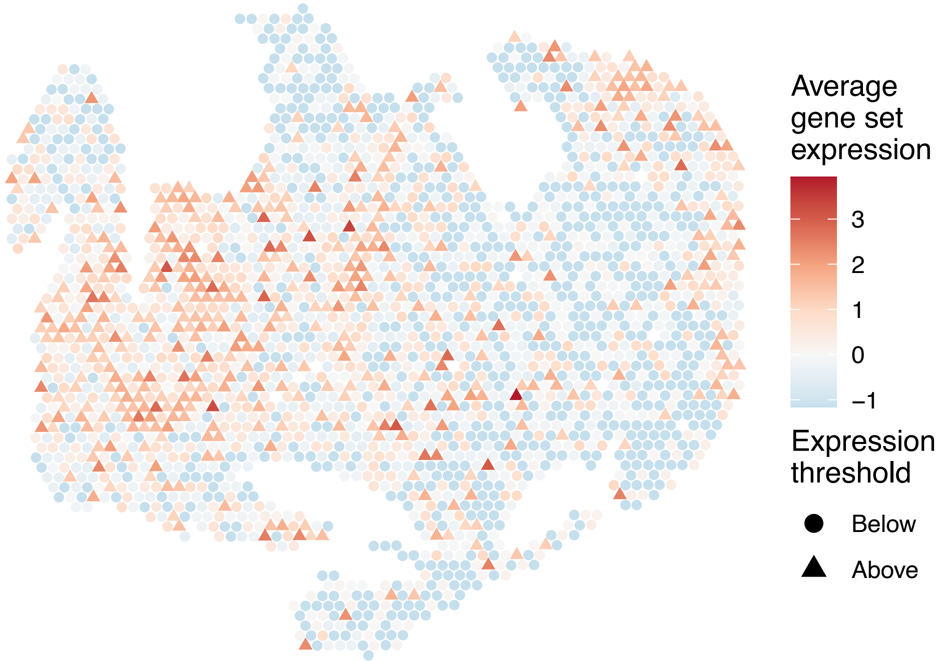


sample8_LMM_B3
KEGG_SMALL_CELL_LUNG_CANCER
Proportion of pathway genes=0.91; Adjusted p−value=0

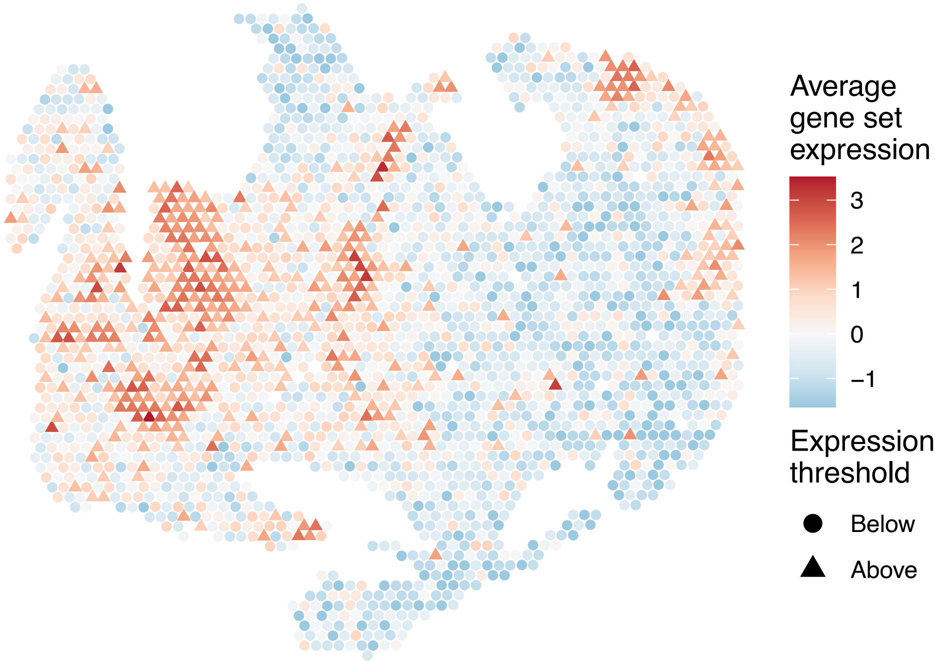

sample8_LMM_B3
KEGG_SNARE_INTERACTIONS_IN_VESICULAR_TRANSPORT
Proportion of pathway genes=0.95; Adjusted p−value=0

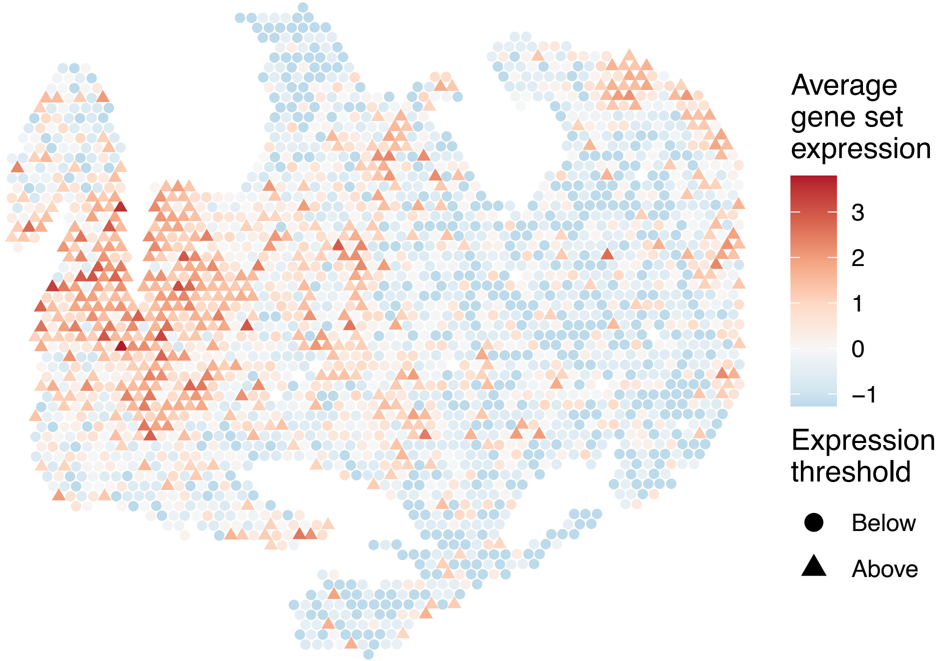


sample8_LMM_B3
KEGG_SPHINGOLIPID_METABOLISM
Proportion of pathway genes=0.84; Adjusted p−value=0

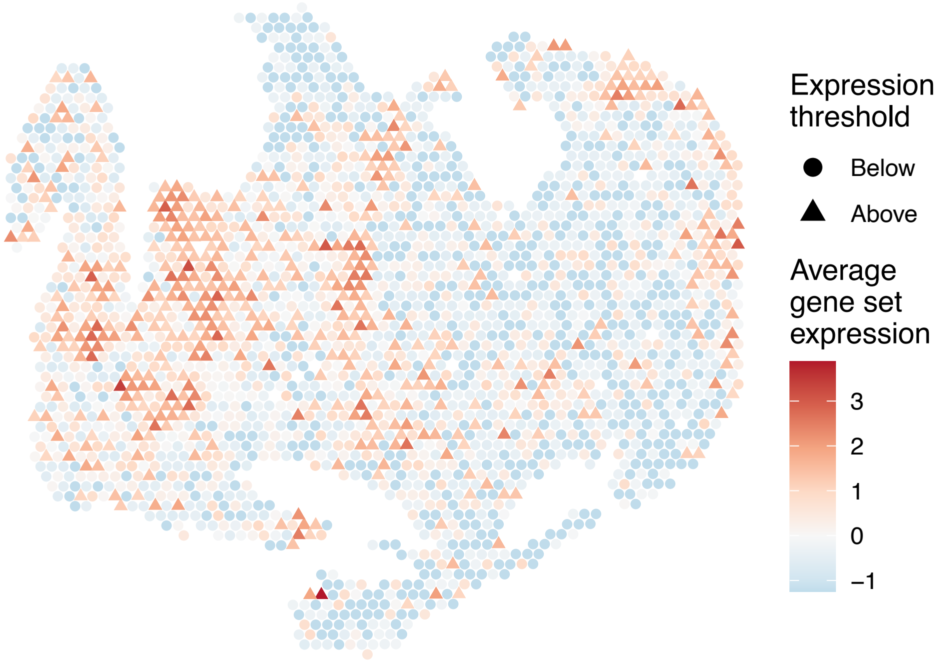


sample8_LMM_B3
KEGG_SPLICEOSOME
Proportion of pathway genes=0.99; Adjusted p−value=0

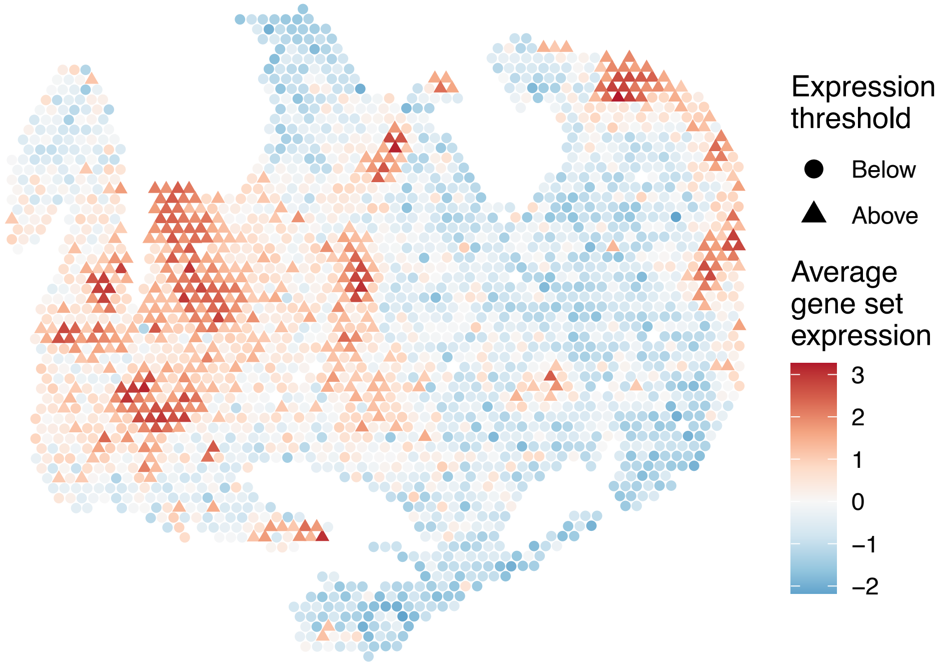


sample8_LMM_B3
KEGG_STARCH_AND_SUCROSE_METABOLISM
Proportion of pathway genes=0.36; Adjusted p−value=0

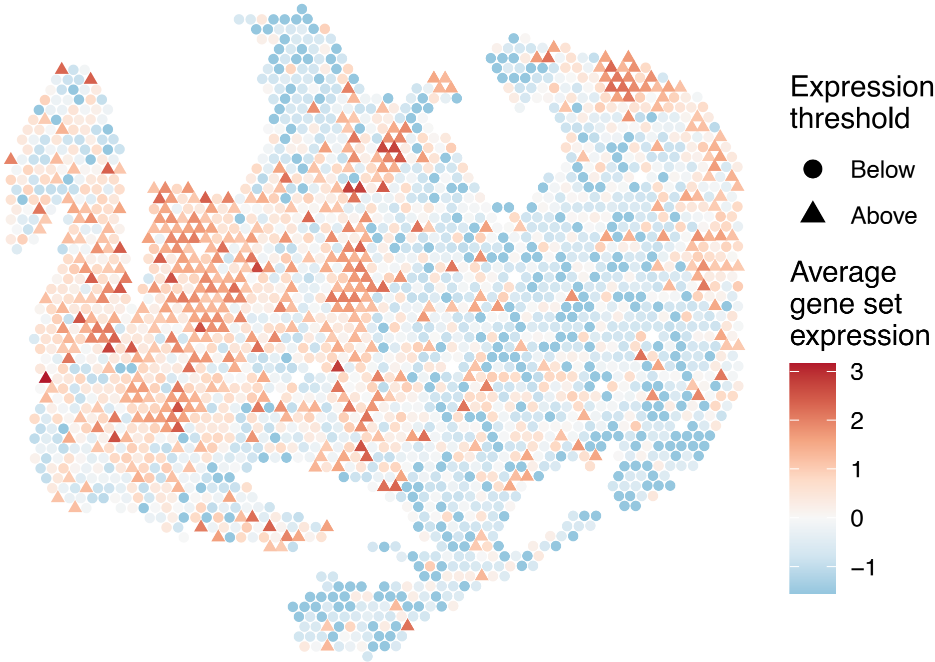

sample8_LMM_B3
KEGG_STEROID_BIOSYNTHESIS
Proportion of pathway genes=0.84; Adjusted p−value=0.0177

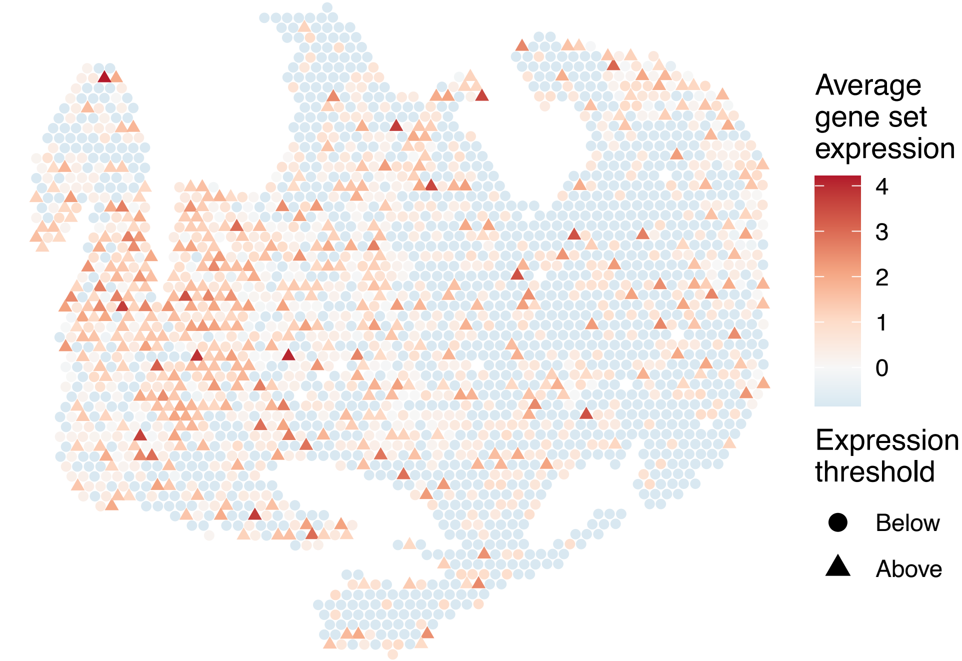


sample8_LMM_B3
KEGG_STEROID_HORMONE_BIOSYNTHESIS
Proportion of pathway genes=0.25; Adjusted p−value=0

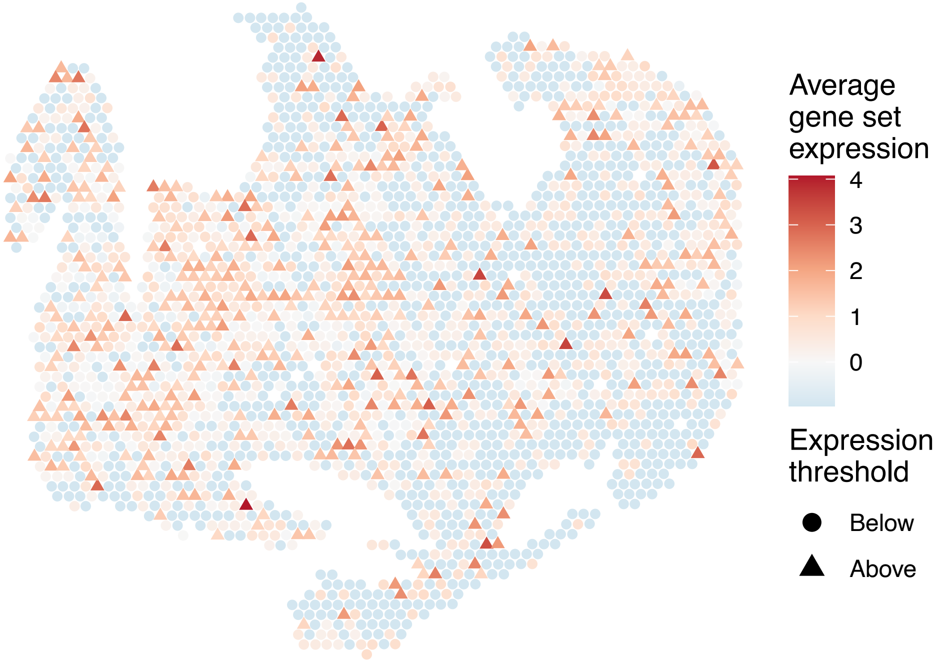


sample8_LMM_B3
KEGG_SULFUR_METABOLISM
Proportion of pathway genes=0.62; Adjusted p−value=NA

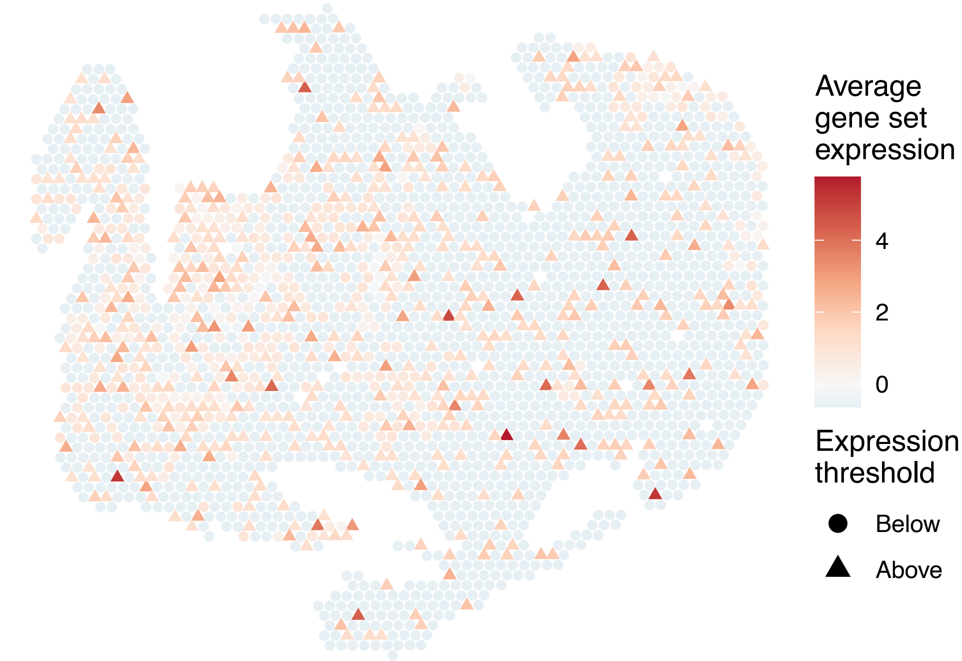


sample8_LMM_B3
KEGG_SYSTEMIC_LUPUS_ERYTHEMATOSUS
Proportion of pathway genes=0.35; Adjusted p−value=0

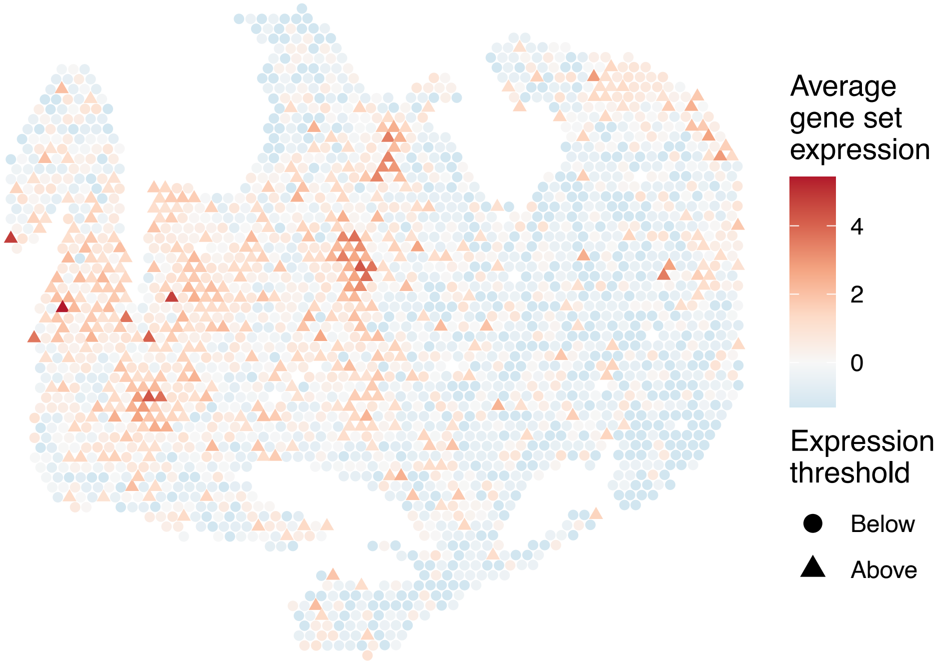

sample8_LMM_B3
KEGG_T_CELL_RECEPTOR_SIGNALING_PATHWAY
Proportion of pathway genes=0.69; Adjusted p−value=0

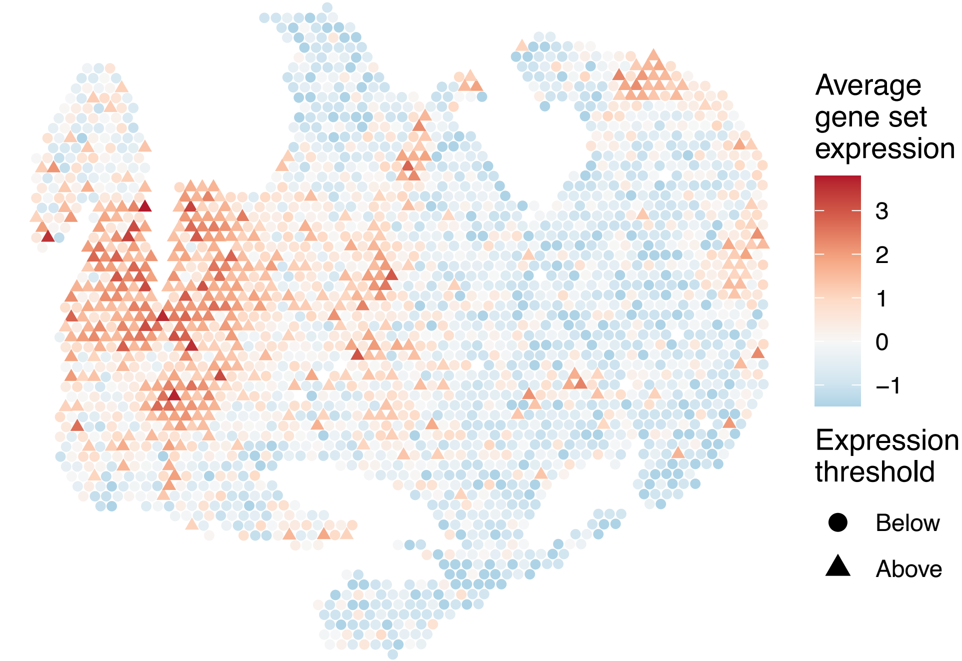


sample8_LMM_B3
KEGG_TERPENOID_BACKBONE_BIOSYNTHESIS
Proportion of pathway genes=0.87; Adjusted p−value=0.1426

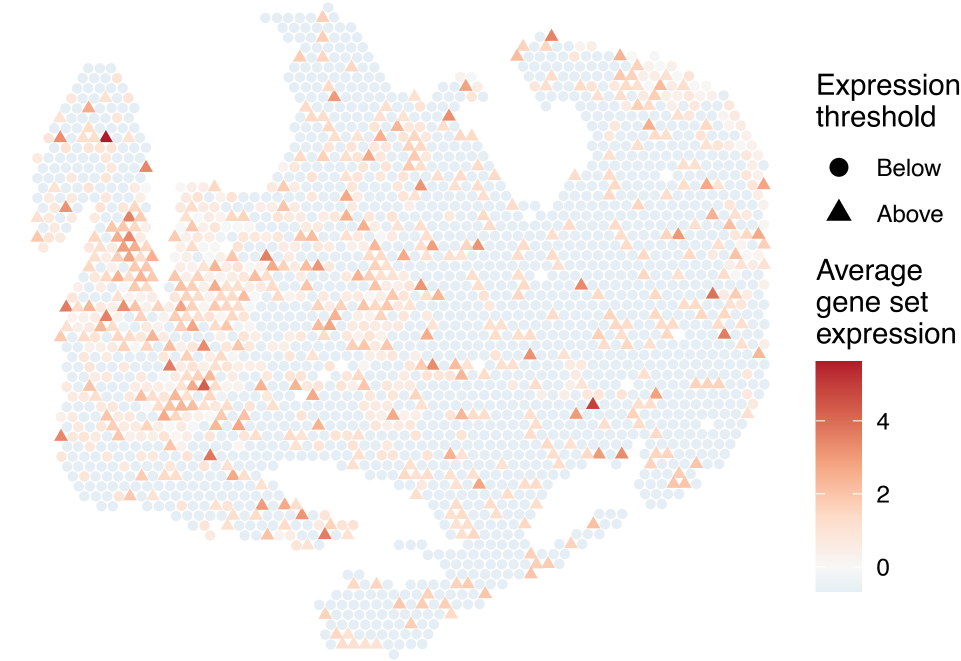


sample8_LMM_B3
KEGG_TGF_BETA_SIGNALING_PATHWAY
Proportion of pathway genes=0.71; Adjusted p−value=0

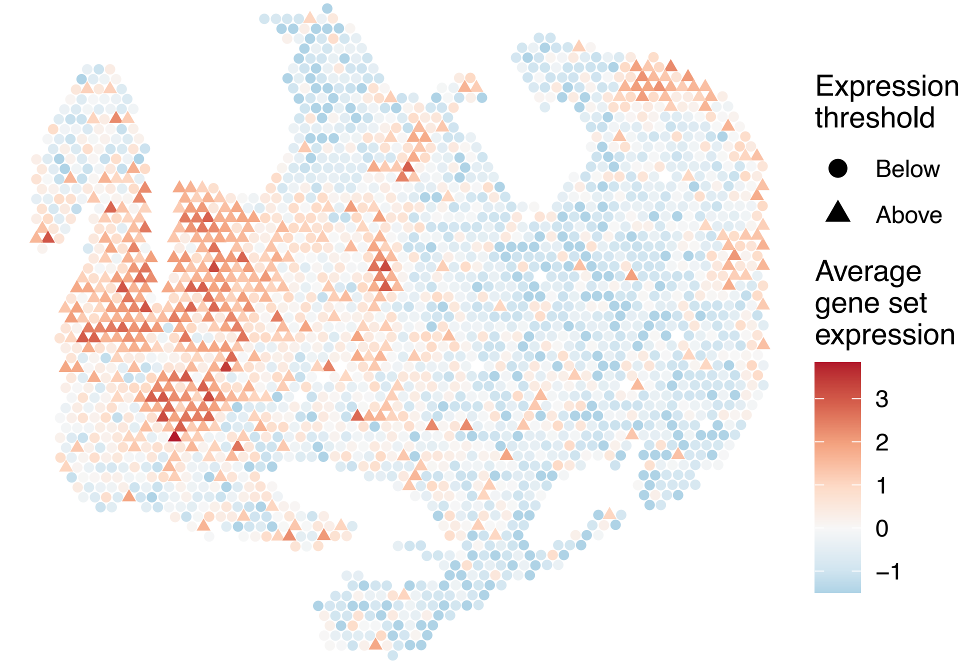


sample8_LMM_B3
KEGG_THYROID_CANCER
Proportion of pathway genes=0.89; Adjusted p−value=0

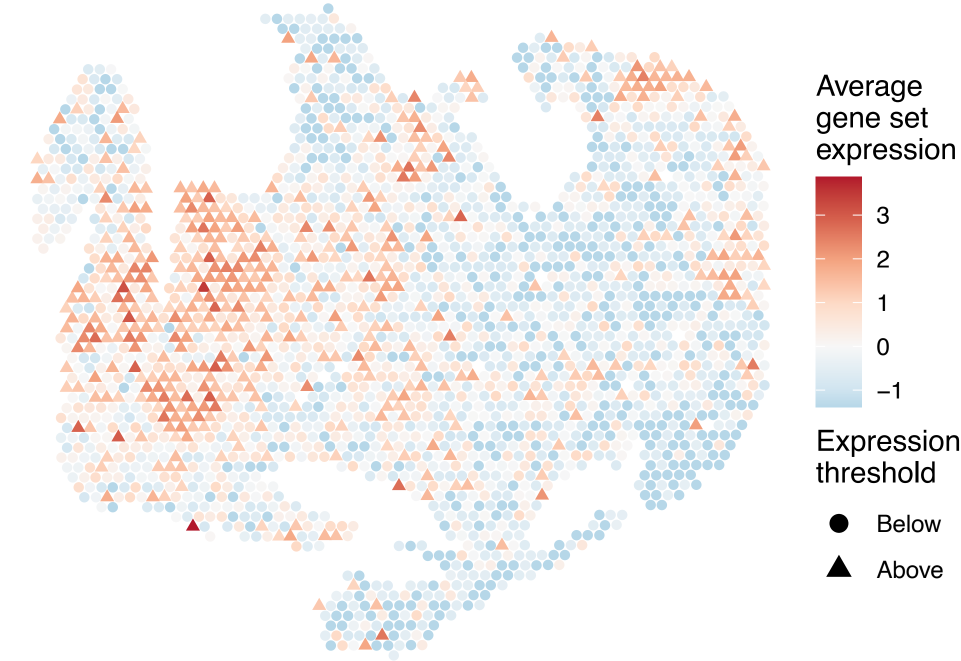

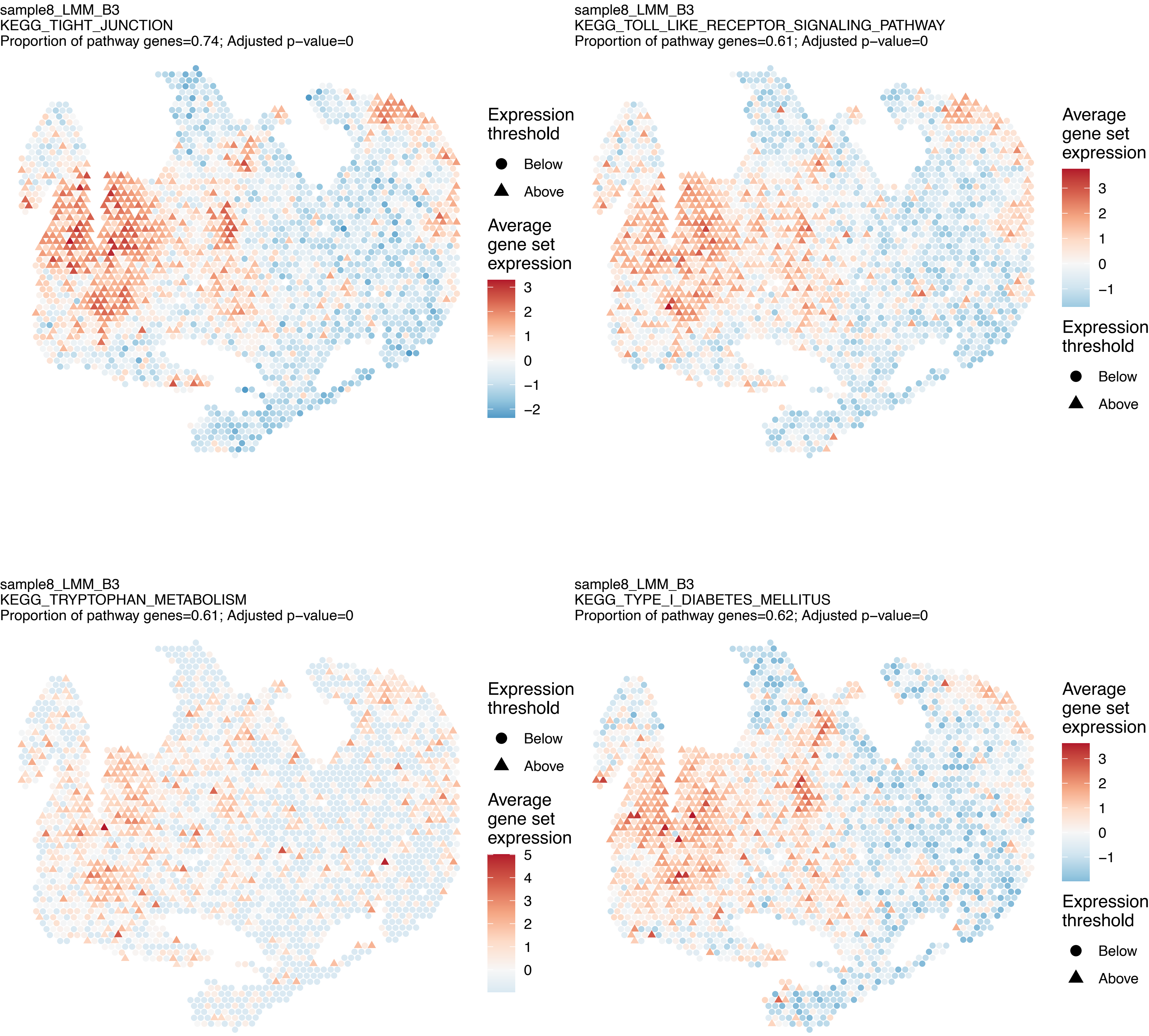
sample8_LMM_B3
KEGG_TIGHT_JUNCTION
Proportion of pathway genes=0.74; Adjusted p−value=0
Expression threshold
Below
Above
Average gene set expression
3
2
1
0
−1
−2
sample8_LMM_B3
KEGG_TOLL_LIKE_RECEPTOR_SIGNALING_PATHWAY
Proportion of pathway genes=0.61; Adjusted p−value=0
Average gene set expression
3
2
1
0
−1
Expression threshold
Below
Above
sample8_LMM_B3
KEGG_TRYPTOPHAN_METABOLISM
Proportion of pathway genes=0.61; Adjusted p−value=0
Expression threshold
Below
Above
Average gene set expression
5
4
3
2
1
0
sample8_LMM_B3
KEGG_TYPE_I_DIABETES_MELLITUS
Proportion of pathway genes=0.62; Adjusted p−value=0
Average gene set expression
3
2
1
0
−1
Expression threshold
Below
Above

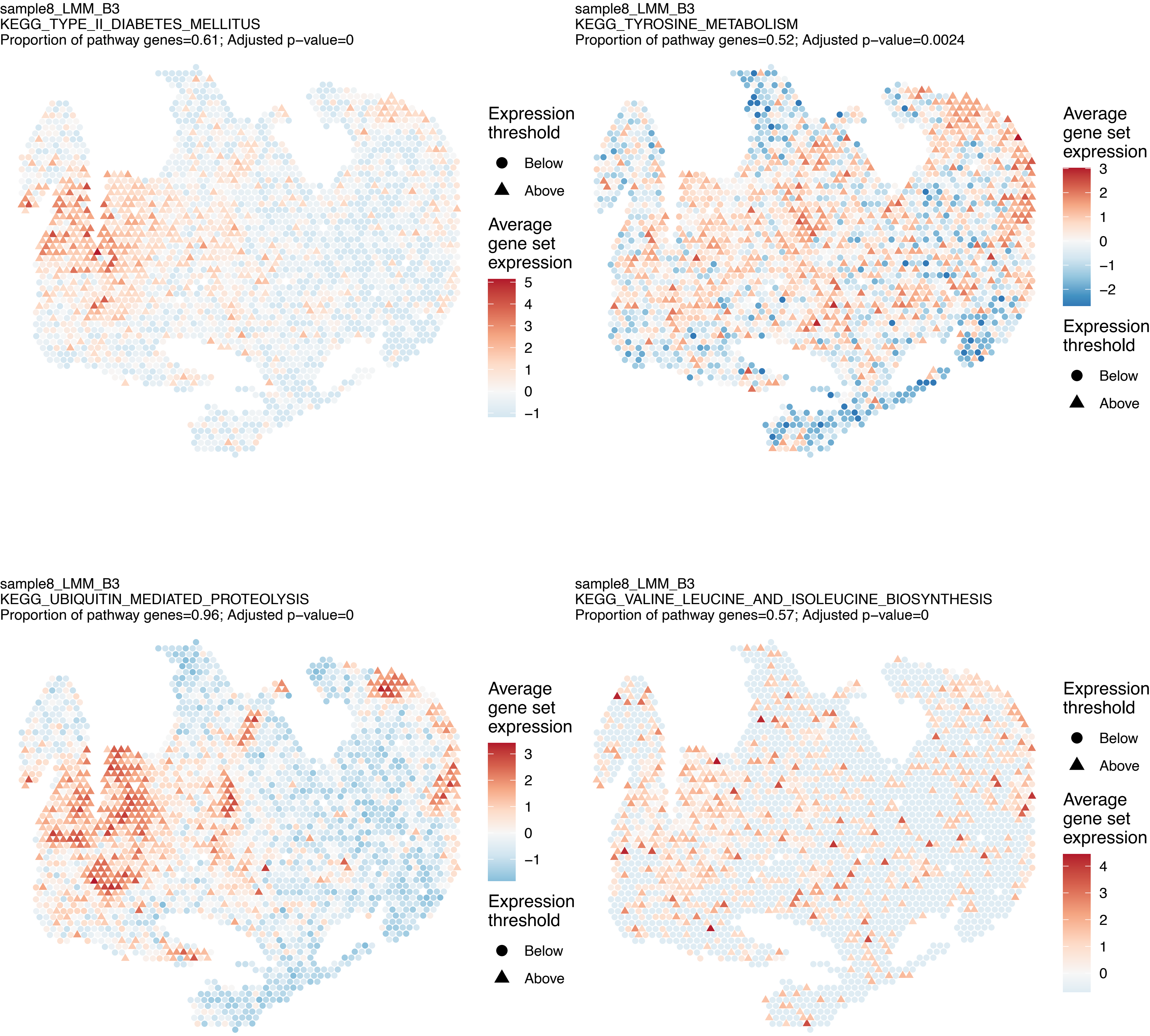
sample8_LMM_B3
KEGG_TYPE_II_DIABETES_MELLITUS
Proportion of pathway genes=0.61; Adjusted p−value=0
Expression threshold
Below
Above
Average gene set expression
5
4
3
2
1
0
−1
sample8_LMM_B3
KEGG_TYROSINE_METABOLISM
Proportion of pathway genes=0.52; Adjusted p−value=0.0024
Average gene set expression
3
2
1
0
−1
−2
Expression threshold
Below
Above
sample8_LMM_B3
KEGG_UBIQUITIN_MEDIATED_PROTEOLYSIS
Proportion of pathway genes=0.96; Adjusted p−value=0
Average gene set expression
3
2
1
0
−1
Expression threshold
Below
Above
sample8_LMM_B3
KEGG_VALINE_LEUCINE_AND_ISOLEUCINE_BIOSYNTHESIS
Proportion of pathway genes=0.57; Adjusted p−value=0
Expression threshold
Below
Above
Average gene set expression
4
3
2
1
0

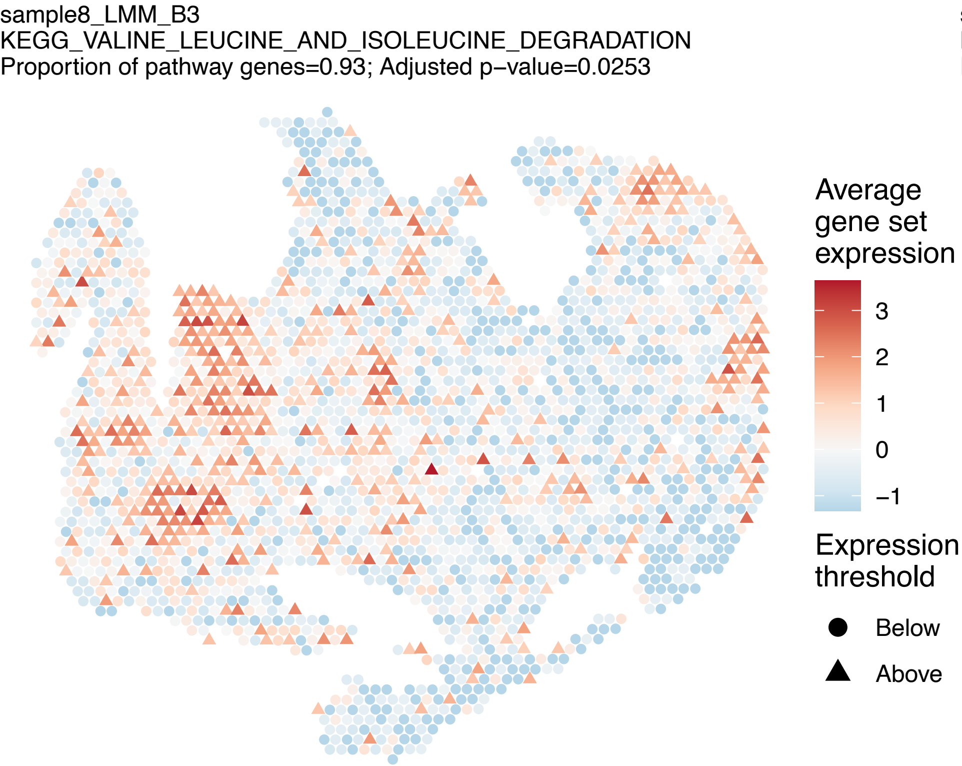
sample8_LMM_B3
KEGG_VALINE_LEUCINE_AND_ISOLEUCINE_DEGRADATION
Proportion of pathway genes=0.93; Adjusted p−value=0.0253
Average gene set expression
3
2
1
0
−1
Expression threshold
Below
Above

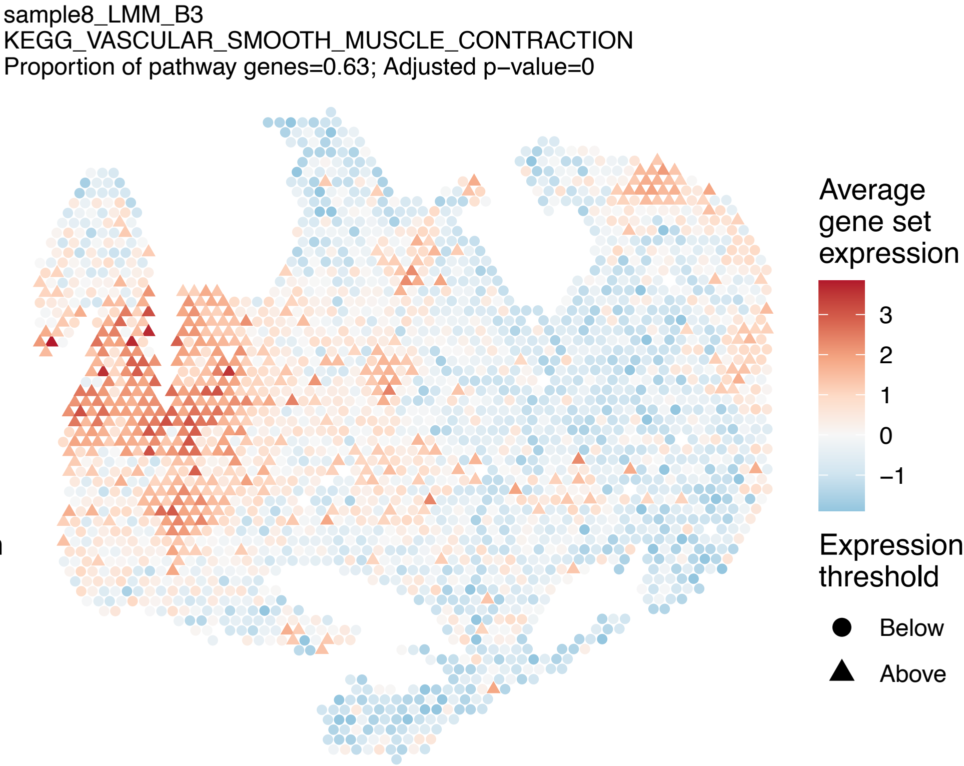
sample8_LMM_B3
KEGG_VASCULAR_SMOOTH_MUSCLE_CONTRACTION
Proportion of pathway genes=0.63; Adjusted p−value=0
Average gene set expression
3
2
1
0
−1
Expression threshold
Below
Above

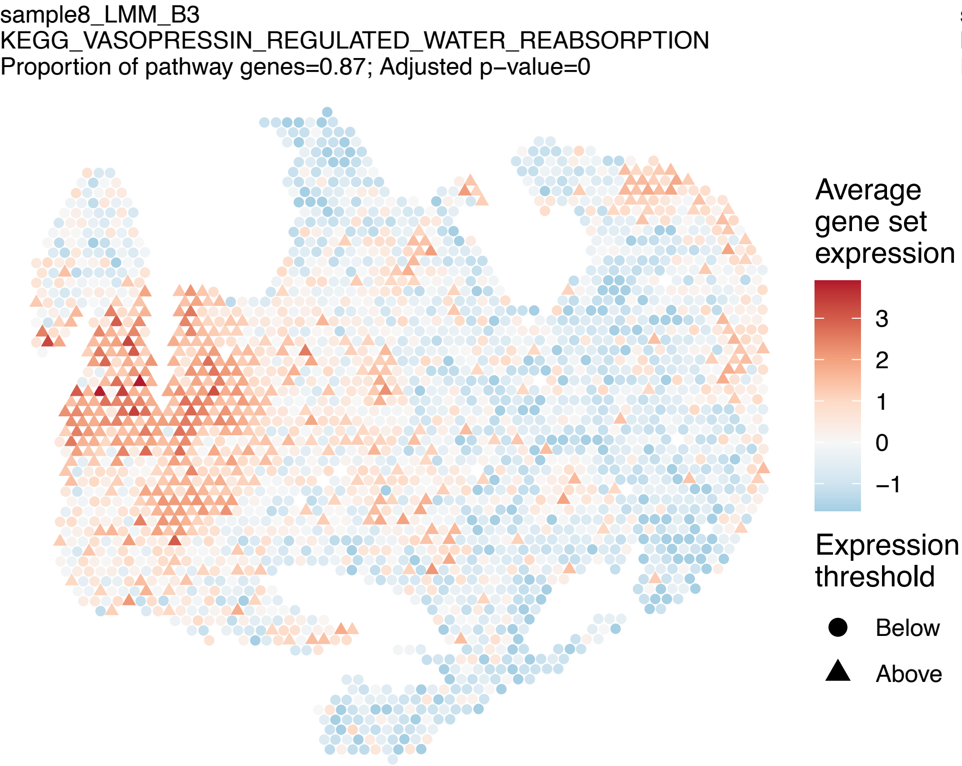
sample8_LMM_B3
KEGG_VASOPRESSIN_REGULATED_WATER_REABSORPTION
Proportion of pathway genes=0.87; Adjusted p−value=0
Average gene set expression
3
2
1
0
−1
Expression threshold
Below
Above

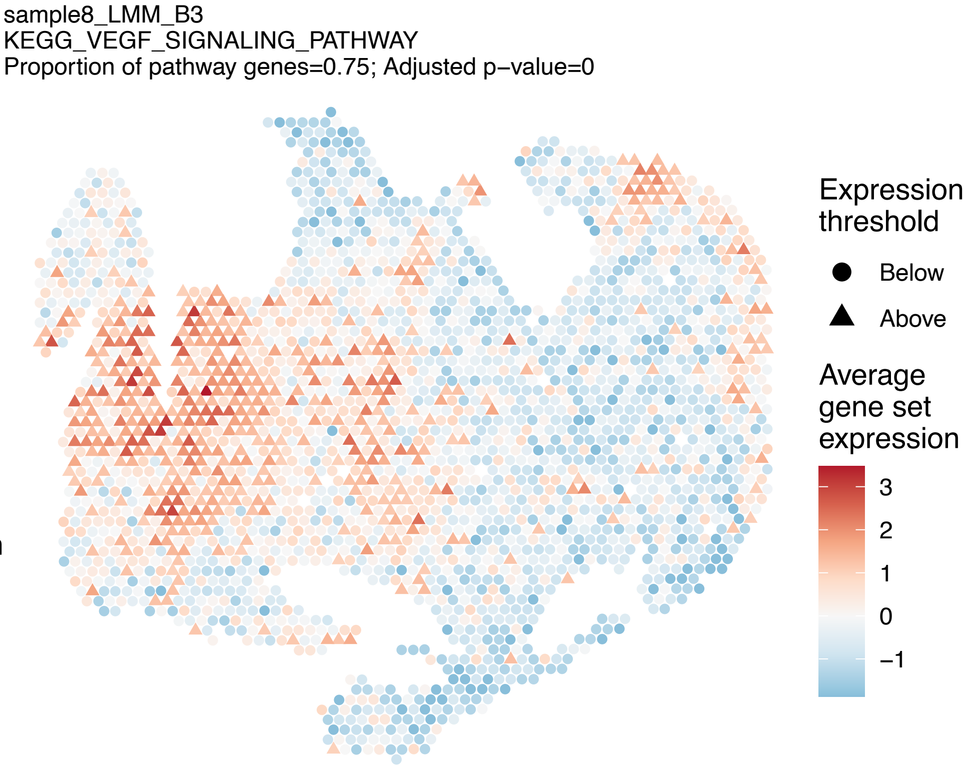
sample8_LMM_B3
KEGG_VEGF_SIGNALING_PATHWAY
Proportion of pathway genes=0.75; Adjusted p−value=0
Expression threshold
Below
Above
Average gene set expression
3
2
1
0
−1

sample8_LMM_B3
KEGG_VIBRIO_CHOLERAE_INFECTION
Proportion of pathway genes=0.84; Adjusted p–value=0

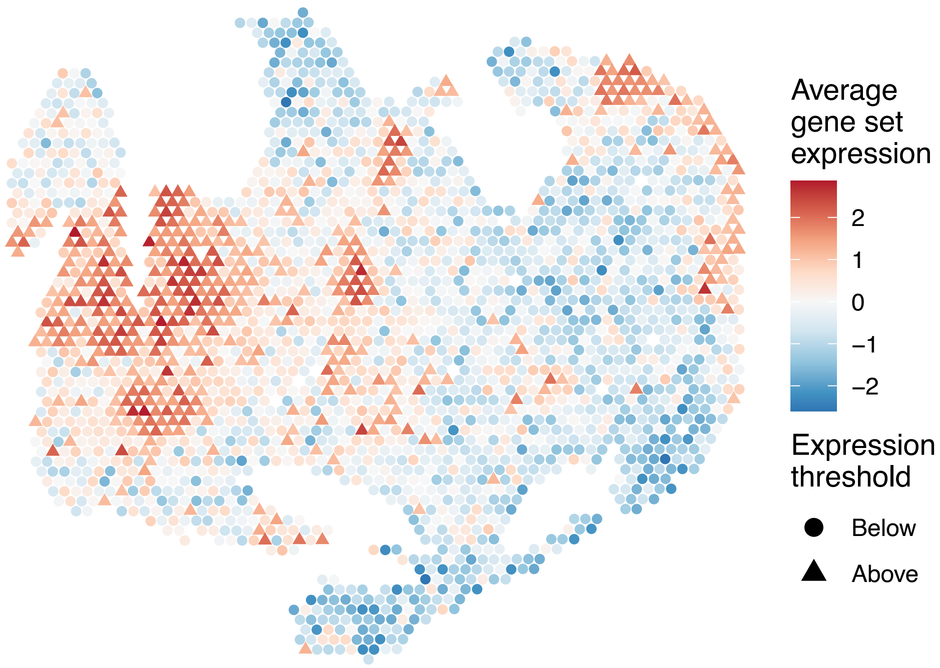


sample8_LMM_B3
KEGG_VIRAL_MYOCARDITIS
Proportion of pathway genes=0.69; Adjusted p–value=0

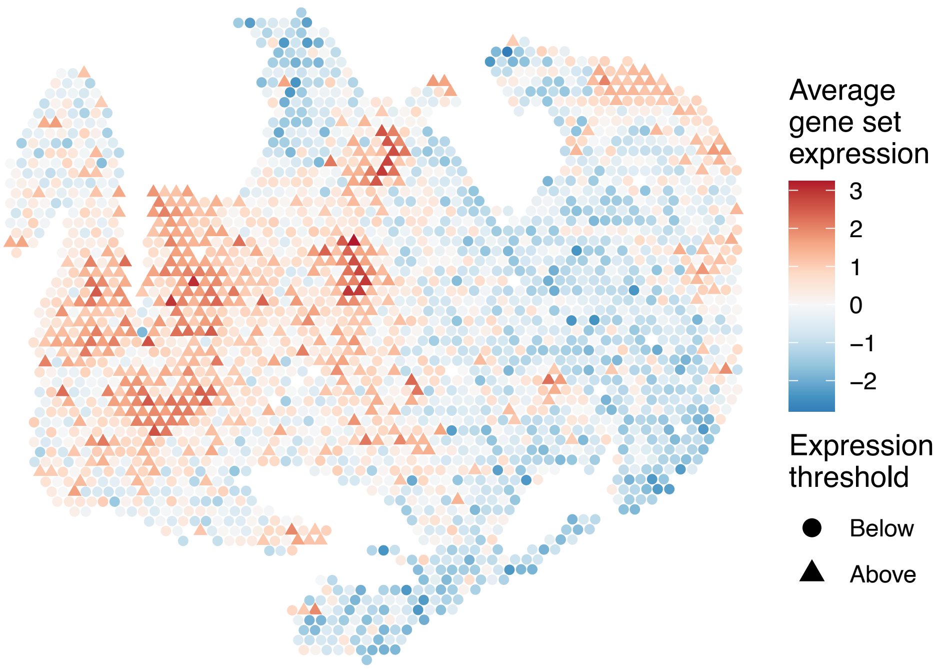


sample8_LMM_B3
KEGG_WNT_SIGNALING_PATHWAY
Proportion of pathway genes=0.8; Adjusted p–value=0

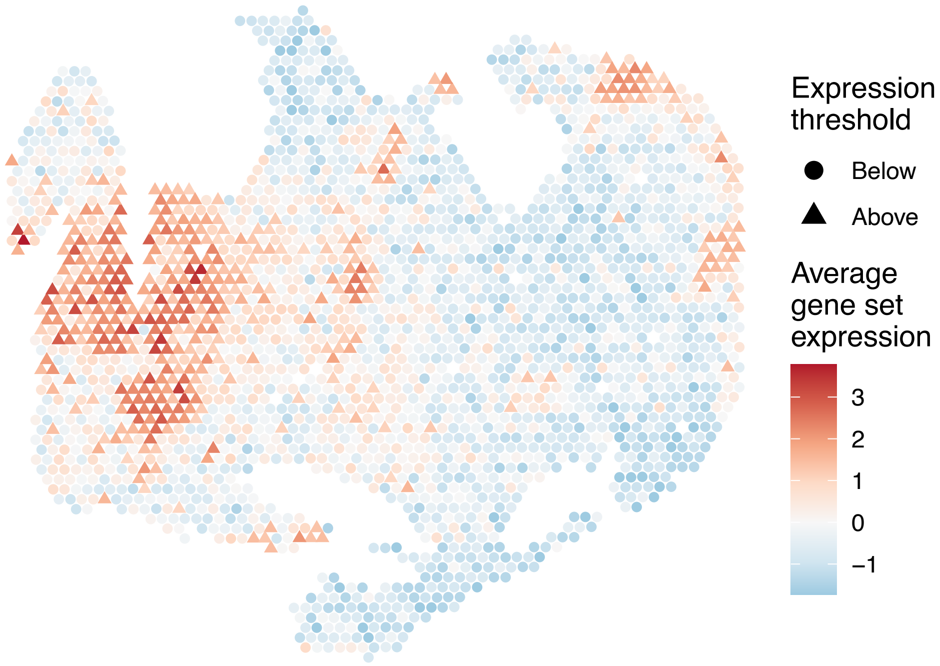


sample8_LMM_B3
KEGG_PENTOSE_AND_GLUCURONATE_INTERCONVERSIONS
Proportion of pathway genes=0.32; Adjusted p–value=0.0315

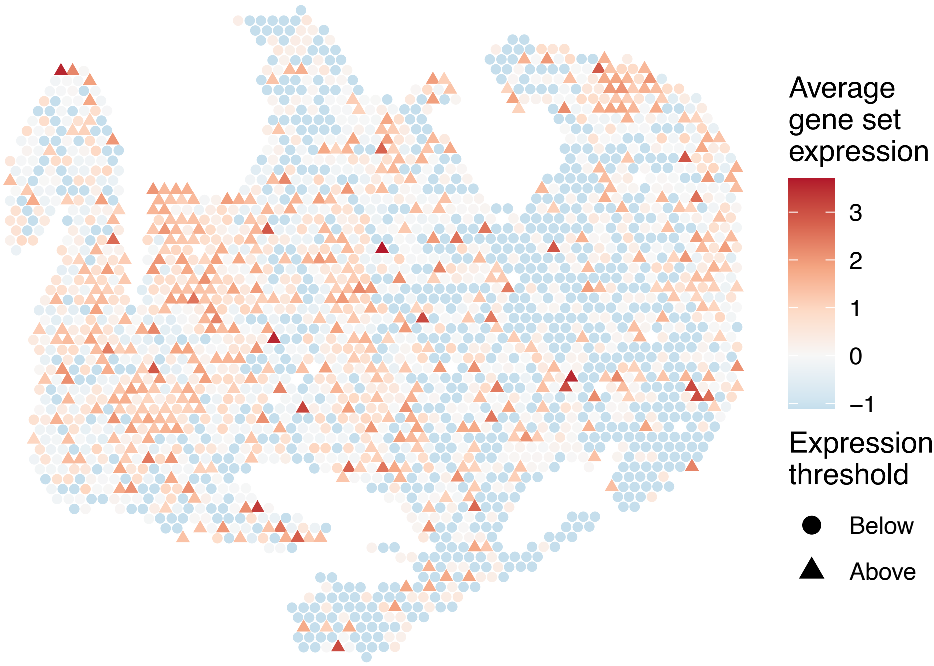

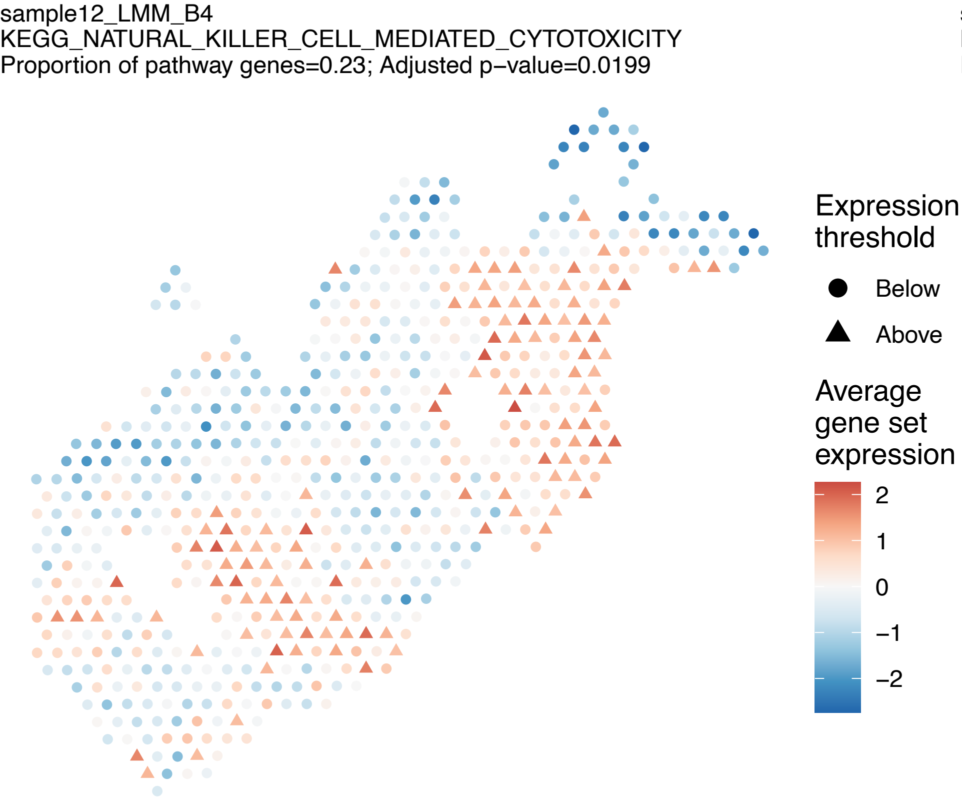
sample12_LMM_B4
KEGG_NATURAL_KILLER_CELL_MEDIATED_CYTOTOXICITY
Proportion of pathway genes=0.23; Adjusted p−value=0.0199
Expression threshold
Below
Above
Average gene set expression
2
1
0
−1
−2

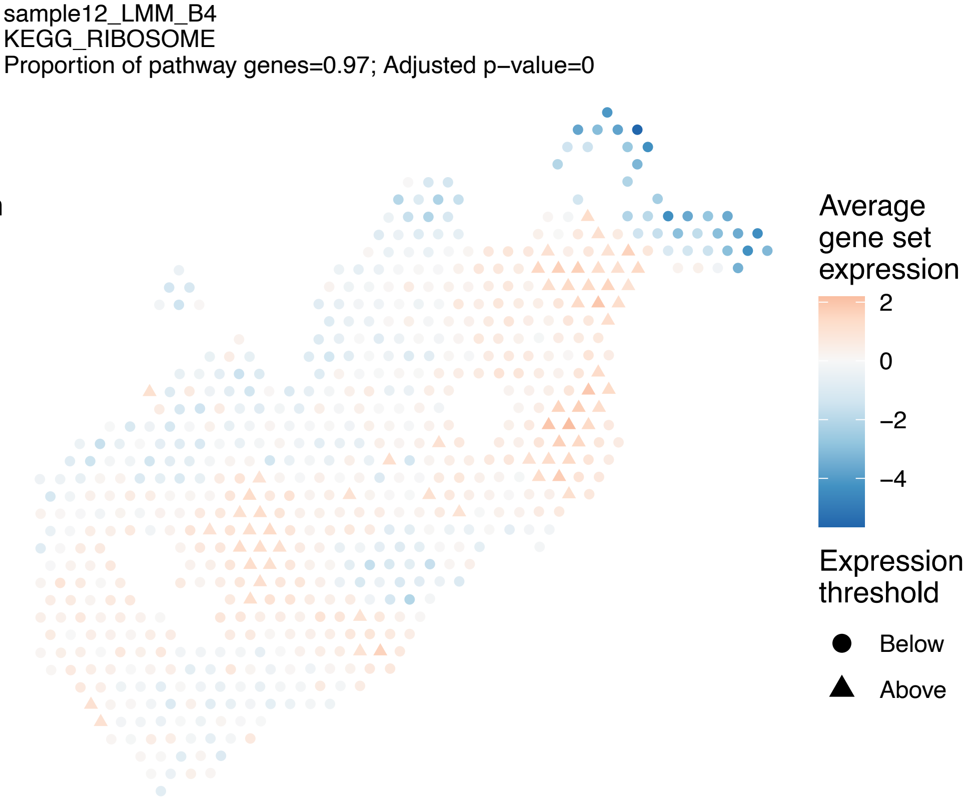
sample12_LMM_B4
KEGG_RIBOSOME
Proportion of pathway genes=0.97; Adjusted p−value=0
Average gene set expression
2
0
−2
−4
Expression threshold
Below
Above

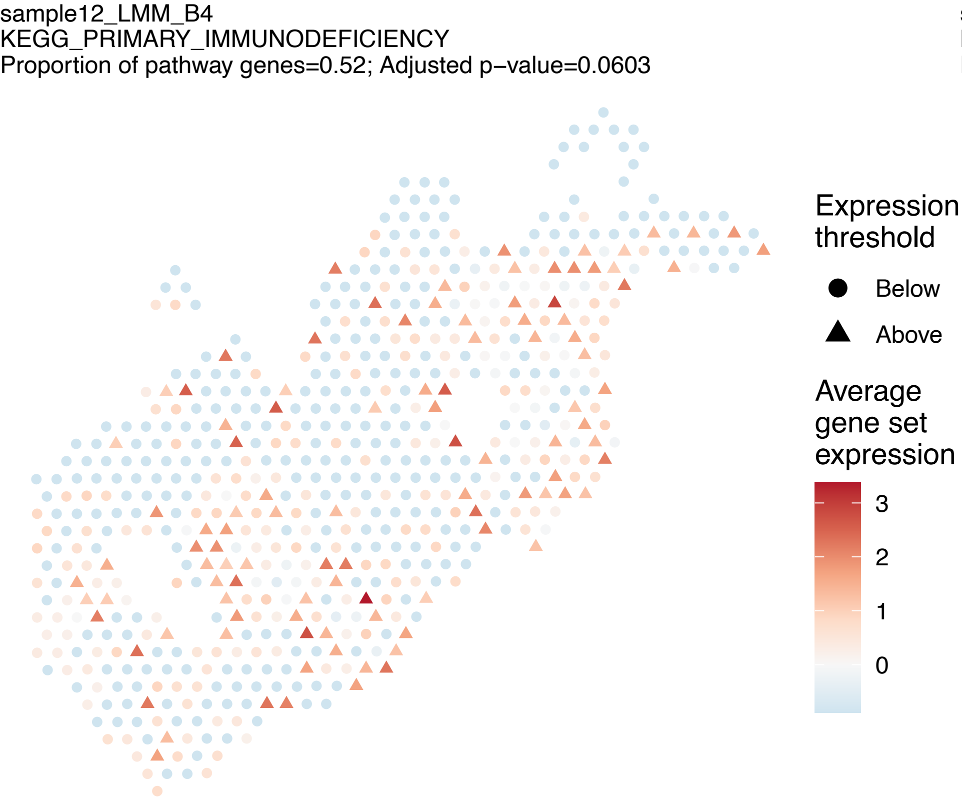
sample12_LMM_B4
KEGG_PRIMARY_IMMUNODEFICIENCY
Proportion of pathway genes=0.52; Adjusted p−value=0.0603
Expression threshold
Below
Above
Average gene set expression
3
2
1
0

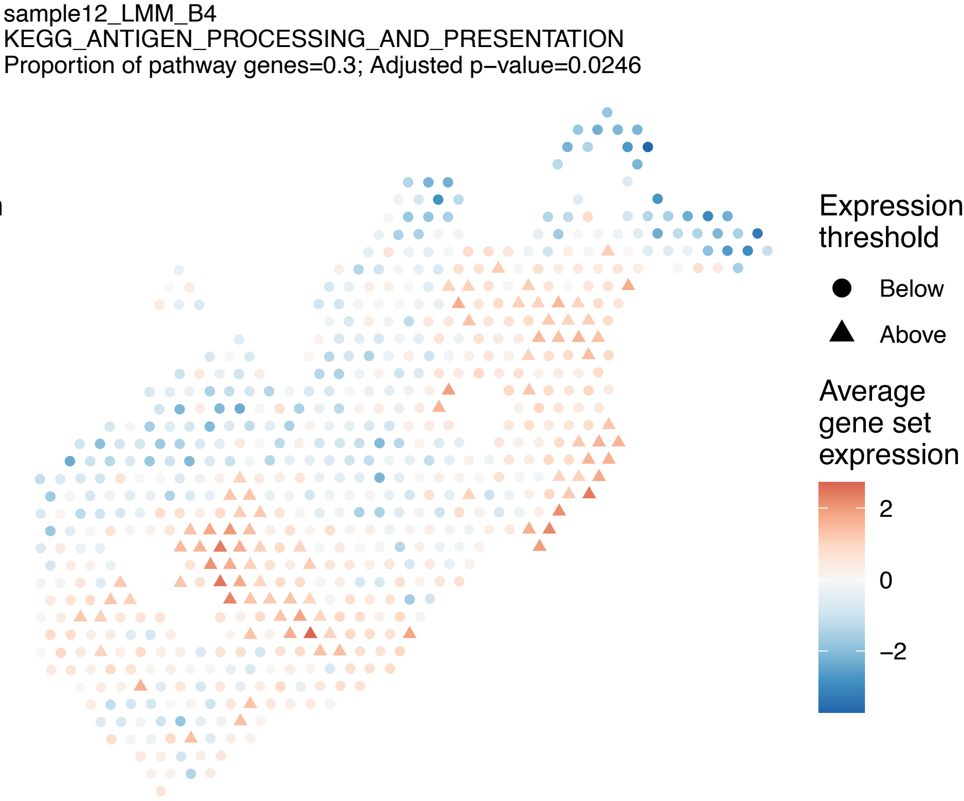
sample12_LMM_B4
KEGG_ANTIGEN_PROCESSING_AND_PRESENTATION
Proportion of pathway genes=0.3; Adjusted p−value=0.0246
Expression threshold
Below
Above
Average gene set expression
2
0
−2

sample12_LMM_B4
KEGG_CYTOKINE_CYTOKINE_RECEPTOR_INTERACTION
Proportion of pathway genes=0.28; Adjusted p−value=0

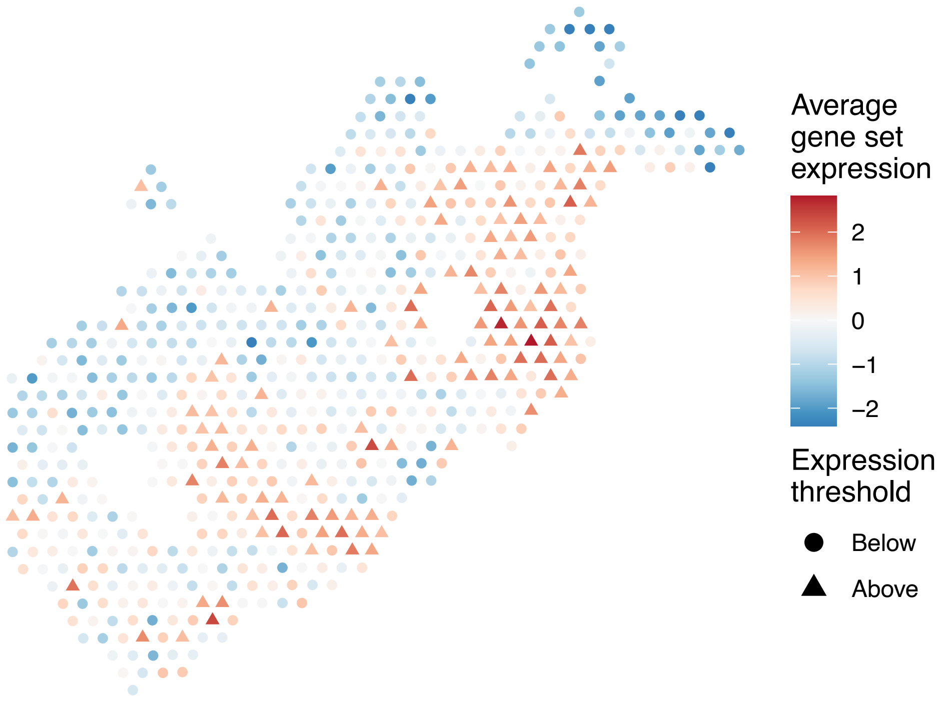


sample12_LMM_B4
KEGG_ABC_TRANSPORTERS
Proportion of pathway genes=0.67; Adjusted p−value=0.0327

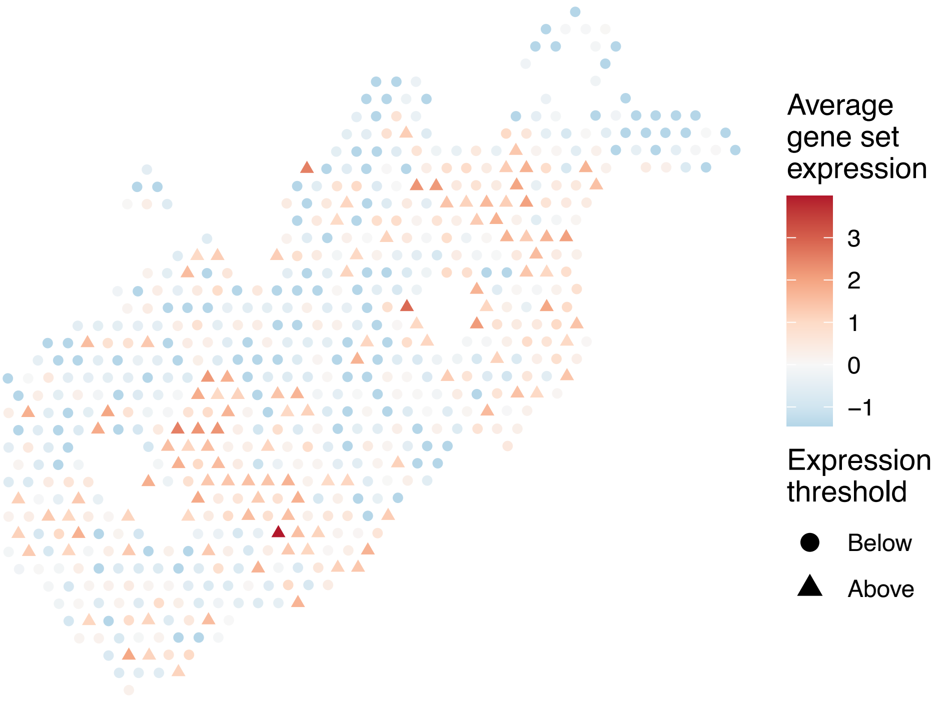


sample12_LMM_B4
KEGG_ACUTE_MYELOID_LEUKEMIA
Proportion of pathway genes=0.88; Adjusted p−value=0.0158

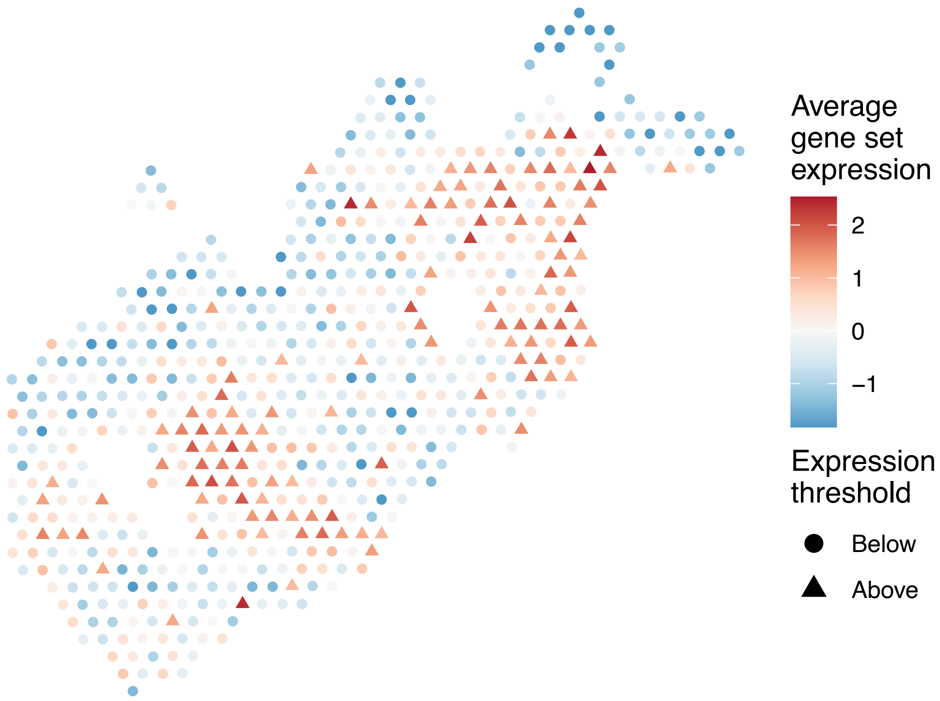


sample12_LMM_B4
KEGG_ADHERENS_JUNCTION
Proportion of pathway genes=0.83; Adjusted p−value=0.2845

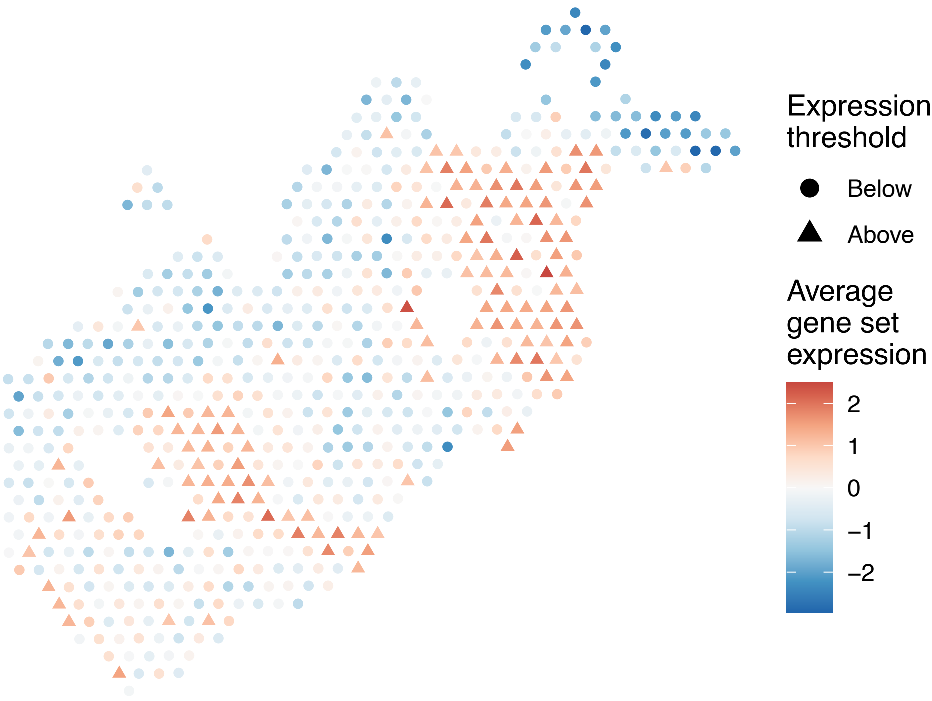

sample12_LMM_B4
KEGG_ADIPOCYTOKINE_SIGNALING_PATHWAY
Proportion of pathway genes=0.69; Adjusted p−value=0.0104

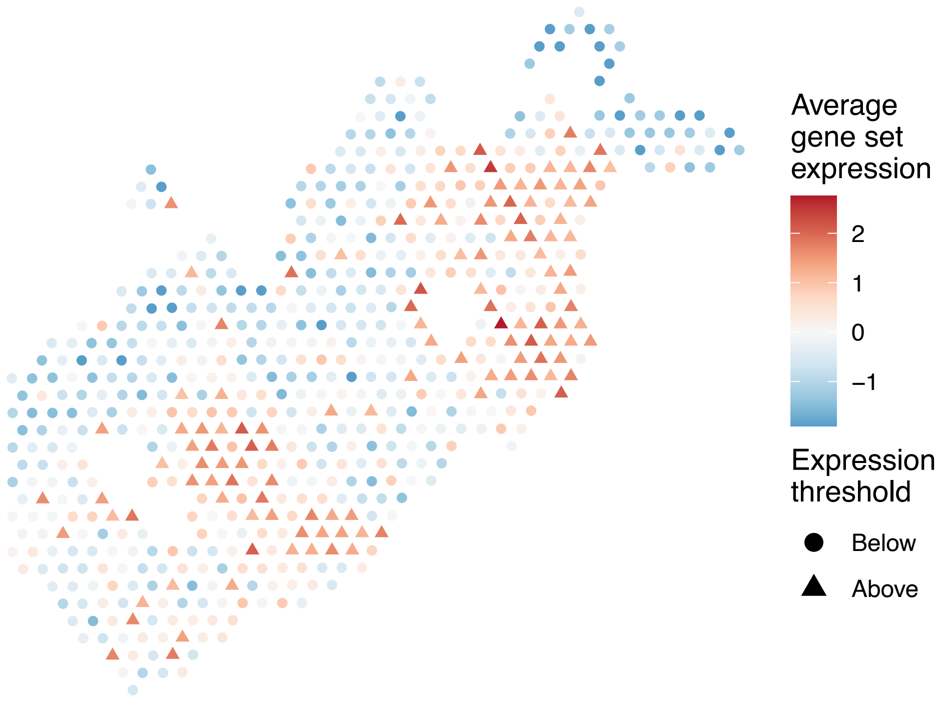


sample12_LMM_B4
KEGG_ALANINE_ASPARTATE_AND_GLUTAMATE_METABOLISM
Proportion of pathway genes=0.55; Adjusted p−value=0.0546

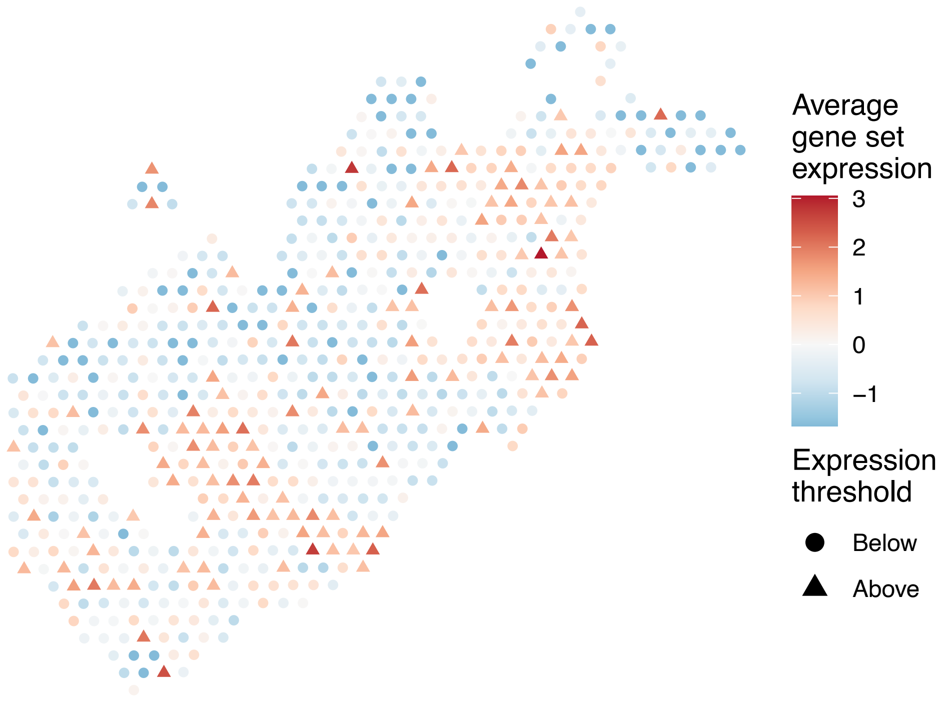


sample12_LMM_B4
KEGG_ALDOSTERONE_REGULATED_SODIUM_REABSORPTION
Proportion of pathway genes=0.52; Adjusted p−value=0.0607

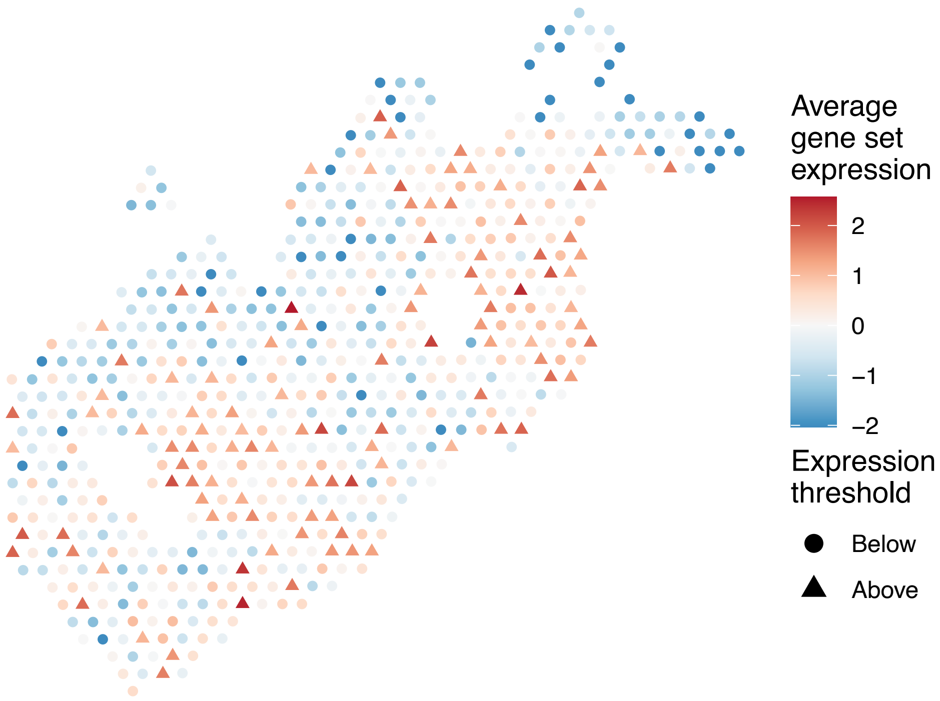


sample12_LMM_B4
KEGG_ALLOGRAFT_REJECTION
Proportion of pathway genes=0.62; Adjusted p−value=0.0236

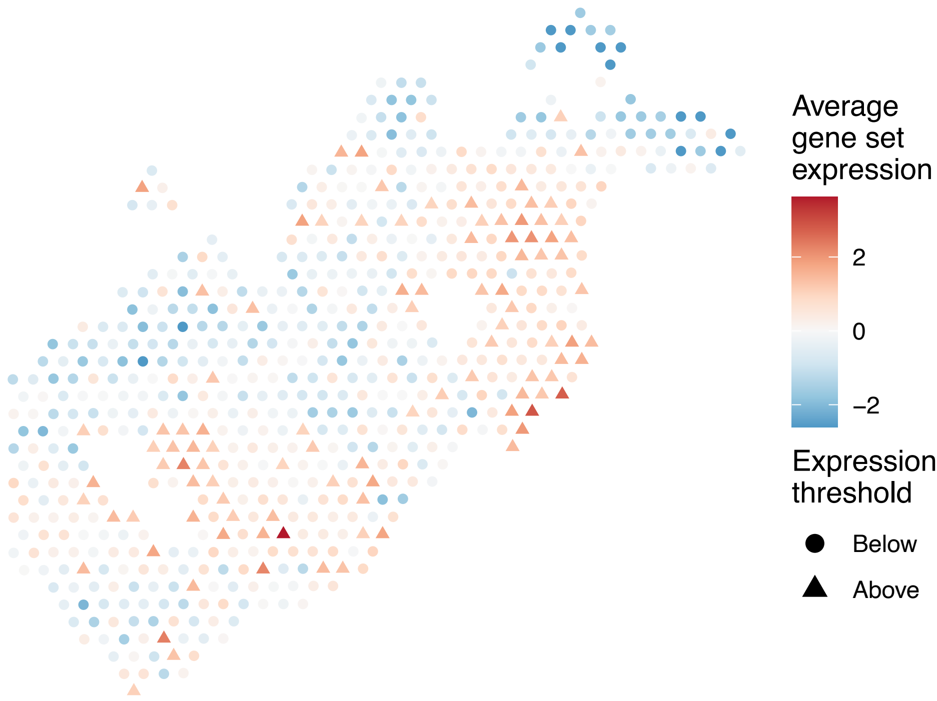

sample12_LMM_B4
KEGG_ALZHEIMERS_DISEASE
Proportion of pathway genes=0.8; Adjusted p−value=0.0335

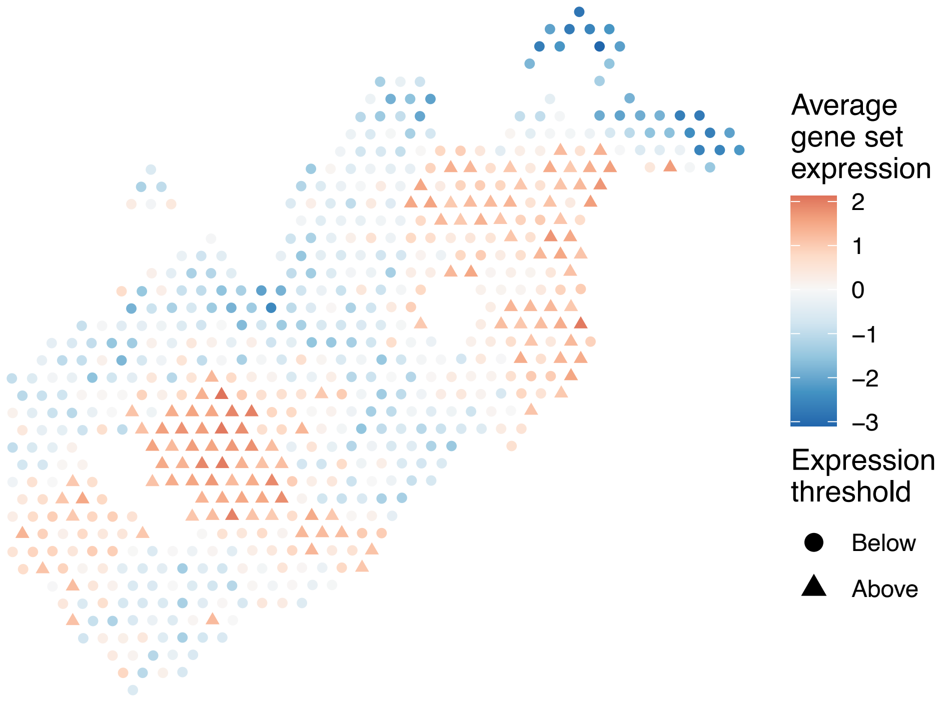


sample12_LMM_B4
KEGG_AMINO_SUGAR_AND_NUCLEOTIDE_SUGAR_METABOLISM
Proportion of pathway genes=0.85; Adjusted p−value=0.0048

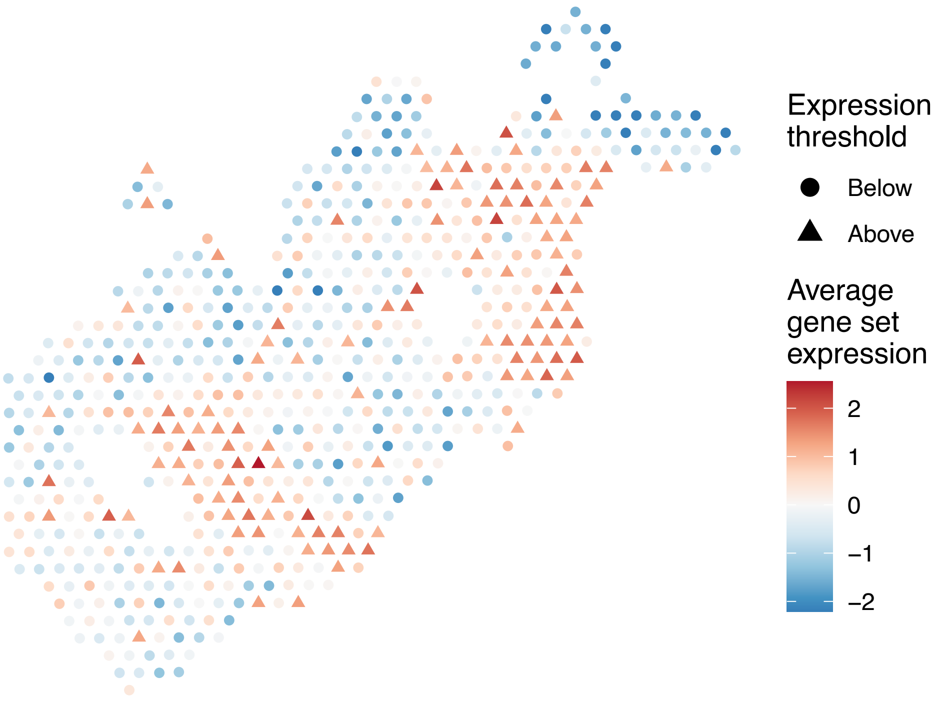


sample12_LMM_B4
KEGG_AMINOACYL_TRNA_BIOSYNTHESIS
Proportion of pathway genes=0.51; Adjusted p−value=0.2821

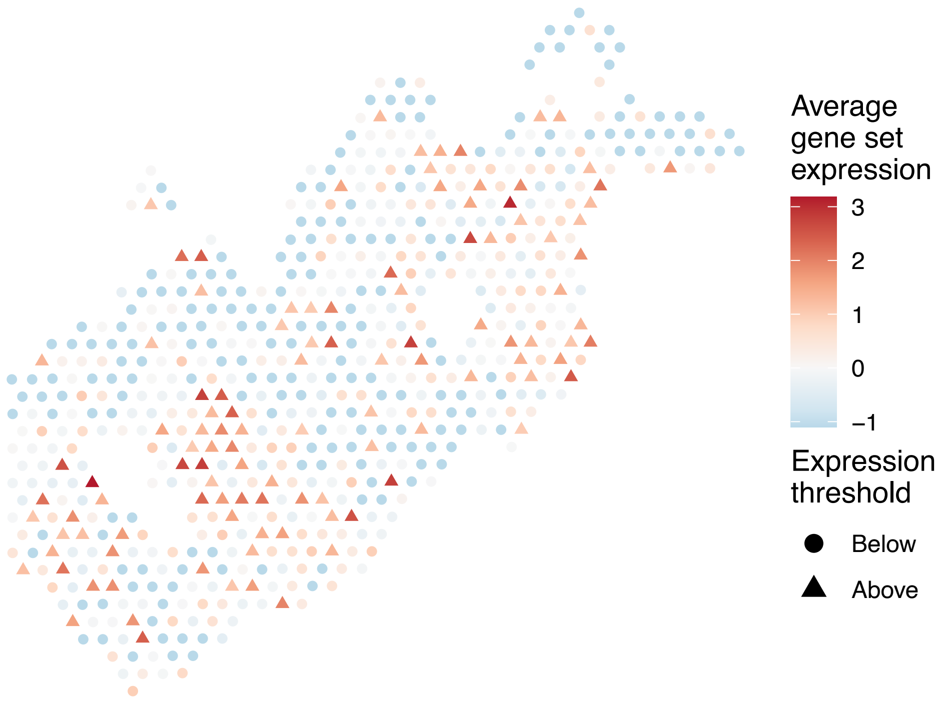


sample12_LMM_B4
KEGG_AMYOTROPHIC_LATERAL_SCLEROSIS_ALS
Proportion of pathway genes=0.62; Adjusted p−value=0.1219

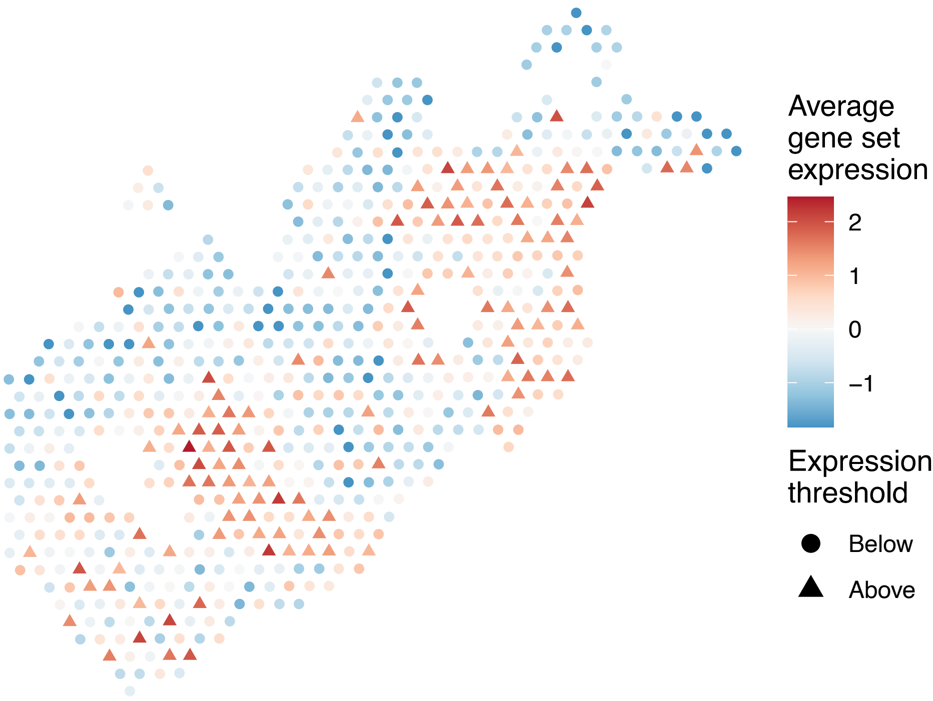

sample12_LMM_B4
KEGG_APOPTOSIS
Proportion of pathway genes=0.68; Adjusted p−value=0

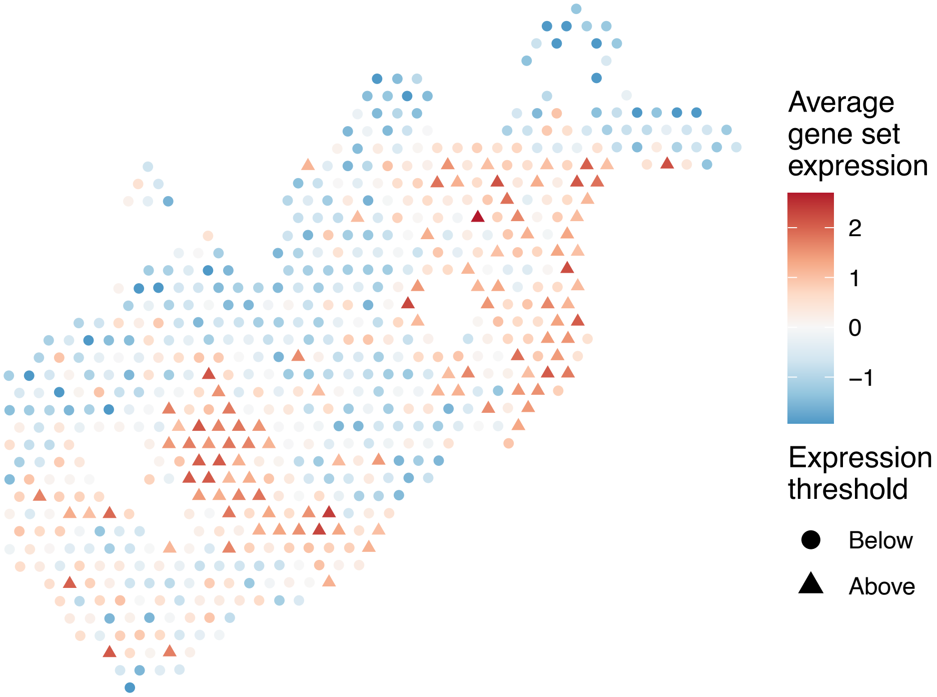


sample12_LMM_B4
KEGG_ARACHIDONIC_ACID_METABOLISM
Proportion of pathway genes=0.3; Adjusted p−value=0.1406

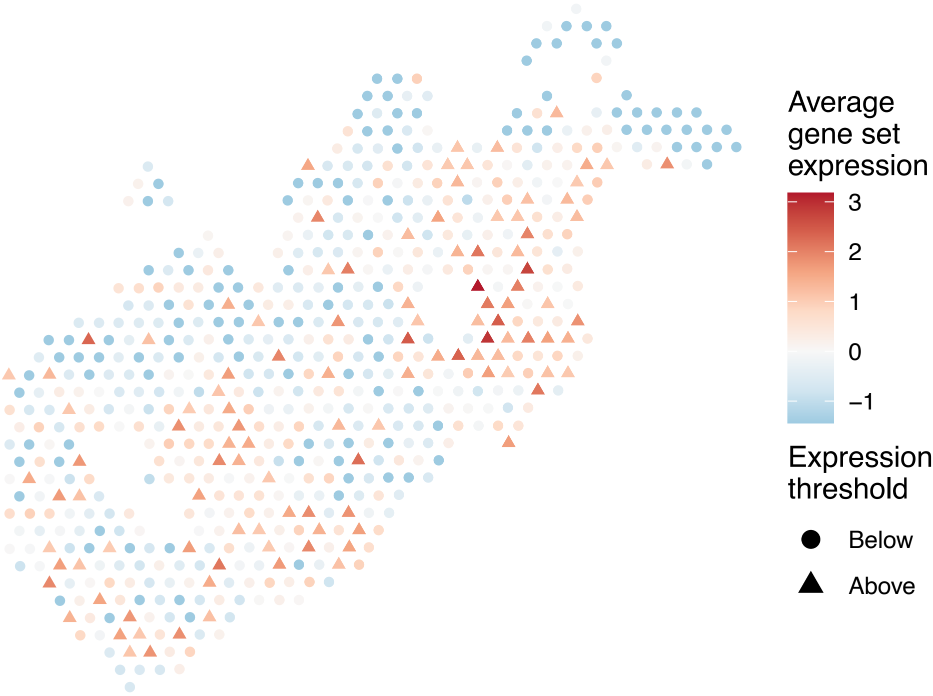


sample12_LMM_B4
KEGG_ARGININE_AND_PROLINE_METABOLISM
Proportion of pathway genes=0.62; Adjusted p−value=0.0826

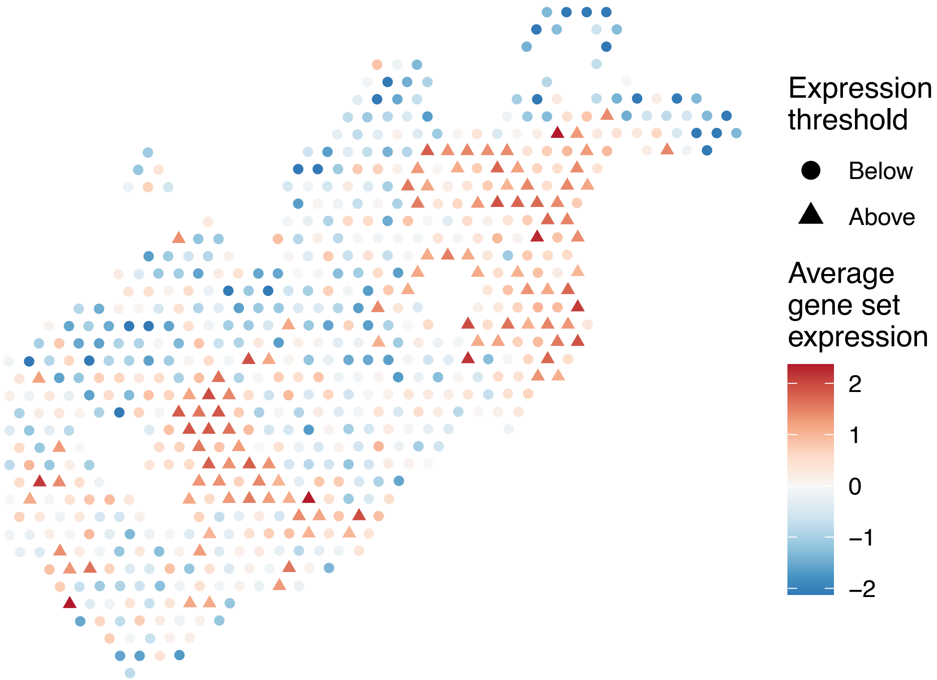


sample12_LMM_B4
KEGG_ARRHYTHMOGENIC_RIGHT_VENTRICULAR_CARDIOMYOPAT
HY_ARVC
Proportion of pathway genes=0.51; Adjusted p−value=0.019

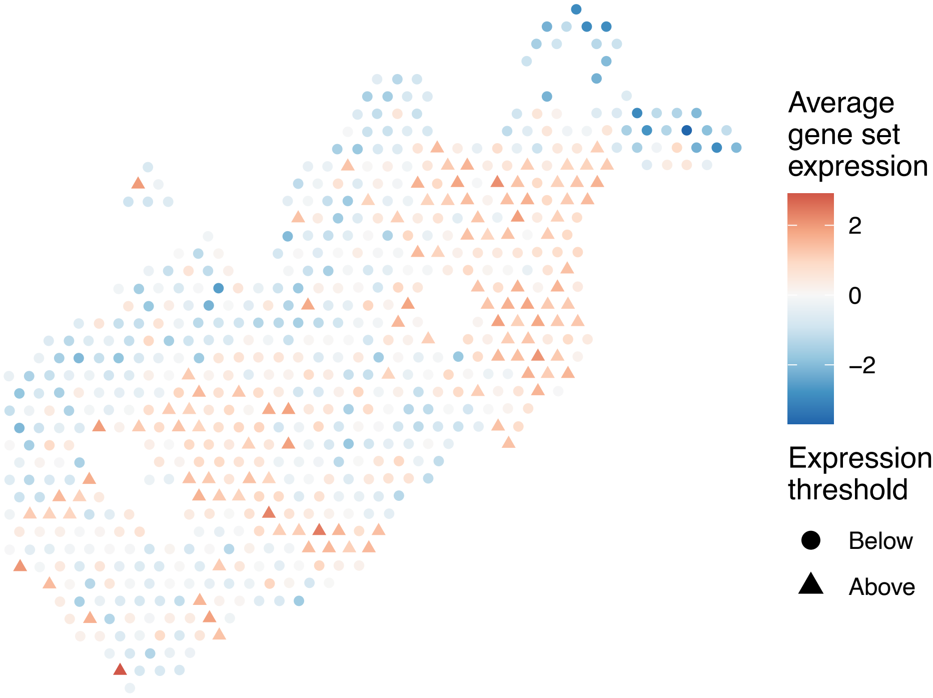

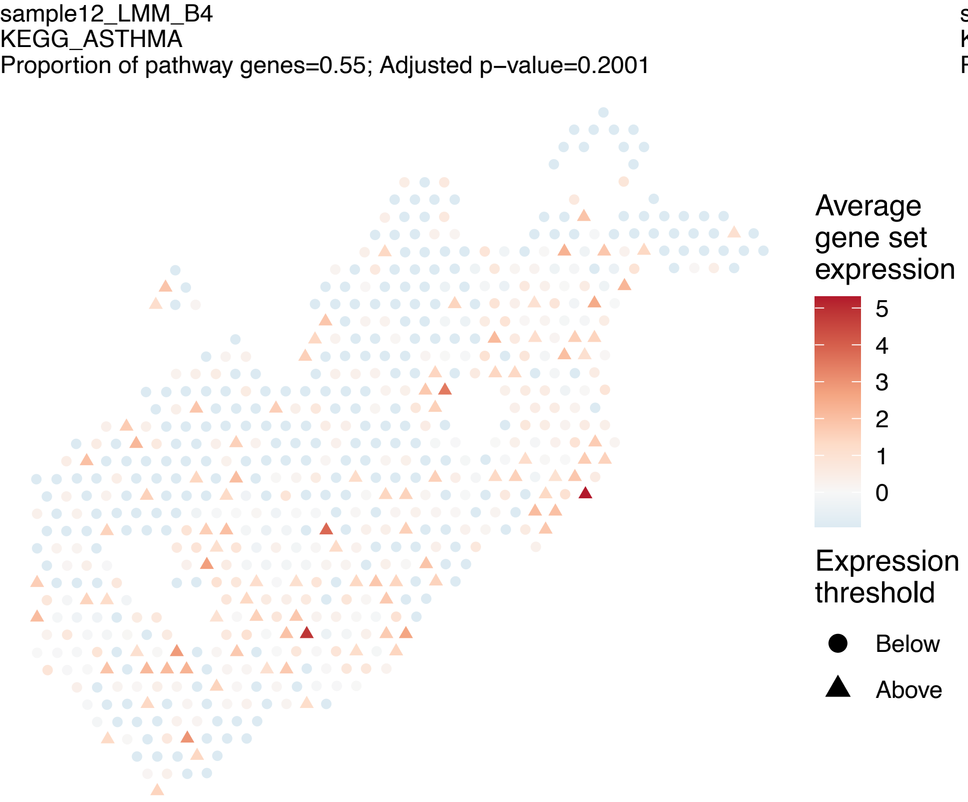
sample12_LMM_B4
KEGG_ASTHMA
Proportion of pathway genes=0.55; Adjusted p−value=0.2001
Average gene set expression
5
4
3
2
1
0
Expression threshold
Below
Above

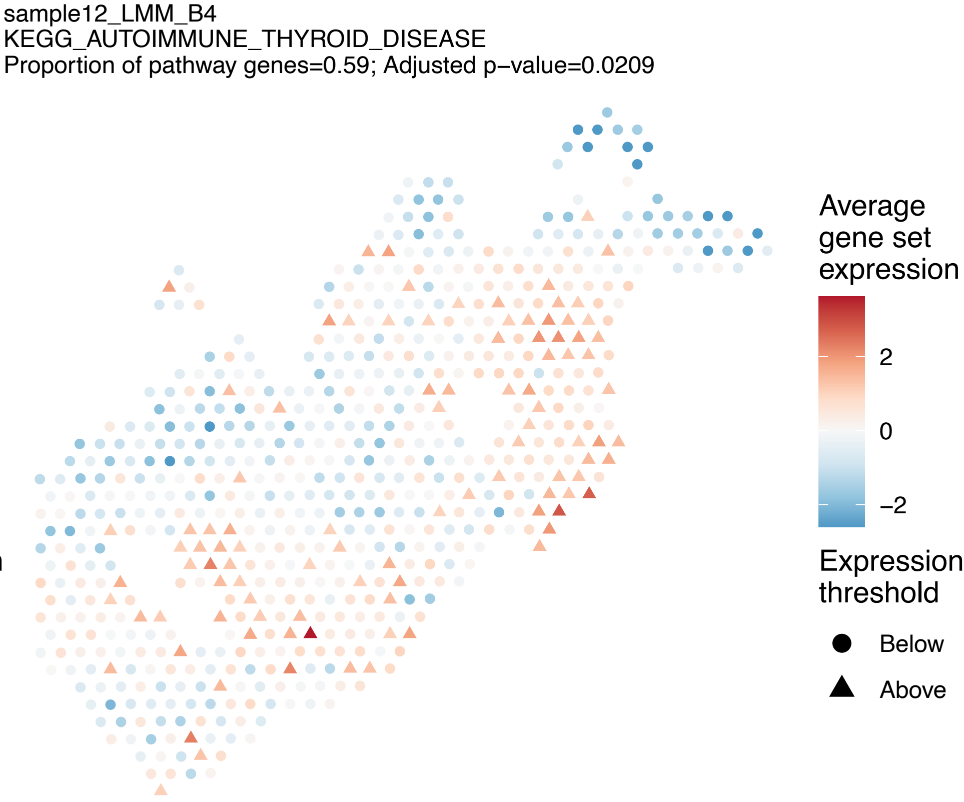
sample12_LMM_B4
KEGG_AUTOIMMUNE_THYROID_DISEASE
Proportion of pathway genes=0.59; Adjusted p−value=0.0209
Average gene set expression
2
0
−2
Expression threshold
Below
Above

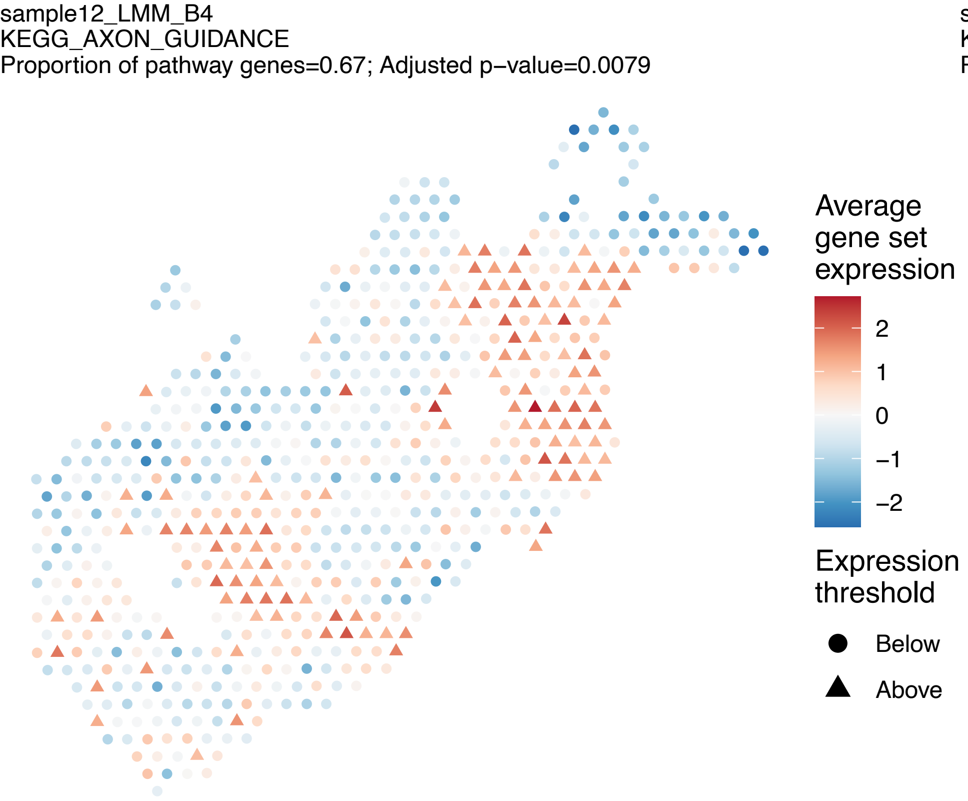
sample12_LMM_B4
KEGG_AXON_GUIDANCE
Proportion of pathway genes=0.67; Adjusted p−value=0.0079
Average gene set expression
2
1
0
−1
−2
Expression threshold
Below
Above

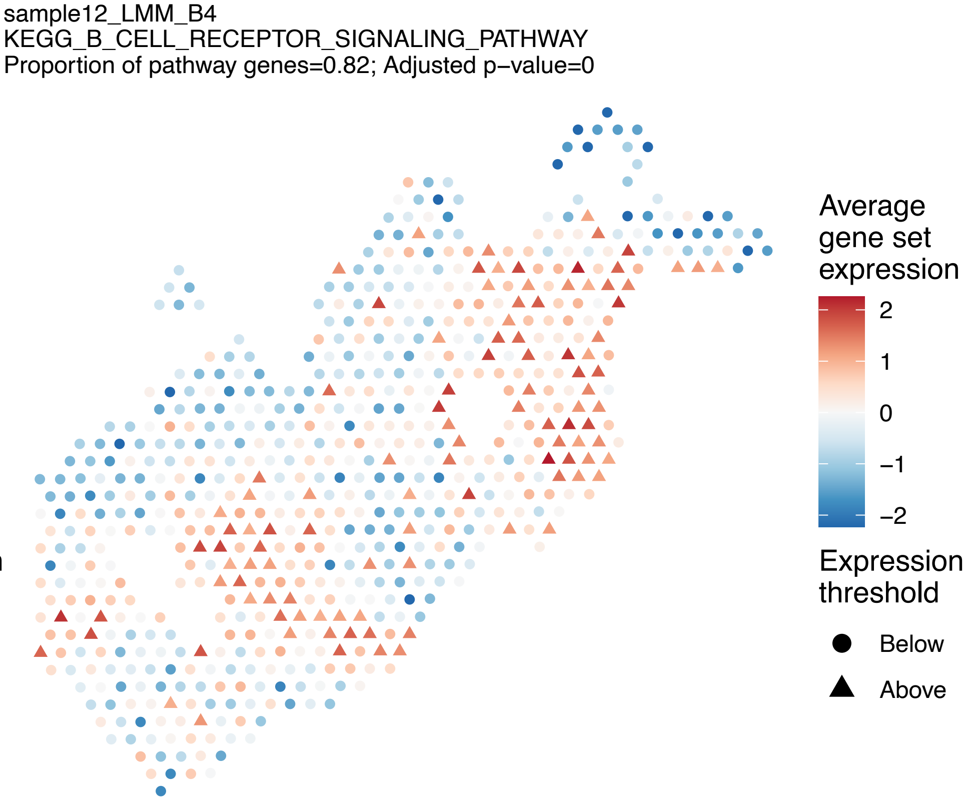
sample12_LMM_B4
KEGG_B_CELL_RECEPTOR_SIGNALING_PATHWAY
Proportion of pathway genes=0.82; Adjusted p−value=0
Average gene set expression
2
1
0
−1
−2
Expression threshold
Below
Above

sample12_LMM_B4
KEGG_BASAL_CELL_CARCINOMA
Proportion of pathway genes=0.48; Adjusted p−value=0.2432

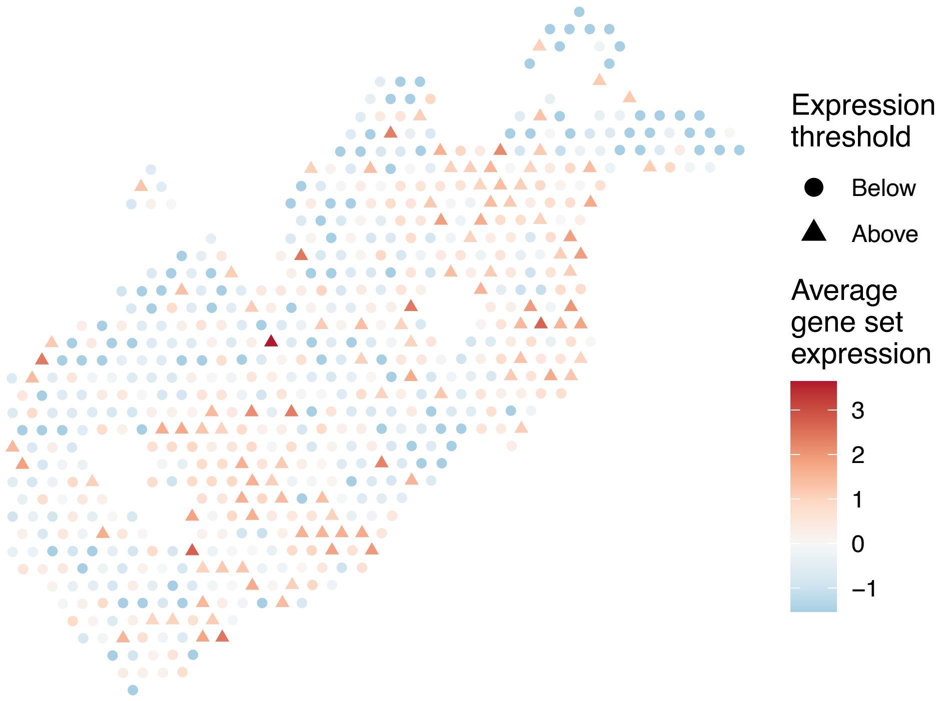


sample12_LMM_B4
KEGG_BASAL_TRANSCRIPTION_FACTORS
Proportion of pathway genes=0.91; Adjusted p−value=0.0129

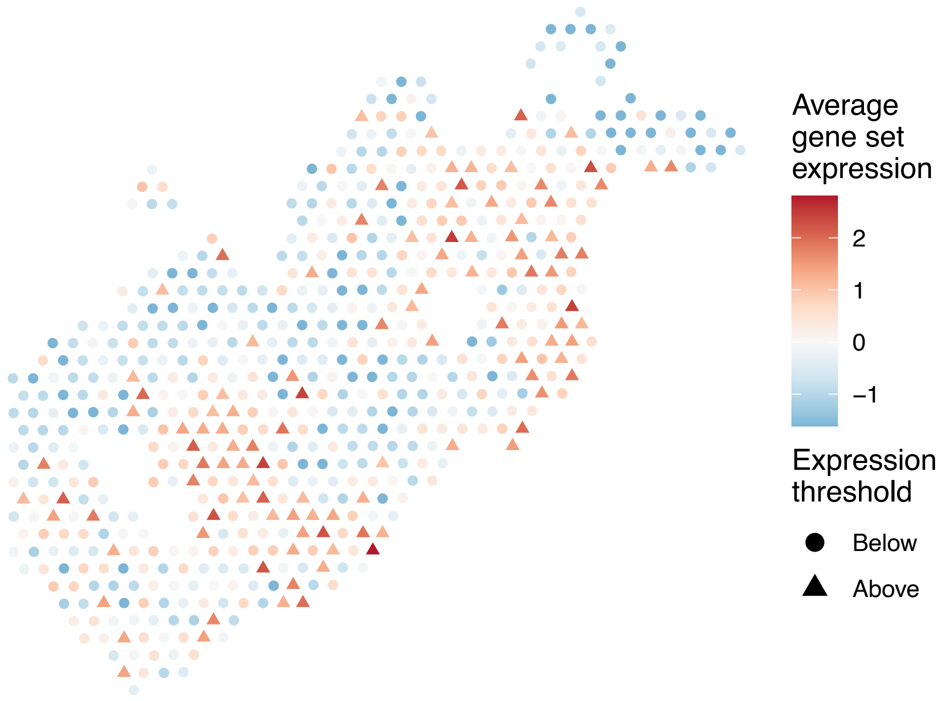


sample12_LMM_B4
KEGG_BASE_EXCISION_REPAIR
Proportion of pathway genes=0.91; Adjusted p−value=0.0199

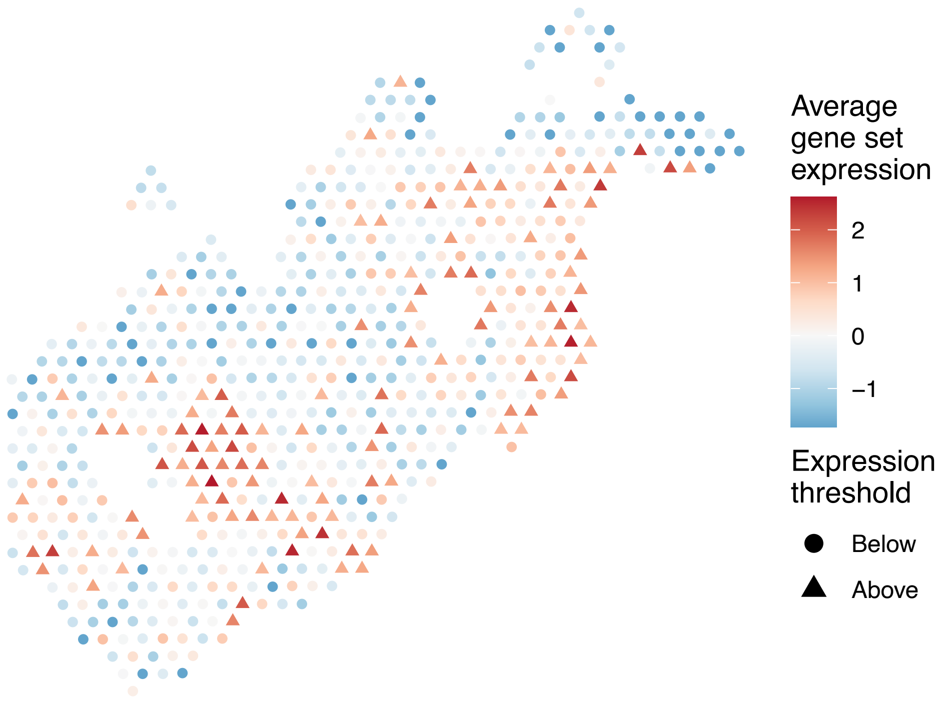


sample12_LMM_B4
KEGG_BETA_ALANINE_METABOLISM
Proportion of pathway genes=0.59; Adjusted p−value=0.0048

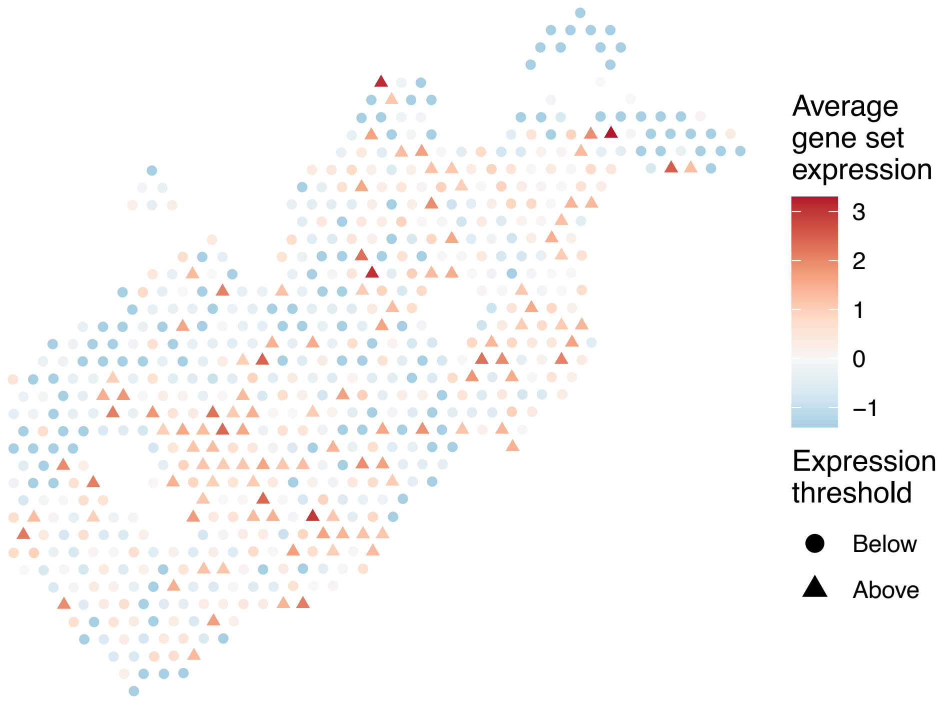

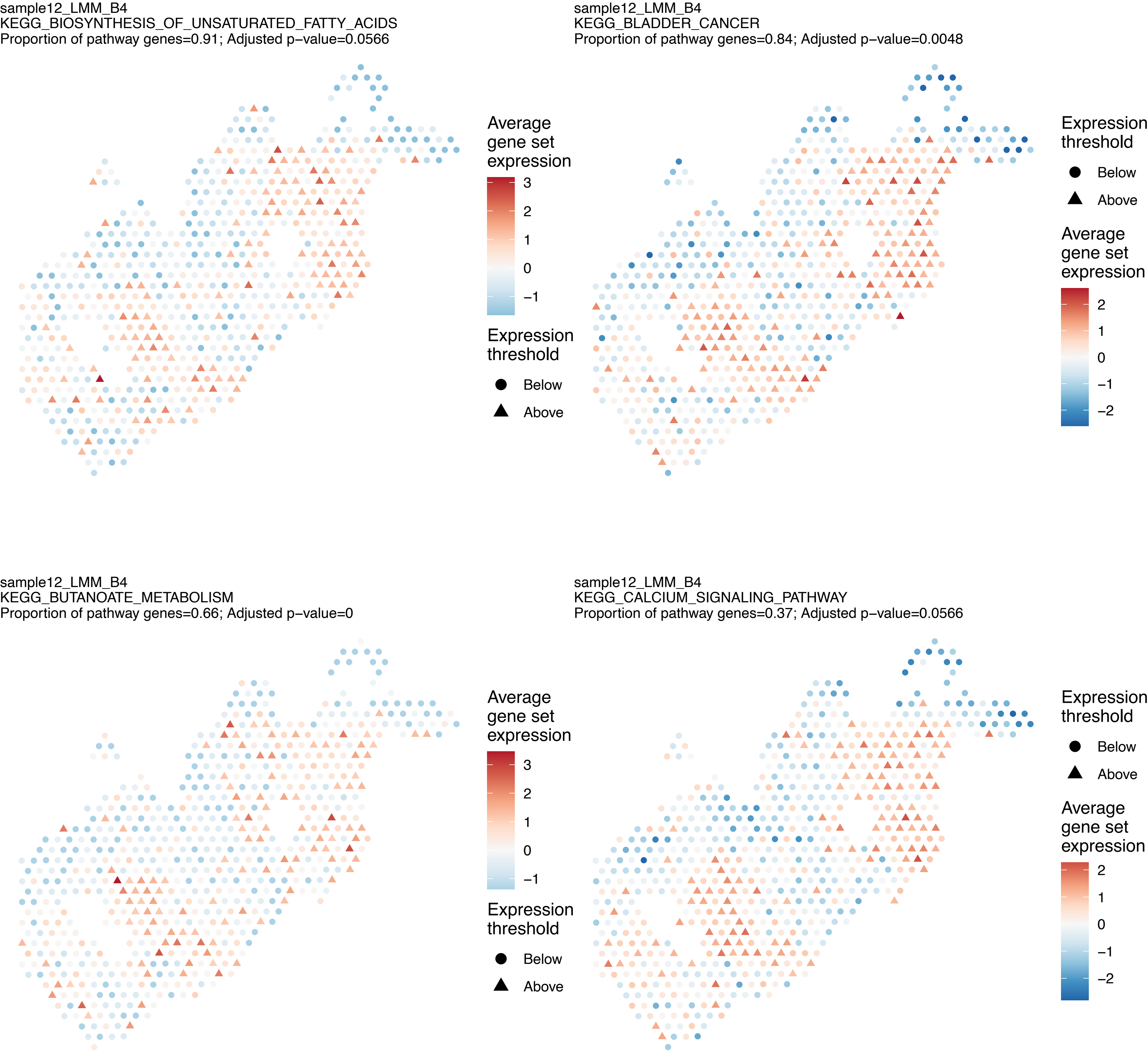
sample12_LMM_B4
KEGG_BIOSYNTHESIS_OF_UNSATURATED_FATTY_ACIDS
Proportion of pathway genes=0.91; Adjusted p−value=0.0566
Average gene set expression
3
2
1
0
−1
Expression threshold
Below
Above
sample12_LMM_B4
KEGG_BLADDER_CANCER
Proportion of pathway genes=0.84; Adjusted p−value=0.0048
Expression threshold
Below
Above
Average gene set expression
2
1
0
−1
−2
sample12_LMM_B4
KEGG_BUTANOATE_METABOLISM
Proportion of pathway genes=0.66; Adjusted p−value=0
Average gene set expression
3
2
1
0
−1
Expression threshold
Below
Above
sample12_LMM_B4
KEGG_CALCIUM_SIGNALING_PATHWAY
Proportion of pathway genes=0.37; Adjusted p−value=0.0566
Expression threshold
Below
Above
Average gene set expression
2
1
0
−1
−2

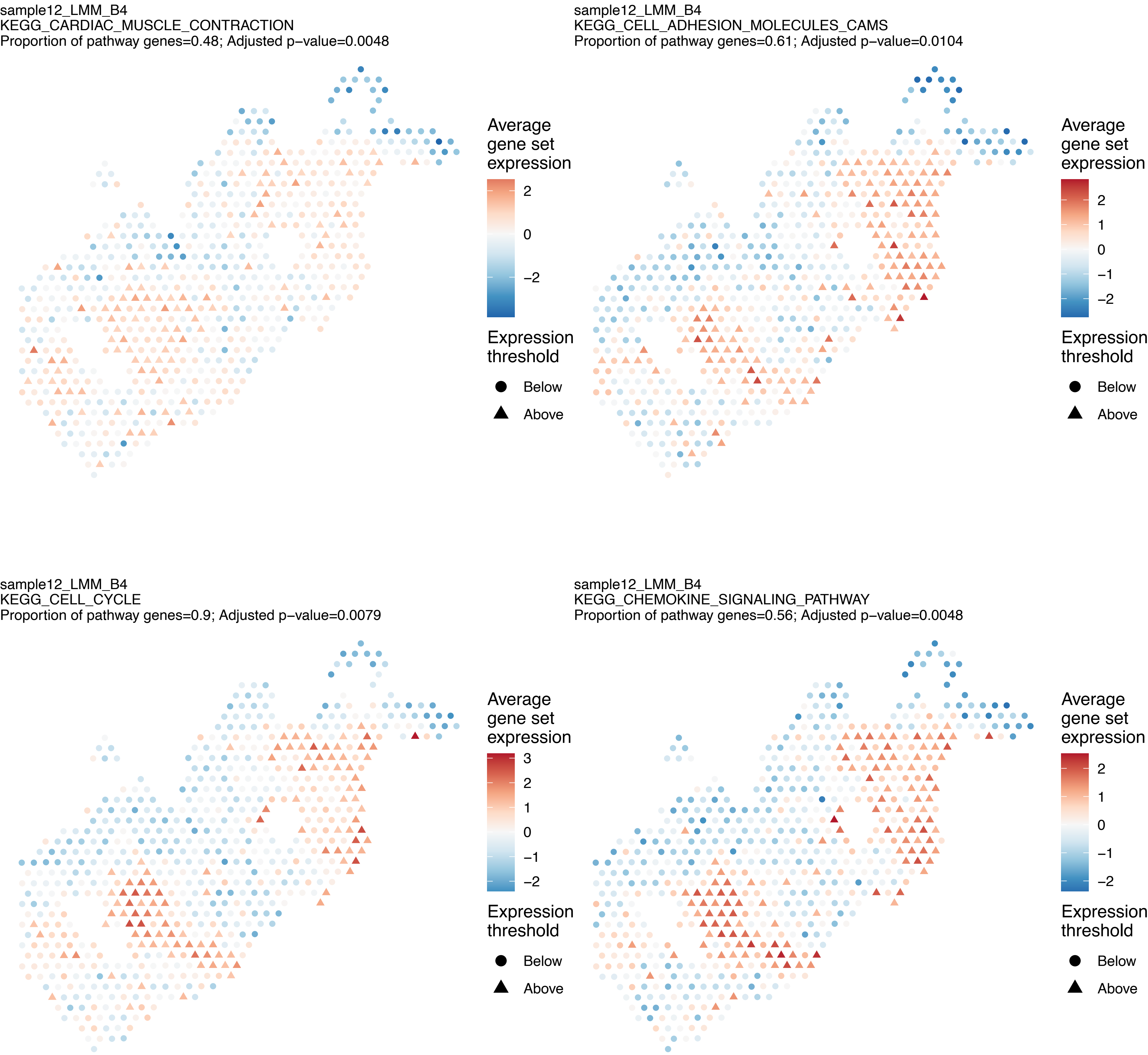

sample12_LMM_B4
KEGG_CARDIAC_MUSCLE_CONTRACTION
Proportion of pathway genes=0.48; Adjusted p−value=0.0048
Average gene set expression
2
0
−2
Expression threshold
Below
Above
sample12_LMM_B4
KEGG_CELL_ADHESION_MOLECULES_CAMS
Proportion of pathway genes=0.61; Adjusted p−value=0.0104
Average gene set expression
2
1
0
−1
−2
Expression threshold
Below
Above
sample12_LMM_B4
KEGG_CELL_CYCLE
Proportion of pathway genes=0.9; Adjusted p−value=0.0079
Average gene set expression
3
2
1
0
−1
−2
Expression threshold
Below
Above
sample12_LMM_B4
KEGG_CHEMOKINE_SIGNALING_PATHWAY
Proportion of pathway genes=0.56; Adjusted p−value=0.0048
Average gene set expression
2
1
0
−1
−2
Expression threshold
Below
Above

sample12_LMM_B4
KEGG_CHRONIC_MYELOID_LEUKEMIA
Proportion of pathway genes=0.91; Adjusted p−value=0

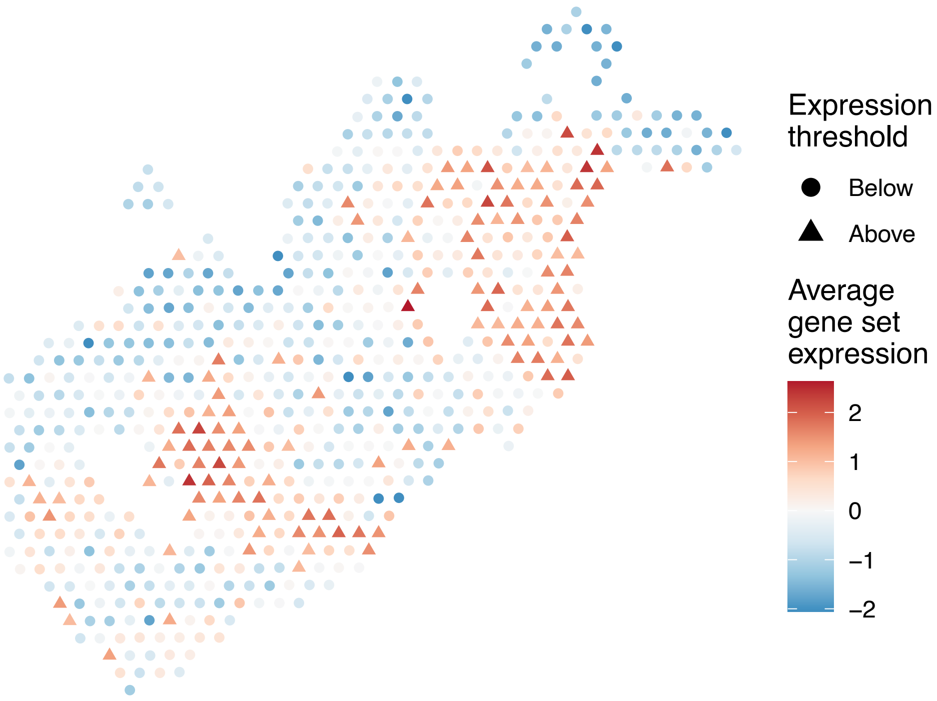


sample12_LMM_B4
KEGG_CIRCADIAN_RHYTHM_MAMMAL
Proportion of pathway genes=0.92; Adjusted p−value=0.0265

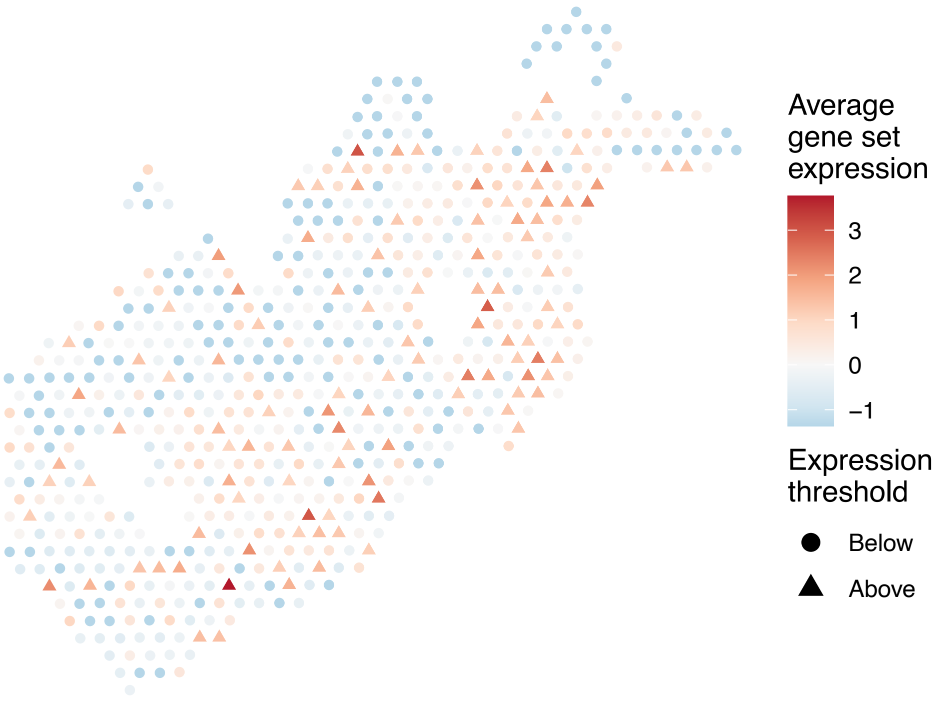


sample12_LMM_B4
KEGG_CITRATE_CYCLE_TCA_CYCLE
Proportion of pathway genes=0.88; Adjusted p−value=0.022

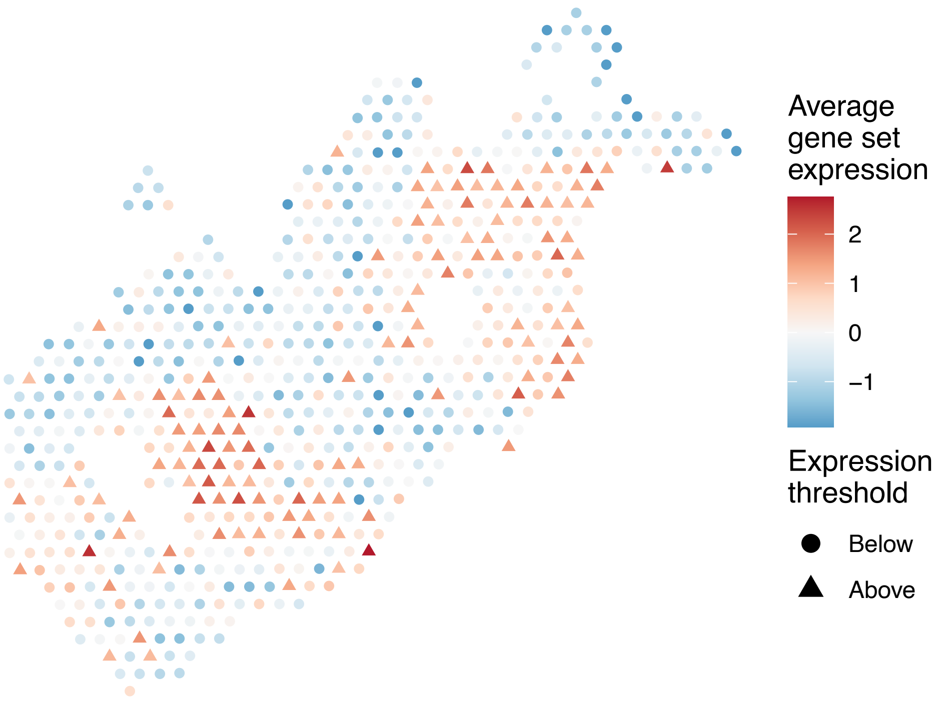


sample12_LMM_B4
KEGG_COLORECTAL_CANCER
Proportion of pathway genes=0.89; Adjusted p−value=0.0174

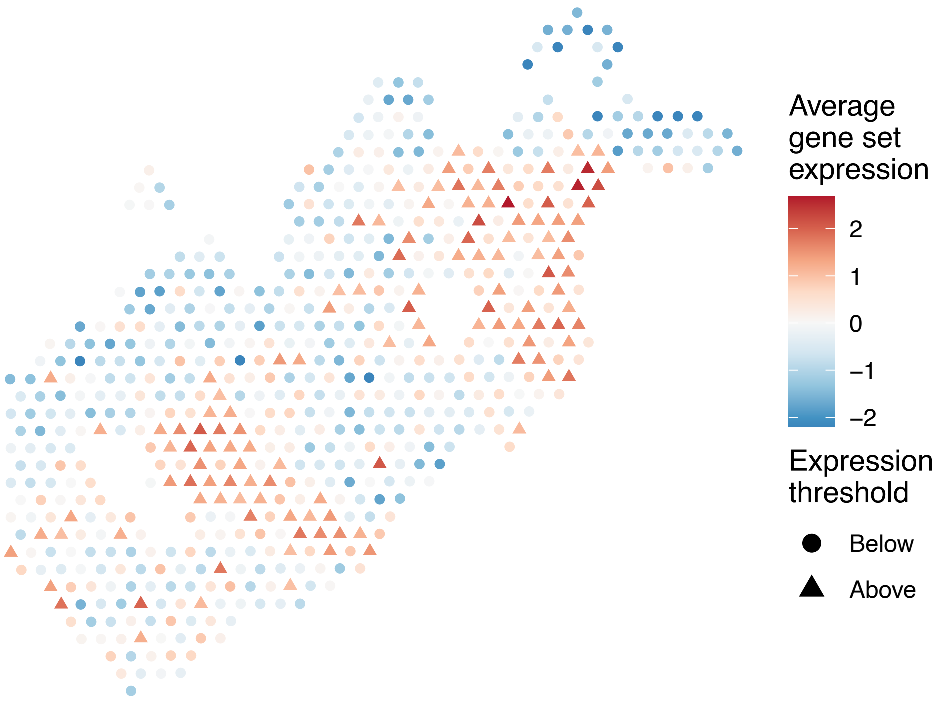

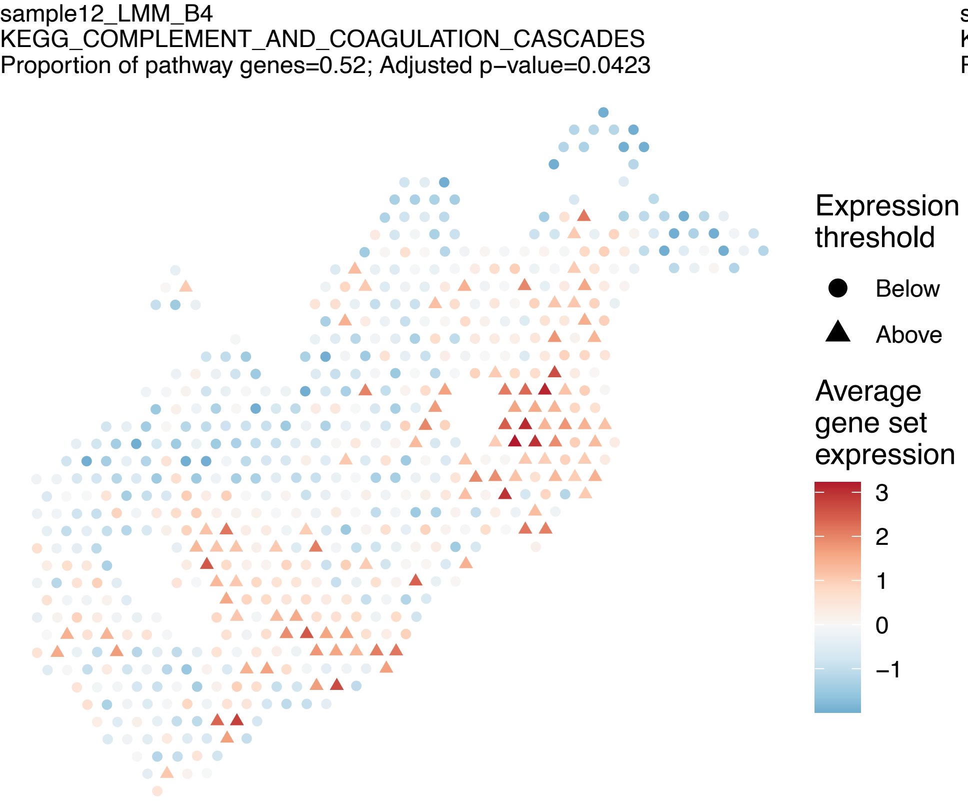
sample12_LMM_B4
KEGG_COMPLEMENT_AND_COAGULATION_CASCADES
Proportion of pathway genes=0.52; Adjusted p−value=0.0423
Expression threshold
Below
Above
Average gene set expression
3
2
1
0
−1

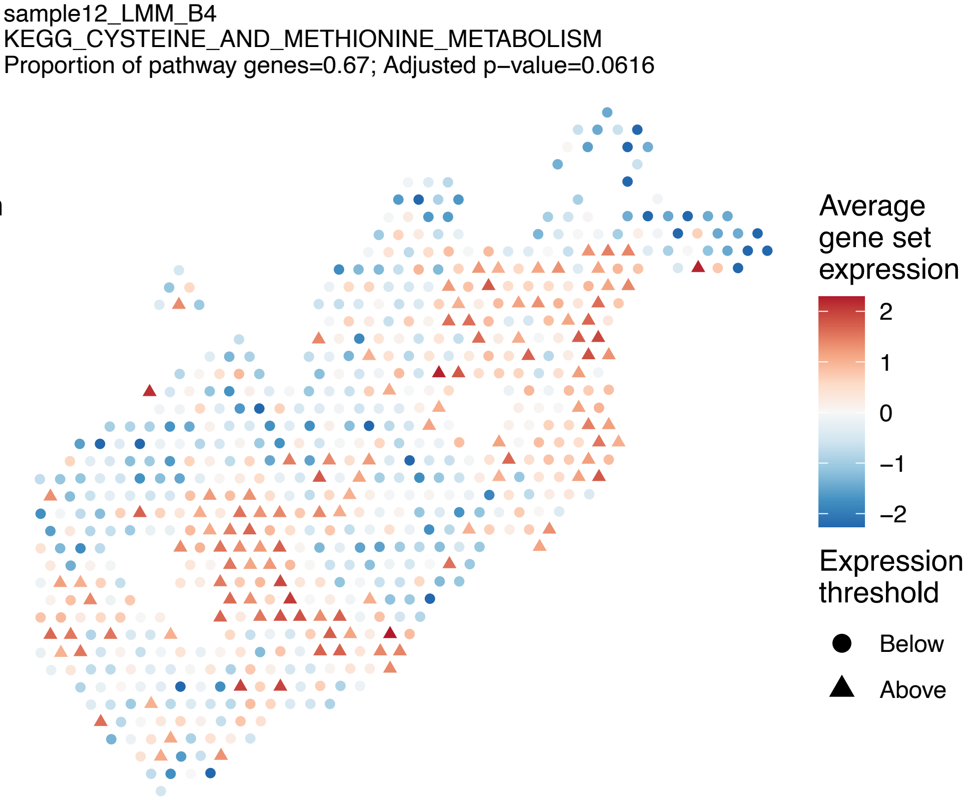
sample12_LMM_B4
KEGG_CYSTEINE_AND_METHIONINE_METABOLISM
Proportion of pathway genes=0.67; Adjusted p−value=0.0616
Average gene set expression
2
1
0
−1
−2
Expression threshold
Below
Above

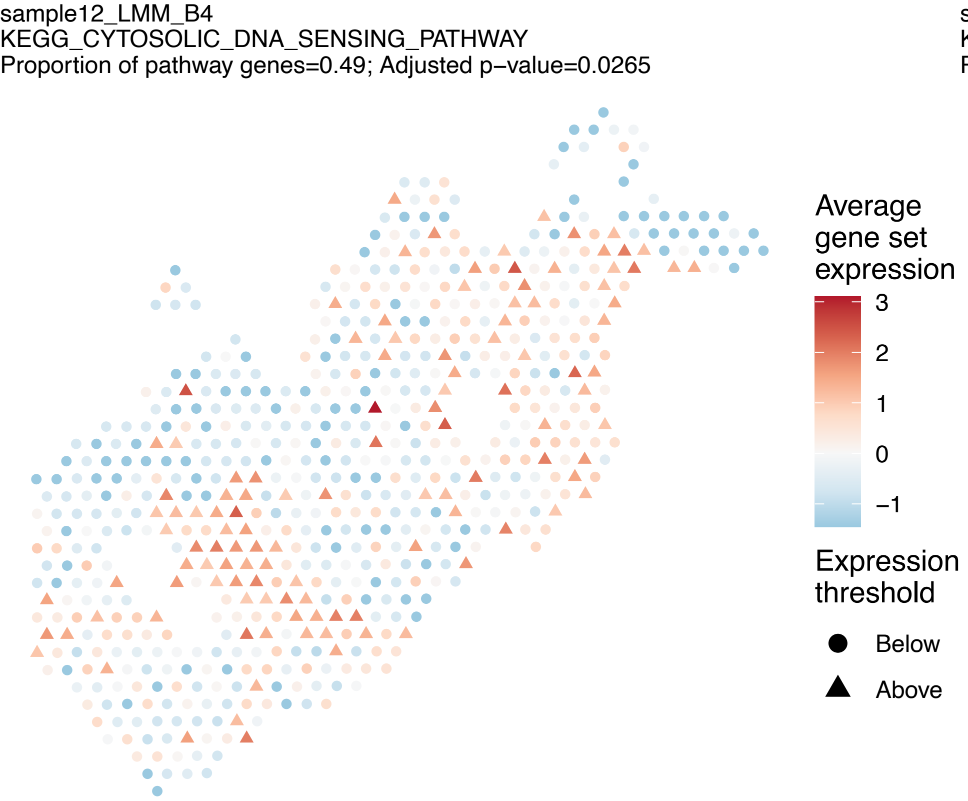
sample12_LMM_B4
KEGG_CYTOSOLIC_DNA_SENSING_PATHWAY
Proportion of pathway genes=0.49; Adjusted p−value=0.0265
Average gene set expression
3
2
1
0
−1
Expression threshold
Below
Above

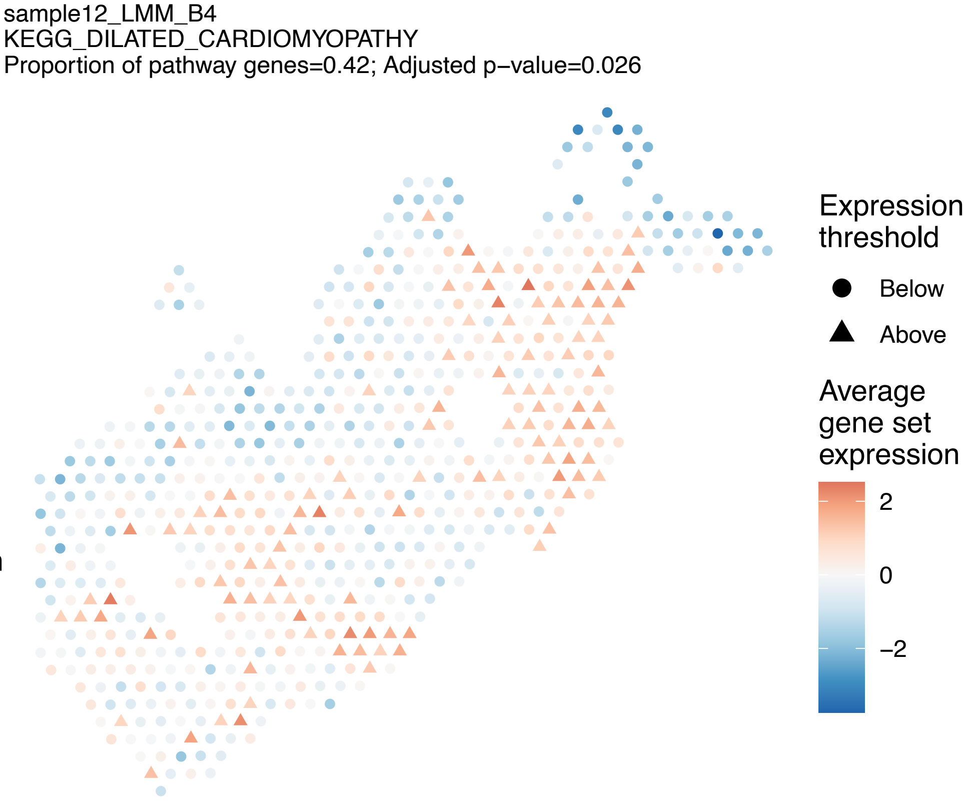
sample12_LMM_B4
KEGG_DILATED_CARDIOMYOPATHY
Proportion of pathway genes=0.42; Adjusted p−value=0.026
Expression threshold
Below
Above
Average gene set expression
2
0
−2

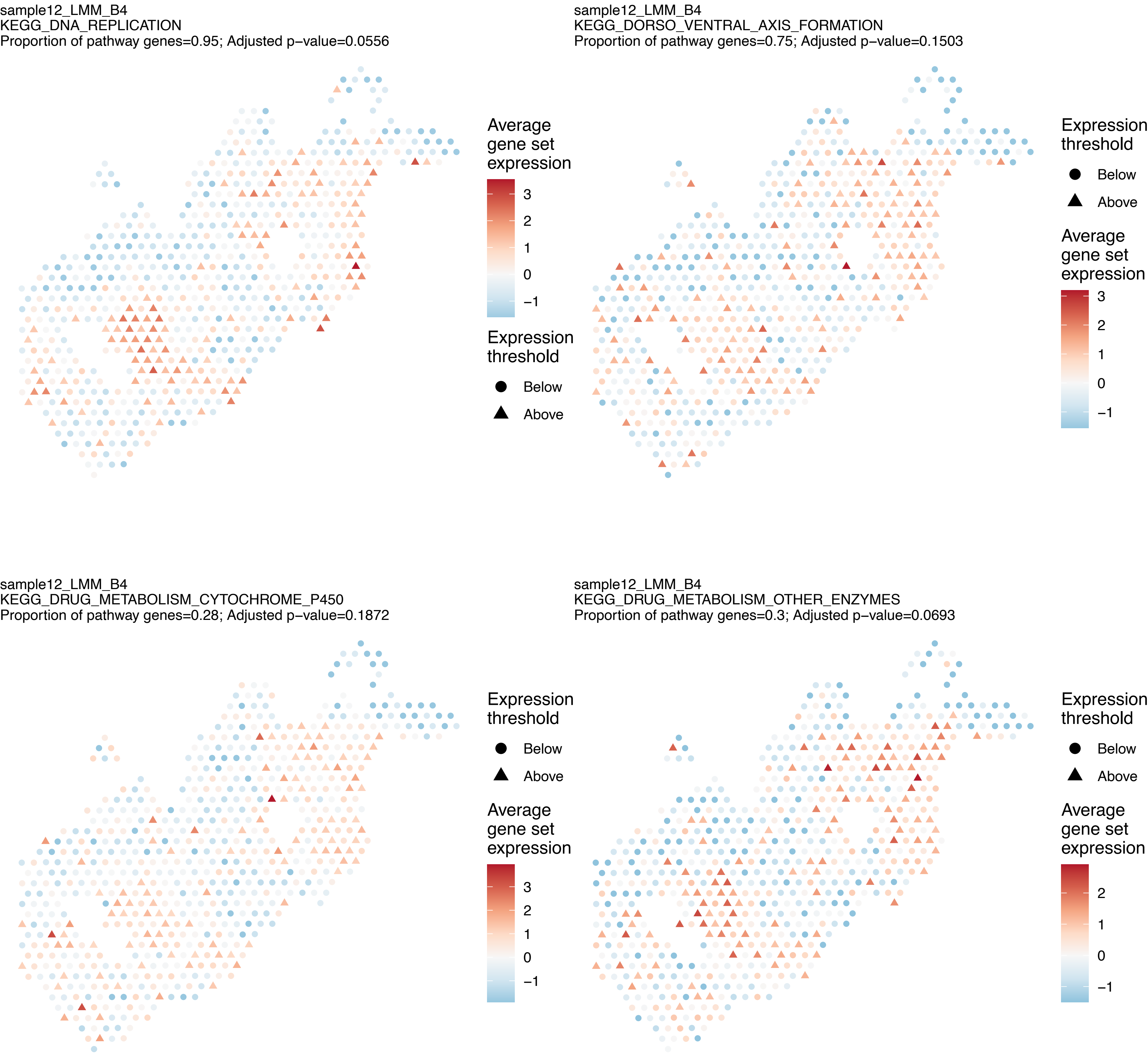
sample12_LMM_B4
KEGG_DNA_REPLICATION
Proportion of pathway genes=0.95; Adjusted p−value=0.0556
Average gene set expression
3
2
1
0
−1
Expression threshold
Below
Above
sample12_LMM_B4
KEGG_DORSO_VENTRAL_AXIS_FORMATION
Proportion of pathway genes=0.75; Adjusted p−value=0.1503
Expression threshold
Below
Above
Average gene set expression
3
2
1
0
−1
sample12_LMM_B4
KEGG_DRUG_METABOLISM_CYTOCHROME_P450
Proportion of pathway genes=0.28; Adjusted p−value=0.1872
Expression threshold
Below
Above
Average gene set expression
3
2
1
0
−1
sample12_LMM_B4
KEGG_DRUG_METABOLISM_OTHER_ENZYMES
Proportion of pathway genes=0.3; Adjusted p−value=0.0693
Expression threshold
Below
Above
Average gene set expression
2
1
0
−1

sample12_LMM_B4
KEGG_ECM_RECEPTOR_INTERACTION
Proportion of pathway genes=0.61; Adjusted p−value=0

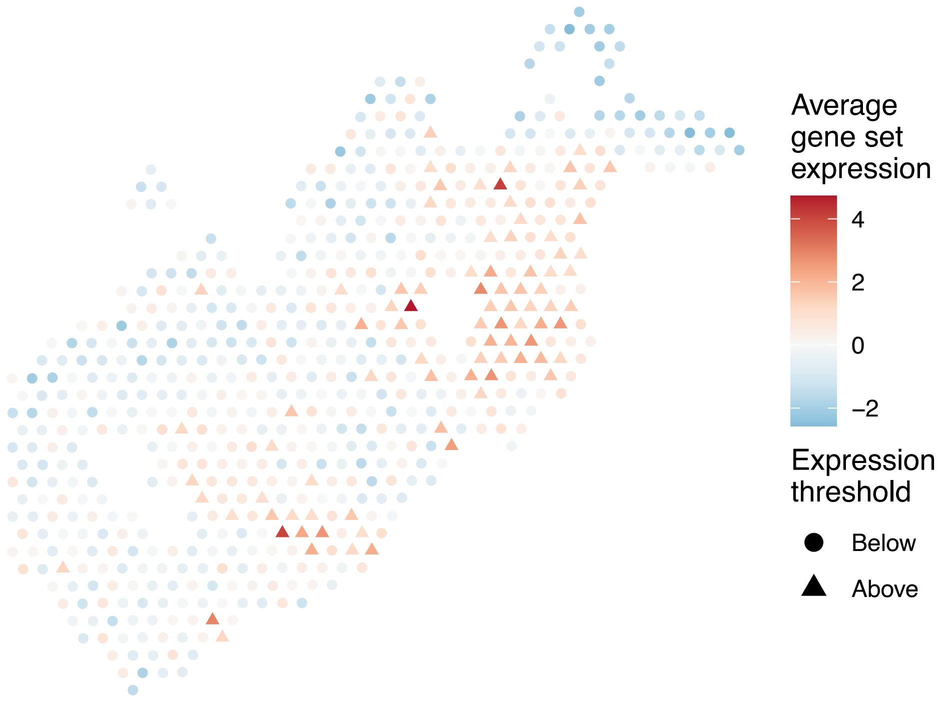


sample12_LMM_B4
KEGG_ENDOCYTOSIS
Proportion of pathway genes=0.8; Adjusted p−value=0

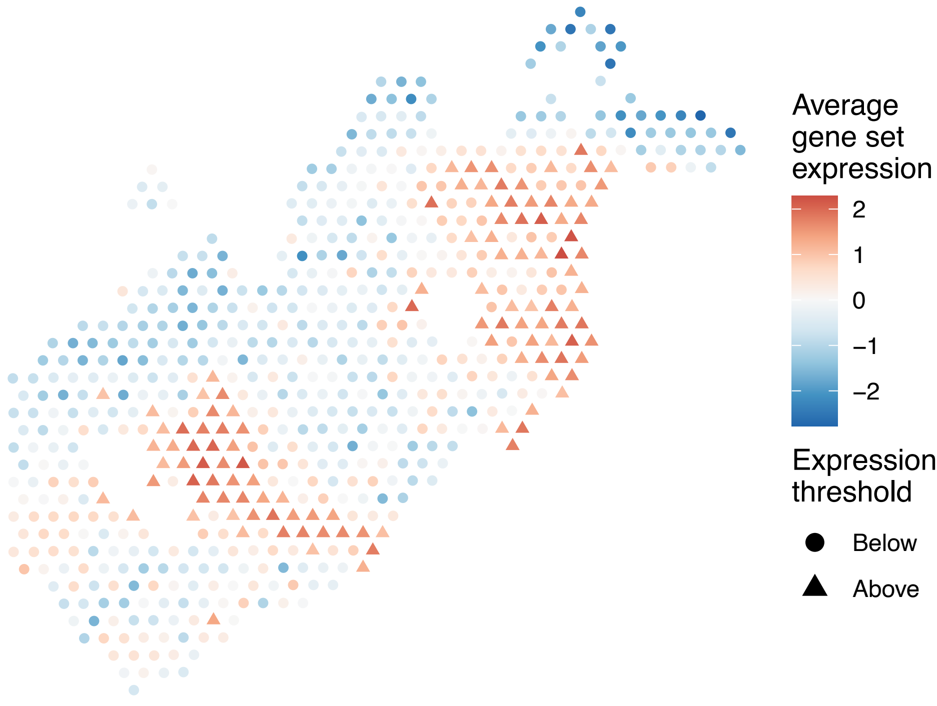


sample12_LMM_B4
KEGG_ENDOMETRIAL_CANCER
Proportion of pathway genes=0.85; Adjusted p−value=0.0104

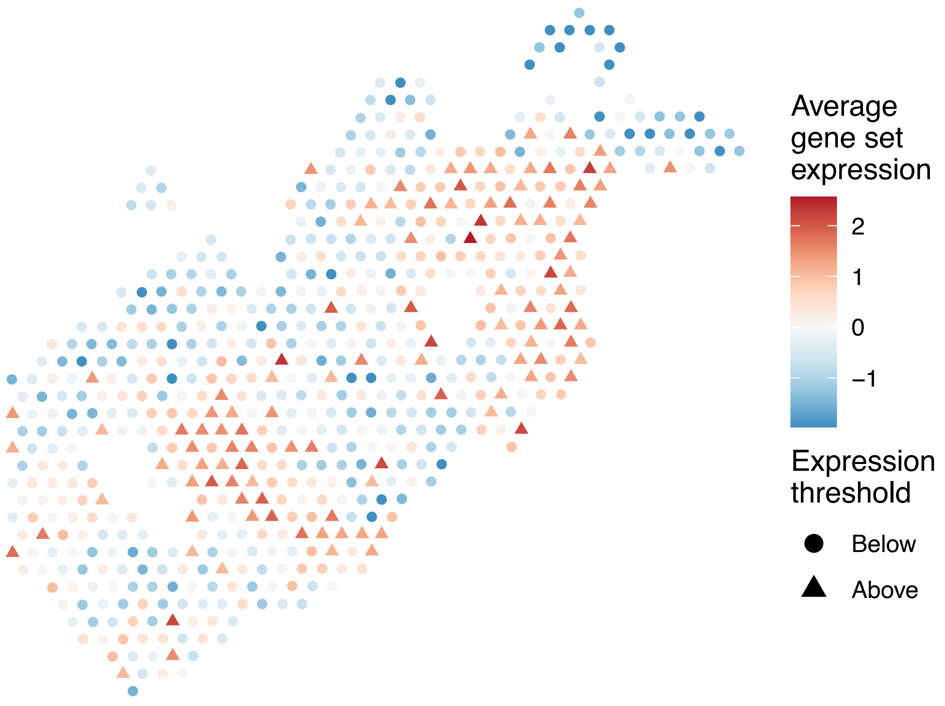


sample12_LMM_B4
KEGG_EPITHELIAL_CELL_SIGNALING_IN_HELICOBACTER_PYL
ORI_INFECTION
Proportion of pathway genes=0.77; Adjusted p−value=0.022

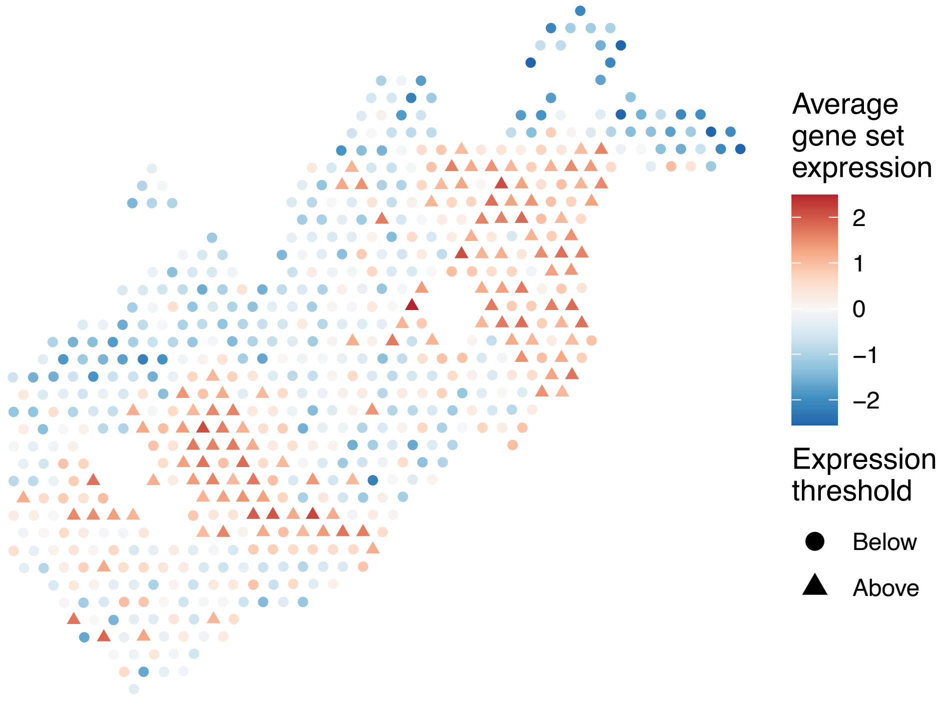

sample12_LMM_B4
KEGG_ERBB_SIGNALING_PATHWAY
Proportion of pathway genes=0.76; Adjusted p−value=0.0079

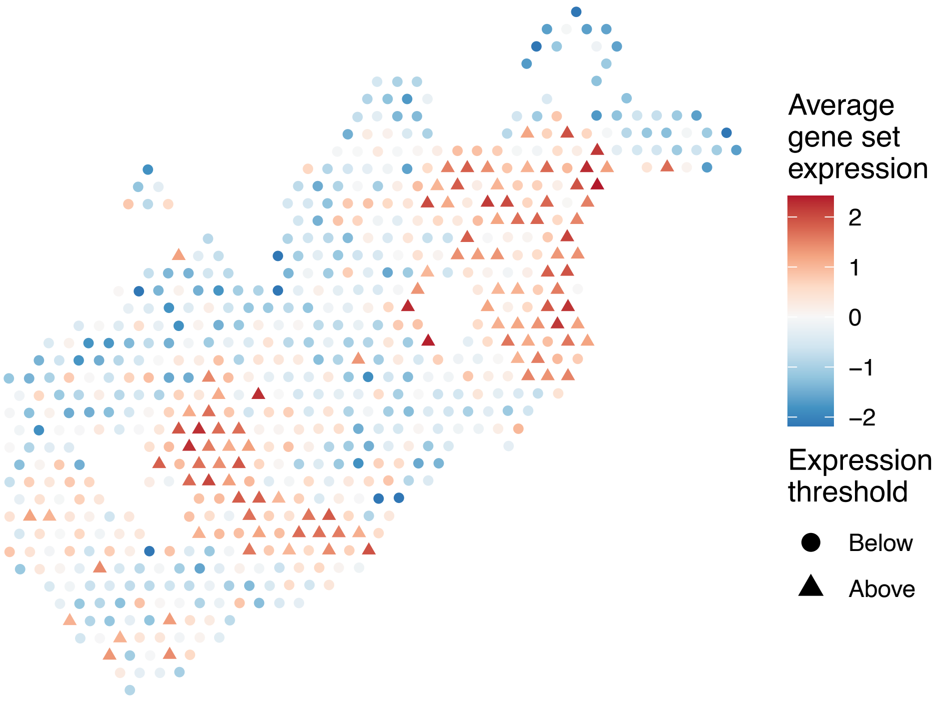


sample12_LMM_B4
KEGG_ETHER_LIPID_METABOLISM
Proportion of pathway genes=0.57; Adjusted p−value=0.0826

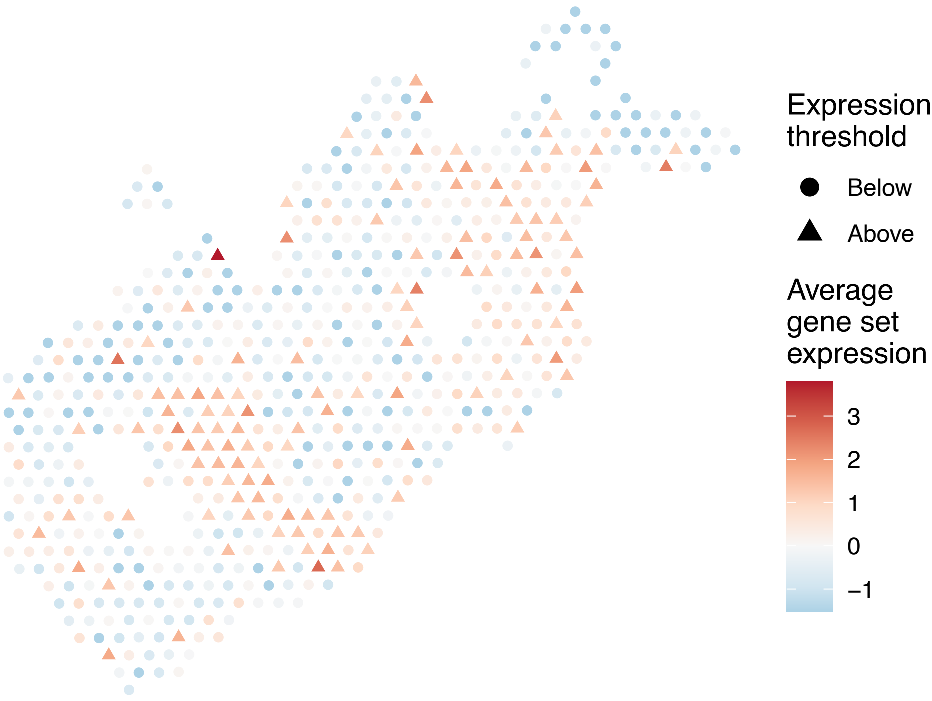


sample12_LMM_B4
KEGG_FATTY_ACID_METABOLISM
Proportion of pathway genes=0.74; Adjusted p−value=0.1392

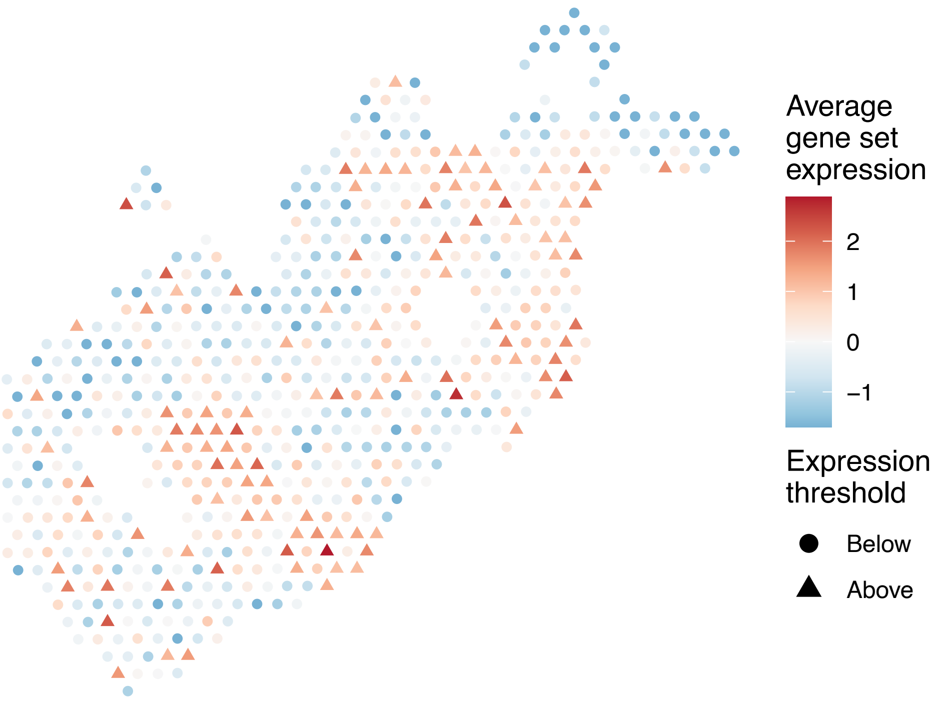


sample12_LMM_B4
KEGG_FC_EPSILON_RI_SIGNALING_PATHWAY
Proportion of pathway genes=0.61; Adjusted p−value=0.1465

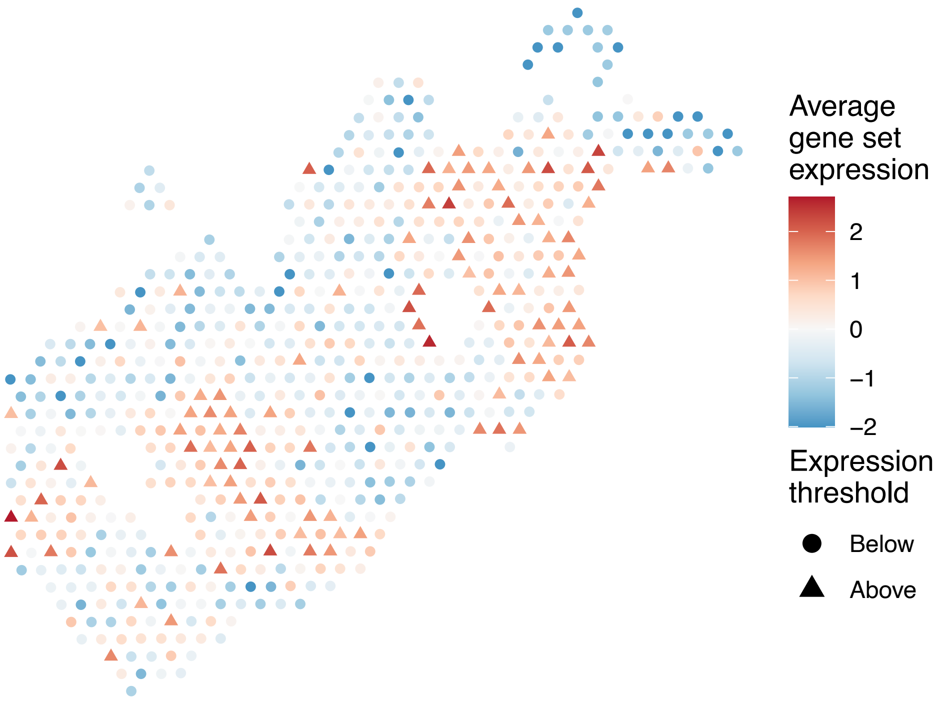

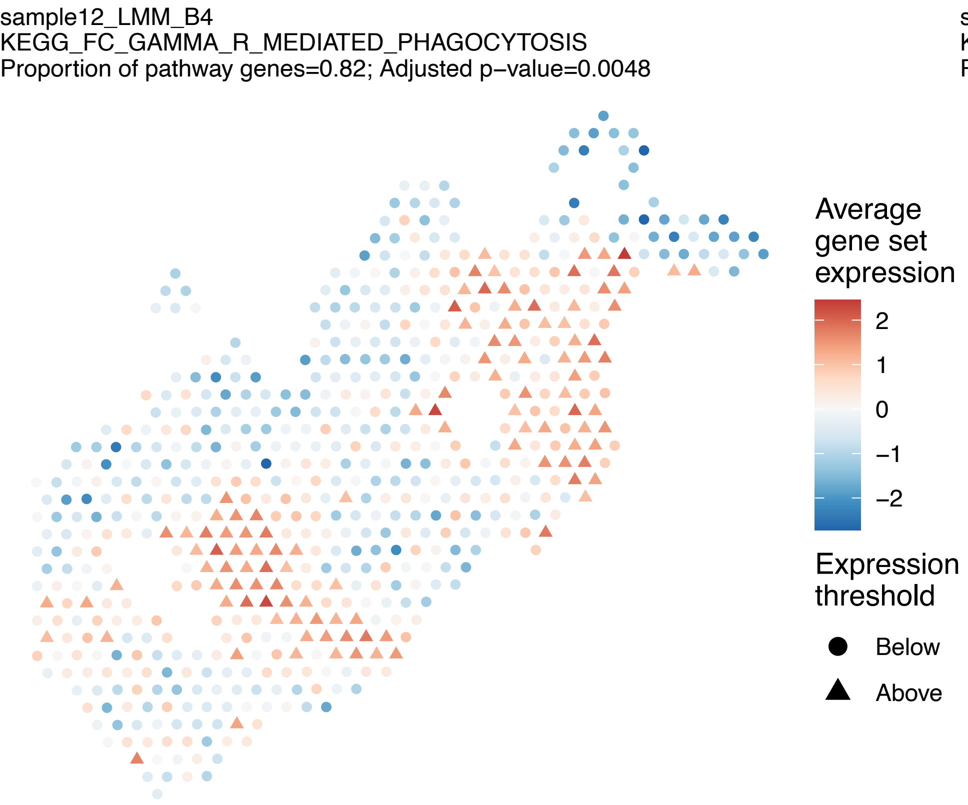
sample12_LMM_B4
KEGG_FC_GAMMA_R_MEDIATED_PHAGOCYTOSIS
Proportion of pathway genes=0.82; Adjusted p−value=0.0048
Average gene set expression
2
1
0
−1
−2
Expression threshold
Below
Above

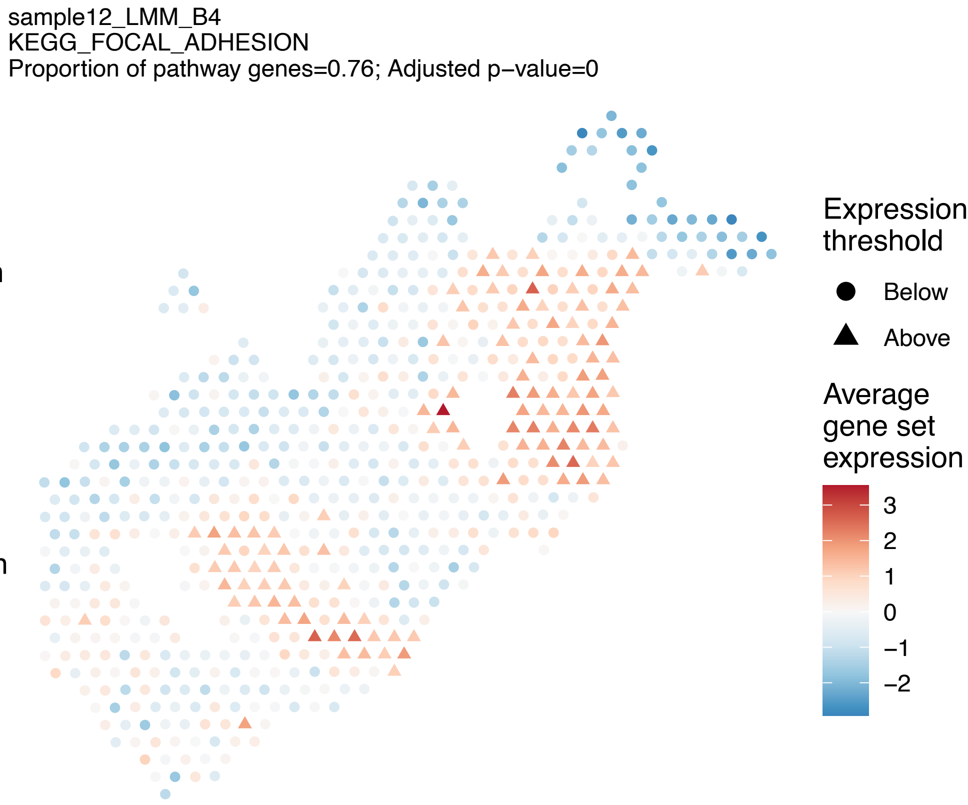
sample12_LMM_B4
KEGG_FOCAL_ADHESION
Proportion of pathway genes=0.76; Adjusted p−value=0
Expression threshold
Below
Above
Average gene set expression
3
2
1
0
−1
−2

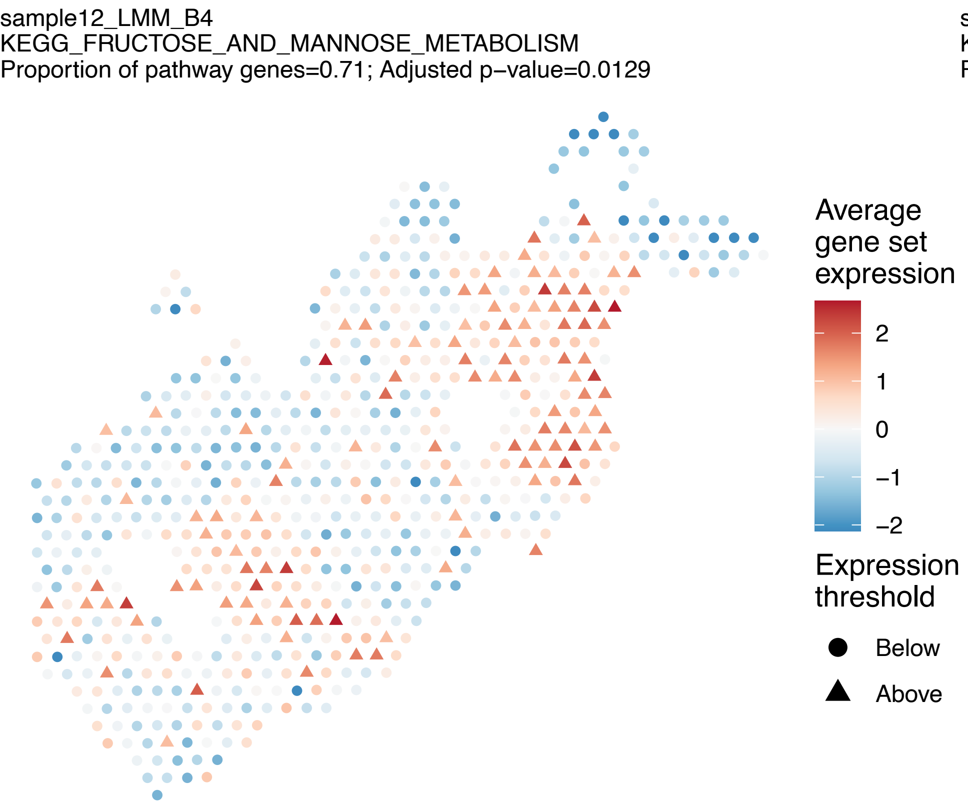
sample12_LMM_B4
KEGG_FRUCTOSE_AND_MANNOSE_METABOLISM
Proportion of pathway genes=0.71; Adjusted p−value=0.0129
Average gene set expression
2
1
0
−1
−2
Expression threshold
Below
Above

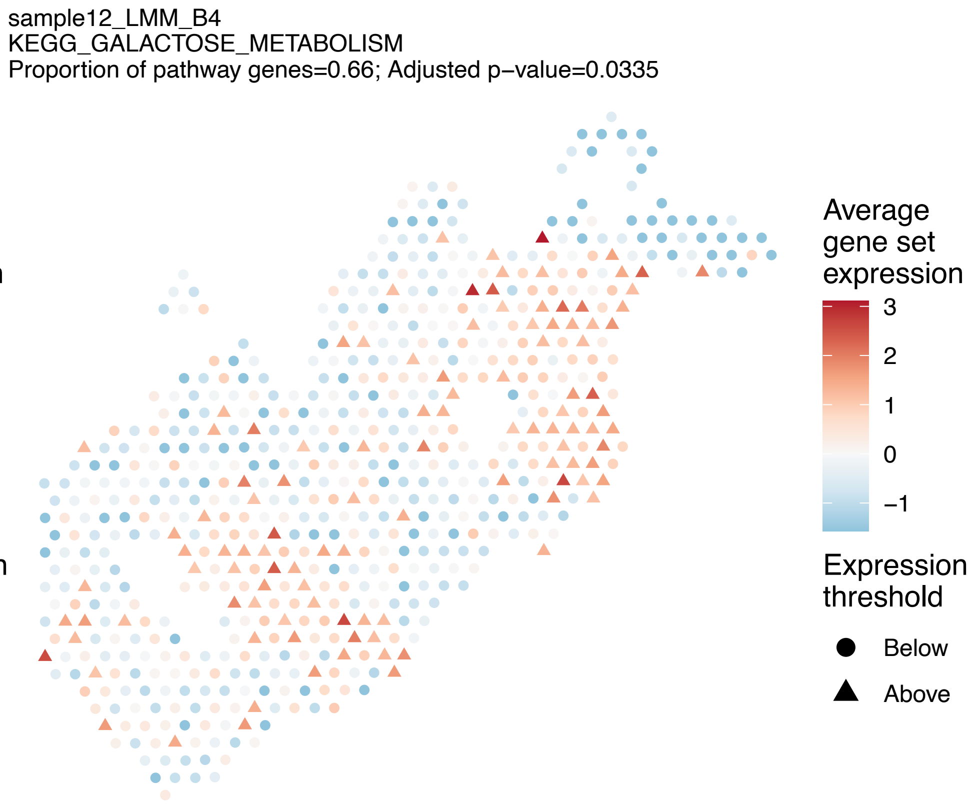
sample12_LMM_B4
KEGG_GALACTOSE_METABOLISM
Proportion of pathway genes=0.66; Adjusted p−value=0.0335
Average gene set expression
3
2
1
0
−1
Expression threshold
Below
Above

sample12_LMM_B4
KEGG_GAP_JUNCTION
Proportion of pathway genes=0.64; Adjusted p−value=0.2316

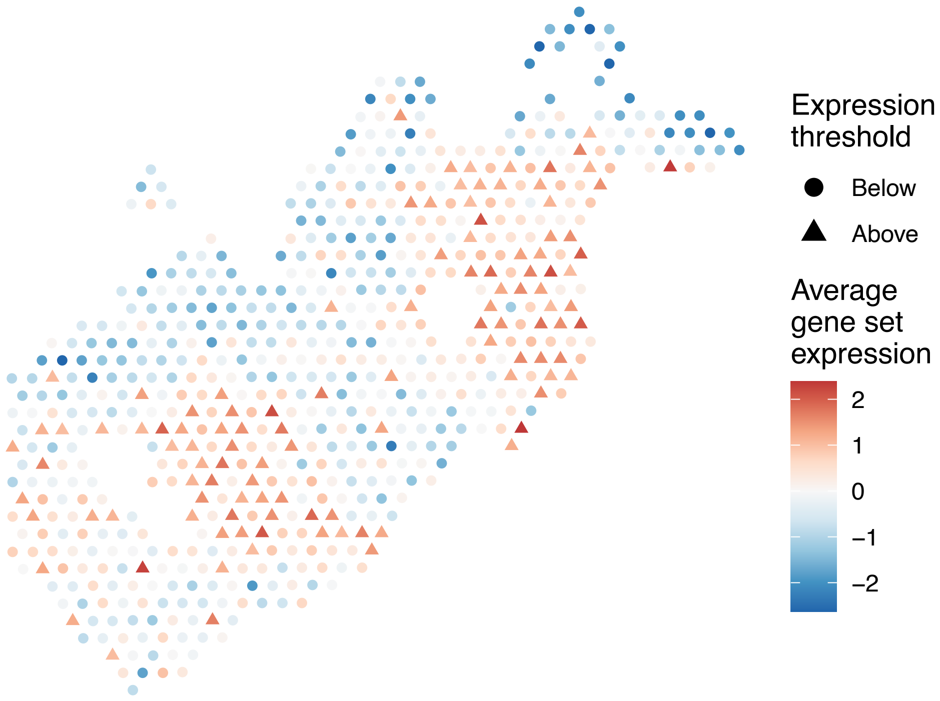


sample12_LMM_B4
KEGG_GLIOMA
Proportion of pathway genes=0.8; Adjusted p−value=0.0048

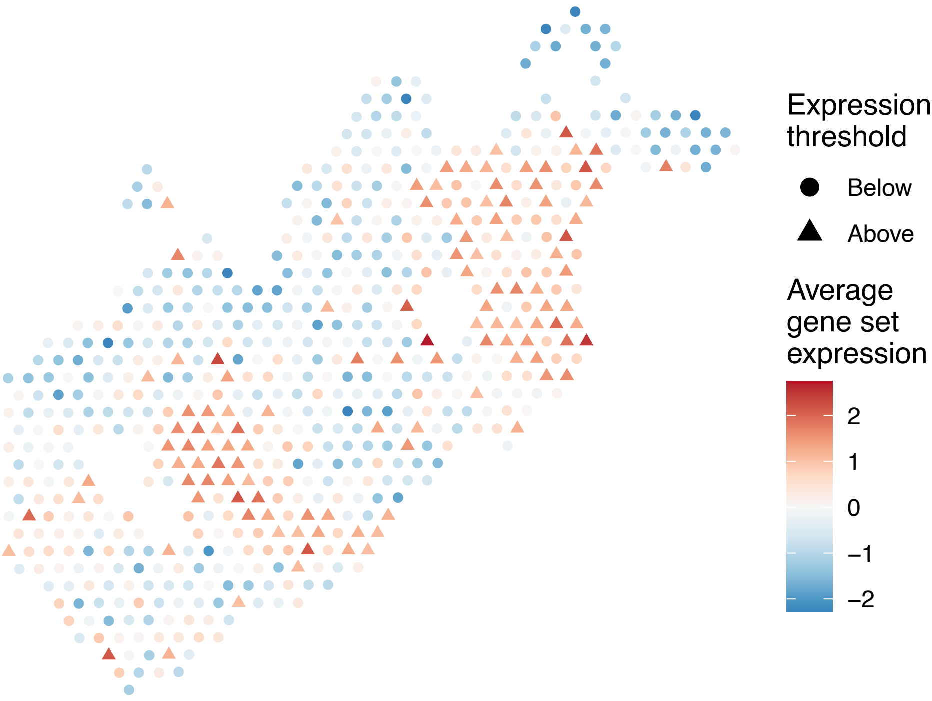


sample12_LMM_B4
KEGG_GLUTATHIONE_METABOLISM
Proportion of pathway genes=0.66; Adjusted p−value=0.0209

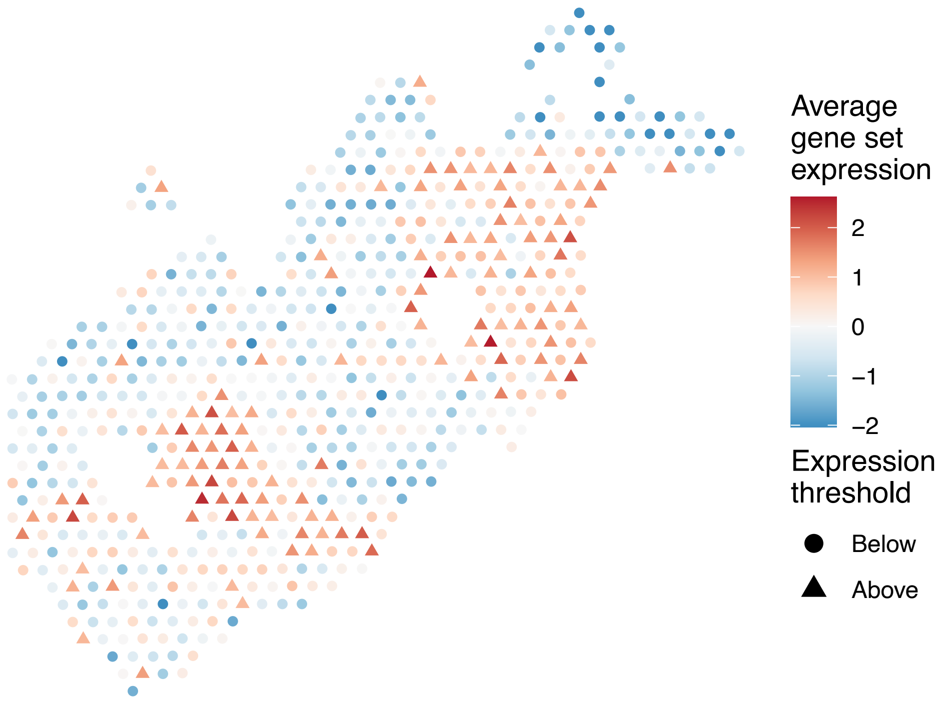


sample12_LMM_B4
KEGG_GLYCEROLIPID_METABOLISM
Proportion of pathway genes=0.71; Adjusted p−value=0.1116

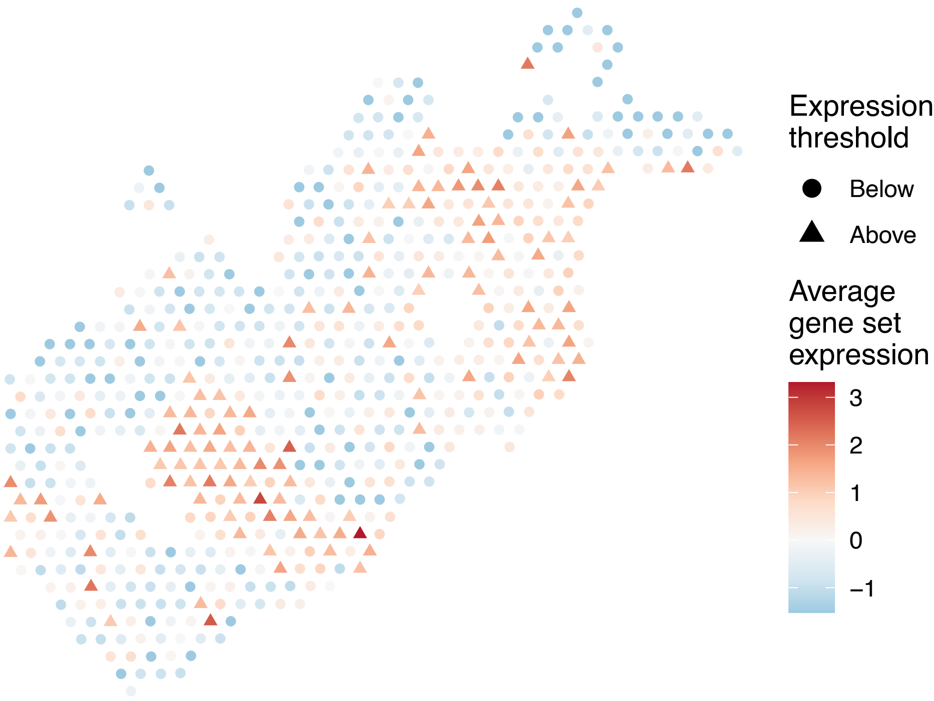

sample12_LMM_B4
KEGG_GLYCEROPHOSPHOLIPID_METABOLISM
Proportion of pathway genes=0.74; Adjusted p−value=0.0209

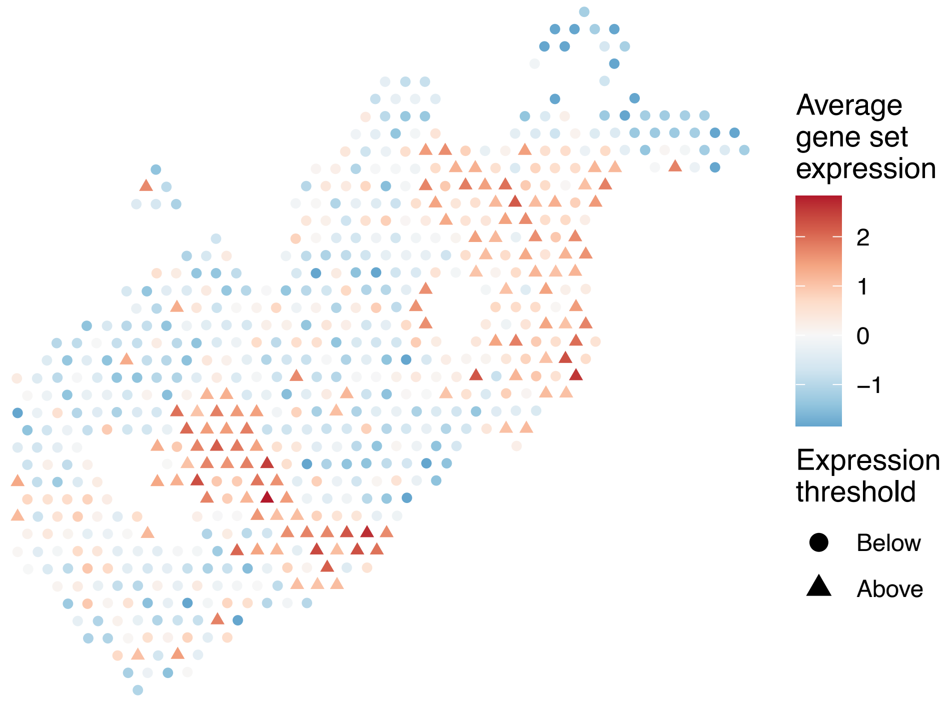


sample12_LMM_B4
KEGG_GLYCINE_SERINE_AND_THREONINE_METABOLISM
Proportion of pathway genes=0.5; Adjusted p−value=0.2432

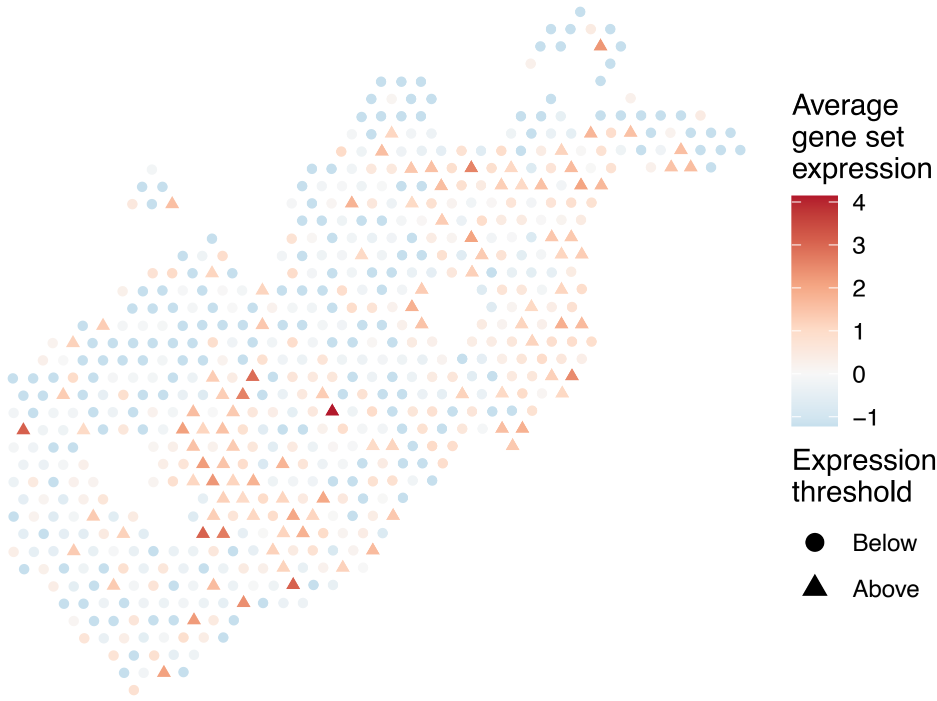


sample12_LMM_B4
KEGG_GLYCOLYSIS_GLUCONEOGENESIS
Proportion of pathway genes=0.64; Adjusted p−value=0

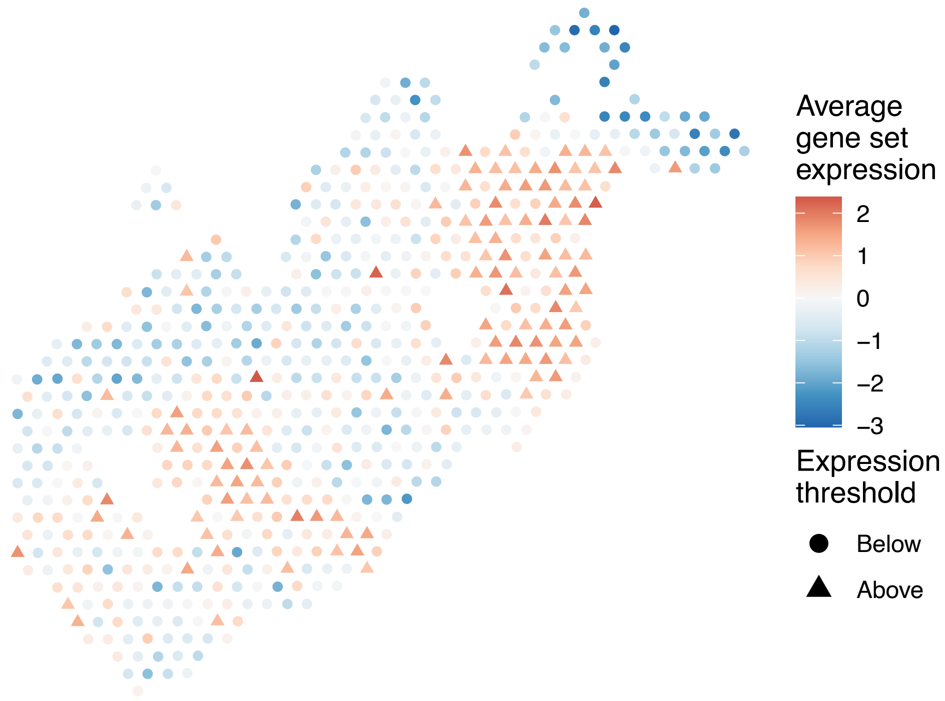


sample12_LMM_B4
KEGG_GLYCOSAMINOGLYCAN_BIOSYNTHESIS_CHONDROITIN_SU
LFATE
Proportion of pathway genes=0.78; Adjusted p−value=0.1478

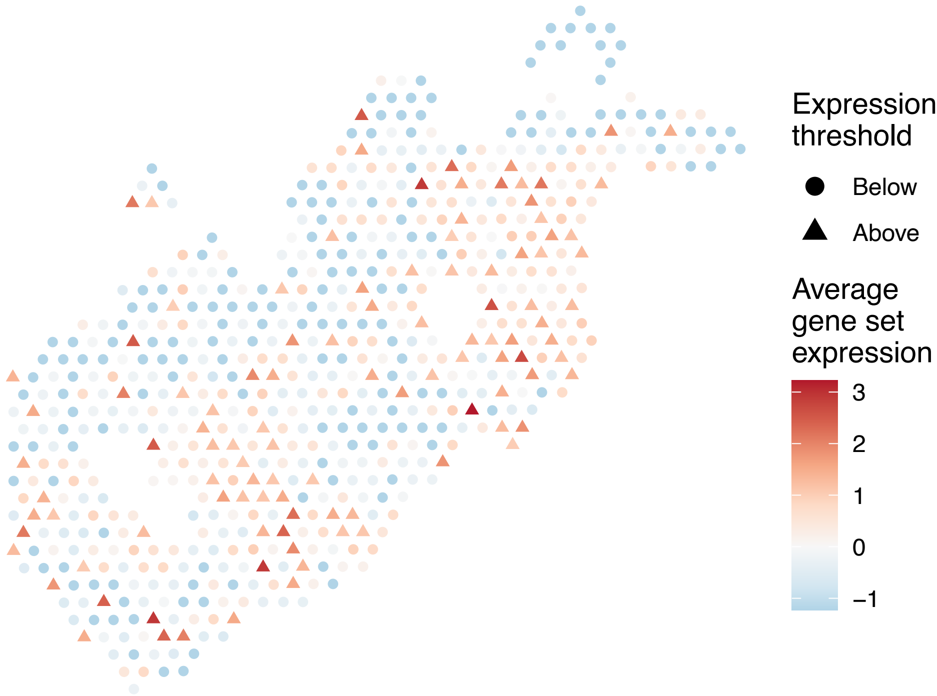

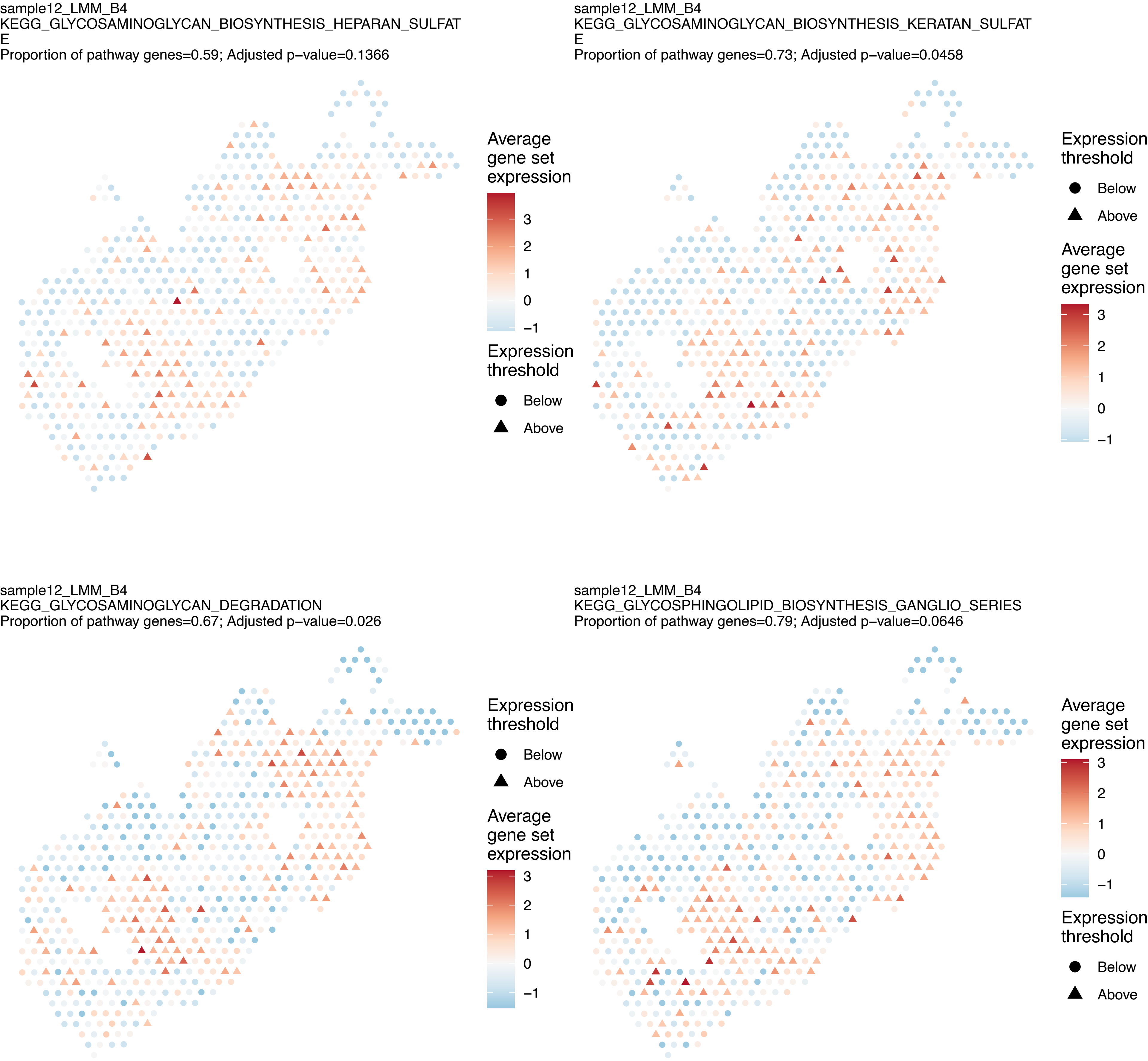
sample12_LMM_B4
KEGG_GLYCOSAMINOGLYCAN_BIOSYNTHESIS_HEPARAN_SULFAT
E
Proportion of pathway genes=0.59; Adjusted p−value=0.1366
Average gene set expression
3
2
1
0
−1
Expression threshold
Below
Above
sample12_LMM_B4
KEGG_GLYCOSAMINOGLYCAN_BIOSYNTHESIS_KERATAN_SULFAT
E
Proportion of pathway genes=0.73; Adjusted p−value=0.0458
Expression threshold
Below
Above
Average gene set expression
3
2
1
0
−1
sample12_LMM_B4
KEGG_GLYCOSAMINOGLYCAN_DEGRADATION
Proportion of pathway genes=0.67; Adjusted p−value=0.026
Expression threshold
Below
Above
Average gene set expression
3
2
1
0
−1
sample12_LMM_B4
KEGG_GLYCOSPHINGOLIPID_BIOSYNTHESIS_GANGLIO_SERIES
Proportion of pathway genes=0.79; Adjusted p−value=0.0646
Average gene set expression
3
2
1
0
−1
Expression threshold
Below
Above

sample12_LMM_B4
KEGG_GLYCOSPHINGOLIPID_BIOSYNTHESIS_GLOBO_SERIES
Proportion of pathway genes=0.5; Adjusted p−value=NA

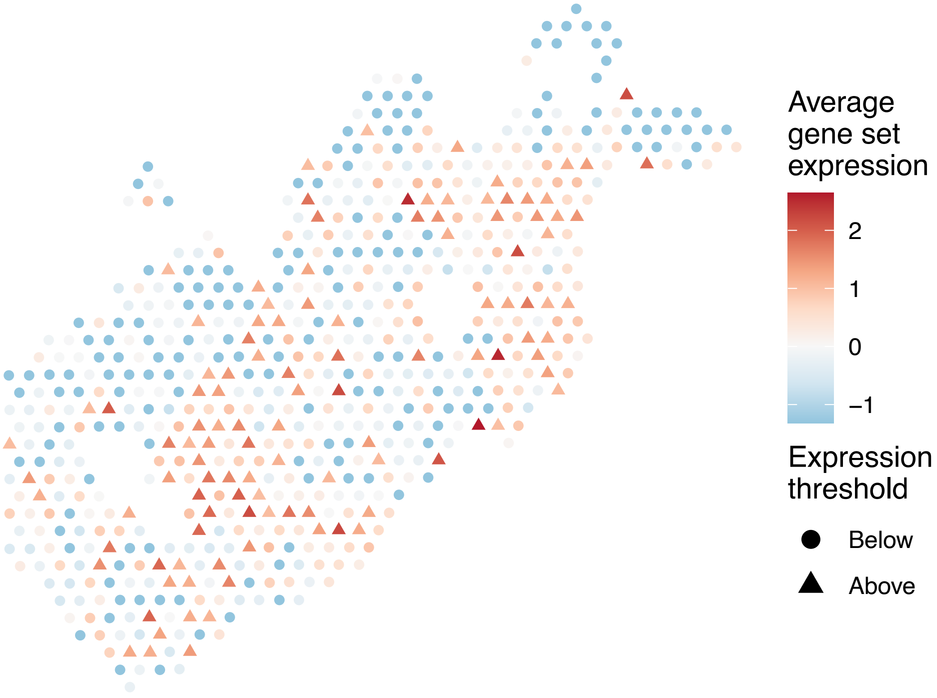


sample12_LMM_B4
KEGG_GLYCOSPHINGOLIPID_BIOSYNTHESIS_LACTO_AND_NEOL
ACTO_SERIES
Proportion of pathway genes=0.41; Adjusted p−value=0.0603

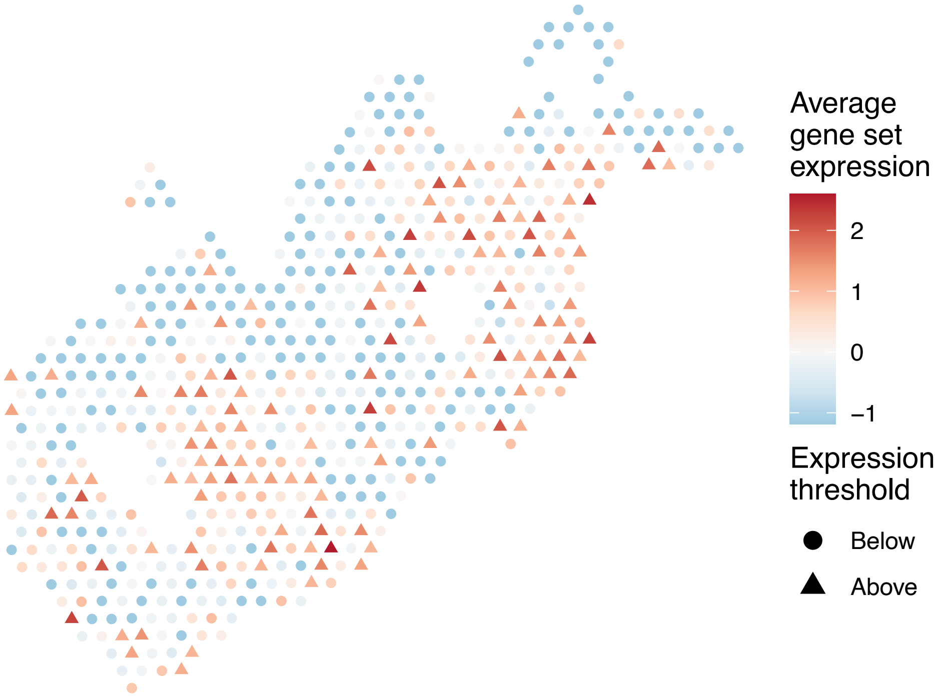


sample12_LMM_B4
KEGG_GLYCOSYLPHOSPHATIDYLINOSITOL_GPI_ANCHOR_BIOSY
NTHESIS
Proportion of pathway genes=0.89; Adjusted p−value=0.0079

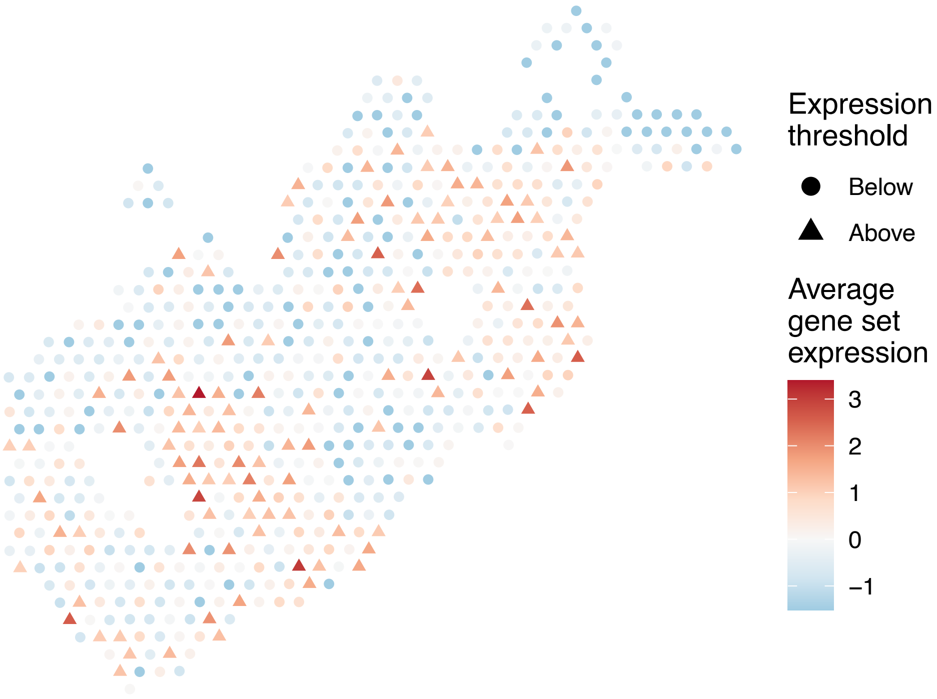


sample12_LMM_B4
KEGG_GLYOXYLATE_AND_DICARBOXYLATE_METABOLISM
Proportion of pathway genes=0.81; Adjusted p−value=0.0104

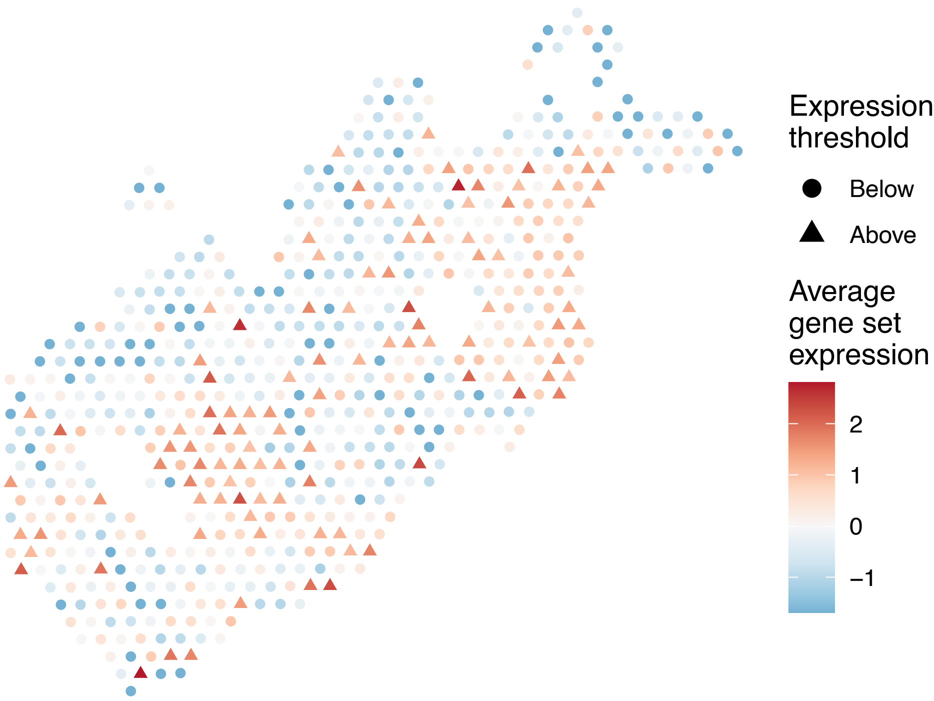

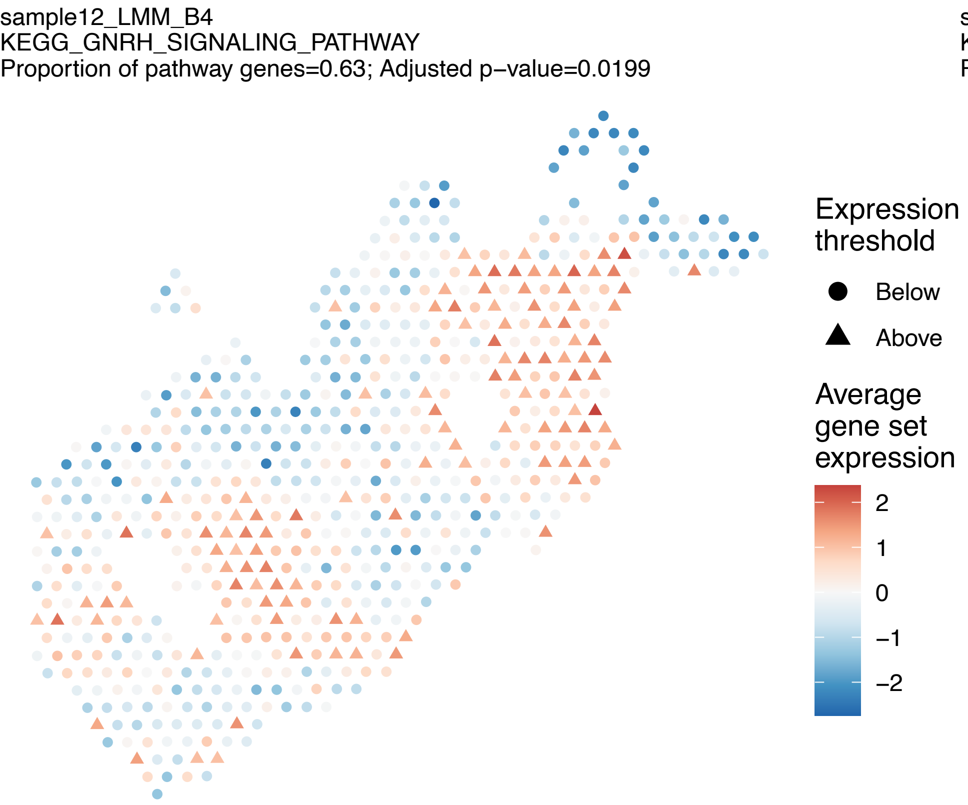
sample12_LMM_B4
KEGG_GNRH_SIGNALING_PATHWAY
Proportion of pathway genes=0.63; Adjusted p−value=0.0199
Expression threshold
Below
Above
Average gene set expression
2
1
0
−1
−2

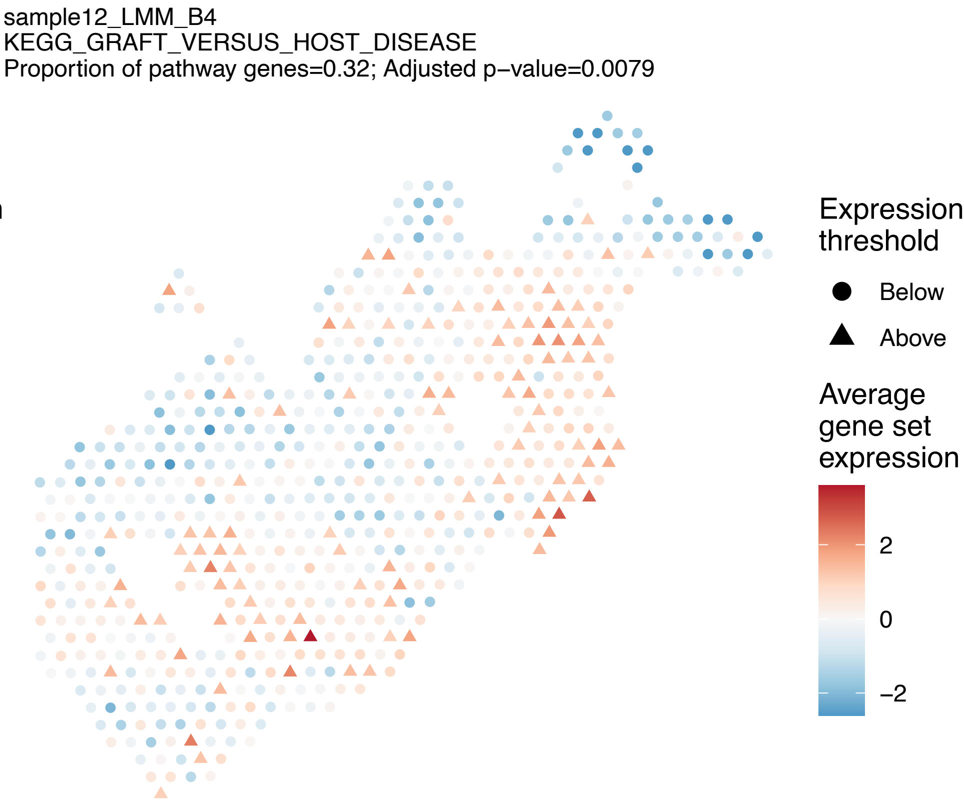
sample12_LMM_B4
KEGG_GRAFT_VERSUS_HOST_DISEASE
Proportion of pathway genes=0.32; Adjusted p−value=0.0079
Expression threshold
Below
Above
Average gene set expression
2
0
−2

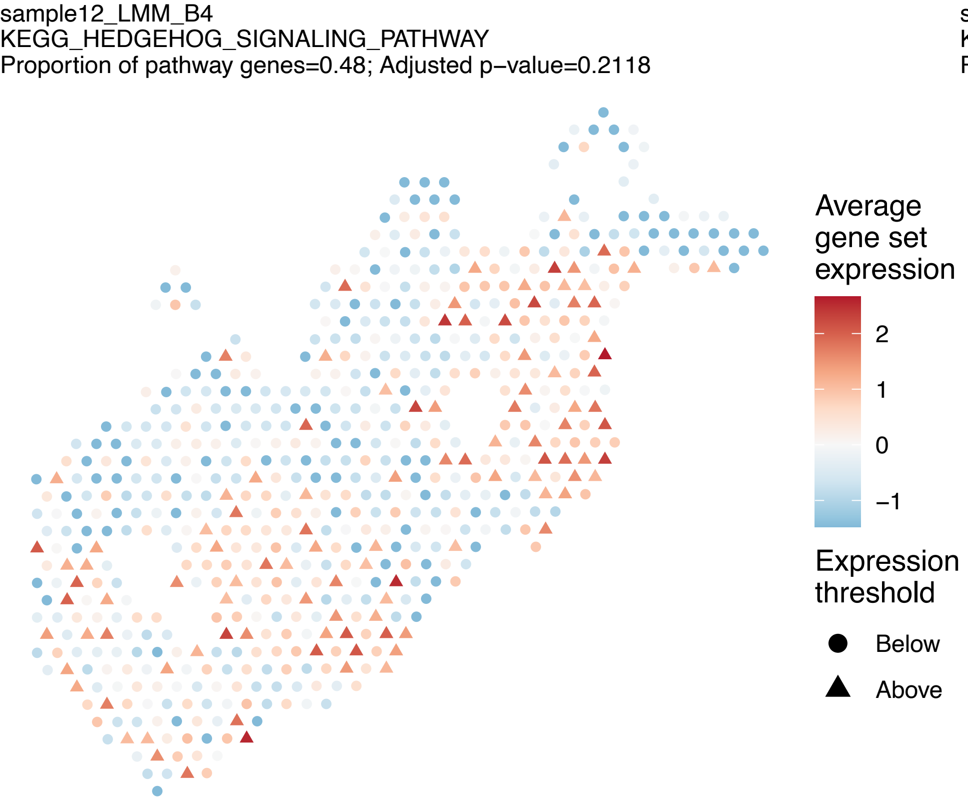
sample12_LMM_B4
KEGG_HEDGEHOG_SIGNALING_PATHWAY
Proportion of pathway genes=0.48; Adjusted p−value=0.2118
Average gene set expression
2
1
0
−1
Expression threshold
Below
Above

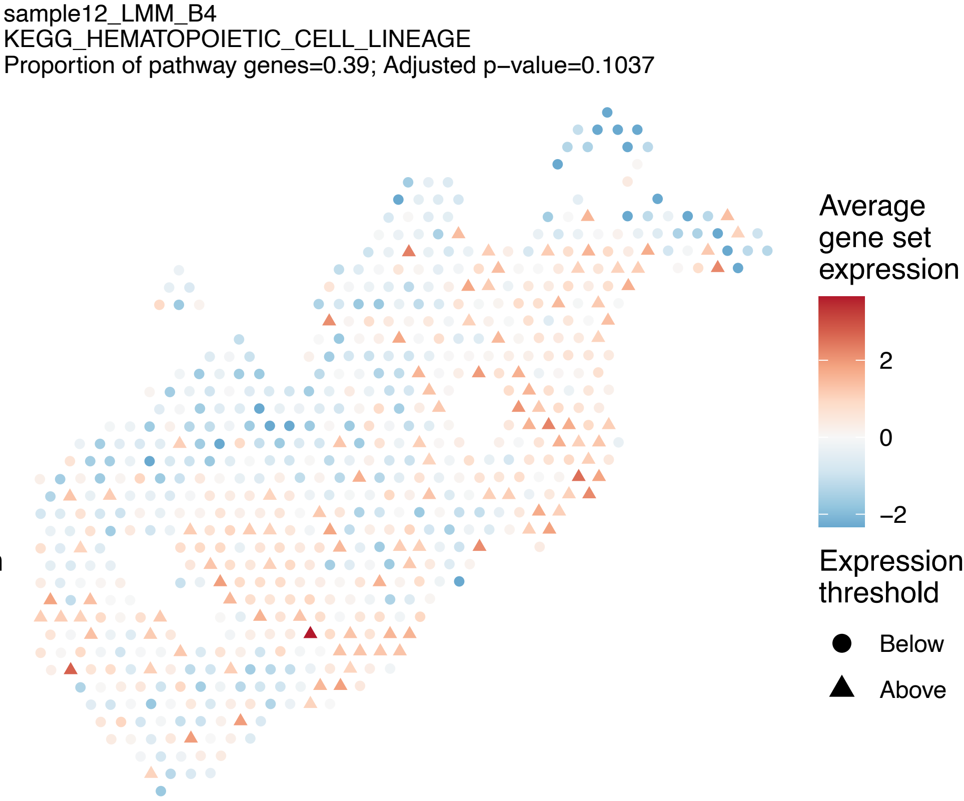
sample12_LMM_B4
KEGG_HEMATOPOIETIC_CELL_LINEAGE
Proportion of pathway genes=0.39; Adjusted p−value=0.1037
Average gene set expression
2
0
−2
Expression threshold
Below
Above

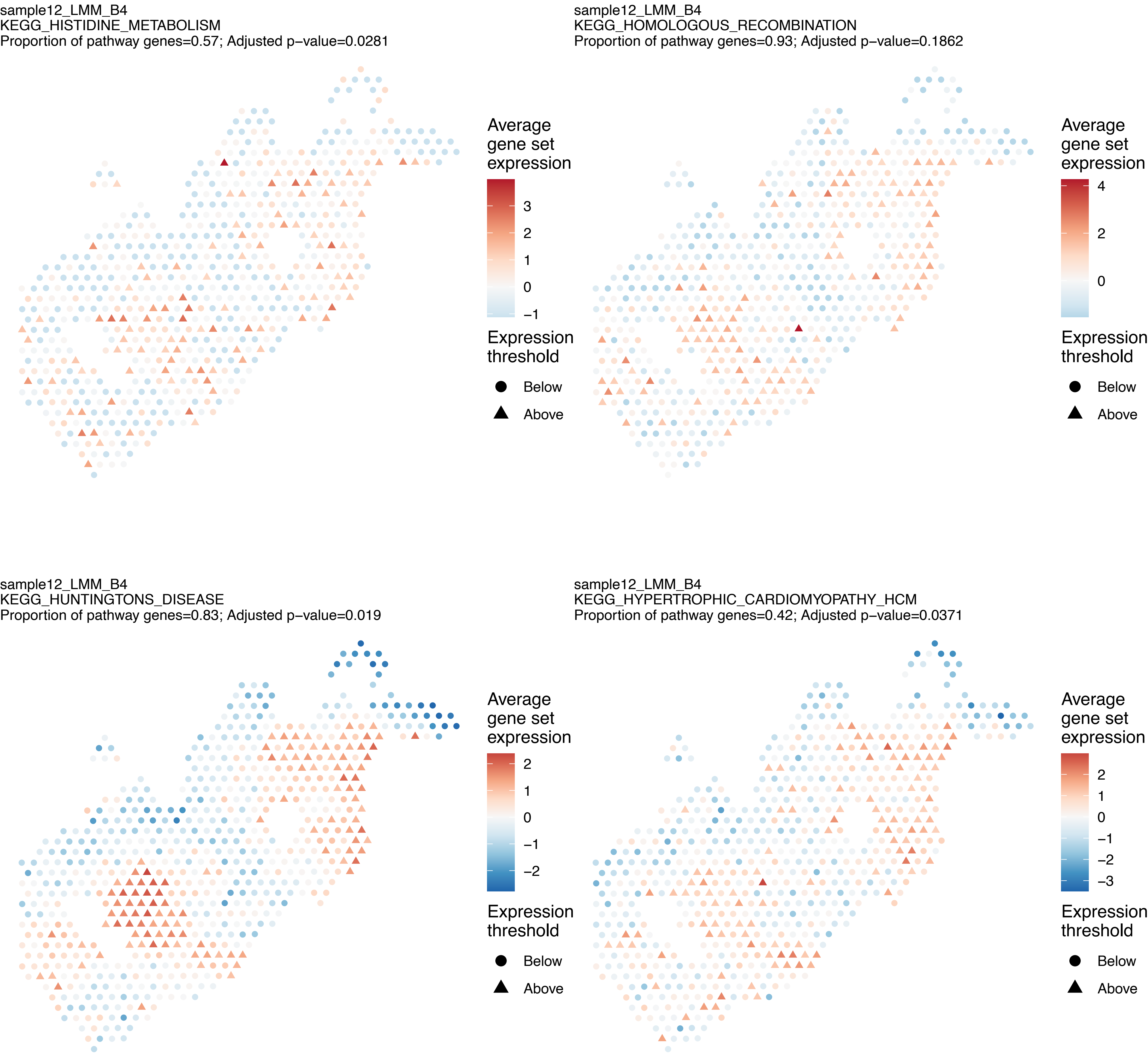
sample12_LMM_B4
KEGG_HISTIDINE_METABOLISM
Proportion of pathway genes=0.57; Adjusted p−value=0.0281
Average gene set expression
3
2
1
0
−1
Expression threshold
Below
Above
sample12_LMM_B4
KEGG_HOMOLOGOUS_RECOMBINATION
Proportion of pathway genes=0.93; Adjusted p−value=0.1862
Average gene set expression
4
2
0
Expression threshold
Below
Above
sample12_LMM_B4
KEGG_HUNTINGTONS_DISEASE
Proportion of pathway genes=0.83; Adjusted p−value=0.019
Average gene set expression
2
1
0
−1
−2
Expression threshold
Below
Above
sample12_LMM_B4
KEGG_HYPERTROPHIC_CARDIOMYOPATHY_HCM
Proportion of pathway genes=0.42; Adjusted p−value=0.0371
Average gene set expression
2
1
0
−1
−2
−3
Expression threshold
Below
Above

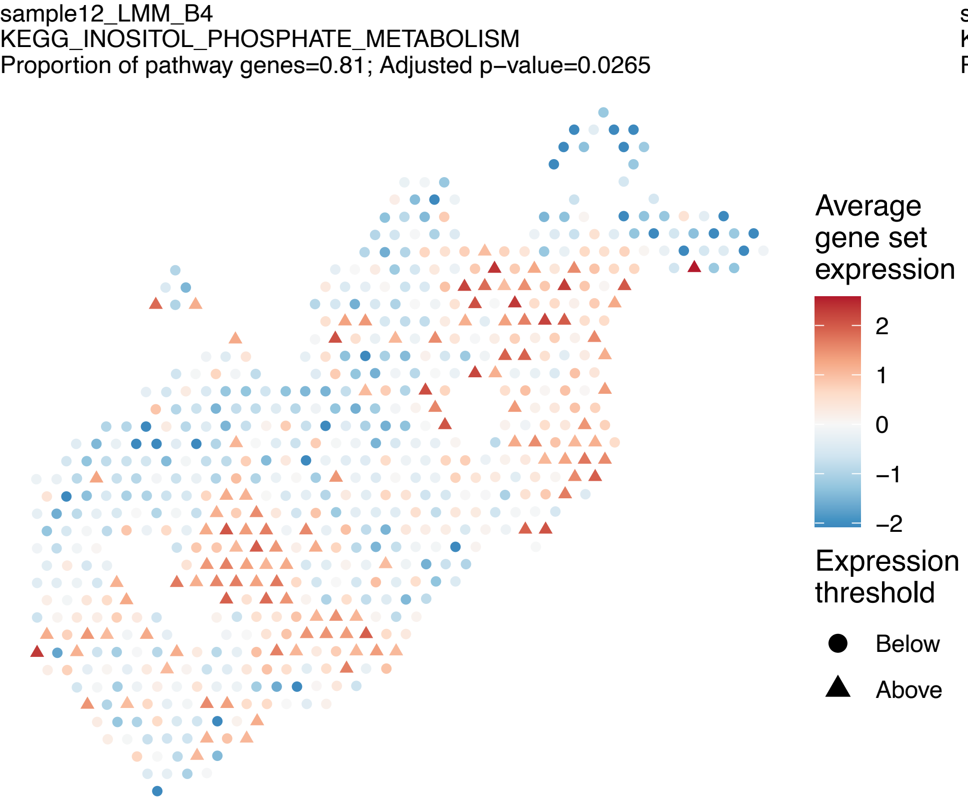
sample12_LMM_B4
KEGG_INOSITOL_PHOSPHATE_METABOLISM
Proportion of pathway genes=0.81; Adjusted p−value=0.0265
Average gene set expression
2
1
0
−1
−2
Expression threshold
Below
Above

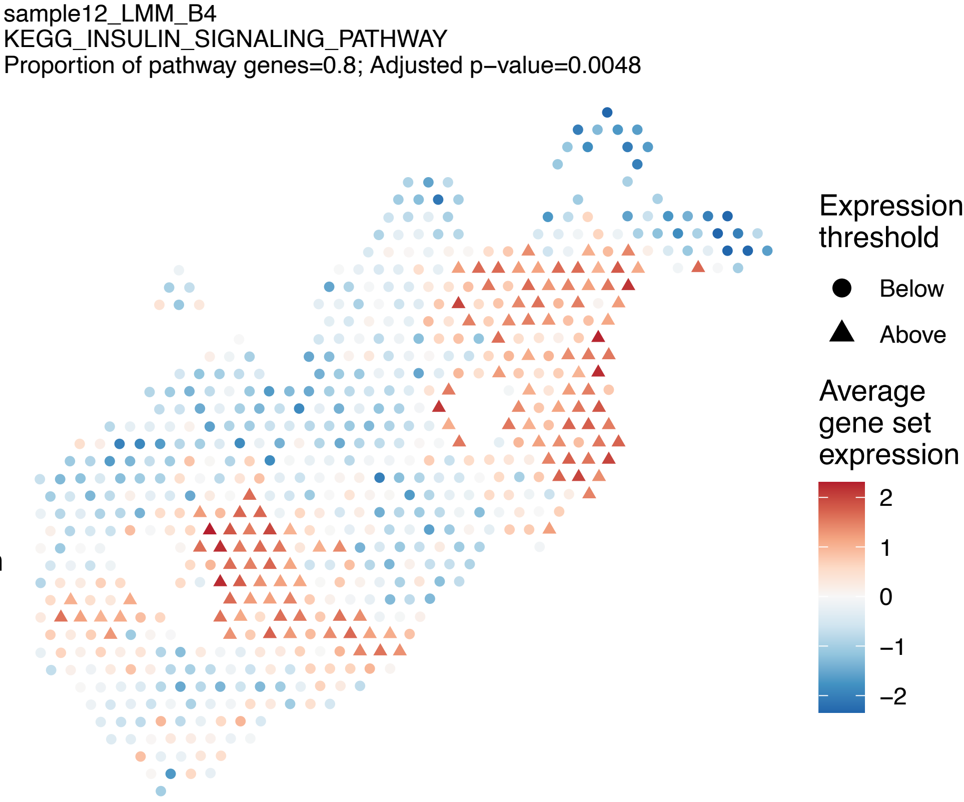
sample12_LMM_B4
KEGG_INSULIN_SIGNALING_PATHWAY
Proportion of pathway genes=0.8; Adjusted p−value=0.0048
Expression threshold
Below
Above
Average gene set expression
2
1
0
−1
−2

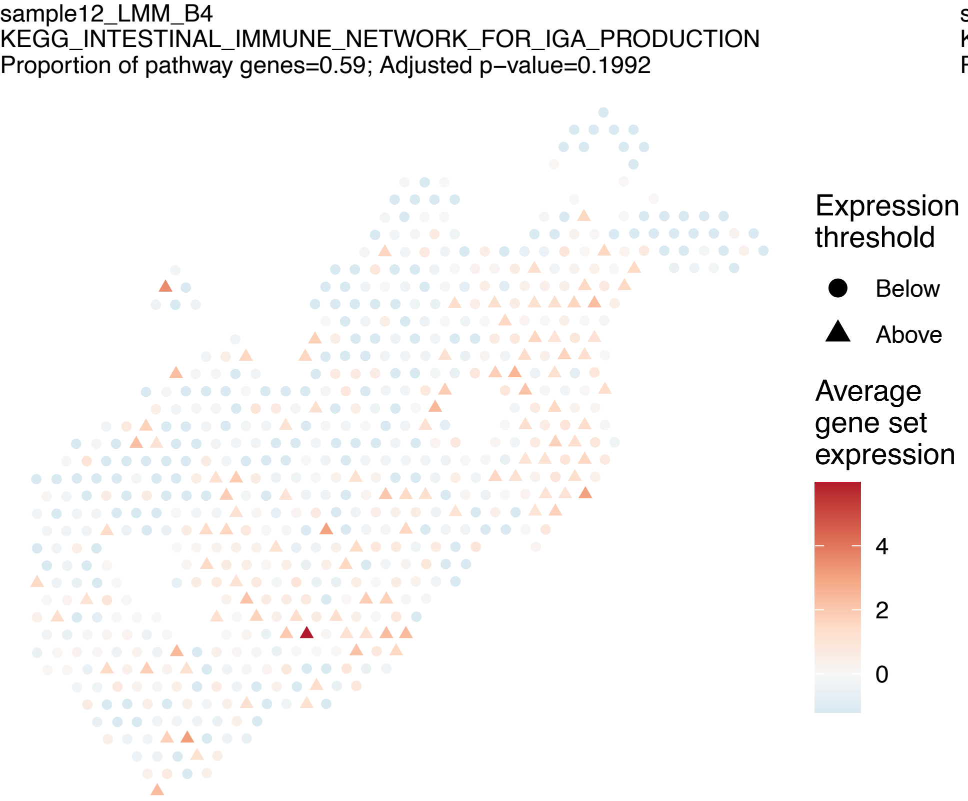
sample12_LMM_B4
KEGG_INTESTINAL_IMMUNE_NETWORK_FOR_IGA_PRODUCTION
Proportion of pathway genes=0.59; Adjusted p−value=0.1992
Expression threshold
Below
Above
Average gene set expression
4
2
0

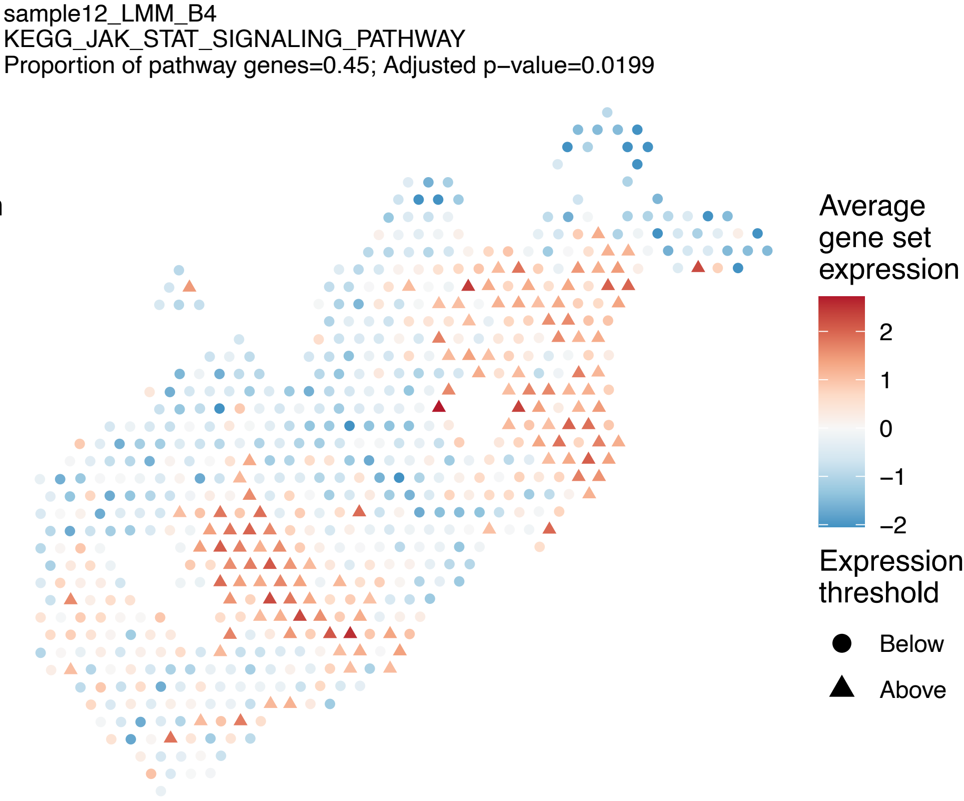
sample12_LMM_B4
KEGG_JAK_STAT_SIGNALING_PATHWAY
Proportion of pathway genes=0.45; Adjusted p−value=0.0199
Average gene set expression
2
1
0
−1
−2
Expression threshold
Below
Above

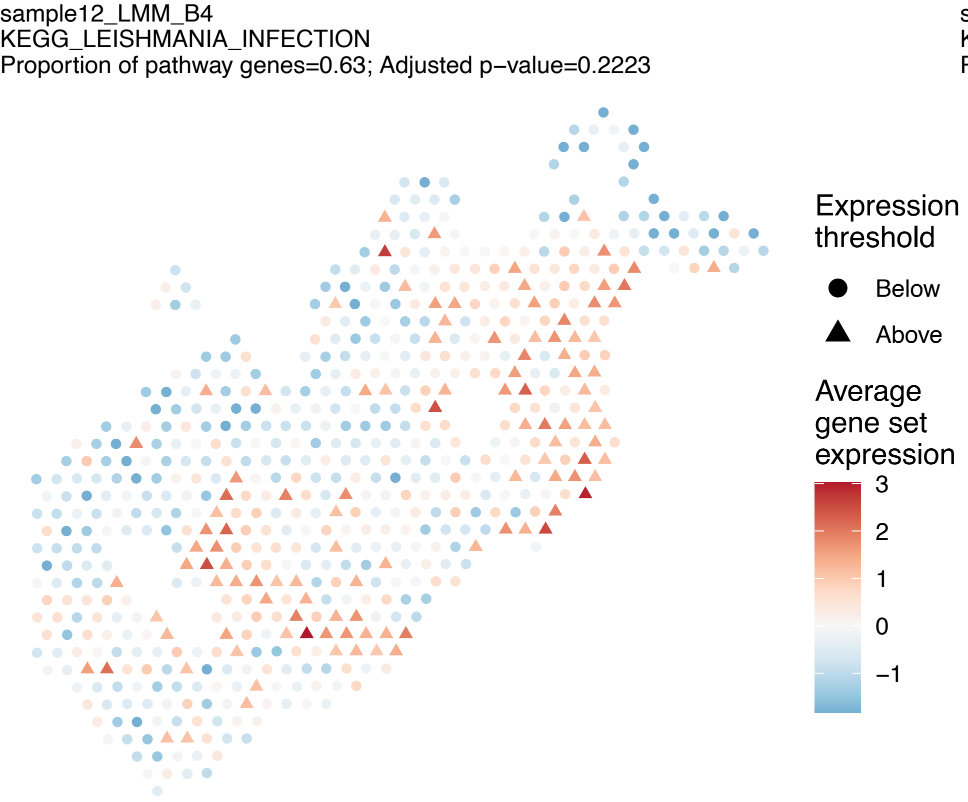
sample12_LMM_B4
KEGG_LEISHMANIA_INFECTION
Proportion of pathway genes=0.63; Adjusted p−value=0.2223
Expression threshold
Below
Above
Average gene set expression
3
2
1
0
−1

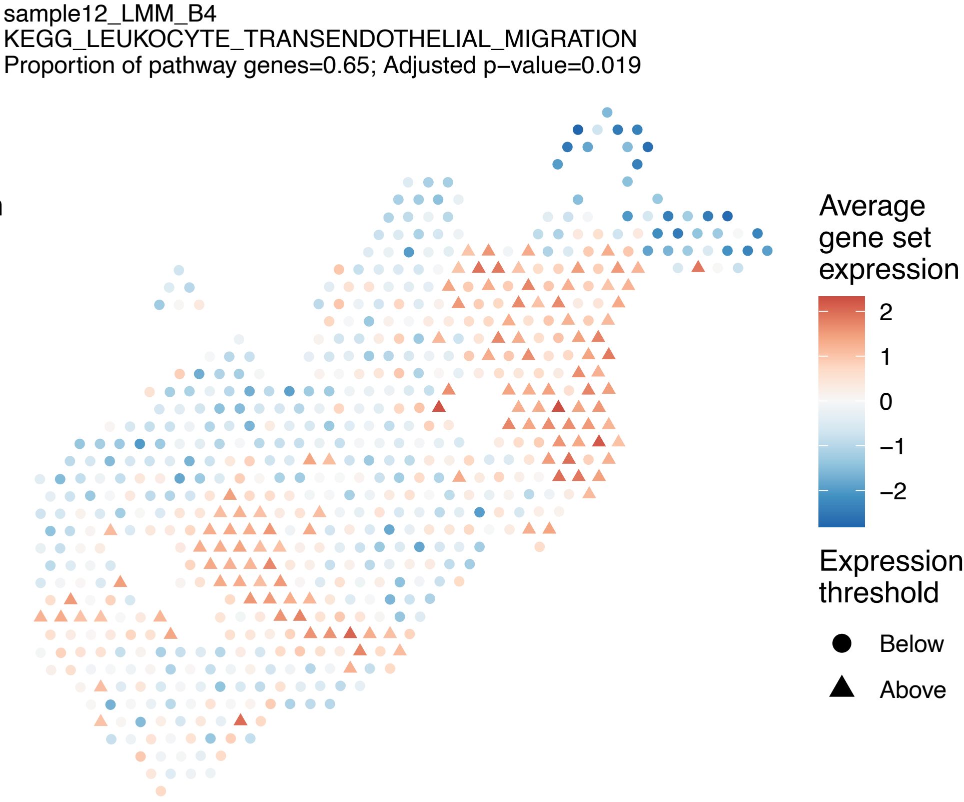
sample12_LMM_B4
KEGG_LEUKOCYTE_TRANSENDOTHELIAL_MIGRATION
Proportion of pathway genes=0.65; Adjusted p−value=0.019
Average gene set expression
2
1
0
−1
−2
Expression threshold
Below
Above

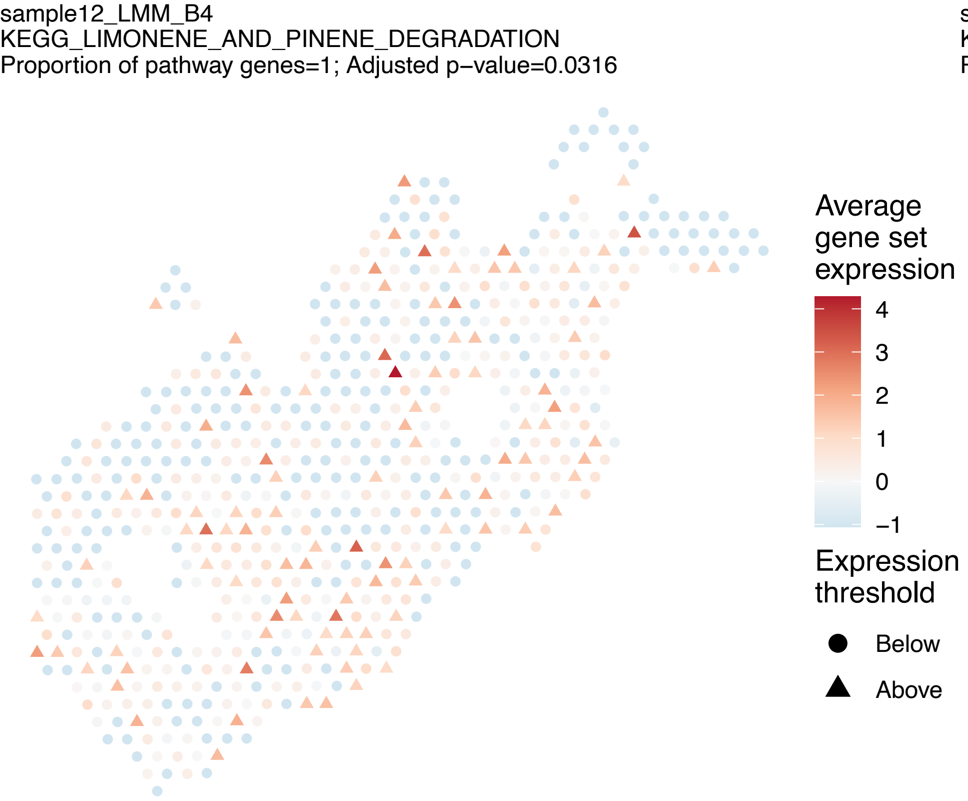
sample12_LMM_B4
KEGG_LIMONENE_AND_PINENE_DEGRADATION
Proportion of pathway genes=1; Adjusted p−value=0.0316
Average gene set expression
4
3
2
1
0
−1
Expression threshold
Below
Above

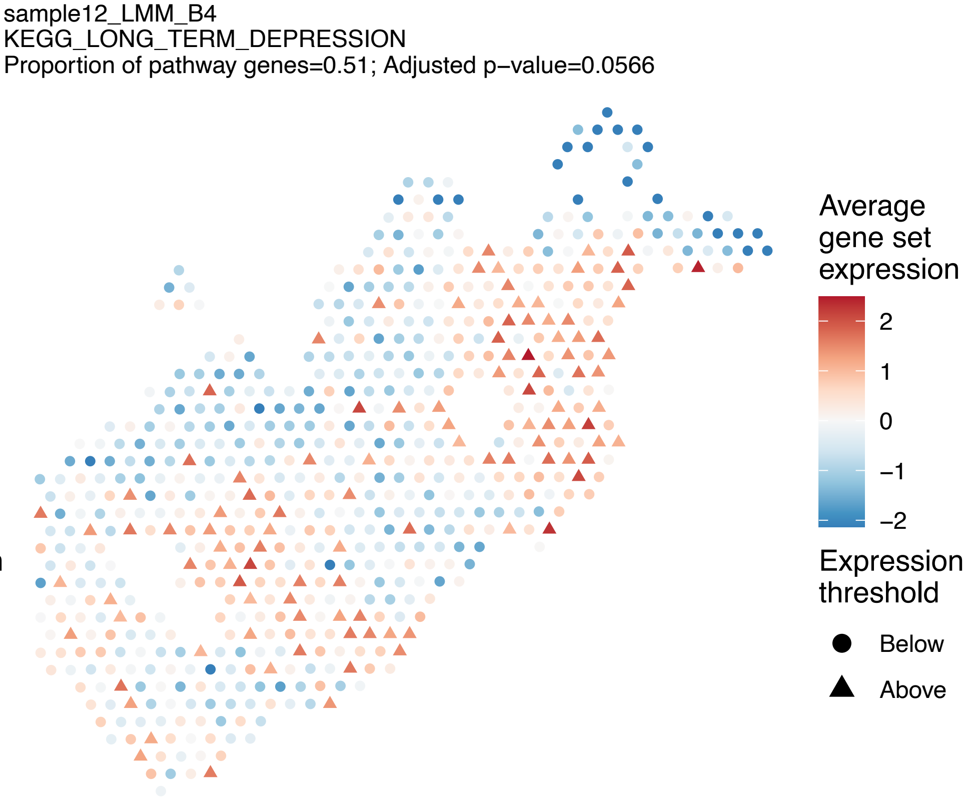
sample12_LMM_B4
KEGG_LONG_TERM_DEPRESSION
Proportion of pathway genes=0.51; Adjusted p−value=0.0566
Average gene set expression
2
1
0
−1
−2
Expression threshold
Below
Above

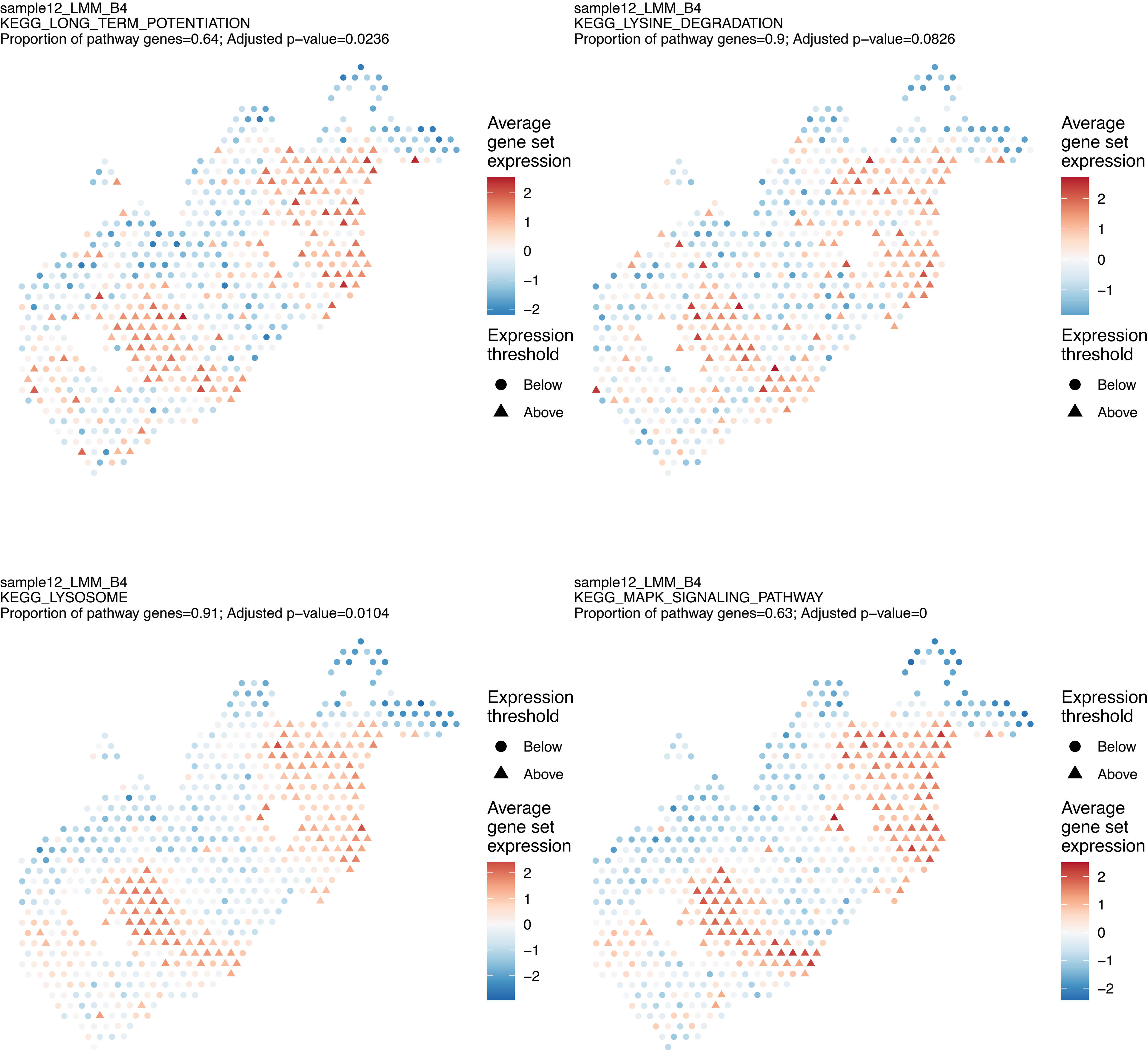
sample12_LMM_B4
KEGG_LONG_TERM_POTENTIATION
Proportion of pathway genes=0.64; Adjusted p−value=0.0236
Average gene set expression
2
1
0
−1
−2
Expression threshold
Below
Above
sample12_LMM_B4
KEGG_LYSINE_DEGRADATION
Proportion of pathway genes=0.9; Adjusted p−value=0.0826
Average gene set expression
2
1
0
−1
Expression threshold
Below
Above
sample12_LMM_B4
KEGG_LYSOSOME
Proportion of pathway genes=0.91; Adjusted p−value=0.0104
Expression threshold
Below
Above
Average gene set expression
2
1
0
−1
−2
sample12_LMM_B4
KEGG_MAPK_SIGNALING_PATHWAY
Proportion of pathway genes=0.63; Adjusted p−value=0
Expression threshold
Below
Above
Average gene set expression
2
1
0
−1
−2

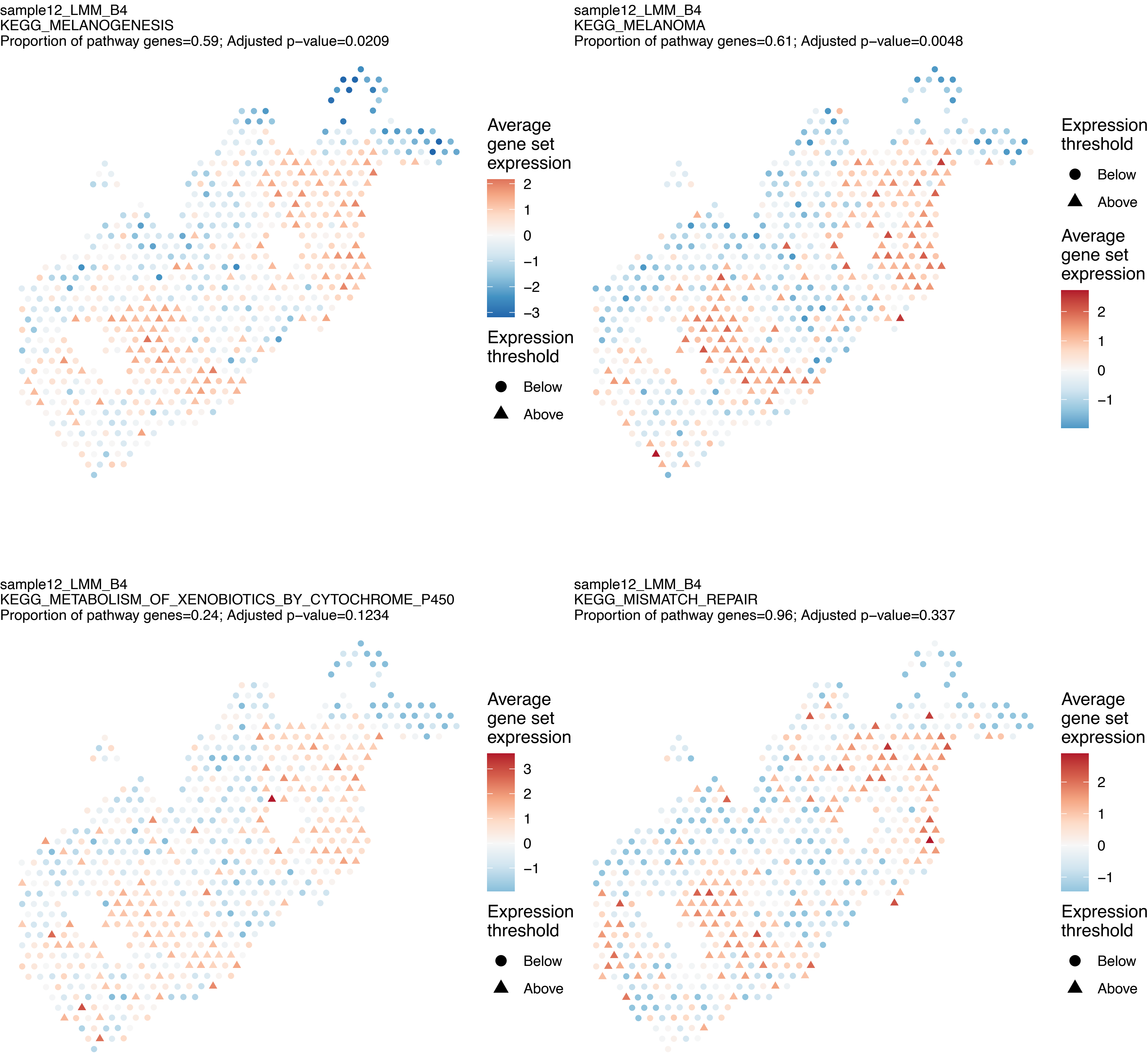

sample12_LMM_B4
KEGG_MELANOGENESIS
Proportion of pathway genes=0.59; Adjusted p−value=0.0209
Average gene set expression
2
1
0
−1
−2
−3
Expression threshold
Below
Above
sample12_LMM_B4
KEGG_MELANOMA
Proportion of pathway genes=0.61; Adjusted p−value=0.0048
Expression threshold
Below
Above
Average gene set expression
2
1
0
−1
sample12_LMM_B4
KEGG_METABOLISM_OF_XENOBIOTICS_BY_CYTOCHROME_P450
Proportion of pathway genes=0.24; Adjusted p−value=0.1234
Average gene set expression
3
2
1
0
−1
Expression threshold
Below
Above
sample12_LMM_B4
KEGG_MISMATCH_REPAIR
Proportion of pathway genes=0.96; Adjusted p−value=0.337
Average gene set expression
2
1
0
−1
Expression threshold
Below
Above

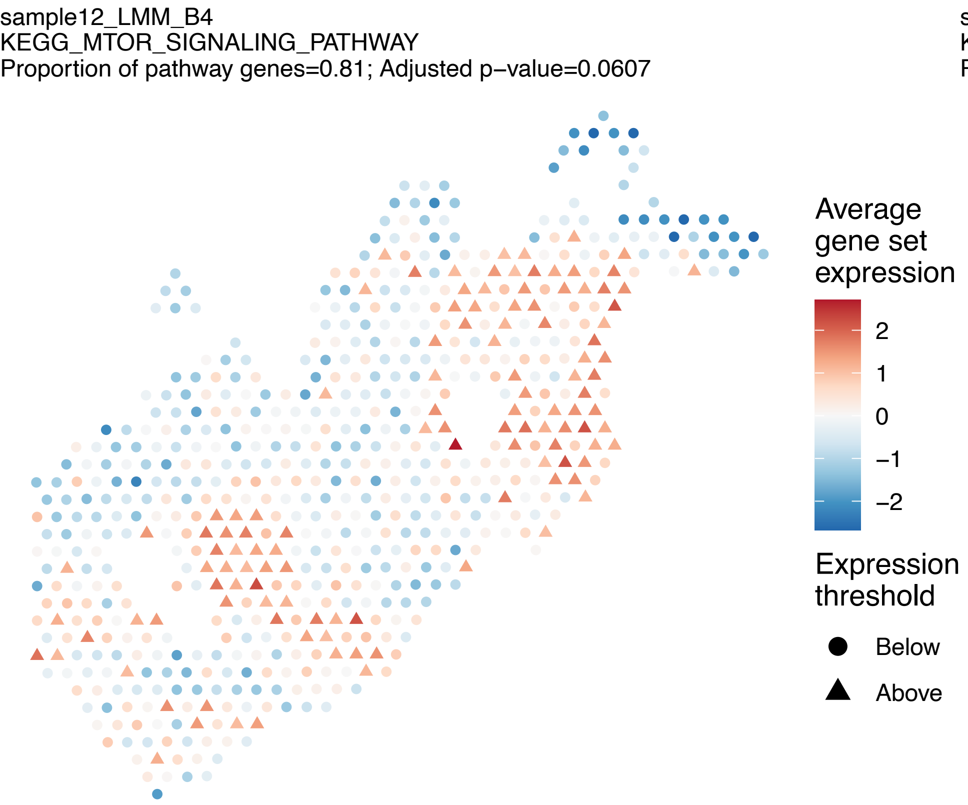
sample12_LMM_B4
KEGG_MTOR_SIGNALING_PATHWAY
Proportion of pathway genes=0.81; Adjusted p−value=0.0607
Average gene set expression
2
1
0
−1
−2
Expression threshold
Below
Above

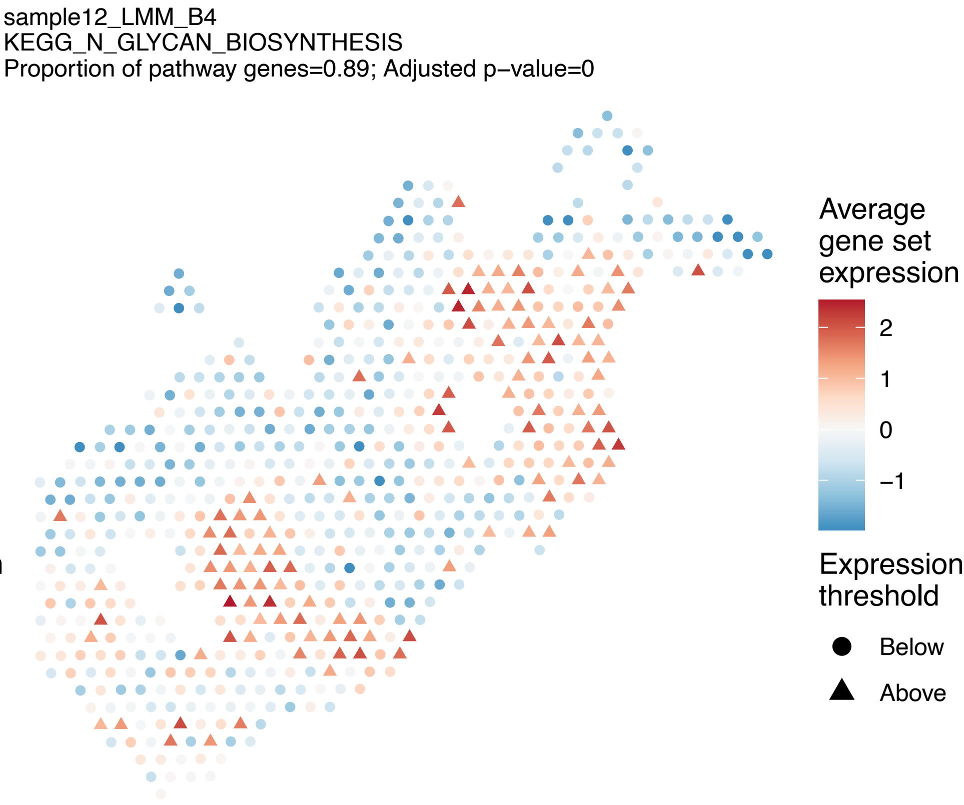
sample12_LMM_B4
KEGG_N_GLYCAN_BIOSYNTHESIS
Proportion of pathway genes=0.89; Adjusted p−value=0
Average gene set expression
2
1
0
−1
Expression threshold
Below
Above

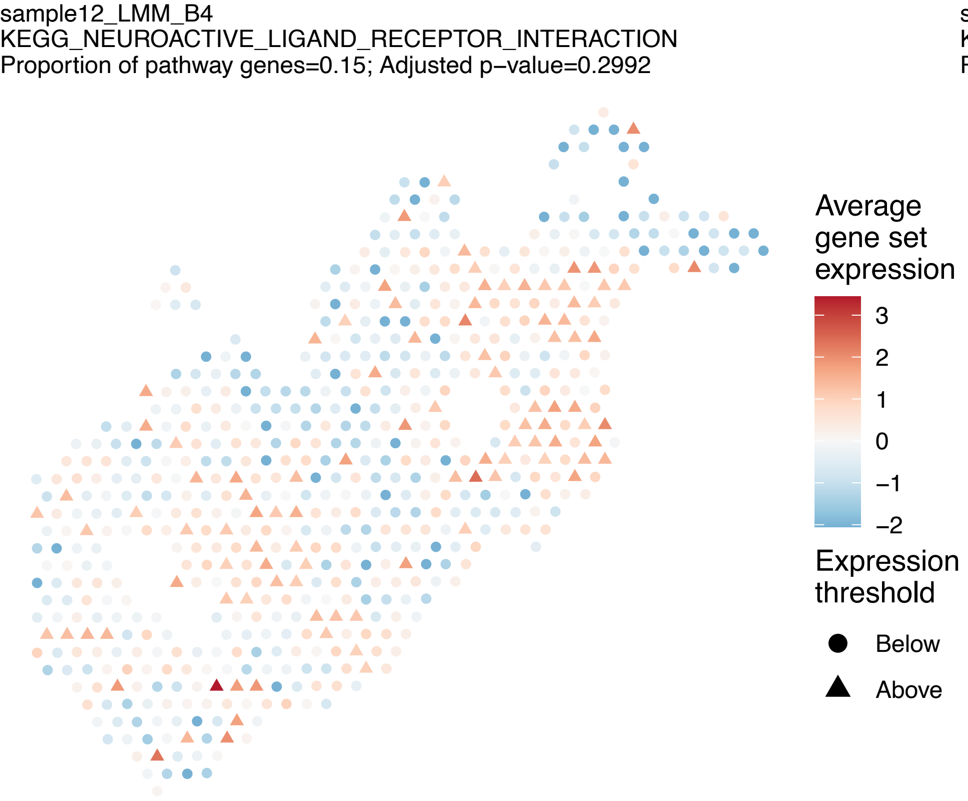
sample12_LMM_B4
KEGG_NEUROACTIVE_LIGAND_RECEPTOR_INTERACTION
Proportion of pathway genes=0.15; Adjusted p−value=0.2992
Average gene set expression
3
2
1
0
−1
−2
Expression threshold
Below
Above

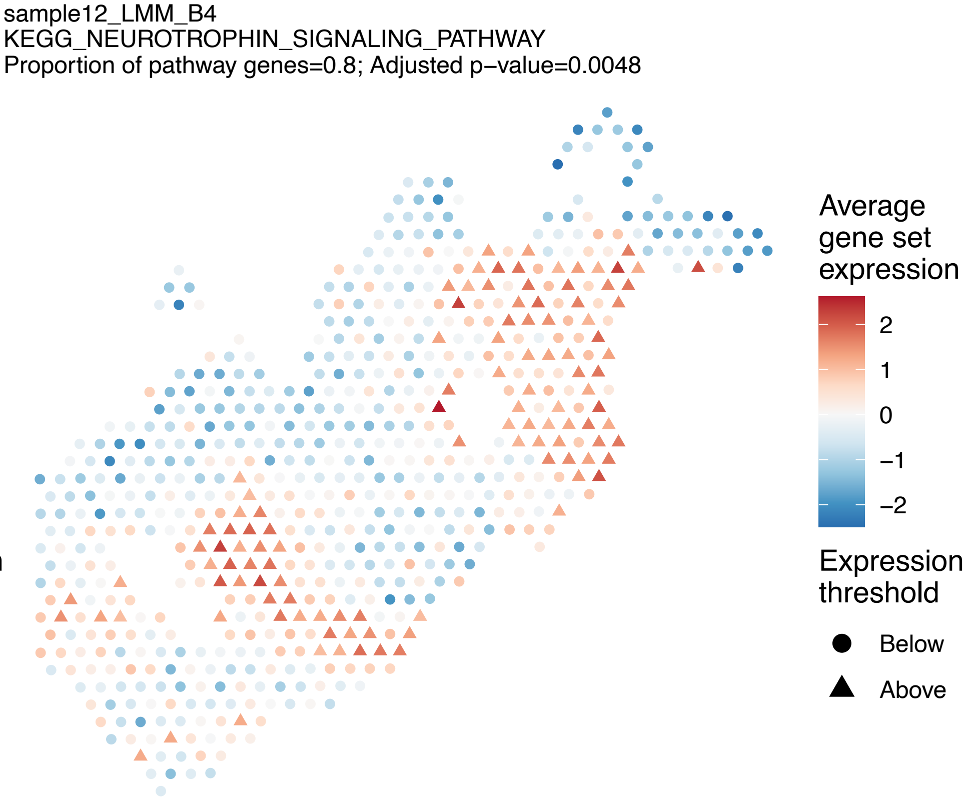
sample12_LMM_B4
KEGG_NEUROTROPHIN_SIGNALING_PATHWAY
Proportion of pathway genes=0.8; Adjusted p−value=0.0048
Average gene set expression
2
1
0
−1
−2
Expression threshold
Below
Above

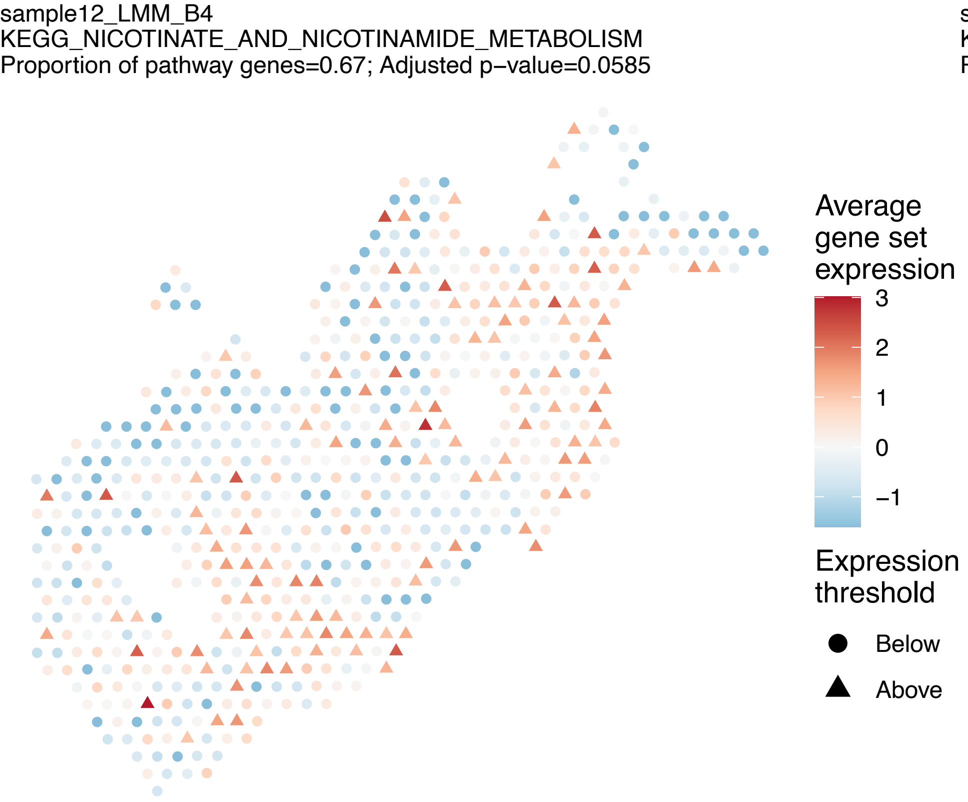
sample12_LMM_B4
KEGG_NICOTINATE_AND_NICOTINAMIDE_METABOLISM
Proportion of pathway genes=0.67; Adjusted p−value=0.0585
Average gene set expression
3
2
1
0
−1
Expression threshold
Below
Above

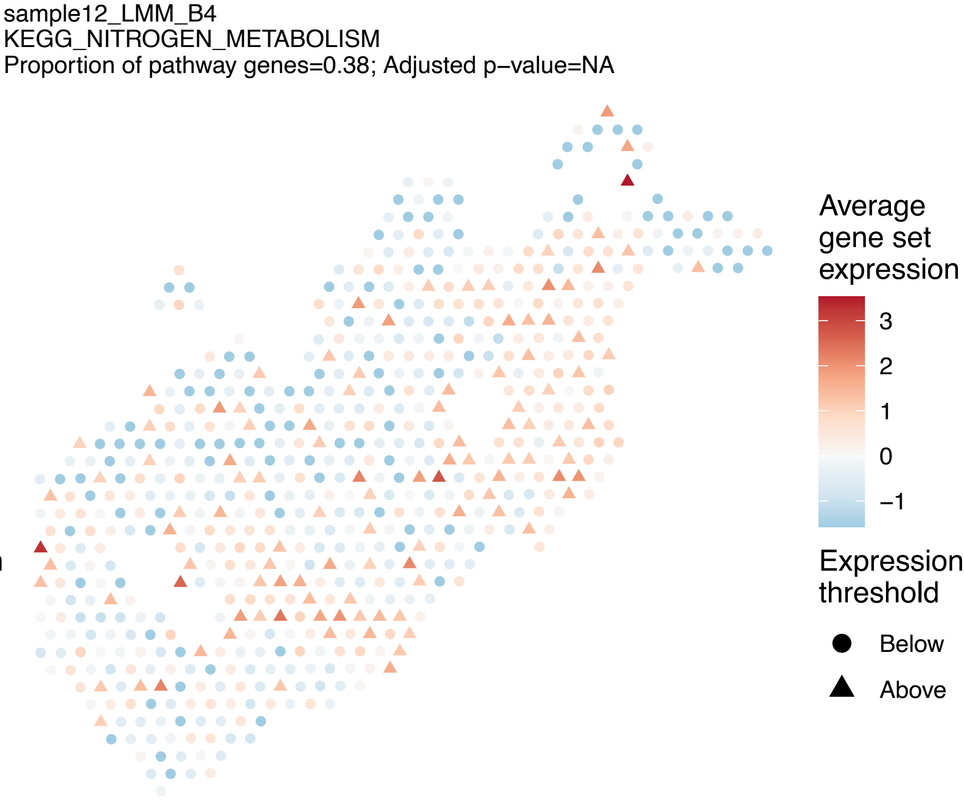
sample12_LMM_B4
KEGG_NITROGEN_METABOLISM
Proportion of pathway genes=0.38; Adjusted p−value=NA
Average gene set expression
3
2
1
0
−1
Expression threshold
Below
Above

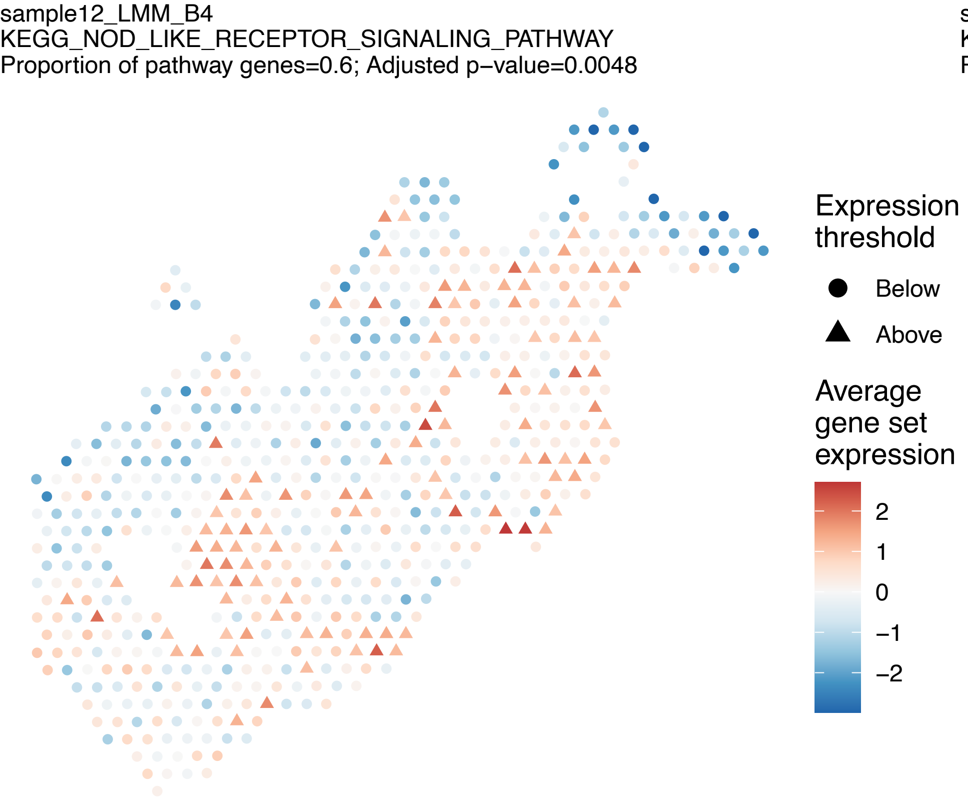
sample12_LMM_B4
KEGG_NOD_LIKE_RECEPTOR_SIGNALING_PATHWAY
Proportion of pathway genes=0.6; Adjusted p−value=0.0048
Expression threshold
Below
Above
Average gene set expression
2
1
0
−1
−2

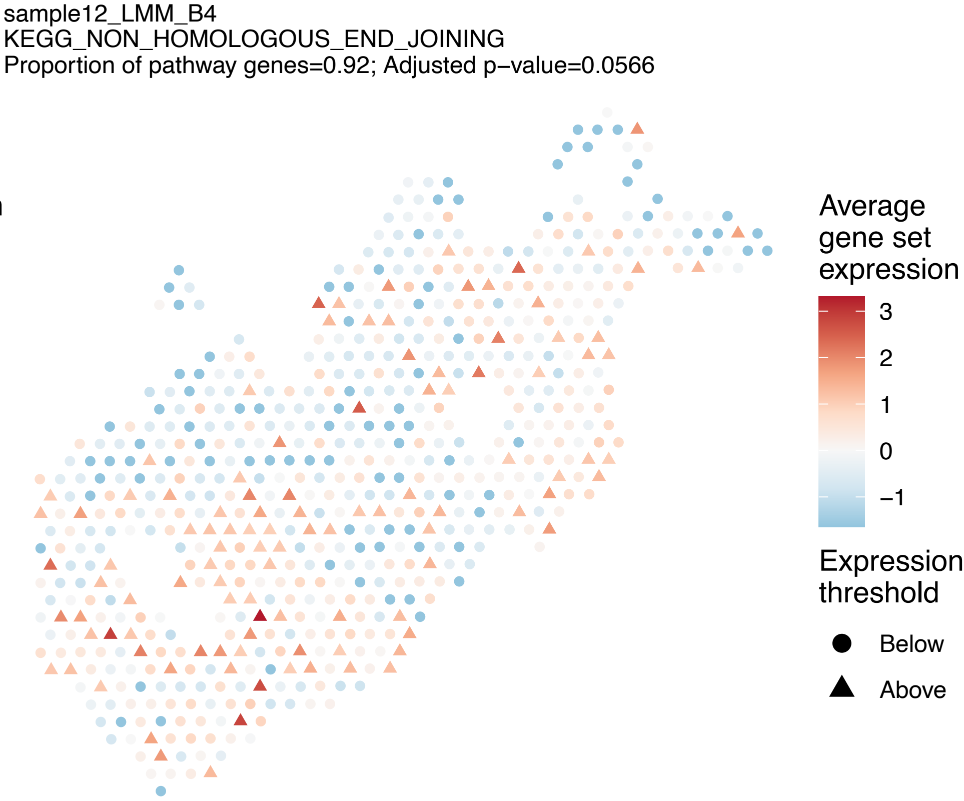
sample12_LMM_B4
KEGG_NON_HOMOLOGOUS_END_JOINING
Proportion of pathway genes=0.92; Adjusted p−value=0.0566
Average gene set expression
3
2
1
0
−1
Expression threshold
Below
Above

sample12_LMM_B4
KEGG_NON_SMALL_CELL_LUNG_CANCER
Proportion of pathway genes=0.87; Adjusted p−value=0.0048

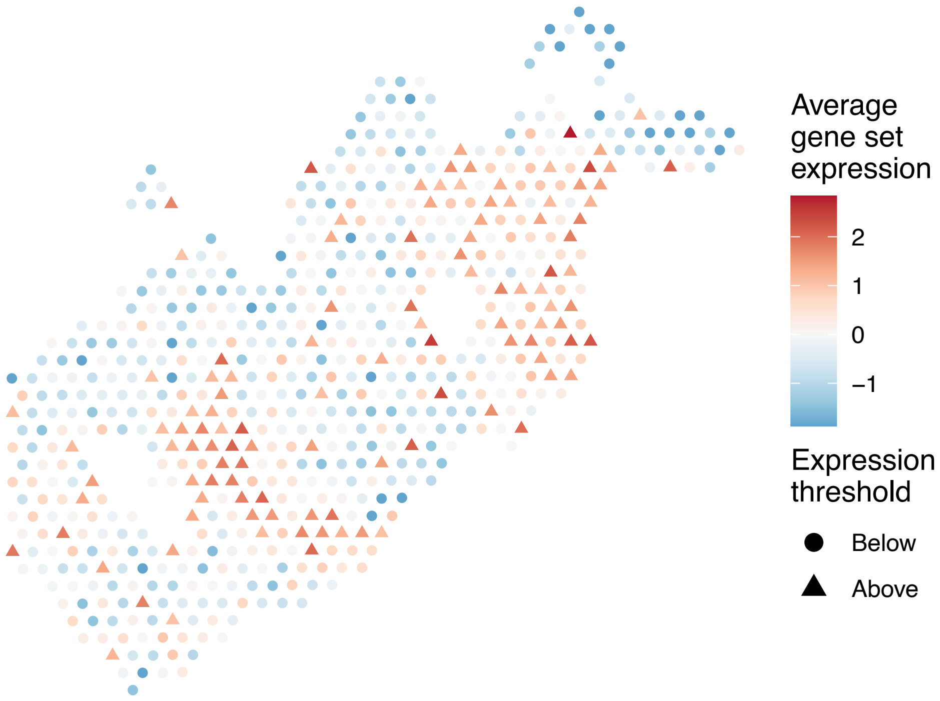


sample12_LMM_B4
KEGG_NOTCH_SIGNALING_PATHWAY
Proportion of pathway genes=0.82; Adjusted p−value=0.019

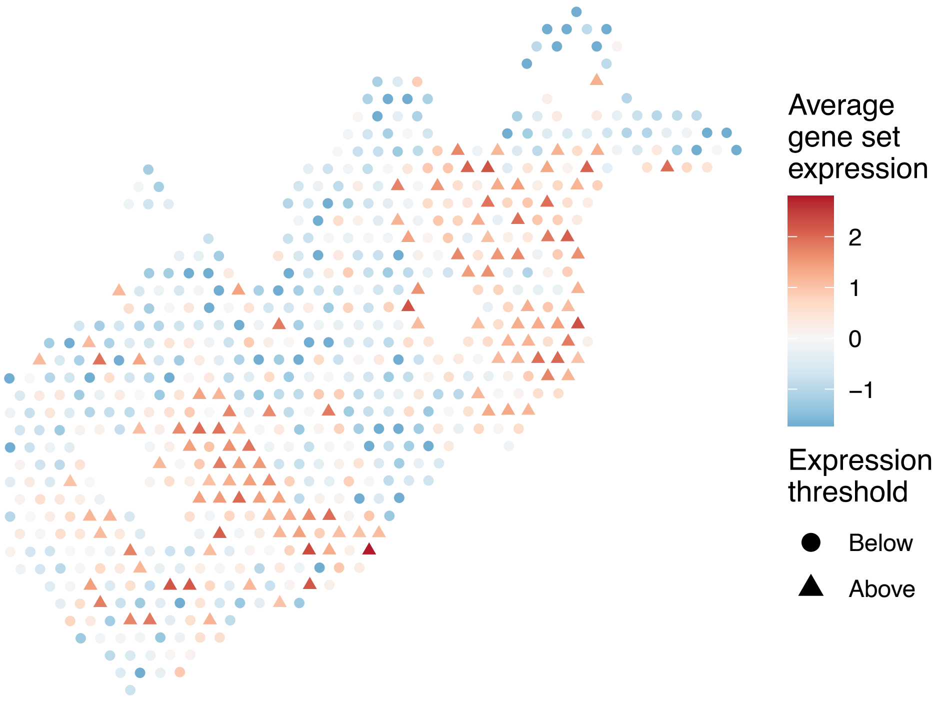


sample12_LMM_B4
KEGG_NUCLEOTIDE_EXCISION_REPAIR
Proportion of pathway genes=0.96; Adjusted p−value=0.0104

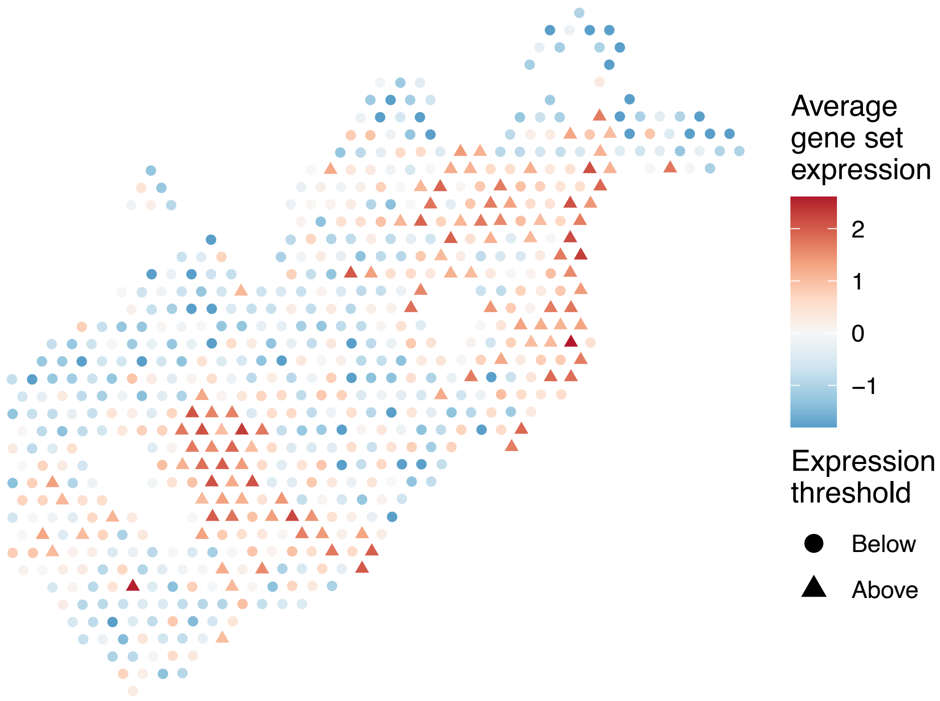


sample12_LMM_B4
KEGG_O_GLYCAN_BIOSYNTHESIS
Proportion of pathway genes=0.48; Adjusted p−value=0.1132

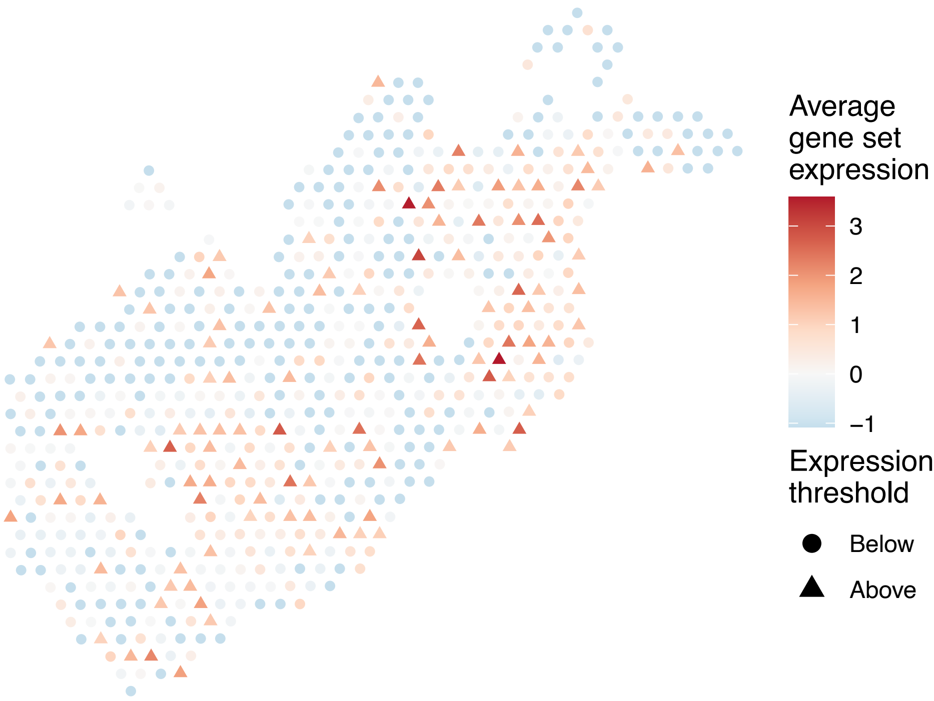

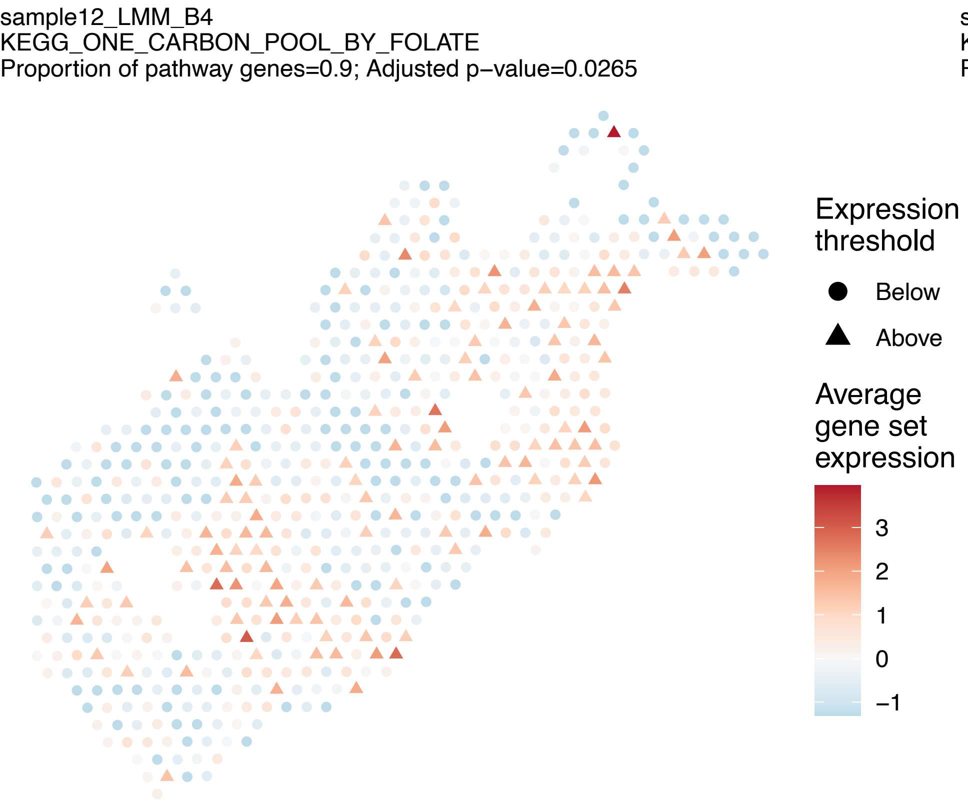
sample12_LMM_B4
KEGG_ONE_CARBON_POOL_BY_FOLATE
Proportion of pathway genes=0.9; Adjusted p−value=0.0265
Expression threshold
Below
Above
Average gene set expression
3
2
1
0
−1

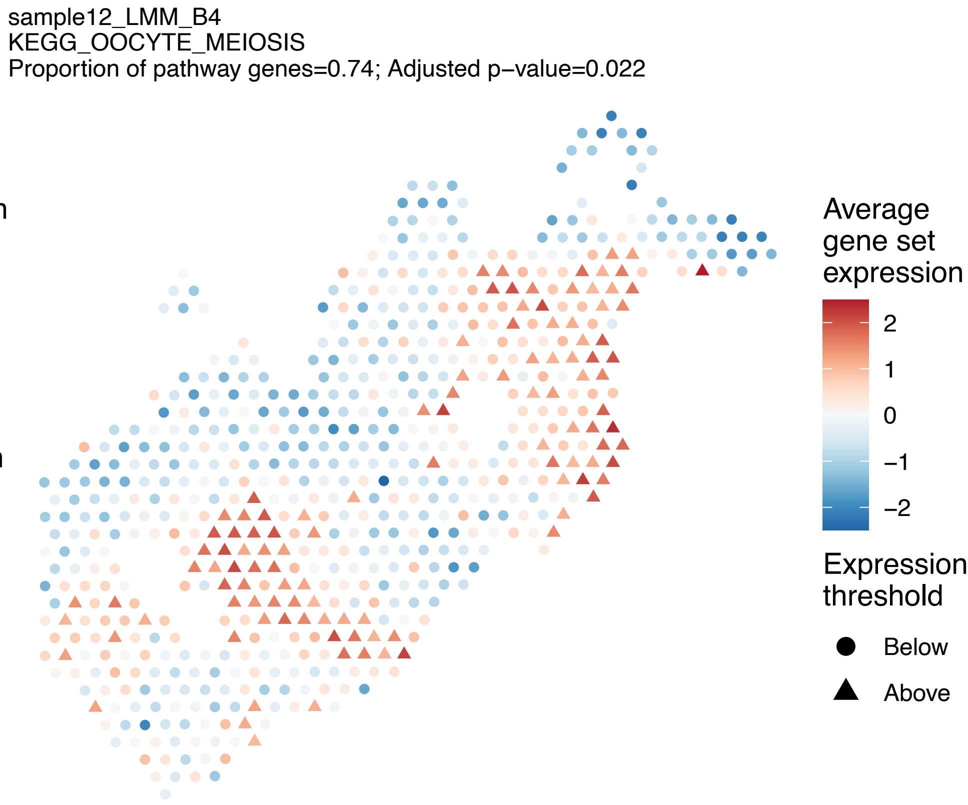
sample12_LMM_B4
KEGG_OOCYTE_MEIOSIS
Proportion of pathway genes=0.74; Adjusted p−value=0.022
Average gene set expression
2
1
0
−1
−2
Expression threshold
Below
Above

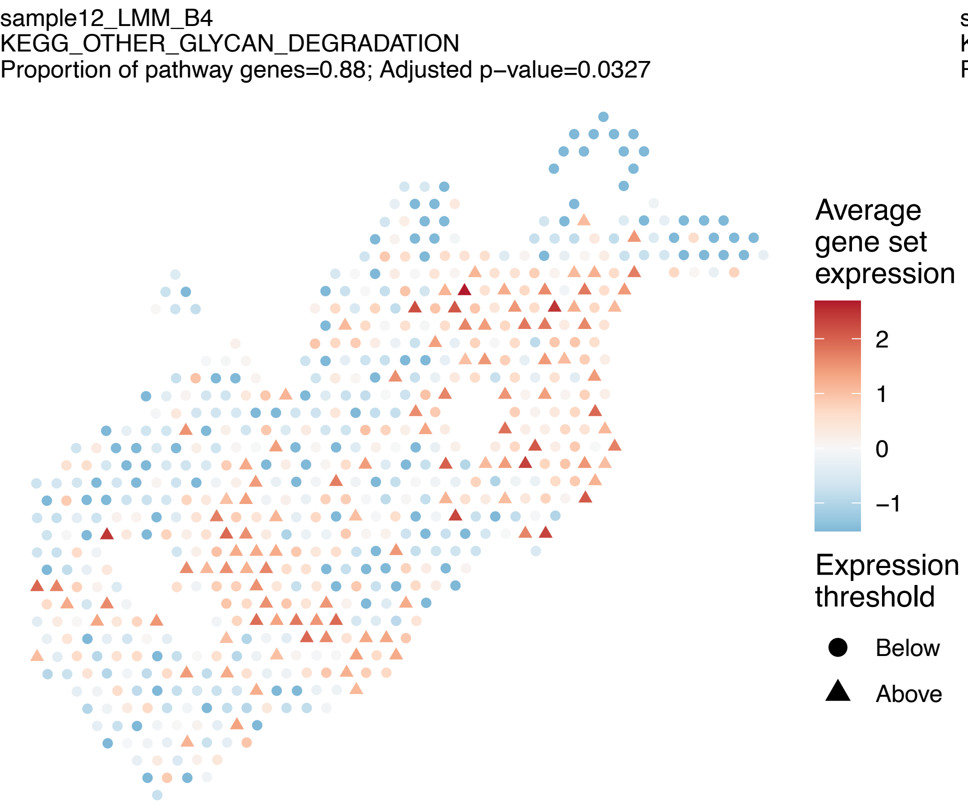
sample12_LMM_B4
KEGG_OTHER_GLYCAN_DEGRADATION
Proportion of pathway genes=0.88; Adjusted p−value=0.0327
Average gene set expression
2
1
0
−1
Expression threshold
Below
Above

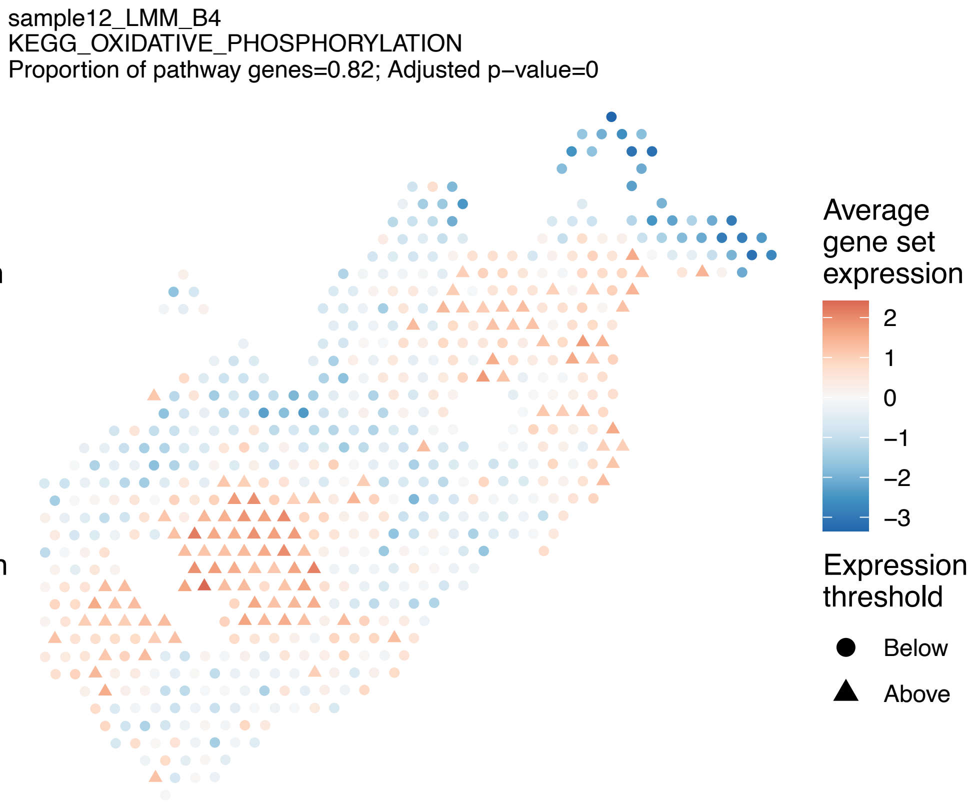
sample12_LMM_B4
KEGG_OXIDATIVE_PHOSPHORYLATION
Proportion of pathway genes=0.82; Adjusted p−value=0
Average gene set expression
2
1
0
−1
−2
−3
Expression threshold
Below
Above

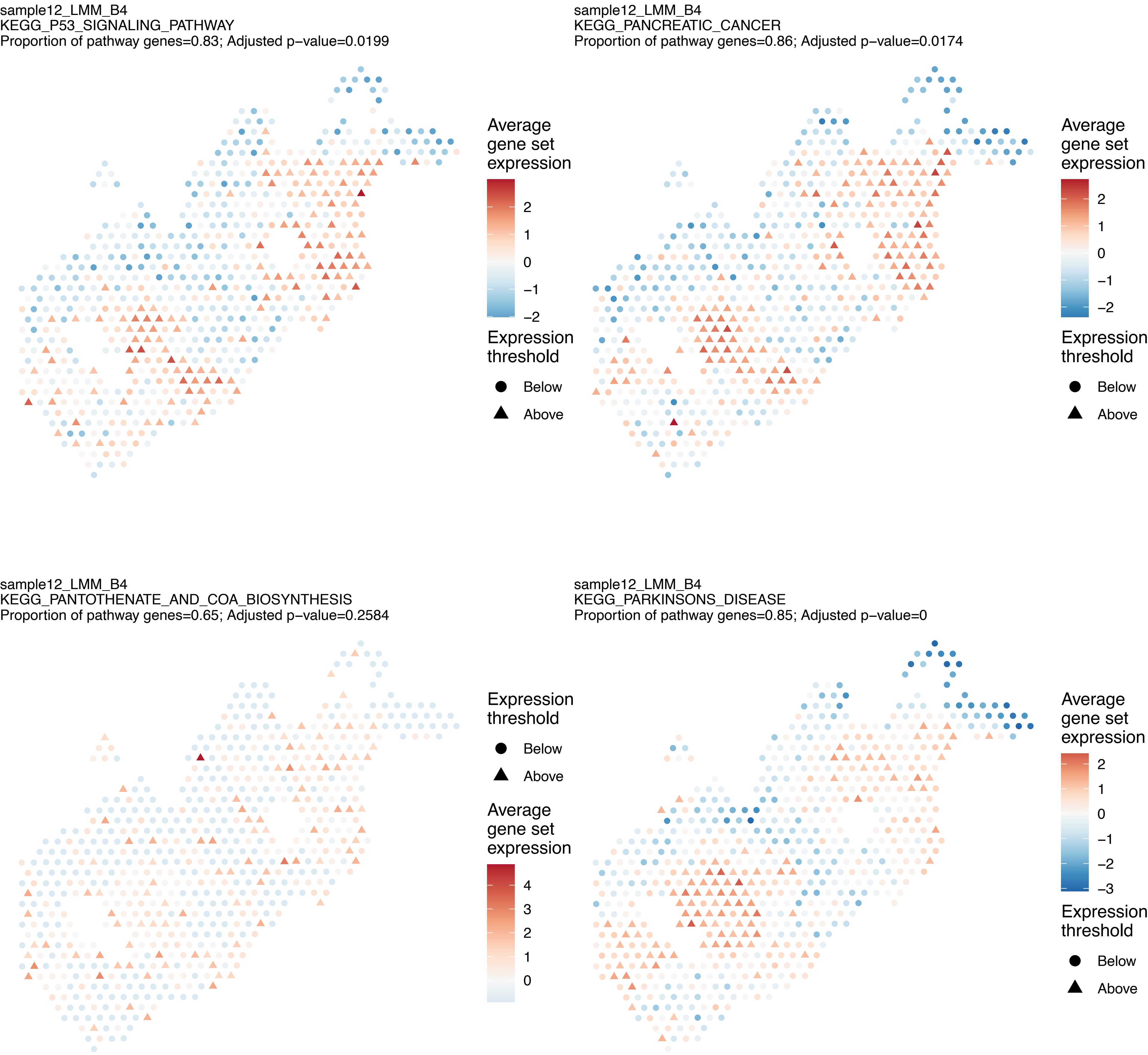
sample12_LMM_B4
KEGG_P53_SIGNALING_PATHWAY
Proportion of pathway genes=0.83; Adjusted p−value=0.0199
Average gene set expression
2
1
0
−1
−2
Expression threshold
Below
Above
sample12_LMM_B4
KEGG_PANCREATIC_CANCER
Proportion of pathway genes=0.86; Adjusted p−value=0.0174
Average gene set expression
2
1
0
−1
−2
Expression threshold
Below
Above
sample12_LMM_B4
KEGG_PANTOTHENATE_AND_COA_BIOSYNTHESIS
Proportion of pathway genes=0.65; Adjusted p−value=0.2584
Expression threshold
Below
Above
Average gene set expression
4
3
2
1
0
sample12_LMM_B4
KEGG_PARKINSONS_DISEASE
Proportion of pathway genes=0.85; Adjusted p−value=0
Average gene set expression
2
1
0
−1
−2
−3
Expression threshold
Below
Above

sample12_LMM_B4
KEGG_PATHOGENIC_ESCHERICHIA_COLI_INFECTION
Proportion of pathway genes=0.75; Adjusted p−value=0.2324

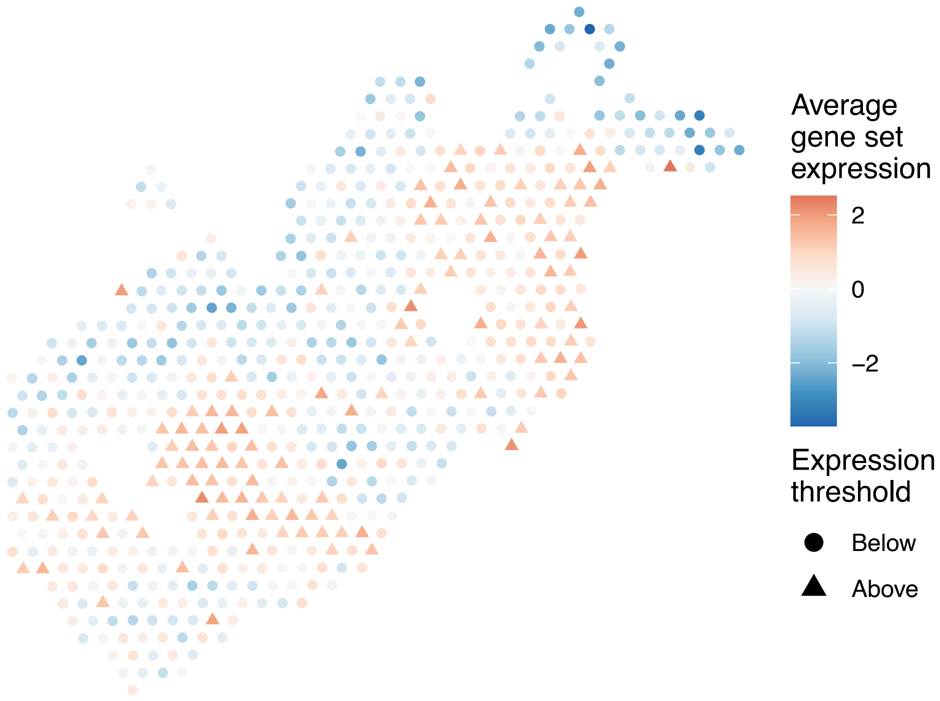


sample12_LMM_B4
KEGG_PATHWAYS_IN_CANCER
Proportion of pathway genes=0.7; Adjusted p−value=0

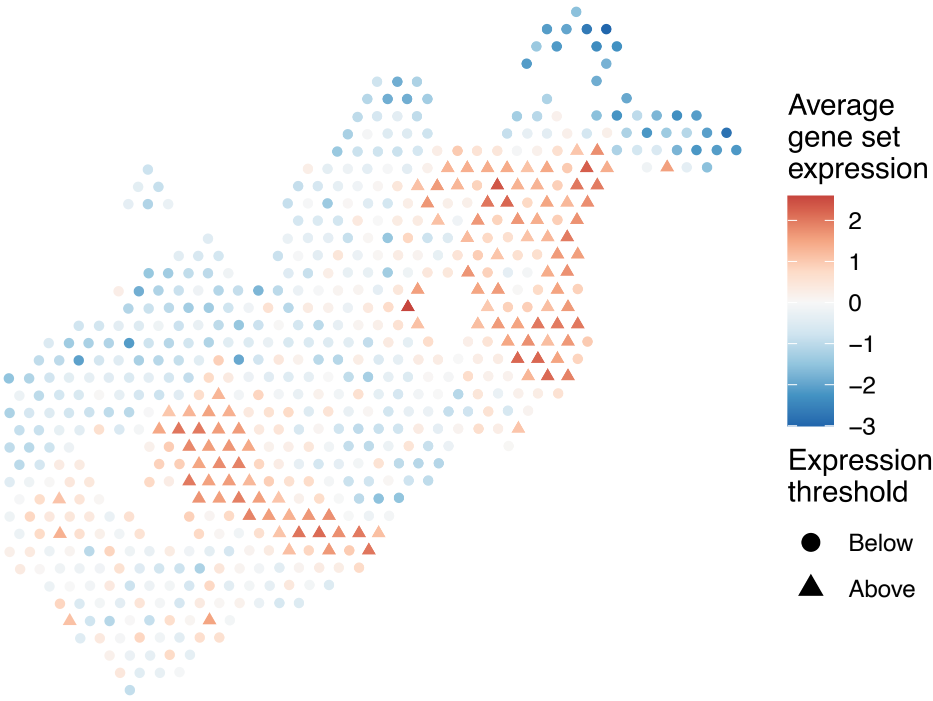


sample12_LMM_B4
KEGG_PENTOSE_PHOSPHATE_PATHWAY
Proportion of pathway genes=0.75; Adjusted p−value=0.0246

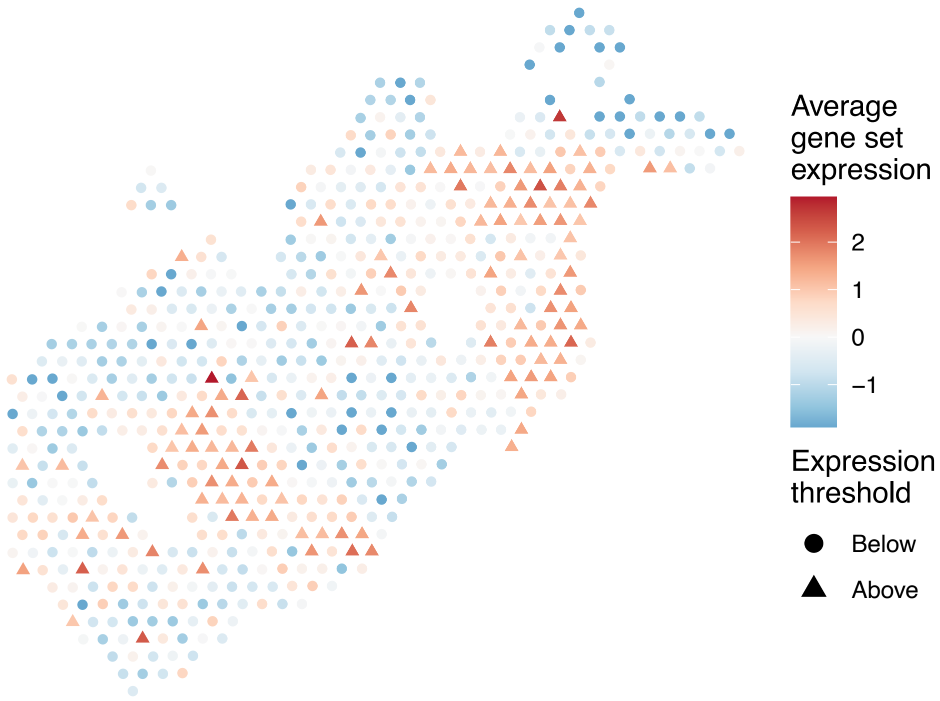


sample12_LMM_B4
KEGG_PEROXISOME
Proportion of pathway genes=0.77; Adjusted p−value=0.0302

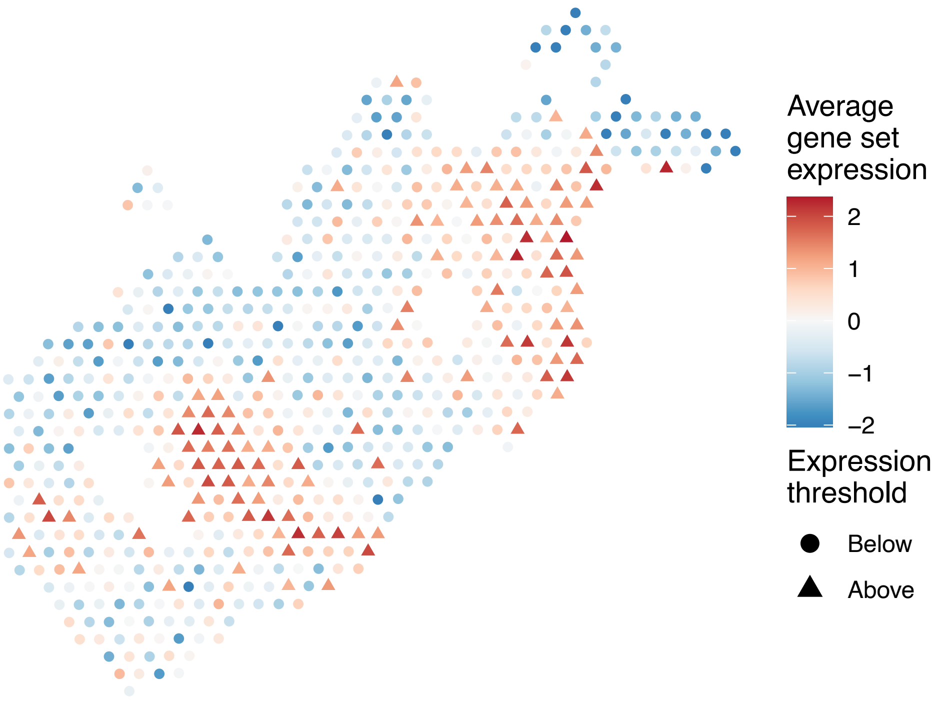

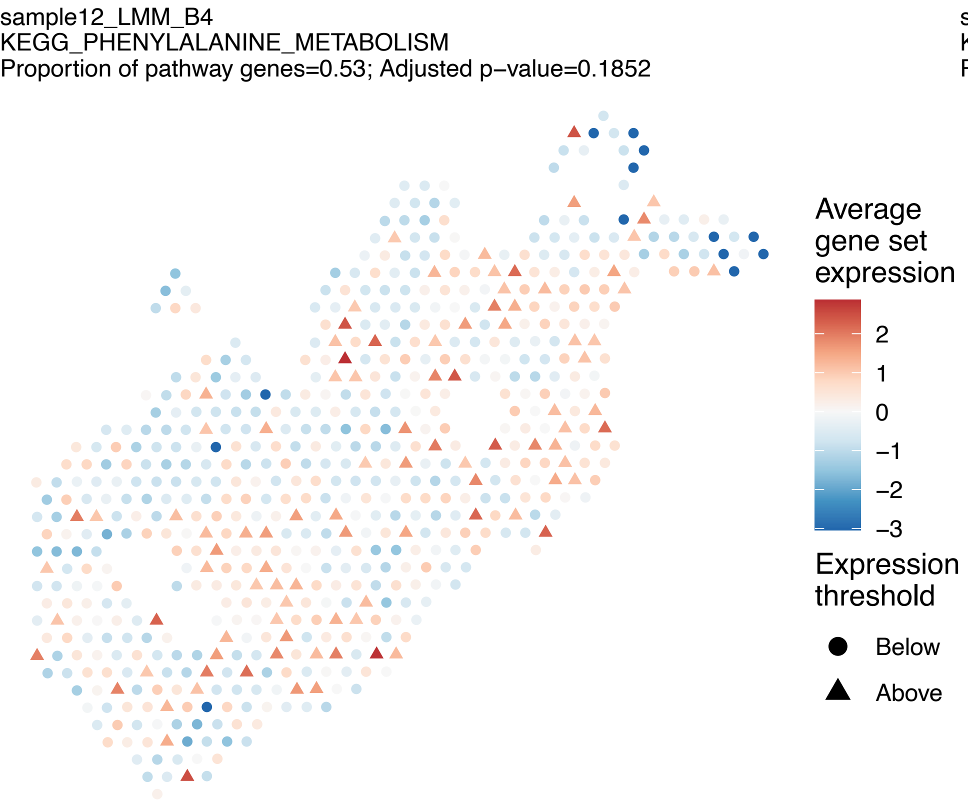
sample12_LMM_B4
KEGG_PHENYLALANINE_METABOLISM
Proportion of pathway genes=0.53; Adjusted p−value=0.1852
Average gene set expression
2
1
0
−1
−2
−3
Expression threshold
Below
Above

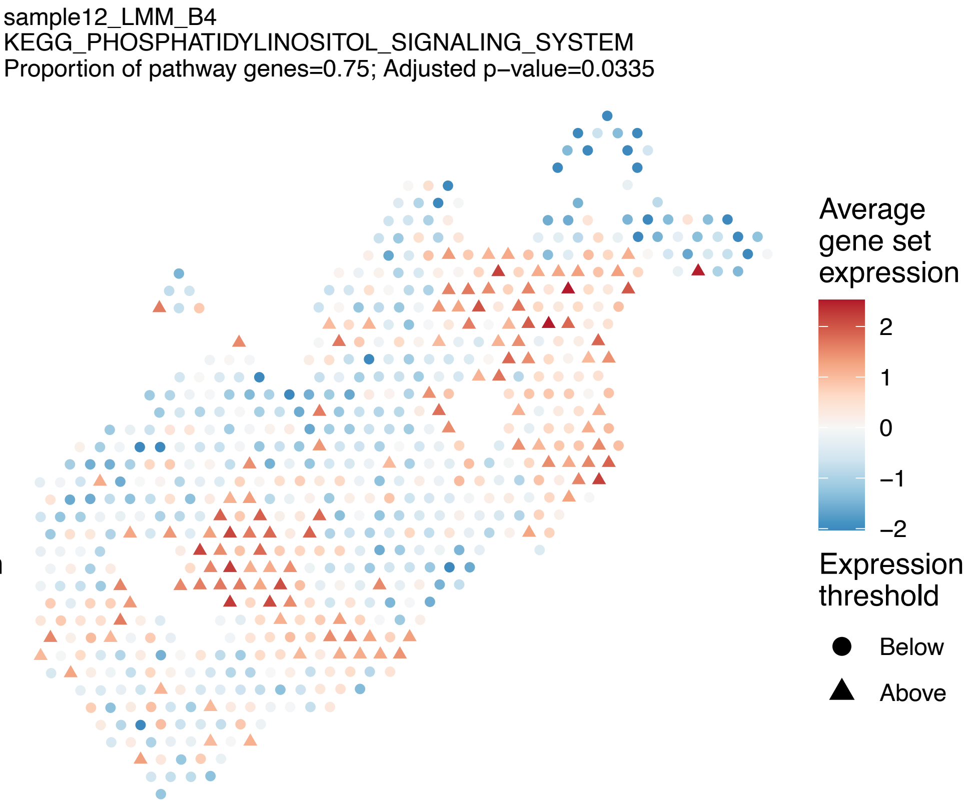
sample12_LMM_B4
KEGG_PHOSPHATIDYLINOSITOL_SIGNALING_SYSTEM
Proportion of pathway genes=0.75; Adjusted p−value=0.0335
Average gene set expression
2
1
0
−1
−2
Expression threshold
Below
Above

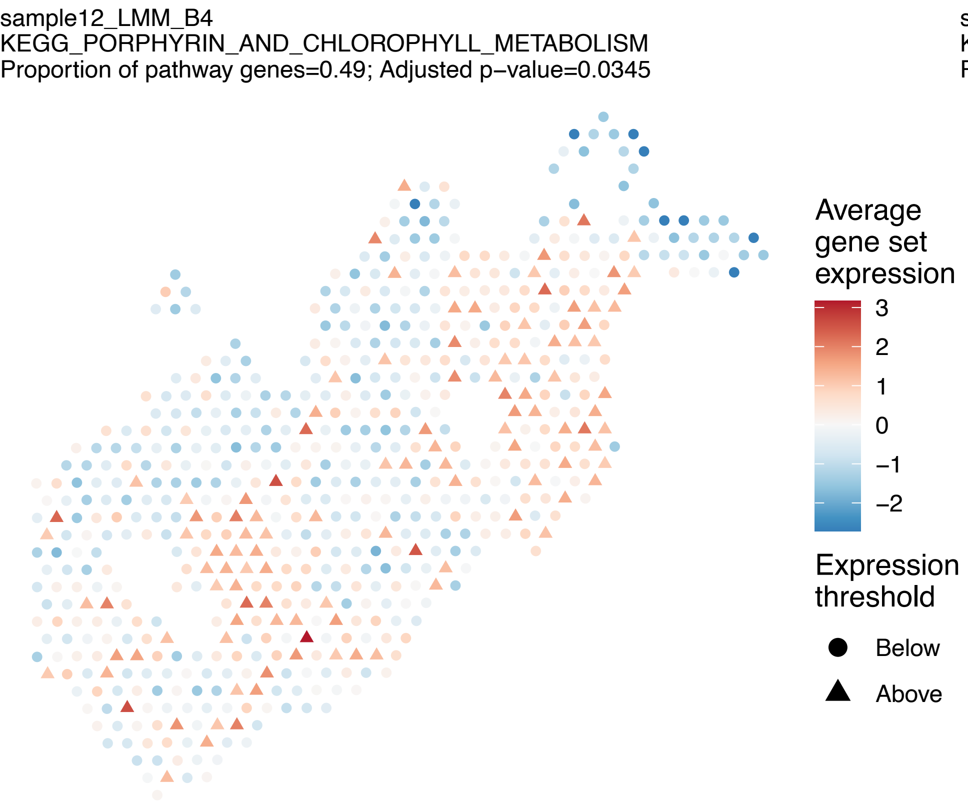
sample12_LMM_B4
KEGG_PORPHYRIN_AND_CHLOROPHYLL_METABOLISM
Proportion of pathway genes=0.49; Adjusted p−value=0.0345
Average gene set expression
3
2
1
0
−1
−2
Expression threshold
Below
Above

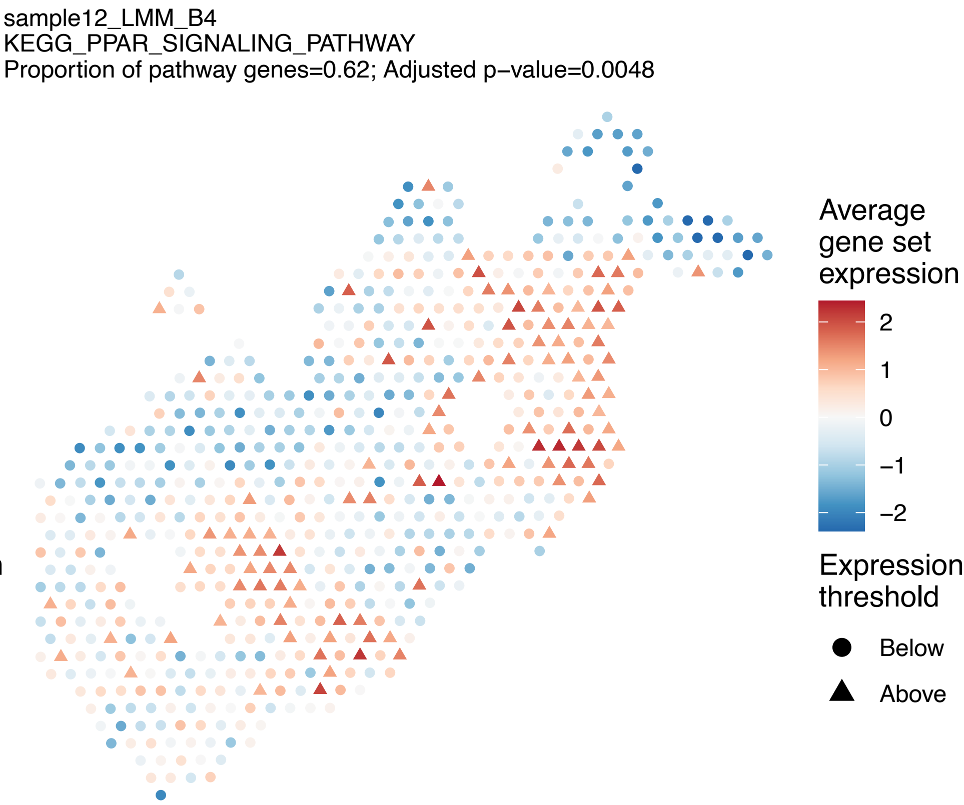
sample12_LMM_B4
KEGG_PPAR_SIGNALING_PATHWAY
Proportion of pathway genes=0.62; Adjusted p−value=0.0048
Average gene set expression
2
1
0
−1
−2
Expression threshold
Below
Above

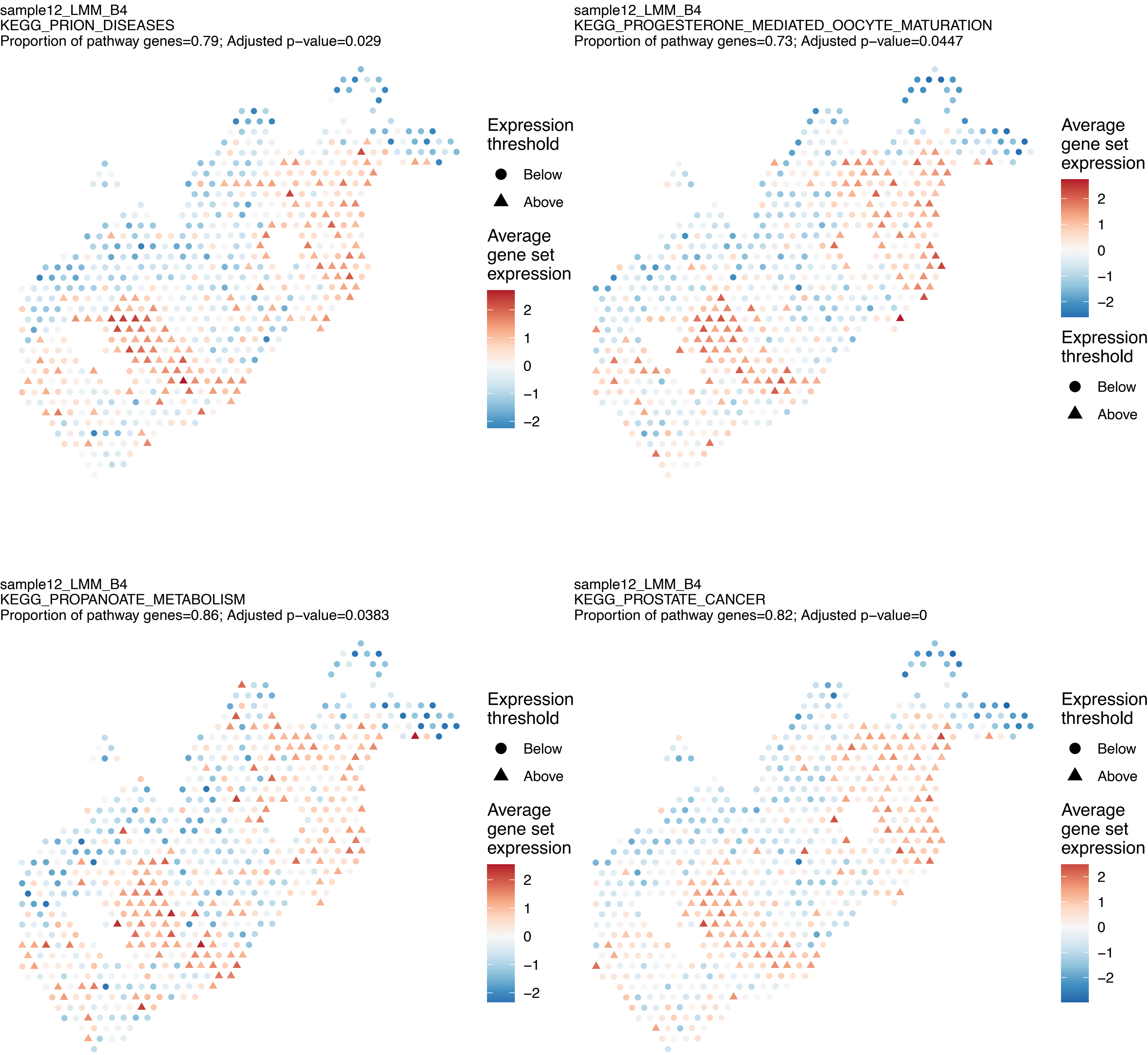
sample12_LMM_B4
KEGG_PRION_DISEASES
Proportion of pathway genes=0.79; Adjusted p−value=0.029
Expression threshold
Below
Above
Average gene set expression
2
1
0
−1
−2
sample12_LMM_B4
KEGG_PROGESTERONE_MEDIATED_OOCYTE_MATURATION
Proportion of pathway genes=0.73; Adjusted p−value=0.0447
Average gene set expression
2
1
0
−1
−2
Expression threshold
Below
Above
sample12_LMM_B4
KEGG_PROPANOATE_METABOLISM
Proportion of pathway genes=0.86; Adjusted p−value=0.0383
Expression threshold
Below
Above
Average gene set expression
2
1
0
−1
−2
sample12_LMM_B4
KEGG_PROSTATE_CANCER
Proportion of pathway genes=0.82; Adjusted p−value=0
Expression threshold
Below
Above
Average gene set expression
2
1
0
−1
−2

sample12_LMM_B4
KEGG_PROTEASOME
Proportion of pathway genes=0.92; Adjusted p−value=0.0435

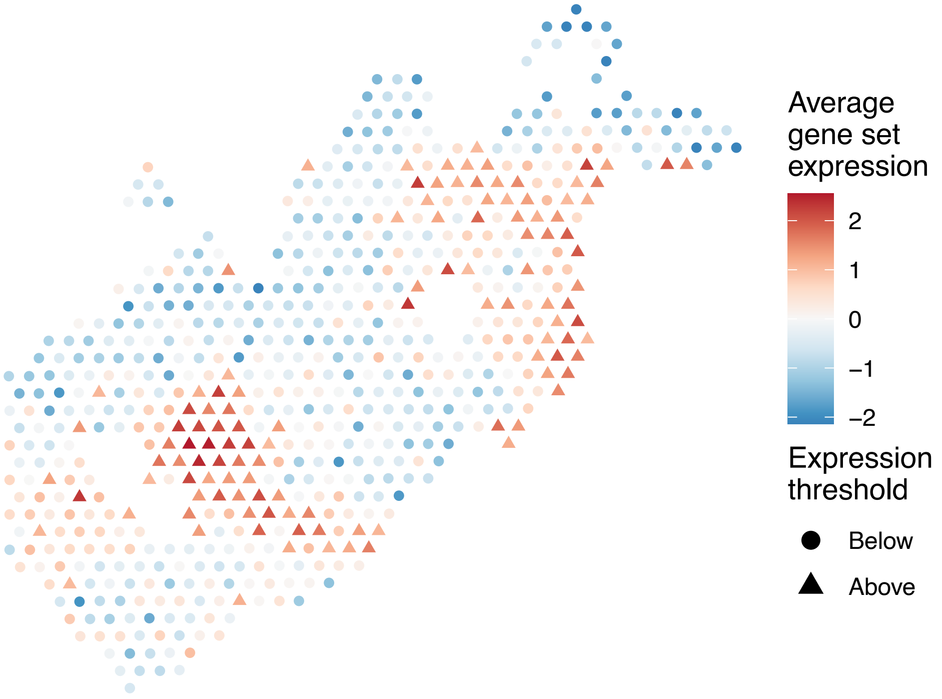


sample12_LMM_B4
KEGG_PROTEIN_EXPORT
Proportion of pathway genes=0.96; Adjusted p−value=0.0174

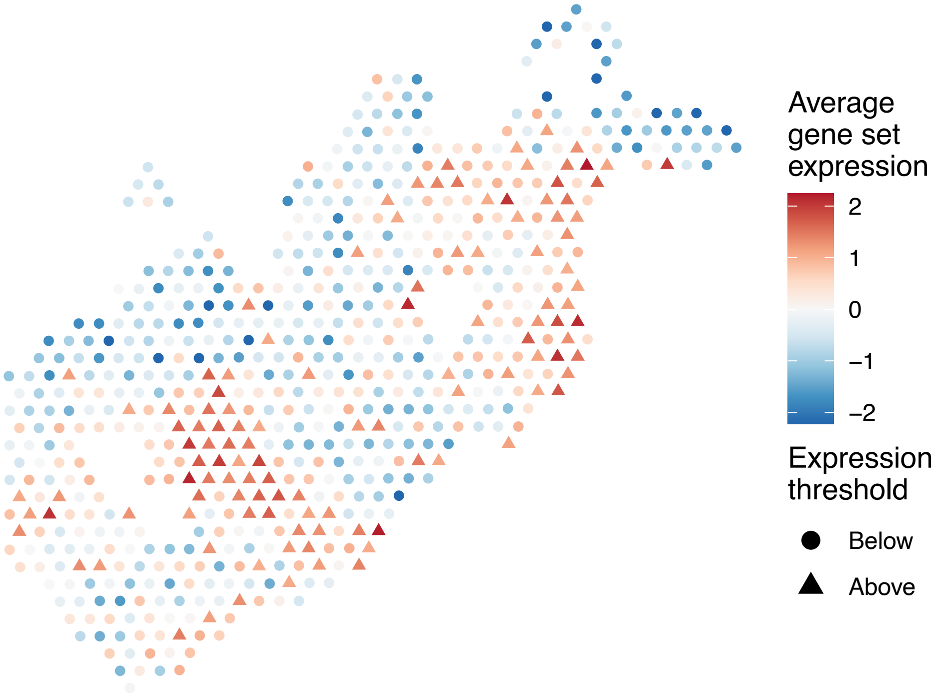


sample12_LMM_B4
KEGG_PROXIMAL_TUBULE_BICARBONATE_RECLAMATION
Proportion of pathway genes=0.54; Adjusted p−value=0.0495

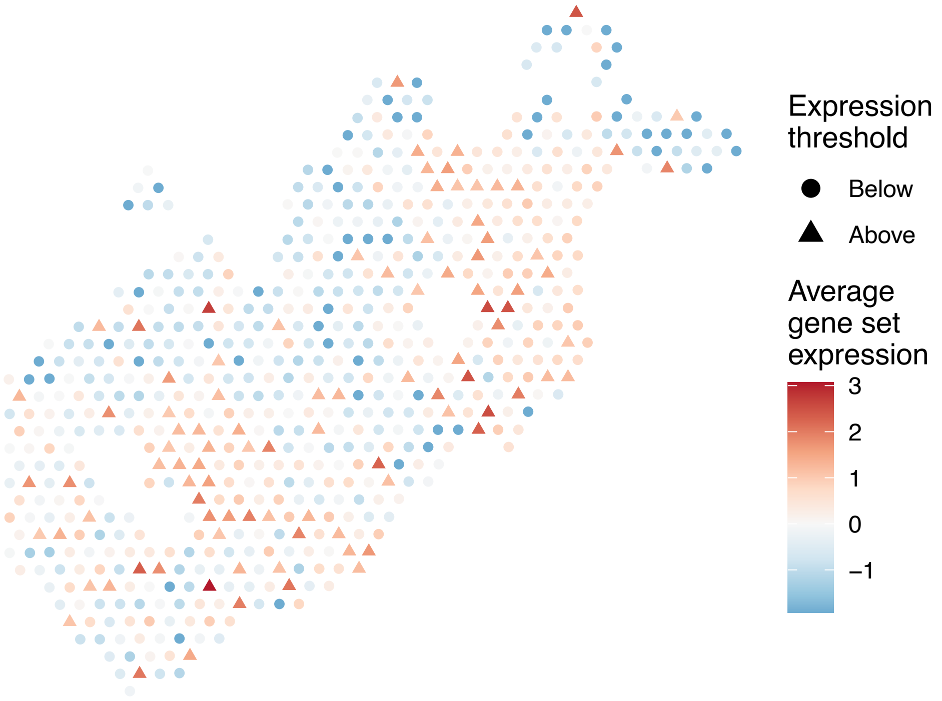


sample12_LMM_B4
KEGG_PURINE_METABOLISM
Proportion of pathway genes=0.69; Adjusted p−value=0.0048

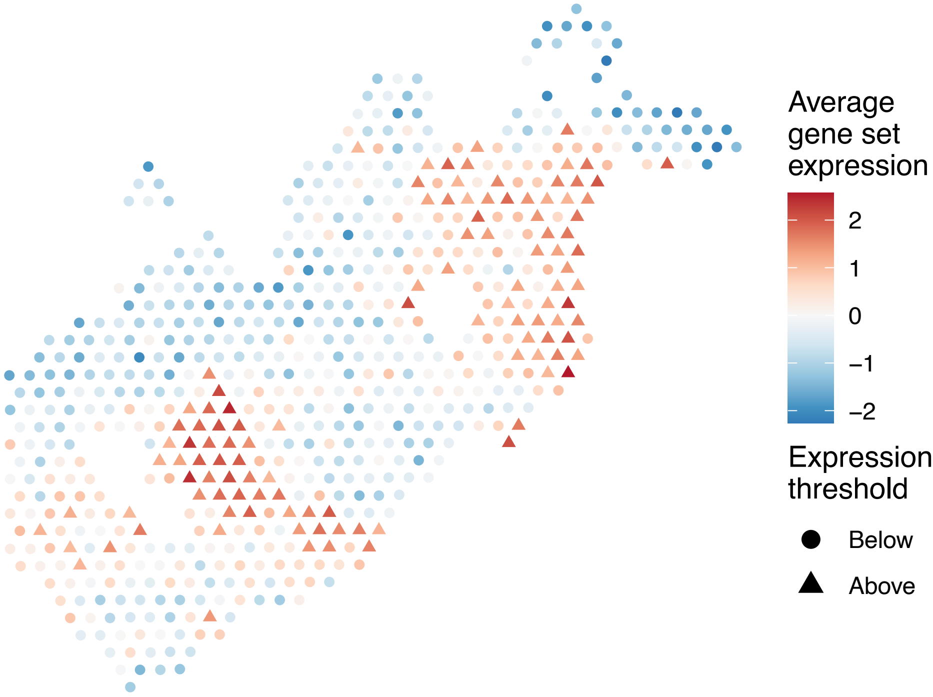

sample12_LMM_B4
KEGG_PYRIMIDINE_METABOLISM
Proportion of pathway genes=0.8; Adjusted p−value=0.0265

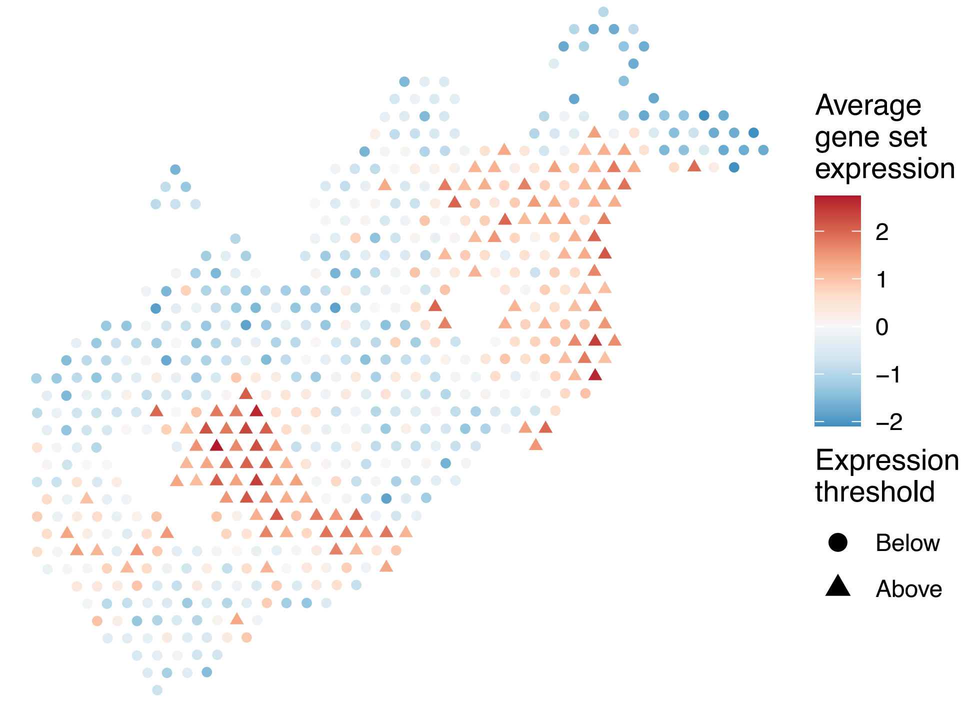


sample12_LMM_B4
KEGG_PYRUVATE_METABOLISM
Proportion of pathway genes=0.82; Adjusted p−value=0.0302

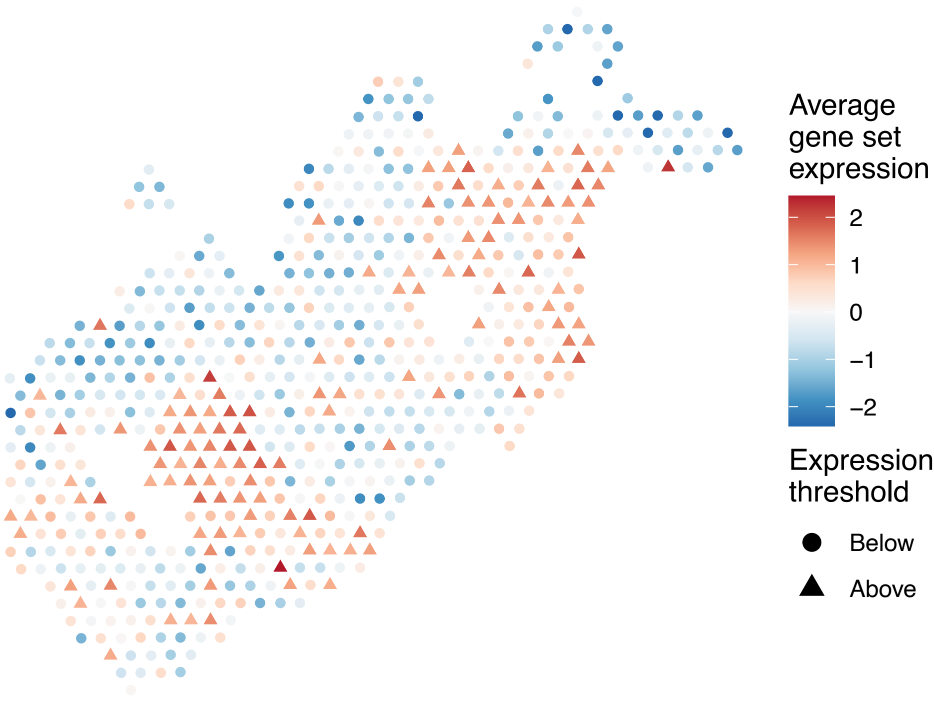


sample12_LMM_B4
KEGG_REGULATION_OF_ACTIN_CYTOSKELETON
Proportion of pathway genes=0.7; Adjusted p−value=0

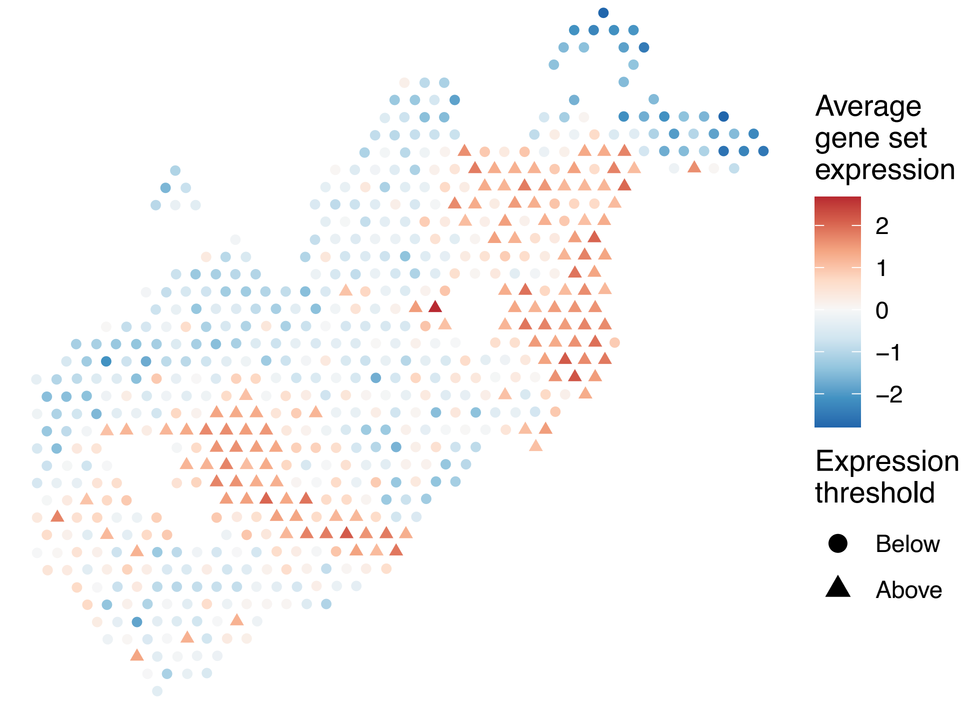


sample12_LMM_B4
KEGG_REGULATION_OF_AUTOPHAGY
Proportion of pathway genes=0.51; Adjusted p−value=0.0535

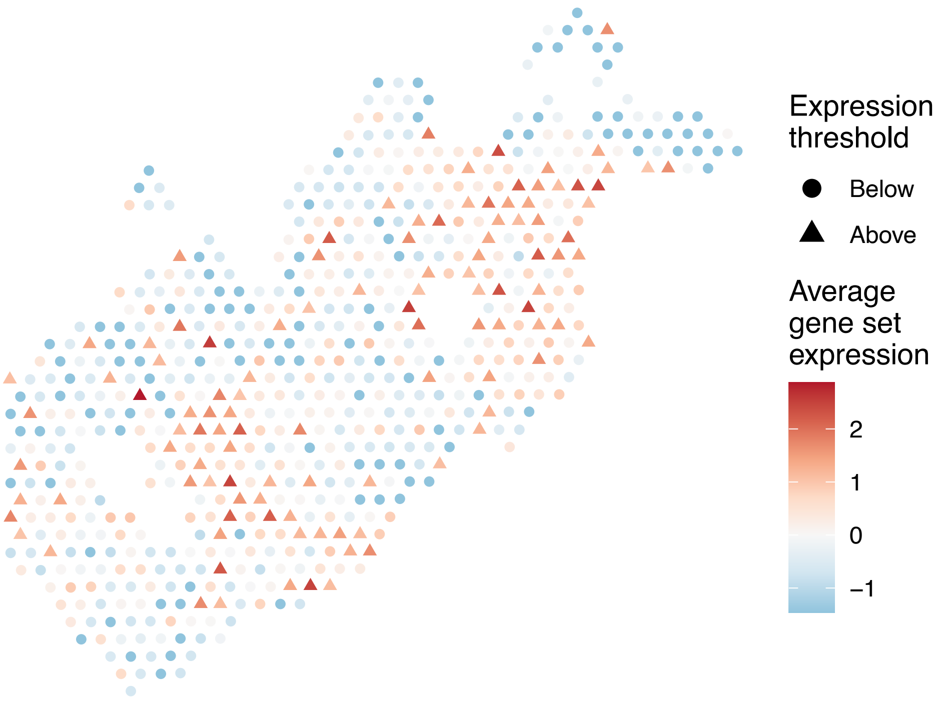

sample12_LMM_B4
KEGG_RENAL_CELL_CARCINOMA
Proportion of pathway genes=0.86; Adjusted p−value=0

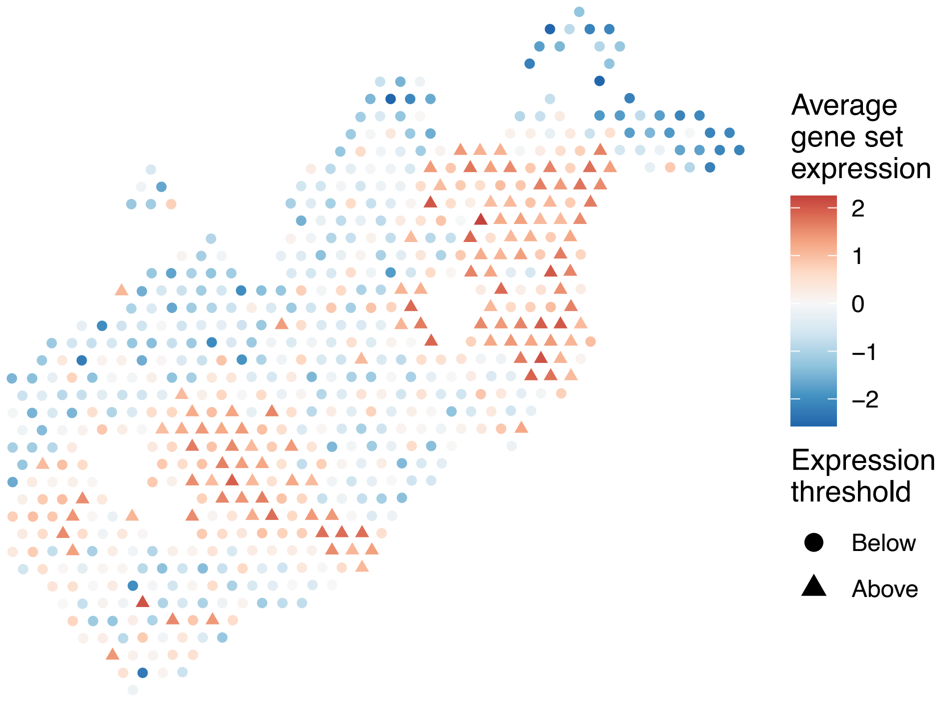


sample12_LMM_B4
KEGG_RETINOL_METABOLISM
Proportion of pathway genes=0.25; Adjusted p−value=0.1112

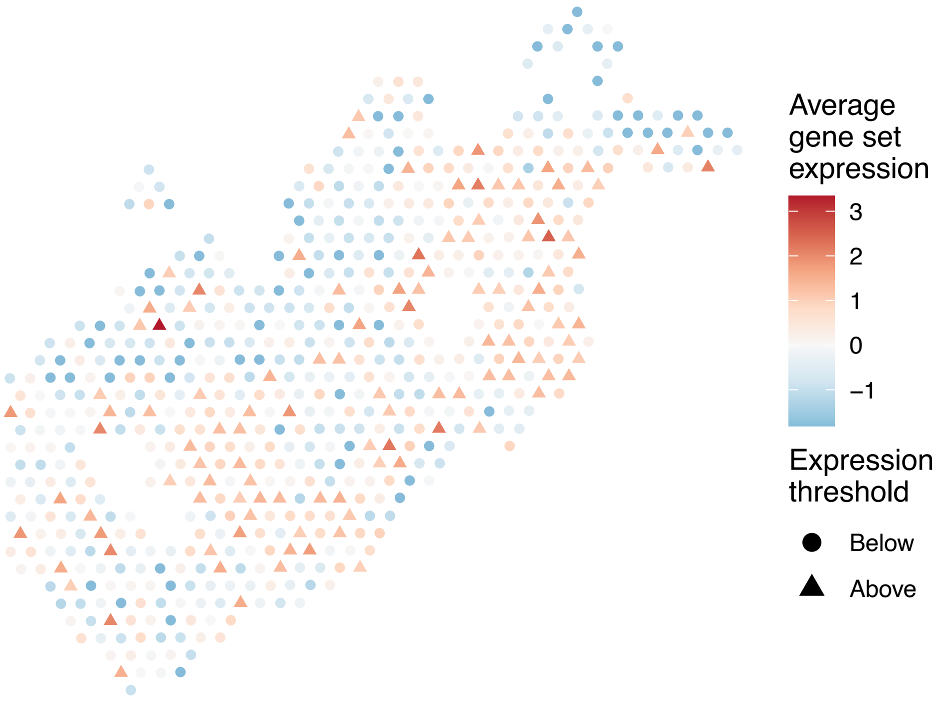


sample12_LMM_B4
KEGG_RIBOFLAVIN_METABOLISM
Proportion of pathway genes=0.75; Adjusted p−value=0.029

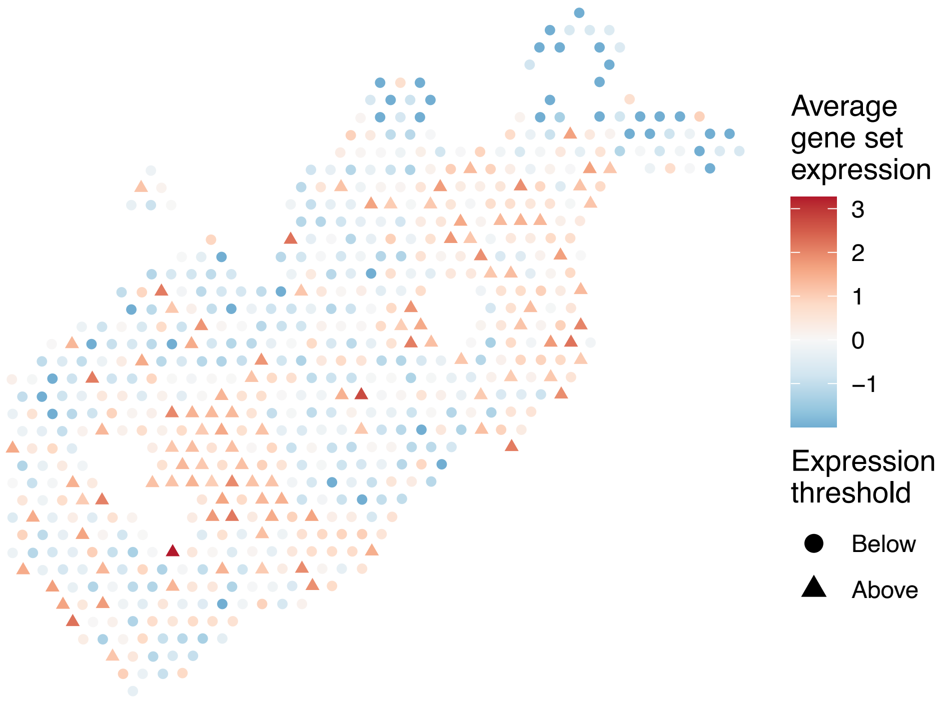


sample12_LMM_B4
KEGG_RIG_I_LIKE_RECEPTOR_SIGNALING_PATHWAY
Proportion of pathway genes=0.56; Adjusted p−value=0.0199

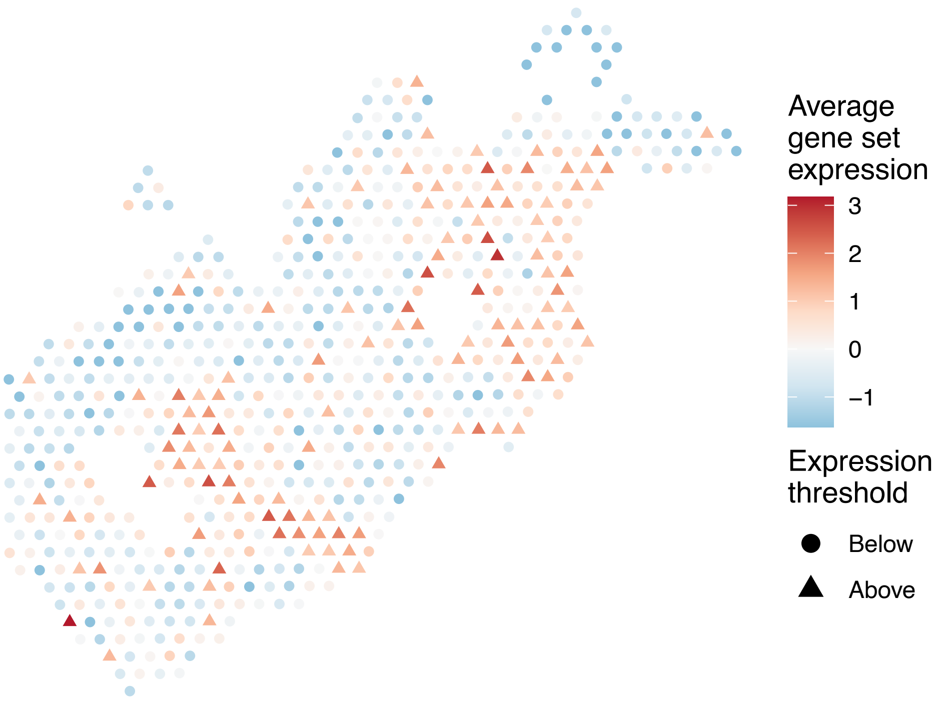

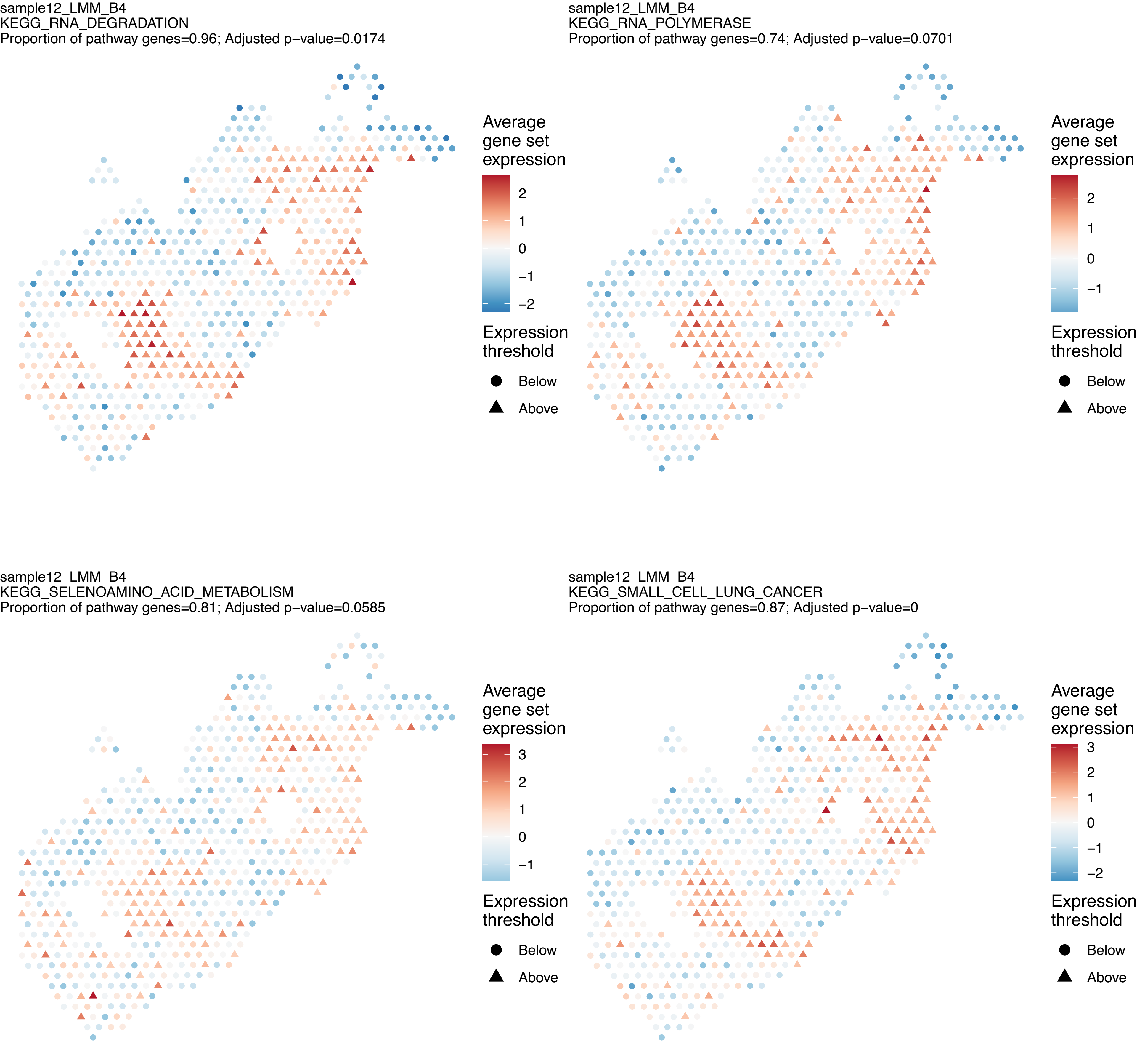
sample12_LMM_B4
KEGG_RNA_DEGRADATION
Proportion of pathway genes=0.96; Adjusted p−value=0.0174
Average gene set expression
2
1
0
−1
−2
Expression threshold
Below
Above
sample12_LMM_B4
KEGG_RNA_POLYMERASE
Proportion of pathway genes=0.74; Adjusted p−value=0.0701
Average gene set expression
2
1
0
−1
Expression threshold
Below
Above
sample12_LMM_B4
KEGG_SELENOAMINO_ACID_METABOLISM
Proportion of pathway genes=0.81; Adjusted p−value=0.0585
Average gene set expression
3
2
1
0
−1
Expression threshold
Below
Above
sample12_LMM_B4
KEGG_SMALL_CELL_LUNG_CANCER
Proportion of pathway genes=0.87; Adjusted p−value=0
Average gene set expression
3
2
1
0
−1
−2
Expression threshold
Below
Above

sample12_LMM_B4
KEGG_SNARE_INTERACTIONS_IN_VESICULAR_TRANSPORT
Proportion of pathway genes=0.87; Adjusted p−value=0.0895

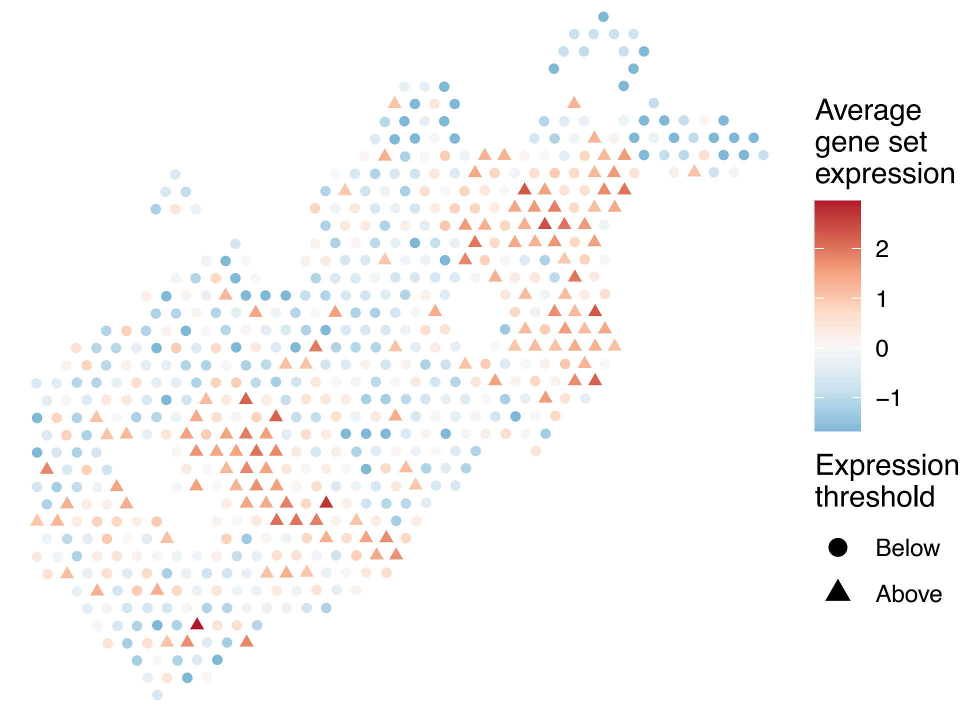


sample12_LMM_B4
KEGG_SPHINGOLIPID_METABOLISM
Proportion of pathway genes=0.76; Adjusted p−value=0.0079

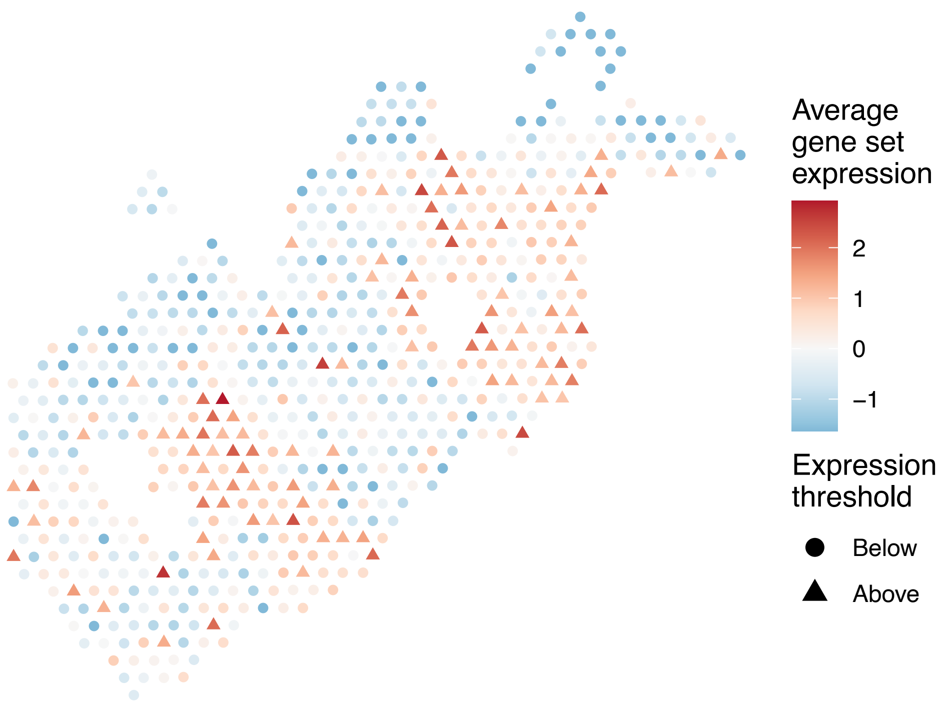


sample12_LMM_B4
KEGG_SPLICEOSOME
Proportion of pathway genes=0.96; Adjusted p−value=0.0345

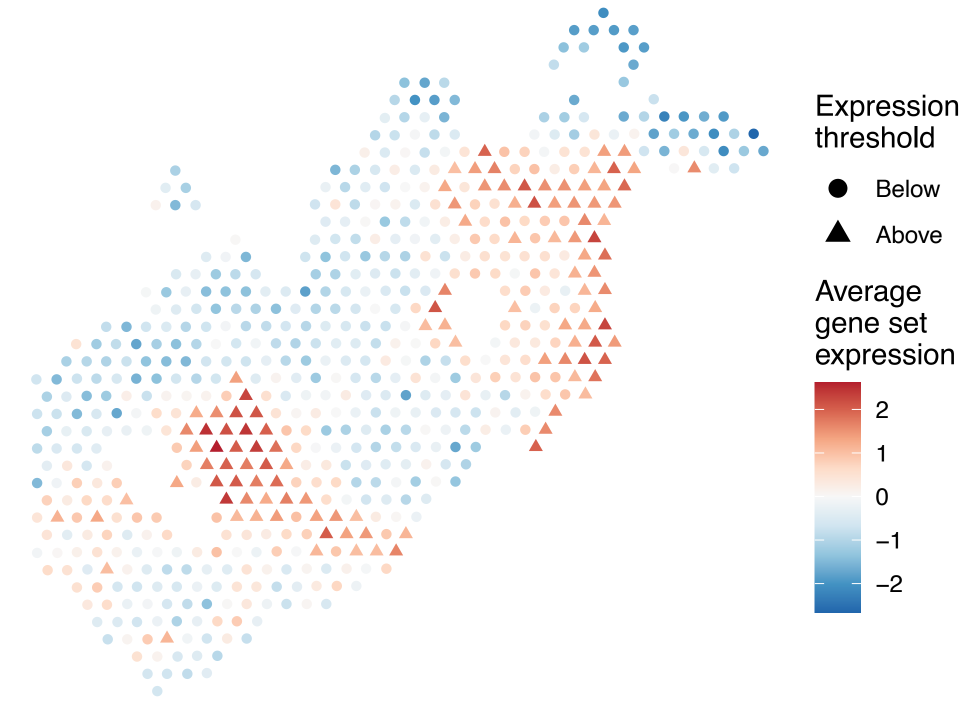


sample12_LMM_B4
KEGG_STARCH_AND_SUCROSE_METABOLISM
Proportion of pathway genes=0.34; Adjusted p−value=0.029

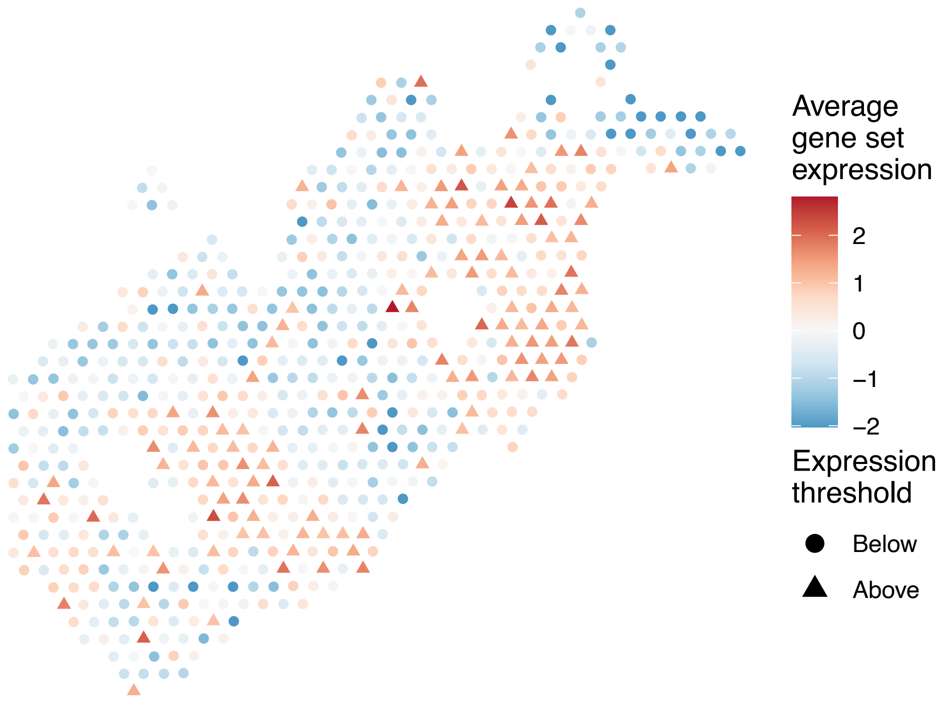

sample12_LMM_B4
KEGG_STEROID_BIOSYNTHESIS
Proportion of pathway genes=0.74; Adjusted p−value=0.3055

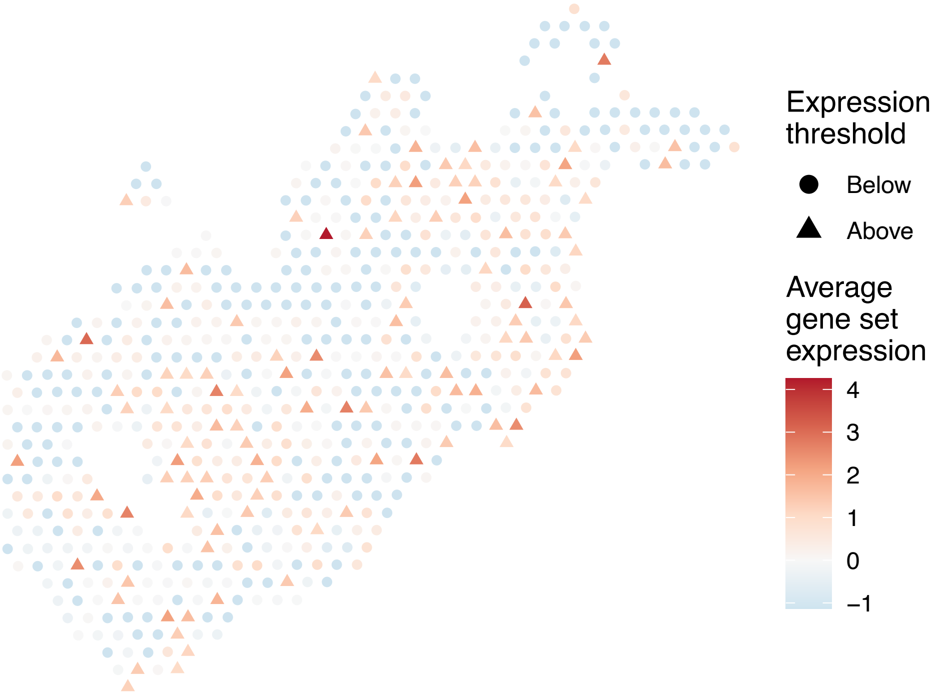


sample12_LMM_B4
KEGG_STEROID_HORMONE_BIOSYNTHESIS
Proportion of pathway genes=0.23; Adjusted p−value=0.1628

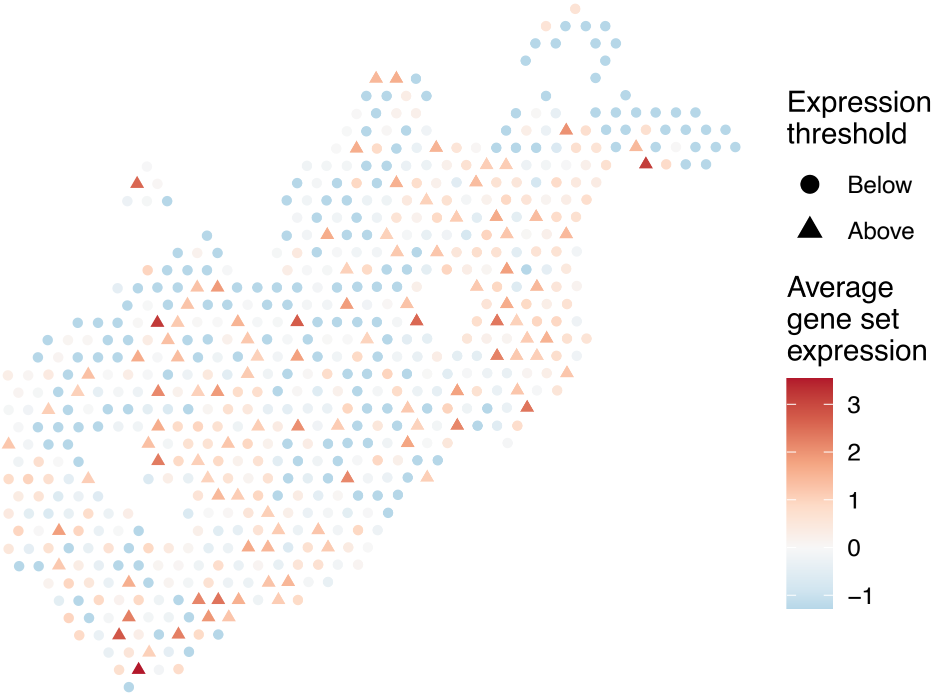


sample12_LMM_B4
KEGG_SULFUR_METABOLISM
Proportion of pathway genes=0.69; Adjusted p−value=NA

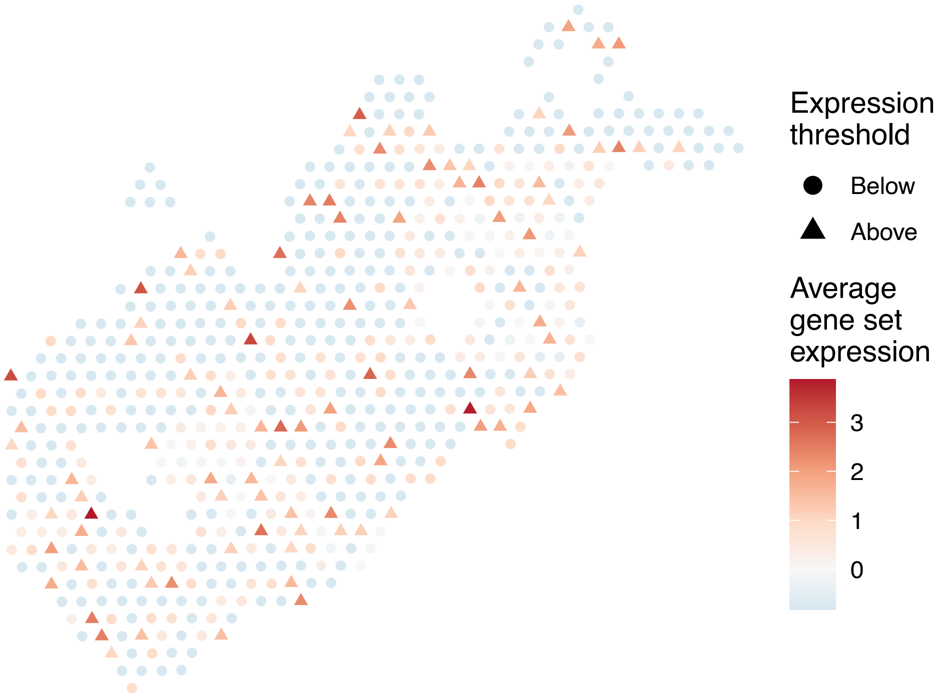


sample12_LMM_B4
KEGG_SYSTEMIC_LUPUS_ERYTHEMATOSUS
Proportion of pathway genes=0.34; Adjusted p−value=0.0246

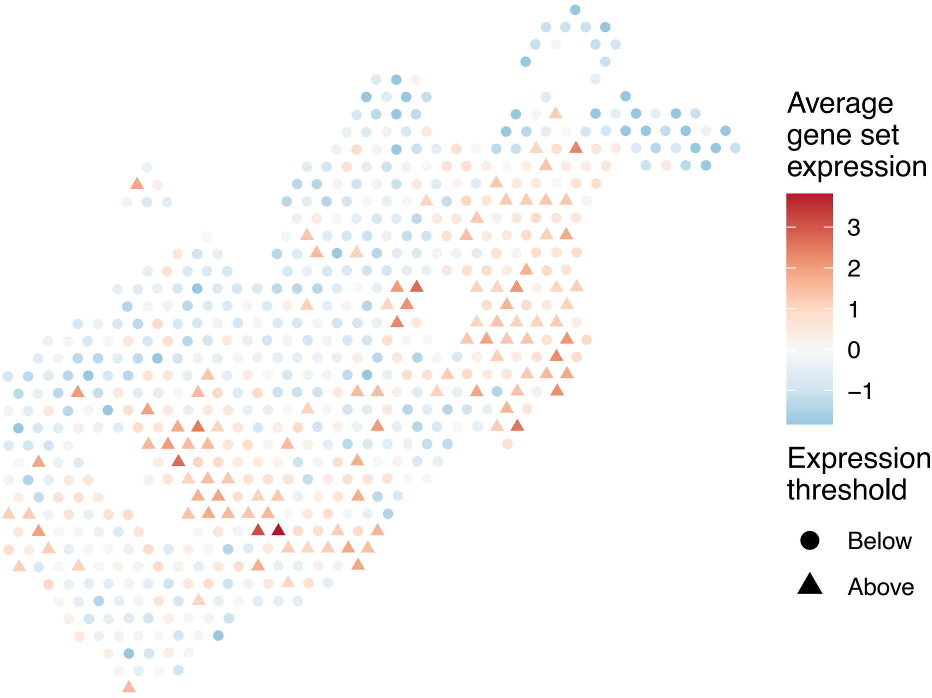

sample12_LMM_B4
KEGG_T_CELL_RECEPTOR_SIGNALING_PATHWAY
Proportion of pathway genes=0.62; Adjusted p−value=0.0423

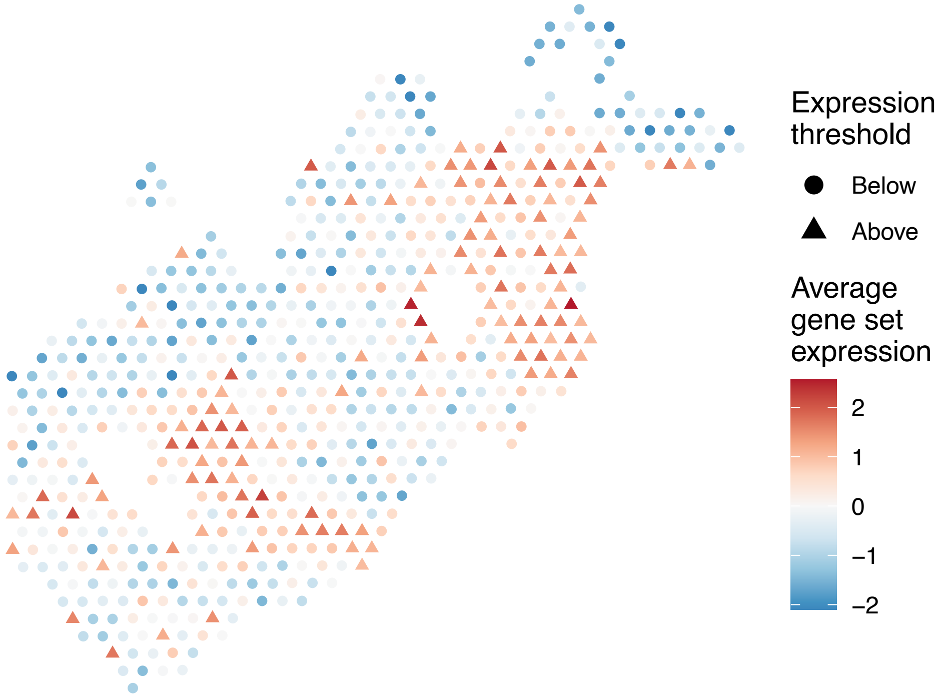


sample12_LMM_B4
KEGG_TERPENOID_BACKBONE_BIOSYNTHESIS
Proportion of pathway genes=0.8; Adjusted p−value=0.2714

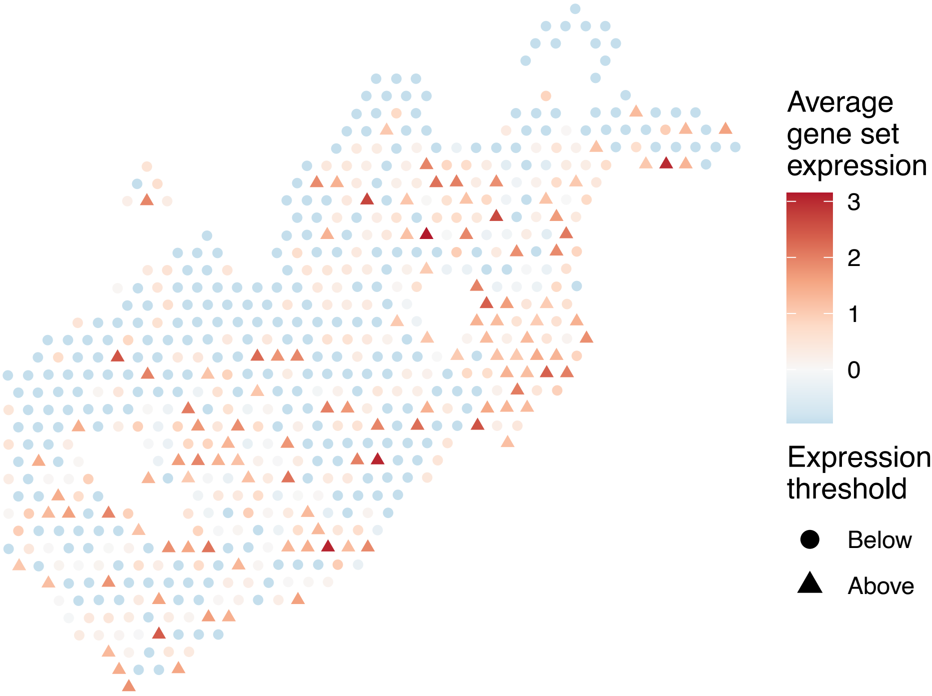


sample12_LMM_B4
KEGG_TGF_BETA_SIGNALING_PATHWAY
Proportion of pathway genes=0.63; Adjusted p−value=0.0048

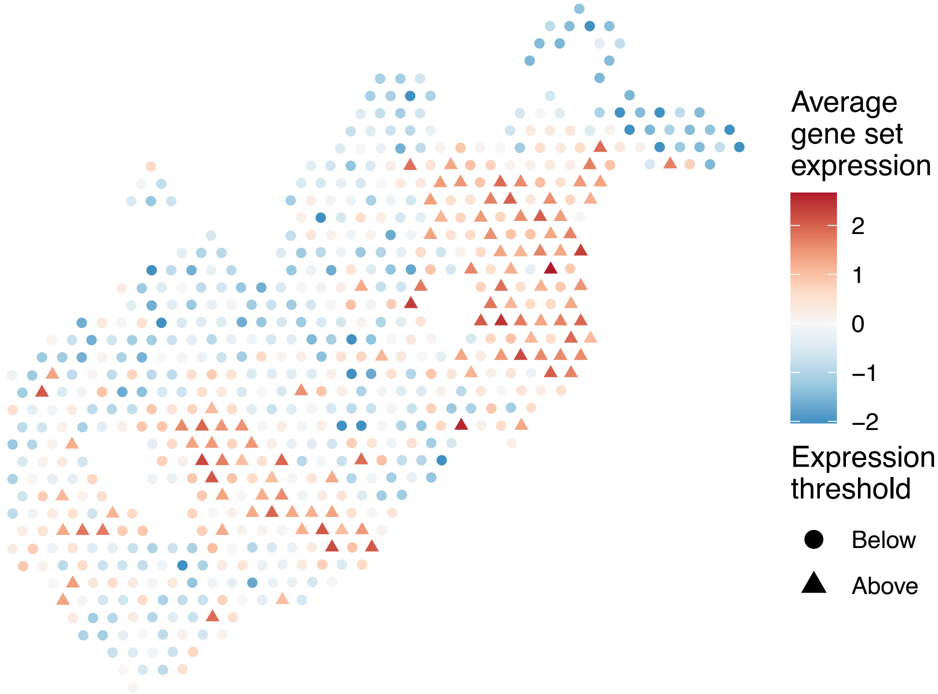


sample12_LMM_B4
KEGG_THYROID_CANCER
Proportion of pathway genes=0.86; Adjusted p−value=0.0174

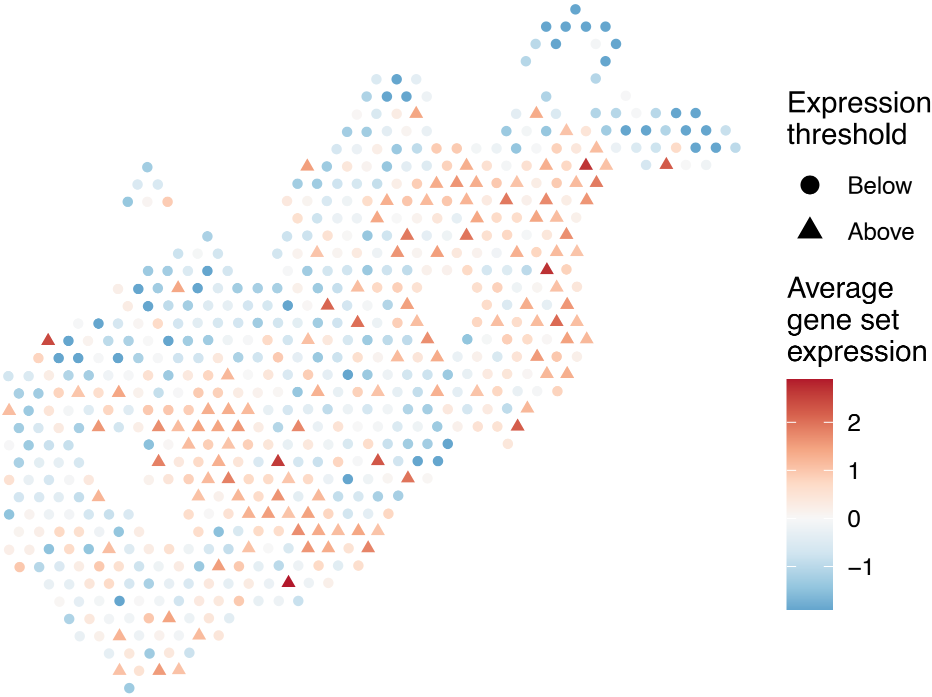

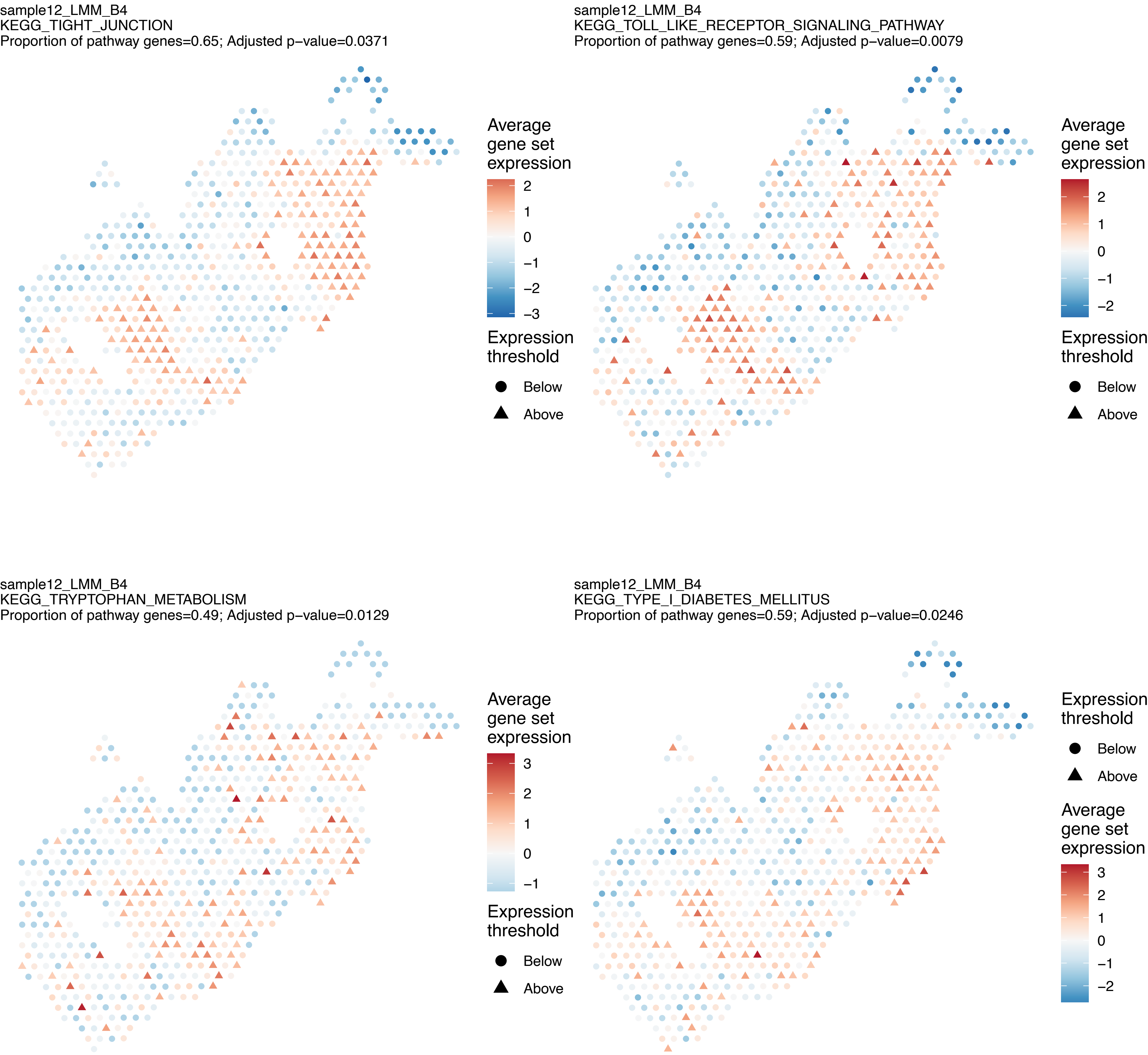
sample12_LMM_B4
KEGG_TIGHT_JUNCTION
Proportion of pathway genes=0.65; Adjusted p−value=0.0371
Average gene set expression
2
1
0
−1
−2
−3
Expression threshold
Below
Above
sample12_LMM_B4
KEGG_TOLL_LIKE_RECEPTOR_SIGNALING_PATHWAY
Proportion of pathway genes=0.59; Adjusted p−value=0.0079
Average gene set expression
2
1
0
−1
−2
Expression threshold
Below
Above
sample12_LMM_B4
KEGG_TRYPTOPHAN_METABOLISM
Proportion of pathway genes=0.49; Adjusted p−value=0.0129
Average gene set expression
3
2
1
0
−1
Expression threshold
Below
Above
sample12_LMM_B4
KEGG_TYPE_I_DIABETES_MELLITUS
Proportion of pathway genes=0.59; Adjusted p−value=0.0246
Expression threshold
Below
Above
Average gene set expression
3
2
1
0
−1
−2

sample12_LMM_B4
KEGG_TYPE_II_DIABETES_MELLITUS
Proportion of pathway genes=0.47; Adjusted p−value=0.0638

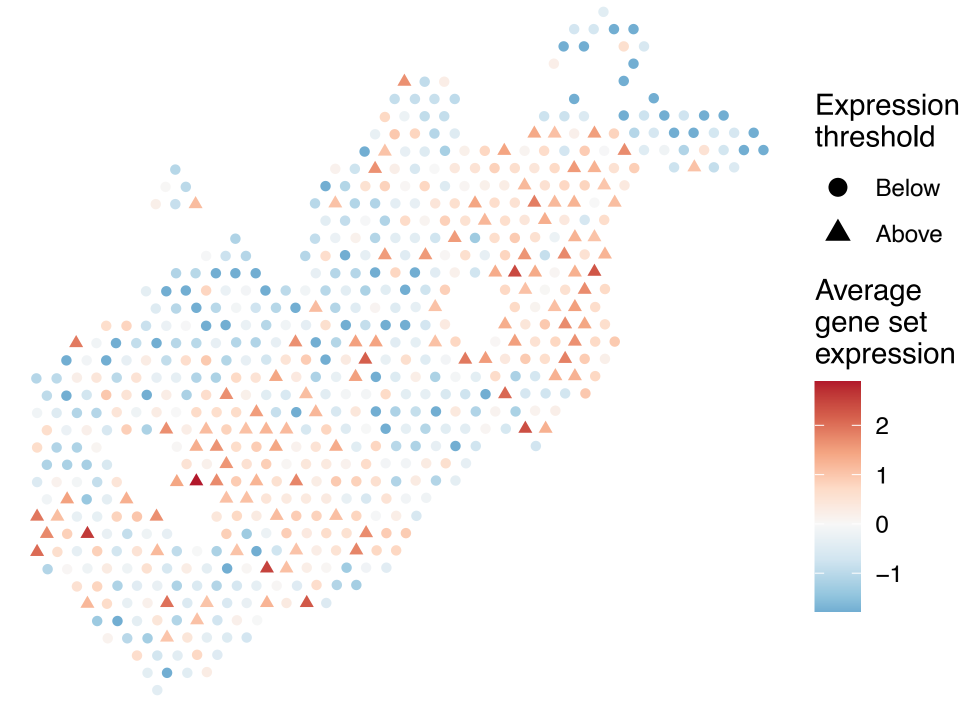


sample12_LMM_B4
KEGG_TYROSINE_METABOLISM
Proportion of pathway genes=0.52; Adjusted p−value=0.1066

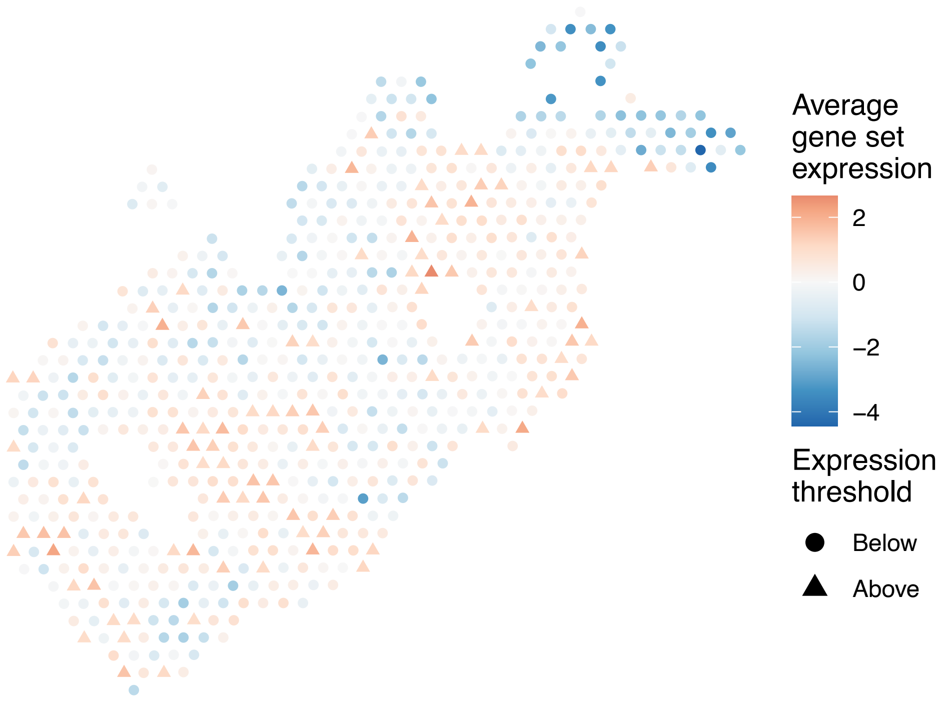


sample12_LMM_B4
KEGG_UBIQUITIN_MEDIATED_PROTEOLYSIS
Proportion of pathway genes=0.91; Adjusted p−value=0

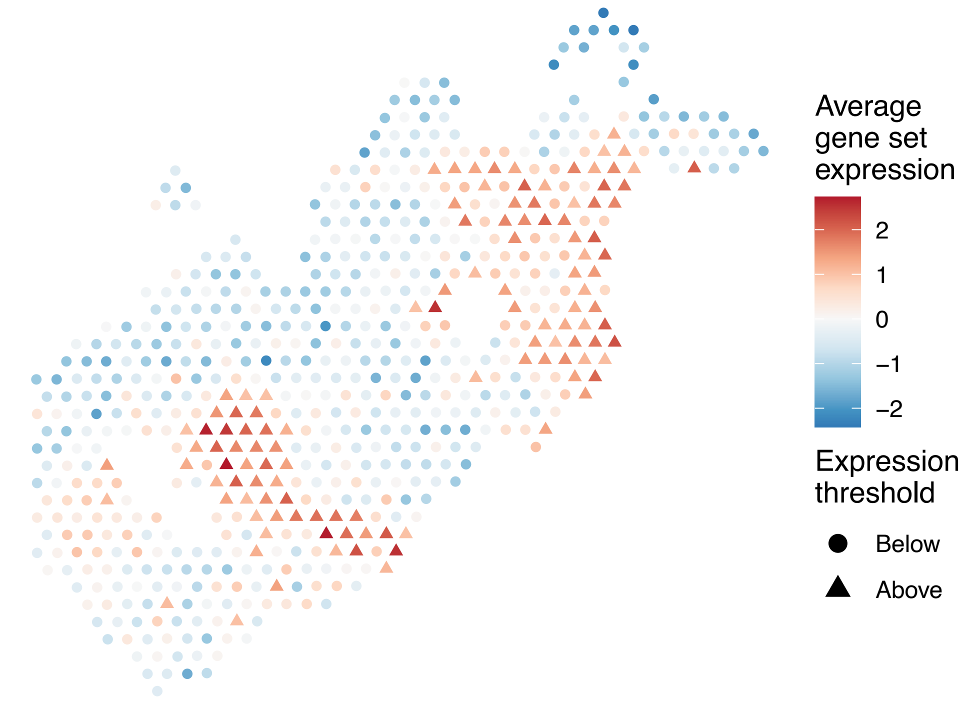


sample12_LMM_B4
KEGG_VALINE_LEUCINE_AND_ISOLEUCINE_BIOSYNTHESIS
Proportion of pathway genes=0.57; Adjusted p−value=0.3098

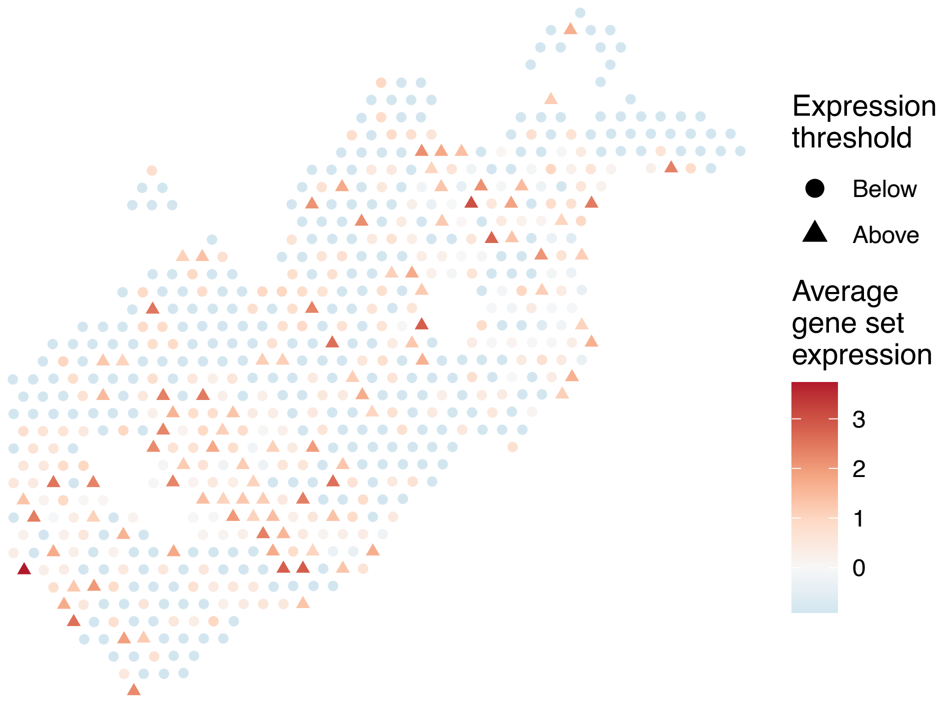

sample12_LMM_B4
KEGG_VALINE_LEUCINE_AND_ISOLEUCINE_DEGRADATION
Proportion of pathway genes=0.91; Adjusted p−value=0.0209

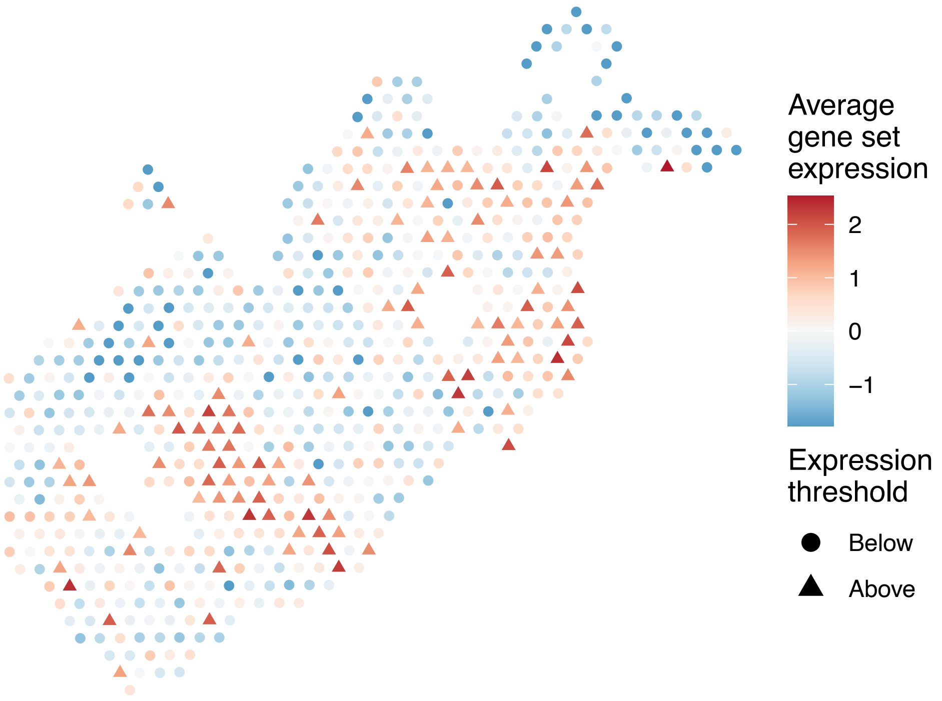


sample12_LMM_B4
KEGG_VASCULAR_SMOOTH_MUSCLE_CONTRACTION
Proportion of pathway genes=0.52; Adjusted p−value=0.0495

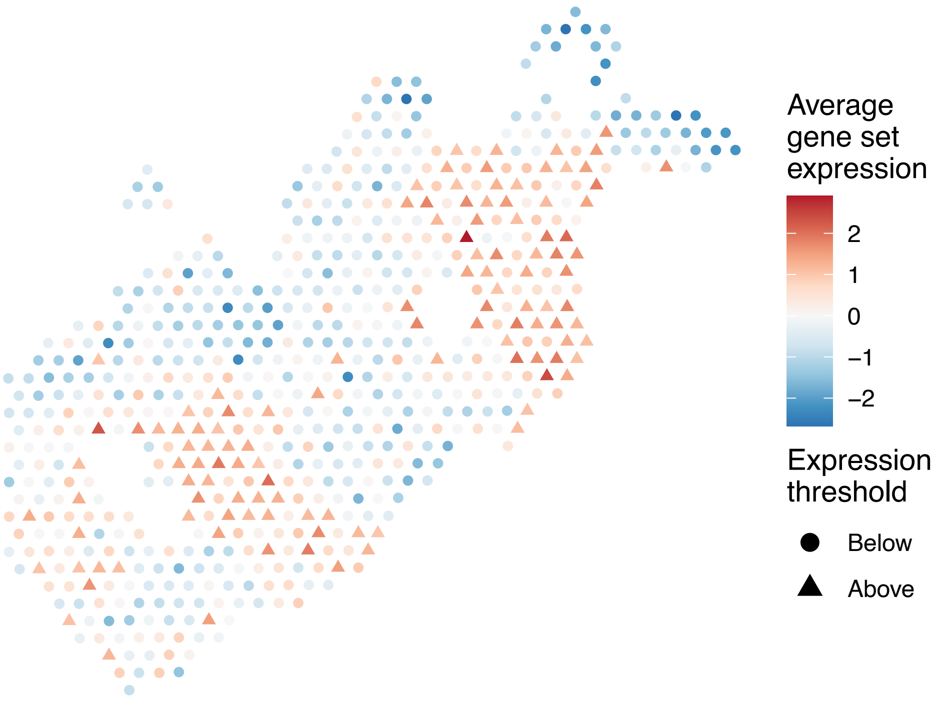


sample12_LMM_B4
KEGG_VASOPRESSIN_REGULATED_WATER_REABSORPTION
Proportion of pathway genes=0.83; Adjusted p−value=0.022

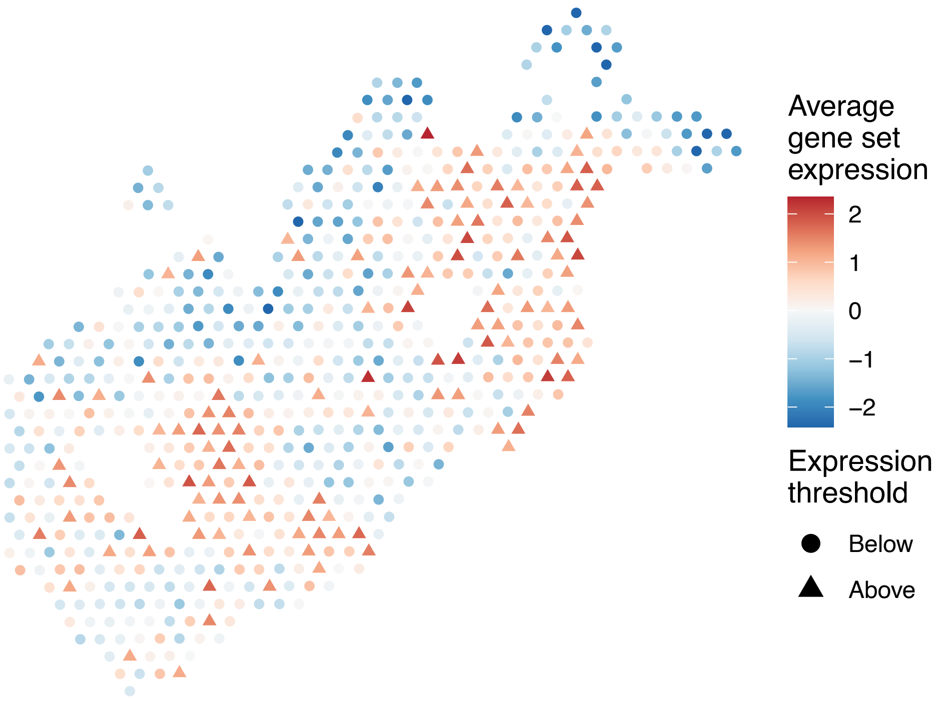


sample12_LMM_B4
KEGG_VEGF_SIGNALING_PATHWAY
Proportion of pathway genes=0.7; Adjusted p−value=0.0826

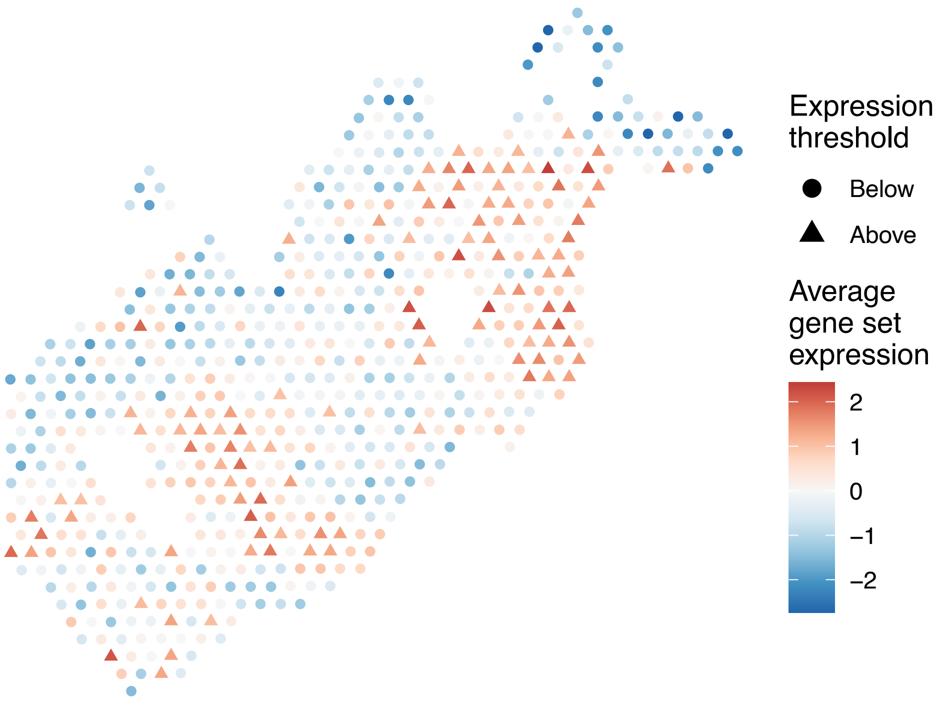

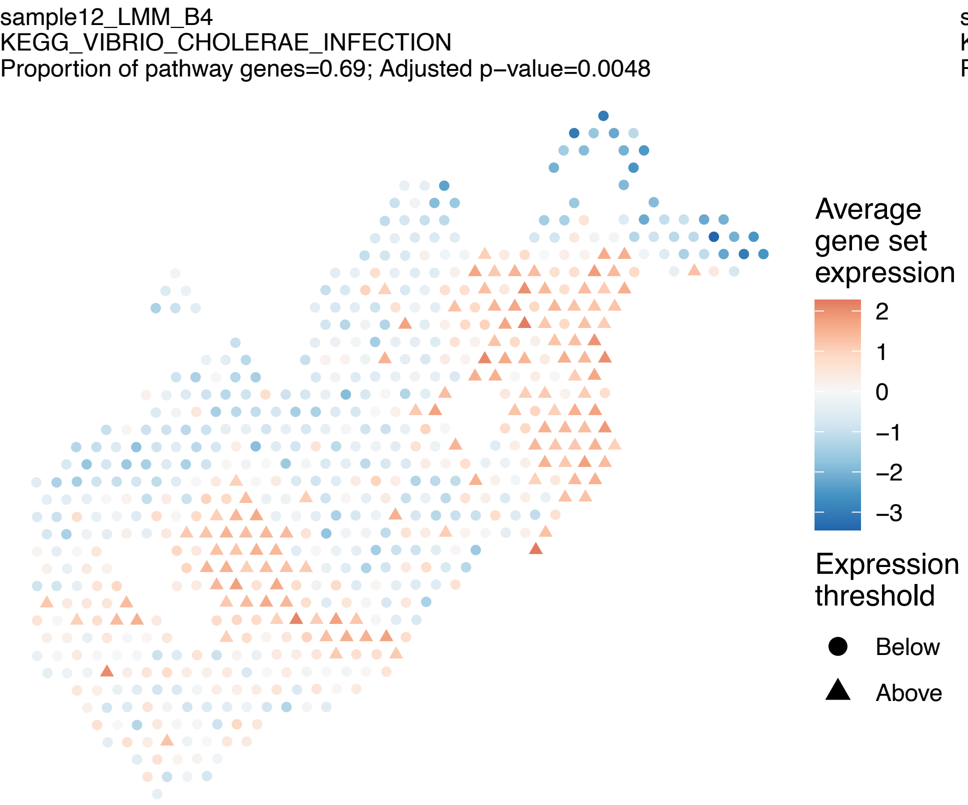
sample12_LMM_B4
KEGG_VIBRIO_CHOLERAE_INFECTION
Proportion of pathway genes=0.69; Adjusted p−value=0.0048
Average gene set expression
2
1
0
−1
−2
−3
Expression threshold
Below
Above

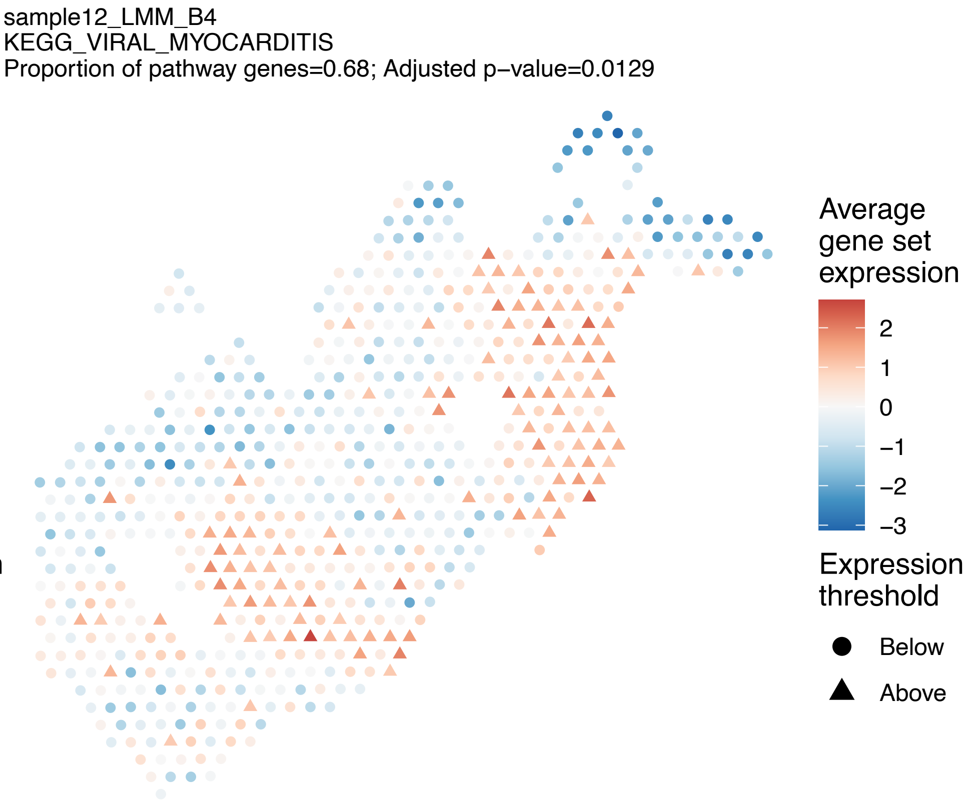
sample12_LMM_B4
KEGG_VIRAL_MYOCARDITIS
Proportion of pathway genes=0.68; Adjusted p−value=0.0129
Average gene set expression
2
1
0
−1
−2
−3
Expression threshold
Below
Above

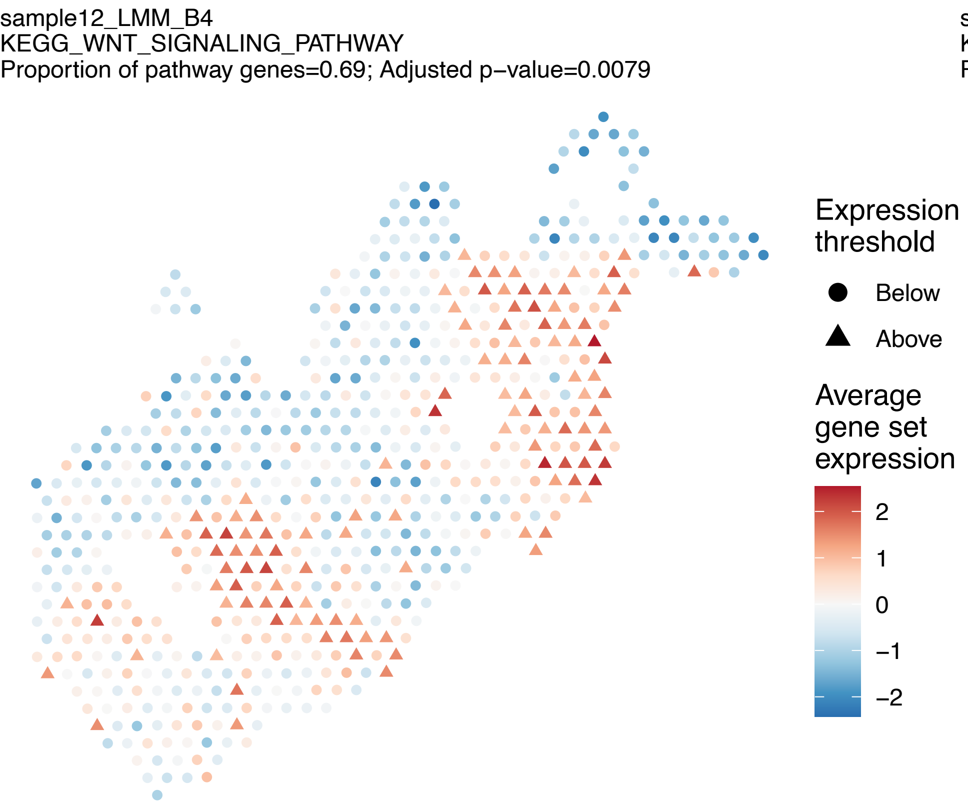
sample12_LMM_B4
KEGG_WNT_SIGNALING_PATHWAY
Proportion of pathway genes=0.69; Adjusted p−value=0.0079
Expression threshold
Below
Above
Average gene set expression
2
1
0
−1
−2

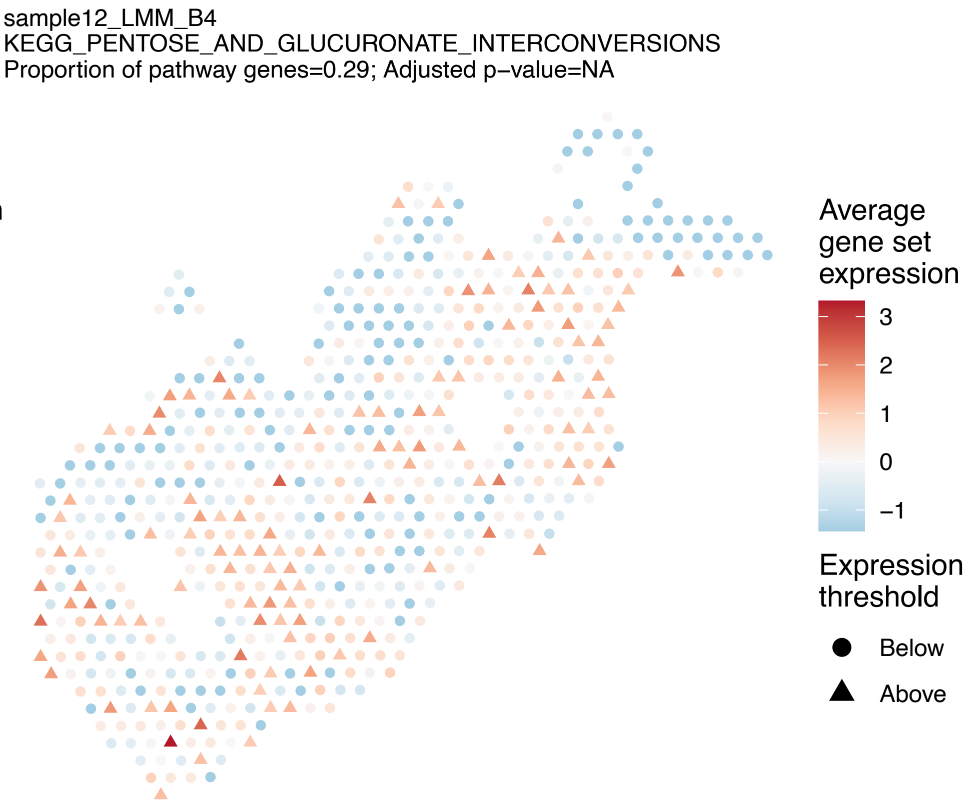
sample12_LMM_B4
KEGG_PENTOSE_AND_GLUCURONATE_INTERCONVERSIONS
Proportion of pathway genes=0.29; Adjusted p−value=NA
Average gene set expression
3
2
1
0
−1
Expression threshold
Below
Above

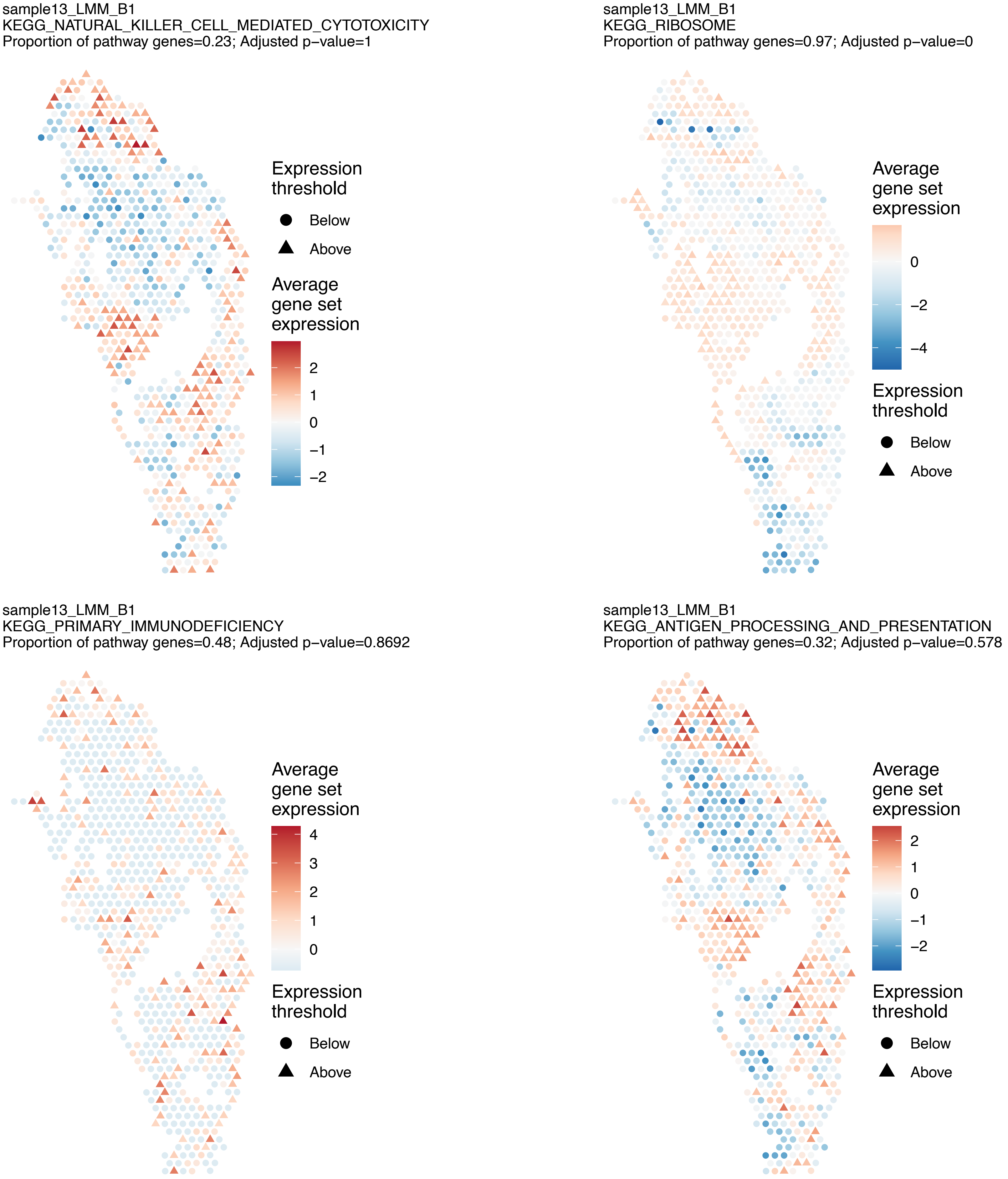
sample13_LMM_B1
KEGG_NATURAL_KILLER_CELL_MEDIATED_CYTOTOXICITY
Proportion of pathway genes=0.23; Adjusted p−value=1
Expression threshold
Below
Above
Average gene set expression
2
1
0
−1
−2
sample13_LMM_B1
KEGG_RIBOSOME
Proportion of pathway genes=0.97; Adjusted p−value=0
Average gene set expression
0
−2
−4
Expression threshold
Below
Above
sample13_LMM_B1
KEGG_PRIMARY_IMMUNODEFICIENCY
Proportion of pathway genes=0.48; Adjusted p−value=0.8692
Average gene set expression
4
3
2
1
0
Expression threshold
Below
Above
sample13_LMM_B1
KEGG_ANTIGEN_PROCESSING_AND_PRESENTATION
Proportion of pathway genes=0.32; Adjusted p−value=0.578
Average gene set expression
2
1
0
−1
−2
Expression threshold
Below
Above

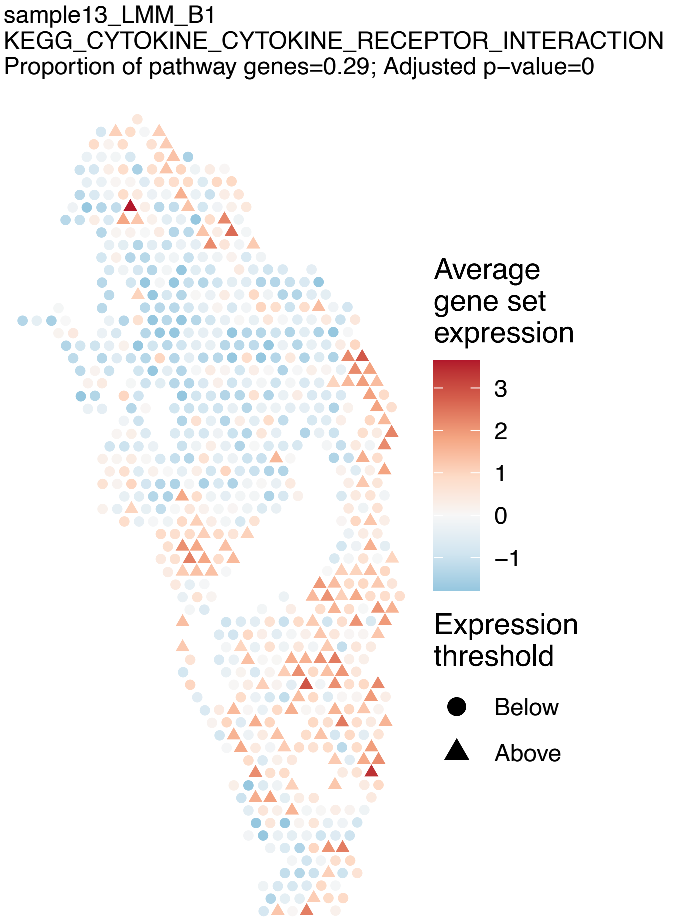
sample13_LMM_B1
KEGG_CYTOKINE_CYTOKINE_RECEPTOR_INTERACTION
Proportion of pathway genes=0.29; Adjusted p−value=0
Average gene set expression
3
2
1
0
−1
Expression threshold
Below
Above

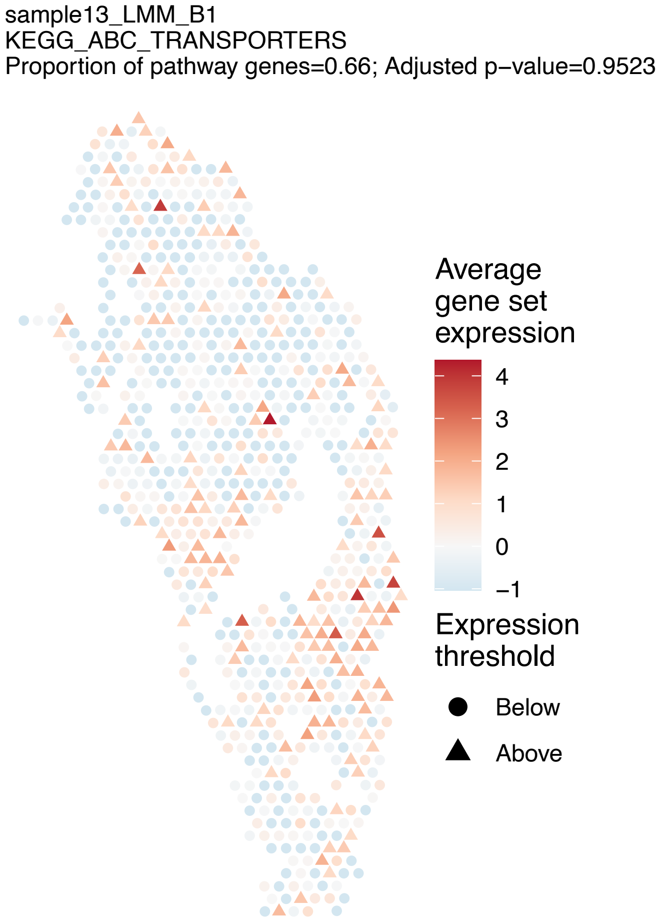
sample13_LMM_B1
KEGG_ABC_TRANSPORTERS
Proportion of pathway genes=0.66; Adjusted p−value=0.9523
Average gene set expression
4
3
2
1
0
−1
Expression threshold
Below
Above

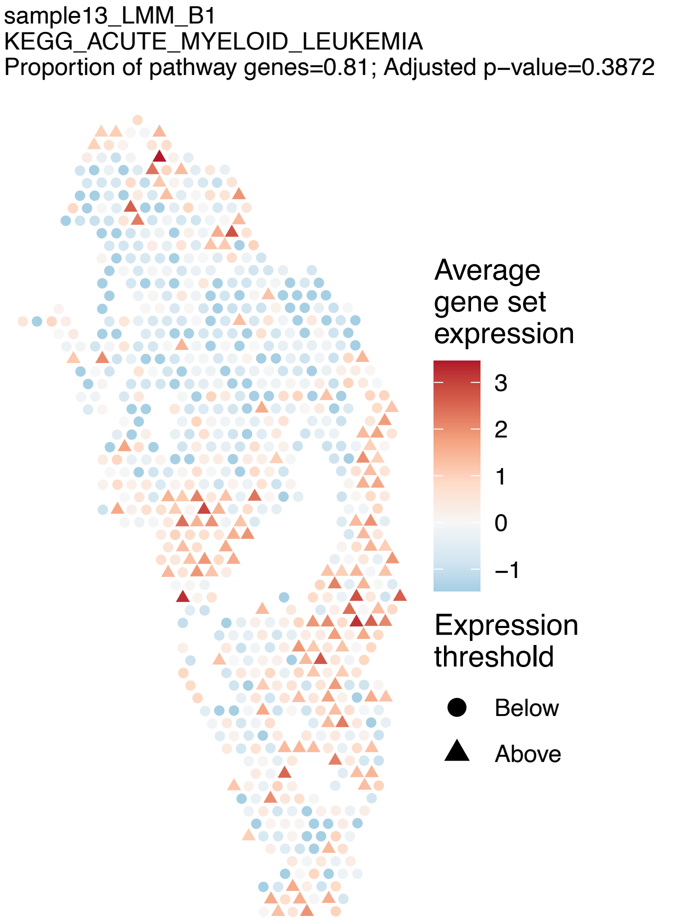
sample13_LMM_B1
KEGG_ACUTE_MYELOID_LEUKEMIA
Proportion of pathway genes=0.81; Adjusted p−value=0.3872
Average gene set expression
3
2
1
0
−1
Expression threshold
Below
Above

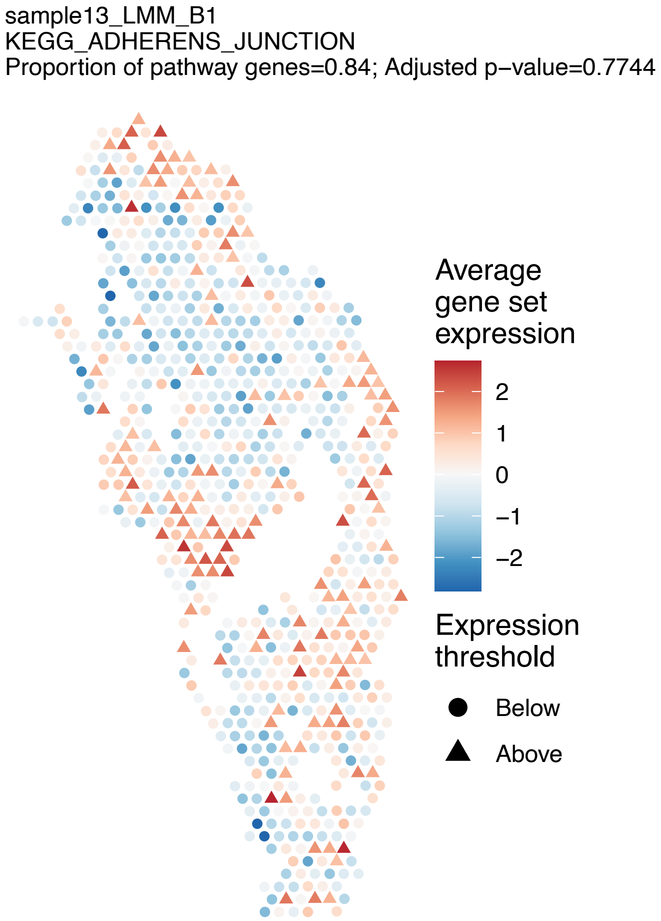
sample13_LMM_B1
KEGG_ADHERENS_JUNCTION
Proportion of pathway genes=0.84; Adjusted p−value=0.7744
Average gene set expression
2
1
0
−1
−2
Expression threshold
Below
Above

sample13_LMM_B1
KEGG_ADIPOCYTOKINE_SIGNALING_PATHWAY
Proportion of pathway genes=0.66; Adjusted p−value=0.0139

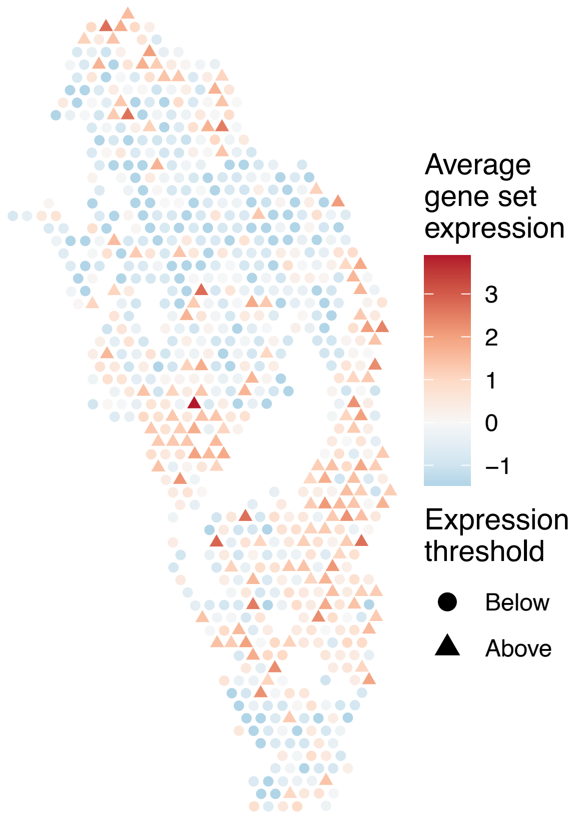


sample13_LMM_B1
KEGG_ALANINE_ASPARTATE_AND_GLUTAMATE_METABOLISM
Proportion of pathway genes=0.52; Adjusted p−value=1

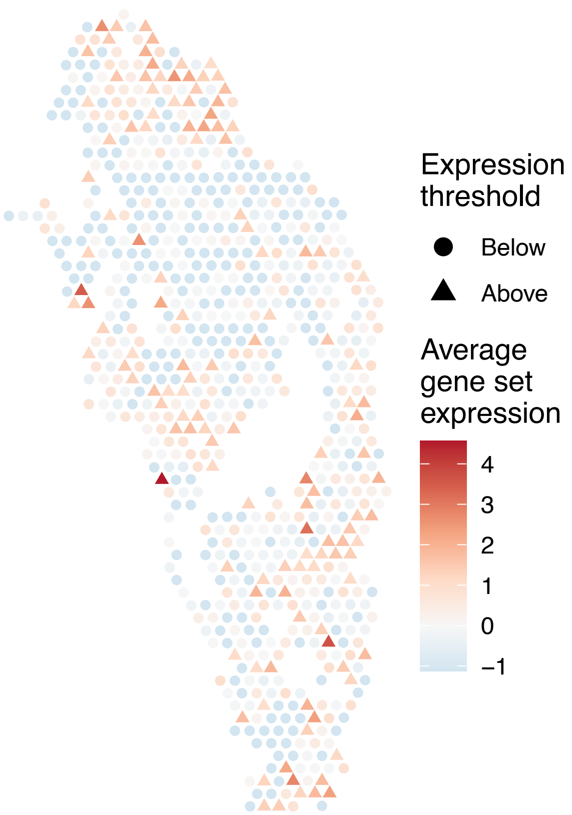


sample13_LMM_B1
KEGG_ALDOSTERONE_REGULATED_SODIUM_REABSORPTION
Proportion of pathway genes=0.5; Adjusted p−value=0.7972

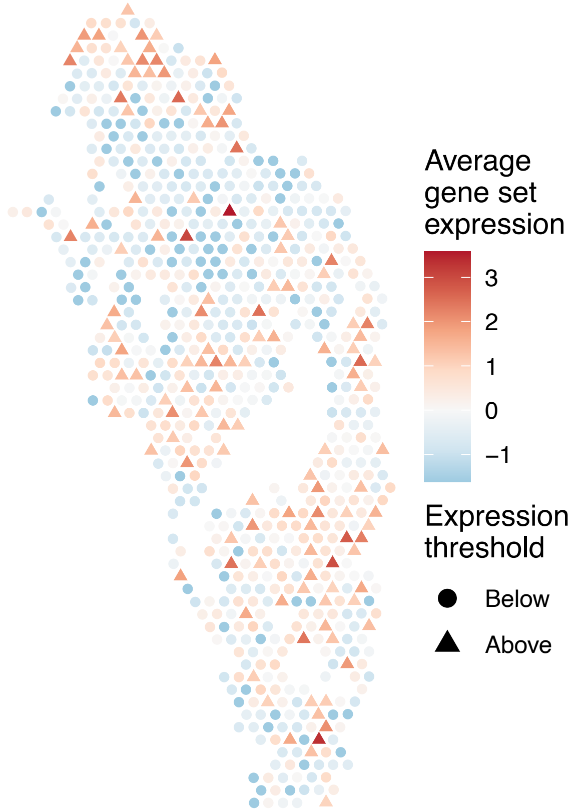


sample13_LMM_B1
KEGG_ALLOGRAFT_REJECTION
Proportion of pathway genes=0.69; Adjusted p−value=0.0239

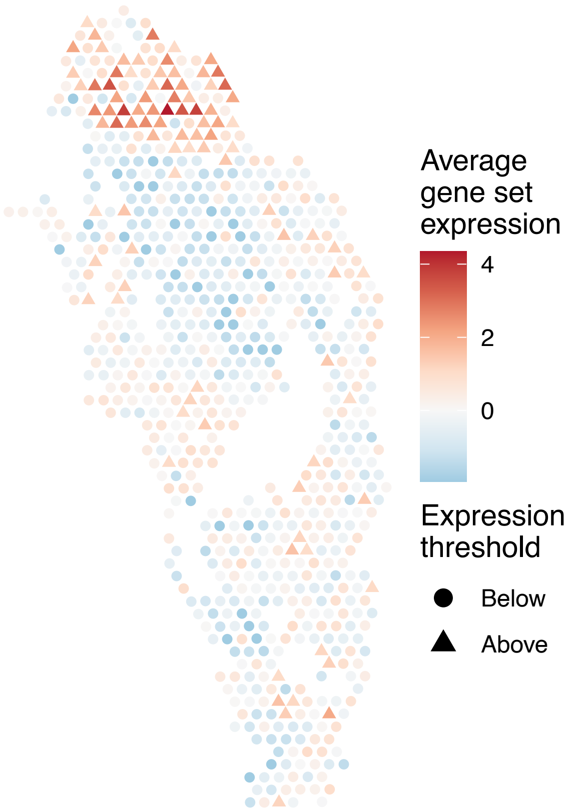

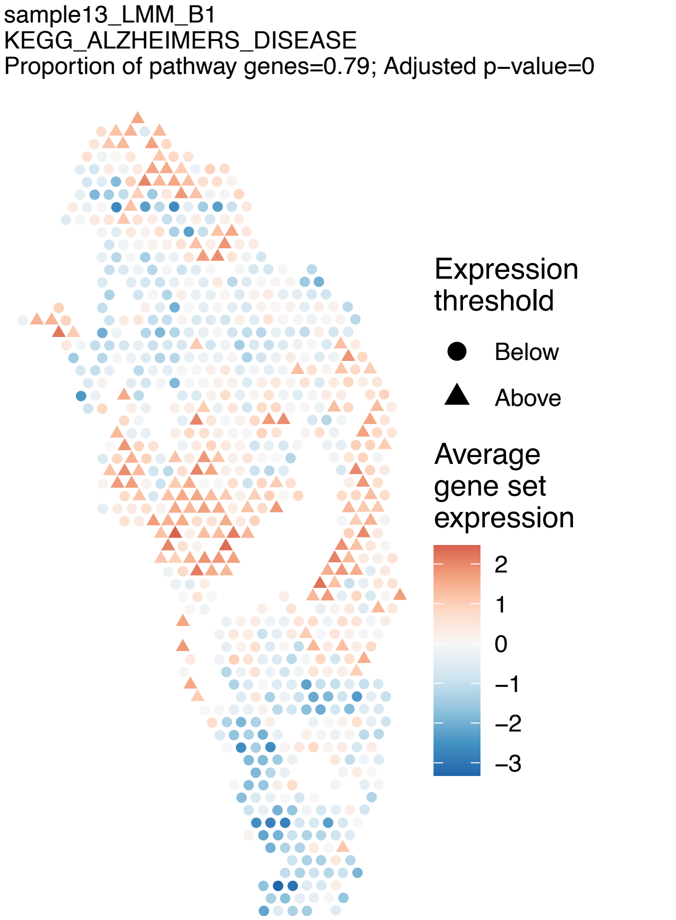
sample13_LMM_B1
KEGG_ALZHEIMERS_DISEASE
Proportion of pathway genes=0.79; Adjusted p−value=0
Expression threshold
Below
Above
Average gene set expression
2
1
0
−1
−2
−3
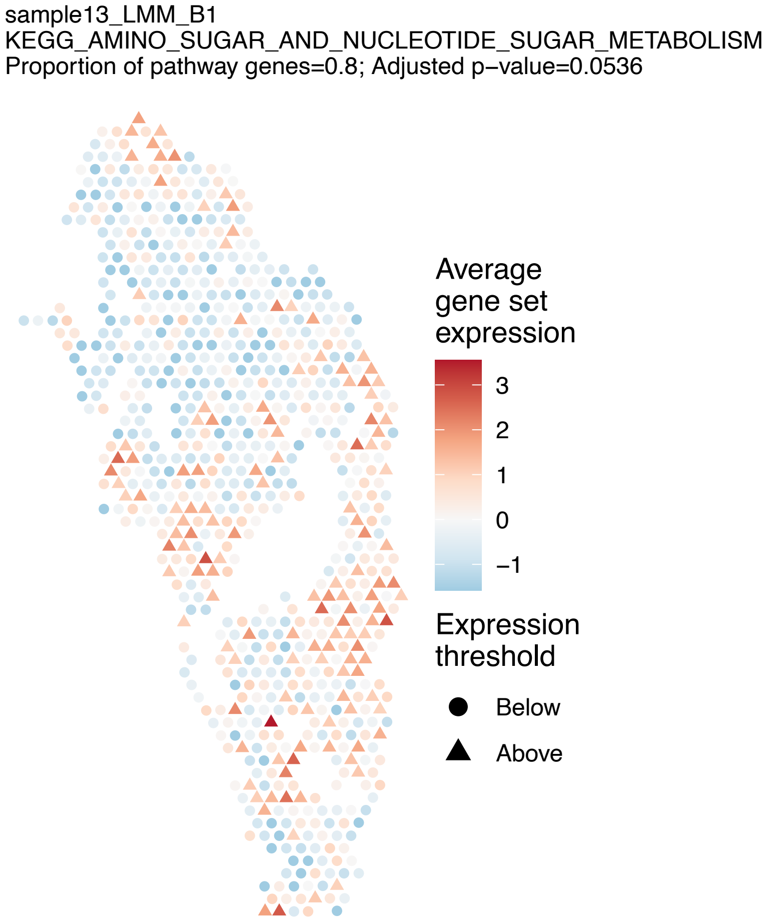
sample13_LMM_B1
KEGG_AMINO_SUGAR_AND_NUCLEOTIDE_SUGAR_METABOLISM
Proportion of pathway genes=0.8; Adjusted p−value=0.0536
Average gene set expression
3
2
1
0
−1
Expression threshold
Below
Above
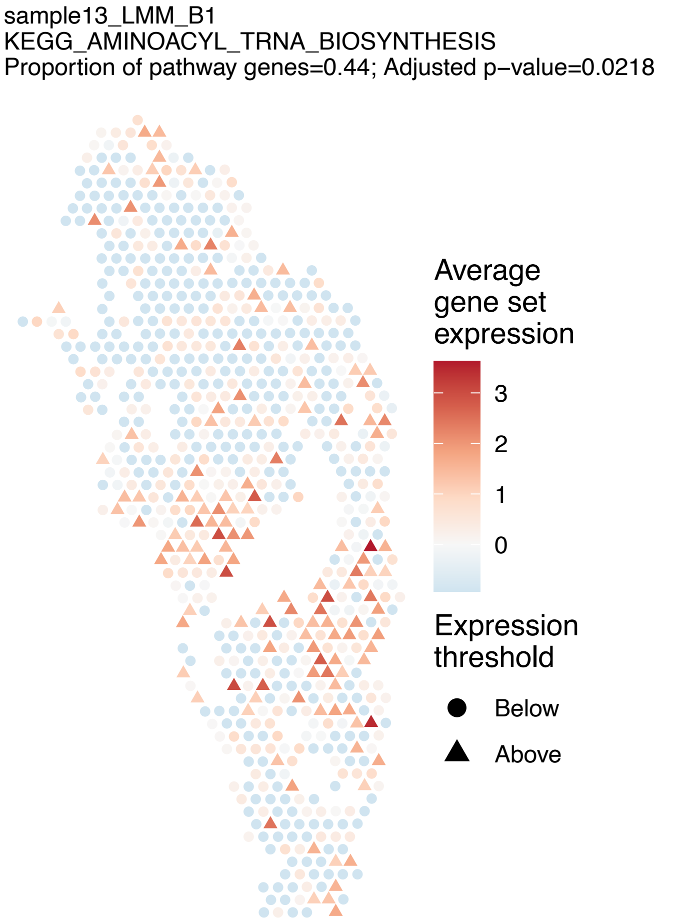
sample13_LMM_B1
KEGG_AMINOACYL_TRNA_BIOSYNTHESIS
Proportion of pathway genes=0.44; Adjusted p−value=0.0218
Average gene set expression
3
2
1
0
Expression threshold
Below
Above
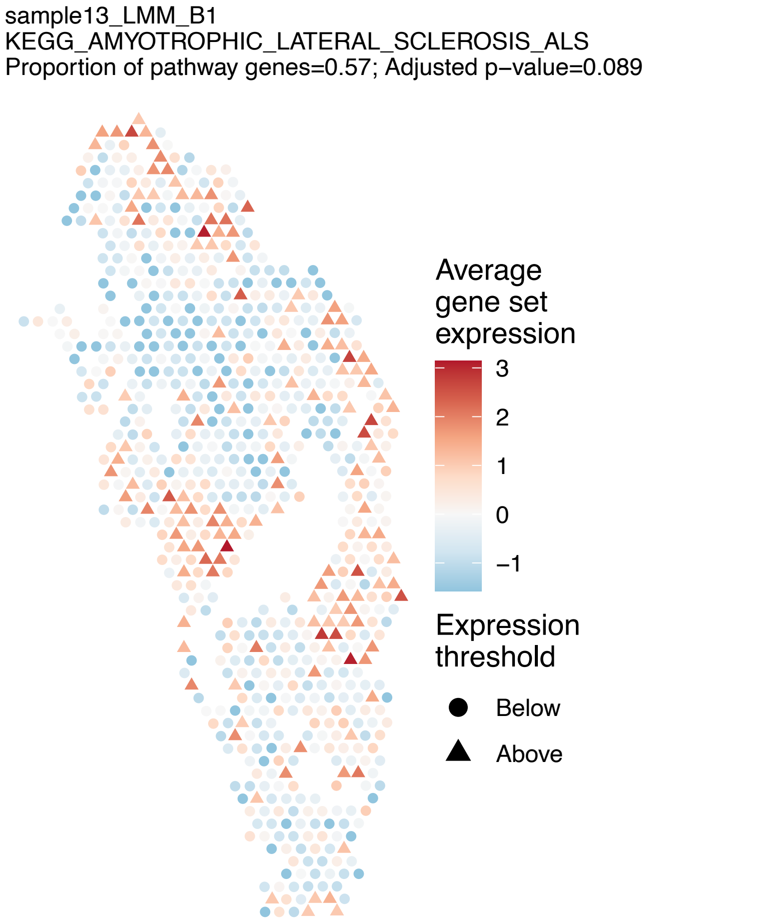
sample13_LMM_B1
KEGG_AMYOTROPHIC_LATERAL_SCLEROSIS_ALS
Proportion of pathway genes=0.57; Adjusted p−value=0.089
Average gene set expression
3
2
1
0
−1
Expression threshold
Below
Above

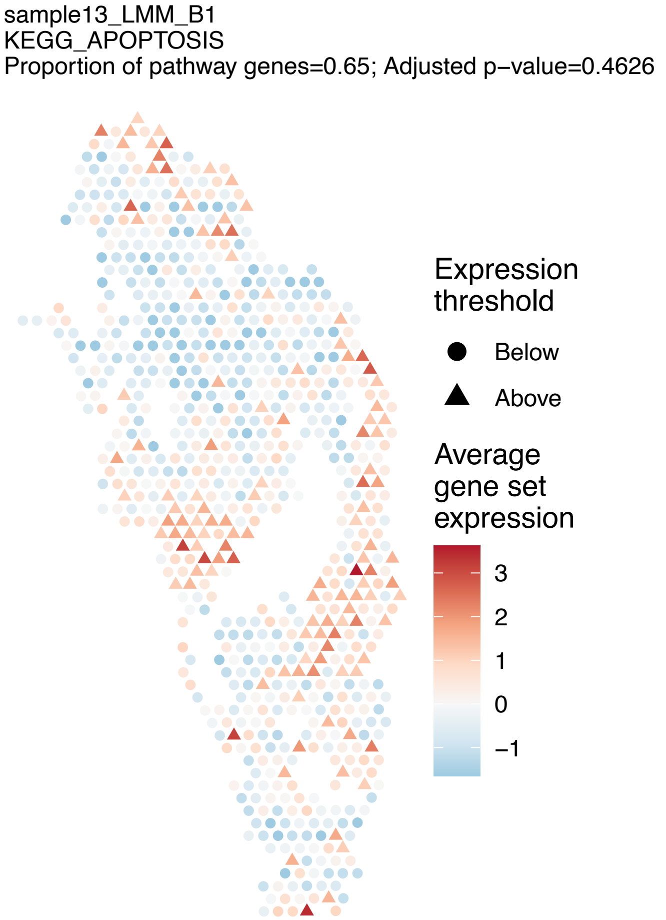
sample13_LMM_B1
KEGG_APOPTOSIS
Proportion of pathway genes=0.65; Adjusted p−value=0.4626
Expression threshold
Below
Above
Average gene set expression
3
2
1
0
−1

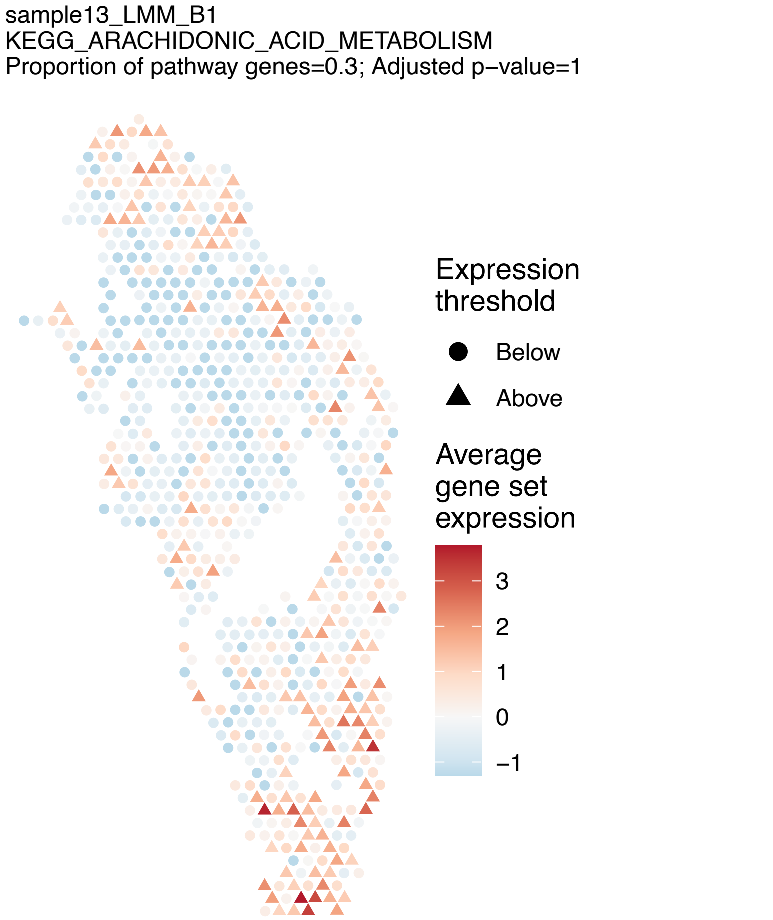
sample13_LMM_B1
KEGG_ARACHIDONIC_ACID_METABOLISM
Proportion of pathway genes=0.3; Adjusted p−value=1
Expression threshold
Below
Above
Average gene set expression
3
2
1
0
−1

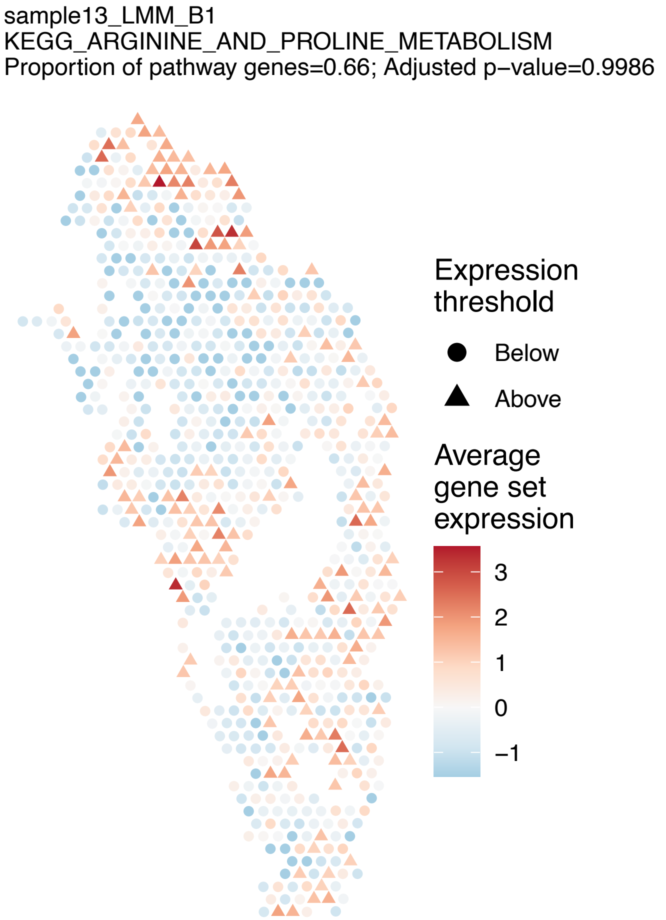
sample13_LMM_B1
KEGG_ARGININE_AND_PROLINE_METABOLISM
Proportion of pathway genes=0.66; Adjusted p−value=0.9986
Expression threshold
Below
Above
Average gene set expression
3
2
1
0
−1

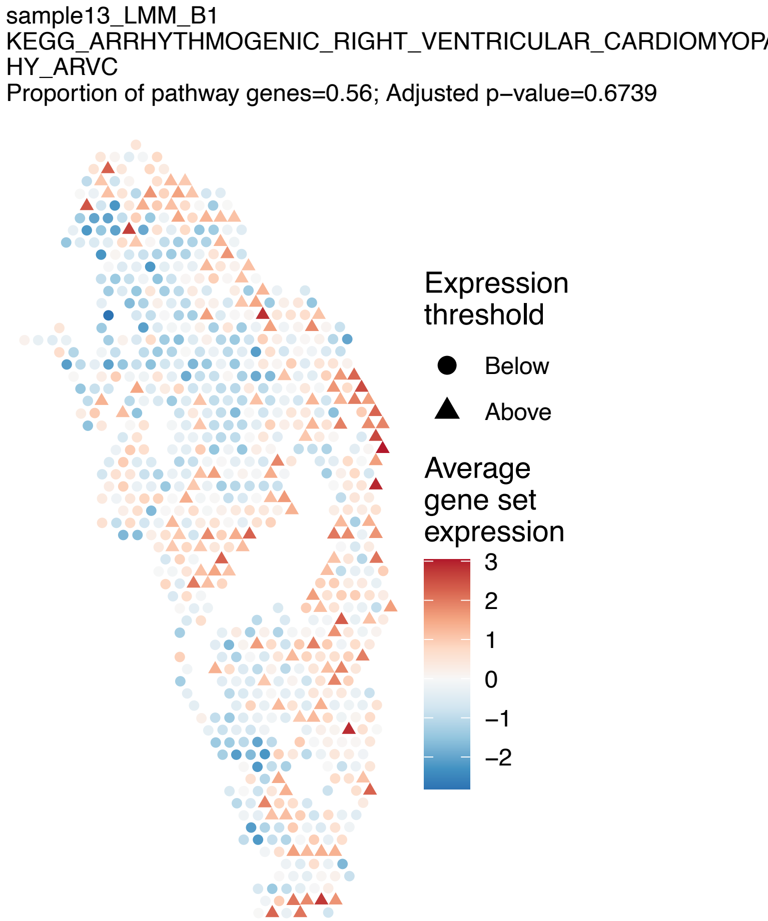
sample13_LMM_B1
KEGG_ARRHYTHMOGENIC_RIGHT_VENTRICULAR_CARDIOMYOP
HY_ARVC
Proportion of pathway genes=0.56; Adjusted p−value=0.6739
Expression threshold
Below
Above
Average gene set expression
3
2
1
0
−1
−2

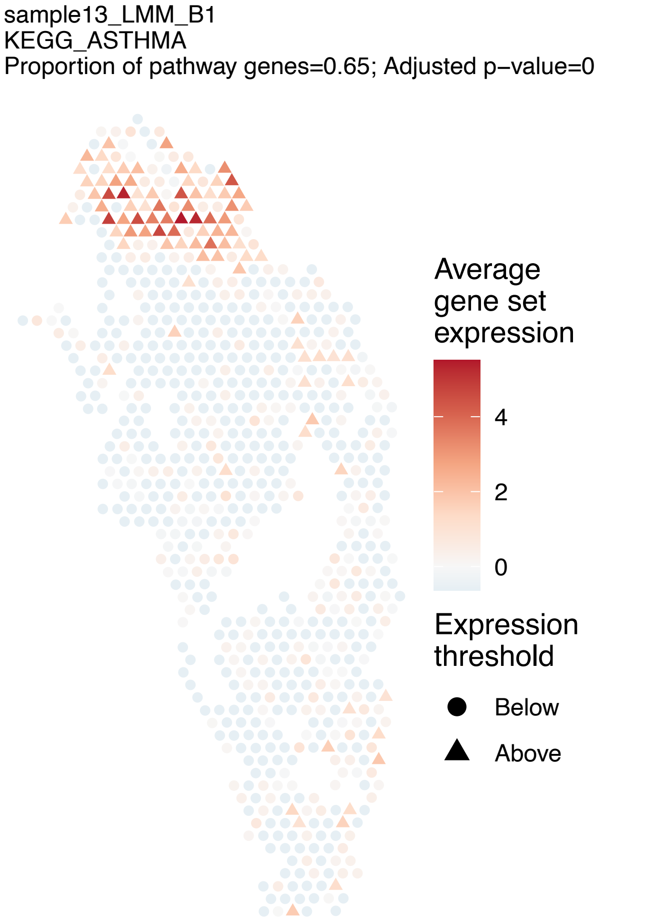
sample13_LMM_B1
KEGG_ASTHMA
Proportion of pathway genes=0.65; Adjusted p−value=0
Average gene set expression
4
2
0
Expression threshold
Below
Above

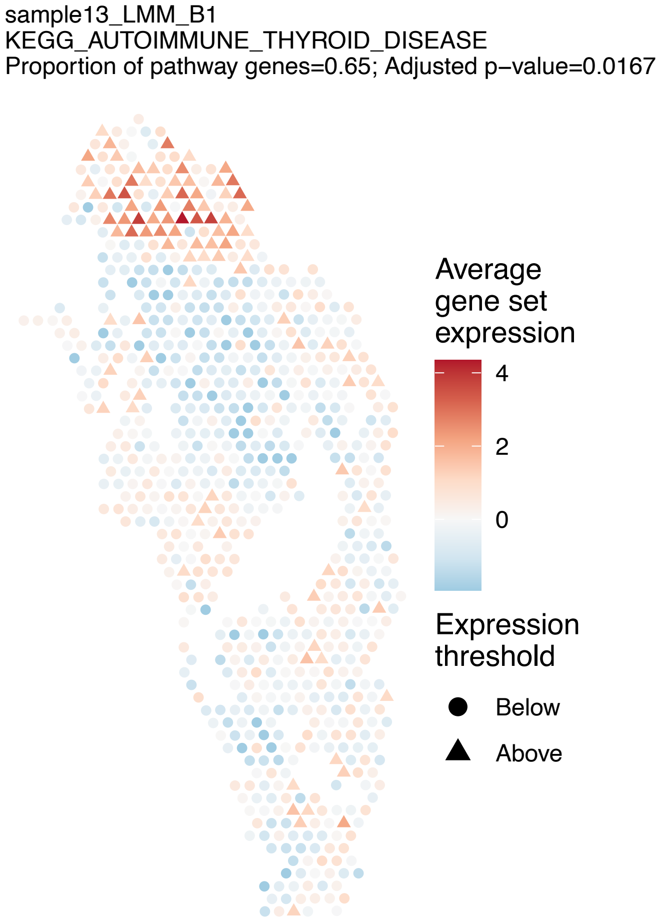
sample13_LMM_B1
KEGG_AUTOIMMUNE_THYROID_DISEASE
Proportion of pathway genes=0.65; Adjusted p−value=0.0167
Average gene set expression
4
2
0
Expression threshold
Below
Above

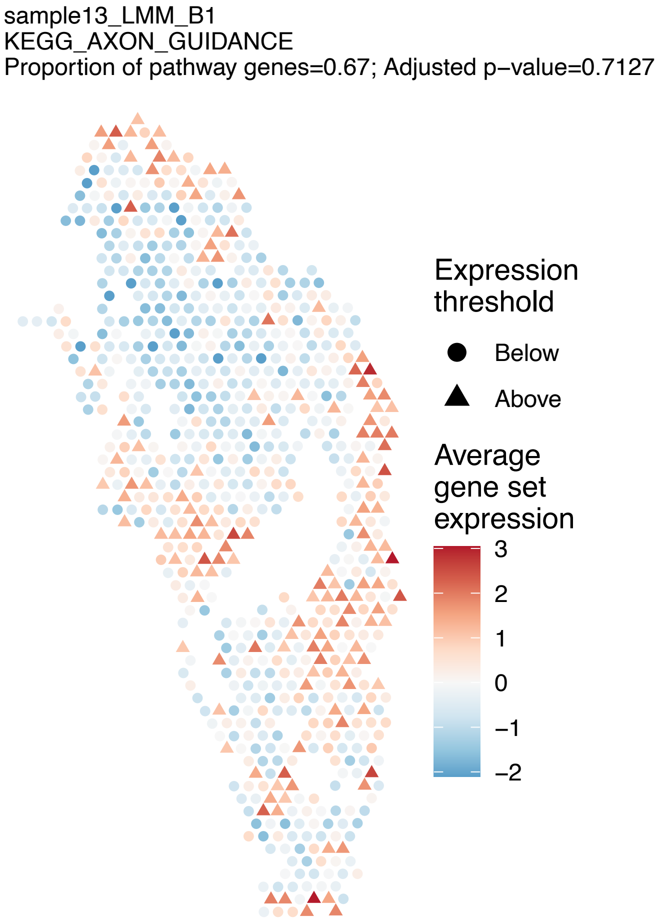
sample13_LMM_B1
KEGG_AXON_GUIDANCE
Proportion of pathway genes=0.67; Adjusted p−value=0.7127
Expression threshold
Below
Above
Average gene set expression
3
2
1
0
−1
−2

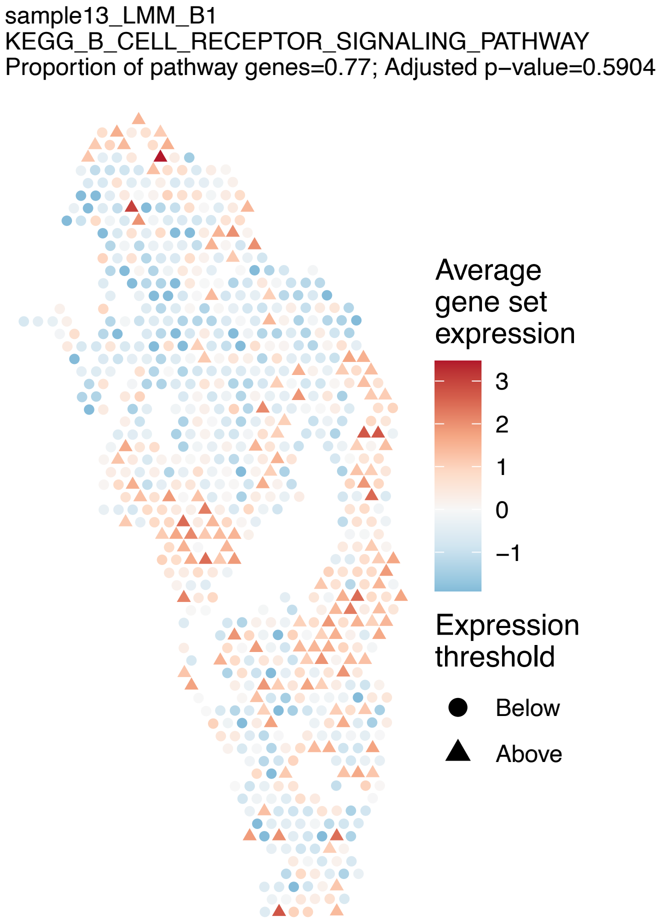
sample13_LMM_B1
KEGG_B_CELL_RECEPTOR_SIGNALING_PATHWAY
Proportion of pathway genes=0.77; Adjusted p−value=0.5904
Average gene set expression
3
2
1
0
−1
Expression threshold
Below
Above

sample13_LMM_B1
KEGG_BASAL_CELL_CARCINOMA
Proportion of pathway genes=0.41; Adjusted p−value=0.1506

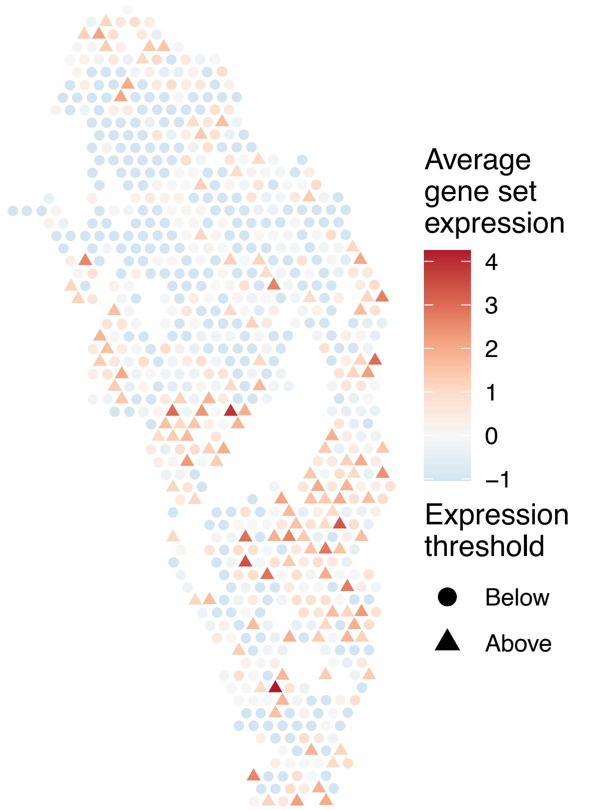


sample13_LMM_B1
KEGG_BASAL_TRANSCRIPTION_FACTORS
Proportion of pathway genes=0.76; Adjusted p−value=0.4477

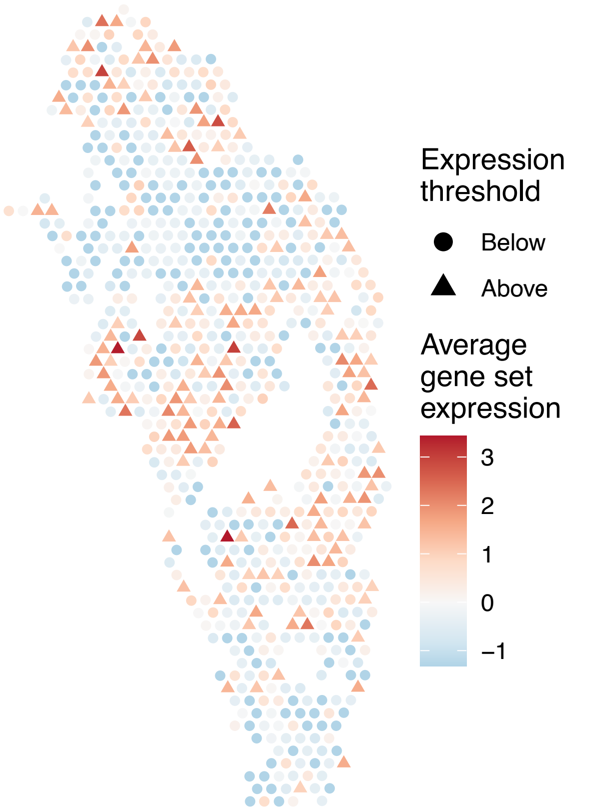


sample13_LMM_B1
KEGG_BASE_EXCISION_REPAIR
Proportion of pathway genes=0.86; Adjusted p−value=0

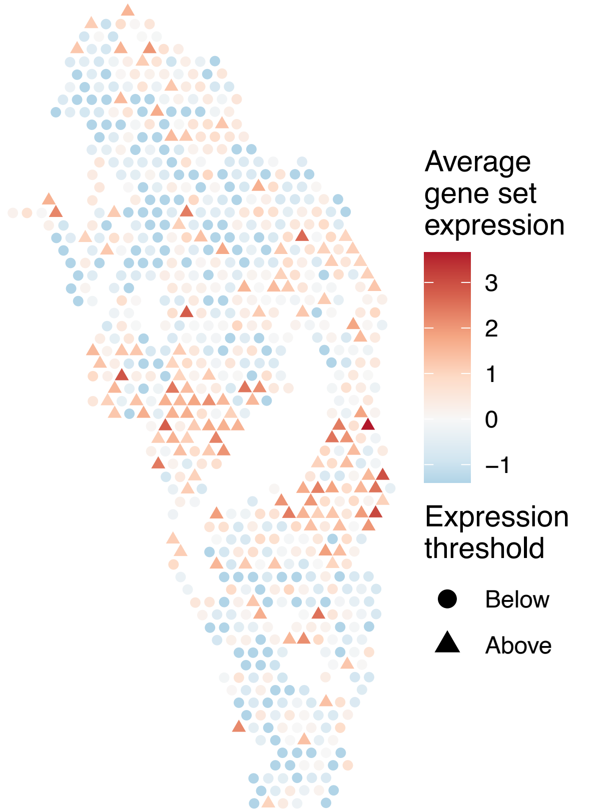


sample13_LMM_B1
KEGG_BETA_ALANINE_METABOLISM
Proportion of pathway genes=0.64; Adjusted p−value=0.0456

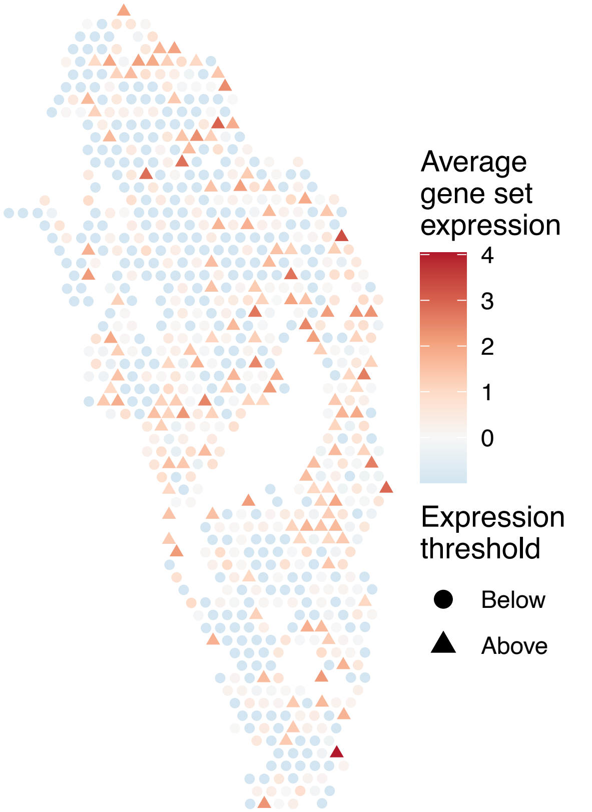

sample13_LMM_B1
KEGG_BIOSYNTHESIS_OF_UNSATURATED_FATTY_ACIDS
Proportion of pathway genes=0.78; Adjusted p−value=0.2815

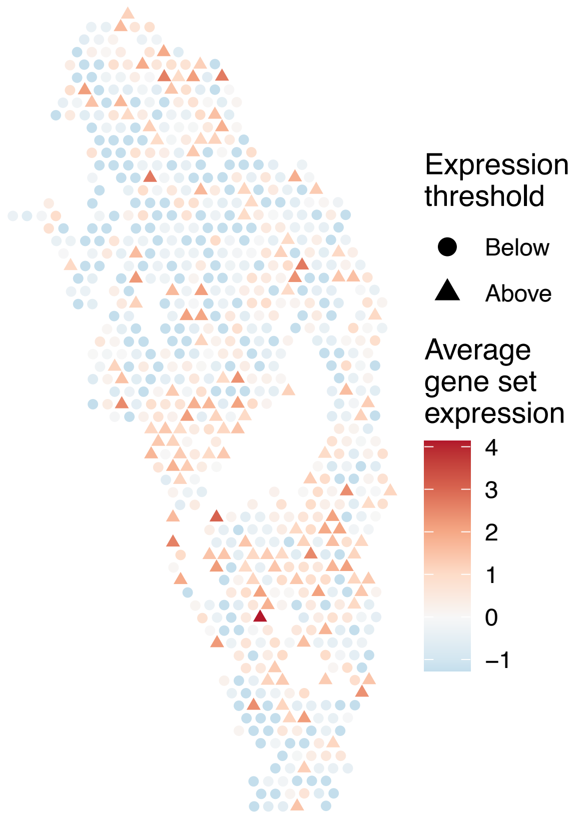


sample13_LMM_B1
KEGG_BLADDER_CANCER
Proportion of pathway genes=0.78; Adjusted p−value=0.2565

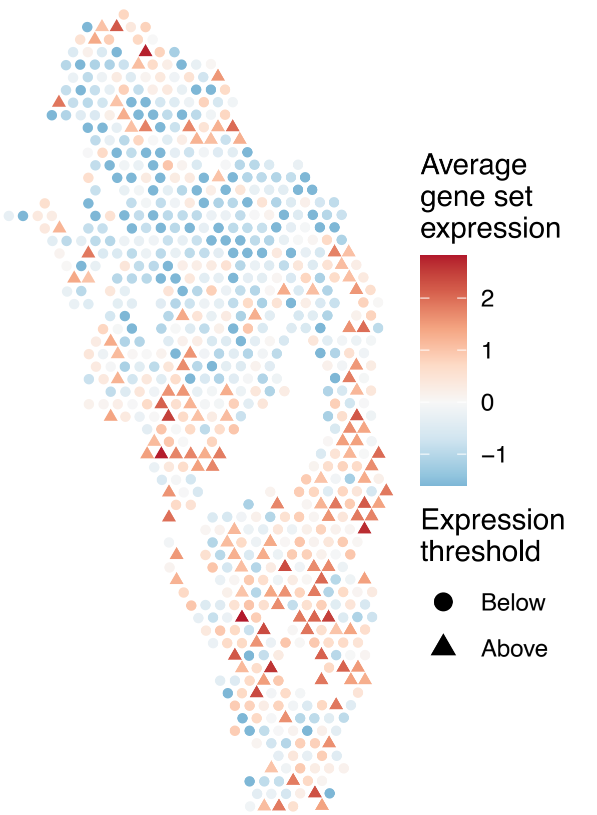


sample13_LMM_B1
KEGG_BUTANOATE_METABOLISM
Proportion of pathway genes=0.57; Adjusted p−value=0.0435

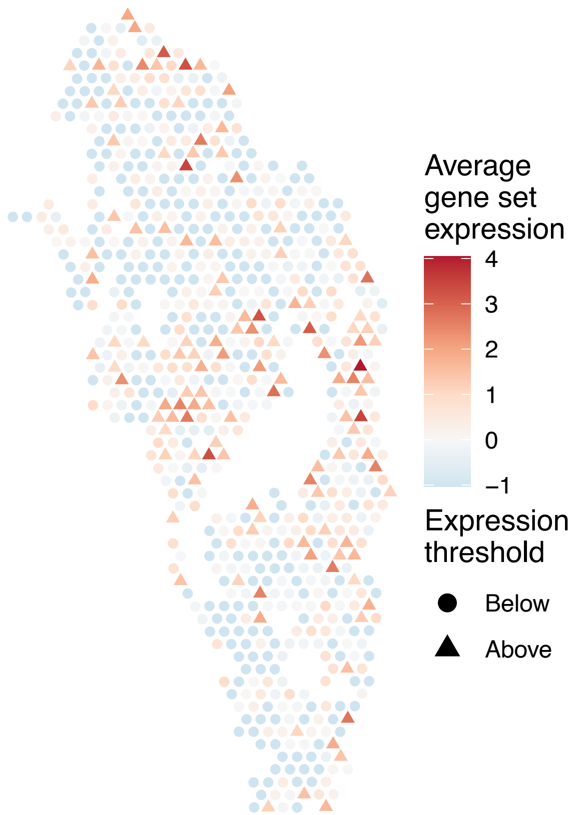


sample13_LMM_B1
KEGG_CALCIUM_SIGNALING_PATHWAY
Proportion of pathway genes=0.36; Adjusted p−value=0.0101

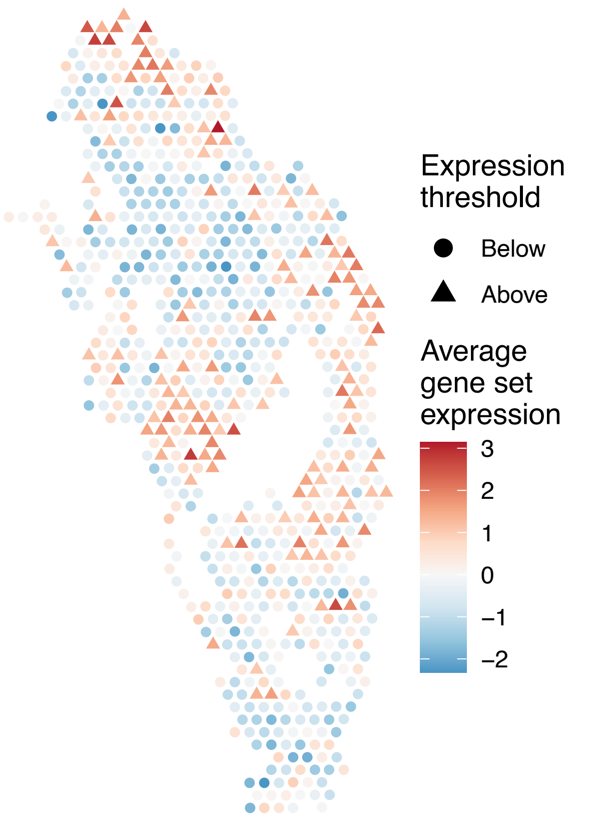

sample13_LMM_B1
KEGG_CARDIAC_MUSCLE_CONTRACTION
Proportion of pathway genes=0.48; Adjusted p−value=0

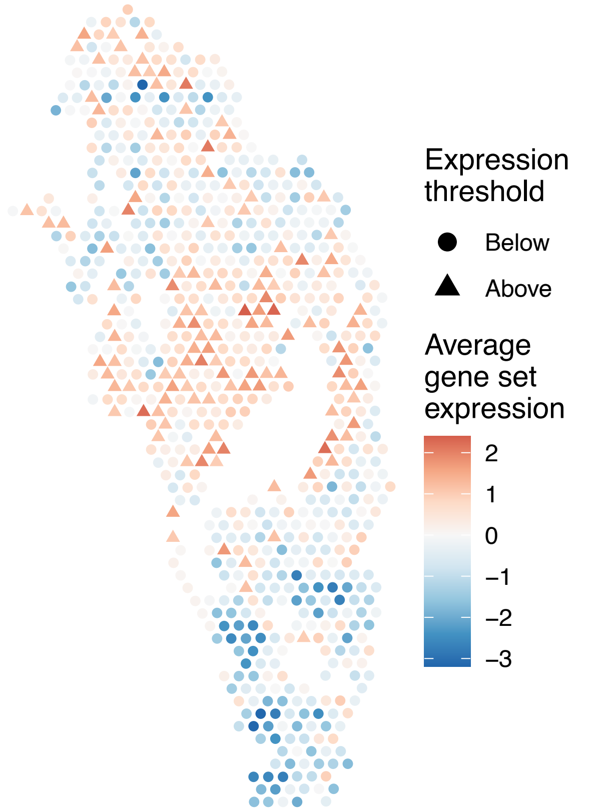


sample13_LMM_B1
KEGG_CELL_ADHESION_MOLECULES_CAMS
Proportion of pathway genes=0.67; Adjusted p−value=0.4721

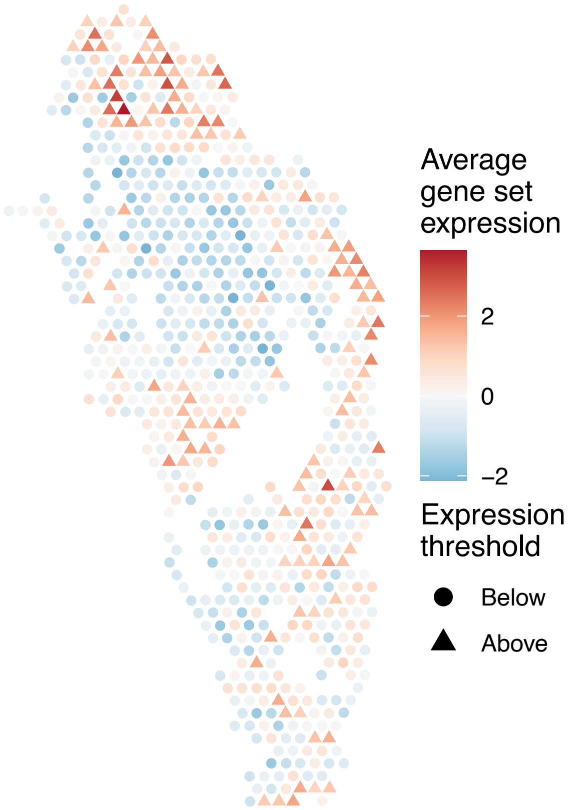


sample13_LMM_B1
KEGG_CELL_CYCLE
Proportion of pathway genes=0.84; Adjusted p−value=0

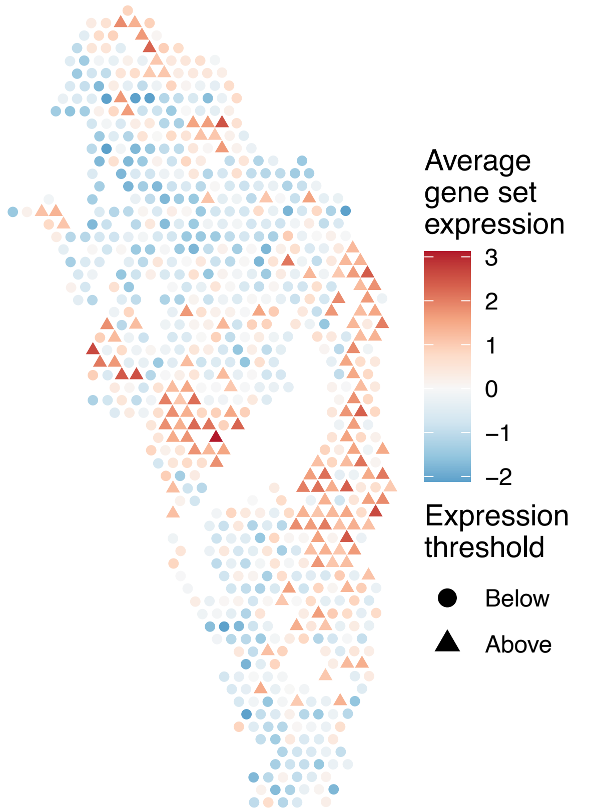


sample13_LMM_B1
KEGG_CHEMOKINE_SIGNALING_PATHWAY
Proportion of pathway genes=0.54; Adjusted p−value=0.7836

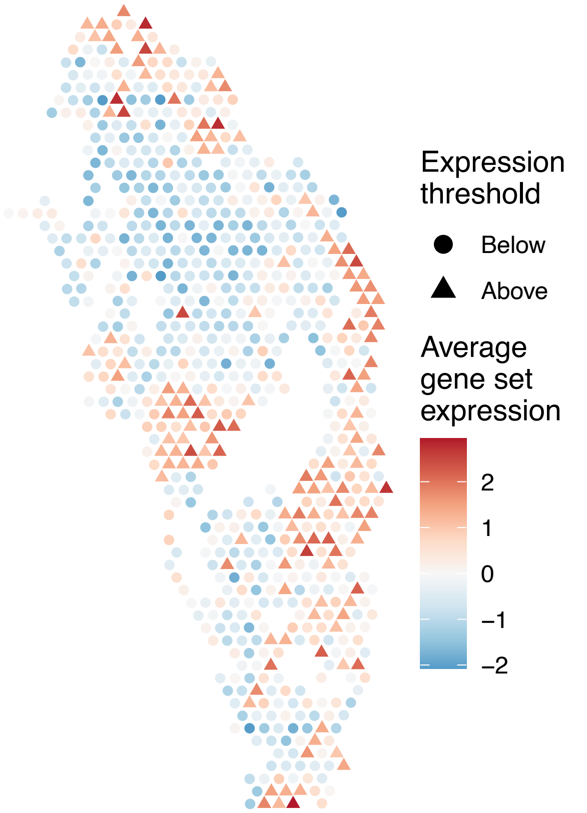

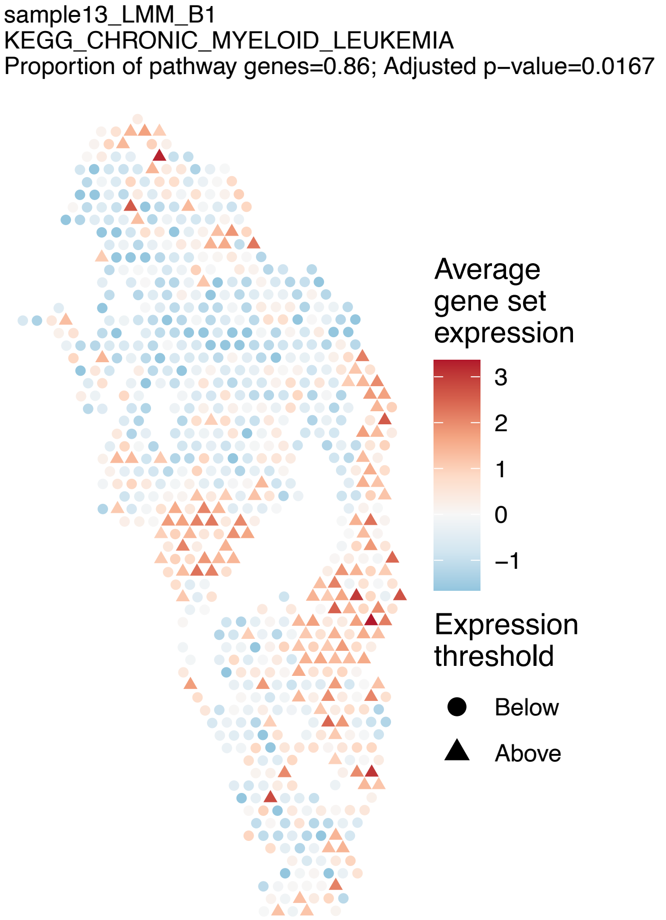
sample13_LMM_B1
KEGG_CHRONIC_MYELOID_LEUKEMIA
Proportion of pathway genes=0.86; Adjusted p−value=0.0167
Average gene set expression
3
2
1
0
−1
Expression threshold
Below
Above

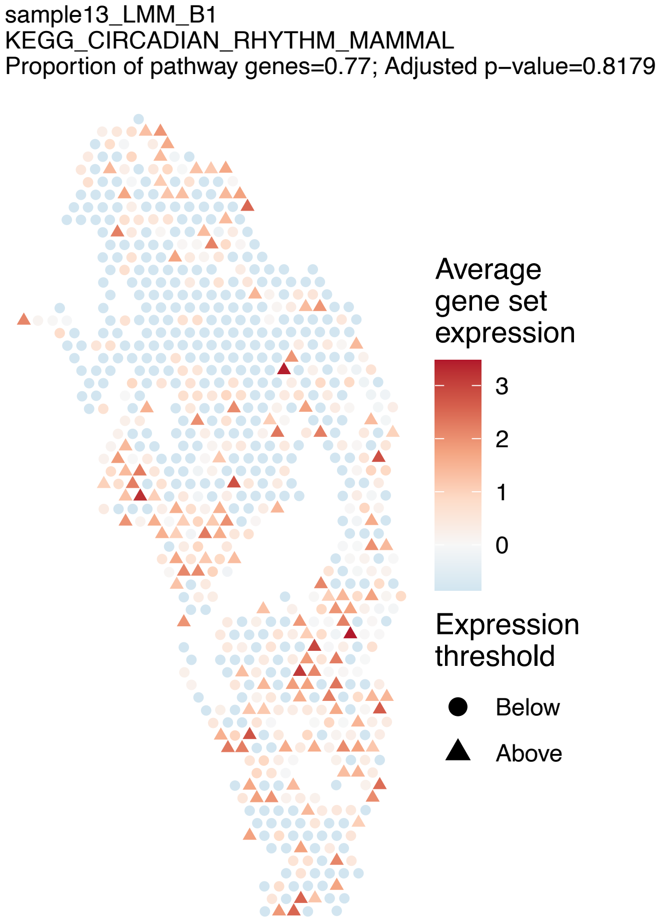
sample13_LMM_B1
KEGG_CIRCADIAN_RHYTHM_MAMMAL
Proportion of pathway genes=0.77; Adjusted p−value=0.8179
Average gene set expression
3
2
1
0
Expression threshold
Below
Above

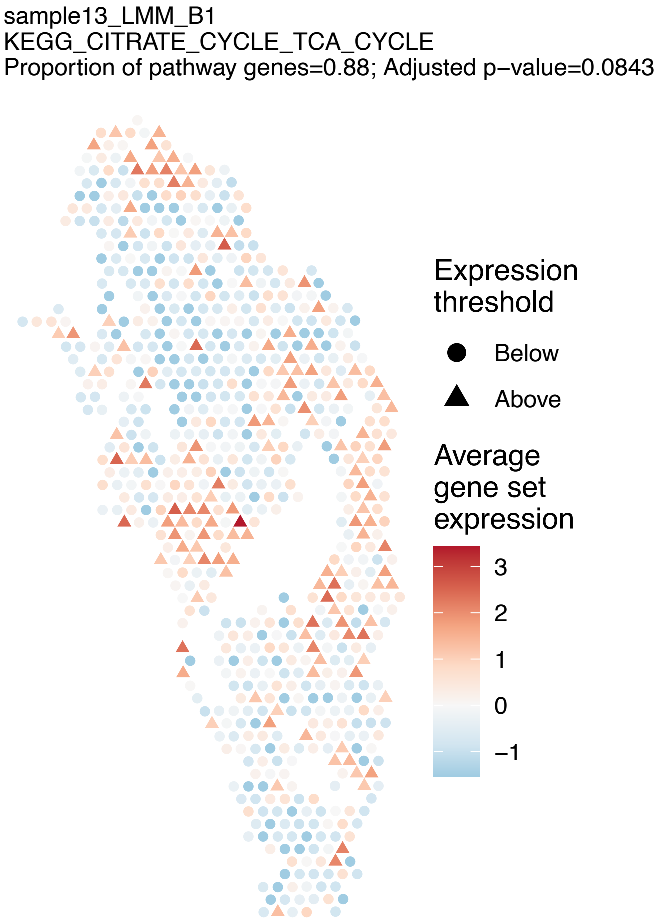
sample13_LMM_B1
KEGG_CITRATE_CYCLE_TCA_CYCLE
Proportion of pathway genes=0.88; Adjusted p−value=0.0843
Expression threshold
Below
Above
Average gene set expression
3
2
1
0
−1

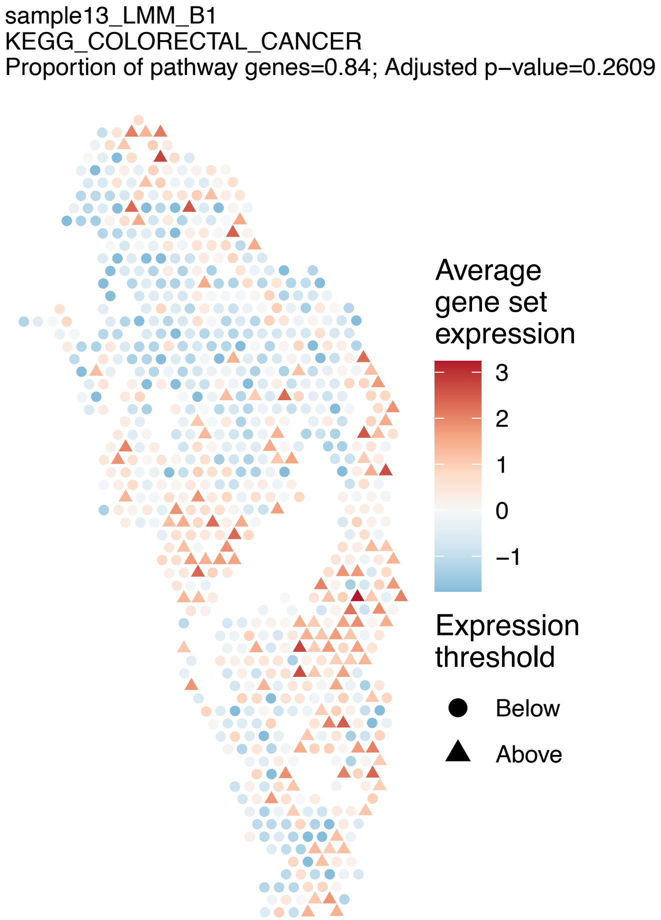
sample13_LMM_B1
KEGG_COLORECTAL_CANCER
Proportion of pathway genes=0.84; Adjusted p−value=0.2609
Average gene set expression
3
2
1
0
−1
Expression threshold
Below
Above

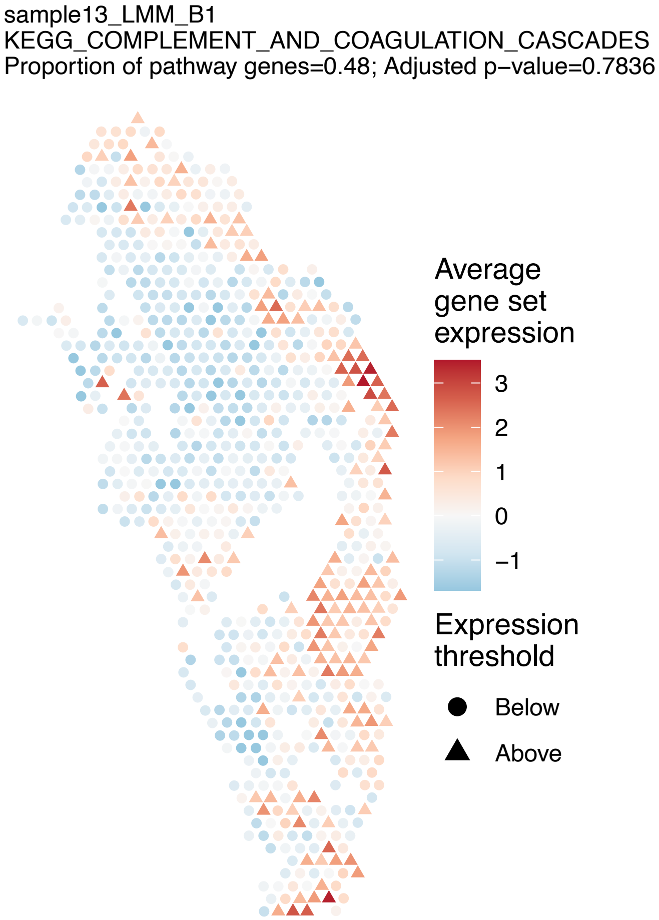
sample13_LMM_B1
KEGG_COMPLEMENT_AND_COAGULATION_CASCADES
Proportion of pathway genes=0.48; Adjusted p−value=0.7836
Average gene set expression
3
2
1
0
−1
Expression threshold
Below
Above

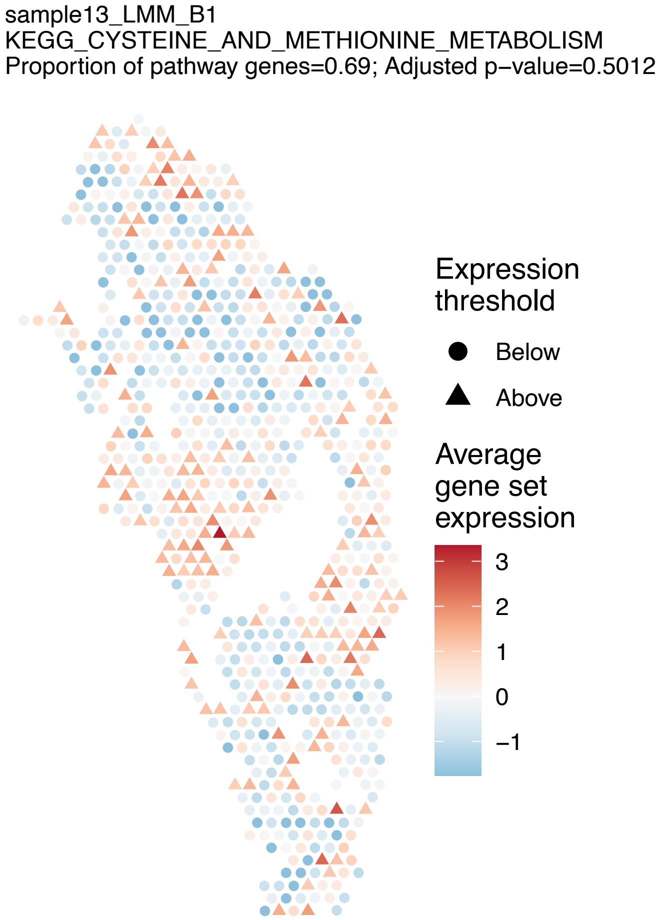
sample13_LMM_B1
KEGG_CYSTEINE_AND_METHIONINE_METABOLISM
Proportion of pathway genes=0.69; Adjusted p−value=0.5012
Expression threshold
Below
Above
Average gene set expression
3
2
1
0
−1

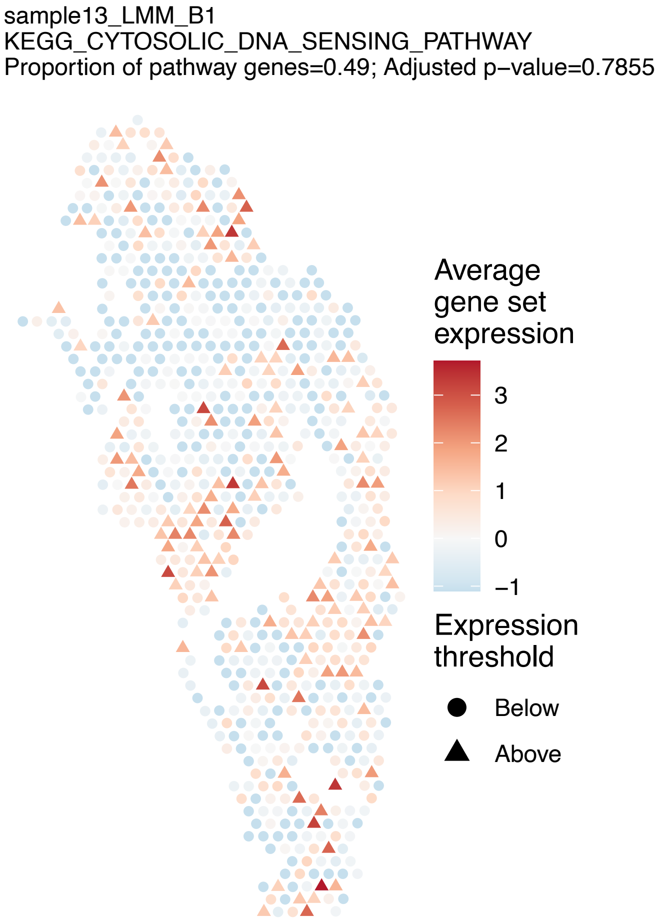
sample13_LMM_B1
KEGG_CYTOSOLIC_DNA_SENSING_PATHWAY
Proportion of pathway genes=0.49; Adjusted p−value=0.7855
Average gene set expression
3
2
1
0
−1
Expression threshold
Below
Above

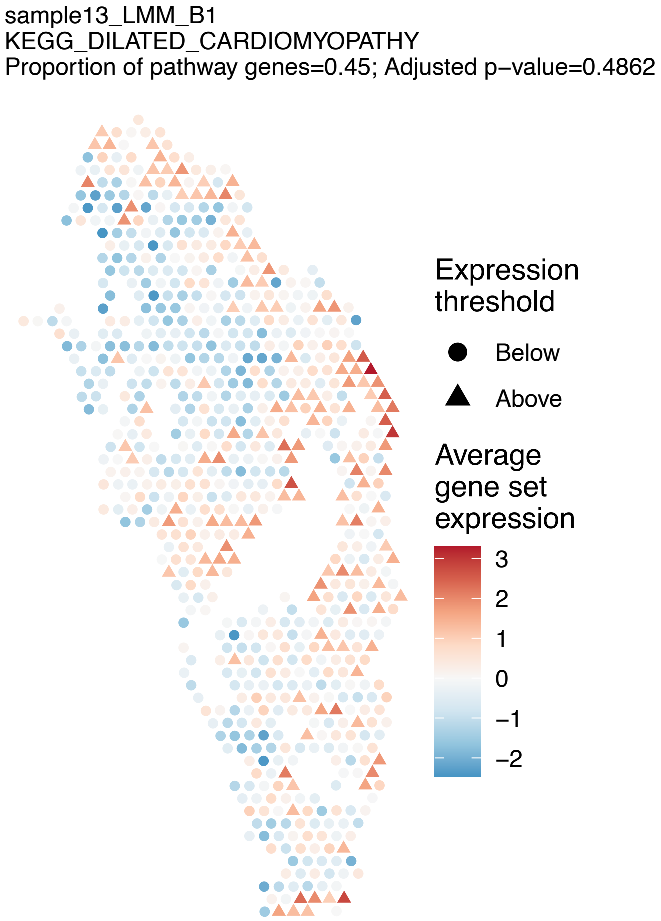
sample13_LMM_B1
KEGG_DILATED_CARDIOMYOPATHY
Proportion of pathway genes=0.45; Adjusted p−value=0.4862
Expression threshold
Below
Above
Average gene set expression
3
2
1
0
−1
−2

sample13_LMM_B1
KEGG_DNA_REPLICATION
Proportion of pathway genes=0.84; Adjusted p−value=0

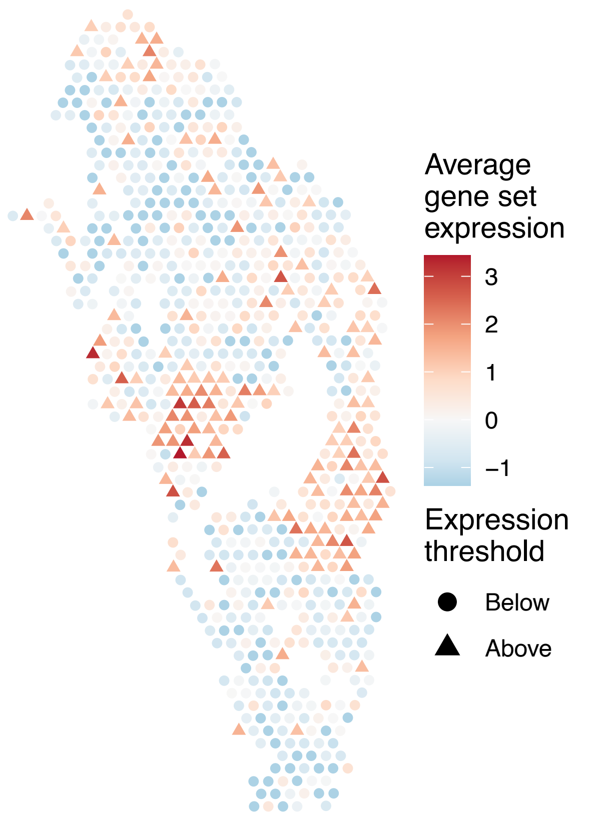


sample13_LMM_B1
KEGG_DORSO_VENTRAL_AXIS_FORMATION
Proportion of pathway genes=0.72; Adjusted p−value=0.114

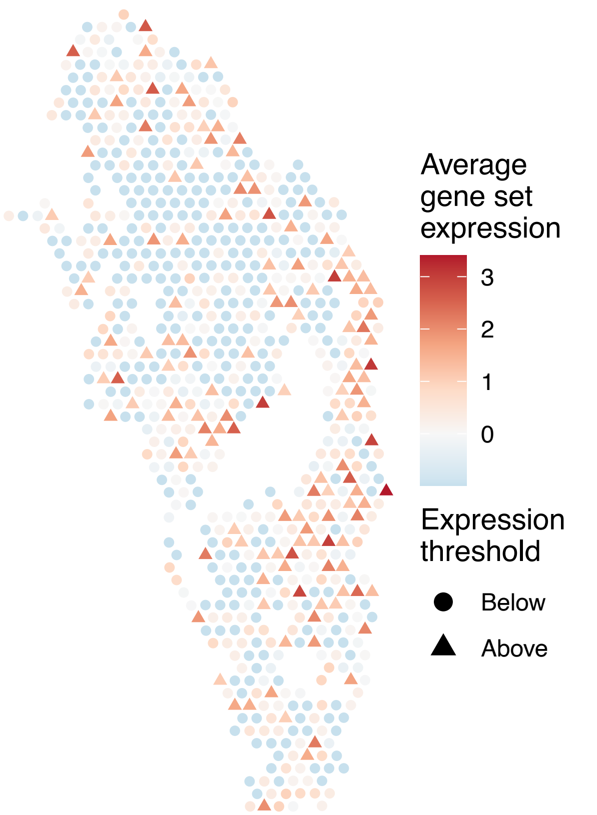


sample13_LMM_B1
KEGG_DRUG_METABOLISM_CYTOCHROME_P450
Proportion of pathway genes=0.27; Adjusted p−value=0.6738

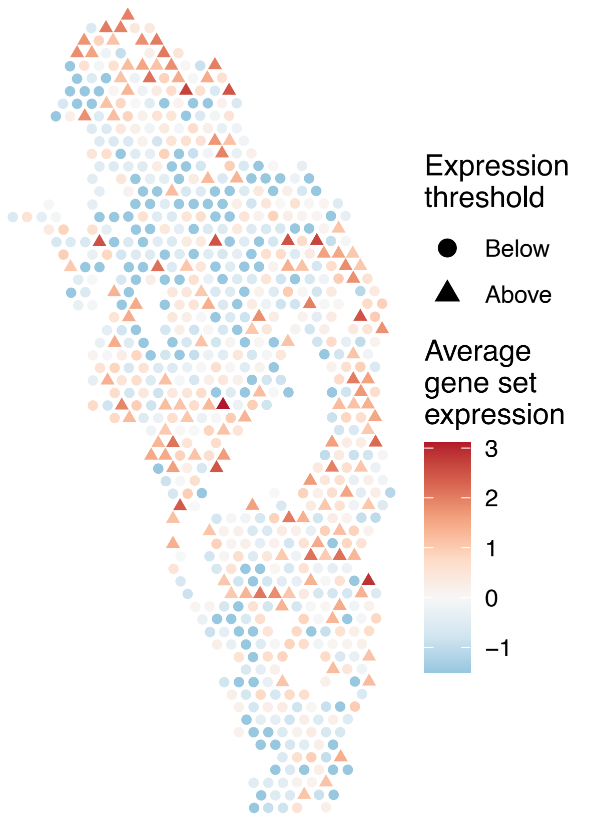


sample13_LMM_B1
KEGG_DRUG_METABOLISM_OTHER_ENZYMES
Proportion of pathway genes=0.3; Adjusted p−value=0.0218

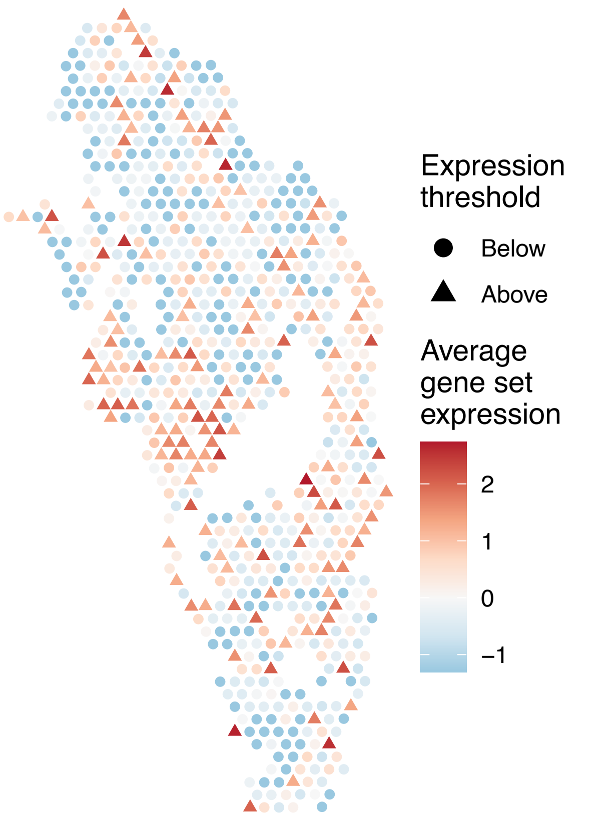

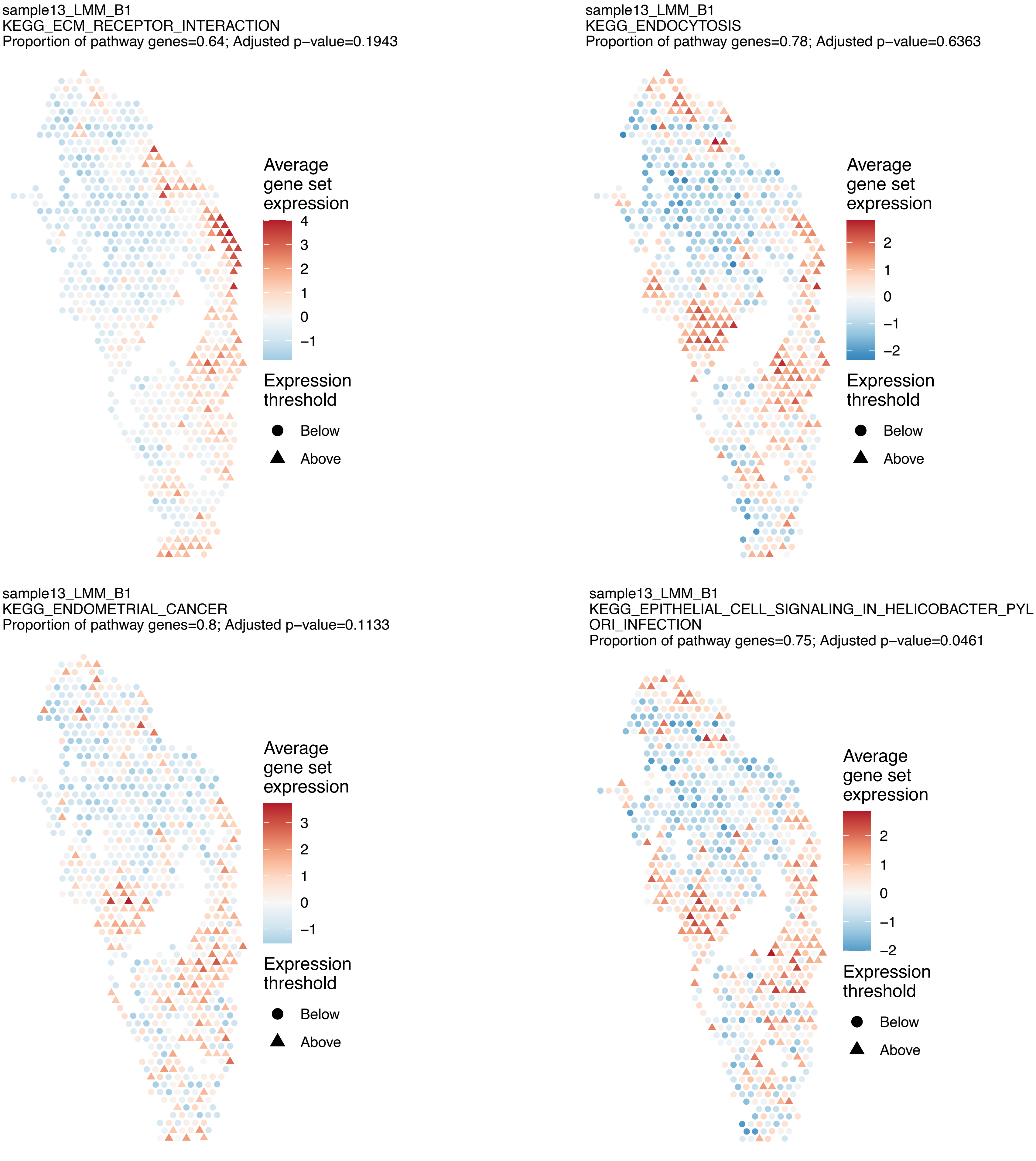
sample13_LMM_B1
KEGG_ECM_RECEPTOR_INTERACTION
Proportion of pathway genes=0.64; Adjusted p−value=0.1943
Average gene set expression
4
3
2
1
0
−1
Expression threshold
Below
Above
sample13_LMM_B1
KEGG_ENDOCYTOSIS
Proportion of pathway genes=0.78; Adjusted p−value=0.6363
Average gene set expression
2
1
0
−1
−2
Expression threshold
Below
Above
sample13_LMM_B1
KEGG_ENDOMETRIAL_CANCER
Proportion of pathway genes=0.8; Adjusted p−value=0.1133
Average gene set expression
3
2
1
0
−1
Expression threshold
Below
Above
sample13_LMM_B1
KEGG_EPITHELIAL_CELL_SIGNALING_IN_HELICOBACTER_PYL
ORI_INFECTION
Proportion of pathway genes=0.75; Adjusted p−value=0.0461
Average gene set expression
2
1
0
−1
−2
Expression threshold
Below
Above

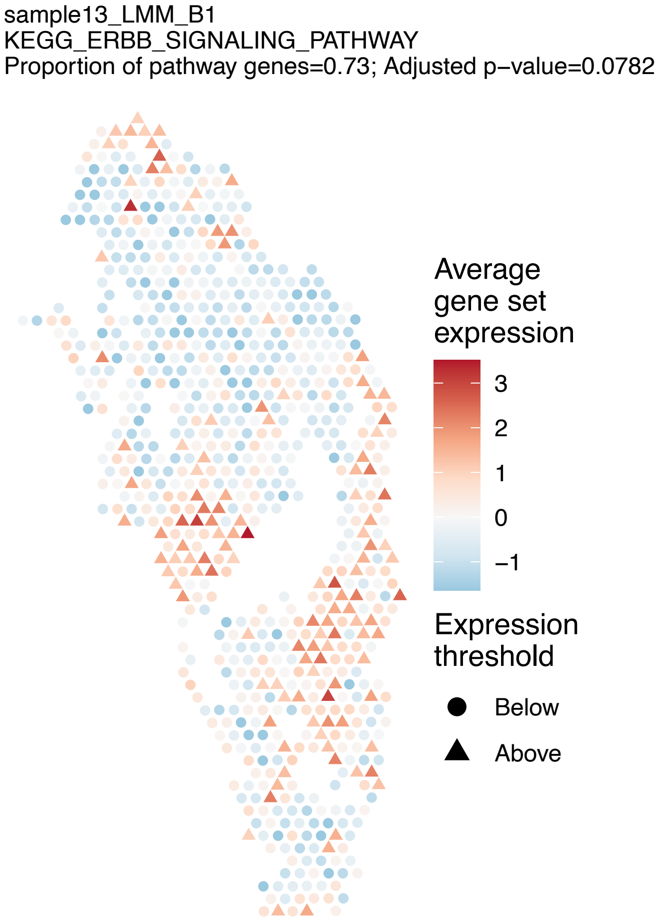
sample13_LMM_B1
KEGG_ERBB_SIGNALING_PATHWAY
Proportion of pathway genes=0.73; Adjusted p−value=0.0782
Average gene set expression
3
2
1
0
−1
Expression threshold
Below
Above

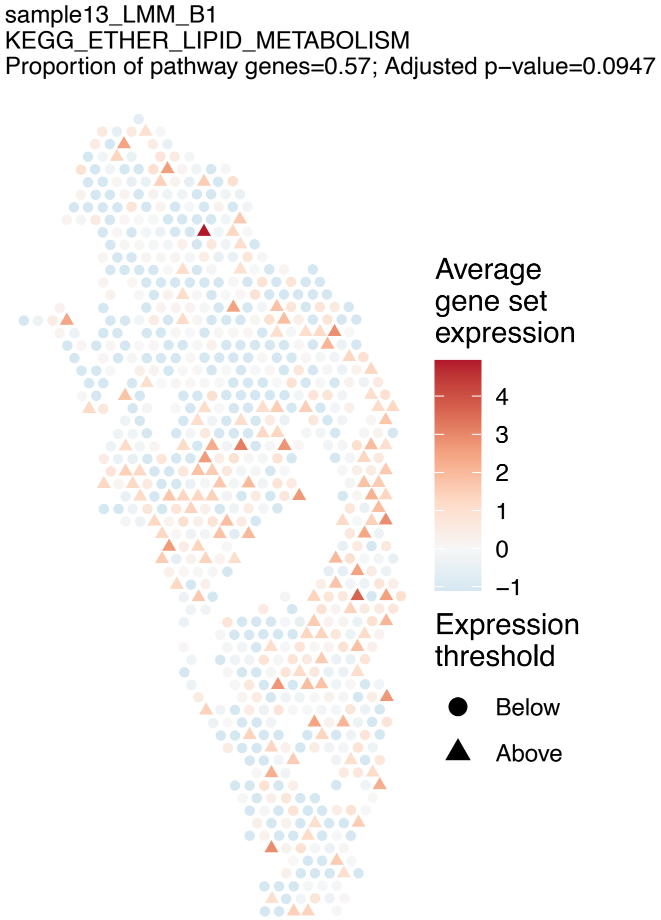
sample13_LMM_B1
KEGG_ETHER_LIPID_METABOLISM
Proportion of pathway genes=0.57; Adjusted p−value=0.0947
Average gene set expression
4
3
2
1
0
−1
Expression threshold
Below
Above

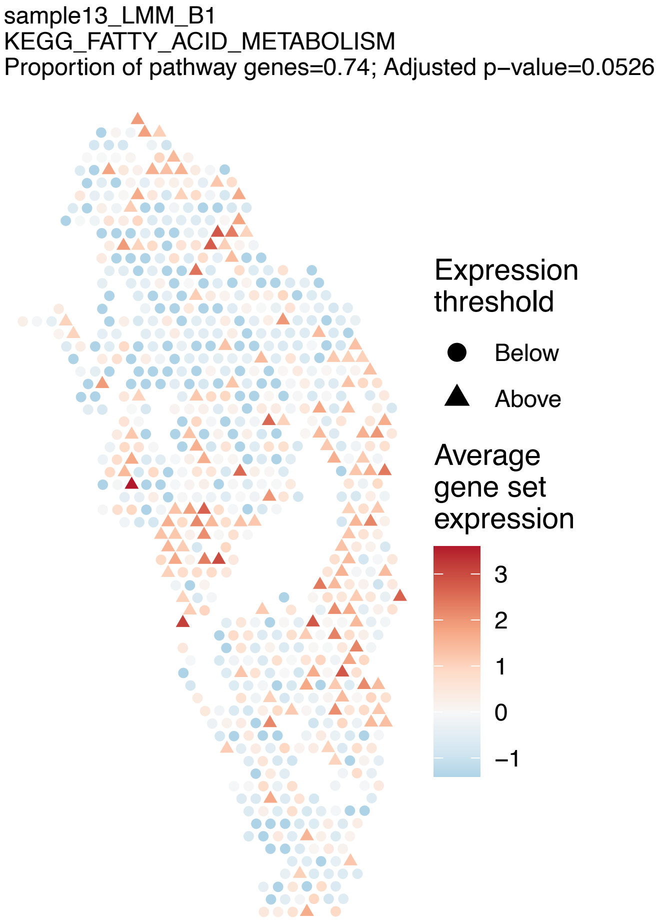
sample13_LMM_B1
KEGG_FATTY_ACID_METABOLISM
Proportion of pathway genes=0.74; Adjusted p−value=0.0526
Expression threshold
Below
Above
Average gene set expression
3
2
1
0
−1

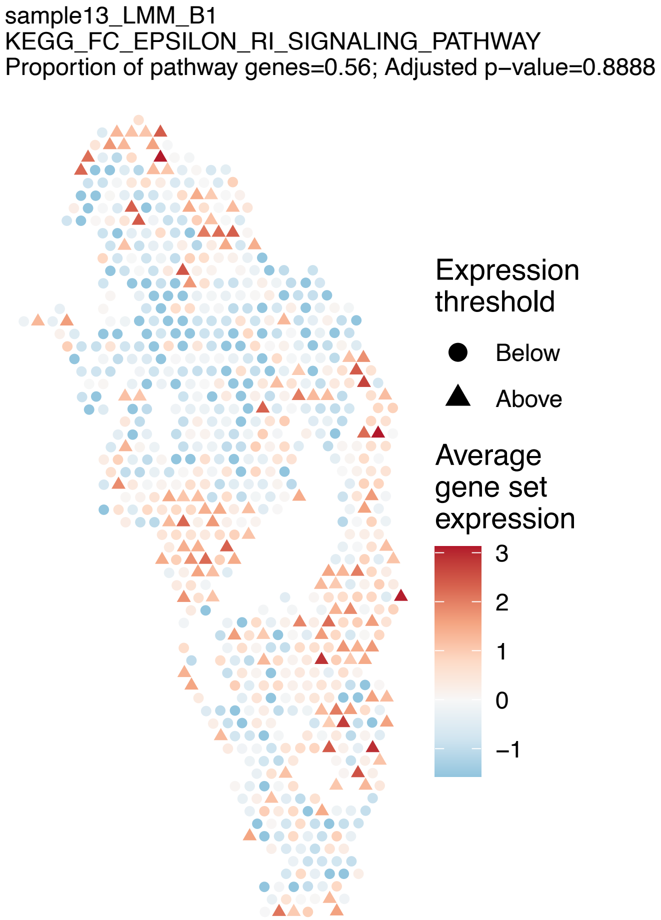
sample13_LMM_B1
KEGG_FC_EPSILON_RI_SIGNALING_PATHWAY
Proportion of pathway genes=0.56; Adjusted p−value=0.8888
Expression threshold
Below
Above
Average gene set expression
3
2
1
0
−1

sample13_LMM_B1
KEGG_FC_GAMMA_R_MEDIATED_PHAGOCYTOSIS
Proportion of pathway genes=0.77; Adjusted p−value=0.5875

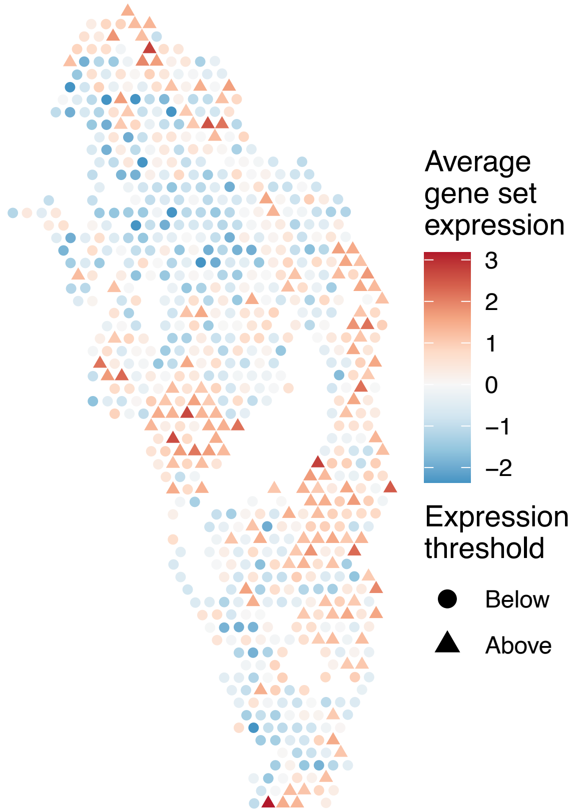


sample13_LMM_B1
KEGG_FOCAL_ADHESION
Proportion of pathway genes=0.78; Adjusted p−value=0.2815

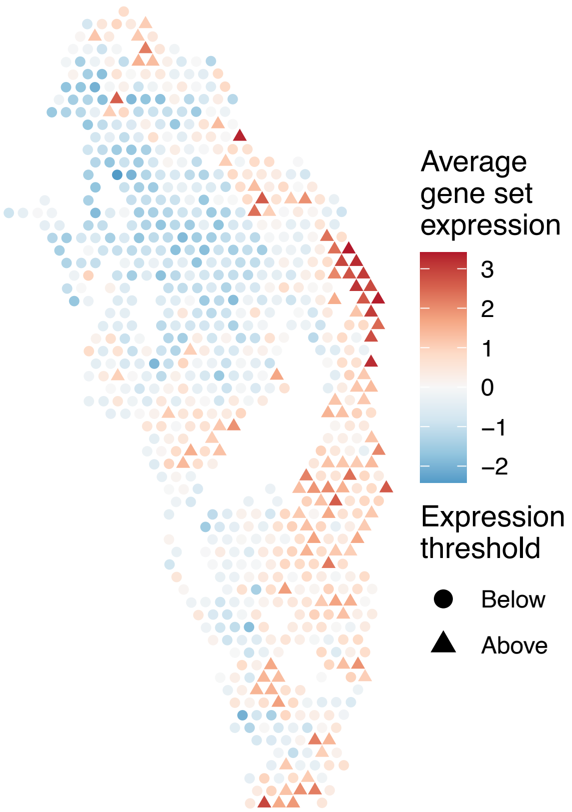


sample13_LMM_B1
KEGG_FRUCTOSE_AND_MANNOSE_METABOLISM
Proportion of pathway genes=0.69; Adjusted p−value=1

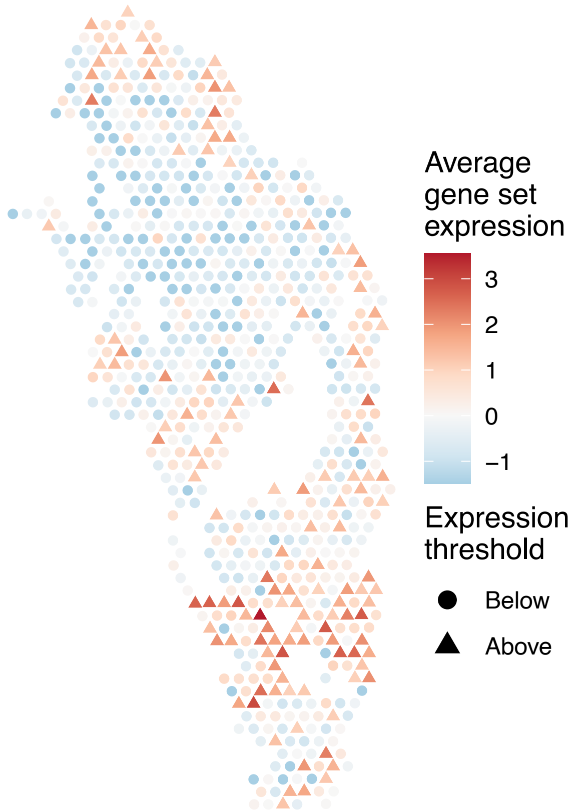


sample13_LMM_B1
KEGG_GALACTOSE_METABOLISM
Proportion of pathway genes=0.66; Adjusted p−value=0.5904

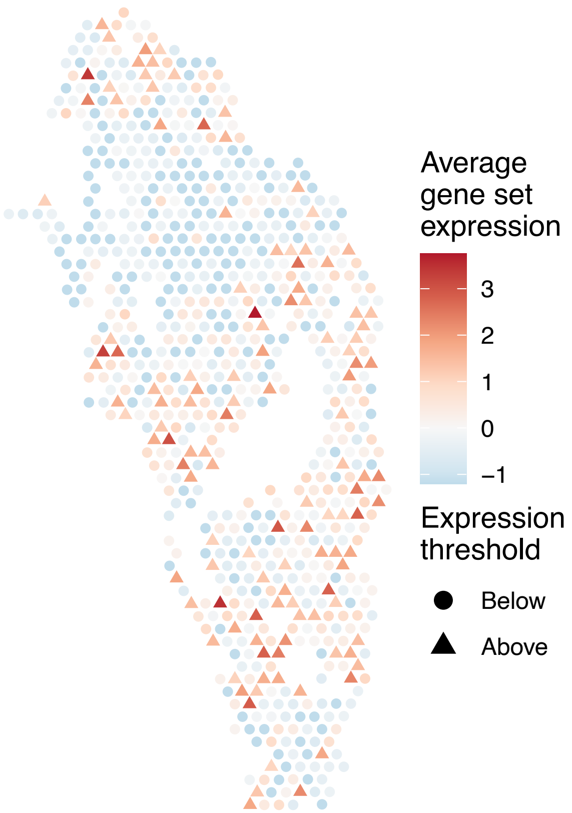

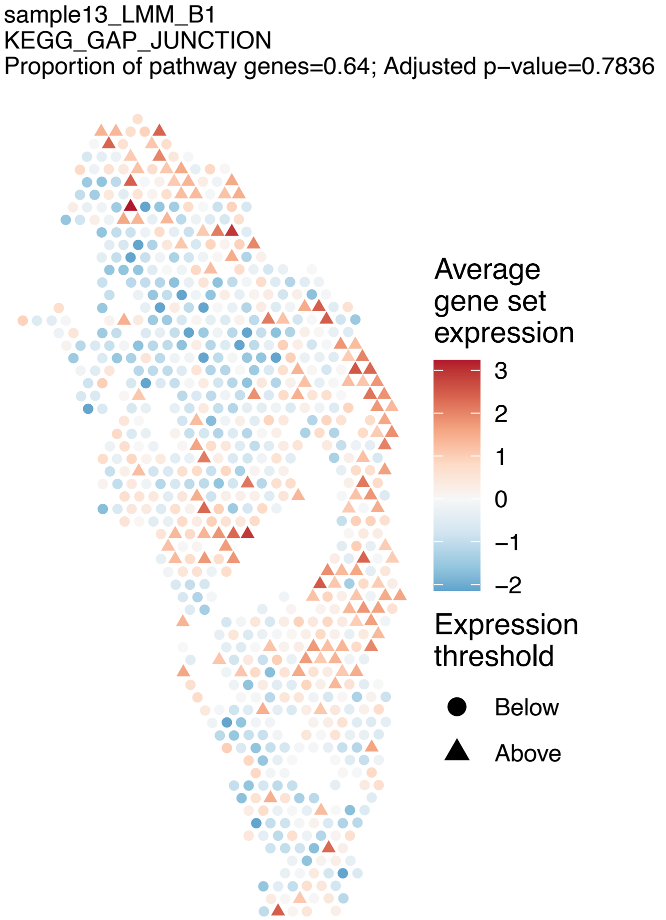
sample13_LMM_B1
KEGG_GAP_JUNCTION
Proportion of pathway genes=0.64; Adjusted p−value=0.7836
Average gene set expression
3
2
1
0
−1
−2
Expression threshold
Below
Above

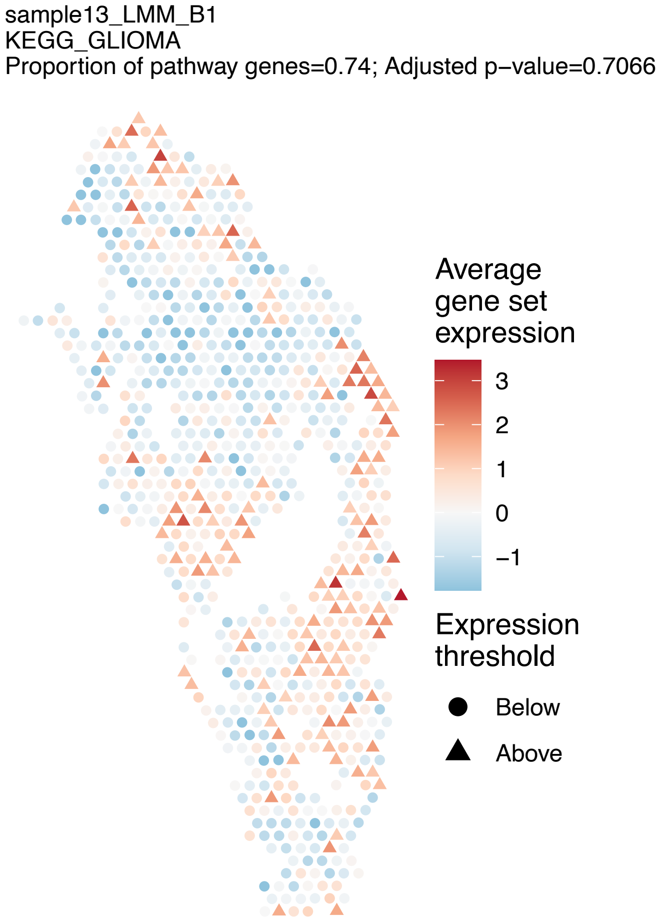
sample13_LMM_B1
KEGG_GLIOMA
Proportion of pathway genes=0.74; Adjusted p−value=0.7066
Average gene set expression
3
2
1
0
−1
Expression threshold
Below
Above

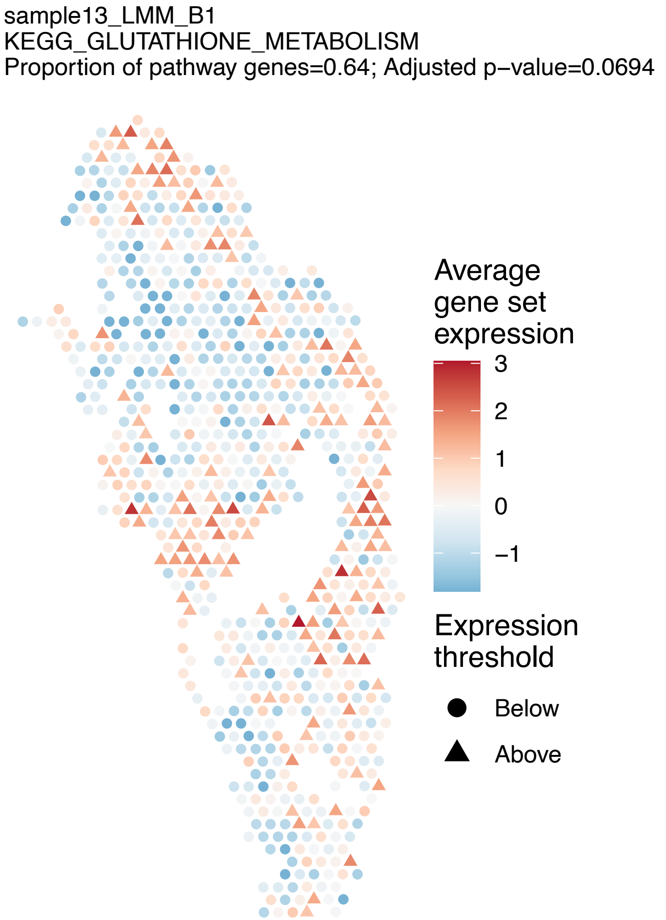
sample13_LMM_B1
KEGG_GLUTATHIONE_METABOLISM
Proportion of pathway genes=0.64; Adjusted p−value=0.0694
Average gene set expression
3
2
1
0
−1
Expression threshold
Below
Above

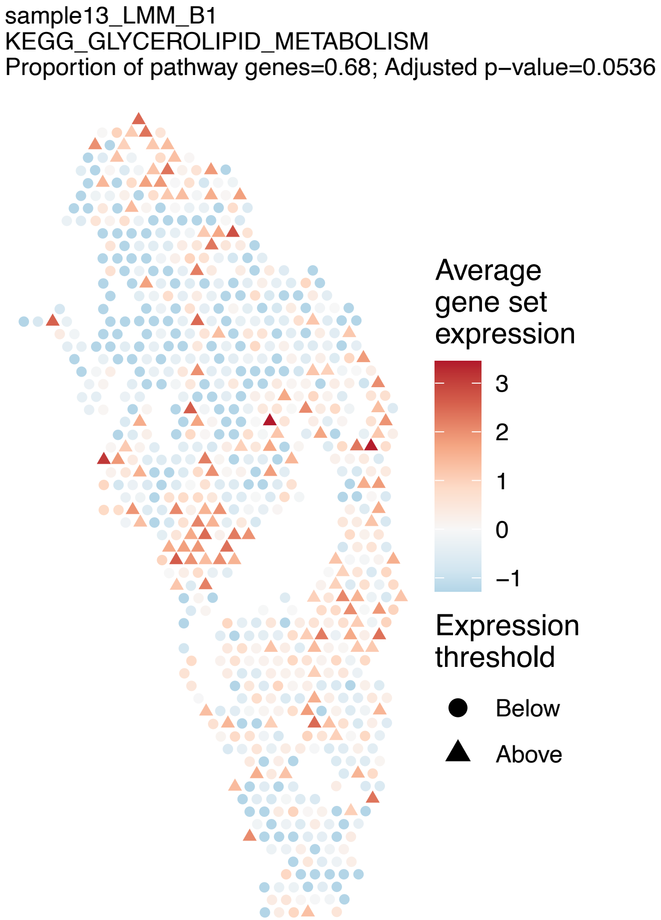
sample13_LMM_B1
KEGG_GLYCEROLIPID_METABOLISM
Proportion of pathway genes=0.68; Adjusted p−value=0.0536
Average gene set expression
3
2
1
0
−1
Expression threshold
Below
Above

sample13_LMM_B1
KEGG_GLYCEROPHOSPHOLIPID_METABOLISM
Proportion of pathway genes=0.72; Adjusted p–value=0.019

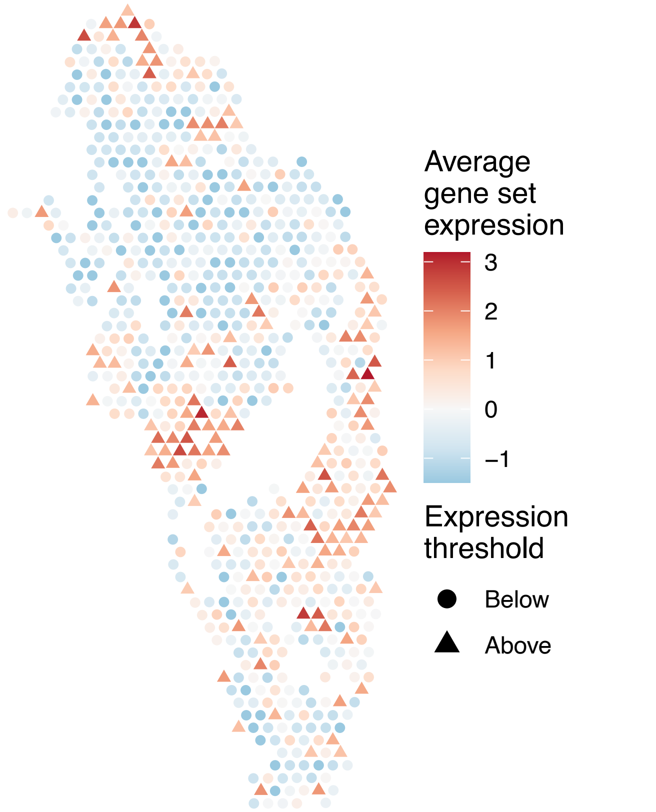


sample13_LMM_B1
KEGG_GLYCINE_SERINE_AND_THREONINE_METABOLISM
Proportion of pathway genes=0.53; Adjusted p–value=0.6724

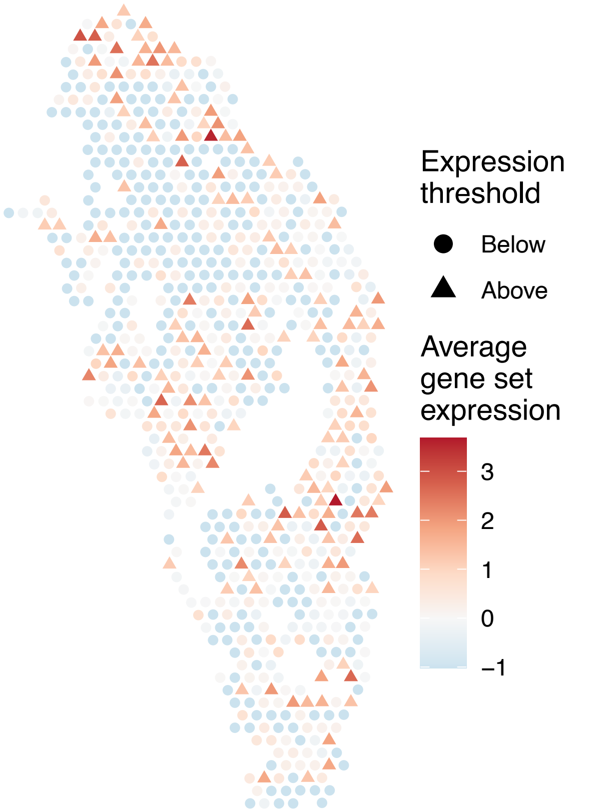


sample13_LMM_B1
KEGG_GLYCOLYSIS_GLUCONEOGENESIS
Proportion of pathway genes=0.63; Adjusted p–value=0.2556

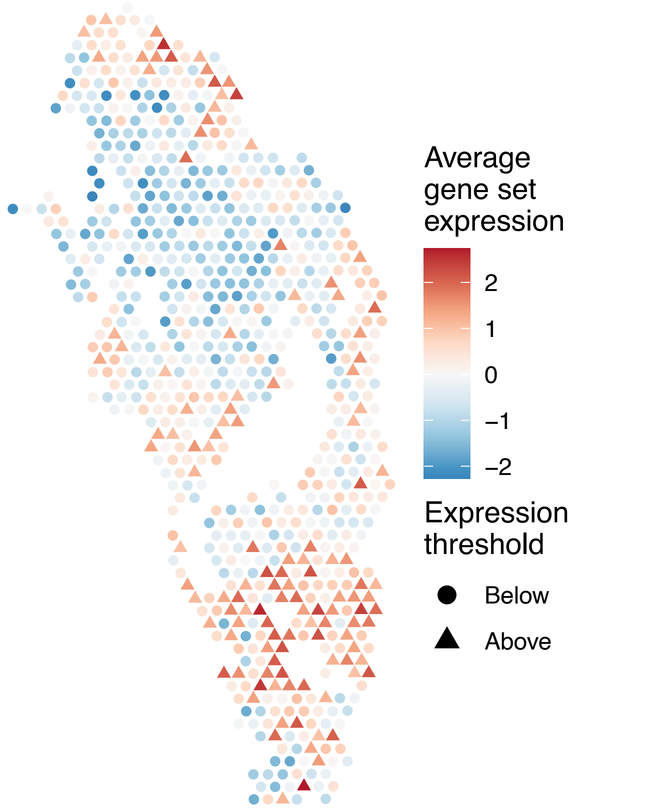


sample13_LMM_B1
KEGG_GLYCOSAMINOGLYCAN_BIOSYNTHESIS_CHONDROITIN_SU
LFATE
Proportion of pathway genes=0.78; Adjusted p–value=0.6025

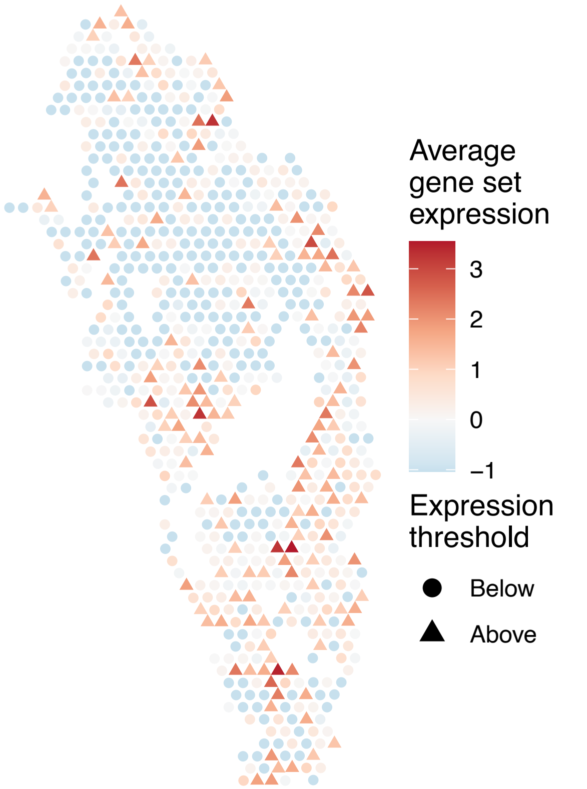

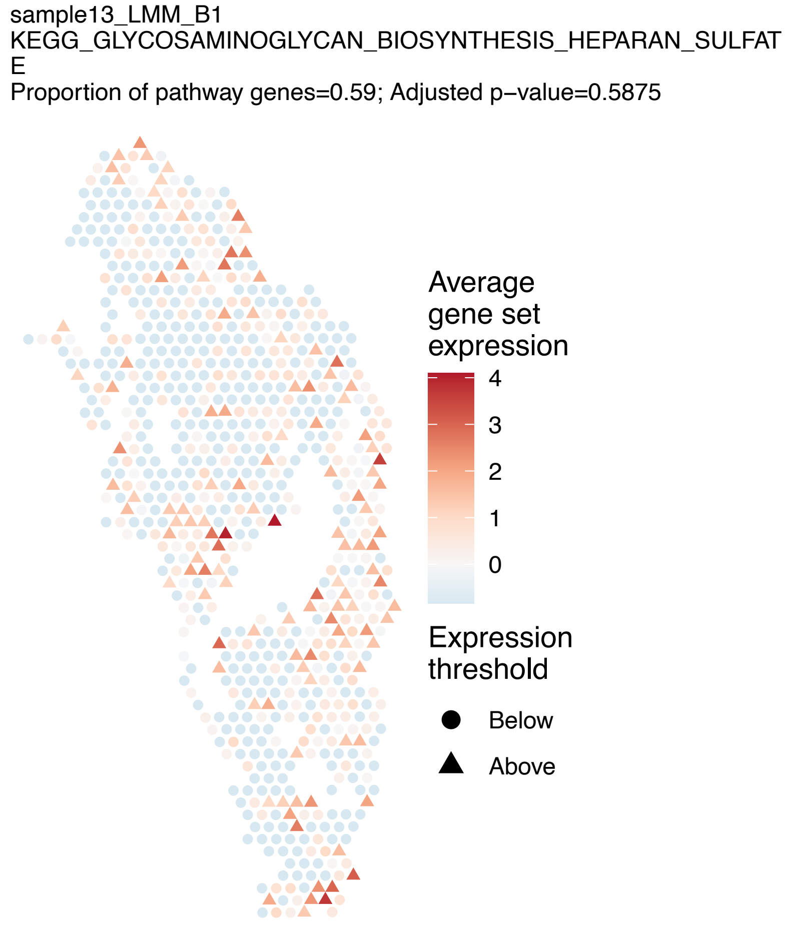
sample13_LMM_B1
KEGG_GLYCOSAMINOGLYCAN_BIOSYNTHESIS_HEPARAN_SULFAT
E
Proportion of pathway genes=0.59; Adjusted p−value=0.5875
Average gene set expression
4
3
2
1
0
Expression threshold
Below
Above

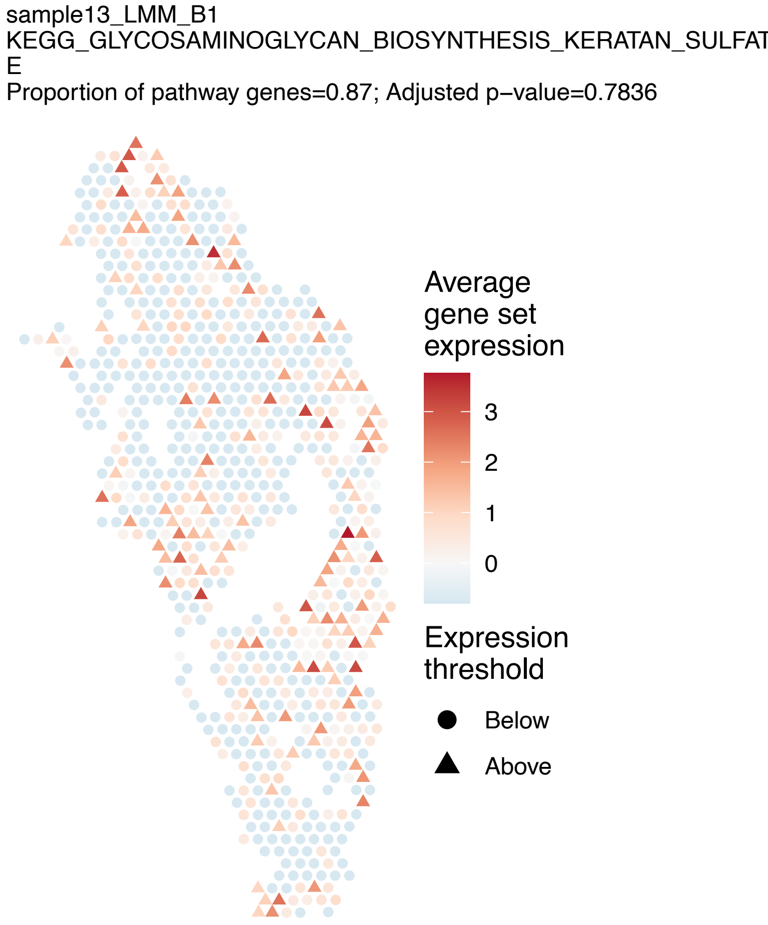
sample13_LMM_B1
KEGG_GLYCOSAMINOGLYCAN_BIOSYNTHESIS_KERATAN_SULFAT
E
Proportion of pathway genes=0.87; Adjusted p−value=0.7836
Average gene set expression
3
2
1
0
Expression threshold
Below
Above

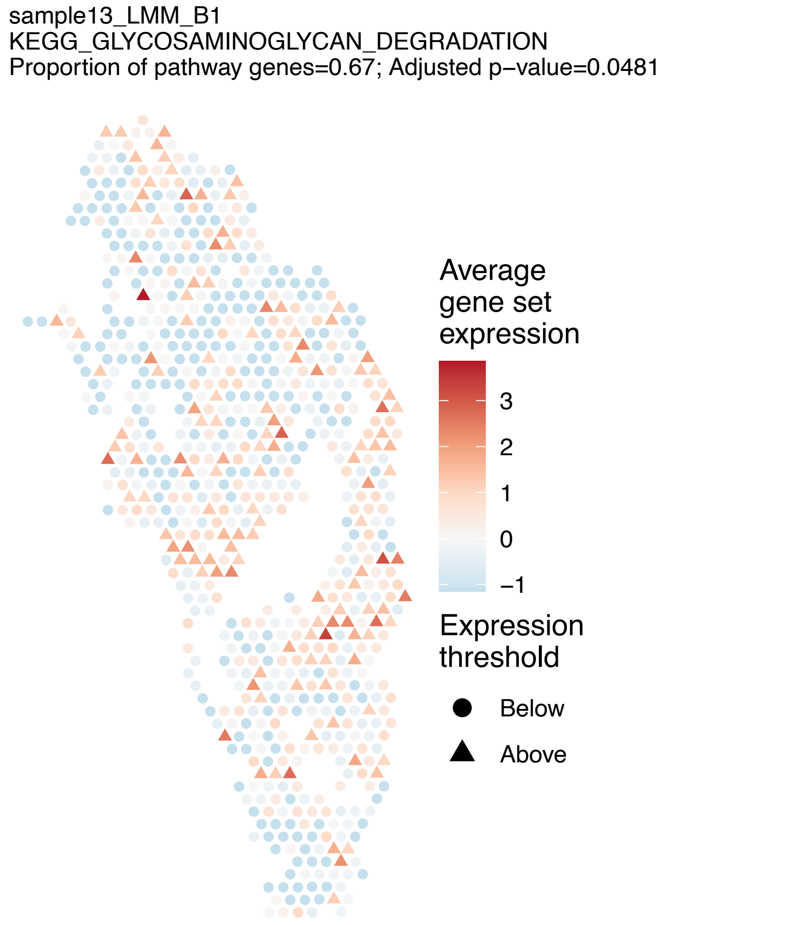
sample13_LMM_B1
KEGG_GLYCOSAMINOGLYCAN_DEGRADATION
Proportion of pathway genes=0.67; Adjusted p−value=0.0481
Average gene set expression
3
2
1
0
−1
Expression threshold
Below
Above

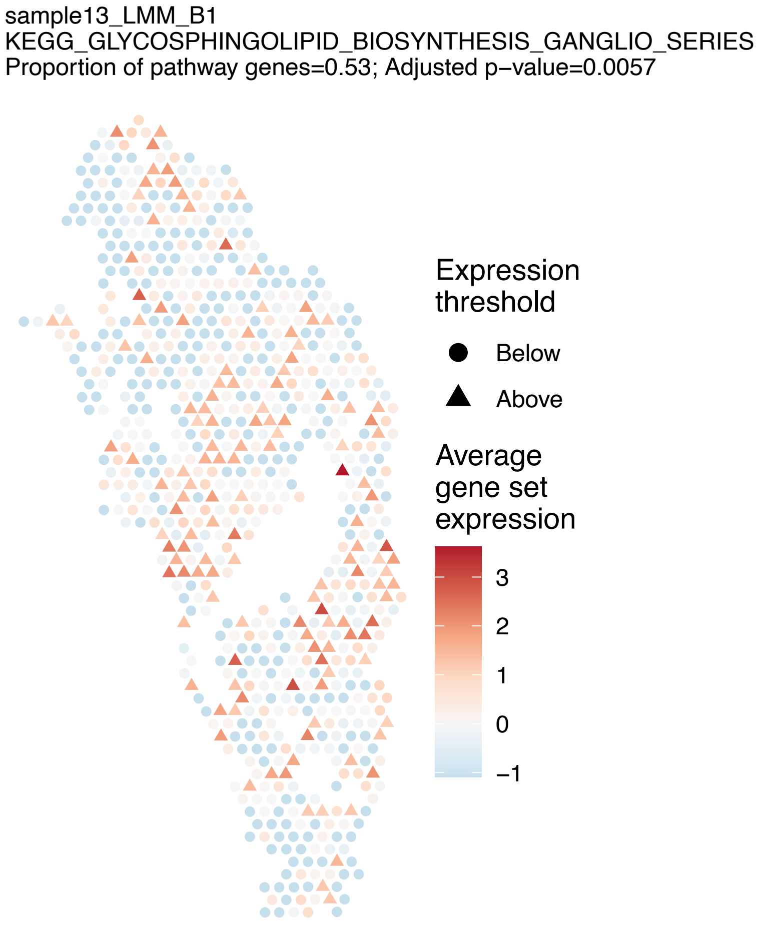
sample13_LMM_B1
KEGG_GLYCOSPHINGOLIPID_BIOSYNTHESIS_GANGLIO_SERIES
Proportion of pathway genes=0.53; Adjusted p−value=0.0057
Expression threshold
Below
Above
Average gene set expression
3
2
1
0
−1

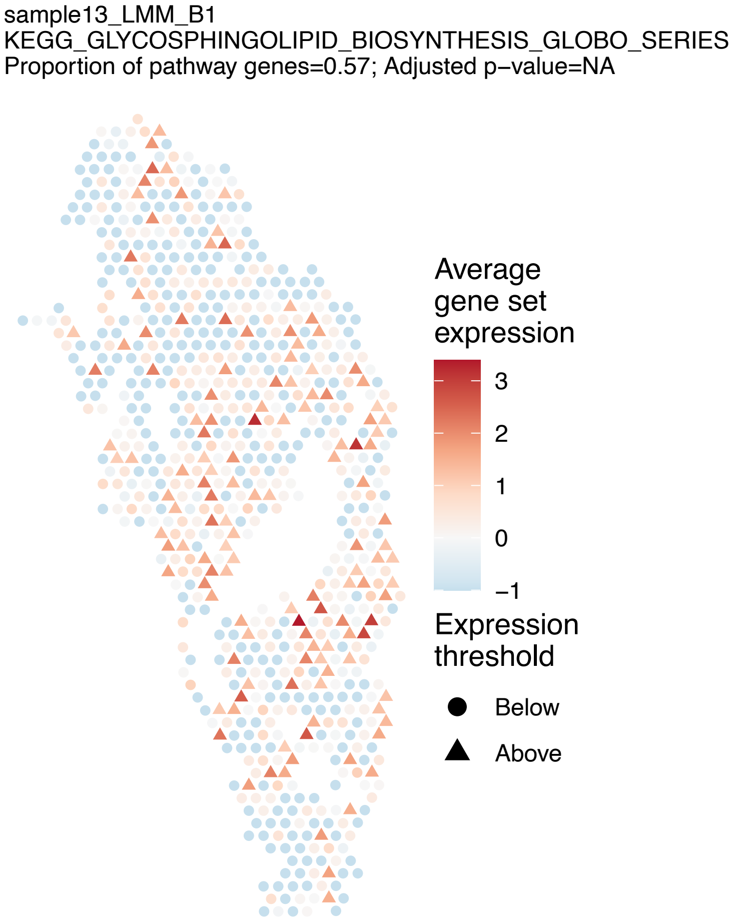
sample13_LMM_B1
KEGG_GLYCOSPHINGOLIPID_BIOSYNTHESIS_GLOBO_SERIES
Proportion of pathway genes=0.57; Adjusted p−value=NA
Average gene set expression
3
2
1
0
−1
Expression threshold
Below
Above

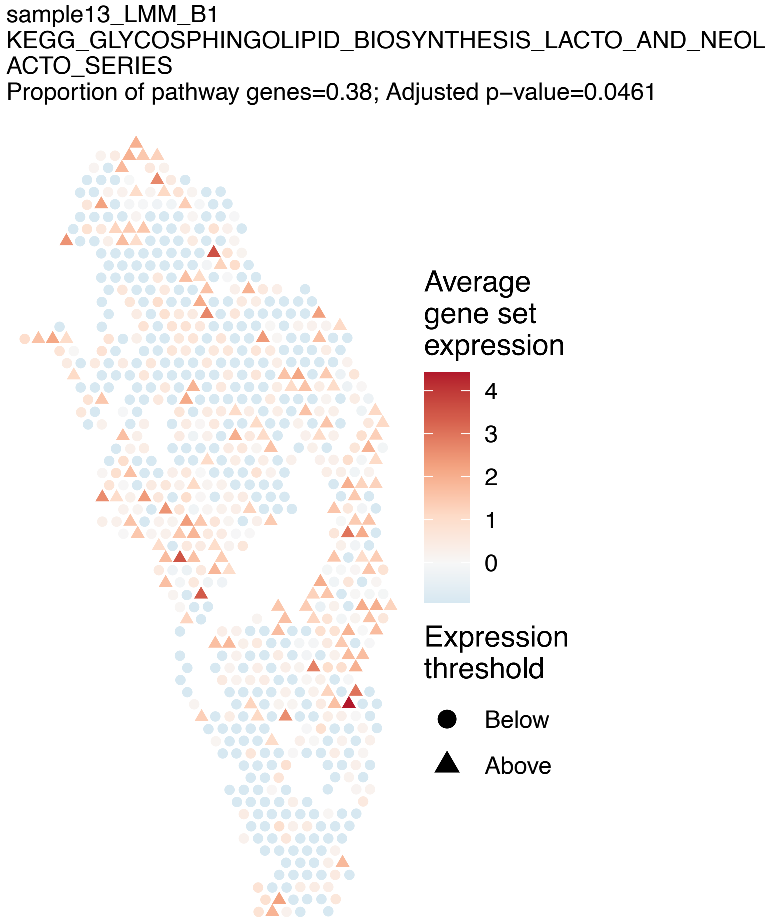
sample13_LMM_B1
KEGG_GLYCOSPHINGOLIPID_BIOSYNTHESIS_LACTO_AND_NEOLACTO_SERIES
Proportion of pathway genes=0.38; Adjusted p−value=0.0461
Average gene set expression
4
3
2
1
0
Expression threshold
Below
Above

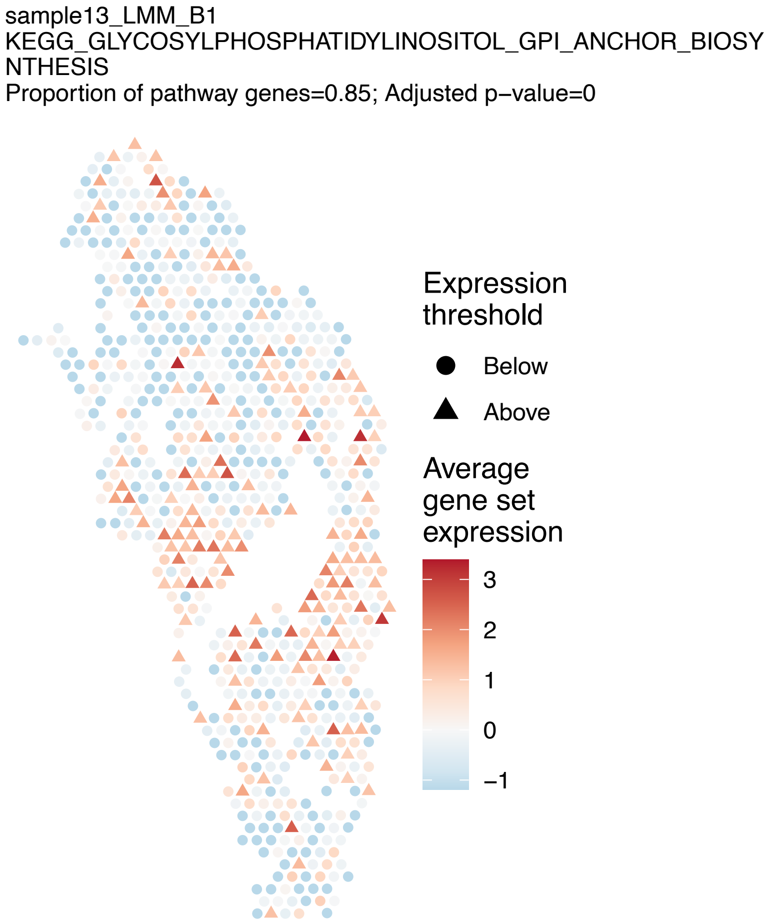
sample13_LMM_B1
KEGG_GLYCOSYLPHOSPHATIDYLINOSITOL_GPI_ANCHOR_BIOSYNTHESIS
Proportion of pathway genes=0.85; Adjusted p−value=0
Expression threshold
Below
Above
Average gene set expression
3
2
1
0
−1

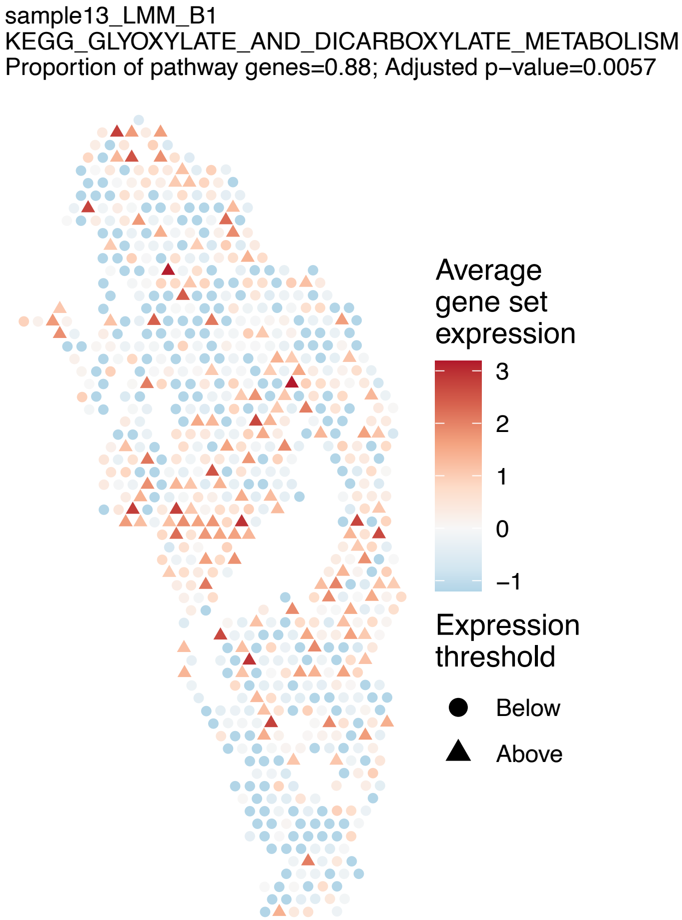
sample13_LMM_B1
KEGG_GLYOXYLATE_AND_DICARBOXYLATE_METABOLISM
Proportion of pathway genes=0.88; Adjusted p−value=0.0057
Average gene set expression
3
2
1
0
−1
Expression threshold
Below
Above

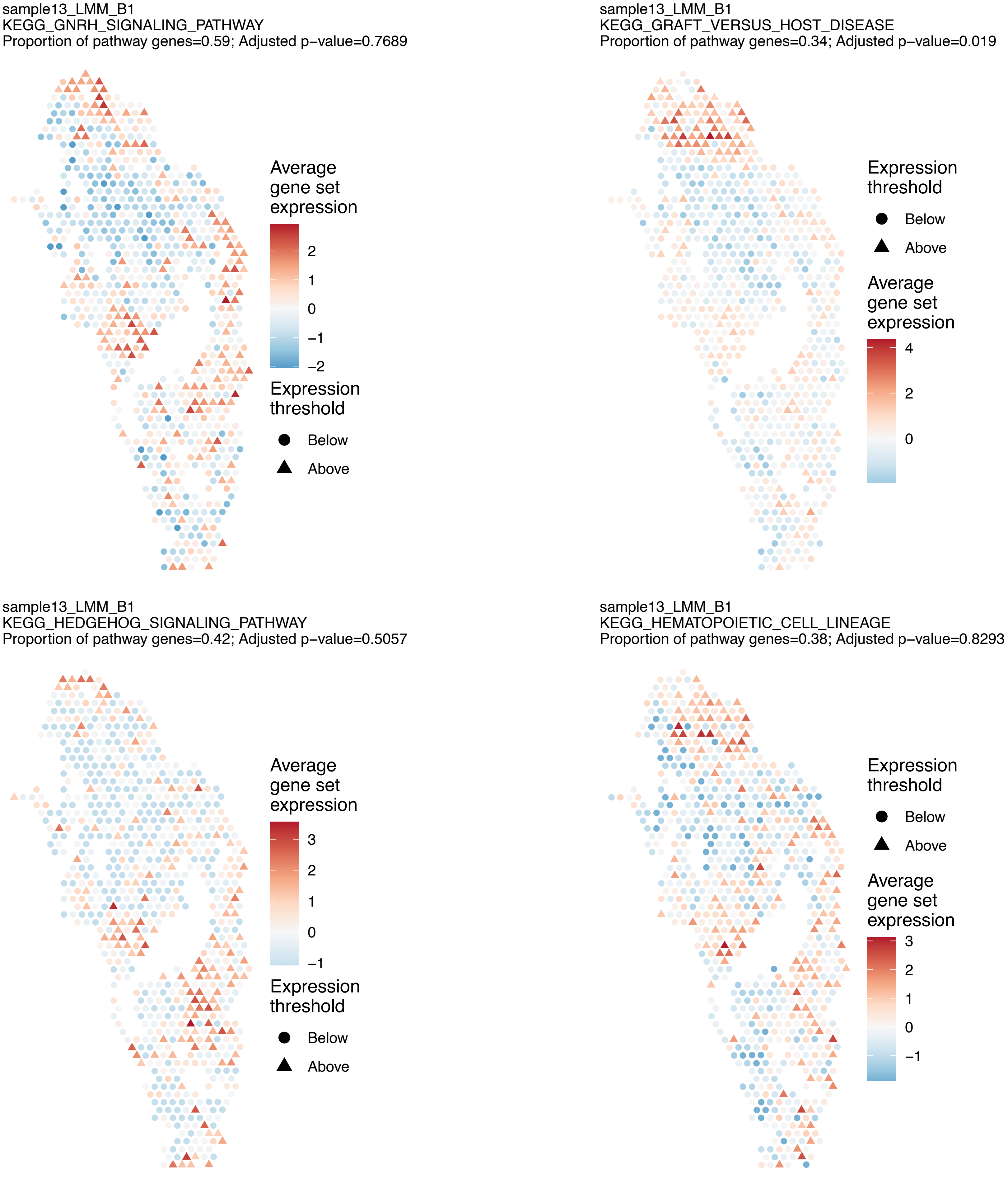
sample13_LMM_B1
KEGG_GNRH_SIGNALING_PATHWAY
Proportion of pathway genes=0.59; Adjusted p−value=0.7689
Average gene set expression
2
1
0
−1
−2
Expression threshold
Below
Above
sample13_LMM_B1
KEGG_GRAFT_VERSUS_HOST_DISEASE
Proportion of pathway genes=0.34; Adjusted p−value=0.019
Expression threshold
Below
Above
Average gene set expression
4
2
0
sample13_LMM_B1
KEGG_HEDGEHOG_SIGNALING_PATHWAY
Proportion of pathway genes=0.42; Adjusted p−value=0.5057
Average gene set expression
3
2
1
0
−1
Expression threshold
Below
Above
sample13_LMM_B1
KEGG_HEMATOPOIETIC_CELL_LINEAGE
Proportion of pathway genes=0.38; Adjusted p−value=0.8293
Expression threshold
Below
Above
Average gene set expression
3
2
1
0
−1

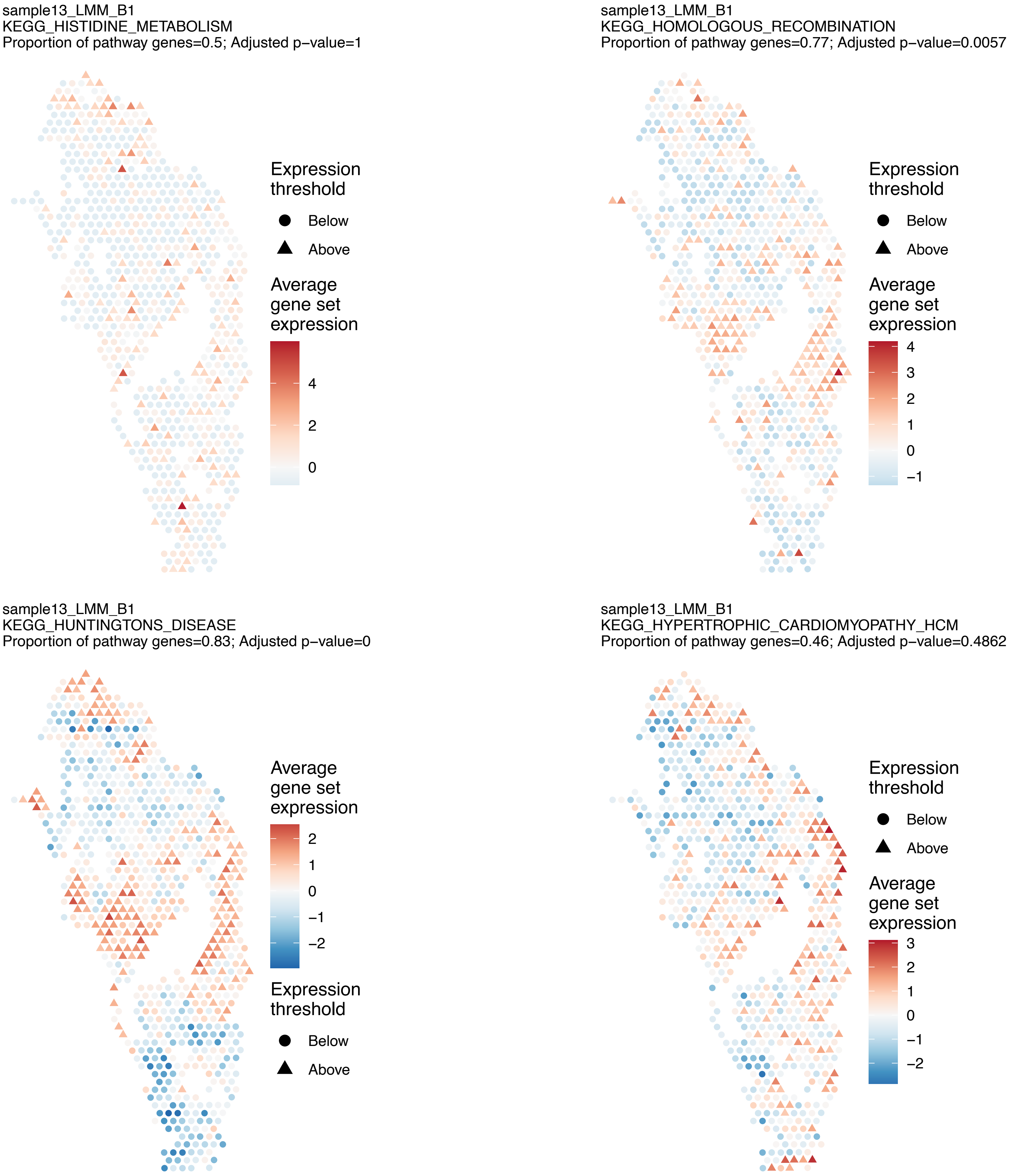
sample13_LMM_B1
KEGG_HISTIDINE_METABOLISM
Proportion of pathway genes=0.5; Adjusted p−value=1
Expression threshold
Below
Above
Average gene set expression
4
2
0
sample13_LMM_B1
KEGG_HOMOLOGOUS_RECOMBINATION
Proportion of pathway genes=0.77; Adjusted p−value=0.0057
Expression threshold
Below
Above
Average gene set expression
4
3
2
1
0
−1
sample13_LMM_B1
KEGG_HUNTINGTONS_DISEASE
Proportion of pathway genes=0.83; Adjusted p−value=0
Average gene set expression
2
1
0
−1
−2
Expression threshold
Below
Above
sample13_LMM_B1
KEGG_HYPERTROPHIC_CARDIOMYOPATHY_HCM
Proportion of pathway genes=0.46; Adjusted p−value=0.4862
Expression threshold
Below
Above
Average gene set expression
3
2
1
0
−1
−2

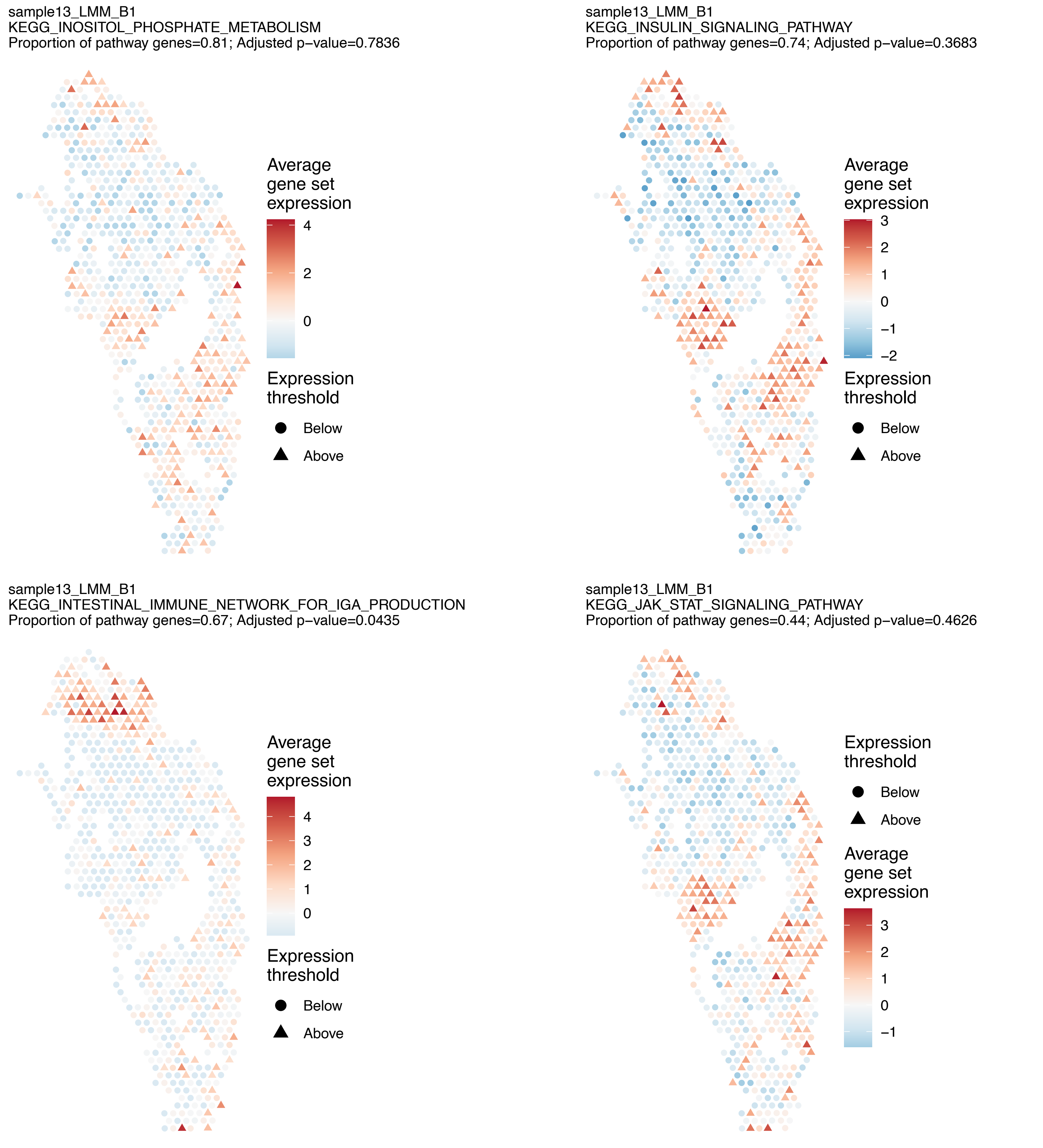
sample13_LMM_B1
KEGG_INOSITOL_PHOSPHATE_METABOLISM
Proportion of pathway genes=0.81; Adjusted p−value=0.7836
Average gene set expression
4
2
0
Expression threshold
Below
Above
sample13_LMM_B1
KEGG_INSULIN_SIGNALING_PATHWAY
Proportion of pathway genes=0.74; Adjusted p−value=0.3683
Average gene set expression
3
2
1
0
−1
−2
Expression threshold
Below
Above
sample13_LMM_B1
KEGG_INTESTINAL_IMMUNE_NETWORK_FOR_IGA_PRODUCTION
Proportion of pathway genes=0.67; Adjusted p−value=0.0435
Average gene set expression
4
3
2
1
0
Expression threshold
Below
Above
sample13_LMM_B1
KEGG_JAK_STAT_SIGNALING_PATHWAY
Proportion of pathway genes=0.44; Adjusted p−value=0.4626
Expression threshold
Below
Above
Average gene set expression
3
2
1
0
−1

sample13_LMM_B1
KEGG_LEISHMANIA_INFECTION
Proportion of pathway genes=0.7; Adjusted p−value=0

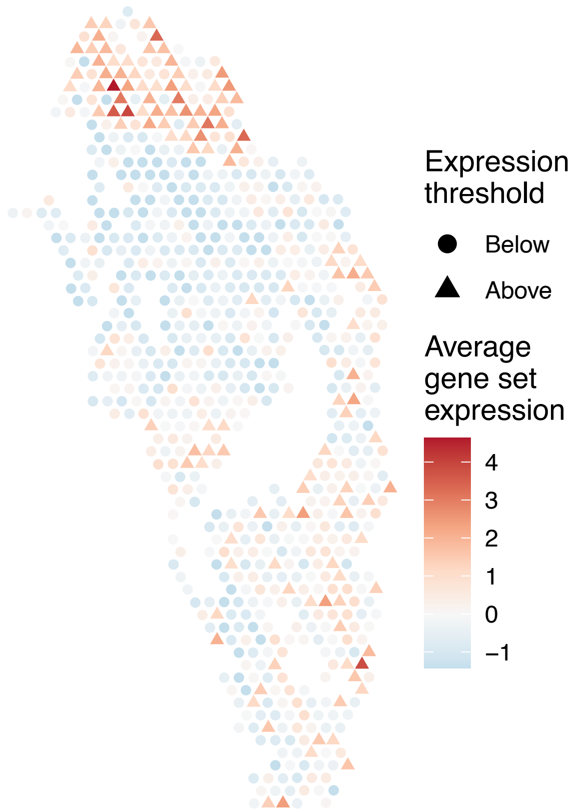


sample13_LMM_B1
KEGG_LEUKOCYTE_TRANSENDOTHELIAL_MIGRATION
Proportion of pathway genes=0.64; Adjusted p−value=0.8307

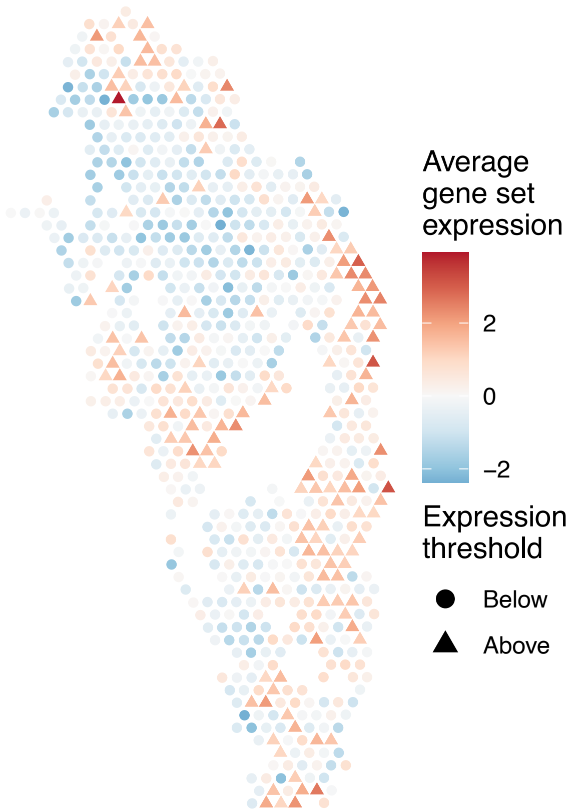


sample13_LMM_B1
KEGG_LIMONENE_AND_PINENE_DEGRADATION
Proportion of pathway genes=0.9; Adjusted p−value=NA

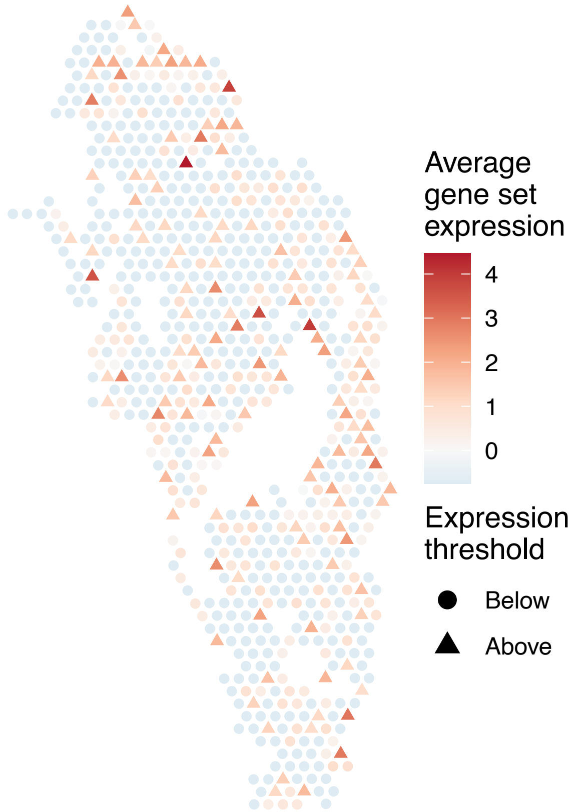


sample13_LMM_B1
KEGG_LONG_TERM_DEPRESSION
Proportion of pathway genes=0.5; Adjusted p−value=0.6592

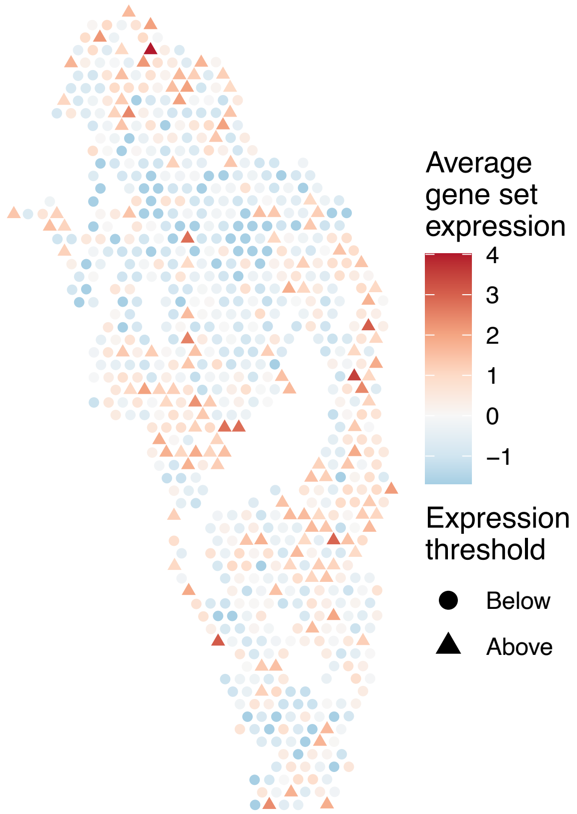

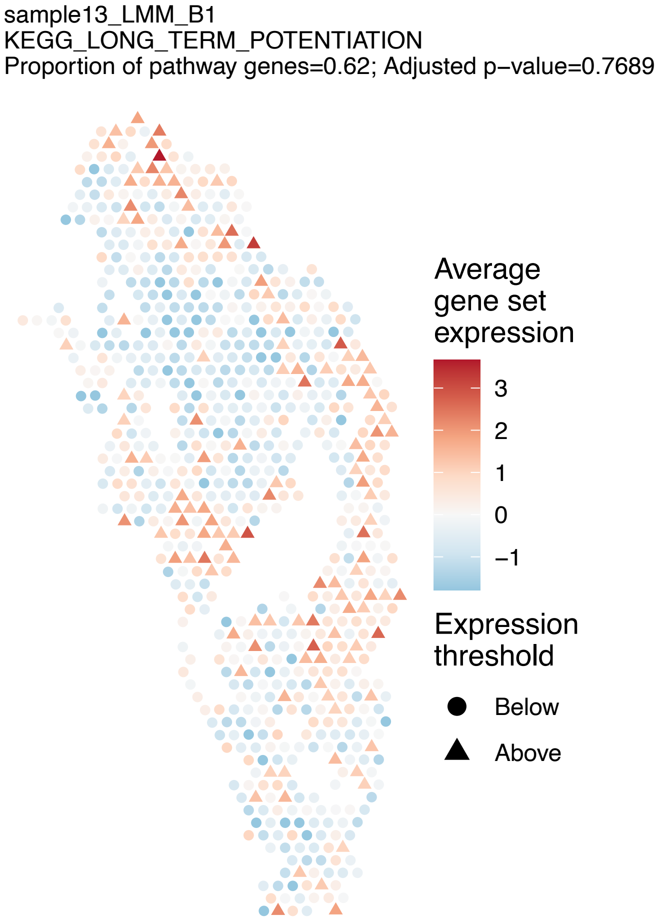
sample13_LMM_B1
KEGG_LONG_TERM_POTENTIATION
Proportion of pathway genes=0.62; Adjusted p−value=0.7689
Average gene set expression
3
2
1
0
−1
Expression threshold
Below
Above

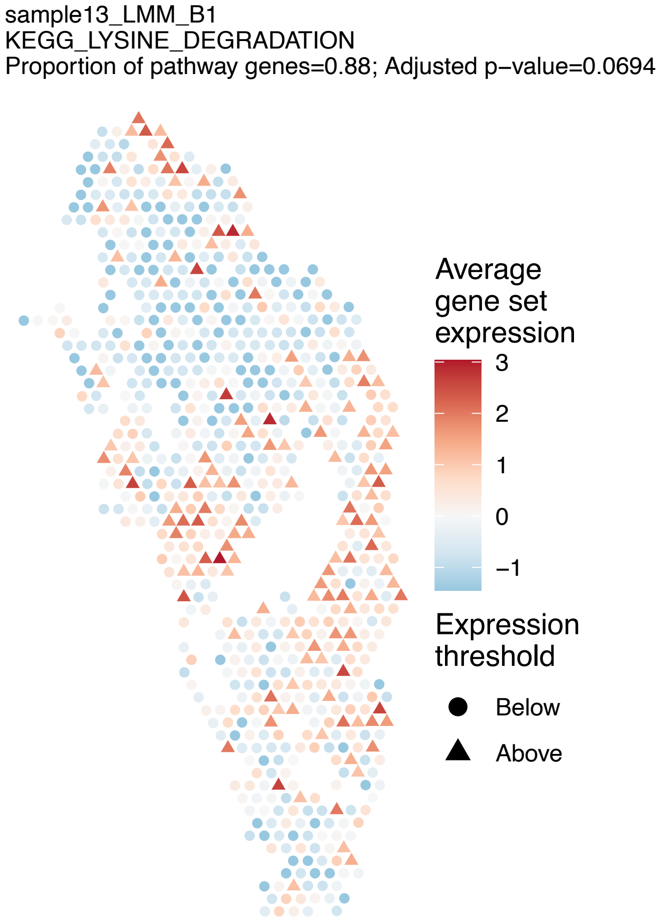
sample13_LMM_B1
KEGG_LYSINE_DEGRADATION
Proportion of pathway genes=0.88; Adjusted p−value=0.0694
Average gene set expression
3
2
1
0
−1
Expression threshold
Below
Above

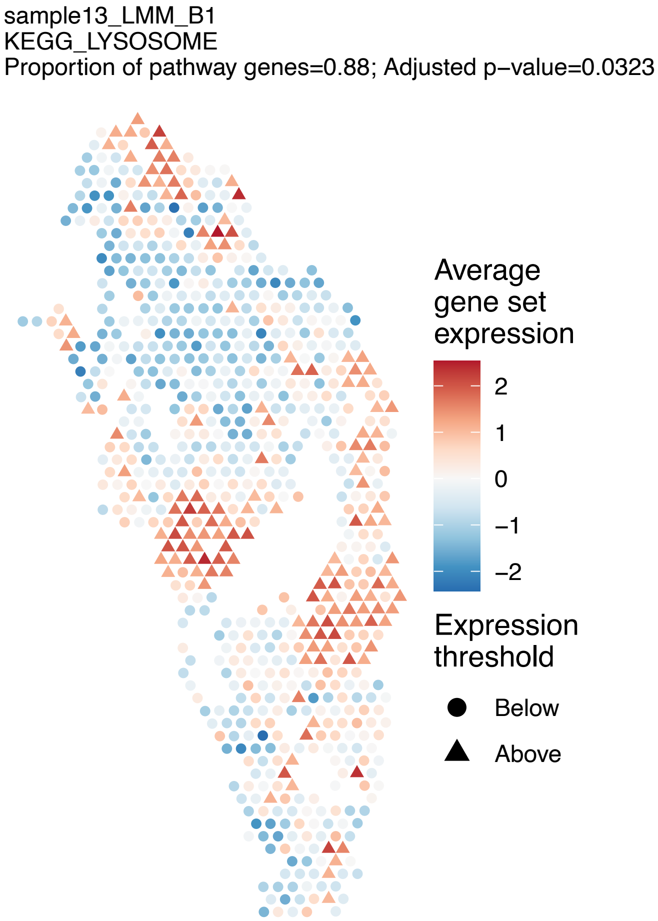
sample13_LMM_B1
KEGG_LYSOSOME
Proportion of pathway genes=0.88; Adjusted p−value=0.0323
Average gene set expression
2
1
0
−1
−2
Expression threshold
Below
Above

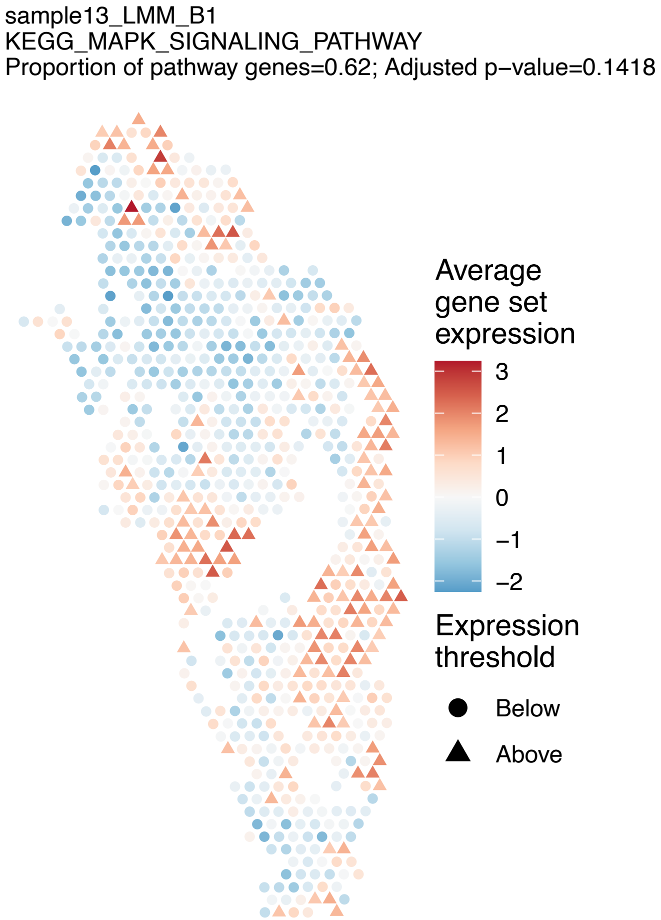
sample13_LMM_B1
KEGG_MAPK_SIGNALING_PATHWAY
Proportion of pathway genes=0.62; Adjusted p−value=0.1418
Average gene set expression
3
2
1
0
−1
−2
Expression threshold
Below
Above

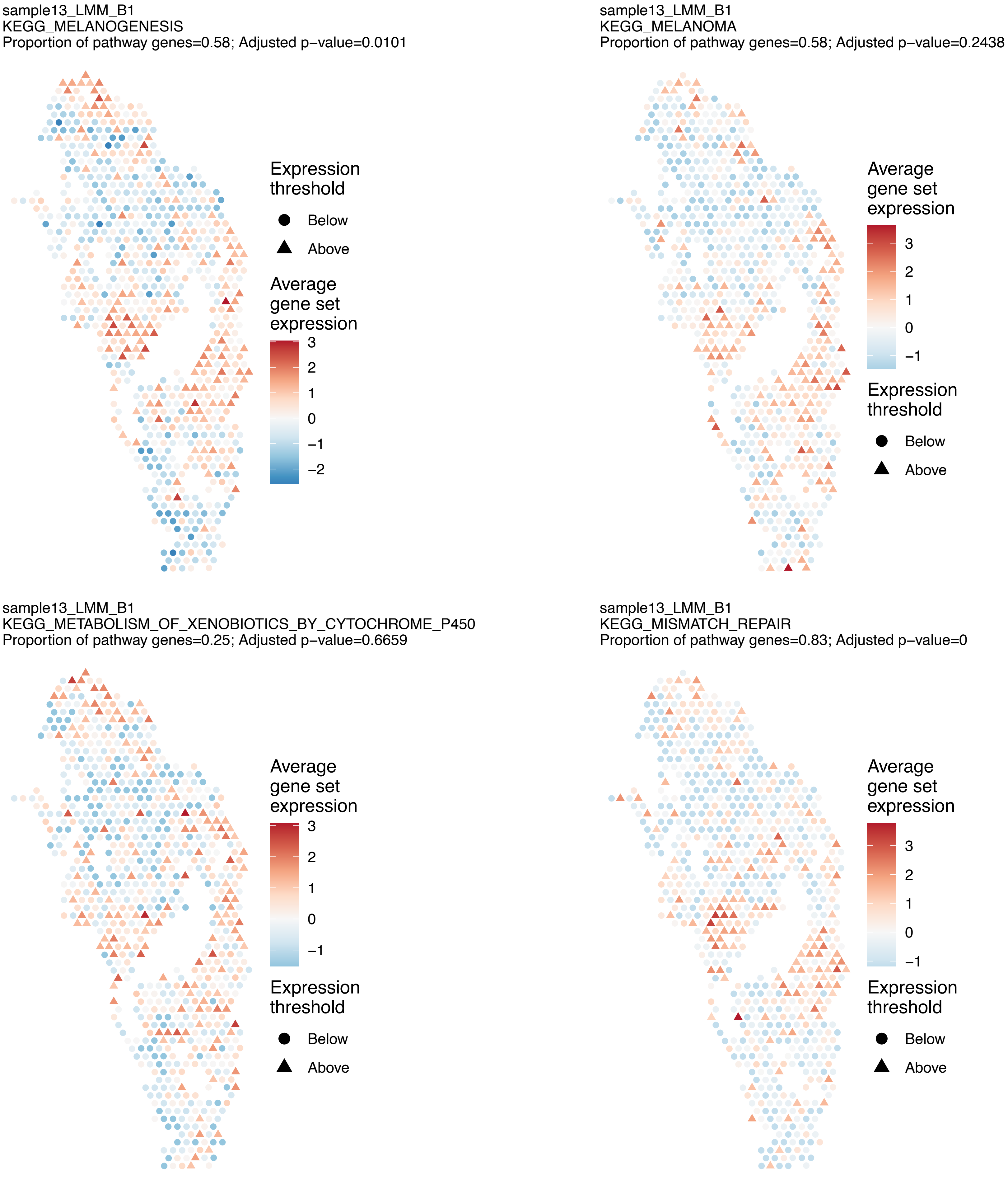
sample13_LMM_B1
KEGG_MELANOGENESIS
Proportion of pathway genes=0.58; Adjusted p−value=0.0101
Expression threshold
Below
Above
Average gene set expression
3
2
1
0
−1
−2
sample13_LMM_B1
KEGG_MELANOMA
Proportion of pathway genes=0.58; Adjusted p−value=0.2438
Average gene set expression
3
2
1
0
−1
Expression threshold
Below
Above
sample13_LMM_B1
KEGG_METABOLISM_OF_XENOBIOTICS_BY_CYTOCHROME_P450
Proportion of pathway genes=0.25; Adjusted p−value=0.6659
Average gene set expression
3
2
1
0
−1
Expression threshold
Below
Above
sample13_LMM_B1
KEGG_MISMATCH_REPAIR
Proportion of pathway genes=0.83; Adjusted p−value=0
Average gene set expression
3
2
1
0
−1
Expression threshold
Below
Above

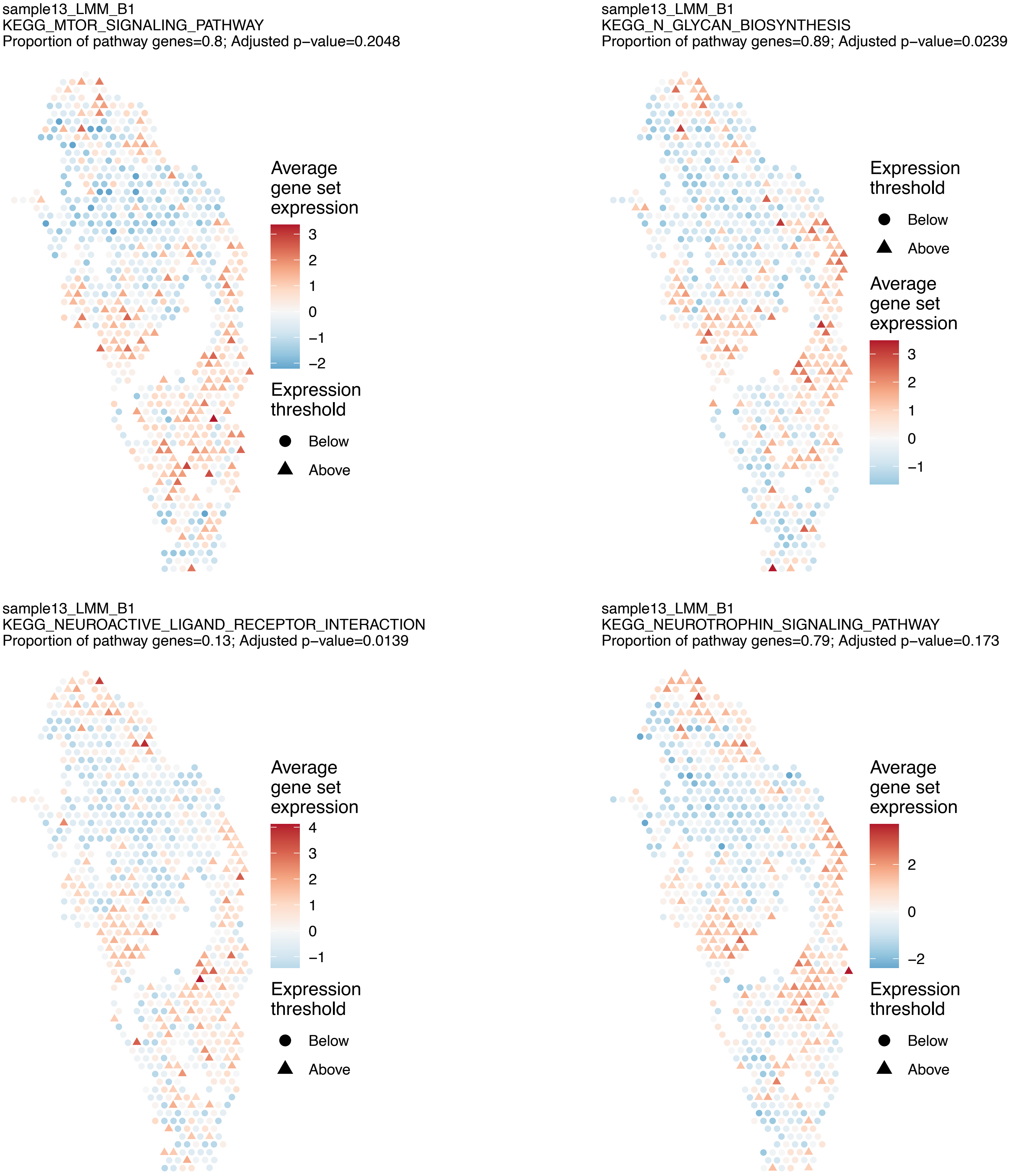
sample13_LMM_B1
KEGG_MTOR_SIGNALING_PATHWAY
Proportion of pathway genes=0.8; Adjusted p−value=0.2048
Average gene set expression
3
2
1
0
−1
−2
Expression threshold
Below
Above
sample13_LMM_B1
KEGG_N_GLYCAN_BIOSYNTHESIS
Proportion of pathway genes=0.89; Adjusted p−value=0.0239
Expression threshold
Below
Above
Average gene set expression
3
2
1
0
−1
sample13_LMM_B1
KEGG_NEUROACTIVE_LIGAND_RECEPTOR_INTERACTION
Proportion of pathway genes=0.13; Adjusted p−value=0.0139
Average gene set expression
4
3
2
1
0
−1
Expression threshold
Below
Above
sample13_LMM_B1
KEGG_NEUROTROPHIN_SIGNALING_PATHWAY
Proportion of pathway genes=0.79; Adjusted p−value=0.173
Average gene set expression
2
0
−2
Expression threshold
Below
Above

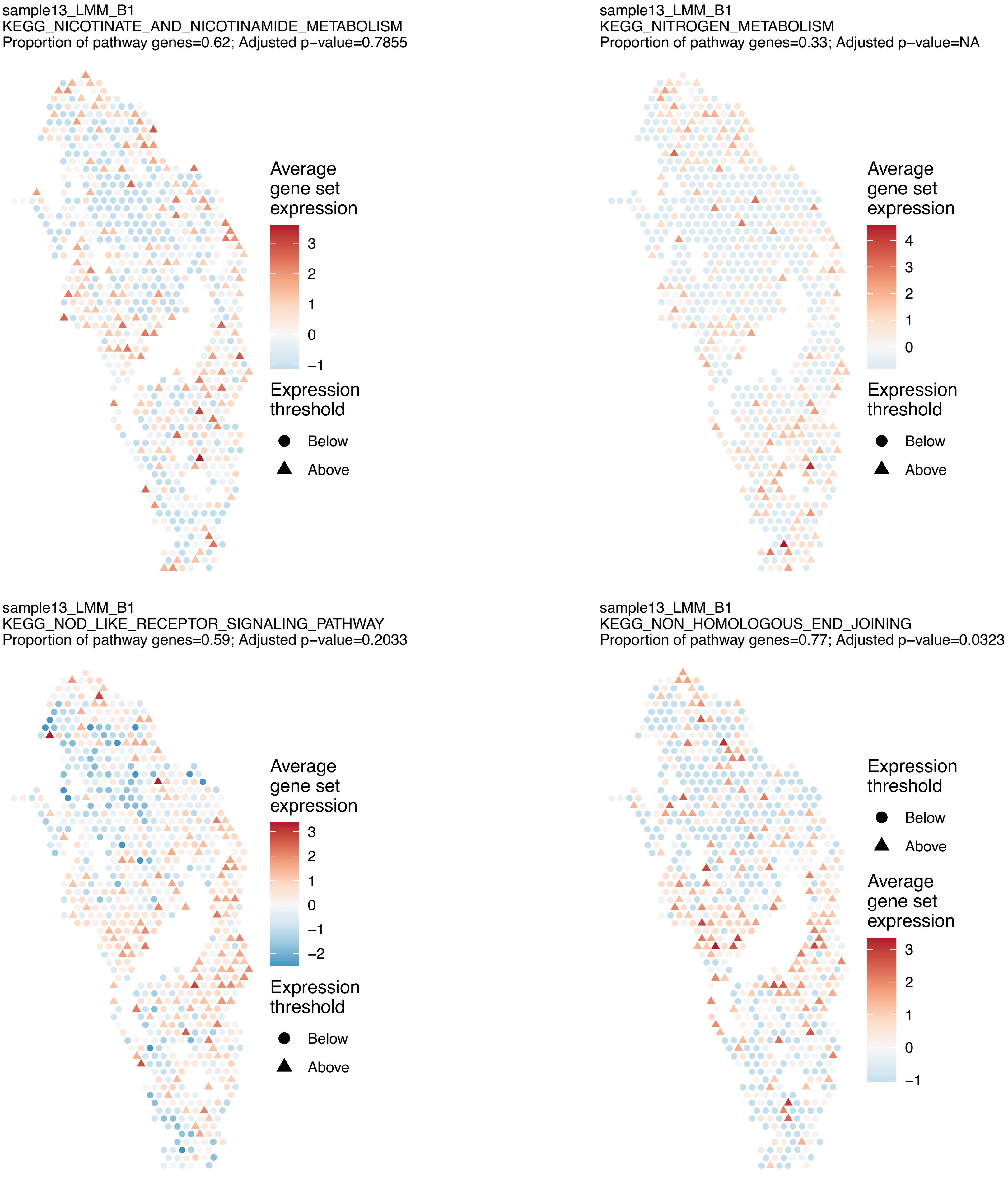

sample13_LMM_B1
KEGG_NICOTINATE_AND_NICOTINAMIDE_METABOLISM
Proportion of pathway genes=0.62; Adjusted p−value=0.7855
Average gene set expression
3
2
1
0
−1
Expression threshold
Below
Above
sample13_LMM_B1
KEGG_NITROGEN_METABOLISM
Proportion of pathway genes=0.33; Adjusted p−value=NA
Average gene set expression
4
3
2
1
0
Expression threshold
Below
Above
sample13_LMM_B1
KEGG_NOD_LIKE_RECEPTOR_SIGNALING_PATHWAY
Proportion of pathway genes=0.59; Adjusted p−value=0.2033
Average gene set expression
3
2
1
0
−1
−2
Expression threshold
Below
Above
sample13_LMM_B1
KEGG_NON_HOMOLOGOUS_END_JOINING
Proportion of pathway genes=0.77; Adjusted p−value=0.0323
Expression threshold
Below
Above
Average gene set expression
3
2
1
0
−1

sample13_LMM_B1
KEGG_NON_SMALL_CELL_LUNG_CANCER
Proportion of pathway genes=0.77; Adjusted p−value=0.1147

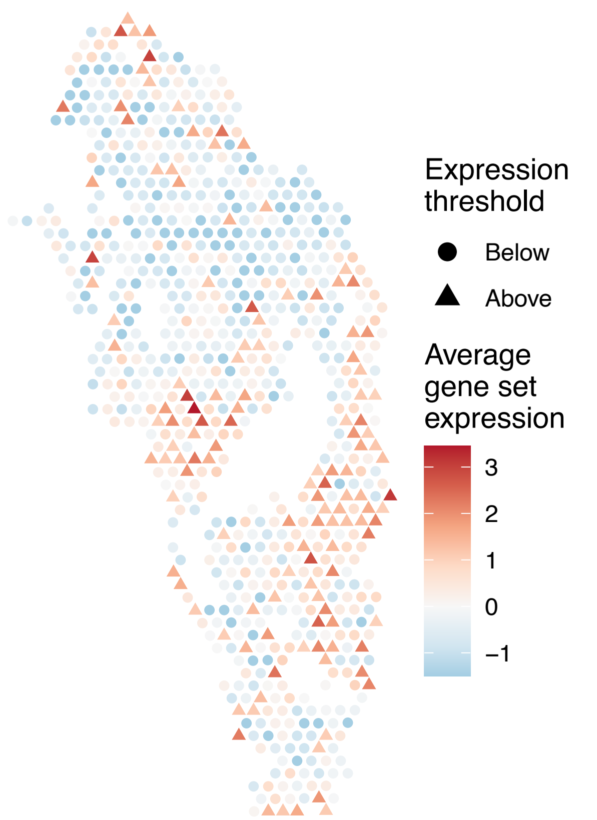


sample13_LMM_B1
KEGG_NOTCH_SIGNALING_PATHWAY
Proportion of pathway genes=0.84; Adjusted p−value=0.0057

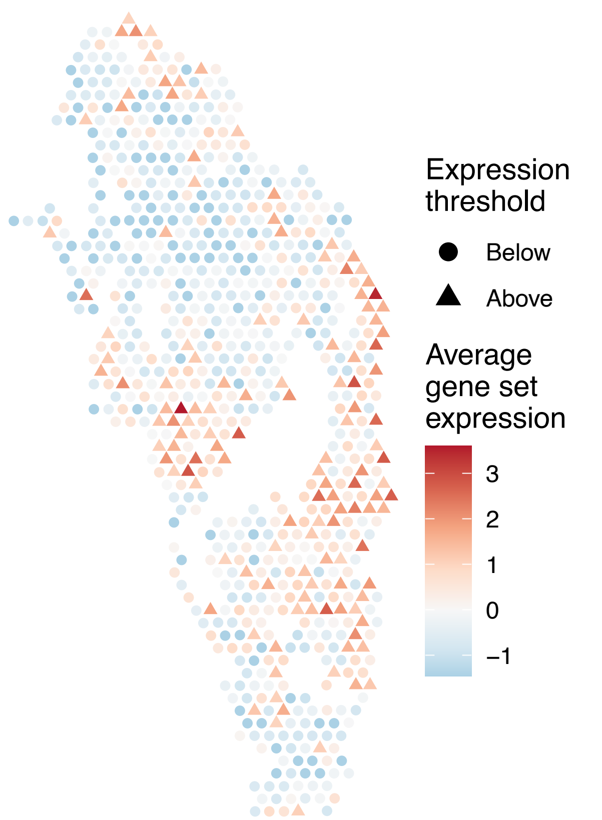


sample13_LMM_B1
KEGG_NUCLEOTIDE_EXCISION_REPAIR
Proportion of pathway genes=0.83; Adjusted p−value=0.0428

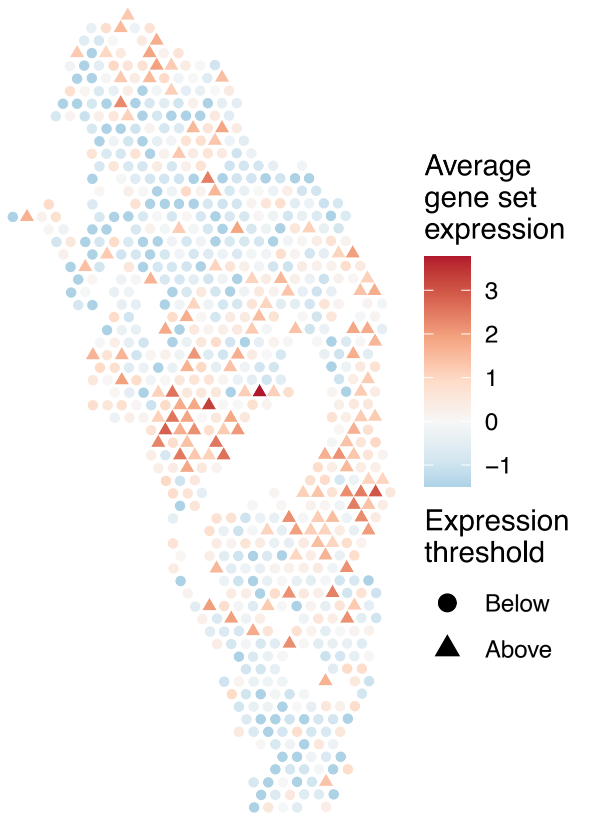


sample13_LMM_B1
KEGG_O_GLYCAN_BIOSYNTHESIS
Proportion of pathway genes=0.45; Adjusted p−value=0.9538

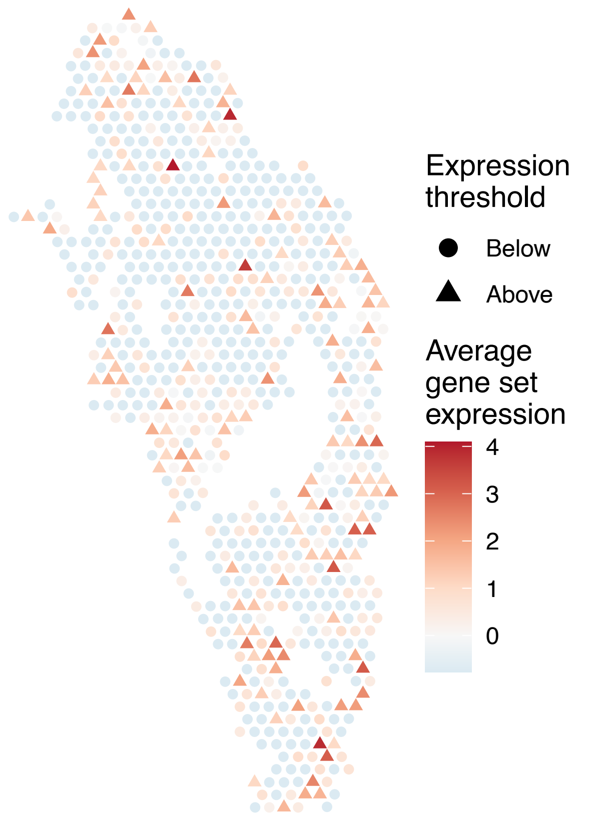

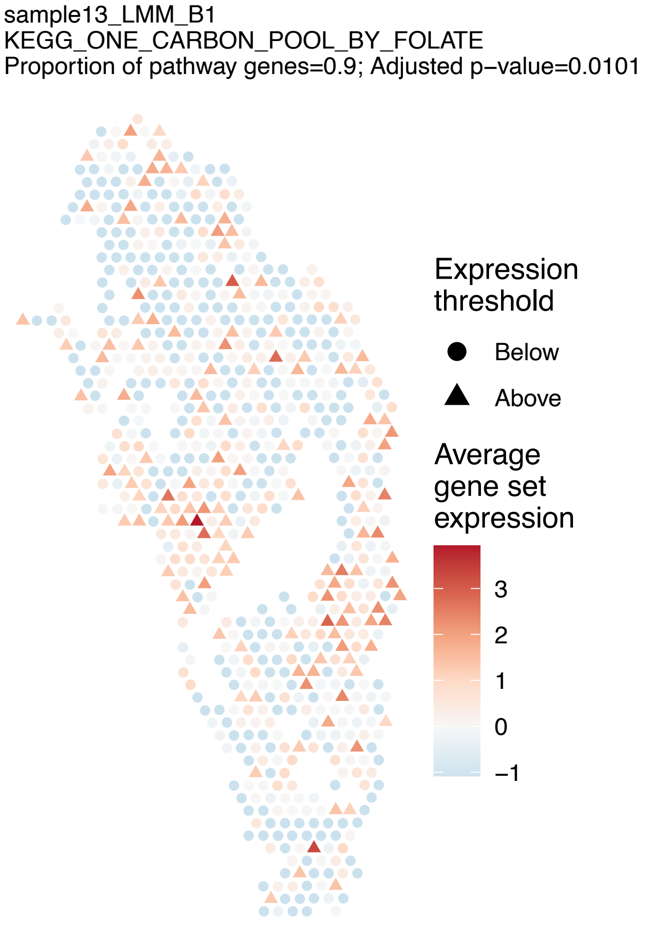
sample13_LMM_B1
KEGG_ONE_CARBON_POOL_BY_FOLATE
Proportion of pathway genes=0.9; Adjusted p−value=0.0101
Expression threshold
Below
Above
Average gene set expression
3
2
1
0
−1

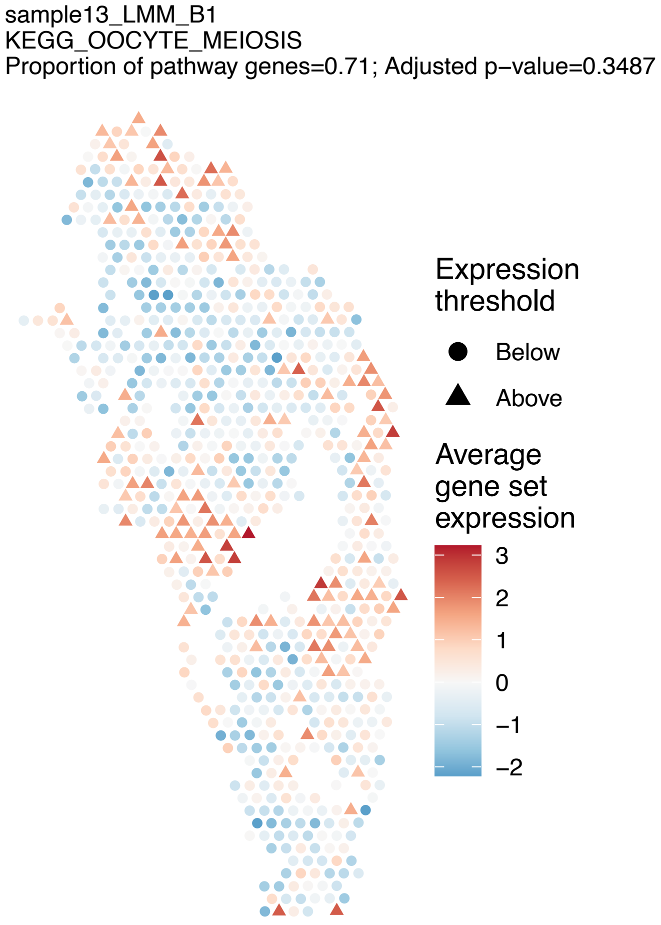
sample13_LMM_B1
KEGG_OOCYTE_MEIOSIS
Proportion of pathway genes=0.71; Adjusted p−value=0.3487
Expression threshold
Below
Above
Average gene set expression
3
2
1
0
−1
−2

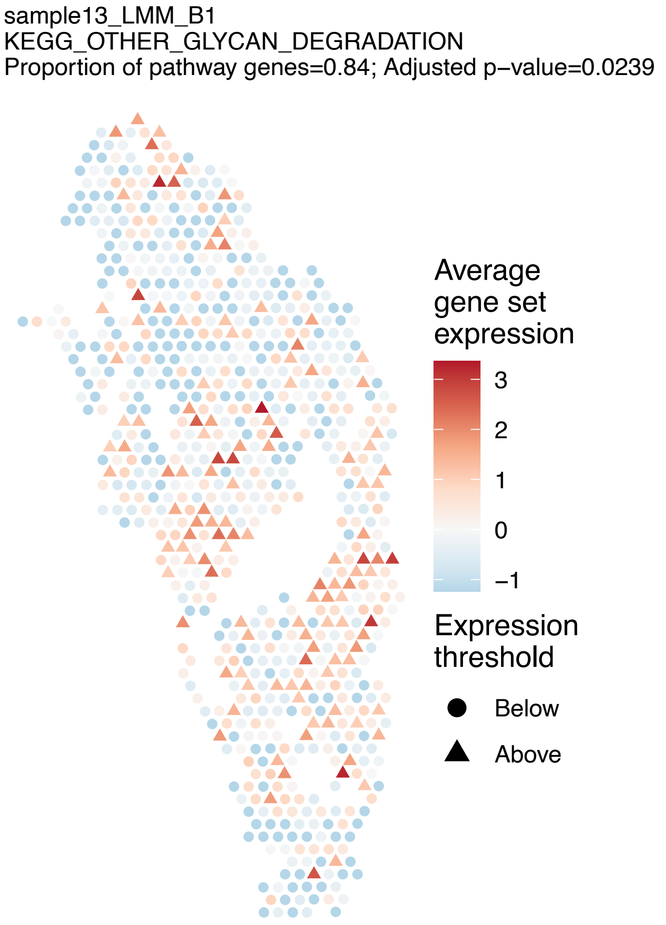
sample13_LMM_B1
KEGG_OTHER_GLYCAN_DEGRADATION
Proportion of pathway genes=0.84; Adjusted p−value=0.0239
Average gene set expression
3
2
1
0
−1
Expression threshold
Below
Above

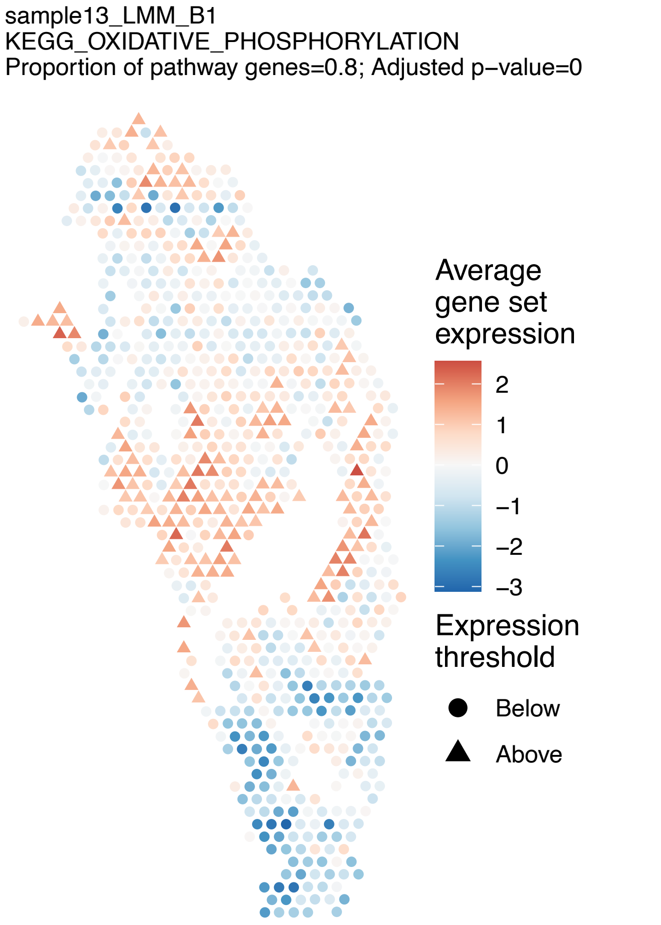
sample13_LMM_B1
KEGG_OXIDATIVE_PHOSPHORYLATION
Proportion of pathway genes=0.8; Adjusted p−value=0
Average gene set expression
2
1
0
−1
−2
−3
Expression threshold
Below
Above

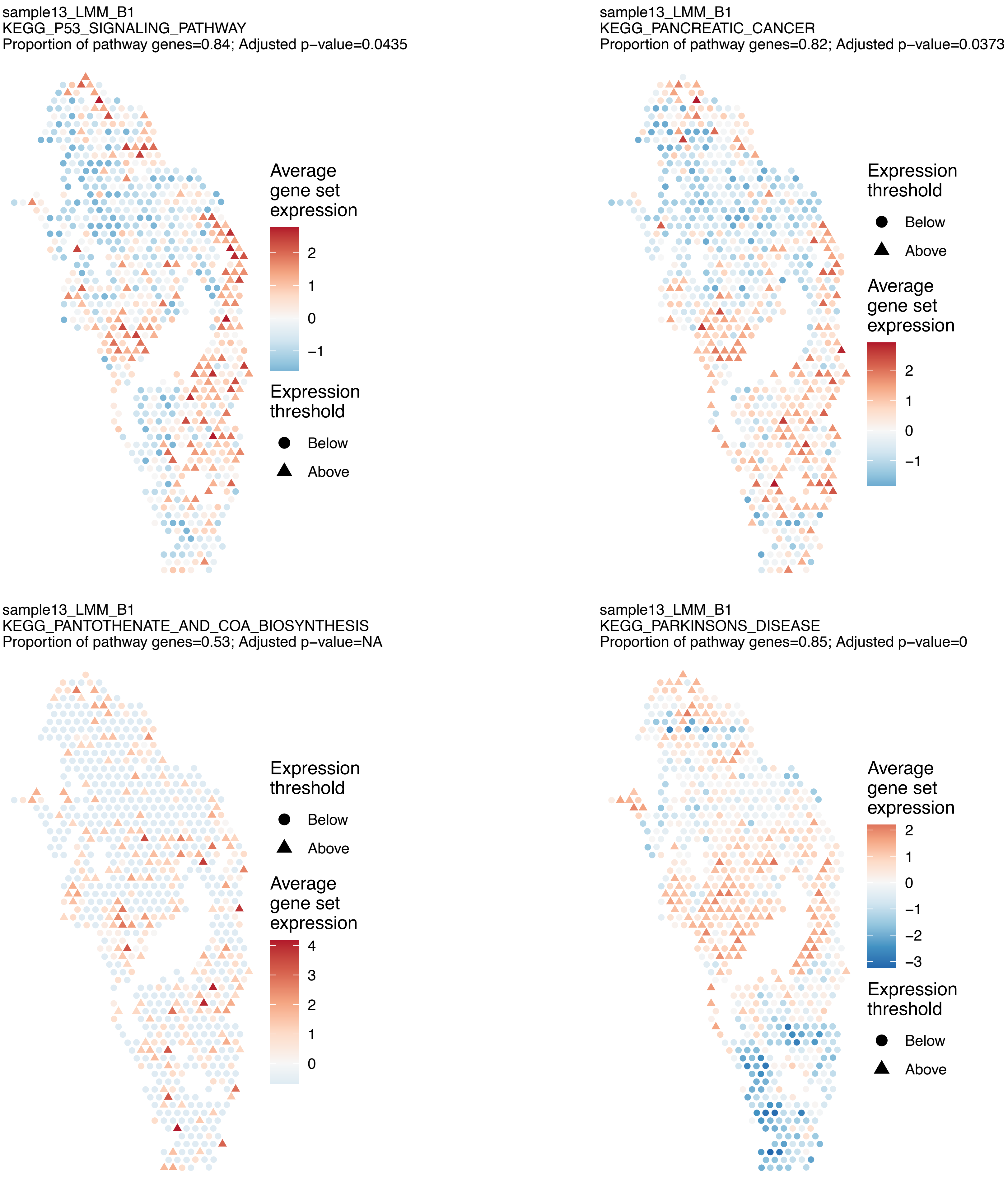

sample13_LMM_B1
KEGG_P53_SIGNALING_PATHWAY
Proportion of pathway genes=0.84; Adjusted p−value=0.0435
Average gene set expression
2
1
0
−1
Expression threshold
Below
Above
sample13_LMM_B1
KEGG_PANCREATIC_CANCER
Proportion of pathway genes=0.82; Adjusted p−value=0.0373
Expression threshold
Below
Above
Average gene set expression
2
1
0
−1
sample13_LMM_B1
KEGG_PANTOTHENATE_AND_COA_BIOSYNTHESIS
Proportion of pathway genes=0.53; Adjusted p−value=NA
Expression threshold
Below
Above
Average gene set expression
4
3
2
1
0
sample13_LMM_B1
KEGG_PARKINSONS_DISEASE
Proportion of pathway genes=0.85; Adjusted p−value=0
Average gene set expression
2
1
0
−1
−2
−3
Expression threshold
Below
Above

sample13_LMM_B1
KEGG_PATHOGENIC_ESCHERICHIA_COLI_INFECTION
Proportion of pathway genes=0.8; Adjusted p−value=0.1463

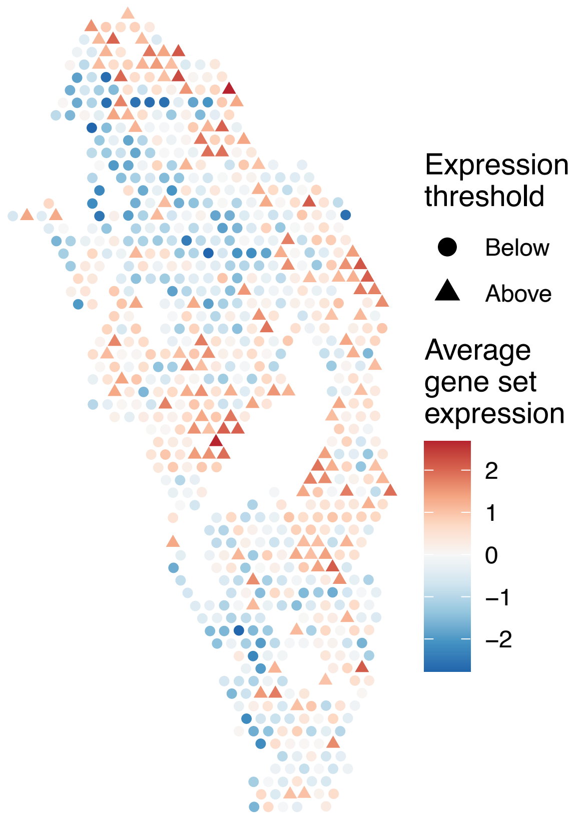


sample13_LMM_B1
KEGG_PATHWAYS_IN_CANCER
Proportion of pathway genes=0.65; Adjusted p−value=0.0268

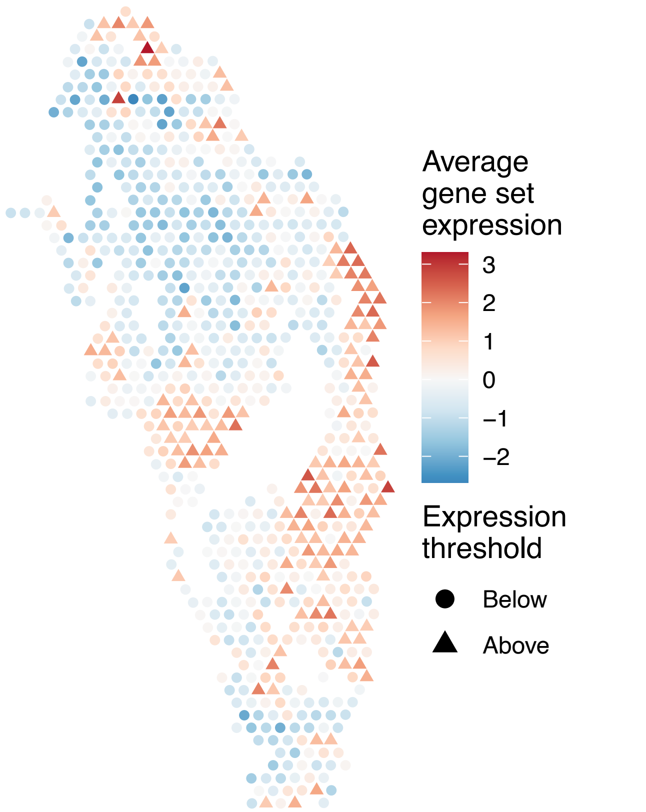


sample13_LMM_B1
KEGG_PENTOSE_PHOSPHATE_PATHWAY
Proportion of pathway genes=0.71; Adjusted p−value=0.3872

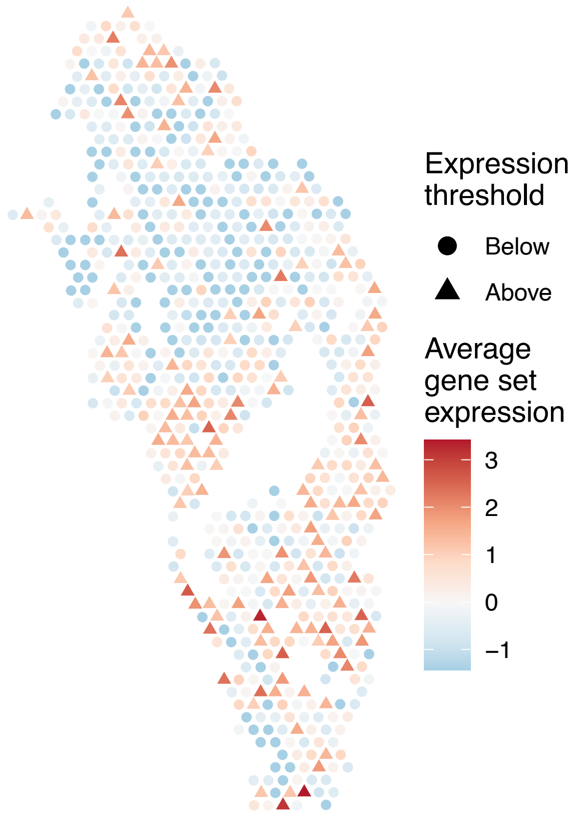


sample13_LMM_B1
KEGG_PEROXISOME
Proportion of pathway genes=0.69; Adjusted p−value=0.019

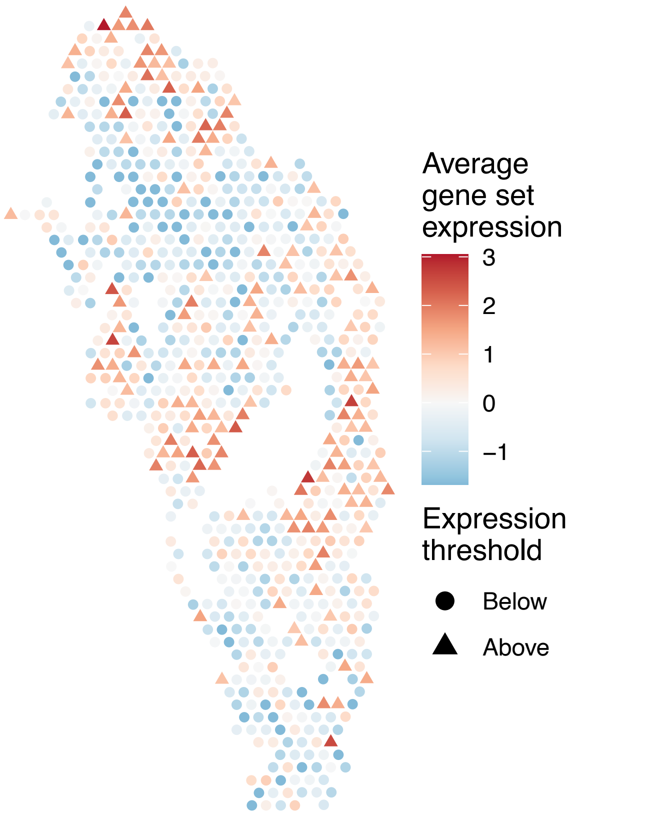

sample13_LMM_B1
KEGG_PHENYLALANINE_METABOLISM
Proportion of pathway genes=0.58; Adjusted p−value=1

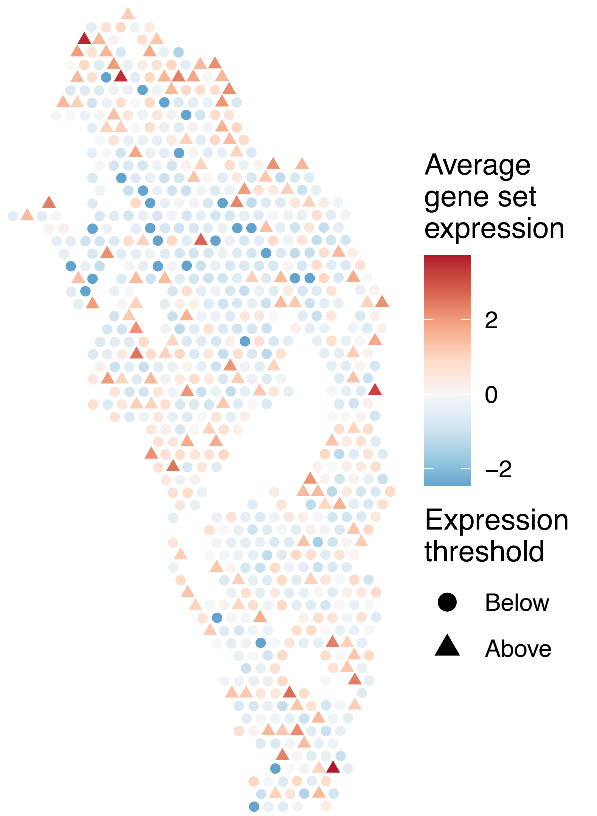


sample13_LMM_B1
KEGG_PHOSPHATIDYLINOSITOL_SIGNALING_SYSTEM
Proportion of pathway genes=0.74; Adjusted p−value=0.4022

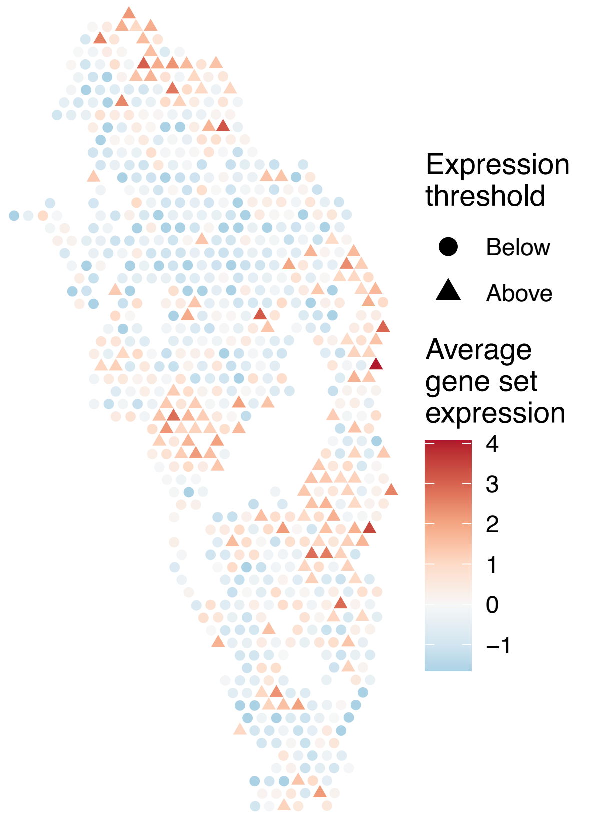


sample13_LMM_B1
KEGG_PORPHYRIN_AND_CHLOROPHYLL_METABOLISM
Proportion of pathway genes=0.47; Adjusted p−value=1

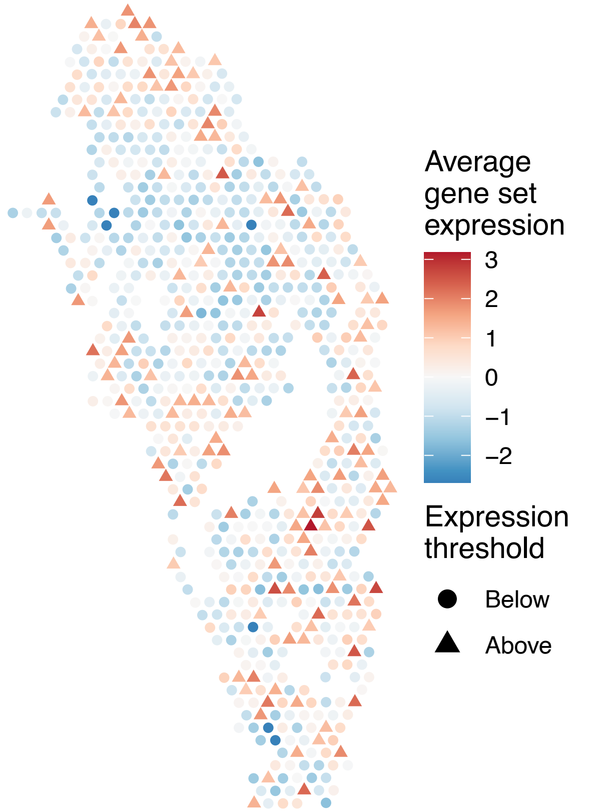


sample13_LMM_B1
KEGG_PPAR_SIGNALING_PATHWAY
Proportion of pathway genes=0.59; Adjusted p−value=0.2033

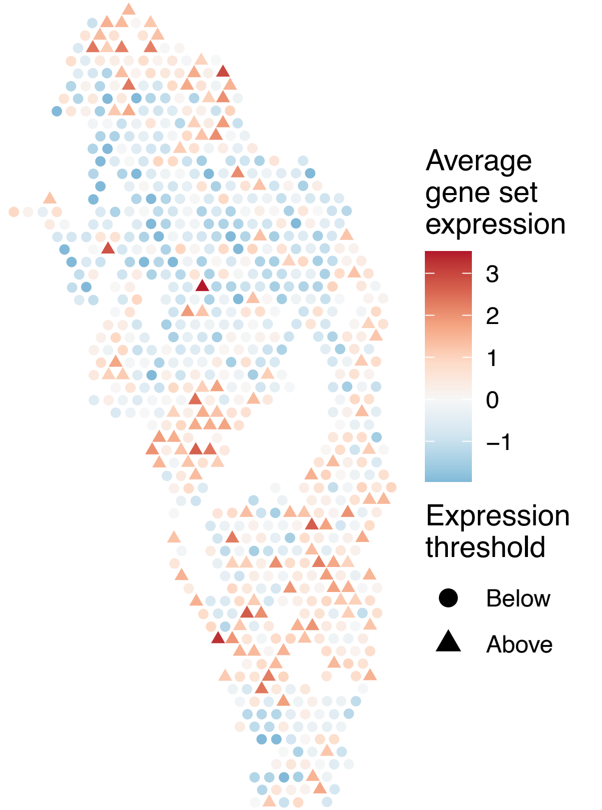

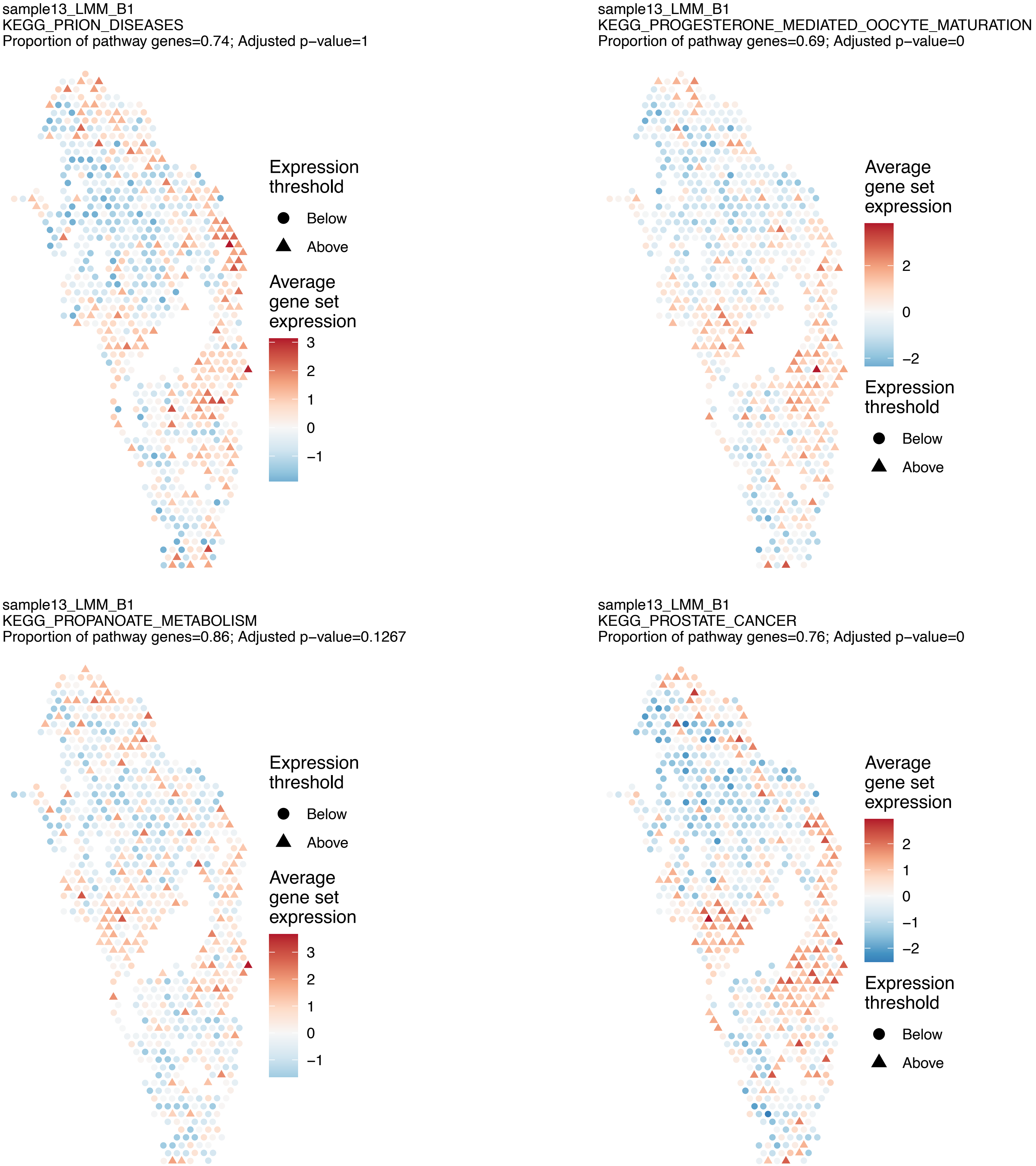
sample13_LMM_B1
KEGG_PRION_DISEASES
Proportion of pathway genes=0.74; Adjusted p−value=1
Expression threshold
Below
Above
Average gene set expression
3
2
1
0
−1
sample13_LMM_B1
KEGG_PROGESTERONE_MEDIATED_OOCYTE_MATURATION
Proportion of pathway genes=0.69; Adjusted p−value=0
Average gene set expression
2
0
−2
Expression threshold
Below
Above
sample13_LMM_B1
KEGG_PROPANOATE_METABOLISM
Proportion of pathway genes=0.86; Adjusted p−value=0.1267
Expression threshold
Below
Above
Average gene set expression
3
2
1
0
−1
sample13_LMM_B1
KEGG_PROSTATE_CANCER
Proportion of pathway genes=0.76; Adjusted p−value=0
Average gene set expression
2
1
0
−1
−2
Expression threshold
Below
Above

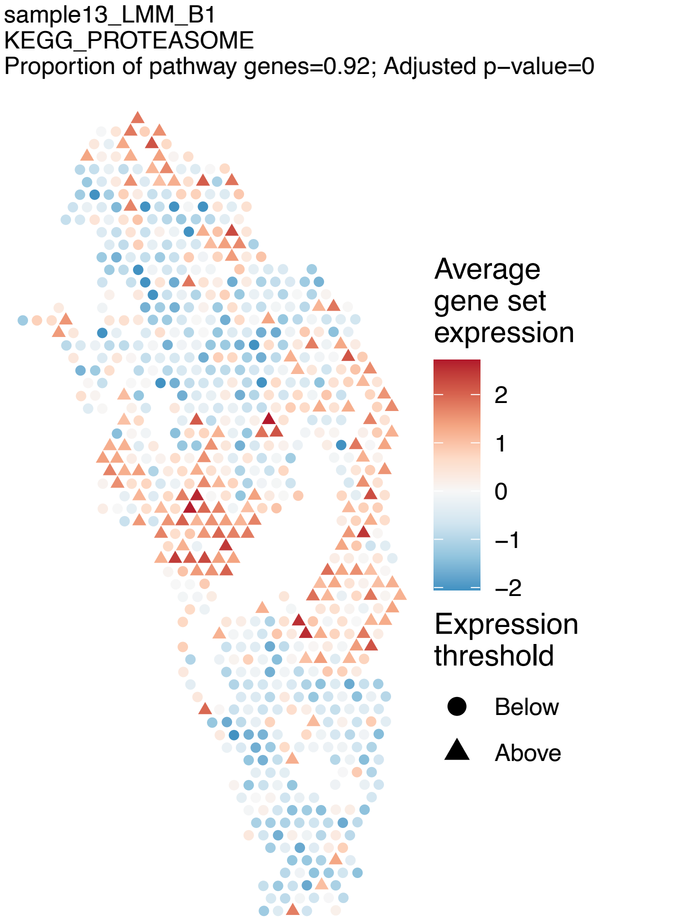
sample13_LMM_B1
KEGG_PROTEASOME
Proportion of pathway genes=0.92; Adjusted p−value=0
Average gene set expression
2
1
0
−1
−2
Expression threshold
Below
Above

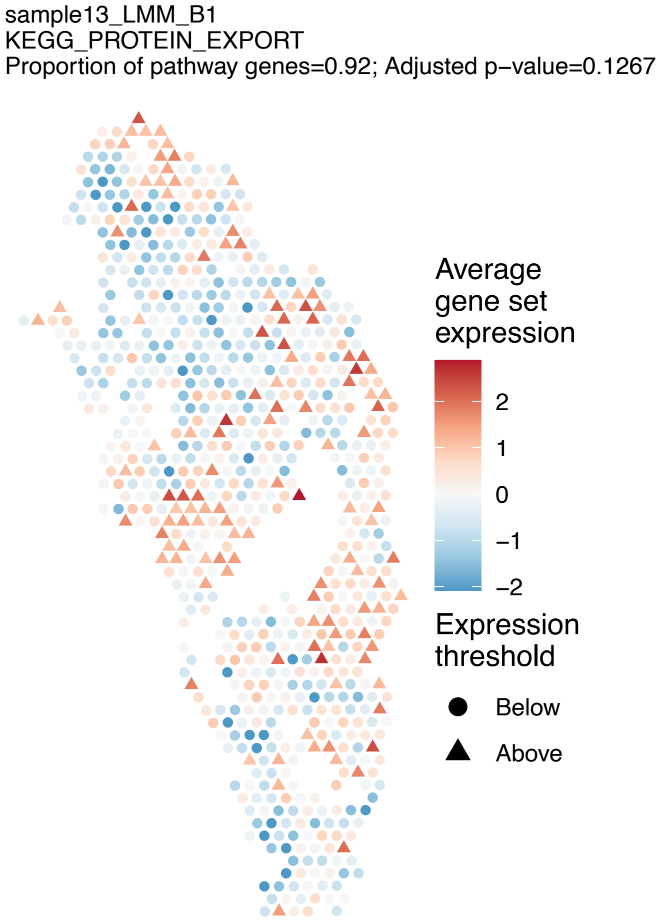
sample13_LMM_B1
KEGG_PROTEIN_EXPORT
Proportion of pathway genes=0.92; Adjusted p−value=0.1267
Average gene set expression
2
1
0
−1
−2
Expression threshold
Below
Above

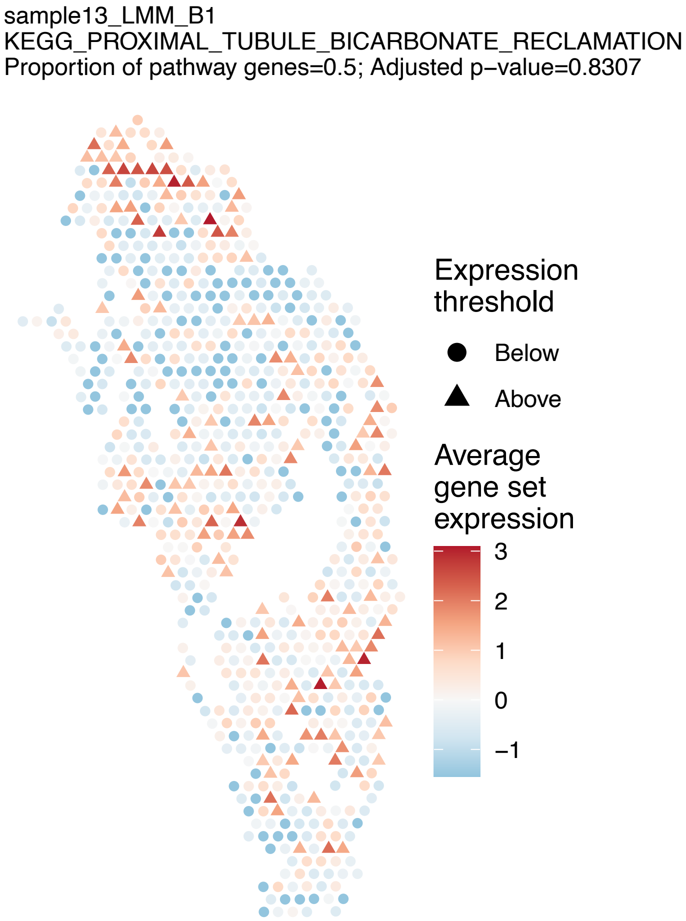
sample13_LMM_B1
KEGG_PROXIMAL_TUBULE_BICARBONATE_RECLAMATION
Proportion of pathway genes=0.5; Adjusted p−value=0.8307
Expression threshold
Below
Above
Average gene set expression
3
2
1
0
−1

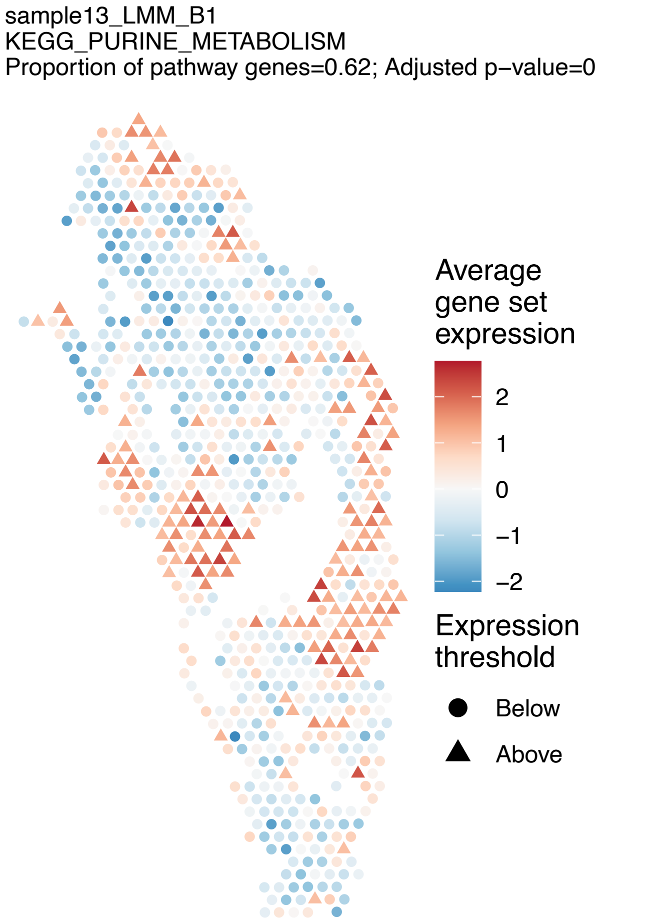
sample13_LMM_B1
KEGG_PURINE_METABOLISM
Proportion of pathway genes=0.62; Adjusted p−value=0
Average gene set expression
2
1
0
−1
−2
Expression threshold
Below
Above

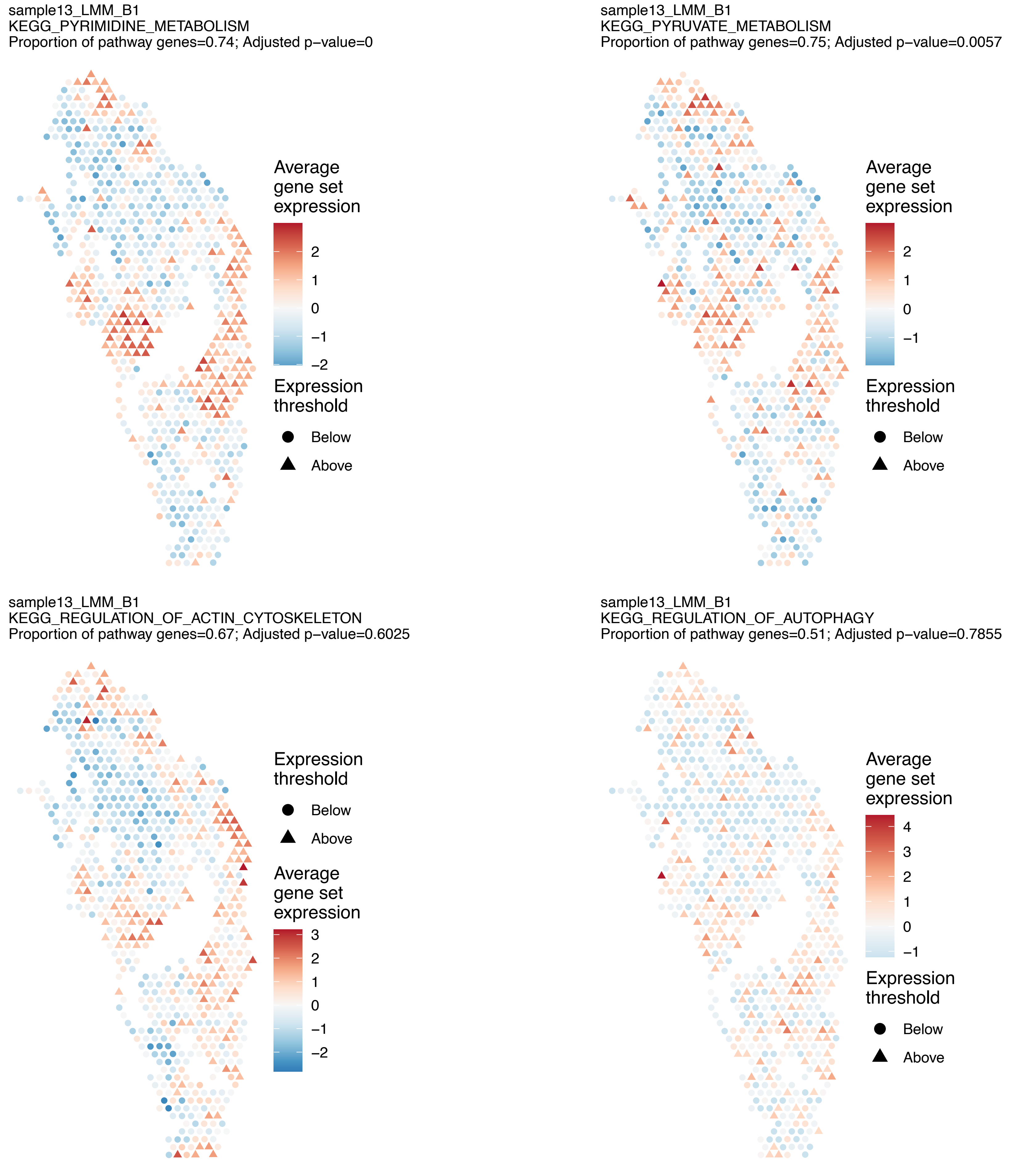
sample13_LMM_B1
KEGG_PYRIMIDINE_METABOLISM
Proportion of pathway genes=0.74; Adjusted p−value=0
Average gene set expression
2
1
0
−1
−2
Expression threshold
Below
Above
sample13_LMM_B1
KEGG_PYRUVATE_METABOLISM
Proportion of pathway genes=0.75; Adjusted p−value=0.0057
Average gene set expression
2
1
0
−1
Expression threshold
Below
Above
sample13_LMM_B1
KEGG_REGULATION_OF_ACTIN_CYTOSKELETON
Proportion of pathway genes=0.67; Adjusted p−value=0.6025
Expression threshold
Below
Above
Average gene set expression
3
2
1
0
−1
−2
sample13_LMM_B1
KEGG_REGULATION_OF_AUTOPHAGY
Proportion of pathway genes=0.51; Adjusted p−value=0.7855
Average gene set expression
4
3
2
1
0
−1
Expression threshold
Below
Above

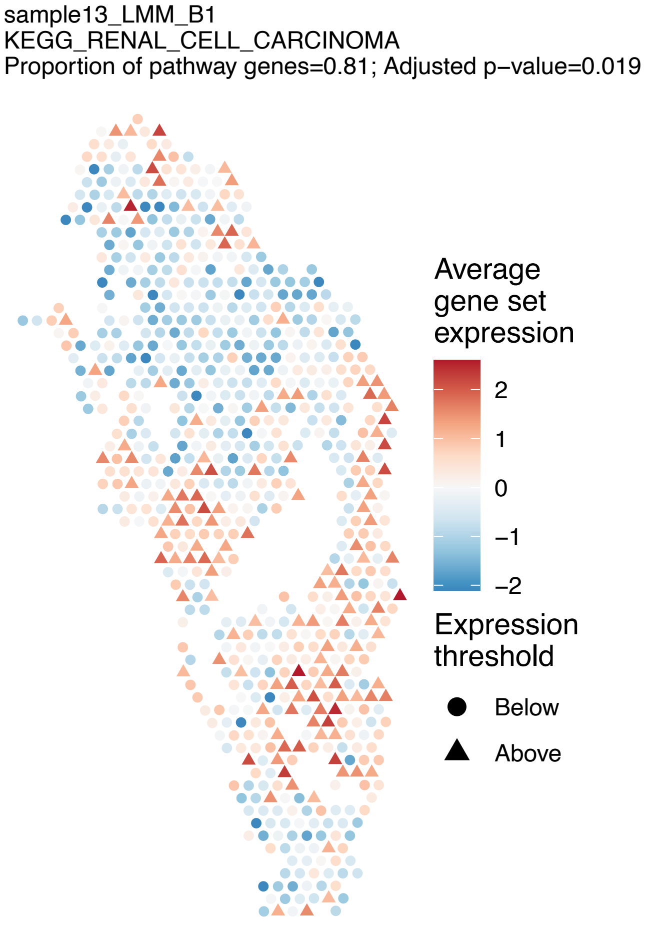

sample13_LMM_B1
KEGG_RENAL_CELL_CARCINOMA
Proportion of pathway genes=0.81; Adjusted p−value=0.019
Average gene set expression
2
1
0
−1
−2
Expression threshold
Below
Above


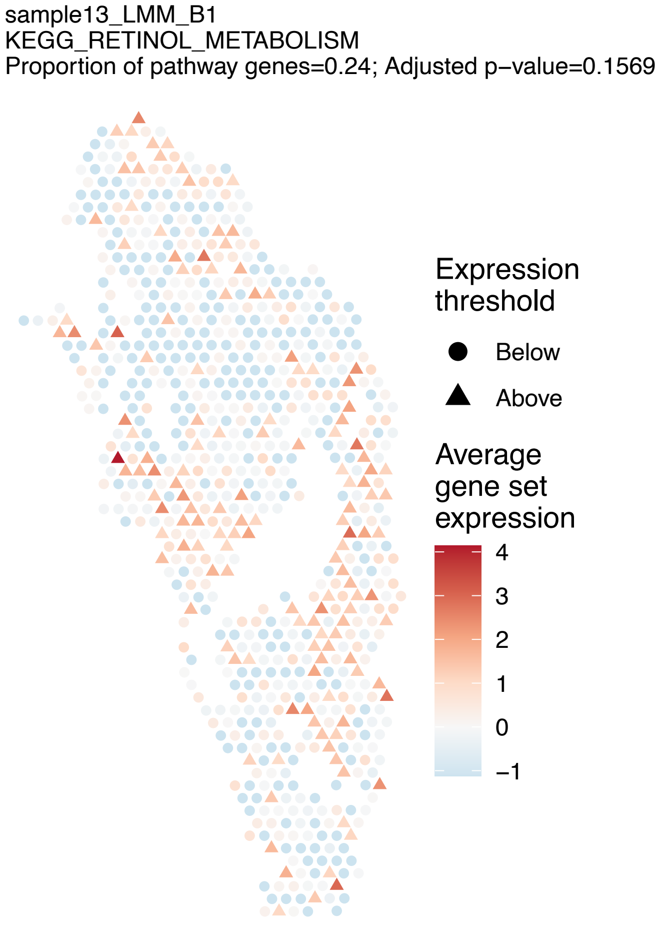

sample13_LMM_B1
KEGG_RETINOL_METABOLISM
Proportion of pathway genes=0.24; Adjusted p−value=0.1569
Expression threshold
Below
Above
Average gene set expression
4
3
2
1
0
−1


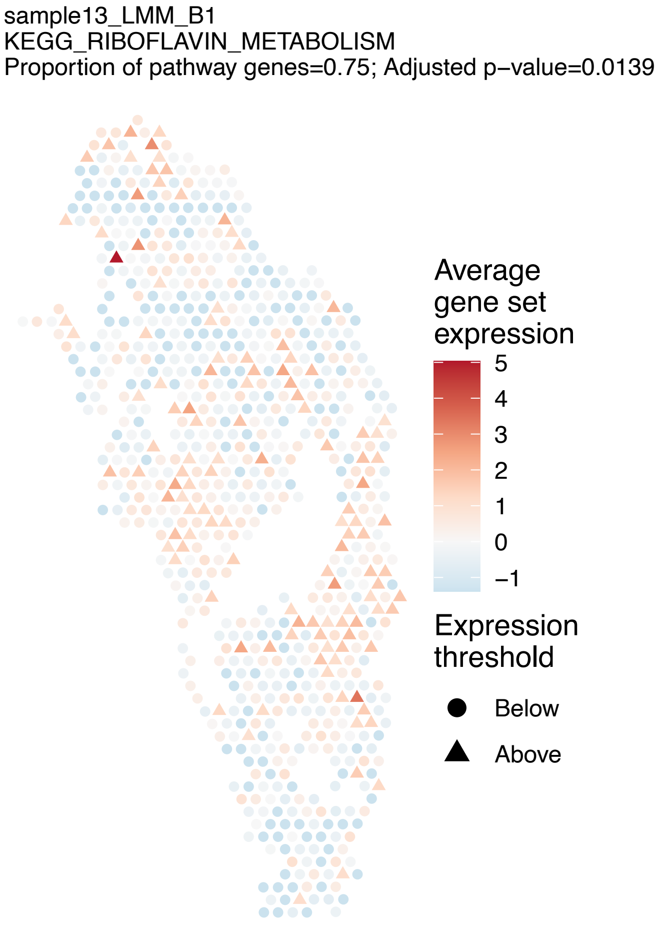

sample13_LMM_B1
KEGG_RIBOFLAVIN_METABOLISM
Proportion of pathway genes=0.75; Adjusted p−value=0.0139
Average gene set expression
5
4
3
2
1
0
−1
Expression threshold
Below
Above


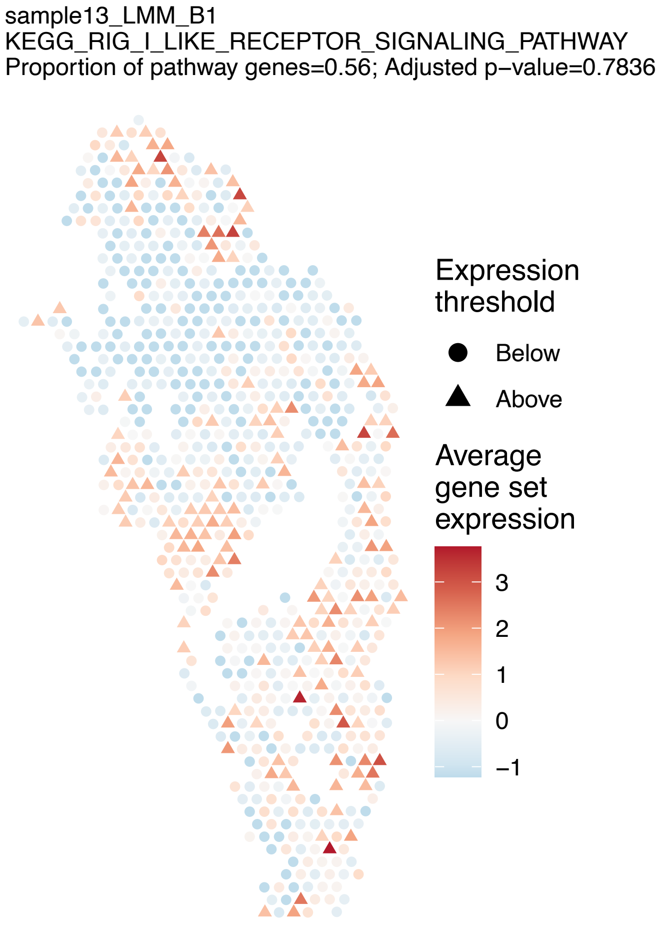

sample13_LMM_B1
KEGG_RIG_I_LIKE_RECEPTOR_SIGNALING_PATHWAY
Proportion of pathway genes=0.56; Adjusted p−value=0.7836
Expression threshold
Below
Above
Average gene set expression
3
2
1
0
−1

sample13_LMM_B1
KEGG_RNA_DEGRADATION
Proportion of pathway genes=0.96; Adjusted p−value=0.0167

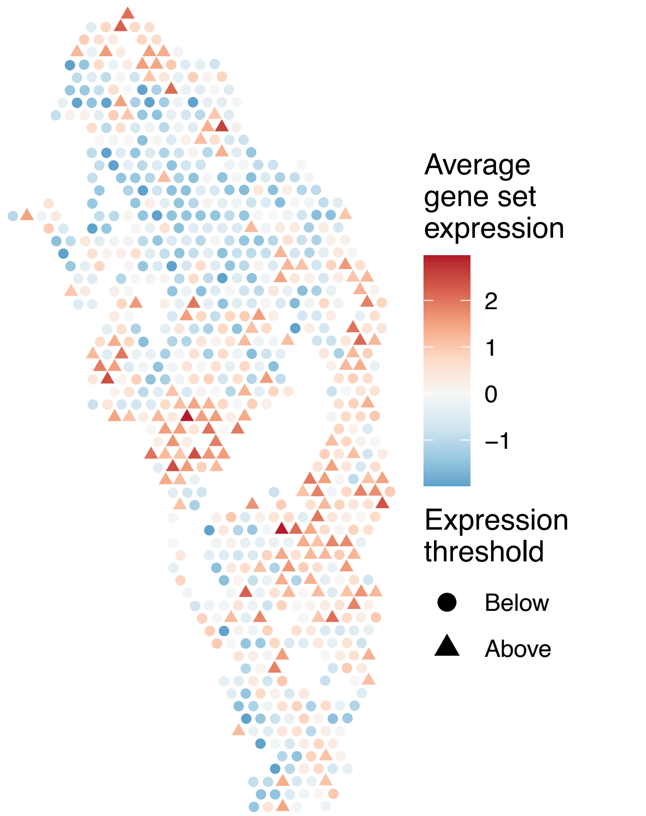


sample13_LMM_B1
KEGG_RNA_POLYMERASE
Proportion of pathway genes=0.69; Adjusted p−value=0

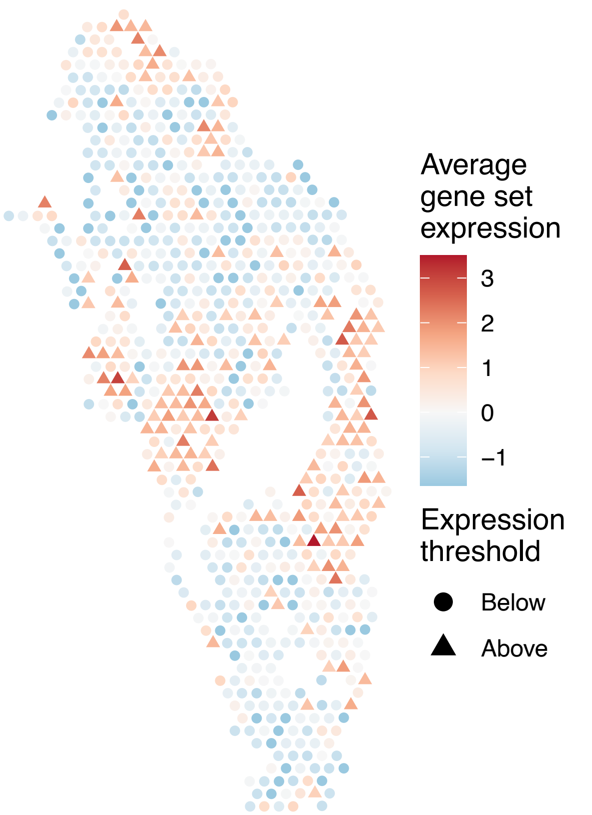


sample13_LMM_B1
KEGG_SELENOAMINO_ACID_METABOLISM
Proportion of pathway genes=0.73; Adjusted p−value=0.3215

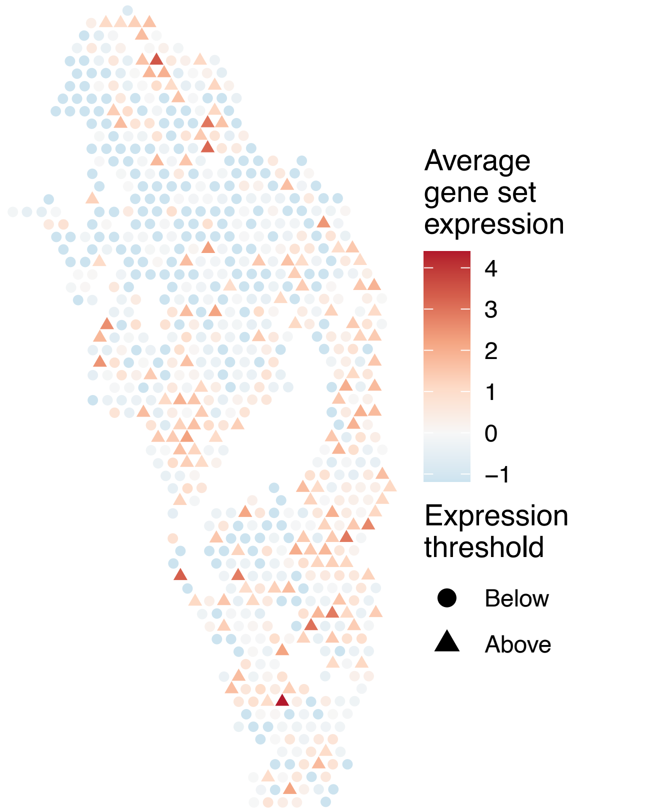


sample13_LMM_B1
KEGG_SMALL_CELL_LUNG_CANCER
Proportion of pathway genes=0.78; Adjusted p−value=0

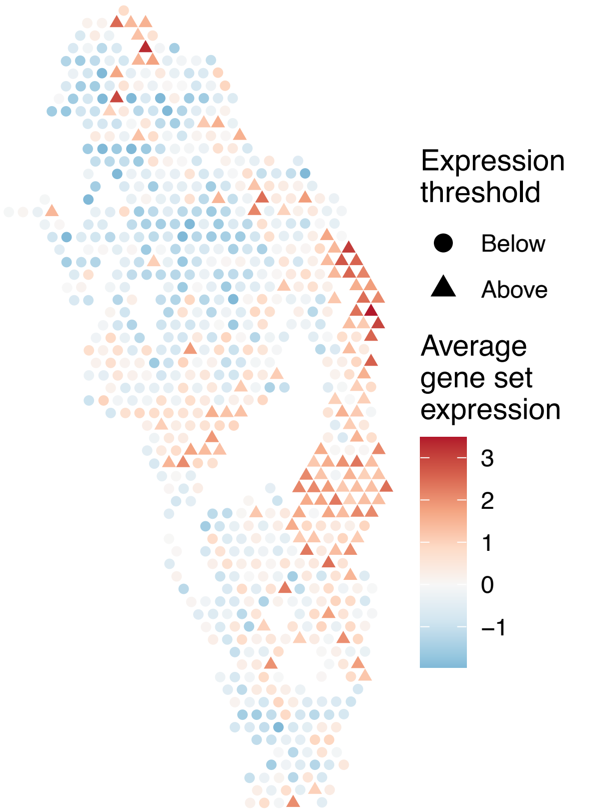

sample13_LMM_B1
KEGG_SNARE_INTERACTIONS_IN_VESICULAR_TRANSPORT
Proportion of pathway genes=0.82; Adjusted p−value=0.9523

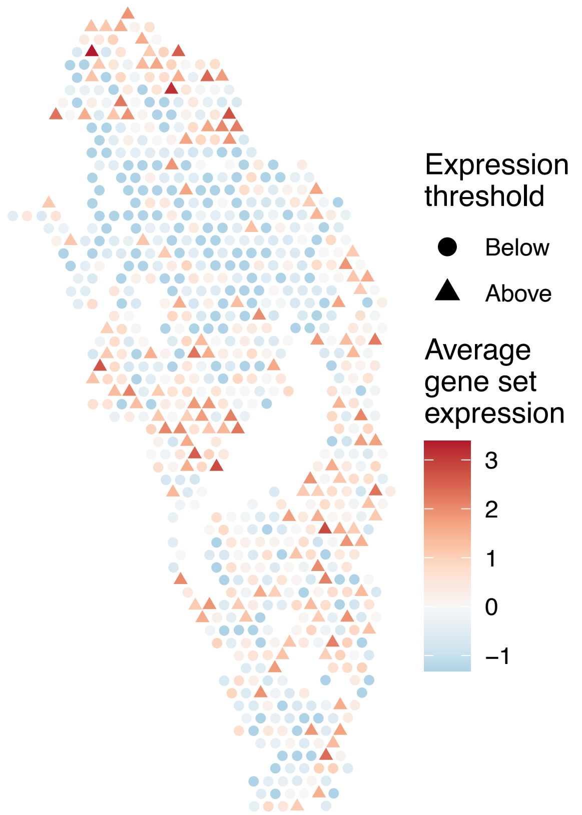


sample13_LMM_B1
KEGG_SPHINGOLIPID_METABOLISM
Proportion of pathway genes=0.7; Adjusted p−value=0.0167

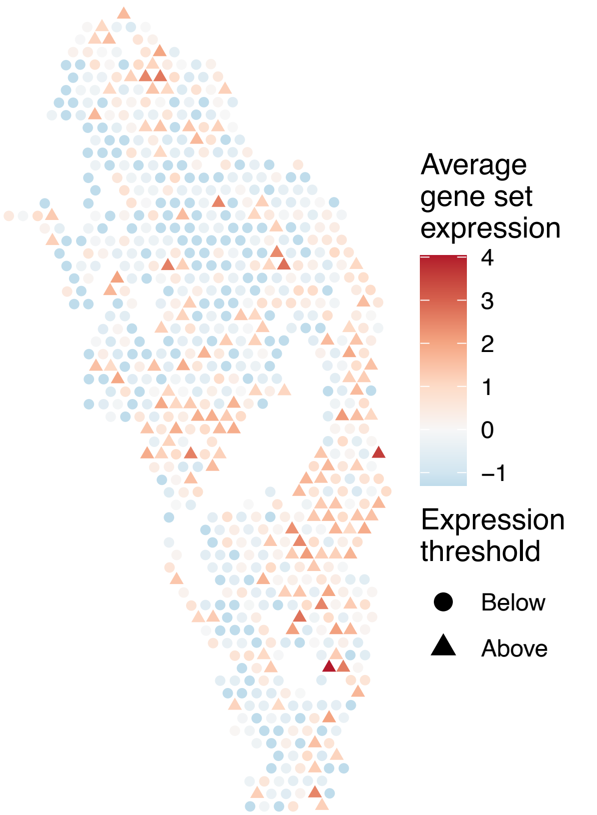


sample13_LMM_B1
KEGG_SPLICEOSOME
Proportion of pathway genes=0.95; Adjusted p−value=0

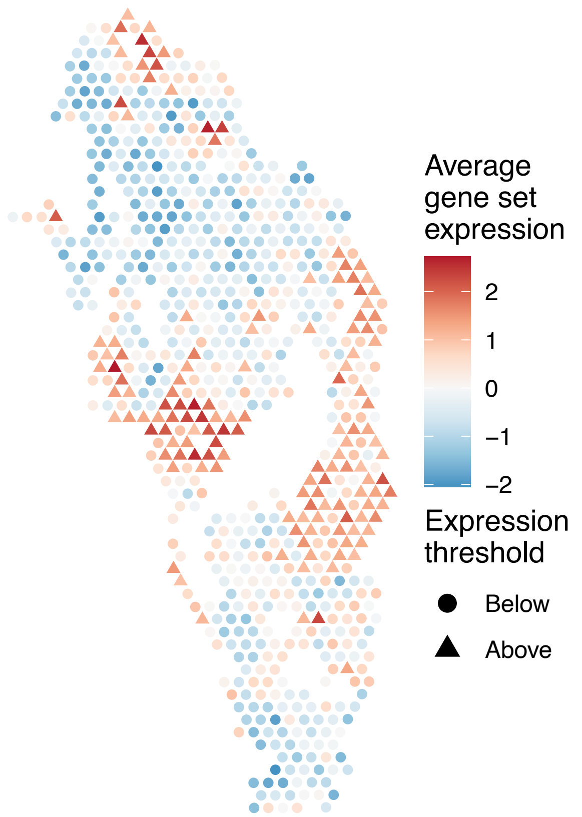


sample13_LMM_B1
KEGG_STARCH_AND_SUCROSE_METABOLISM
Proportion of pathway genes=0.34; Adjusted p−value=0.9523

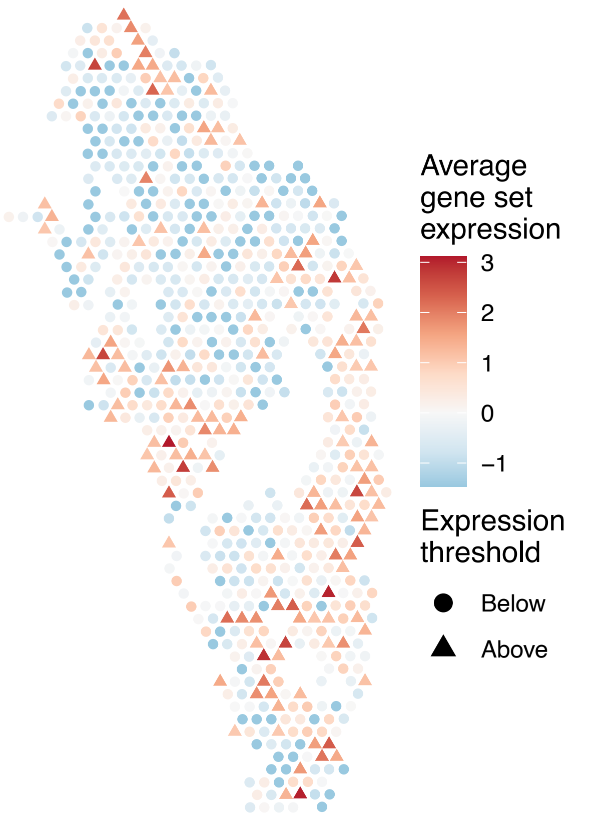

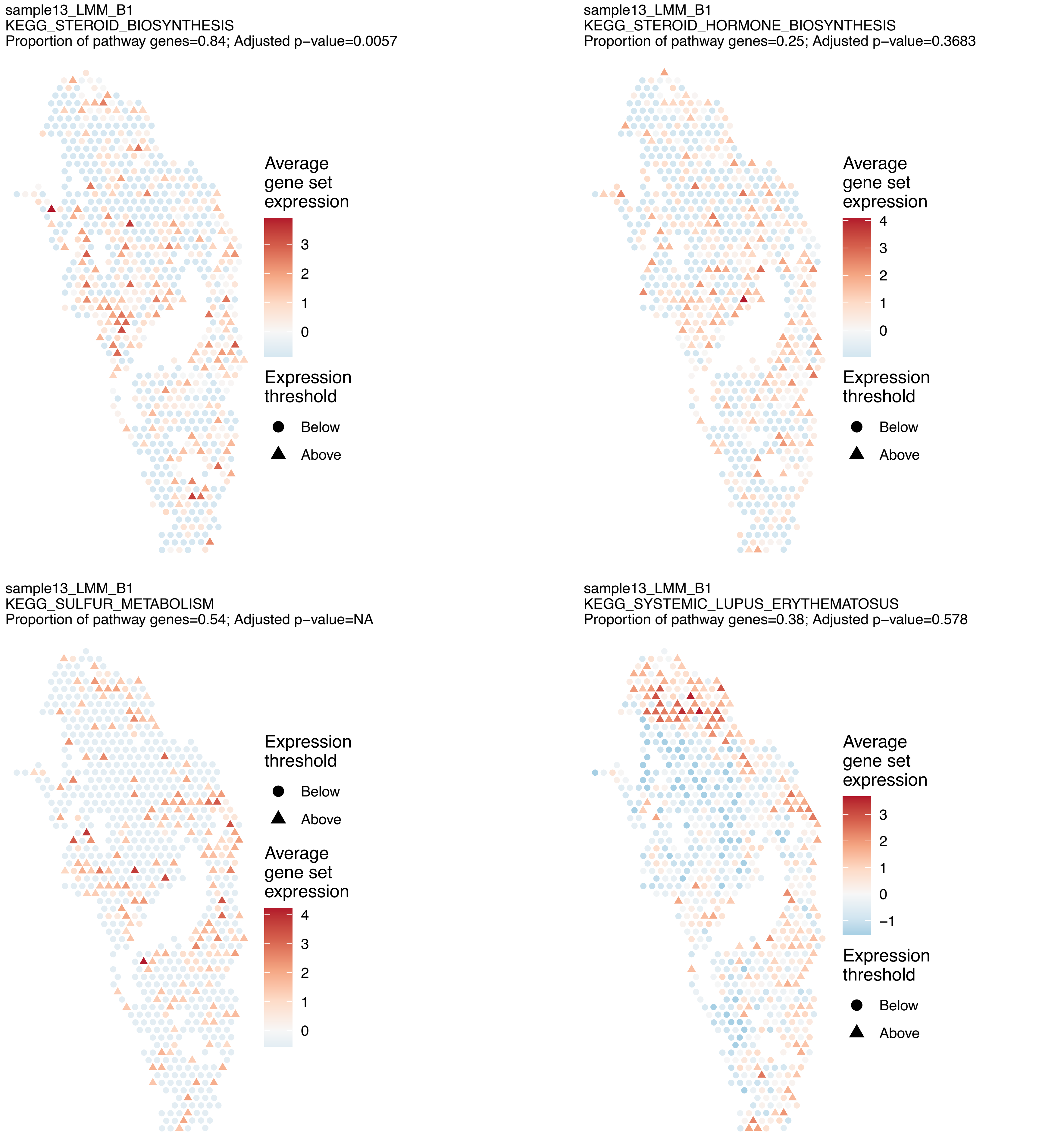
sample13_LMM_B1
KEGG_STEROID_BIOSYNTHESIS
Proportion of pathway genes=0.84; Adjusted p−value=0.0057
Average gene set expression
3
2
1
0
Expression threshold
Below
Above
sample13_LMM_B1
KEGG_STEROID_HORMONE_BIOSYNTHESIS
Proportion of pathway genes=0.25; Adjusted p−value=0.3683
Average gene set expression
4
3
2
1
0
Expression threshold
Below
Above
sample13_LMM_B1
KEGG_SULFUR_METABOLISM
Proportion of pathway genes=0.54; Adjusted p−value=NA
Expression threshold
Below
Above
Average gene set expression
4
3
2
1
0
sample13_LMM_B1
KEGG_SYSTEMIC_LUPUS_ERYTHEMATOSUS
Proportion of pathway genes=0.38; Adjusted p−value=0.578
Average gene set expression
3
2
1
0
−1
Expression threshold
Below
Above

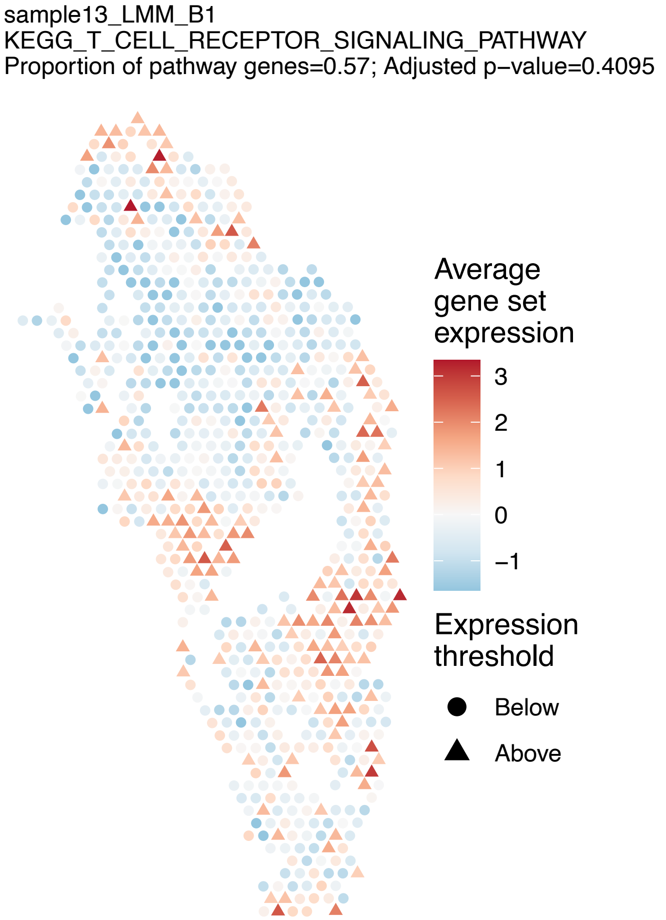
sample13_LMM_B1
KEGG_T_CELL_RECEPTOR_SIGNALING_PATHWAY
Proportion of pathway genes=0.57; Adjusted p−value=0.4095
Average gene set expression
3
2
1
0
−1
Expression threshold
Below
Above

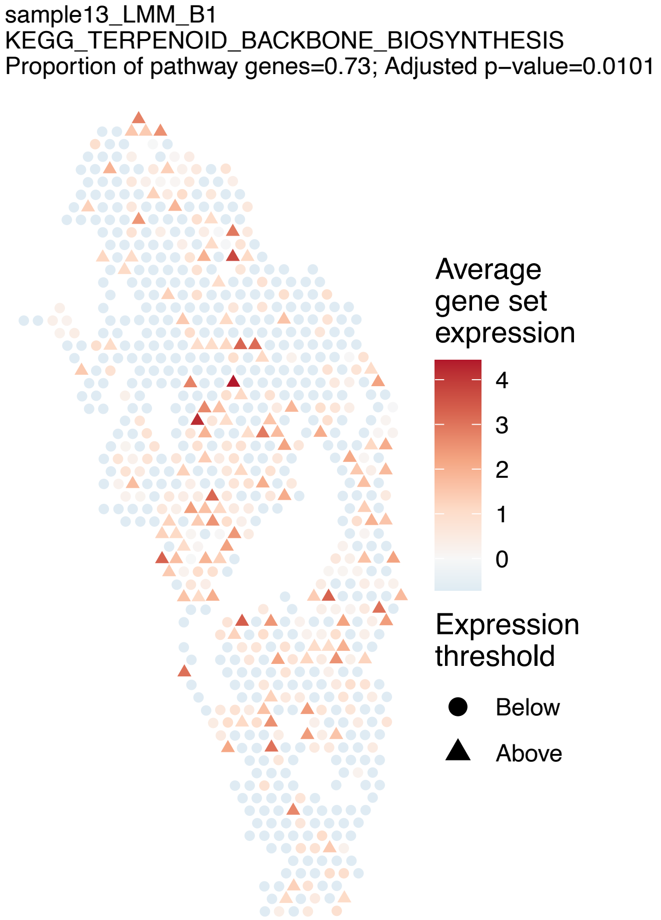
sample13_LMM_B1
KEGG_TERPENOID_BACKBONE_BIOSYNTHESIS
Proportion of pathway genes=0.73; Adjusted p−value=0.0101
Average gene set expression
4
3
2
1
0
Expression threshold
Below
Above

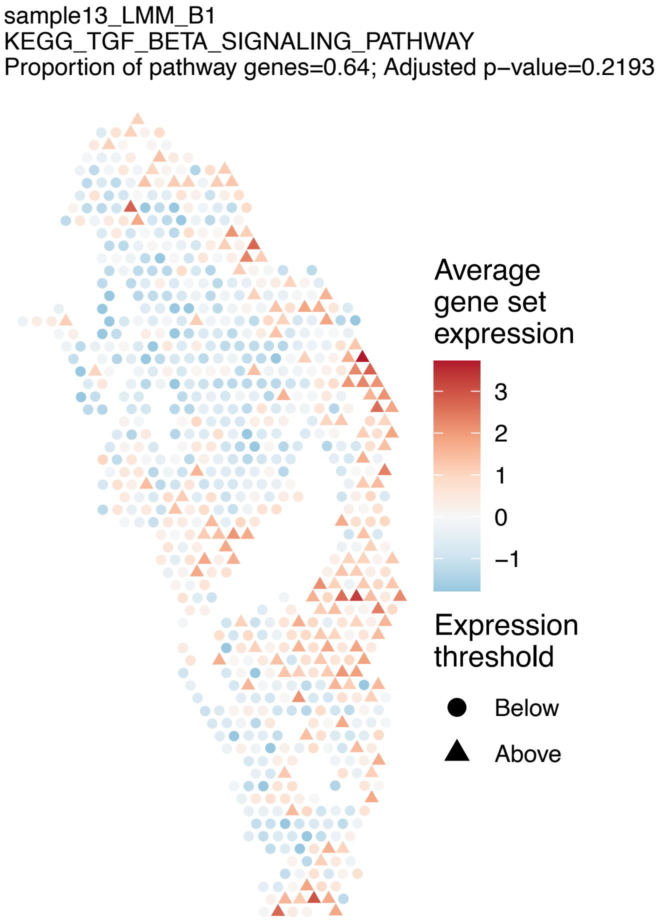
sample13_LMM_B1
KEGG_TGF_BETA_SIGNALING_PATHWAY
Proportion of pathway genes=0.64; Adjusted p−value=0.2193
Average gene set expression
3
2
1
0
−1
Expression threshold
Below
Above

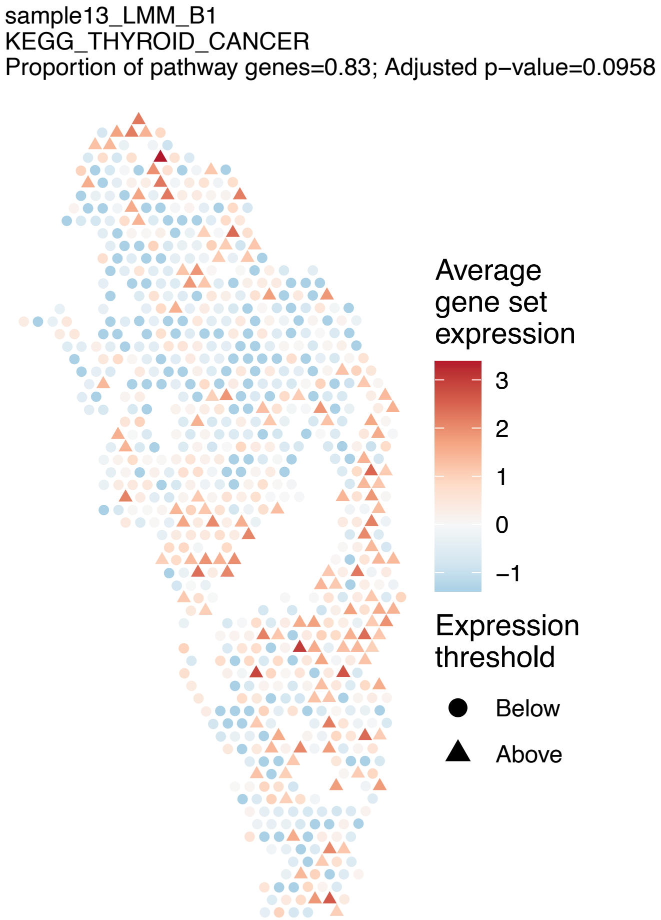
sample13_LMM_B1
KEGG_THYROID_CANCER
Proportion of pathway genes=0.83; Adjusted p−value=0.0958
Average gene set expression
3
2
1
0
−1
Expression threshold
Below
Above

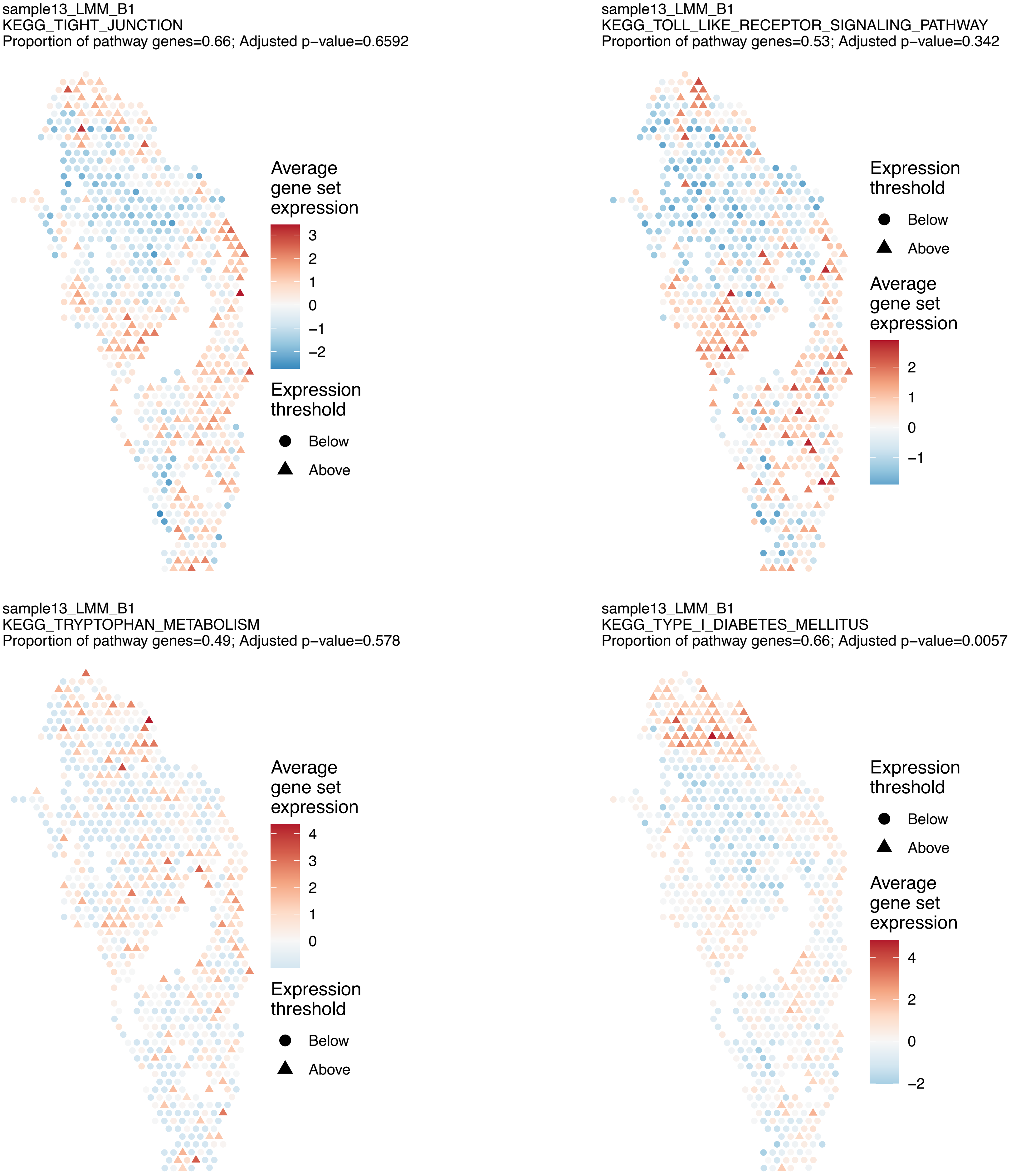
sample13_LMM_B1
KEGG_TIGHT_JUNCTION
Proportion of pathway genes=0.66; Adjusted p−value=0.6592
Average gene set expression
3
2
1
0
−1
−2
Expression threshold
Below
Above
sample13_LMM_B1
KEGG_TOLL_LIKE_RECEPTOR_SIGNALING_PATHWAY
Proportion of pathway genes=0.53; Adjusted p−value=0.342
Expression threshold
Below
Above
Average gene set expression
2
1
0
−1
sample13_LMM_B1
KEGG_TRYPTOPHAN_METABOLISM
Proportion of pathway genes=0.49; Adjusted p−value=0.578
Average gene set expression
4
3
2
1
0
Expression threshold
Below
Above
sample13_LMM_B1
KEGG_TYPE_I_DIABETES_MELLITUS
Proportion of pathway genes=0.66; Adjusted p−value=0.0057
Expression threshold
Below
Above
Average gene set expression
4
2
0
−2

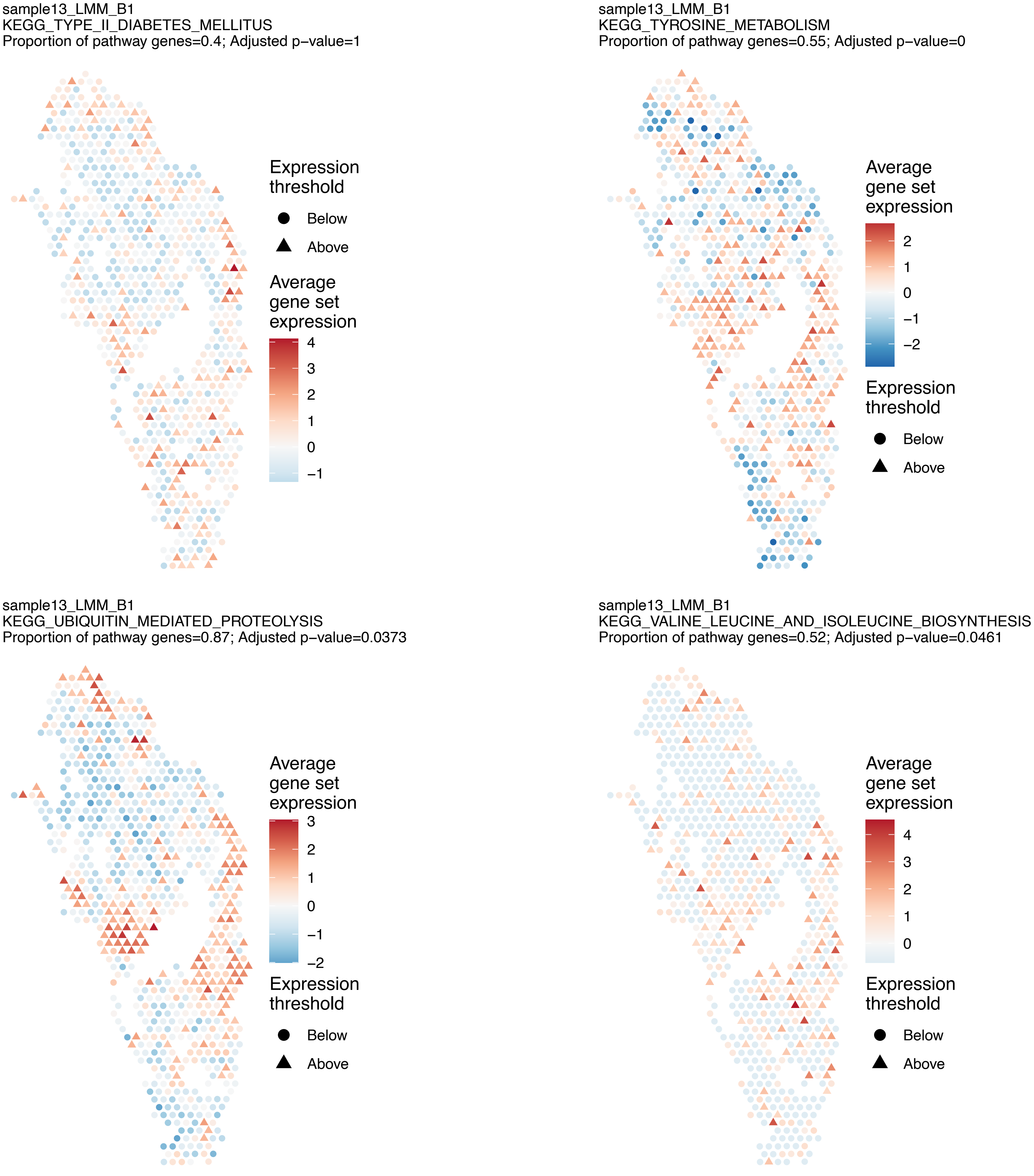
sample13_LMM_B1
KEGG_TYPE_II_DIABETES_MELLITUS
Proportion of pathway genes=0.4; Adjusted p−value=1
Expression threshold
Below
Above
Average gene set expression
4
3
2
1
0
−1
sample13_LMM_B1
KEGG_TYROSINE_METABOLISM
Proportion of pathway genes=0.55; Adjusted p−value=0
Average gene set expression
2
1
0
−1
−2
Expression threshold
Below
Above
sample13_LMM_B1
KEGG_UBIQUITIN_MEDIATED_PROTEOLYSIS
Proportion of pathway genes=0.87; Adjusted p−value=0.0373
Average gene set expression
3
2
1
0
−1
−2
Expression threshold
Below
Above
sample13_LMM_B1
KEGG_VALINE_LEUCINE_AND_ISOLEUCINE_BIOSYNTHESIS
Proportion of pathway genes=0.52; Adjusted p−value=0.0461
Average gene set expression
4
3
2
1
0
Expression threshold
Below
Above

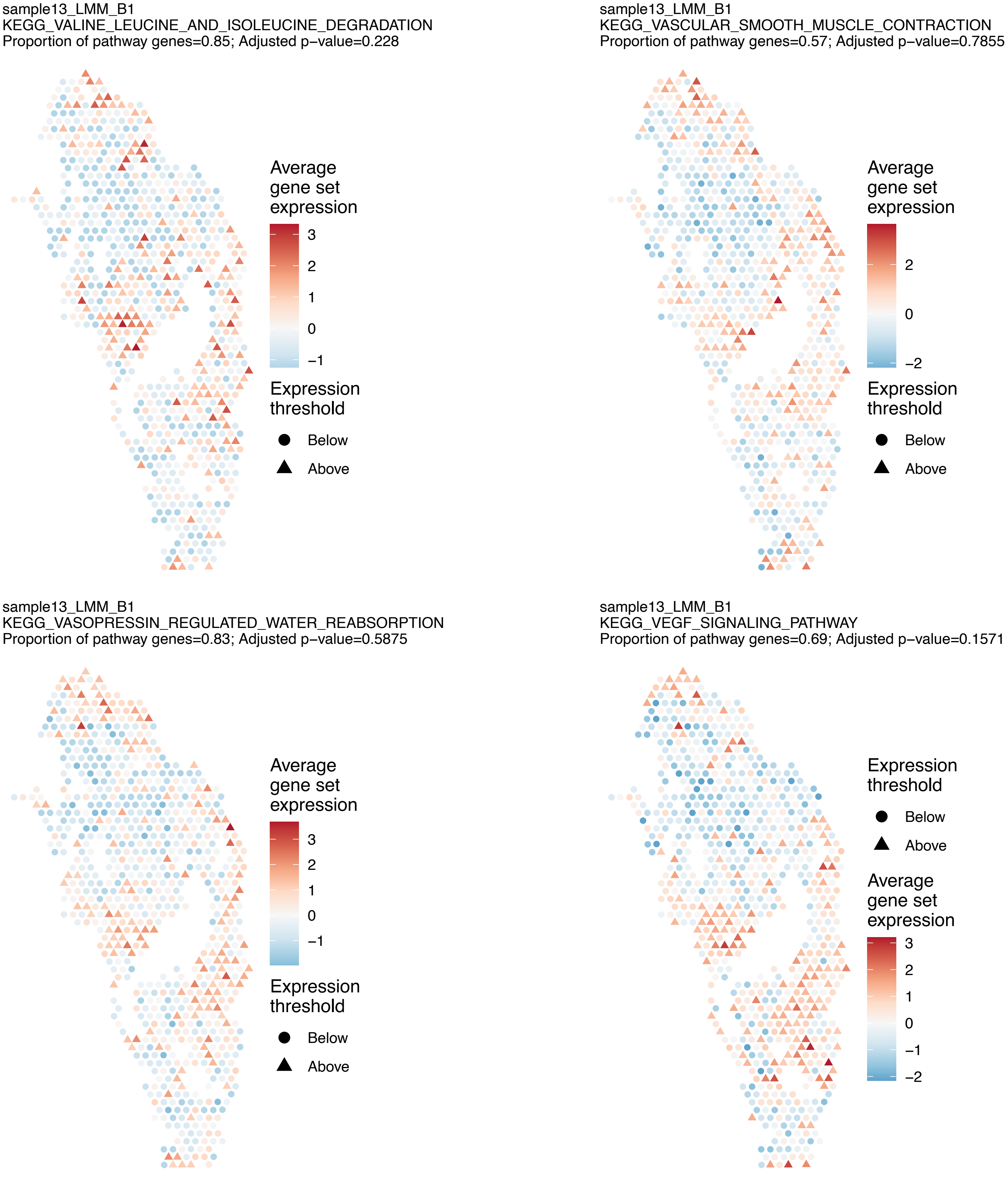
sample13_LMM_B1
KEGG_VALINE_LEUCINE_AND_ISOLEUCINE_DEGRADATION
Proportion of pathway genes=0.85; Adjusted p−value=0.228
Average gene set expression
3
2
1
0
−1
Expression threshold
Below
Above
sample13_LMM_B1
KEGG_VASCULAR_SMOOTH_MUSCLE_CONTRACTION
Proportion of pathway genes=0.57; Adjusted p−value=0.7855
Average gene set expression
2
0
−2
Expression threshold
Below
Above
sample13_LMM_B1
KEGG_VASOPRESSIN_REGULATED_WATER_REABSORPTION
Proportion of pathway genes=0.83; Adjusted p−value=0.5875
Average gene set expression
3
2
1
0
−1
Expression threshold
Below
Above
sample13_LMM_B1
KEGG_VEGF_SIGNALING_PATHWAY
Proportion of pathway genes=0.69; Adjusted p−value=0.1571
Expression threshold
Below
Above
Average gene set expression
3
2
1
0
−1
−2

sample13_LMM_B1
KEGG_VIBRIO_CHOLERAE_INFECTION
Proportion of pathway genes=0.63; Adjusted p−value=0.0879

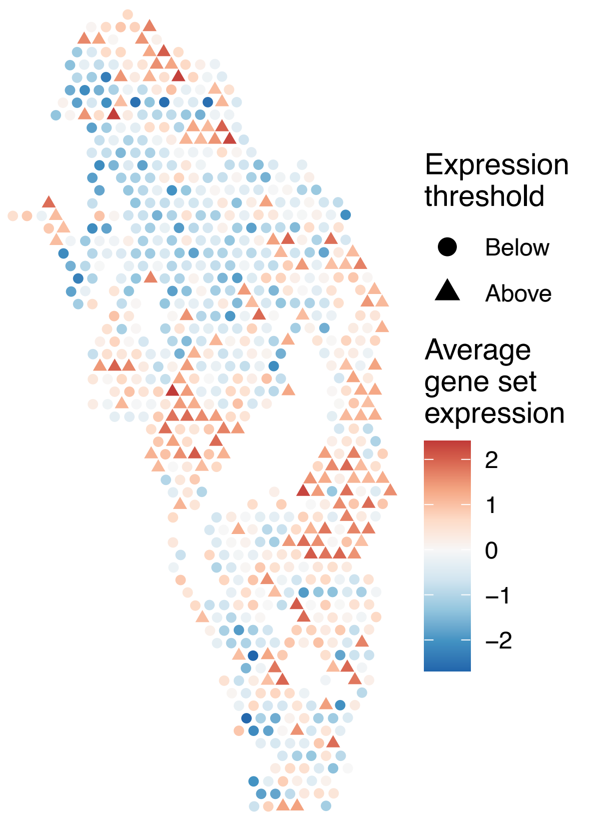


sample13_LMM_B1
KEGG_VIRAL_MYOCARDITIS
Proportion of pathway genes=0.74; Adjusted p−value=0.1102

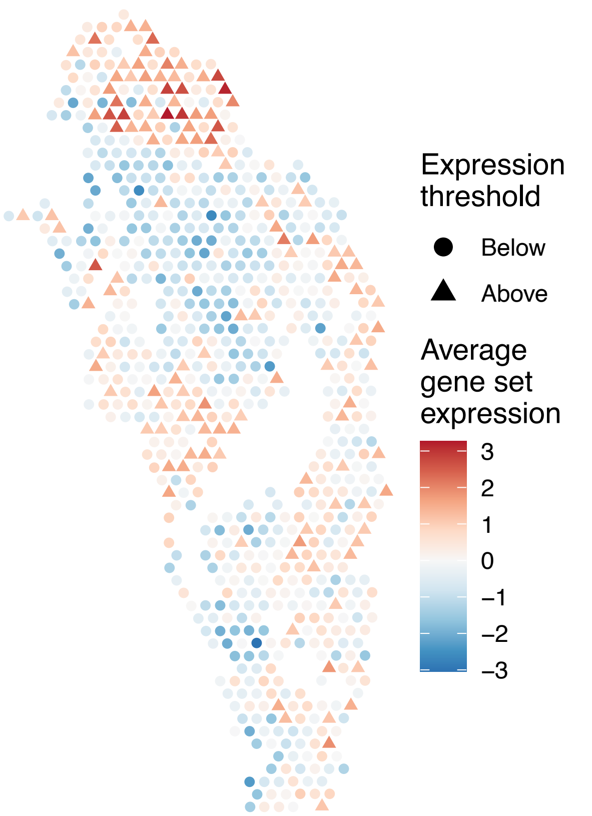


sample13_LMM_B1
KEGG_WNT_SIGNALING_PATHWAY
Proportion of pathway genes=0.67; Adjusted p−value=0.578

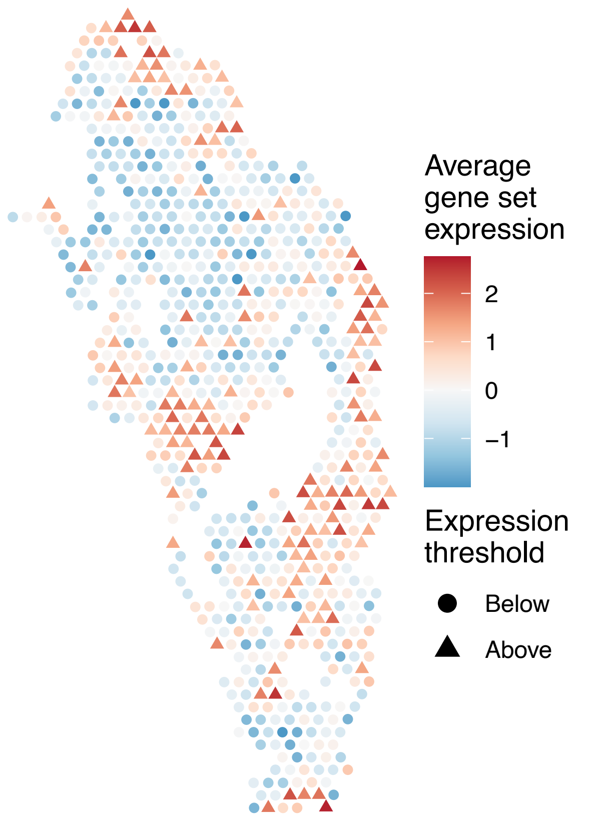


sample13_LMM_B1
KEGG_PENTOSE_AND_GLUCURONATE_INTERCONVERSIONS
Proportion of pathway genes=0.29; Adjusted p−value=NA

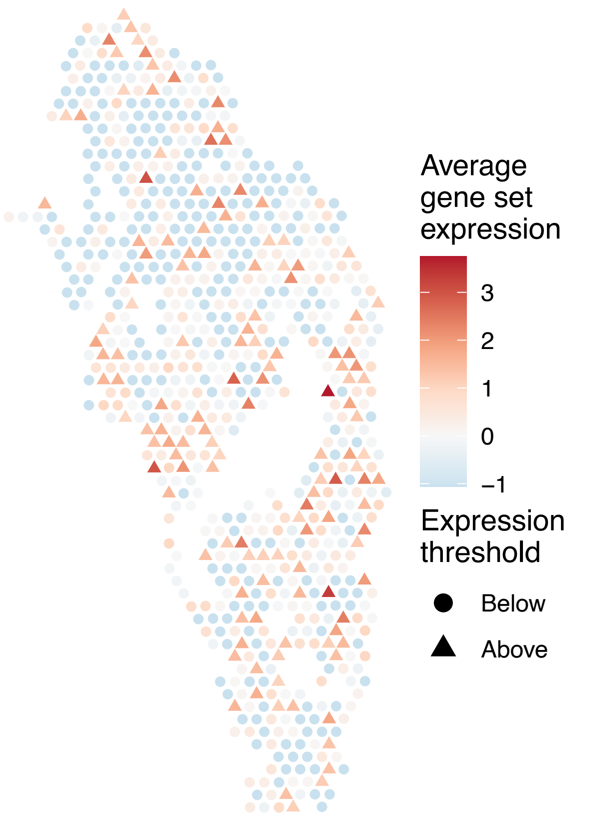

sample14_LMM_B5
KEGG_NATURAL_KILLER_CELL_MEDIATED_CYTOTOXICITY
Proportion of pathway genes=0.23; Adjusted p−value=0.0259

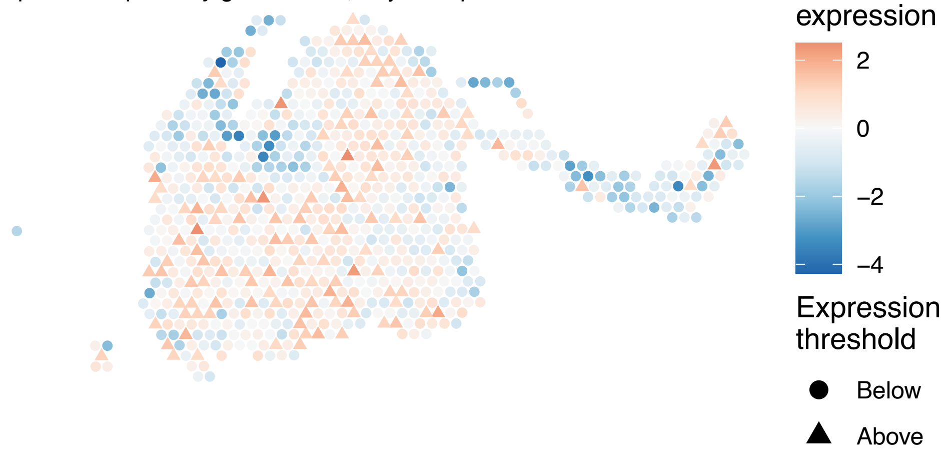


sample14_LMM_B5
KEGG_RIBOSOME
Proportion of pathway genes=0.97; Adjusted p−value=0

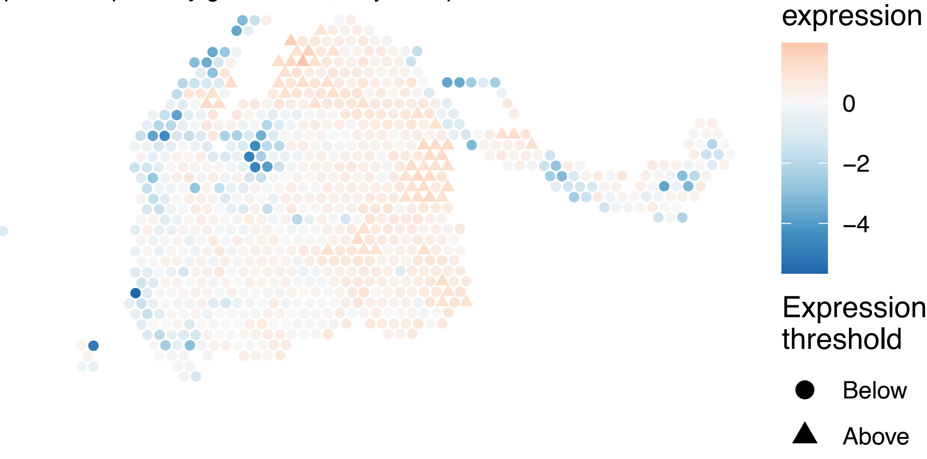


sample14_LMM_B5
KEGG_PRIMARY_IMMUNODEFICIENCY
Proportion of pathway genes=0.46; Adjusted p−value=0.3815

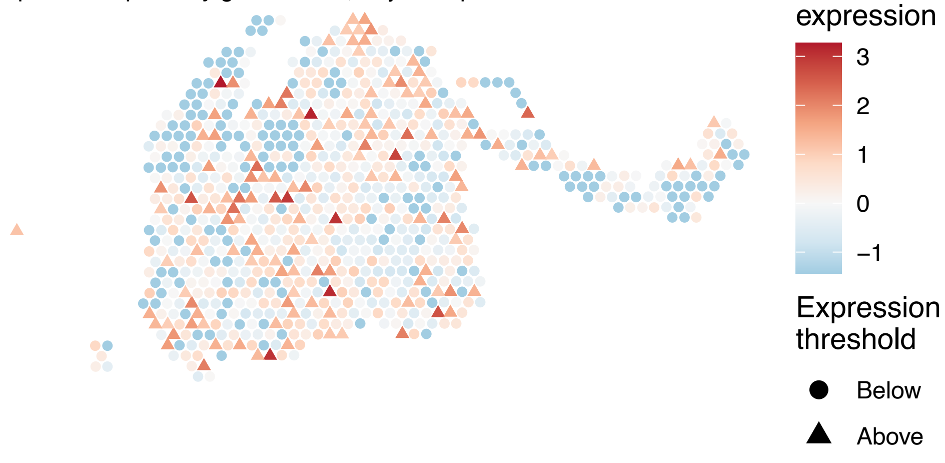


sample14_LMM_B5
KEGG_ANTIGEN_PROCESSING_AND_PRESENTATION
Proportion of pathway genes=0.28; Adjusted p−value=0

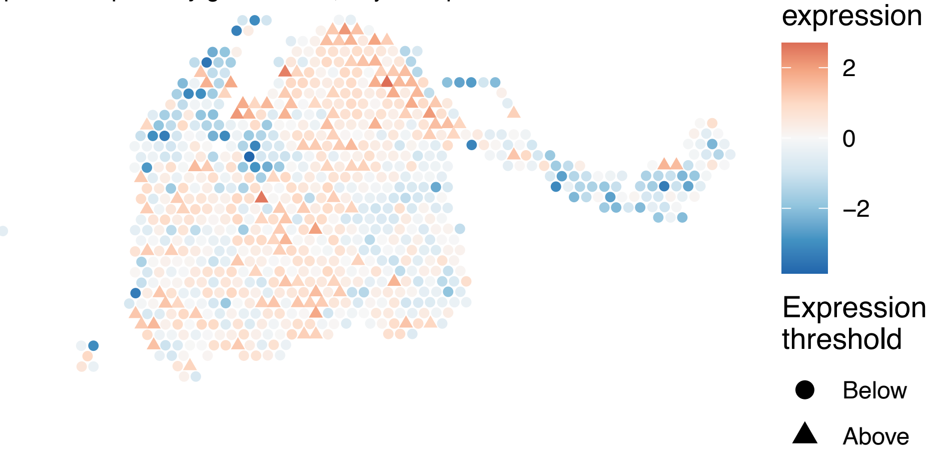

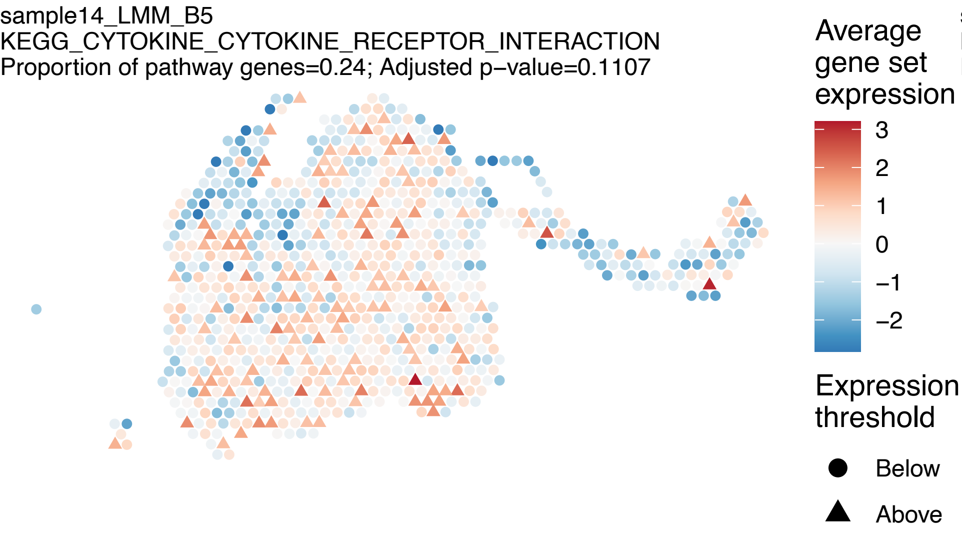
sample14_LMM_B5
KEGG_CYTOKINE_CYTOKINE_RECEPTOR_INTERACTION
Proportion of pathway genes=0.24; Adjusted p−value=0.1107
Average gene set expression
3
2
1
0
−1
−2
Expression threshold
Below
Above

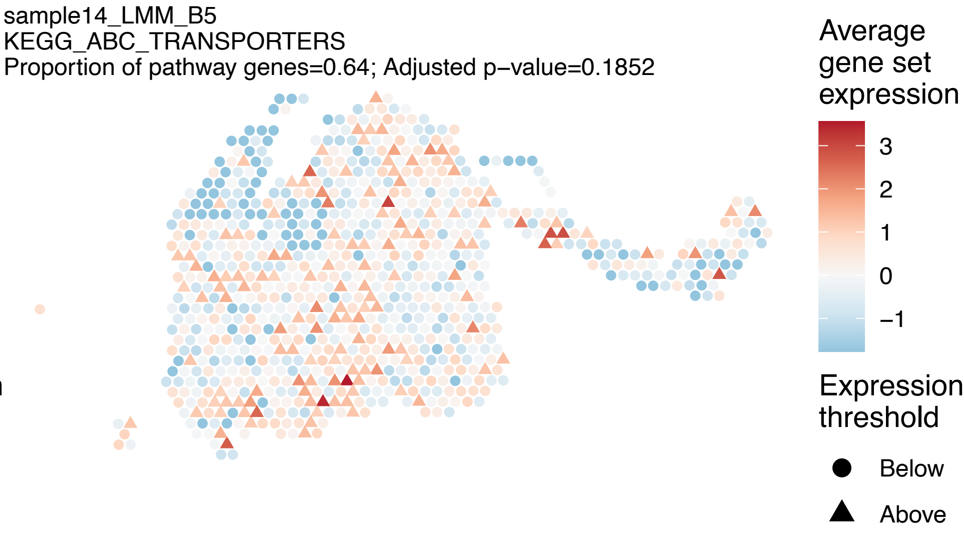
sample14_LMM_B5
KEGG_ABC_TRANSPORTERS
Proportion of pathway genes=0.64; Adjusted p−value=0.1852
Average gene set expression
3
2
1
0
−1
Expression threshold
Below
Above

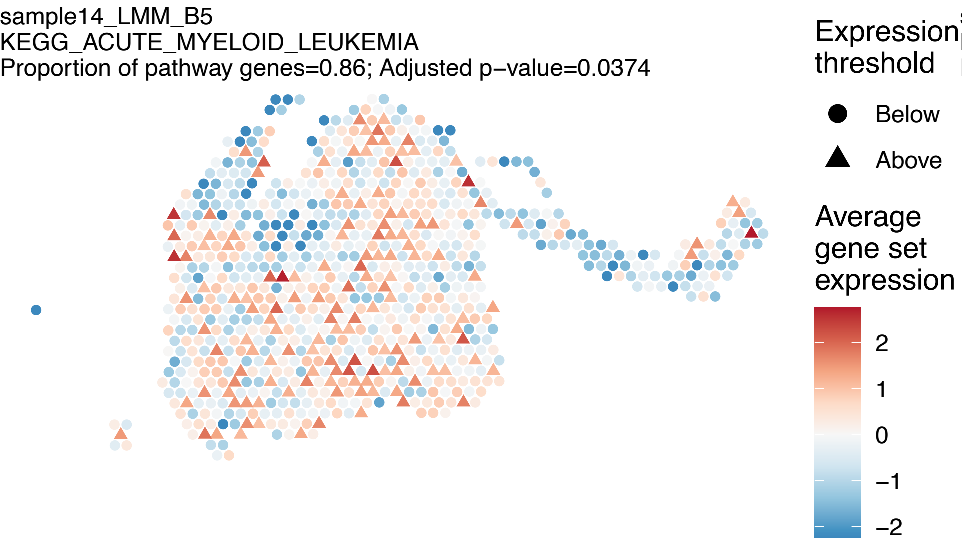
sample14_LMM_B5
KEGG_ACUTE_MYELOID_LEUKEMIA
Proportion of pathway genes=0.86; Adjusted p−value=0.0374
Expression threshold
Below
Above
Average gene set expression
2
1
0
−1
−2

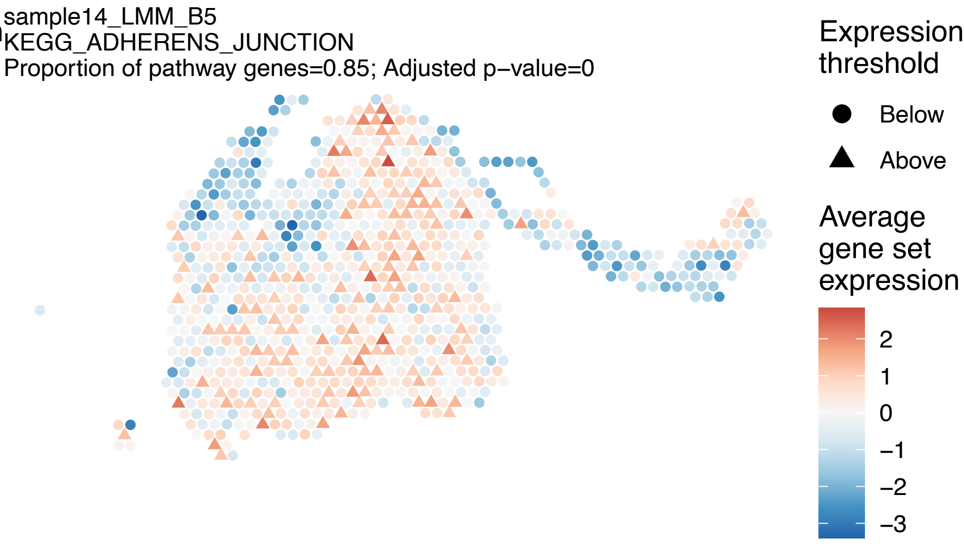
sample14_LMM_B5
KEGG_ADHERENS_JUNCTION
Proportion of pathway genes=0.85; Adjusted p−value=0
Expression threshold
Below
Above
Average gene set expression
2
1
0
−1
−2
−3

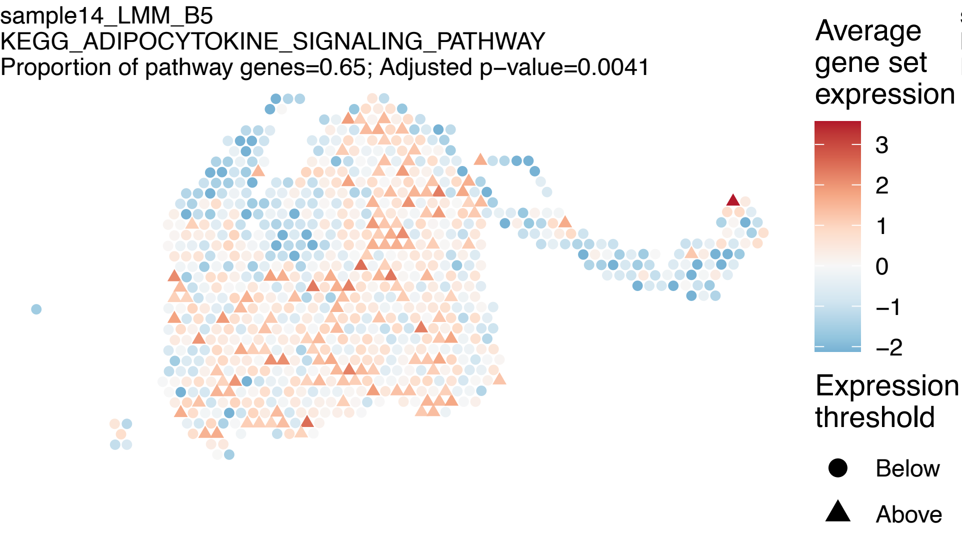
sample14_LMM_B5
KEGG_ADIPOCYTOKINE_SIGNALING_PATHWAY
Proportion of pathway genes=0.65; Adjusted p−value=0.0041
Average gene set expression
3
2
1
0
−1
−2
Expression threshold
Below
Above

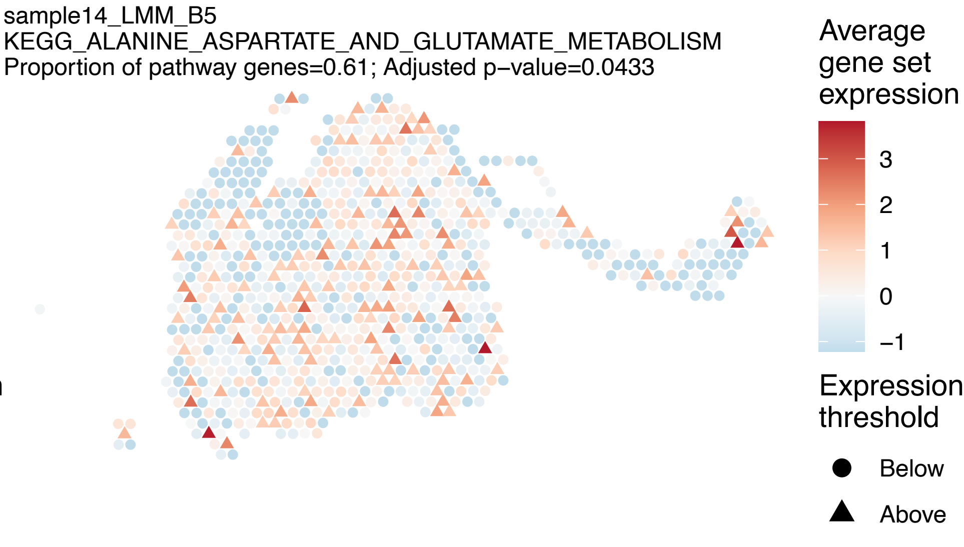
sample14_LMM_B5
KEGG_ALANINE_ASPARTATE_AND_GLUTAMATE_METABOLISM
Proportion of pathway genes=0.61; Adjusted p−value=0.0433
Average gene set expression
3
2
1
0
−1
Expression threshold
Below
Above

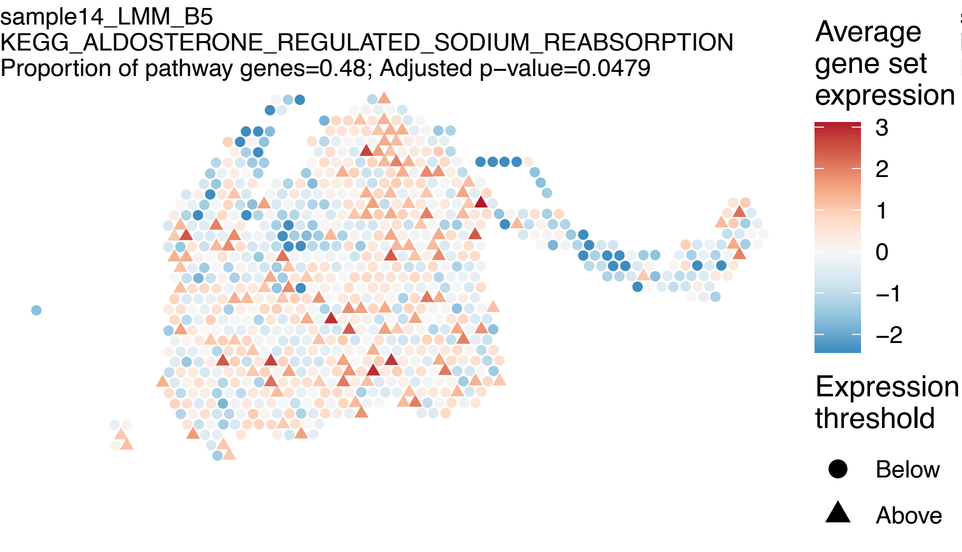
sample14_LMM_B5
KEGG_ALDOSTERONE_REGULATED_SODIUM_REABSORPTION
Proportion of pathway genes=0.48; Adjusted p−value=0.0479
Average gene set expression
3
2
1
0
−1
−2
Expression threshold
Below
Above

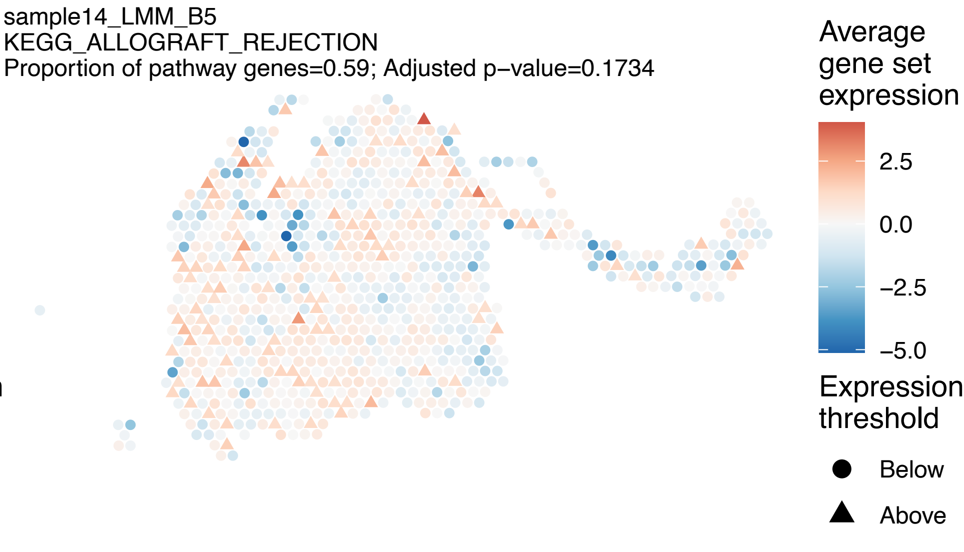
sample14_LMM_B5
KEGG_ALLOGRAFT_REJECTION
Proportion of pathway genes=0.59; Adjusted p−value=0.1734
Average gene set expression
2.5
0.0
−2.5
−5.0
Expression threshold
Below
Above

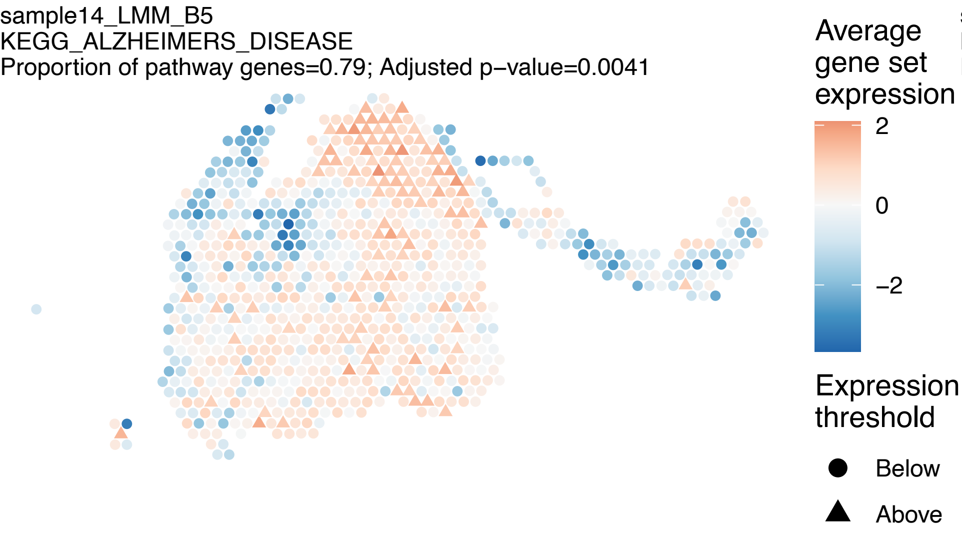
sample14_LMM_B5
KEGG_ALZHEIMERS_DISEASE
Proportion of pathway genes=0.79; Adjusted p−value=0.0041
Average gene set expression
2
0
−2
Expression threshold
Below
Above

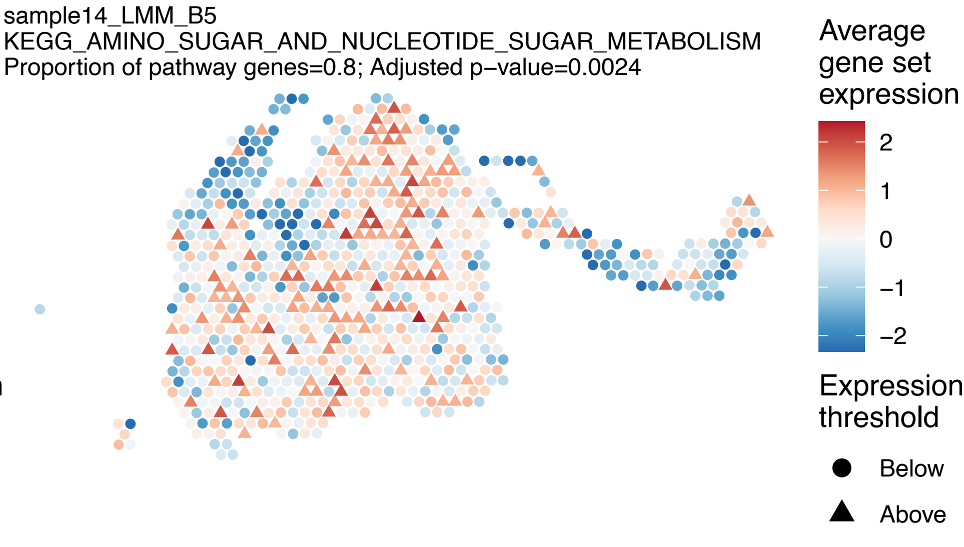
sample14_LMM_B5
KEGG_AMINO_SUGAR_AND_NUCLEOTIDE_SUGAR_METABOLISM
Proportion of pathway genes=0.8; Adjusted p−value=0.0024
Average gene set expression
2
1
0
−1
−2
Expression threshold
Below
Above

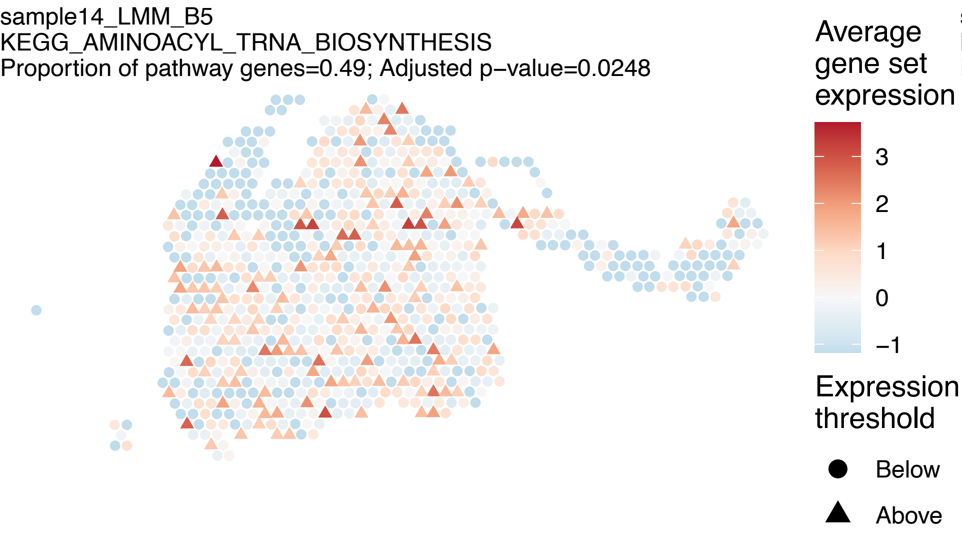
sample14_LMM_B5
KEGG_AMINOACYL_TRNA_BIOSYNTHESIS
Proportion of pathway genes=0.49; Adjusted p−value=0.0248
Average gene set expression
3
2
1
0
−1
Expression threshold
Below
Above

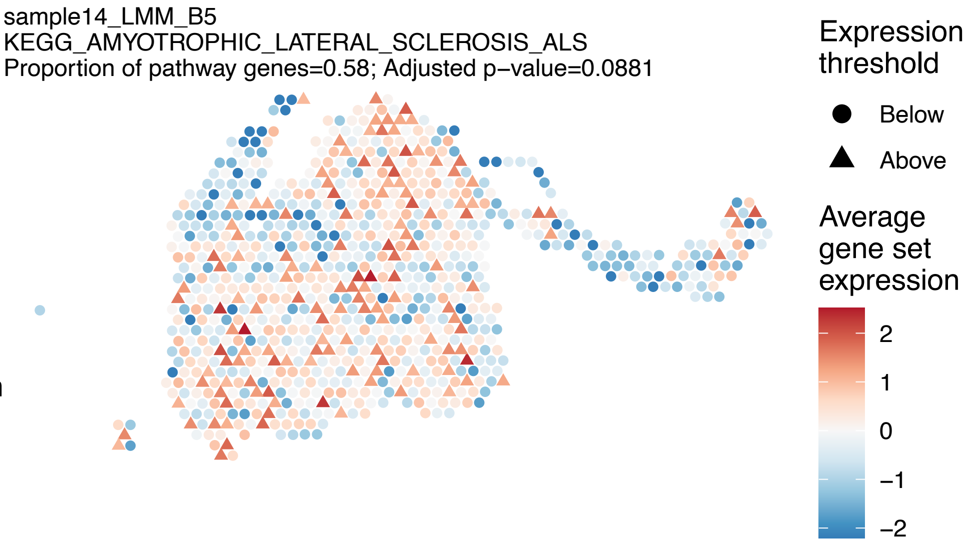
sample14_LMM_B5
KEGG_AMYOTROPHIC_LATERAL_SCLEROSIS_ALS
Proportion of pathway genes=0.58; Adjusted p−value=0.0881
Expression threshold
Below
Above
Average gene set expression
2
1
0
−1
−2

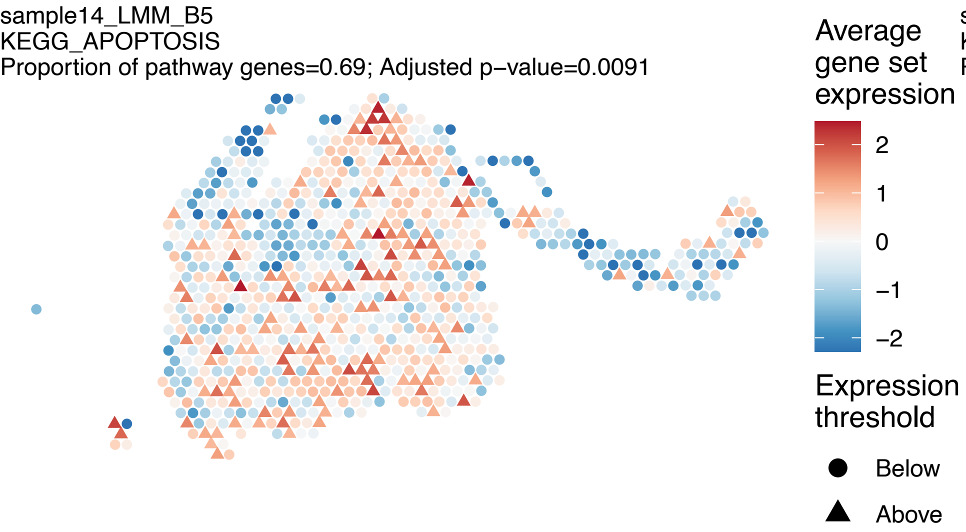
sample14_LMM_B5
KEGG_APOPTOSIS
Proportion of pathway genes=0.69; Adjusted p−value=0.0091
Average gene set expression
2
1
0
−1
−2
Expression threshold
Below
Above

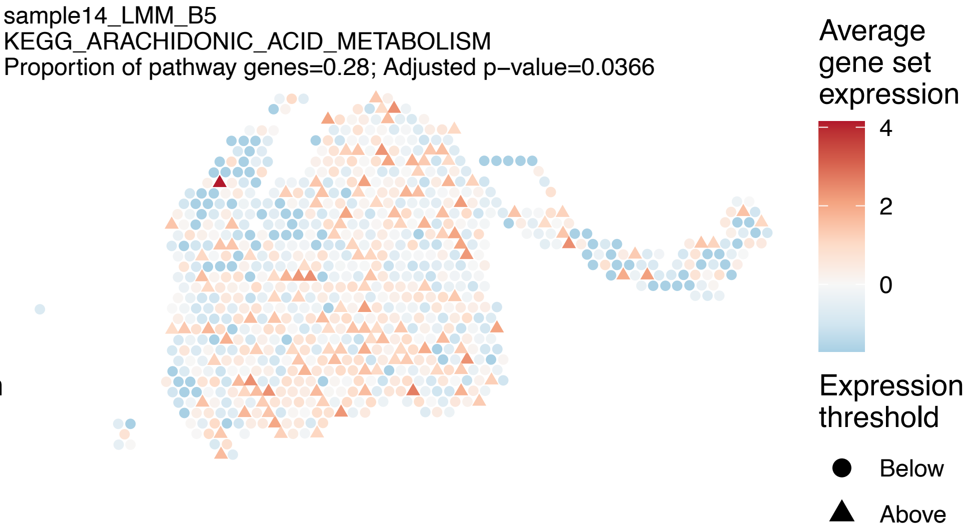
sample14_LMM_B5
KEGG_ARACHIDONIC_ACID_METABOLISM
Proportion of pathway genes=0.28; Adjusted p−value=0.0366
Average gene set expression
4
2
0
Expression threshold
Below
Above

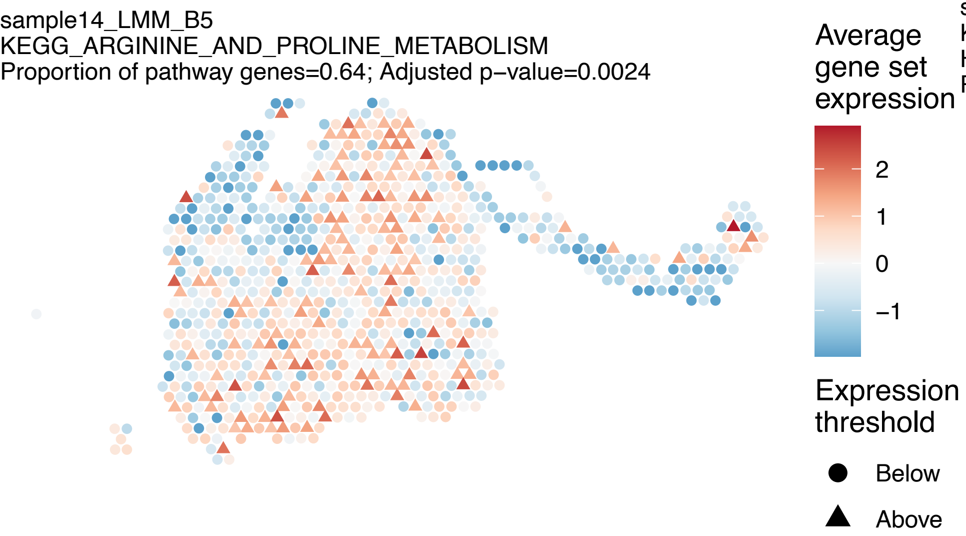
sample14_LMM_B5
KEGG_ARGININE_AND_PROLINE_METABOLISM
Proportion of pathway genes=0.64; Adjusted p−value=0.0024
Average gene set expression
2
1
0
−1
Expression threshold
Below
Above

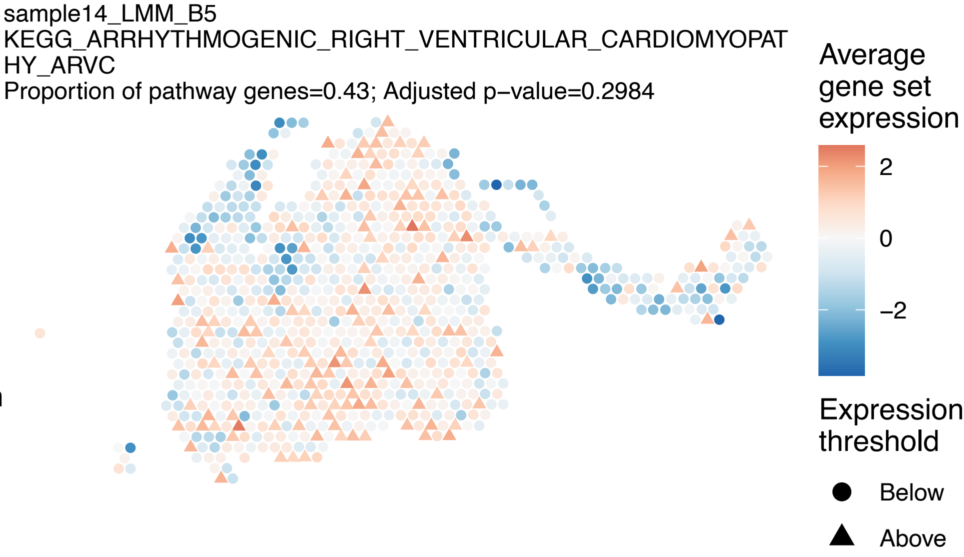
sample14_LMM_B5
KEGG_ARRHYTHMOGENIC_RIGHT_VENTRICULAR_CARDIOMYOPAT
HY_ARVC
Proportion of pathway genes=0.43; Adjusted p−value=0.2984
Average gene set expression
2
0
−2
Expression threshold
Below
Above

sample14_LMM_B5
KEGG_ASTHMA
Proportion of pathway genes=0.5; Adjusted p−value=0.5586

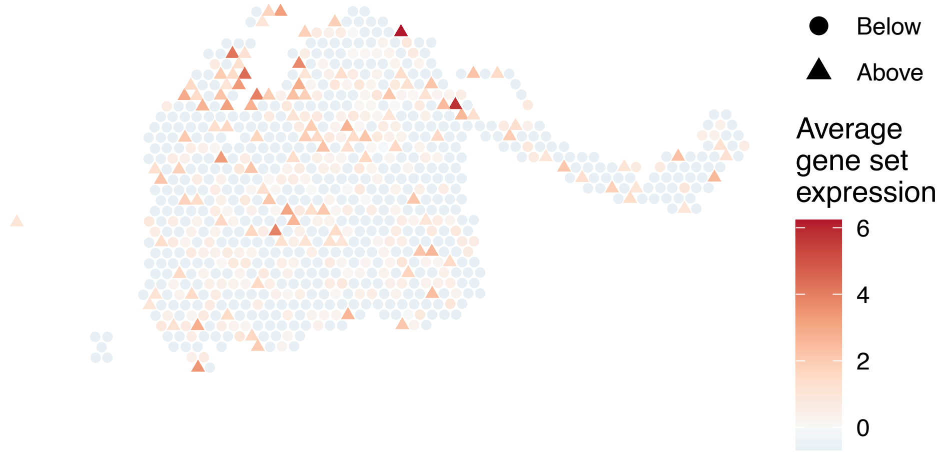


sample14_LMM_B5
KEGG_AUTOIMMUNE_THYROID_DISEASE
Proportion of pathway genes=0.55; Adjusted p−value=0.1734

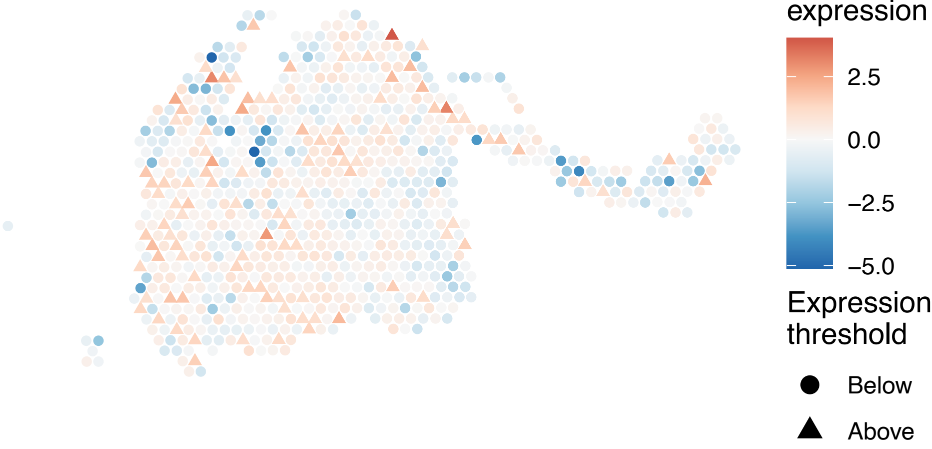


sample14_LMM_B5
KEGG_AXON_GUIDANCE
Proportion of pathway genes=0.62; Adjusted p−value=0

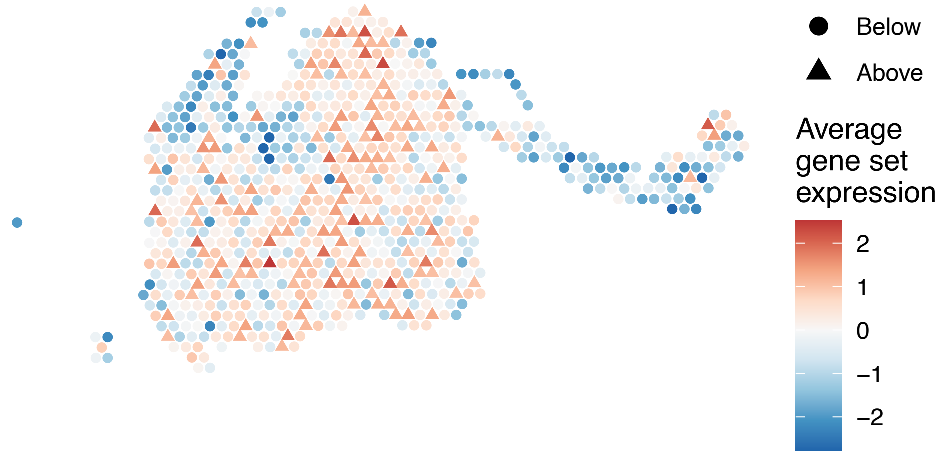


sample14_LMM_B5
KEGG_B_CELL_RECEPTOR_SIGNALING_PATHWAY
Proportion of pathway genes=0.75; Adjusted p−value=0.0024

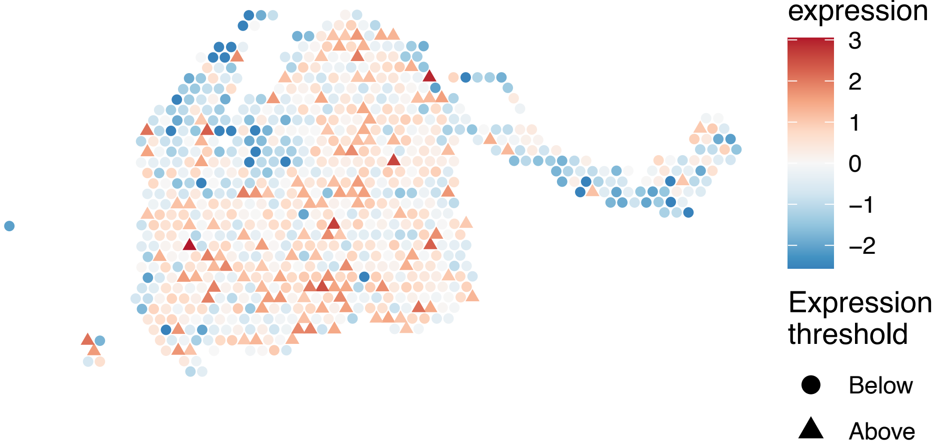

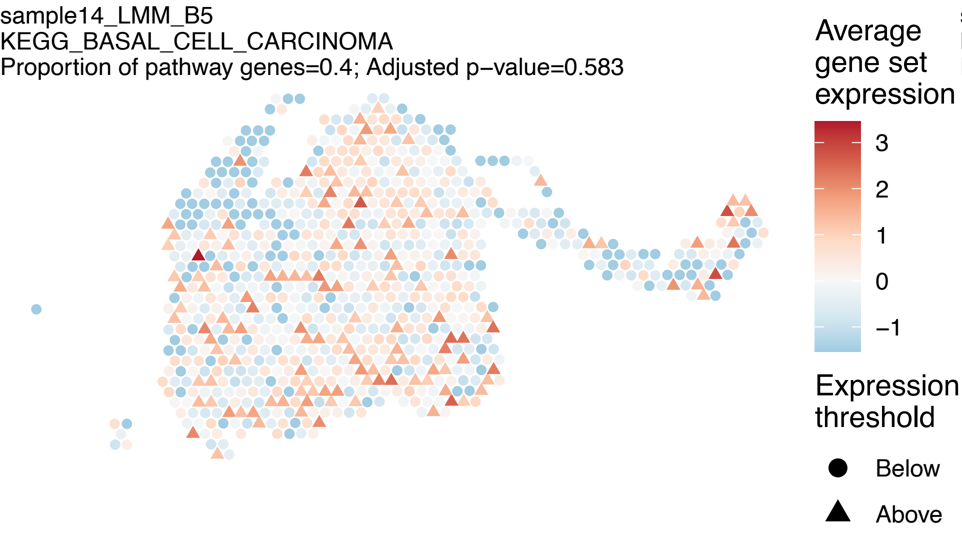
sample14_LMM_B5
KEGG_BASAL_CELL_CARCINOMA
Proportion of pathway genes=0.4; Adjusted p−value=0.583
Average gene set expression
3
2
1
0
−1
Expression threshold
Below
Above

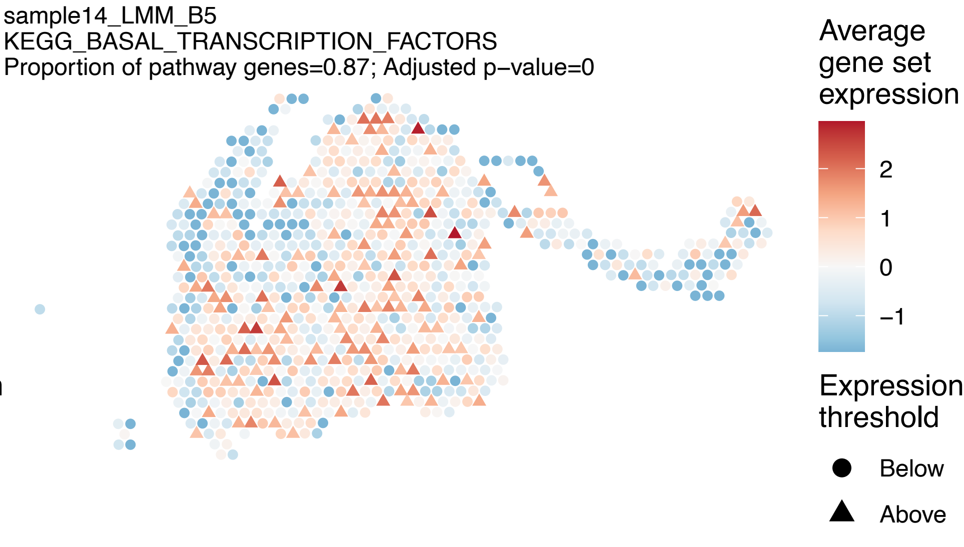
sample14_LMM_B5
KEGG_BASAL_TRANSCRIPTION_FACTORS
Proportion of pathway genes=0.87; Adjusted p−value=0
Average gene set expression
2
1
0
−1
Expression threshold
Below
Above

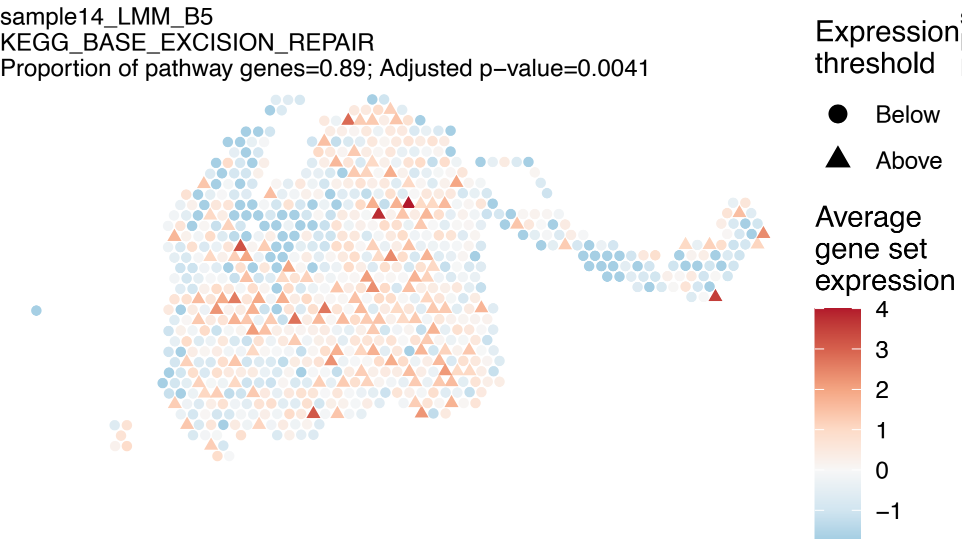
sample14_LMM_B5
KEGG_BASE_EXCISION_REPAIR
Proportion of pathway genes=0.89; Adjusted p−value=0.0041
Expression threshold
Below
Above
Average gene set expression
4
3
2
1
0
−1

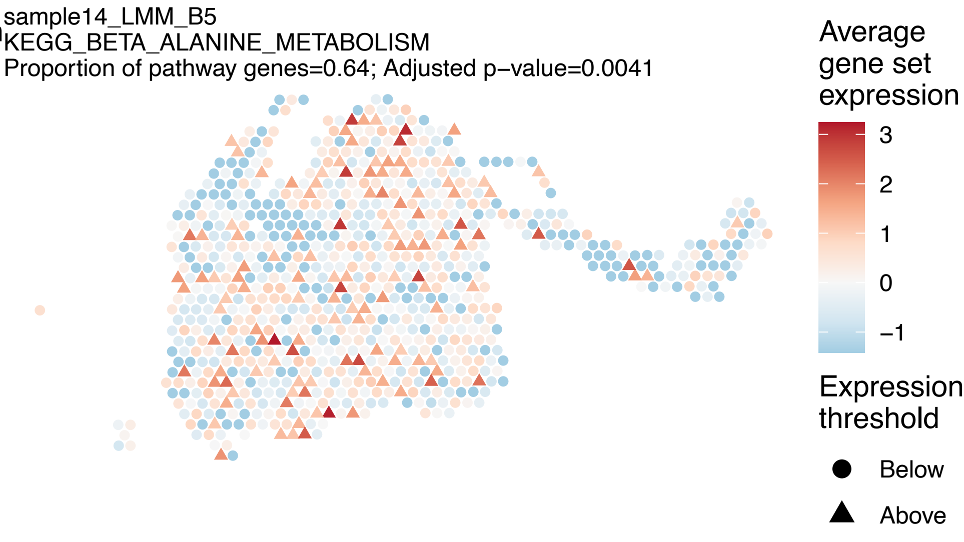
sample14_LMM_B5
KEGG_BETA_ALANINE_METABOLISM
Proportion of pathway genes=0.64; Adjusted p−value=0.0041
Average gene set expression
3
2
1
0
−1
Expression threshold
Below
Above

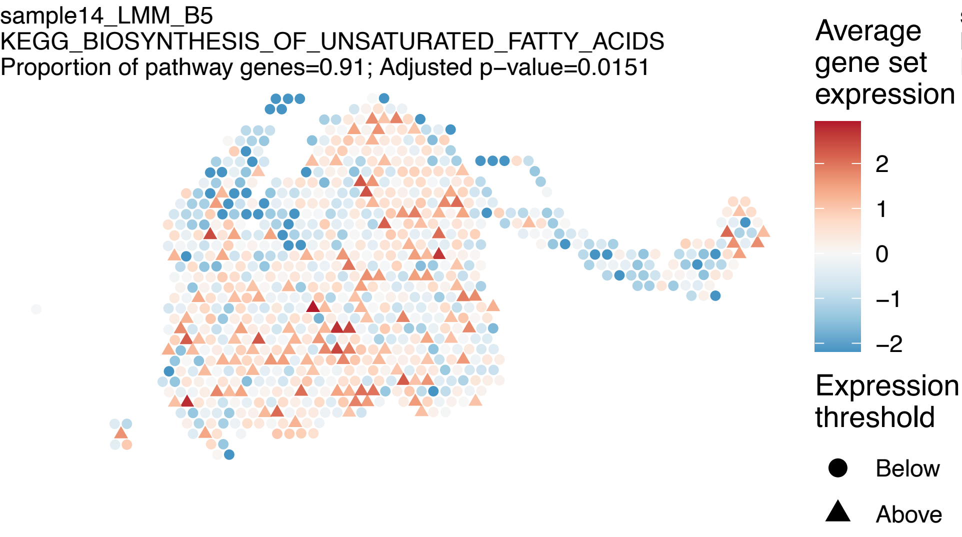
sample14_LMM_B5
KEGG_BIOSYNTHESIS_OF_UNSATURATED_FATTY_ACIDS
Proportion of pathway genes=0.91; Adjusted p−value=0.0151
Average gene set expression
2
1
0
−1
−2
Expression threshold
Below
Above

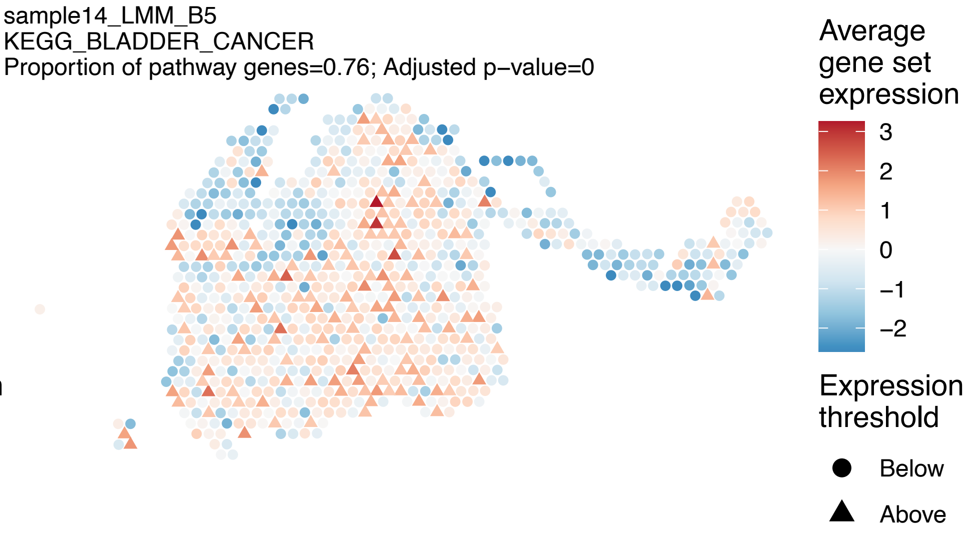
sample14_LMM_B5
KEGG_BLADDER_CANCER
Proportion of pathway genes=0.76; Adjusted p−value=0
Average gene set expression
3
2
1
0
−1
−2
Expression threshold
Below
Above

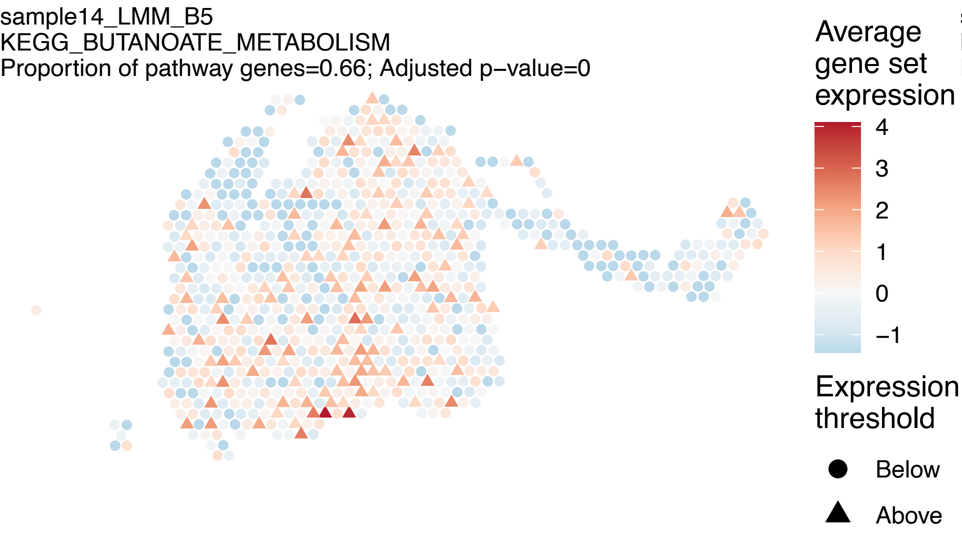
sample14_LMM_B5
KEGG_BUTANOATE_METABOLISM
Proportion of pathway genes=0.66; Adjusted p−value=0
Average gene set expression
4
3
2
1
0
−1
Expression threshold
Below
Above

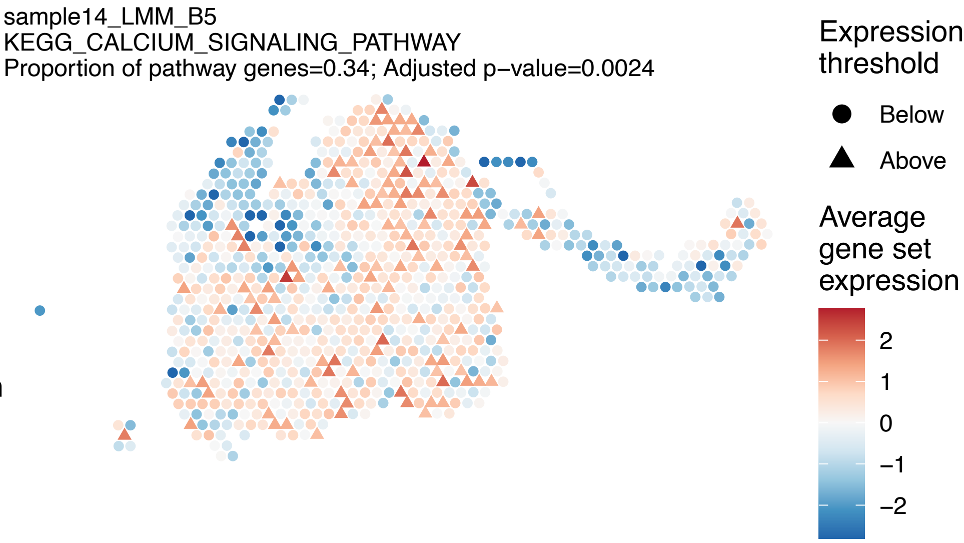
sample14_LMM_B5
KEGG_CALCIUM_SIGNALING_PATHWAY
Proportion of pathway genes=0.34; Adjusted p−value=0.0024
Expression threshold
Below
Above
Average gene set expression
2
1
0
−1
−2

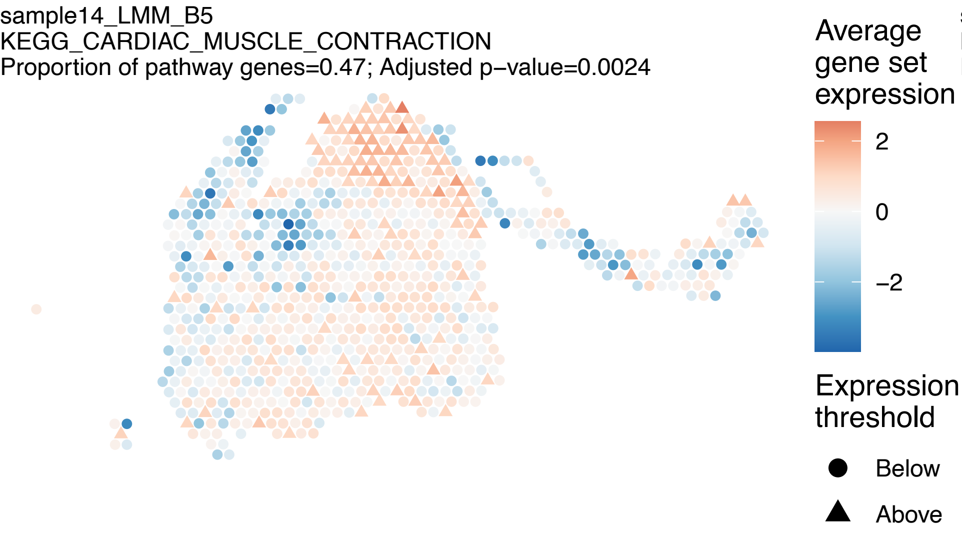
sample14_LMM_B5
KEGG_CARDIAC_MUSCLE_CONTRACTION
Proportion of pathway genes=0.47; Adjusted p−value=0.0024
Average gene set expression
2
0
−2
Expression threshold
Below
Above

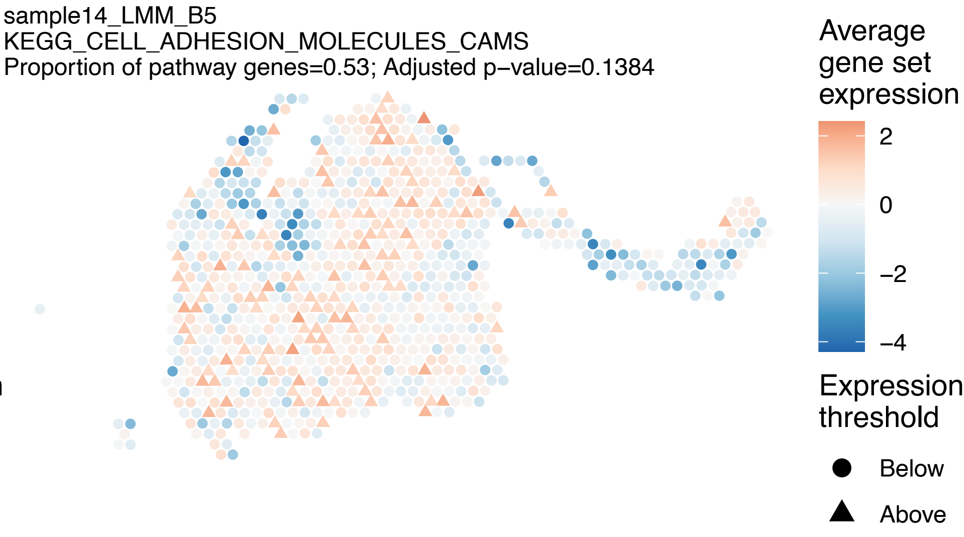
sample14_LMM_B5
KEGG_CELL_ADHESION_MOLECULES_CAMS
Proportion of pathway genes=0.53; Adjusted p−value=0.1384
Average gene set expression
2
0
−2
−4
Expression threshold
Below
Above

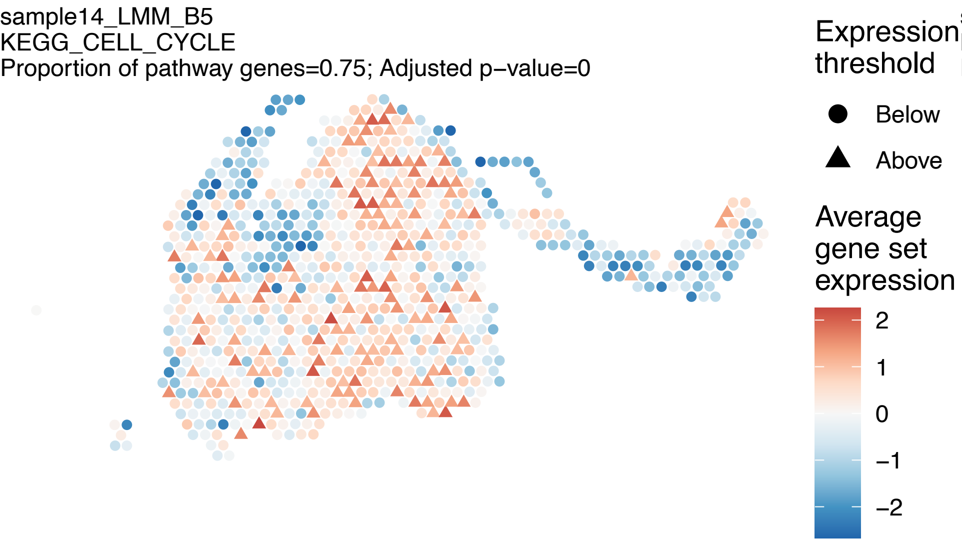
sample14_LMM_B5
KEGG_CELL_CYCLE
Proportion of pathway genes=0.75; Adjusted p−value=0
Expression threshold
Below
Above
Average gene set expression
2
1
0
−1
−2

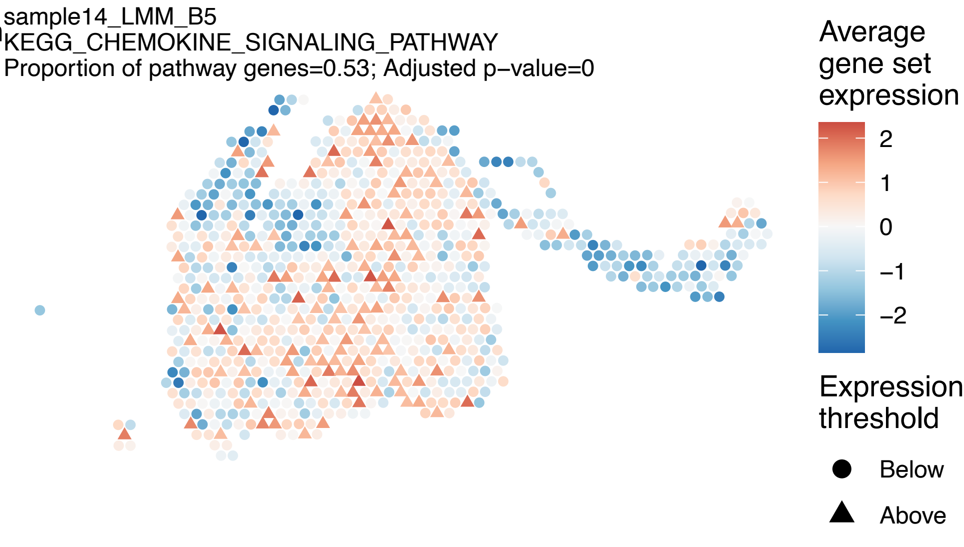
sample14_LMM_B5
KEGG_CHEMOKINE_SIGNALING_PATHWAY
Proportion of pathway genes=0.53; Adjusted p−value=0
Average gene set expression
2
1
0
−1
−2
Expression threshold
Below
Above

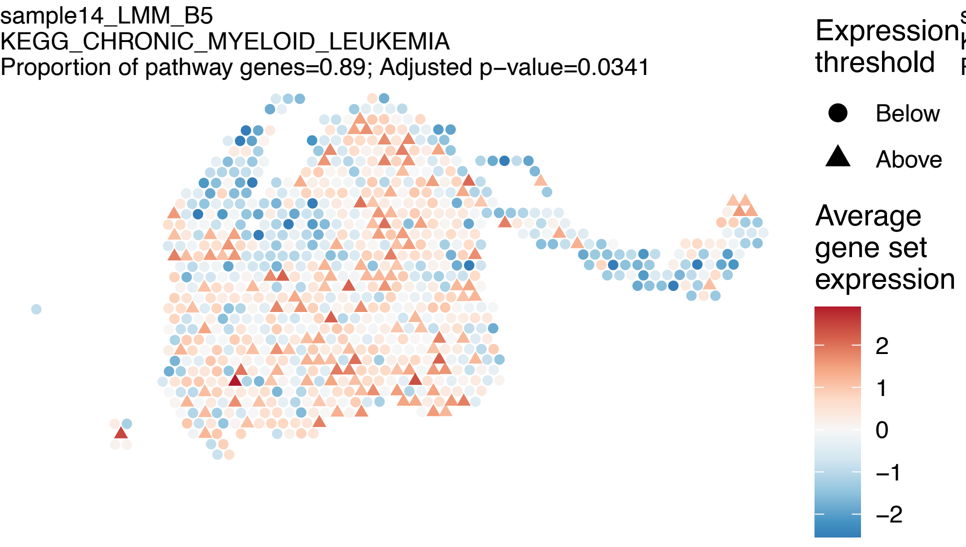
sample14_LMM_B5
KEGG_CHRONIC_MYELOID_LEUKEMIA
Proportion of pathway genes=0.89; Adjusted p−value=0.0341
Expression threshold
Below
Above
Average gene set expression
2
1
0
−1
−2

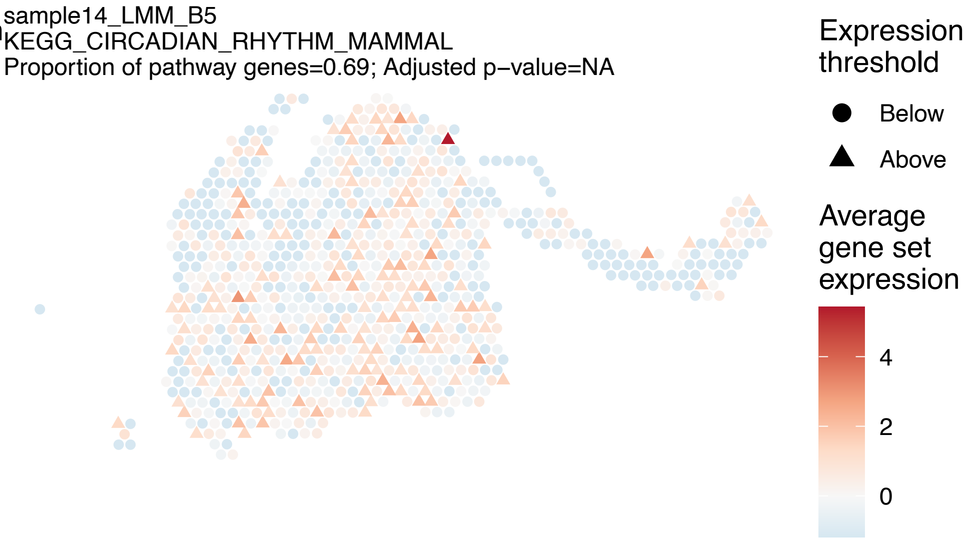
sample14_LMM_B5
KEGG_CIRCADIAN_RHYTHM_MAMMAL
Proportion of pathway genes=0.69; Adjusted p−value=NA
Expression threshold
Below
Above
Average gene set expression
4
2
0

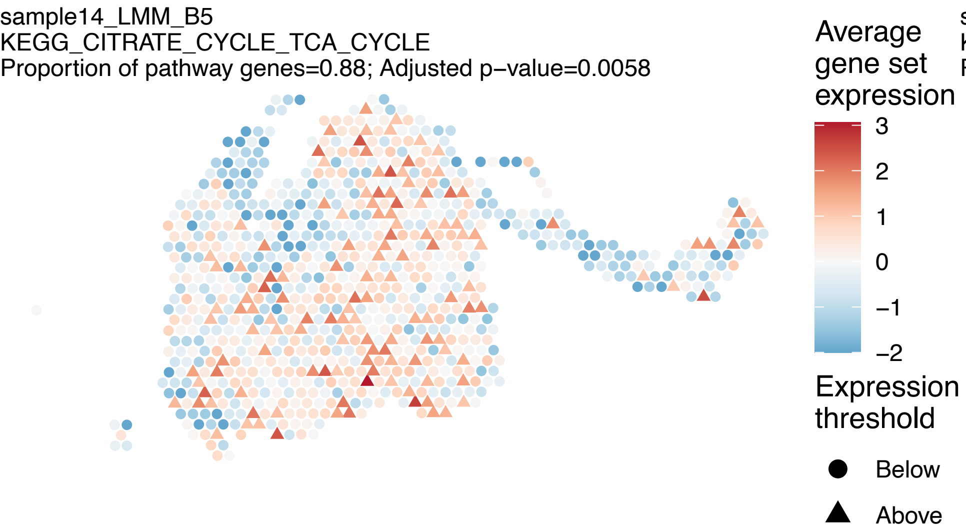
sample14_LMM_B5
KEGG_CITRATE_CYCLE_TCA_CYCLE
Proportion of pathway genes=0.88; Adjusted p−value=0.0058
Average gene set expression
3
2
1
0
−1
−2
Expression threshold
Below
Above

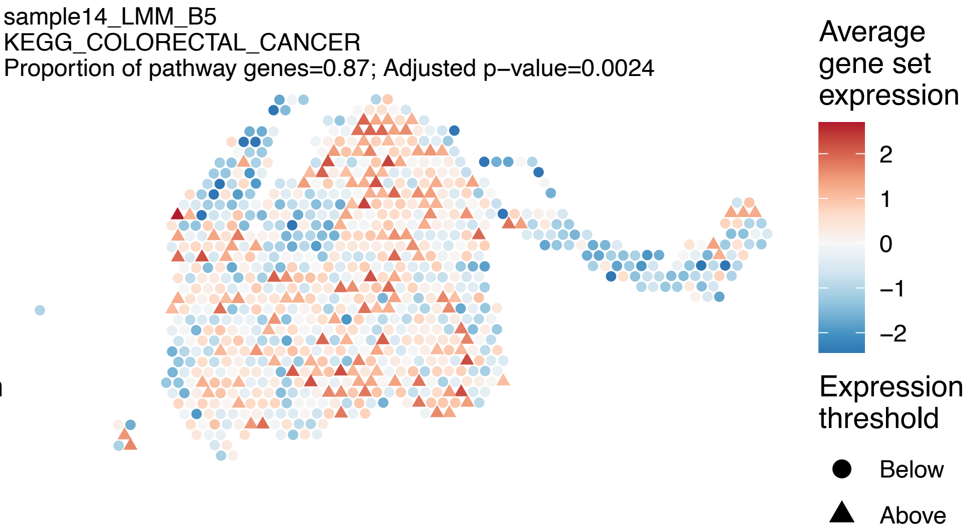
sample14_LMM_B5
KEGG_COLORECTAL_CANCER
Proportion of pathway genes=0.87; Adjusted p−value=0.0024
Average gene set expression
2
1
0
−1
−2
Expression threshold
Below
Above

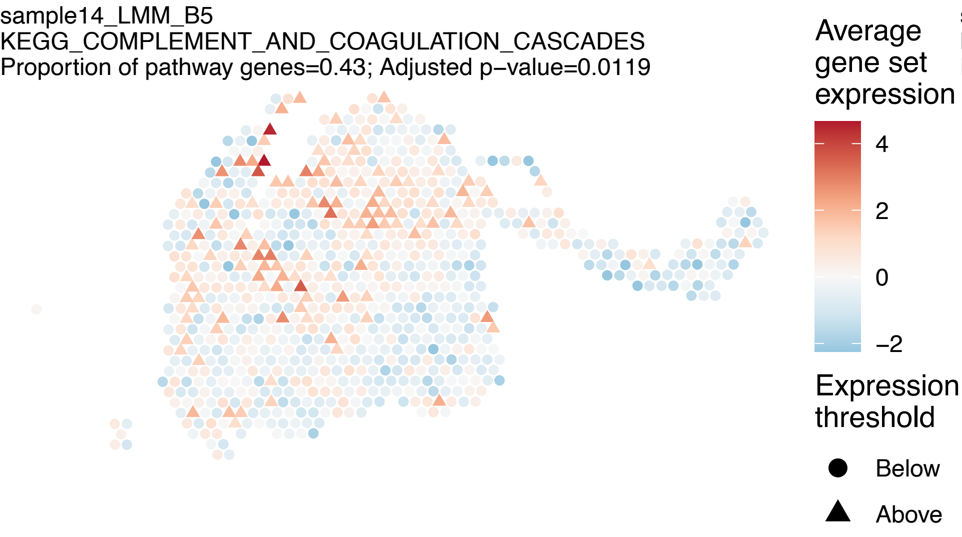
sample14_LMM_B5
KEGG_COMPLEMENT_AND_COAGULATION_CASCADES
Proportion of pathway genes=0.43; Adjusted p−value=0.0119
Average gene set expression
4
2
0
−2
Expression threshold
Below
Above

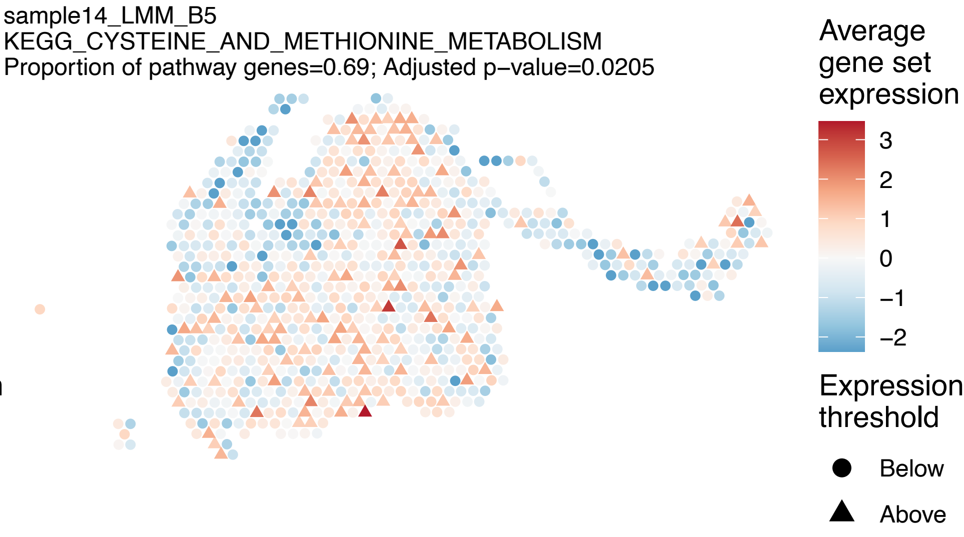
sample14_LMM_B5
KEGG_CYSTEINE_AND_METHIONINE_METABOLISM
Proportion of pathway genes=0.69; Adjusted p−value=0.0205
Average gene set expression
3
2
1
0
−1
−2
Expression threshold
Below
Above

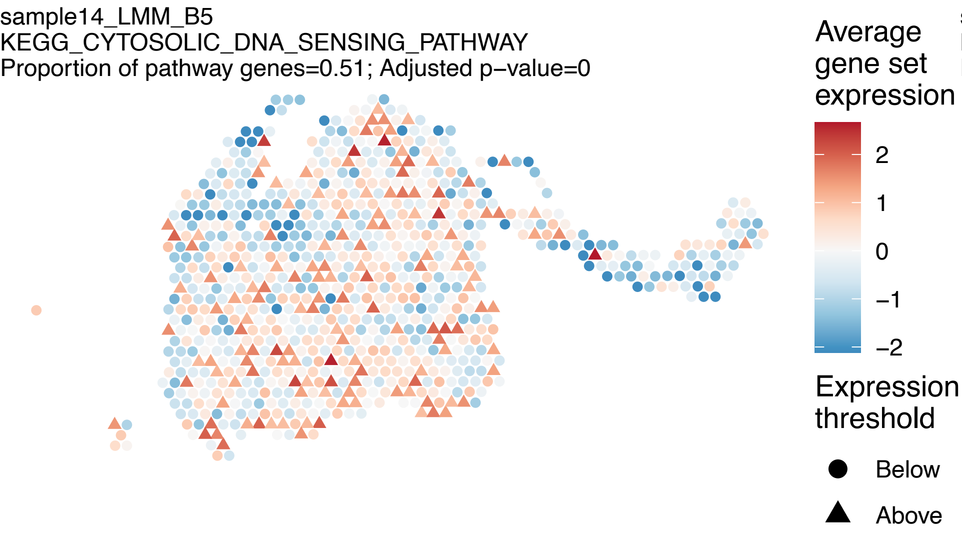
sample14_LMM_B5
KEGG_CYTOSOLIC_DNA_SENSING_PATHWAY
Proportion of pathway genes=0.51; Adjusted p−value=0
Average gene set expression
2
1
0
−1
−2
Expression threshold
Below
Above

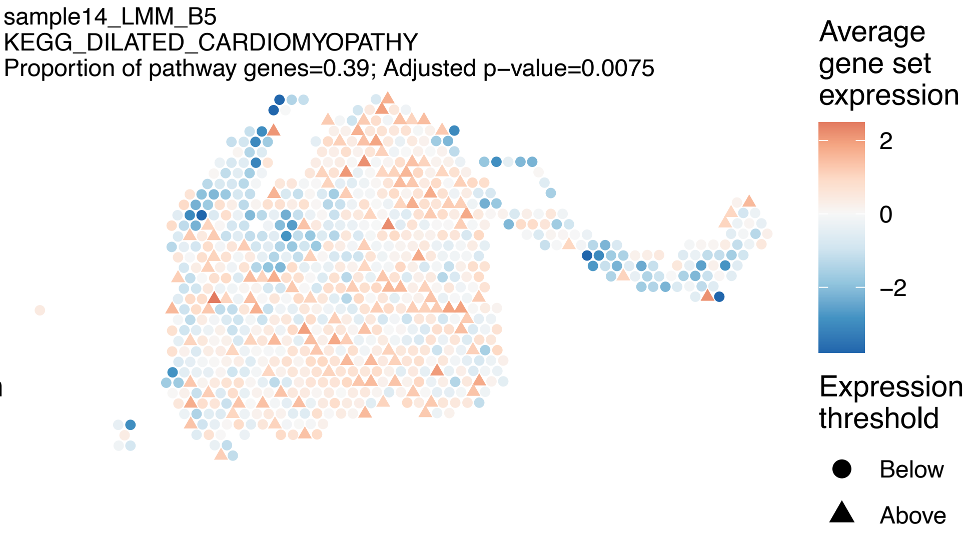
sample14_LMM_B5
KEGG_DILATED_CARDIOMYOPATHY
Proportion of pathway genes=0.39; Adjusted p−value=0.0075
Average gene set expression
2
0
−2
Expression threshold
Below
Above

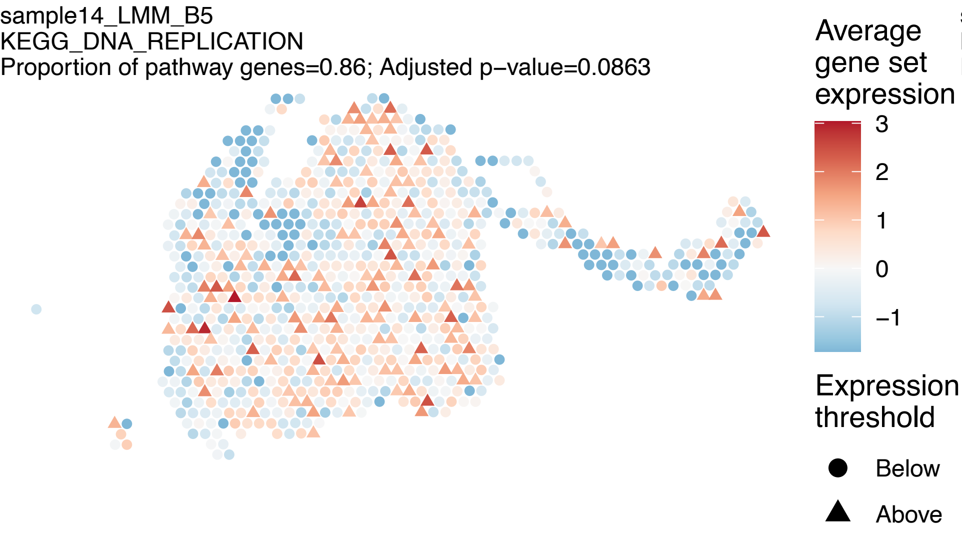
sample14_LMM_B5
KEGG_DNA_REPLICATION
Proportion of pathway genes=0.86; Adjusted p−value=0.0863
Average gene set expression
3
2
1
0
−1
Expression threshold
Below
Above

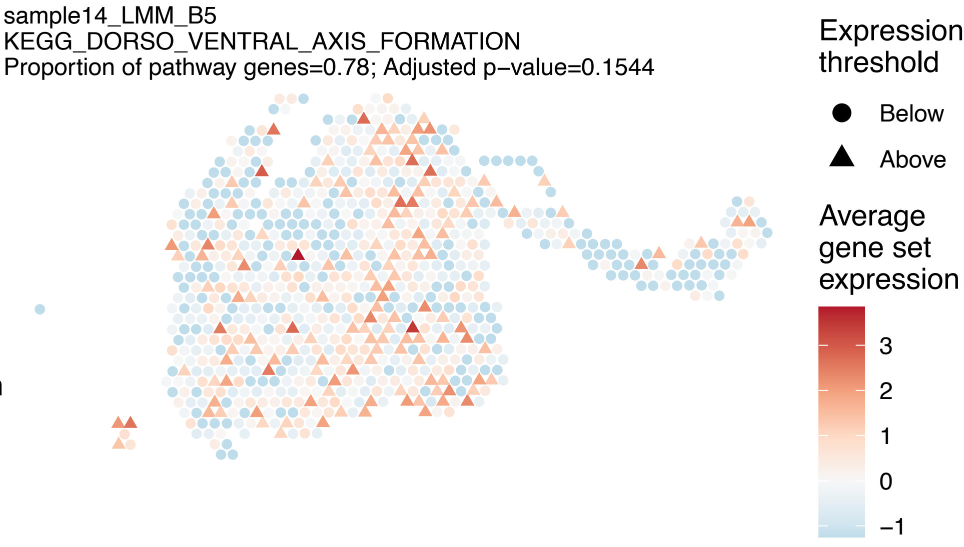
sample14_LMM_B5
KEGG_DORSO_VENTRAL_AXIS_FORMATION
Proportion of pathway genes=0.78; Adjusted p−value=0.1544
Expression threshold
Below
Above
Average gene set expression
3
2
1
0
−1

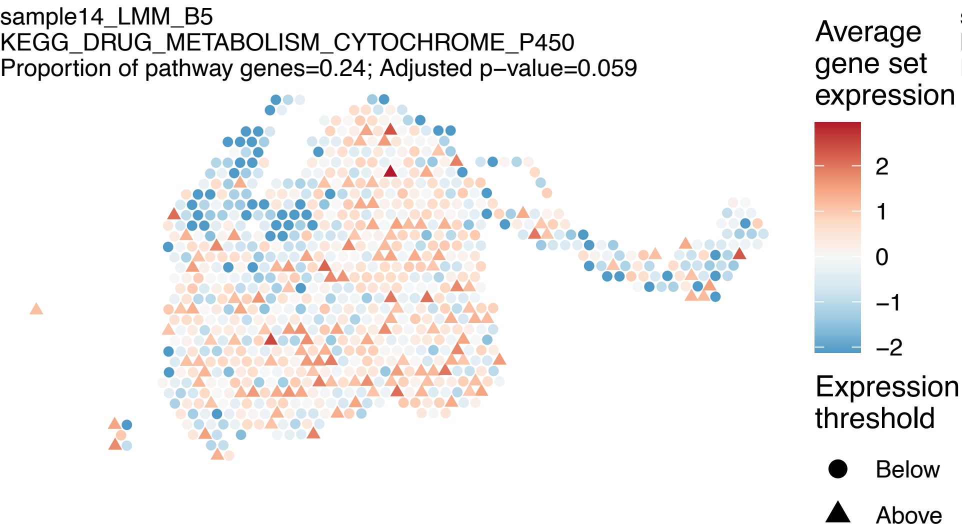
sample14_LMM_B5
KEGG_DRUG_METABOLISM_CYTOCHROME_P450
Proportion of pathway genes=0.24; Adjusted p−value=0.059
Average gene set expression
2
1
0
−1
−2
Expression threshold
Below
Above

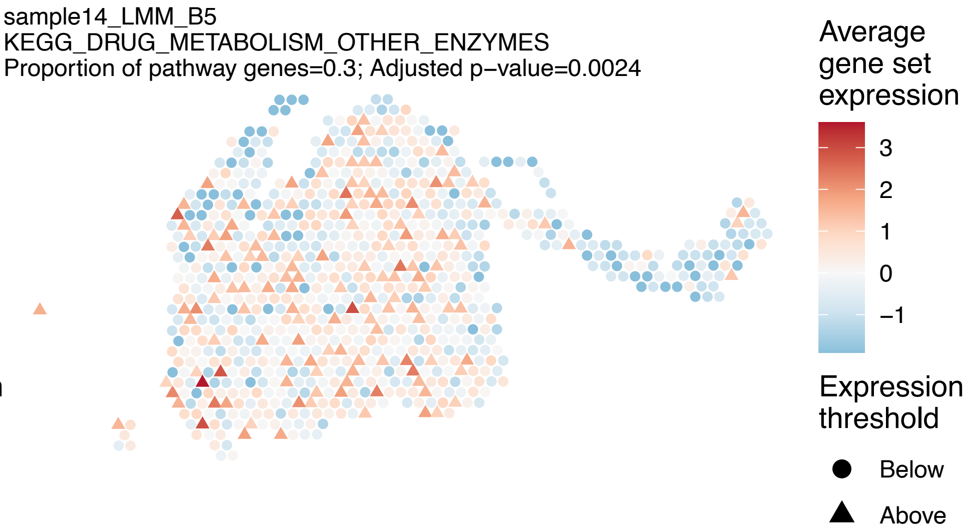
sample14_LMM_B5
KEGG_DRUG_METABOLISM_OTHER_ENZYMES
Proportion of pathway genes=0.3; Adjusted p−value=0.0024
Average gene set expression
3
2
1
0
−1
Expression threshold
Below
Above

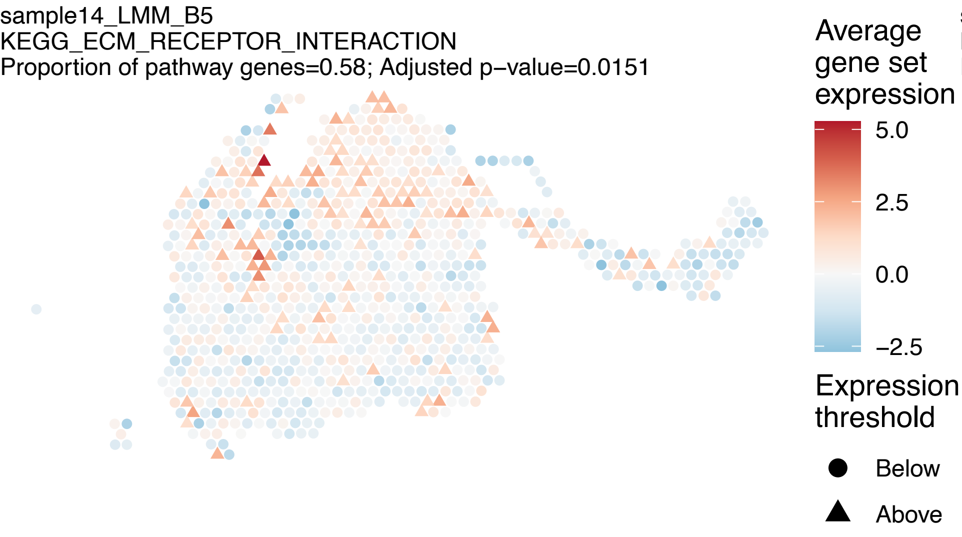
sample14_LMM_B5
KEGG_ECM_RECEPTOR_INTERACTION
Proportion of pathway genes=0.58; Adjusted p−value=0.0151
Average gene set expression
5.0
2.5
0.0
−2.5
Expression threshold
Below
Above

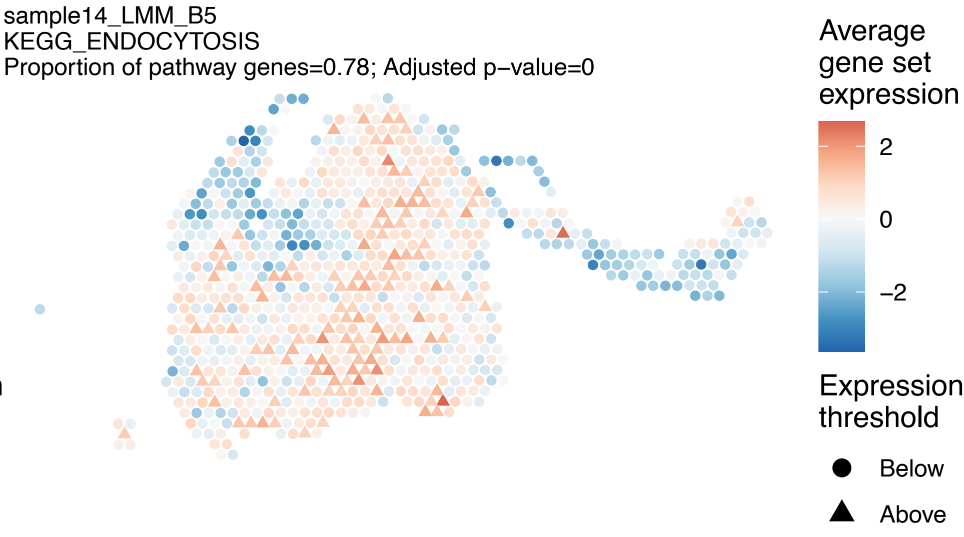
sample14_LMM_B5
KEGG_ENDOCYTOSIS
Proportion of pathway genes=0.78; Adjusted p−value=0
Average gene set expression
2
0
−2
Expression threshold
Below
Above

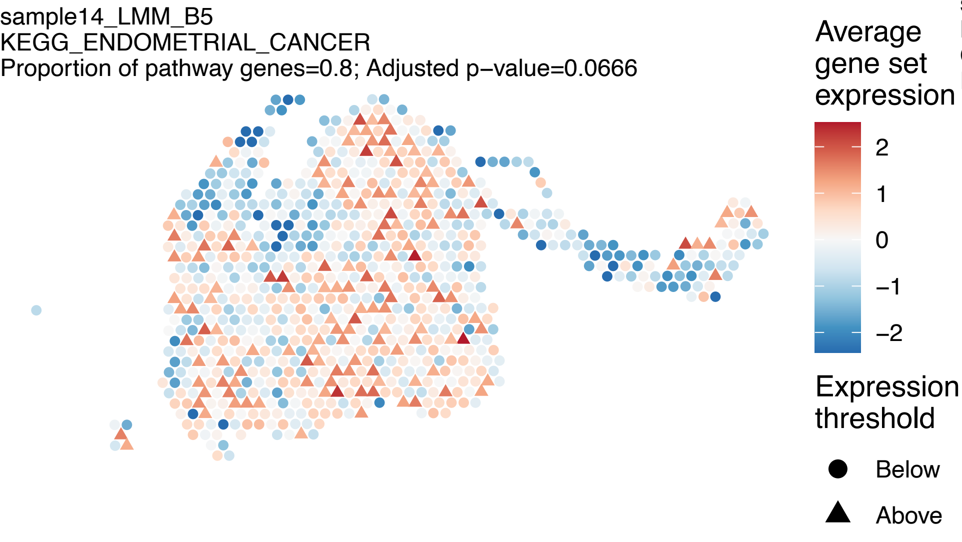
sample14_LMM_B5
KEGG_ENDOMETRIAL_CANCER
Proportion of pathway genes=0.8; Adjusted p−value=0.0666
Average gene set expression
2
1
0
−1
−2
Expression threshold
Below
Above

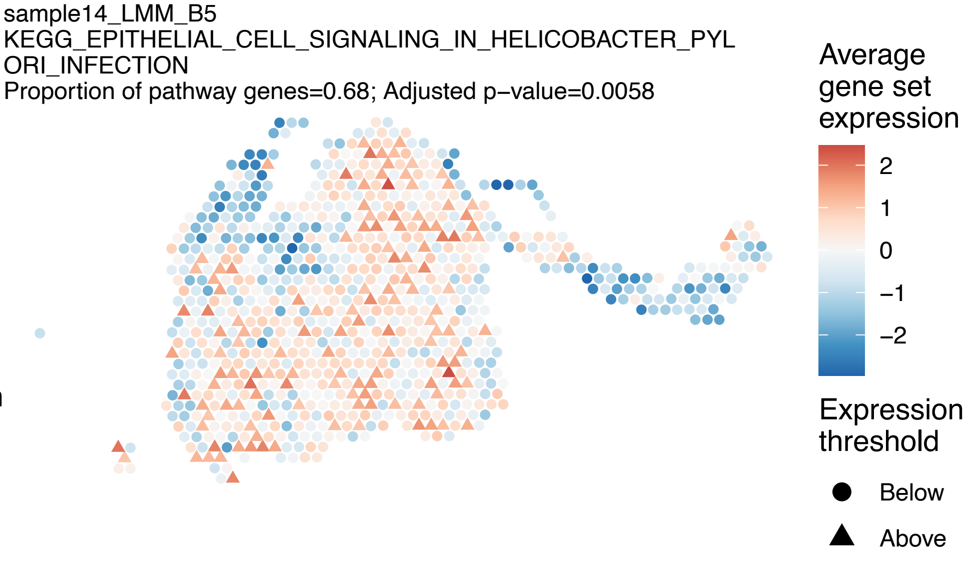
sample14_LMM_B5
KEGG_EPITHELIAL_CELL_SIGNALING_IN_HELICOBACTER_PYLORI_INFECTION
Proportion of pathway genes=0.68; Adjusted p−value=0.0058
Average gene set expression
2
1
0
−1
−2
Expression threshold
Below
Above

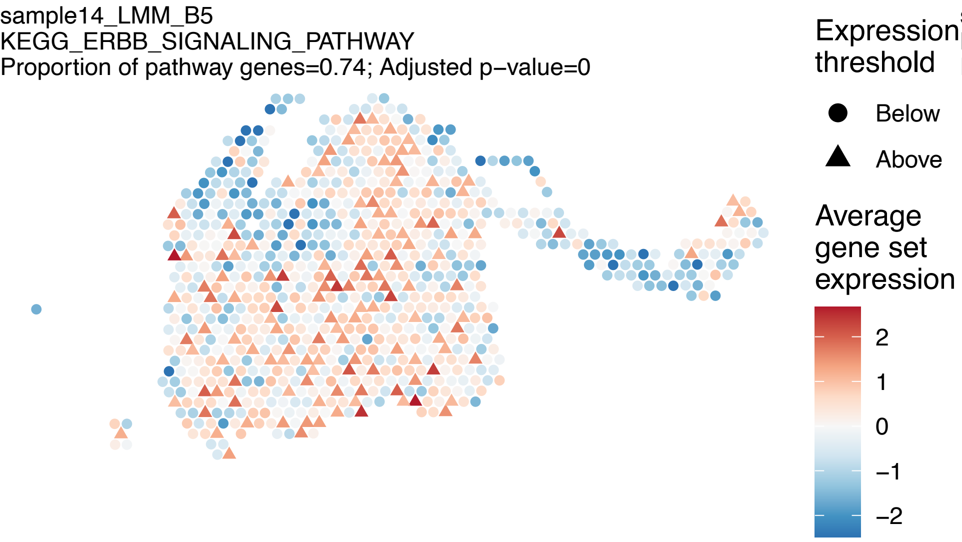

sample14_LMM_B5
KEGG_ERBB_SIGNALING_PATHWAY
Proportion of pathway genes=0.74; Adjusted p−value=0
Expression threshold
Below
Above
Average gene set expression
2
1
0
−1
−2


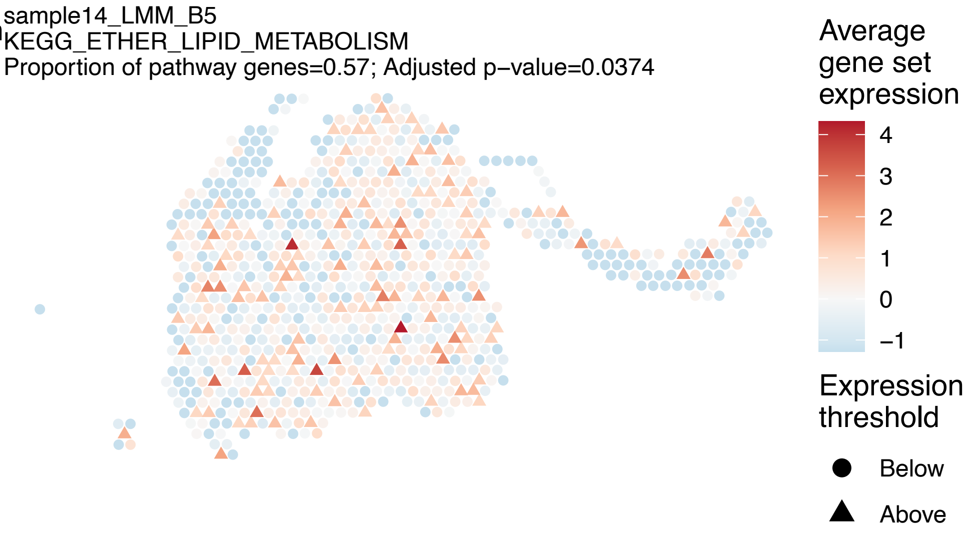

sample14_LMM_B5
KEGG_ETHER_LIPID_METABOLISM
Proportion of pathway genes=0.57; Adjusted p−value=0.0374
Average gene set expression
4
3
2
1
0
−1
Expression threshold
Below
Above


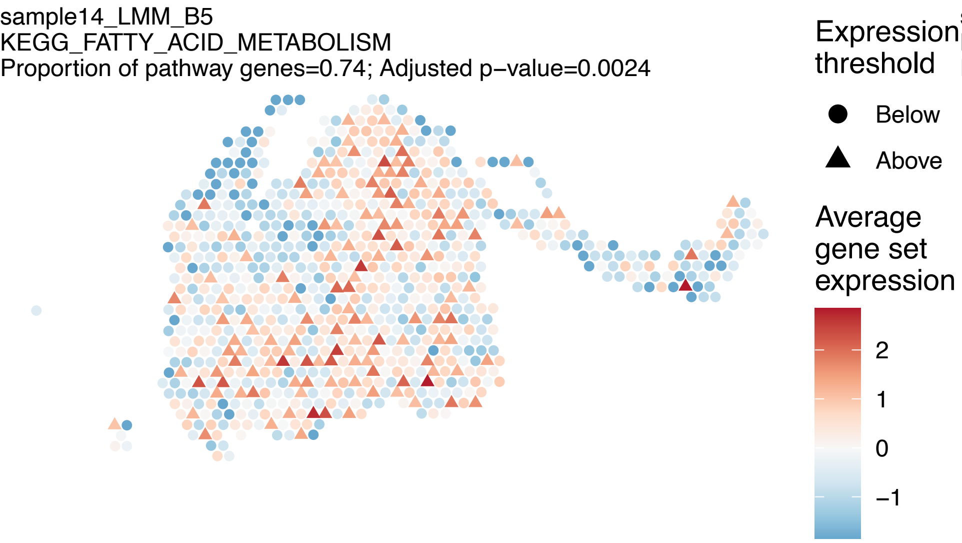

sample14_LMM_B5
KEGG_FATTY_ACID_METABOLISM
Proportion of pathway genes=0.74; Adjusted p−value=0.0024
Expression threshold
Below
Above
Average gene set expression
2
1
0
−1


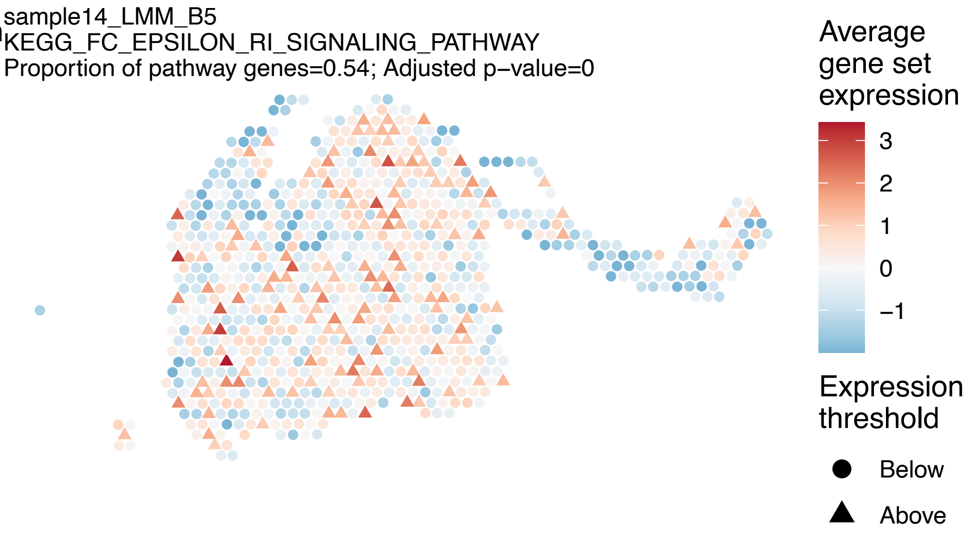

sample14_LMM_B5
KEGG_FC_EPSILON_RI_SIGNALING_PATHWAY
Proportion of pathway genes=0.54; Adjusted p−value=0
Average gene set expression
3
2
1
0
−1
Expression threshold
Below
Above

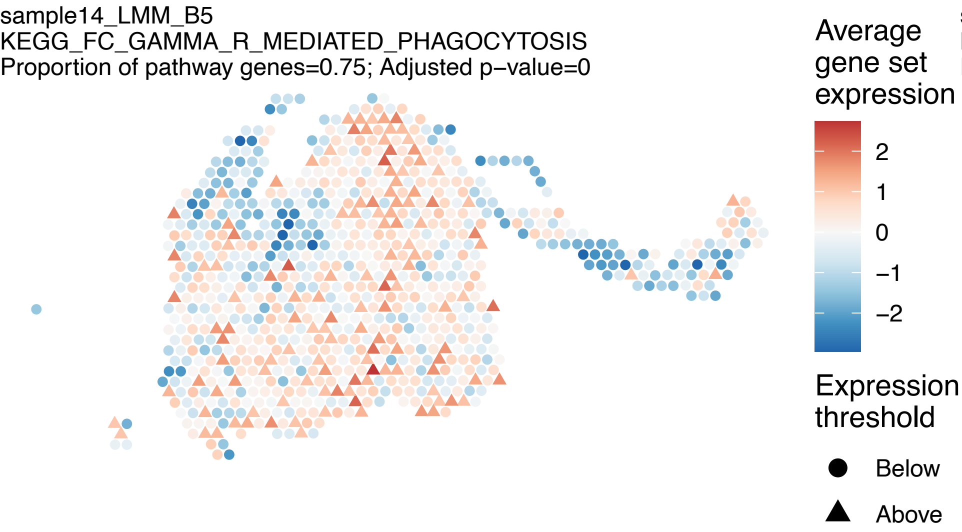
sample14_LMM_B5
KEGG_FC_GAMMA_R_MEDIATED_PHAGOCYTOSIS
Proportion of pathway genes=0.75; Adjusted p−value=0
Average gene set expression
2
1
0
−1
−2
Expression threshold
Below
Above

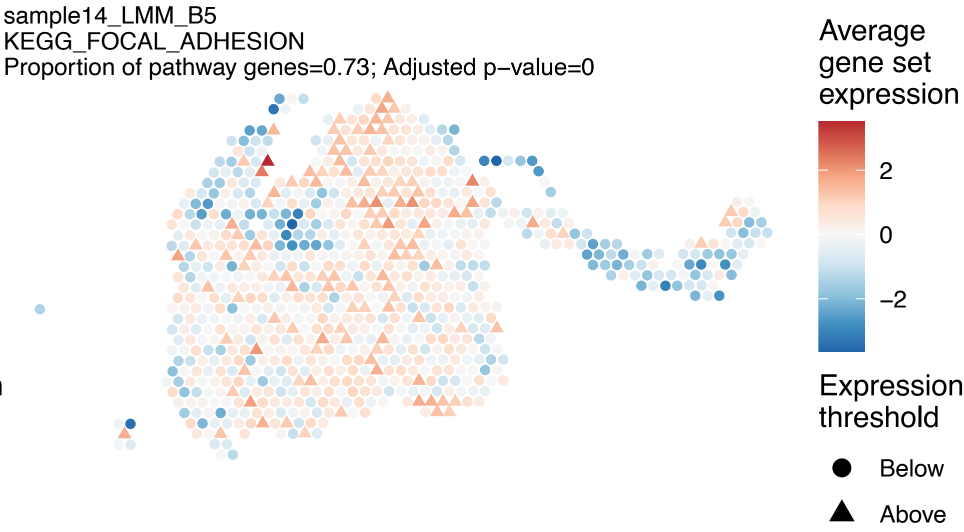
sample14_LMM_B5
KEGG_FOCAL_ADHESION
Proportion of pathway genes=0.73; Adjusted p−value=0
Average gene set expression
2
0
−2
Expression threshold
Below
Above

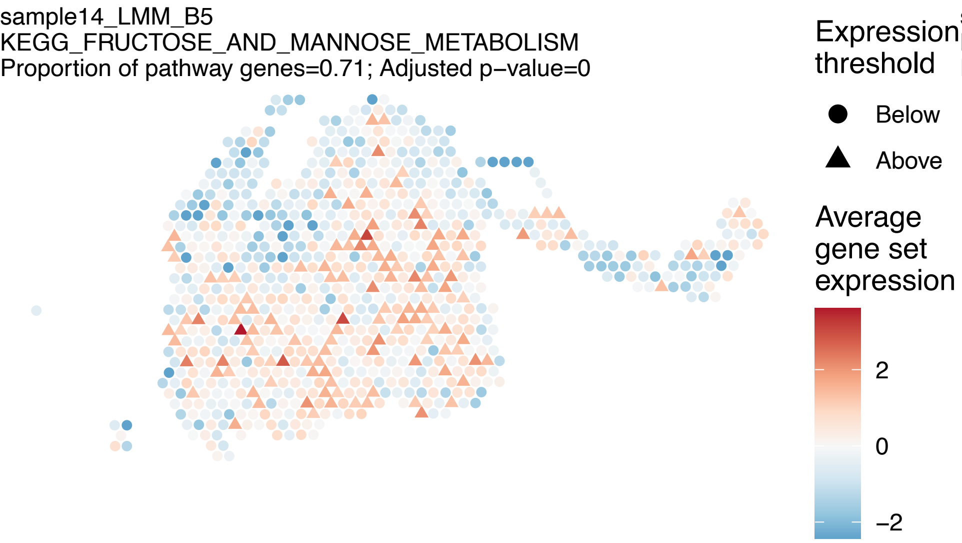
sample14_LMM_B5
KEGG_FRUCTOSE_AND_MANNOSE_METABOLISM
Proportion of pathway genes=0.71; Adjusted p−value=0
Expression threshold
Below
Above
Average gene set expression
2
0
−2

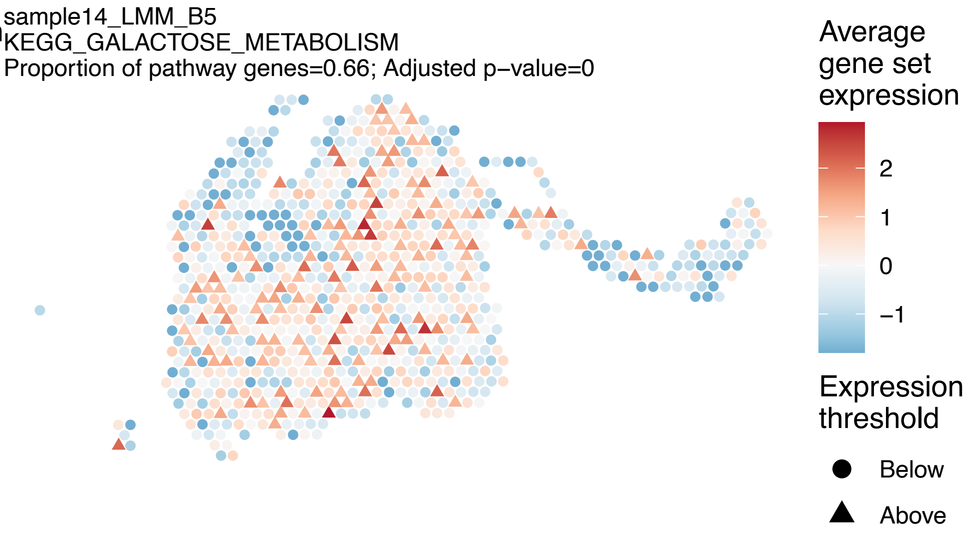
sample14_LMM_B5
KEGG_GALACTOSE_METABOLISM
Proportion of pathway genes=0.66; Adjusted p−value=0
Average gene set expression
2
1
0
−1
Expression threshold
Below
Above

sample14_LMM_B5
KEGG_GAP_JUNCTION
Proportion of pathway genes=0.61; Adjusted p−value=0.5714

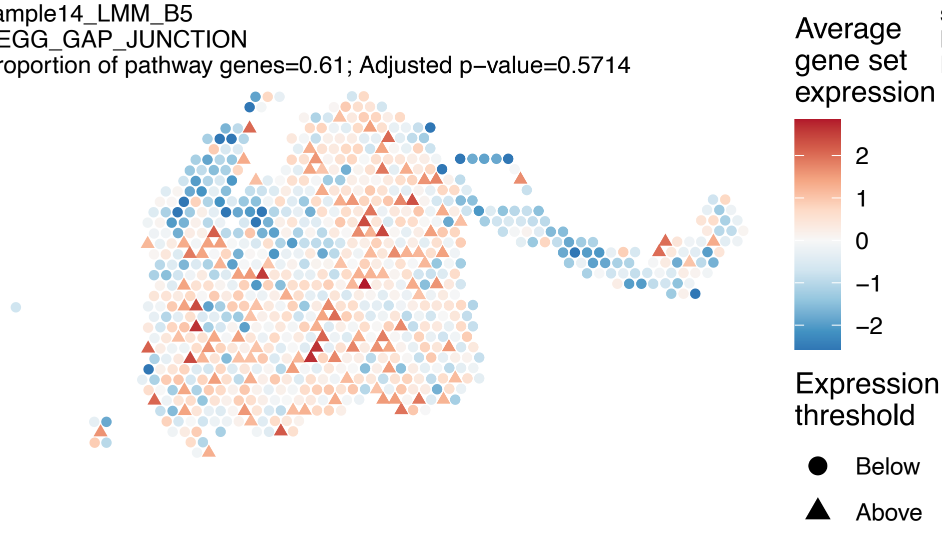


sample14_LMM_B5
KEGG_GLIOMA
Proportion of pathway genes=0.7; Adjusted p−value=0.0489

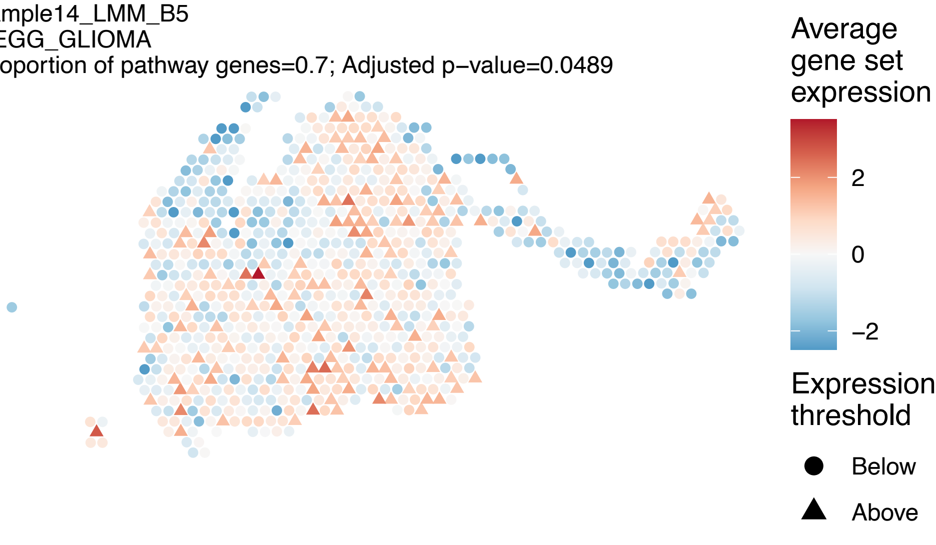


sample14_LMM_B5
KEGG_GLUTATHIONE_METABOLISM
Proportion of pathway genes=0.6; Adjusted p−value=0

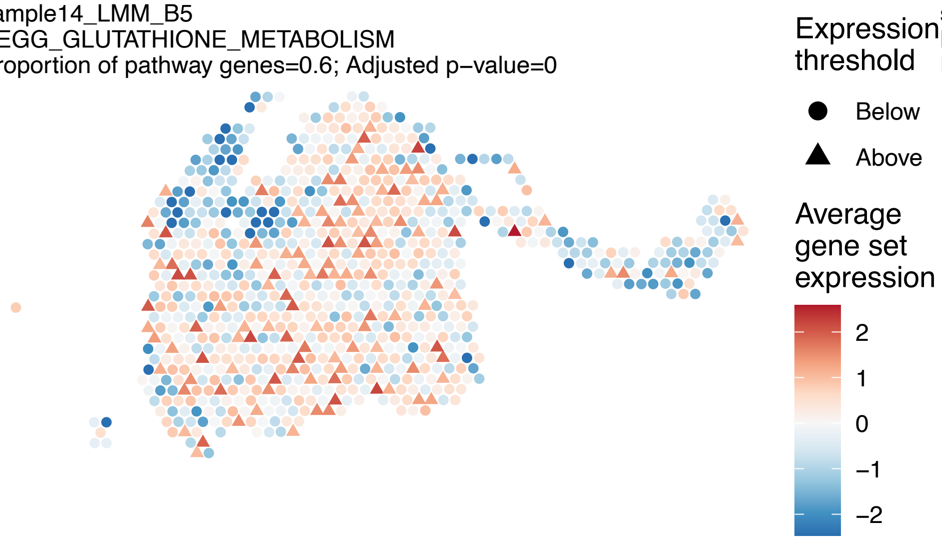


sample14_LMM_B5
KEGG_GLYCEROLIPID_METABOLISM
Proportion of pathway genes=0.73; Adjusted p−value=0.0166

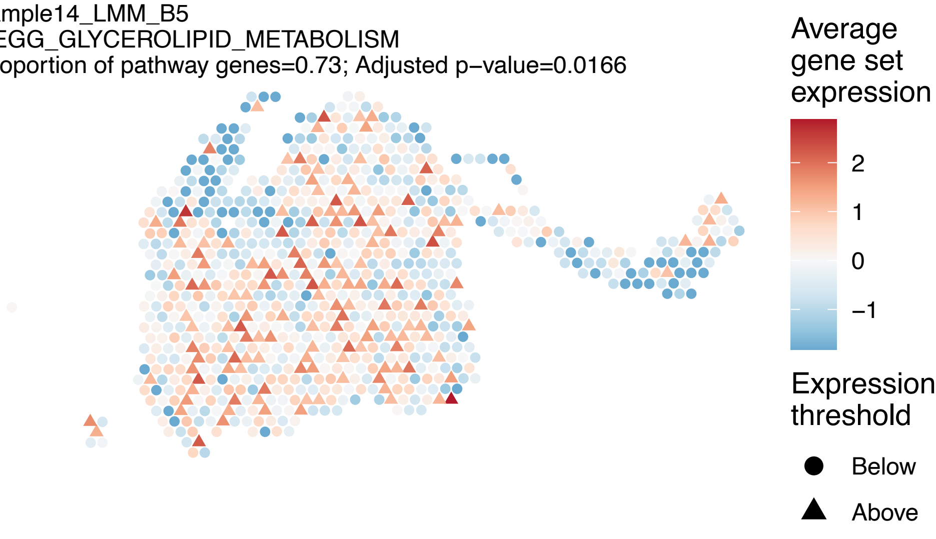

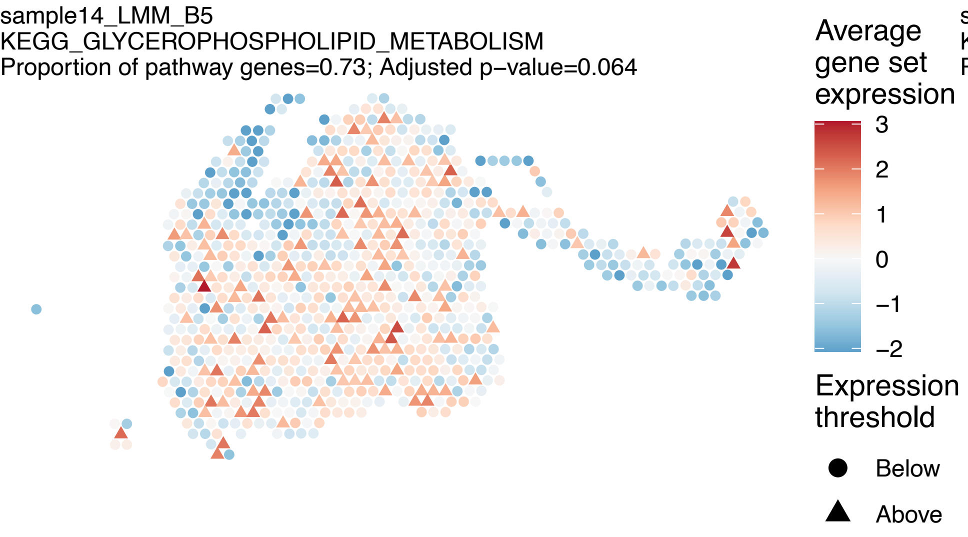
sample14_LMM_B5
KEGG_GLYCEROPHOSPHOLIPID_METABOLISM
Proportion of pathway genes=0.73; Adjusted p−value=0.064
Average gene set expression
3
2
1
0
−1
−2
Expression threshold
Below
Above

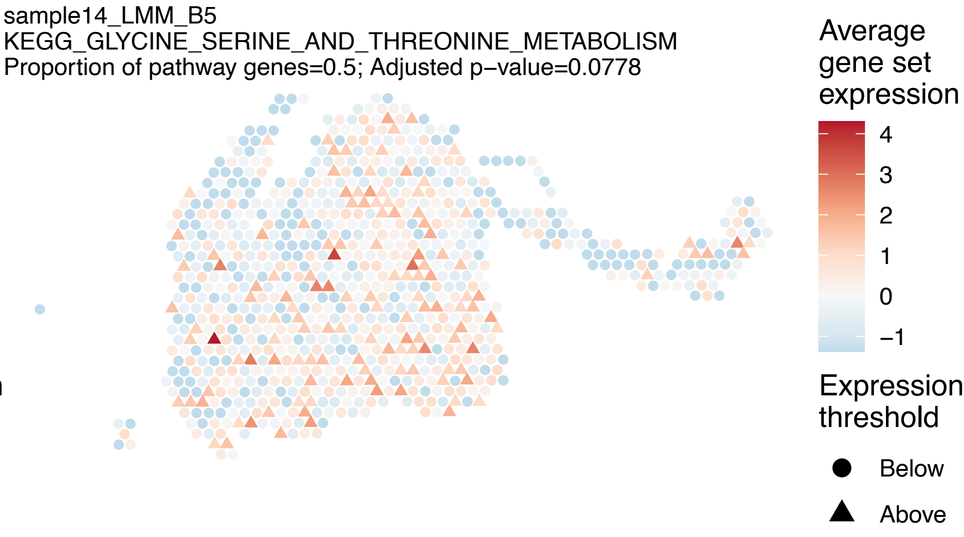
sample14_LMM_B5
KEGG_GLYCINE_SERINE_AND_THREONINE_METABOLISM
Proportion of pathway genes=0.5; Adjusted p−value=0.0778
Average gene set expression
4
3
2
1
0
−1
Expression threshold
Below
Above

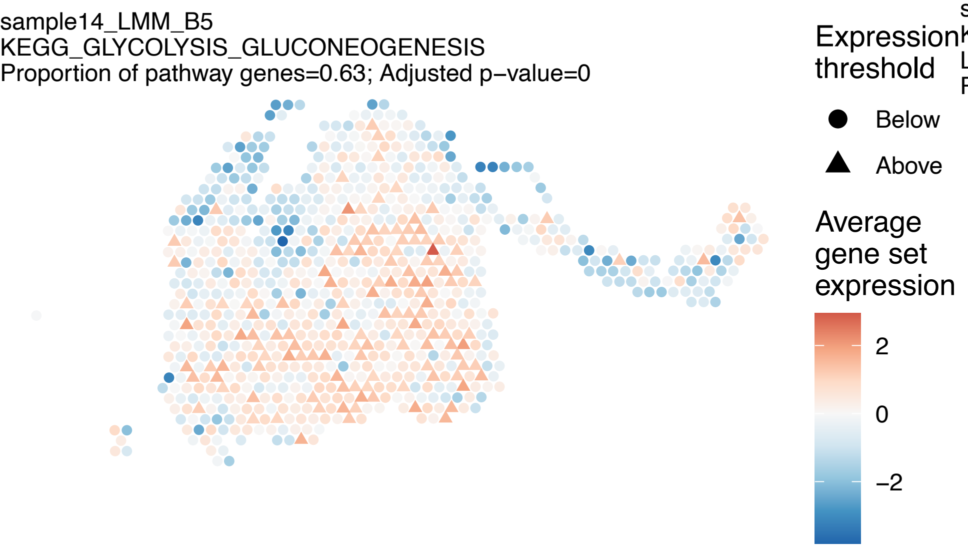
sample14_LMM_B5
KEGG_GLYCOLYSIS_GLUCONEOGENESIS
Proportion of pathway genes=0.63; Adjusted p−value=0
Expression threshold
Below
Above
Average gene set expression
2
0
−2

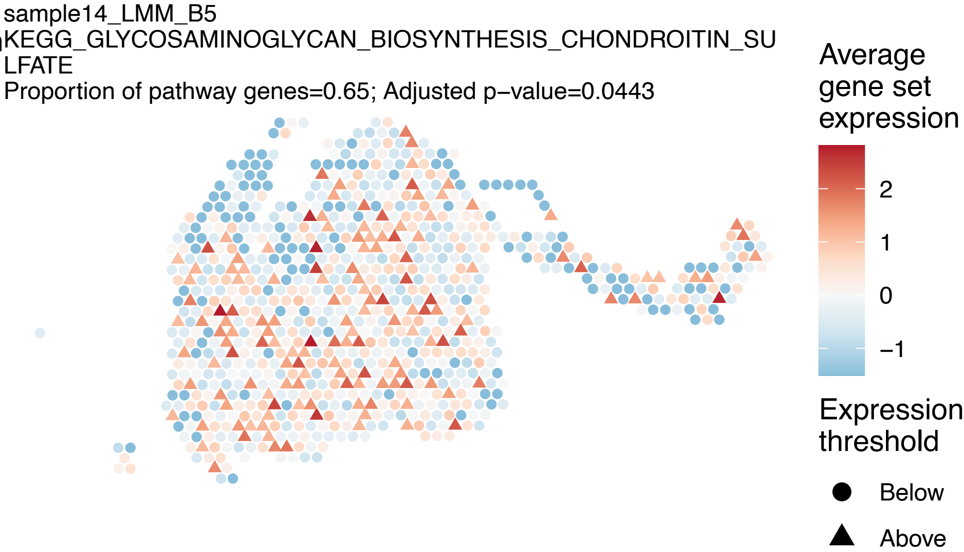
sample14_LMM_B5
KEGG_GLYCOSAMINOGLYCAN_BIOSYNTHESIS_CHONDROITIN_SU
LFATE
Proportion of pathway genes=0.65; Adjusted p−value=0.0443
Average gene set expression
2
1
0
−1
Expression threshold
Below
Above

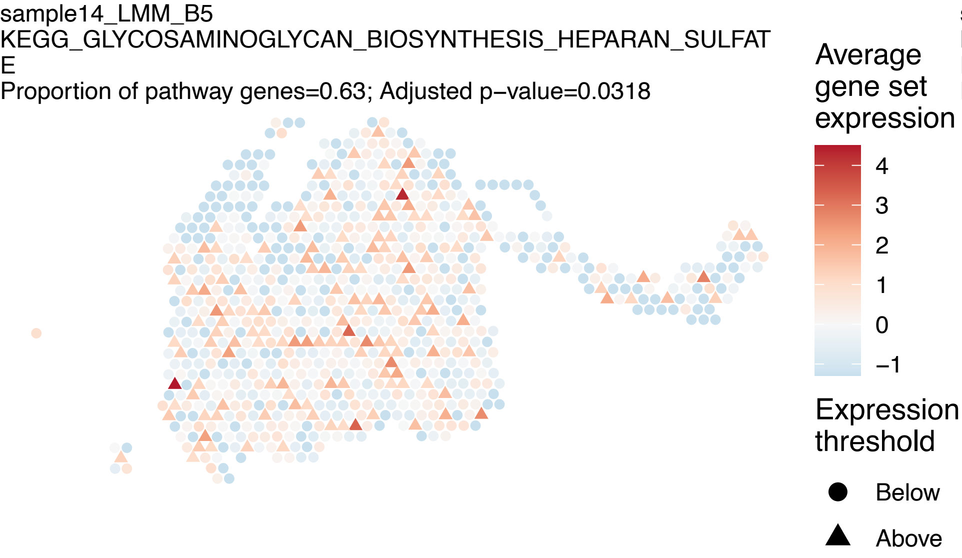
sample14_LMM_B5
KEGG_GLYCOSAMINOGLYCAN_BIOSYNTHESIS_HEPARAN_SULFAT
E
Proportion of pathway genes=0.63; Adjusted p−value=0.0318
Average gene set expression
4
3
2
1
0
−1
Expression threshold
Below
Above

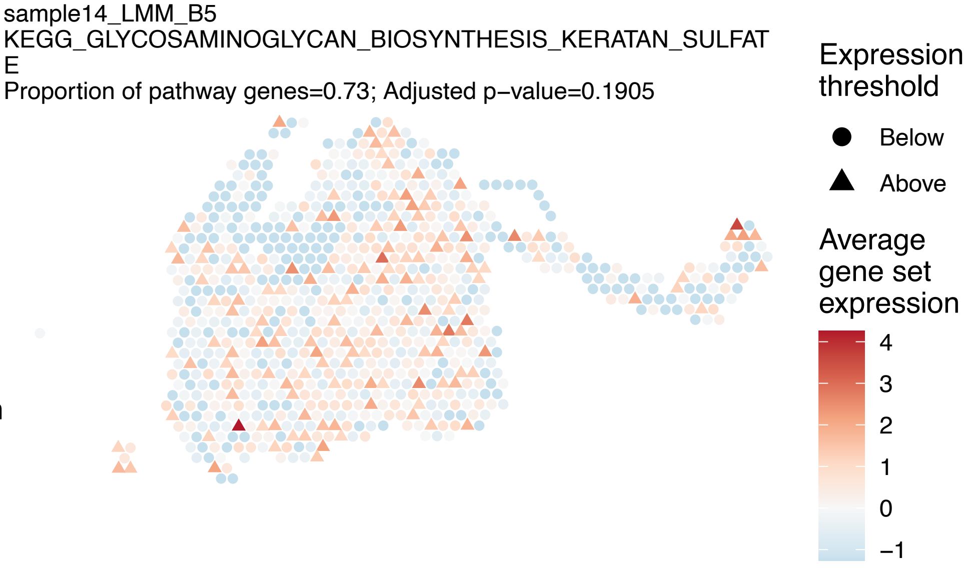
sample14_LMM_B5
KEGG_GLYCOSAMINOGLYCAN_BIOSYNTHESIS_KERATAN_SULFAT
E
Proportion of pathway genes=0.73; Adjusted p−value=0.1905
Expression threshold
Below
Above
Average gene set expression
4
3
2
1
0
−1

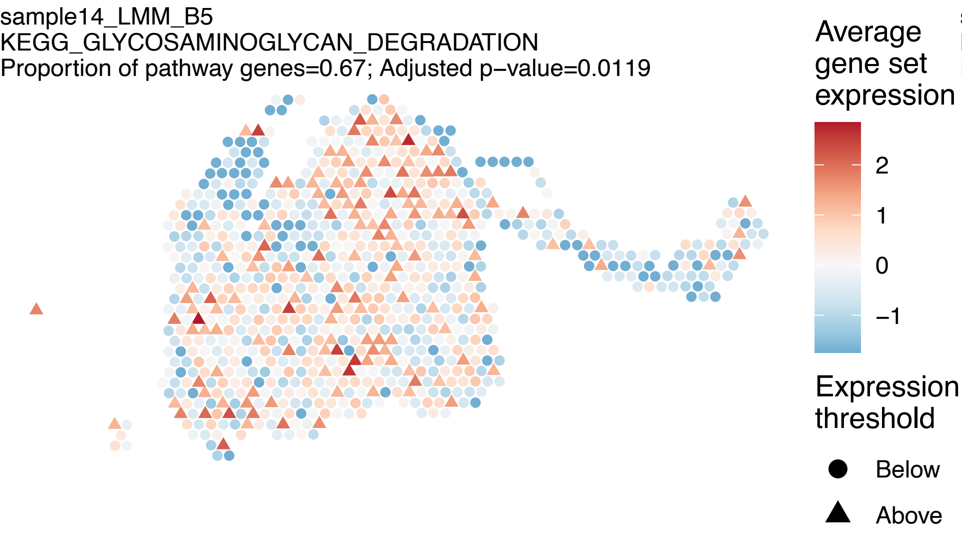
sample14_LMM_B5
KEGG_GLYCOSAMINOGLYCAN_DEGRADATION
Proportion of pathway genes=0.67; Adjusted p−value=0.0119
Average gene set expression
2
1
0
−1
Expression threshold
Below
Above

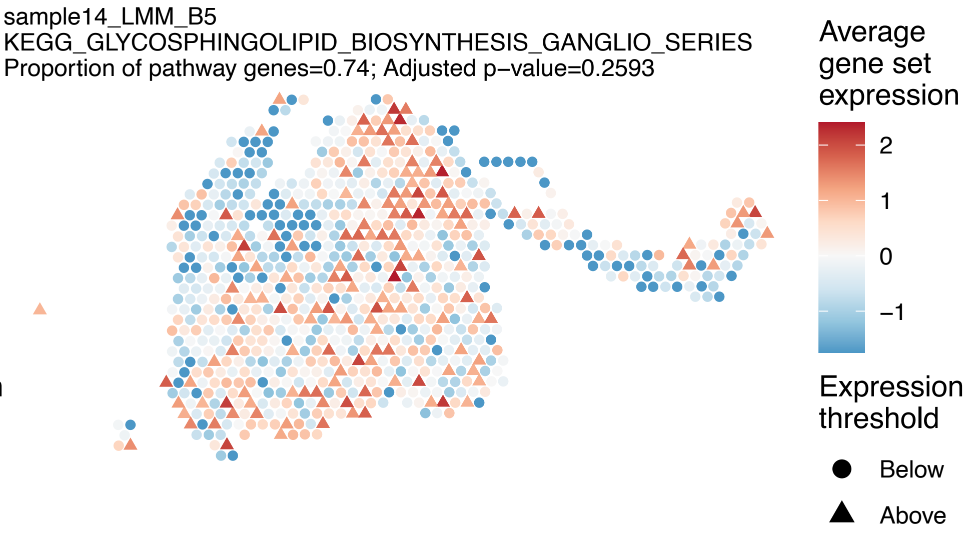
sample14_LMM_B5
KEGG_GLYCOSPHINGOLIPID_BIOSYNTHESIS_GANGLIO_SERIES
Proportion of pathway genes=0.74; Adjusted p−value=0.2593
Average gene set expression
2
1
0
−1
Expression threshold
Below
Above

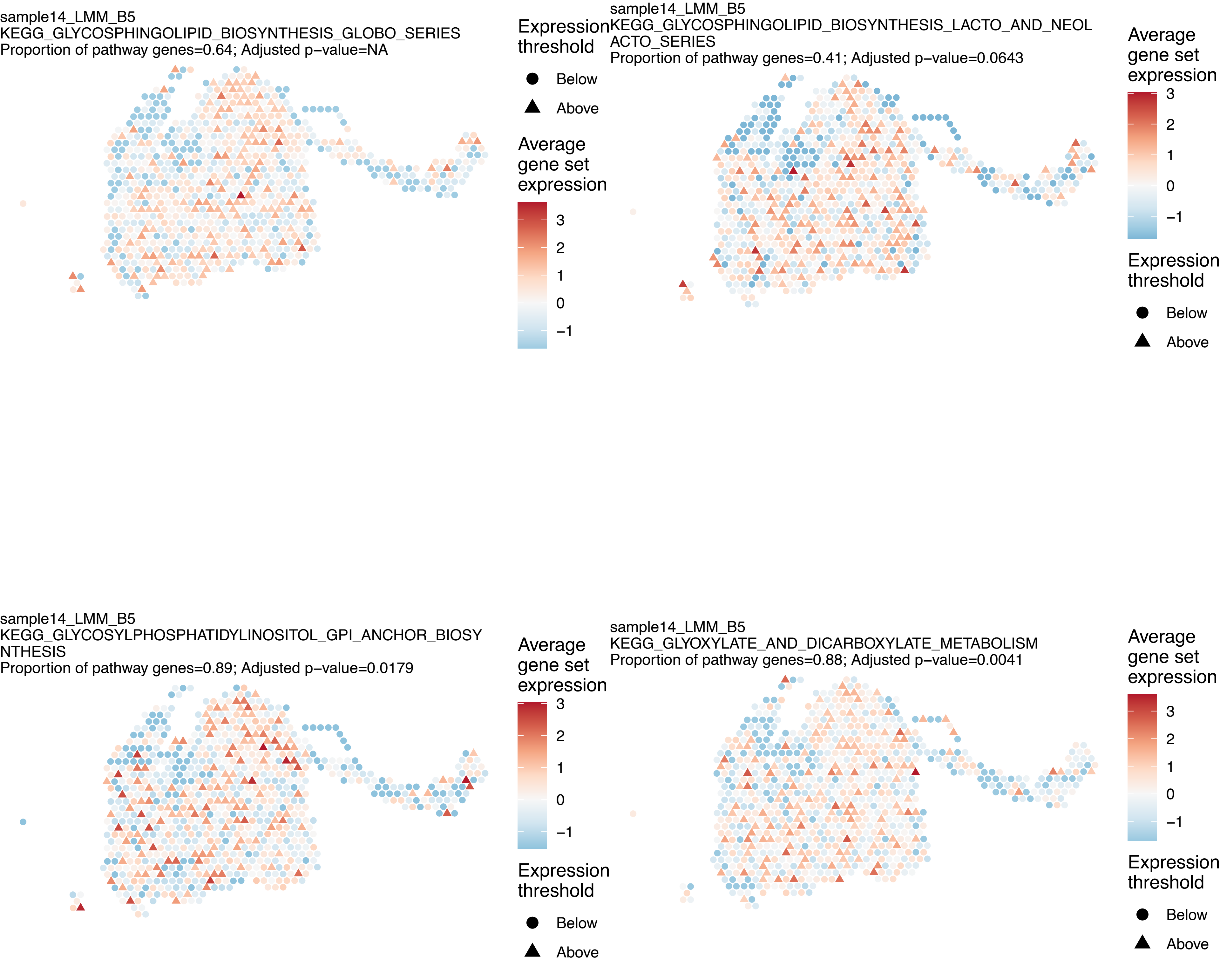

sample14_LMM_B5
KEGG_GLYCOSPHINGOLIPID_BIOSYNTHESIS_GLOBO_SERIES
Proportion of pathway genes=0.64; Adjusted p−value=NA
Expression threshold
Below
Above
Average gene set expression
3
2
1
0
−1
sample14_LMM_B5
KEGG_GLYCOSPHINGOLIPID_BIOSYNTHESIS_LACTO_AND_NEOL
ACTO_SERIES
Proportion of pathway genes=0.41; Adjusted p−value=0.0643
Average gene set expression
3
2
1
0
−1
Expression threshold
Below
Above
sample14_LMM_B5
KEGG_GLYCOSYLPHOSPHATIDYLINOSITOL_GPI_ANCHOR_BIOSY
NTHESIS
Proportion of pathway genes=0.89; Adjusted p−value=0.0179
Average gene set expression
3
2
1
0
−1
Expression threshold
Below
Above
sample14_LMM_B5
KEGG_GLYOXYLATE_AND_DICARBOXYLATE_METABOLISM
Proportion of pathway genes=0.88; Adjusted p−value=0.0041
Average gene set expression
3
2
1
0
−1
Expression threshold
Below
Above

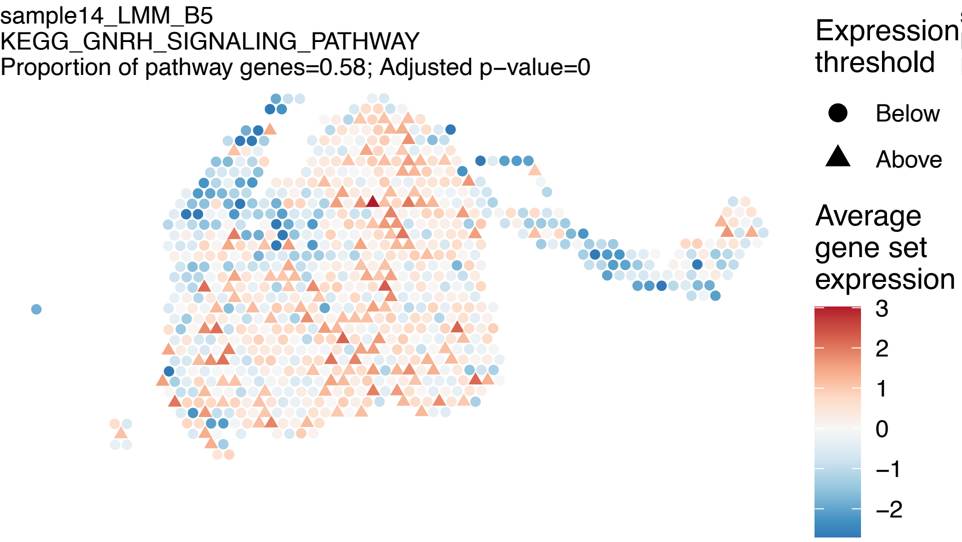
sample14_LMM_B5
KEGG_GNRH_SIGNALING_PATHWAY
Proportion of pathway genes=0.58; Adjusted p−value=0
Expression threshold
Below
Above
Average gene set expression
3
2
1
0
−1
−2

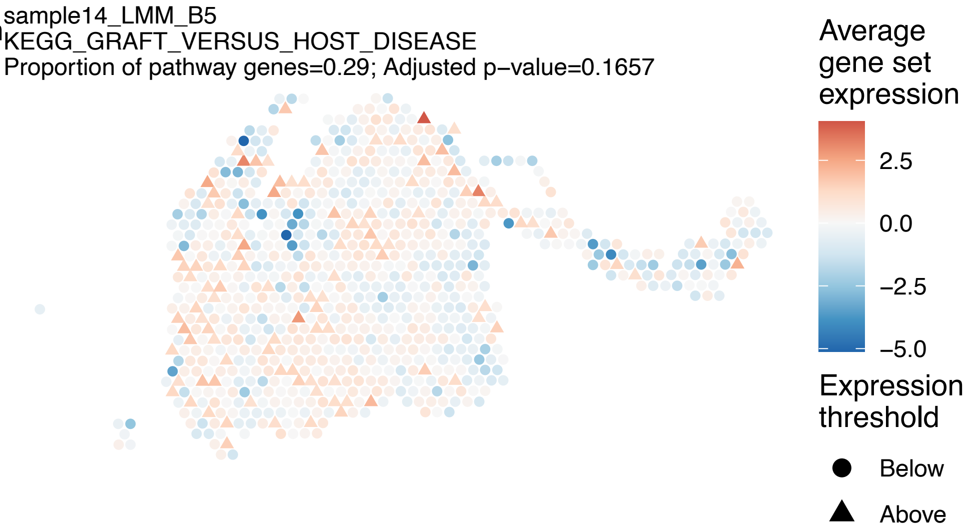
sample14_LMM_B5
KEGG_GRAFT_VERSUS_HOST_DISEASE
Proportion of pathway genes=0.29; Adjusted p−value=0.1657
Average gene set expression
2.5
0.0
−2.5
−5.0
Expression threshold
Below
Above

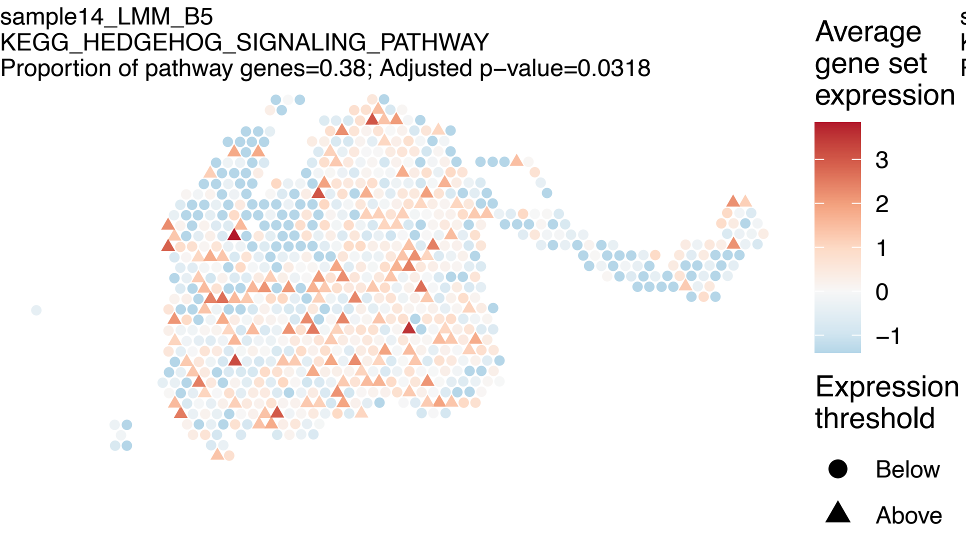
sample14_LMM_B5
KEGG_HEDGEHOG_SIGNALING_PATHWAY
Proportion of pathway genes=0.38; Adjusted p−value=0.0318
Average gene set expression
3
2
1
0
−1
Expression threshold
Below
Above

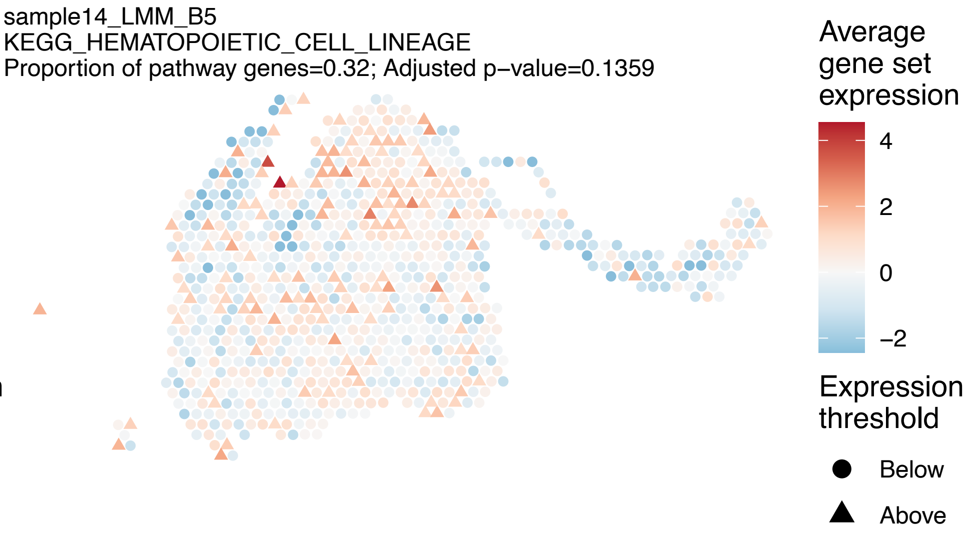
sample14_LMM_B5
KEGG_HEMATOPOIETIC_CELL_LINEAGE
Proportion of pathway genes=0.32; Adjusted p−value=0.1359
Average gene set expression
4
2
0
−2
Expression threshold
Below
Above

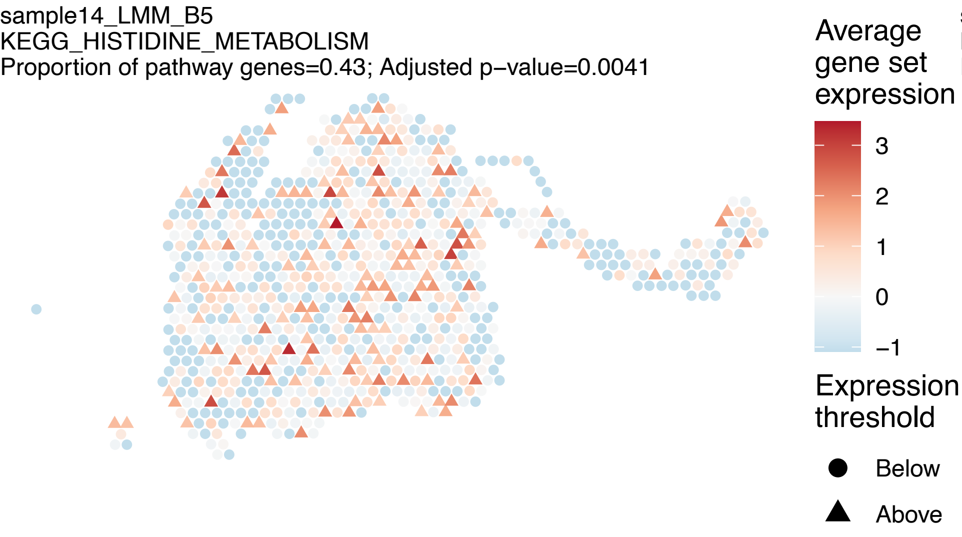
sample14_LMM_B5
KEGG_HISTIDINE_METABOLISM
Proportion of pathway genes=0.43; Adjusted p−value=0.0041
Average gene set expression
3
2
1
0
−1
Expression threshold
Below
Above

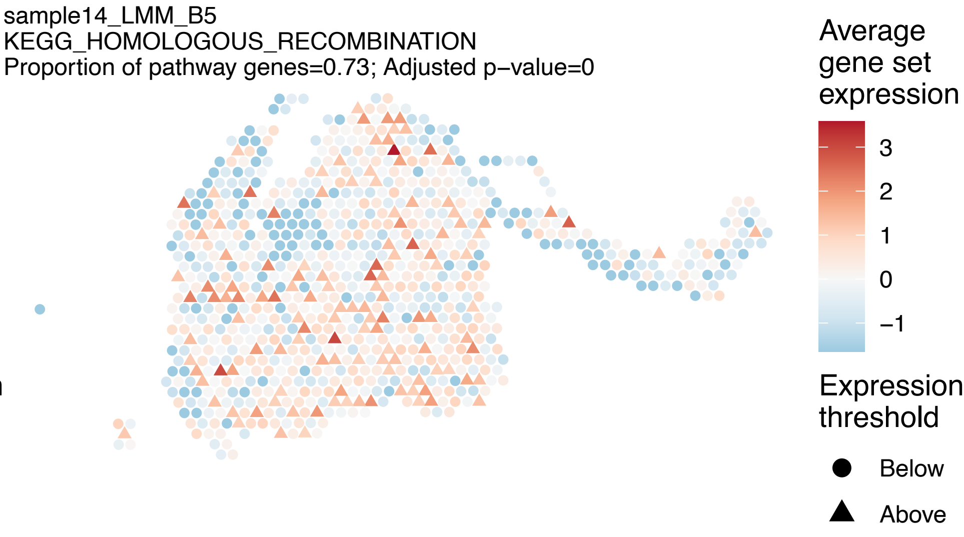
sample14_LMM_B5
KEGG_HOMOLOGOUS_RECOMBINATION
Proportion of pathway genes=0.73; Adjusted p−value=0
Average gene set expression
3
2
1
0
−1
Expression threshold
Below
Above

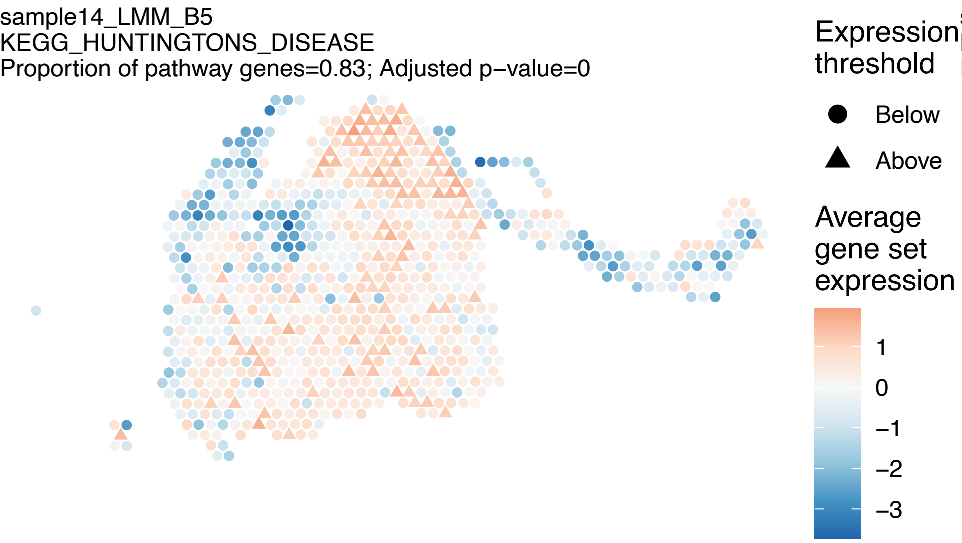
sample14_LMM_B5
KEGG_HUNTINGTONS_DISEASE
Proportion of pathway genes=0.83; Adjusted p−value=0
Expression threshold
Below
Above
Average gene set expression
1
0
−1
−2
−3

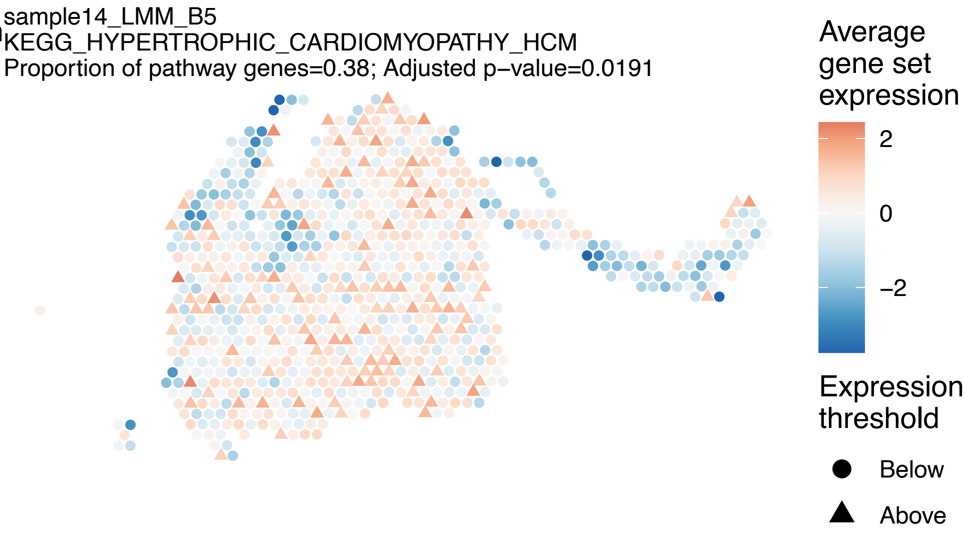
sample14_LMM_B5
KEGG_HYPERTROPHIC_CARDIOMYOPATHY_HCM
Proportion of pathway genes=0.38; Adjusted p−value=0.0191
Average gene set expression
2
0
−2
Expression threshold
Below
Above

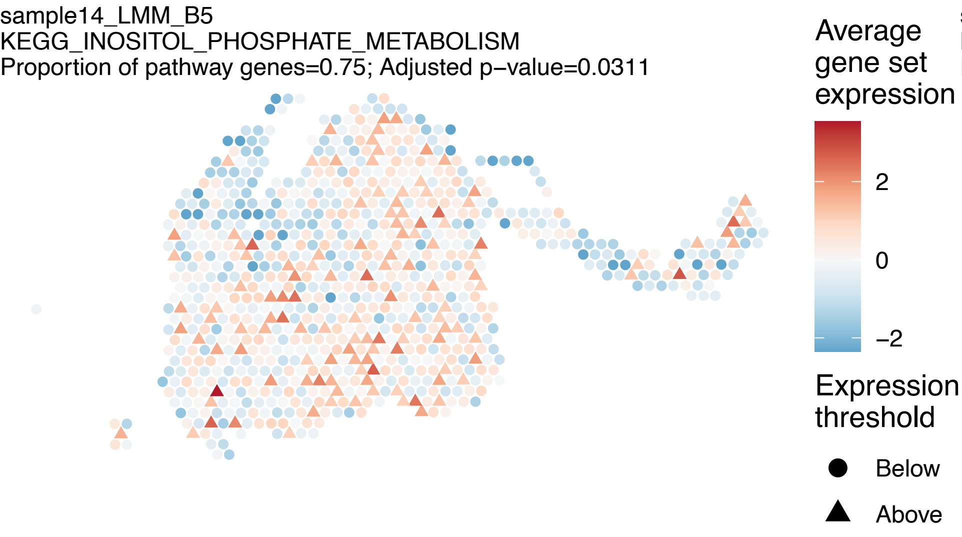
sample14_LMM_B5
KEGG_INOSITOL_PHOSPHATE_METABOLISM
Proportion of pathway genes=0.75; Adjusted p−value=0.0311
Average gene set expression
2
0
−2
Expression threshold
Below
Above

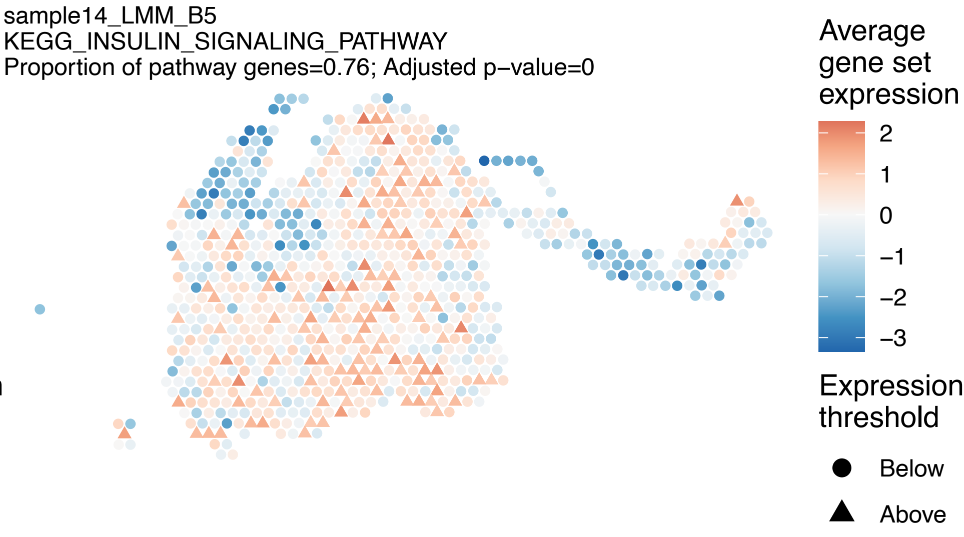
sample14_LMM_B5
KEGG_INSULIN_SIGNALING_PATHWAY
Proportion of pathway genes=0.76; Adjusted p−value=0
Average gene set expression
2
1
0
−1
−2
−3
Expression threshold
Below
Above

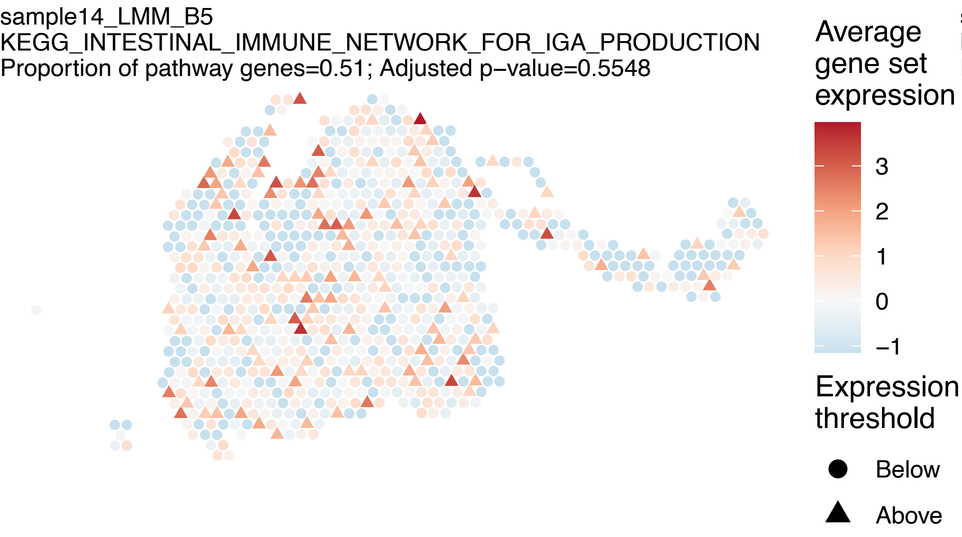
sample14_LMM_B5
KEGG_INTESTINAL_IMMUNE_NETWORK_FOR_IGA_PRODUCTION
Proportion of pathway genes=0.51; Adjusted p−value=0.5548
Average gene set expression
3
2
1
0
−1
Expression threshold
Below
Above

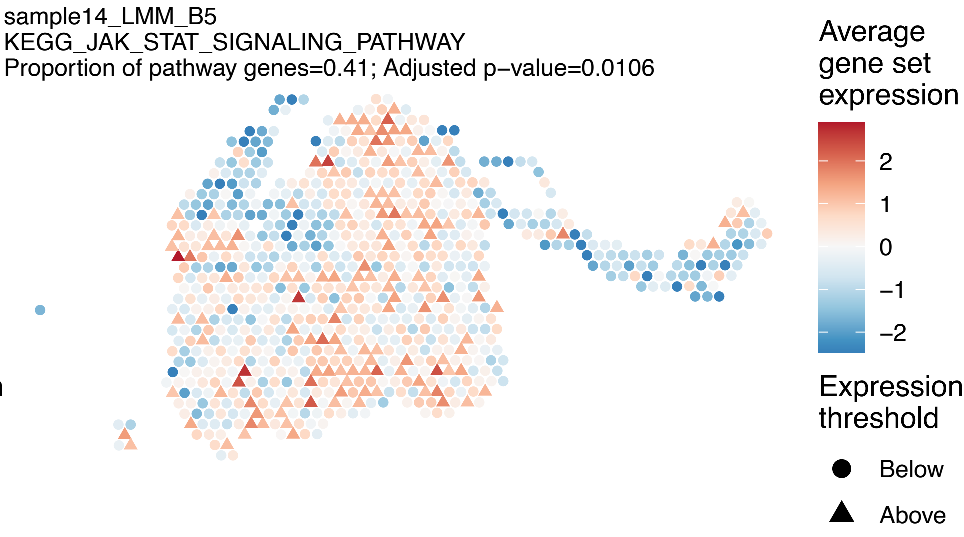
sample14_LMM_B5
KEGG_JAK_STAT_SIGNALING_PATHWAY
Proportion of pathway genes=0.41; Adjusted p−value=0.0106
Average gene set expression
2
1
0
−1
−2
Expression threshold
Below
Above

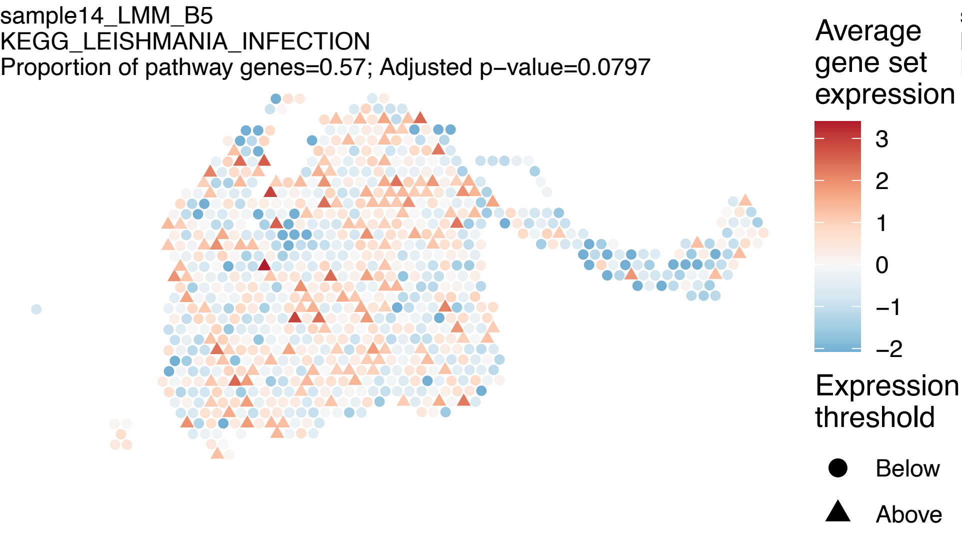
sample14_LMM_B5
KEGG_LEISHMANIA_INFECTION
Proportion of pathway genes=0.57; Adjusted p−value=0.0797
Average gene set expression
3
2
1
0
−1
−2
Expression threshold
Below
Above

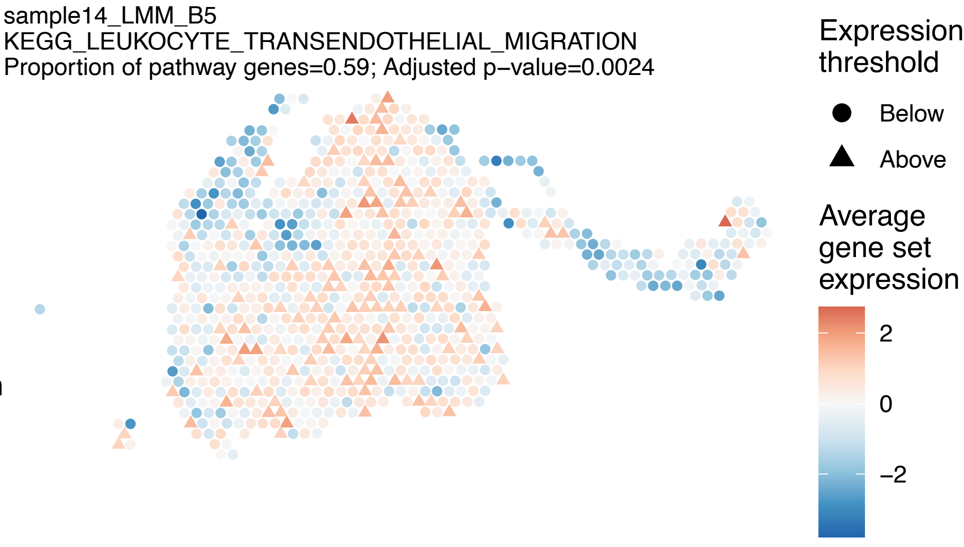
sample14_LMM_B5
KEGG_LEUKOCYTE_TRANSENDOTHELIAL_MIGRATION
Proportion of pathway genes=0.59; Adjusted p−value=0.0024
Expression threshold
Below
Above
Average gene set expression
2
0
−2

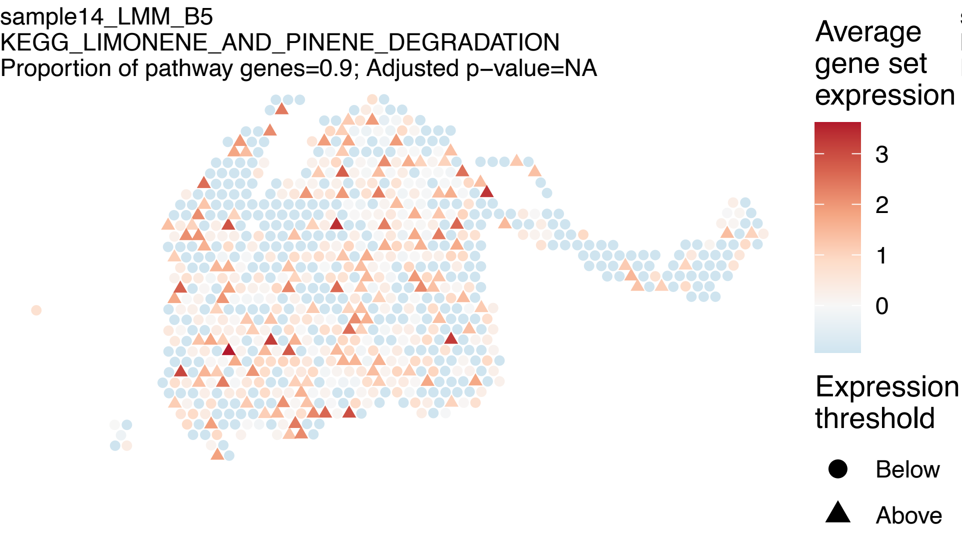
sample14_LMM_B5
KEGG_LIMONENE_AND_PINENE_DEGRADATION
Proportion of pathway genes=0.9; Adjusted p−value=NA
Average gene set expression
3
2
1
0
Expression threshold
Below
Above

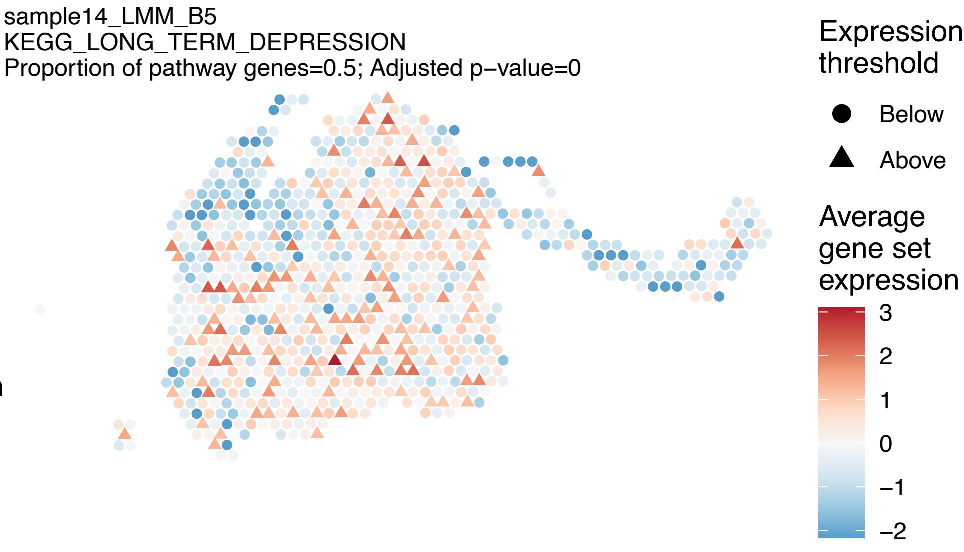
sample14_LMM_B5
KEGG_LONG_TERM_DEPRESSION
Proportion of pathway genes=0.5; Adjusted p−value=0
Expression threshold
Below
Above
Average gene set expression
3
2
1
0
−1
−2

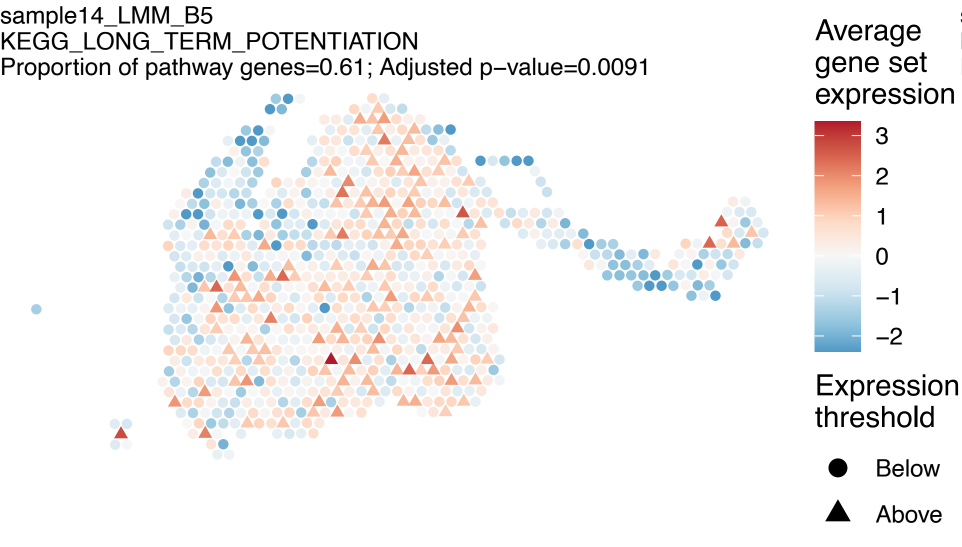
sample14_LMM_B5
KEGG_LONG_TERM_POTENTIATION
Proportion of pathway genes=0.61; Adjusted p−value=0.0091
Average gene set expression
3
2
1
0
−1
−2
Expression threshold
Below
Above

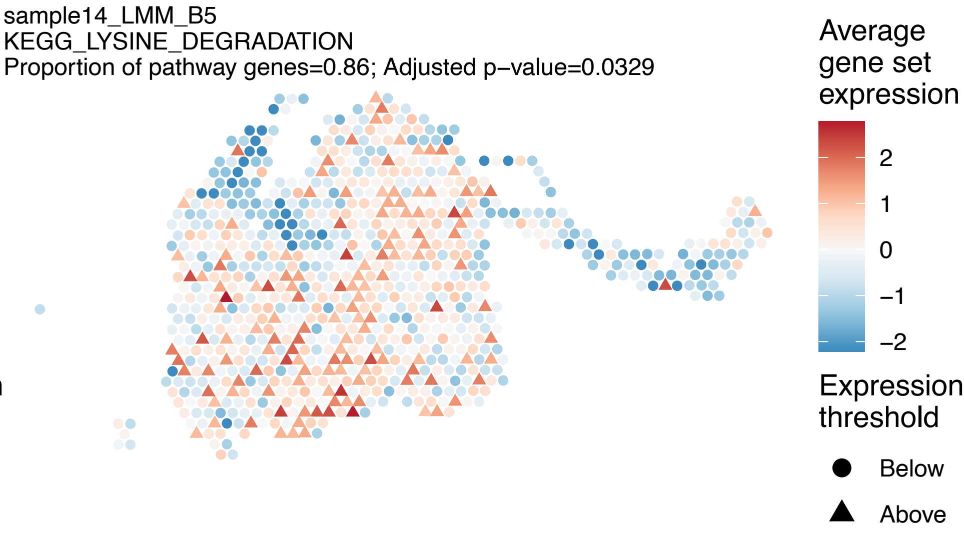
sample14_LMM_B5
KEGG_LYSINE_DEGRADATION
Proportion of pathway genes=0.86; Adjusted p−value=0.0329
Average gene set expression
2
1
0
−1
−2
Expression threshold
Below
Above

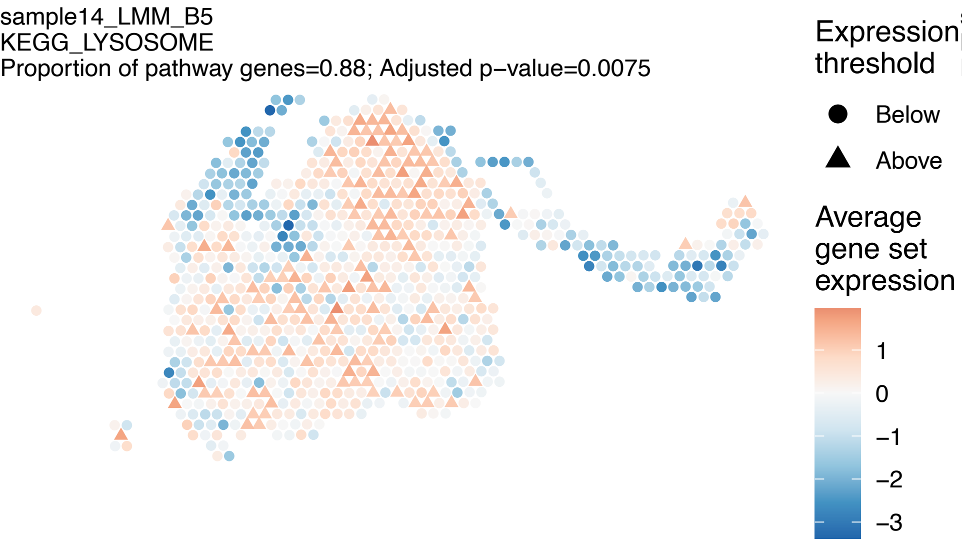
sample14_LMM_B5
KEGG_LYSOSOME
Proportion of pathway genes=0.88; Adjusted p−value=0.0075
Expression threshold
Below
Above
Average gene set expression
1
0
−1
−2
−3

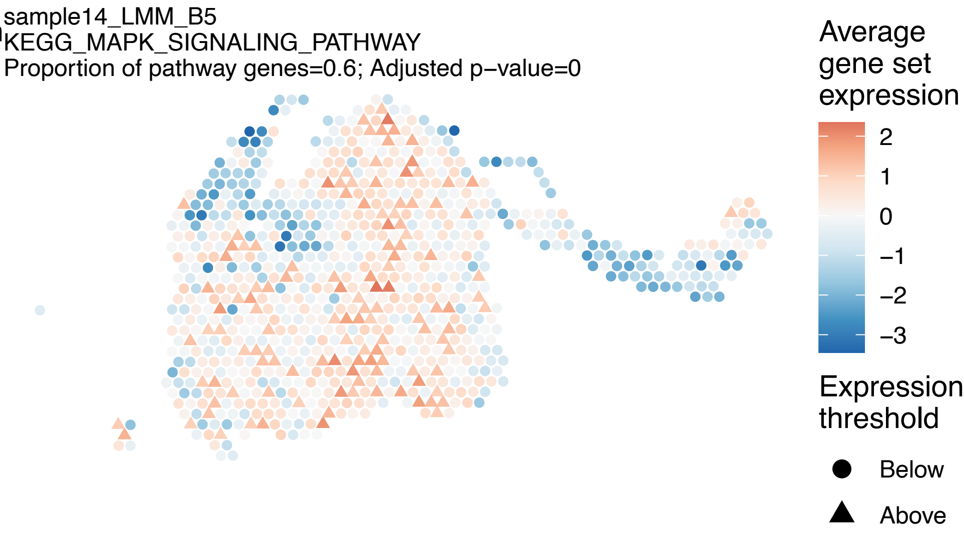
sample14_LMM_B5
KEGG_MAPK_SIGNALING_PATHWAY
Proportion of pathway genes=0.6; Adjusted p−value=0
Average gene set expression
2
1
0
−1
−2
−3
Expression threshold
Below
Above

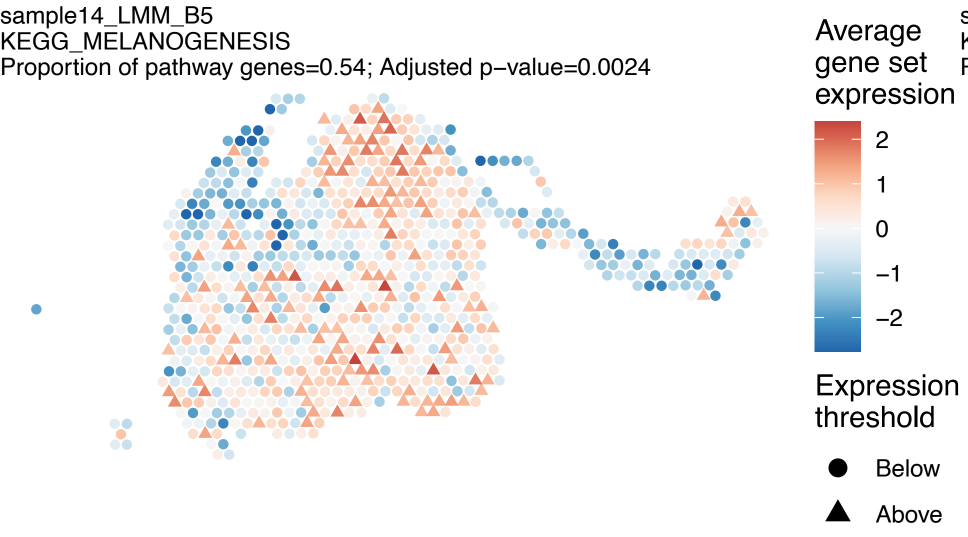

sample14_LMM_B5
KEGG_MELANOGENESIS
Proportion of pathway genes=0.54; Adjusted p−value=0.0024
Average gene set expression
2
1
0
−1
−2
Expression threshold
Below
Above


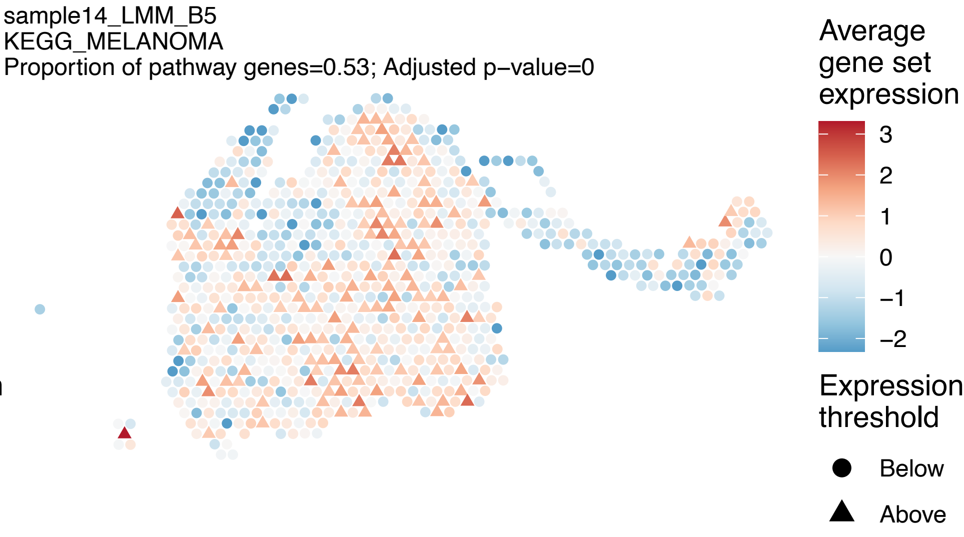

sample14_LMM_B5
KEGG_MELANOMA
Proportion of pathway genes=0.53; Adjusted p−value=0
Average gene set expression
3
2
1
0
−1
−2
Expression threshold
Below
Above


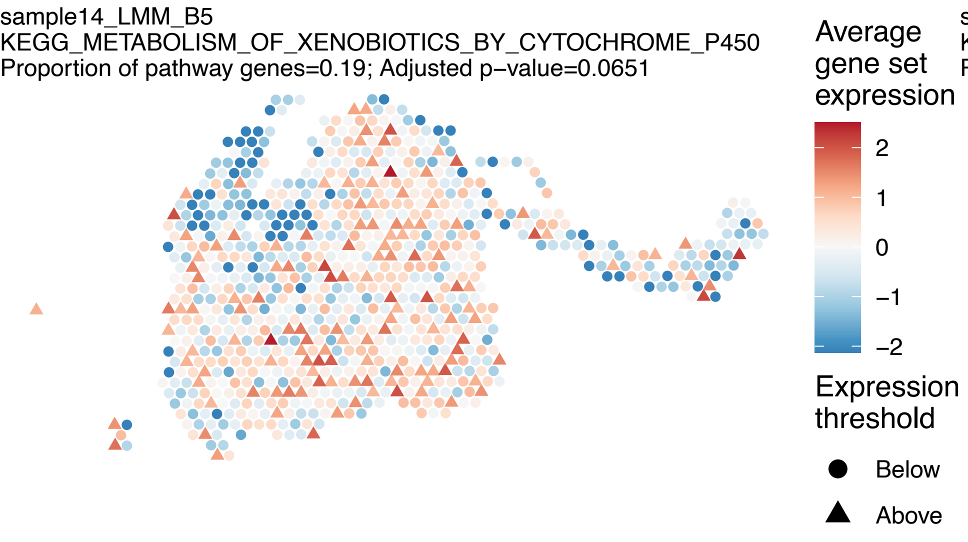

sample14_LMM_B5
KEGG_METABOLISM_OF_XENOBIOTICS_BY_CYTOCHROME_P450
Proportion of pathway genes=0.19; Adjusted p−value=0.0651
Average gene set expression
2
1
0
−1
−2
Expression threshold
Below
Above


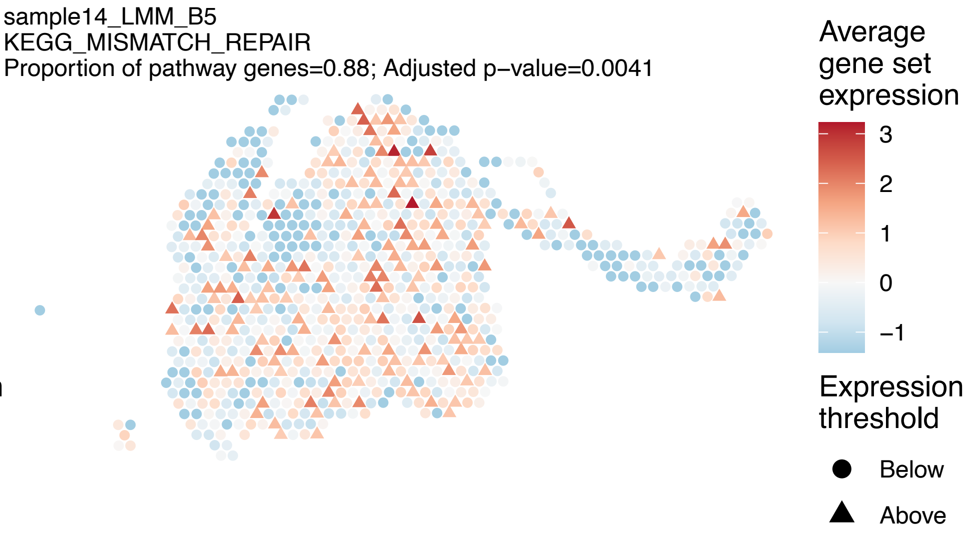

sample14_LMM_B5
KEGG_MISMATCH_REPAIR
Proportion of pathway genes=0.88; Adjusted p−value=0.0041
Average gene set expression
3
2
1
0
−1
Expression threshold
Below
Above

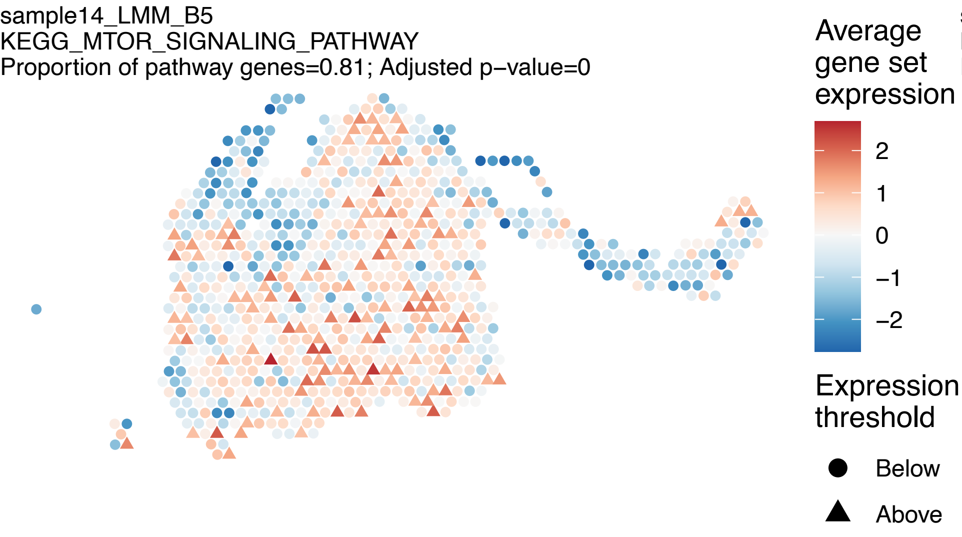
sample14_LMM_B5
KEGG_MTOR_SIGNALING_PATHWAY
Proportion of pathway genes=0.81; Adjusted p−value=0
Average gene set expression
2
1
0
−1
−2
Expression threshold
Below
Above

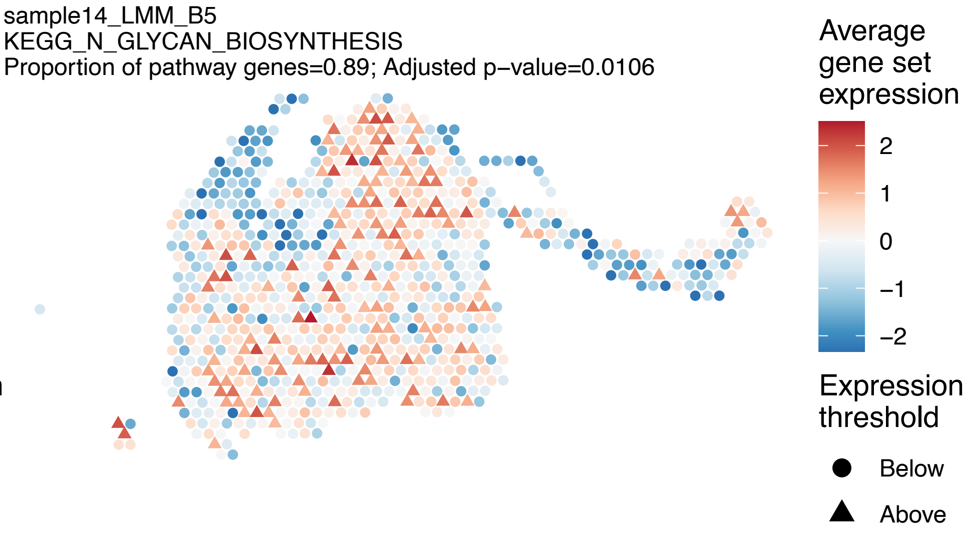
sample14_LMM_B5
KEGG_N_GLYCAN_BIOSYNTHESIS
Proportion of pathway genes=0.89; Adjusted p−value=0.0106
Average gene set expression
2
1
0
−1
−2
Expression threshold
Below
Above

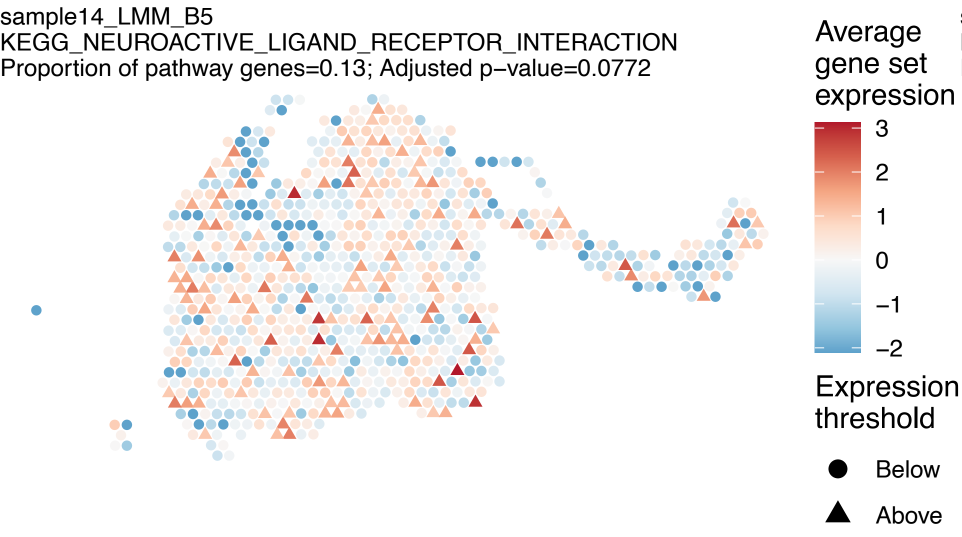
sample14_LMM_B5
KEGG_NEUROACTIVE_LIGAND_RECEPTOR_INTERACTION
Proportion of pathway genes=0.13; Adjusted p−value=0.0772
Average gene set expression
3
2
1
0
−1
−2
Expression threshold
Below
Above

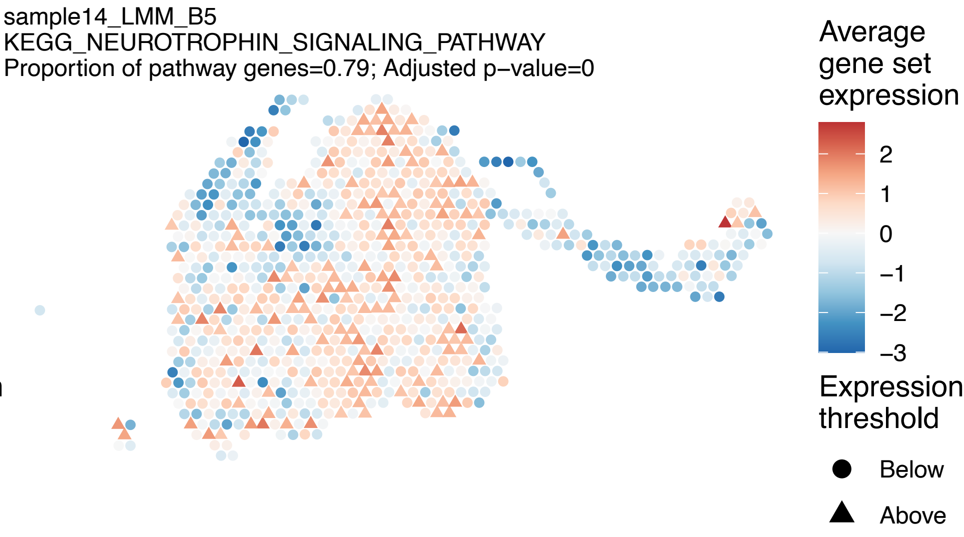
sample14_LMM_B5
KEGG_NEUROTROPHIN_SIGNALING_PATHWAY
Proportion of pathway genes=0.79; Adjusted p−value=0
Average gene set expression
2
1
0
−1
−2
−3
Expression threshold
Below
Above

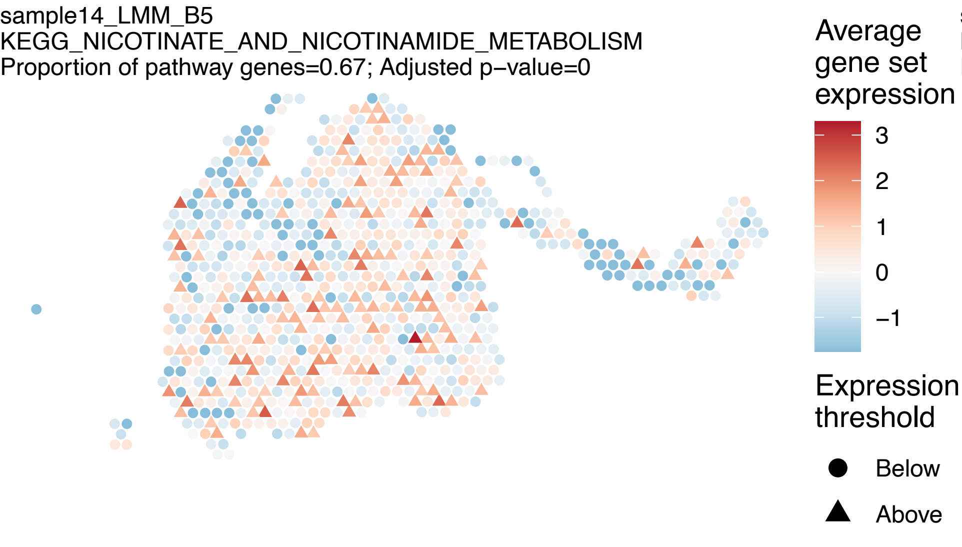
sample14_LMM_B5
KEGG_NICOTINATE_AND_NICOTINAMIDE_METABOLISM
Proportion of pathway genes=0.67; Adjusted p−value=0
Average gene set expression
3
2
1
0
−1
Expression threshold
Below
Above

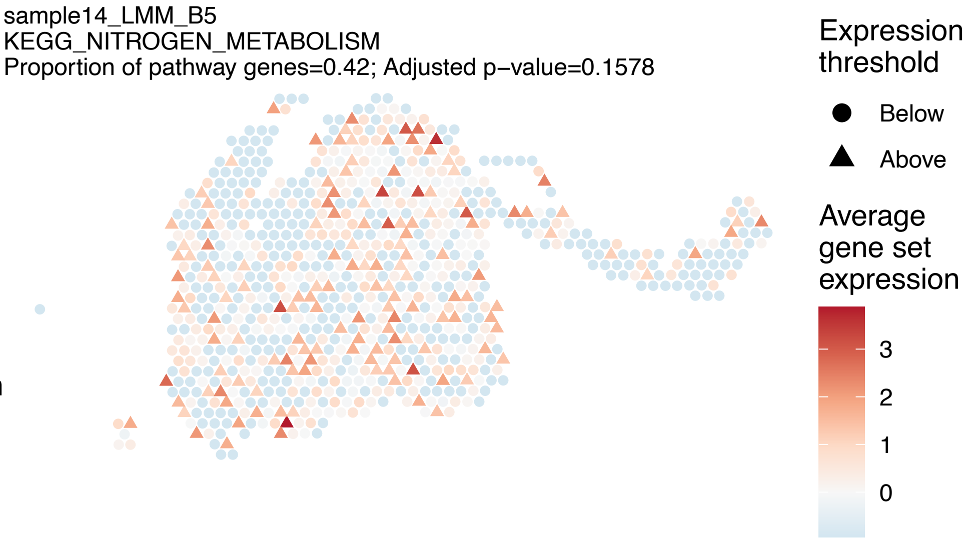
sample14_LMM_B5
KEGG_NITROGEN_METABOLISM
Proportion of pathway genes=0.42; Adjusted p−value=0.1578
Expression threshold
Below
Above
Average gene set expression
3
2
1
0

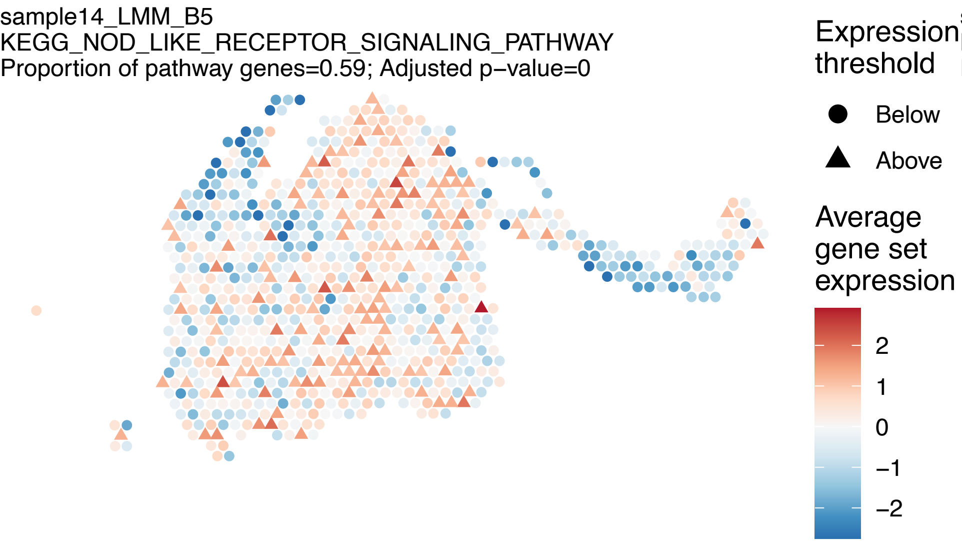
sample14_LMM_B5
KEGG_NOD_LIKE_RECEPTOR_SIGNALING_PATHWAY
Proportion of pathway genes=0.59; Adjusted p−value=0
Expression threshold
Below
Above
Average gene set expression
2
1
0
−1
−2

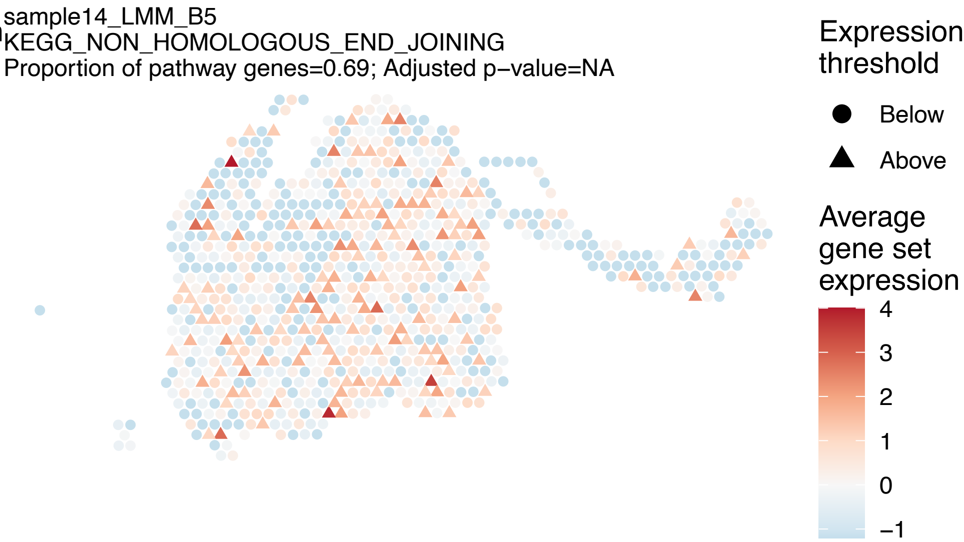
sample14_LMM_B5
KEGG_NON_HOMOLOGOUS_END_JOINING
Proportion of pathway genes=0.69; Adjusted p−value=NA
Expression threshold
Below
Above
Average gene set expression
4
3
2
1
0
−1

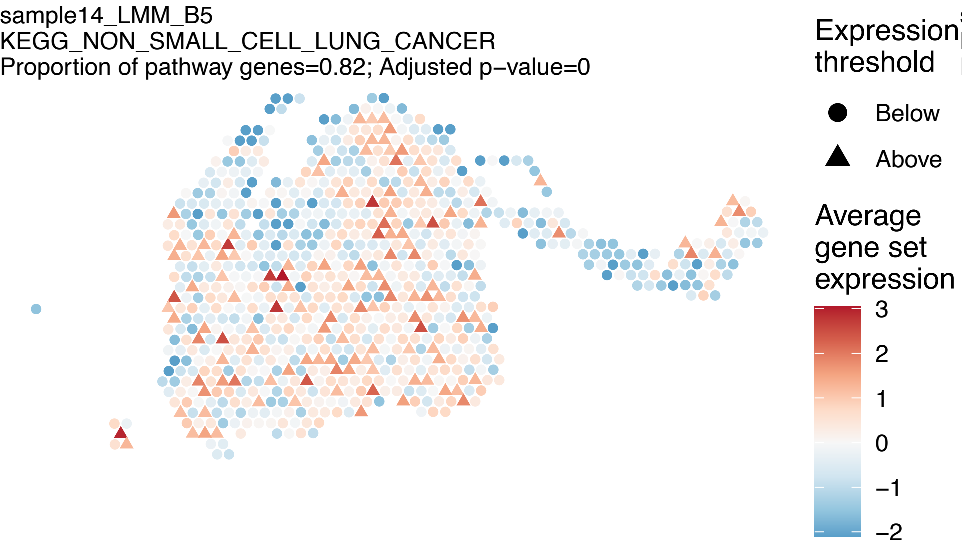
sample14_LMM_B5
KEGG_NON_SMALL_CELL_LUNG_CANCER
Proportion of pathway genes=0.82; Adjusted p−value=0
Expression threshold
Below
Above
Average gene set expression
3
2
1
0
−1
−2

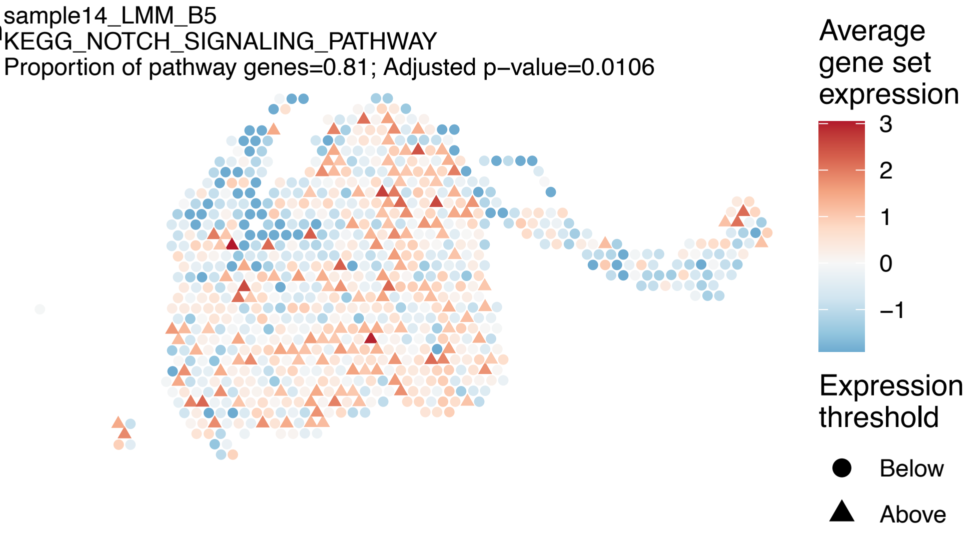
sample14_LMM_B5
KEGG_NOTCH_SIGNALING_PATHWAY
Proportion of pathway genes=0.81; Adjusted p−value=0.0106
Average gene set expression
3
2
1
0
−1
Expression threshold
Below
Above

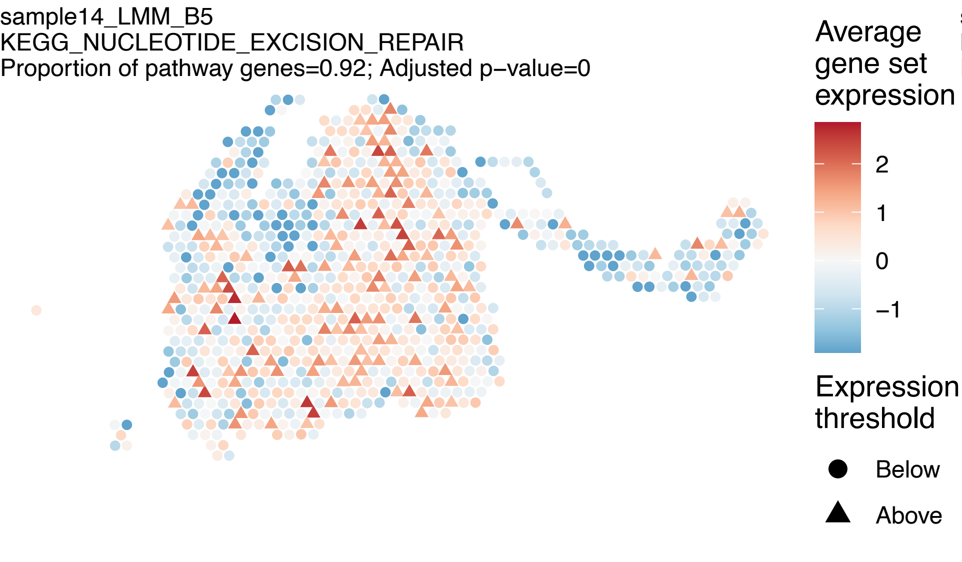
sample14_LMM_B5
KEGG_NUCLEOTIDE_EXCISION_REPAIR
Proportion of pathway genes=0.92; Adjusted p−value=0
Average gene set expression
2
1
0
−1
Expression threshold
Below
Above

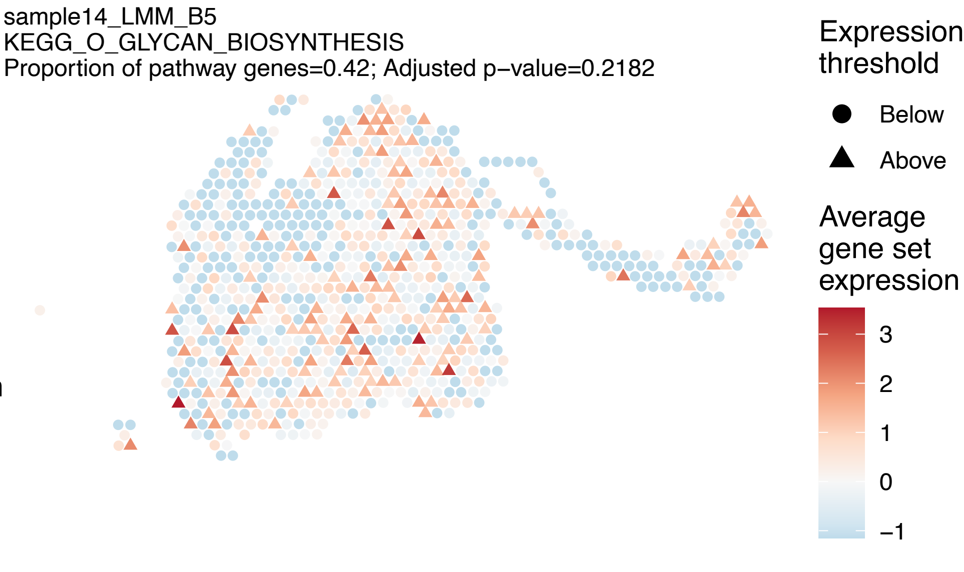
sample14_LMM_B5
KEGG_O_GLYCAN_BIOSYNTHESIS
Proportion of pathway genes=0.42; Adjusted p−value=0.2182
Expression threshold
Below
Above
Average gene set expression
3
2
1
0
−1

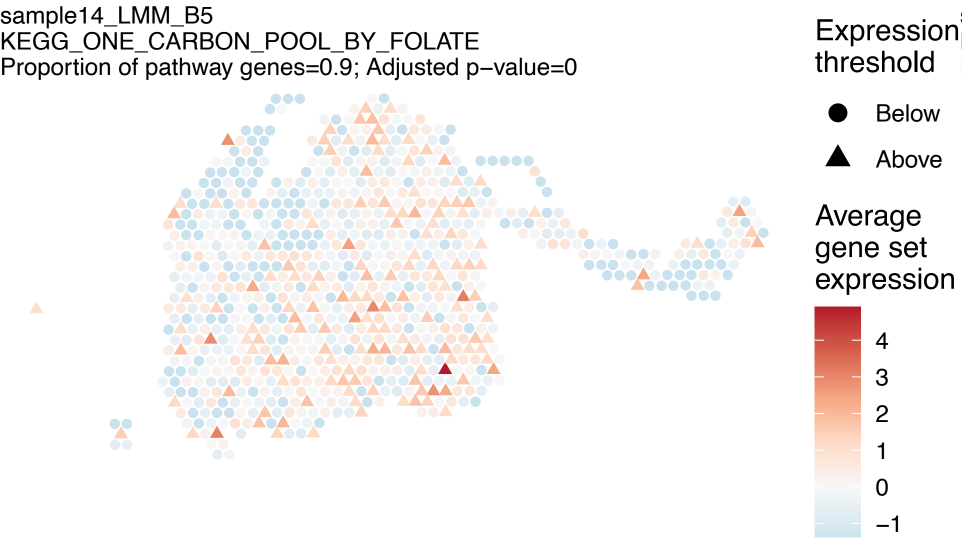
sample14_LMM_B5
KEGG_ONE_CARBON_POOL_BY_FOLATE
Proportion of pathway genes=0.9; Adjusted p−value=0
Expression threshold
Below
Above
Average gene set expression
4
3
2
1
0
−1

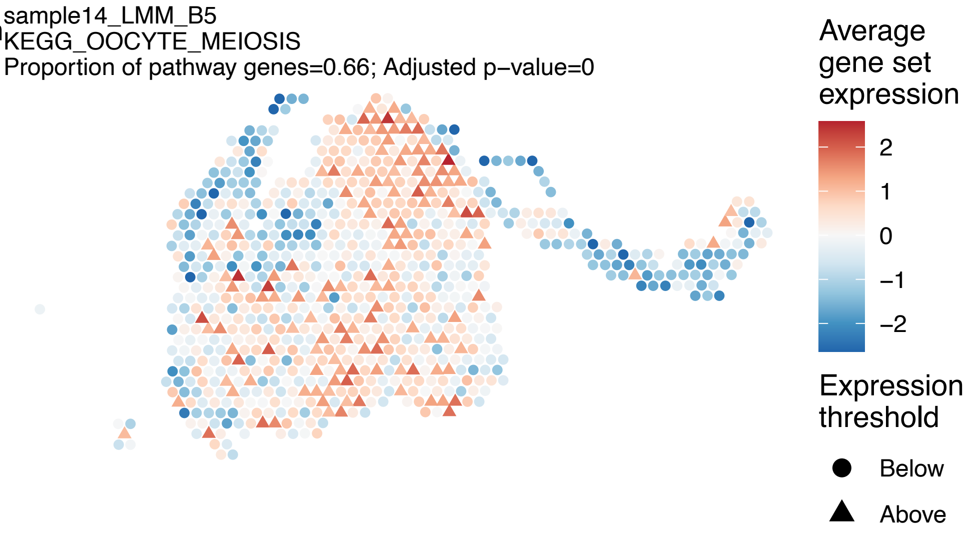
sample14_LMM_B5
KEGG_OOCYTE_MEIOSIS
Proportion of pathway genes=0.66; Adjusted p−value=0
Average gene set expression
2
1
0
−1
−2
Expression threshold
Below
Above

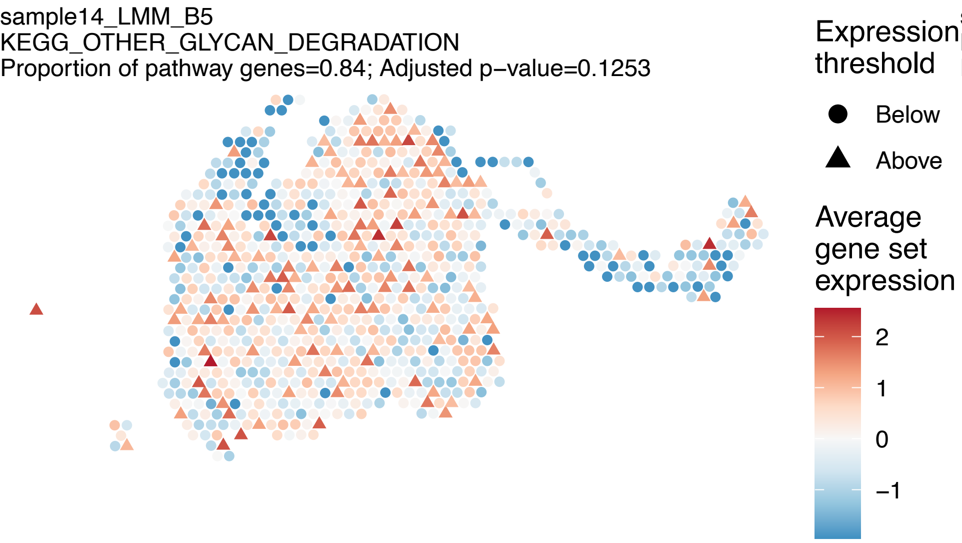
sample14_LMM_B5
KEGG_OTHER_GLYCAN_DEGRADATION
Proportion of pathway genes=0.84; Adjusted p−value=0.1253
Expression threshold
Below
Above
Average gene set expression
2
1
0
−1

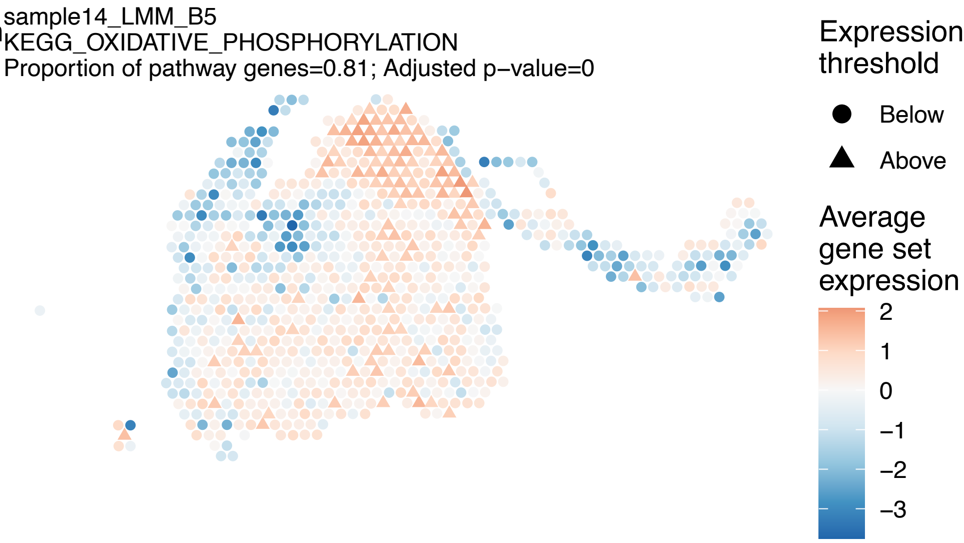
sample14_LMM_B5
KEGG_OXIDATIVE_PHOSPHORYLATION
Proportion of pathway genes=0.81; Adjusted p−value=0
Expression threshold
Below
Above
Average gene set expression
2
1
0
−1
−2
−3

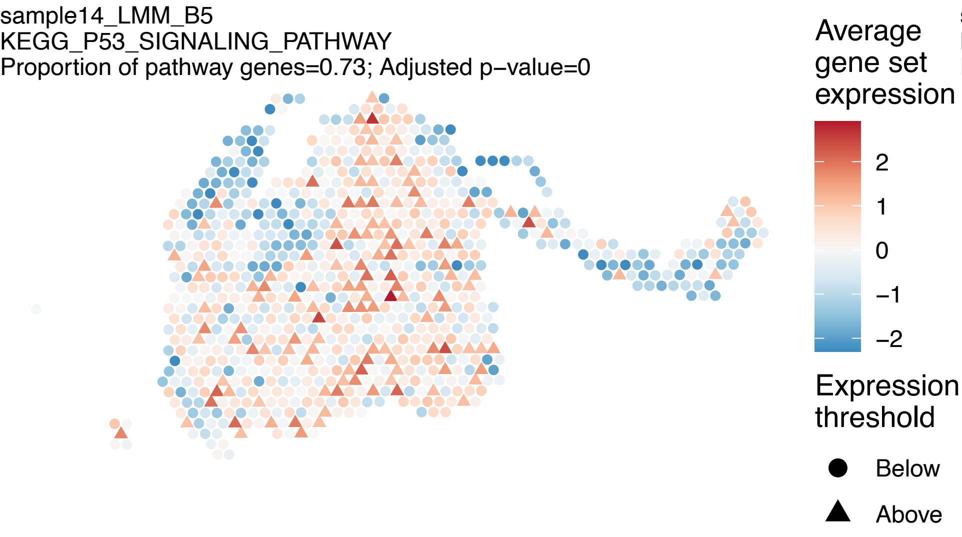
sample14_LMM_B5
KEGG_P53_SIGNALING_PATHWAY
Proportion of pathway genes=0.73; Adjusted p−value=0
Average gene set expression
2
1
0
−1
−2
Expression threshold
Below
Above

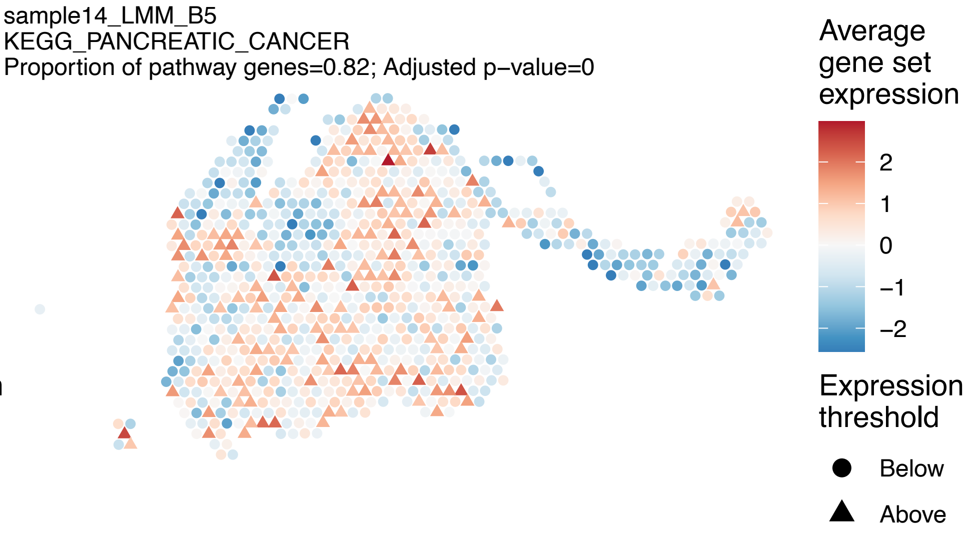
sample14_LMM_B5
KEGG_PANCREATIC_CANCER
Proportion of pathway genes=0.82; Adjusted p−value=0
Average gene set expression
2
1
0
−1
−2
Expression threshold
Below
Above

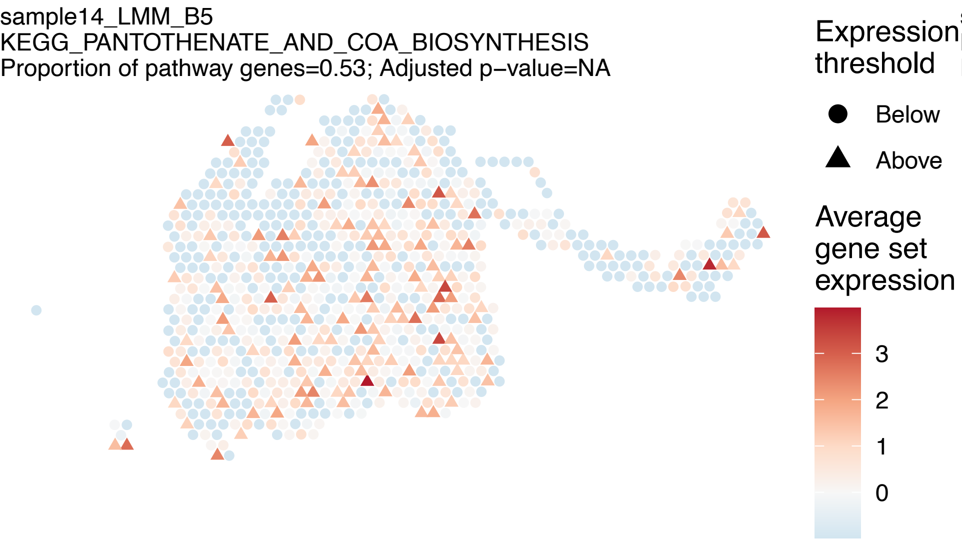
sample14_LMM_B5
KEGG_PANTOTHENATE_AND_COA_BIOSYNTHESIS
Proportion of pathway genes=0.53; Adjusted p−value=NA
Expression threshold
Below
Above
Average gene set expression
3
2
1
0

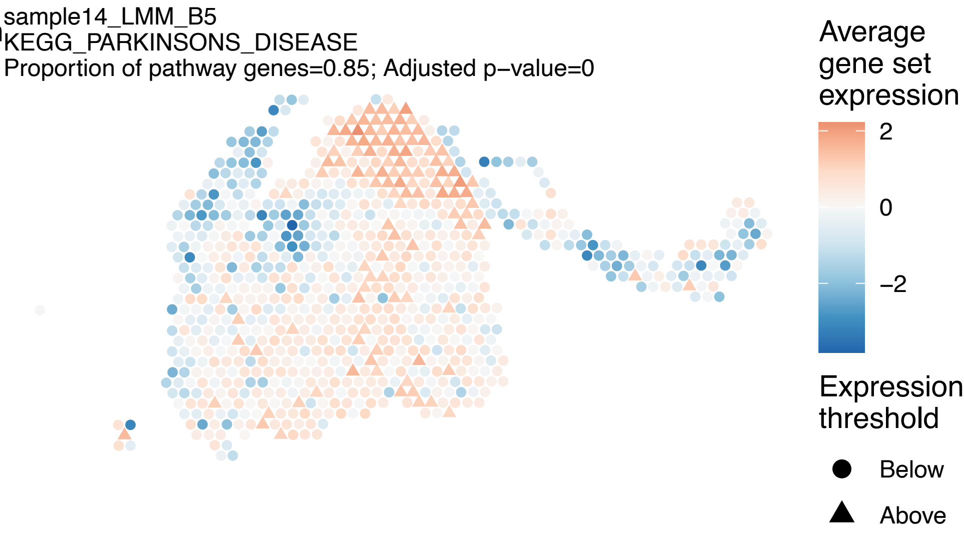
sample14_LMM_B5
KEGG_PARKINSONS_DISEASE
Proportion of pathway genes=0.85; Adjusted p−value=0
Average gene set expression
2
0
−2
Expression threshold
Below
Above

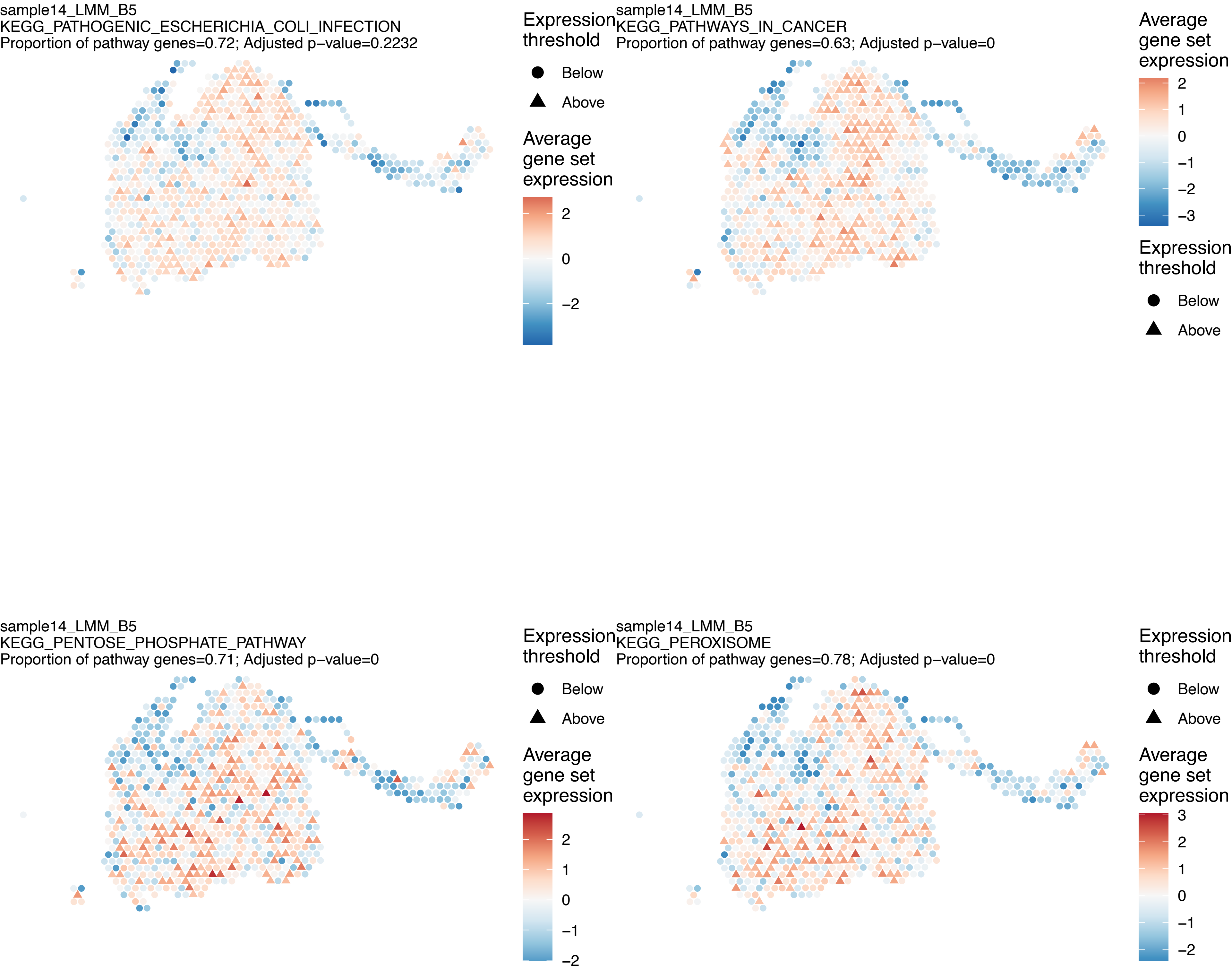
sample14_LMM_B5
KEGG_PATHOGENIC_ESCHERICHIA_COLI_INFECTION
Proportion of pathway genes=0.72; Adjusted p−value=0.2232
Expression threshold
Below
Above
Average gene set expression
2
0
−2
sample14_LMM_B5
KEGG_PATHWAYS_IN_CANCER
Proportion of pathway genes=0.63; Adjusted p−value=0
Average gene set expression
2
1
0
−1
−2
−3
Expression threshold
Below
Above
sample14_LMM_B5
KEGG_PENTOSE_PHOSPHATE_PATHWAY
Proportion of pathway genes=0.71; Adjusted p−value=0
Expression threshold
Below
Above
Average gene set expression
2
1
0
−1
−2
sample14_LMM_B5
KEGG_PEROXISOME
Proportion of pathway genes=0.78; Adjusted p−value=0
Expression threshold
Below
Above
Average gene set expression
3
2
1
0
−1
−2

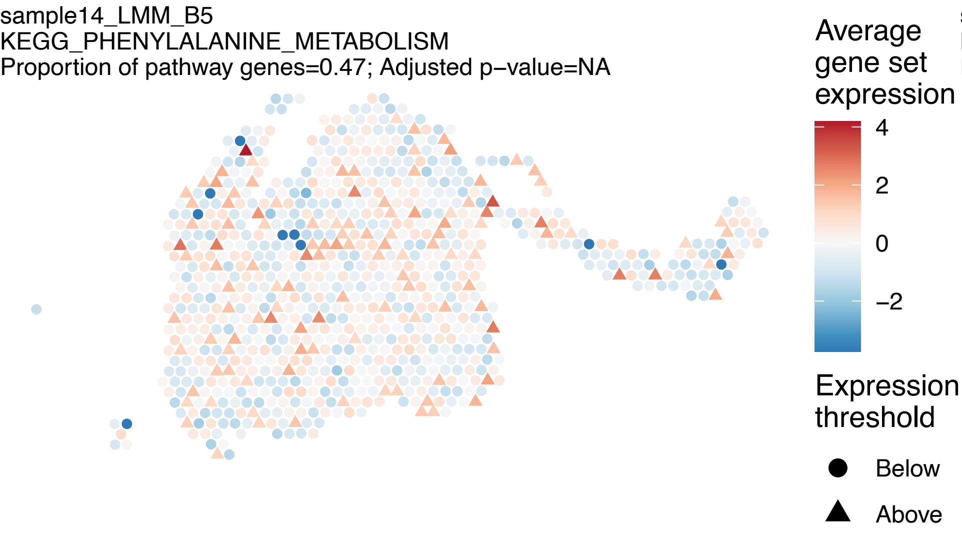
sample14_LMM_B5
KEGG_PHENYLALANINE_METABOLISM
Proportion of pathway genes=0.47; Adjusted p−value=NA
Average gene set expression
4
2
0
−2
Expression threshold
Below
Above

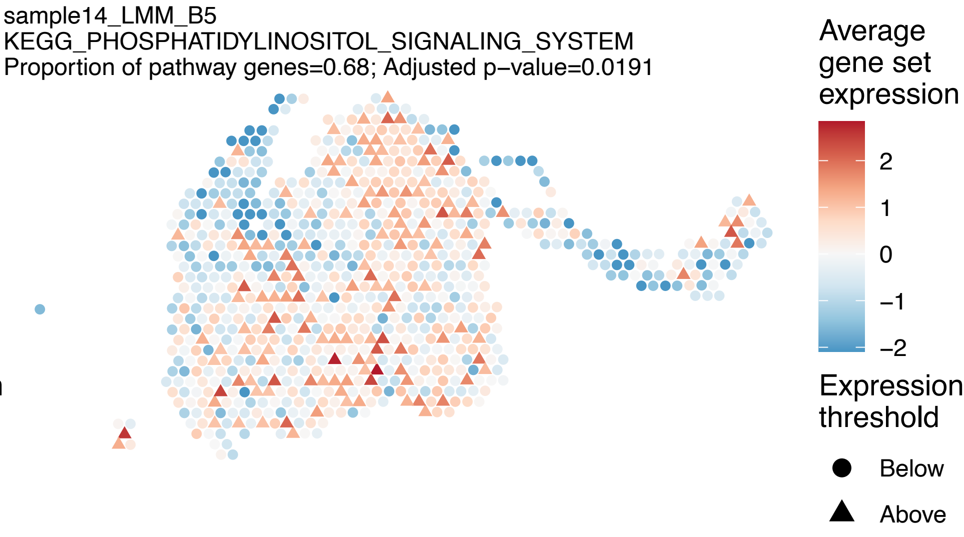
sample14_LMM_B5
KEGG_PHOSPHATIDYLINOSITOL_SIGNALING_SYSTEM
Proportion of pathway genes=0.68; Adjusted p−value=0.0191
Average gene set expression
2
1
0
−1
−2
Expression threshold
Below
Above

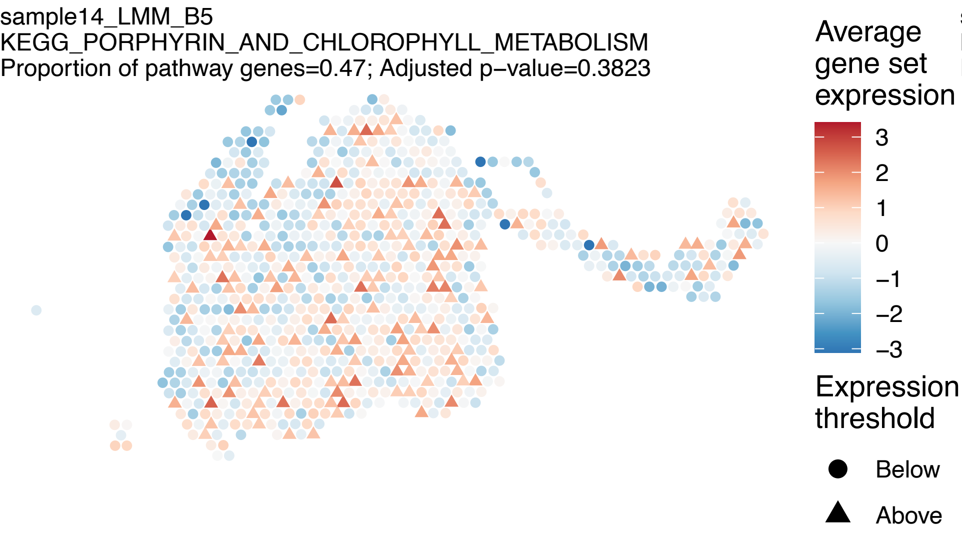
sample14_LMM_B5
KEGG_PORPHYRIN_AND_CHLOROPHYLL_METABOLISM
Proportion of pathway genes=0.47; Adjusted p−value=0.3823
Average gene set expression
3
2
1
0
−1
−2
−3
Expression threshold
Below
Above

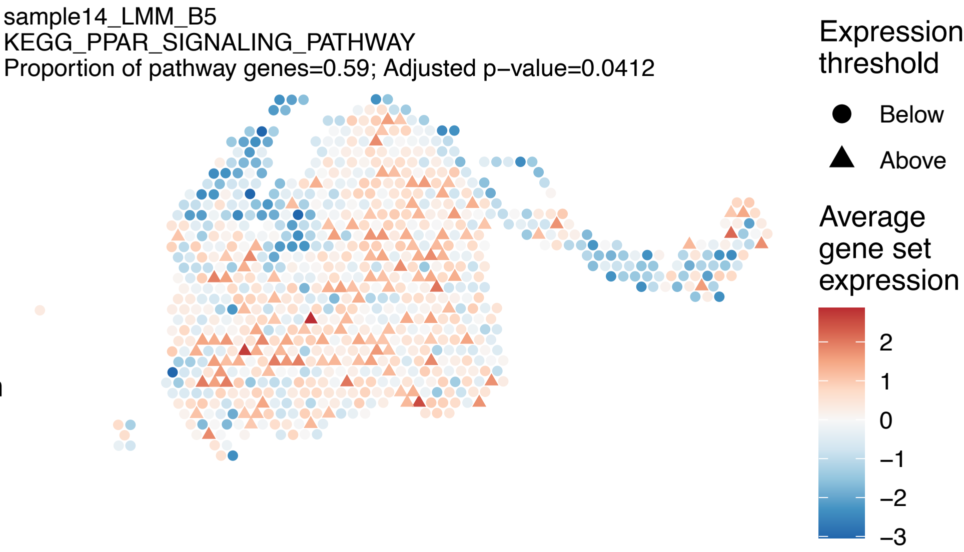
sample14_LMM_B5
KEGG_PPAR_SIGNALING_PATHWAY
Proportion of pathway genes=0.59; Adjusted p−value=0.0412
Expression threshold
Below
Above
Average gene set expression
2
1
0
−1
−2
−3

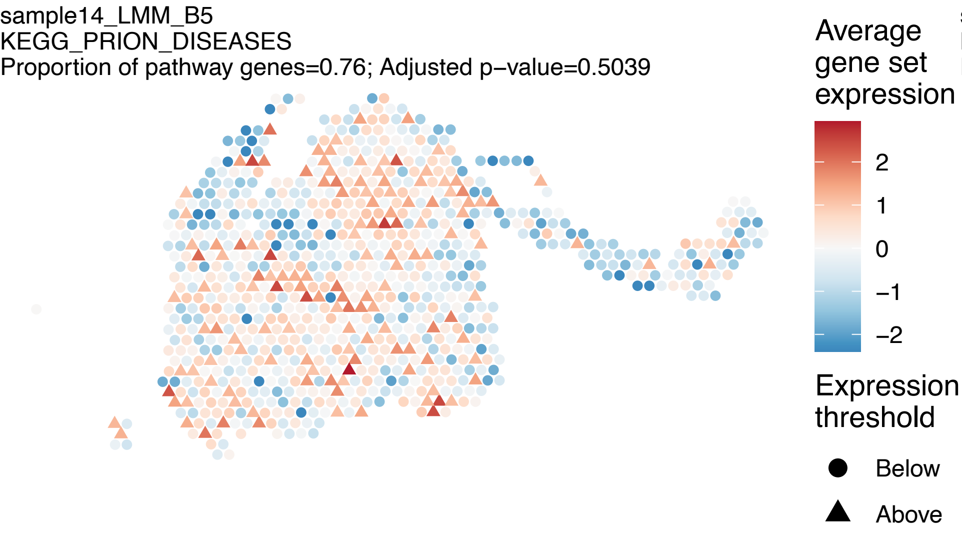
sample14_LMM_B5
KEGG_PRION_DISEASES
Proportion of pathway genes=0.76; Adjusted p−value=0.5039
Average gene set expression
2
1
0
−1
−2
Expression threshold
Below
Above

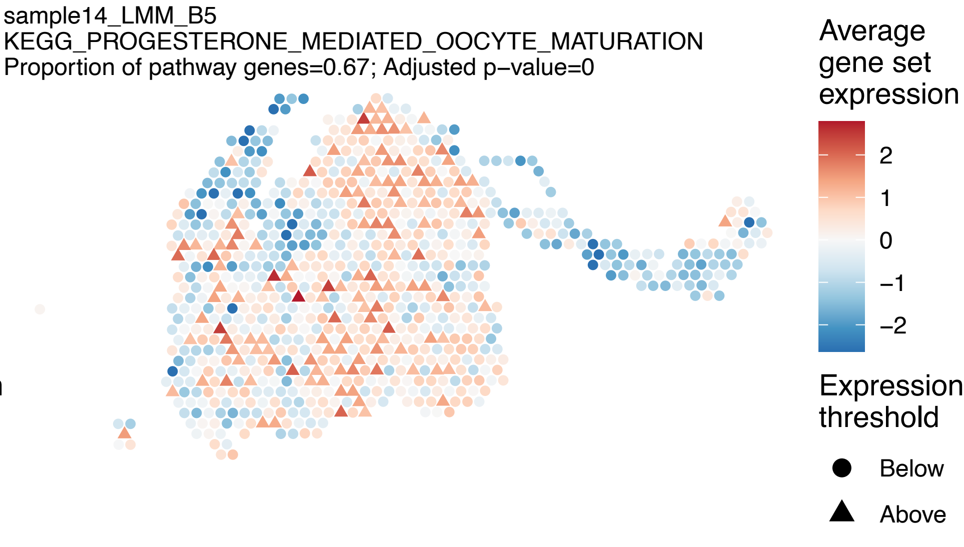
sample14_LMM_B5
KEGG_PROGESTERONE_MEDIATED_OOCYTE_MATURATION
Proportion of pathway genes=0.67; Adjusted p−value=0
Average gene set expression
2
1
0
−1
−2
Expression threshold
Below
Above

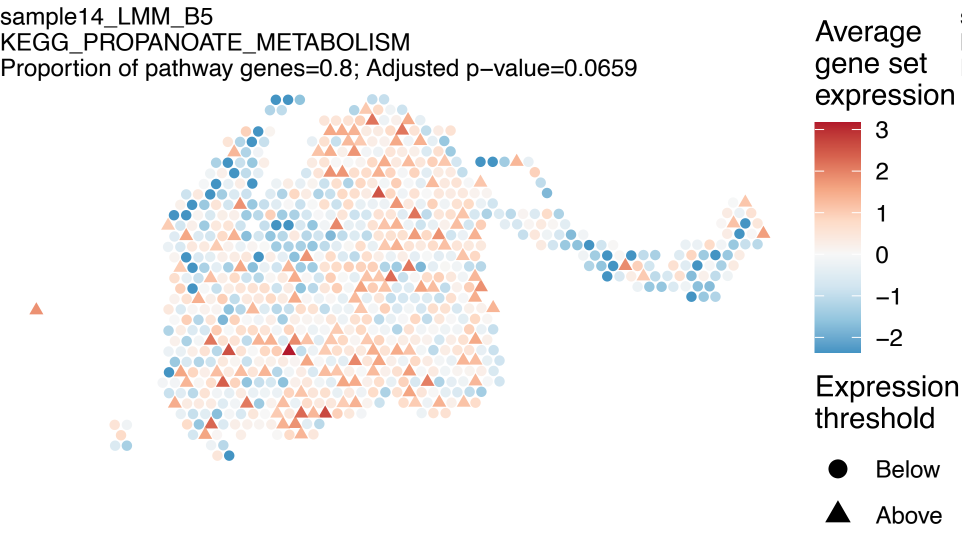
sample14_LMM_B5
KEGG_PROPANOATE_METABOLISM
Proportion of pathway genes=0.8; Adjusted p−value=0.0659
Average gene set expression
3
2
1
0
−1
−2
Expression threshold
Below
Above

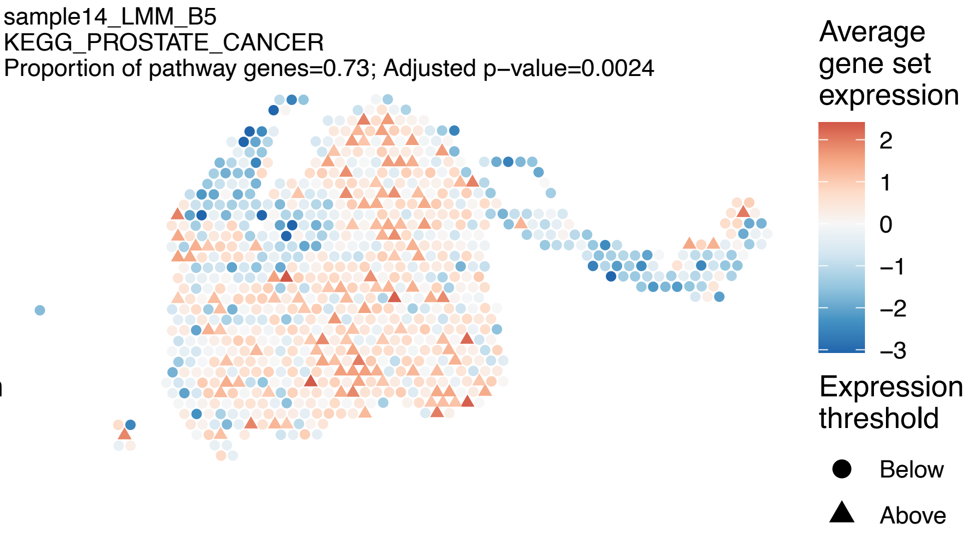
sample14_LMM_B5
KEGG_PROSTATE_CANCER
Proportion of pathway genes=0.73; Adjusted p−value=0.0024
Average gene set expression
2
1
0
−1
−2
−3
Expression threshold
Below
Above

sample14_LMM_B5
KEGG_PROTEASOME
Proportion of pathway genes=0.92; Adjusted p−value=0.0075

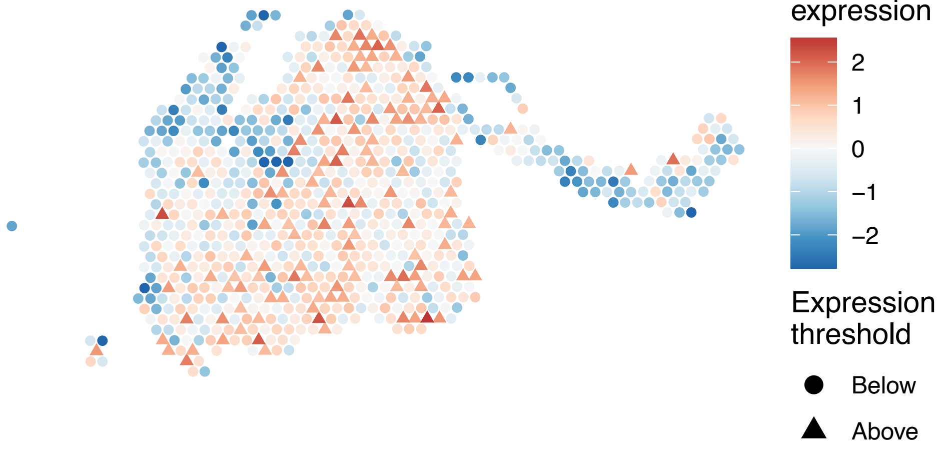


sample14_LMM_B5
KEGG_PROTEIN_EXPORT
Proportion of pathway genes=0.96; Adjusted p−value=0

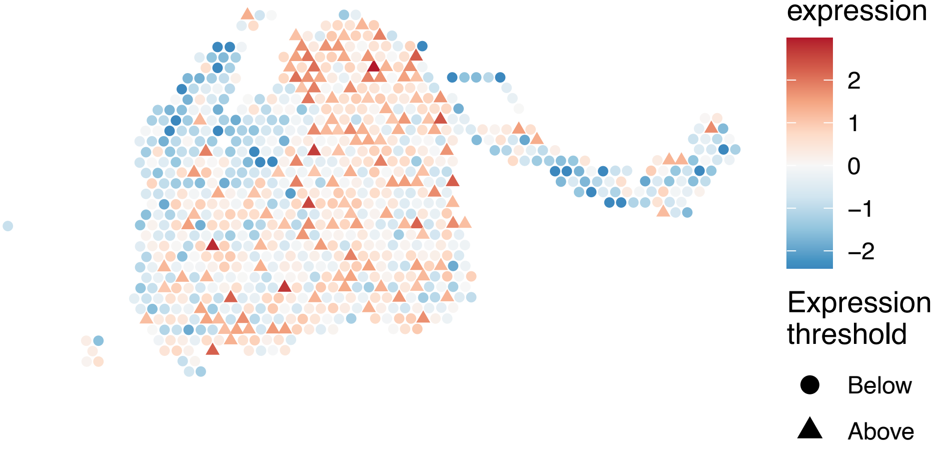


sample14_LMM_B5
KEGG_PROXIMAL_TUBULE_BICARBONATE_RECLAMATION
Proportion of pathway genes=0.5; Adjusted p−value=0.0041

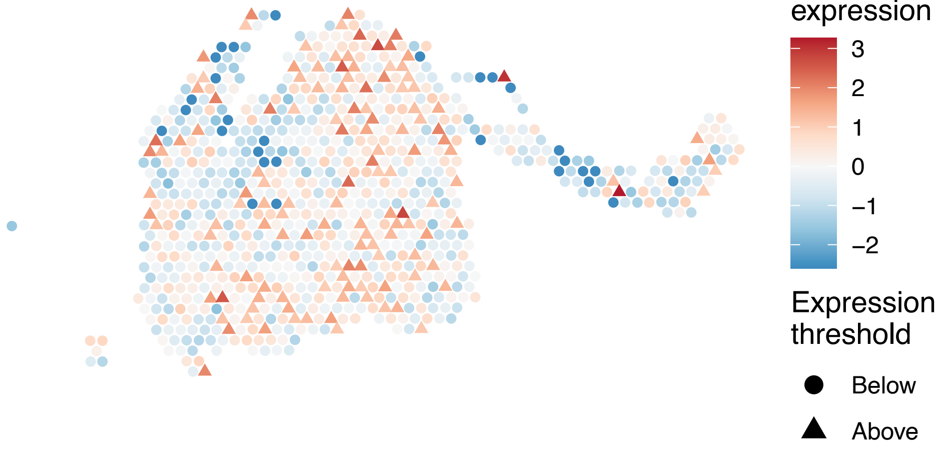


sample14_LMM_B5
KEGG_PURINE_METABOLISM
Proportion of pathway genes=0.65; Adjusted p−value=0

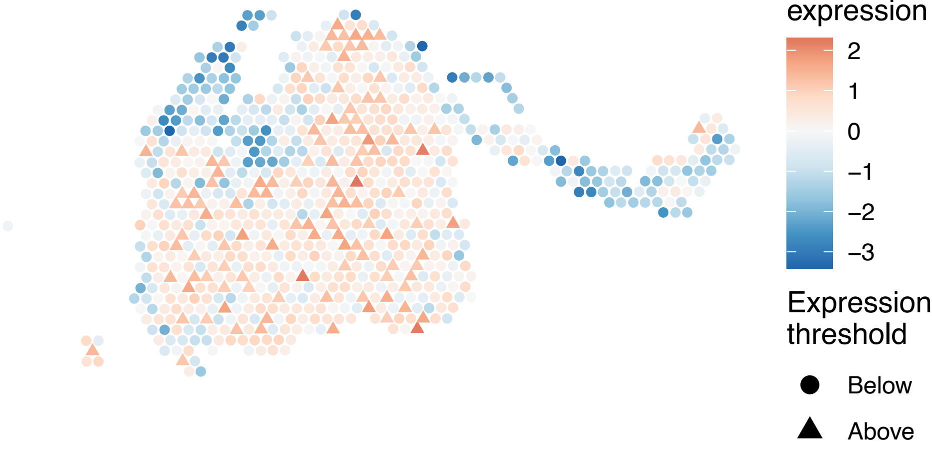

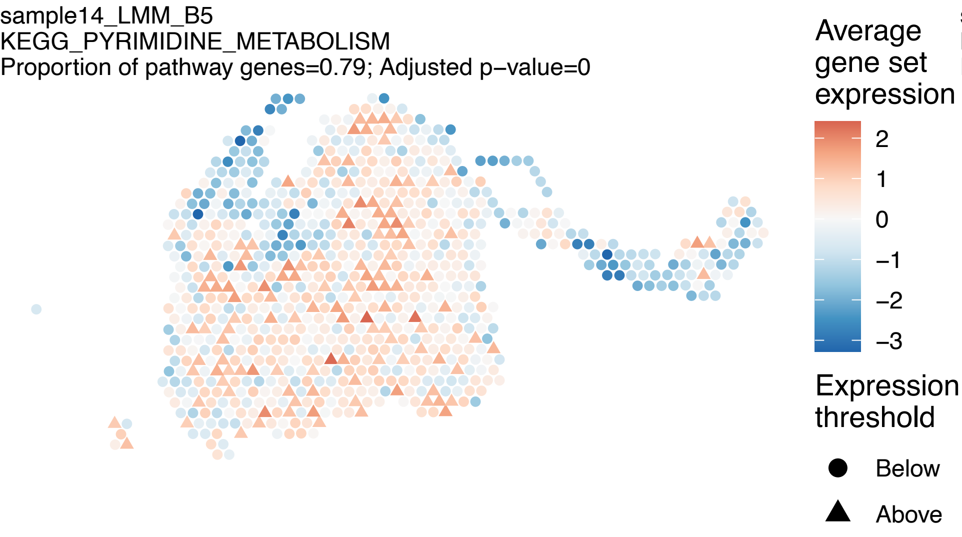
sample14_LMM_B5
KEGG_PYRIMIDINE_METABOLISM
Proportion of pathway genes=0.79; Adjusted p−value=0
Average gene set expression
2
1
0
−1
−2
−3
Expression threshold
Below
Above

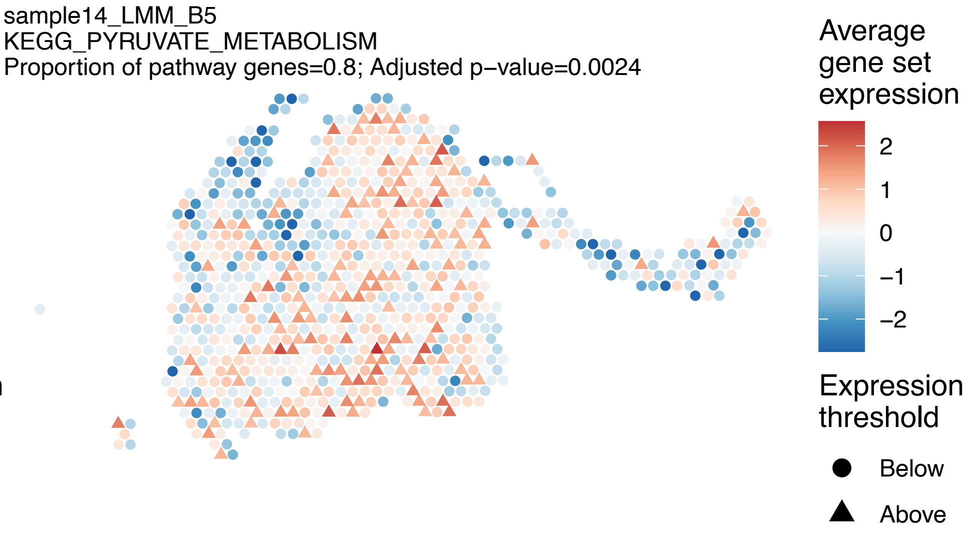
sample14_LMM_B5
KEGG_PYRUVATE_METABOLISM
Proportion of pathway genes=0.8; Adjusted p−value=0.0024
Average gene set expression
2
1
0
−1
−2
Expression threshold
Below
Above

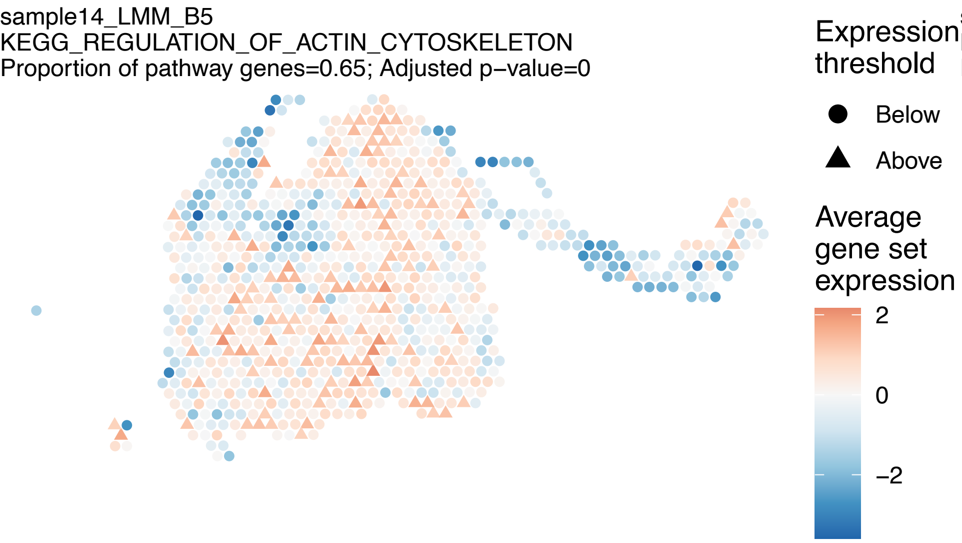
sample14_LMM_B5
KEGG_REGULATION_OF_ACTIN_CYTOSKELETON
Proportion of pathway genes=0.65; Adjusted p−value=0
Expression threshold
Below
Above
Average gene set expression
2
0
−2

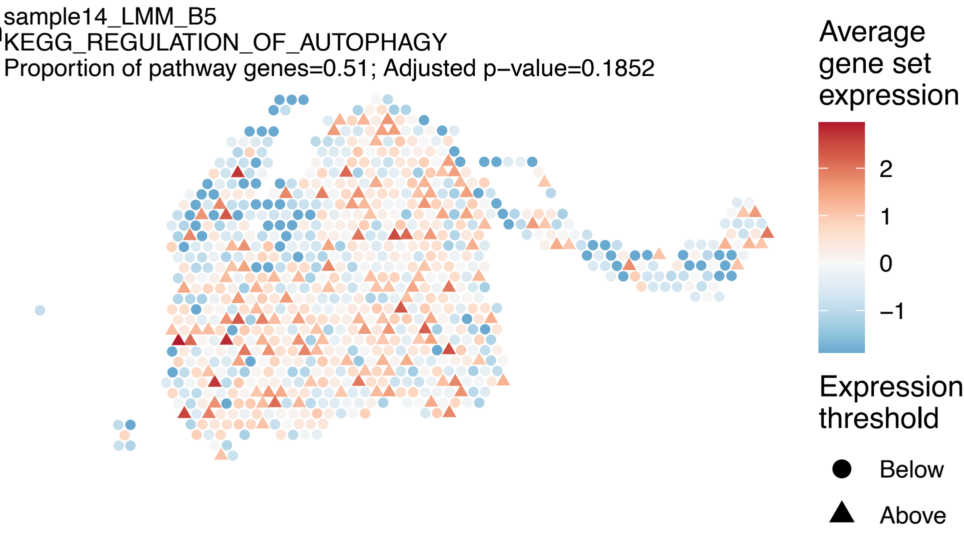
sample14_LMM_B5
KEGG_REGULATION_OF_AUTOPHAGY
Proportion of pathway genes=0.51; Adjusted p−value=0.1852
Average gene set expression
2
1
0
−1
Expression threshold
Below
Above

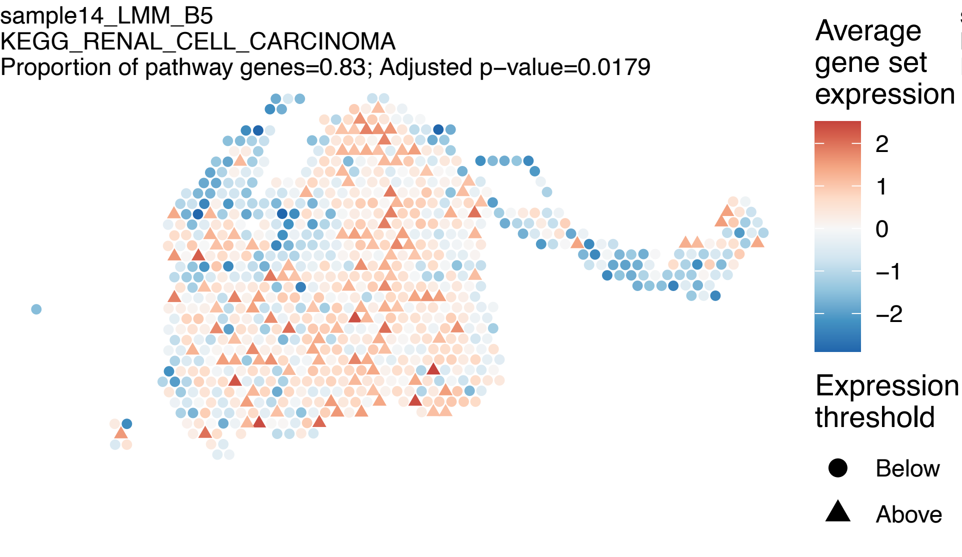
sample14_LMM_B5
KEGG_RENAL_CELL_CARCINOMA
Proportion of pathway genes=0.83; Adjusted p−value=0.0179
Average gene set expression
2
1
0
−1
−2
Expression threshold
Below
Above

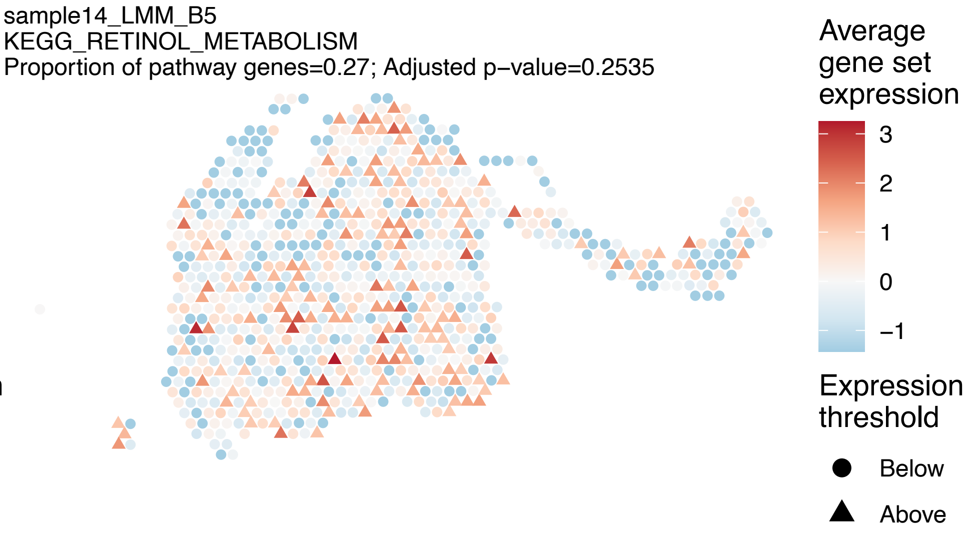
sample14_LMM_B5
KEGG_RETINOL_METABOLISM
Proportion of pathway genes=0.27; Adjusted p−value=0.2535
Average gene set expression
3
2
1
0
−1
Expression threshold
Below
Above

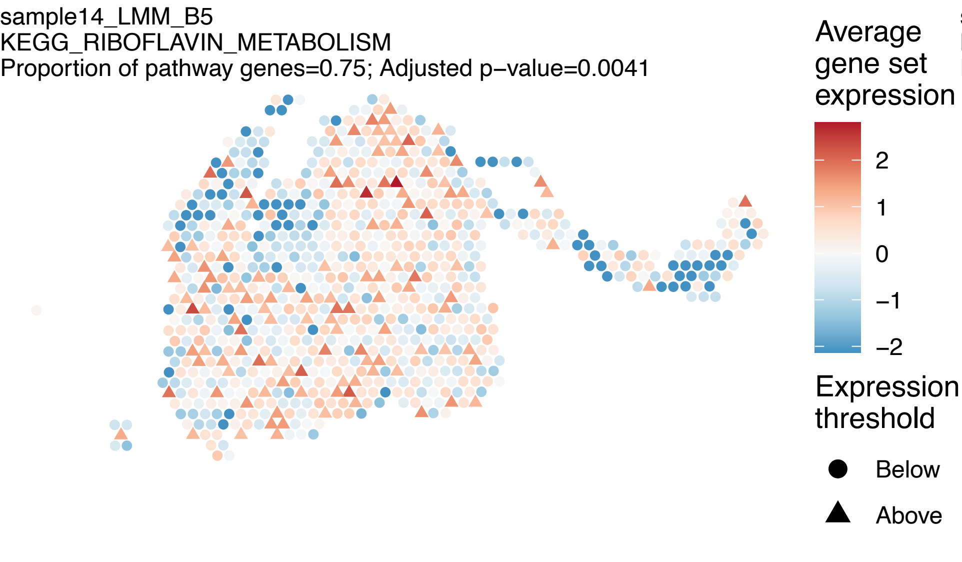
sample14_LMM_B5
KEGG_RIBOFLAVIN_METABOLISM
Proportion of pathway genes=0.75; Adjusted p−value=0.0041
Average gene set expression
2
1
0
−1
−2
Expression threshold
Below
Above

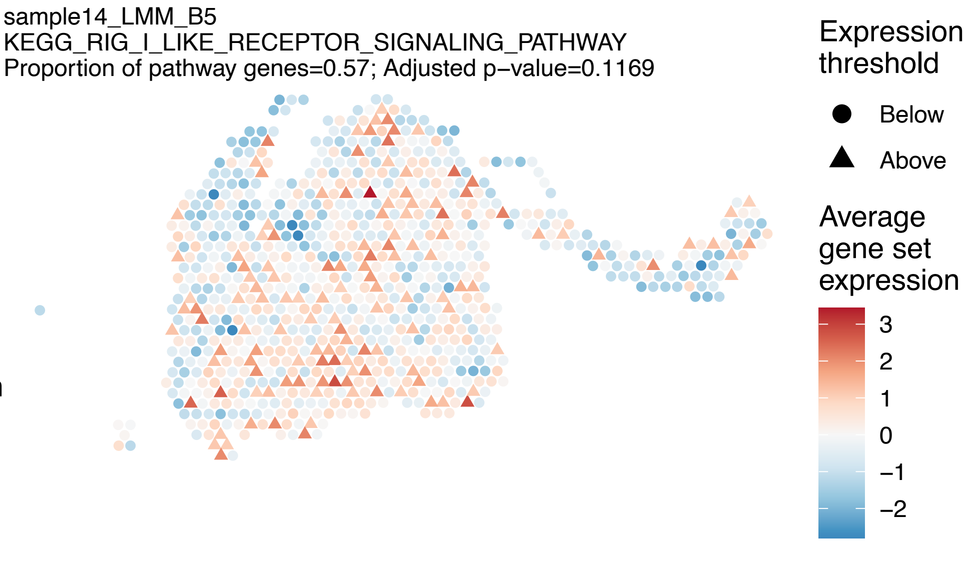
sample14_LMM_B5
KEGG_RIG_I_LIKE_RECEPTOR_SIGNALING_PATHWAY
Proportion of pathway genes=0.57; Adjusted p−value=0.1169
Expression threshold
Below
Above
Average gene set expression
3
2
1
0
−1
−2

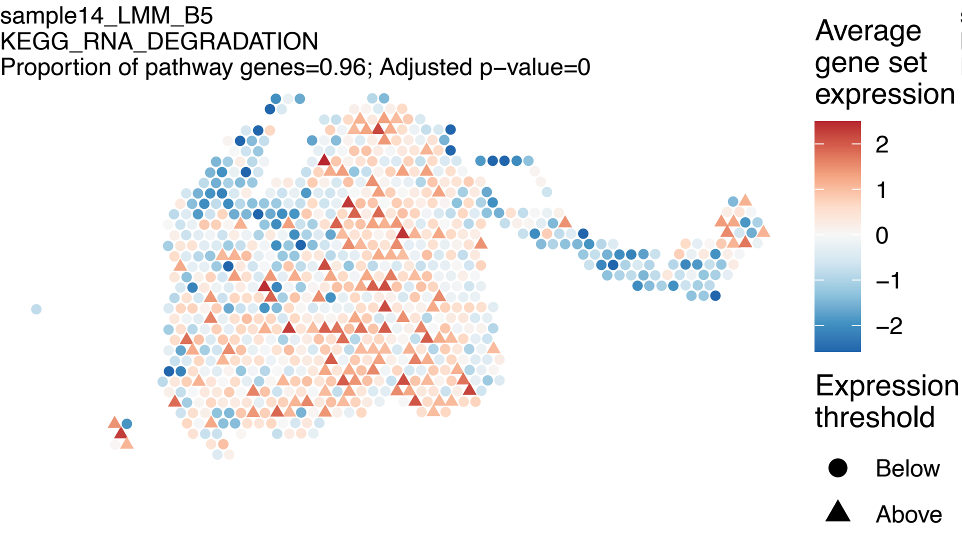
sample14_LMM_B5
KEGG_RNA_DEGRADATION
Proportion of pathway genes=0.96; Adjusted p−value=0
Average gene set expression
2
1
0
−1
−2
Expression threshold
Below
Above

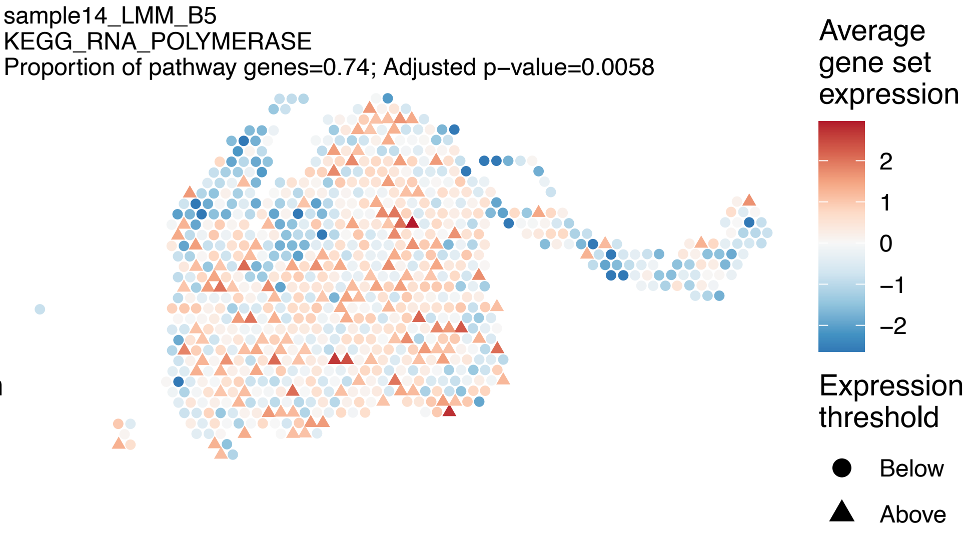
sample14_LMM_B5
KEGG_RNA_POLYMERASE
Proportion of pathway genes=0.74; Adjusted p−value=0.0058
Average gene set expression
2
1
0
−1
−2
Expression threshold
Below
Above

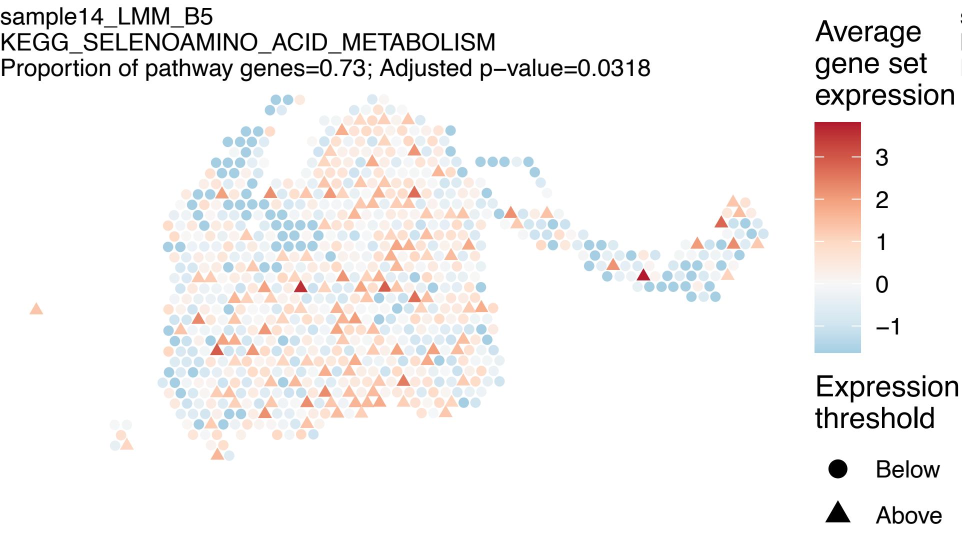
sample14_LMM_B5
KEGG_SELENOAMINO_ACID_METABOLISM
Proportion of pathway genes=0.73; Adjusted p−value=0.0318
Average gene set expression
3
2
1
0
−1
Expression threshold
Below
Above

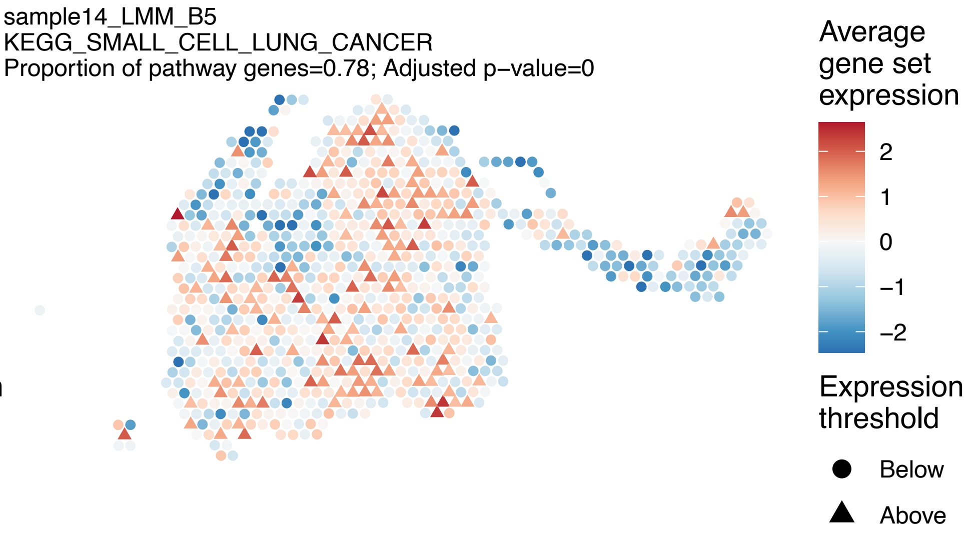
sample14_LMM_B5
KEGG_SMALL_CELL_LUNG_CANCER
Proportion of pathway genes=0.78; Adjusted p−value=0
Average gene set expression
2
1
0
−1
−2
Expression threshold
Below
Above

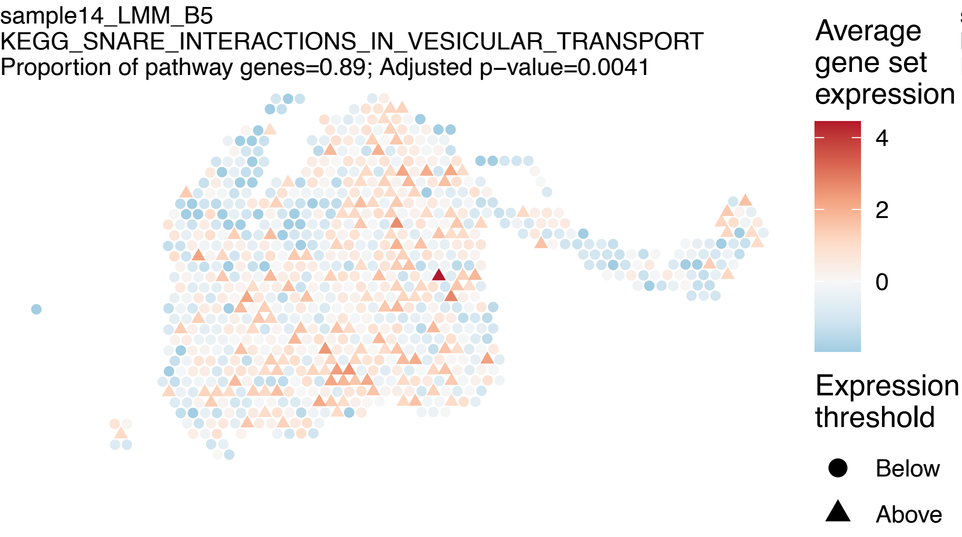
sample14_LMM_B5
KEGG_SNARE_INTERACTIONS_IN_VESICULAR_TRANSPORT
Proportion of pathway genes=0.89; Adjusted p−value=0.0041
Average gene set expression
4
2
0
Expression threshold
Below
Above

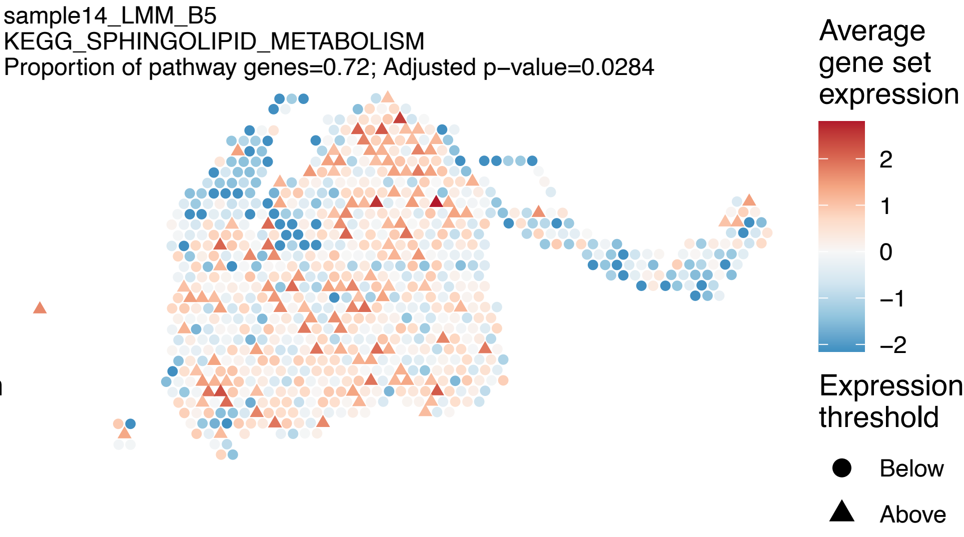
sample14_LMM_B5
KEGG_SPHINGOLIPID_METABOLISM
Proportion of pathway genes=0.72; Adjusted p−value=0.0284
Average gene set expression
2
1
0
−1
−2
Expression threshold
Below
Above

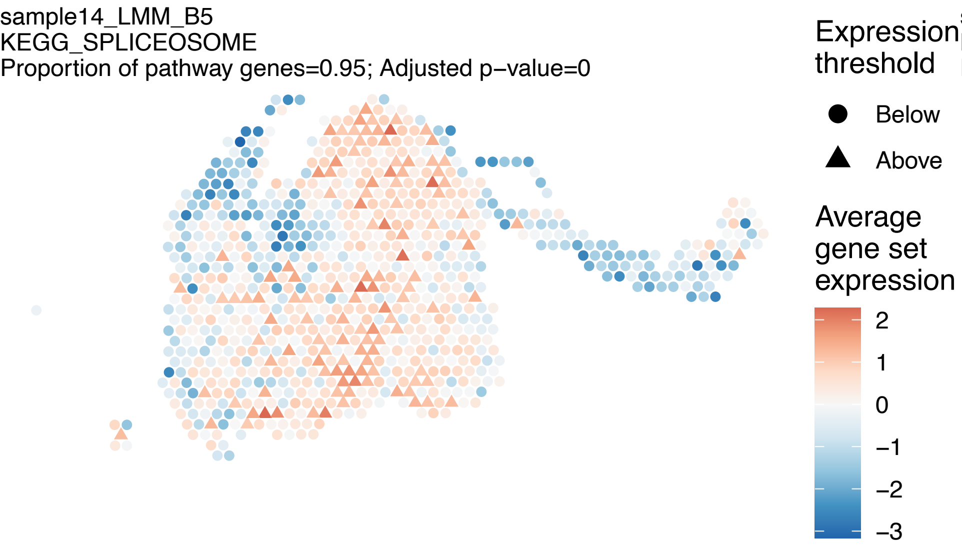
sample14_LMM_B5
KEGG_SPLICEOSOME
Proportion of pathway genes=0.95; Adjusted p−value=0
Expression threshold
Below
Above
Average gene set expression
2
1
0
−1
−2
−3

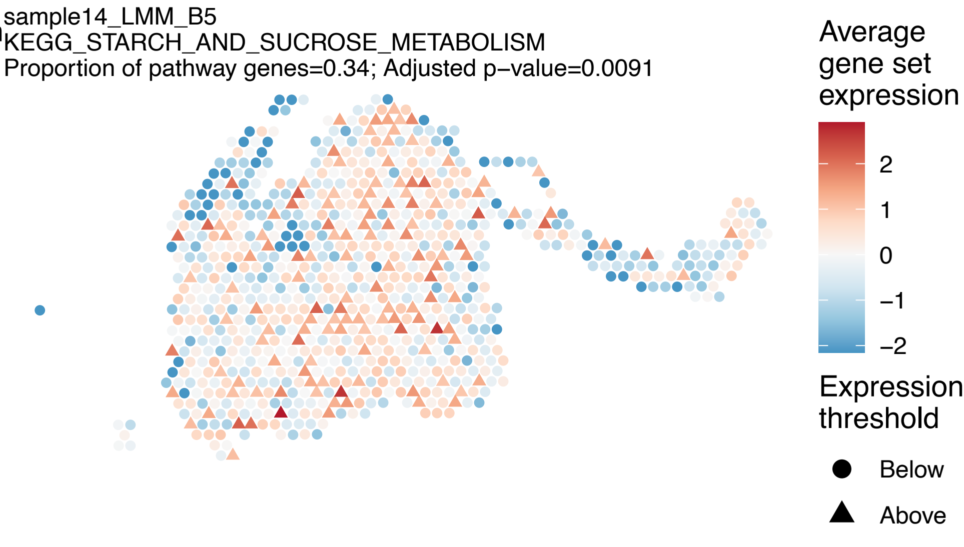
sample14_LMM_B5
KEGG_STARCH_AND_SUCROSE_METABOLISM
Proportion of pathway genes=0.34; Adjusted p−value=0.0091
Average gene set expression
2
1
0
−1
−2
Expression threshold
Below
Above

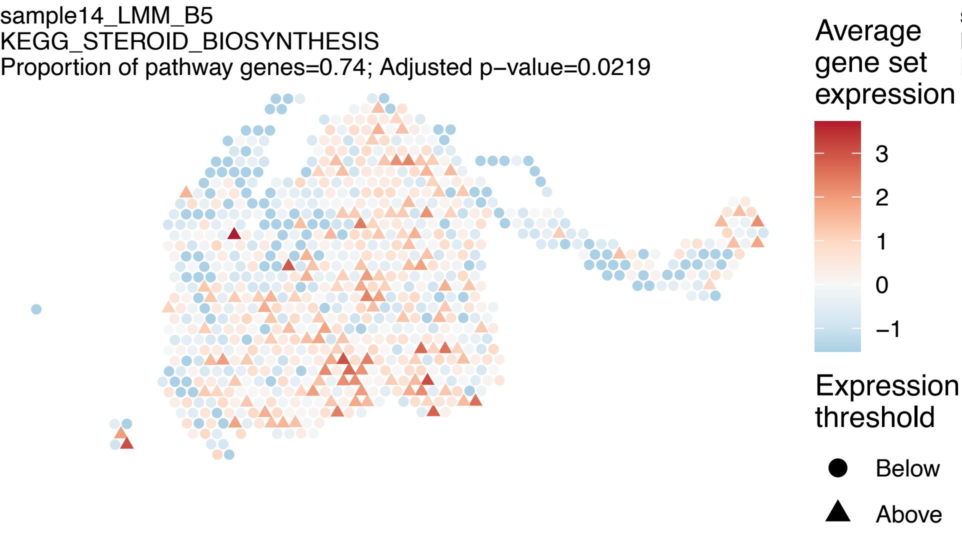
sample14_LMM_B5
KEGG_STEROID_BIOSYNTHESIS
Proportion of pathway genes=0.74; Adjusted p−value=0.0219
Average gene set expression
3
2
1
0
−1
Expression threshold
Below
Above

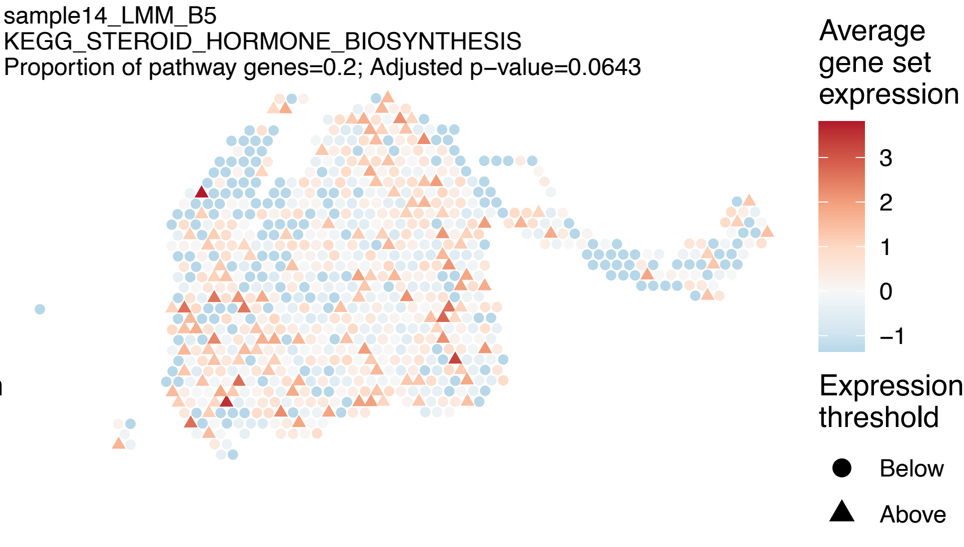
sample14_LMM_B5
KEGG_STEROID_HORMONE_BIOSYNTHESIS
Proportion of pathway genes=0.2; Adjusted p−value=0.0643
Average gene set expression
3
2
1
0
−1
Expression threshold
Below
Above

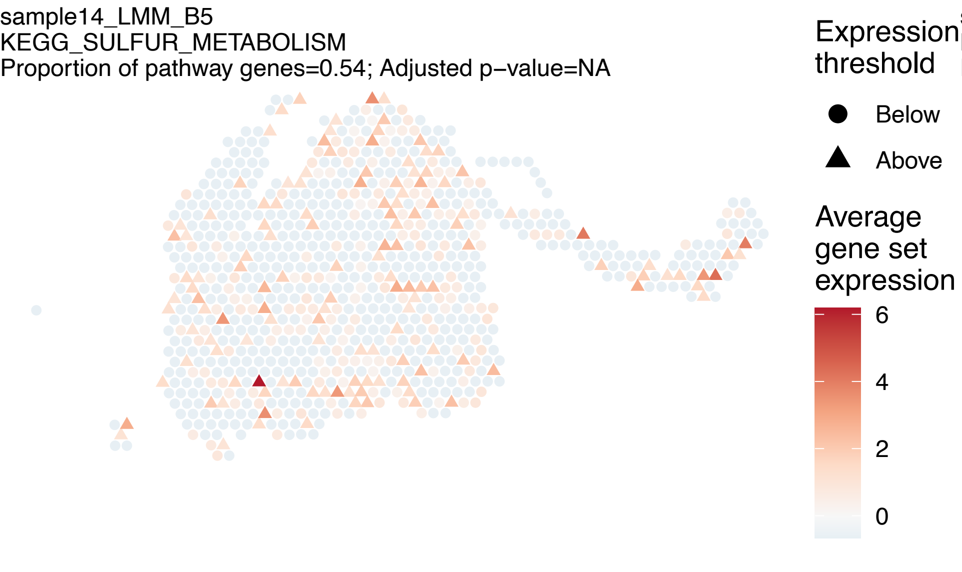
sample14_LMM_B5
KEGG_SULFUR_METABOLISM
Proportion of pathway genes=0.54; Adjusted p−value=NA
Expression threshold
Below
Above
Average gene set expression
6
4
2
0

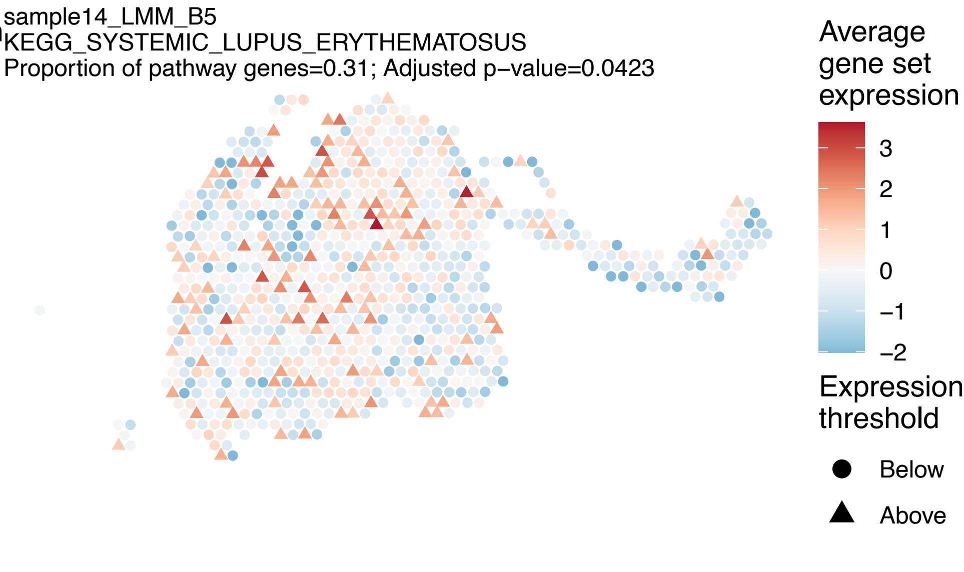
sample14_LMM_B5
KEGG_SYSTEMIC_LUPUS_ERYTHEMATOSUS
Proportion of pathway genes=0.31; Adjusted p−value=0.0423
Average gene set expression
3
2
1
0
−1
−2
Expression threshold
Below
Above

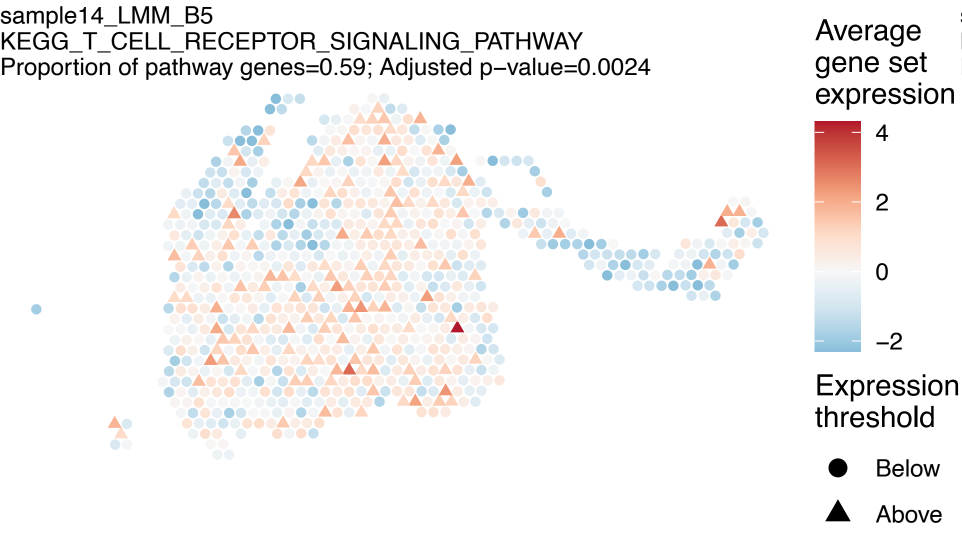
sample14_LMM_B5
KEGG_T_CELL_RECEPTOR_SIGNALING_PATHWAY
Proportion of pathway genes=0.59; Adjusted p−value=0.0024
Average gene set expression
4
2
0
−2
Expression threshold
Below
Above

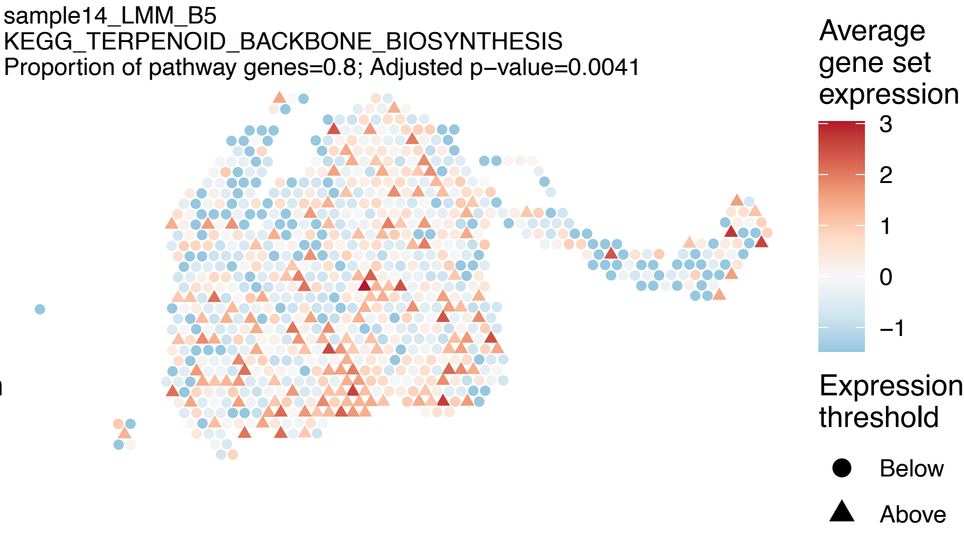
sample14_LMM_B5
KEGG_TERPENOID_BACKBONE_BIOSYNTHESIS
Proportion of pathway genes=0.8; Adjusted p−value=0.0041
Average gene set expression
3
2
1
0
−1
Expression threshold
Below
Above

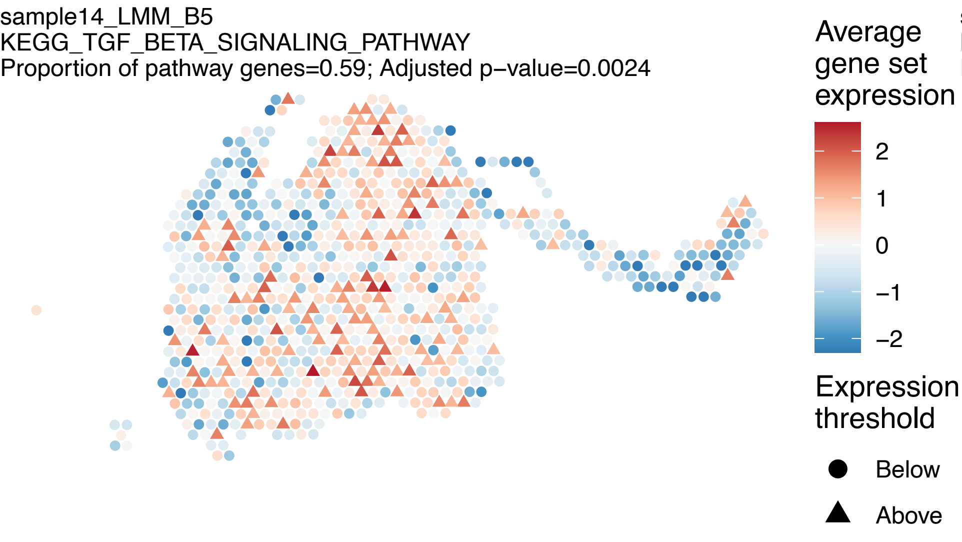
sample14_LMM_B5
KEGG_TGF_BETA_SIGNALING_PATHWAY
Proportion of pathway genes=0.59; Adjusted p−value=0.0024
Average gene set expression
2
1
0
−1
−2
Expression threshold
Below
Above

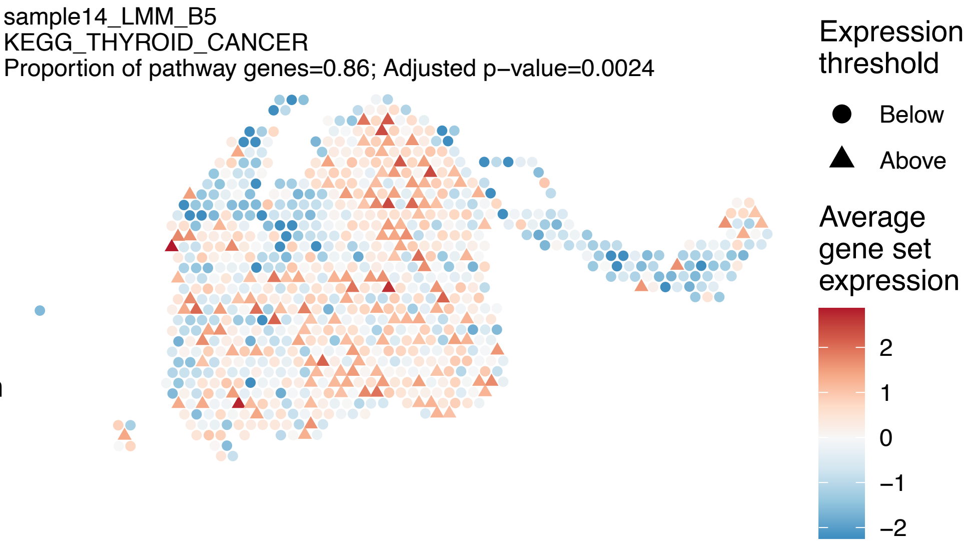
sample14_LMM_B5
KEGG_THYROID_CANCER
Proportion of pathway genes=0.86; Adjusted p−value=0.0024
Expression threshold
Below
Above
Average gene set expression
2
1
0
−1
−2

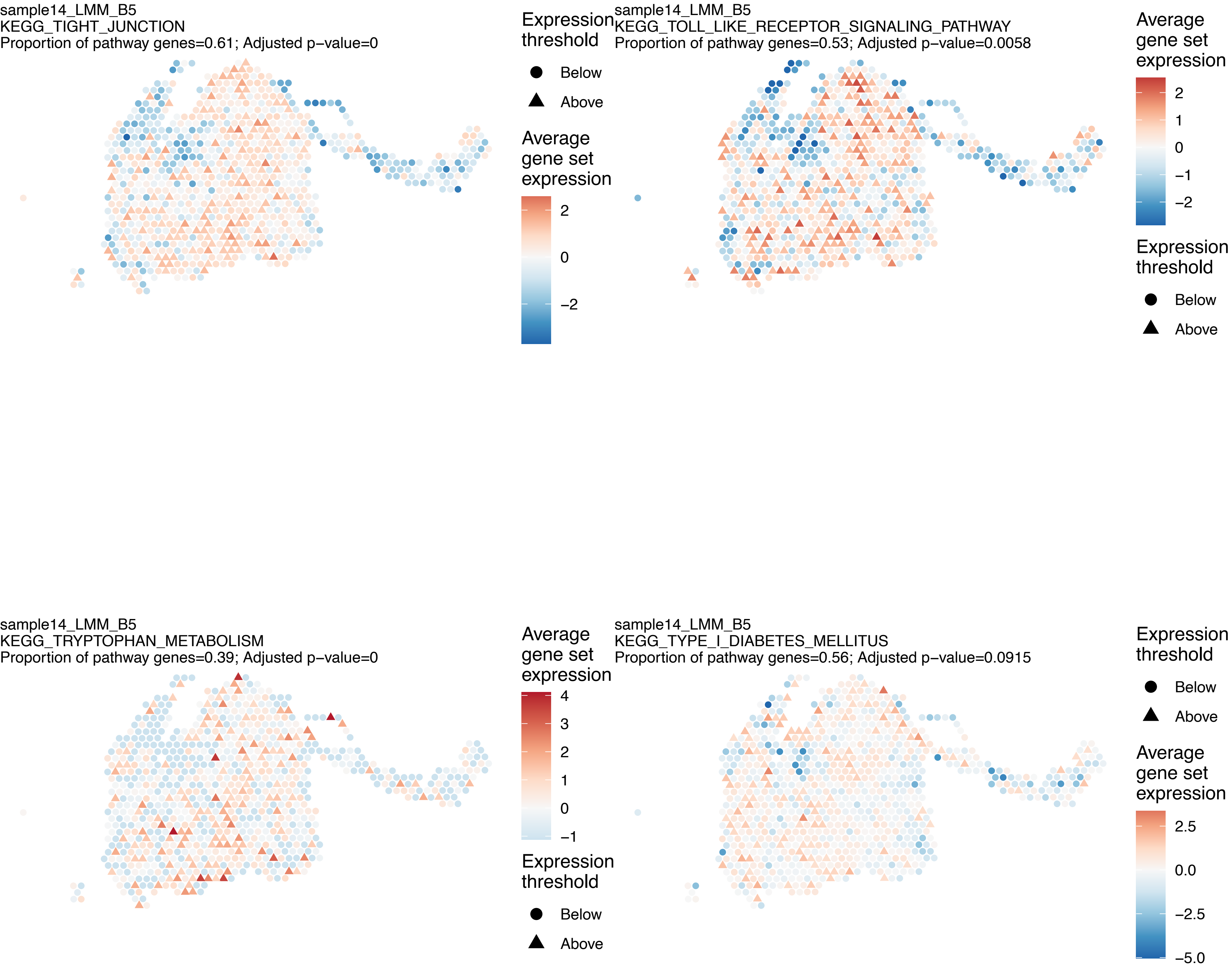
sample14_LMM_B5
KEGG_TIGHT_JUNCTION
Proportion of pathway genes=0.61; Adjusted p−value=0
Expression threshold
Below
Above
Average gene set expression
2
0
−2
sample14_LMM_B5
KEGG_TOLL_LIKE_RECEPTOR_SIGNALING_PATHWAY
Proportion of pathway genes=0.53; Adjusted p−value=0.0058
Average gene set expression
2
1
0
−1
−2
Expression threshold
Below
Above
sample14_LMM_B5
KEGG_TRYPTOPHAN_METABOLISM
Proportion of pathway genes=0.39; Adjusted p−value=0
Average gene set expression
4
3
2
1
0
−1
Expression threshold
Below
Above
sample14_LMM_B5
KEGG_TYPE_I_DIABETES_MELLITUS
Proportion of pathway genes=0.56; Adjusted p−value=0.0915
Expression threshold
Below
Above
Average gene set expression
2.5
0.0
−2.5
−5.0

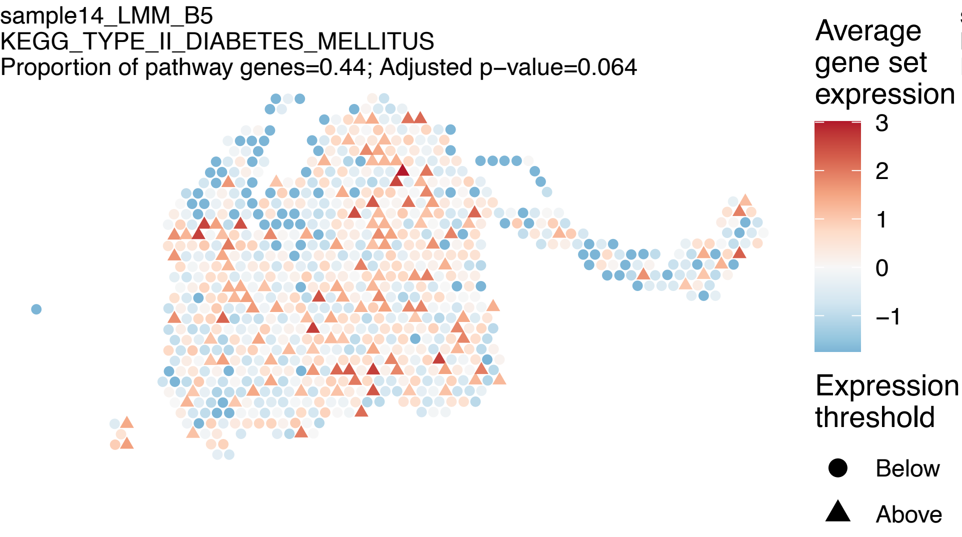
sample14_LMM_B5
KEGG_TYPE_II_DIABETES_MELLITUS
Proportion of pathway genes=0.44; Adjusted p−value=0.064
Average gene set expression
3
2
1
0
−1
Expression threshold
Below
Above

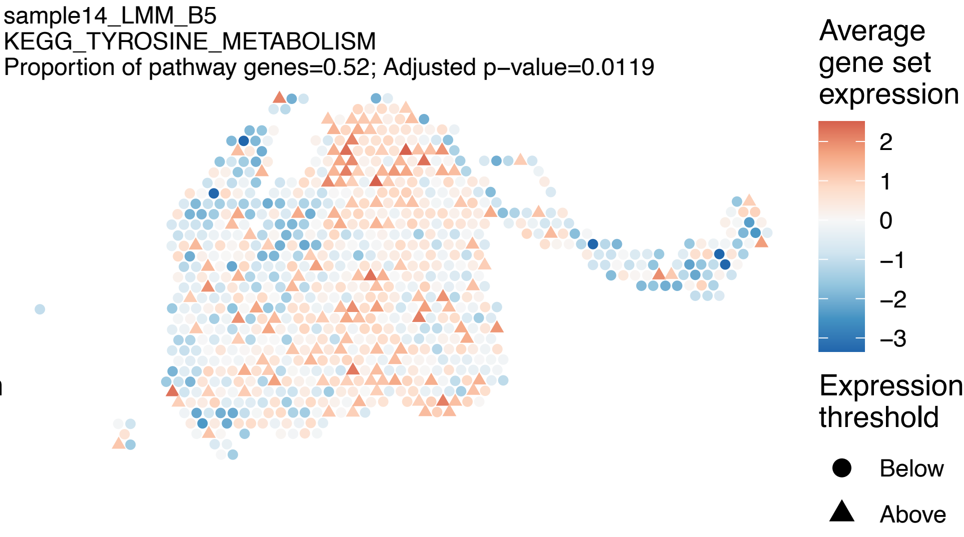
sample14_LMM_B5
KEGG_TYROSINE_METABOLISM
Proportion of pathway genes=0.52; Adjusted p−value=0.0119
Average gene set expression
2
1
0
−1
−2
−3
Expression threshold
Below
Above

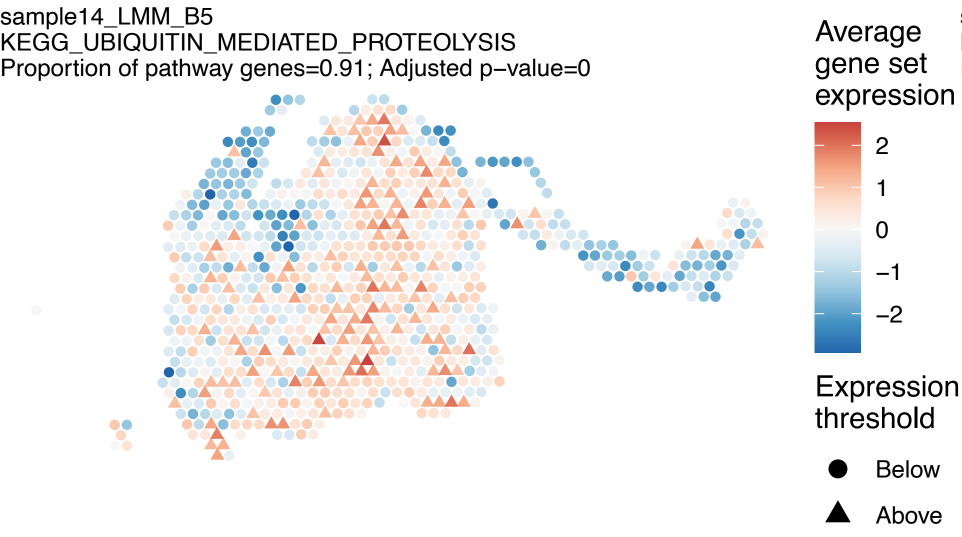
sample14_LMM_B5
KEGG_UBIQUITIN_MEDIATED_PROTEOLYSIS
Proportion of pathway genes=0.91; Adjusted p−value=0
Average gene set expression
2
1
0
−1
−2
Expression threshold
Below
Above

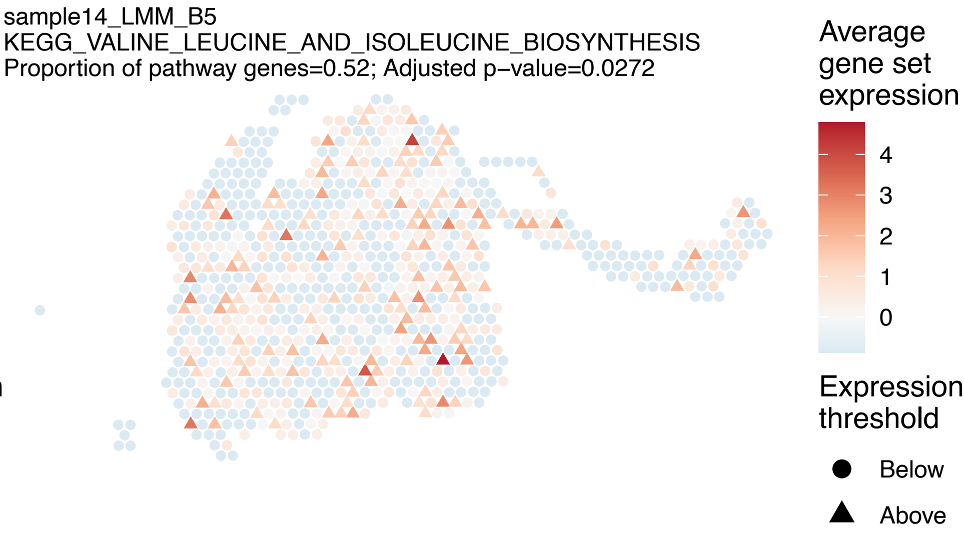
sample14_LMM_B5
KEGG_VALINE_LEUCINE_AND_ISOLEUCINE_BIOSYNTHESIS
Proportion of pathway genes=0.52; Adjusted p−value=0.0272
Average gene set expression
4
3
2
1
0
Expression threshold
Below
Above

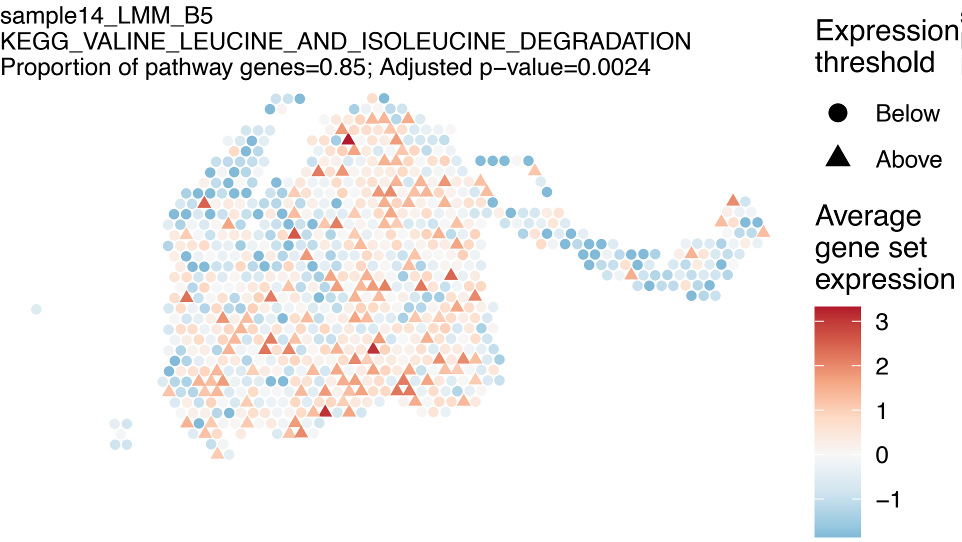
sample14_LMM_B5
KEGG_VALINE_LEUCINE_AND_ISOLEUCINE_DEGRADATION
Proportion of pathway genes=0.85; Adjusted p−value=0.0024
Expression threshold
Below
Above
Average gene set expression
3
2
1
0
−1

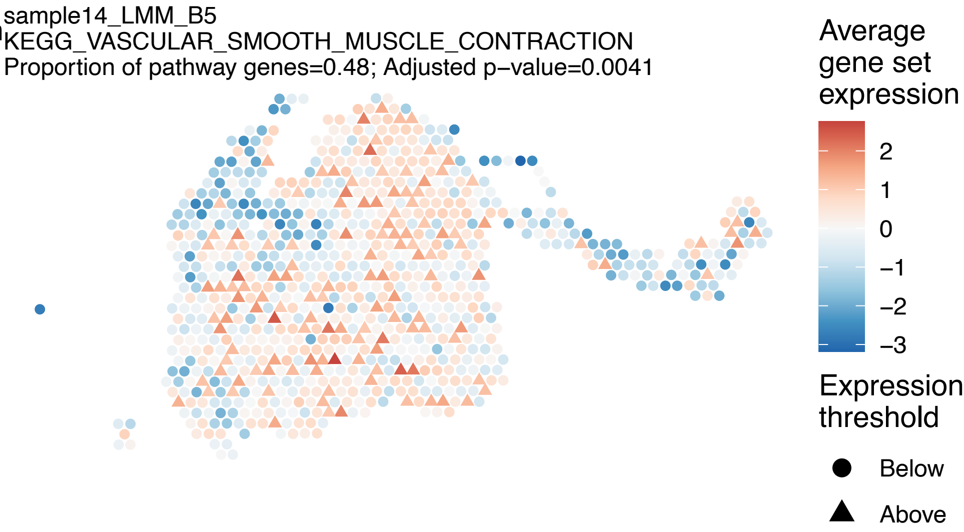
sample14_LMM_B5
KEGG_VASCULAR_SMOOTH_MUSCLE_CONTRACTION
Proportion of pathway genes=0.48; Adjusted p−value=0.0041
Average gene set expression
2
1
0
−1
−2
−3
Expression threshold
Below
Above

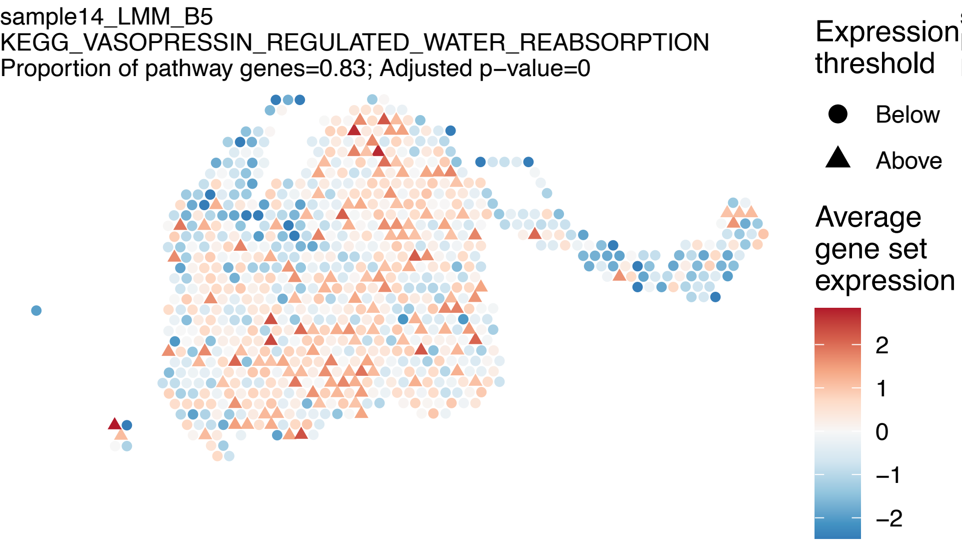
sample14_LMM_B5
KEGG_VASOPRESSIN_REGULATED_WATER_REABSORPTION
Proportion of pathway genes=0.83; Adjusted p−value=0
Expression threshold
Below
Above
Average gene set expression
2
1
0
−1
−2

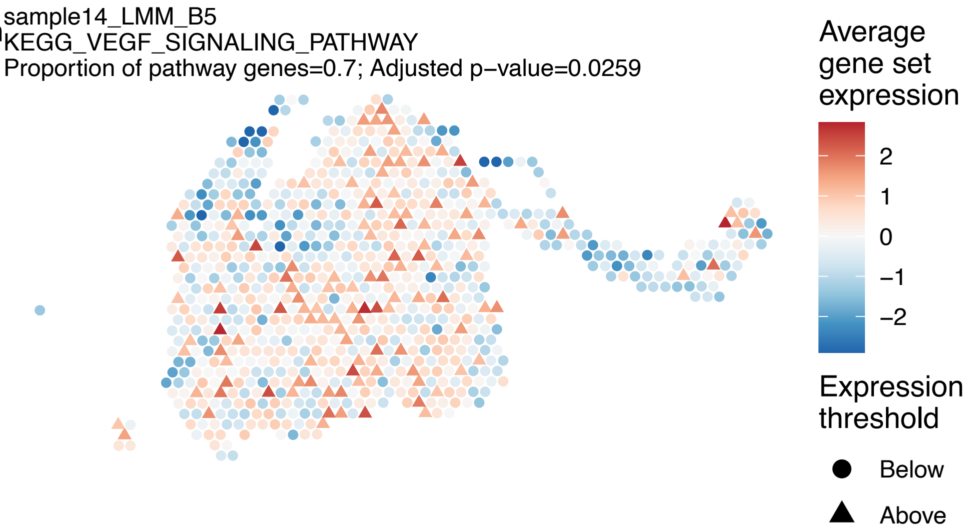
sample14_LMM_B5
KEGG_VEGF_SIGNALING_PATHWAY
Proportion of pathway genes=0.7; Adjusted p−value=0.0259
Average gene set expression
2
1
0
−1
−2
Expression threshold
Below
Above

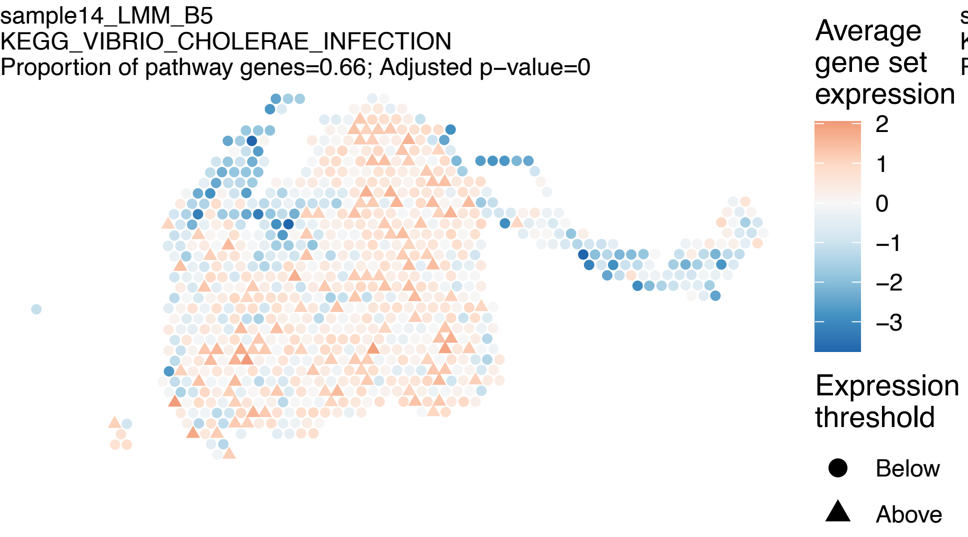
sample14_LMM_B5
KEGG_VIBRIO_CHOLERAE_INFECTION
Proportion of pathway genes=0.66; Adjusted p−value=0
Average gene set expression
2
1
0
−1
−2
−3
Expression threshold
Below
Above

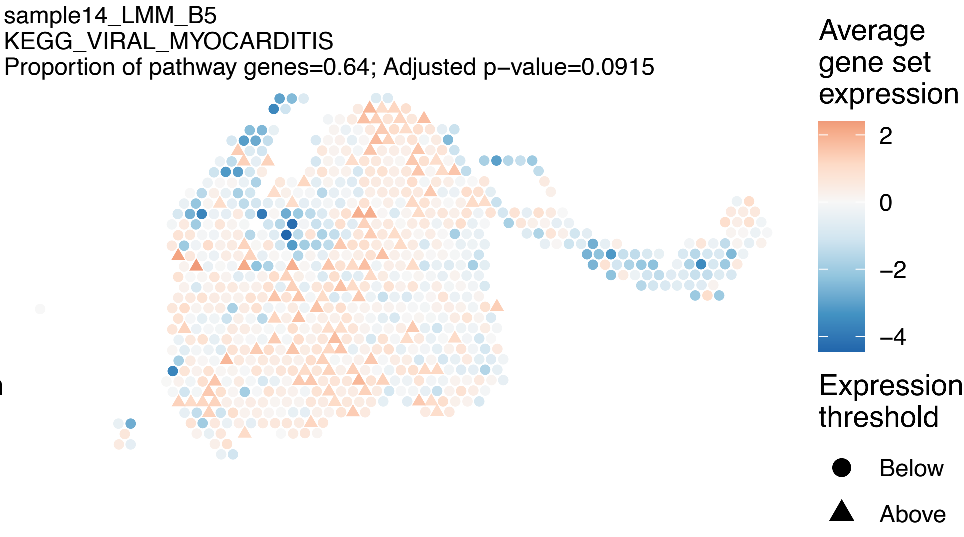
sample14_LMM_B5
KEGG_VIRAL_MYOCARDITIS
Proportion of pathway genes=0.64; Adjusted p−value=0.0915
Average gene set expression
2
0
−2
−4
Expression threshold
Below
Above

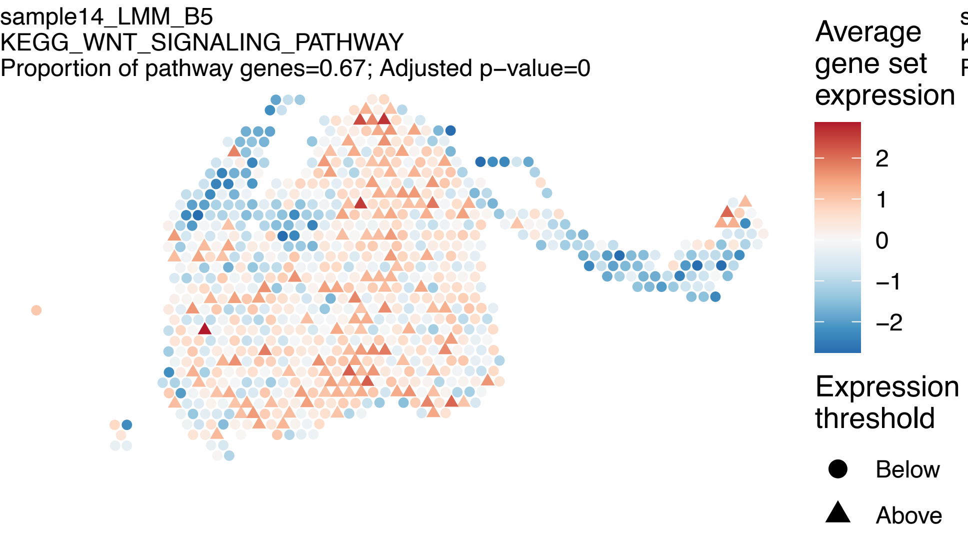
sample14_LMM_B5
KEGG_WNT_SIGNALING_PATHWAY
Proportion of pathway genes=0.67; Adjusted p−value=0
Average gene set expression
2
1
0
−1
−2
Expression threshold
Below
Above

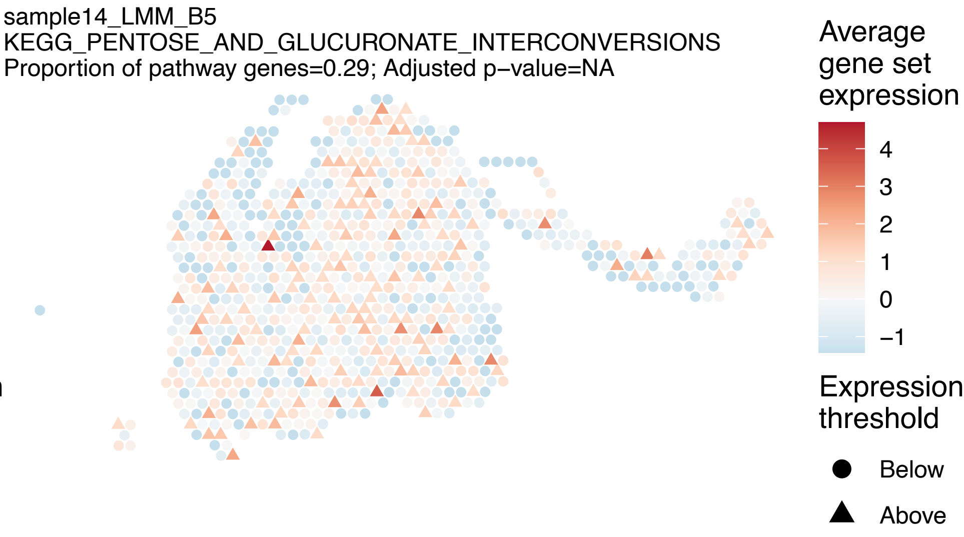
sample14_LMM_B5
KEGG_PENTOSE_AND_GLUCURONATE_INTERCONVERSIONS
Proportion of pathway genes=0.29; Adjusted p−value=NA
Average gene set expression
4
3
2
1
0
−1
Expression threshold
Below
Above

sample15_LMM_B6
KEGG_NATURAL_KILLER_CELL_MEDIATED_CYTOTOXICITY
Proportion of pathway genes=0.24; Adjusted p−value=0

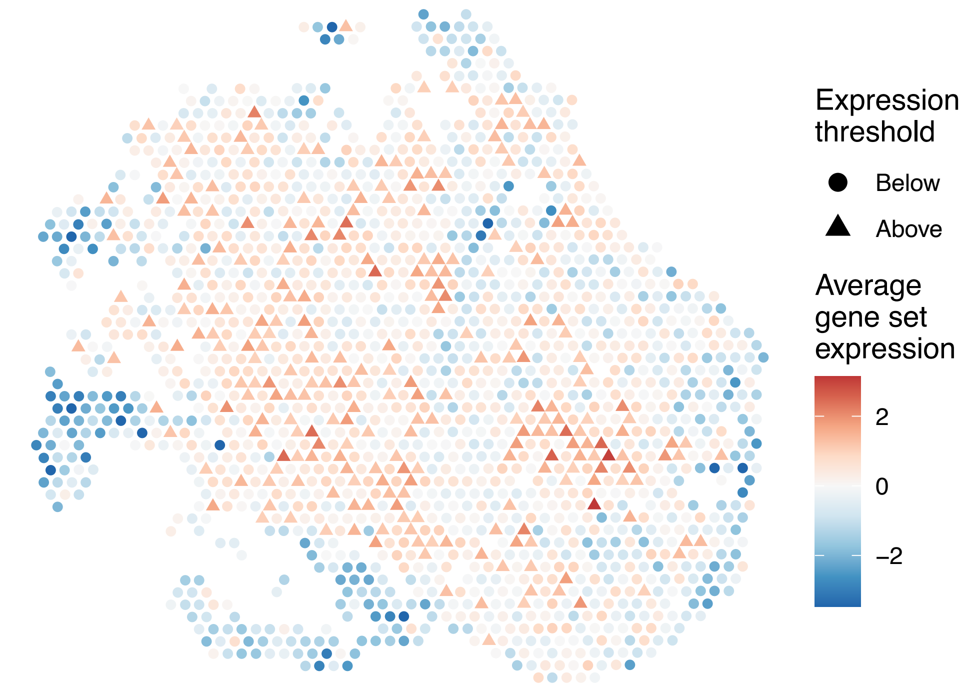


sample15_LMM_B6
KEGG_RIBOSOME
Proportion of pathway genes=0.97; Adjusted p−value=0.2474

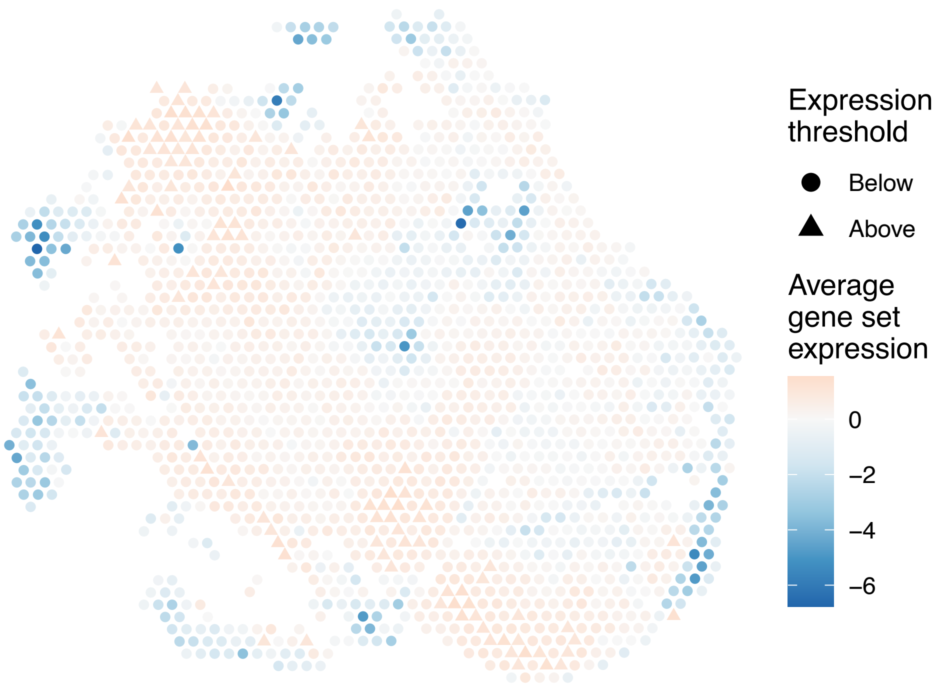


sample15_LMM_B6
KEGG_PRIMARY_IMMUNODEFICIENCY
Proportion of pathway genes=0.58; Adjusted p−value=0

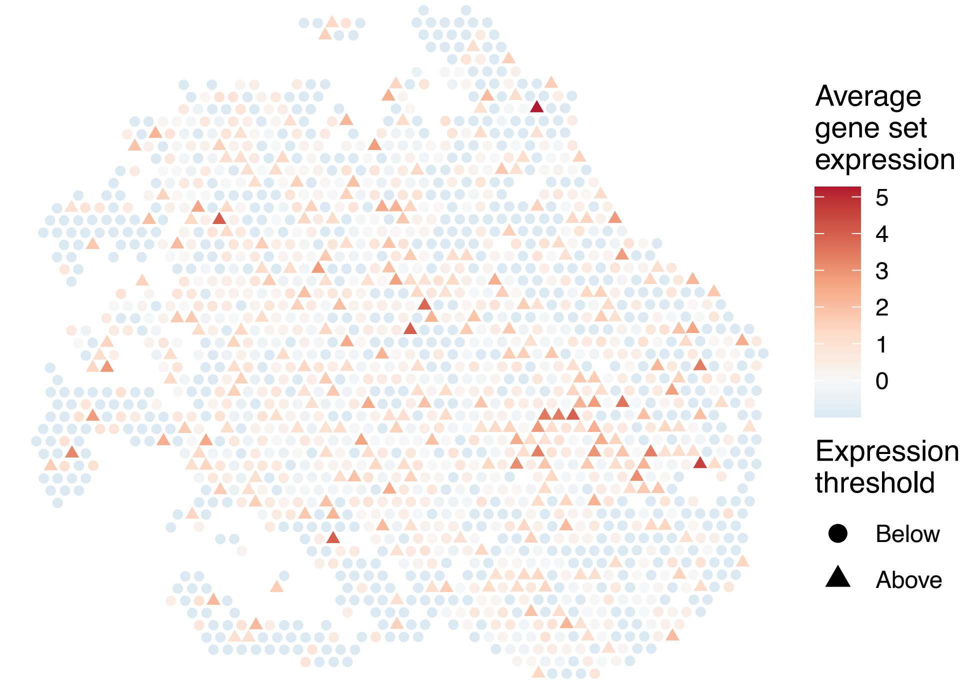


sample15_LMM_B6
KEGG_ANTIGEN_PROCESSING_AND_PRESENTATION
Proportion of pathway genes=0.3; Adjusted p−value=0

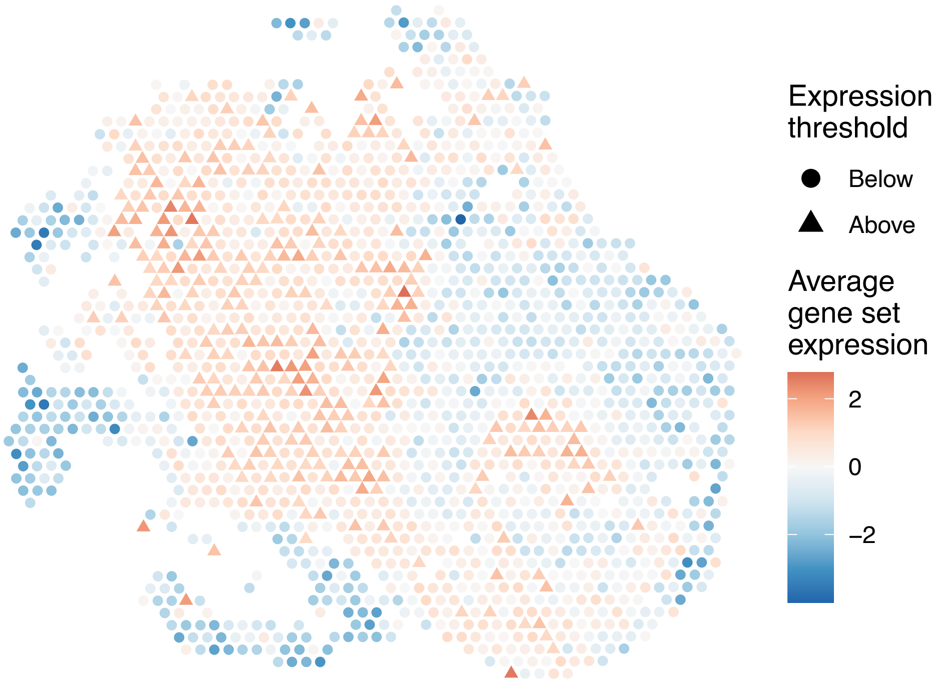

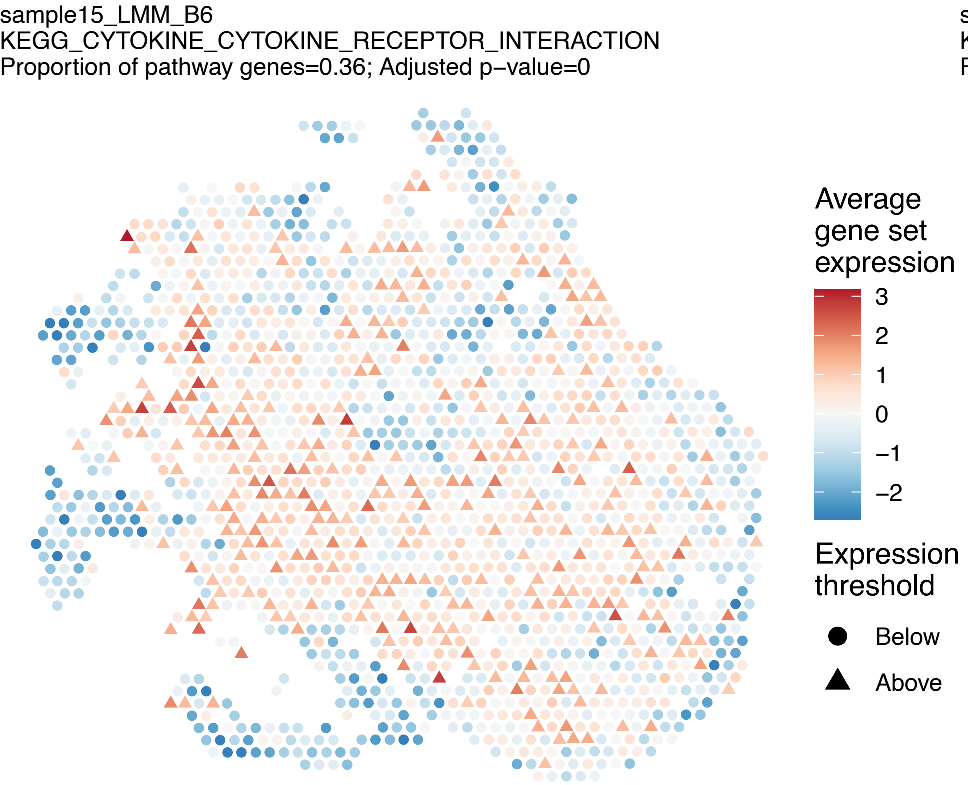
sample15_LMM_B6
KEGG_CYTOKINE_CYTOKINE_RECEPTOR_INTERACTION
Proportion of pathway genes=0.36; Adjusted p−value=0
Average gene set expression
3
2
1
0
−1
−2
Expression threshold
Below
Above

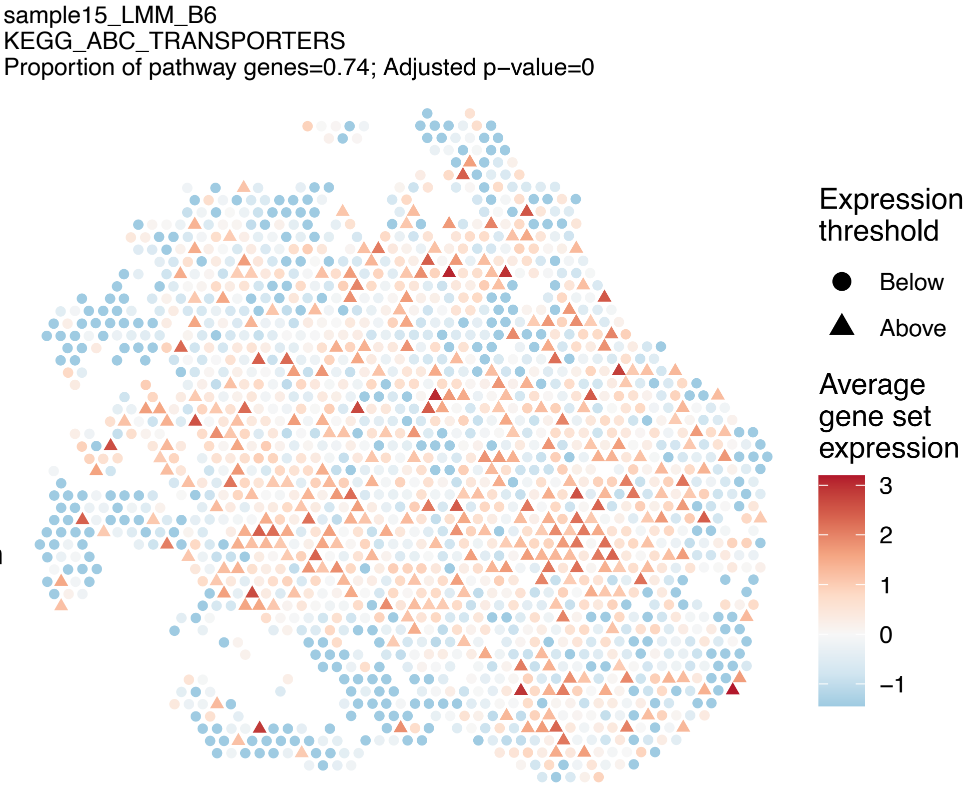
sample15_LMM_B6
KEGG_ABC_TRANSPORTERS
Proportion of pathway genes=0.74; Adjusted p−value=0
Expression threshold
Below
Above
Average gene set expression
3
2
1
0
−1

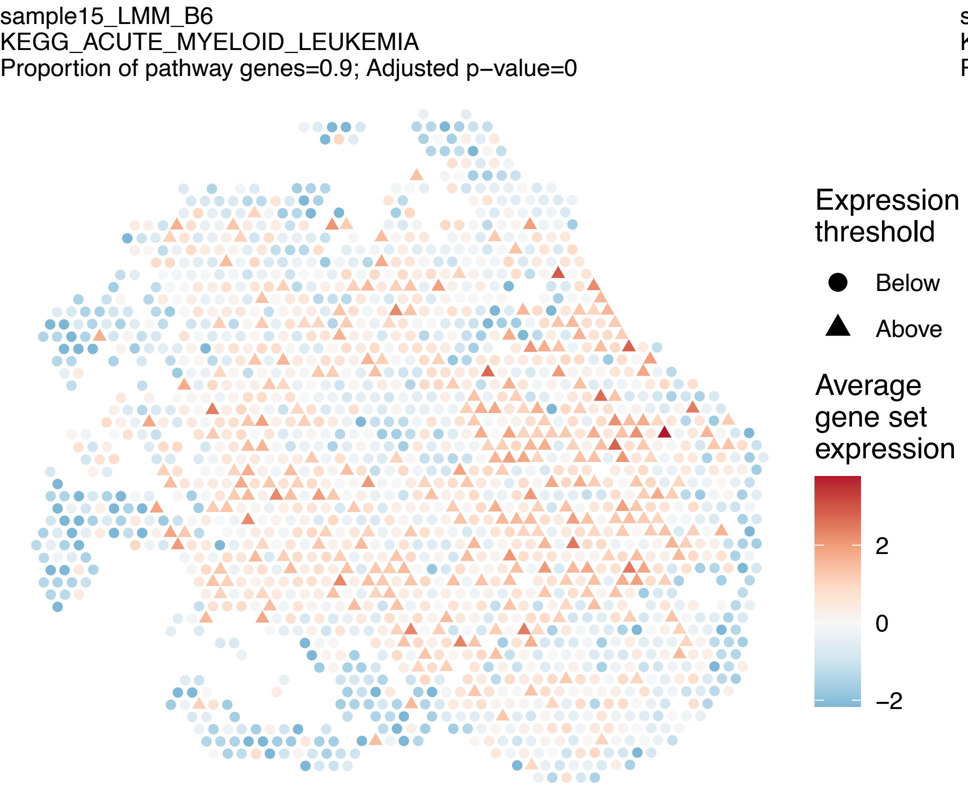
sample15_LMM_B6
KEGG_ACUTE_MYELOID_LEUKEMIA
Proportion of pathway genes=0.9; Adjusted p−value=0
Expression threshold
Below
Above
Average gene set expression
2
0
−2

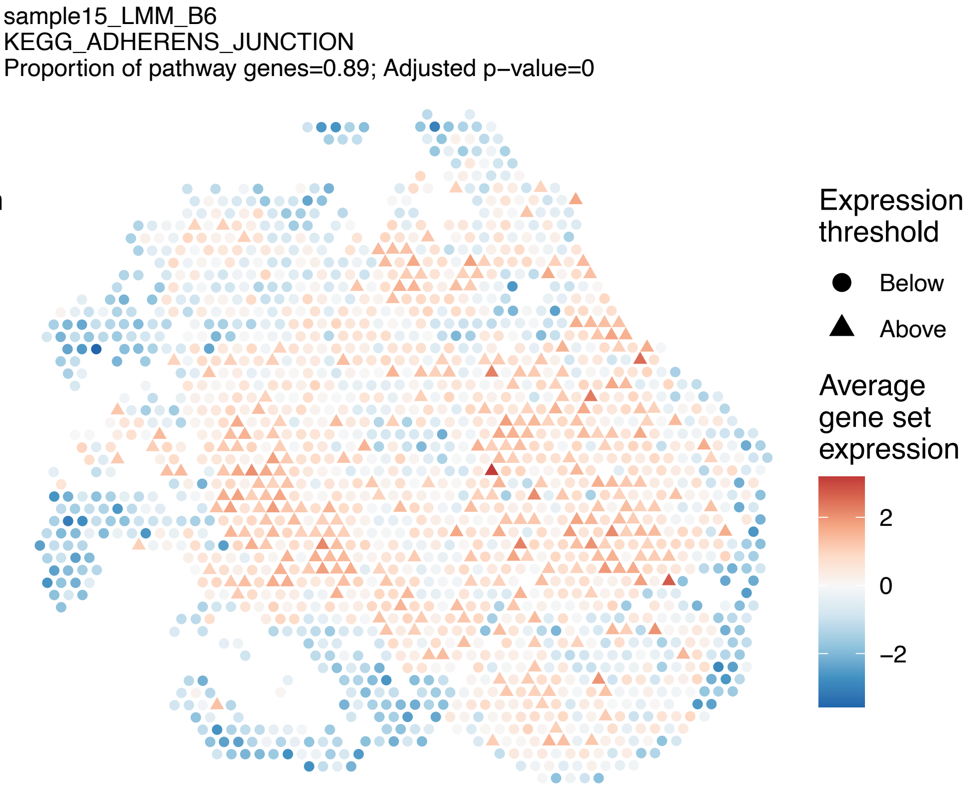
sample15_LMM_B6
KEGG_ADHERENS_JUNCTION
Proportion of pathway genes=0.89; Adjusted p−value=0
Expression threshold
Below
Above
Average gene set expression
2
0
−2

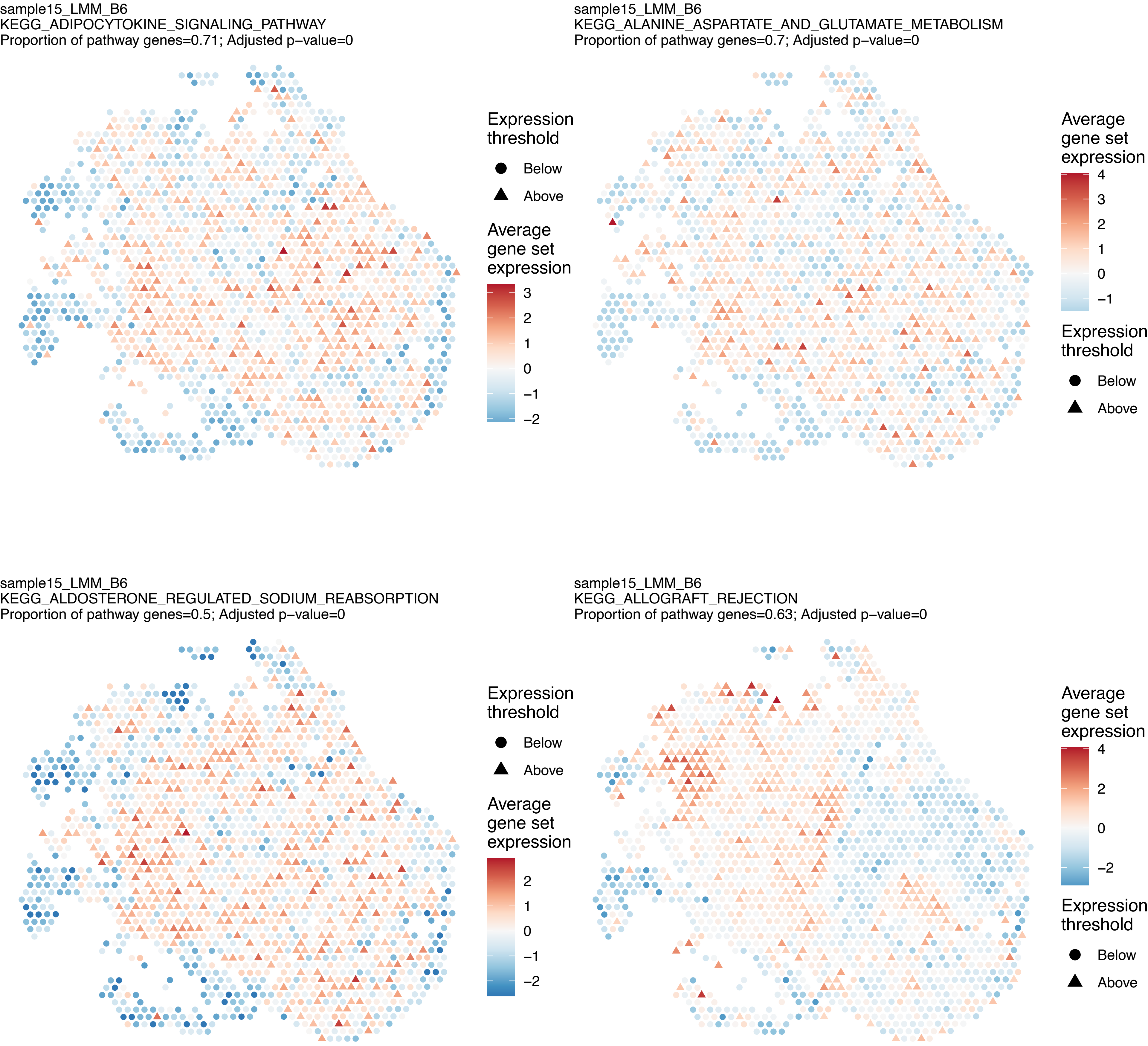

sample15_LMM_B6
KEGG_ADIPOCYTOKINE_SIGNALING_PATHWAY
Proportion of pathway genes=0.71; Adjusted p−value=0
Expression threshold
Below
Above
Average gene set expression
3
2
1
0
−1
−2
sample15_LMM_B6
KEGG_ALANINE_ASPARTATE_AND_GLUTAMATE_METABOLISM
Proportion of pathway genes=0.7; Adjusted p−value=0
Average gene set expression
4
3
2
1
0
−1
Expression threshold
Below
Above
sample15_LMM_B6
KEGG_ALDOSTERONE_REGULATED_SODIUM_REABSORPTION
Proportion of pathway genes=0.5; Adjusted p−value=0
Expression threshold
Below
Above
Average gene set expression
2
1
0
−1
−2
sample15_LMM_B6
KEGG_ALLOGRAFT_REJECTION
Proportion of pathway genes=0.63; Adjusted p−value=0
Average gene set expression
4
2
0
−2
Expression threshold
Below
Above

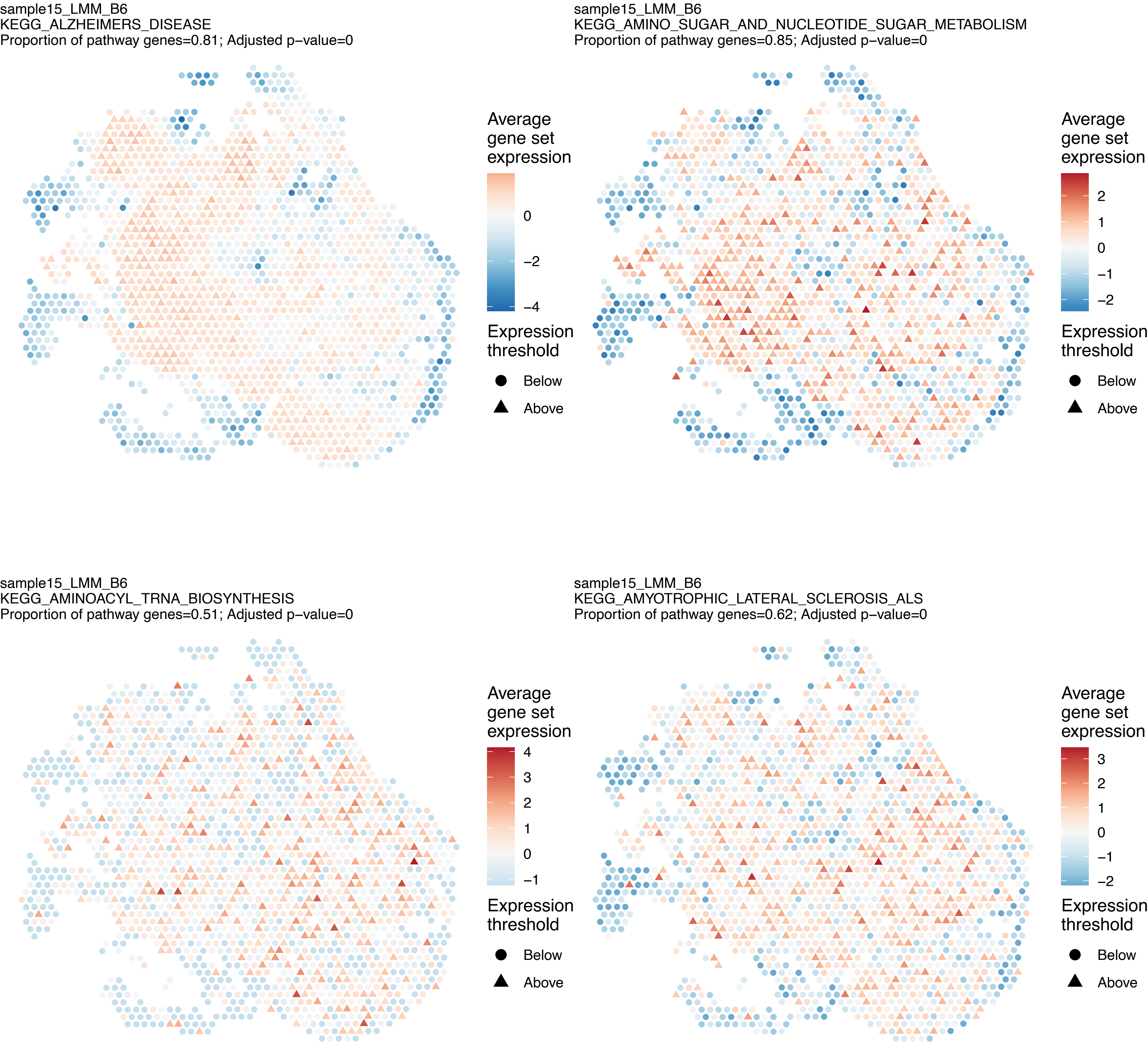
sample15_LMM_B6
KEGG_ALZHEIMERS_DISEASE
Proportion of pathway genes=0.81; Adjusted p−value=0
Average gene set expression
0
−2
−4
Expression threshold
Below
Above
sample15_LMM_B6
KEGG_AMINO_SUGAR_AND_NUCLEOTIDE_SUGAR_METABOLISM
Proportion of pathway genes=0.85; Adjusted p−value=0
Average gene set expression
2
1
0
−1
−2
Expression threshold
Below
Above
sample15_LMM_B6
KEGG_AMINOACYL_TRNA_BIOSYNTHESIS
Proportion of pathway genes=0.51; Adjusted p−value=0
Average gene set expression
4
3
2
1
0
−1
Expression threshold
Below
Above
sample15_LMM_B6
KEGG_AMYOTROPHIC_LATERAL_SCLEROSIS_ALS
Proportion of pathway genes=0.62; Adjusted p−value=0
Average gene set expression
3
2
1
0
−1
−2
Expression threshold
Below
Above

sample15_LMM_B6
KEGG_APOPTOSIS
Proportion of pathway genes=0.76; Adjusted p−value=0

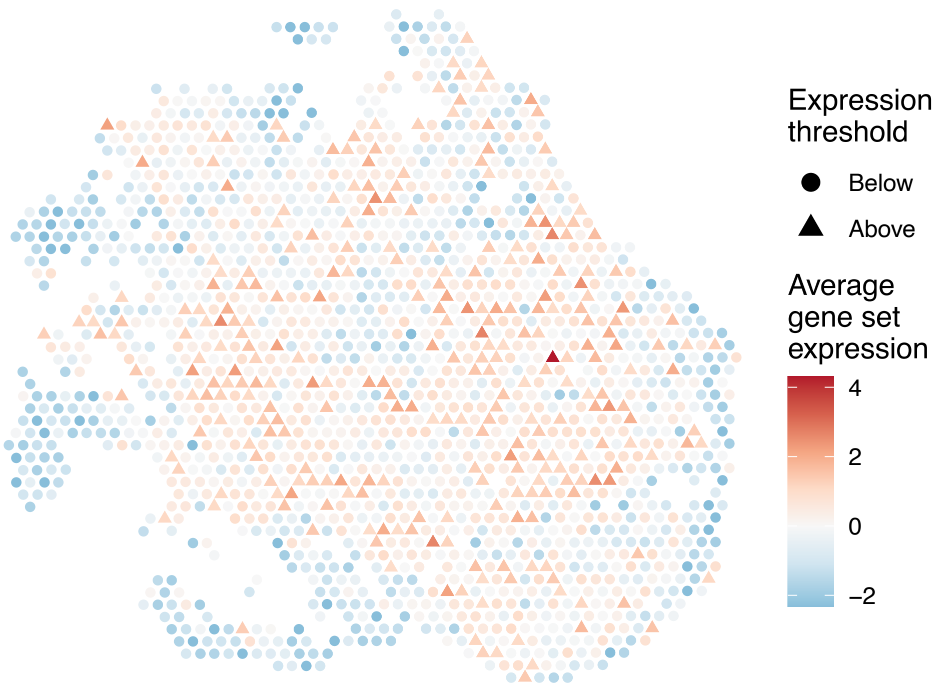


sample15_LMM_B6
KEGG_ARACHIDONIC_ACID_METABOLISM
Proportion of pathway genes=0.45; Adjusted p−value=1

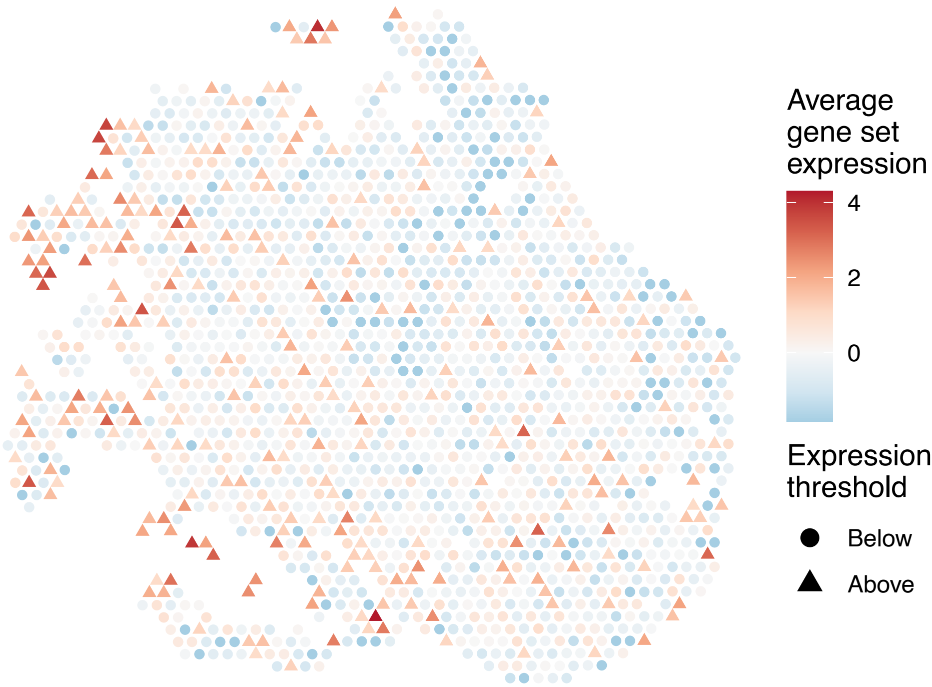


sample15_LMM_B6
KEGG_ARGININE_AND_PROLINE_METABOLISM
Proportion of pathway genes=0.73; Adjusted p−value=0

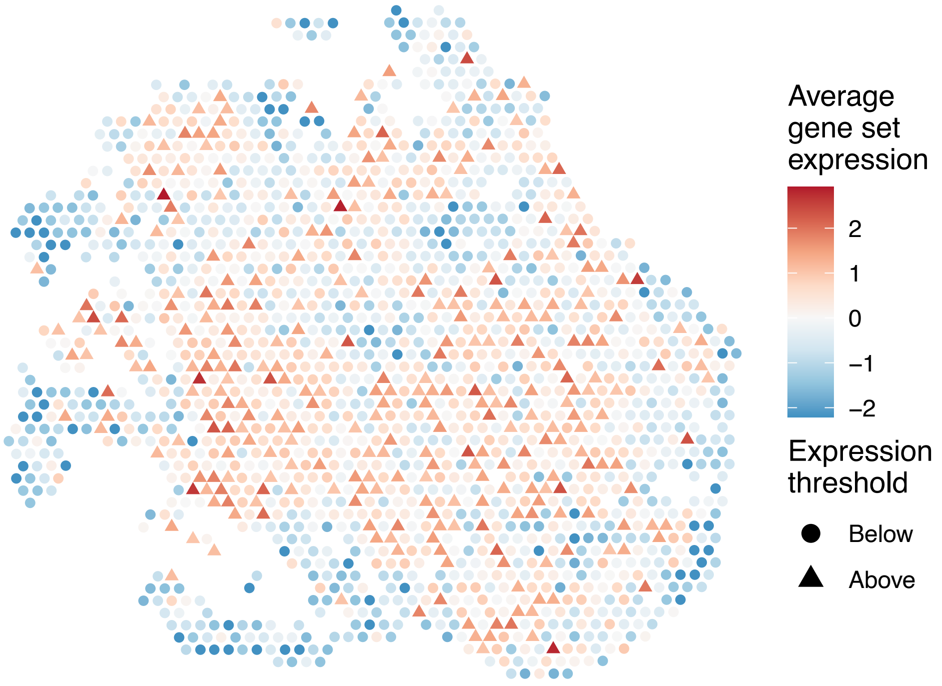


sample15_LMM_B6
KEGG_ARRHYTHMOGENIC_RIGHT_VENTRICULAR_CARDIOMYOPAT
HY_ARVC
Proportion of pathway genes=0.51; Adjusted p−value=0

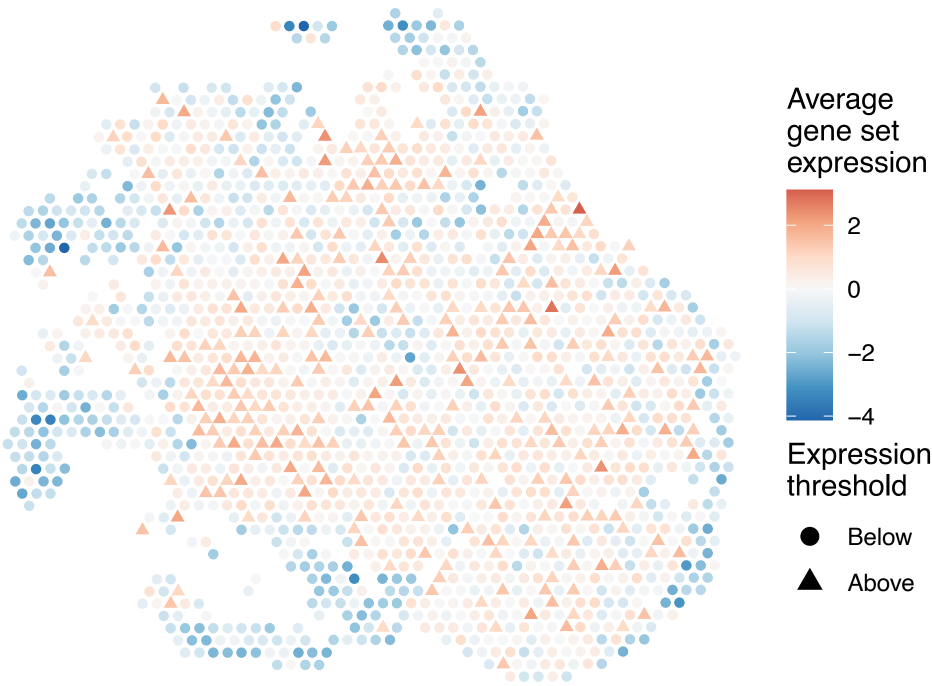

sample15_LMM_B6
KEGG_ASTHMA
Proportion of pathway genes=0.57; Adjusted p−value=0.6549

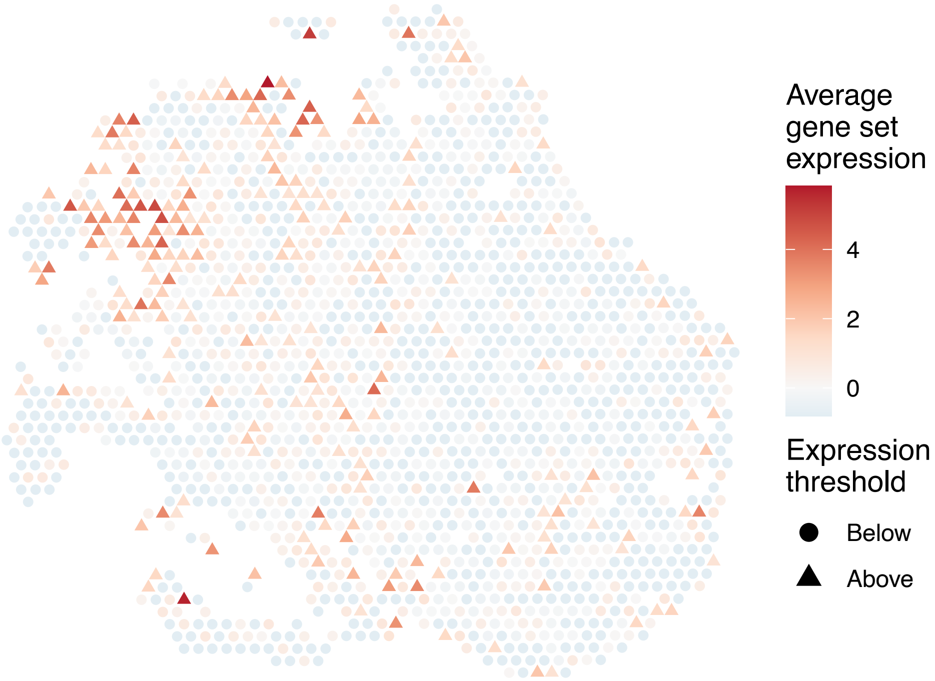


sample15_LMM_B6
KEGG_AUTOIMMUNE_THYROID_DISEASE
Proportion of pathway genes=0.6; Adjusted p−value=0

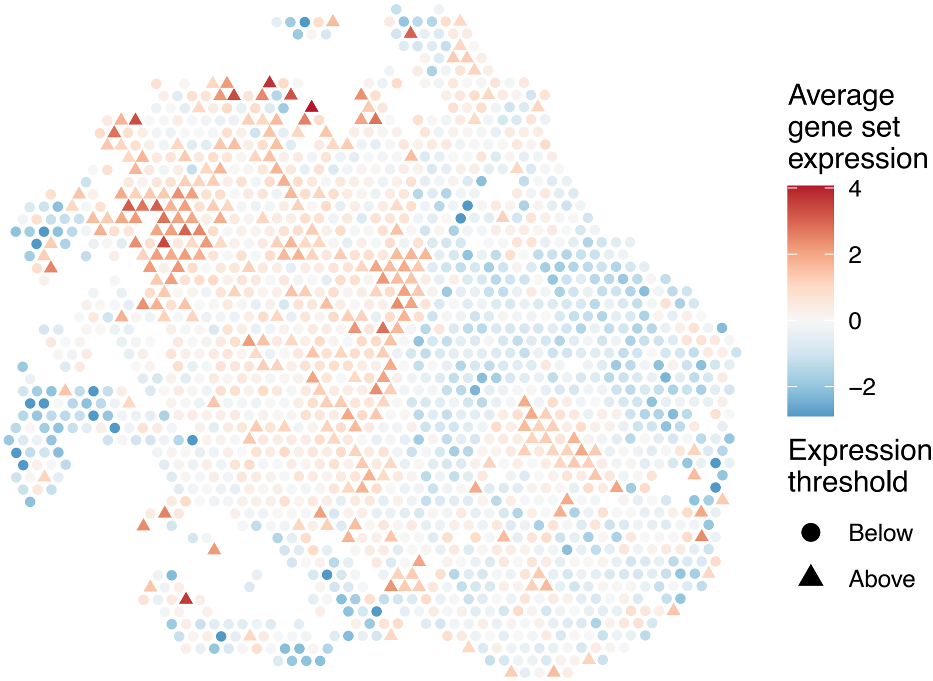


sample15_LMM_B6
KEGG_AXON_GUIDANCE
Proportion of pathway genes=0.72; Adjusted p−value=0

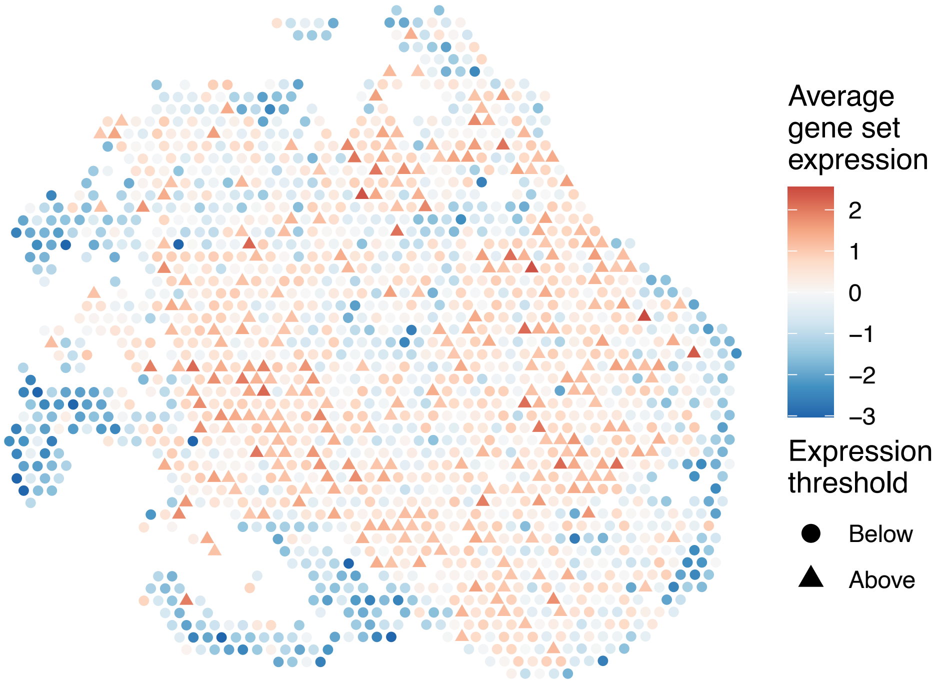


sample15_LMM_B6
KEGG_B_CELL_RECEPTOR_SIGNALING_PATHWAY
Proportion of pathway genes=0.83; Adjusted p−value=0

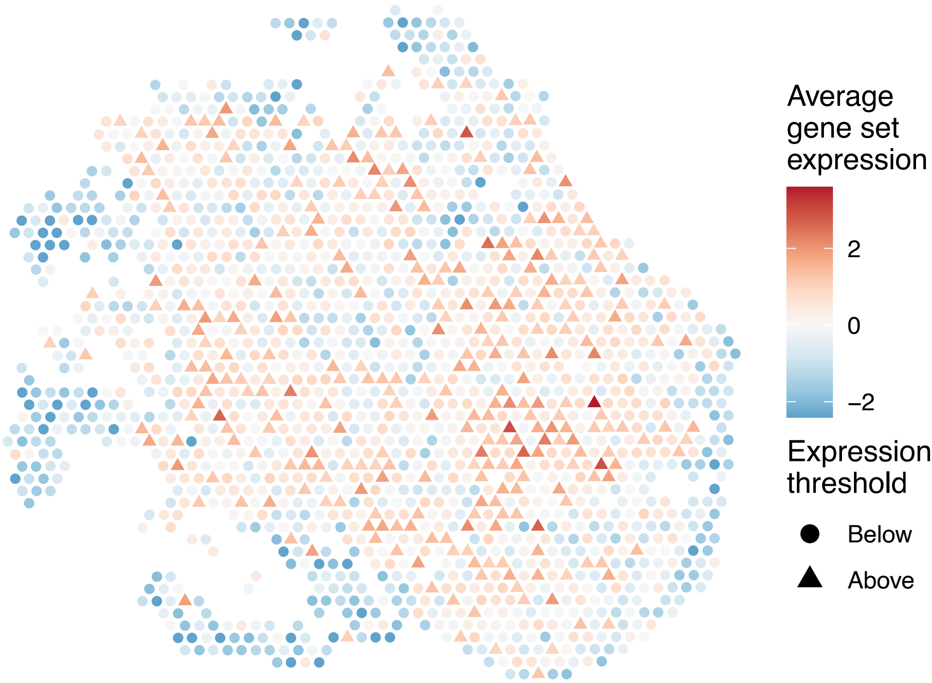

sample15_LMM_B6
KEGG_BASAL_CELL_CARCINOMA
Proportion of pathway genes=0.52; Adjusted p–value=0

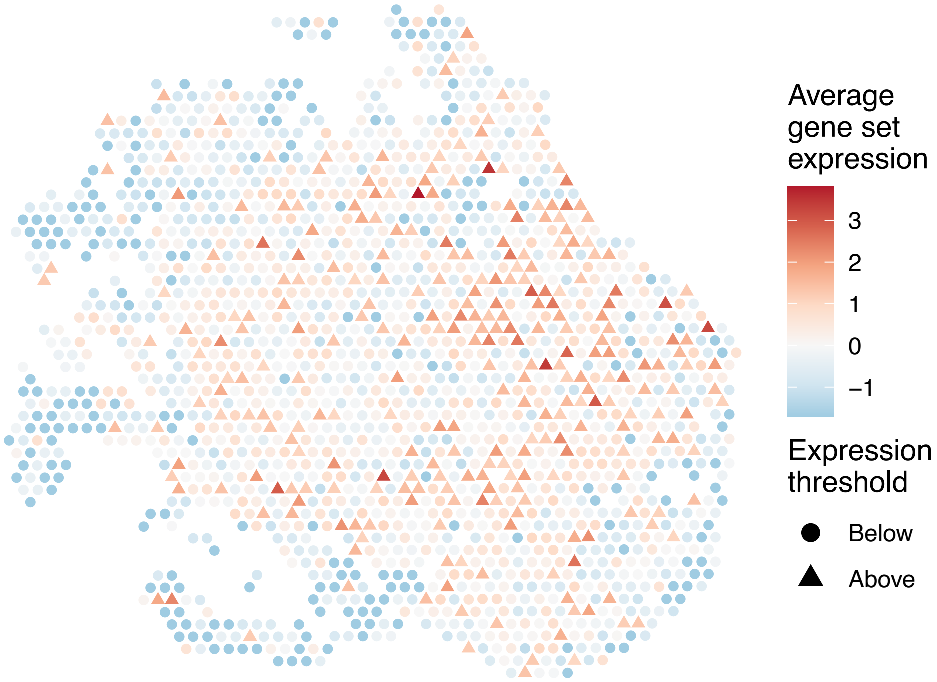


sample15_LMM_B6
KEGG_BASAL_TRANSCRIPTION_FACTORS
Proportion of pathway genes=0.87; Adjusted p–value=0

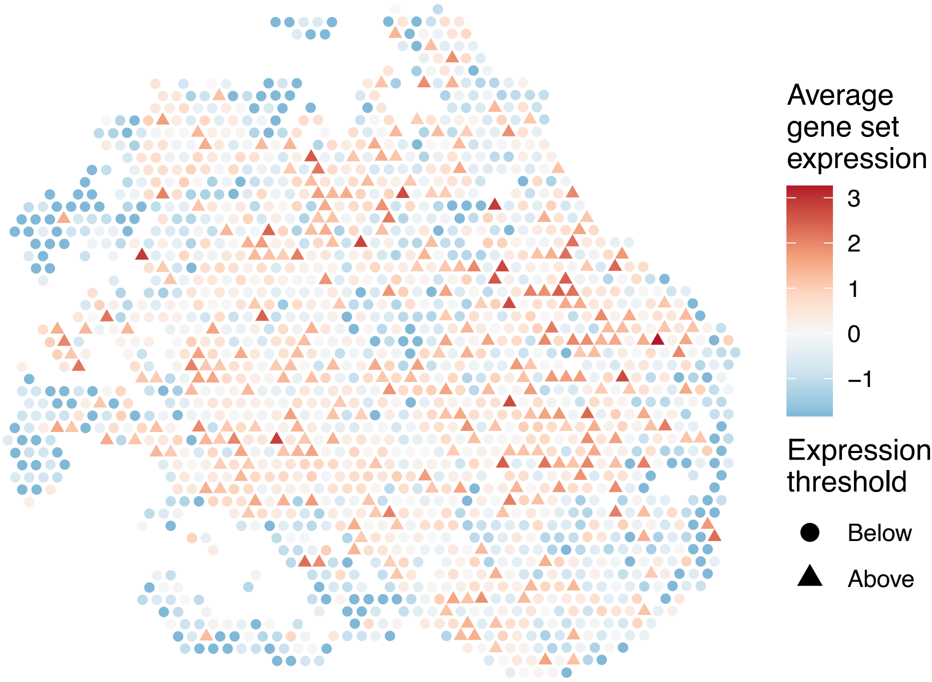


sample15_LMM_B6
KEGG_BASE_EXCISION_REPAIR
Proportion of pathway genes=0.89; Adjusted p–value=0

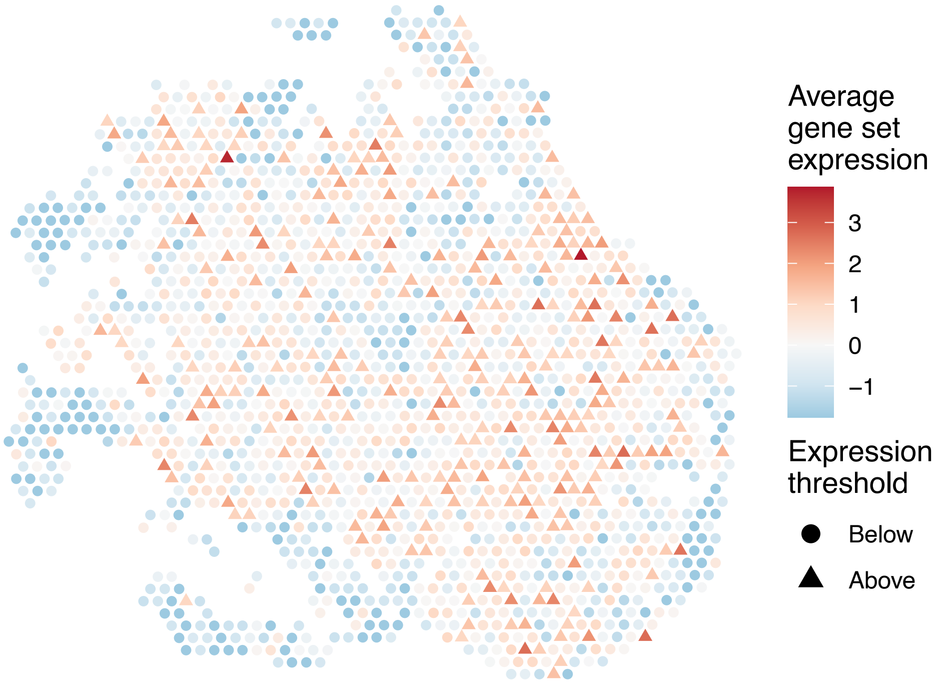


sample15_LMM_B6
KEGG_BETA_ALANINE_METABOLISM
Proportion of pathway genes=0.68; Adjusted p–value=0

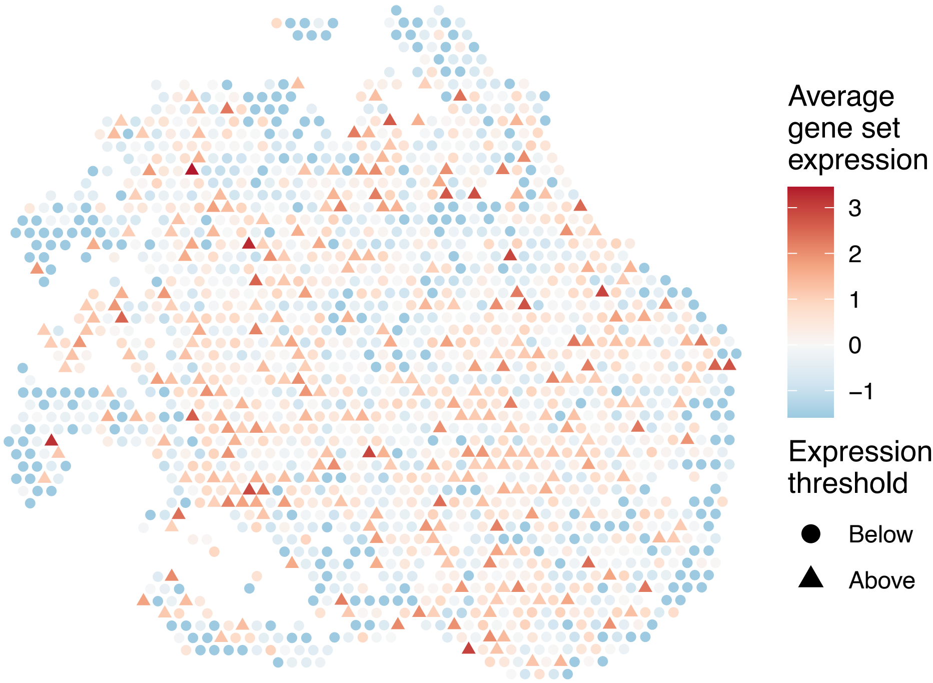

sample15_LMM_B6
KEGG_BIOSYNTHESIS_OF_UNSATURATED_FATTY_ACIDS
Proportion of pathway genes=0.87; Adjusted p−value=0

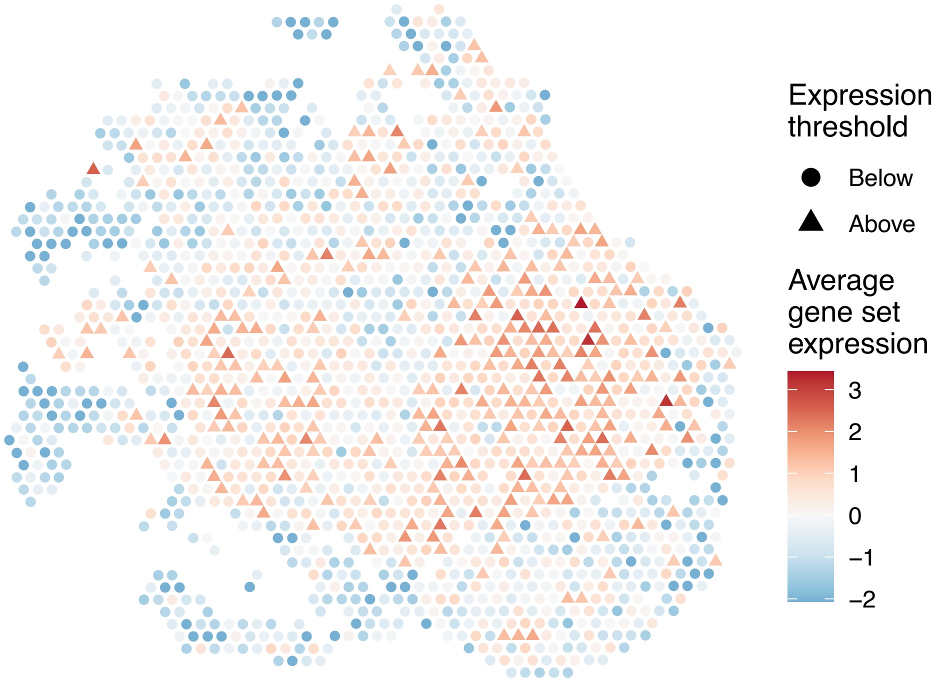


sample15_LMM_B6
KEGG_BLADDER_CANCER
Proportion of pathway genes=0.8; Adjusted p−value=0

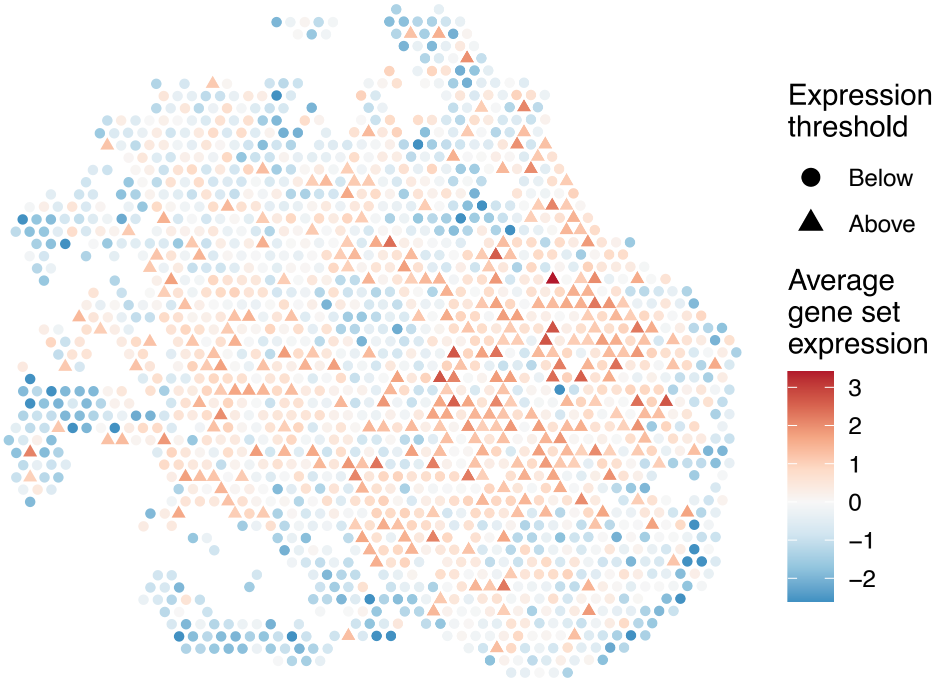


sample15_LMM_B6
KEGG_BUTANOATE_METABOLISM
Proportion of pathway genes=0.66; Adjusted p−value=0

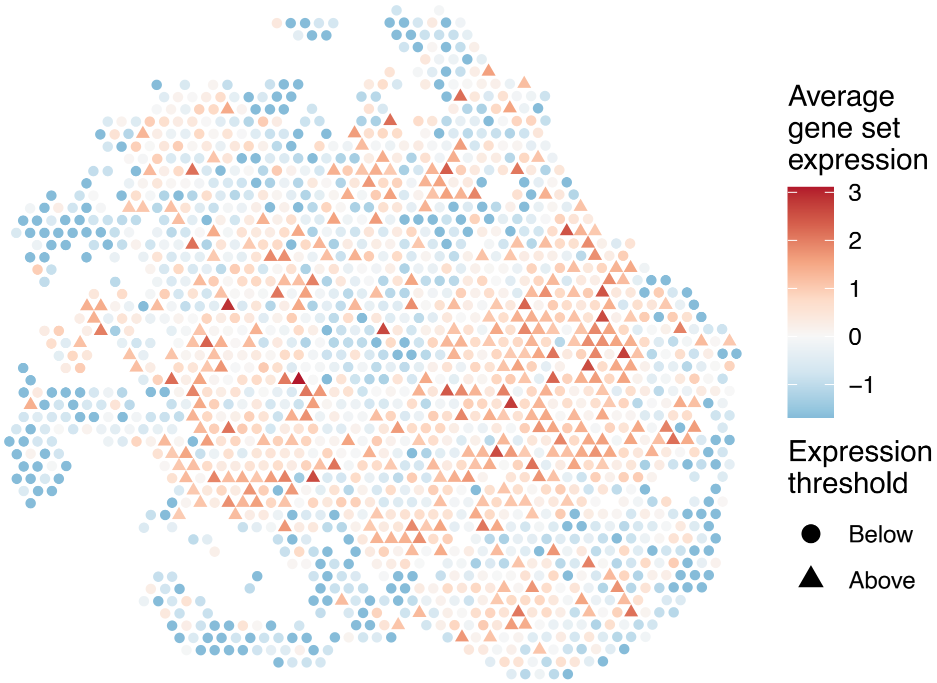


sample15_LMM_B6
KEGG_CALCIUM_SIGNALING_PATHWAY
Proportion of pathway genes=0.44; Adjusted p−value=0

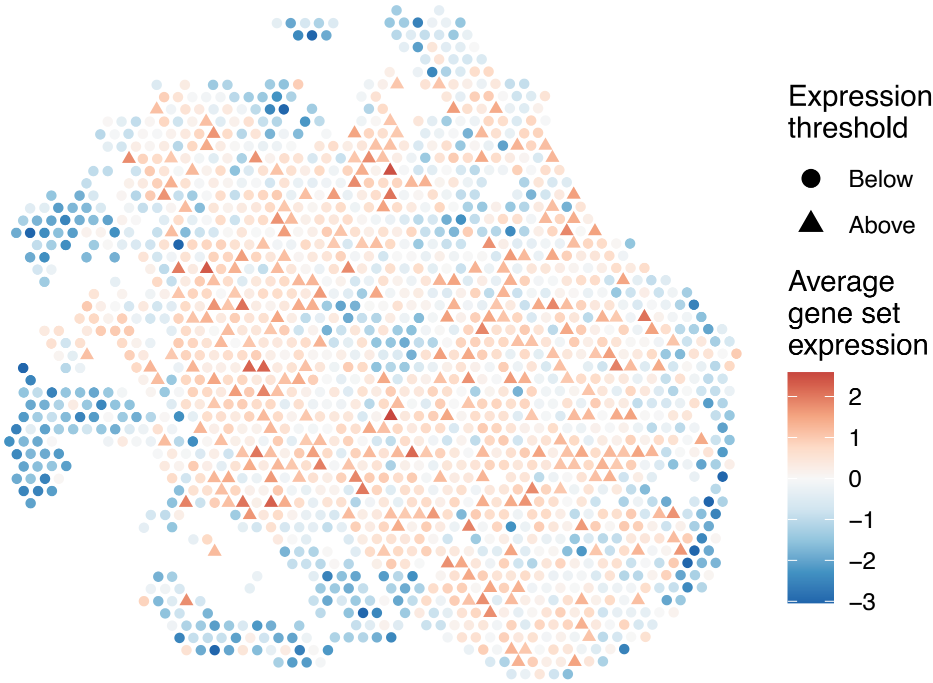

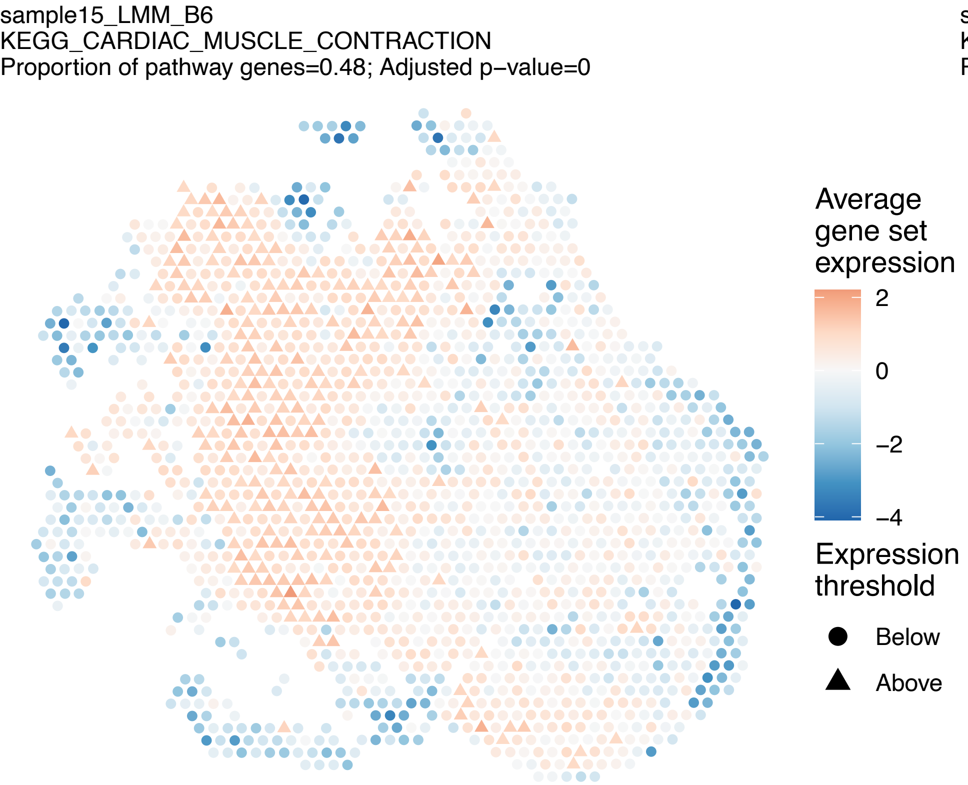
sample15_LMM_B6
KEGG_CARDIAC_MUSCLE_CONTRACTION
Proportion of pathway genes=0.48; Adjusted p−value=0
Average gene set expression
2
0
−2
−4
Expression threshold
Below
Above

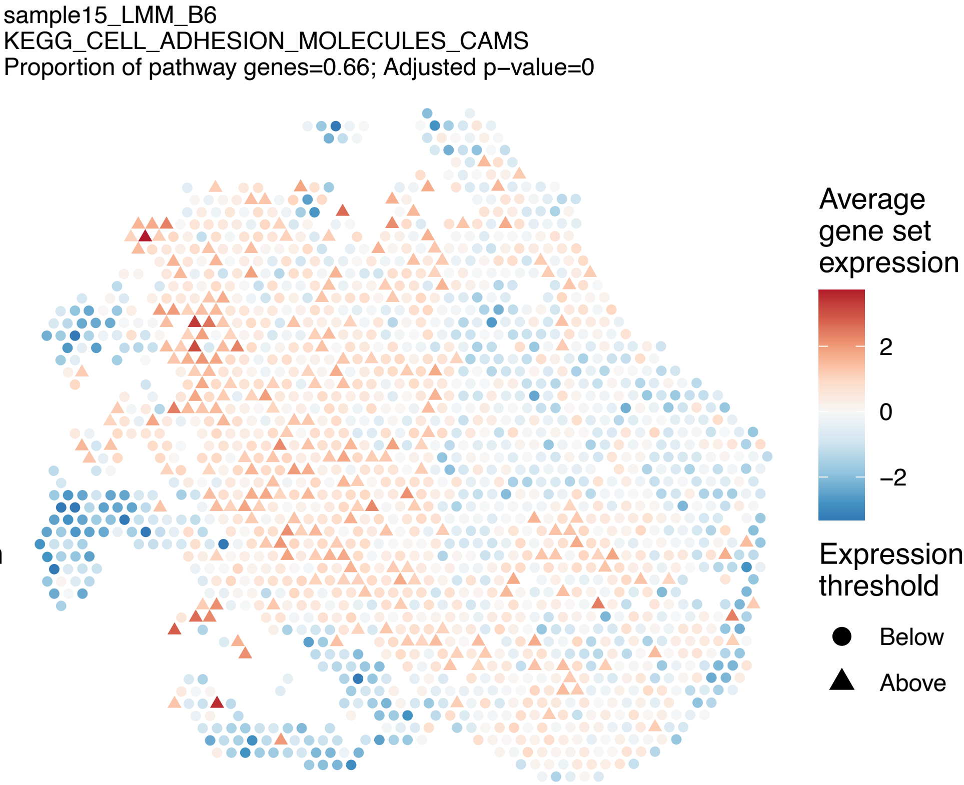
sample15_LMM_B6
KEGG_CELL_ADHESION_MOLECULES_CAMS
Proportion of pathway genes=0.66; Adjusted p−value=0
Average gene set expression
2
0
−2
Expression threshold
Below
Above

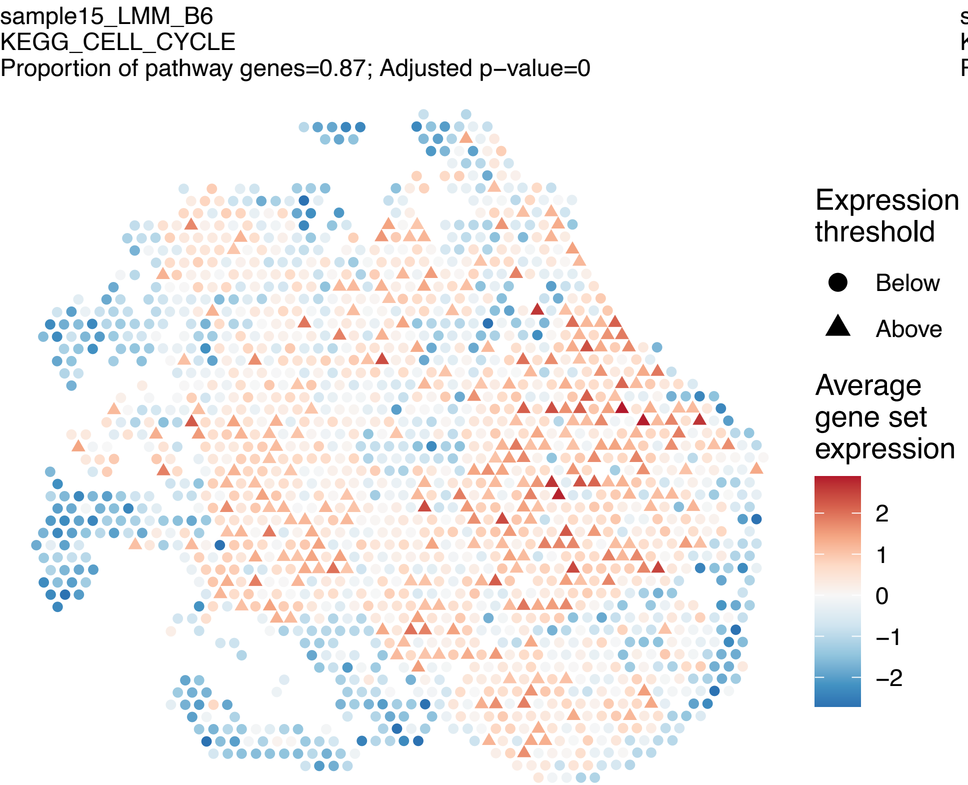
sample15_LMM_B6
KEGG_CELL_CYCLE
Proportion of pathway genes=0.87; Adjusted p−value=0
Expression threshold
Below
Above
Average gene set expression
2
1
0
−1
−2

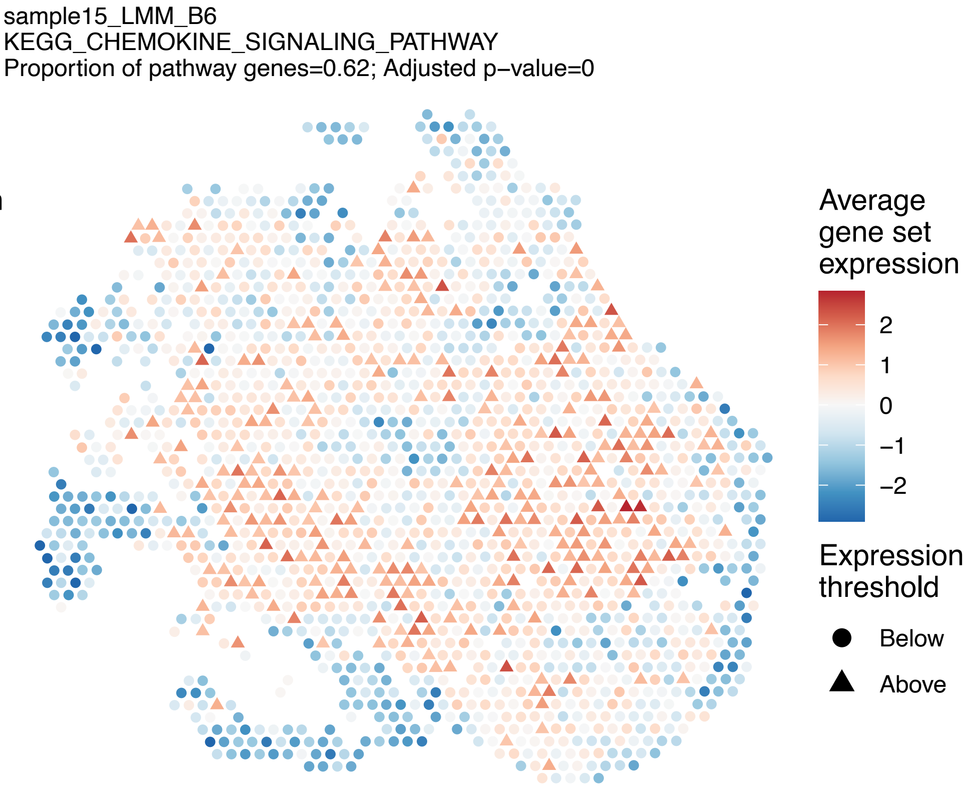
sample15_LMM_B6
KEGG_CHEMOKINE_SIGNALING_PATHWAY
Proportion of pathway genes=0.62; Adjusted p−value=0
Average gene set expression
2
1
0
−1
−2
Expression threshold
Below
Above

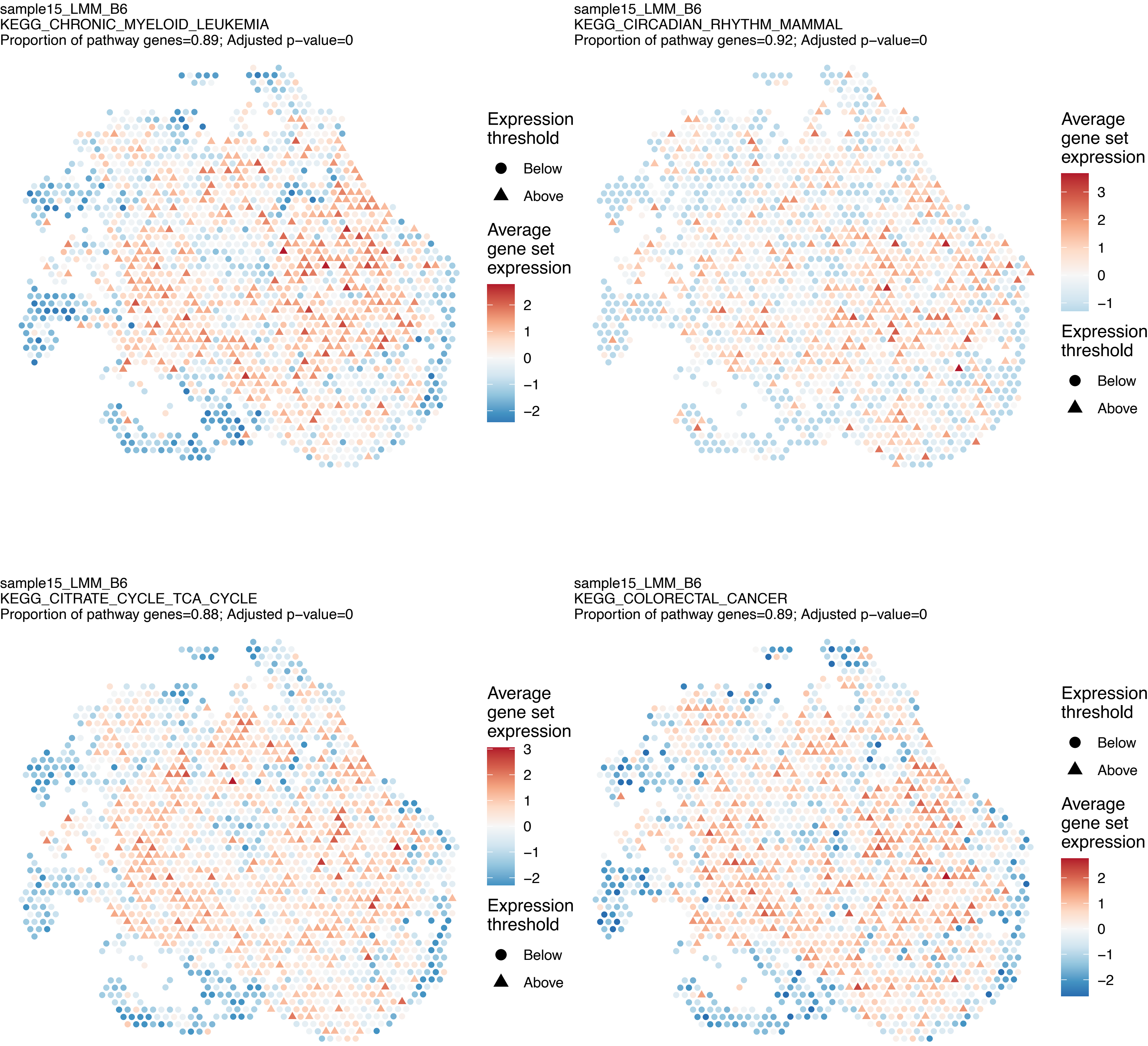
sample15_LMM_B6
KEGG_CHRONIC_MYELOID_LEUKEMIA
Proportion of pathway genes=0.89; Adjusted p−value=0
Expression threshold
Below
Above
Average gene set expression
2
1
0
−1
−2
sample15_LMM_B6
KEGG_CIRCADIAN_RHYTHM_MAMMAL
Proportion of pathway genes=0.92; Adjusted p−value=0
Average gene set expression
3
2
1
0
−1
Expression threshold
Below
Above
sample15_LMM_B6
KEGG_CITRATE_CYCLE_TCA_CYCLE
Proportion of pathway genes=0.88; Adjusted p−value=0
Average gene set expression
3
2
1
0
−1
−2
Expression threshold
Below
Above
sample15_LMM_B6
KEGG_COLORECTAL_CANCER
Proportion of pathway genes=0.89; Adjusted p−value=0
Expression threshold
Below
Above
Average gene set expression
2
1
0
−1
−2

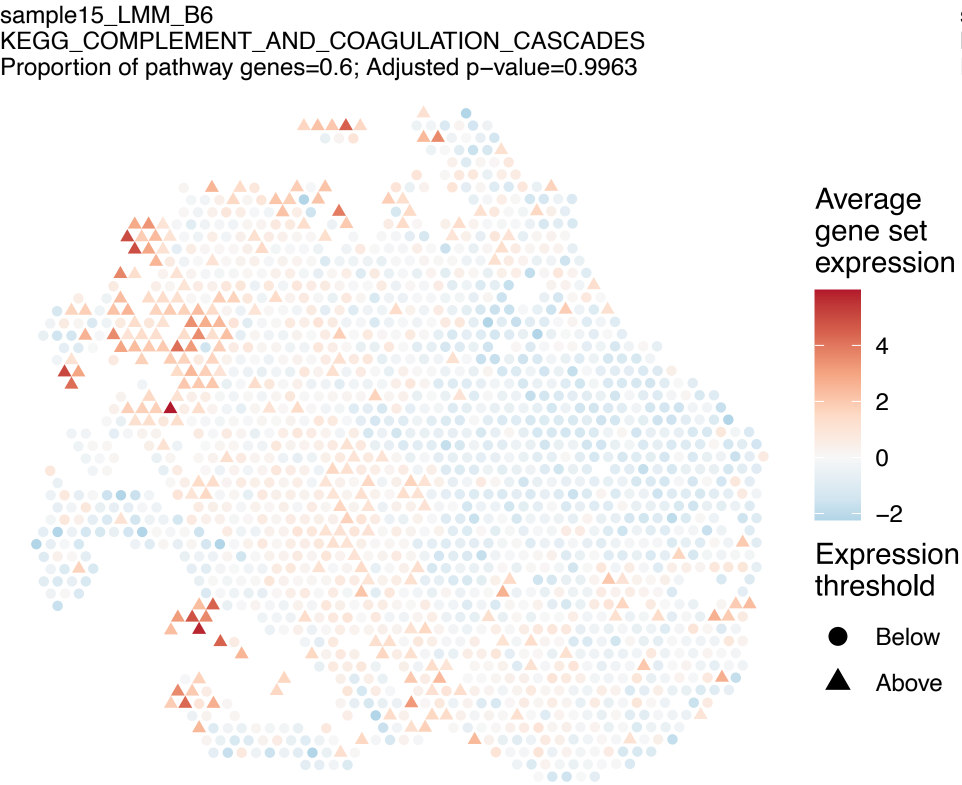
sample15_LMM_B6
KEGG_COMPLEMENT_AND_COAGULATION_CASCADES
Proportion of pathway genes=0.6; Adjusted p–value=0.9963
Average gene set expression
4
2
0
−2
Expression threshold
Below
Above

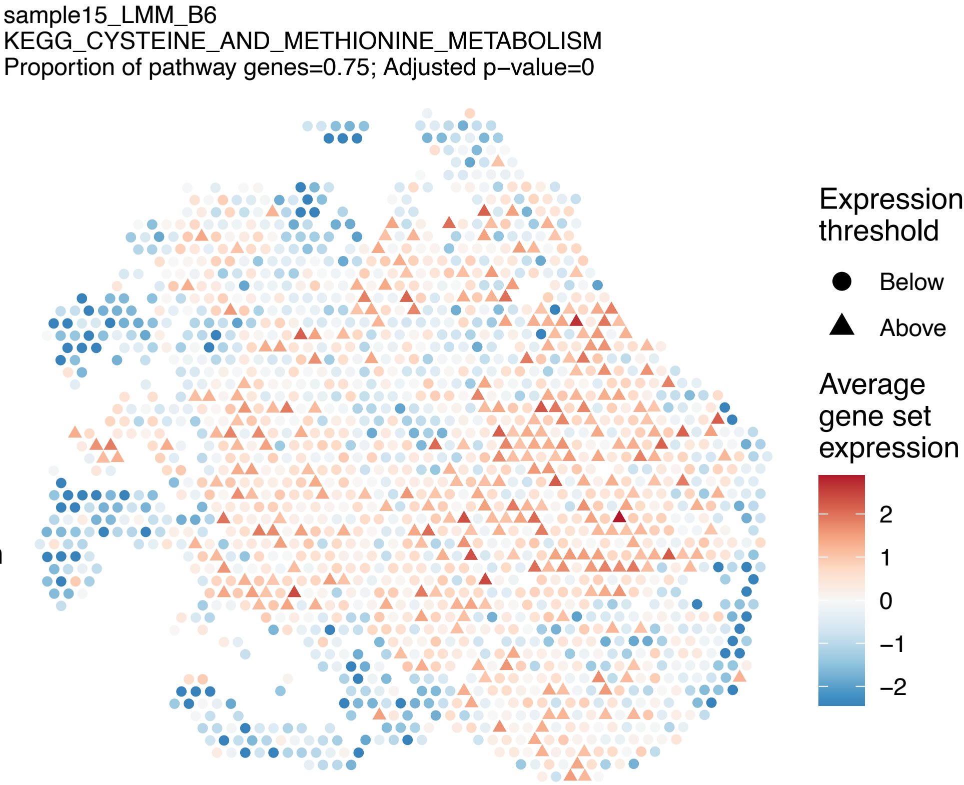
sample15_LMM_B6
KEGG_CYSTEINE_AND_METHIONINE_METABOLISM
Proportion of pathway genes=0.75; Adjusted p–value=0
Expression threshold
Below
Above
Average gene set expression
2
1
0
−1
−2

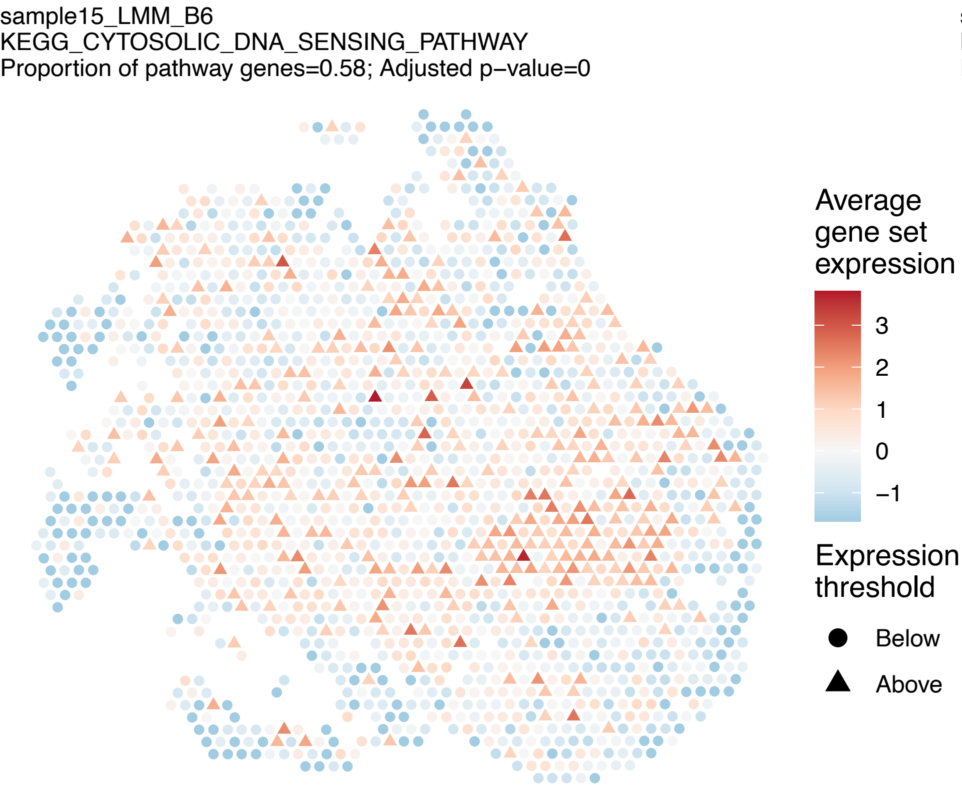
sample15_LMM_B6
KEGG_CYTOSOLIC_DNA_SENSING_PATHWAY
Proportion of pathway genes=0.58; Adjusted p–value=0
Average gene set expression
3
2
1
0
−1
Expression threshold
Below
Above

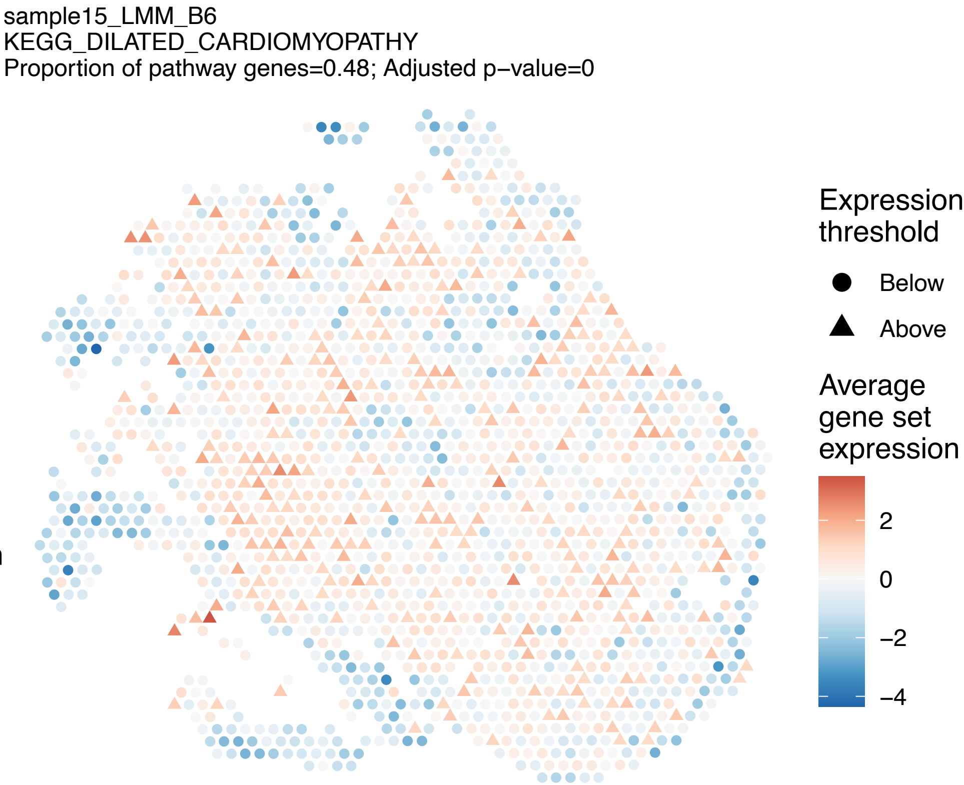
sample15_LMM_B6
KEGG_DILATED_CARDIOMYOPATHY
Proportion of pathway genes=0.48; Adjusted p–value=0
Expression threshold
Below
Above
Average gene set expression
2
0
−2
−4

sample15_LMM_B6
KEGG_DNA_REPLICATION
Proportion of pathway genes=0.92; Adjusted p–value=0

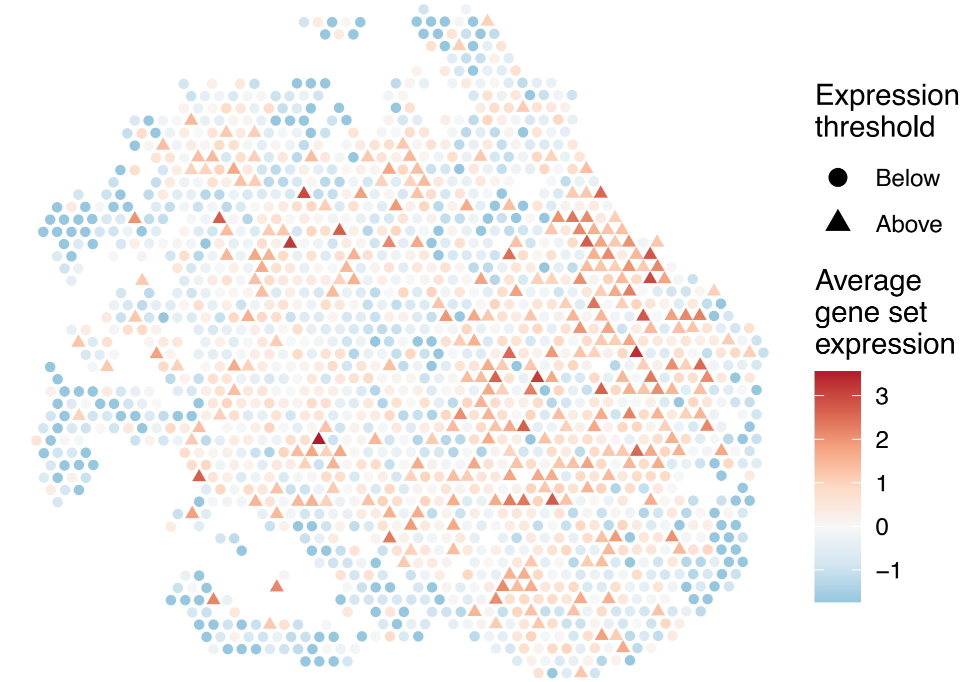


sample15_LMM_B6
KEGG_DORSO_VENTRAL_AXIS_FORMATION
Proportion of pathway genes=0.78; Adjusted p–value=0

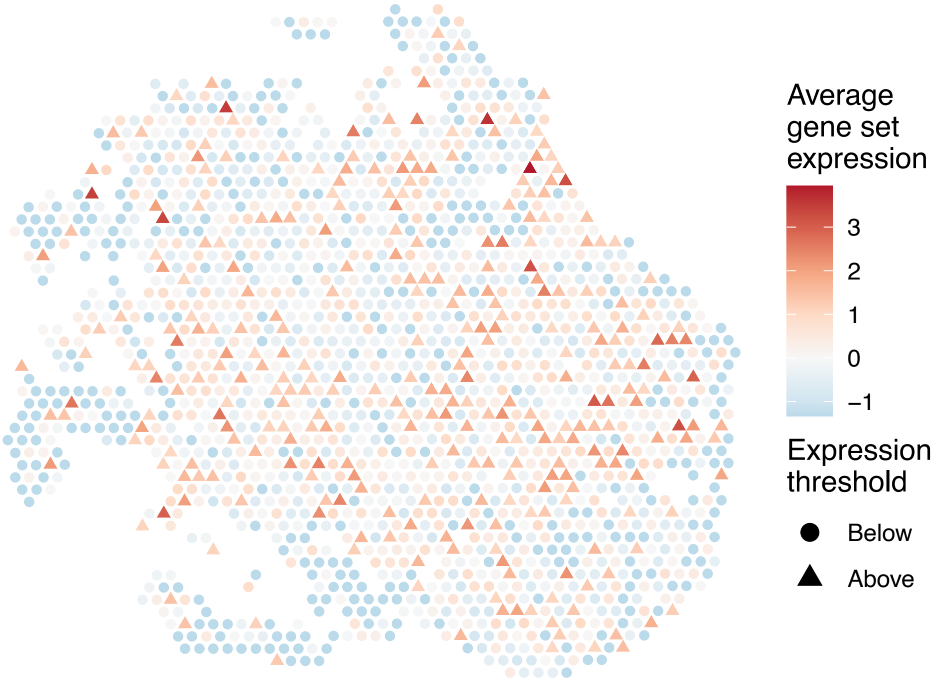


sample15_LMM_B6
KEGG_DRUG_METABOLISM_CYTOCHROME_P450
Proportion of pathway genes=0.37; Adjusted p–value=0

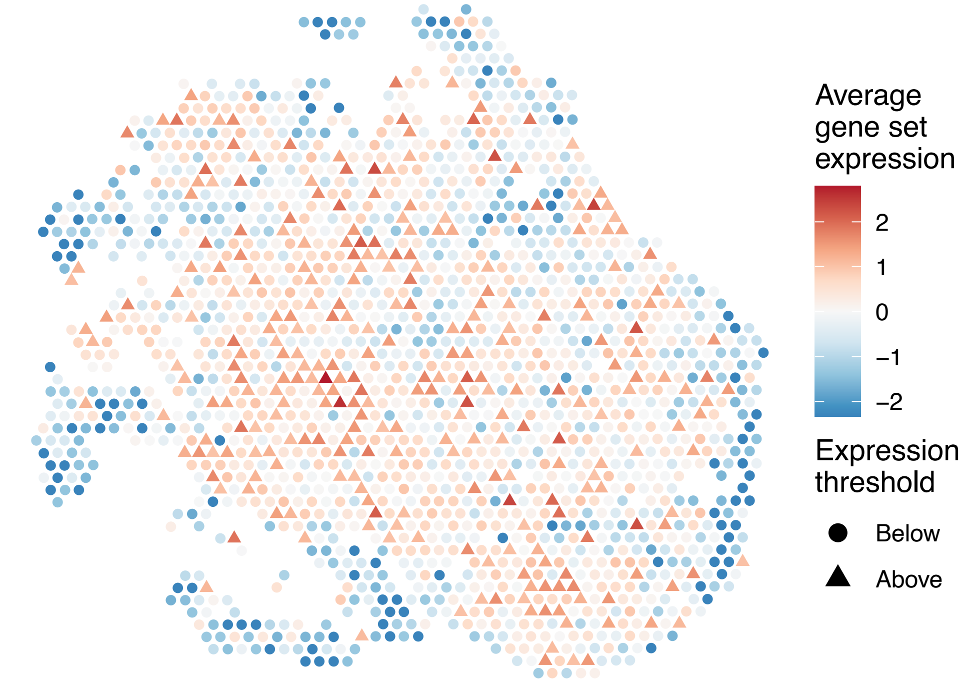


sample15_LMM_B6
KEGG_DRUG_METABOLISM_OTHER_ENZYMES
Proportion of pathway genes=0.34; Adjusted p–value=0

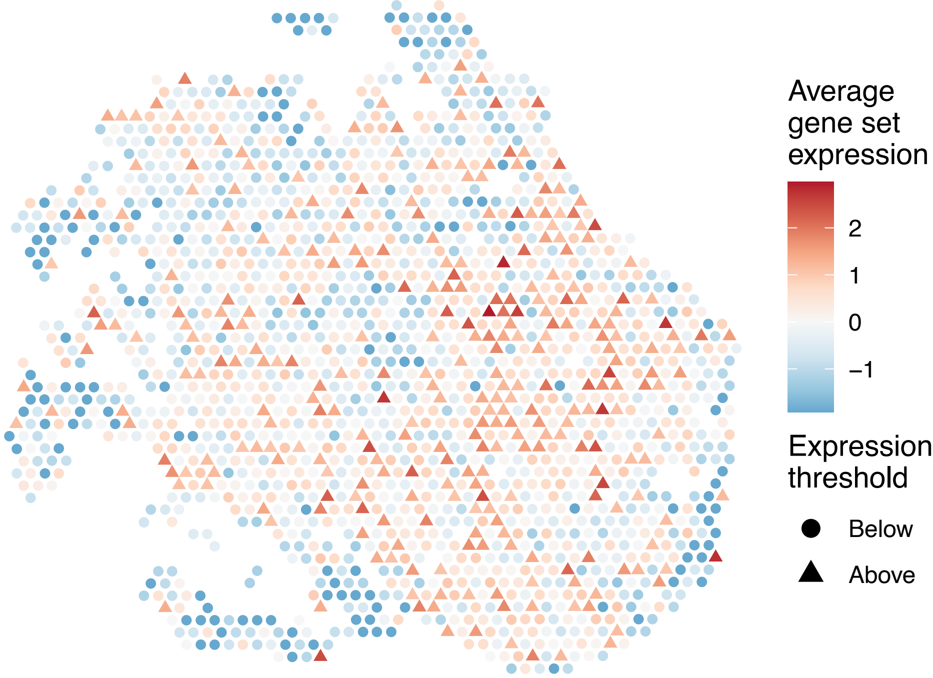

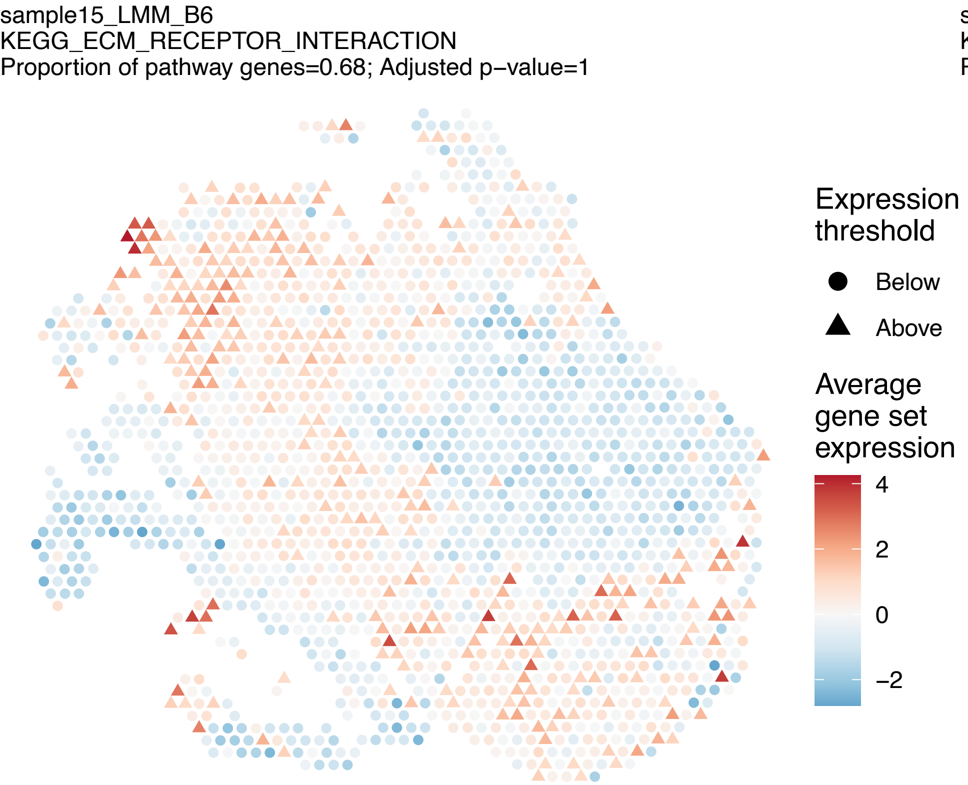
sample15_LMM_B6
KEGG_ECM_RECEPTOR_INTERACTION
Proportion of pathway genes=0.68; Adjusted p−value=1
Expression threshold
Below
Above
Average gene set expression
4
2
0
−2

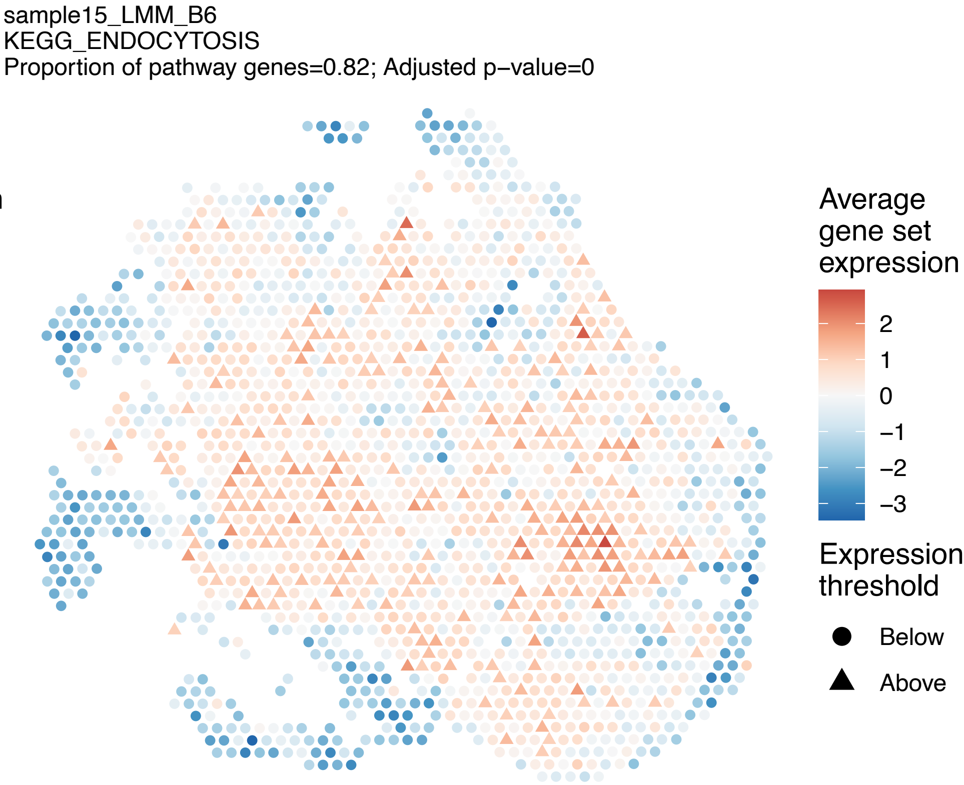
sample15_LMM_B6
KEGG_ENDOCYTOSIS
Proportion of pathway genes=0.82; Adjusted p−value=0
Average gene set expression
2
1
0
−1
−2
−3
Expression threshold
Below
Above

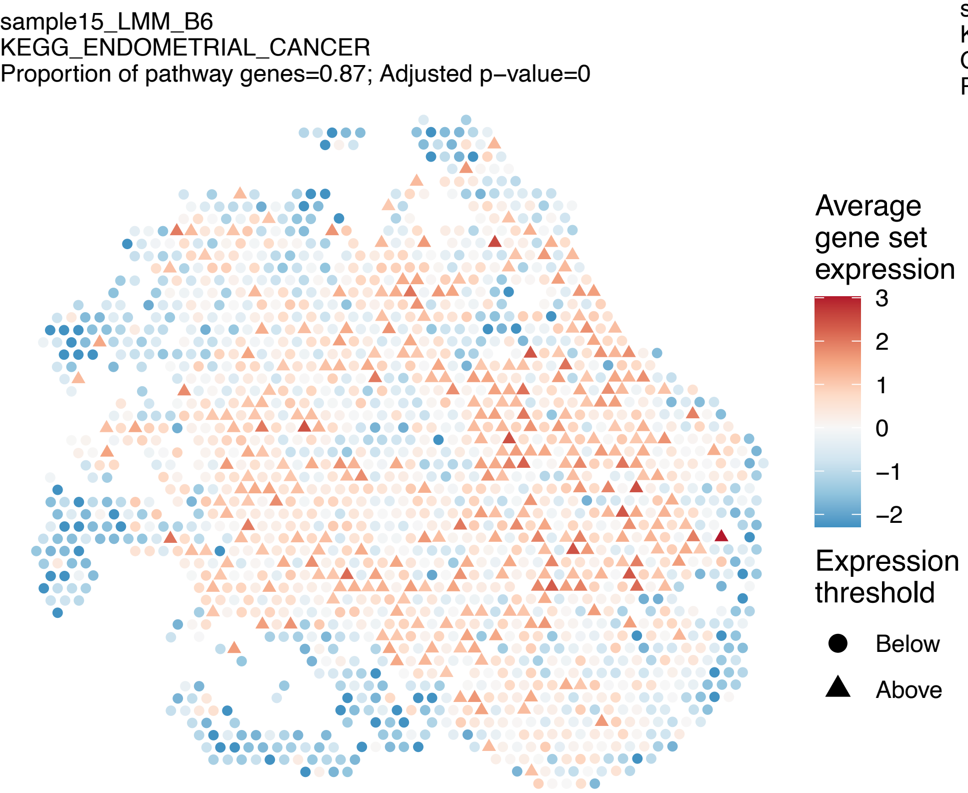
sample15_LMM_B6
KEGG_ENDOMETRIAL_CANCER
Proportion of pathway genes=0.87; Adjusted p−value=0
Average gene set expression
3
2
1
0
−1
−2
Expression threshold
Below
Above

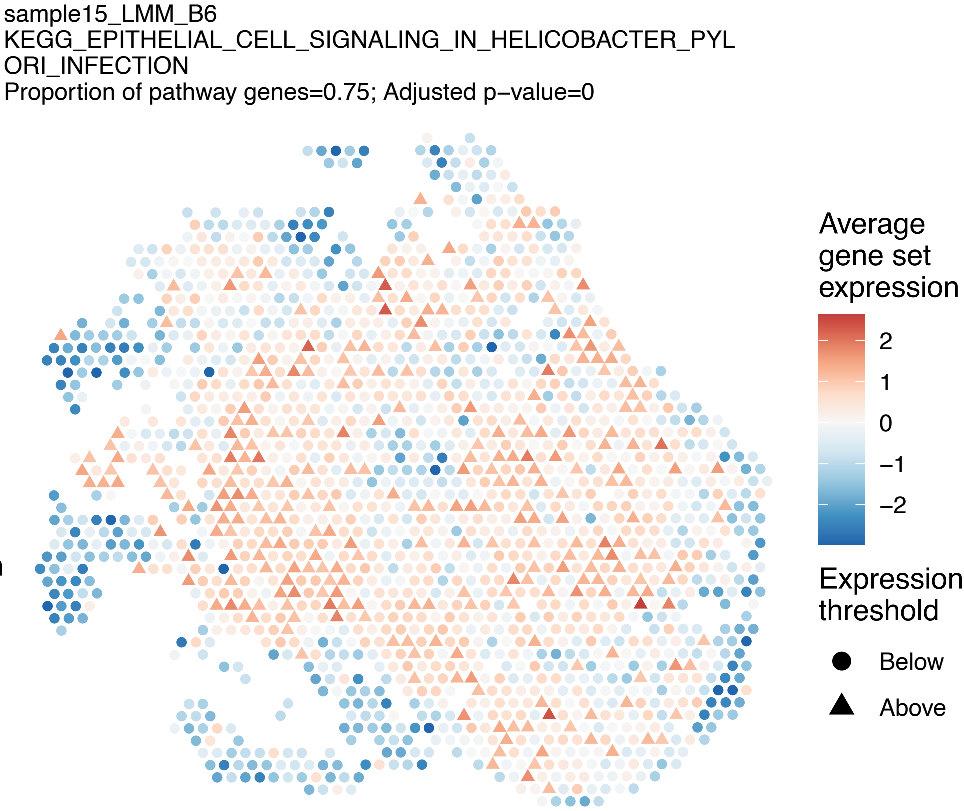
sample15_LMM_B6
KEGG_EPITHELIAL_CELL_SIGNALING_IN_HELICOBACTER_PYL
ORI_INFECTION
Proportion of pathway genes=0.75; Adjusted p−value=0
Average gene set expression
2
1
0
−1
−2
Expression threshold
Below
Above

sample15_LMM_B6
KEGG_ERBB_SIGNALING_PATHWAY
Proportion of pathway genes=0.79; Adjusted p−value=0

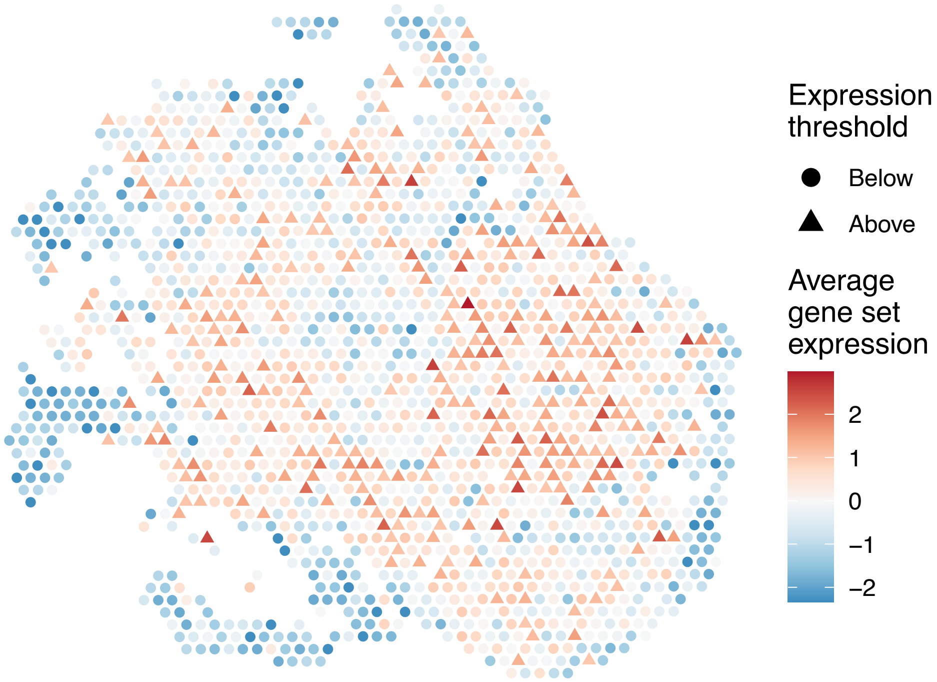


sample15_LMM_B6
KEGG_ETHER_LIPID_METABOLISM
Proportion of pathway genes=0.63; Adjusted p−value=0

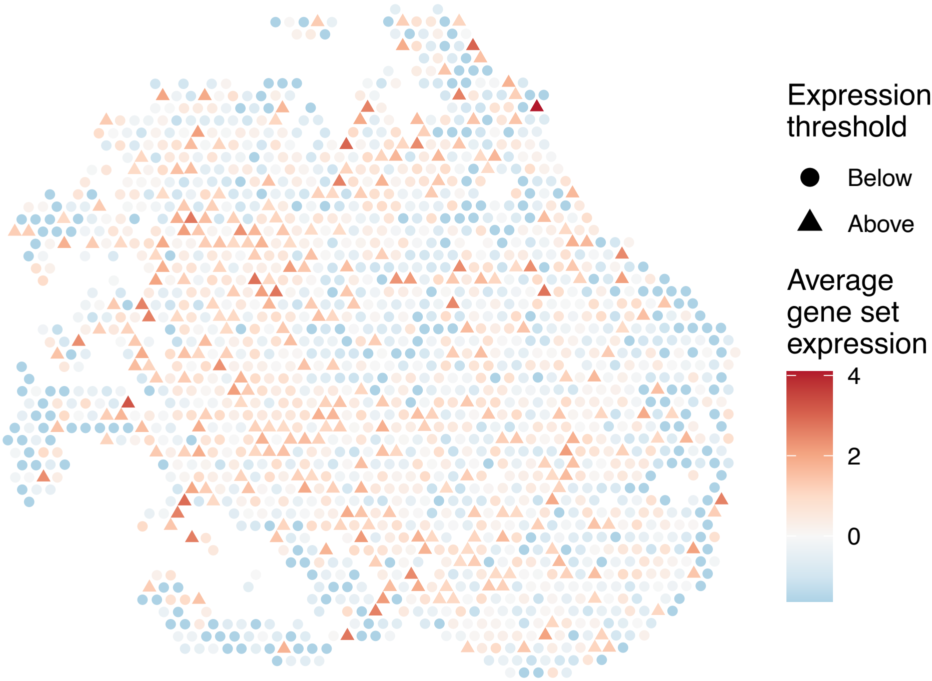


sample15_LMM_B6
KEGG_FATTY_ACID_METABOLISM
Proportion of pathway genes=0.79; Adjusted p−value=0

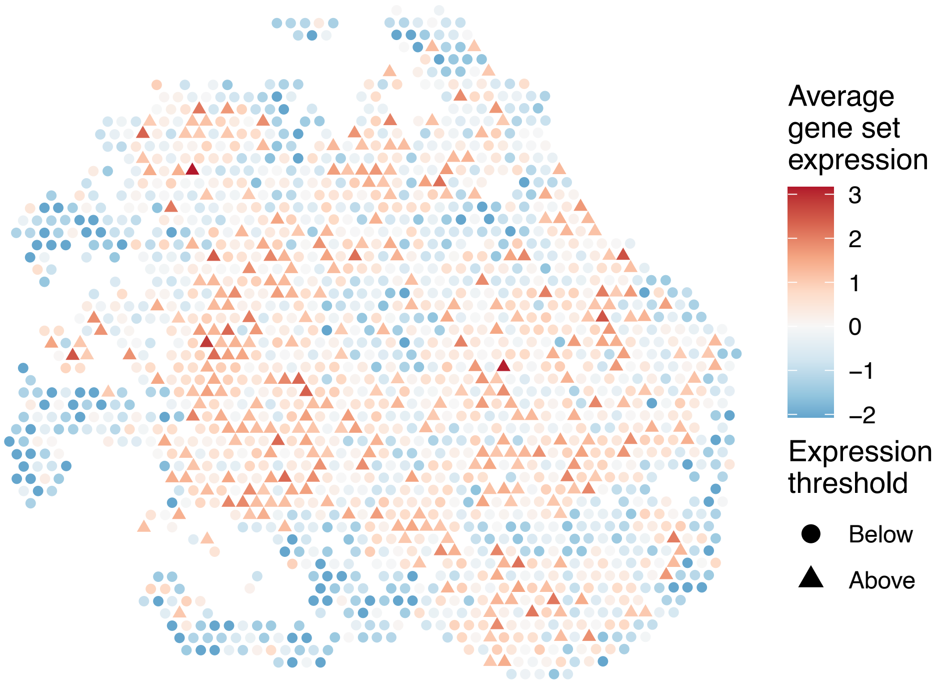


sample15_LMM_B6
KEGG_FC_EPSILON_RI_SIGNALING_PATHWAY
Proportion of pathway genes=0.62; Adjusted p−value=0

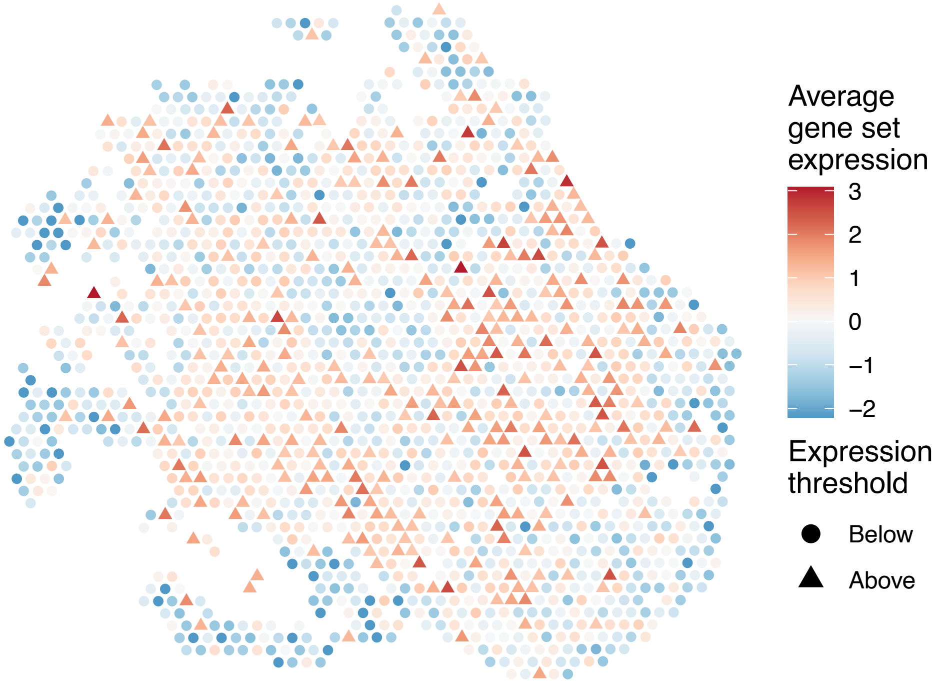

sample15_LMM_B6
KEGG_FC_GAMMA_R_MEDIATED_PHAGOCYTOSIS
Proportion of pathway genes=0.82; Adjusted p−value=0

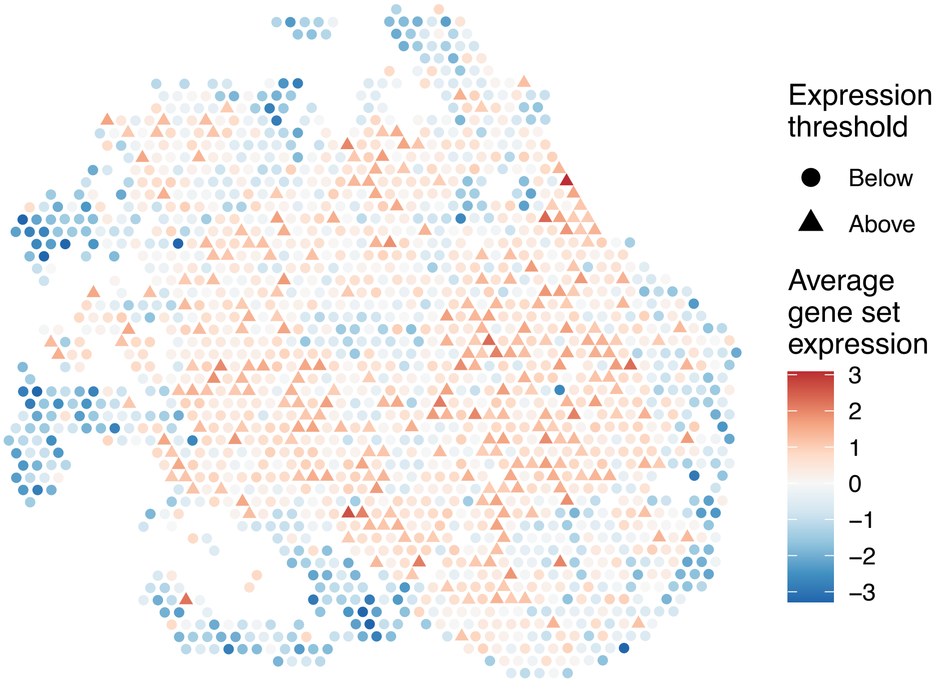


sample15_LMM_B6
KEGG_FOCAL_ADHESION
Proportion of pathway genes=0.81; Adjusted p−value=0.0117

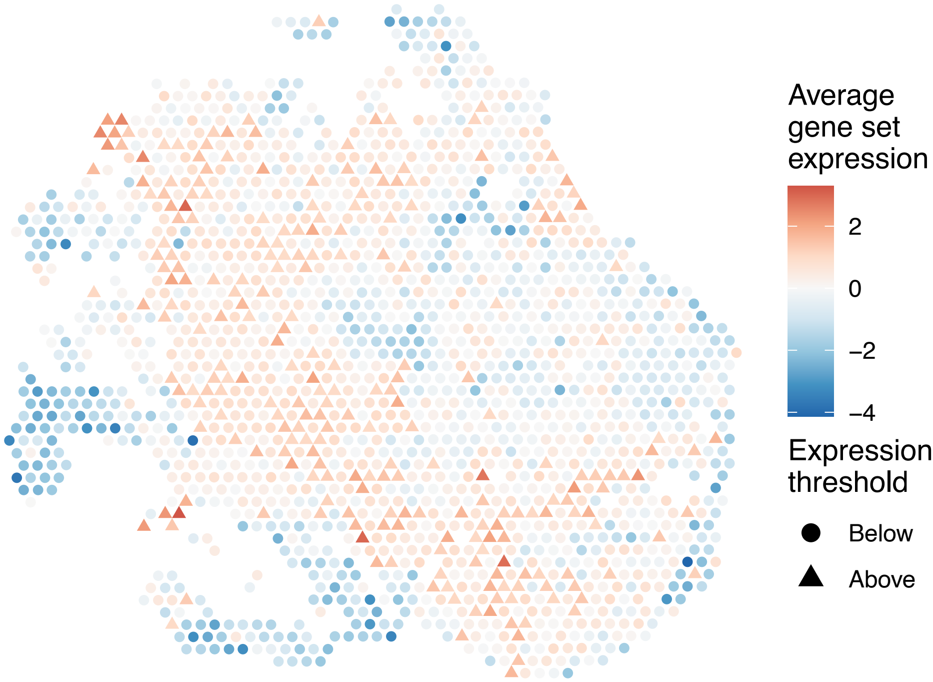


sample15_LMM_B6
KEGG_FRUCTOSE_AND_MANNOSE_METABOLISM
Proportion of pathway genes=0.71; Adjusted p−value=0

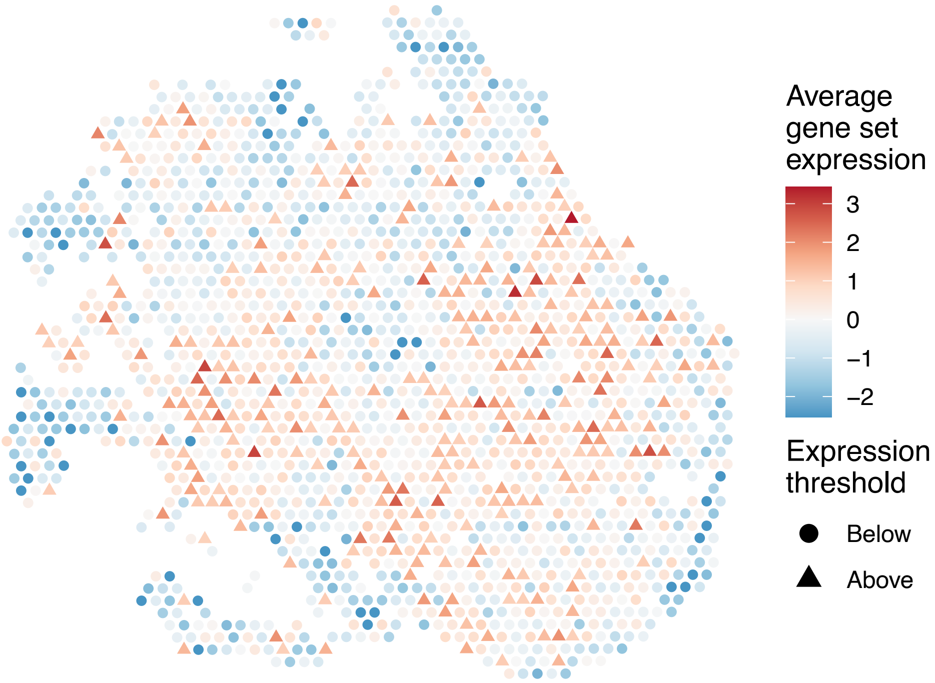


sample15_LMM_B6
KEGG_GALACTOSE_METABOLISM
Proportion of pathway genes=0.66; Adjusted p−value=0

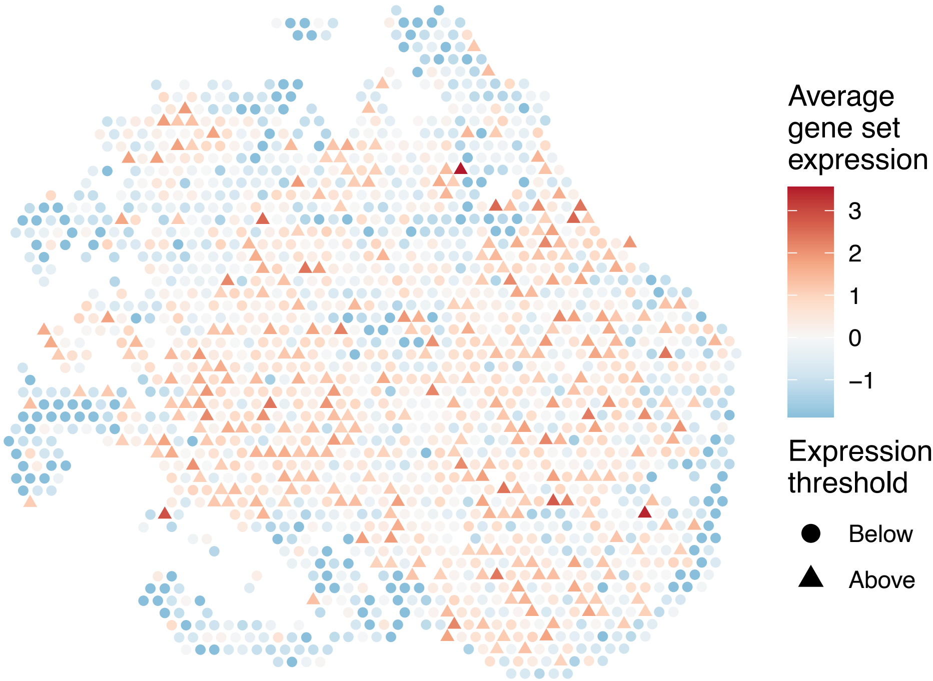

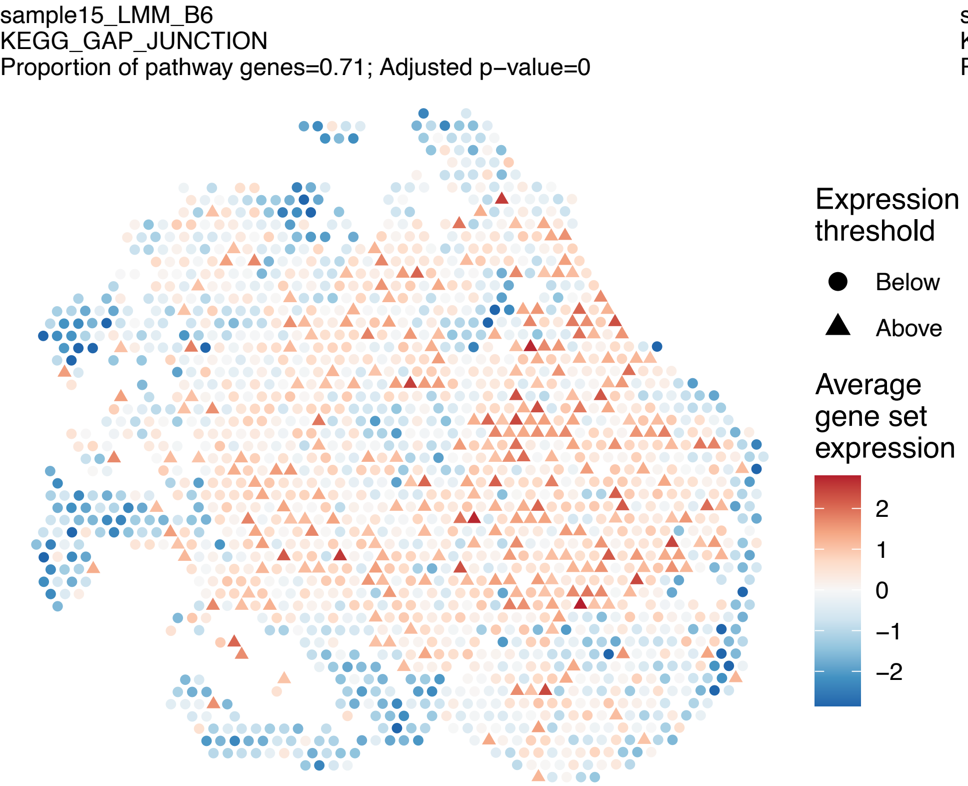
sample15_LMM_B6
KEGG_GAP_JUNCTION
Proportion of pathway genes=0.71; Adjusted p−value=0
Expression threshold
Below
Above
Average gene set expression
2
1
0
−1
−2

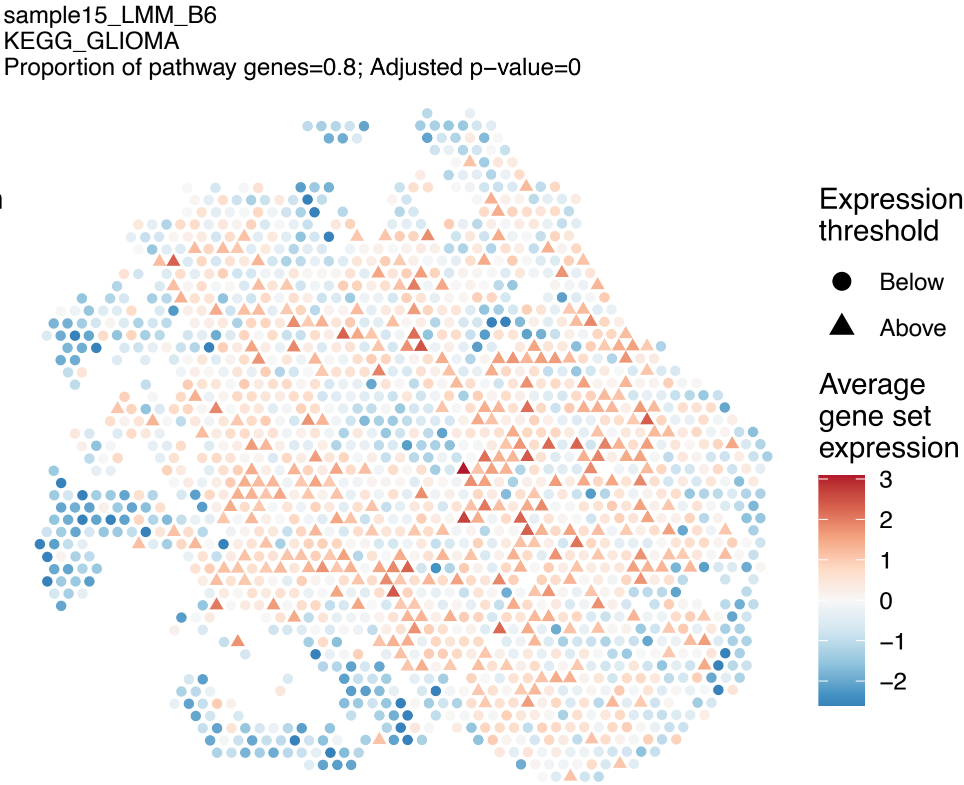
sample15_LMM_B6
KEGG_GLIOMA
Proportion of pathway genes=0.8; Adjusted p−value=0
Expression threshold
Below
Above
Average gene set expression
3
2
1
0
−1
−2

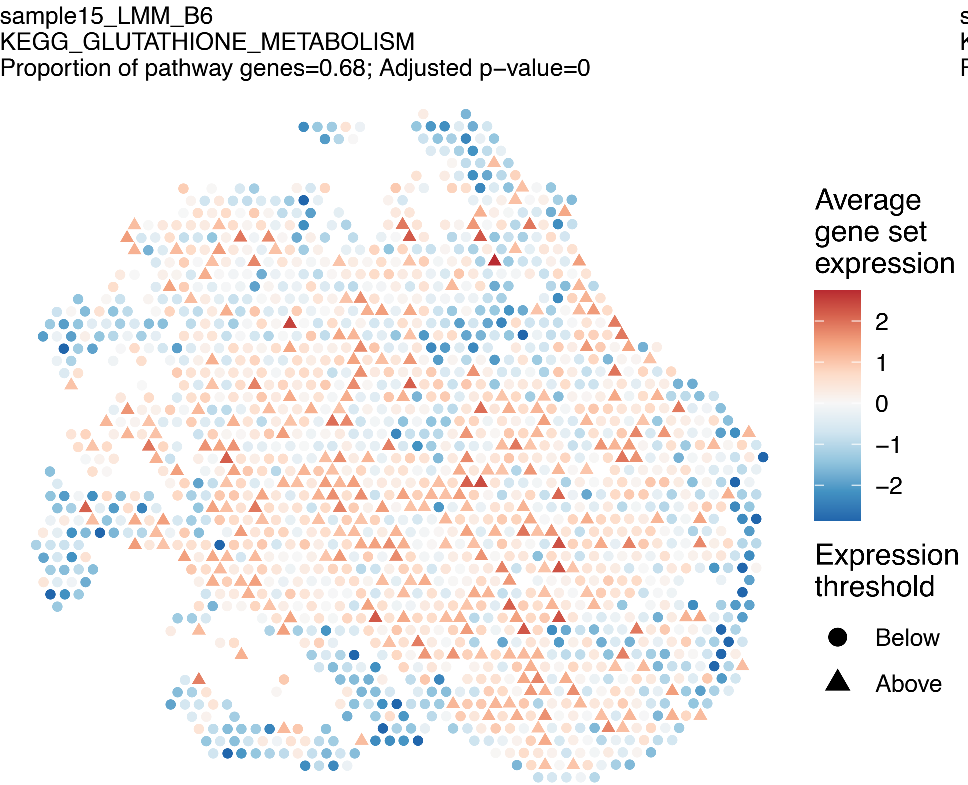
sample15_LMM_B6
KEGG_GLUTATHIONE_METABOLISM
Proportion of pathway genes=0.68; Adjusted p−value=0
Average gene set expression
2
1
0
−1
−2
Expression threshold
Below
Above

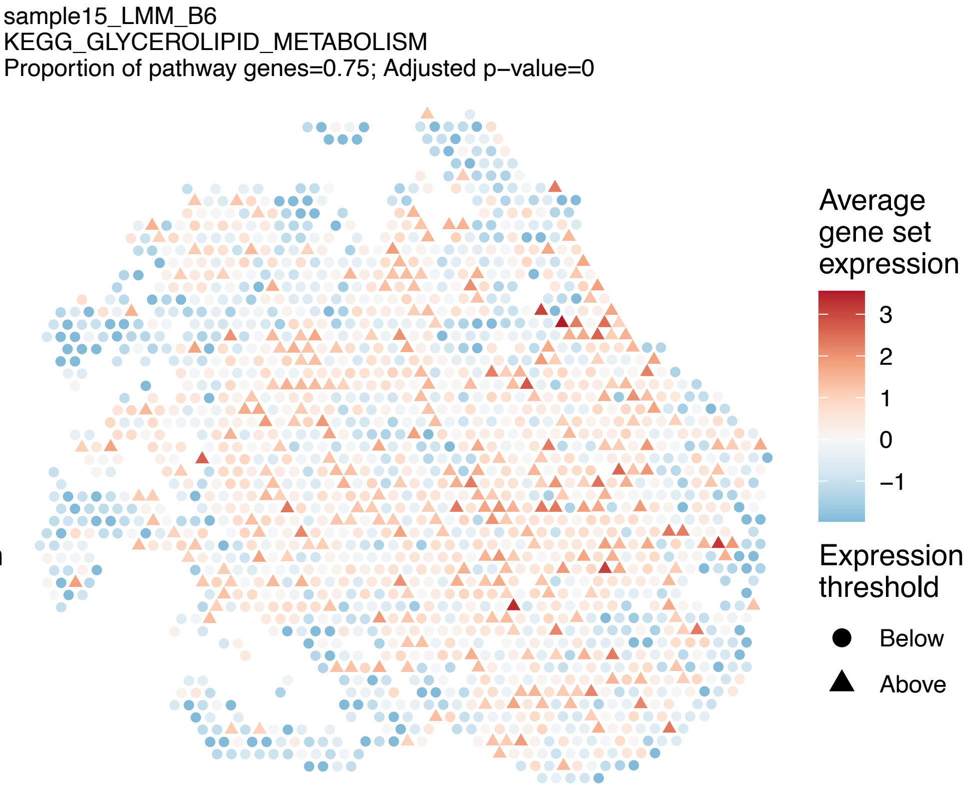
sample15_LMM_B6
KEGG_GLYCEROLIPID_METABOLISM
Proportion of pathway genes=0.75; Adjusted p−value=0
Average gene set expression
3
2
1
0
−1
Expression threshold
Below
Above

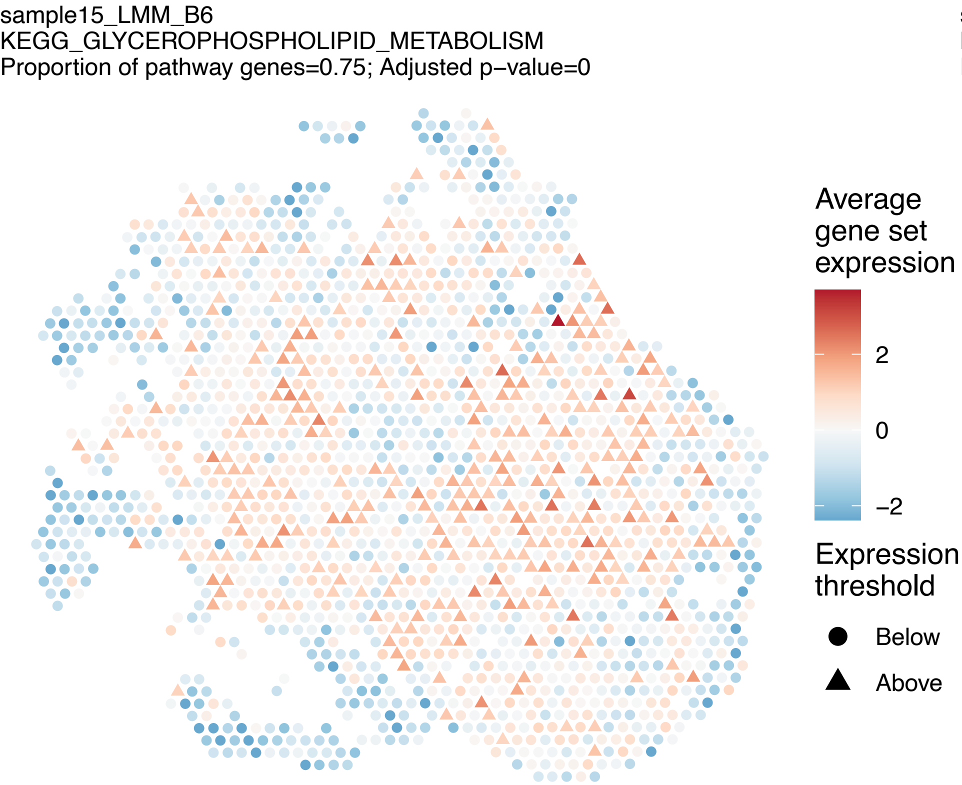
sample15_LMM_B6
KEGG_GLYCEROPHOSPHOLIPID_METABOLISM
Proportion of pathway genes=0.75; Adjusted p−value=0
Average gene set expression
2
0
−2
Expression threshold
Below
Above

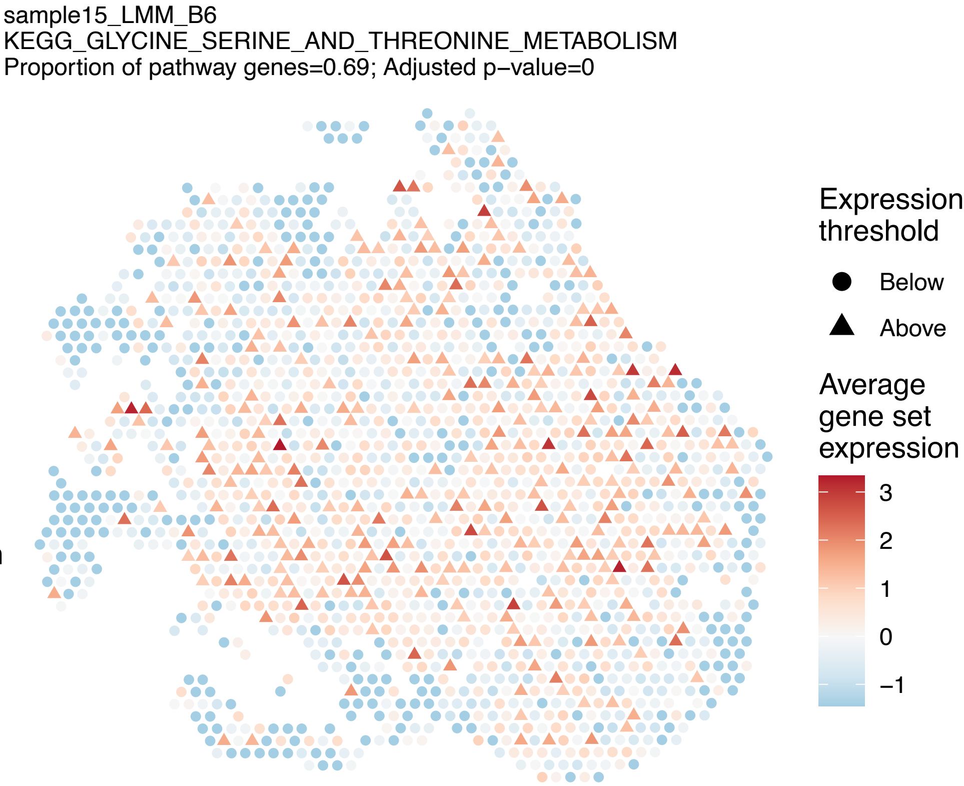
sample15_LMM_B6
KEGG_GLYCINE_SERINE_AND_THREONINE_METABOLISM
Proportion of pathway genes=0.69; Adjusted p−value=0
Expression threshold
Below
Above
Average gene set expression
3
2
1
0
−1

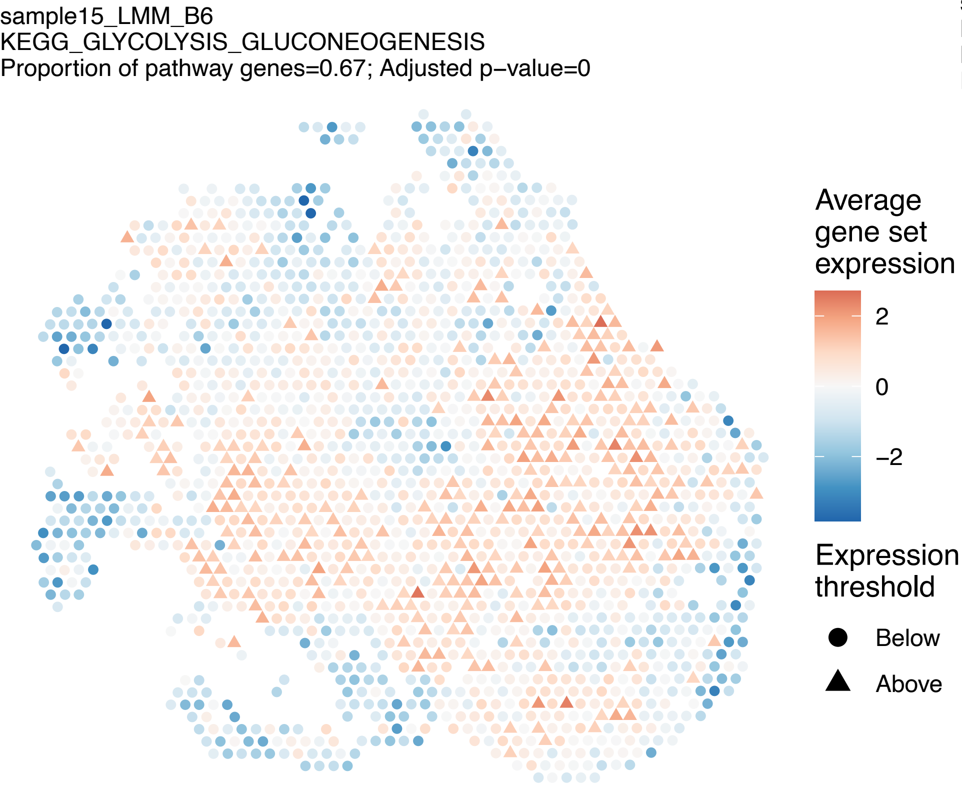
sample15_LMM_B6
KEGG_GLYCOLYSIS_GLUCONEOGENESIS
Proportion of pathway genes=0.67; Adjusted p−value=0
Average gene set expression
2
0
−2
Expression threshold
Below
Above

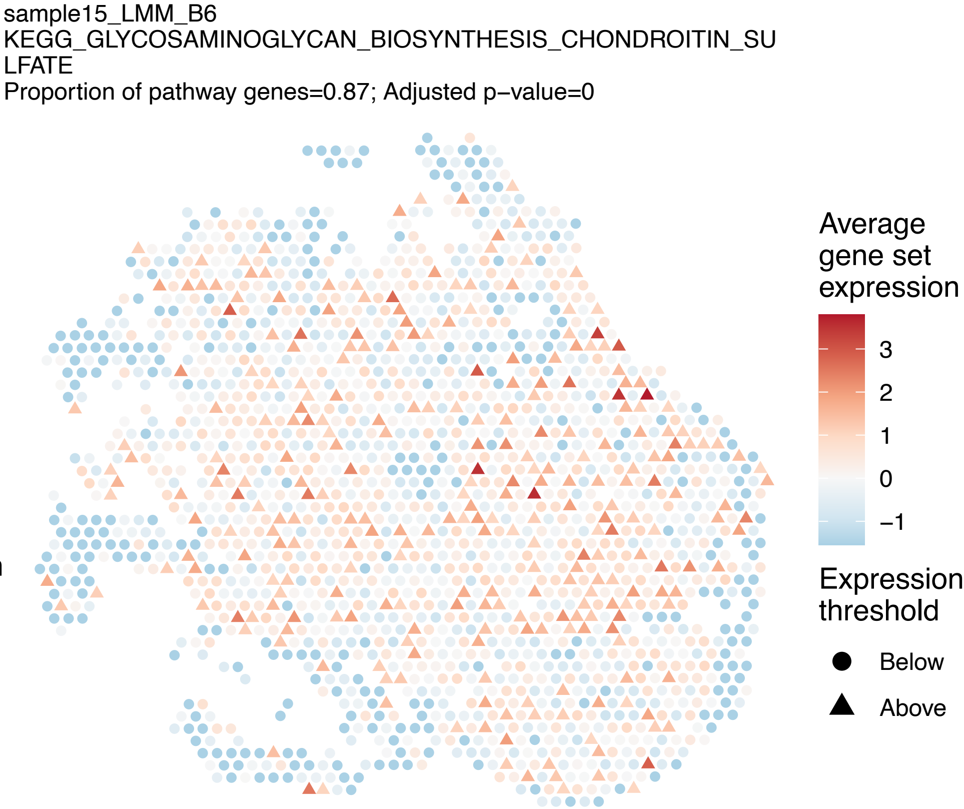
sample15_LMM_B6
KEGG_GLYCOSAMINOGLYCAN_BIOSYNTHESIS_CHONDROITIN_SU
LFATE
Proportion of pathway genes=0.87; Adjusted p−value=0
Average gene set expression
3
2
1
0
−1
Expression threshold
Below
Above

sample15_LMM_B6
KEGG_GLYCOSAMINOGLYCAN_BIOSYNTHESIS_HEPARAN_SULFAT
E
Proportion of pathway genes=0.63; Adjusted p−value=0

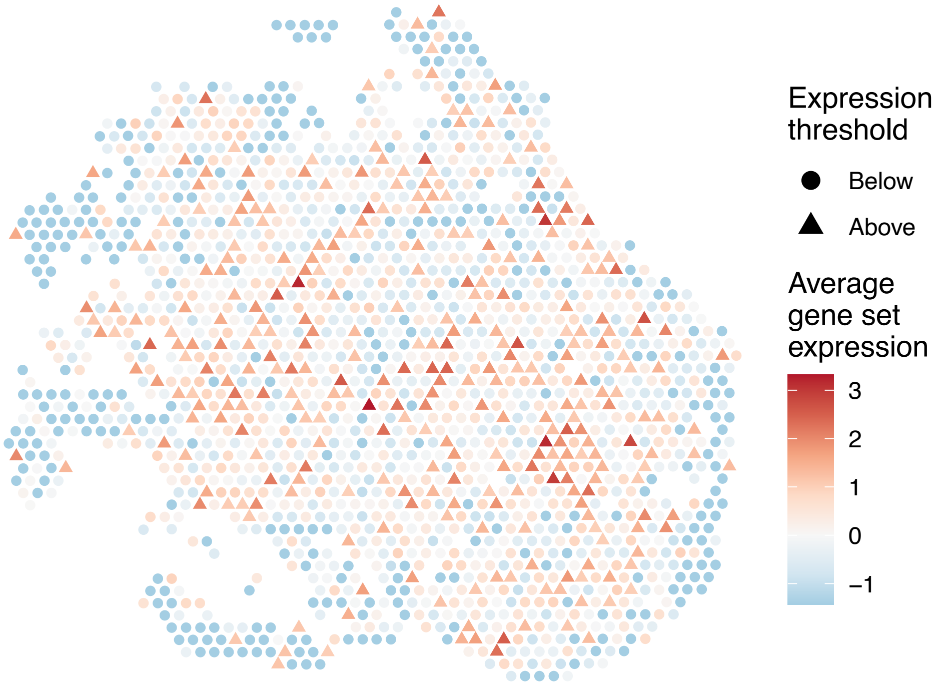


sample15_LMM_B6
KEGG_GLYCOSAMINOGLYCAN_BIOSYNTHESIS_KERATAN_SULFAT
E
Proportion of pathway genes=0.93; Adjusted p−value=0

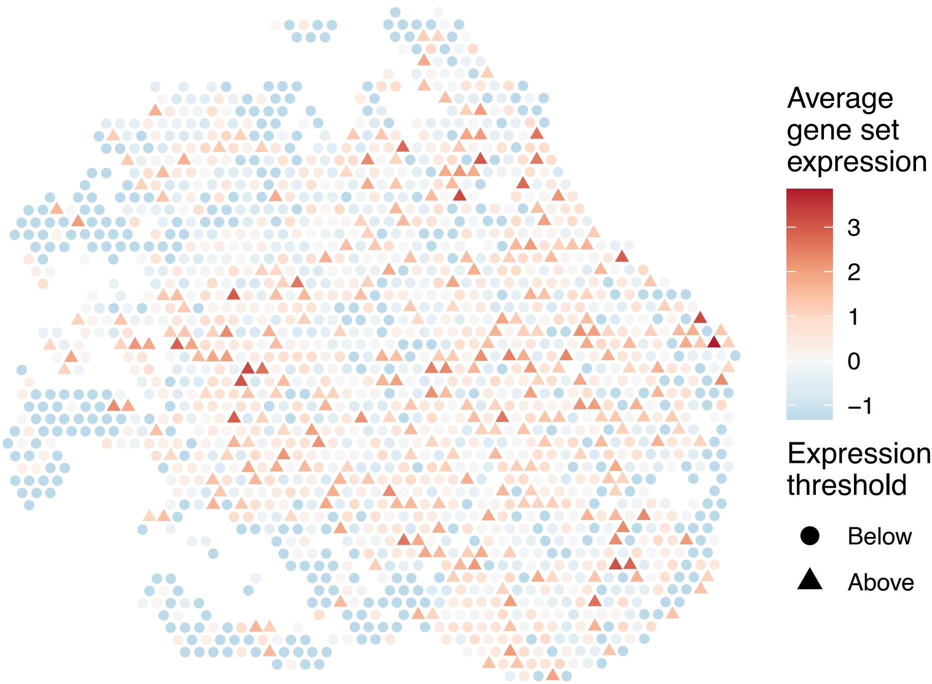


sample15_LMM_B6
KEGG_GLYCOSAMINOGLYCAN_DEGRADATION
Proportion of pathway genes=0.71; Adjusted p−value=0

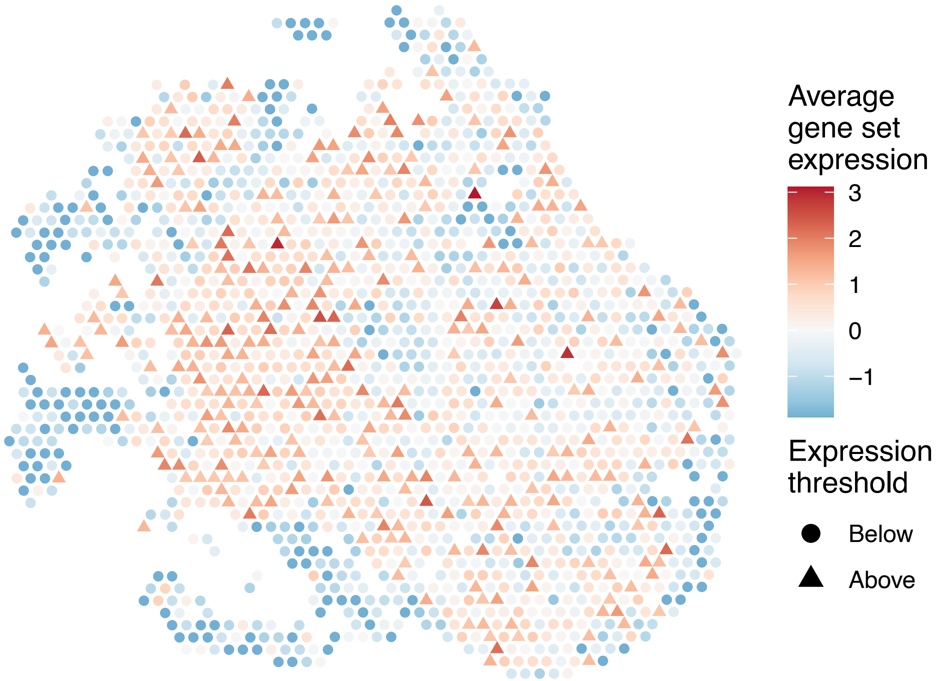


sample15_LMM_B6
KEGG_GLYCOSPHINGOLIPID_BIOSYNTHESIS_GANGLIO_SERIES
Proportion of pathway genes=0.79; Adjusted p−value=0

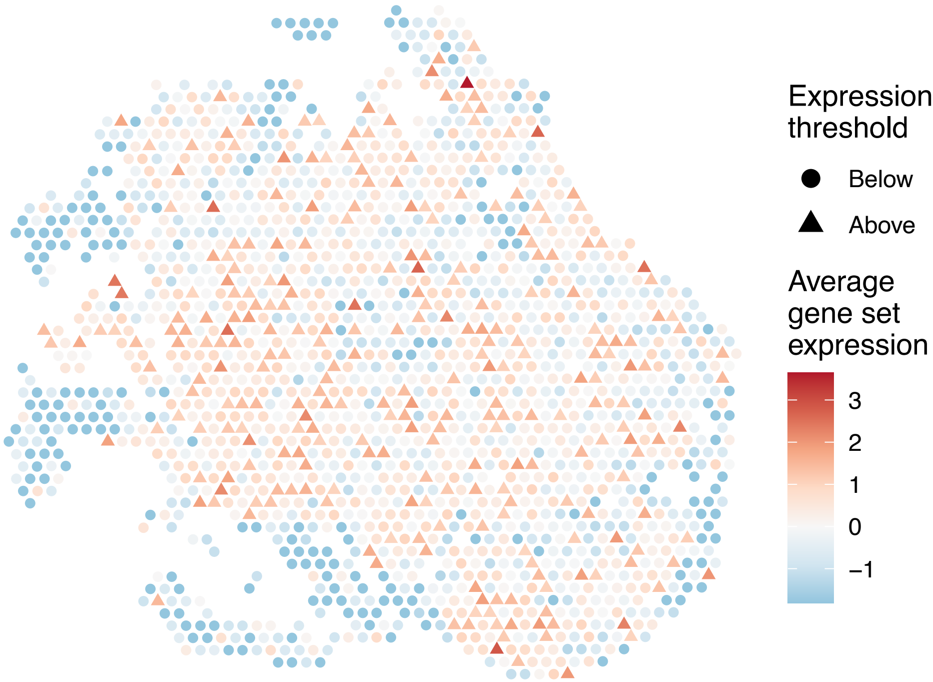

sample15_LMM_B6
KEGG_GLYCOSPHINGOLIPID_BIOSYNTHESIS_GLOBO_SERIES
Proportion of pathway genes=0.71; Adjusted p−value=0

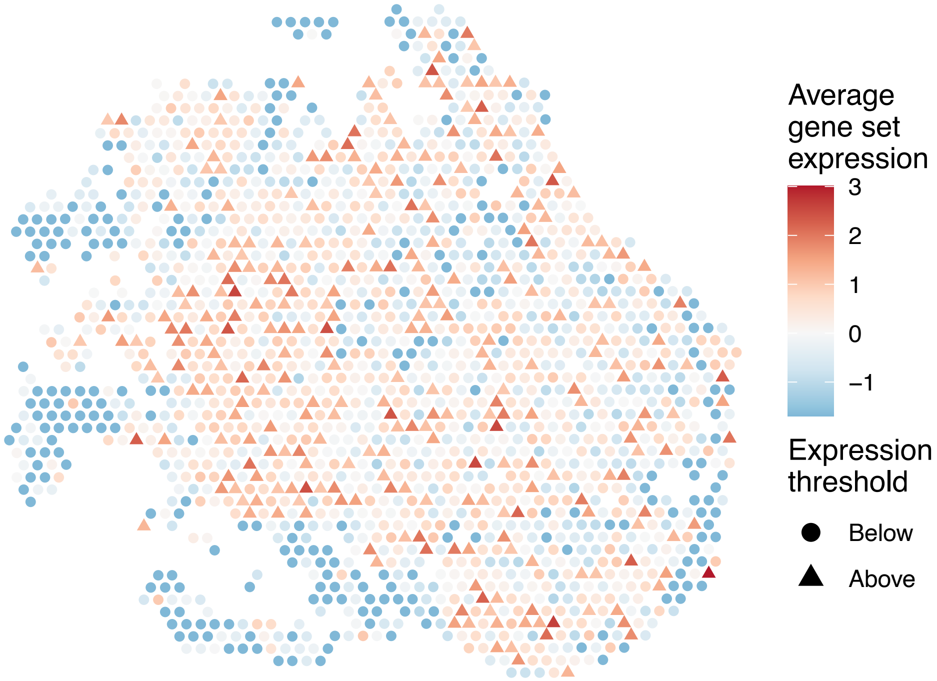


sample15_LMM_B6
KEGG_GLYCOSPHINGOLIPID_BIOSYNTHESIS_LACTO_AND_NEOL
ACTO_SERIES
Proportion of pathway genes=0.52; Adjusted p−value=0

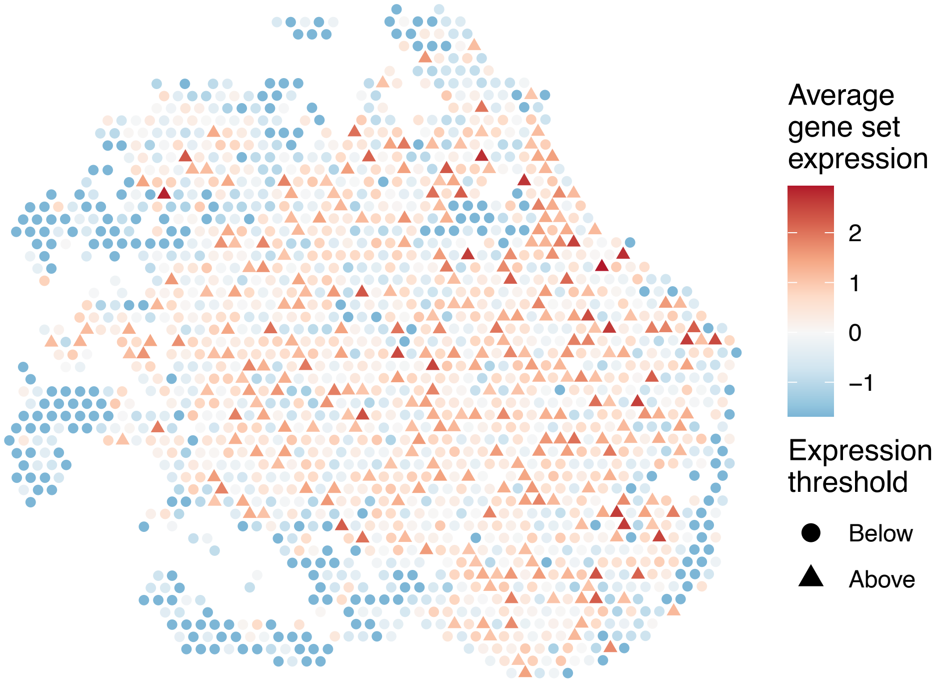


sample15_LMM_B6
KEGG_GLYCOSYLPHOSPHATIDYLINOSITOL_GPI_ANCHOR_BIOSY
NTHESIS
Proportion of pathway genes=0.93; Adjusted p−value=0.0011

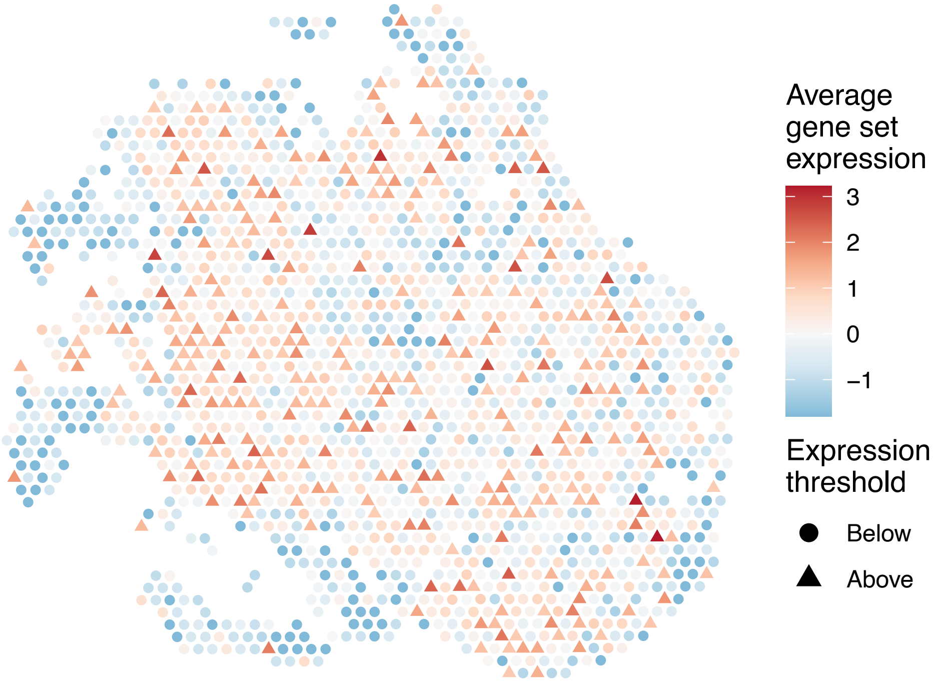


sample15_LMM_B6
KEGG_GLYOXYLATE_AND_DICARBOXYLATE_METABOLISM
Proportion of pathway genes=0.88; Adjusted p−value=0

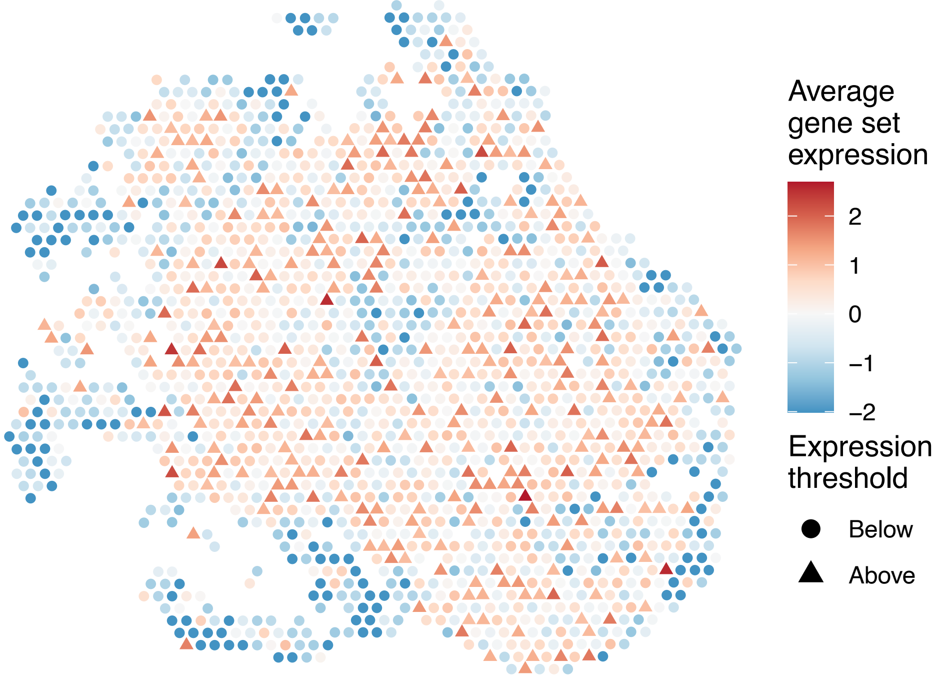

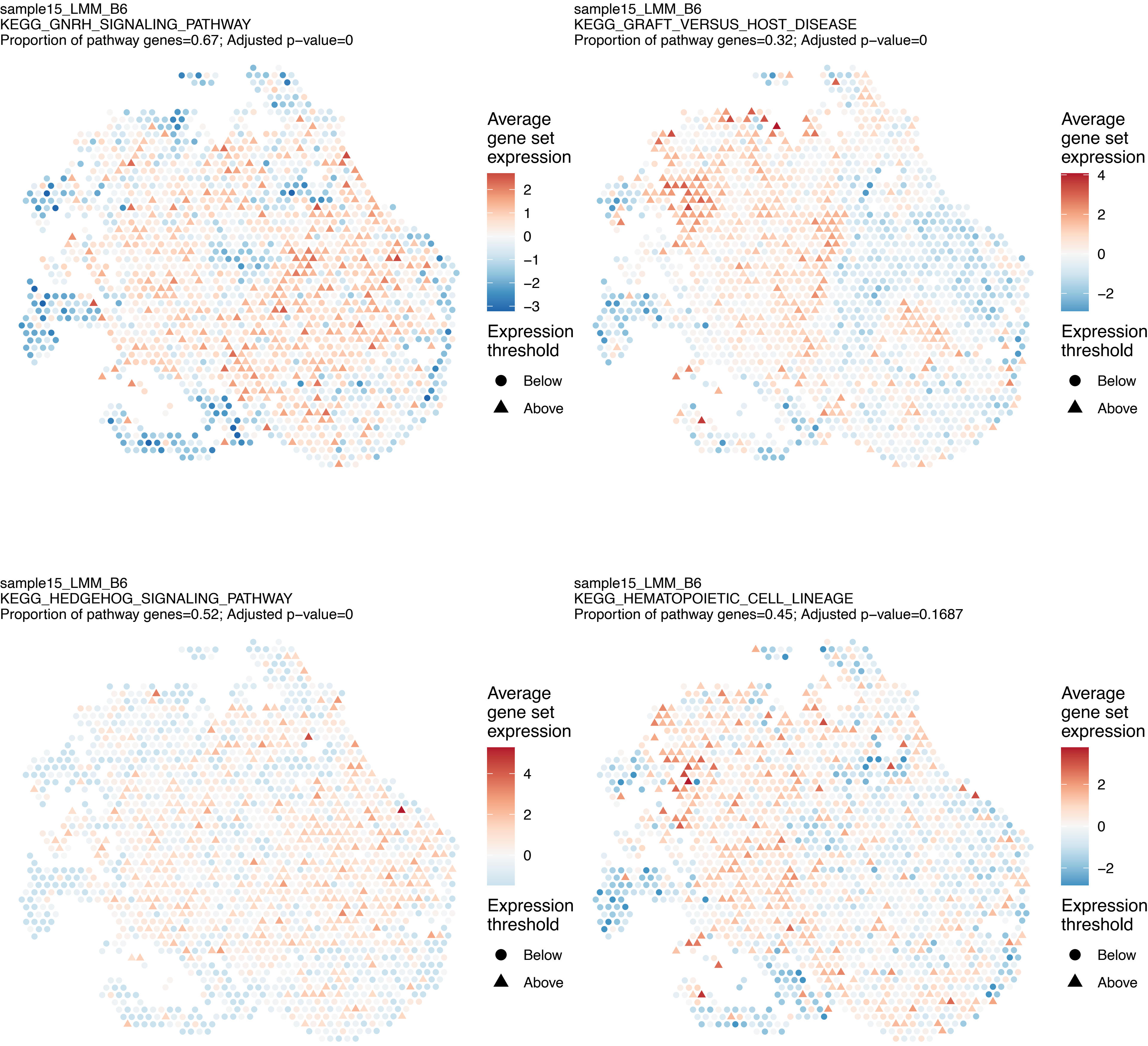

sample15_LMM_B6
KEGG_GNRH_SIGNALING_PATHWAY
Proportion of pathway genes=0.67; Adjusted p−value=0
Average gene set expression
2
1
0
−1
−2
−3
Expression threshold
Below
Above
sample15_LMM_B6
KEGG_GRAFT_VERSUS_HOST_DISEASE
Proportion of pathway genes=0.32; Adjusted p−value=0
Average gene set expression
4
2
0
−2
Expression threshold
Below
Above
sample15_LMM_B6
KEGG_HEDGEHOG_SIGNALING_PATHWAY
Proportion of pathway genes=0.52; Adjusted p−value=0
Average gene set expression
4
2
0
Expression threshold
Below
Above
sample15_LMM_B6
KEGG_HEMATOPOIETIC_CELL_LINEAGE
Proportion of pathway genes=0.45; Adjusted p−value=0.1687
Average gene set expression
2
0
−2
Expression threshold
Below
Above

sample15_LMM_B6
KEGG_HISTIDINE_METABOLISM
Proportion of pathway genes=0.63; Adjusted p−value=0

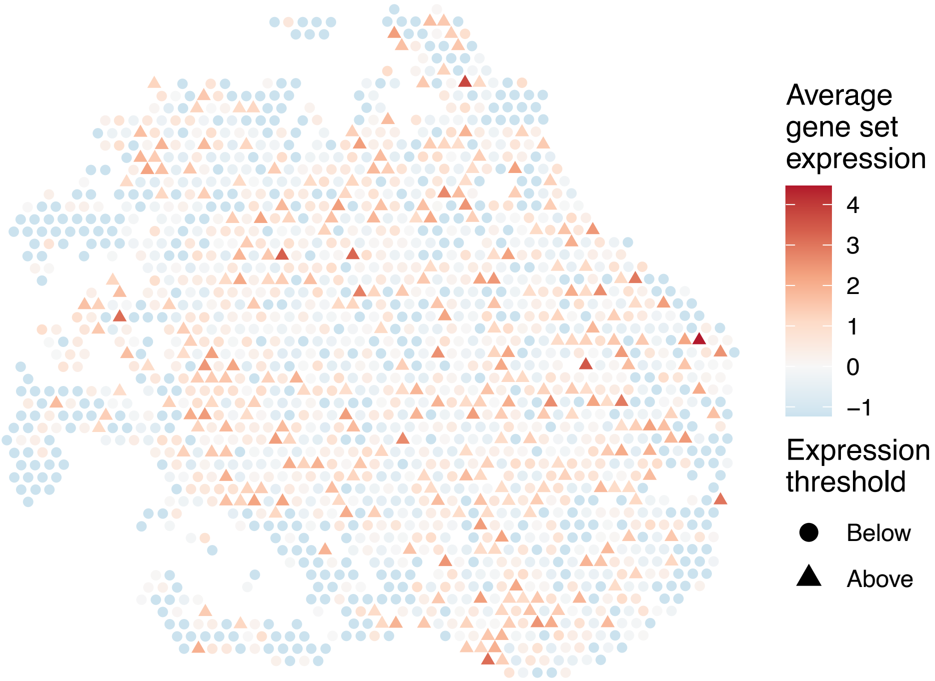


sample15_LMM_B6
KEGG_HOMOLOGOUS_RECOMBINATION
Proportion of pathway genes=0.9; Adjusted p−value=0

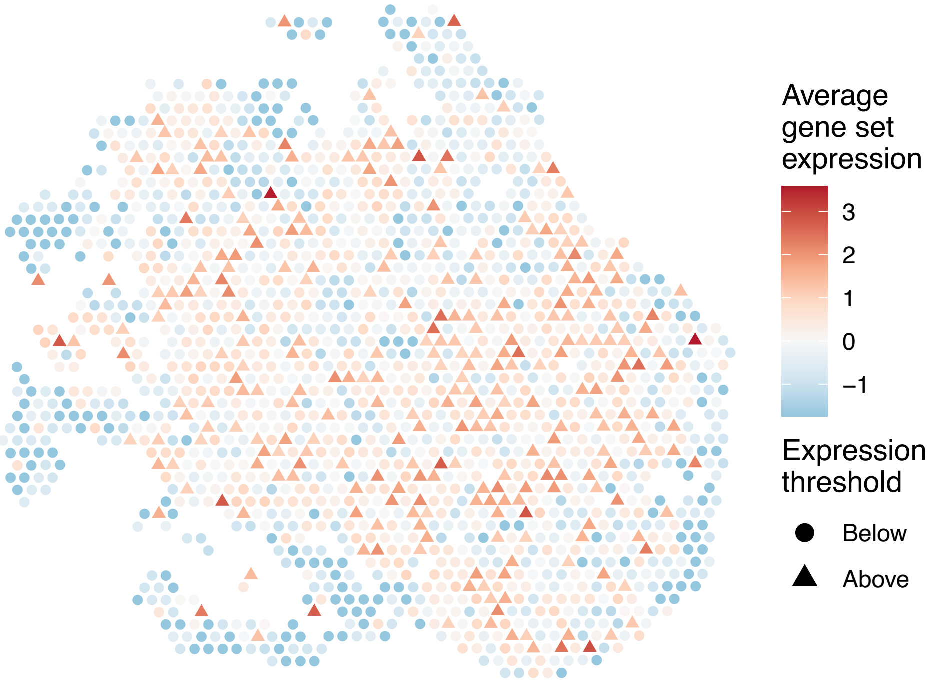


sample15_LMM_B6
KEGG_HUNTINGTONS_DISEASE
Proportion of pathway genes=0.87; Adjusted p−value=0

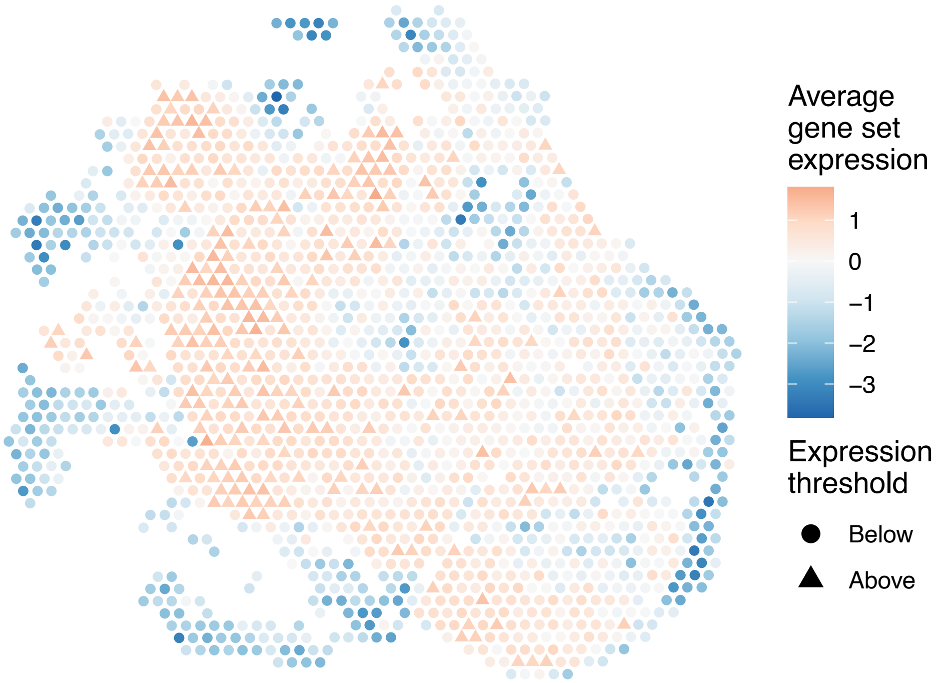


sample15_LMM_B6
KEGG_HYPERTROPHIC_CARDIOMYOPATHY_HCM
Proportion of pathway genes=0.47; Adjusted p−value=0

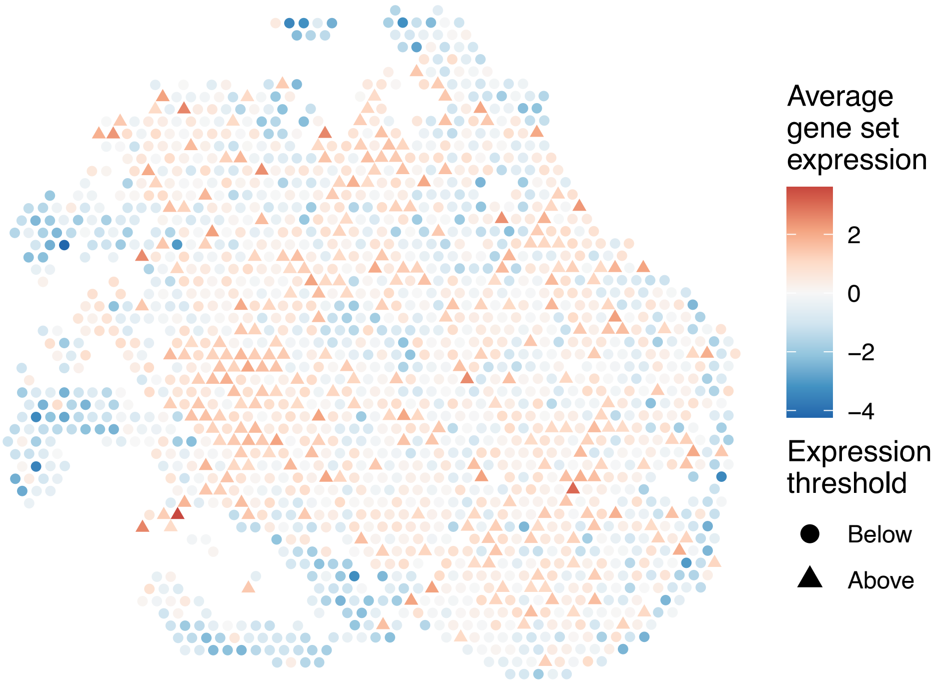

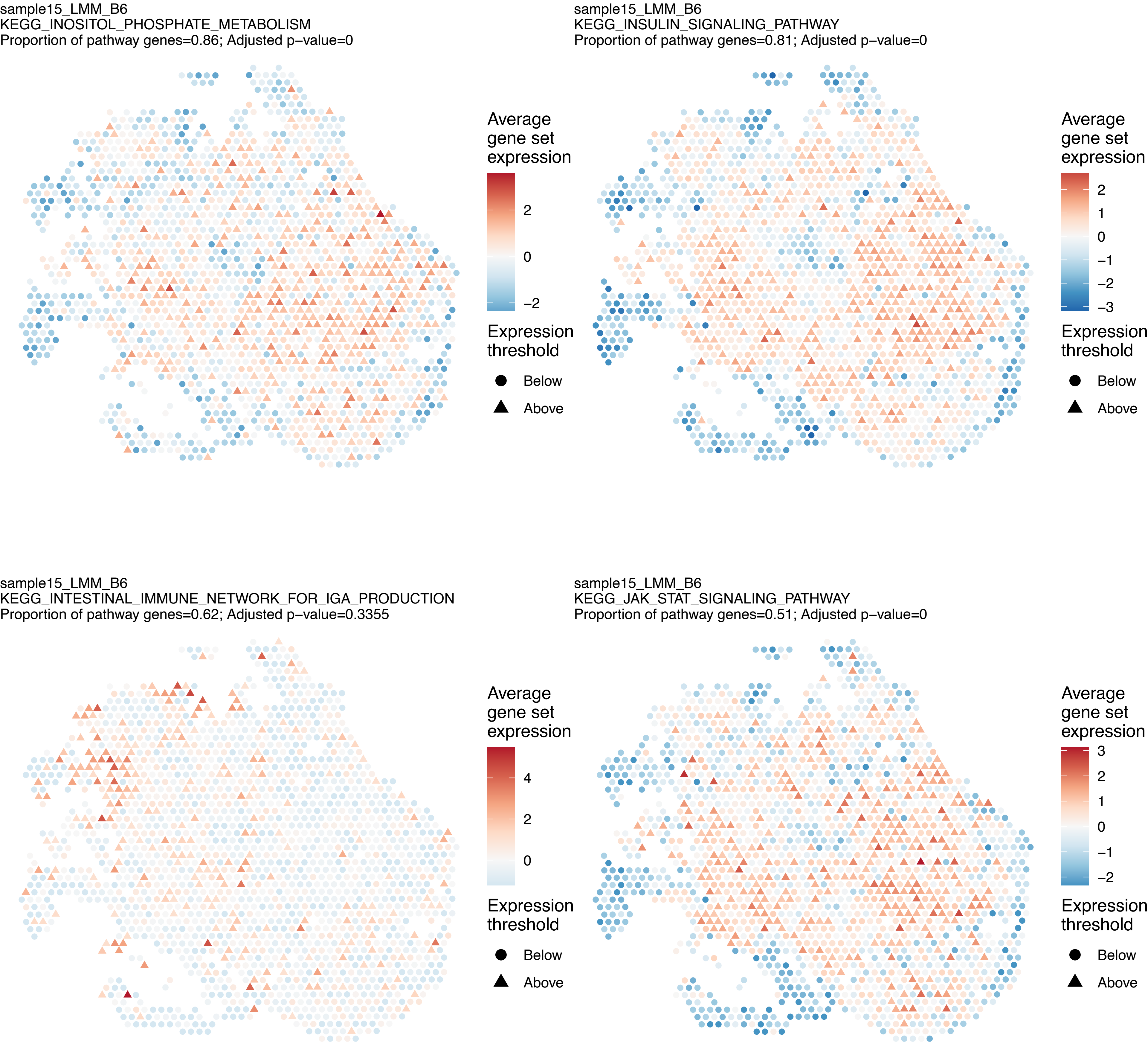
sample15_LMM_B6
KEGG_INOSITOL_PHOSPHATE_METABOLISM
Proportion of pathway genes=0.86; Adjusted p–value=0
Average gene set expression
2
0
−2
Expression threshold
Below
Above
sample15_LMM_B6
KEGG_INSULIN_SIGNALING_PATHWAY
Proportion of pathway genes=0.81; Adjusted p–value=0
Average gene set expression
2
1
0
−1
−2
−3
Expression threshold
Below
Above
sample15_LMM_B6
KEGG_INTESTINAL_IMMUNE_NETWORK_FOR_IGA_PRODUCTION
Proportion of pathway genes=0.62; Adjusted p–value=0.3355
Average gene set expression
4
2
0
Expression threshold
Below
Above
sample15_LMM_B6
KEGG_JAK_STAT_SIGNALING_PATHWAY
Proportion of pathway genes=0.51; Adjusted p–value=0
Average gene set expression
3
2
1
0
−1
−2
Expression threshold
Below
Above

sample15_LMM_B6
KEGG_LEISHMANIA_INFECTION
Proportion of pathway genes=0.67; Adjusted p−value=0.0482

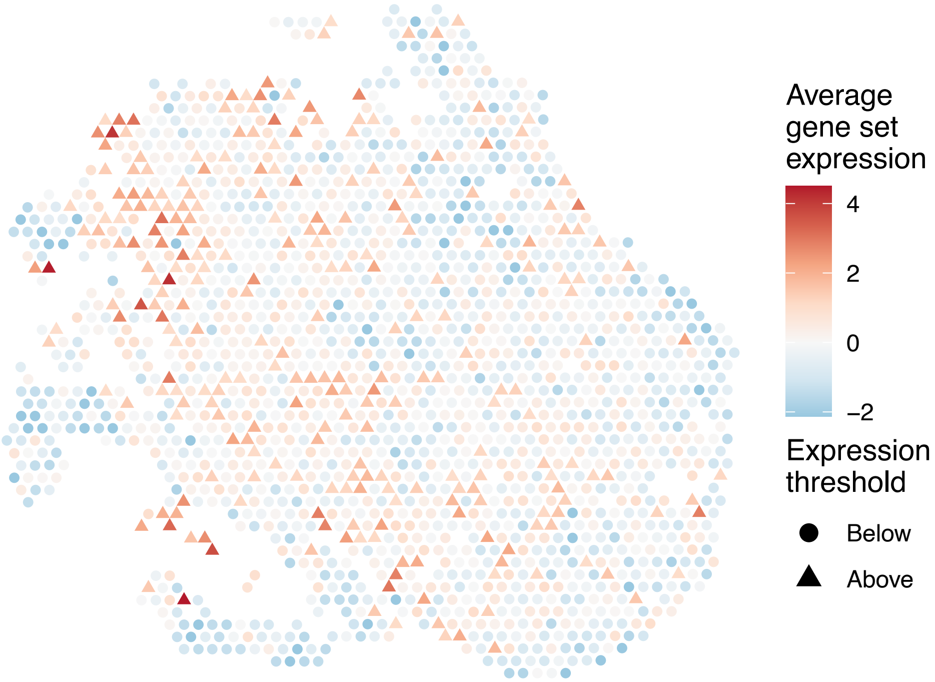


sample15_LMM_B6
KEGG_LEUKOCYTE_TRANSENDOTHELIAL_MIGRATION
Proportion of pathway genes=0.69; Adjusted p−value=0

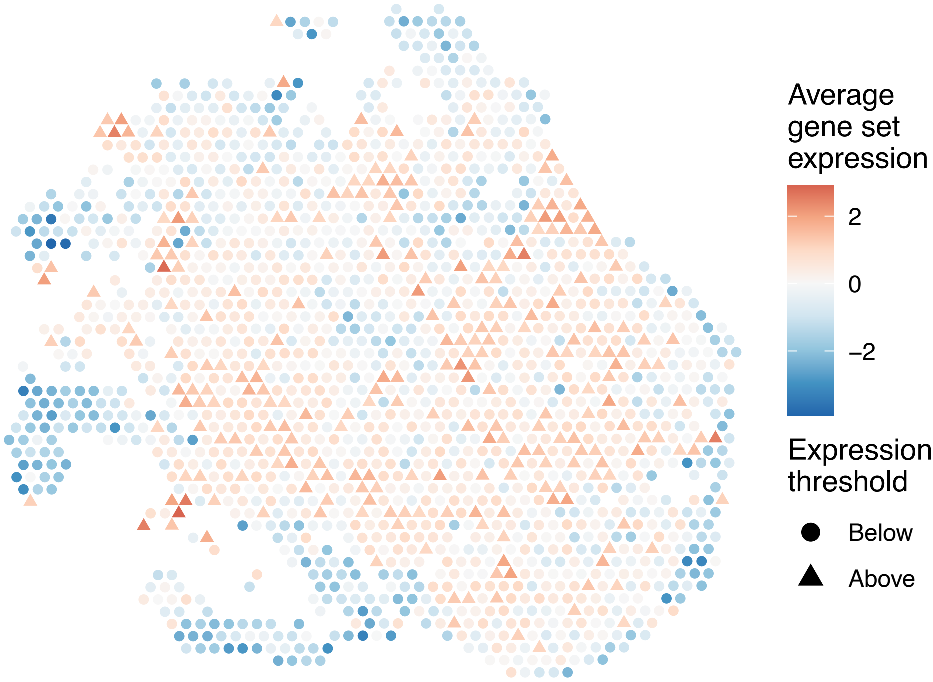


sample15_LMM_B6
KEGG_LIMONENE_AND_PINENE_DEGRADATION
Proportion of pathway genes=1; Adjusted p−value=0

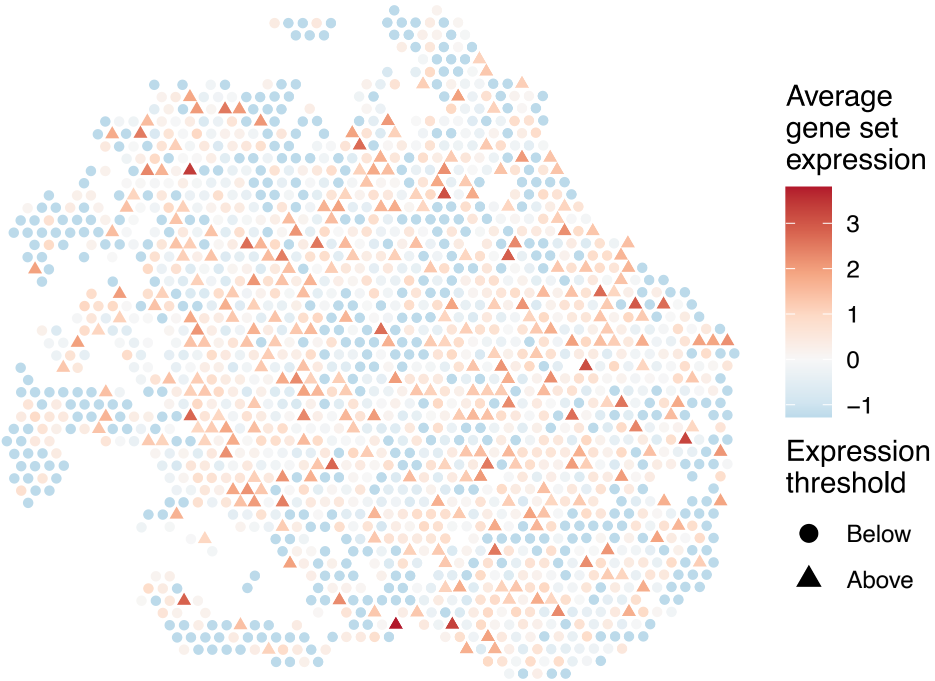


sample15_LMM_B6
KEGG_LONG_TERM_DEPRESSION
Proportion of pathway genes=0.59; Adjusted p−value=0

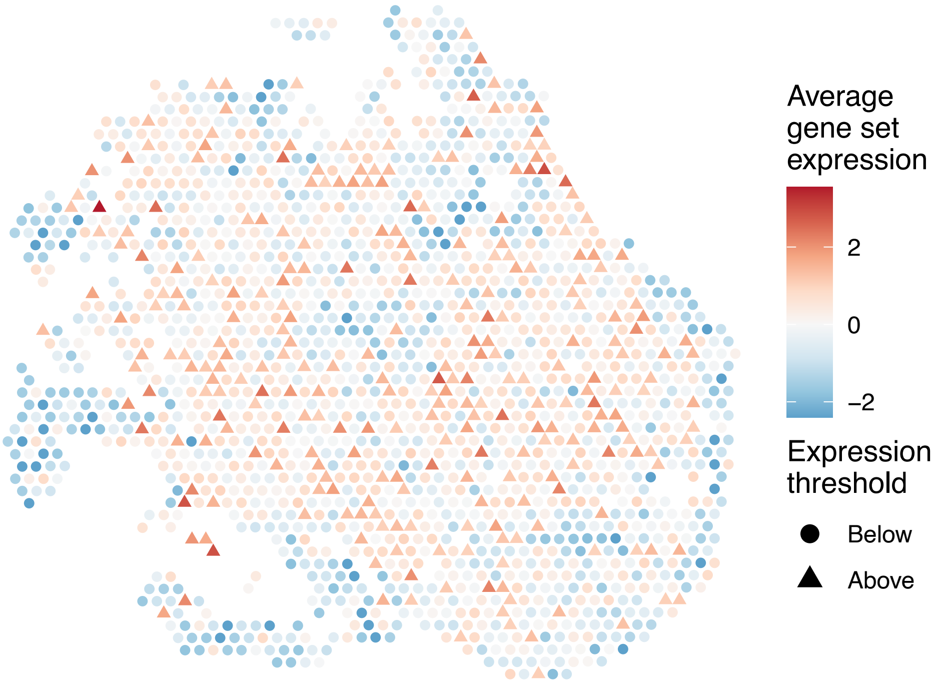

sample15_LMM_B6
KEGG_LONG_TERM_POTENTIATION
Proportion of pathway genes=0.68; Adjusted p−value=0

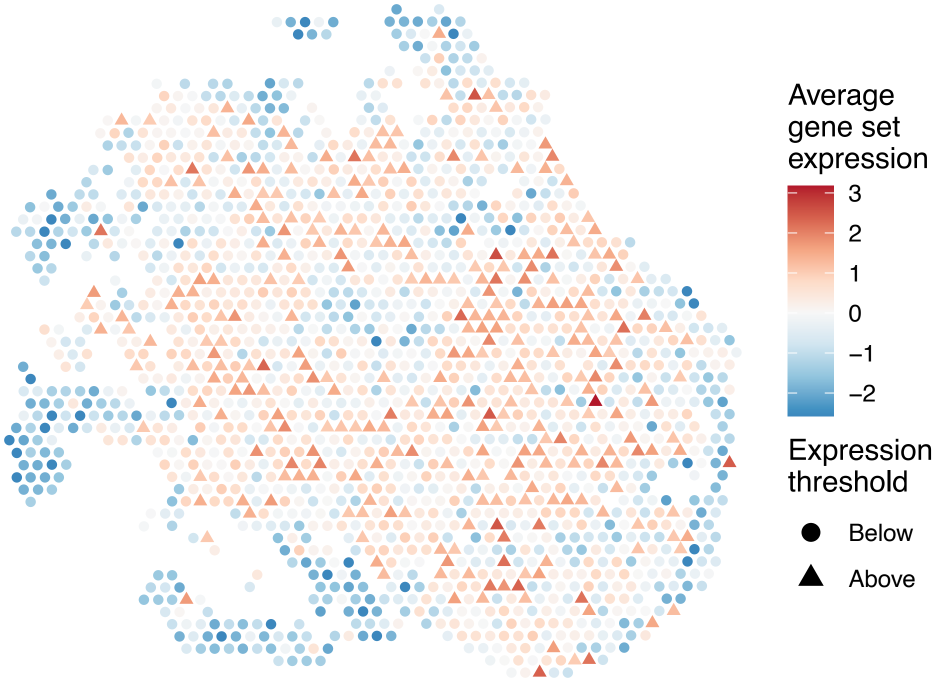


sample15_LMM_B6
KEGG_LYSINE_DEGRADATION
Proportion of pathway genes=0.92; Adjusted p−value=0

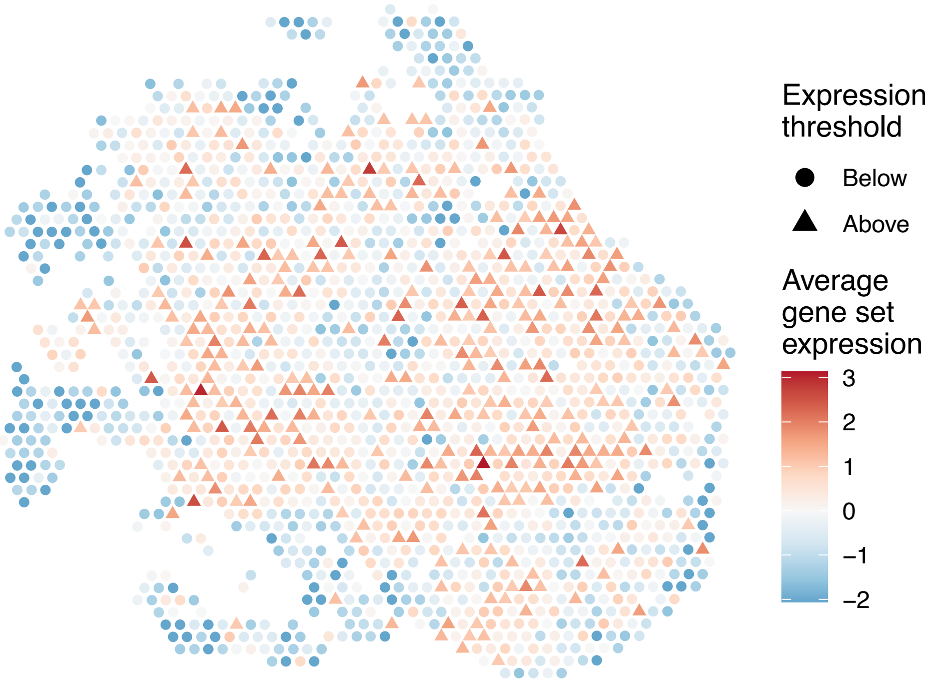


sample15_LMM_B6
KEGG_LYSOSOME
Proportion of pathway genes=0.92; Adjusted p−value=0

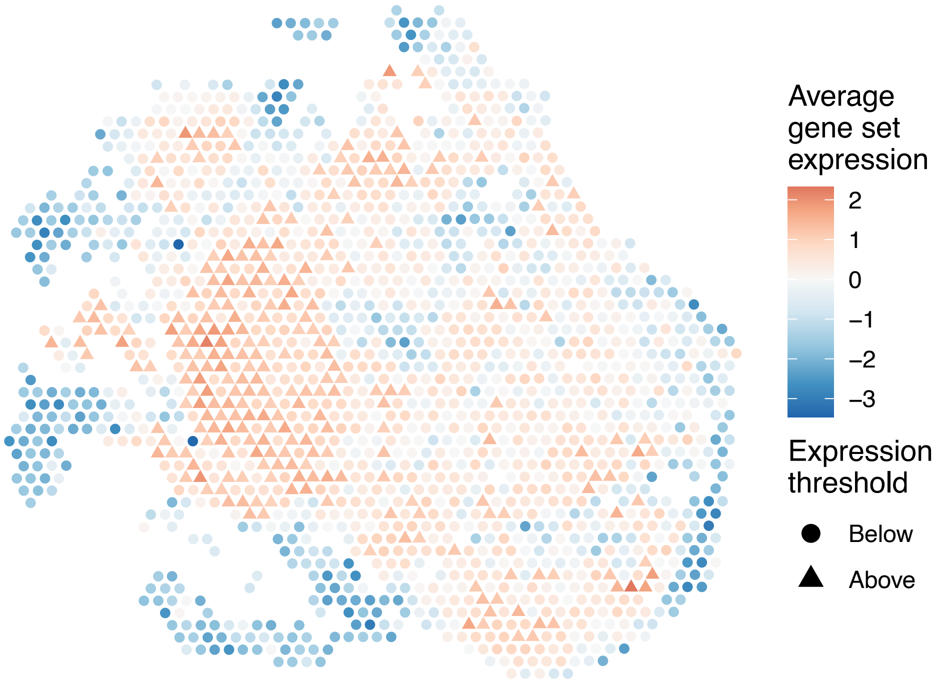


sample15_LMM_B6
KEGG_MAPK_SIGNALING_PATHWAY
Proportion of pathway genes=0.66; Adjusted p−value=0

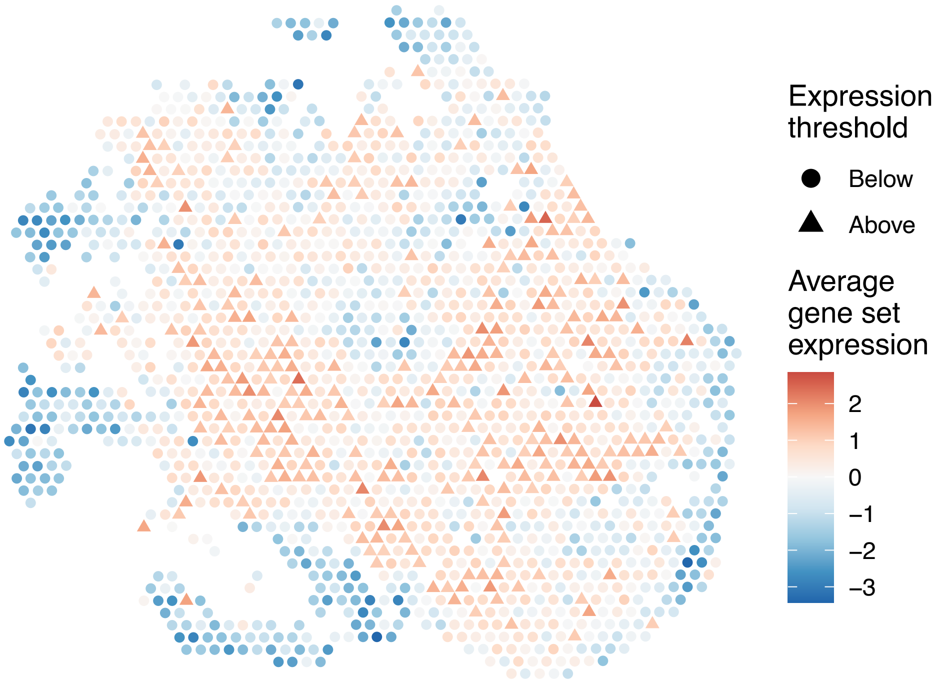

sample15_LMM_B6
KEGG_MELANOGENESIS
Proportion of pathway genes=0.63; Adjusted p−value=0

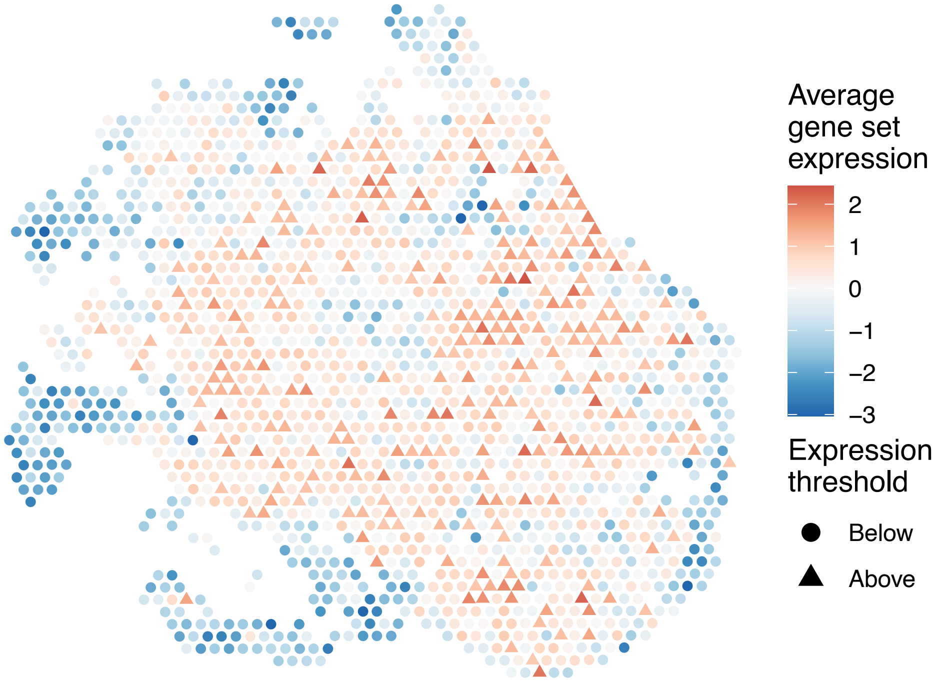


sample15_LMM_B6
KEGG_MELANOMA
Proportion of pathway genes=0.64; Adjusted p−value=0

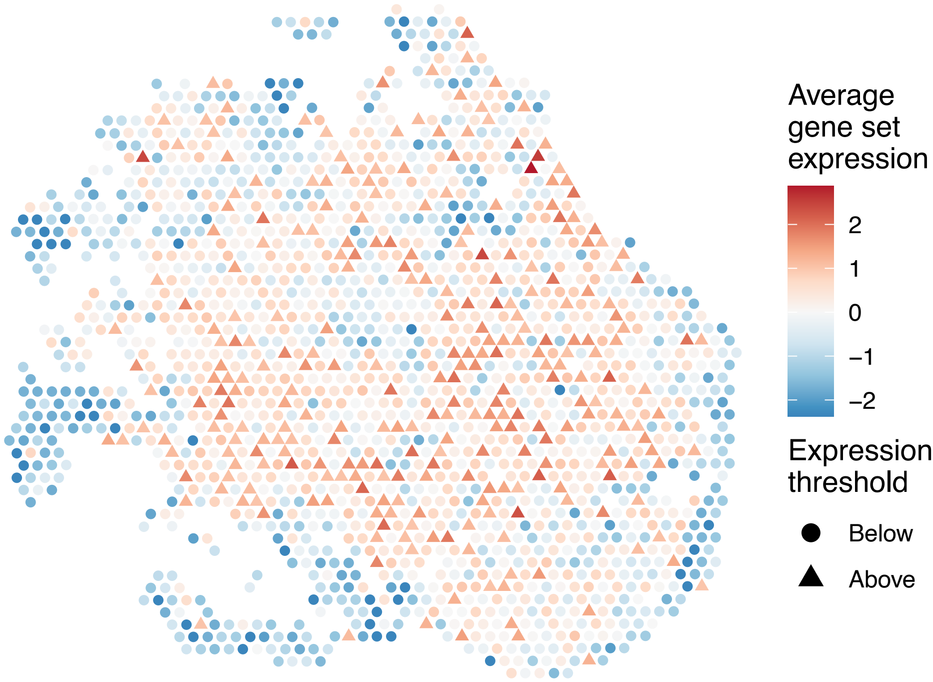


sample15_LMM_B6
KEGG_METABOLISM_OF_XENOBIOTICS_BY_CYTOCHROME_P450
Proportion of pathway genes=0.28; Adjusted p−value=0

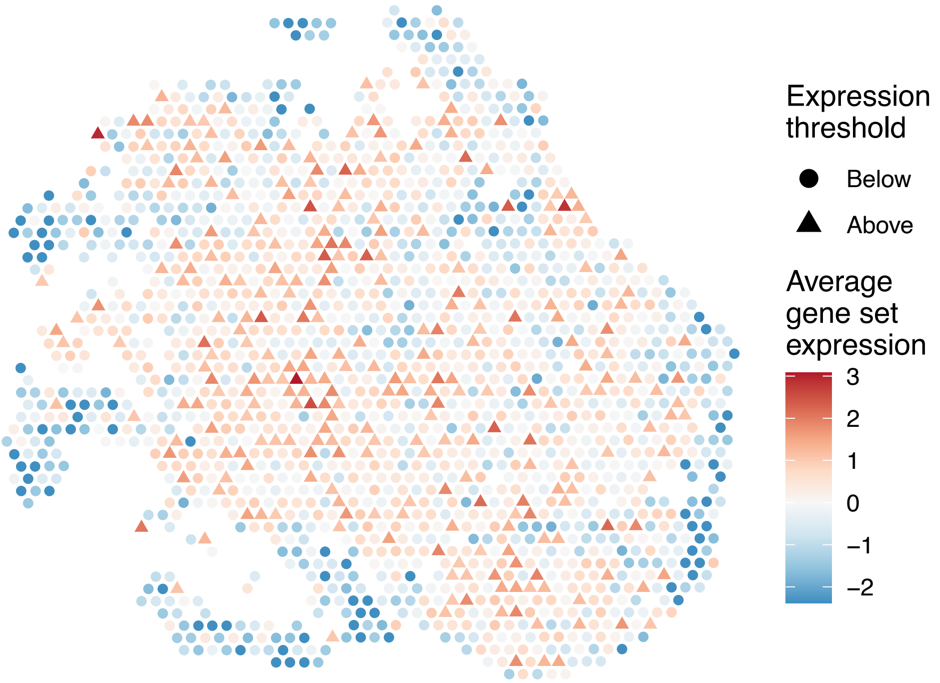


sample15_LMM_B6
KEGG_MISMATCH_REPAIR
Proportion of pathway genes=0.92; Adjusted p−value=0

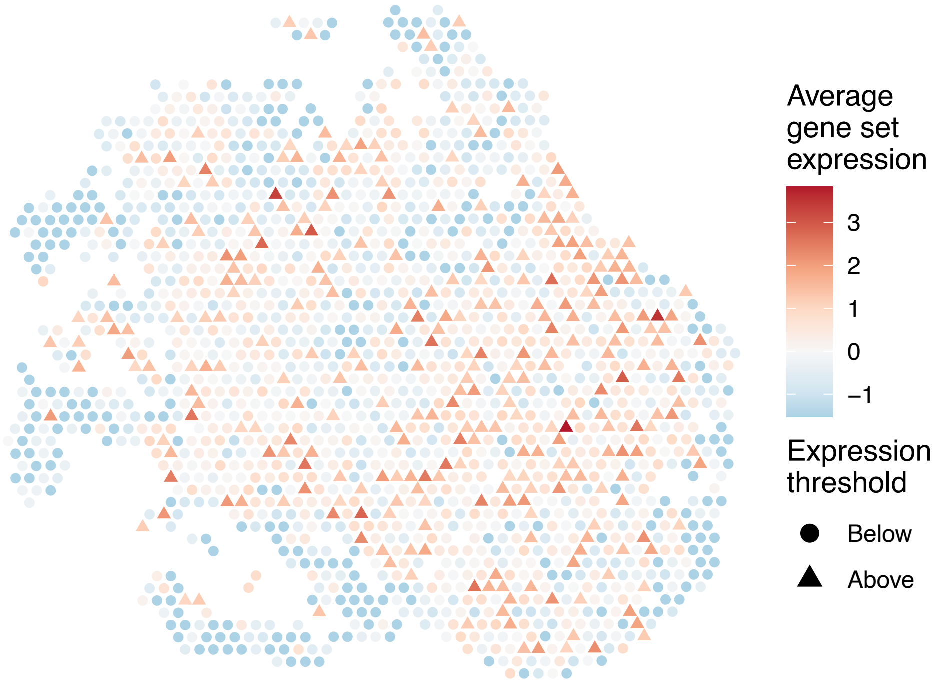

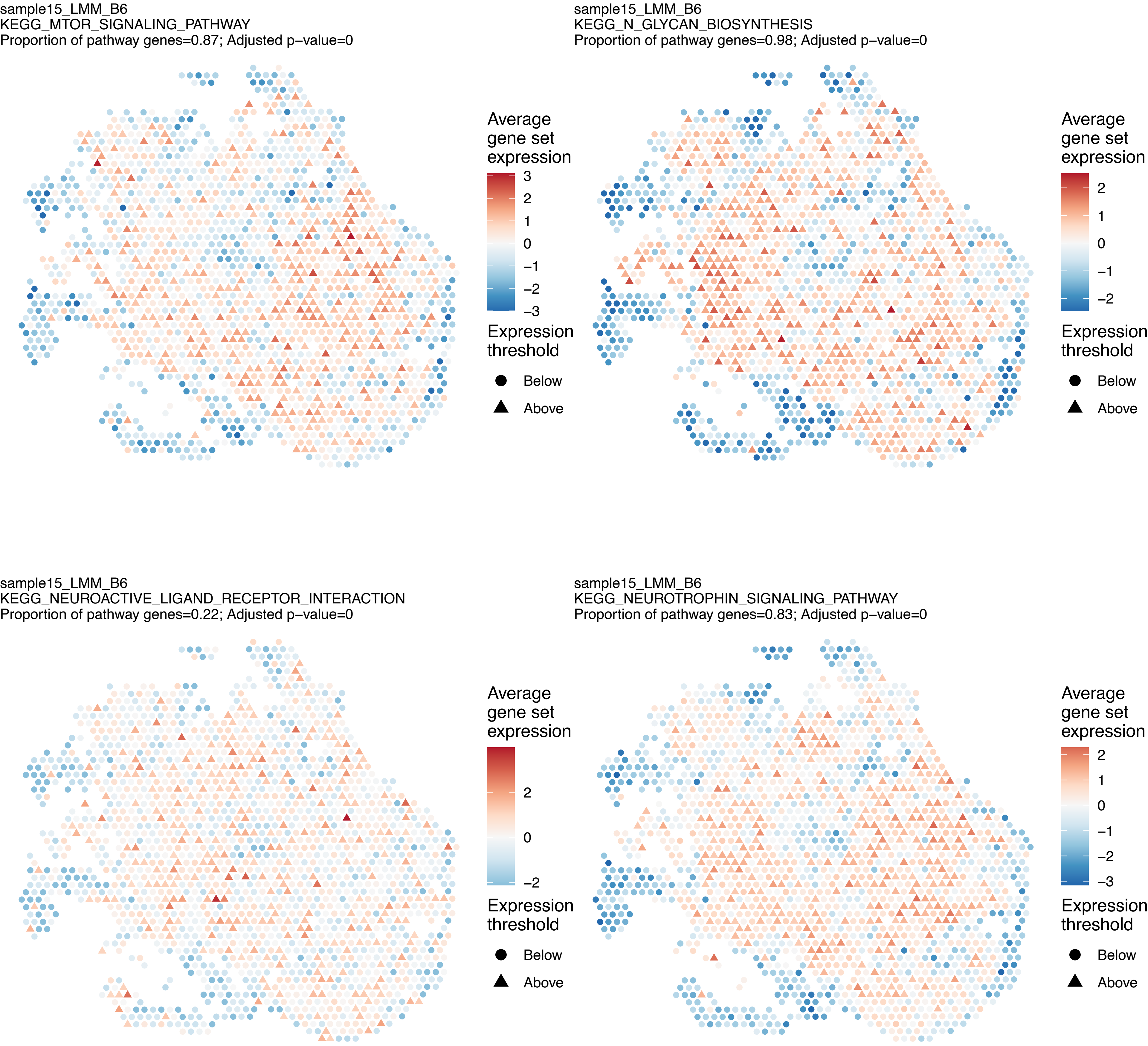
sample15_LMM_B6
KEGG_MTOR_SIGNALING_PATHWAY
Proportion of pathway genes=0.87; Adjusted p−value=0
Average gene set expression
3
2
1
0
−1
−2
−3
Expression threshold
Below
Above
sample15_LMM_B6
KEGG_N_GLYCAN_BIOSYNTHESIS
Proportion of pathway genes=0.98; Adjusted p−value=0
Average gene set expression
2
1
0
−1
−2
Expression threshold
Below
Above
sample15_LMM_B6
KEGG_NEUROACTIVE_LIGAND_RECEPTOR_INTERACTION
Proportion of pathway genes=0.22; Adjusted p−value=0
Average gene set expression
2
0
−2
Expression threshold
Below
Above
sample15_LMM_B6
KEGG_NEUROTROPHIN_SIGNALING_PATHWAY
Proportion of pathway genes=0.83; Adjusted p−value=0
Average gene set expression
2
1
0
−1
−2
−3
Expression threshold
Below
Above

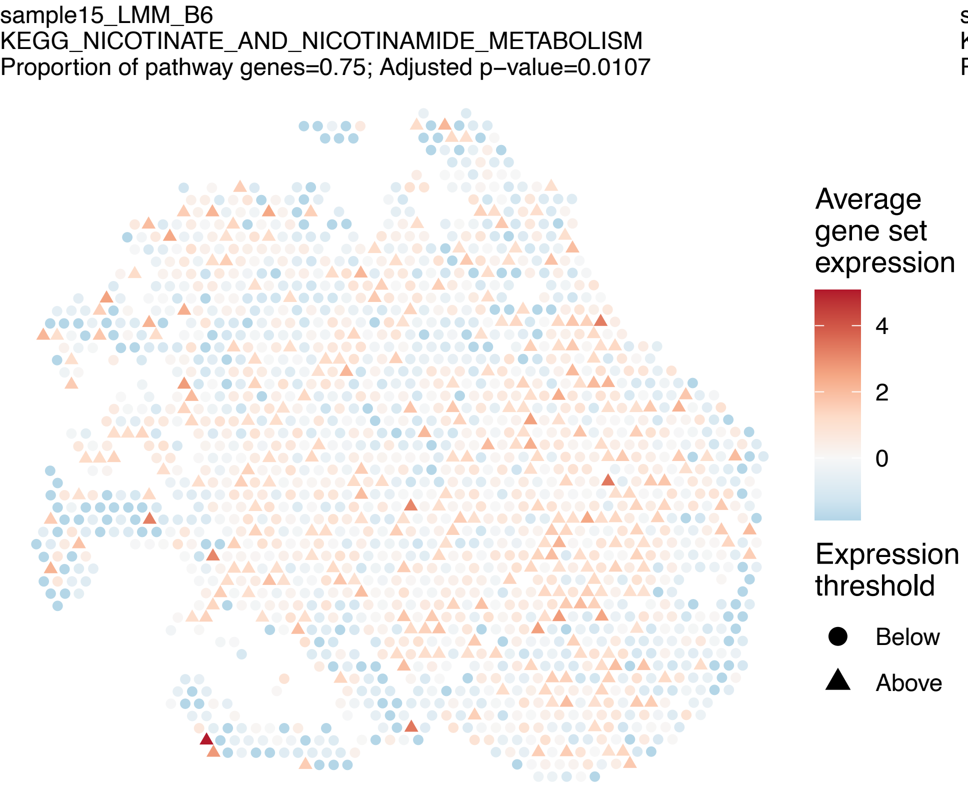
sample15_LMM_B6
KEGG_NICOTINATE_AND_NICOTINAMIDE_METABOLISM
Proportion of pathway genes=0.75; Adjusted p−value=0.0107
Average gene set expression
4
2
0
Expression threshold
Below
Above

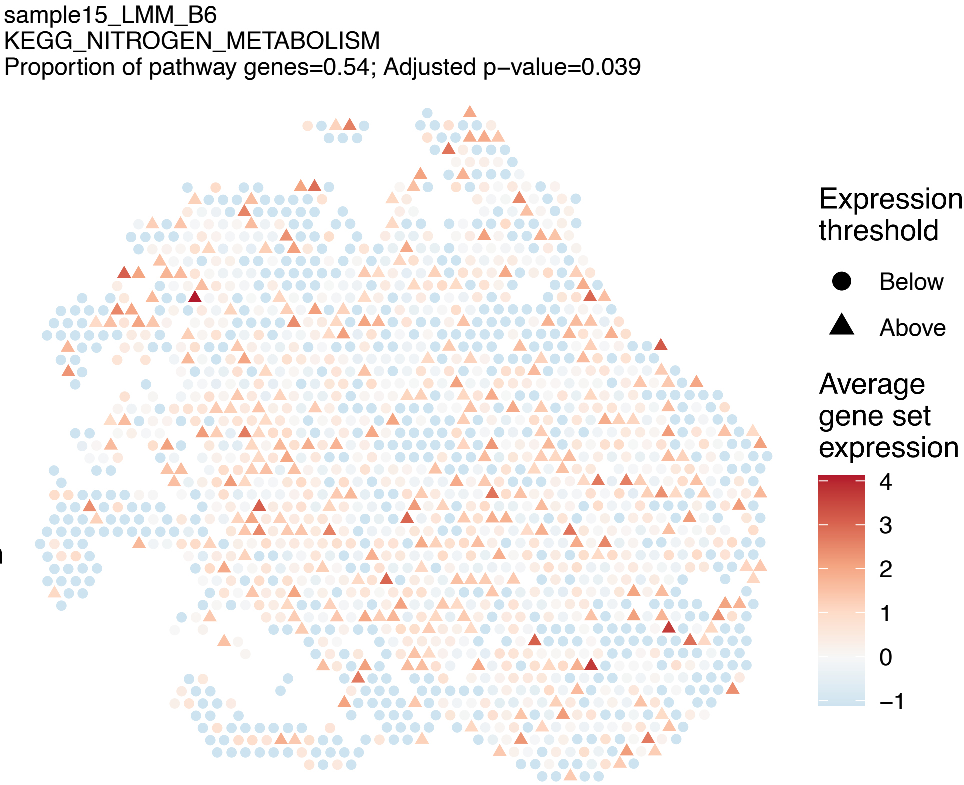
sample15_LMM_B6
KEGG_NITROGEN_METABOLISM
Proportion of pathway genes=0.54; Adjusted p−value=0.039
Expression threshold
Below
Above
Average gene set expression
4
3
2
1
0
−1

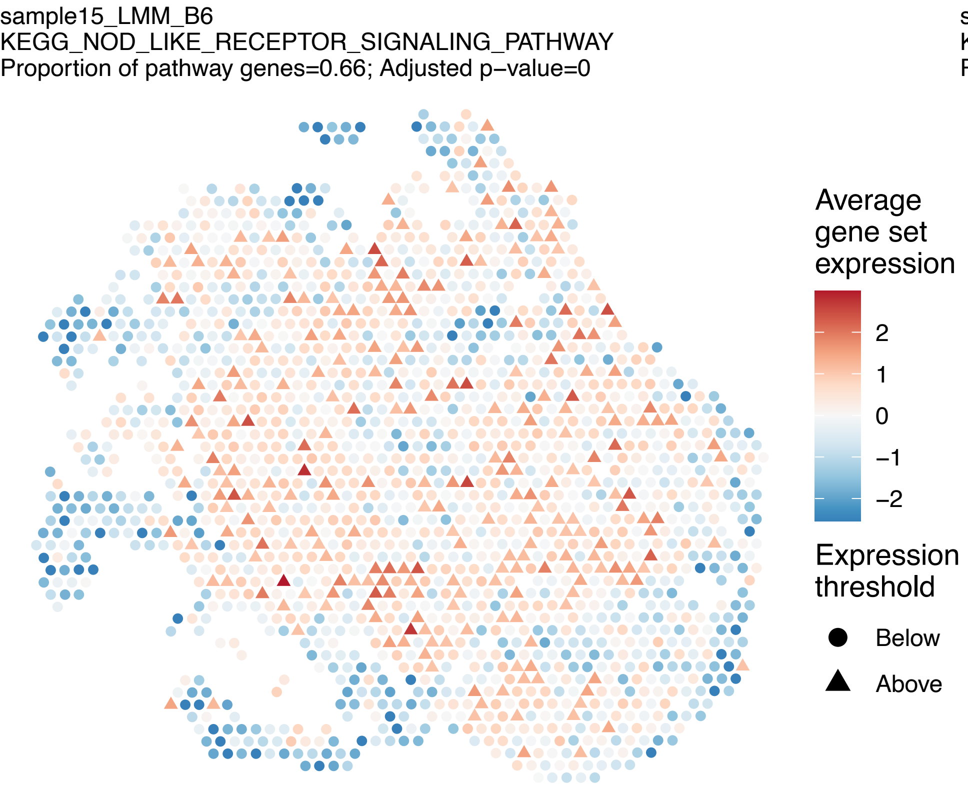
sample15_LMM_B6
KEGG_NOD_LIKE_RECEPTOR_SIGNALING_PATHWAY
Proportion of pathway genes=0.66; Adjusted p−value=0
Average gene set expression
2
1
0
−1
−2
Expression threshold
Below
Above

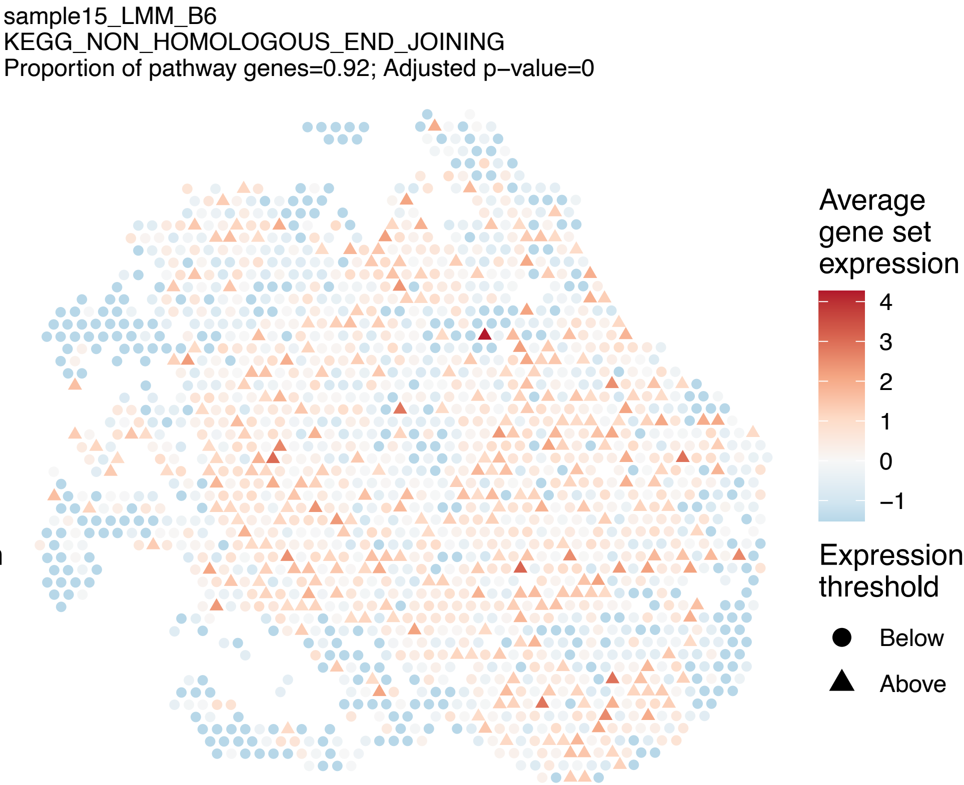
sample15_LMM_B6
KEGG_NON_HOMOLOGOUS_END_JOINING
Proportion of pathway genes=0.92; Adjusted p−value=0
Average gene set expression
4
3
2
1
0
−1
Expression threshold
Below
Above

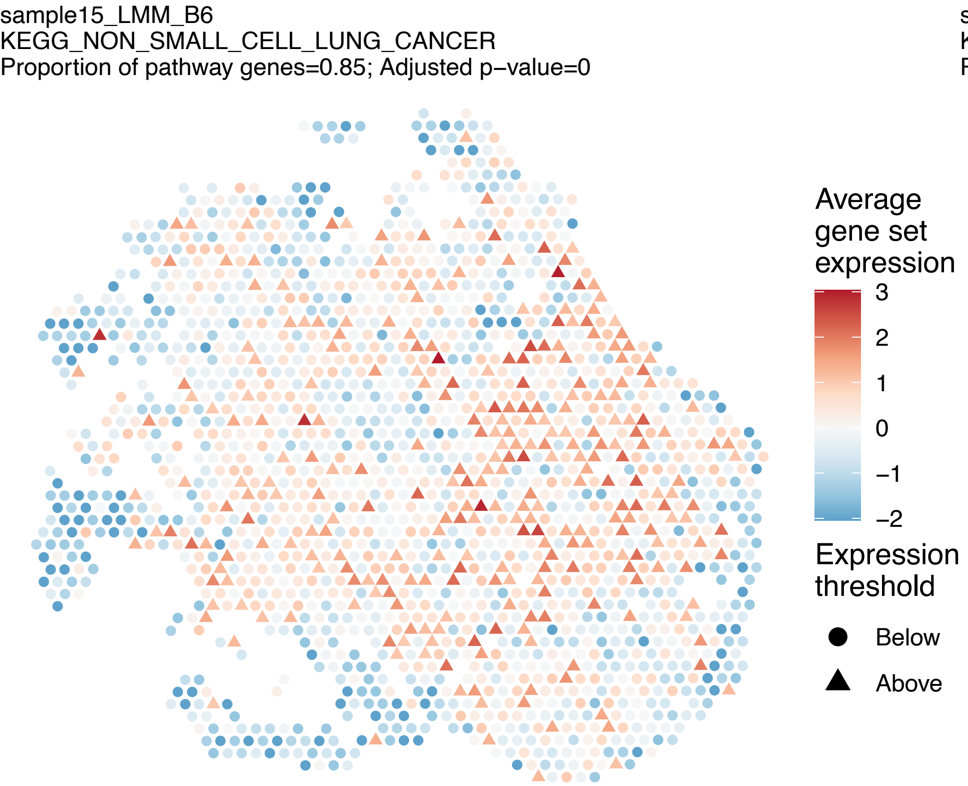
sample15_LMM_B6
KEGG_NON_SMALL_CELL_LUNG_CANCER
Proportion of pathway genes=0.85; Adjusted p–value=0
Average gene set expression
3
2
1
0
−1
−2
Expression threshold
Below
Above

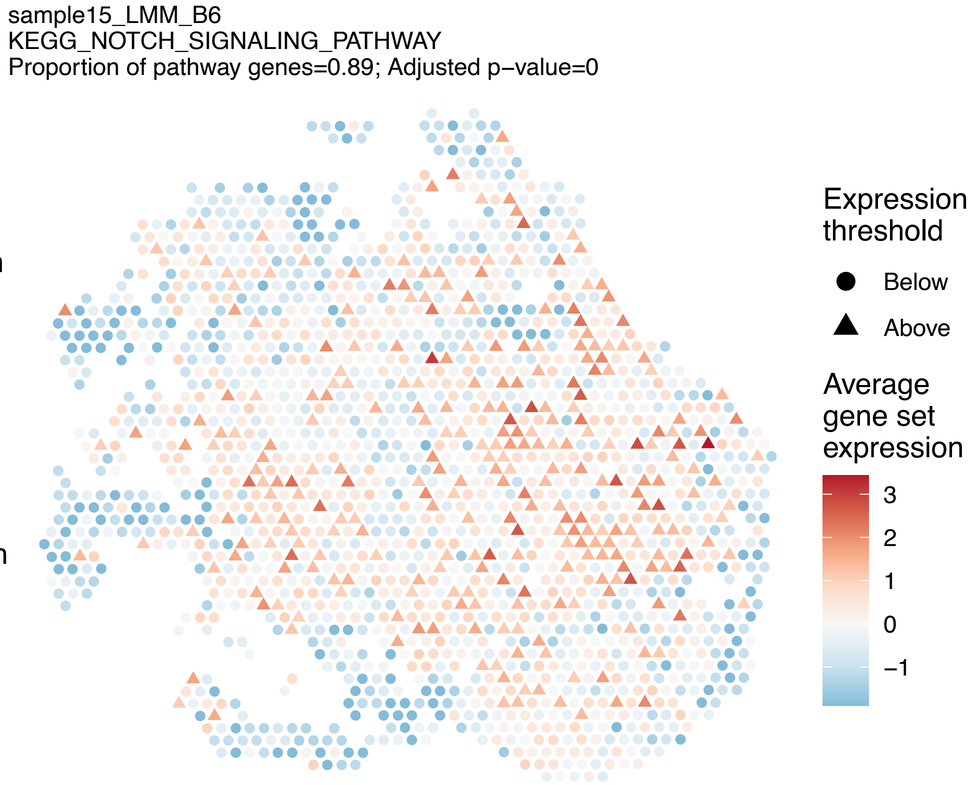
sample15_LMM_B6
KEGG_NOTCH_SIGNALING_PATHWAY
Proportion of pathway genes=0.89; Adjusted p–value=0
Expression threshold
Below
Above
Average gene set expression
3
2
1
0
−1

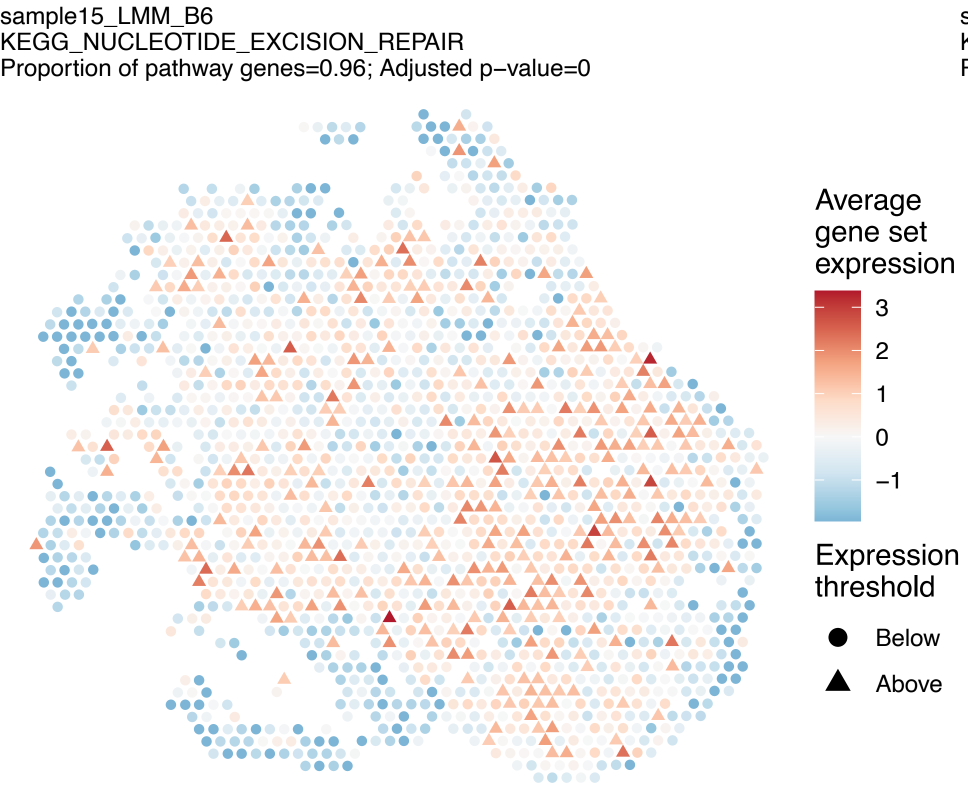
sample15_LMM_B6
KEGG_NUCLEOTIDE_EXCISION_REPAIR
Proportion of pathway genes=0.96; Adjusted p–value=0
Average gene set expression
3
2
1
0
−1
Expression threshold
Below
Above

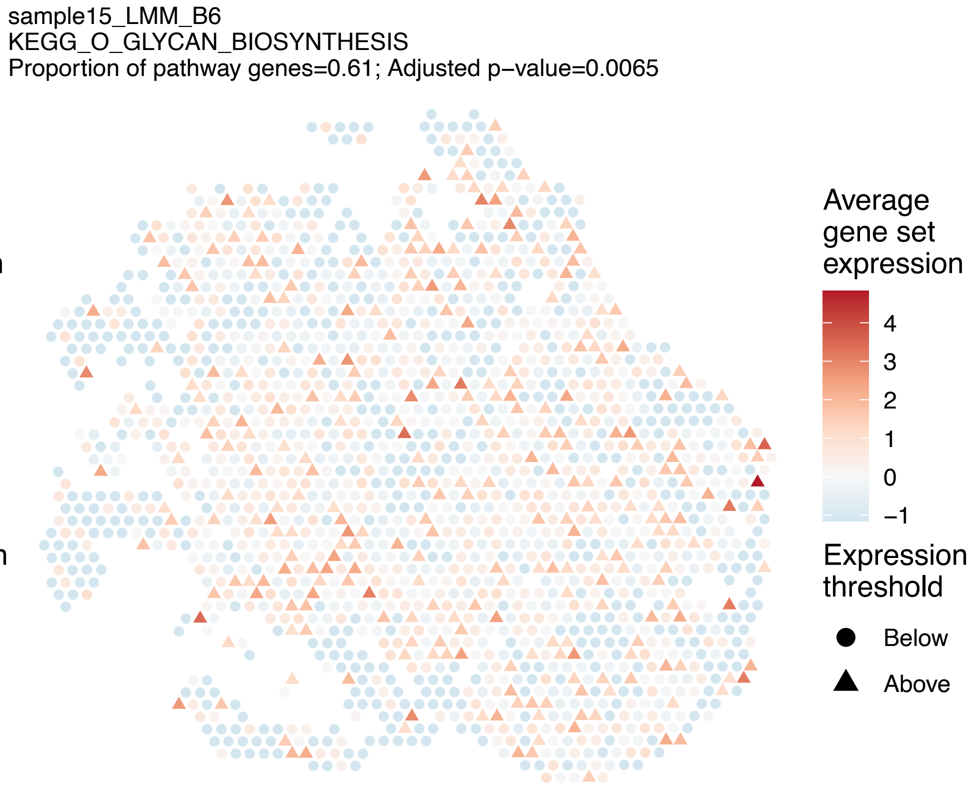
sample15_LMM_B6
KEGG_O_GLYCAN_BIOSYNTHESIS
Proportion of pathway genes=0.61; Adjusted p–value=0.0065
Average gene set expression
4
3
2
1
0
−1
Expression threshold
Below
Above

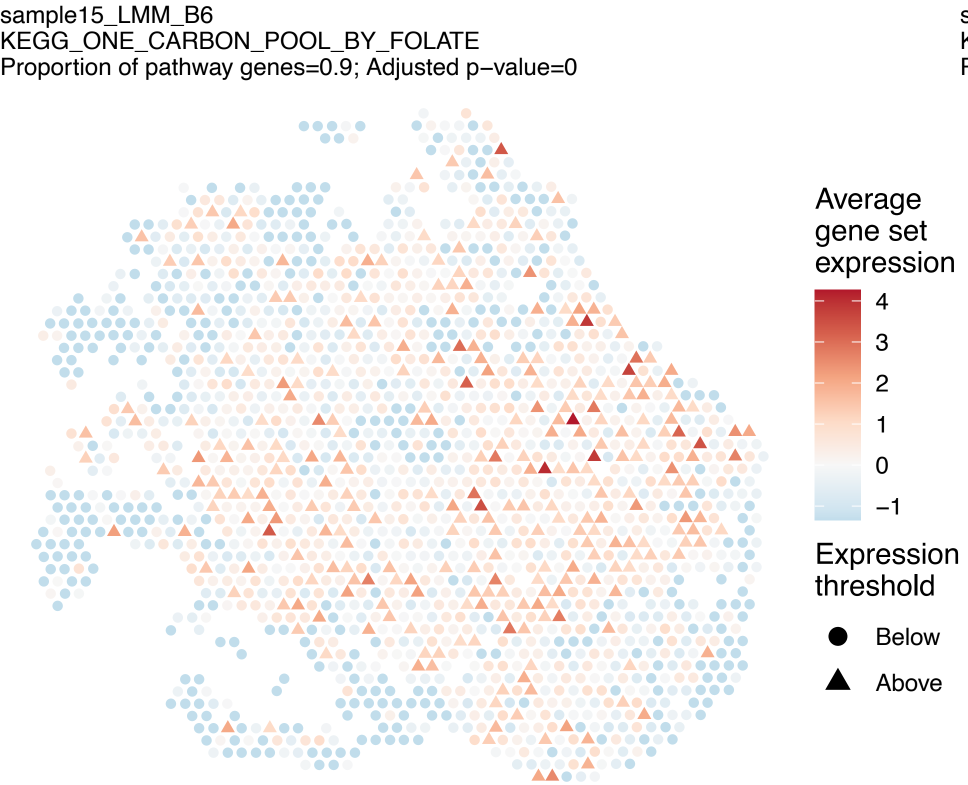
sample15_LMM_B6
KEGG_ONE_CARBON_POOL_BY_FOLATE
Proportion of pathway genes=0.9; Adjusted p−value=0
Average gene set expression
4
3
2
1
0
−1
Expression threshold
Below
Above

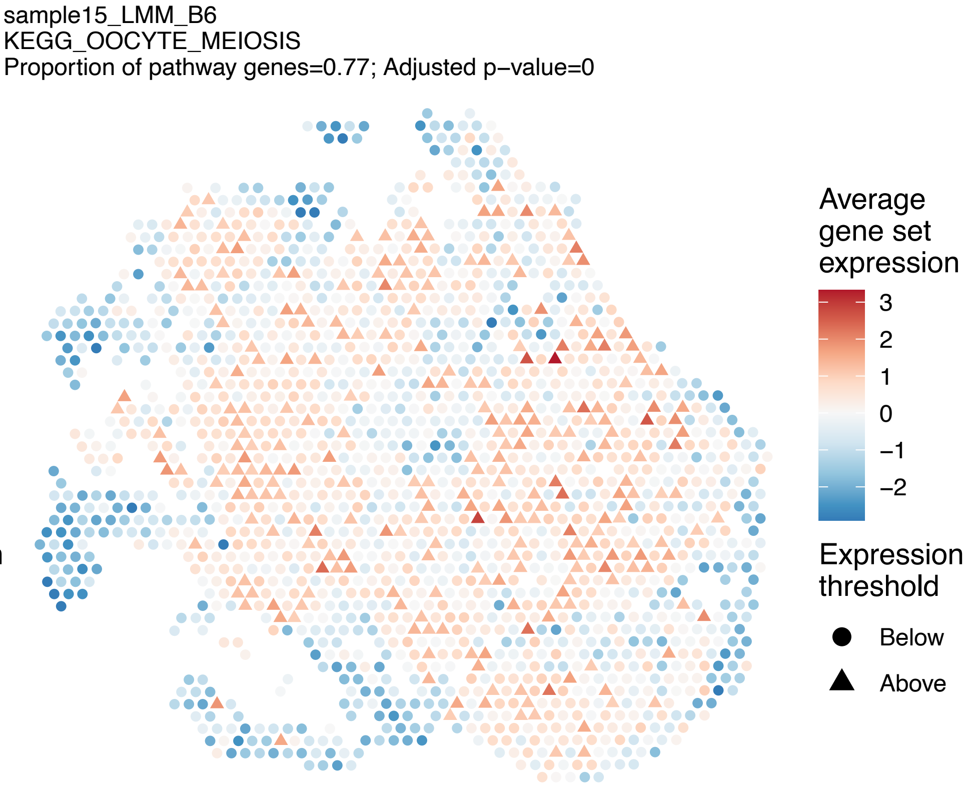
sample15_LMM_B6
KEGG_OOCYTE_MEIOSIS
Proportion of pathway genes=0.77; Adjusted p−value=0
Average gene set expression
3
2
1
0
−1
−2
Expression threshold
Below
Above

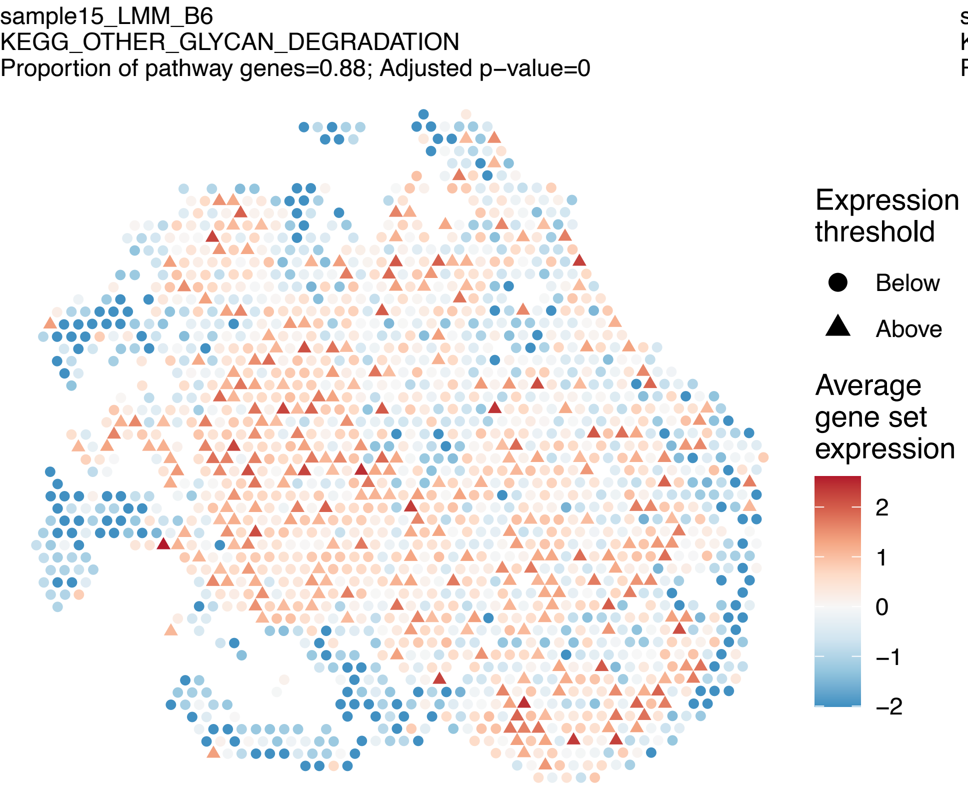
sample15_LMM_B6
KEGG_OTHER_GLYCAN_DEGRADATION
Proportion of pathway genes=0.88; Adjusted p−value=0
Expression threshold
Below
Above
Average gene set expression
2
1
0
−1
−2

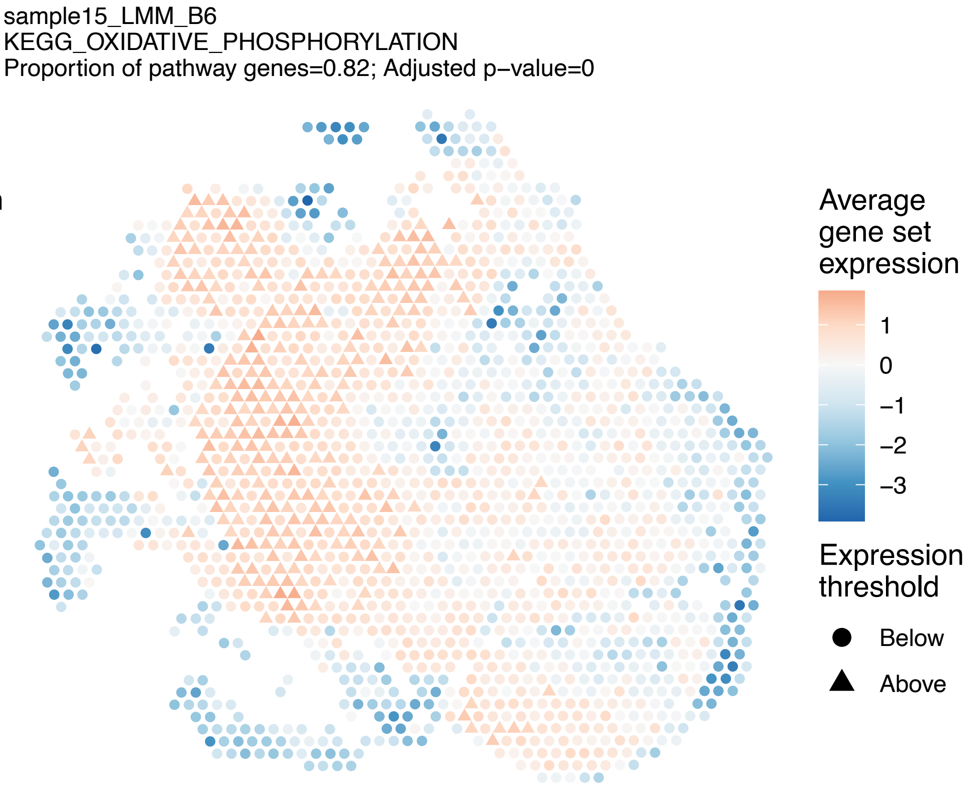
sample15_LMM_B6
KEGG_OXIDATIVE_PHOSPHORYLATION
Proportion of pathway genes=0.82; Adjusted p−value=0
Average gene set expression
1
0
−1
−2
−3
Expression threshold
Below
Above

sample15_LMM_B6
KEGG_P53_SIGNALING_PATHWAY
Proportion of pathway genes=0.89; Adjusted p−value=0

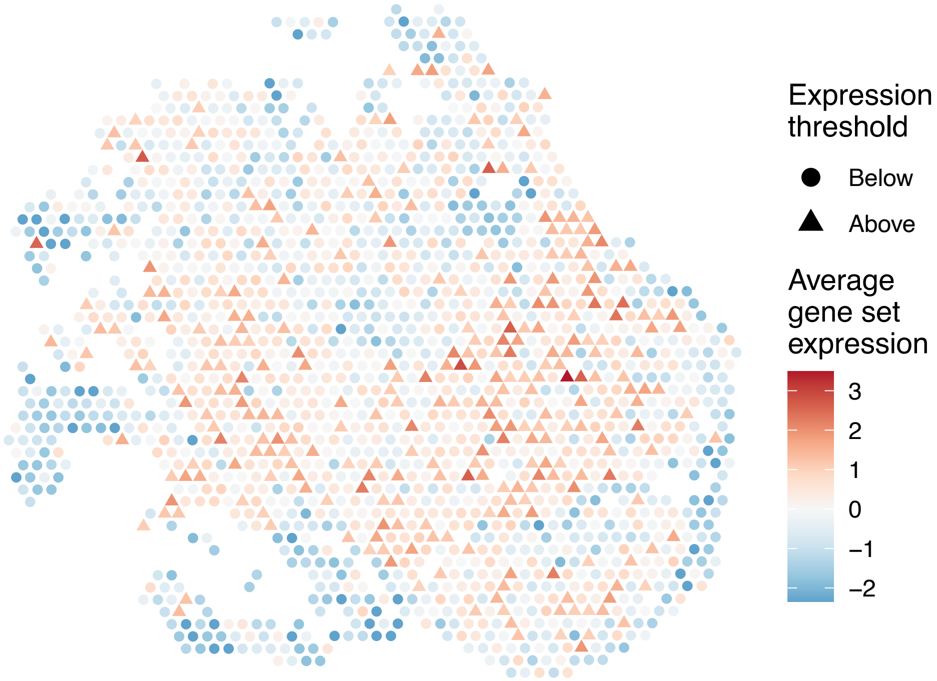


sample15_LMM_B6
KEGG_PANCREATIC_CANCER
Proportion of pathway genes=0.86; Adjusted p−value=0

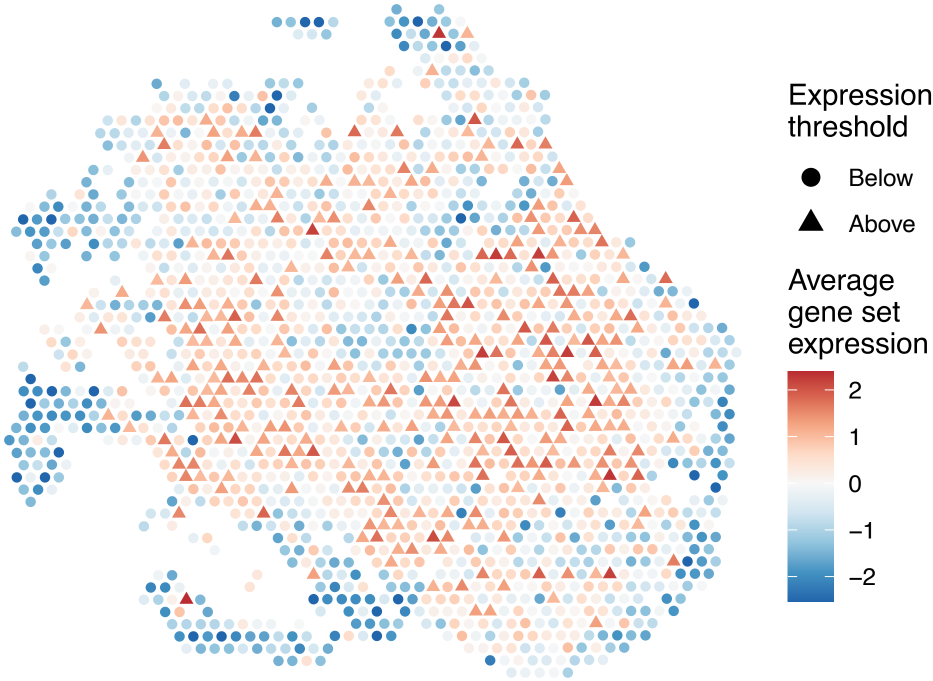


sample15_LMM_B6
KEGG_PANTOTHENATE_AND_COA_BIOSYNTHESIS
Proportion of pathway genes=0.76; Adjusted p−value=0

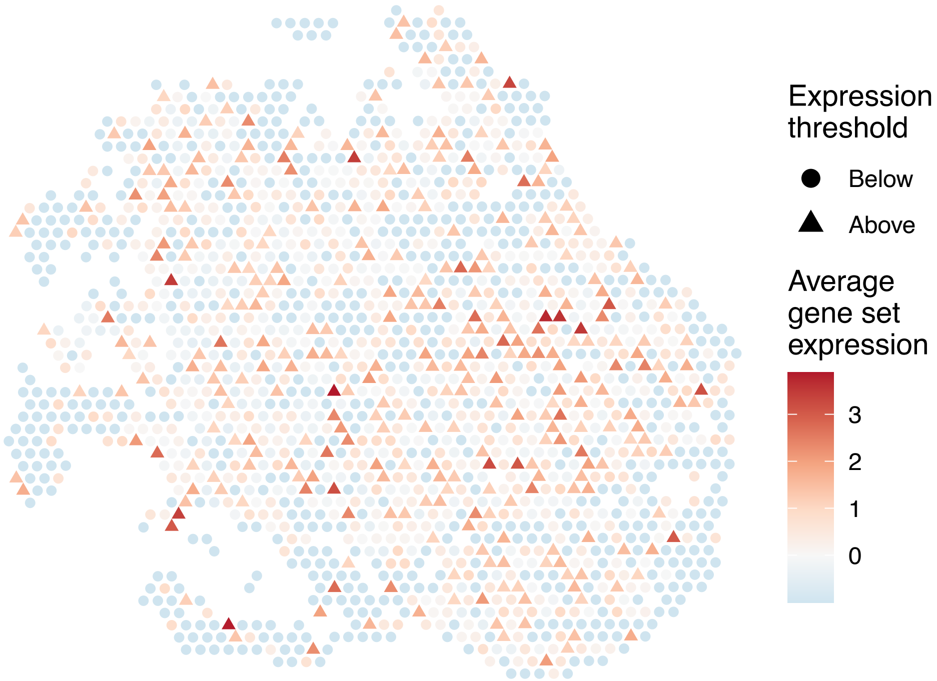


sample15_LMM_B6
KEGG_PARKINSONS_DISEASE
Proportion of pathway genes=0.86; Adjusted p−value=0

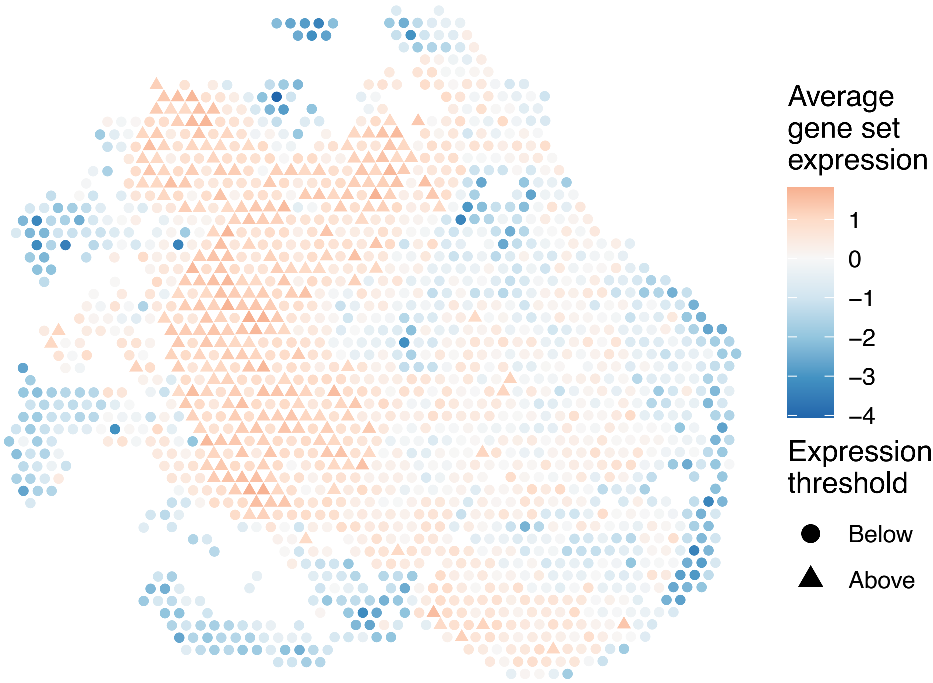

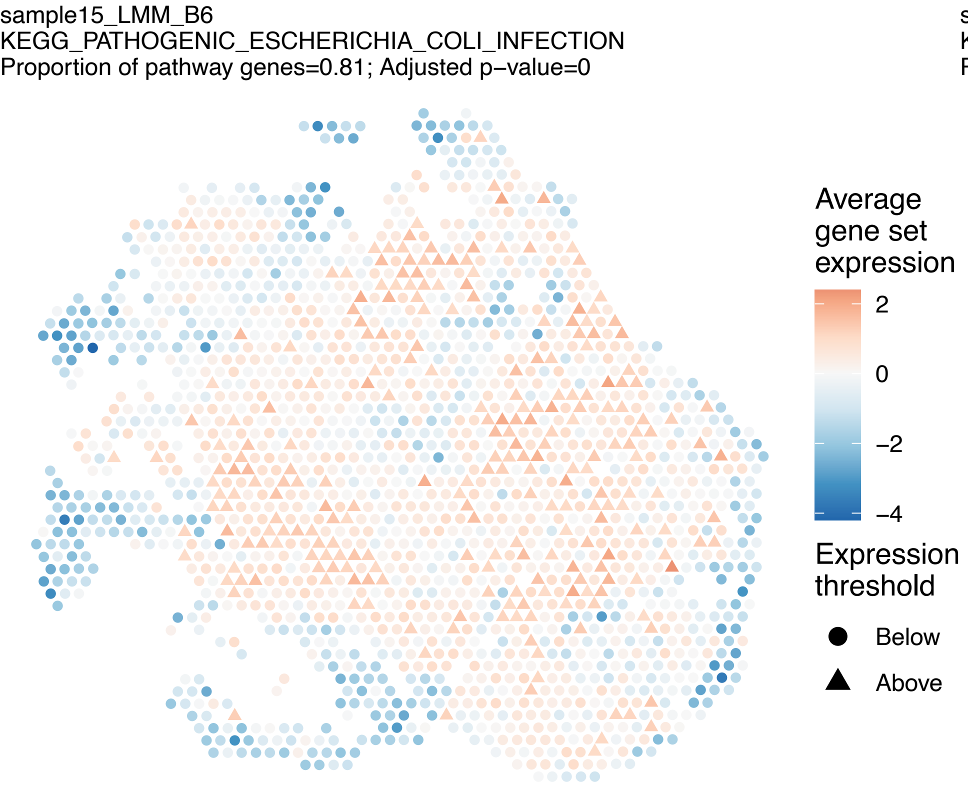
sample15_LMM_B6
KEGG_PATHOGENIC_ESCHERICHIA_COLI_INFECTION
Proportion of pathway genes=0.81; Adjusted p−value=0
Average gene set expression
2
0
−2
−4
Expression threshold
Below
Above

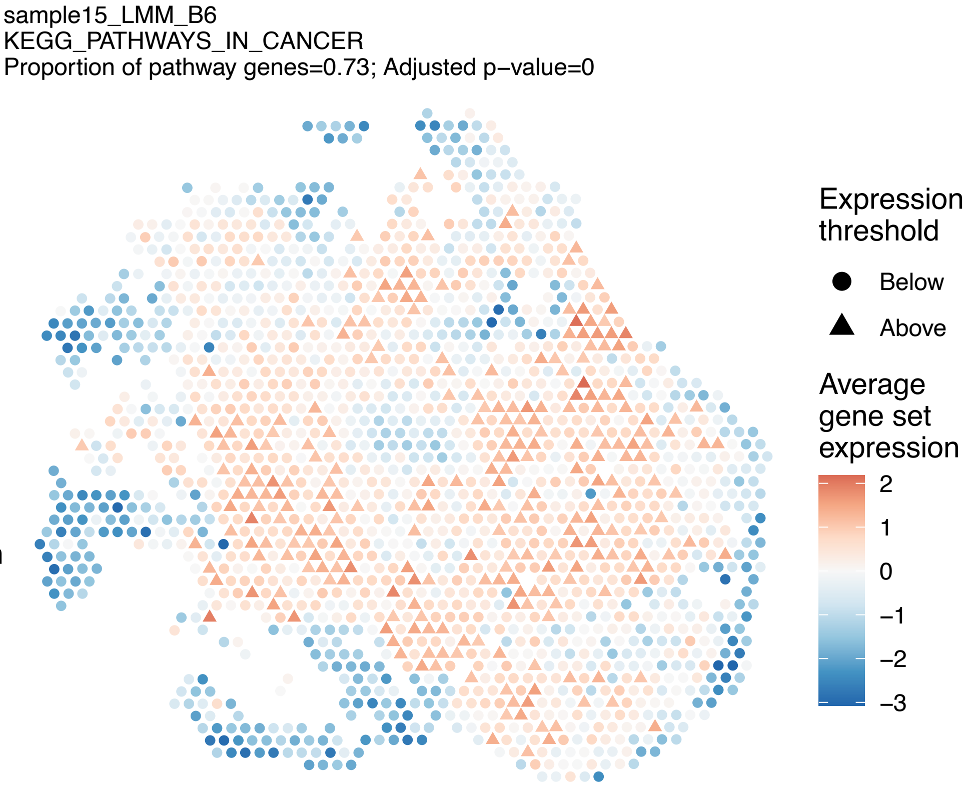
sample15_LMM_B6
KEGG_PATHWAYS_IN_CANCER
Proportion of pathway genes=0.73; Adjusted p−value=0
Expression threshold
Below
Above
Average gene set expression
2
1
0
−1
−2
−3

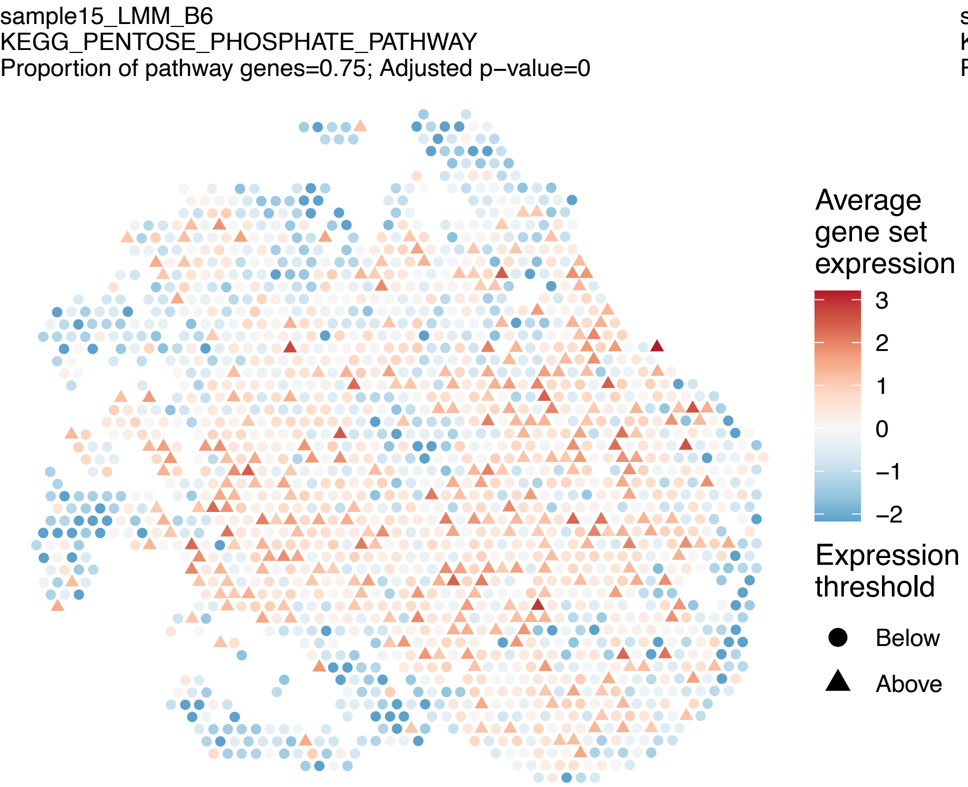
sample15_LMM_B6
KEGG_PENTOSE_PHOSPHATE_PATHWAY
Proportion of pathway genes=0.75; Adjusted p−value=0
Average gene set expression
3
2
1
0
−1
−2
Expression threshold
Below
Above

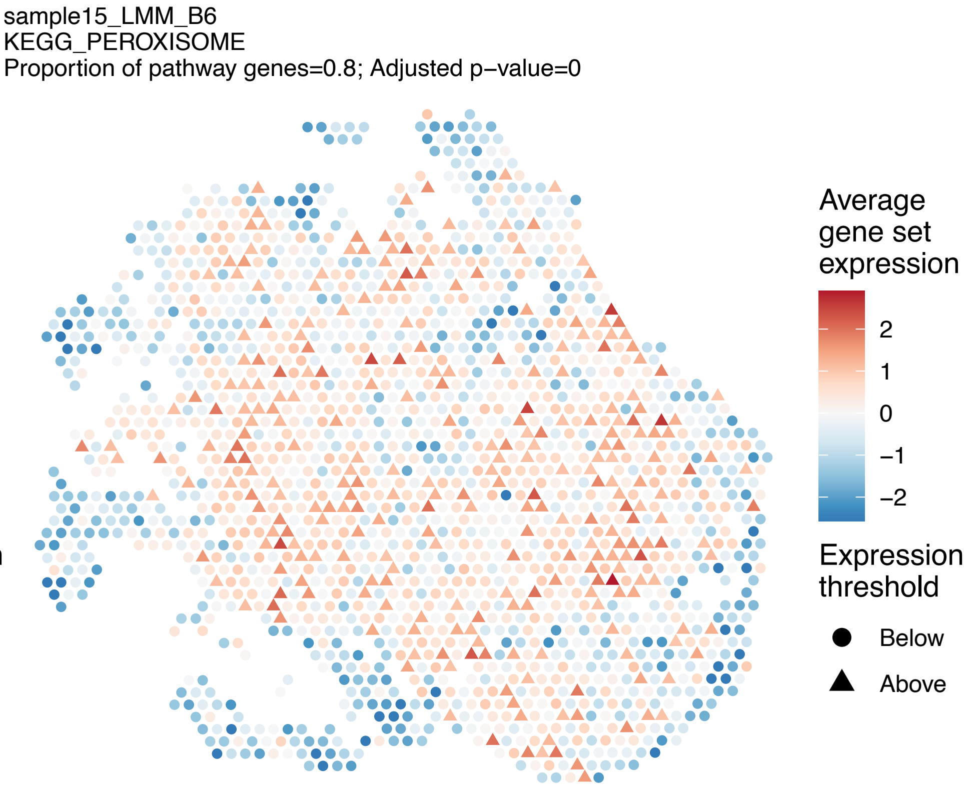
sample15_LMM_B6
KEGG_PEROXISOME
Proportion of pathway genes=0.8; Adjusted p−value=0
Average gene set expression
2
1
0
−1
−2
Expression threshold
Below
Above

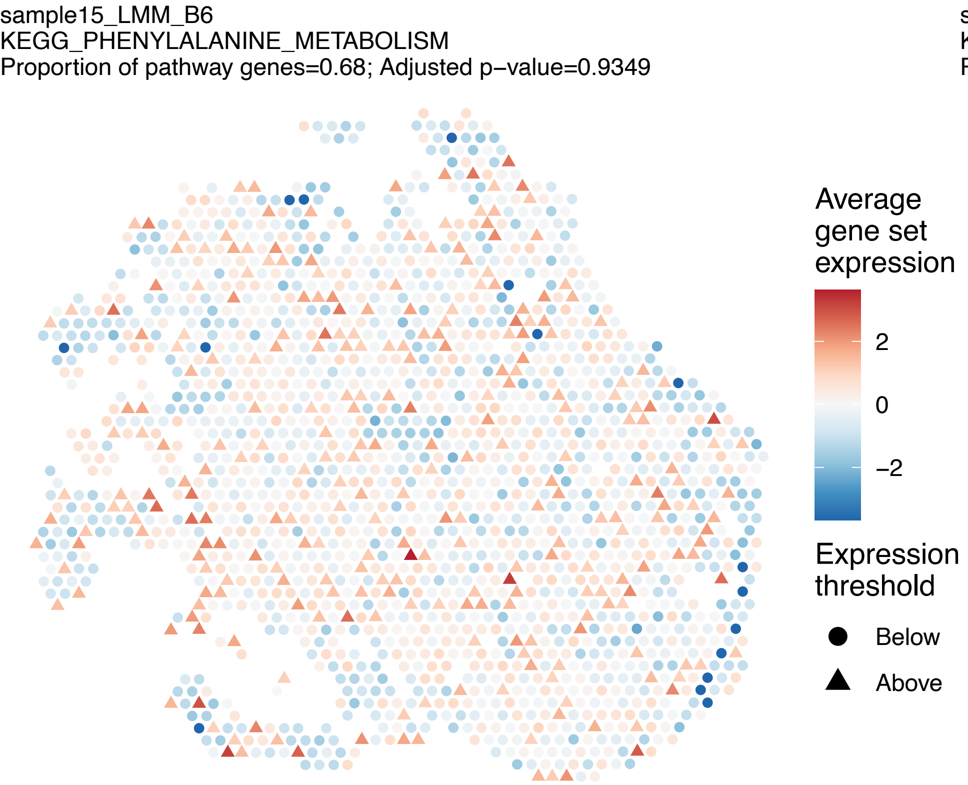

sample15_LMM_B6
KEGG_PHENYLALANINE_METABOLISM
Proportion of pathway genes=0.68; Adjusted p−value=0.9349
Average gene set expression
2
0
−2
Expression threshold
Below
Above


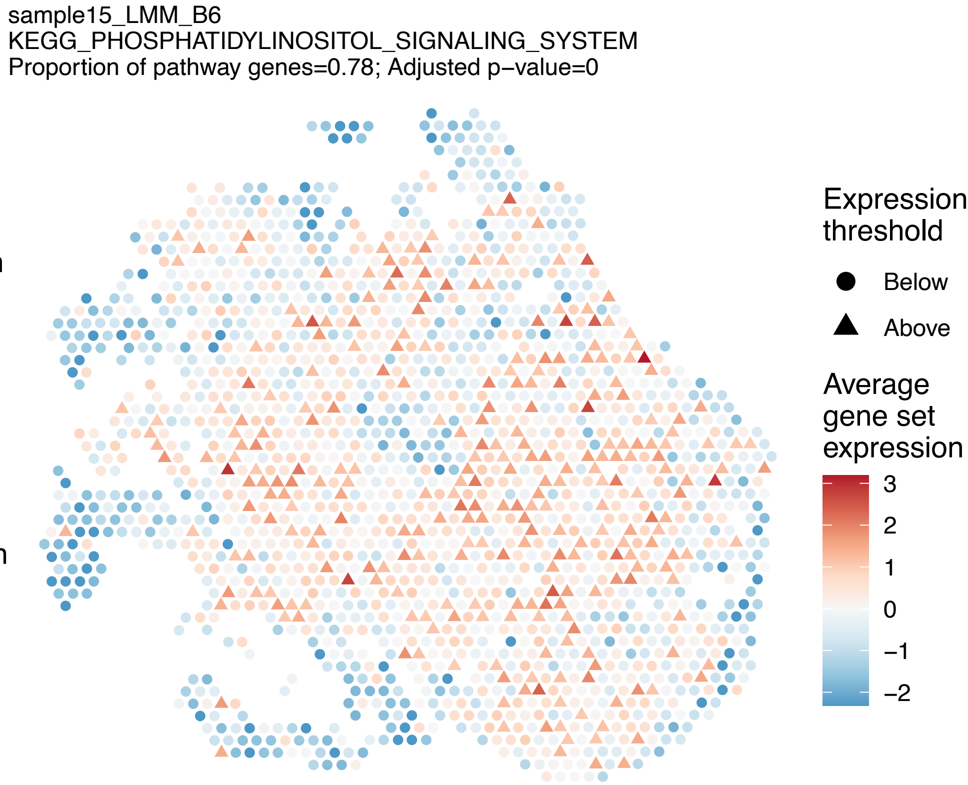

sample15_LMM_B6
KEGG_PHOSPHATIDYLINOSITOL_SIGNALING_SYSTEM
Proportion of pathway genes=0.78; Adjusted p−value=0
Expression threshold
Below
Above
Average gene set expression
3
2
1
0
−1
−2


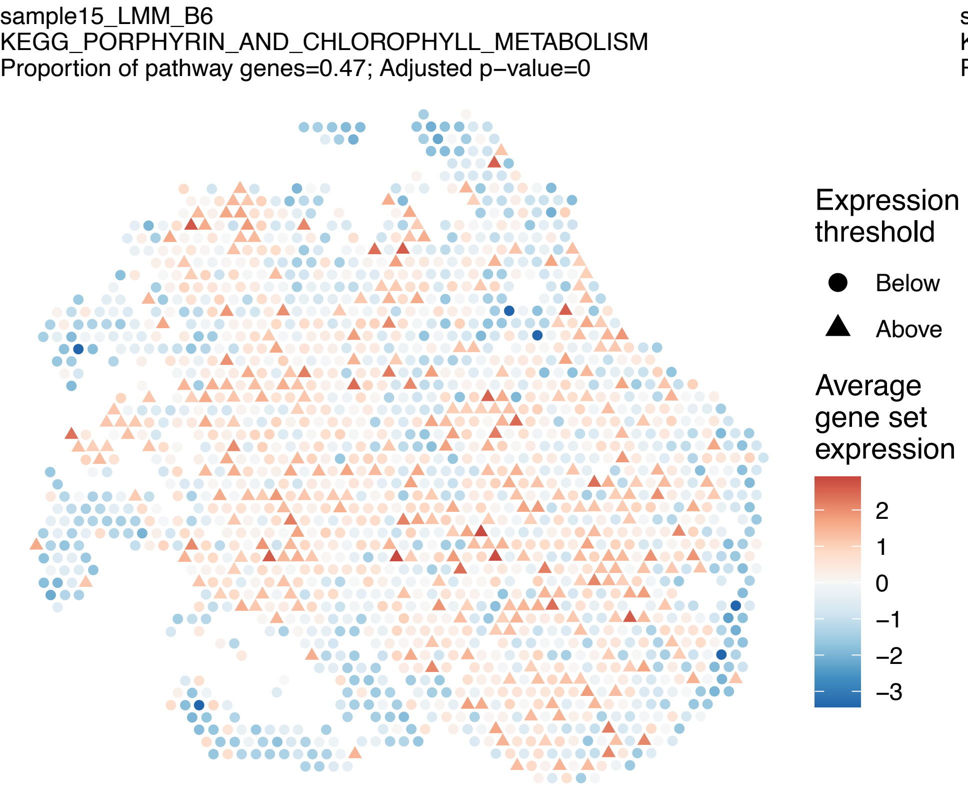

sample15_LMM_B6
KEGG_PORPHYRIN_AND_CHLOROPHYLL_METABOLISM
Proportion of pathway genes=0.47; Adjusted p−value=0
Expression threshold
Below
Above
Average gene set expression
2
1
0
−1
−2
−3


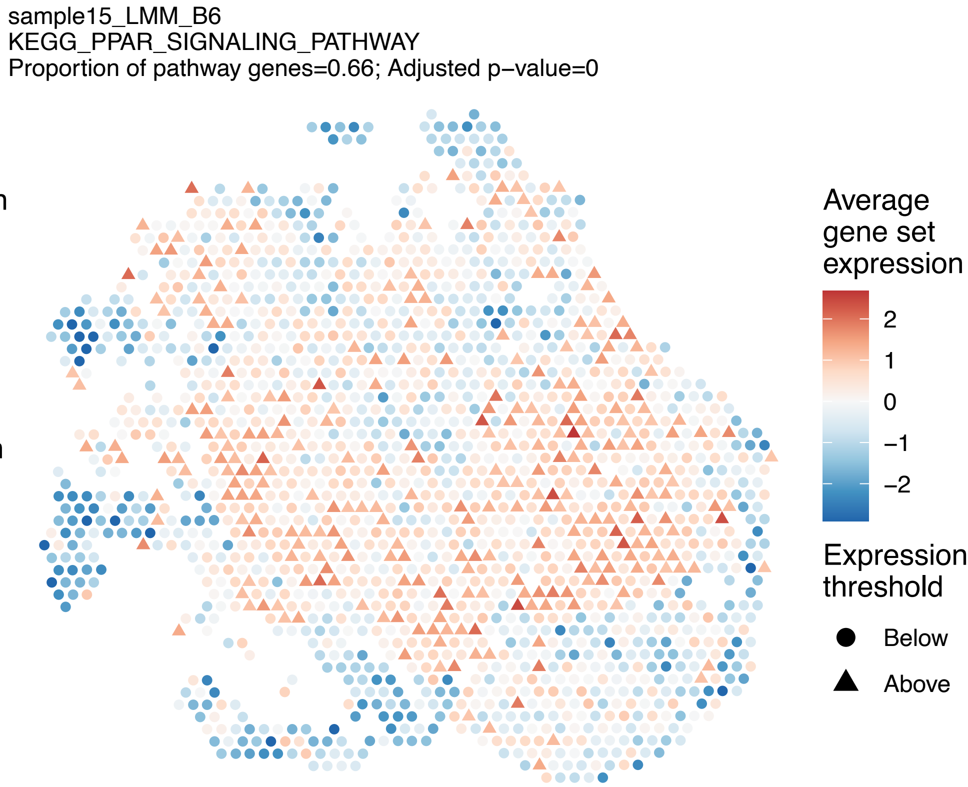

sample15_LMM_B6
KEGG_PPAR_SIGNALING_PATHWAY
Proportion of pathway genes=0.66; Adjusted p−value=0
Average gene set expression
2
1
0
−1
−2
Expression threshold
Below
Above

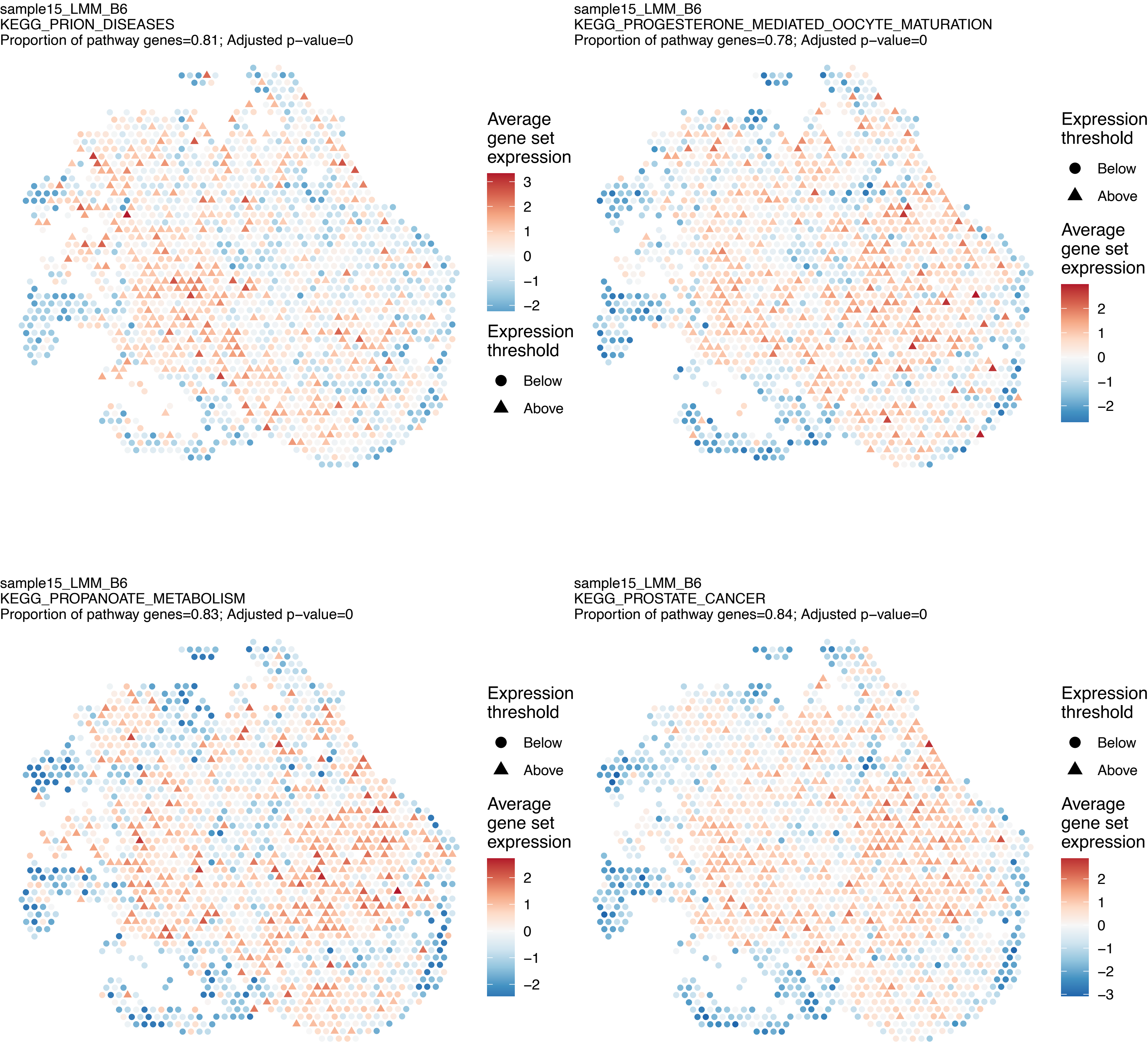
sample15_LMM_B6
KEGG_PRION_DISEASES
Proportion of pathway genes=0.81; Adjusted p−value=0
Average gene set expression
3
2
1
0
−1
−2
Expression threshold
Below
Above
sample15_LMM_B6
KEGG_PROGESTERONE_MEDIATED_OOCYTE_MATURATION
Proportion of pathway genes=0.78; Adjusted p−value=0
Expression threshold
Below
Above
Average gene set expression
2
1
0
−1
−2
sample15_LMM_B6
KEGG_PROPANOATE_METABOLISM
Proportion of pathway genes=0.83; Adjusted p−value=0
Expression threshold
Below
Above
Average gene set expression
2
1
0
−1
−2
sample15_LMM_B6
KEGG_PROSTATE_CANCER
Proportion of pathway genes=0.84; Adjusted p−value=0
Expression threshold
Below
Above
Average gene set expression
2
1
0
−1
−2
−3

sample15_LMM_B6
KEGG_PROTEASOME
Proportion of pathway genes=0.92; Adjusted p−value=0

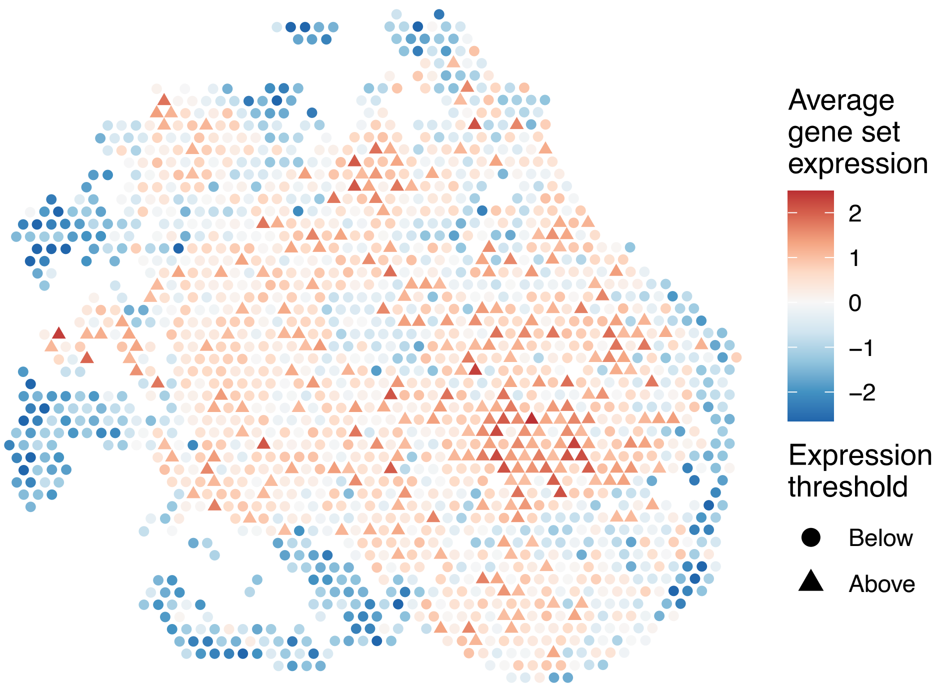


sample15_LMM_B6
KEGG_PROTEIN_EXPORT
Proportion of pathway genes=0.96; Adjusted p−value=0

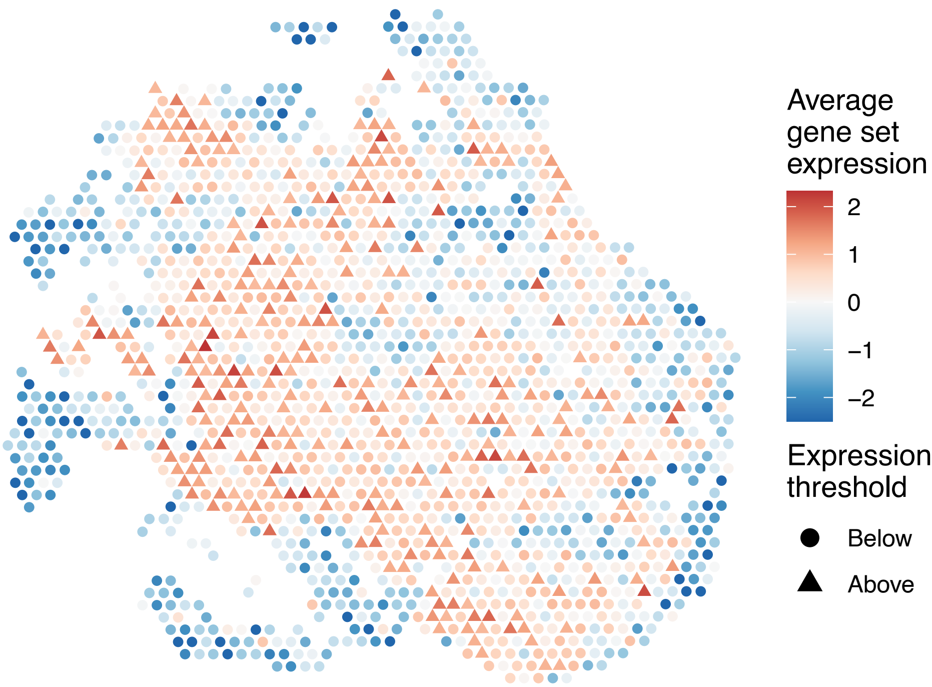


sample15_LMM_B6
KEGG_PROXIMAL_TUBULE_BICARBONATE_RECLAMATION
Proportion of pathway genes=0.54; Adjusted p−value=0

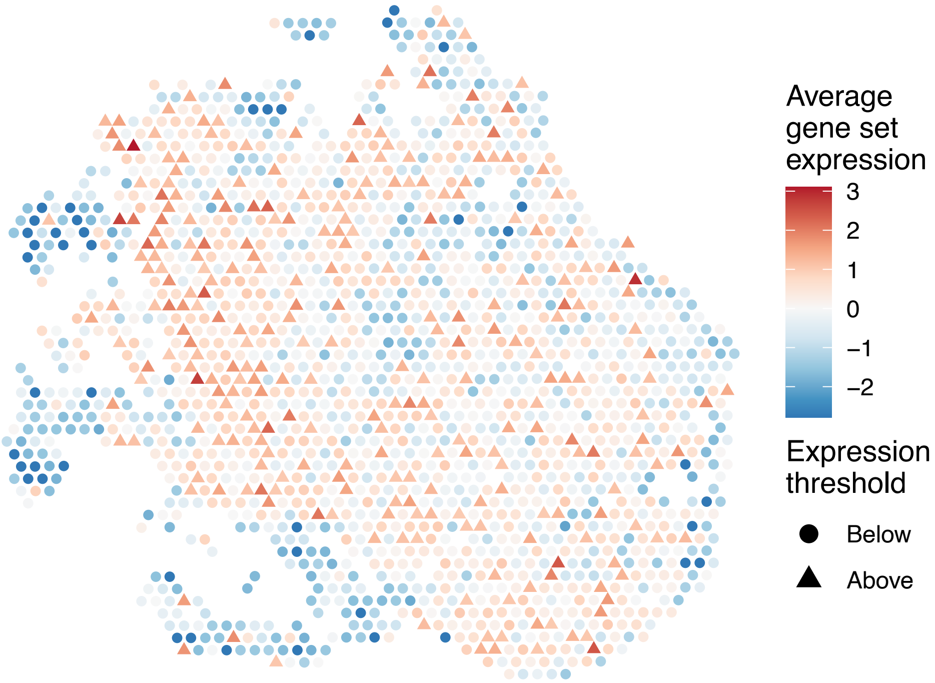


sample15_LMM_B6
KEGG_PURINE_METABOLISM
Proportion of pathway genes=0.71; Adjusted p−value=0

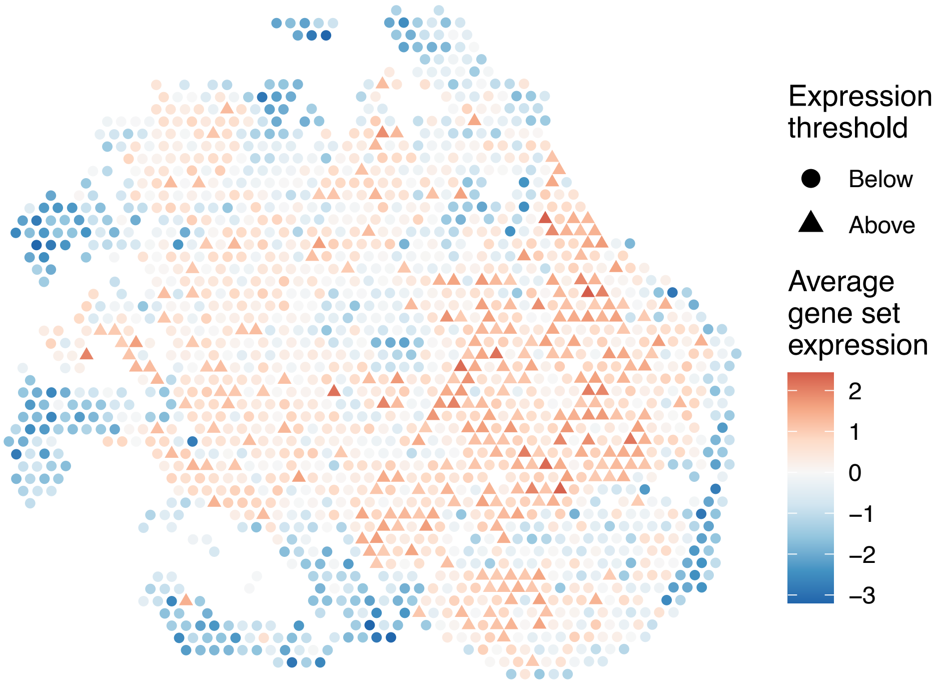

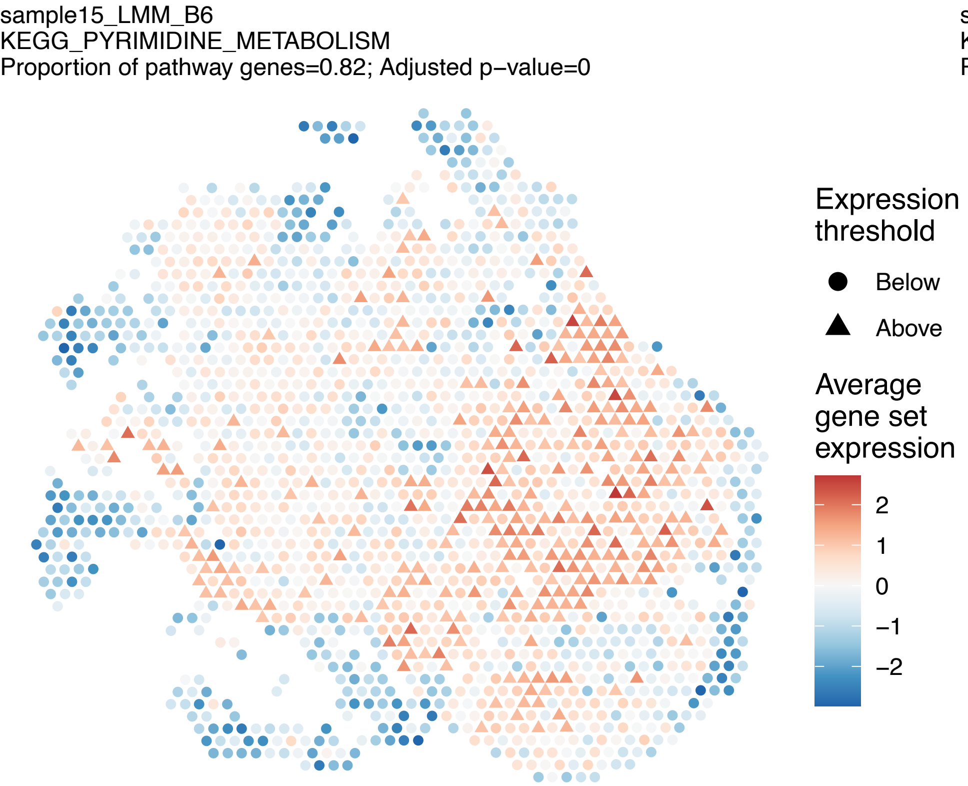
sample15_LMM_B6
KEGG_PYRIMIDINE_METABOLISM
Proportion of pathway genes=0.82; Adjusted p−value=0
Expression threshold
Below
Above
Average gene set expression
2
1
0
−1
−2

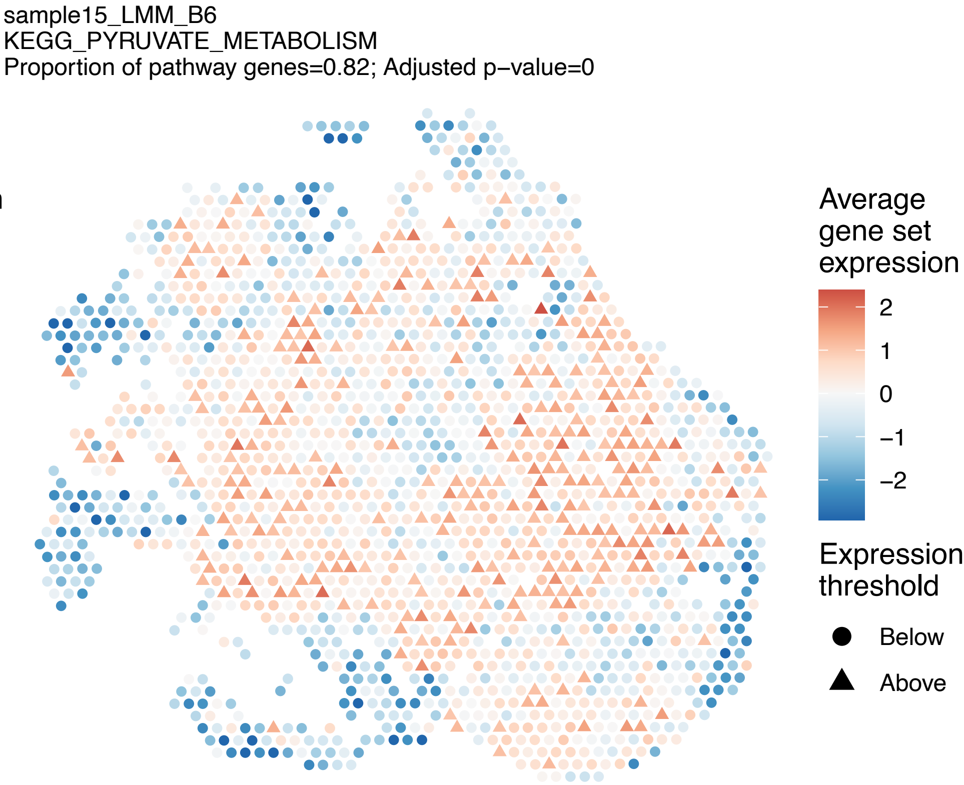
sample15_LMM_B6
KEGG_PYRUVATE_METABOLISM
Proportion of pathway genes=0.82; Adjusted p−value=0
Average gene set expression
2
1
0
−1
−2
Expression threshold
Below
Above

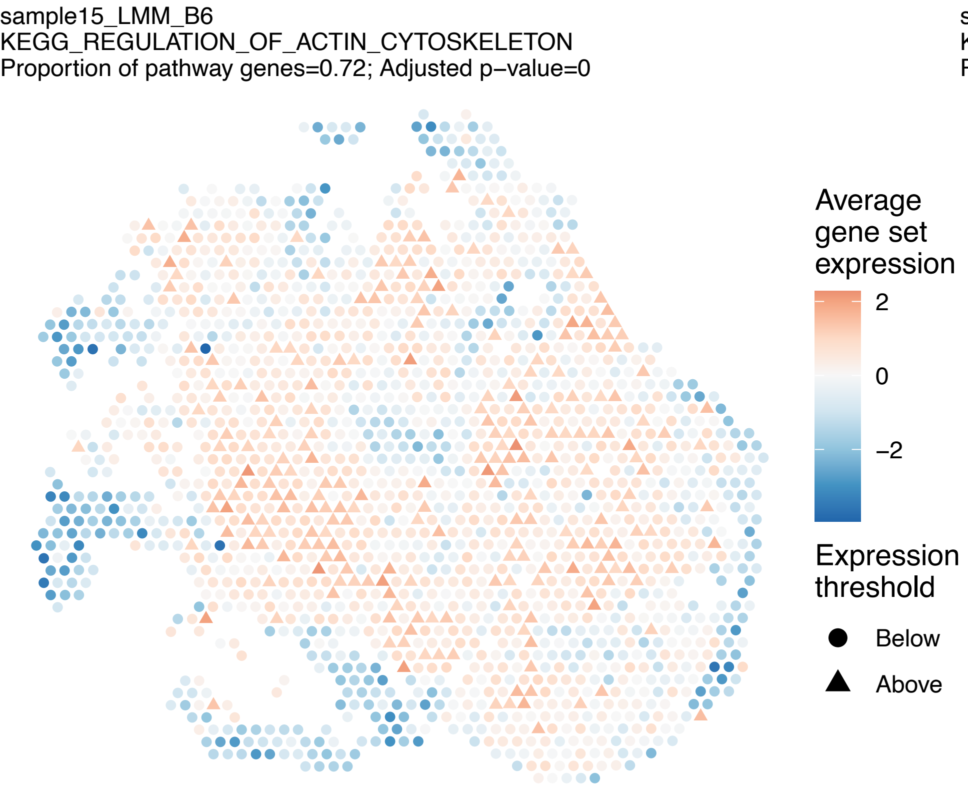
sample15_LMM_B6
KEGG_REGULATION_OF_ACTIN_CYTOSKELETON
Proportion of pathway genes=0.72; Adjusted p−value=0
Average gene set expression
2
0
−2
Expression threshold
Below
Above

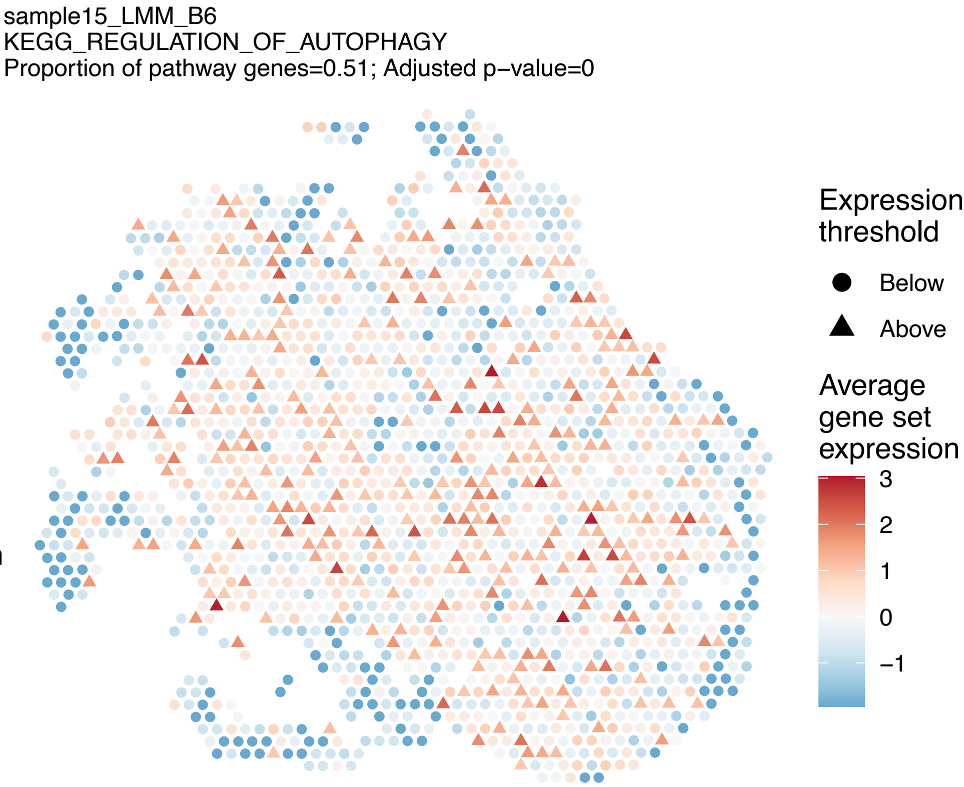
sample15_LMM_B6
KEGG_REGULATION_OF_AUTOPHAGY
Proportion of pathway genes=0.51; Adjusted p−value=0
Expression threshold
Below
Above
Average gene set expression
3
2
1
0
−1

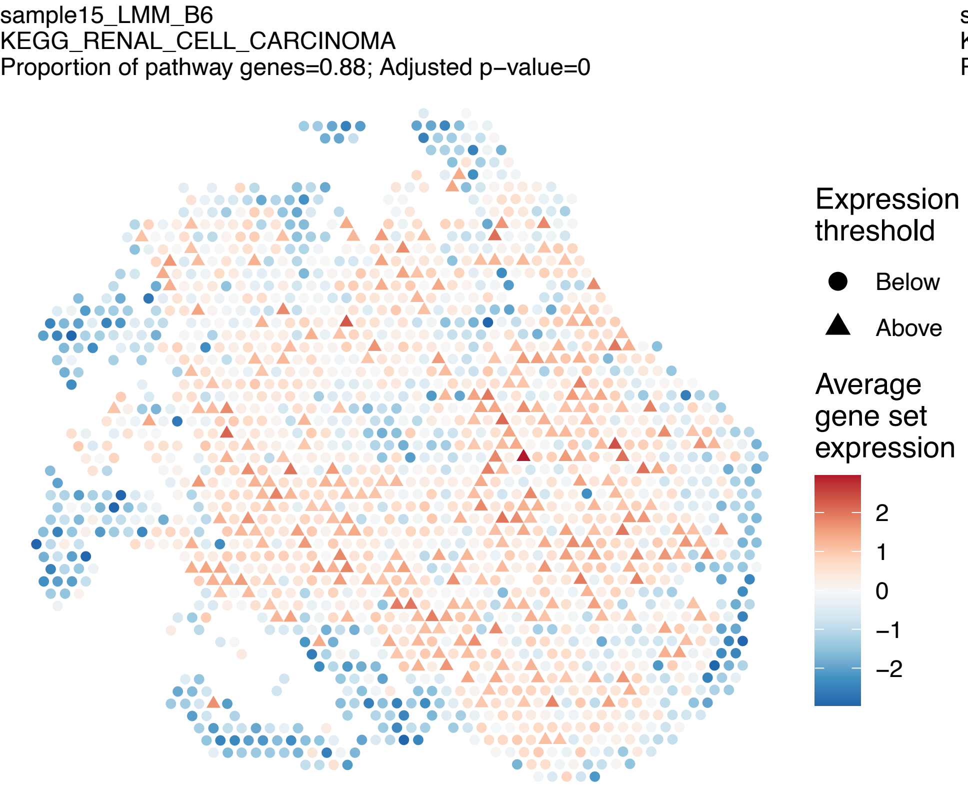
sample15_LMM_B6
KEGG_RENAL_CELL_CARCINOMA
Proportion of pathway genes=0.88; Adjusted p−value=0
Expression threshold
Below
Above
Average gene set expression
2
1
0
−1
−2

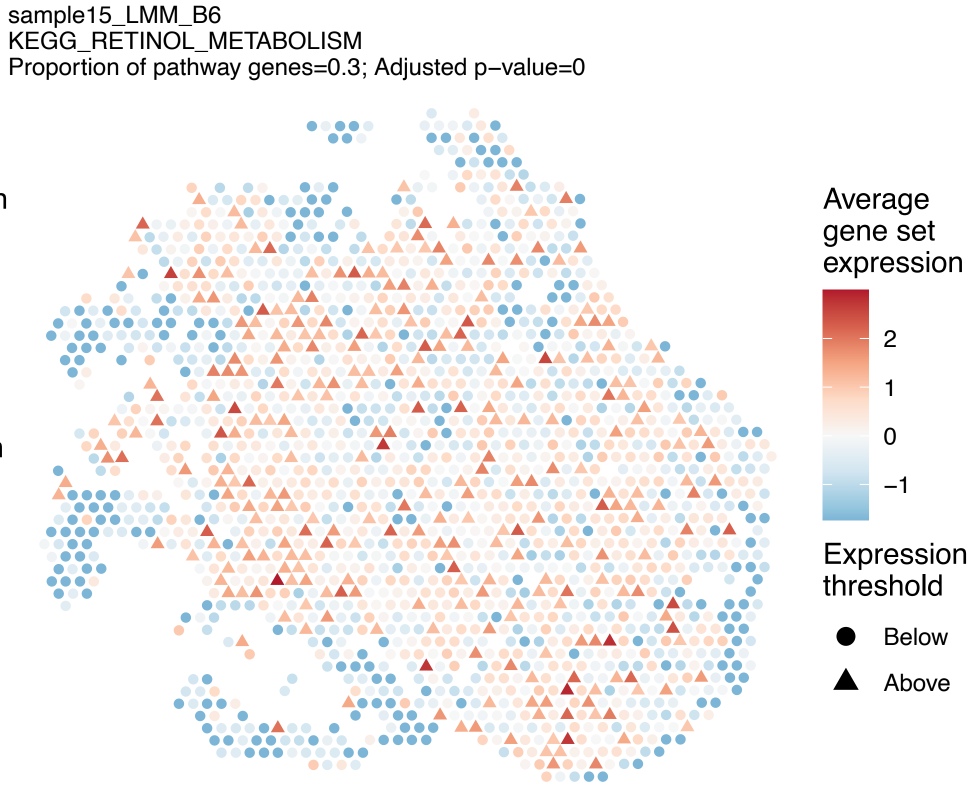
sample15_LMM_B6
KEGG_RETINOL_METABOLISM
Proportion of pathway genes=0.3; Adjusted p−value=0
Average gene set expression
2
1
0
−1
Expression threshold
Below
Above

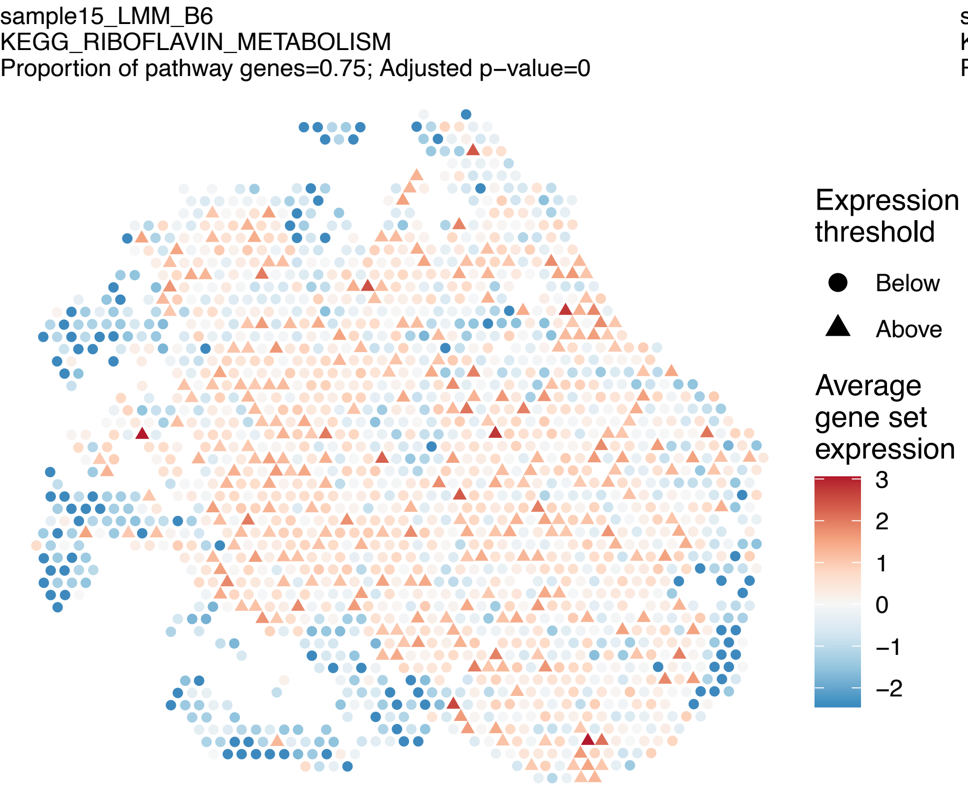
sample15_LMM_B6
KEGG_RIBOFLAVIN_METABOLISM
Proportion of pathway genes=0.75; Adjusted p−value=0
Expression threshold
Below
Above
Average gene set expression
3
2
1
0
−1
−2

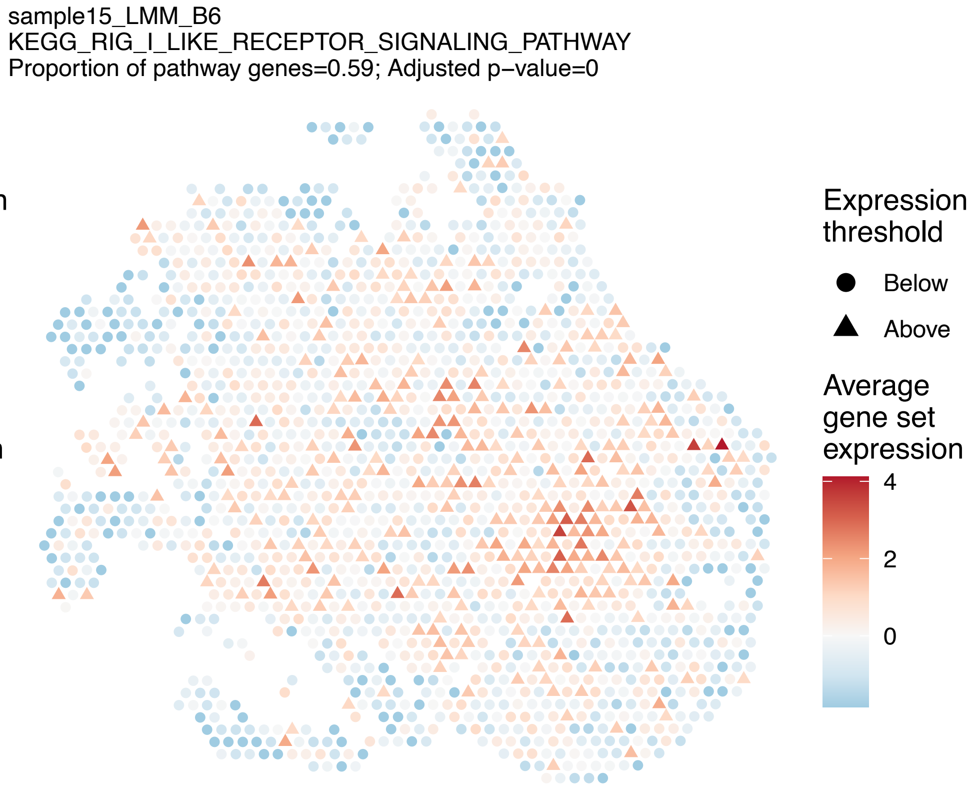
sample15_LMM_B6
KEGG_RIG_I_LIKE_RECEPTOR_SIGNALING_PATHWAY
Proportion of pathway genes=0.59; Adjusted p−value=0
Expression threshold
Below
Above
Average gene set expression
4
2
0

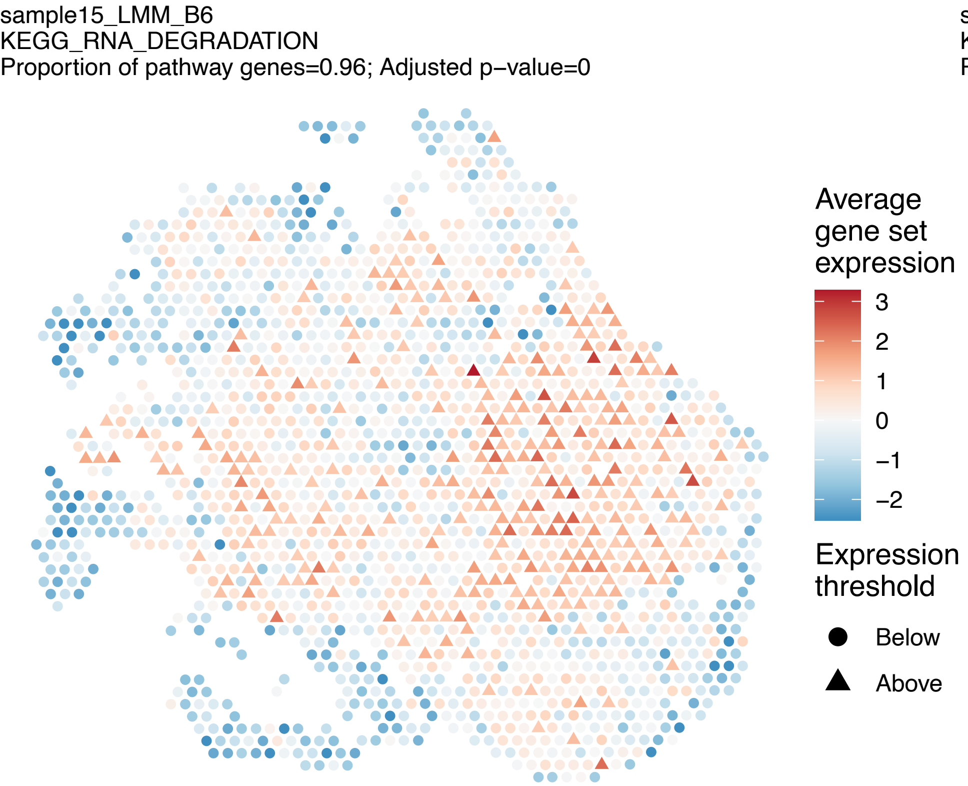
sample15_LMM_B6
KEGG_RNA_DEGRADATION
Proportion of pathway genes=0.96; Adjusted p−value=0
Average gene set expression
3
2
1
0
−1
−2
Expression threshold
Below
Above

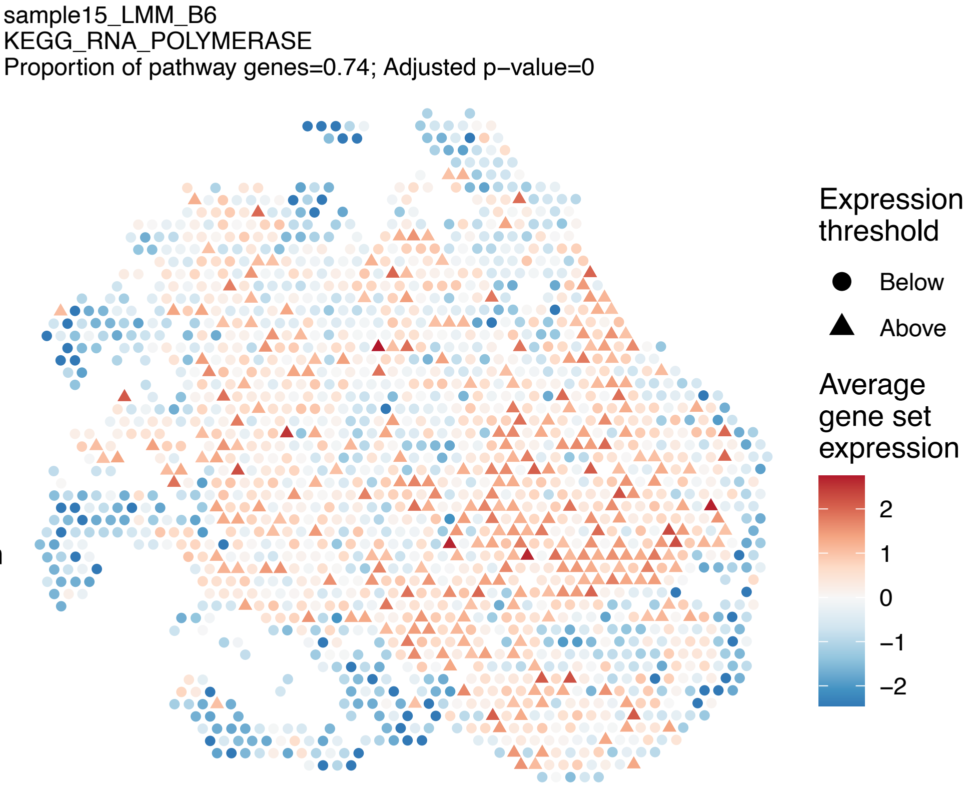
sample15_LMM_B6
KEGG_RNA_POLYMERASE
Proportion of pathway genes=0.74; Adjusted p−value=0
Expression threshold
Below
Above
Average gene set expression
2
1
0
−1
−2

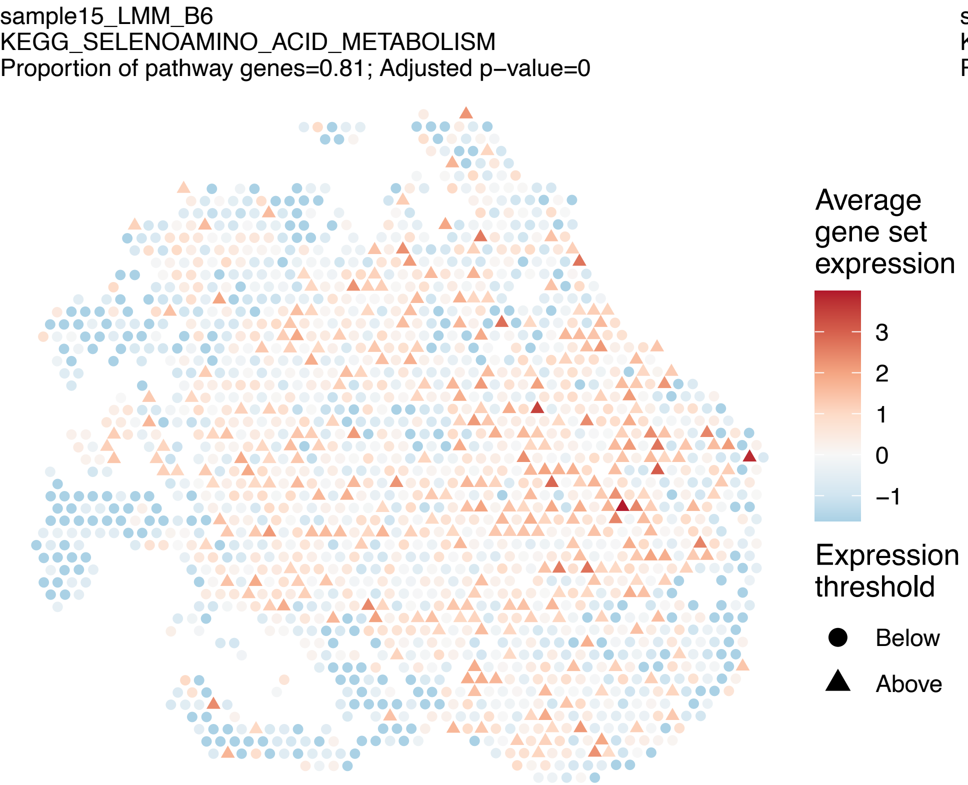
sample15_LMM_B6
KEGG_SELENOAMINO_ACID_METABOLISM
Proportion of pathway genes=0.81; Adjusted p−value=0
Average gene set expression
3
2
1
0
−1
Expression threshold
Below
Above

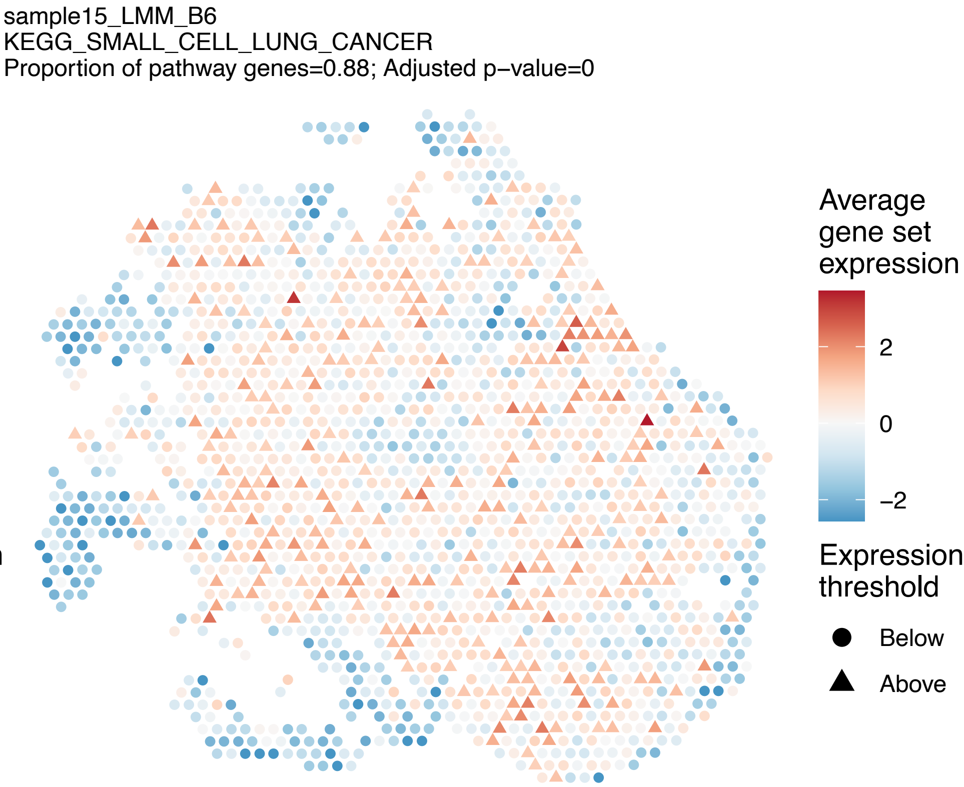
sample15_LMM_B6
KEGG_SMALL_CELL_LUNG_CANCER
Proportion of pathway genes=0.88; Adjusted p−value=0
Average gene set expression
2
0
−2
Expression threshold
Below
Above

sample15_LMM_B6
KEGG_SNARE_INTERACTIONS_IN_VESICULAR_TRANSPORT
Proportion of pathway genes=0.89; Adjusted p−value=0

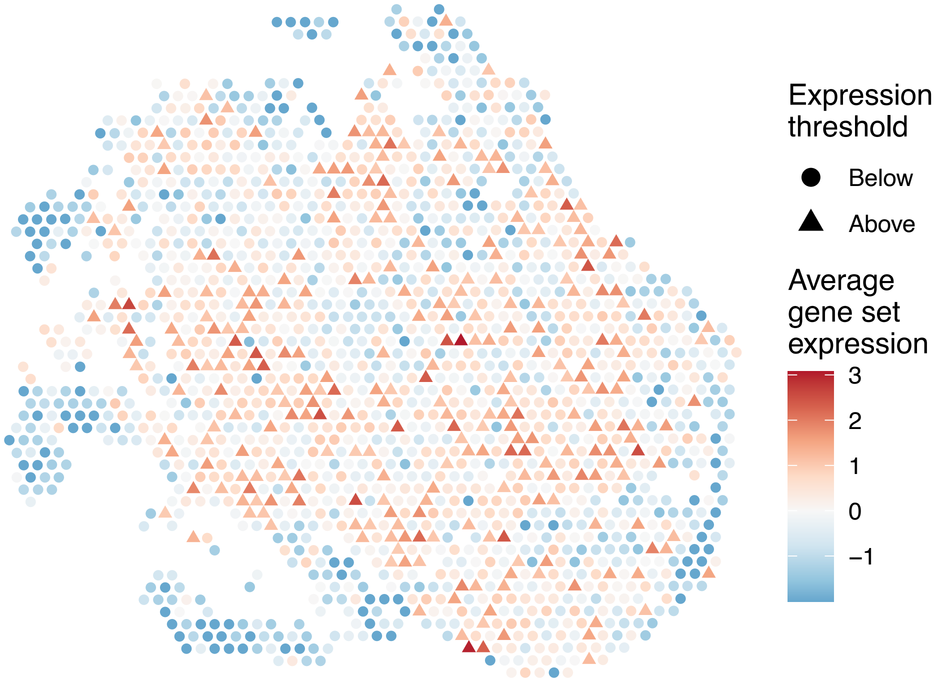


sample15_LMM_B6
KEGG_SPHINGOLIPID_METABOLISM
Proportion of pathway genes=0.76; Adjusted p−value=0

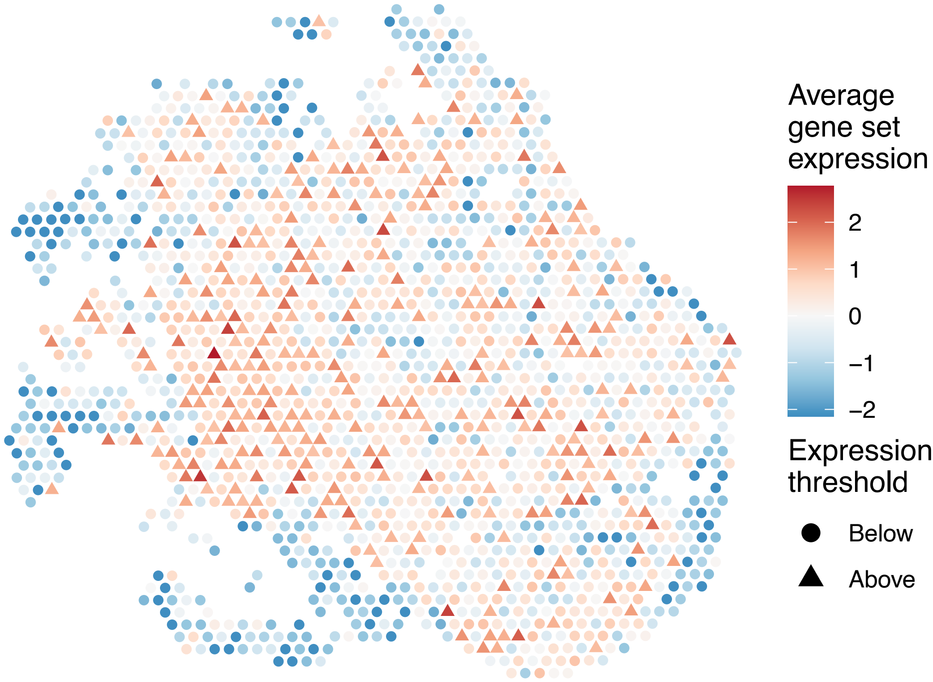


sample15_LMM_B6
KEGG_SPLICEOSOME
Proportion of pathway genes=0.96; Adjusted p−value=0

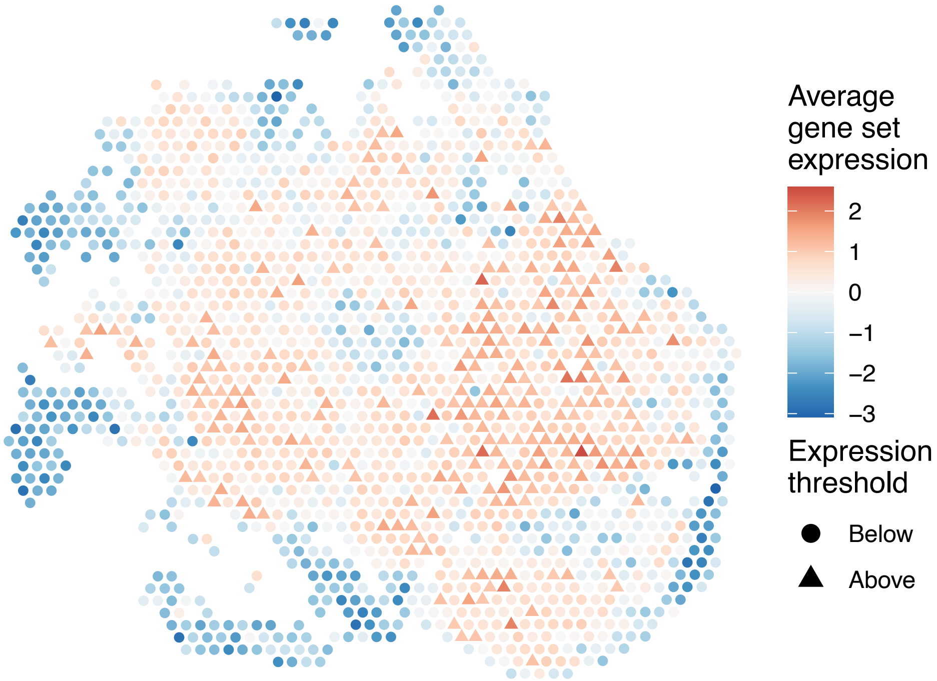


sample15_LMM_B6
KEGG_STARCH_AND_SUCROSE_METABOLISM
Proportion of pathway genes=0.34; Adjusted p−value=0

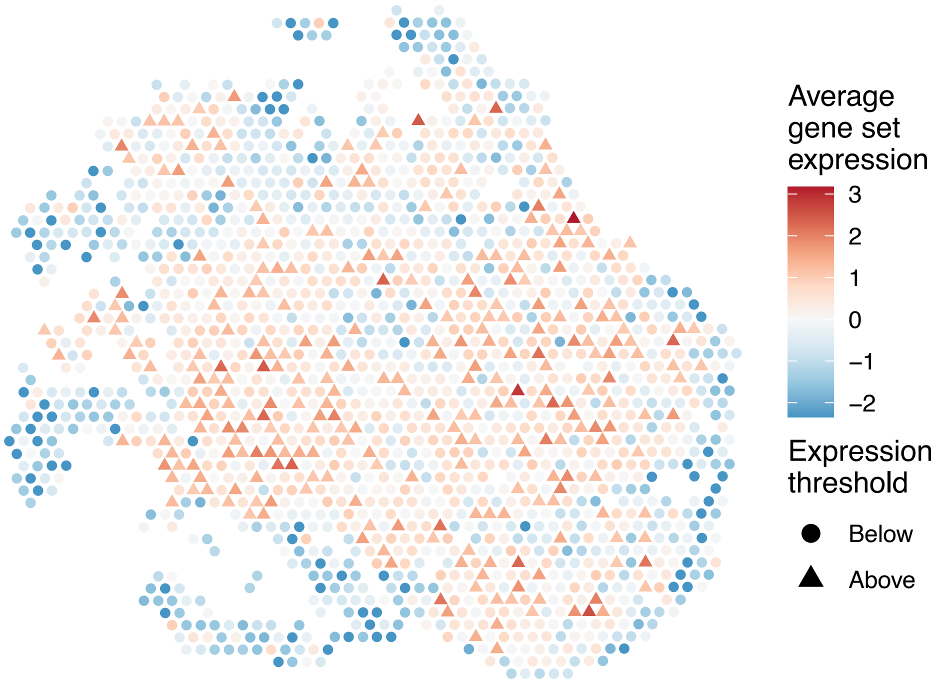

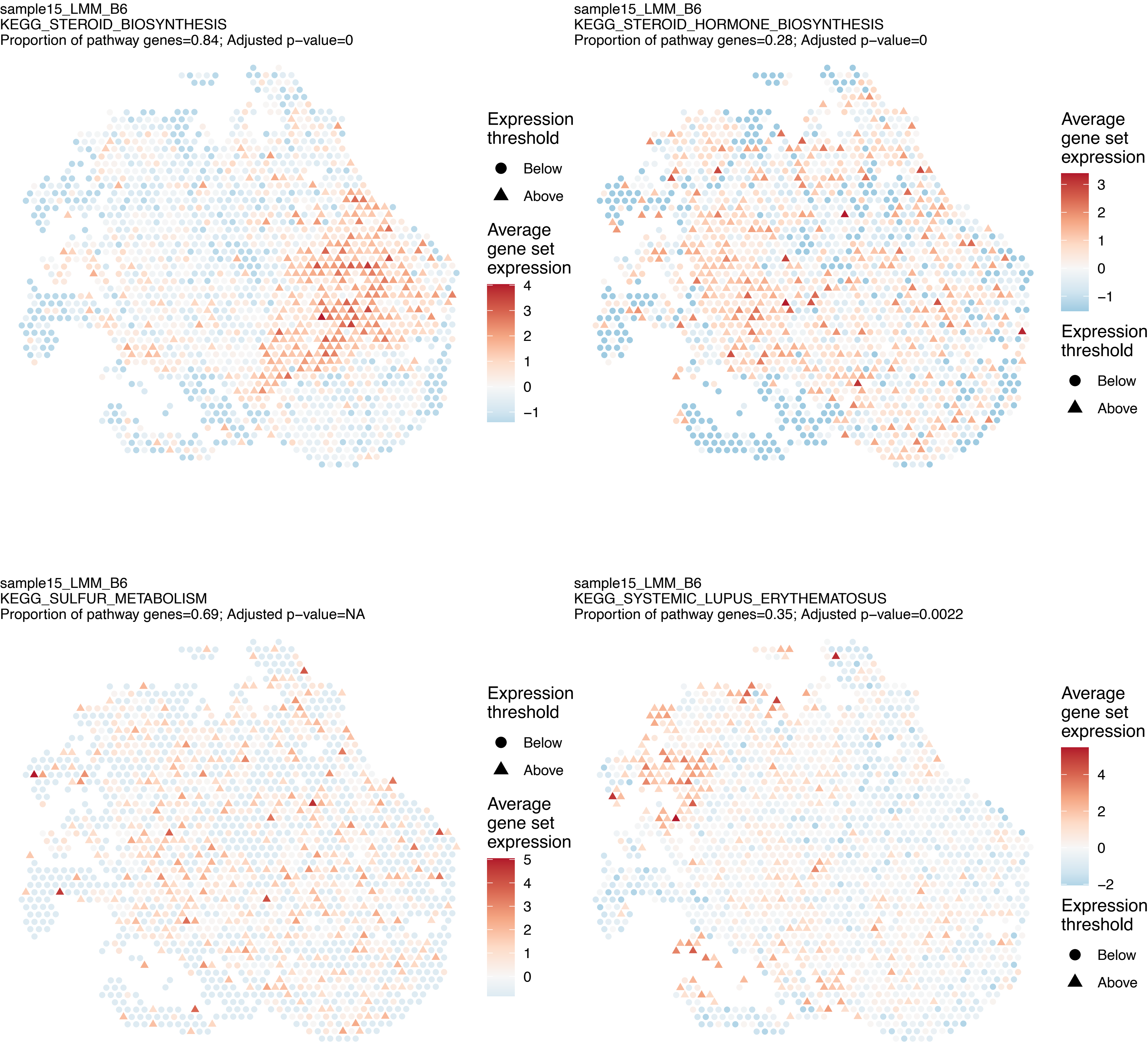
sample15_LMM_B6
KEGG_STEROID_BIOSYNTHESIS
Proportion of pathway genes=0.84; Adjusted p−value=0
Expression threshold
Below
Above
Average gene set expression
4
3
2
1
0
−1
sample15_LMM_B6
KEGG_STEROID_HORMONE_BIOSYNTHESIS
Proportion of pathway genes=0.28; Adjusted p−value=0
Average gene set expression
3
2
1
0
−1
Expression threshold
Below
Above
sample15_LMM_B6
KEGG_SULFUR_METABOLISM
Proportion of pathway genes=0.69; Adjusted p−value=NA
Expression threshold
Below
Above
Average gene set expression
5
4
3
2
1
0
sample15_LMM_B6
KEGG_SYSTEMIC_LUPUS_ERYTHEMATOSUS
Proportion of pathway genes=0.35; Adjusted p−value=0.0022
Average gene set expression
4
2
0
−2
Expression threshold
Below
Above

sample15_LMM_B6
KEGG_T_CELL_RECEPTOR_SIGNALING_PATHWAY
Proportion of pathway genes=0.65; Adjusted p−value=0

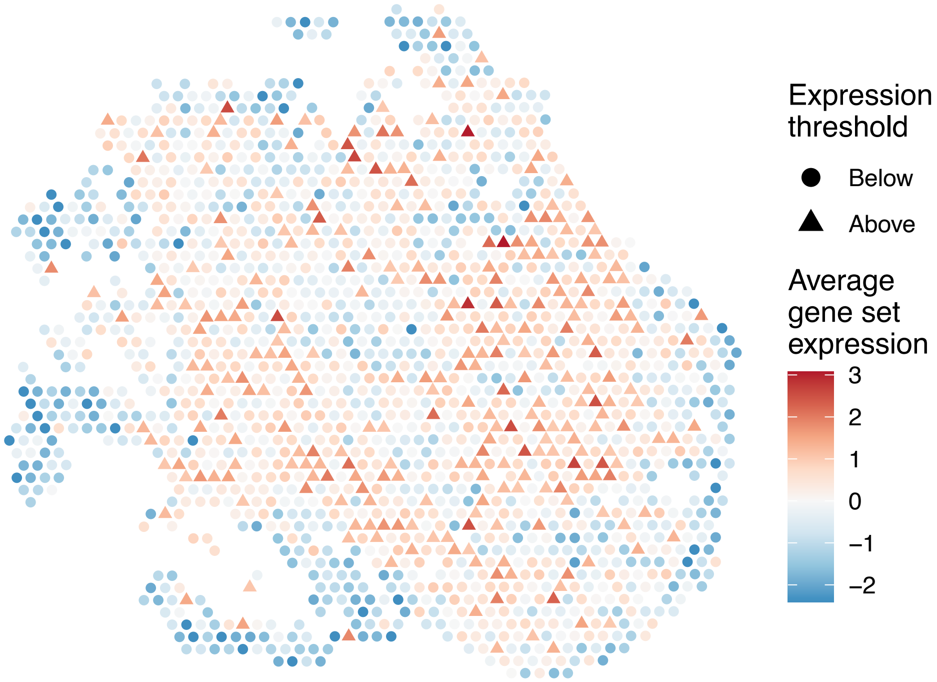


sample15_LMM_B6
KEGG_TERPENOID_BACKBONE_BIOSYNTHESIS
Proportion of pathway genes=0.87; Adjusted p−value=0

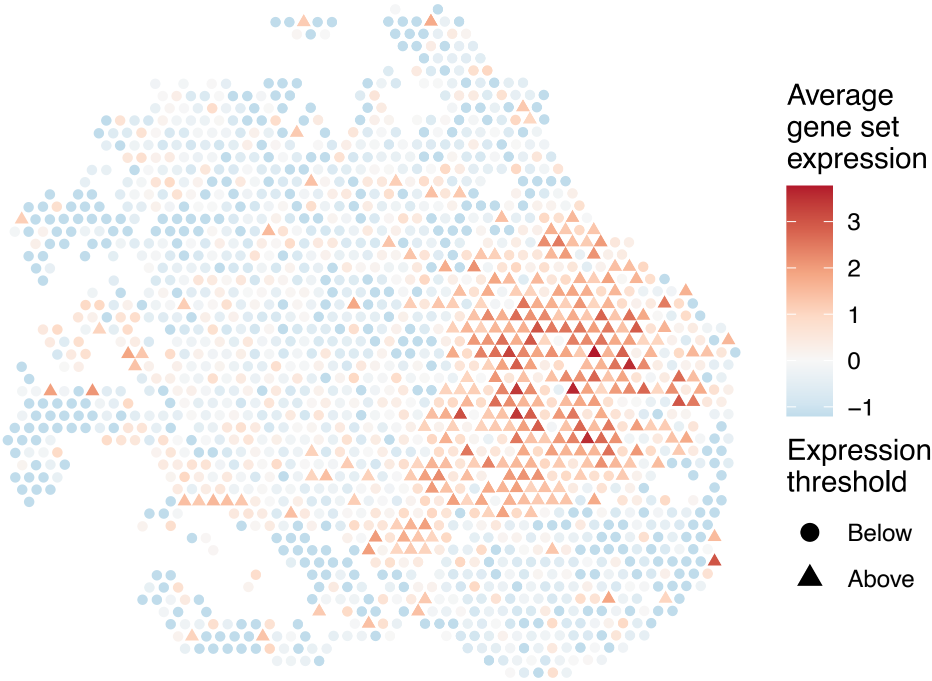


sample15_LMM_B6
KEGG_TGF_BETA_SIGNALING_PATHWAY
Proportion of pathway genes=0.68; Adjusted p−value=0

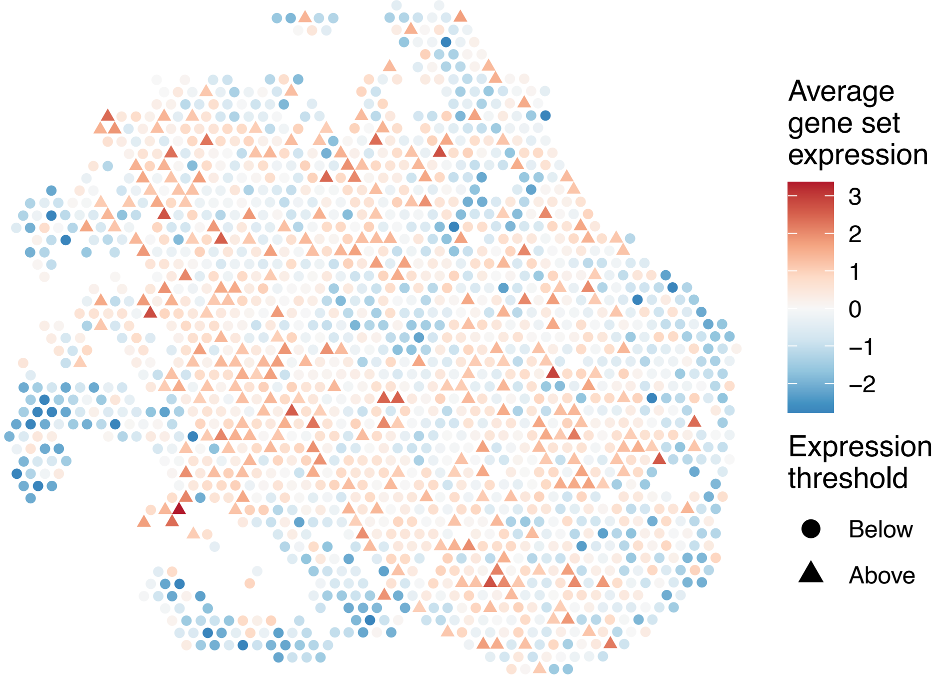


sample15_LMM_B6
KEGG_THYROID_CANCER
Proportion of pathway genes=0.89; Adjusted p−value=0

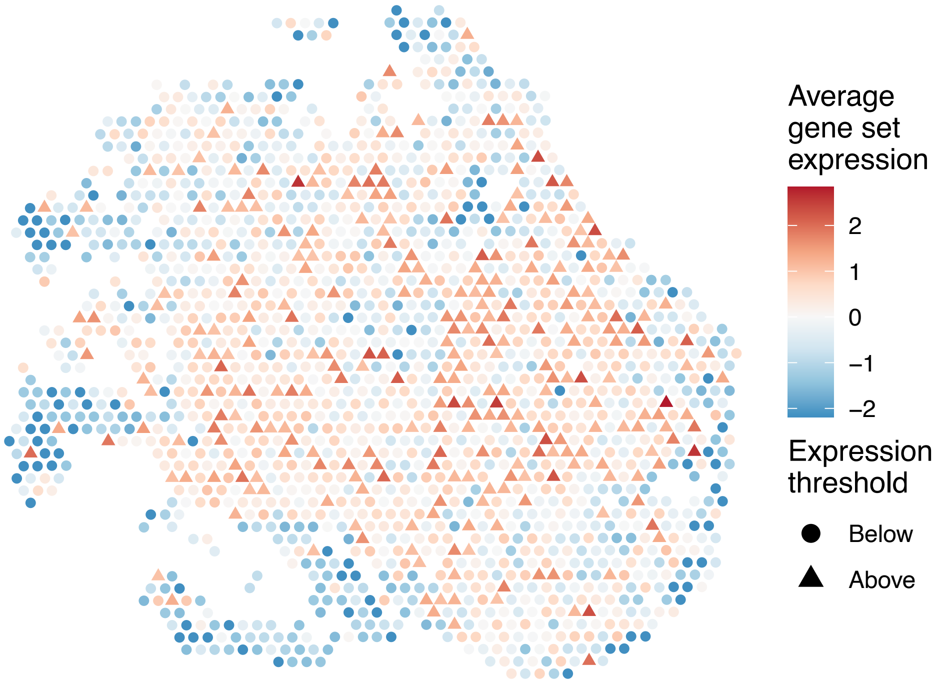

sample15_LMM_B6
KEGG_TIGHT_JUNCTION
Proportion of pathway genes=0.69; Adjusted p−value=0

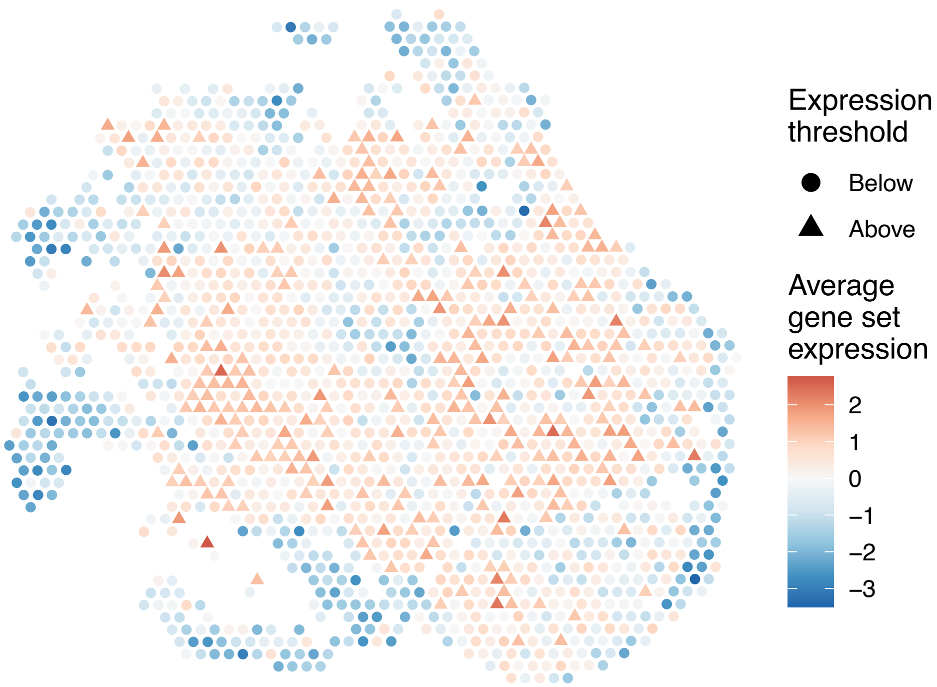


sample15_LMM_B6
KEGG_TOLL_LIKE_RECEPTOR_SIGNALING_PATHWAY
Proportion of pathway genes=0.67; Adjusted p−value=0

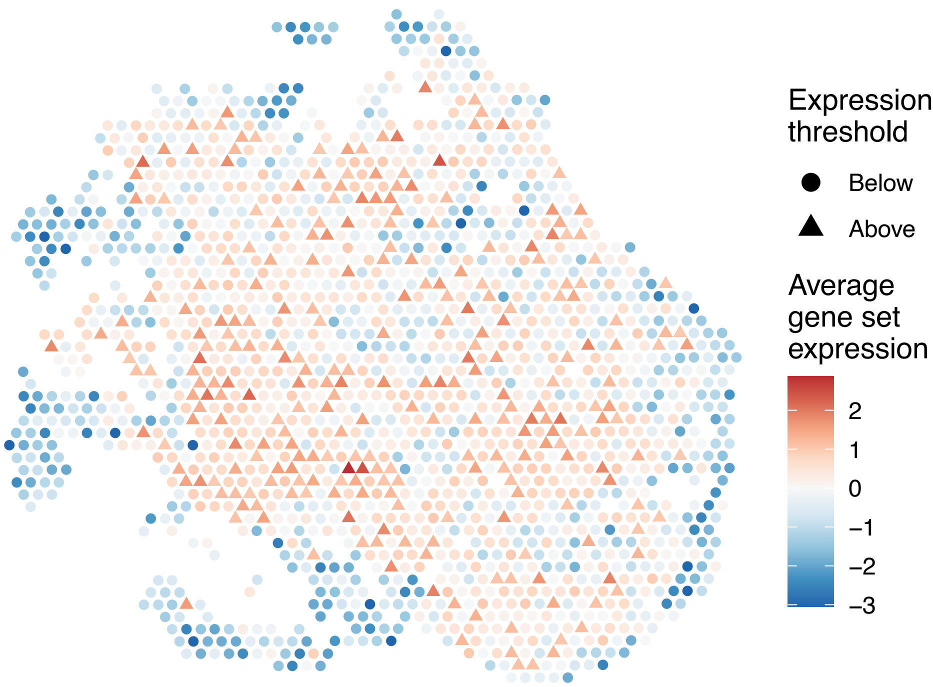


sample15_LMM_B6
KEGG_TRYPTOPHAN_METABOLISM
Proportion of pathway genes=0.59; Adjusted p−value=0

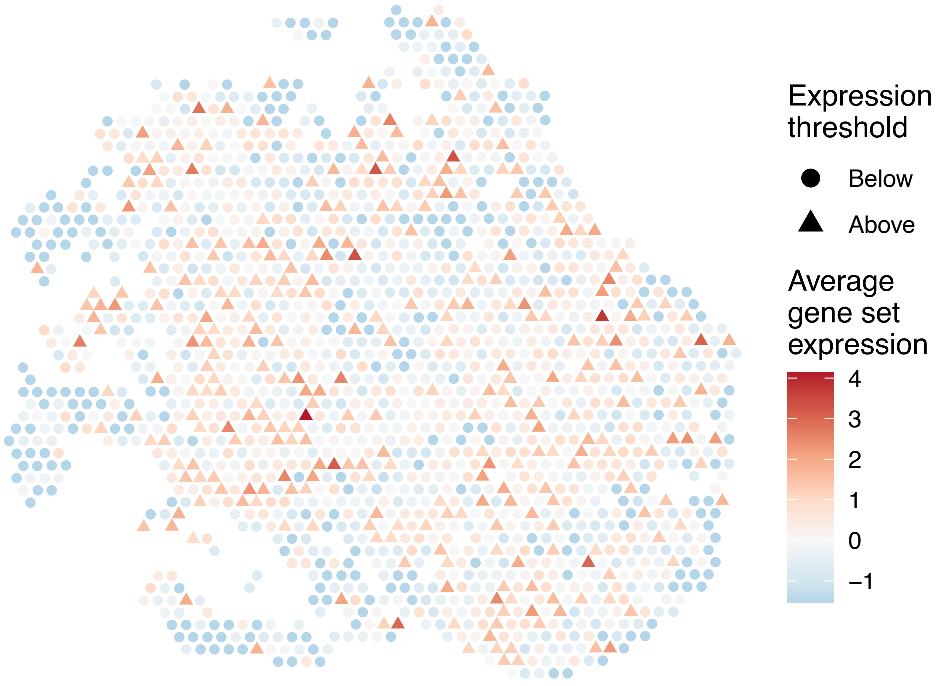


sample15_LMM_B6
KEGG_TYPE_I_DIABETES_MELLITUS
Proportion of pathway genes=0.59; Adjusted p−value=0

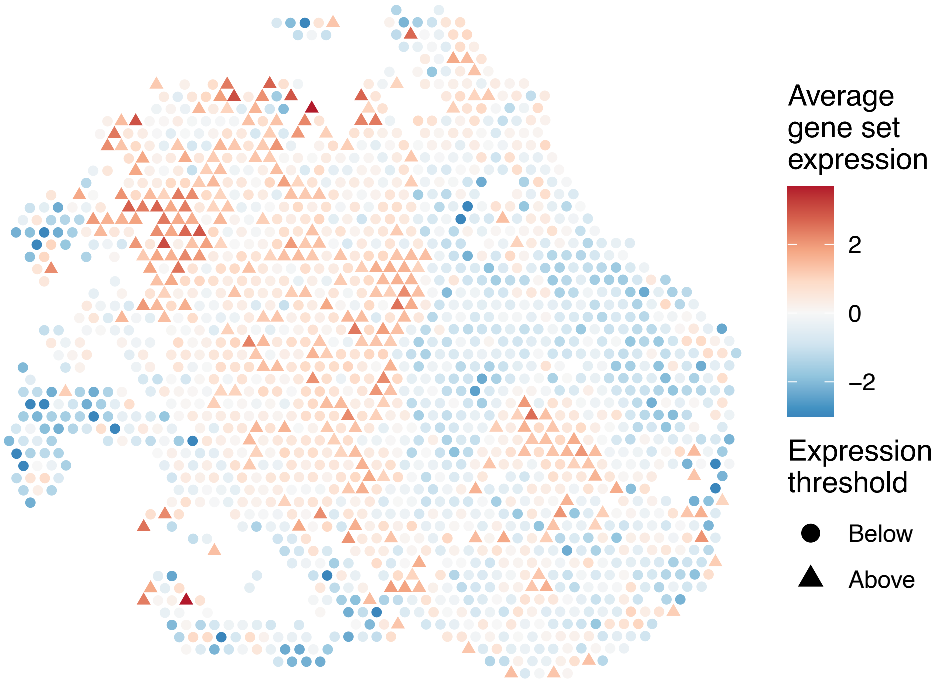

sample15_LMM_B6
KEGG_TYPE_II_DIABETES_MELLITUS
Proportion of pathway genes=0.51; Adjusted p–value=0

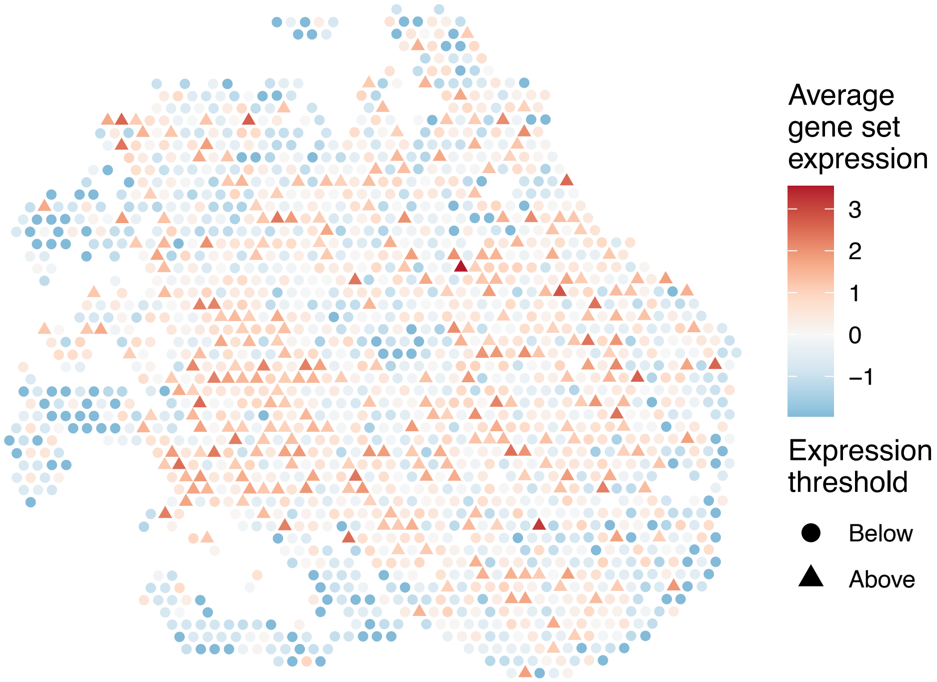


sample15_LMM_B6
KEGG_TYROSINE_METABOLISM
Proportion of pathway genes=0.64; Adjusted p–value=0

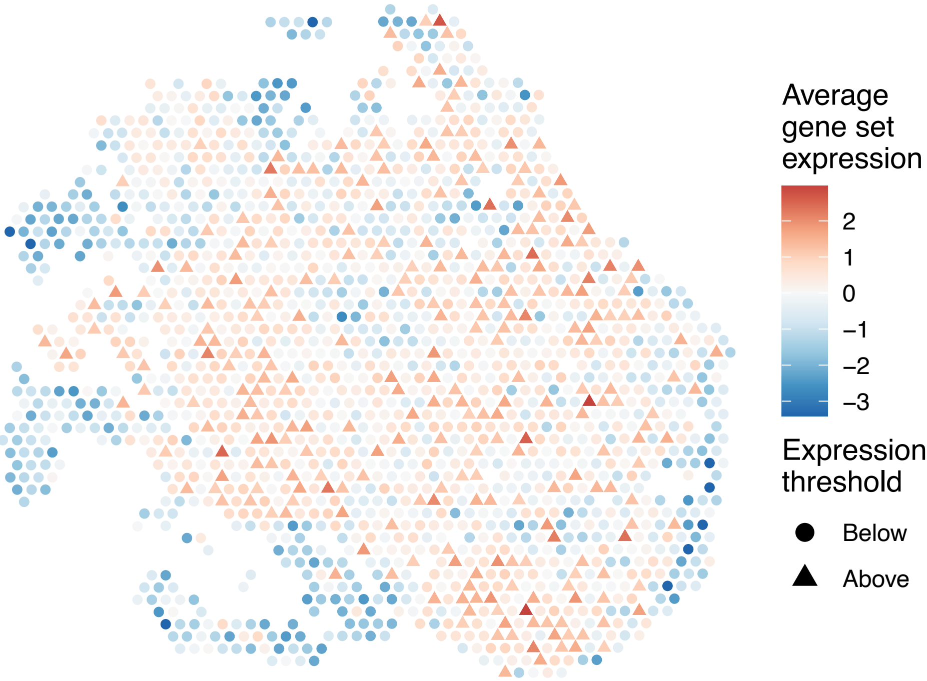


sample15_LMM_B6
KEGG_UBIQUITIN_MEDIATED_PROTEOLYSIS
Proportion of pathway genes=0.94; Adjusted p–value=0

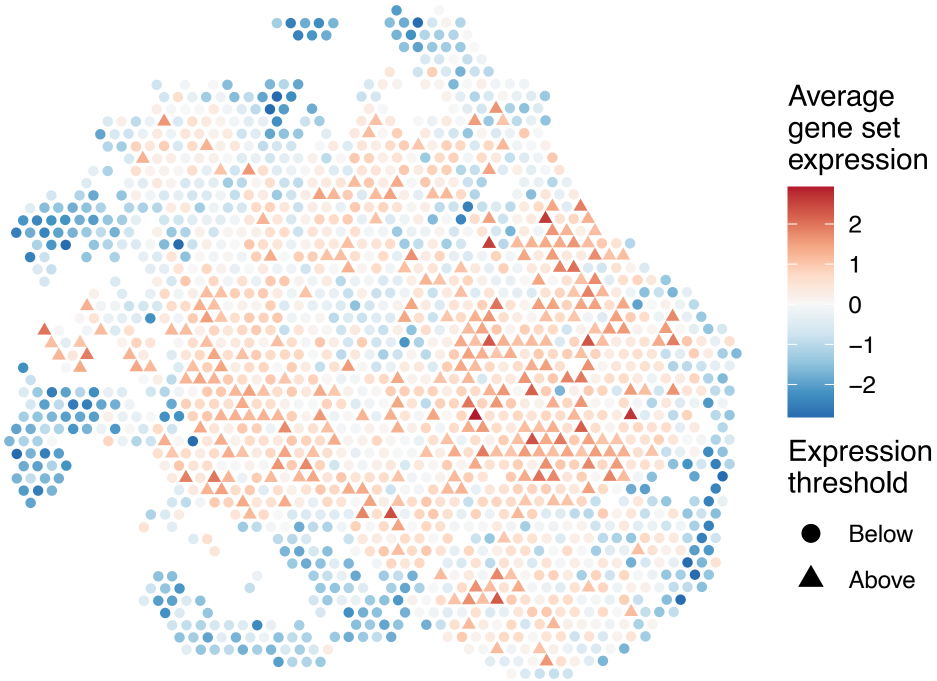


sample15_LMM_B6
KEGG_VALINE_LEUCINE_AND_ISOLEUCINE_BIOSYNTHESIS
Proportion of pathway genes=0.57; Adjusted p–value=0

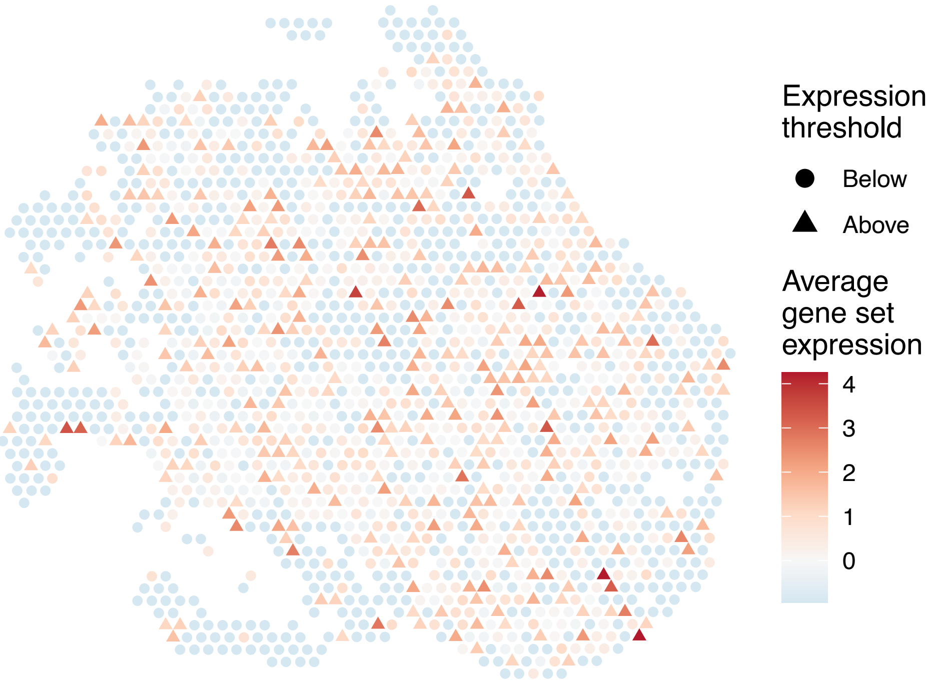

sample15_LMM_B6
KEGG_VALINE_LEUCINE_AND_ISOLEUCINE_DEGRADATION
Proportion of pathway genes=0.93; Adjusted p−value=0

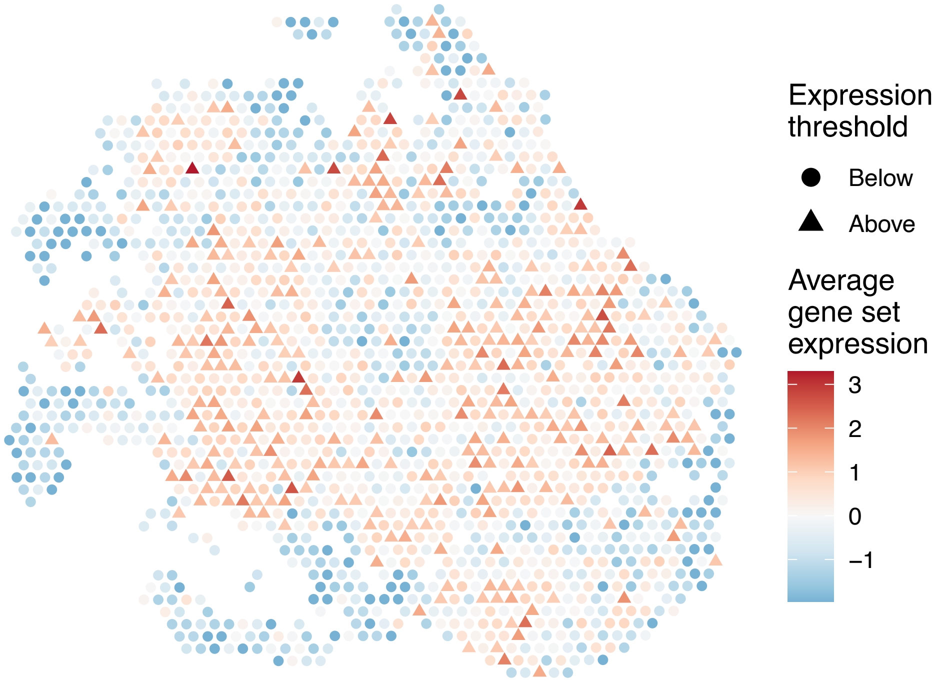


sample15_LMM_B6
KEGG_VASCULAR_SMOOTH_MUSCLE_CONTRACTION
Proportion of pathway genes=0.59; Adjusted p−value=0

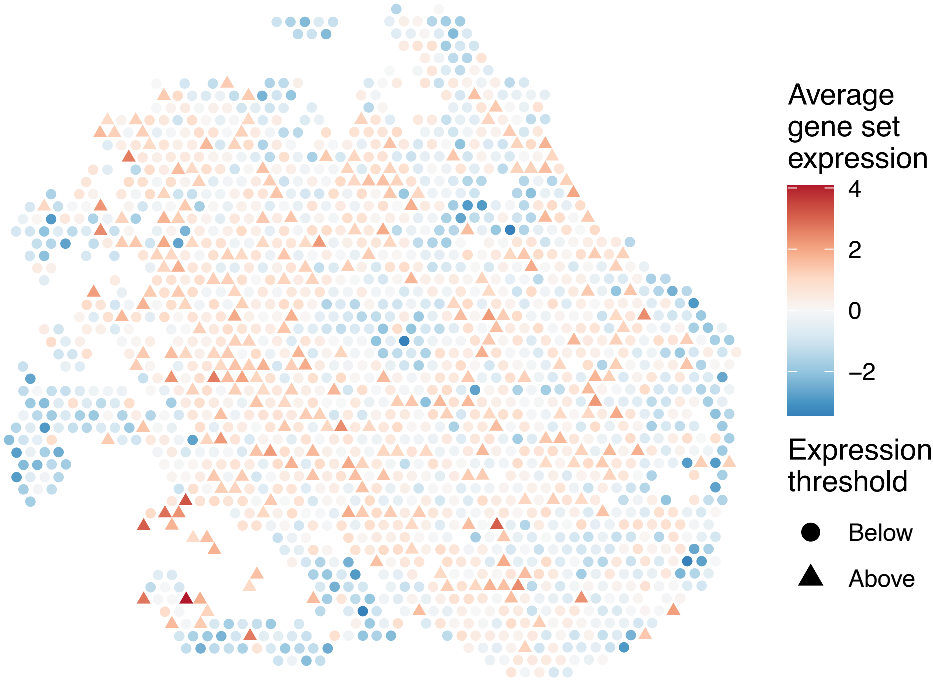


sample15_LMM_B6
KEGG_VASOPRESSIN_REGULATED_WATER_REABSORPTION
Proportion of pathway genes=0.85; Adjusted p−value=0

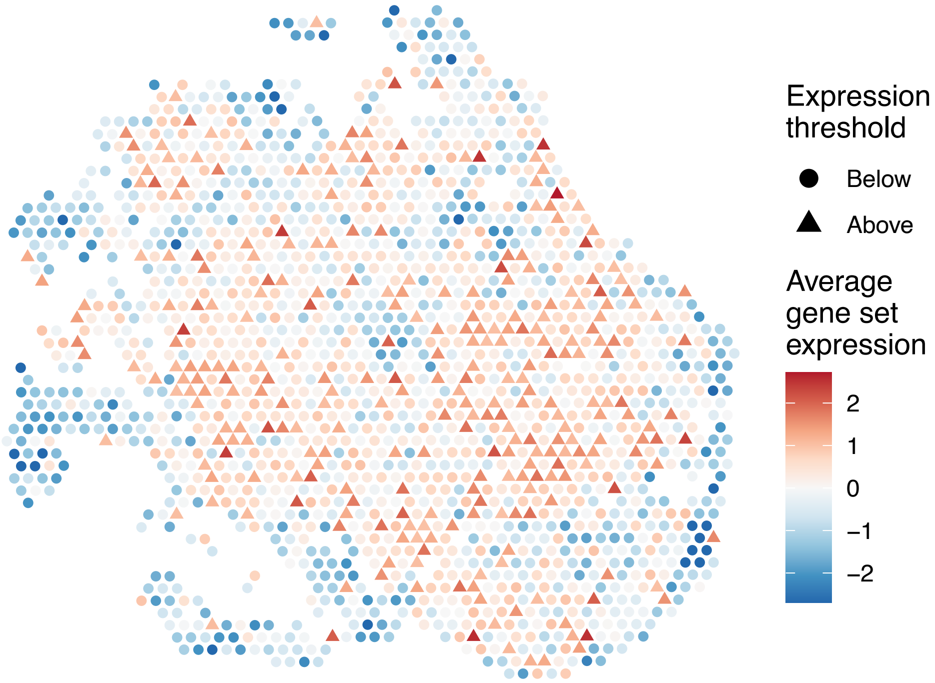


sample15_LMM_B6
KEGG_VEGF_SIGNALING_PATHWAY
Proportion of pathway genes=0.75; Adjusted p−value=0

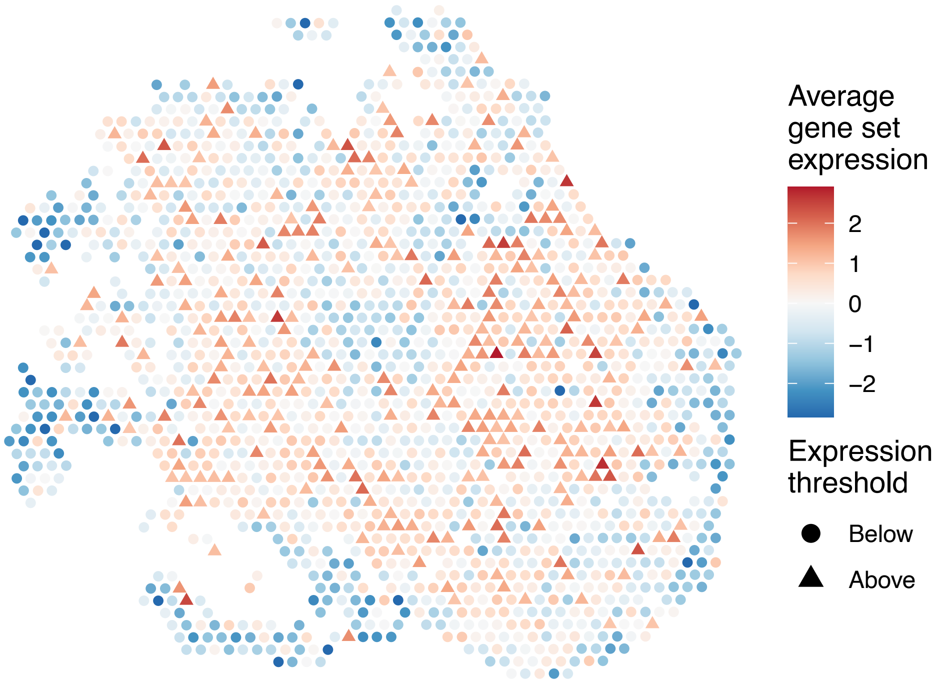

sample15_LMM_B6
KEGG_VIBRIO_CHOLERAE_INFECTION
Proportion of pathway genes=0.67; Adjusted p−value=0

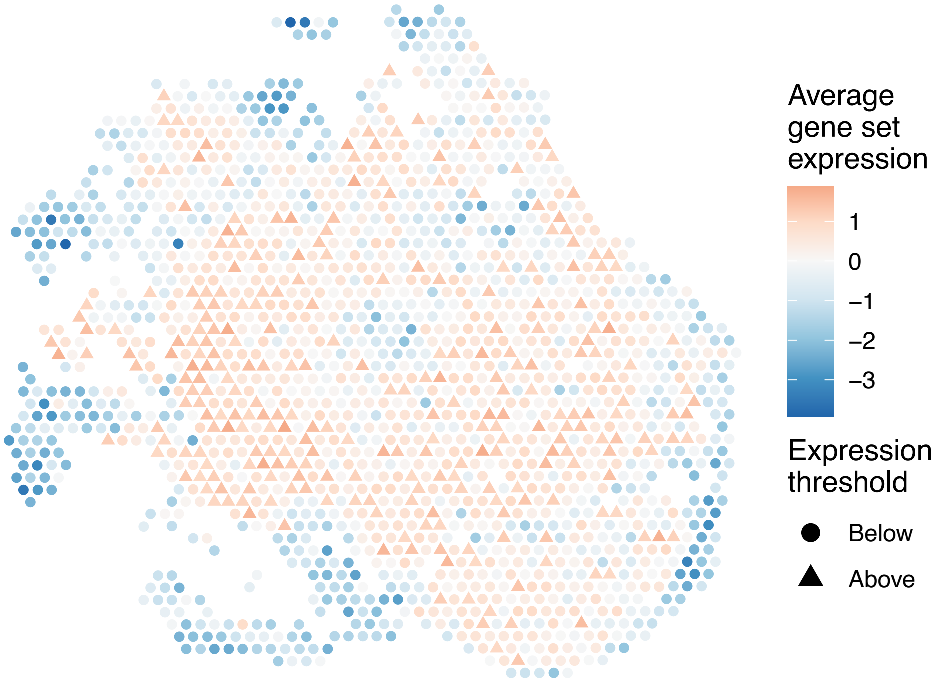


sample15_LMM_B6
KEGG_VIRAL_MYOCARDITIS
Proportion of pathway genes=0.69; Adjusted p−value=0

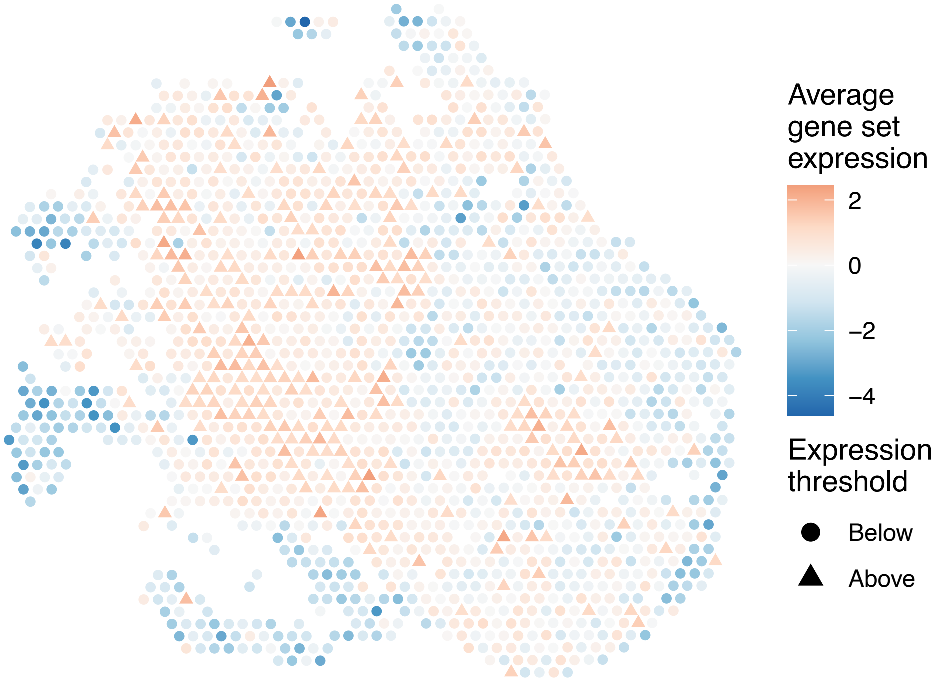


sample15_LMM_B6
KEGG_WNT_SIGNALING_PATHWAY
Proportion of pathway genes=0.75; Adjusted p−value=0

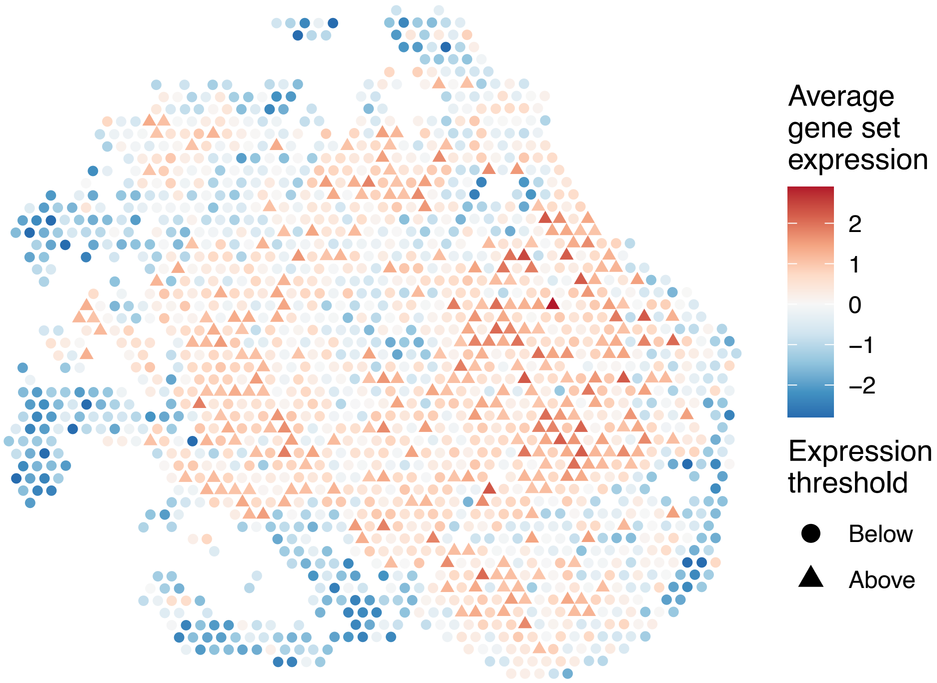


sample15_LMM_B6
KEGG_PENTOSE_AND_GLUCURONATE_INTERCONVERSIONS
Proportion of pathway genes=0.29; Adjusted p−value=NA

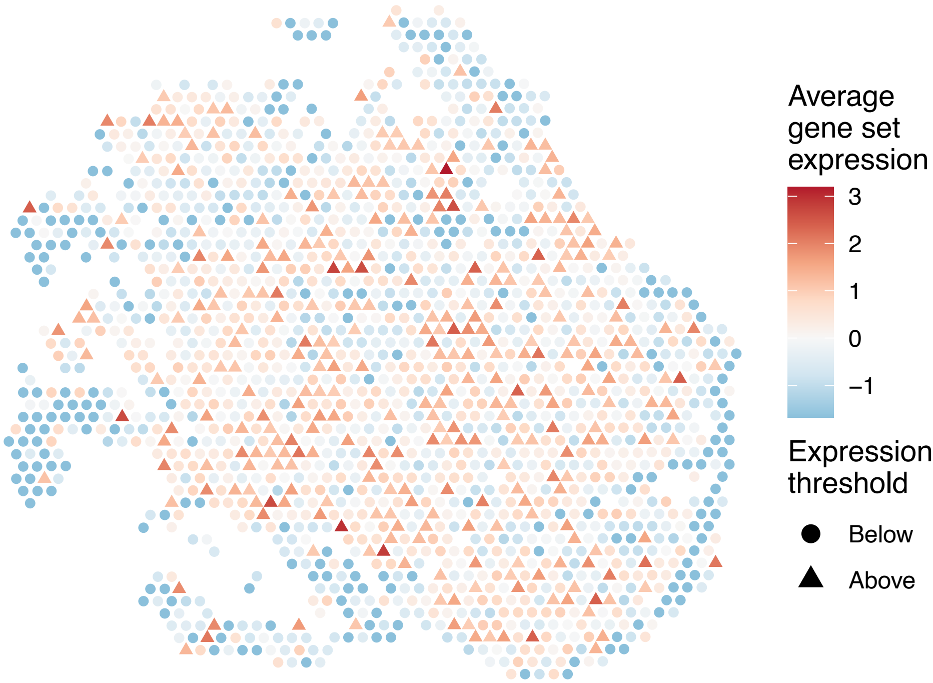

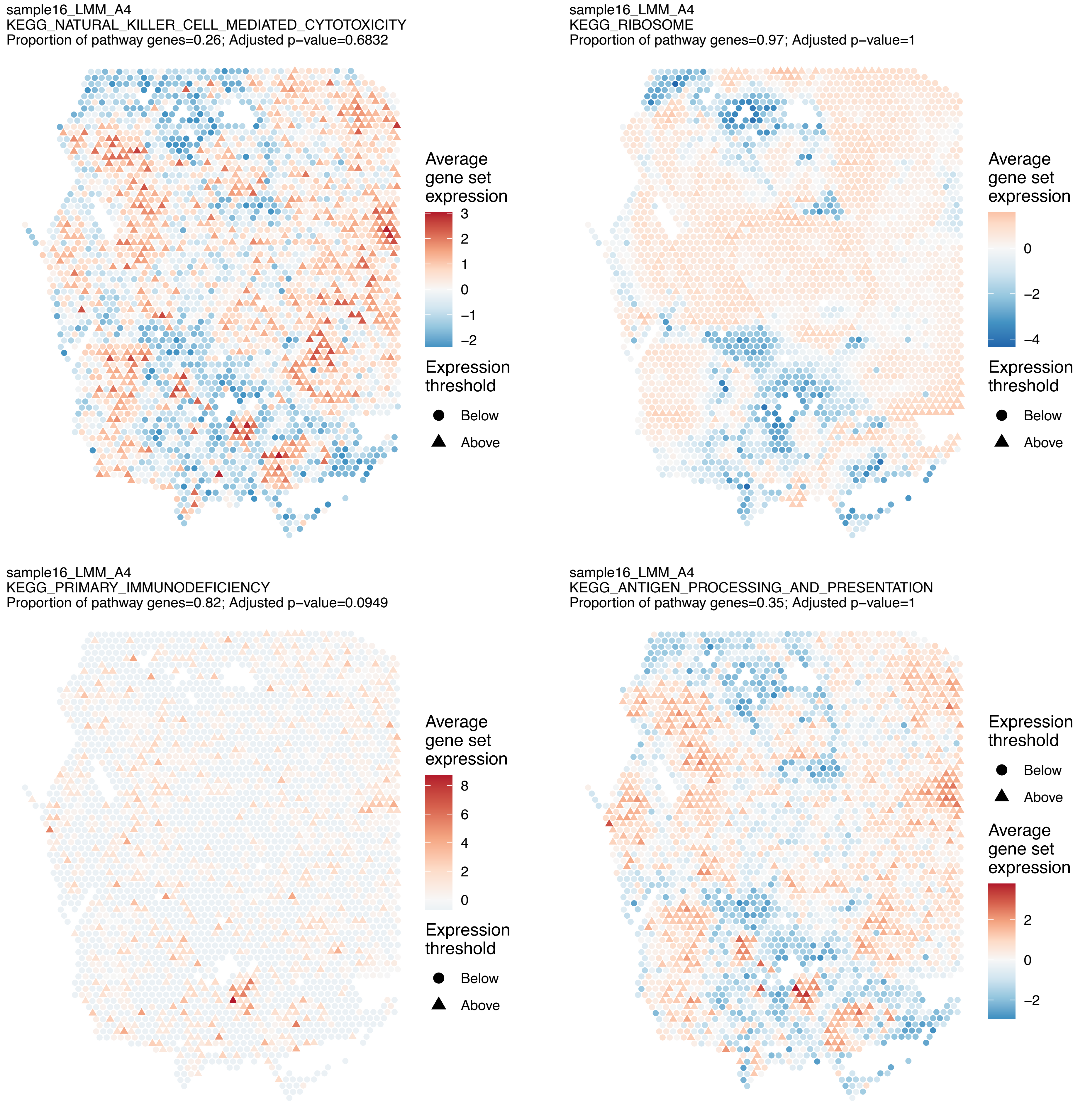
sample16_LMM_A4
KEGG_NATURAL_KILLER_CELL_MEDIATED_CYTOTOXICITY
Proportion of pathway genes=0.26; Adjusted p–value=0.6832
Average gene set expression
3
2
1
0
−1
−2
Expression threshold
Below
Above
sample16_LMM_A4
KEGG_RIBOSOME
Proportion of pathway genes=0.97; Adjusted p–value=1
Average gene set expression
0
−2
−4
Expression threshold
Below
Above
sample16_LMM_A4
KEGG_PRIMARY_IMMUNODEFICIENCY
Proportion of pathway genes=0.82; Adjusted p–value=0.0949
Average gene set expression
8
6
4
2
0
Expression threshold
Below
Above
sample16_LMM_A4
KEGG_ANTIGEN_PROCESSING_AND_PRESENTATION
Proportion of pathway genes=0.35; Adjusted p–value=1
Expression threshold
Below
Above
Average gene set expression
2
0
−2

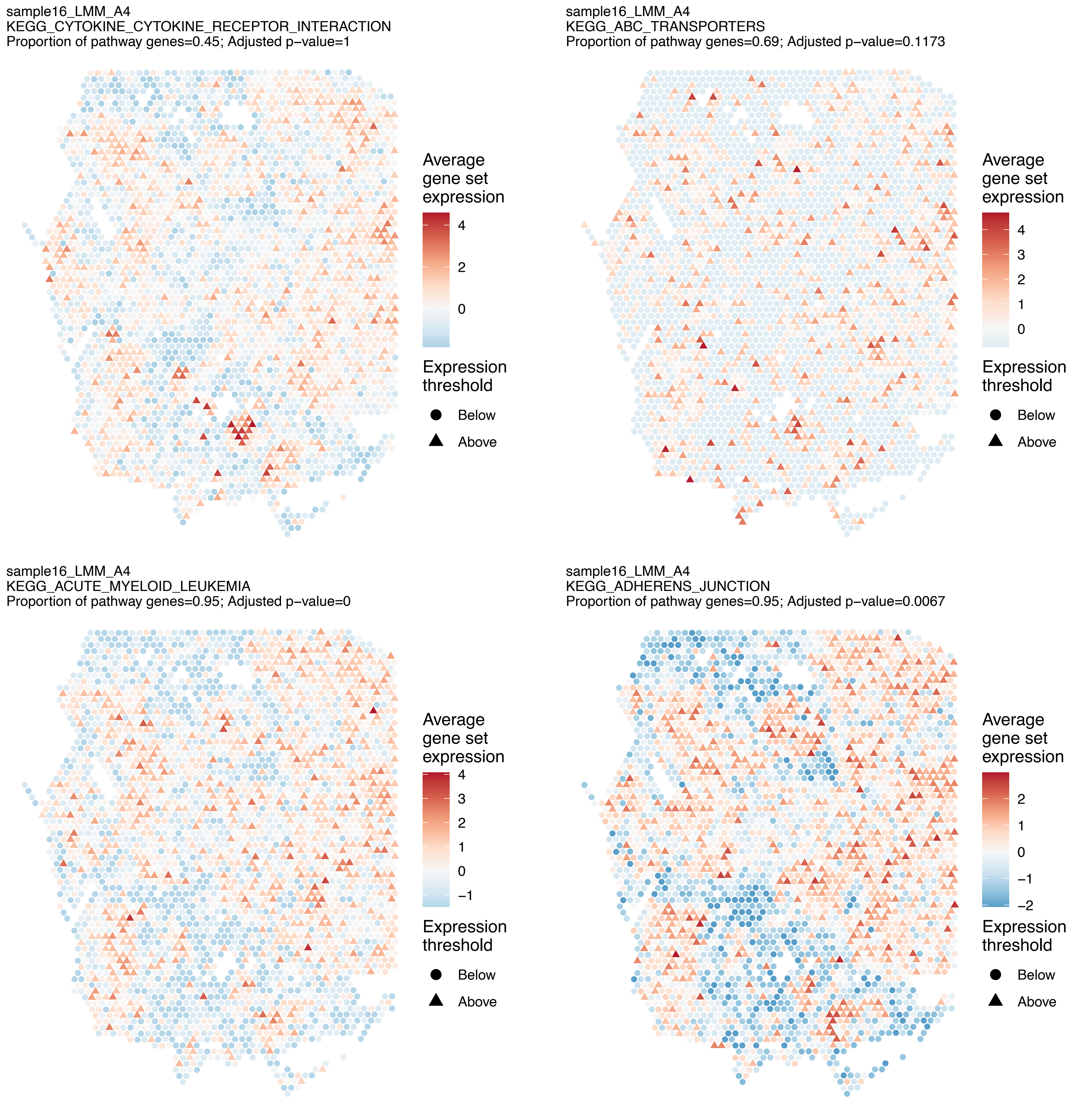
sample16_LMM_A4
KEGG_CYTOKINE_CYTOKINE_RECEPTOR_INTERACTION
Proportion of pathway genes=0.45; Adjusted p−value=1
Average gene set expression
4
2
0
Expression threshold
Below
Above
sample16_LMM_A4
KEGG_ABC_TRANSPORTERS
Proportion of pathway genes=0.69; Adjusted p−value=0.1173
Average gene set expression
4
3
2
1
0
Expression threshold
Below
Above
sample16_LMM_A4
KEGG_ACUTE_MYELOID_LEUKEMIA
Proportion of pathway genes=0.95; Adjusted p−value=0
Average gene set expression
4
3
2
1
0
−1
Expression threshold
Below
Above
sample16_LMM_A4
KEGG_ADHERENS_JUNCTION
Proportion of pathway genes=0.95; Adjusted p−value=0.0067
Average gene set expression
2
1
0
−1
−2
Expression threshold
Below
Above

sample16_LMM_A4
KEGG_ADIPOCYTOKINE_SIGNALING_PATHWAY
Proportion of pathway genes=0.72; Adjusted p−value=0.0219

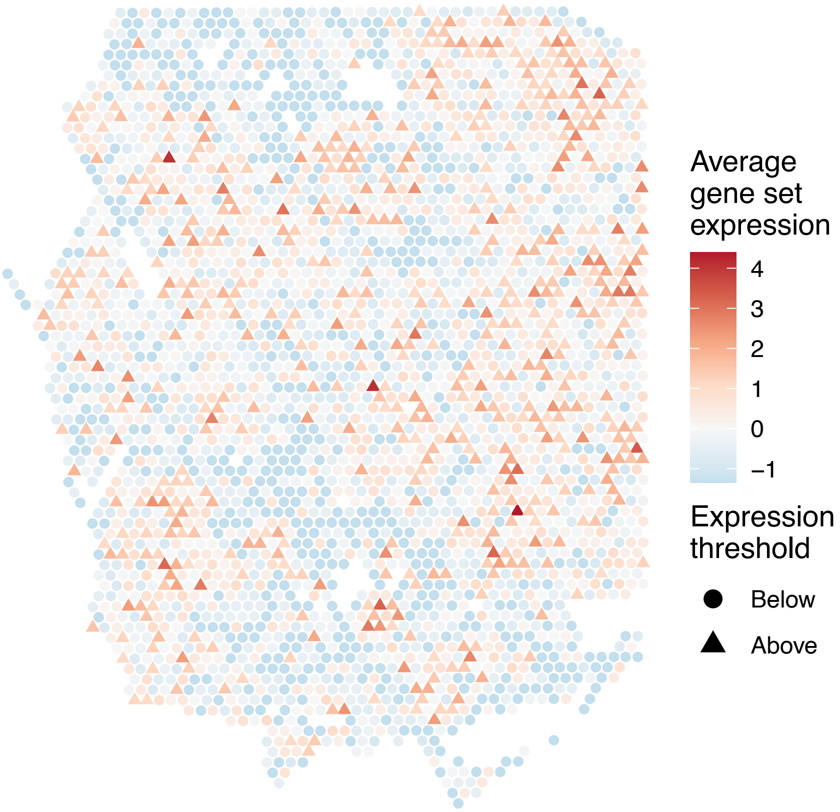


sample16_LMM_A4
KEGG_ALANINE_ASPARTATE_AND_GLUTAMATE_METABOLISM
Proportion of pathway genes=0.64; Adjusted p−value=0.4905

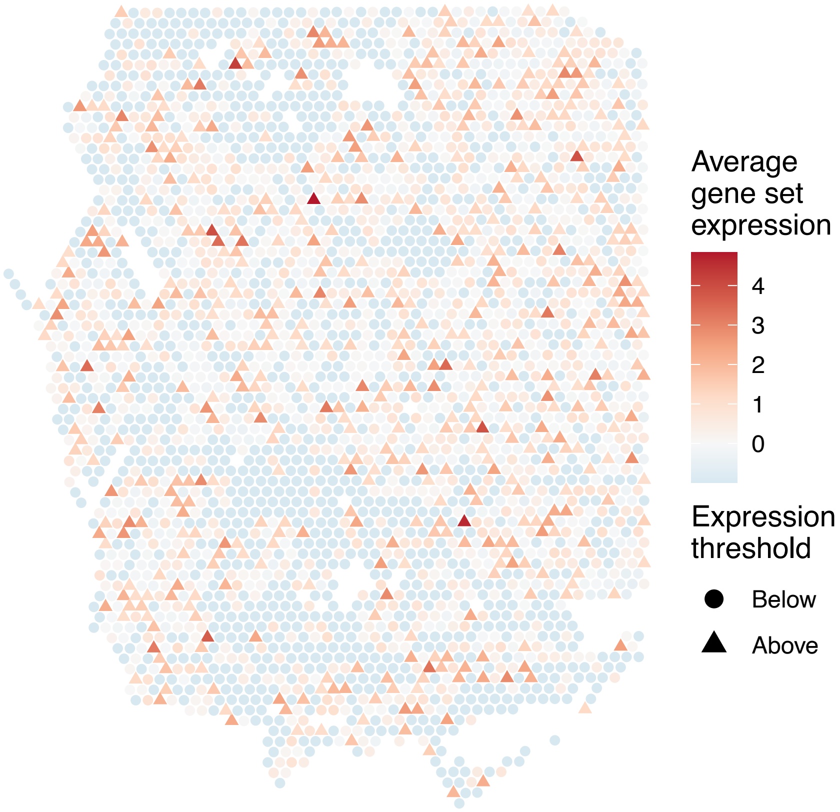


sample16_LMM_A4
KEGG_ALDOSTERONE_REGULATED_SODIUM_REABSORPTION
Proportion of pathway genes=0.62; Adjusted p−value=0.0106

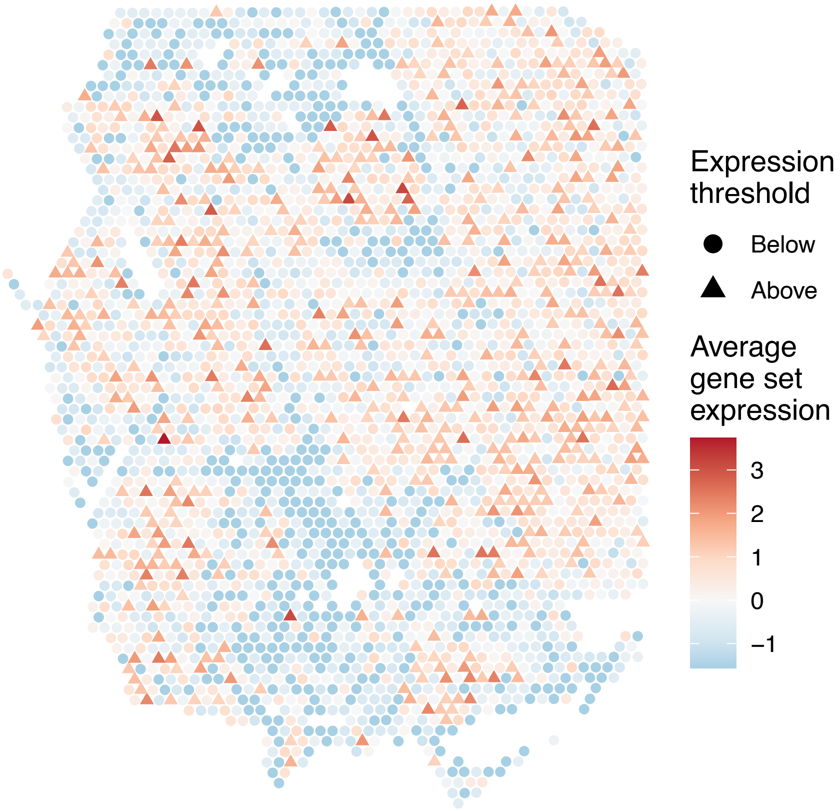


sample16_LMM_A4
KEGG_ALLOGRAFT_REJECTION
Proportion of pathway genes=0.75; Adjusted p−value=1

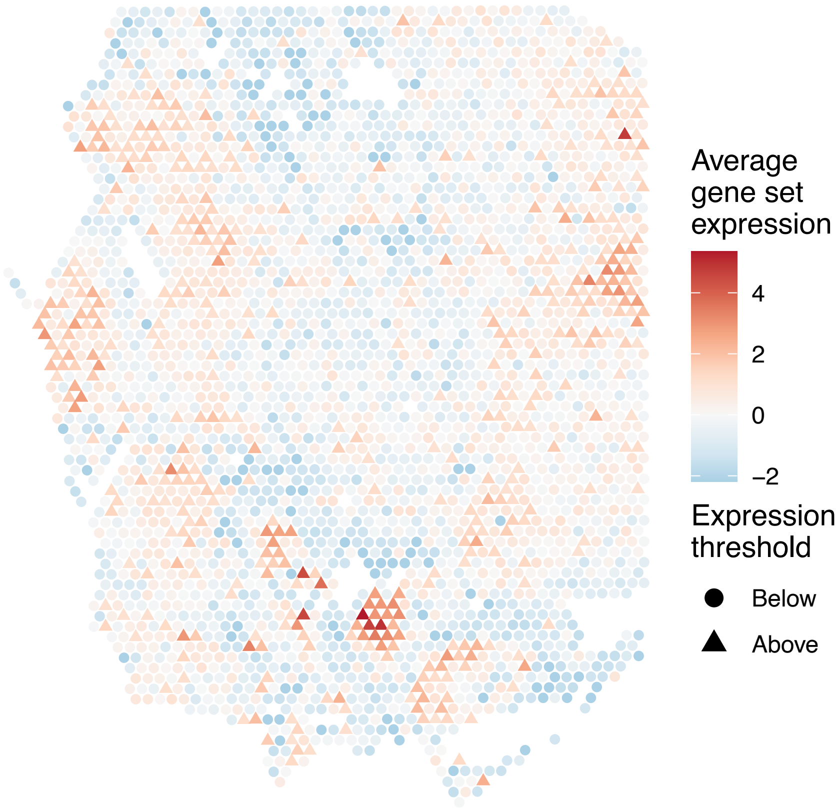

sample16_LMM_A4
KEGG_ALZHEIMERS_DISEASE
Proportion of pathway genes=0.8; Adjusted p−value=0

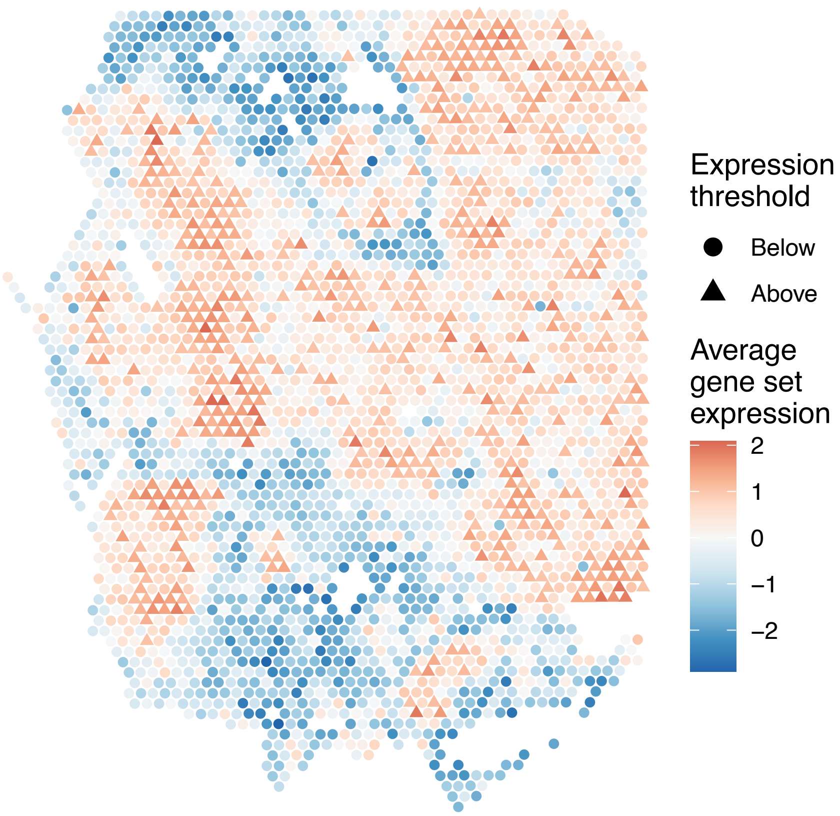


sample16_LMM_A4
KEGG_AMINO_SUGAR_AND_NUCLEOTIDE_SUGAR_METABOLISM
Proportion of pathway genes=0.89; Adjusted p−value=0.7472

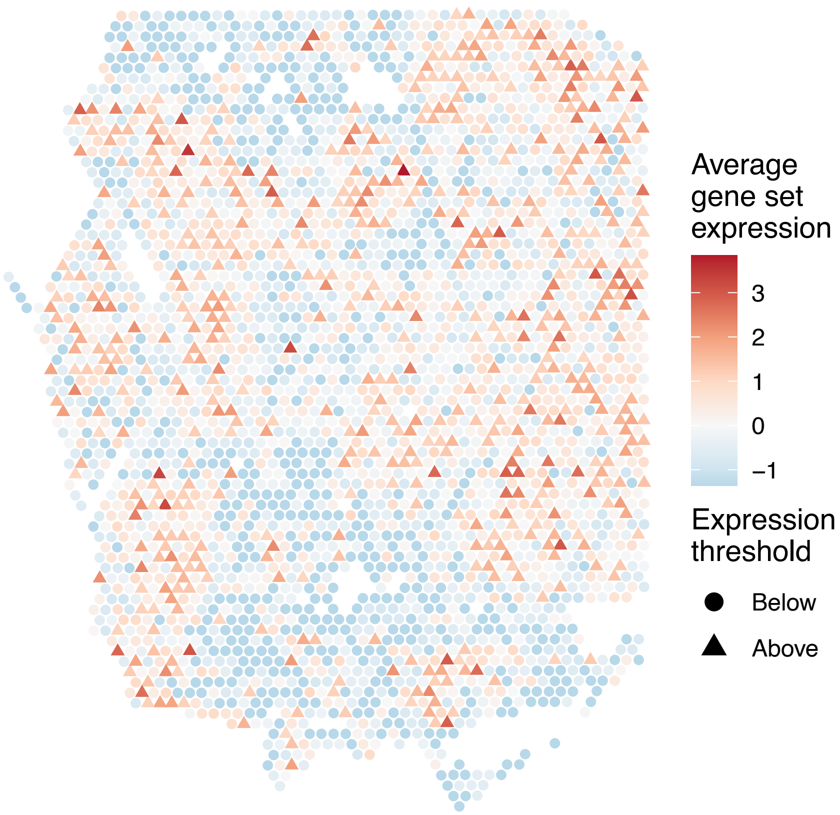


sample16_LMM_A4
KEGG_AMINOACYL_TRNA_BIOSYNTHESIS
Proportion of pathway genes=0.51; Adjusted p−value=0.2892

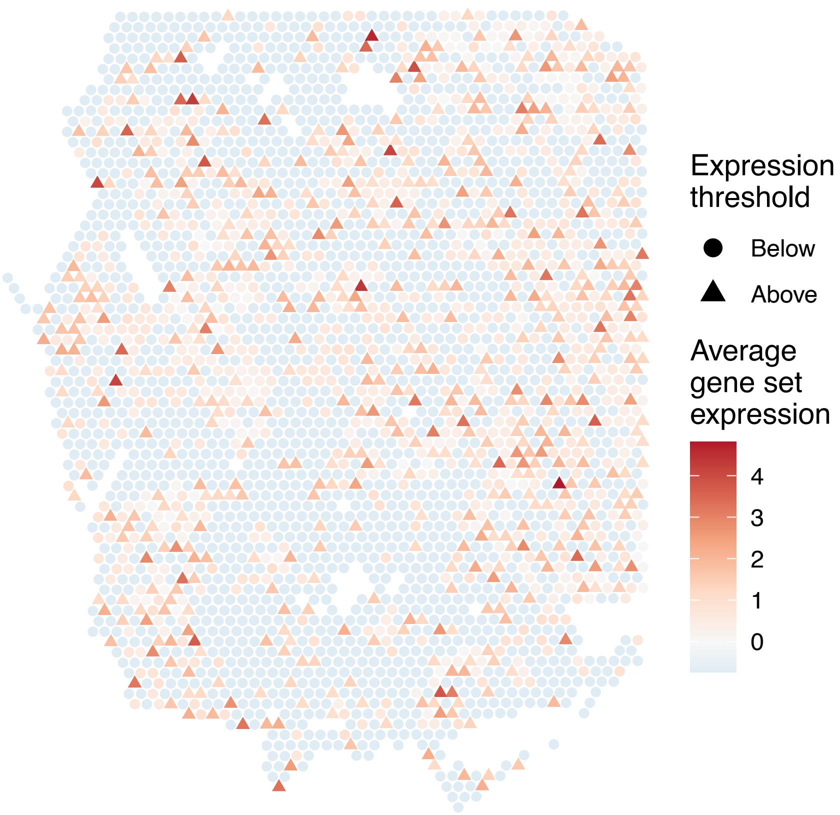


sample16_LMM_A4
KEGG_AMYOTROPHIC_LATERAL_SCLEROSIS_ALS
Proportion of pathway genes=0.65; Adjusted p−value=0.261

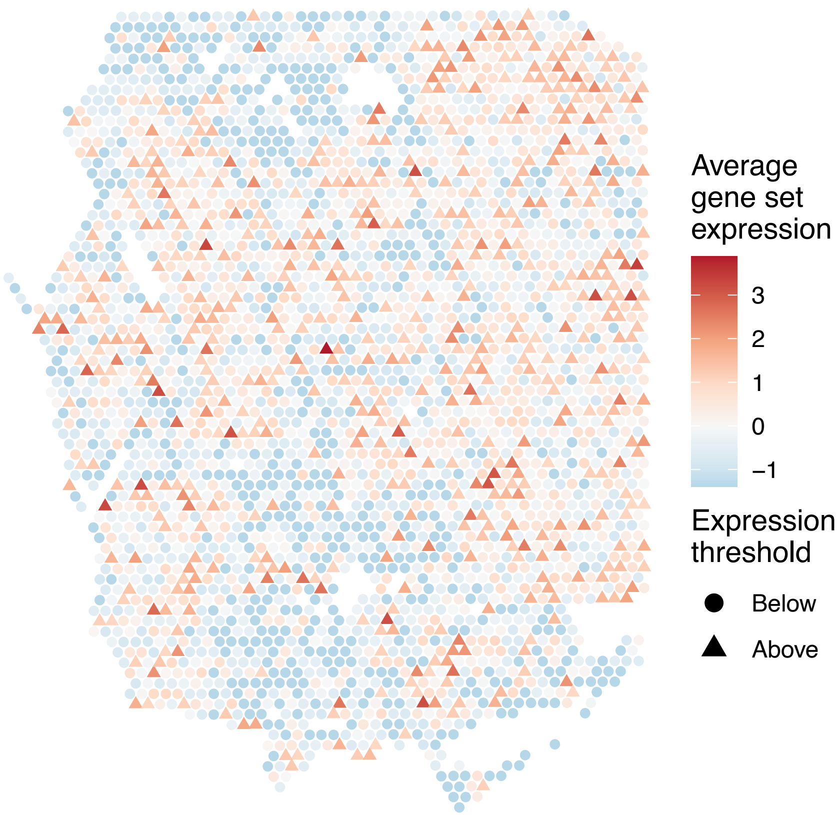

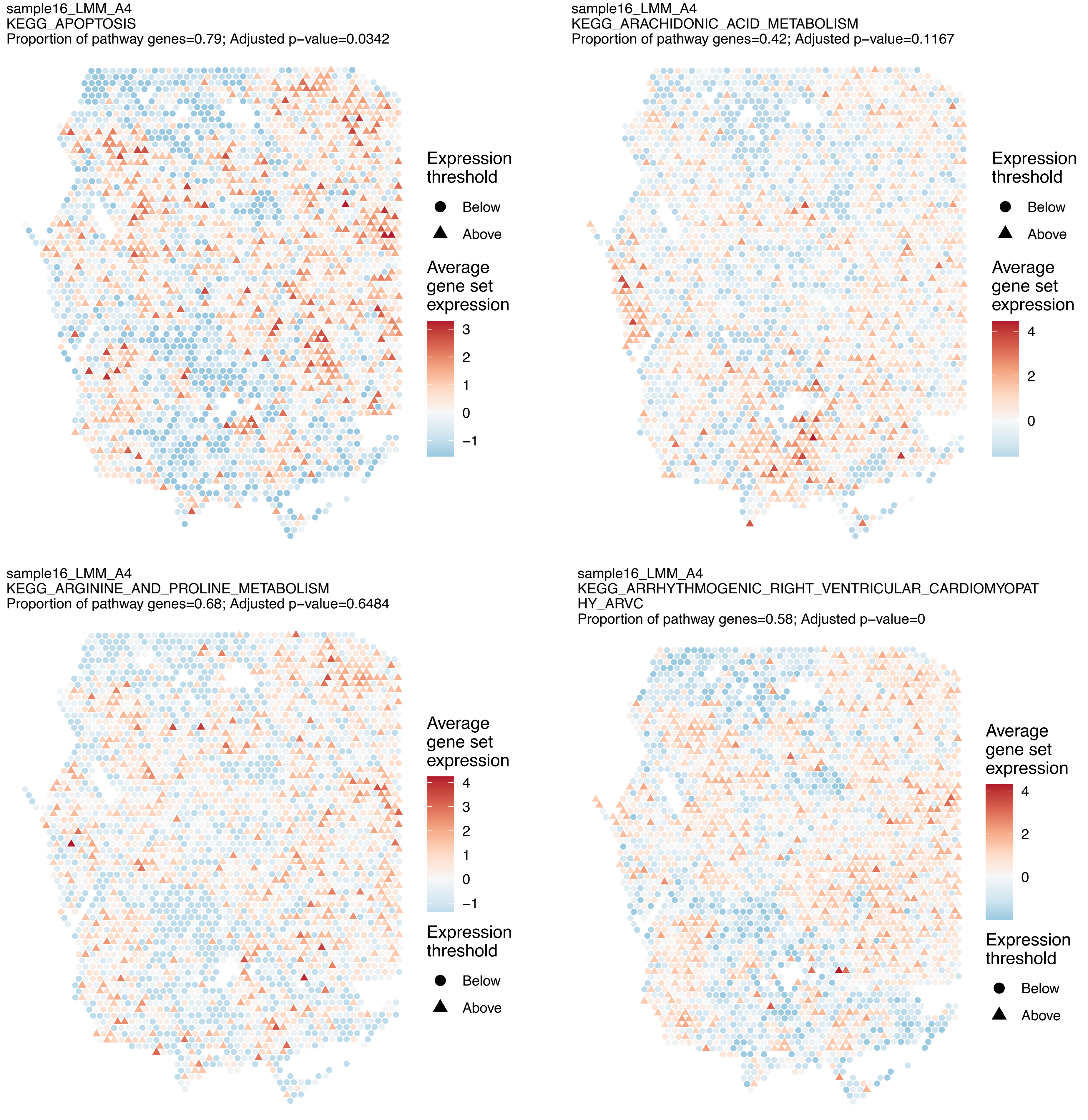
sample16_LMM_A4
KEGG_APOPTOSIS
Proportion of pathway genes=0.79; Adjusted p−value=0.0342
Expression threshold
Below
Above
Average gene set expression
3
2
1
0
−1
sample16_LMM_A4
KEGG_ARACHIDONIC_ACID_METABOLISM
Proportion of pathway genes=0.42; Adjusted p−value=0.1167
Expression threshold
Below
Above
Average gene set expression
4
2
0
sample16_LMM_A4
KEGG_ARGININE_AND_PROLINE_METABOLISM
Proportion of pathway genes=0.68; Adjusted p−value=0.6484
Average gene set expression
4
3
2
1
0
−1
Expression threshold
Below
Above
sample16_LMM_A4
KEGG_ARRHYTHMOGENIC_RIGHT_VENTRICULAR_CARDIOMYOPAT
HY_ARVC
Proportion of pathway genes=0.58; Adjusted p−value=0
Average gene set expression
4
2
0
Expression threshold
Below
Above

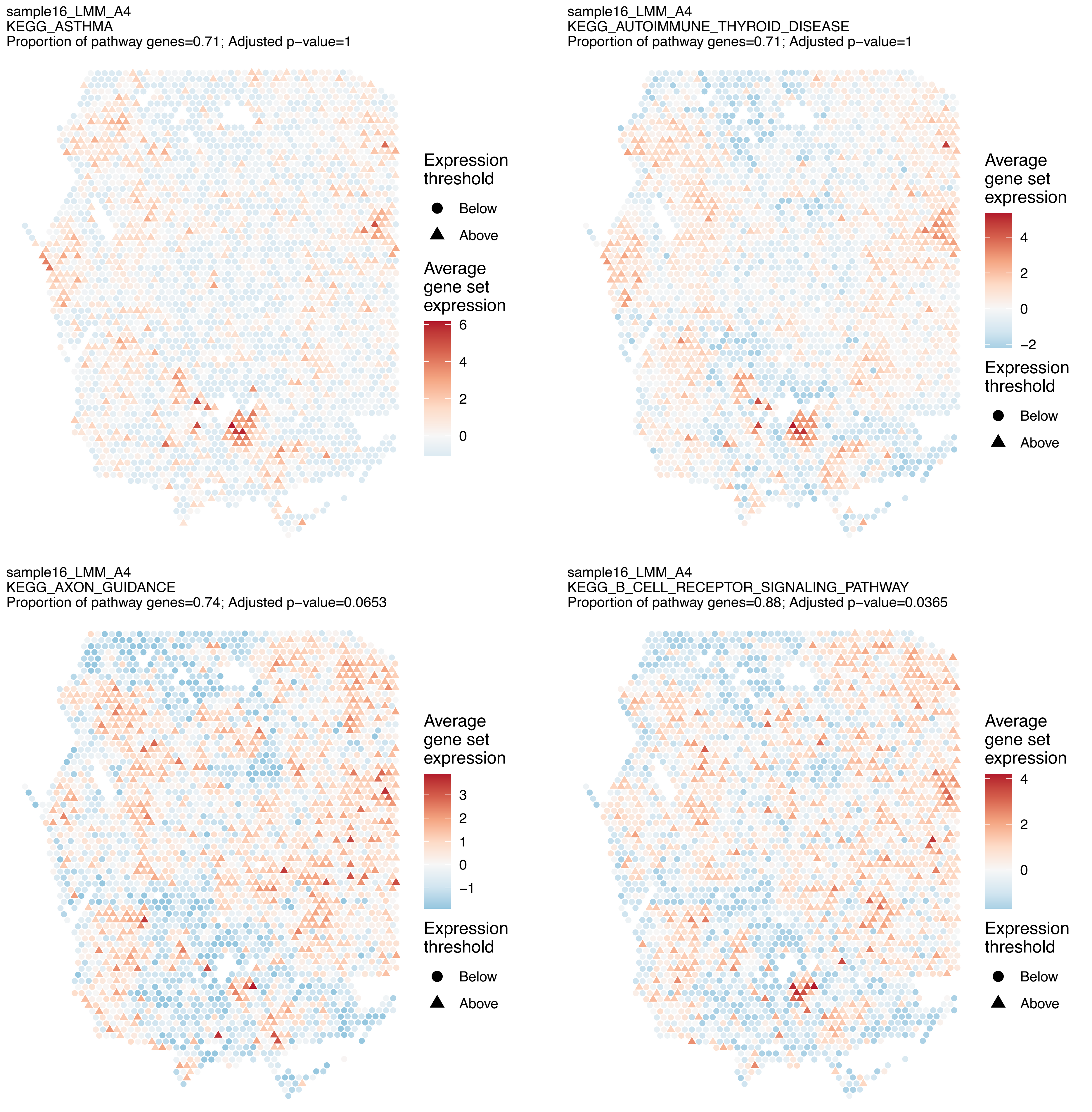
sample16_LMM_A4
KEGG_ASTHMA
Proportion of pathway genes=0.71; Adjusted p−value=1
Expression threshold
Below
Above
Average gene set expression
6
4
2
0
sample16_LMM_A4
KEGG_AUTOIMMUNE_THYROID_DISEASE
Proportion of pathway genes=0.71; Adjusted p−value=1
Average gene set expression
4
2
0
−2
Expression threshold
Below
Above
sample16_LMM_A4
KEGG_AXON_GUIDANCE
Proportion of pathway genes=0.74; Adjusted p−value=0.0653
Average gene set expression
3
2
1
0
−1
Expression threshold
Below
Above
sample16_LMM_A4
KEGG_B_CELL_RECEPTOR_SIGNALING_PATHWAY
Proportion of pathway genes=0.88; Adjusted p−value=0.0365
Average gene set expression
4
2
0
Expression threshold
Below
Above

sample16_LMM_A4
KEGG_BASAL_CELL_CARCINOMA
Proportion of pathway genes=0.62; Adjusted p−value=0.0574

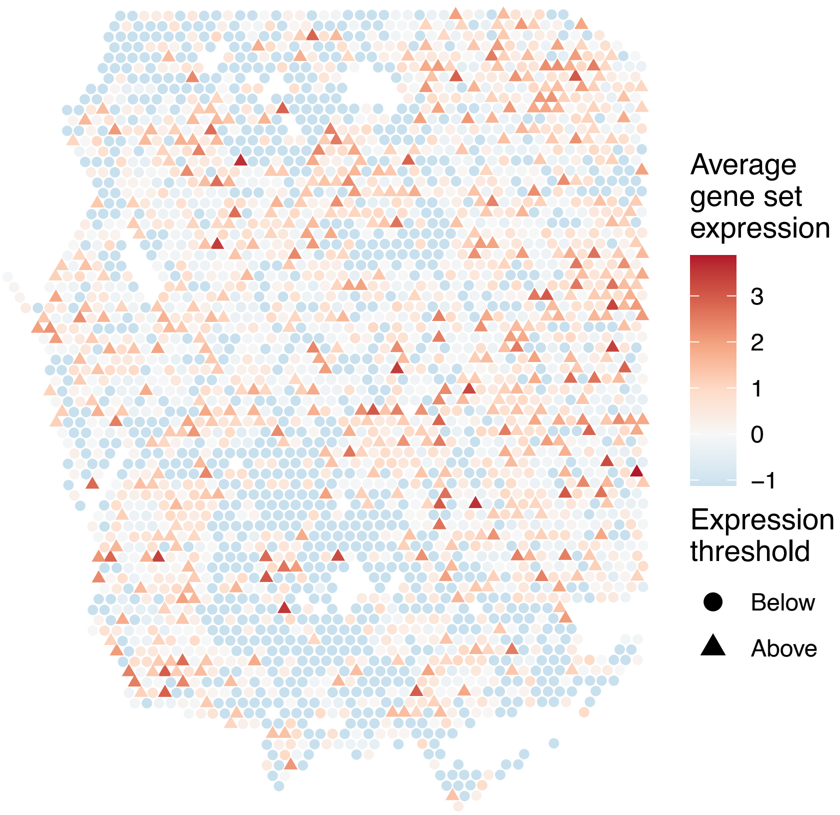


sample16_LMM_A4
KEGG_BASAL_TRANSCRIPTION_FACTORS
Proportion of pathway genes=0.91; Adjusted p−value=0.0979

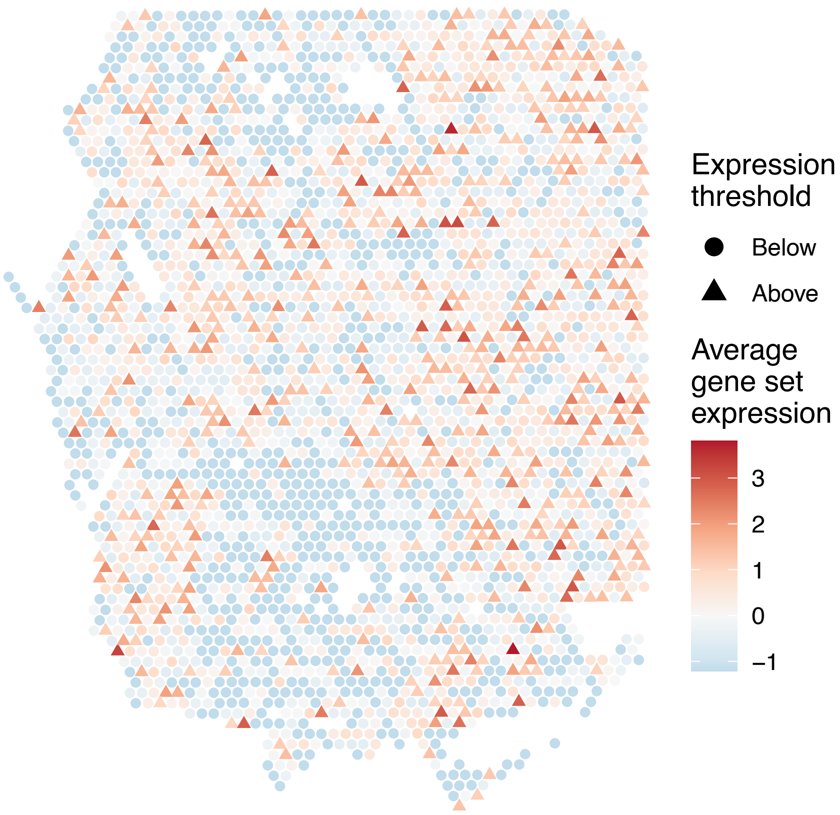


sample16_LMM_A4
KEGG_BASE_EXCISION_REPAIR
Proportion of pathway genes=0.89; Adjusted p−value=0.0731

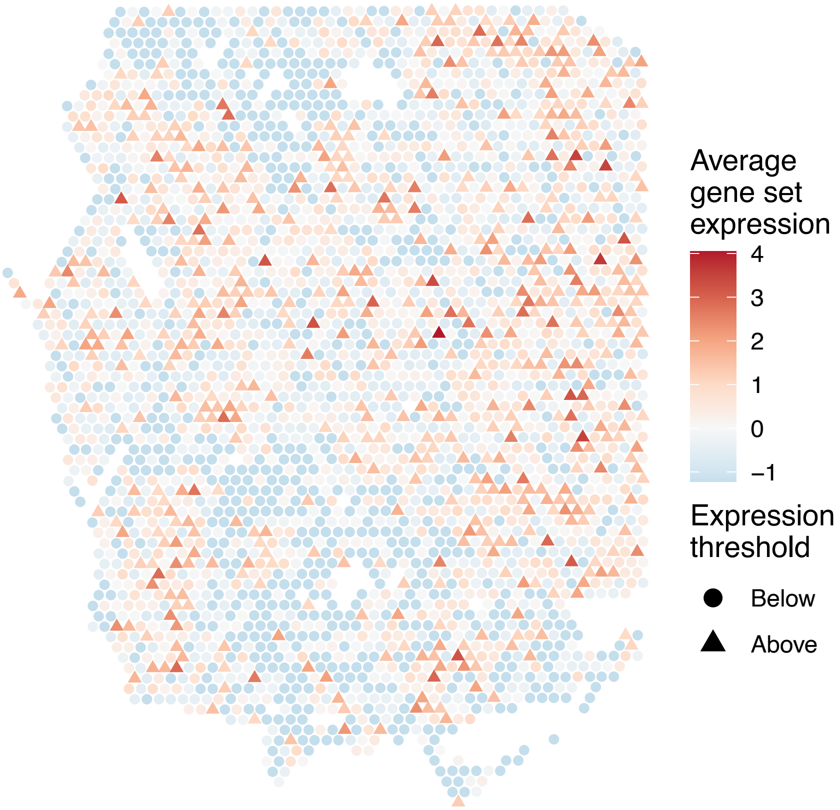


sample16_LMM_A4
KEGG_BETA_ALANINE_METABOLISM
Proportion of pathway genes=0.77; Adjusted p−value=0.4382

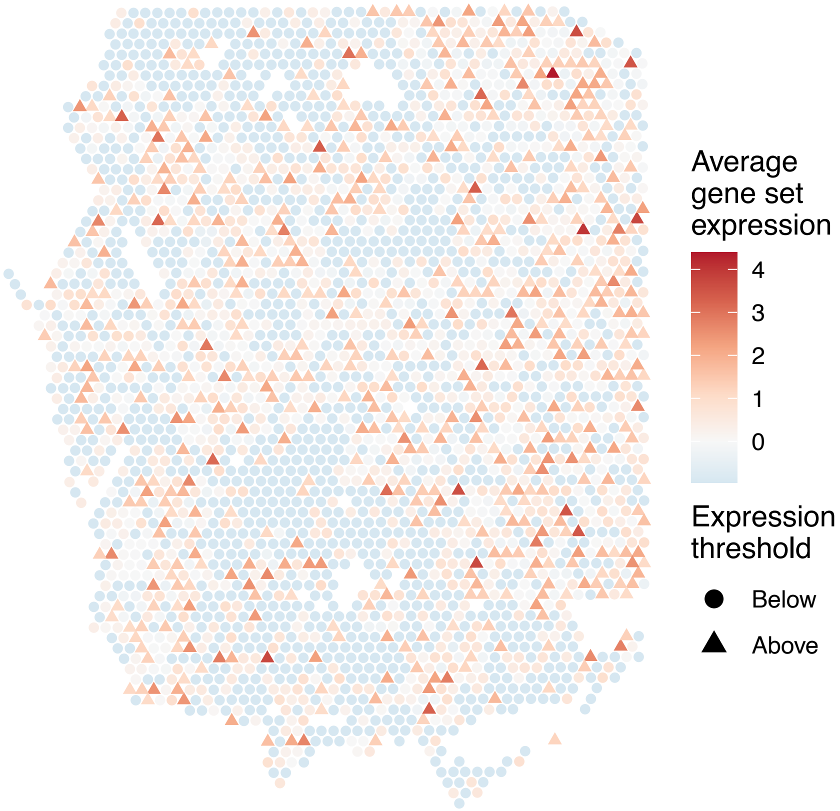

sample16_LMM_A4
KEGG_BIOSYNTHESIS_OF_UNSATURATED_FATTY_ACIDS
Proportion of pathway genes=0.78; Adjusted p−value=0.025

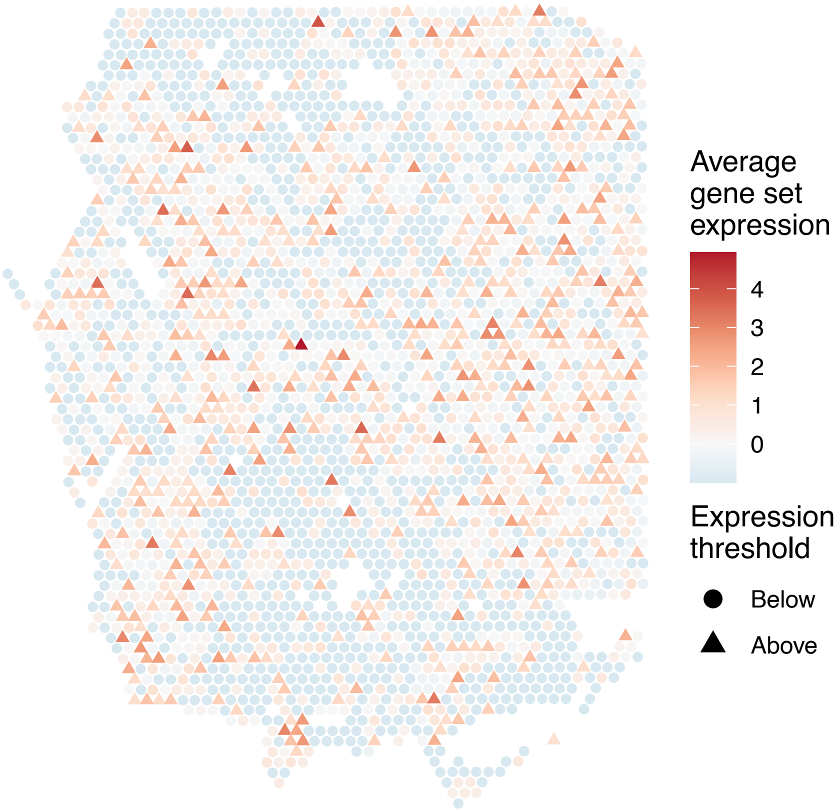


sample16_LMM_A4
KEGG_BLADDER_CANCER
Proportion of pathway genes=0.8; Adjusted p−value=1

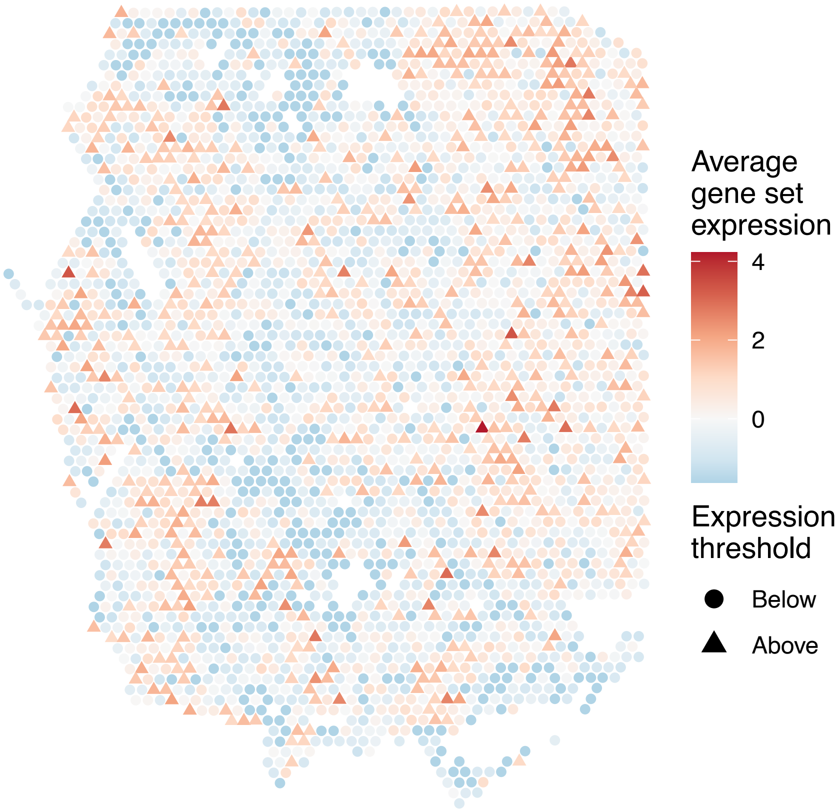


sample16_LMM_A4
KEGG_BUTANOATE_METABOLISM
Proportion of pathway genes=0.74; Adjusted p−value=0.1559

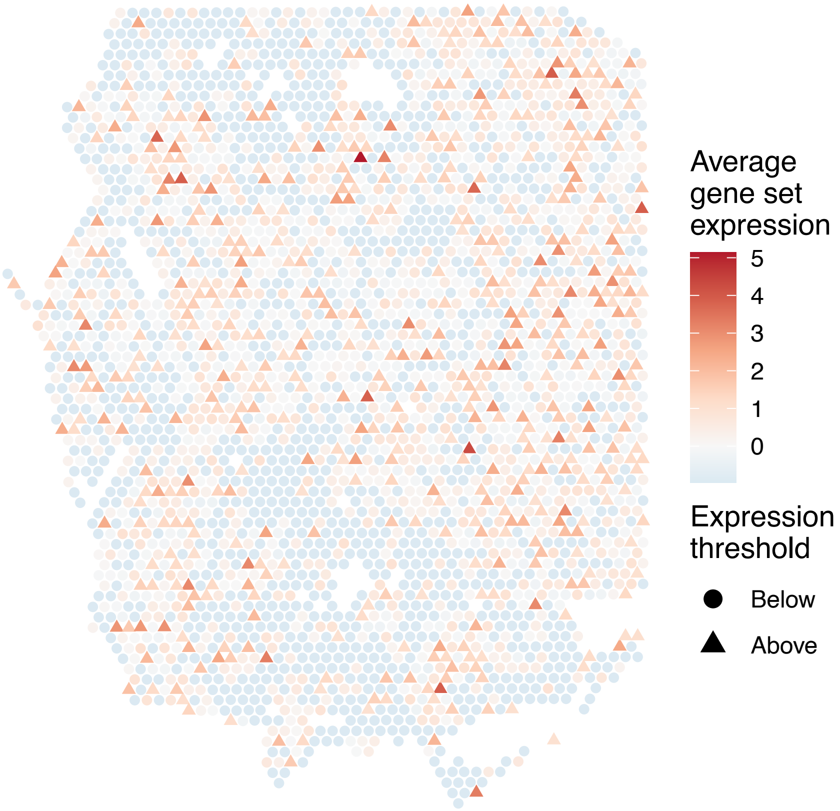


sample16_LMM_A4
KEGG_CALCIUM_SIGNALING_PATHWAY
Proportion of pathway genes=0.46; Adjusted p−value=0.0067

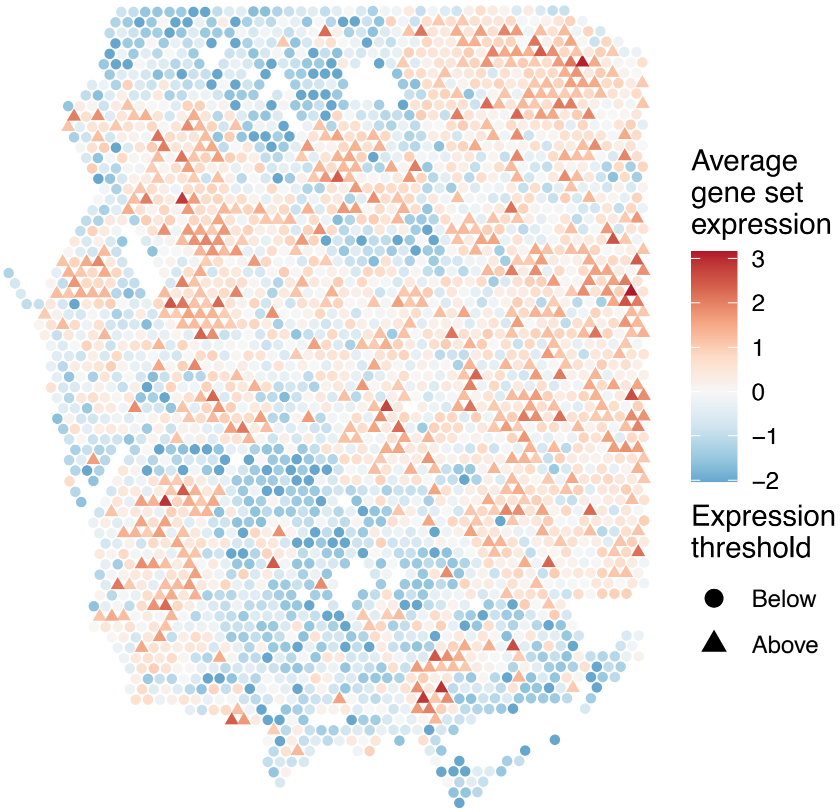

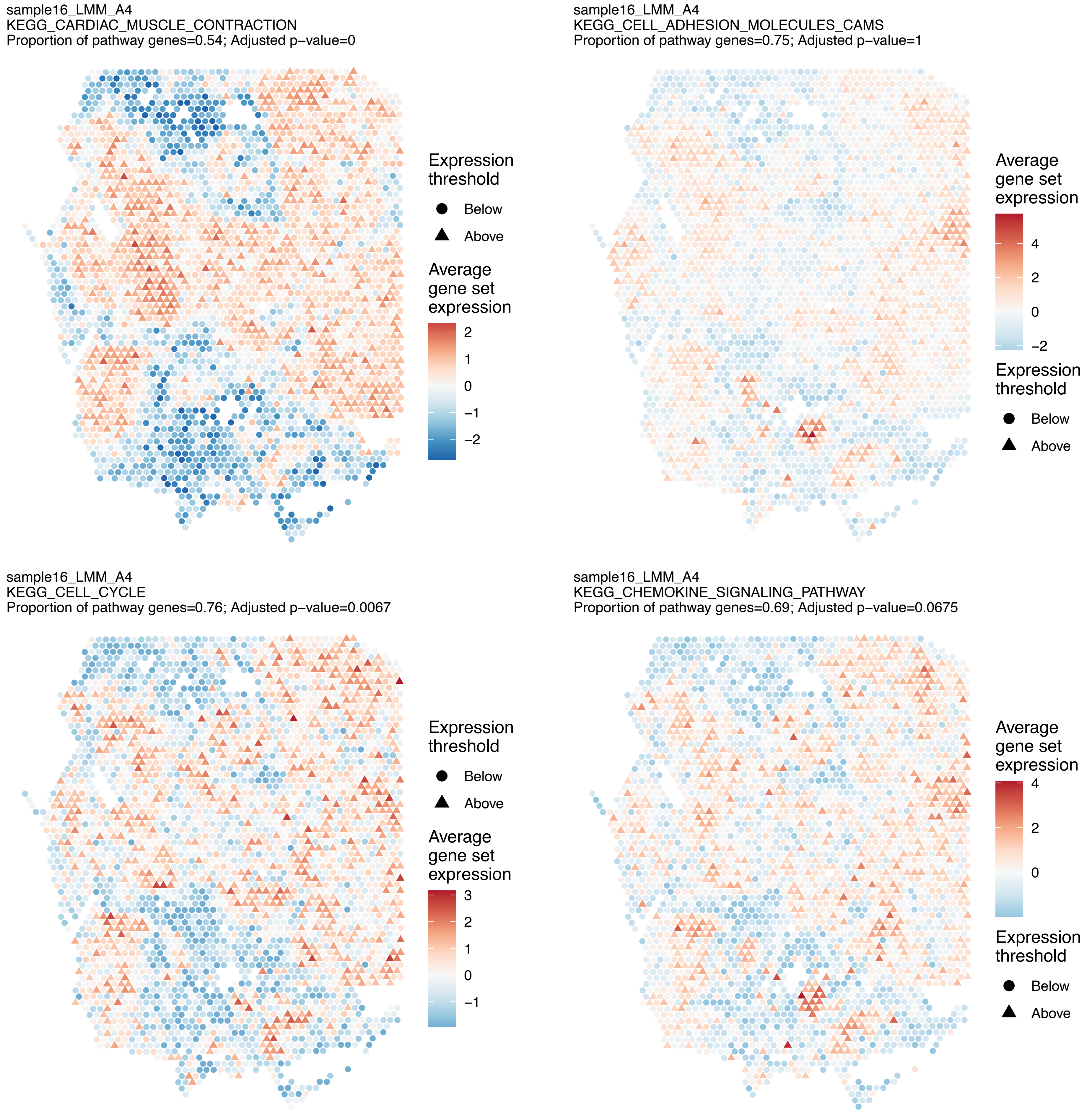
sample16_LMM_A4
KEGG_CARDIAC_MUSCLE_CONTRACTION
Proportion of pathway genes=0.54; Adjusted p−value=0
Expression threshold
Below
Above
Average gene set expression
2
1
0
−1
−2
sample16_LMM_A4
KEGG_CELL_ADHESION_MOLECULES_CAMS
Proportion of pathway genes=0.75; Adjusted p−value=1
Average gene set expression
4
2
0
−2
Expression threshold
Below
Above
sample16_LMM_A4
KEGG_CELL_CYCLE
Proportion of pathway genes=0.76; Adjusted p−value=0.0067
Expression threshold
Below
Above
Average gene set expression
3
2
1
0
−1
sample16_LMM_A4
KEGG_CHEMOKINE_SIGNALING_PATHWAY
Proportion of pathway genes=0.69; Adjusted p−value=0.0675
Average gene set expression
4
2
0
Expression threshold
Below
Above

sample16_LMM_A4
KEGG_CHRONIC_MYELOID_LEUKEMIA
Proportion of pathway genes=0.89; Adjusted p−value=0.1071

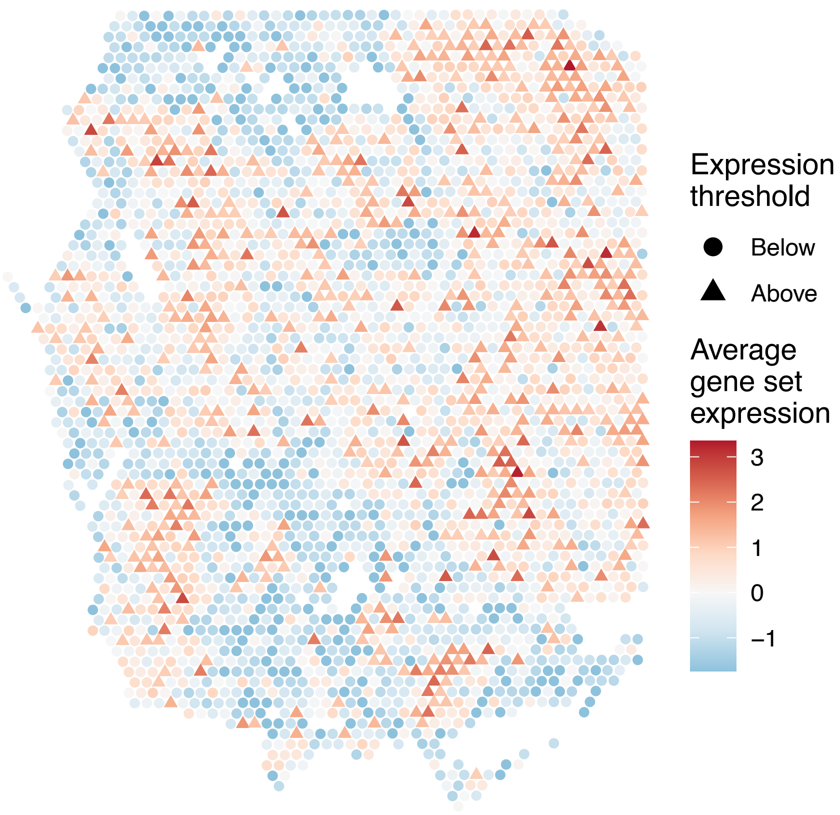


sample16_LMM_A4
KEGG_CIRCADIAN_RHYTHM_MAMMAL
Proportion of pathway genes=1; Adjusted p−value=0.6633

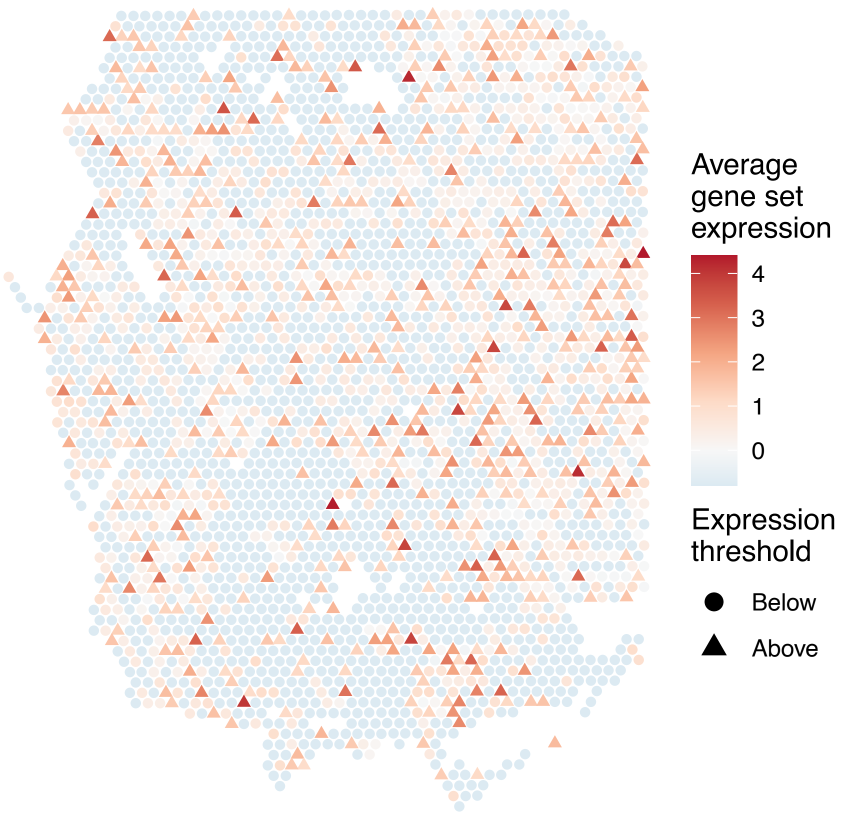


sample16_LMM_A4
KEGG_CITRATE_CYCLE_TCA_CYCLE
Proportion of pathway genes=0.88; Adjusted p−value=0.2211

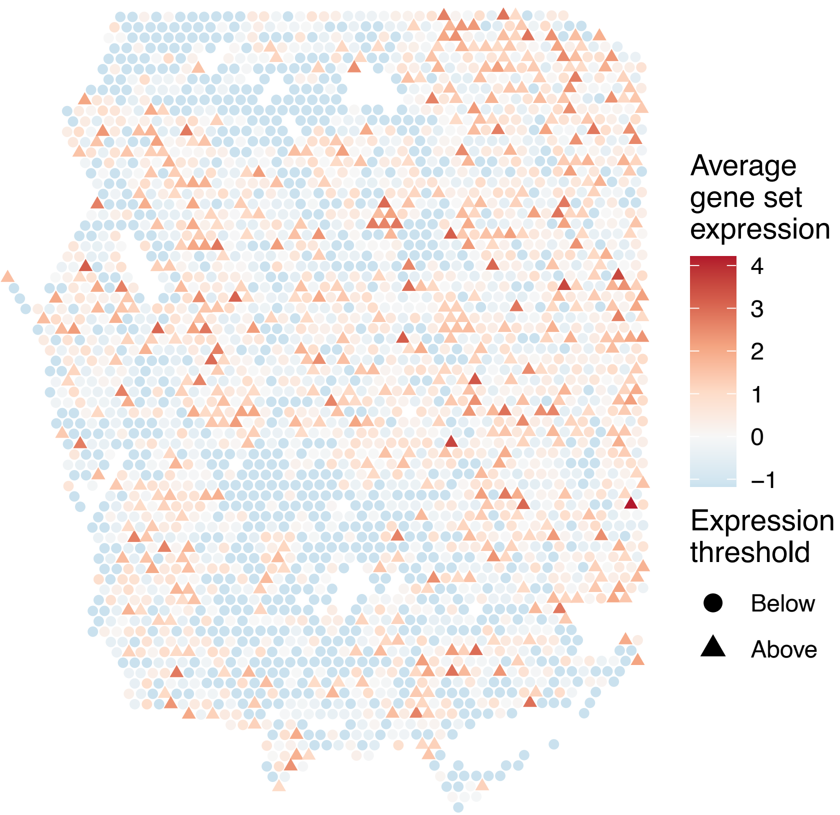


sample16_LMM_A4
KEGG_COLORECTAL_CANCER
Proportion of pathway genes=0.9; Adjusted p−value=0.0106

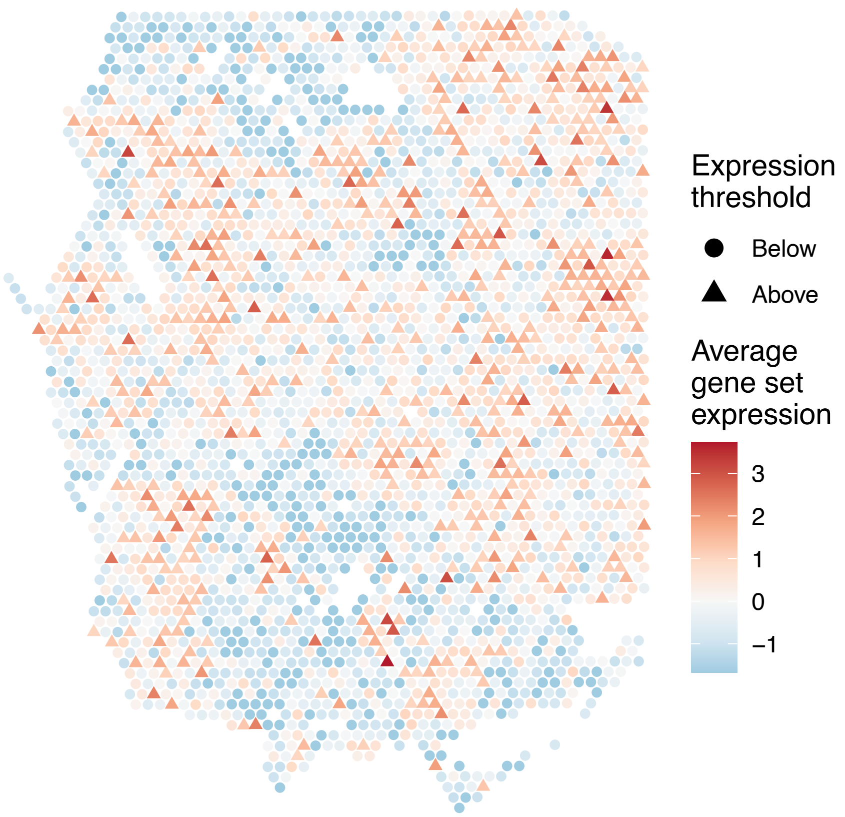

sample16_LMM_A4
KEGG_COMPLEMENT_AND_COAGULATION_CASCADES
Proportion of pathway genes=0.55; Adjusted p−value=0.7778

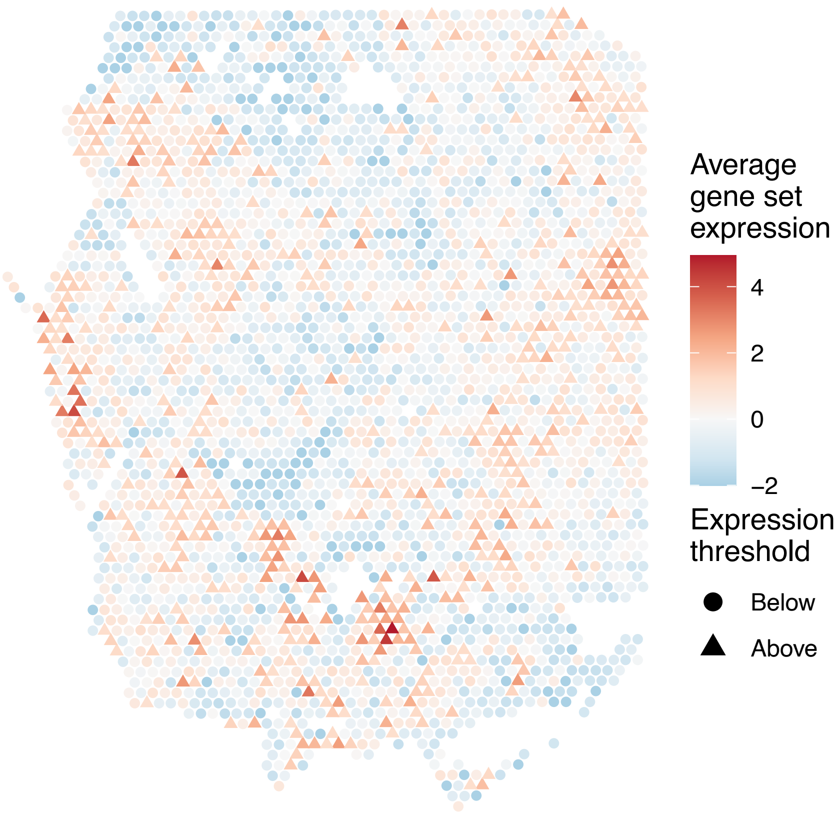


sample16_LMM_A4
KEGG_CYSTEINE_AND_METHIONINE_METABOLISM
Proportion of pathway genes=0.69; Adjusted p−value=0.1485

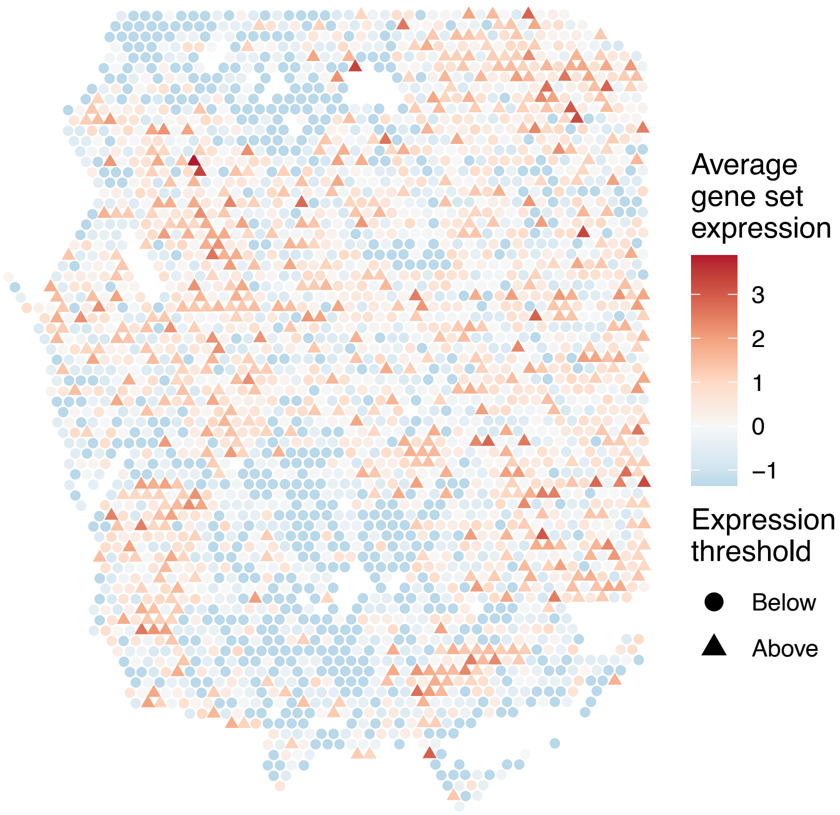


sample16_LMM_A4
KEGG_CYTOSOLIC_DNA_SENSING_PATHWAY
Proportion of pathway genes=0.65; Adjusted p−value=0.9479

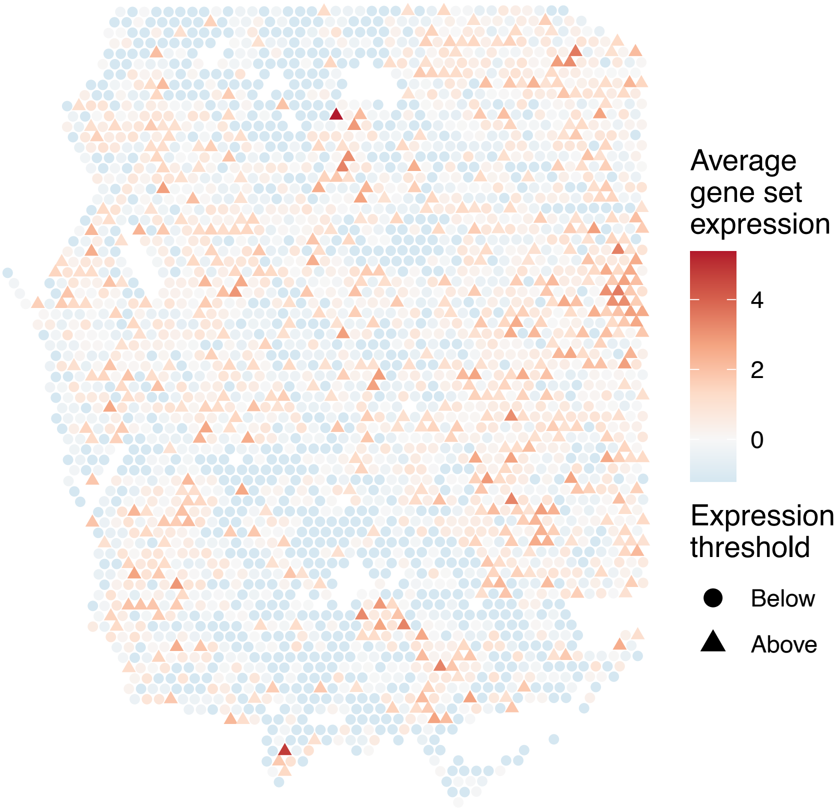


sample16_LMM_A4
KEGG_DILATED_CARDIOMYOPATHY
Proportion of pathway genes=0.51; Adjusted p−value=0.0528

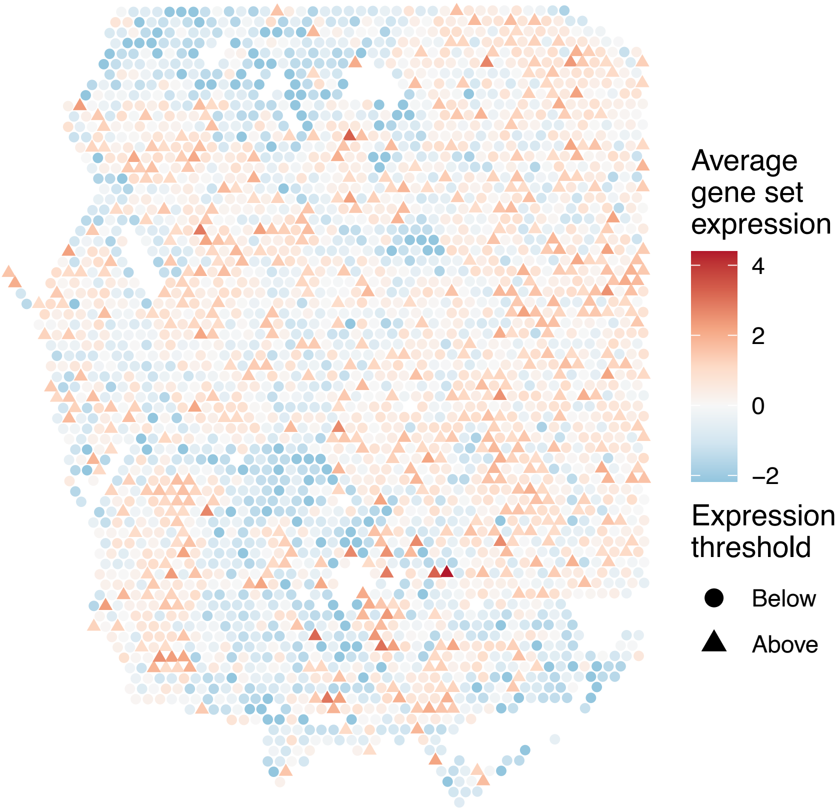

sample16_LMM_A4
KEGG_DNA_REPLICATION
Proportion of pathway genes=0.92; Adjusted p−value=0.2524

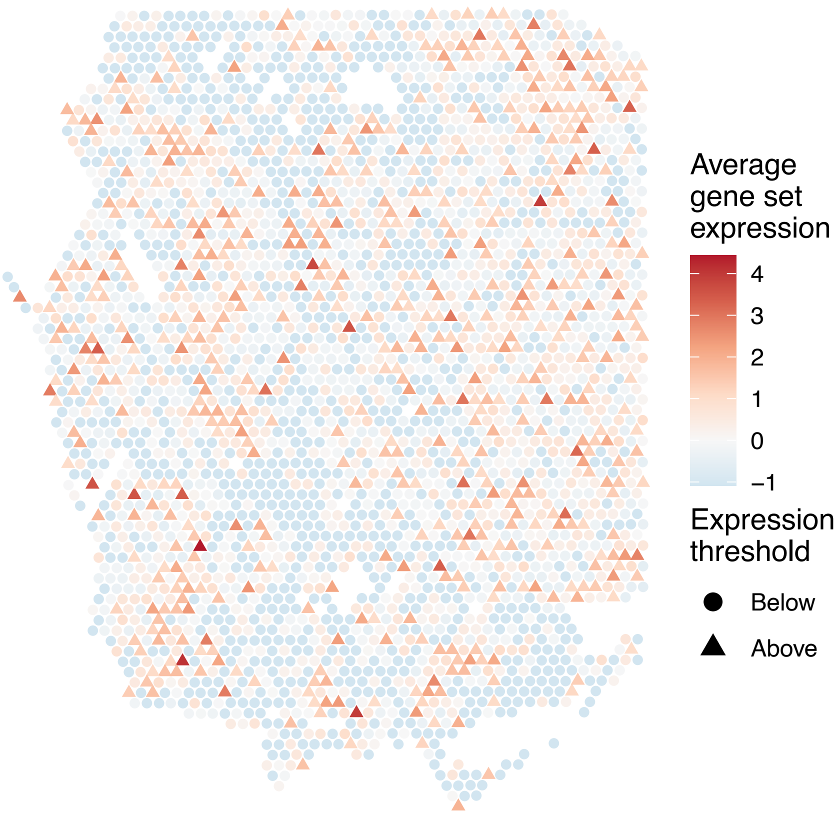


sample16_LMM_A4
KEGG_DORSO_VENTRAL_AXIS_FORMATION
Proportion of pathway genes=0.84; Adjusted p−value=0.2524

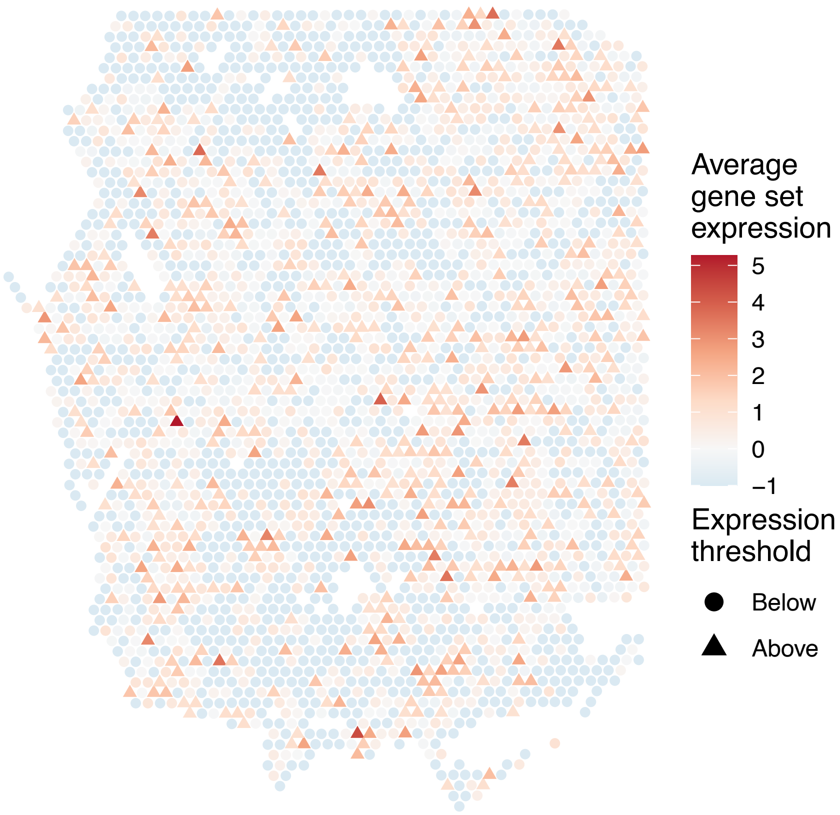


sample16_LMM_A4
KEGG_DRUG_METABOLISM_CYTOCHROME_P450
Proportion of pathway genes=0.21; Adjusted p−value=0.2693

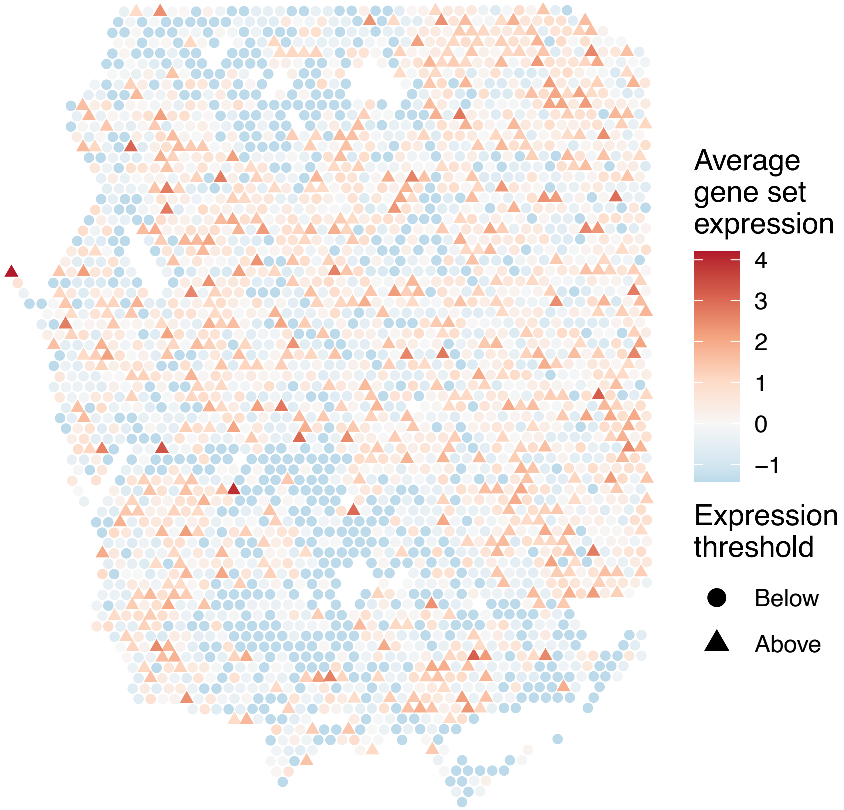


sample16_LMM_A4
KEGG_DRUG_METABOLISM_OTHER_ENZYMES
Proportion of pathway genes=0.32; Adjusted p−value=0.4382

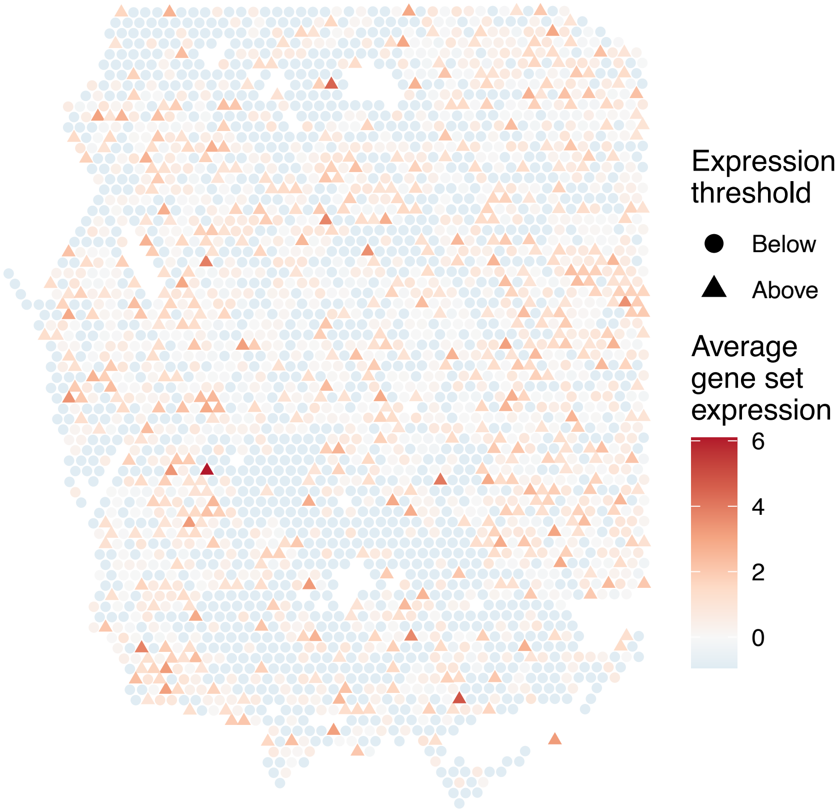

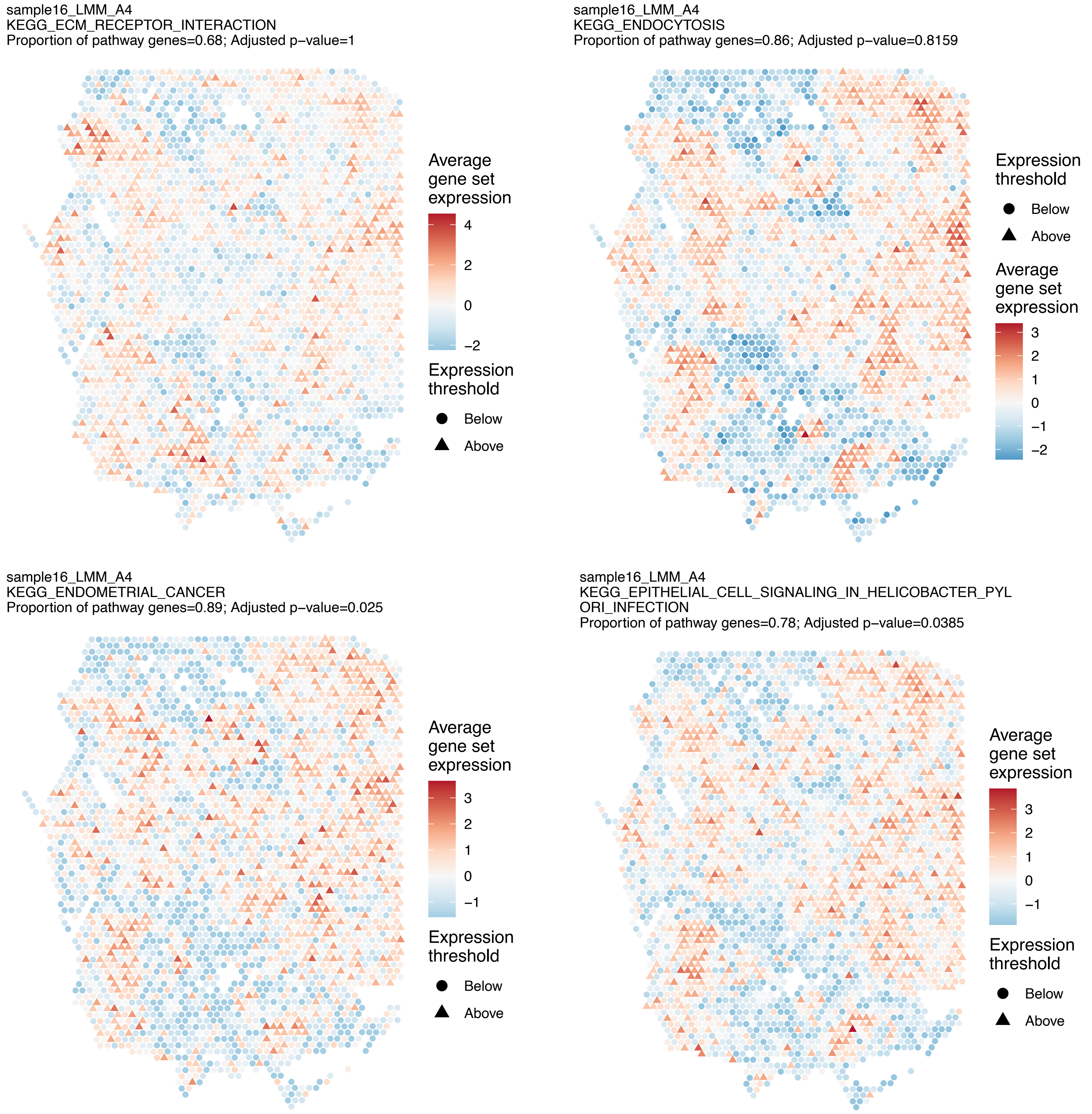
sample16_LMM_A4
KEGG_ECM_RECEPTOR_INTERACTION
Proportion of pathway genes=0.68; Adjusted p−value=1
Average gene set expression
4
2
0
−2
Expression threshold
Below
Above
sample16_LMM_A4
KEGG_ENDOCYTOSIS
Proportion of pathway genes=0.86; Adjusted p−value=0.8159
Expression threshold
Below
Above
Average gene set expression
3
2
1
0
−1
−2
sample16_LMM_A4
KEGG_ENDOMETRIAL_CANCER
Proportion of pathway genes=0.89; Adjusted p−value=0.025
Average gene set expression
3
2
1
0
−1
Expression threshold
Below
Above
sample16_LMM_A4
KEGG_EPITHELIAL_CELL_SIGNALING_IN_HELICOBACTER_PYL
ORI_INFECTION
Proportion of pathway genes=0.78; Adjusted p−value=0.0385
Average gene set expression
3
2
1
0
−1
Expression threshold
Below
Above

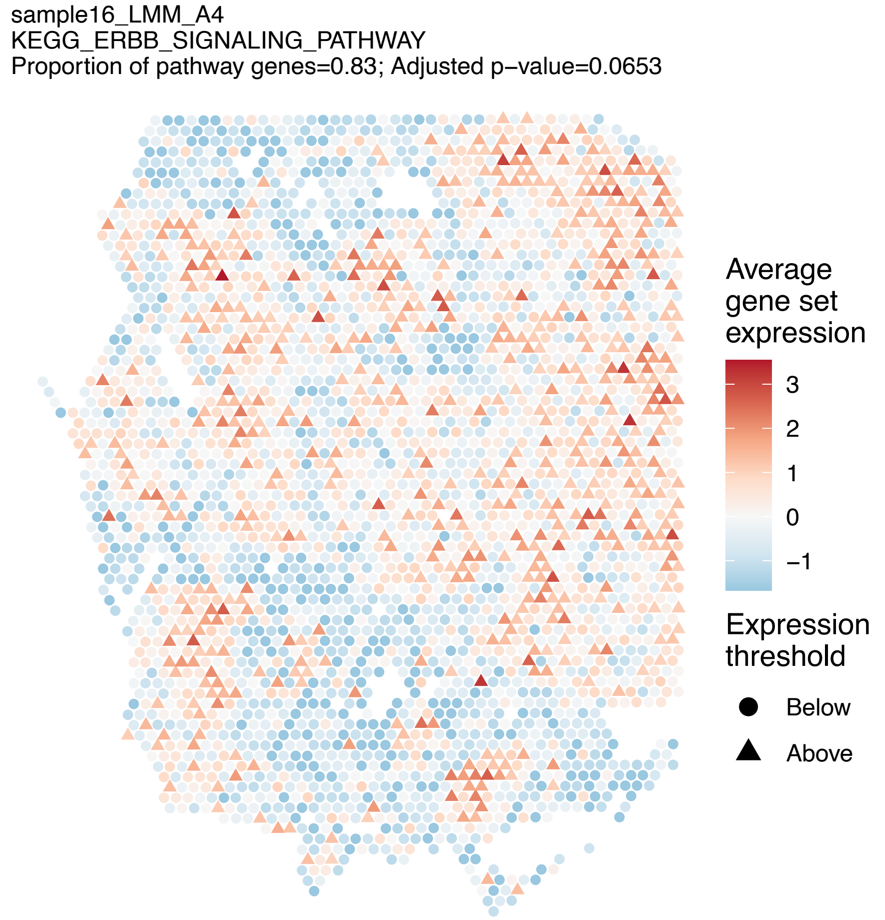
sample16_LMM_A4
KEGG_ERBB_SIGNALING_PATHWAY
Proportion of pathway genes=0.83; Adjusted p−value=0.0653
Average gene set expression
3
2
1
0
−1
Expression threshold
Below
Above

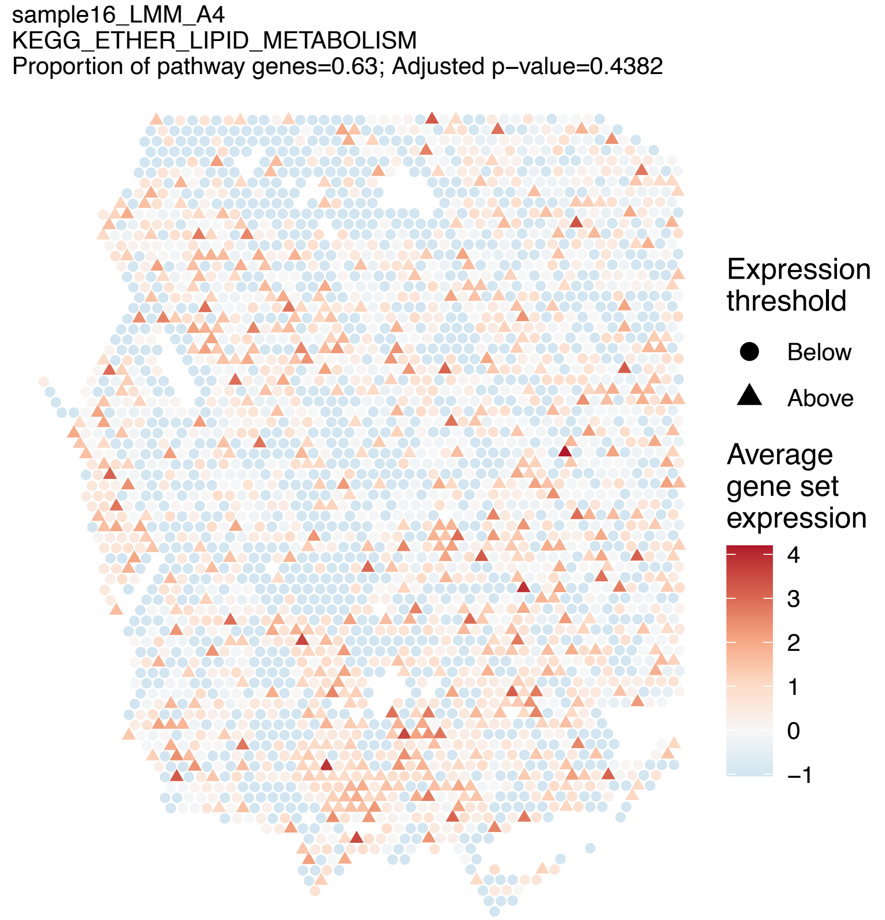
sample16_LMM_A4
KEGG_ETHER_LIPID_METABOLISM
Proportion of pathway genes=0.63; Adjusted p−value=0.4382
Expression threshold
Below
Above
Average gene set expression
4
3
2
1
0
−1

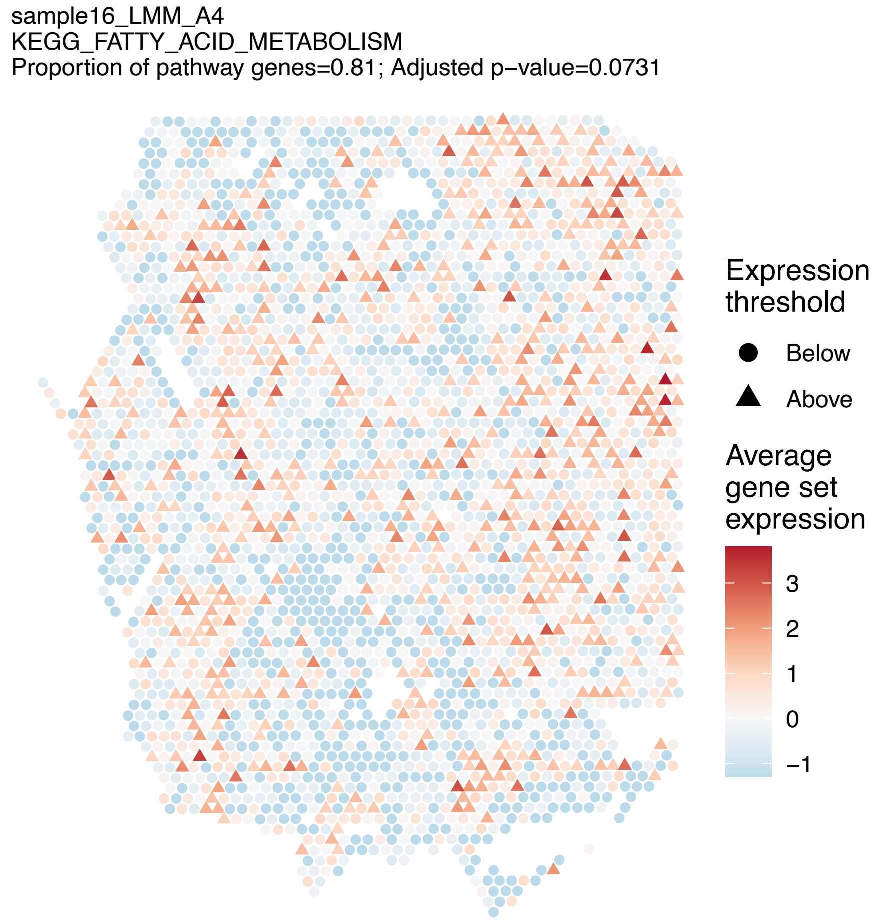
sample16_LMM_A4
KEGG_FATTY_ACID_METABOLISM
Proportion of pathway genes=0.81; Adjusted p−value=0.0731
Expression threshold
Below
Above
Average gene set expression
3
2
1
0
−1

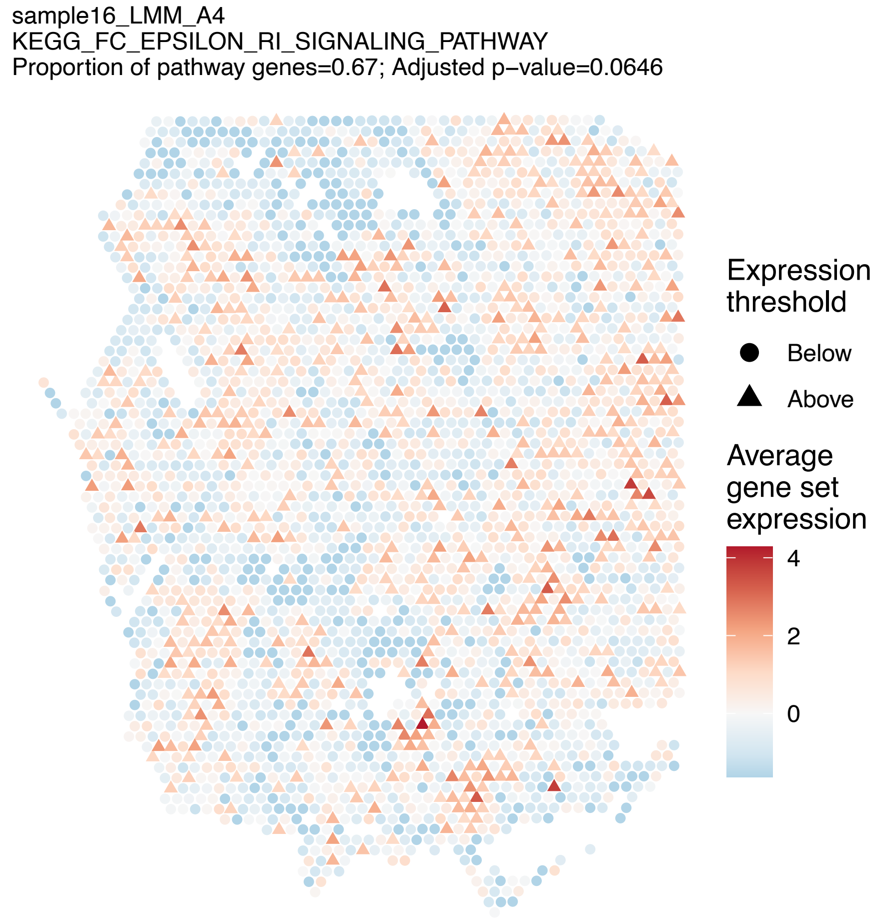
sample16_LMM_A4
KEGG_FC_EPSILON_RI_SIGNALING_PATHWAY
Proportion of pathway genes=0.67; Adjusted p−value=0.0646
Expression threshold
Below
Above
Average gene set expression
4
2
0

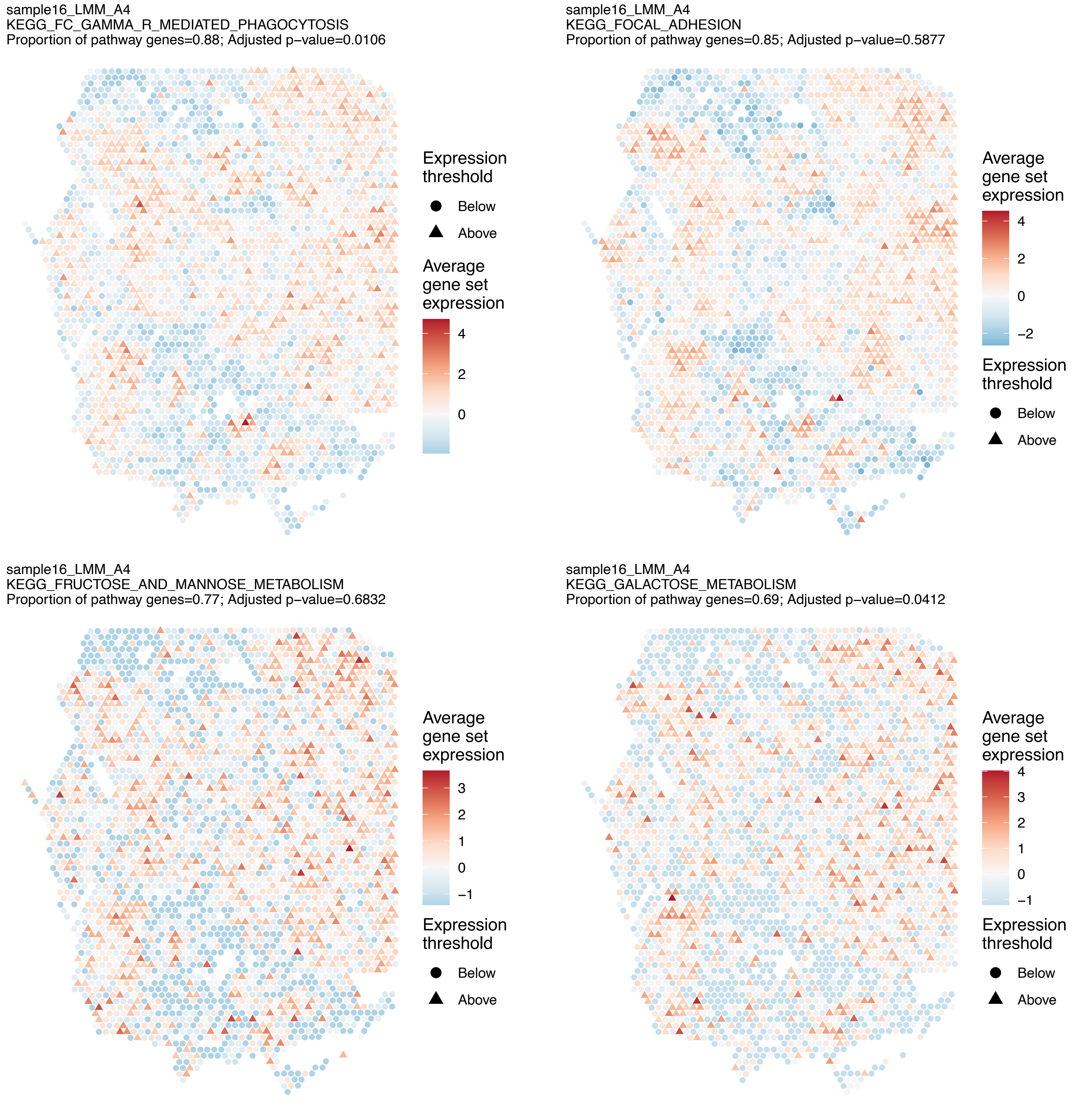
sample16_LMM_A4
KEGG_FC_GAMMA_R_MEDIATED_PHAGOCYTOSIS
Proportion of pathway genes=0.88; Adjusted p−value=0.0106
Expression threshold
Below
Above
Average gene set expression
4
2
0
sample16_LMM_A4
KEGG_FOCAL_ADHESION
Proportion of pathway genes=0.85; Adjusted p−value=0.5877
Average gene set expression
4
2
0
−2
Expression threshold
Below
Above
sample16_LMM_A4
KEGG_FRUCTOSE_AND_MANNOSE_METABOLISM
Proportion of pathway genes=0.77; Adjusted p−value=0.6832
Average gene set expression
3
2
1
0
−1
Expression threshold
Below
Above
sample16_LMM_A4
KEGG_GALACTOSE_METABOLISM
Proportion of pathway genes=0.69; Adjusted p−value=0.0412
Average gene set expression
4
3
2
1
0
−1
Expression threshold
Below
Above

sample16_LMM_A4
KEGG_GAP_JUNCTION
Proportion of pathway genes=0.7; Adjusted p−value=0.0695

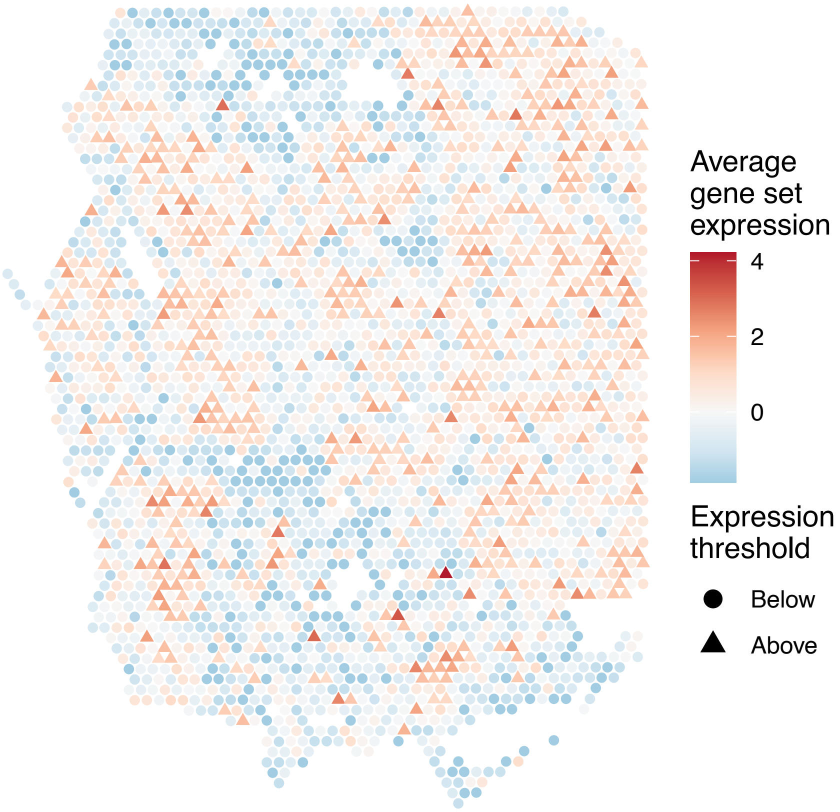


sample16_LMM_A4
KEGG_GLIOMA
Proportion of pathway genes=0.81; Adjusted p−value=0.0067

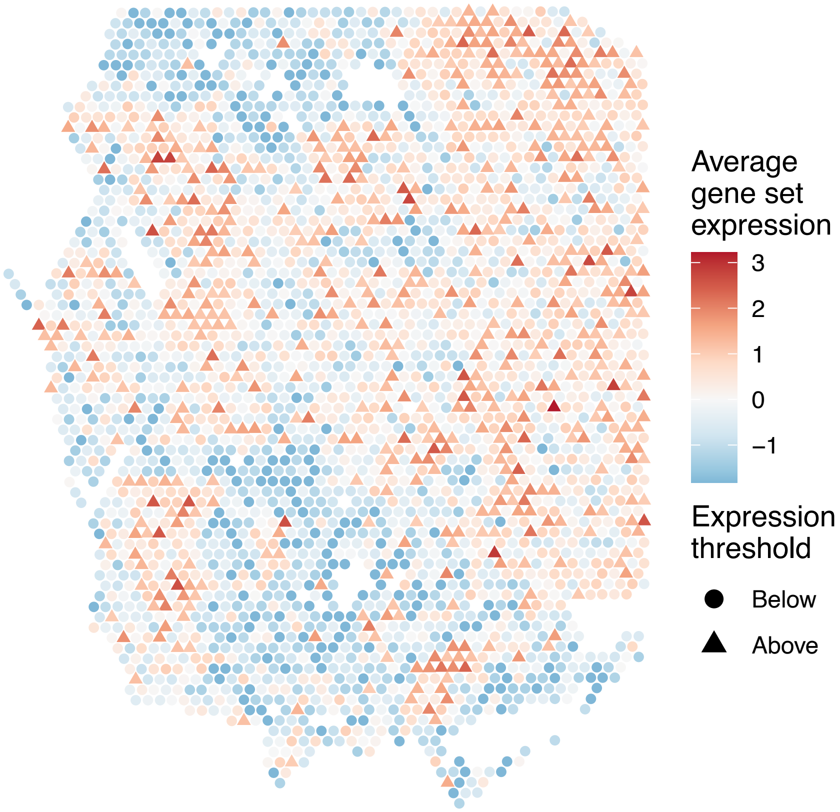


sample16_LMM_A4
KEGG_GLUTATHIONE_METABOLISM
Proportion of pathway genes=0.64; Adjusted p−value=0.6484

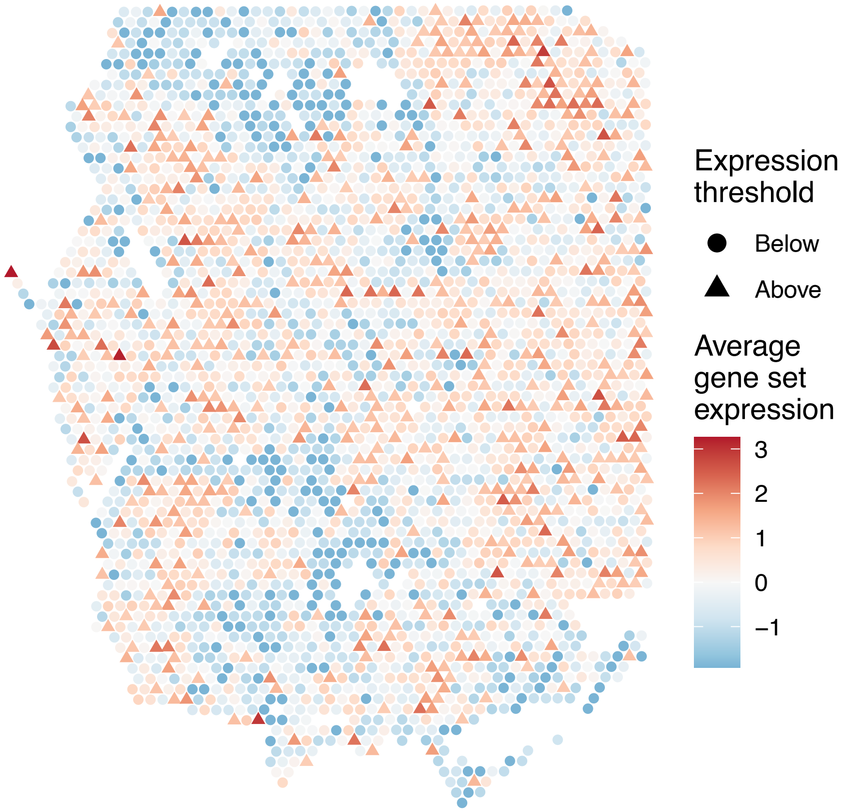


sample16_LMM_A4
KEGG_GLYCEROLIPID_METABOLISM
Proportion of pathway genes=0.78; Adjusted p−value=0.65

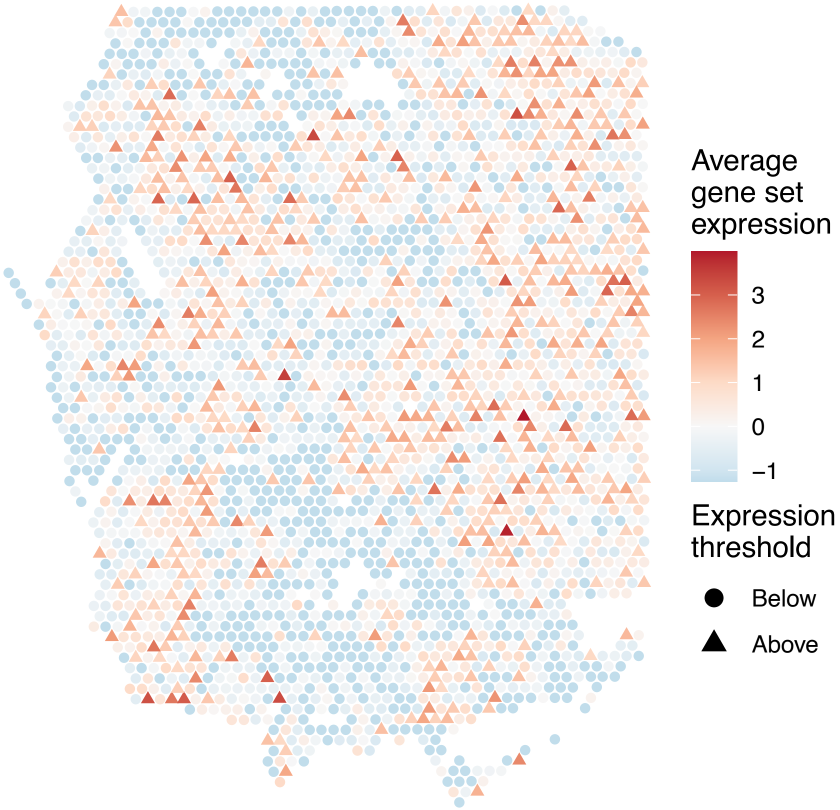

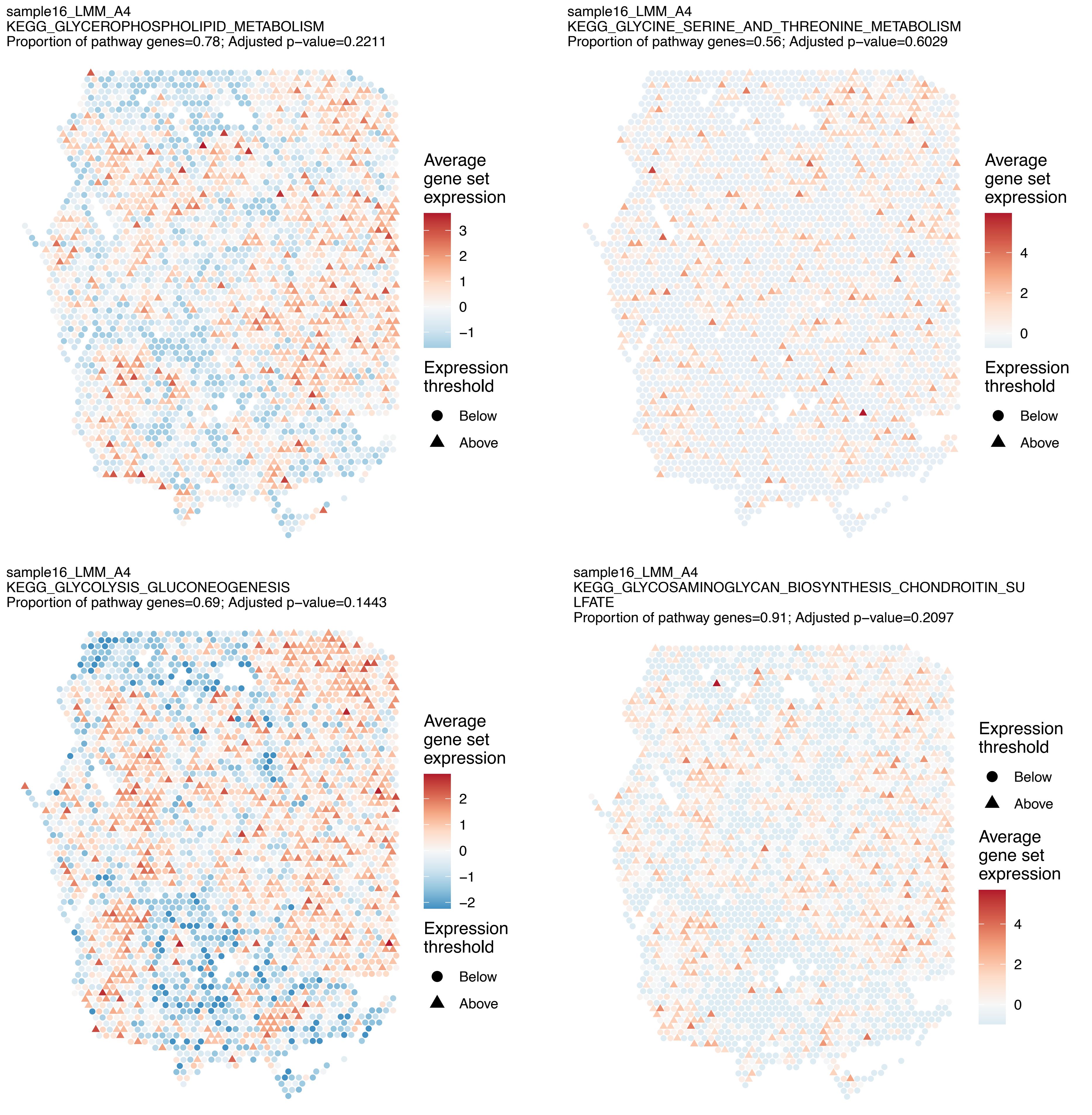
sample16_LMM_A4
KEGG_GLYCEROPHOSPHOLIPID_METABOLISM
Proportion of pathway genes=0.78; Adjusted p−value=0.2211
Average gene set expression
3
2
1
0
−1
Expression threshold
Below
Above
sample16_LMM_A4
KEGG_GLYCINE_SERINE_AND_THREONINE_METABOLISM
Proportion of pathway genes=0.56; Adjusted p−value=0.6029
Average gene set expression
4
2
0
Expression threshold
Below
Above
sample16_LMM_A4
KEGG_GLYCOLYSIS_GLUCONEOGENESIS
Proportion of pathway genes=0.69; Adjusted p−value=0.1443
Average gene set expression
2
1
0
−1
−2
Expression threshold
Below
Above
sample16_LMM_A4
KEGG_GLYCOSAMINOGLYCAN_BIOSYNTHESIS_CHONDROITIN_SU
LFATE
Proportion of pathway genes=0.91; Adjusted p−value=0.2097
Expression threshold
Below
Above
Average gene set expression
4
2
0

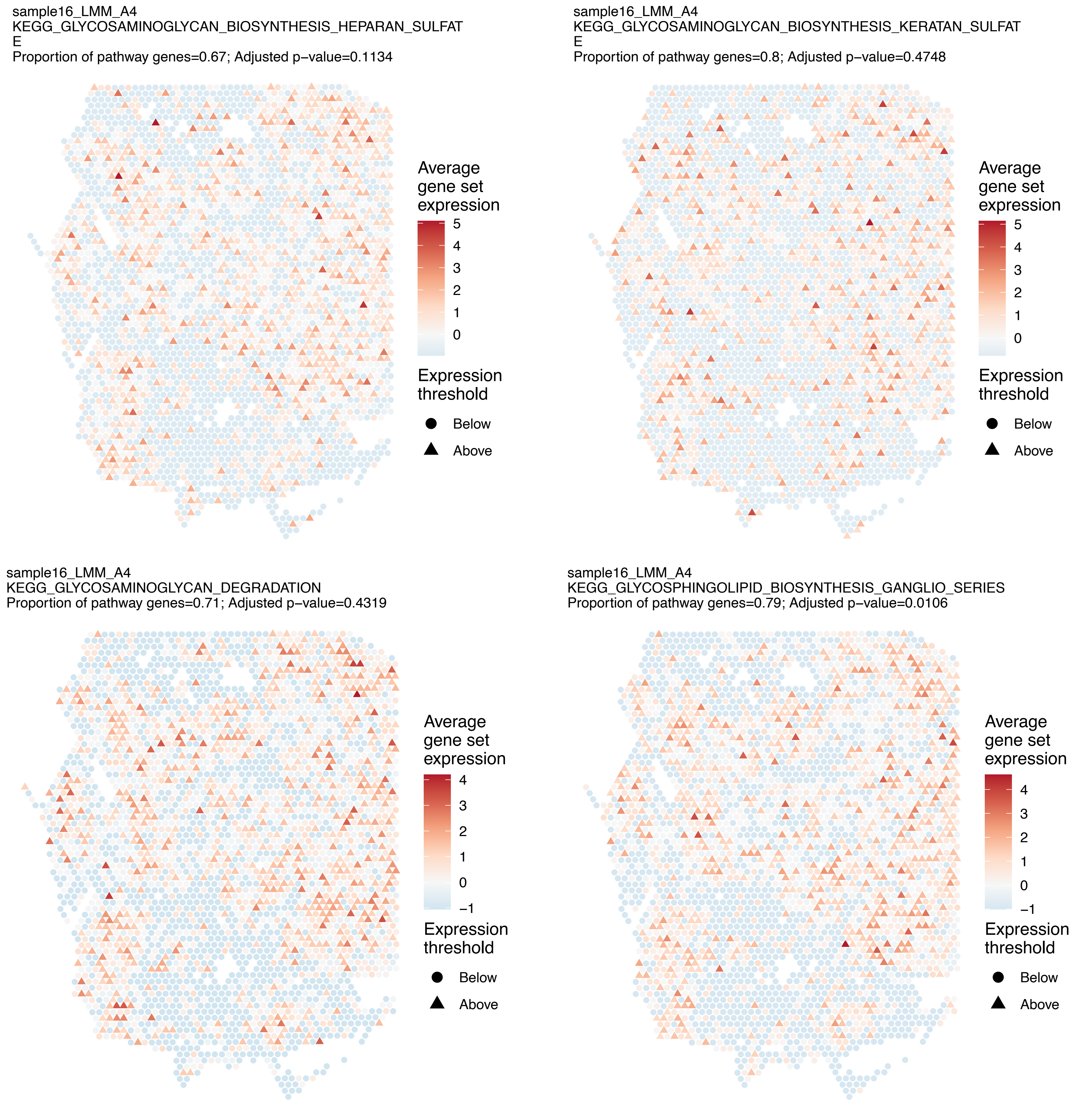
sample16_LMM_A4
KEGG_GLYCOSAMINOGLYCAN_BIOSYNTHESIS_HEPARAN_SULFAT
E
Proportion of pathway genes=0.67; Adjusted p−value=0.1134
Average gene set expression
5
4
3
2
1
0
Expression threshold
Below
Above
sample16_LMM_A4
KEGG_GLYCOSAMINOGLYCAN_BIOSYNTHESIS_KERATAN_SULFAT
E
Proportion of pathway genes=0.8; Adjusted p−value=0.4748
Average gene set expression
5
4
3
2
1
0
Expression threshold
Below
Above
sample16_LMM_A4
KEGG_GLYCOSAMINOGLYCAN_DEGRADATION
Proportion of pathway genes=0.71; Adjusted p−value=0.4319
Average gene set expression
4
3
2
1
0
−1
Expression threshold
Below
Above
sample16_LMM_A4
KEGG_GLYCOSPHINGOLIPID_BIOSYNTHESIS_GANGLIO_SERIES
Proportion of pathway genes=0.79; Adjusted p−value=0.0106
Average gene set expression
4
3
2
1
0
−1
Expression threshold
Below
Above

sample16_LMM_A4
KEGG_GLYCOSPHINGOLIPID_BIOSYNTHESIS_GLOBO_SERIES
Proportion of pathway genes=0.79; Adjusted p−value=0.0146

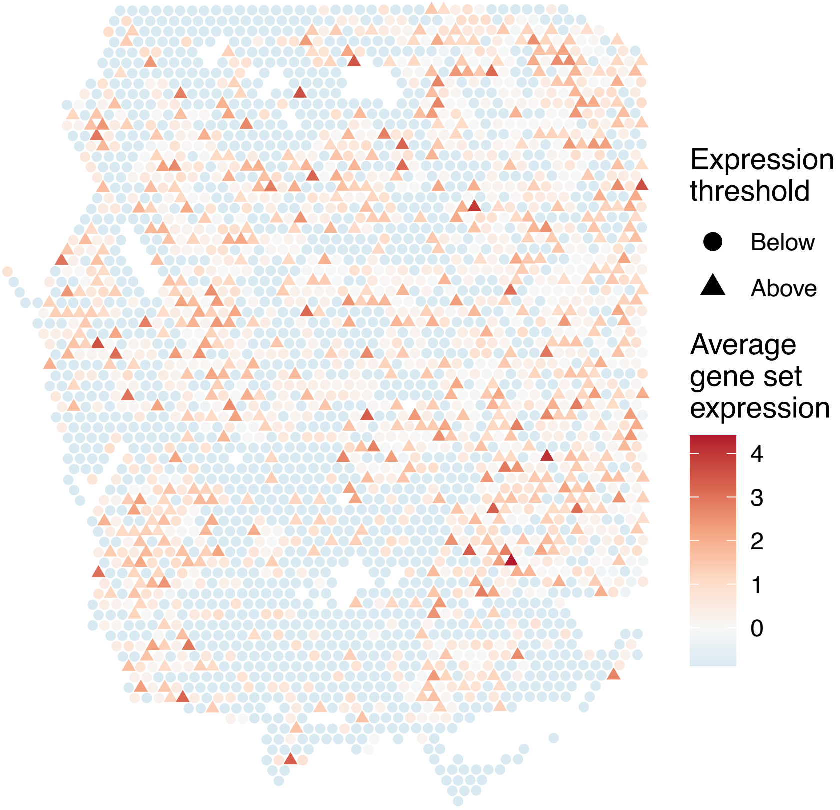


sample16_LMM_A4
KEGG_GLYCOSPHINGOLIPID_BIOSYNTHESIS_LACTO_AND_NEOL
ACTO_SERIES
Proportion of pathway genes=0.55; Adjusted p−value=0.0574

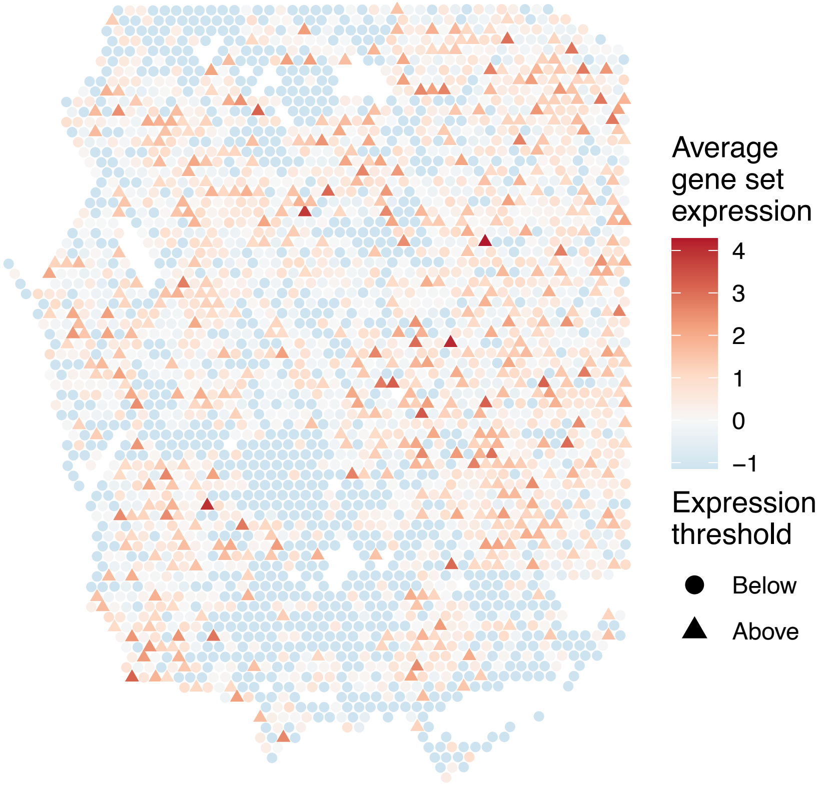


sample16_LMM_A4
KEGG_GLYCOSYLPHOSPHATIDYLINOSITOL_GPI_ANCHOR_BIOSY
NTHESIS
Proportion of pathway genes=0.89; Adjusted p−value=0.227

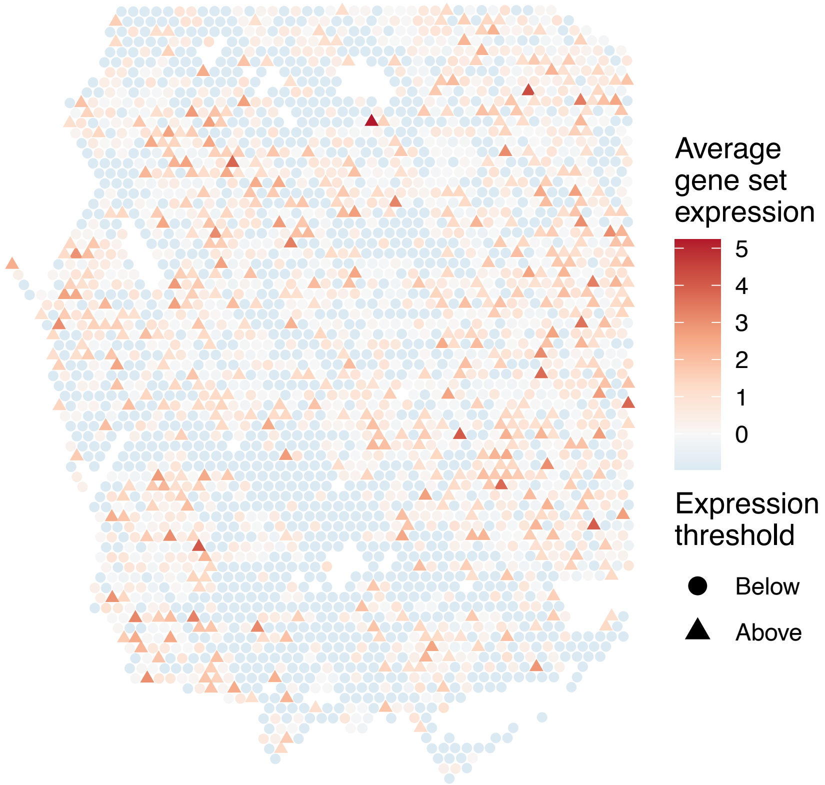


sample16_LMM_A4
KEGG_GLYOXYLATE_AND_DICARBOXYLATE_METABOLISM
Proportion of pathway genes=0.81; Adjusted p−value=0.0531

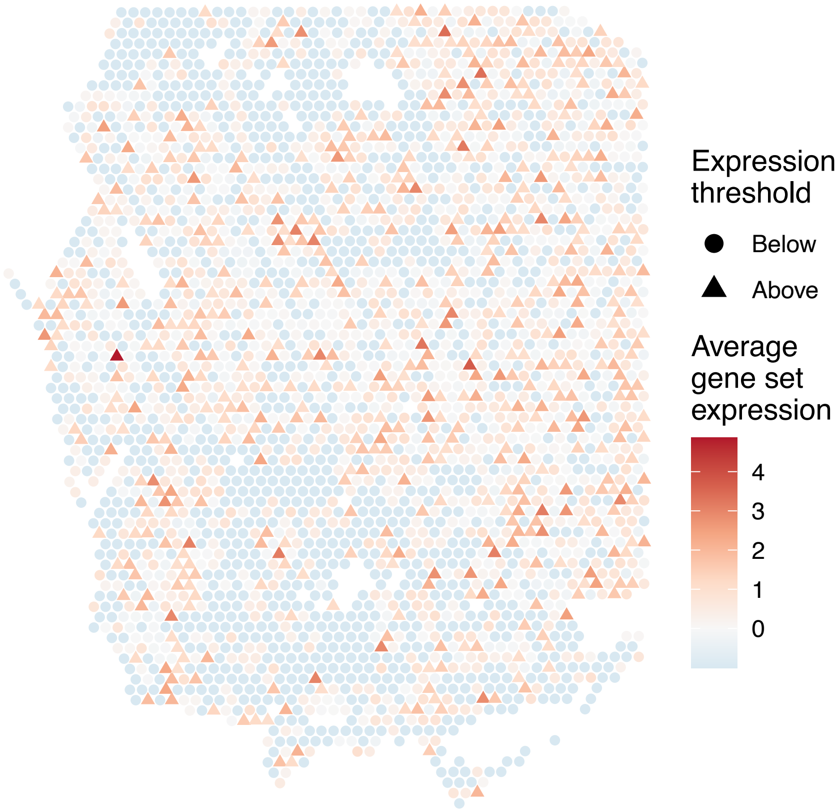

sample16_LMM_A4
KEGG_GNRH_SIGNALING_PATHWAY
Proportion of pathway genes=0.7; Adjusted p−value=0.5526

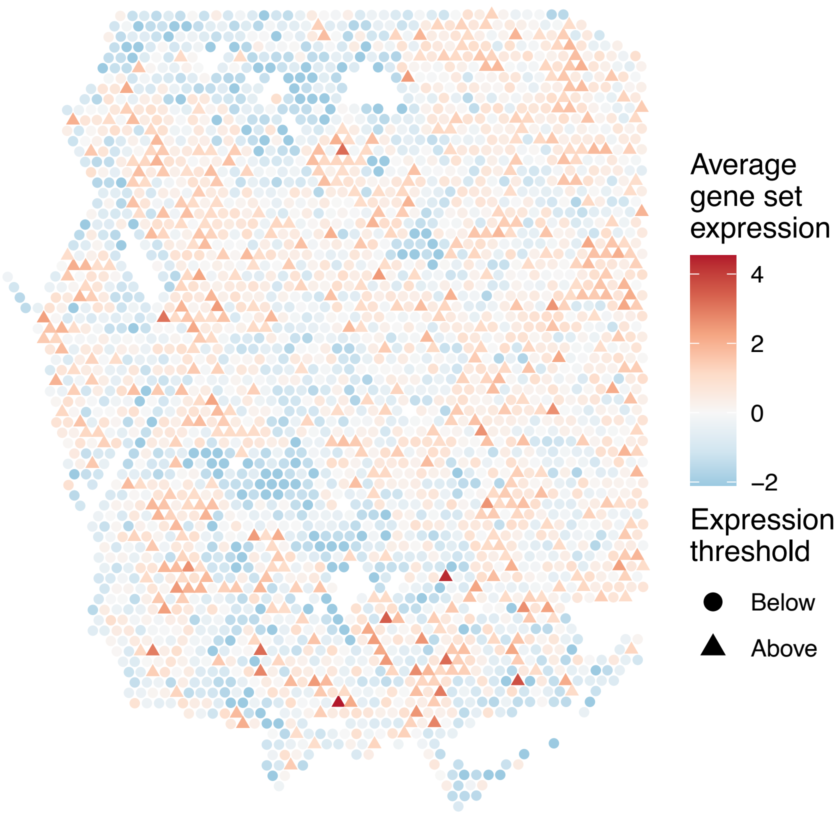


sample16_LMM_A4
KEGG_GRAFT_VERSUS_HOST_DISEASE
Proportion of pathway genes=0.37; Adjusted p−value=1

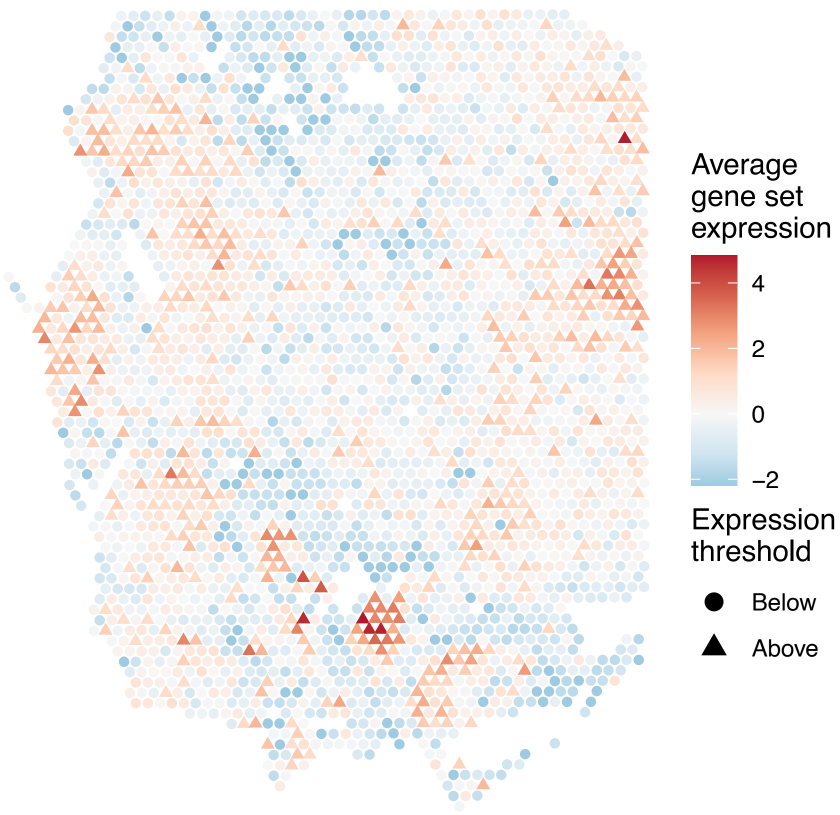


sample16_LMM_A4
KEGG_HEDGEHOG_SIGNALING_PATHWAY
Proportion of pathway genes=0.53; Adjusted p−value=0.0744

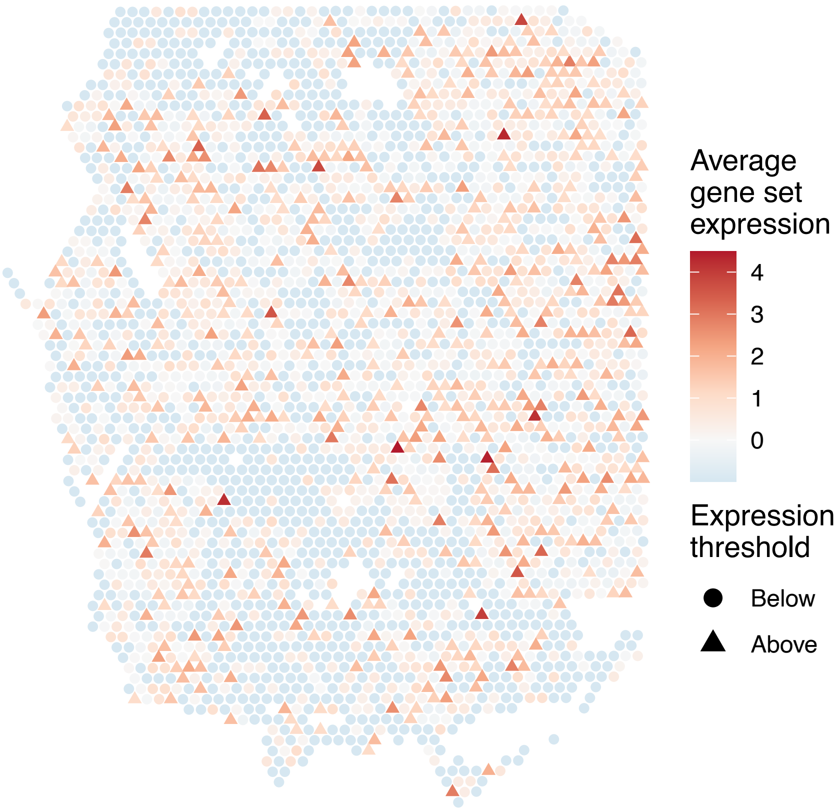


sample16_LMM_A4
KEGG_HEMATOPOIETIC_CELL_LINEAGE
Proportion of pathway genes=0.57; Adjusted p−value=1

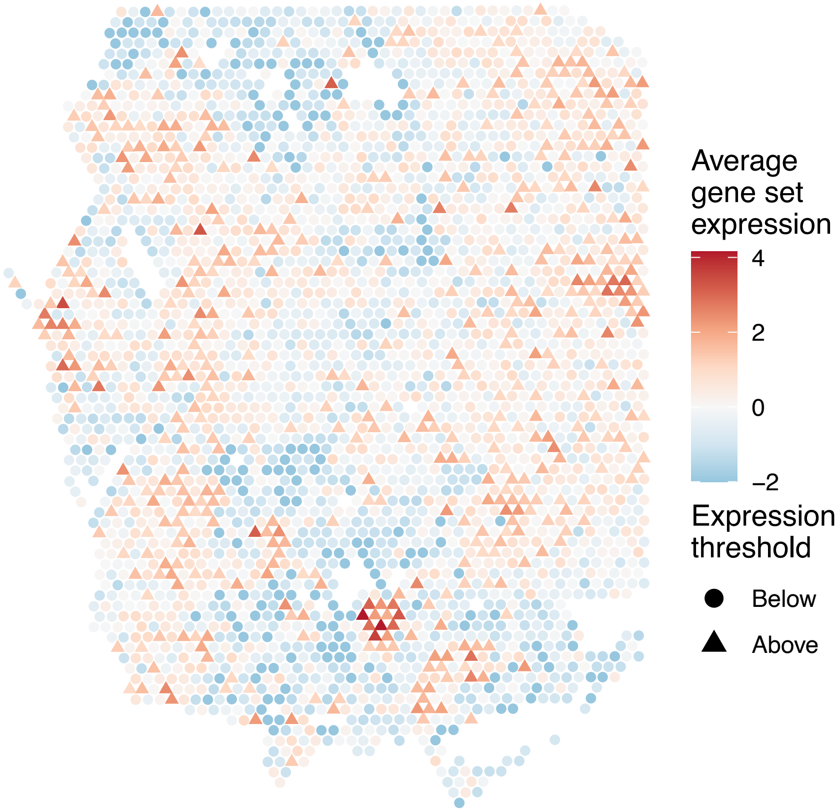

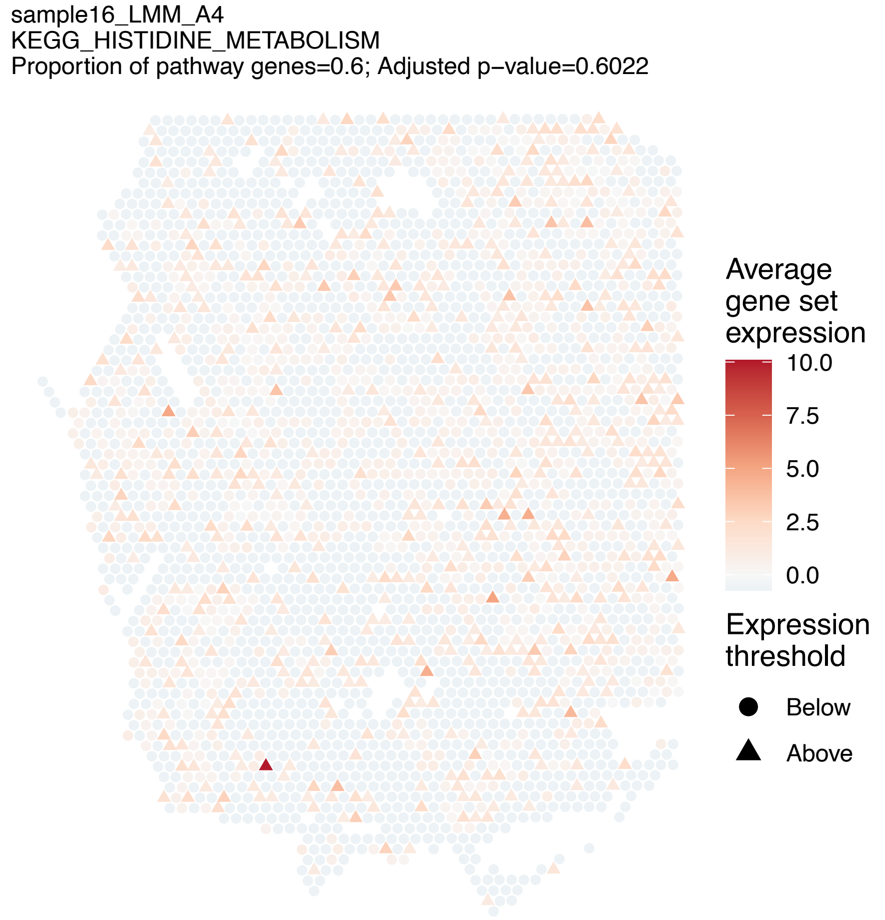
sample16_LMM_A4
KEGG_HISTIDINE_METABOLISM
Proportion of pathway genes=0.6; Adjusted p−value=0.6022
Average gene set expression
10.0
7.5
5.0
2.5
0.0
Expression threshold
Below
Above

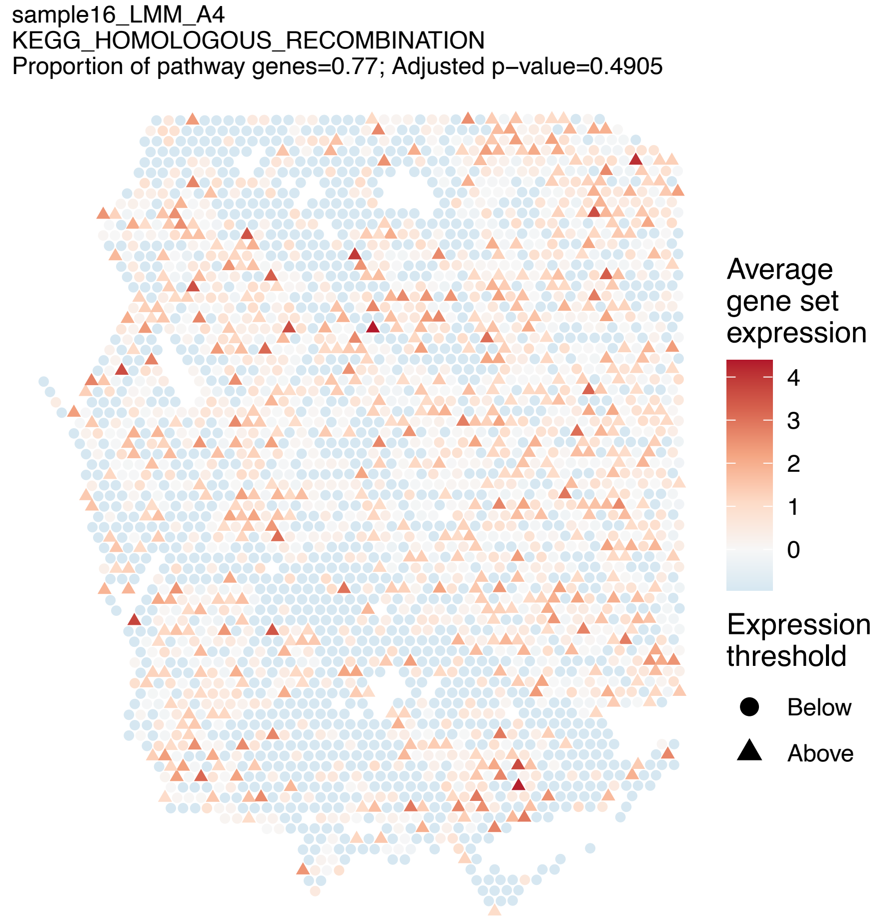
sample16_LMM_A4
KEGG_HOMOLOGOUS_RECOMBINATION
Proportion of pathway genes=0.77; Adjusted p−value=0.4905
Average gene set expression
4
3
2
1
0
Expression threshold
Below
Above

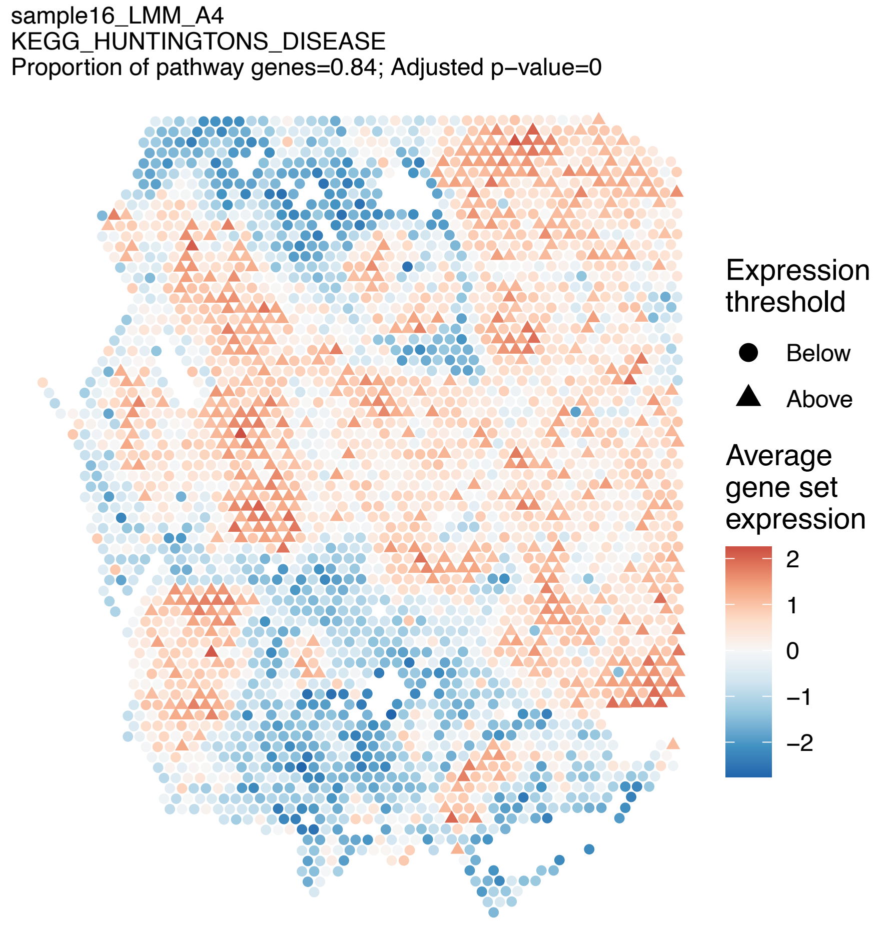
sample16_LMM_A4
KEGG_HUNTINGTONS_DISEASE
Proportion of pathway genes=0.84; Adjusted p−value=0
Expression threshold
Below
Above
Average gene set expression
2
1
0
−1
−2

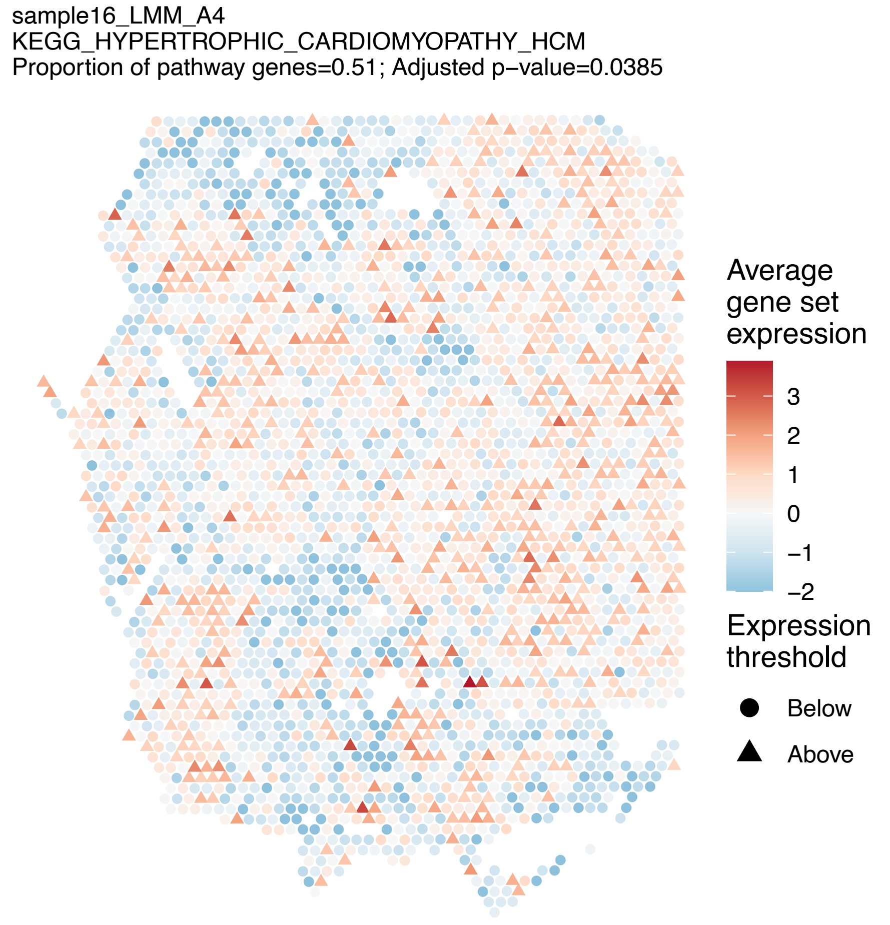
sample16_LMM_A4
KEGG_HYPERTROPHIC_CARDIOMYOPATHY_HCM
Proportion of pathway genes=0.51; Adjusted p−value=0.0385
Average gene set expression
3
2
1
0
−1
−2
Expression threshold
Below
Above

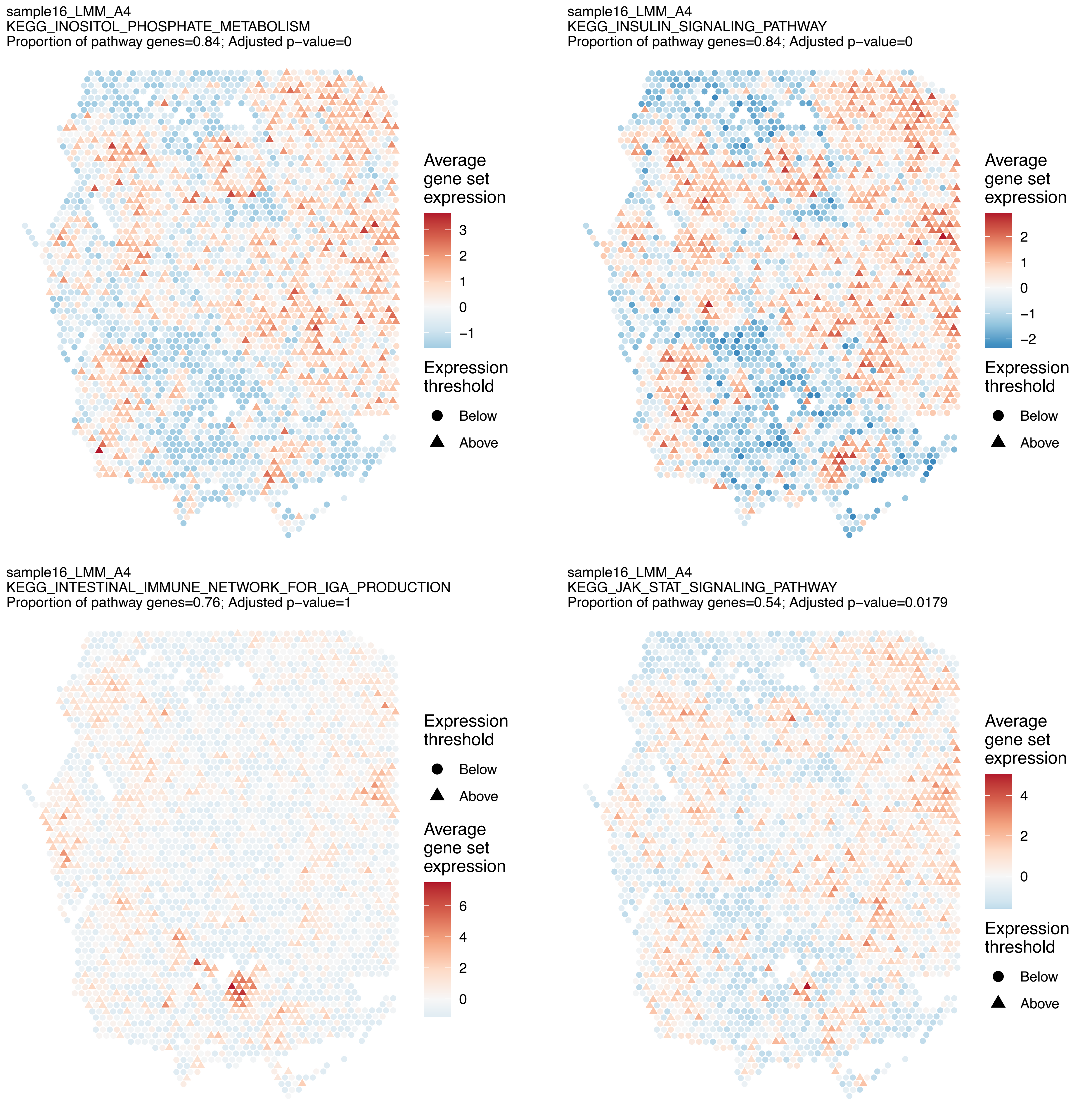
sample16_LMM_A4
KEGG_INOSITOL_PHOSPHATE_METABOLISM
Proportion of pathway genes=0.84; Adjusted p−value=0
Average gene set expression
3
2
1
0
−1
Expression threshold
Below
Above
sample16_LMM_A4
KEGG_INSULIN_SIGNALING_PATHWAY
Proportion of pathway genes=0.84; Adjusted p−value=0
Average gene set expression
2
1
0
−1
−2
Expression threshold
Below
Above
sample16_LMM_A4
KEGG_INTESTINAL_IMMUNE_NETWORK_FOR_IGA_PRODUCTION
Proportion of pathway genes=0.76; Adjusted p−value=1
Expression threshold
Below
Above
Average gene set expression
6
4
2
0
sample16_LMM_A4
KEGG_JAK_STAT_SIGNALING_PATHWAY
Proportion of pathway genes=0.54; Adjusted p−value=0.0179
Average gene set expression
4
2
0
Expression threshold
Below
Above

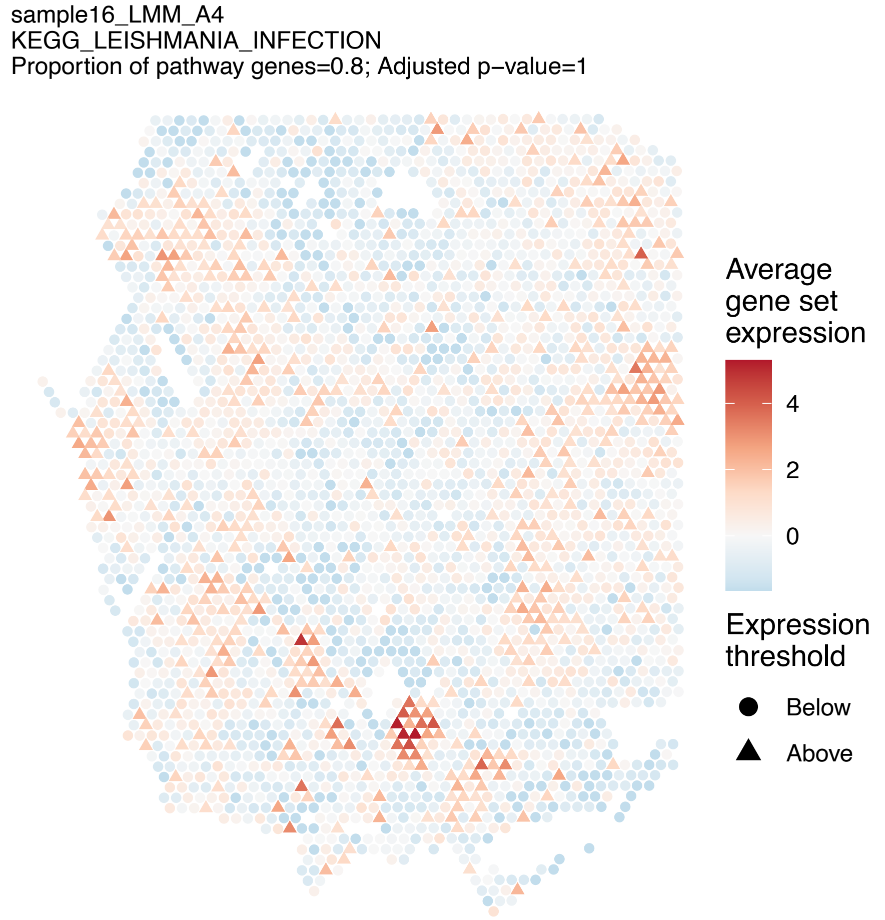
sample16_LMM_A4
KEGG_LEISHMANIA_INFECTION
Proportion of pathway genes=0.8; Adjusted p−value=1
Average gene set expression
4
2
0
Expression threshold
Below
Above

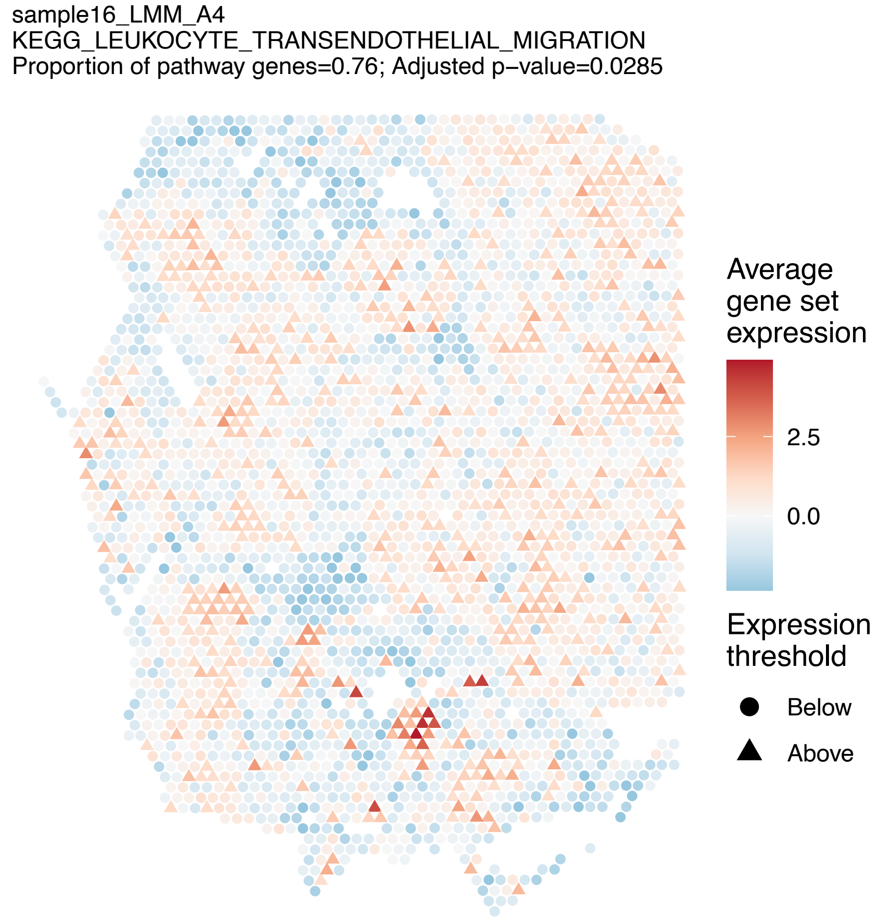
sample16_LMM_A4
KEGG_LEUKOCYTE_TRANSENDOTHELIAL_MIGRATION
Proportion of pathway genes=0.76; Adjusted p−value=0.0285
Average gene set expression
2.5
0.0
Expression threshold
Below
Above

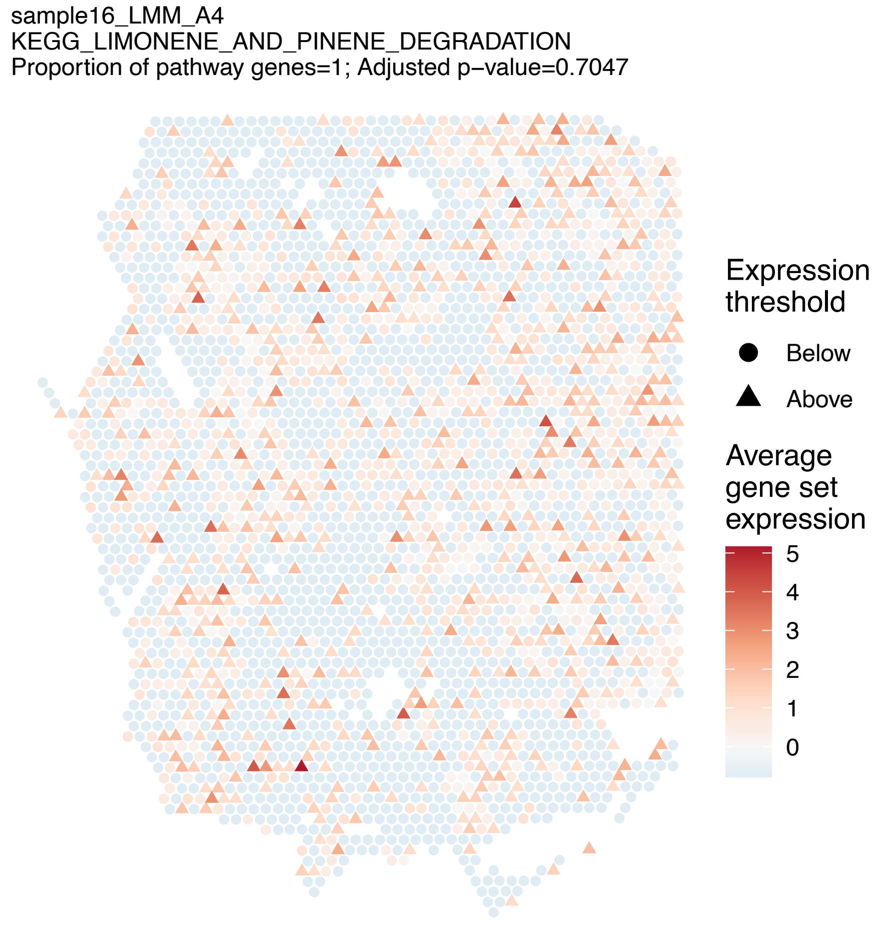
sample16_LMM_A4
KEGG_LIMONENE_AND_PINENE_DEGRADATION
Proportion of pathway genes=1; Adjusted p−value=0.7047
Expression threshold
Below
Above
Average gene set expression
5
4
3
2
1
0

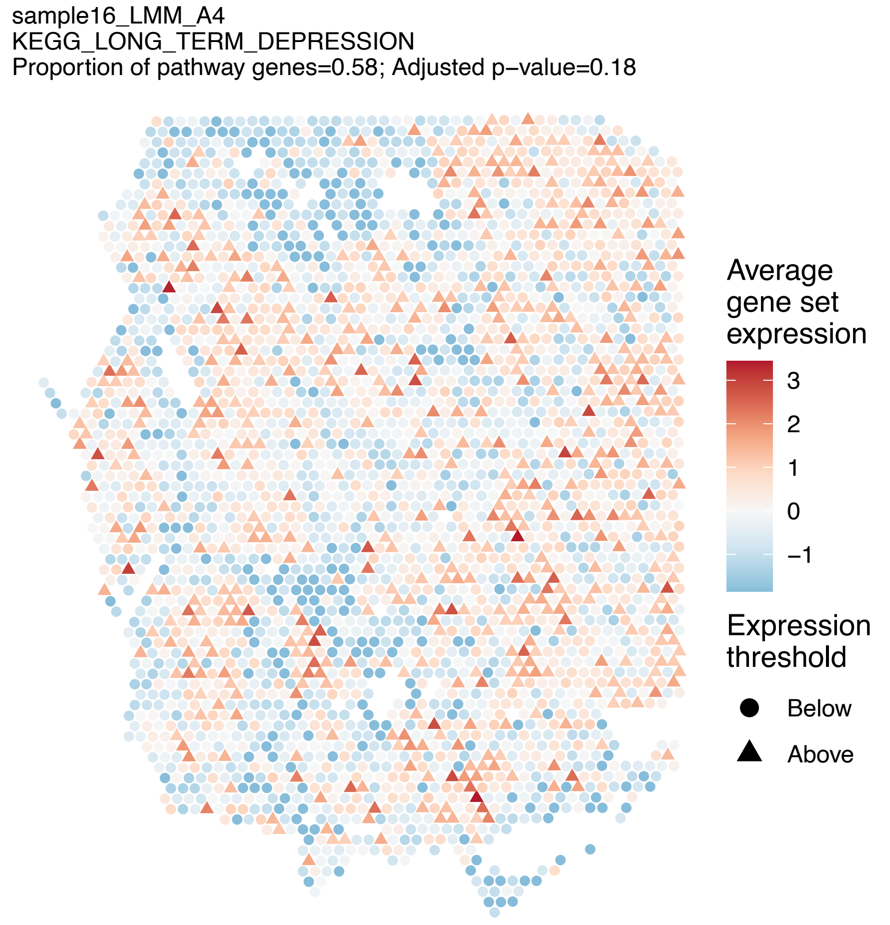
sample16_LMM_A4
KEGG_LONG_TERM_DEPRESSION
Proportion of pathway genes=0.58; Adjusted p−value=0.18
Average gene set expression
3
2
1
0
−1
Expression threshold
Below
Above

sample16_LMM_A4
KEGG_LONG_TERM_POTENTIATION
Proportion of pathway genes=0.68; Adjusted p−value=0.0146

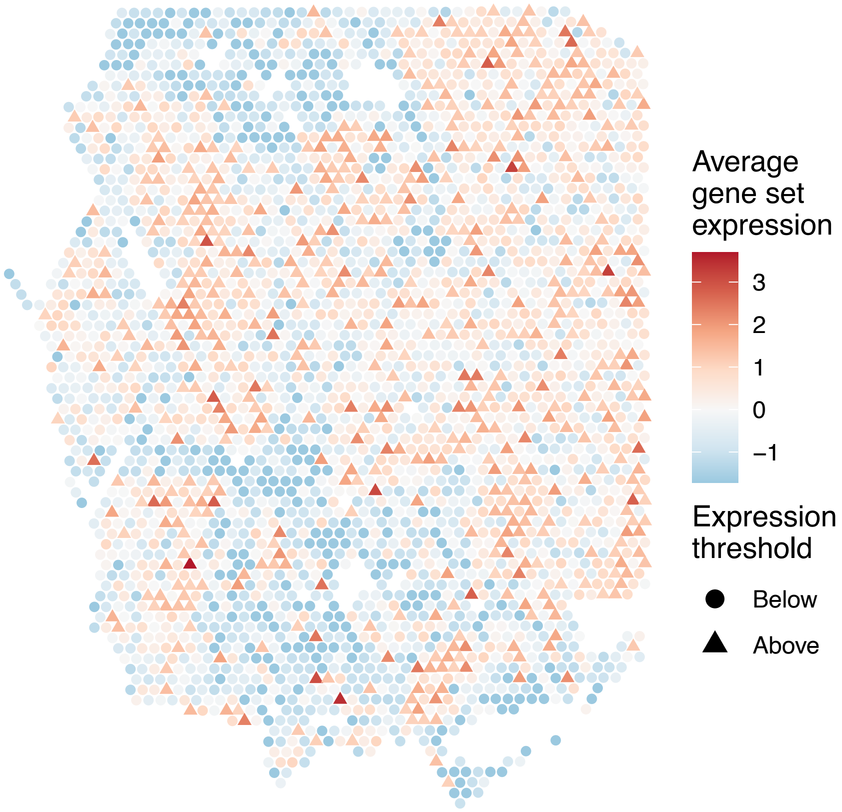


sample16_LMM_A4
KEGG_LYSINE_DEGRADATION
Proportion of pathway genes=0.92; Adjusted p−value=0.0675

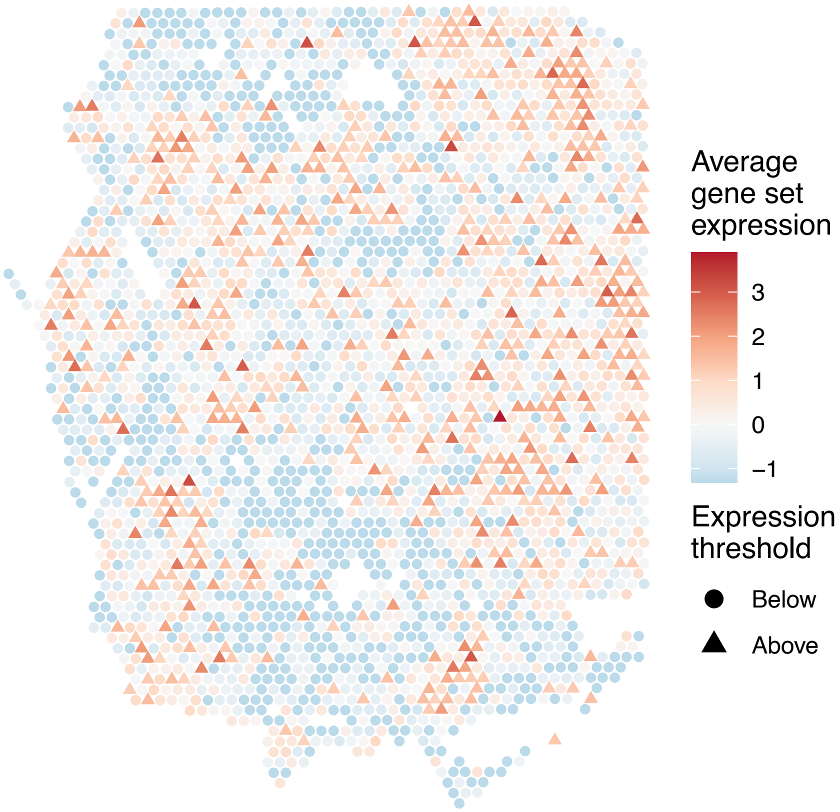


sample16_LMM_A4
KEGG_LYSOSOME
Proportion of pathway genes=0.92; Adjusted p−value=0.0574

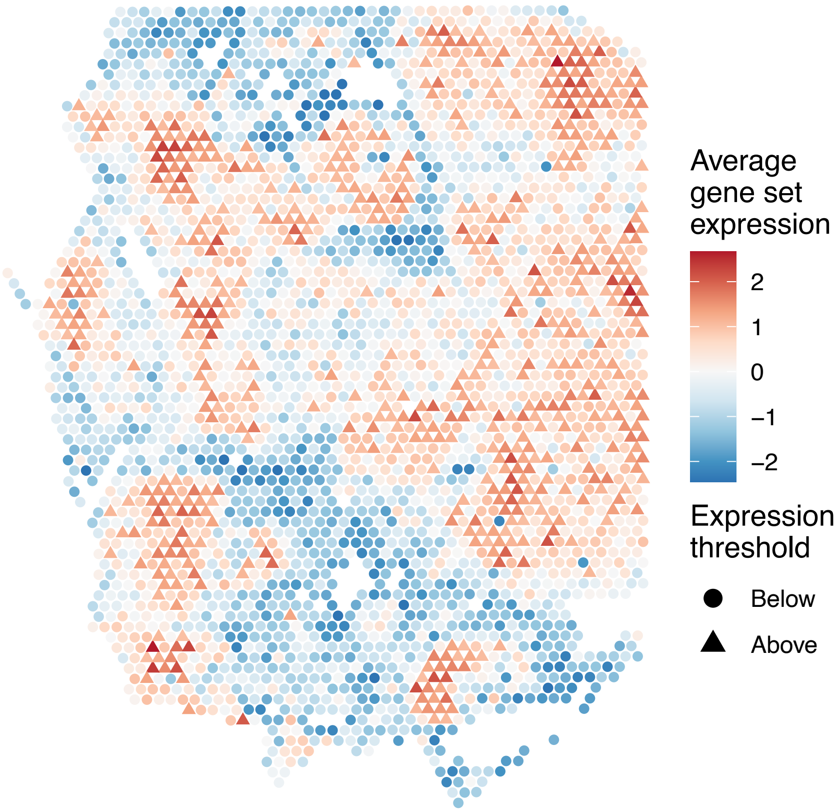


sample16_LMM_A4
KEGG_MAPK_SIGNALING_PATHWAY
Proportion of pathway genes=0.72; Adjusted p−value=0.9798

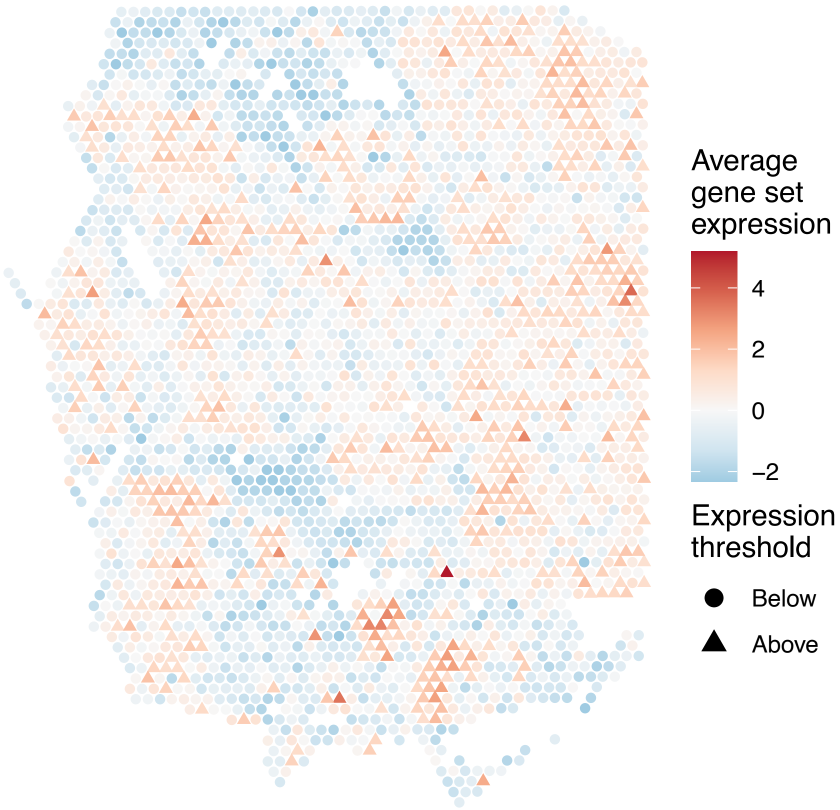

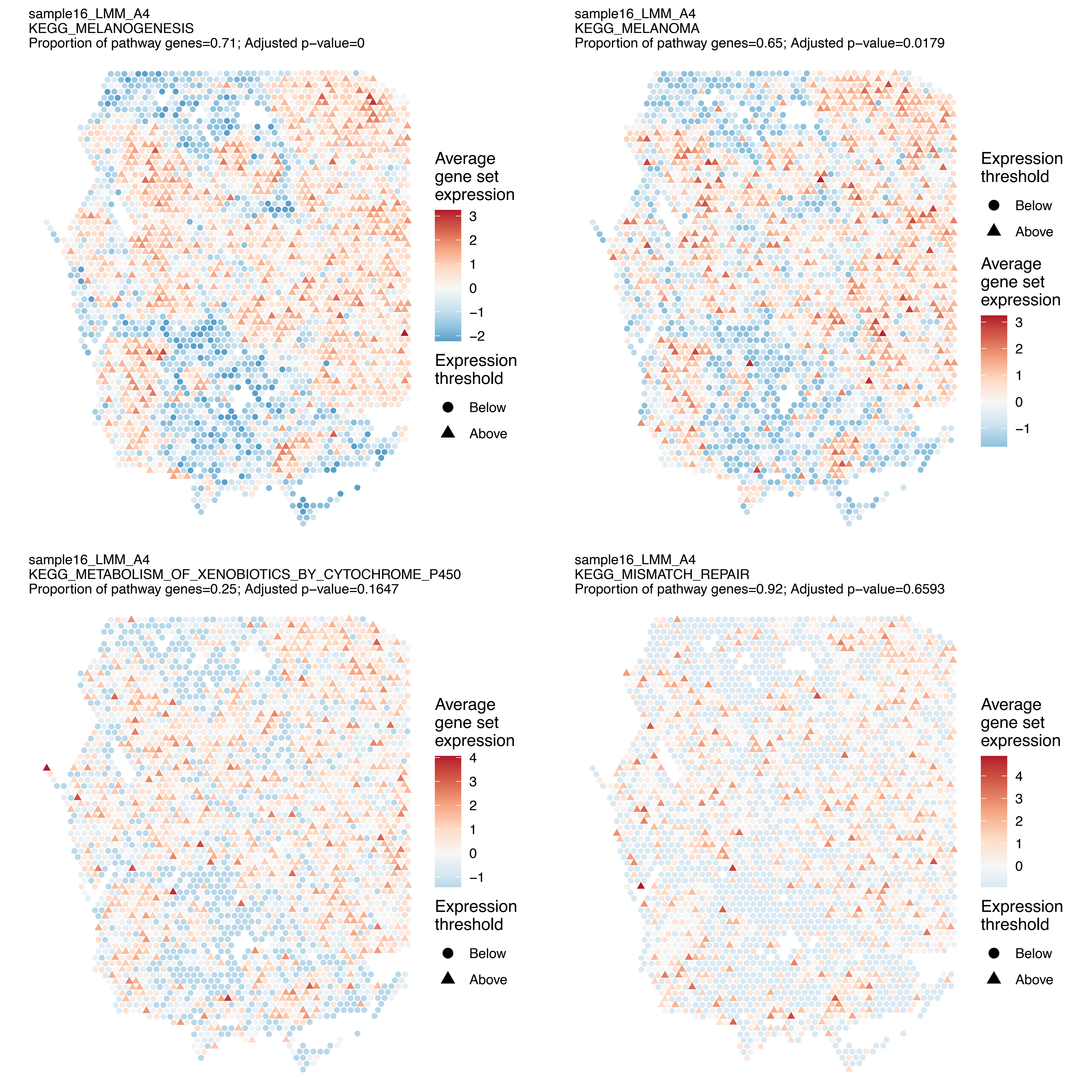
sample16_LMM_A4
KEGG_MELANOGENESIS
Proportion of pathway genes=0.71; Adjusted p−value=0
Average gene set expression
3
2
1
0
−1
−2
Expression threshold
Below
Above
sample16_LMM_A4
KEGG_MELANOMA
Proportion of pathway genes=0.65; Adjusted p−value=0.0179
Expression threshold
Below
Above
Average gene set expression
3
2
1
0
−1
sample16_LMM_A4
KEGG_METABOLISM_OF_XENOBIOTICS_BY_CYTOCHROME_P450
Proportion of pathway genes=0.25; Adjusted p−value=0.1647
Average gene set expression
4
3
2
1
0
−1
Expression threshold
Below
Above
sample16_LMM_A4
KEGG_MISMATCH_REPAIR
Proportion of pathway genes=0.92; Adjusted p−value=0.6593
Average gene set expression
4
3
2
1
0
Expression threshold
Below
Above

sample16_LMM_A4
KEGG_MTOR_SIGNALING_PATHWAY
Proportion of pathway genes=0.93; Adjusted p−value=0.1078

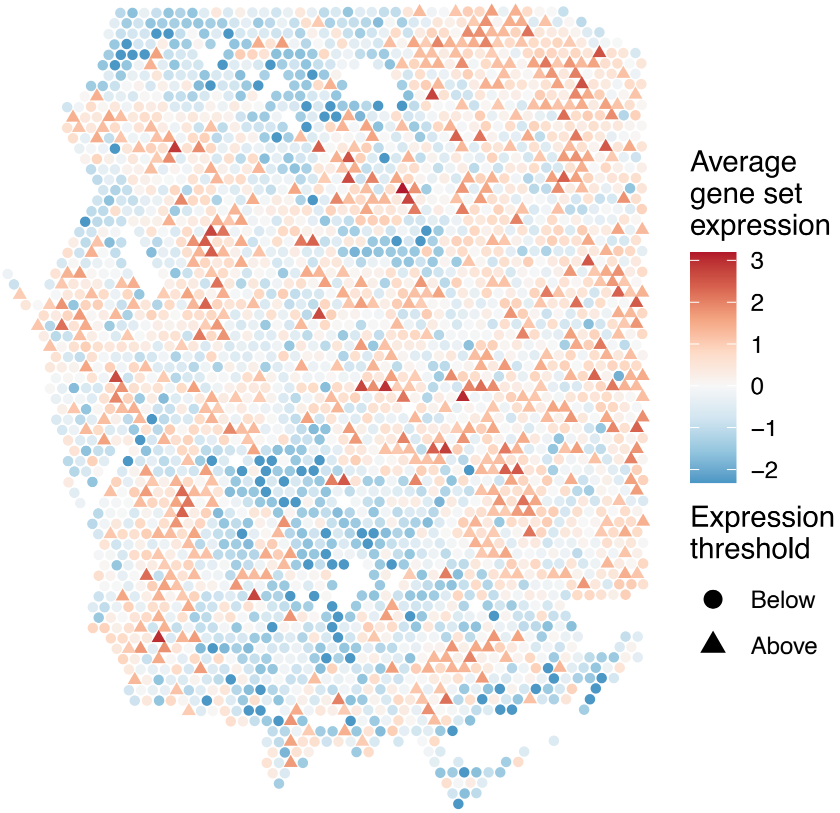


sample16_LMM_A4
KEGG_N_GLYCAN_BIOSYNTHESIS
Proportion of pathway genes=0.94; Adjusted p−value=0.0731

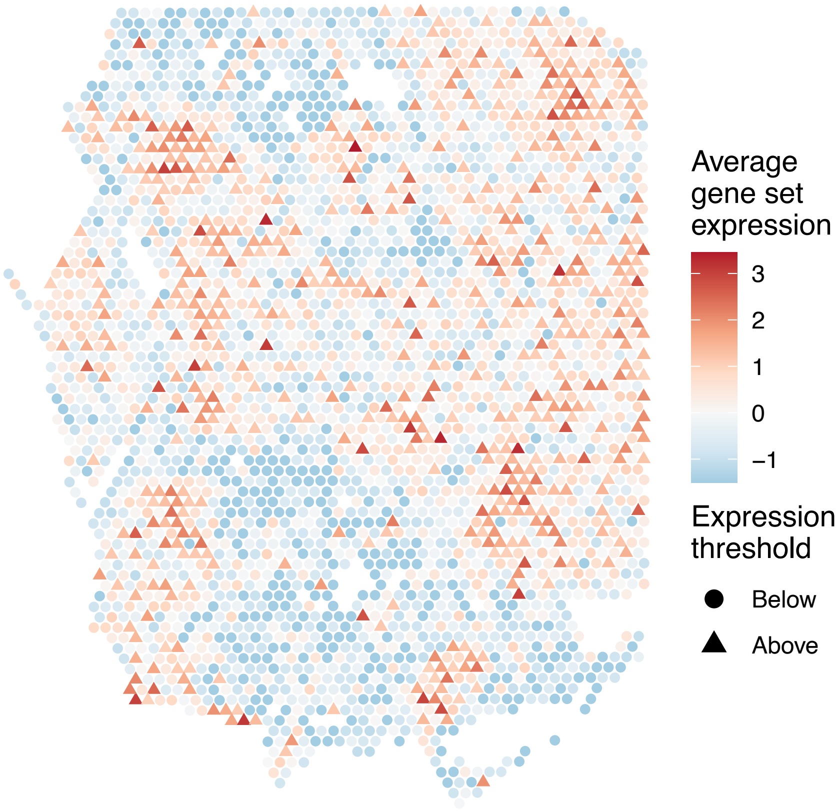


sample16_LMM_A4
KEGG_NEUROACTIVE_LIGAND_RECEPTOR_INTERACTION
Proportion of pathway genes=0.24; Adjusted p−value=0.1559

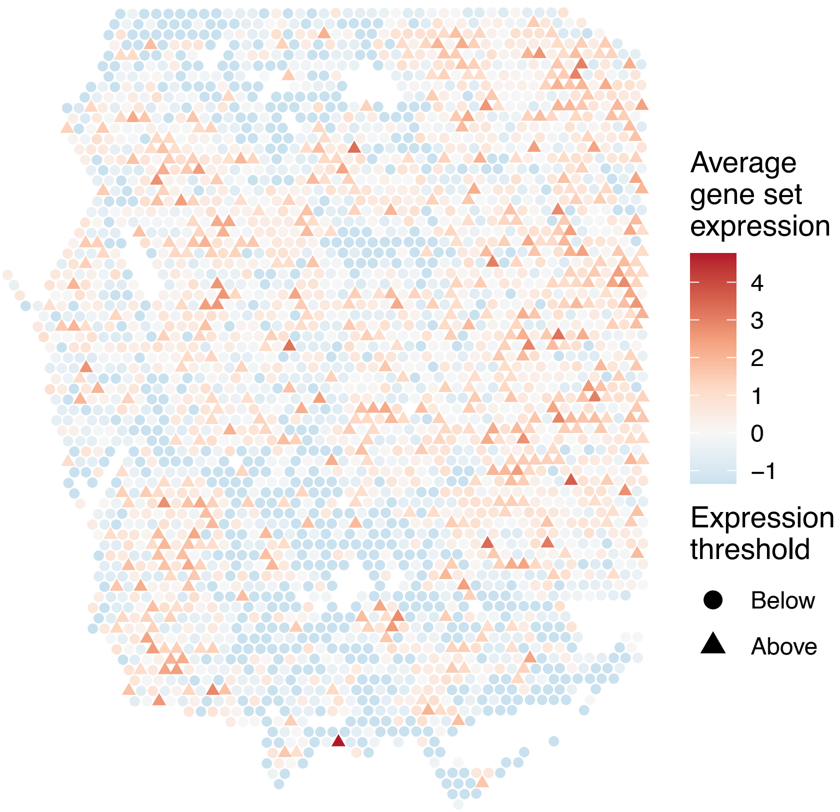


sample16_LMM_A4
KEGG_NEUROTROPHIN_SIGNALING_PATHWAY
Proportion of pathway genes=0.84; Adjusted p−value=0.0067

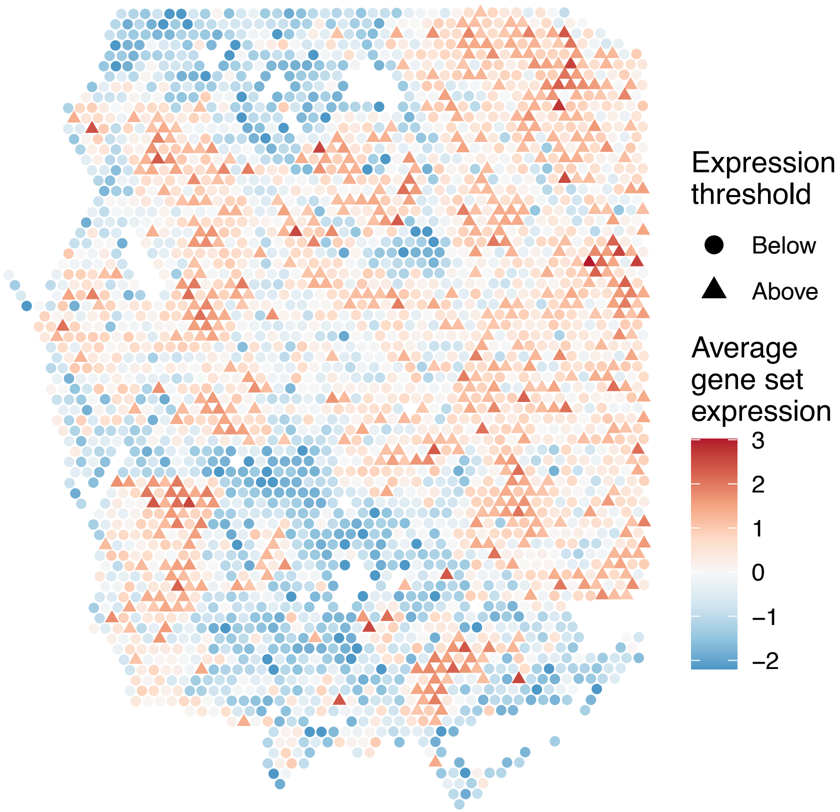

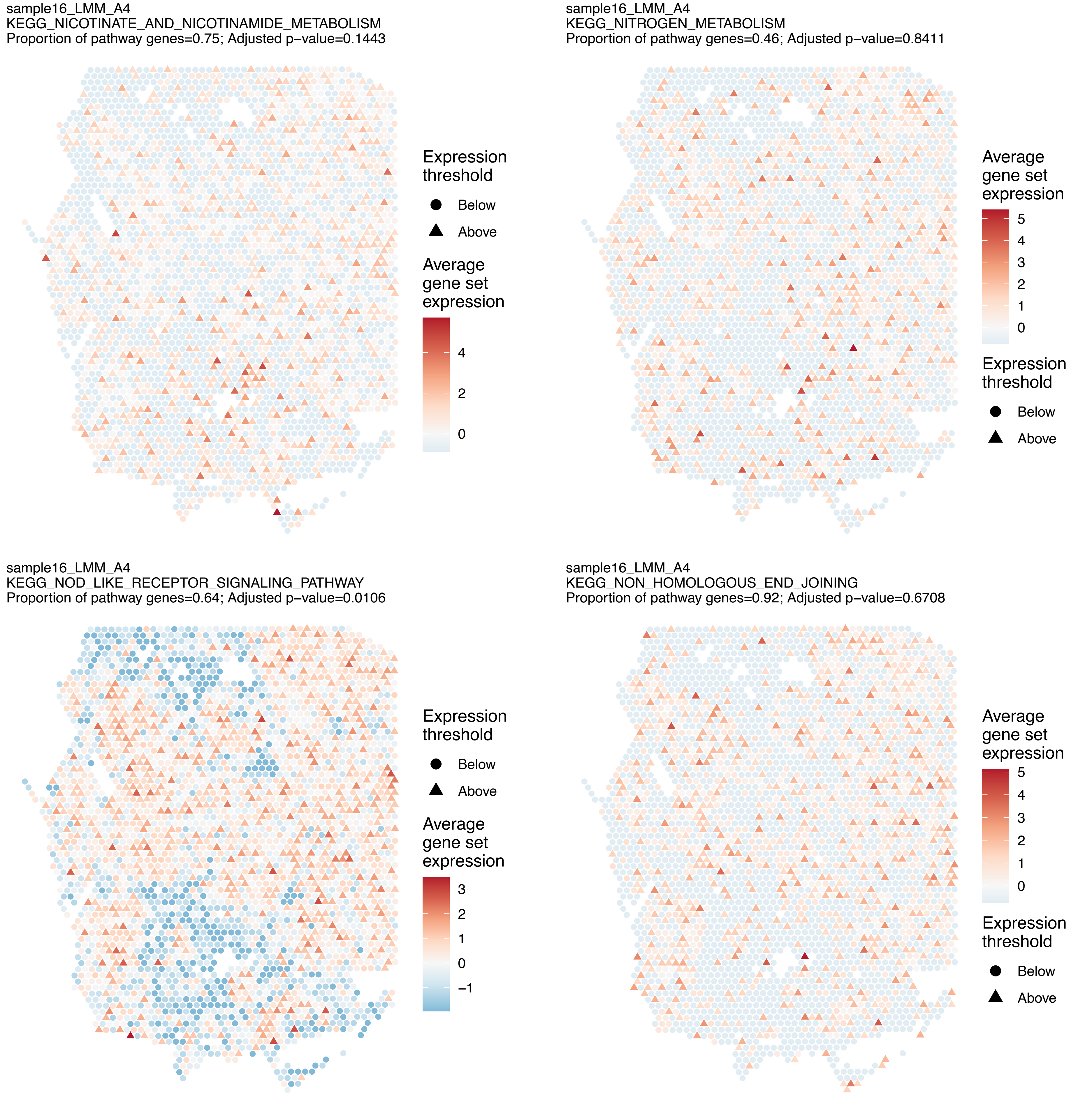
sample16_LMM_A4
KEGG_NICOTINATE_AND_NICOTINAMIDE_METABOLISM
Proportion of pathway genes=0.75; Adjusted p−value=0.1443
Expression threshold
Below
Above
Average gene set expression
4
2
0
sample16_LMM_A4
KEGG_NITROGEN_METABOLISM
Proportion of pathway genes=0.46; Adjusted p−value=0.8411
Average gene set expression
5
4
3
2
1
0
Expression threshold
Below
Above
sample16_LMM_A4
KEGG_NOD_LIKE_RECEPTOR_SIGNALING_PATHWAY
Proportion of pathway genes=0.64; Adjusted p−value=0.0106
Expression threshold
Below
Above
Average gene set expression
3
2
1
0
−1
sample16_LMM_A4
KEGG_NON_HOMOLOGOUS_END_JOINING
Proportion of pathway genes=0.92; Adjusted p−value=0.6708
Average gene set expression
5
4
3
2
1
0
Expression threshold
Below
Above

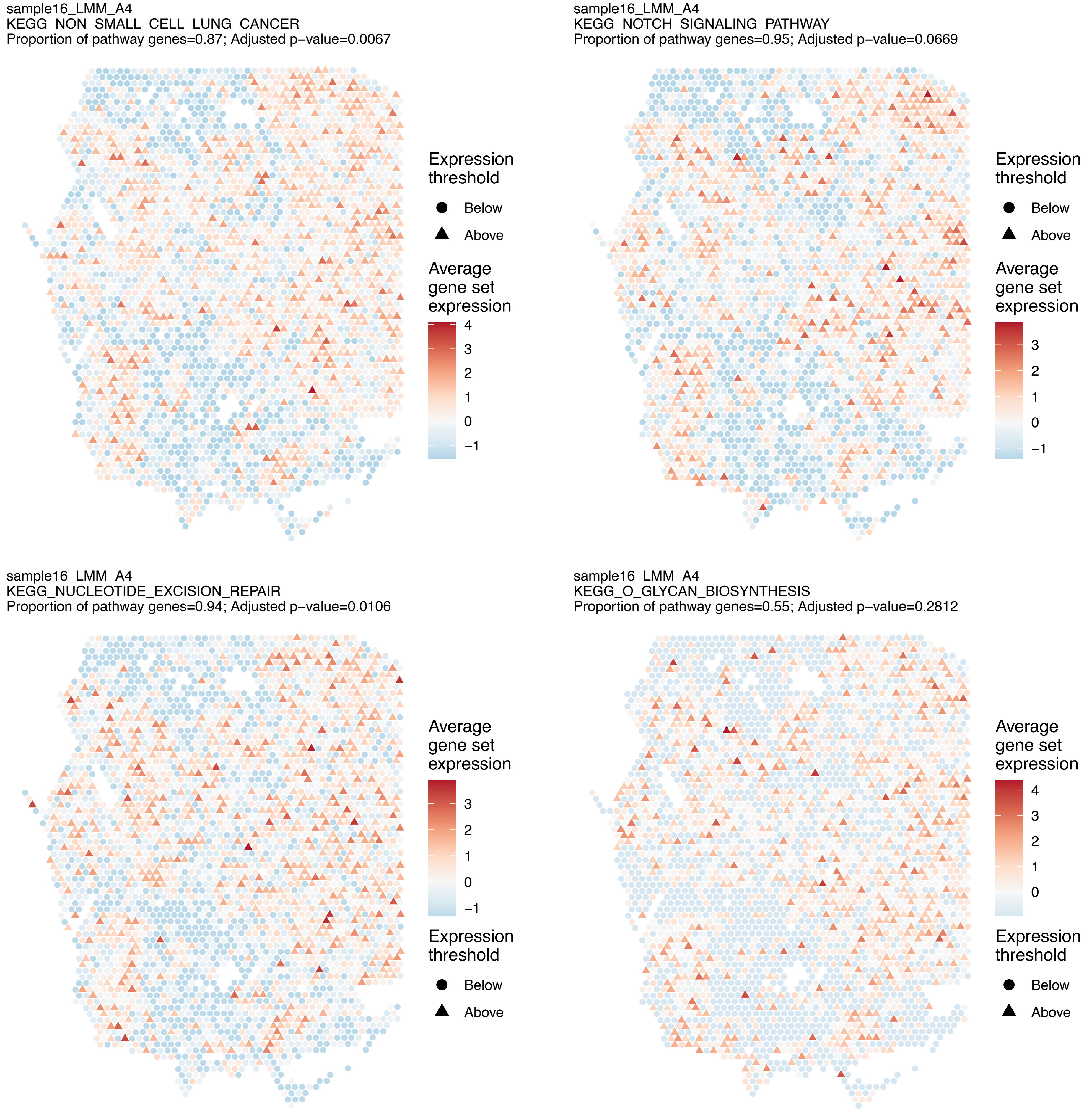
sample16_LMM_A4
KEGG_NON_SMALL_CELL_LUNG_CANCER
Proportion of pathway genes=0.87; Adjusted p–value=0.0067
Expression threshold
Below
Above
Average gene set expression
4
3
2
1
0
−1
sample16_LMM_A4
KEGG_NOTCH_SIGNALING_PATHWAY
Proportion of pathway genes=0.95; Adjusted p–value=0.0669
Expression threshold
Below
Above
Average gene set expression
3
2
1
0
−1
sample16_LMM_A4
KEGG_NUCLEOTIDE_EXCISION_REPAIR
Proportion of pathway genes=0.94; Adjusted p–value=0.0106
Average gene set expression
3
2
1
0
−1
Expression threshold
Below
Above
sample16_LMM_A4
KEGG_O_GLYCAN_BIOSYNTHESIS
Proportion of pathway genes=0.55; Adjusted p–value=0.2812
Average gene set expression
4
3
2
1
0
Expression threshold
Below
Above

sample16_LMM_A4
KEGG_ONE_CARBON_POOL_BY_FOLATE
Proportion of pathway genes=0.95; Adjusted p−value=0.5937

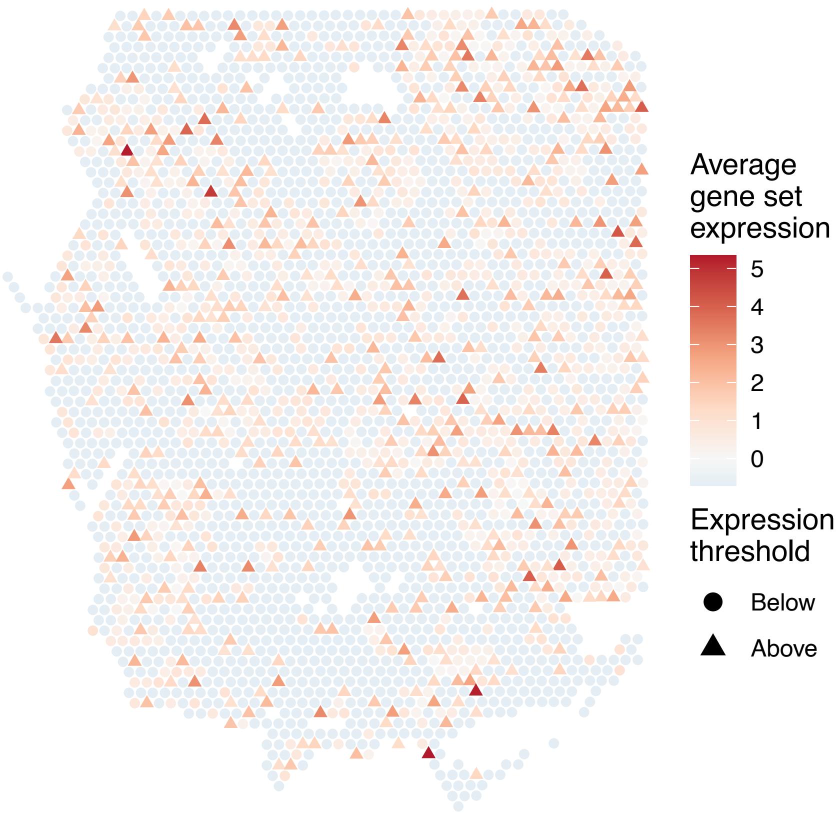


sample16_LMM_A4
KEGG_OOCYTE_MEIOSIS
Proportion of pathway genes=0.71; Adjusted p−value=0

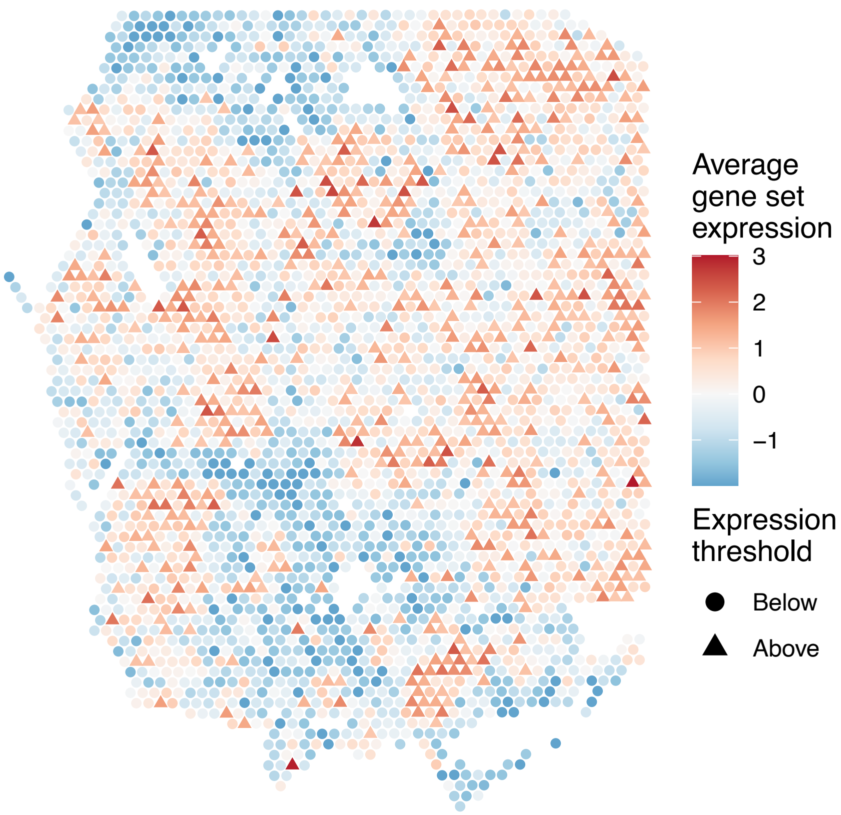


sample16_LMM_A4
KEGG_OTHER_GLYCAN_DEGRADATION
Proportion of pathway genes=0.88; Adjusted p−value=0

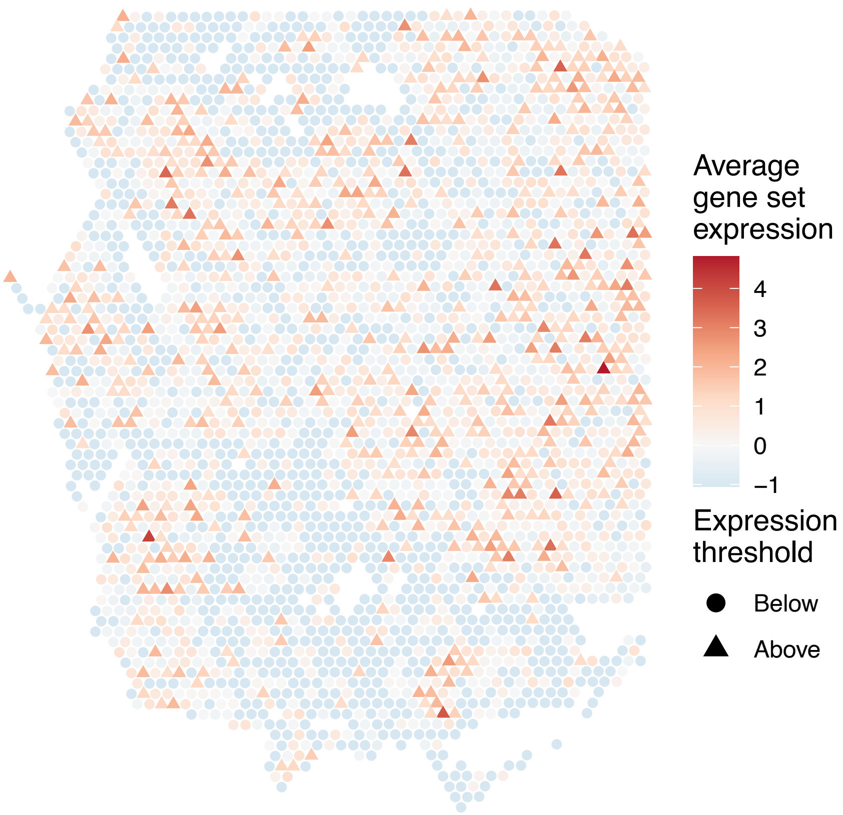


sample16_LMM_A4
KEGG_OXIDATIVE_PHOSPHORYLATION
Proportion of pathway genes=0.83; Adjusted p−value=0

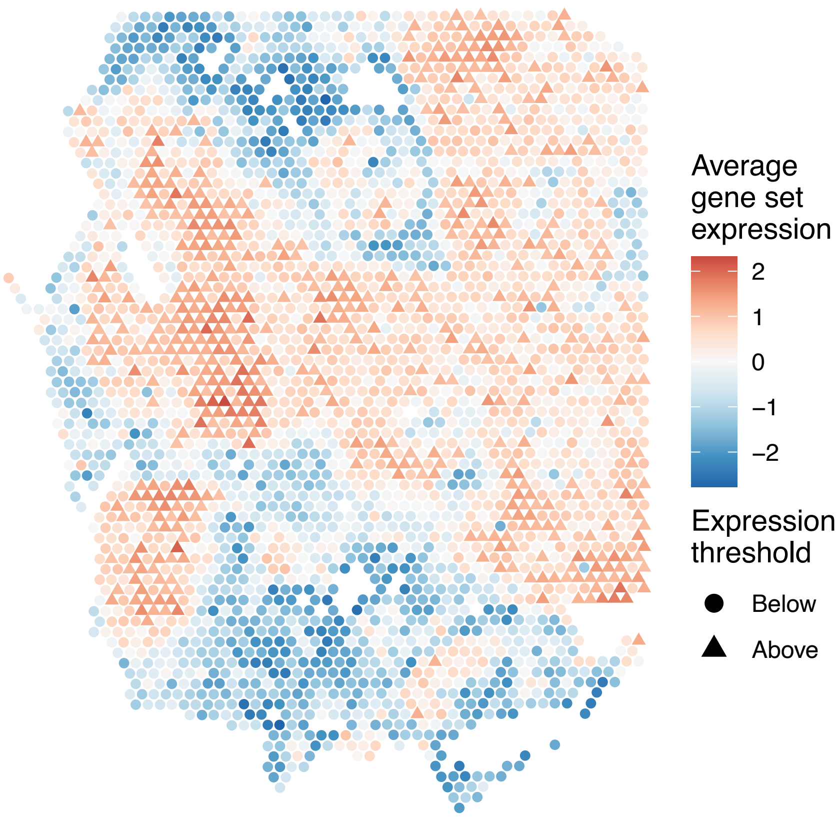

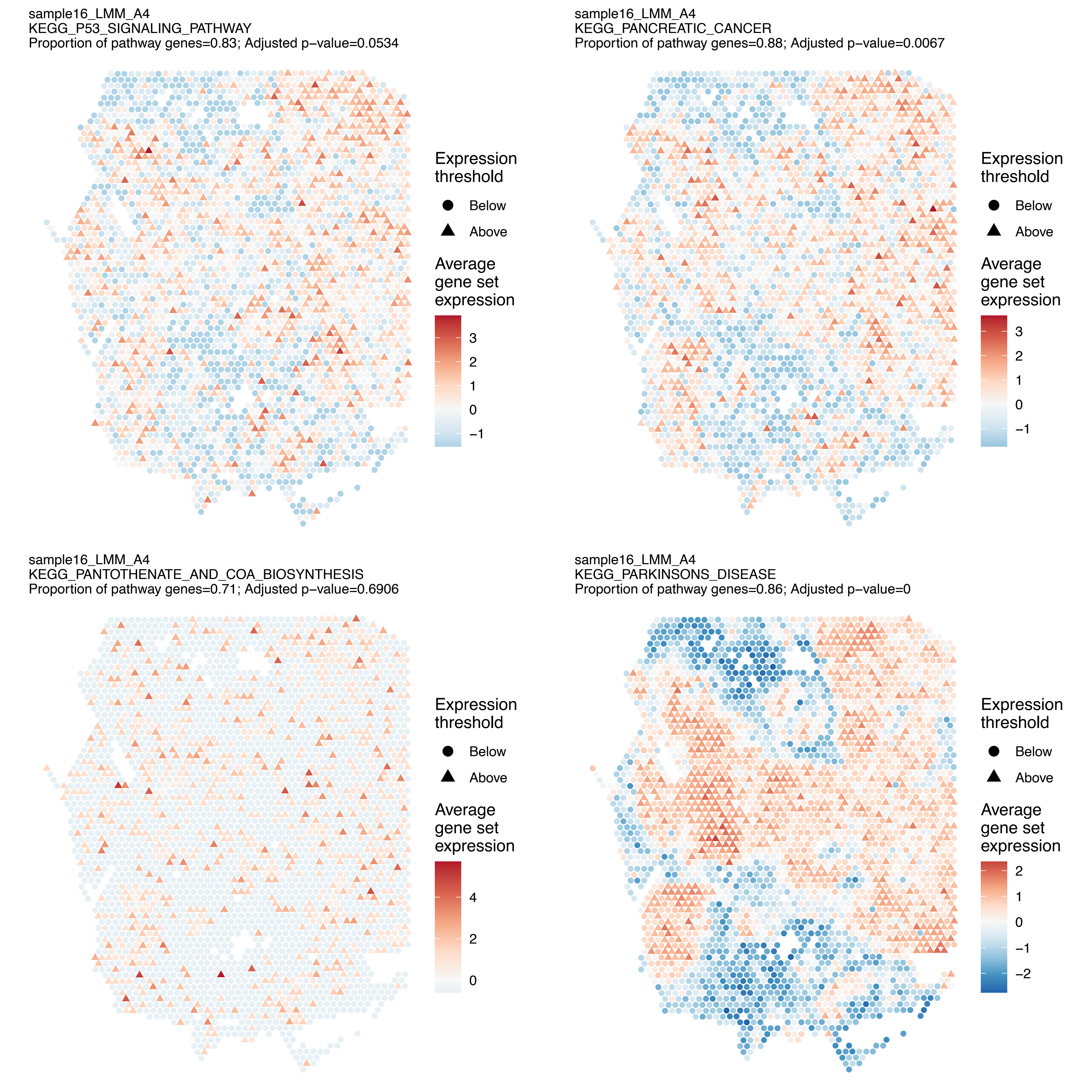
sample16_LMM_A4
KEGG_P53_SIGNALING_PATHWAY
Proportion of pathway genes=0.83; Adjusted p−value=0.0534
Expression threshold
Below
Above
Average gene set expression
3
2
1
0
−1
sample16_LMM_A4
KEGG_PANCREATIC_CANCER
Proportion of pathway genes=0.88; Adjusted p−value=0.0067
Expression threshold
Below
Above
Average gene set expression
3
2
1
0
−1
sample16_LMM_A4
KEGG_PANTOTHENATE_AND_COA_BIOSYNTHESIS
Proportion of pathway genes=0.71; Adjusted p−value=0.6906
Expression threshold
Below
Above
Average gene set expression
4
2
0
sample16_LMM_A4
KEGG_PARKINSONS_DISEASE
Proportion of pathway genes=0.86; Adjusted p−value=0
Expression threshold
Below
Above
Average gene set expression
2
1
0
−1
−2

sample16_LMM_A4
KEGG_PATHOGENIC_ESCHERICHIA_COLI_INFECTION
Proportion of pathway genes=0.83; Adjusted p−value=0.1482

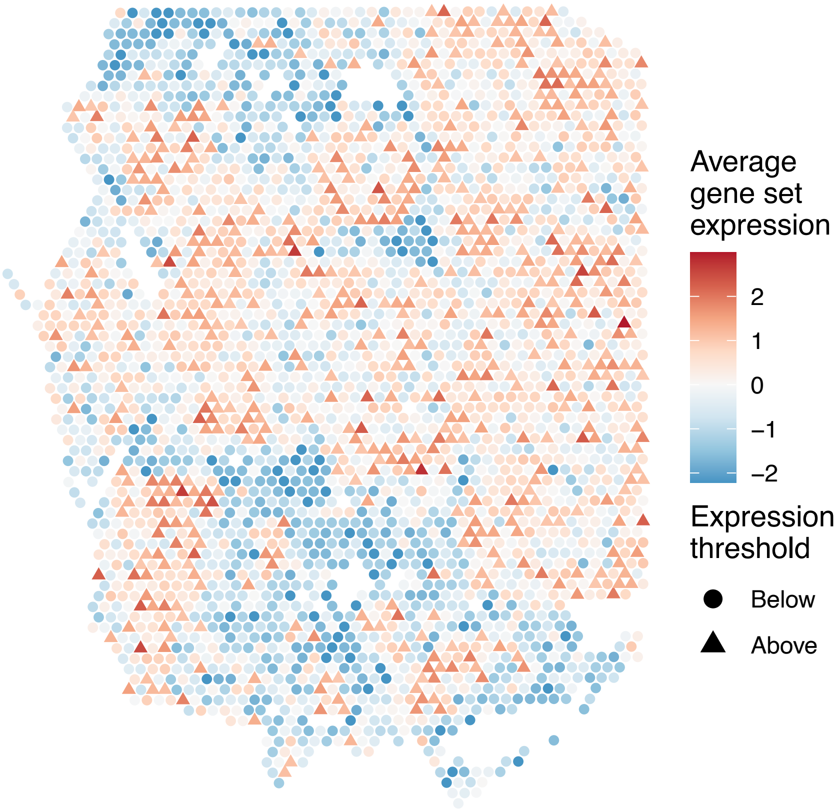


sample16_LMM_A4
KEGG_PATHWAYS_IN_CANCER
Proportion of pathway genes=0.75; Adjusted p−value=0.0695

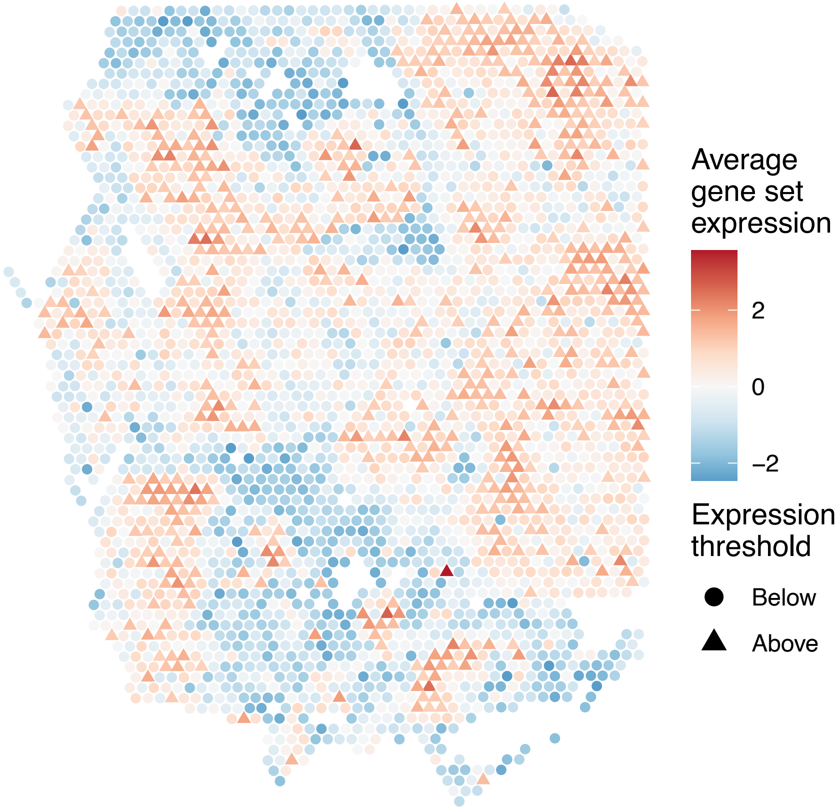


sample16_LMM_A4
KEGG_PENTOSE_PHOSPHATE_PATHWAY
Proportion of pathway genes=0.79; Adjusted p−value=0.0342

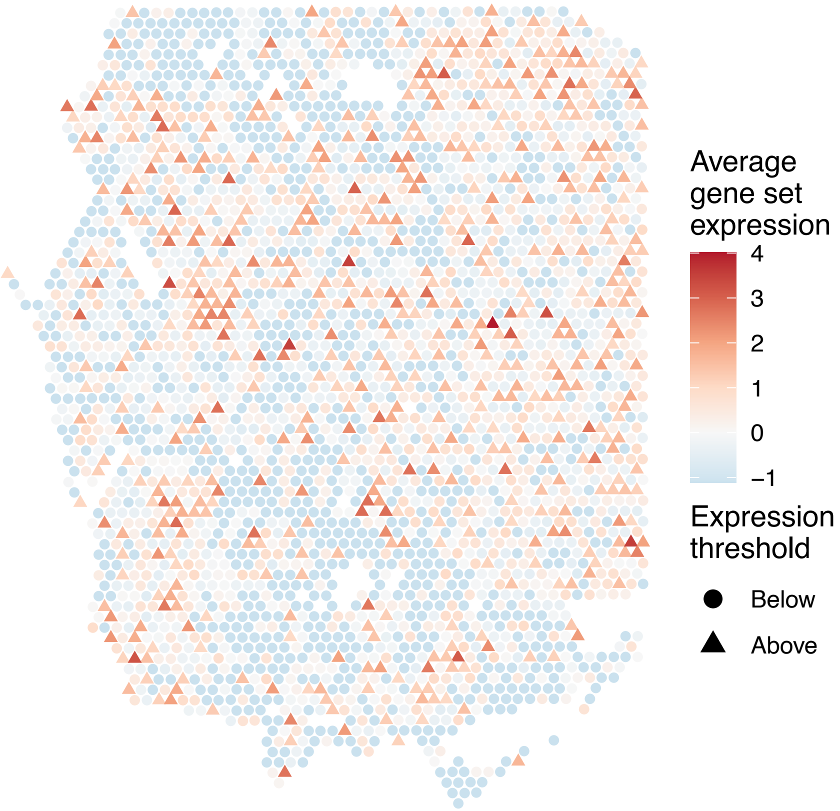


sample16_LMM_A4
KEGG_PEROXISOME
Proportion of pathway genes=0.85; Adjusted p−value=0.0146

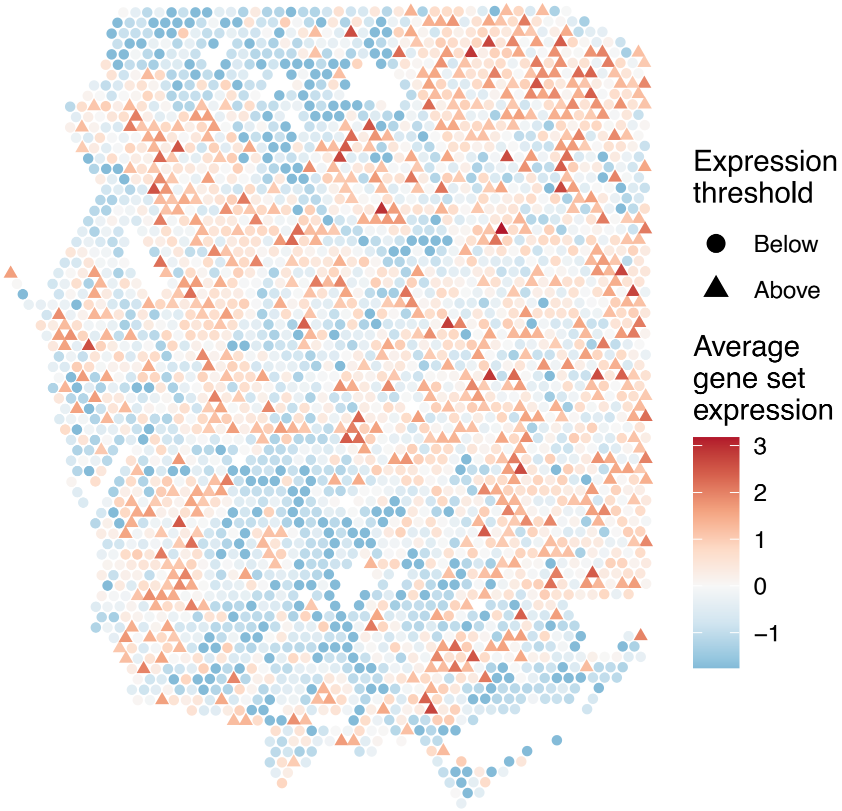

sample16_LMM_A4
KEGG_PHENYLALANINE_METABOLISM
Proportion of pathway genes=0.47; Adjusted p−value=NA

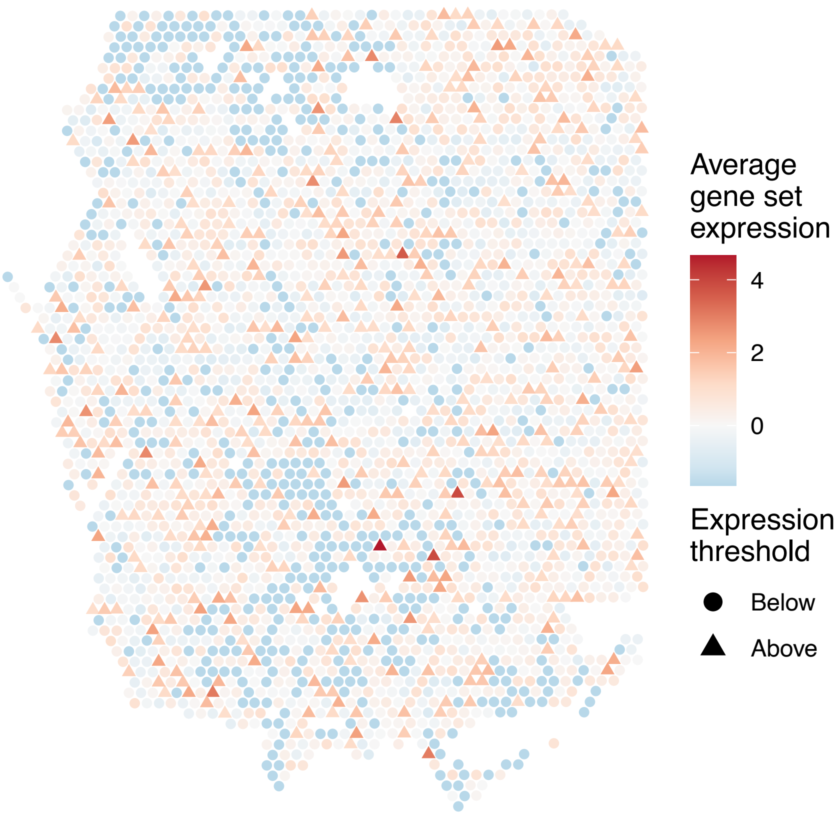


sample16_LMM_A4
KEGG_PHOSPHATIDYLINOSITOL_SIGNALING_SYSTEM
Proportion of pathway genes=0.83; Adjusted p−value=0

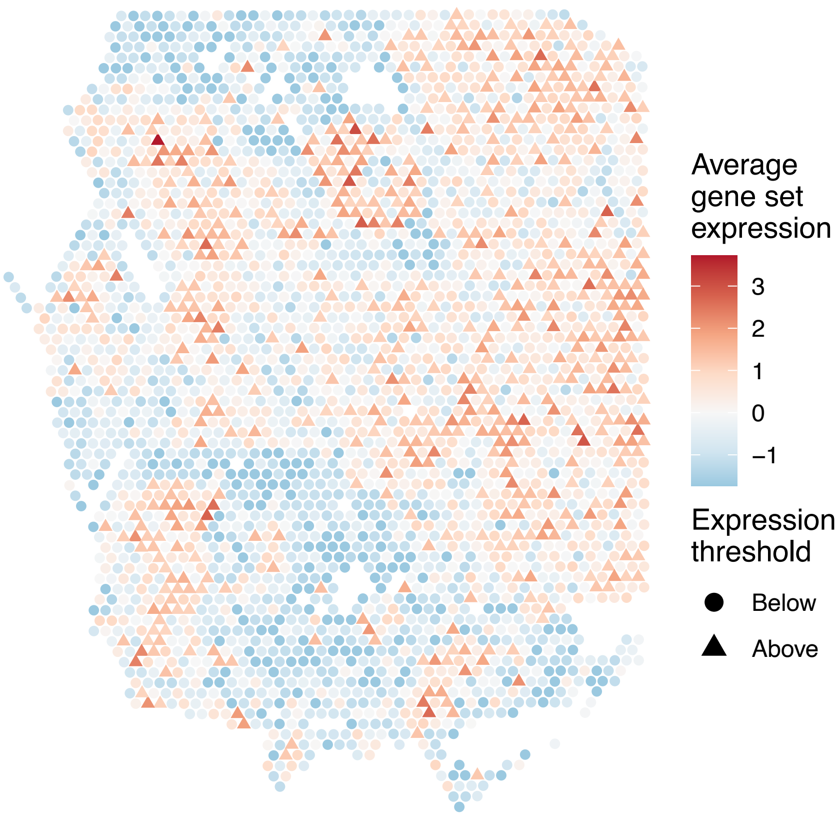


sample16_LMM_A4
KEGG_PORPHYRIN_AND_CHLOROPHYLL_METABOLISM
Proportion of pathway genes=0.44; Adjusted p−value=0.2716

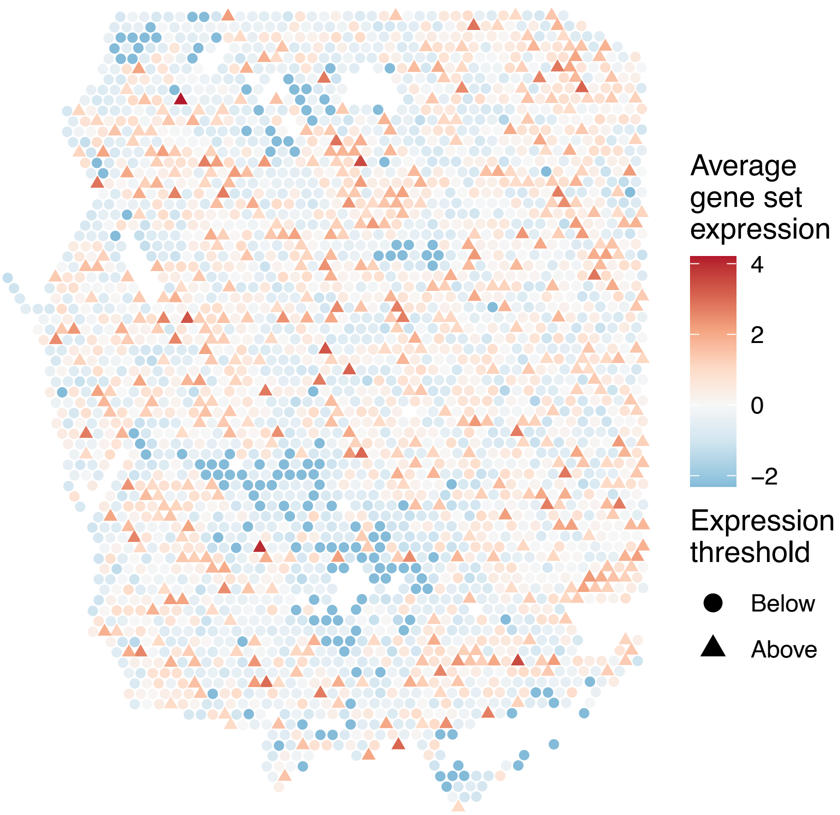


sample16_LMM_A4
KEGG_PPAR_SIGNALING_PATHWAY
Proportion of pathway genes=0.64; Adjusted p−value=0.0106

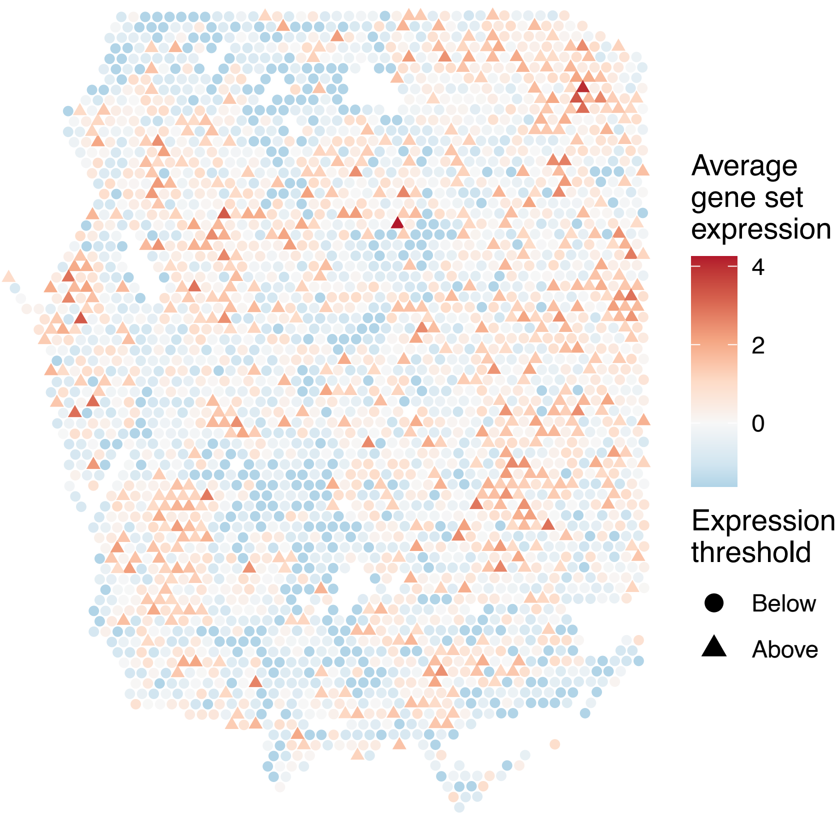

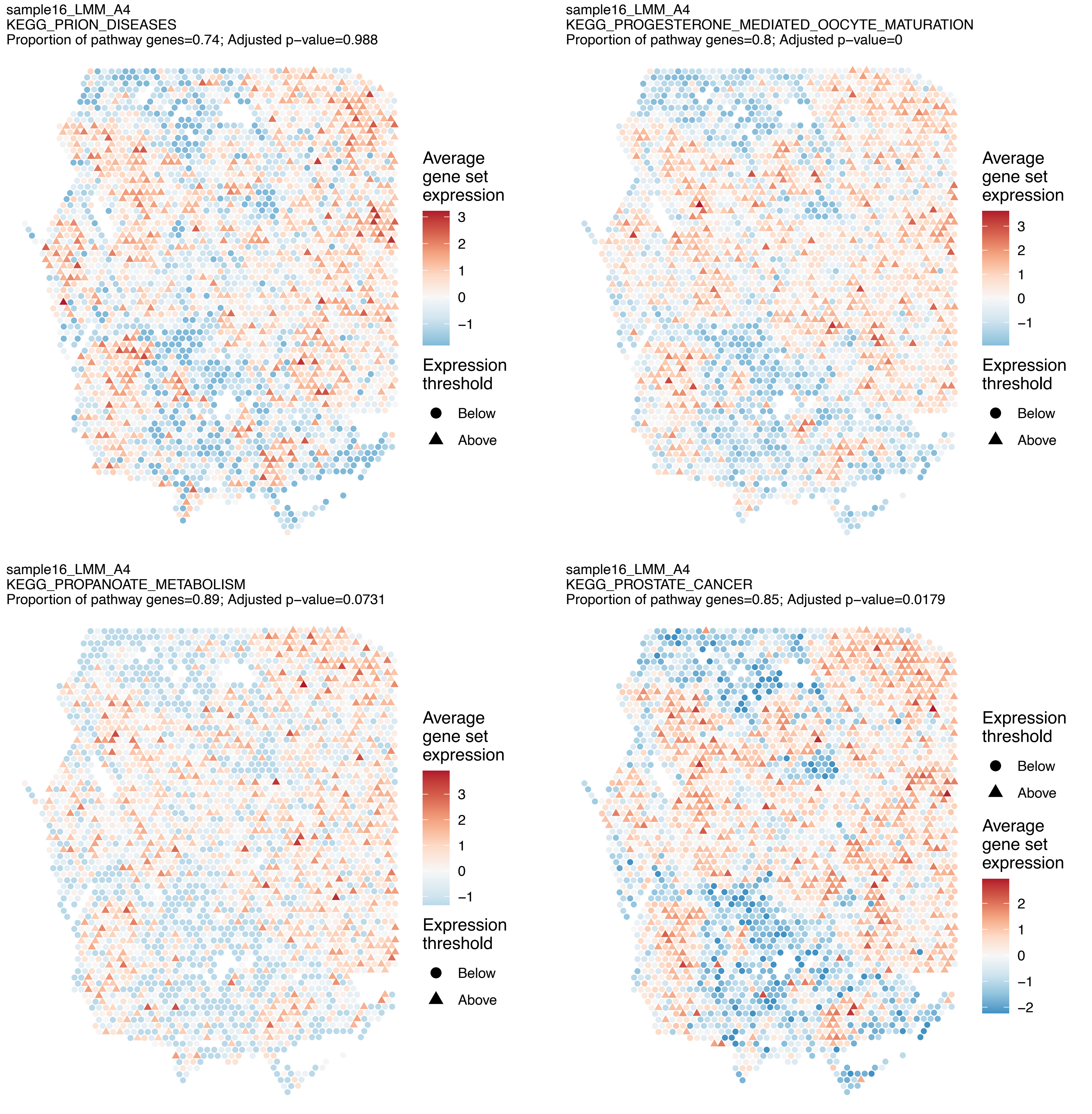
sample16_LMM_A4
KEGG_PRION_DISEASES
Proportion of pathway genes=0.74; Adjusted p−value=0.988
Average gene set expression
3
2
1
0
−1
Expression threshold
Below
Above
sample16_LMM_A4
KEGG_PROGESTERONE_MEDIATED_OOCYTE_MATURATION
Proportion of pathway genes=0.8; Adjusted p−value=0
Average gene set expression
3
2
1
0
−1
Expression threshold
Below
Above
sample16_LMM_A4
KEGG_PROPANOATE_METABOLISM
Proportion of pathway genes=0.89; Adjusted p−value=0.0731
Average gene set expression
3
2
1
0
−1
Expression threshold
Below
Above
sample16_LMM_A4
KEGG_PROSTATE_CANCER
Proportion of pathway genes=0.85; Adjusted p−value=0.0179
Expression threshold
Below
Above
Average gene set expression
2
1
0
−1
−2

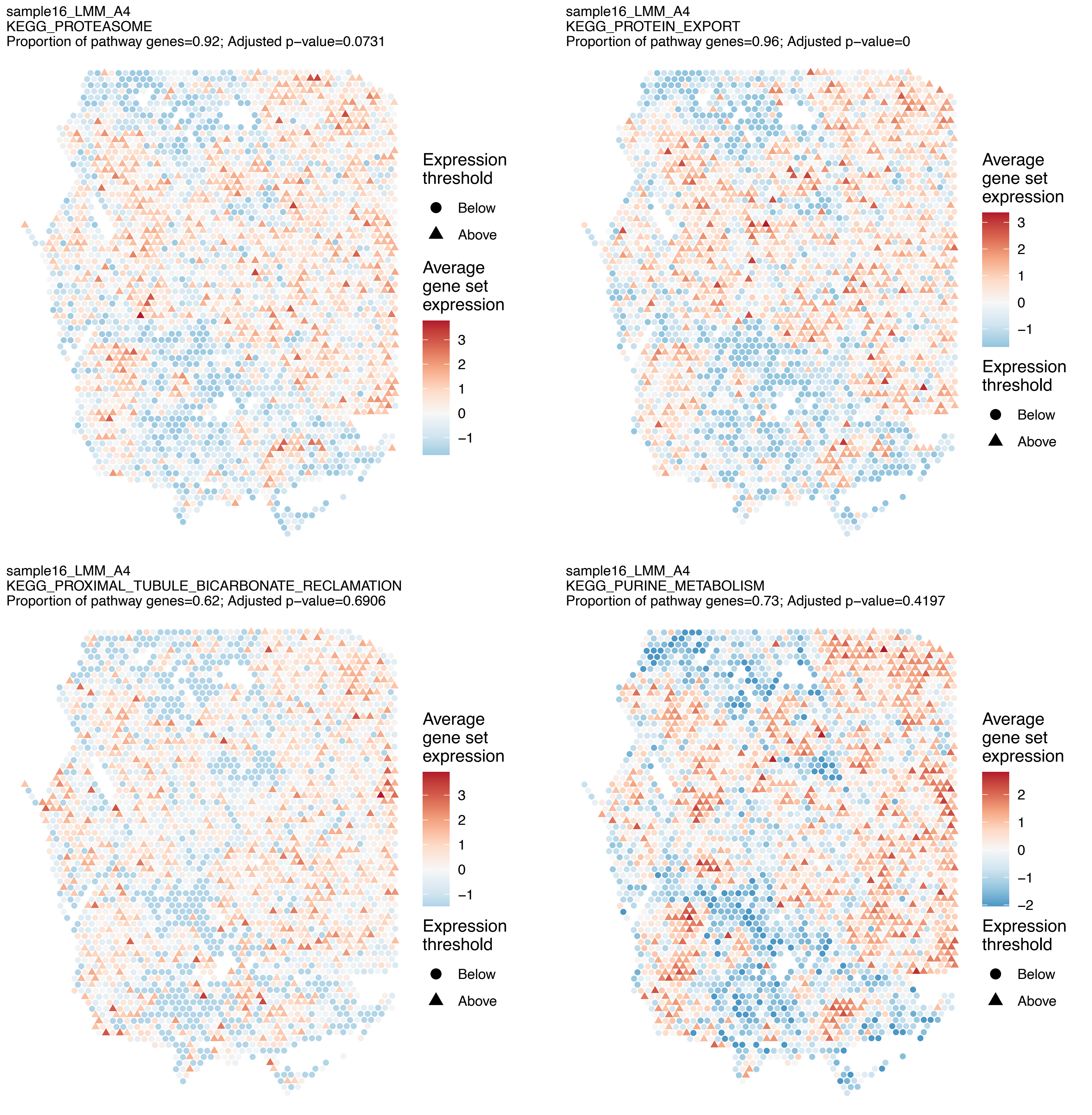
sample16_LMM_A4
KEGG_PROTEASOME
Proportion of pathway genes=0.92; Adjusted p−value=0.0731
Expression threshold
Below
Above
Average gene set expression
3
2
1
0
−1
sample16_LMM_A4
KEGG_PROTEIN_EXPORT
Proportion of pathway genes=0.96; Adjusted p−value=0
Average gene set expression
3
2
1
0
−1
Expression threshold
Below
Above
sample16_LMM_A4
KEGG_PROXIMAL_TUBULE_BICARBONATE_RECLAMATION
Proportion of pathway genes=0.62; Adjusted p−value=0.6906
Average gene set expression
3
2
1
0
−1
Expression threshold
Below
Above
sample16_LMM_A4
KEGG_PURINE_METABOLISM
Proportion of pathway genes=0.73; Adjusted p−value=0.4197
Average gene set expression
2
1
0
−1
−2
Expression threshold
Below
Above

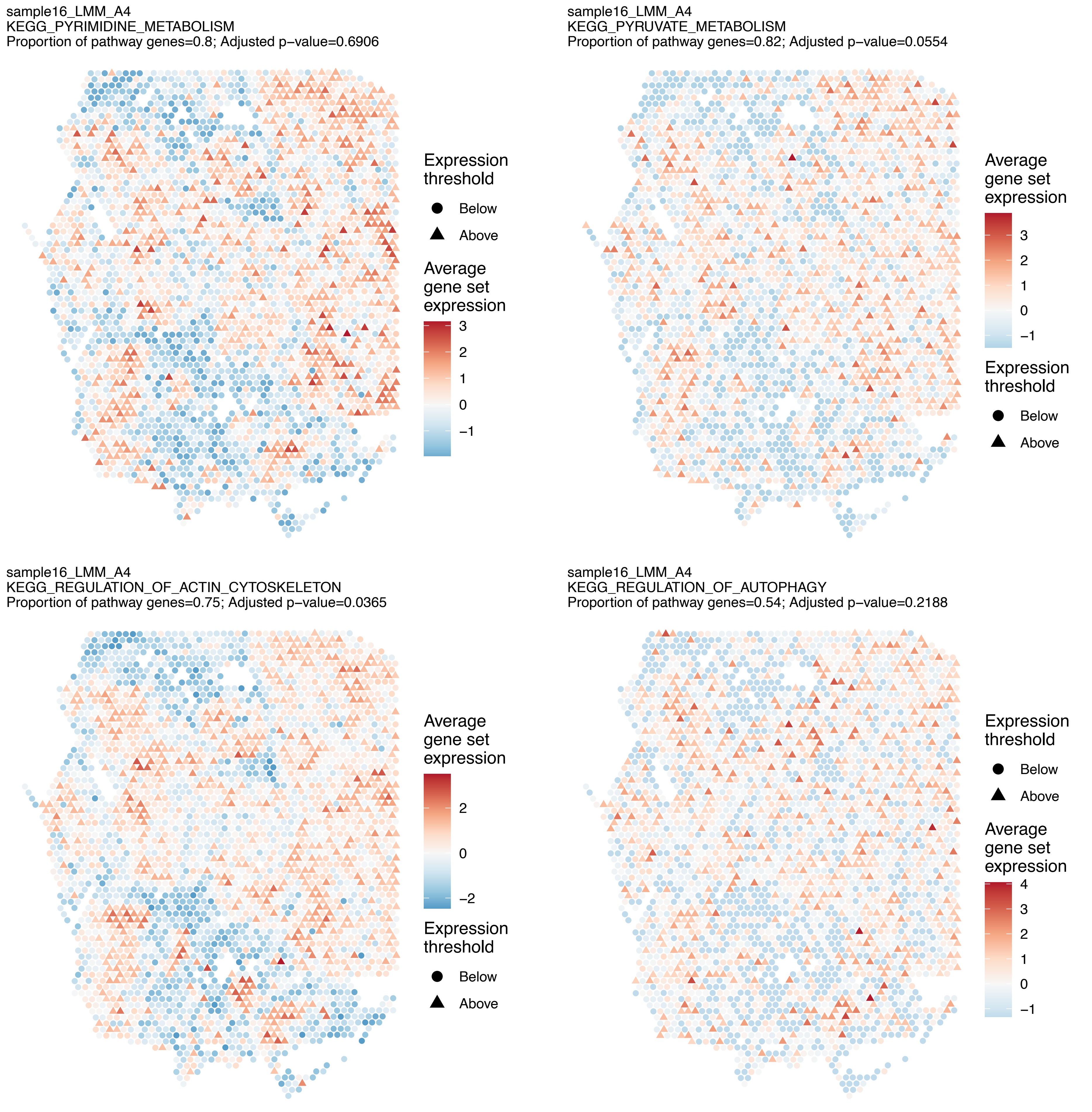
sample16_LMM_A4
KEGG_PYRIMIDINE_METABOLISM
Proportion of pathway genes=0.8; Adjusted p−value=0.6906
Expression threshold
Below
Above
Average gene set expression
3
2
1
0
−1
sample16_LMM_A4
KEGG_PYRUVATE_METABOLISM
Proportion of pathway genes=0.82; Adjusted p−value=0.0554
Average gene set expression
3
2
1
0
−1
Expression threshold
Below
Above
sample16_LMM_A4
KEGG_REGULATION_OF_ACTIN_CYTOSKELETON
Proportion of pathway genes=0.75; Adjusted p−value=0.0365
Average gene set expression
2
0
−2
Expression threshold
Below
Above
sample16_LMM_A4
KEGG_REGULATION_OF_AUTOPHAGY
Proportion of pathway genes=0.54; Adjusted p−value=0.2188
Expression threshold
Below
Above
Average gene set expression
4
3
2
1
0
−1

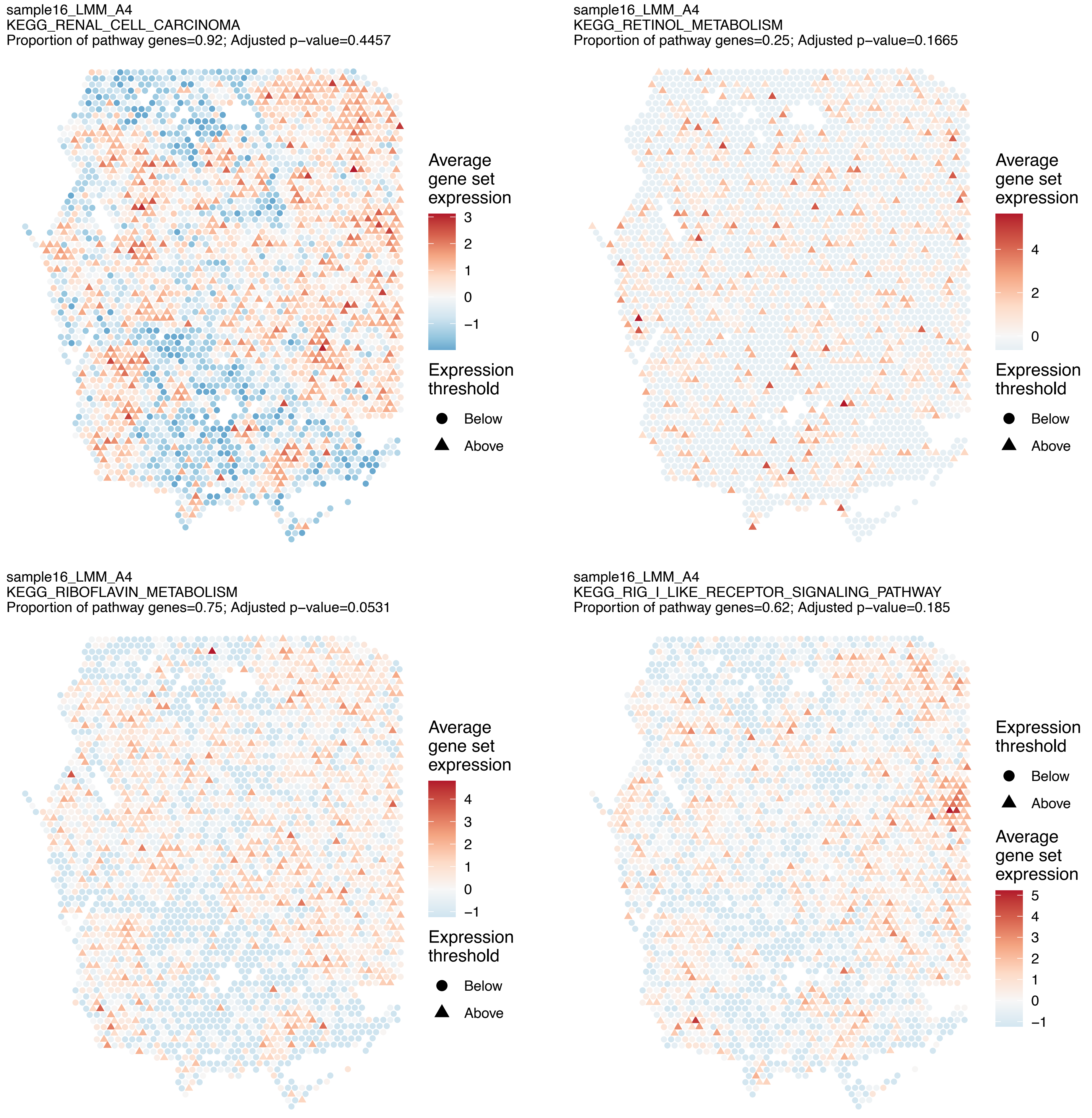
sample16_LMM_A4
KEGG_RENAL_CELL_CARCINOMA
Proportion of pathway genes=0.92; Adjusted p−value=0.4457
Average gene set expression
3
2
1
0
−1
Expression threshold
Below
Above
sample16_LMM_A4
KEGG_RETINOL_METABOLISM
Proportion of pathway genes=0.25; Adjusted p−value=0.1665
Average gene set expression
4
2
0
Expression threshold
Below
Above
sample16_LMM_A4
KEGG_RIBOFLAVIN_METABOLISM
Proportion of pathway genes=0.75; Adjusted p−value=0.0531
Average gene set expression
4
3
2
1
0
−1
Expression threshold
Below
Above
sample16_LMM_A4
KEGG_RIG_I_LIKE_RECEPTOR_SIGNALING_PATHWAY
Proportion of pathway genes=0.62; Adjusted p−value=0.185
Expression threshold
Below
Above
Average gene set expression
5
4
3
2
1
0
−1

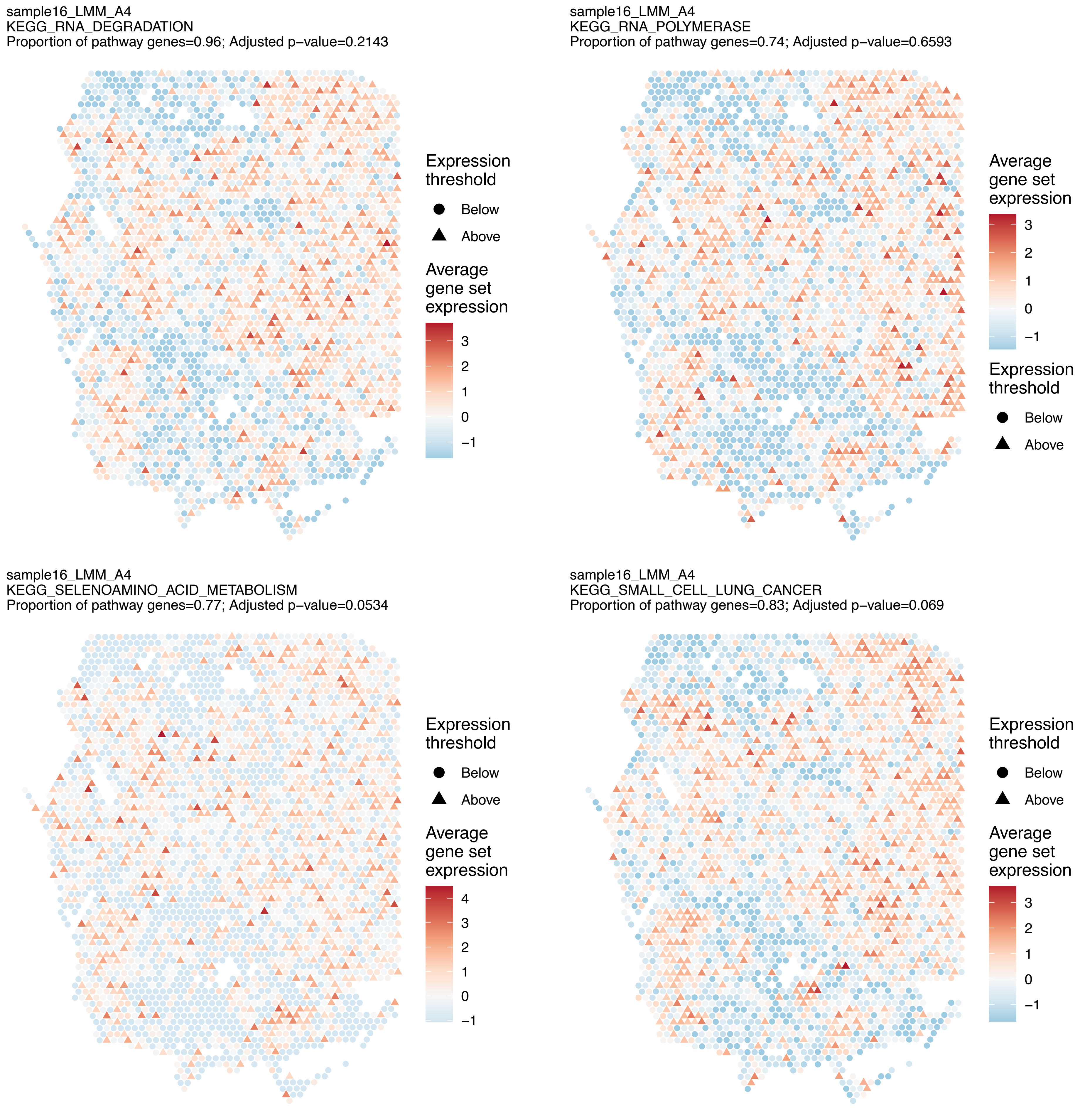
sample16_LMM_A4
KEGG_RNA_DEGRADATION
Proportion of pathway genes=0.96; Adjusted p−value=0.2143
Expression threshold
Below
Above
Average gene set expression
3
2
1
0
−1
sample16_LMM_A4
KEGG_RNA_POLYMERASE
Proportion of pathway genes=0.74; Adjusted p−value=0.6593
Average gene set expression
3
2
1
0
−1
Expression threshold
Below
Above
sample16_LMM_A4
KEGG_SELENOAMINO_ACID_METABOLISM
Proportion of pathway genes=0.77; Adjusted p−value=0.0534
Expression threshold
Below
Above
Average gene set expression
4
3
2
1
0
−1
sample16_LMM_A4
KEGG_SMALL_CELL_LUNG_CANCER
Proportion of pathway genes=0.83; Adjusted p−value=0.069
Expression threshold
Below
Above
Average gene set expression
3
2
1
0
−1

sample16_LMM_A4
KEGG_SNARE_INTERACTIONS_IN_VESICULAR_TRANSPORT
Proportion of pathway genes=0.89; Adjusted p−value=0.1443

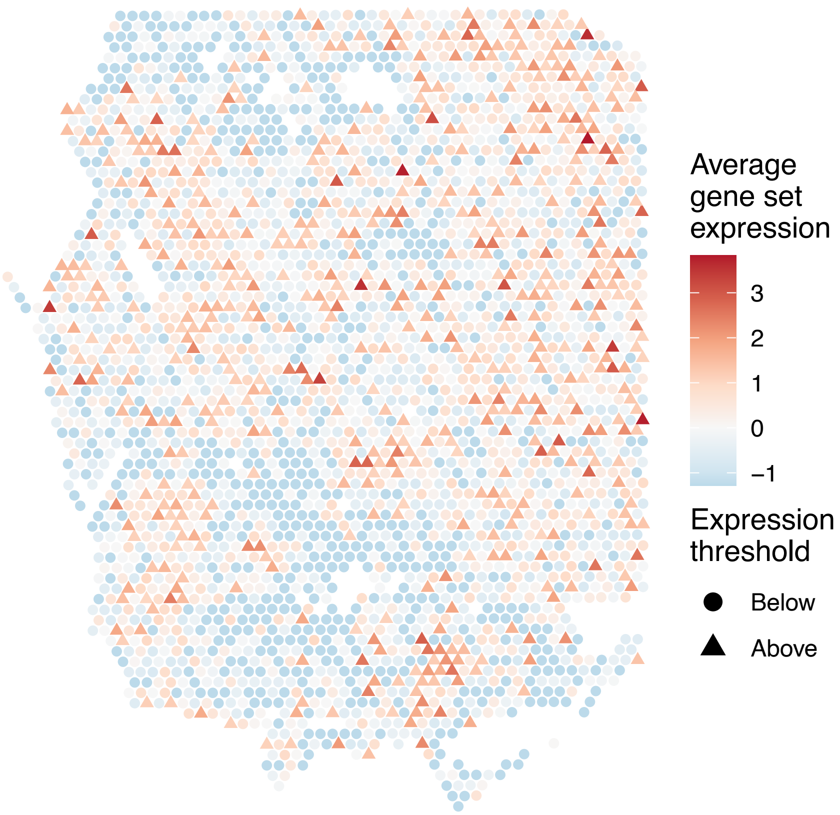


sample16_LMM_A4
KEGG_SPHINGOLIPID_METABOLISM
Proportion of pathway genes=0.76; Adjusted p−value=0.0067

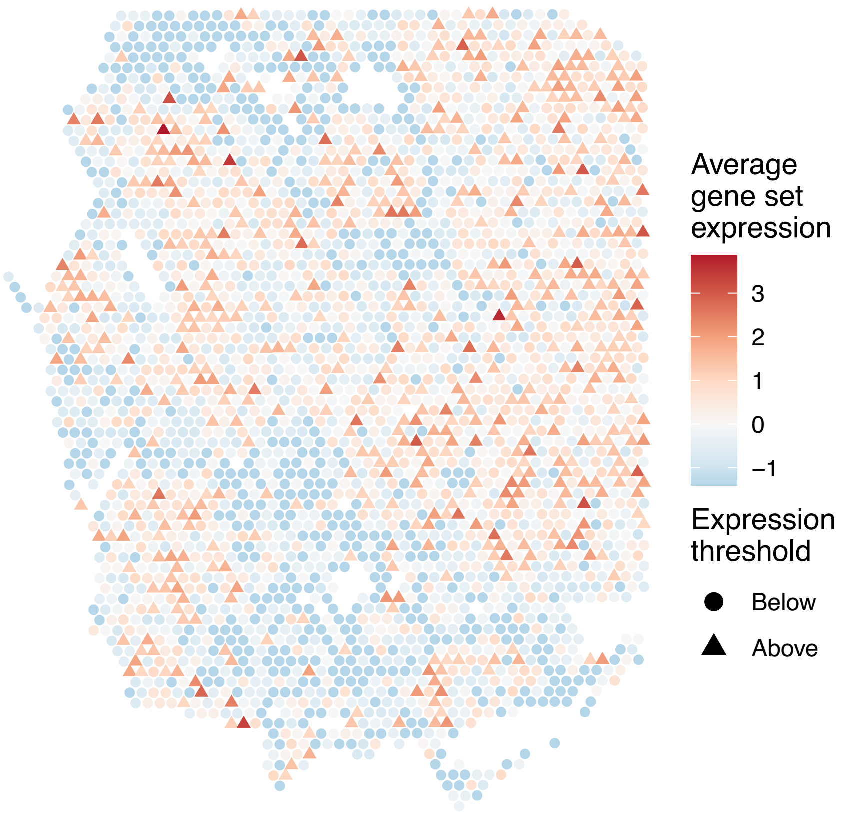


sample16_LMM_A4
KEGG_SPLICEOSOME
Proportion of pathway genes=0.99; Adjusted p−value=0.0067

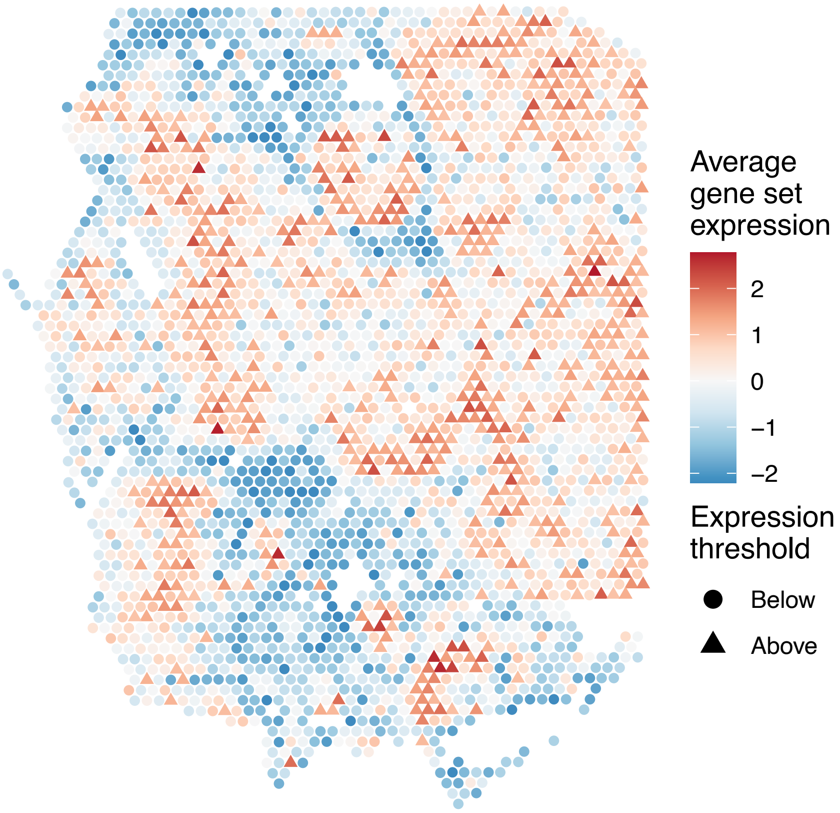


sample16_LMM_A4
KEGG_STARCH_AND_SUCROSE_METABOLISM
Proportion of pathway genes=0.37; Adjusted p−value=0.1408

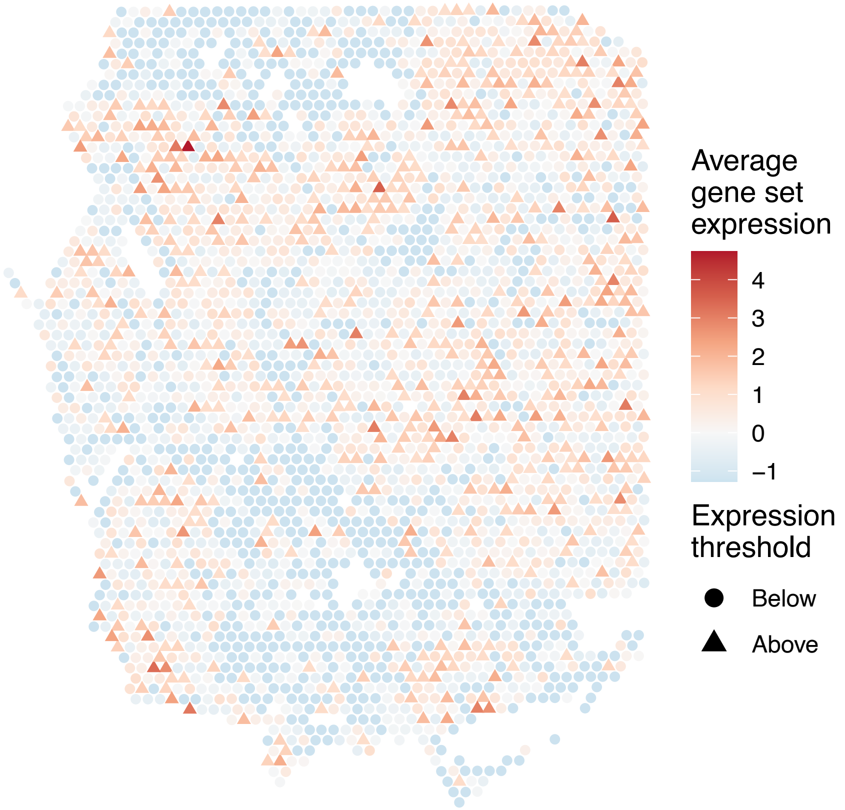

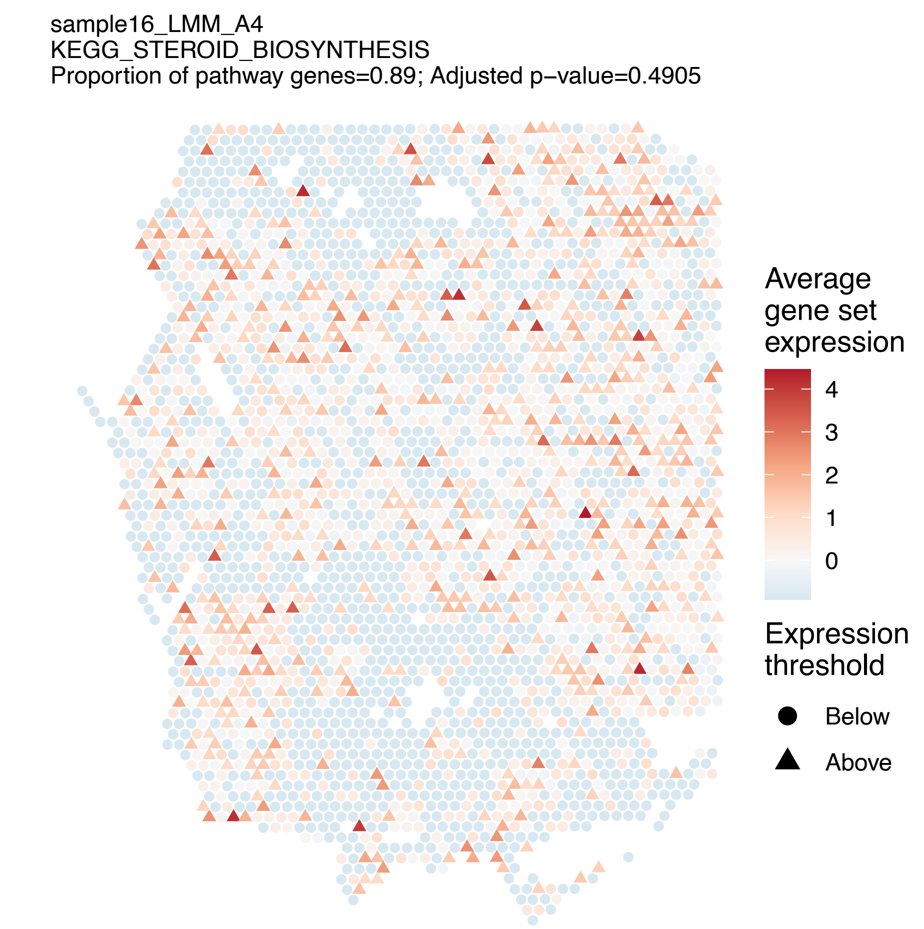
sample16_LMM_A4
KEGG_STEROID_BIOSYNTHESIS
Proportion of pathway genes=0.89; Adjusted p−value=0.4905
Average gene set expression
4
3
2
1
0
Expression threshold
Below
Above

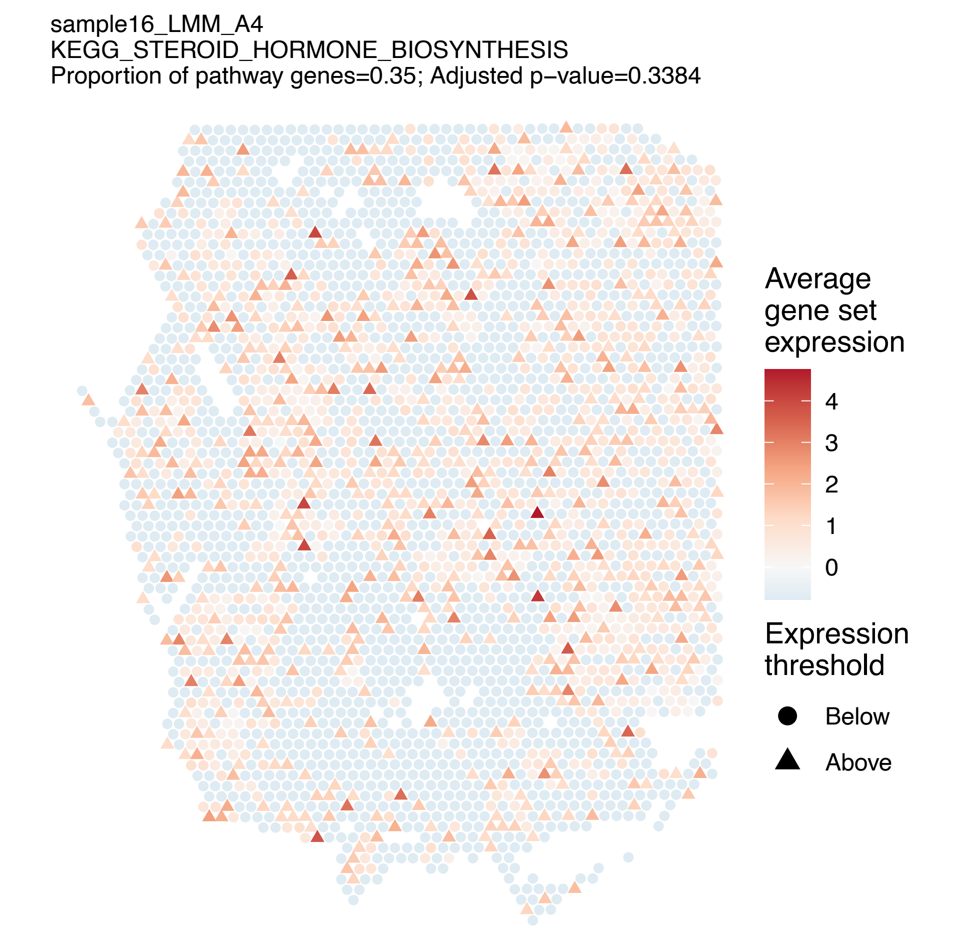
sample16_LMM_A4
KEGG_STEROID_HORMONE_BIOSYNTHESIS
Proportion of pathway genes=0.35; Adjusted p−value=0.3384
Average gene set expression
4
3
2
1
0
Expression threshold
Below
Above

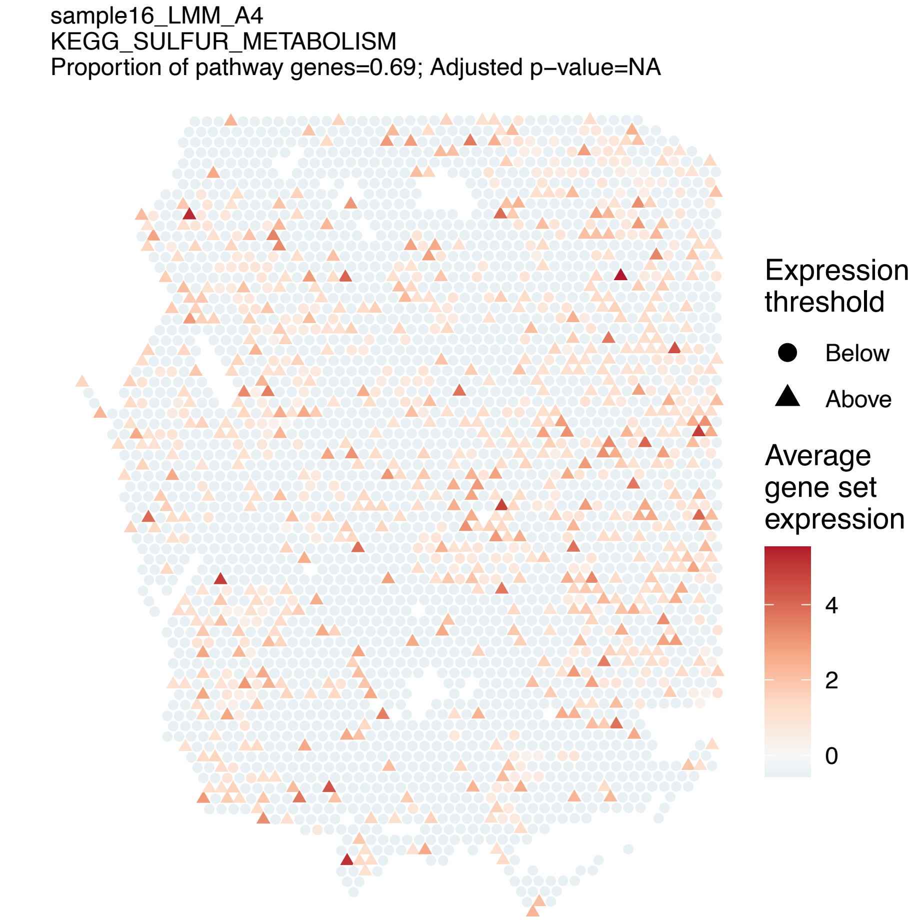
sample16_LMM_A4
KEGG_SULFUR_METABOLISM
Proportion of pathway genes=0.69; Adjusted p−value=NA
Expression threshold
Below
Above
Average gene set expression
4
2
0

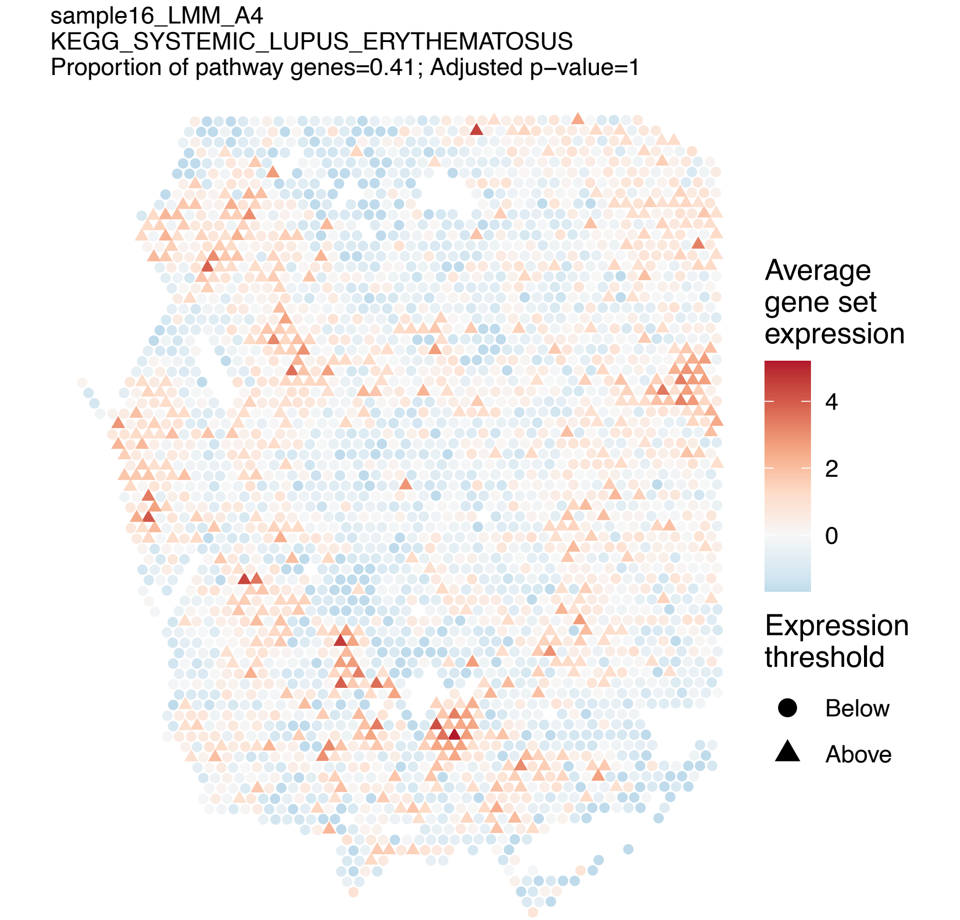
sample16_LMM_A4
KEGG_SYSTEMIC_LUPUS_ERYTHEMATOSUS
Proportion of pathway genes=0.41; Adjusted p−value=1
Average gene set expression
4
2
0
Expression threshold
Below
Above

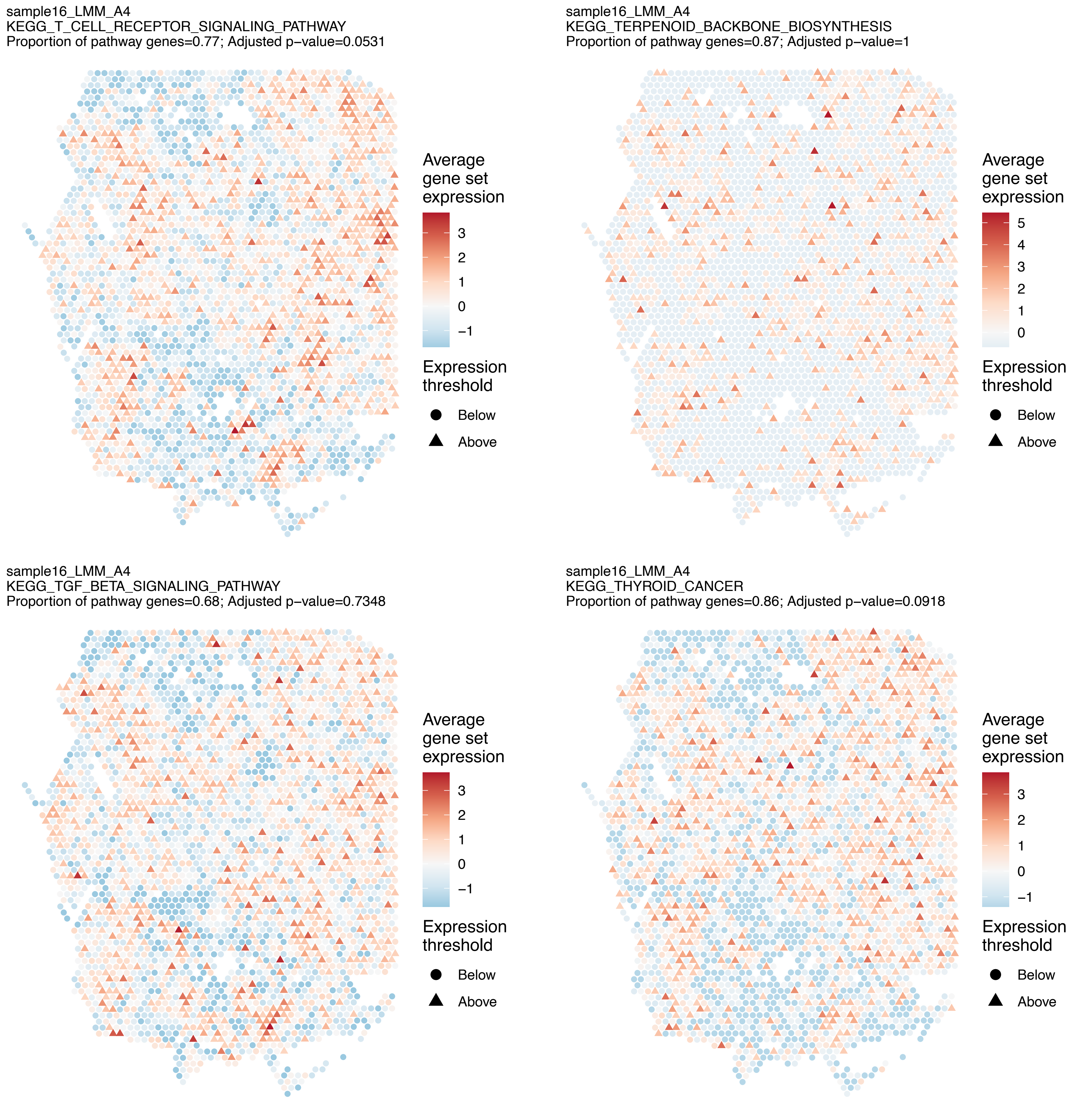
sample16_LMM_A4
KEGG_T_CELL_RECEPTOR_SIGNALING_PATHWAY
Proportion of pathway genes=0.77; Adjusted p−value=0.0531
Average gene set expression
3
2
1
0
−1
Expression threshold
Below
Above
sample16_LMM_A4
KEGG_TERPENOID_BACKBONE_BIOSYNTHESIS
Proportion of pathway genes=0.87; Adjusted p−value=1
Average gene set expression
5
4
3
2
1
0
Expression threshold
Below
Above
sample16_LMM_A4
KEGG_TGF_BETA_SIGNALING_PATHWAY
Proportion of pathway genes=0.68; Adjusted p−value=0.7348
Average gene set expression
3
2
1
0
−1
Expression threshold
Below
Above
sample16_LMM_A4
KEGG_THYROID_CANCER
Proportion of pathway genes=0.86; Adjusted p−value=0.0918
Average gene set expression
3
2
1
0
−1
Expression threshold
Below
Above

sample16_LMM_A4
KEGG_TIGHT_JUNCTION
Proportion of pathway genes=0.76; Adjusted p−value=0.0534

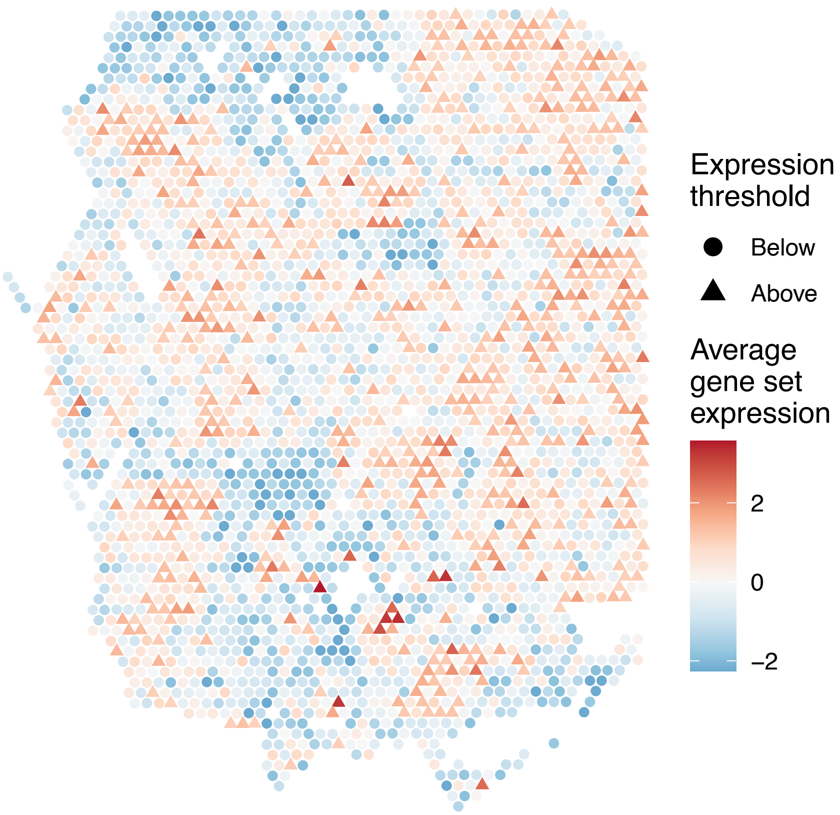


sample16_LMM_A4
KEGG_TOLL_LIKE_RECEPTOR_SIGNALING_PATHWAY
Proportion of pathway genes=0.66; Adjusted p−value=0.3383

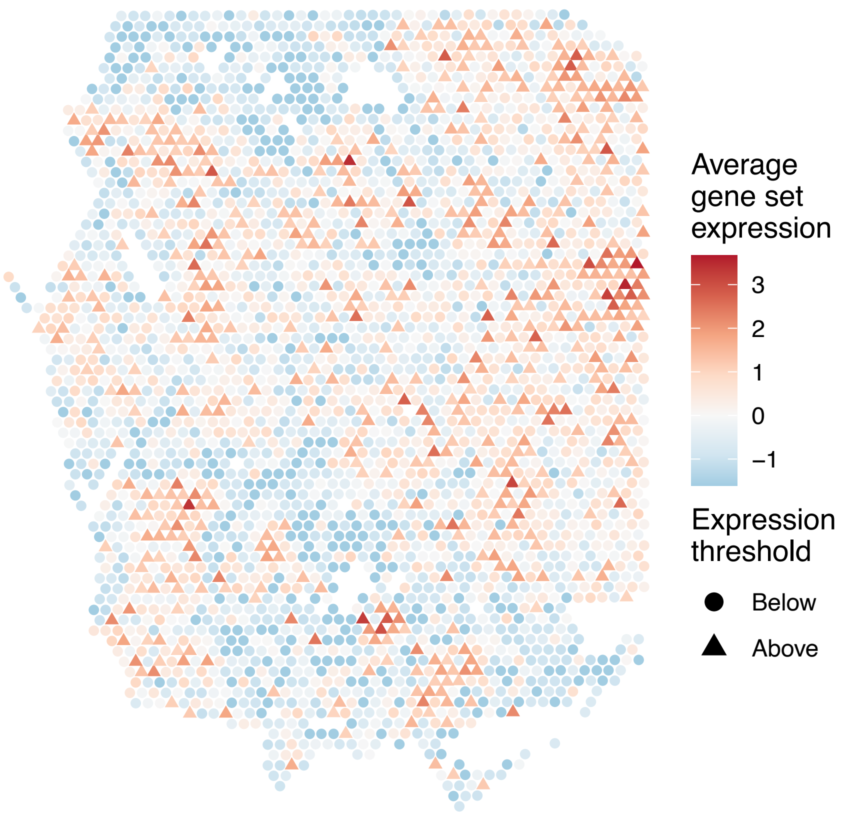


sample16_LMM_A4
KEGG_TRYPTOPHAN_METABOLISM
Proportion of pathway genes=0.49; Adjusted p−value=0.0471

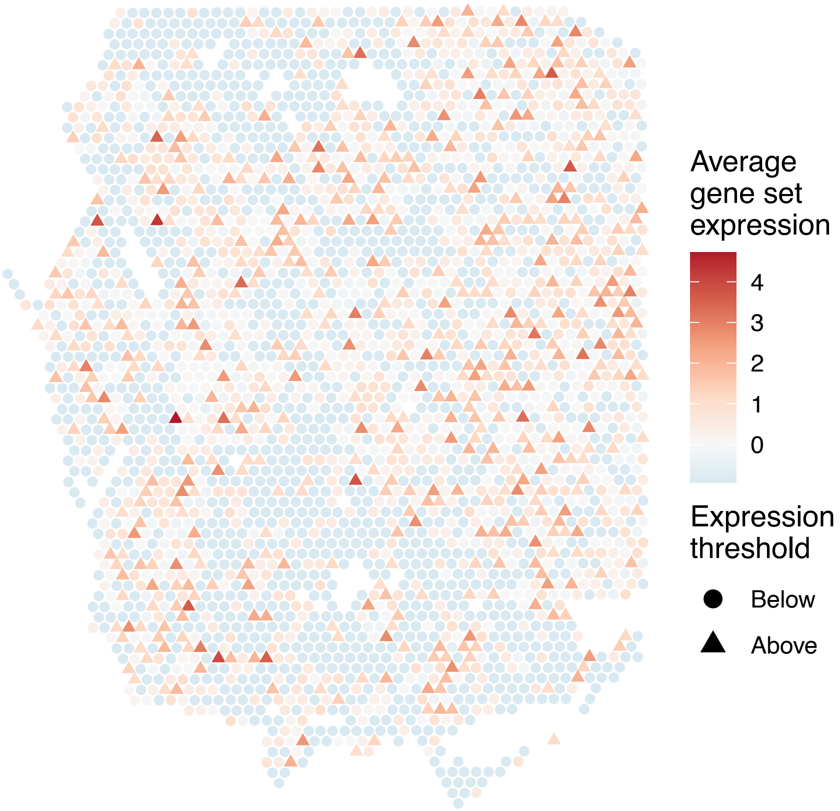


sample16_LMM_A4
KEGG_TYPE_I_DIABETES_MELLITUS
Proportion of pathway genes=0.72; Adjusted p−value=1

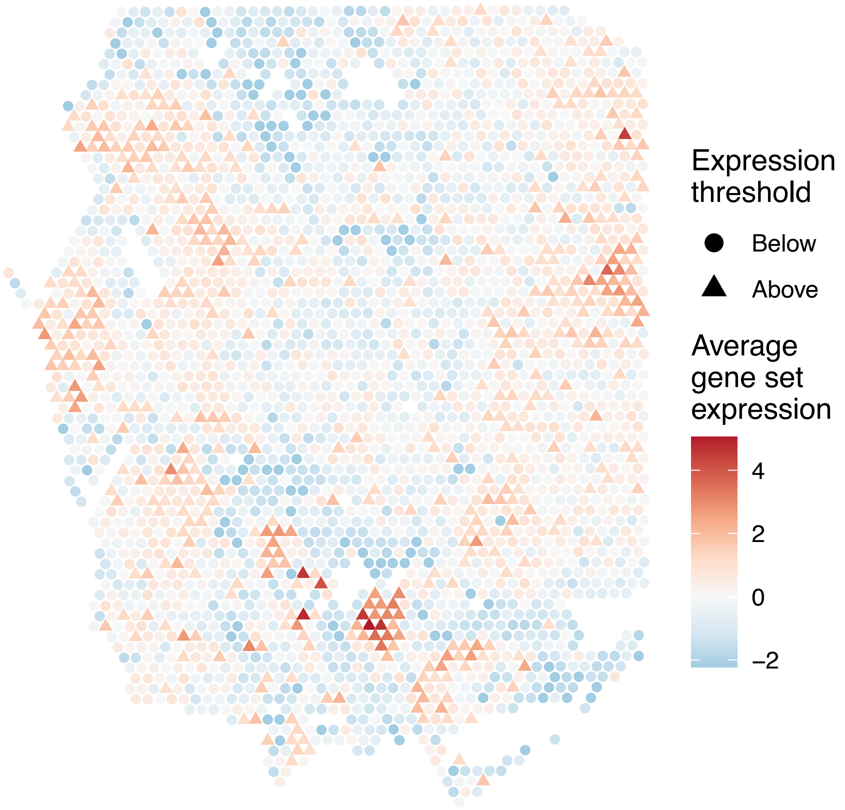

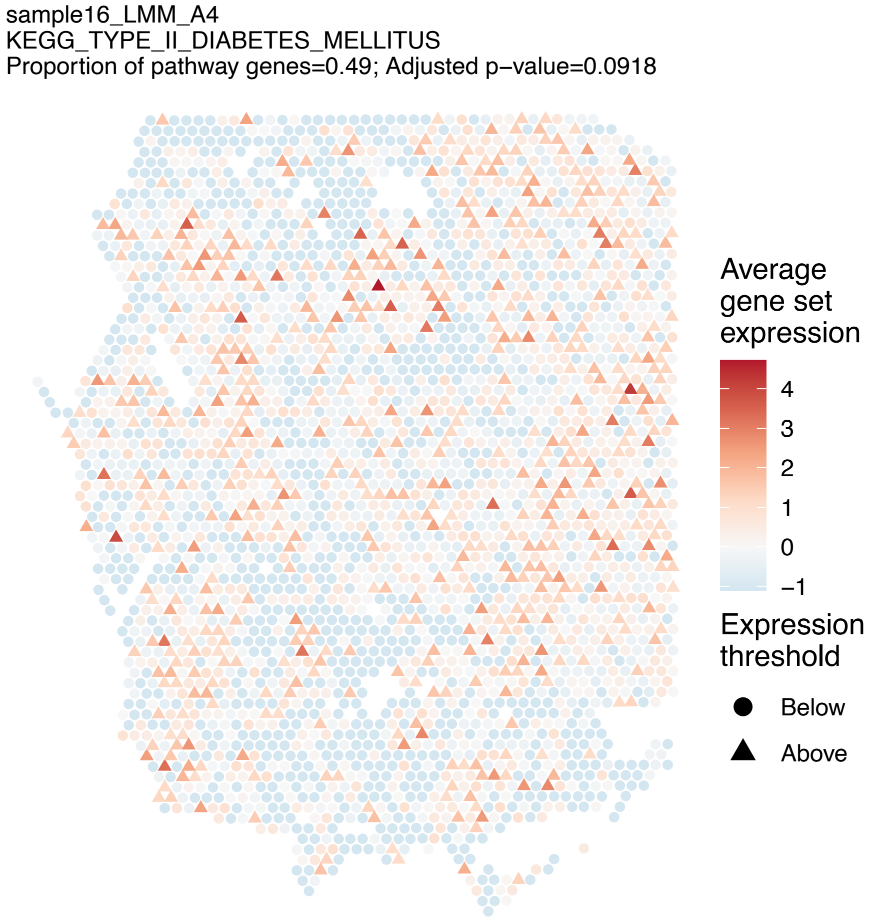
sample16_LMM_A4
KEGG_TYPE_II_DIABETES_MELLITUS
Proportion of pathway genes=0.49; Adjusted p−value=0.0918
Average gene set expression
4
3
2
1
0
−1
Expression threshold
Below
Above

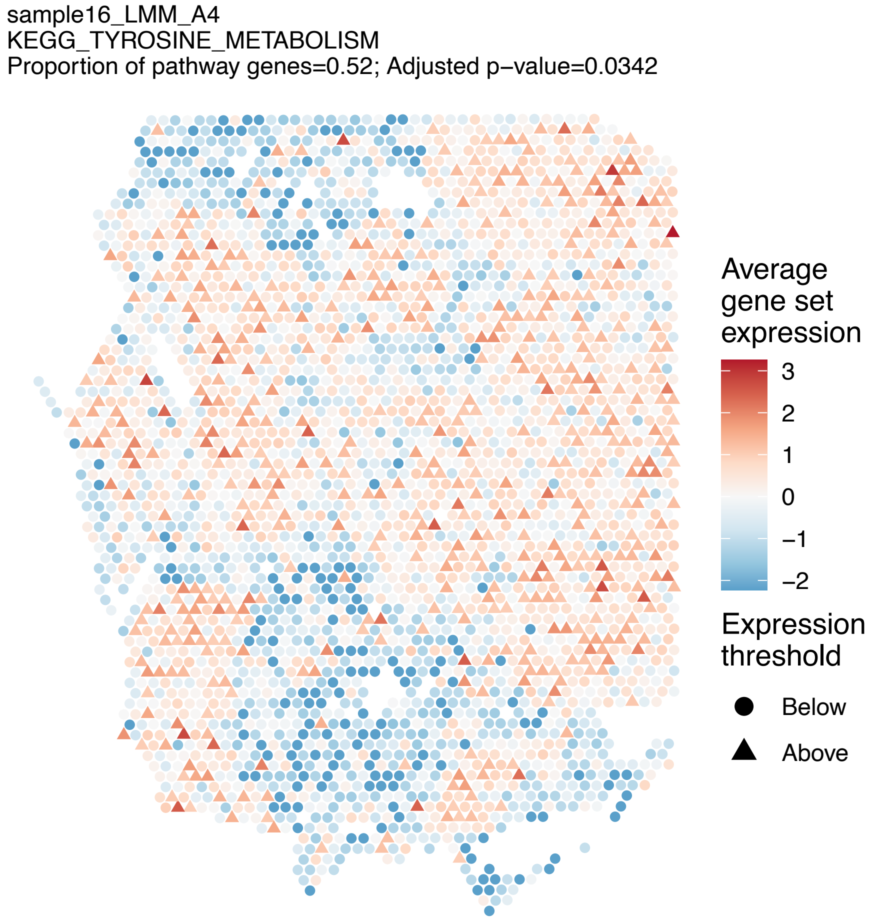
sample16_LMM_A4
KEGG_TYROSINE_METABOLISM
Proportion of pathway genes=0.52; Adjusted p−value=0.0342
Average gene set expression
3
2
1
0
−1
−2
Expression threshold
Below
Above

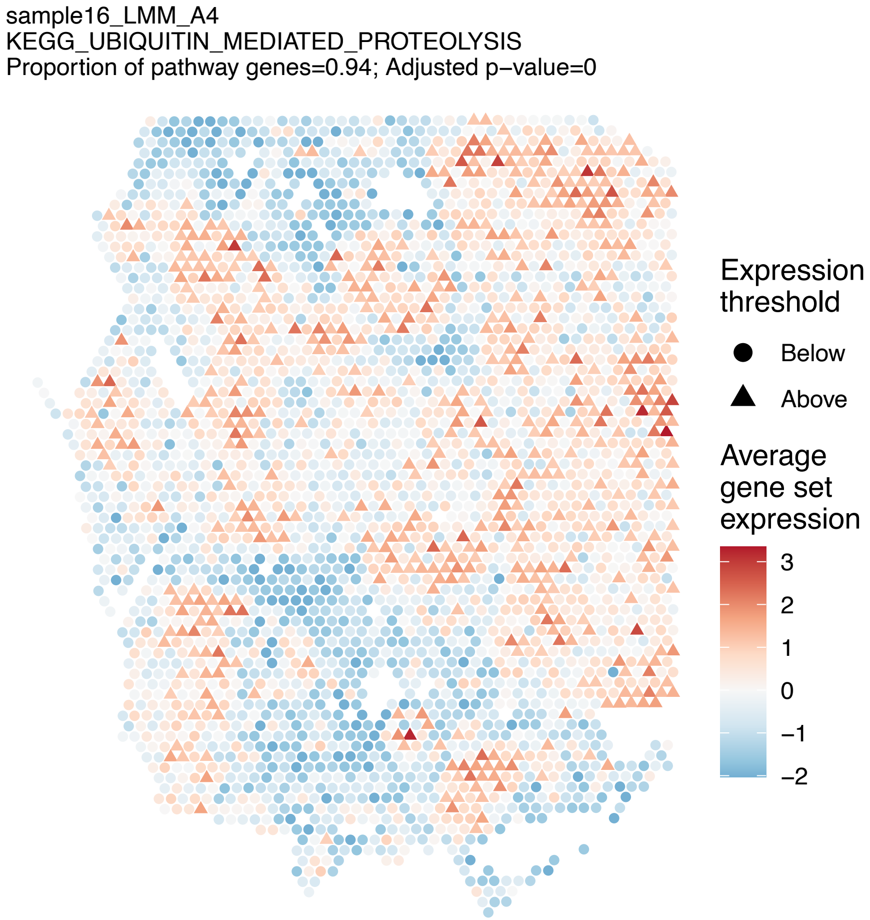
sample16_LMM_A4
KEGG_UBIQUITIN_MEDIATED_PROTEOLYSIS
Proportion of pathway genes=0.94; Adjusted p−value=0
Expression threshold
Below
Above
Average gene set expression
3
2
1
0
−1
−2

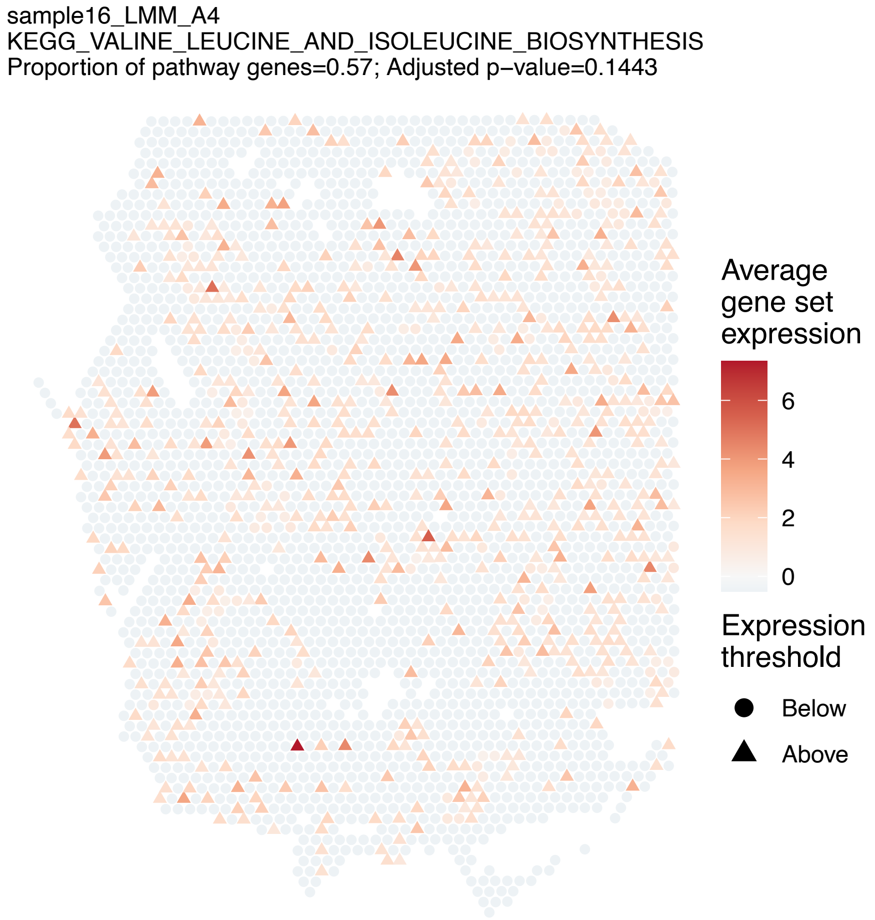
sample16_LMM_A4
KEGG_VALINE_LEUCINE_AND_ISOLEUCINE_BIOSYNTHESIS
Proportion of pathway genes=0.57; Adjusted p−value=0.1443
Average gene set expression
6
4
2
0
Expression threshold
Below
Above

sample16_LMM_A4
KEGG_VALINE_LEUCINE_AND_ISOLEUCINE_DEGRADATION
Proportion of pathway genes=0.93; Adjusted p−value=0.1086

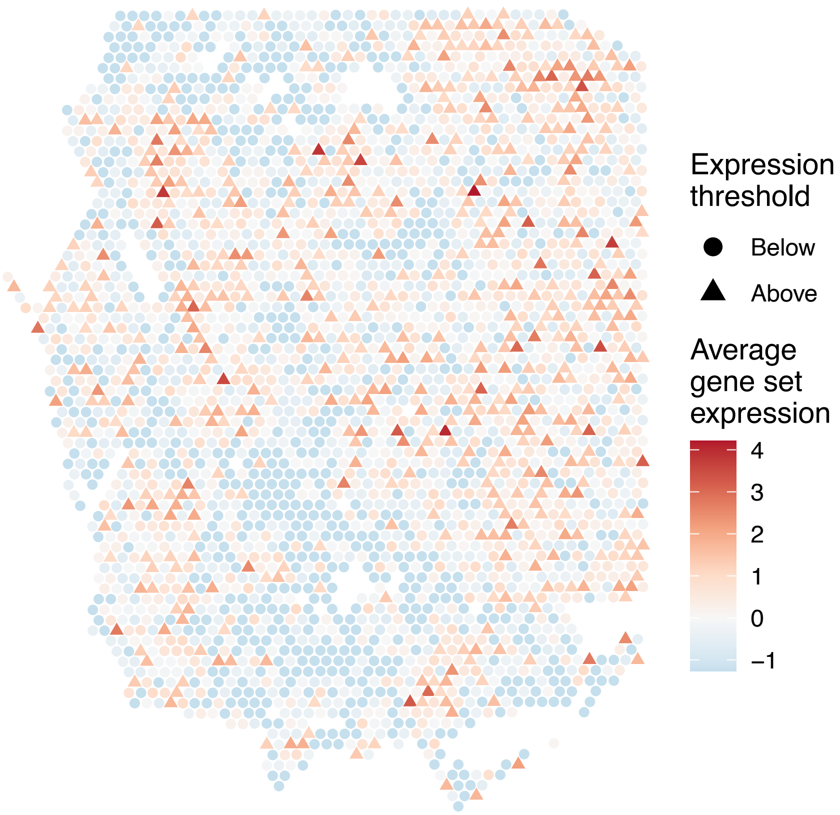


sample16_LMM_A4
KEGG_VASCULAR_SMOOTH_MUSCLE_CONTRACTION
Proportion of pathway genes=0.62; Adjusted p−value=0.3221

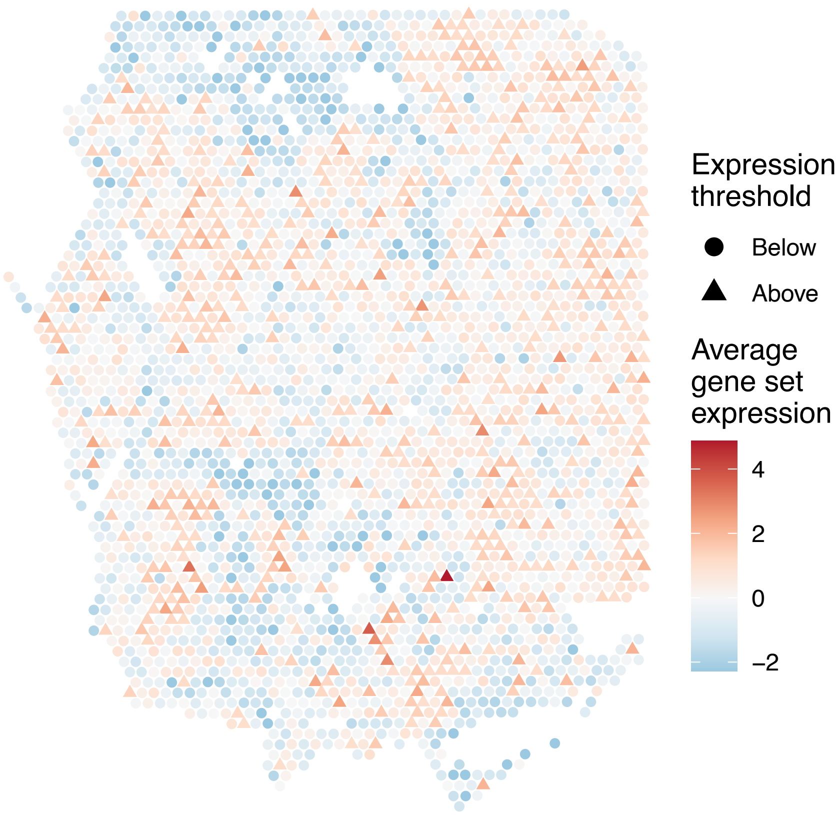


sample16_LMM_A4
KEGG_VASOPRESSIN_REGULATED_WATER_REABSORPTION
Proportion of pathway genes=0.85; Adjusted p−value=0.0574

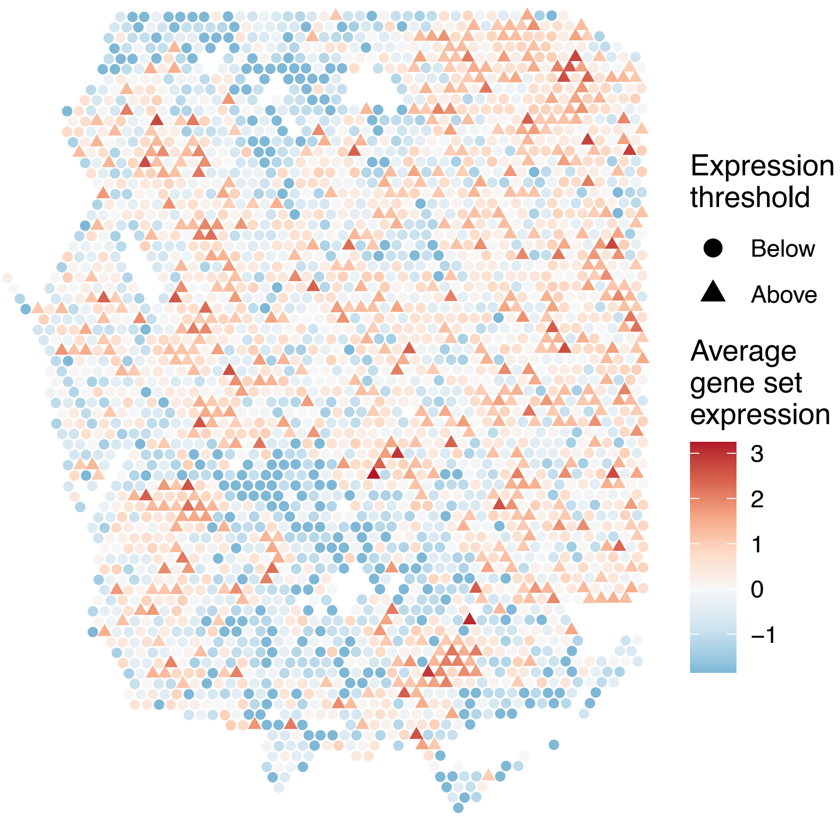


sample16_LMM_A4
KEGG_VEGF_SIGNALING_PATHWAY
Proportion of pathway genes=0.79; Adjusted p−value=0.3629

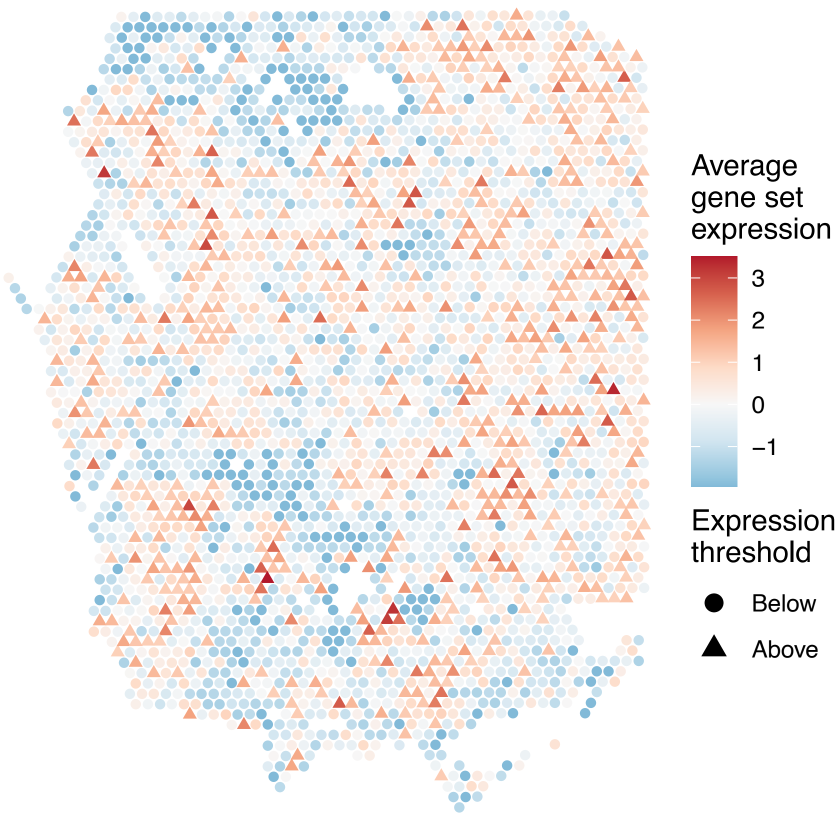

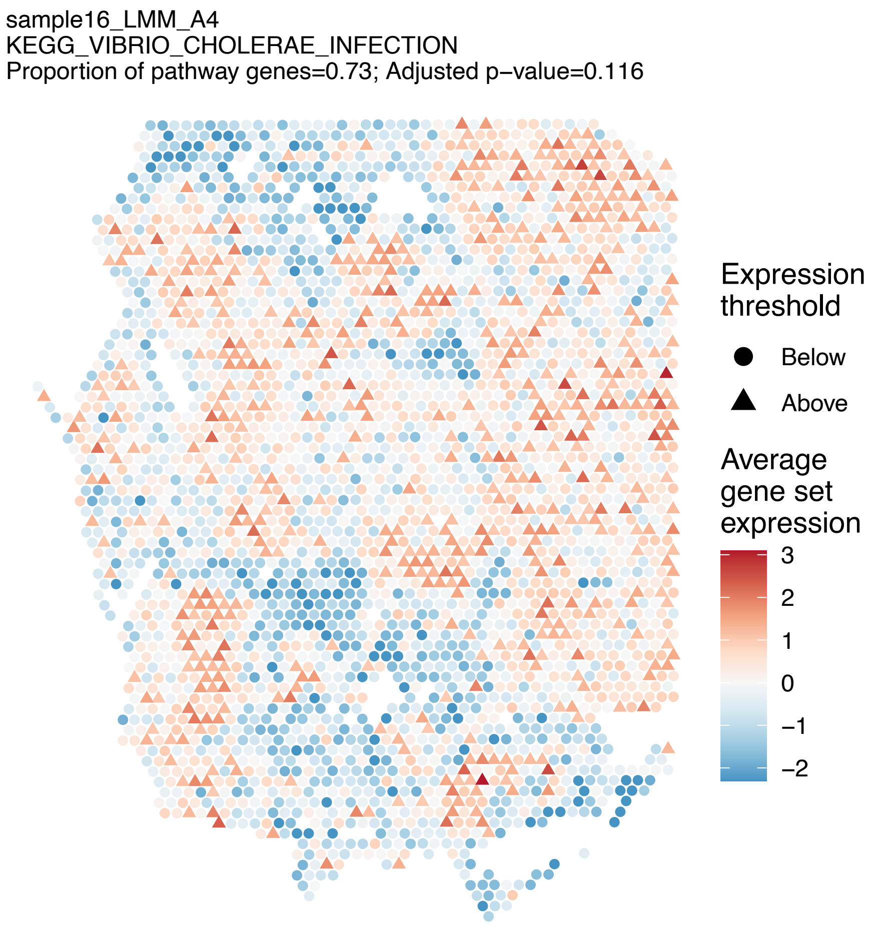
sample16_LMM_A4
KEGG_VIBRIO_CHOLERAE_INFECTION
Proportion of pathway genes=0.73; Adjusted p−value=0.116
Expression threshold
Below
Above
Average gene set expression
3
2
1
0
−1
−2

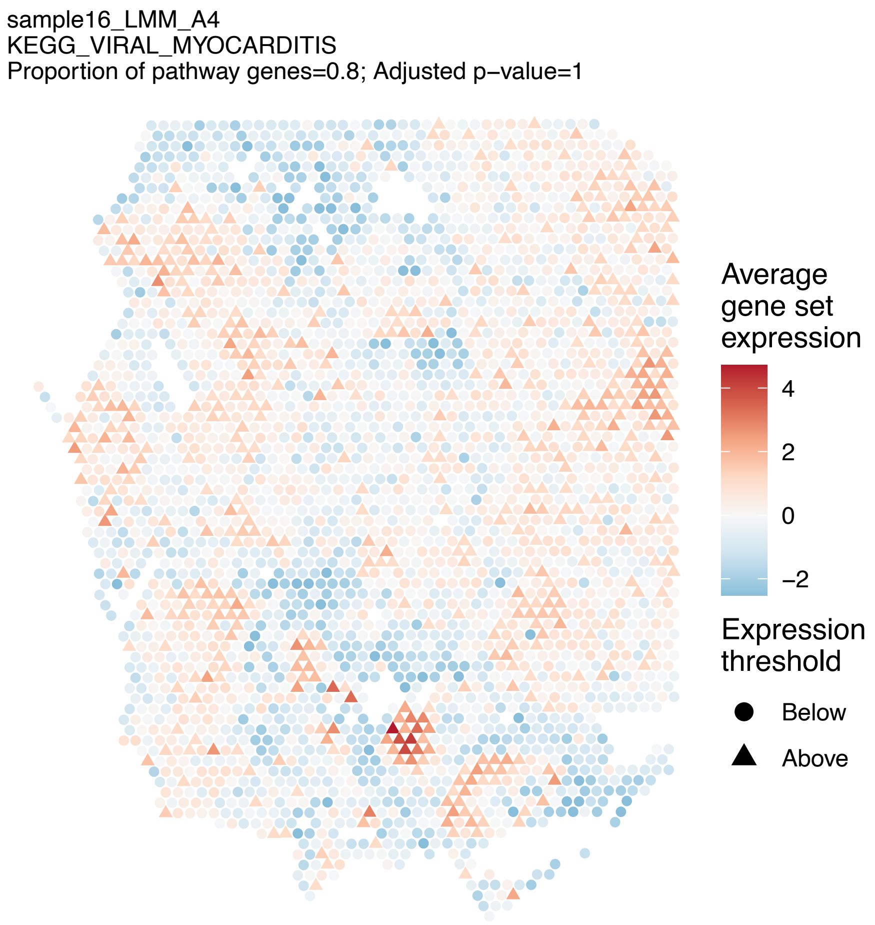
sample16_LMM_A4
KEGG_VIRAL_MYOCARDITIS
Proportion of pathway genes=0.8; Adjusted p−value=1
Average gene set expression
4
2
0
−2
Expression threshold
Below
Above

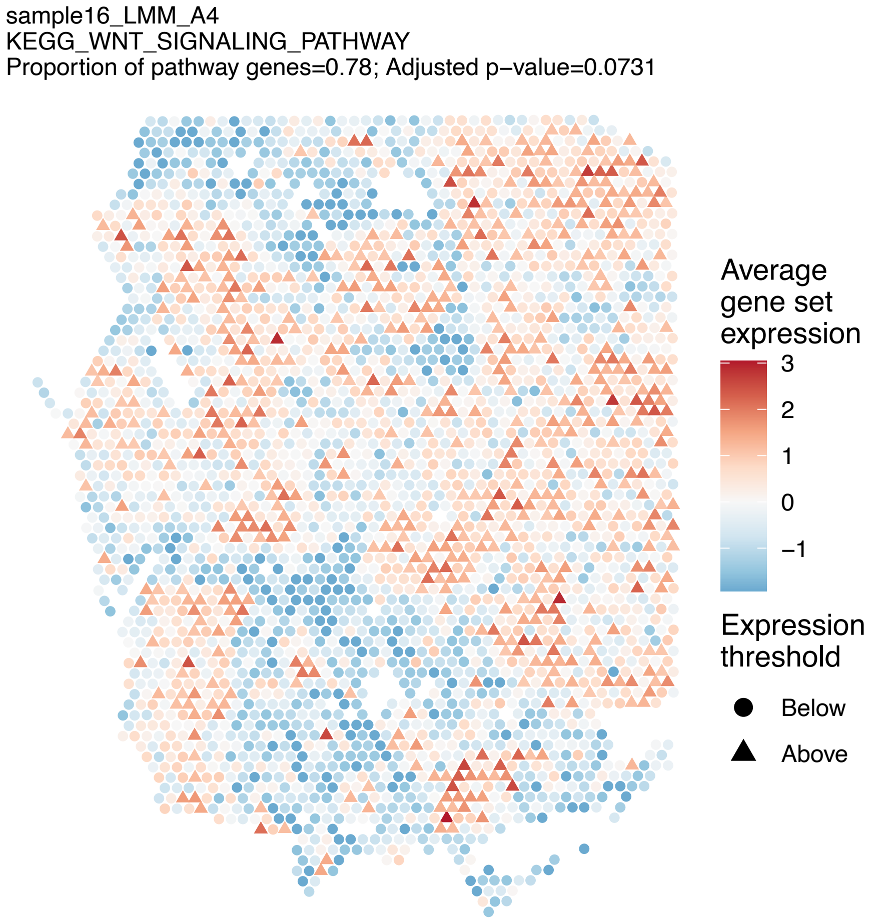
sample16_LMM_A4
KEGG_WNT_SIGNALING_PATHWAY
Proportion of pathway genes=0.78; Adjusted p−value=0.0731
Average gene set expression
3
2
1
0
−1
Expression threshold
Below
Above

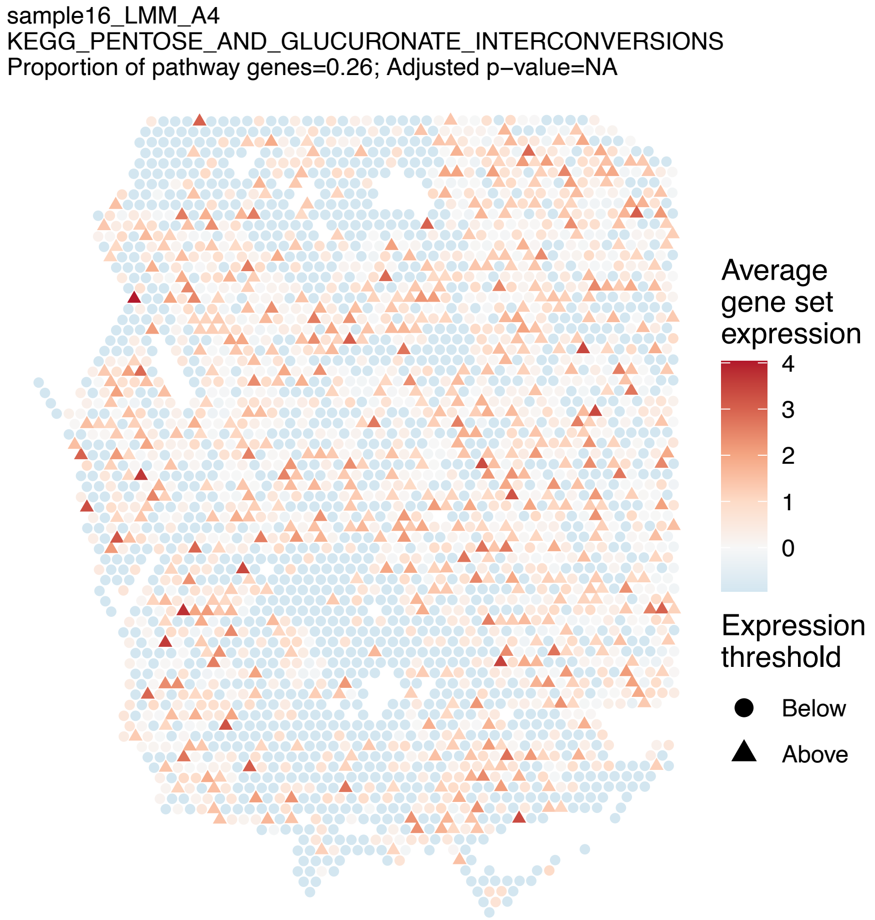
sample16_LMM_A4
KEGG_PENTOSE_AND_GLUCURONATE_INTERCONVERSIONS
Proportion of pathway genes=0.26; Adjusted p−value=NA
Average gene set expression
4
3
2
1
0
Expression threshold
Below
Above
